# Understanding Substance Dependence: What Differentiates Addictive from Non-Addictive Drugs?

**DOI:** 10.64898/2026.05.05.723067

**Authors:** Hargobind Singh, Hongyi Zhou, Samuel Skolnick, Jeffrey Skolnick

## Abstract

Addiction is a global health challenge, yet the molecular features that distinguish addictive from non-addictive drugs remain incompletely understood at the pathway and circuit levels. Here, we present a systematic computational framework that integrates drug-target binding predictions (provided by FINDSITE^comb2.0^) with brain-region-specific protein expression to compare addictive and non-addictive compounds. We analyzed 457 addictive and 1,774 non-addictive blood-brain barrier permeable drugs and mapped their predicted targets and associated pathways onto proteins expressed across 120 addiction-relevant brain regions. This analysis reveals widespread convergence between the two classes (addictive and non-addictive drugs) on shared molecular pathways, accompanied by distinct patterns of target and pathway engagement. Functional annotation of differentially engaged targets highlights biases toward plasticity-associated components for addictive drugs. In contrast, non-addictive drugs interact with both plasticity-associated proteins and proteins within the same molecular complex that have addiction suppression, regulatory, and homeostatic functions. Notably, both target classes co-localize within the same addiction-relevant circuits and form an integrated protein-protein interaction network. Together, these results define a differential engagement landscape that links chemical interactions to pathway-level utilization in the brain, revealing molecular features associated with differences in addiction propensity.

**Significance Statement:** Addiction is a global health crisis, yet the molecular features that distinguish addictive from non-addictive drugs remain poorly defined. By systematically comparing drug–target and pathway engagement across shared addiction-relevant brain circuits, this study identifies distinct signatures in which addictive compounds preferentially engage plasticity- and stress-associated targets, while non-addictive compounds engage these as well as addiction suppression, regulatory, transport, and metabolic pathways. These results provide a circuit- and pathway-level framework for interpreting addiction liability and guiding the design of safer therapeutics.

## Introduction

Substance use disorders (SUDs) represent a major and escalating global health crisis (3). Recent estimates indicate that nearly 292 million people worldwide, approximately 5.6% of the population aged 15-64, used drugs in 2022 (3, 4). SUDs are associated with a wide spectrum of adverse outcomes, including mental health disorders, infectious diseases, and chronic medical conditions, and they impose a substantial economic toll on healthcare and productivity (3, 4). Despite extensive research in addiction, effective therapeutic interventions remain limited, and relapse rates persistently range between 40-60% according to the National Institute of Drug and Abuse (NIDA) (5, 6).

Addiction is a neurobiological disorder involving interactions across molecular, cellular, and neural circuit levels (7, 8). Central to the development of addiction is the dysregulation of dopaminergic signaling within the mesolimbic pathway, which reinforces reward-seeking and motivational drivers (7, 8). Neuroadaptations in glutamatergic, GABAergic, and serotonergic systems further modulate craving, reinforcement, and emotional responses, while intracellular signaling cascades such as MAPK, cAMP/PKA, and mTOR pathways, along with epigenetic remodeling, contribute to persistent changes in neural plasticity (7–10). Chronic substance use also activates neuroimmune and stress response systems, exacerbating vulnerability to relapses and loss of control (7, 11). While the literature describes these addiction-promoting processes, comparatively less attention has been given to understanding how molecular engagement patterns differ between addictive and non-addictive compounds at the systems level.

Although individual addiction-related targets have been identified, systematic characterization of pathway and circuit-level engagement patterns that distinguish addictive potential remains incomplete. These observations raise broader questions about how molecular engagement/interaction patterns relate to variability in addiction risk. There is substantial evidence in the literature that reveals that, despite exposure to addictive substances, only a minority experience addiction (6, 7).

The availability of large-scale molecular datasets enables systematic analysis of addiction mechanisms. The Human Protein Atlas (HPA) maps proteins across 193 brain subregions, while virtual ligand screening algorithms like FINDSITE^comb2.0^ can often reliably predict exome-wide protein-ligand interactions (1, 2). FINDSITE^comb2.0^ is an advanced, structure-based virtual screening tool that helps predict protein-ligand interactions and identify potential drug targets. Pathway databases provide functional integration of individual protein target-level information, enabling the interpretation of biological processes and molecular mechanisms (12). We hypothesize that drugs differentially engage different molecular mechanisms in the brain, which can be identified through an integrated approach that involves: (a) Differential analysis of protein-ligand binding between addictive and non-addictive compounds, (b) Brain-region specific expression profiling, (c) Pathway engagement analysis to pinpoint the functionally relevant targets, pathways, and regions of interest. These can reveal molecular and pathway-level engagement patterns that distinguish addictive and non-addictive compounds.

## Results

An overview of the methodology is provided in Figure 1.

**Figure 1.**
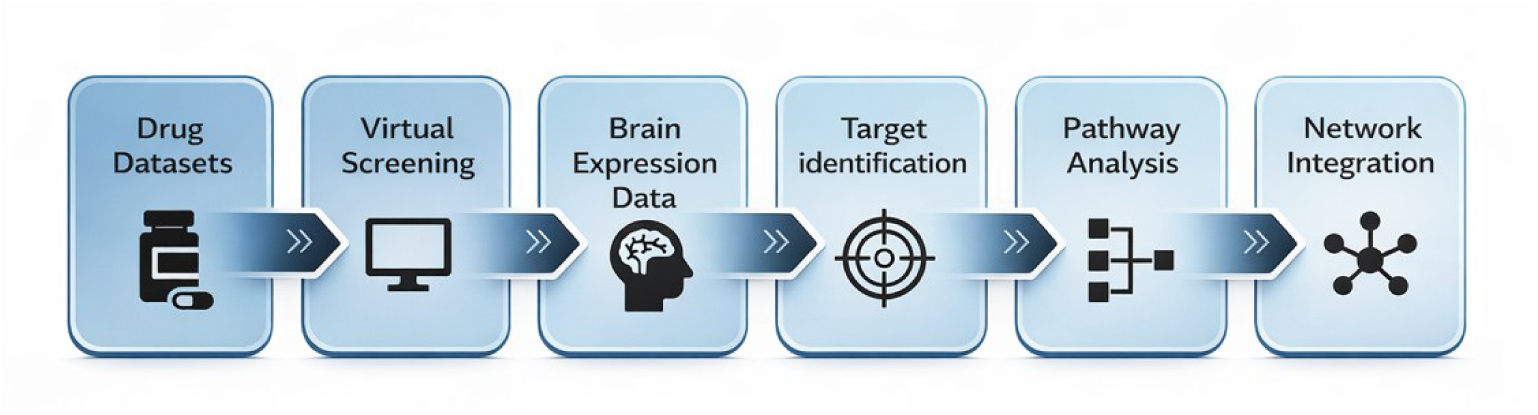
Computational workflow. Addictive and non-addictive drugs (n = 457 and n = 1,774) were analyzed using FINDSITE^comb2.0^ to derive enrichment factors (EF > 2 for addictive and EF < 0.5 for non-addictive) (1). Enriched targets were mapped onto HPA brain regions (the top 5 brain regions for each target were selected from 193 subregions) (2). This yielded 120 unique overlapping brain regions for addictive and non-addictive targets. Pathway engagement was performed to identify the pathways specifically associated with addictive drugs or non-addictive drugs. The protein-protein interaction network was created to determine if the predicted addictive and non-addictive targets and their associated networks are independent or coupled.

**Table 1.** The number of brain regions in which targets enriched for addictive and non-addictive drugs are co-expressed, as well as regions uniquely associated with each target class.

| Target Set | # of Overlapping Regions | # of Addictive-Specific Brain regions | # of Non-Addictive Specific Brain regions | Total Unique Regions |
| --- | --- | --- | --- | --- |
| Tier 1<br>(High Confidence) | 45 | 81 | 2 | 128 |
| Tier 2<br>(High Consensus) | 120 | 10 | 32 | 162 |

### Addictive and Non-Addictive Drugs Classification

To ensure that the two classes of drugs differ in chemical nature and capture distinct signals, compounds were assembled from Drug Enforcement Administration (DEA) scheduling lists and bioactivity records in ChEMBL (13, 14). Next, chemical similarity was assessed using the Morgan Fingerprint (radius = 2, corresponding to a circular neighborhood extending 2 bonds from each atom, 2048 bits), and compounds were clustered using the Butina Algorithm with different Tanimoto thresholds (15). A threshold of 0.95 Tanimoto similarity was used to collapse near analog compounds, while a similarity threshold of 0.80 captured broader chemotype-level similarity. For a 0.80 threshold, 2,689 total drugs remain, with 462 addictive drugs and 2,227 non-addictive drugs.

For analysis, we use the set generated with a 0.80 threshold of Tanimoto similarity to ensure that the full analysis remains robust. After clustering, using RDKit, we used blood-brain barrier (BBB) descriptors to make sure that drugs have the potential for brain exposure. Of the total of 2,227 non-addictive drugs, 1,774 showed blood-brain barrier permeability, and of the 462 addictive drugs, 457 can cross the blood-brain barrier. These drugs were then used for downstream analysis. The addictive and non-addictive drug lists can be found in Supplementary files S1 and S2.

### Target Classification for Drugs

To ensure chemical diversity, we restricted our analysis to BBB-permeable drugs clustered with a Tanimoto coefficient of 0.80. Proteome-wide binding probabilities of the drug binding to a particular protein were predicted using FINDSITE^comb2.0^, retaining interactions with a predicted binding probability ≥ 0.70 (1). These predicted protein targets were subsequently filtered for brain-relevance using Human Protein Atlas expression data (pTPM (protein transcripts per million) >= 1) (2). Drugs with ≥ 1 interaction in the brain were retained. A total of 421 addictive drugs out of 457 blood-brain barrier permeable drugs and 1,448 out of 1,774 non-addictive drugs had at least one interaction with a protein found in the brain. These drugs collectively engaged (i.e., interacted with) 4,076 unique protein targets in the addictive class and 13,123 unique protein targets in the non-addictive class. On average, addictive drugs interacted with more protein targets per compound than non-addictive drugs (mean 167.9 vs 112.6 proteins per drug).

We calculated the Enrichment Factor (EF) (see Equation 1, Methods) for each target from the blood-brain barrier universe. We assessed statistical significance using a two-sided Fisher’s exact test with the Benjamini-Hochberg (False Discovery Rate) correction. To maximize the recovery of biologically relevant signals, we defined two complementary target sets and begin by considering Tier 1 (high confidence) targets that were selected based on strict statistical criteria: Addictive: EF > 2, FDR ≤ 0.05, N_add (number of addictive targets) = 815, Non-addictive: EF < 0.5, FDR ≤ 0.05, N_non (number of non-addictive targets) = 42.

The marked asymmetry in the Tier 1 target counts between addictive and non-addictive drugs reflects a fundamental difference in the number of distinct proteins that they bind. While non-addictive drugs exhibited extensive absolute class-level target coverage, binding to a widely distributed set of ∼13,123 different proteins compared to the more focused set of proteins that bind addictive drugs of 4,076. Individually, addictive drugs show high target coverage (the average number of distinct proteins bound per addictive drug) of 168 protein targets compared to the non-addictive drugs (average number of distinct proteins bound per non-addictive drug) of 112. To quantify the degree of overlap among shared targets within each class, we calculated the pairwise Jaccard index. Non-addictive drugs show a mean Jaccard index of 0.013 while addictive drugs show a Jaccard index of 0.082, a sixfold difference. Inter-class comparisons yielded an intermediate but still modest mean Jaccard index of 0.022, indicating partial overlap in target space but much stronger convergence within the addictive class. This indicates that not only do non-addictive drugs collectively engage a large and diverse target space, but individual compounds within this class interact with rather distinct sets of proteins. In contrast, addictive drugs converge on a shared subset of targets, despite their individually broad interaction profiles.

This fundamental asymmetry in binding behavior imposed a severe multiple-hypothesis-testing penalty on the non-addictive class under strict FDR thresholds, masking biologically relevant consensus targets. To address this limitation, we implemented a dual-tier approach and next considered Tier 2 (high consensus) targets defined by interaction consistency, independent of statistical significance: EF thresholds met; n ≥ 10 supporting compounds; N_add = 941; N_non = 1,299. Analyzing Tier 2 proteins, we recovered high-consensus targets consistently engaged by diverse chemotypes but diluted by the statistical background of the expanded landscape.

Polypharmacology (the number of protein targets bound per drug), target convergence, and class-level target diversity of both drug classes are summarized in Figure 2. Together, these analyses reveal a clear distinction: addictive compounds exhibit high convergent polypharmacology, preferentially engaging a shared subset of protein targets, while non-addictive compounds display broad, distributed interactions across a diverse and largely non-overlapping target space.

**Figure 2.**
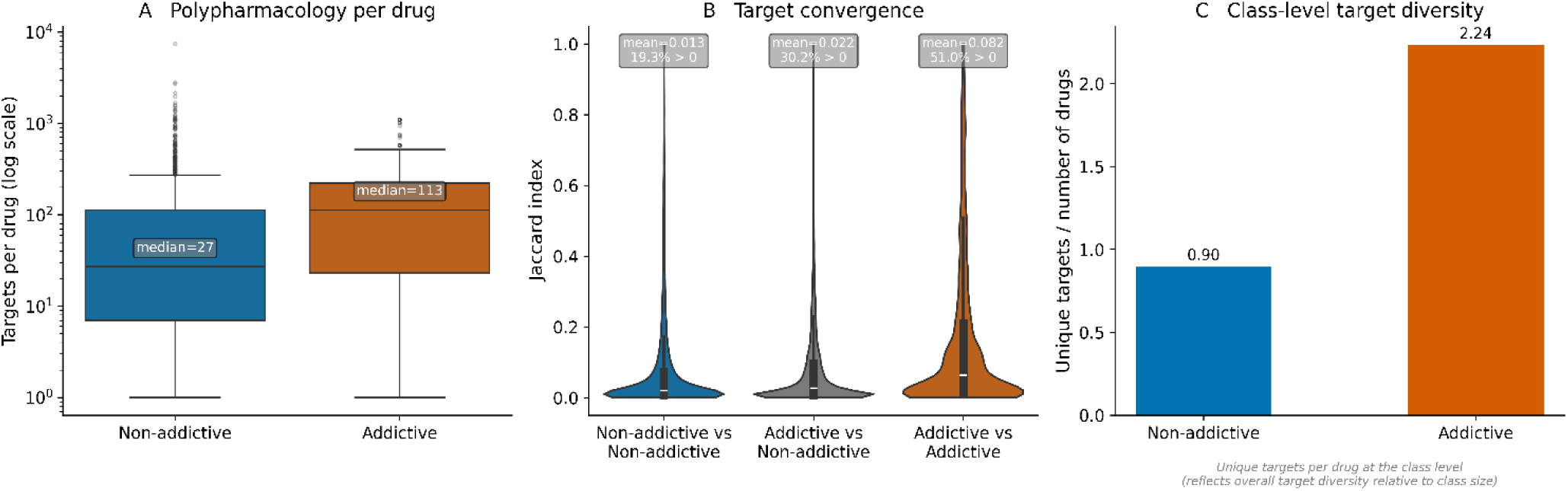
Distinct polypharmacology and target engagement patterns differentiate addictive and non-addictive drugs. (A) Distribution of the number of predicted protein targets per drug for non-addictive and addictive compounds, shown on a logarithmic scale. Addictive drugs exhibit increased polypharmacology, with a higher median number of targets bound per compound. (B) The Jaccard index quantifies pairwise target convergence for all three drug pairs that share at least one target. While both classes show substantial overlap in protein target space, addictive drugs show markedly higher within-class convergence (mean Jaccard = 0.082). (C) Class-level target diversity, defined as the number of unique targets normalized by the number of drugs in each class. Addictive drugs collectively engage a more diverse target repertoire relative to non-addictive drugs. Together, these results indicate that although addictive and non-addictive compounds share substantial overlap in their molecular targets, addictive drugs are characterized by increased polypharmacology and expanded target diversity, suggesting differential engagement of the shared biological network.

Together, these results demonstrate that both classes exhibit polypharmacology but differ in organization. Non-addictive drugs show a more diverse profile of protein targets while addictive drugs exhibit convergent polypharmacology. The list of Tier 2 addictive and non-addictive enriched targets can be found in Supplementary files S3 and S4. In the next section, we examine whether the putative Tier 2 enriched targets are consistent with the literature.

### Literature Validation of Predicted Targets

To assess the biological plausibility of the predicted targets, we performed literature-based validation on a representative sample of 100 targets from each group (addictive and non-addictive). An LLM augmented evaluation pipeline was used (GPT-5.4 thinking with medium reasoning); see Methods. The results have a striking asymmetry between the two groups. Among the 100 addictive drug targets, 25% received a “likely True” verdict with literature support, while 44% returned “Mixed evidence”. This reflects partial, indirect, or context-dependent associations, while 31% returned “No evidence.” This likely reflects the presence of novel or unknown addiction-linked targets in the addiction-determining networks. If correct, these could increase our understanding of addiction and provide additional targets for addiction suppression. In contrast, the non-addictive target set showed markedly stronger literature support. Of the 100 claims evaluated, 95% received a positive verdict, with 5% returning “Mixed Evidence.” This near-complete validation confirms the presence of proteins with well-established roles in neural homeostasis, signal regulation, and cellular maintenance. The claim files can be found in the Supplementary Folder S5. Taken together, these validation rates support the biological coherence of both target sets. In the next section, we look at the brain sub-regions in which Tier 2 targets are being expressed.

### Brain Region Identification of Addictive and Non-addictive Protein Targets

To identify the anatomical brain loci of primary biological relevance, we determined the regions of interest for each target class. For every protein in the Tier 1 and Tier 2 sets, we selected the top 5 brain regions exhibiting the highest transcript expression (pTPM). We then performed a set overlap analysis on these regions to identify overlapping regions (anatomical structures shared by both classes) and class-specific brain regions (structures uniquely prioritized by one class). This approach highlights the candidate brain loci at which the predicted targets are spatially permissible and anatomically convergent, thereby providing a regional context for downstream analysis. The total unique regions represent the union of all regions identified across both target classes after deduplication of shared regions. This provides a comprehensive anatomical map of predicted target engagement.

Overlapping regions represent anatomical structures prioritized by both target sets, whereas class-specific regions indicate differential regional engagement.

In the high confidence Tier 1 set, the 45 overlapping regions correspond to the canonical addiction-associated anatomical core. Both drug classes converged on the mesolimbic dopamine system, specifically the nucleus accumbens, ventral tegmental area (VTA), and substantia nigra (7, 16, 17). Crucially, this overlap extended to key modulatory regions, including the noradrenergic locus coeruleus and serotonergic dorsal and median raphe nuclei (18, 19). We also observed shared engagement of the habenula, a critical node in negative reward prediction error, and the hippocampus, implicating shared impacts on reward, processing, arousal, and contextual memory (20, 21).

In Tier 2, the network was observed to extend to 120 brain regions (Figure 3), encompassing structures associated with the transition from goal-directed to habitual behavior and withdrawal-induced negative effects. Tier 2 added dorsal striatum (caudate nucleus, putamen) and the extended amygdala (bed nucleus of the stria terminalis), regions central to habit formation and stress responsiveness (22, 23). Furthermore, Tier 2 overlap included the anterior and posterior insular cortex and the claustrum, suggesting that broadly engaged targets modulate interoceptive processing and conscious integration of drug states (24–26).

**Figure 3.**
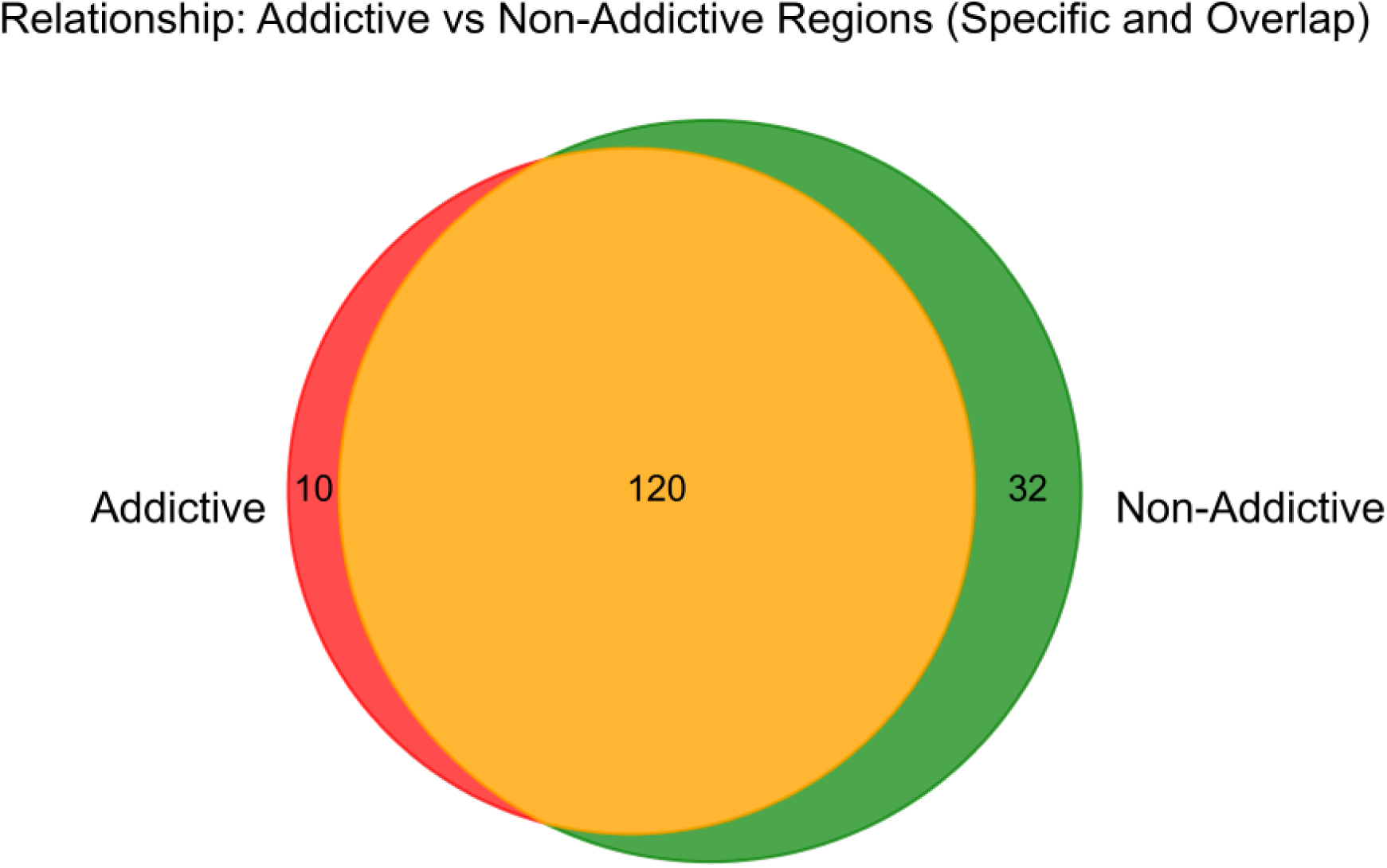
Brain-region overlap and specificity between targets enriched for addictive and non-addictive drugs. Brain regions were identified based on the highest transcript expression levels (pTPM) of tier 2 protein targets in the Human Protein Atlas. Regions shared between both target classes are shown in orange (n=120), while regions uniquely associated with addictive enriched targets are shown in red (n=10) and those uniquely associated with non-addictive enriched targets are shown in green (n=32). These results indicate substantial anatomical overlap between the two target classes, alongside a smaller subset of class-specific regions, providing spatial context for the potential neuroanatomical localization of addictive and non-addictive drug target engagement (i.e., binding).

Regional engagement was not limited to the classical addiction circuit. Both target classes showed high expression in circuits not traditionally associated with addiction. Among these are sensory processing (temporal gyrus, area striata) and autonomic brainstem nuclei (area postrema, nucleus ambiguus, pontine nuclei). Additionally, the white matter tracts occur in both tiers, including the corpus callosum and cerebellar white matter. Anterior/dorsal funiculi are also observed in the Tier 2 list, but not in the Tier 1 list. While addiction research has traditionally emphasized gray matter circuitry, neuroimaging studies have consistently reported white matter integrity changes in substance use disorders (27, 28). Our findings indicate that predicted targets for both addictive and non-addictive compounds are highly expressed in both regions, offering a potential molecular context for reported structural deficits. These regions contribute to the broader anatomical distribution of predicted non-addictive drug target binding, although further experimental validation is required. The widespread anatomical distribution suggests that the addiction liability is not determined solely by regional localization but rather by functional differences in how they are engaged. The list of regions can be found in Supplementary Files S6 and S7. In the next section, we look at the functional association patterns of these drugs.

### Pathway Occurrence in Addictive and Non-addictive Protein Targets

Given that regional analyses revealed substantial overlap between addictive and non-addictive protein targets, we next examined pathway-level interactions to identify functional differences that were not evident from brain-region analysis alone. Because pathways often contain both addiction-enriched and non-addiction-enriched targets, simple counts cannot distinguish whether drugs preferentially interact with one group or the other.

To resolve this, we developed normalized engagement metrics (per-drug engagement score, compositional engagement score, and pathway tilt; see Methods). These metrics quantify whether drugs preferentially bind addiction-enriched or non-addiction-enriched targets within the same pathway, while controlling for unequal target representation and assessing the uniformity of this bias across the drug class. Positive scores indicate a bias toward addiction-enriched targets, whereas negative scores indicate a bias toward non-addictive-enriched targets.

From the previous analysis for Tier 2 drugs, we identified 120 regions that have both addictive and non-addictive protein targets. We mapped these targets to the pathways and found 1,864 pathways for the Tier 2 protein target set. When the same analysis was done for 10 addictive and 32 non-addictive specific regions, we found 1,256 pathways in the addiction-specific areas and 1,664 pathways in the non-addictive specific regions. The specific region pathways represent the set of pathways associated with targets expressed in class-specific regions. Target selection for pathway annotation was restricted to proteins with an expression level of pTPM >= 1 (proteins per million) within the corresponding regions, a criterion distinct from the whole-brain expression filter applied in earlier analyses. The pathways for the overlapping regions are provided in Supplementary File S8.

Among the 1,864 pathways identified in the overlapping regions, 937 pathways were addiction-leaning (mean composition score > 0, fraction of drugs with positive composition score > 0.5, FDR < 0.05) and 7 were non-addictive-leaning (mean composition score < 0, fraction of drugs with positive composition score < 0.5, FDR < 0.05). The pathway composition is shown in Figure 4.

**Figure 4.**
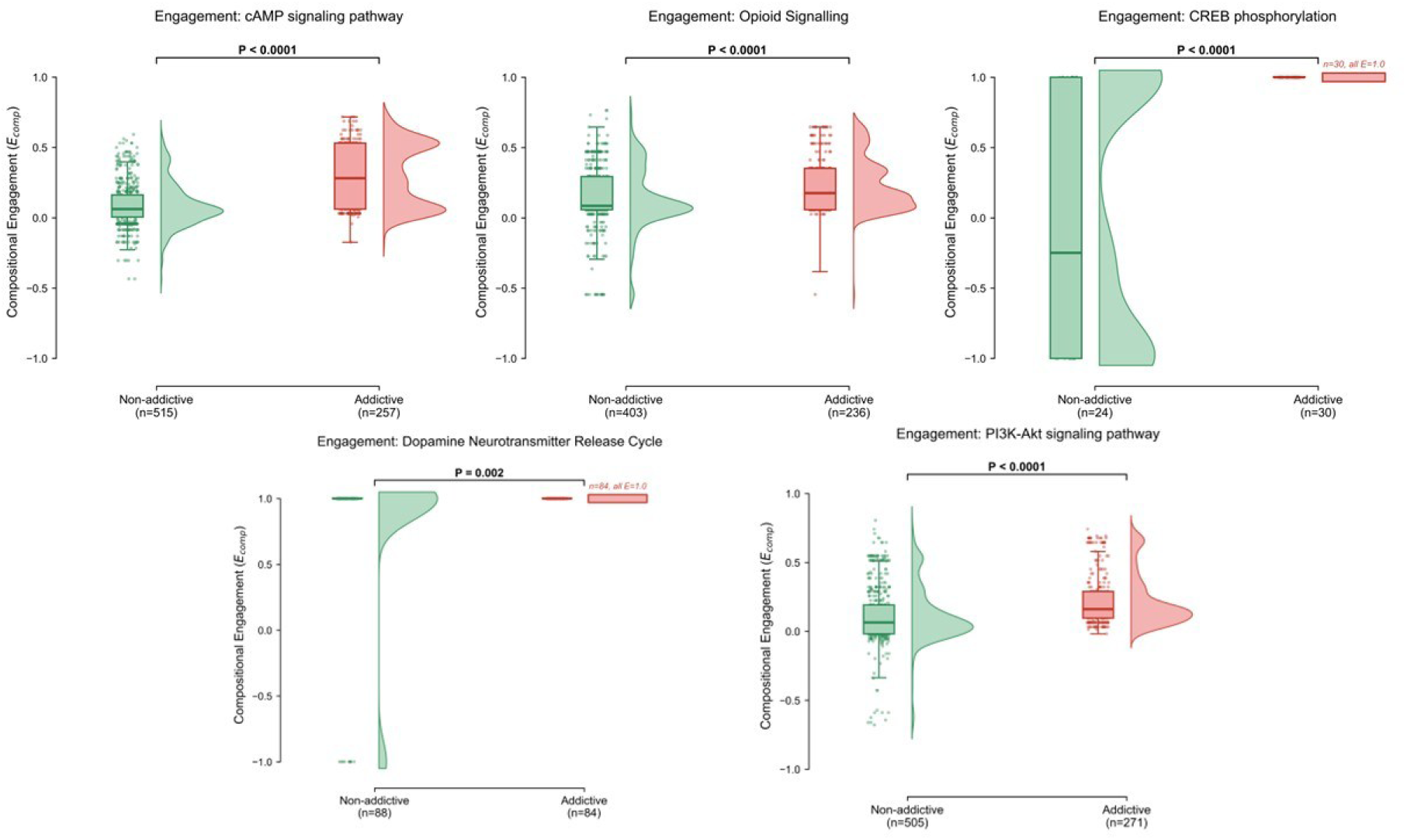
Differential pathway engagement for addictive and non-addictive drug targets as assessed by their Compositional Engagement score. Each panel displays a raincloud plot combining a half-violin (density distribution), box plot (median and IQR) and individual data points. The y-axis represents the compositional engagement score (E_comp) in all panels, quantifying the normalized pathway engagement per drug. (A) cAMP signaling pathway (n_add = 257, n_non = 515; P < 0.0001, Mann–Whitney U test). (B) Opioid signaling pathway (n_add = 236, n_non = 403; P < 0.0001, Mann–Whitney U test). (C) CREB phosphorylation pathway (n_add = 30, n_non = 24; P < 0.0001, Mann–Whitney U test). All 30 addictive drugs show maximal engagement (E_comp = 1.0), indicating complete and uniform engagement of this pathway. (D) Dopamine neurotransmitter release cycle (n_add = 84, n_non = 88; P = 0.002, Mann–Whitney U test). This pathway contains three proteins, one addictive and two non-addictive; addictive drugs interact exclusively with the addictive target, while non-addictive drugs interact with both target types. (E) PI3K-Akt signaling pathway (n_add = 271, n_non = 505; P < 0.0001, Mann–Whitney U test). Across all pathways, addictive drugs consistently show significantly higher engagement compared to non-addictive drugs.

As an illustrative example, we examined the addiction-dominant cAMP signaling (Figure 4A) pathway, which is a canonical addiction pathway. When we looked at the target-level information for this pathway, we found classic addiction-related proteins, such as BDNF, CREB1, GRIN2/3, and multiple GPCRs (NPY1R, OXTR, SSTRs, etc.). BDNF showed a high probability of predicted interactions with potent psychostimulants and synthetic opioids. Specifically, methamphetamine, mephedrone, and fenproporex, alongside some fentanyl analogs, were found. CREB1 was seen to be heavily targeted by heroin, morphine, oxycodone, and hydromorphone. It was also seen to be targeted by some non-addictive drugs, primarily opioid antagonists or partial agonists such as naloxone and naltrexone.

In contrast, non-addictive drugs engaging the cAMP pathway preferentially interact with the downstream signal termination and execution machinery of that pathway. PDE4A, a phosphodiesterase responsible for cAMP degradation and signal attenuation, was predicted to interact with several non-addictive drugs that would modulate cAMP tone without directly engaging reward circuitry. Similarly, downstream effector kinases, including PKA subunits, AKT, and MAPK family members, which execute rather than initiate cAMP signaling, were disproportionately represented among non-addictive target profiles. This suggests that non-addictive drugs engage cAMP signaling at a homeostatic, regulatory downstream level rather than at the receptor-transcription factor interface where addiction-associated perturbations originate. Thus, they act to attenuate the addiction response.

This divergence also illustrates a key principle of our framework: shared target engagement does not imply shared addiction liability. DRD1, canonically associated with dopaminergic reward signaling, is engaged by both addictive and non-addictive compounds in our dataset, but the nature of engagement differs fundamentally. Addictive agents engaging DRD1, including LSD, ibogaine, and related hallucinogens and stimulants, do so in the context of broad, high-affinity psychoactive target profiles that converge on reward circuitry. The 133 non-addictive compounds predicted to engage DRD1 are dominated by cardiovascular agents such as atenolol, metoprolol, and verapamil. When we investigated how these drugs affect addiction, we came across some surprising results. Atenolol was found to suppress certain aspects of addiction. It was shown to reduce alcohol withdrawal severity, attenuate alcohol craving, and lower autonomic arousal during withdrawal (29). Metoprolol, sharing the same β1-selective mechanism, has similarly been evaluated as an adjuvant in alcohol withdrawal management (30). Verapamil has pre-clinical evidence that has shown reduced morphine tolerance and dependence, and lower alcohol consumption, with some evidence suggesting disruption of drug-induced neuroplasticity (31).

Taken together, these findings reveal that pathway co-engagement masks a deeper asymmetry: addictive drugs concentrate interactions at nodes implicated in addiction-associated neuroplasticity. Non-addictive drugs distribute across the signal-routing and termination machinery that modulates an addiction response, while also showing interaction with addiction nodes.

### Network Level Organization of Addictive and Non-addictive Targets

To determine whether addictive or non-addictive targets form distinct or shared functional modules, we constructed a protein-protein interaction network comprising both target sets from the Tier 2 list, which helped determine whether these targets are integrated into shared networks or segregated into isolated modules. This network was constructed using STRINGdb on Cytoscape with high confidence (>= 0.7) (32, 33). STRINGdb was selected because it provides proteome-wide, multi-evidence, confidence-scored data and captures functional associations, which were required to test the functional module hypothesis.

To make the network more robust, we constructed the entire protein-protein interaction (PPI) network (red nodes represent addictive targets, green nodes represent non-addictive targets) comprising 973 proteins and 2,068 interactions. Cross-class interactions represent edges between only addictive and non-addictive nodes. The goal is to capture the functional coupling between the two classes, which suggests potential functional relations. The resulting network shows a clustering coefficient of zero, indicating that no closed triangles form exclusively between same-class nodes, consistent with the interdigitated rather than segregated organization of addictive and non-addictive targets. Despite its sparse density (0.004), the network formed a single connected component, indicating interconnectedness through direct protein-protein interactions rather than segregation into subnetworks, even in the cross-class network. Figure 5 shows the integrated network between the addictive and non-addictive targets.

**Figure 5.**
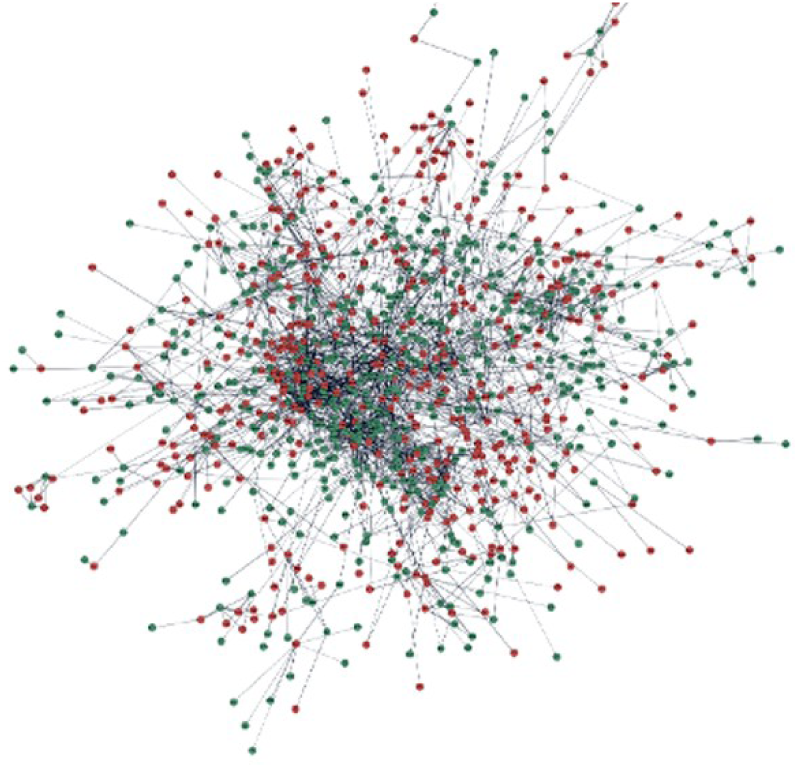
An interconnected protein-protein graph of addictive (red) and non-addictive (green) nodes representing functional coupling between the targets. The interactions between the nodes are STRING high confidence (>= 0.70) between Tier 2 addictive enriched and non-addictive enriched targets (32). The resulting graph forms a single connected component with low density and short average path length, indicating widespread intermixing. The absence of class-specific clustering suggests that addictive and non-addictive targets participate within shared molecular networks.

The average number of neighbors for this network is 4.25, and the characteristic path length is 5.708. The network diameter is 16, which indicates that connections between addictive and non-addictive targets span multiple interaction steps. Cross-class connectivity shows a broad, distributed interaction structure. Thus, addiction and non-addictive protein targets are highly interdigitated. To dig deeper into this mechanism, we analyzed how the addictive and non-addictive drug classes, on average, interact with the proteins of interest.

### First-order and second-order interactions of drugs

Many of the addictive drugs, on average, don’t bind non-addictive protein targets even at individual levels. In contrast, non-addictive drugs interact with both addictive and non-addictive targets simultaneously. About 978 out of 1,448 blood-brain barrier-permeable, non-addictive drugs were found to interact with both addictive and non-addictive targets. To quantify this interaction pattern, we took a ratio of addictive/non-addictive targets for both drug classes. Addictive drugs showed a mean ratio of 92 with a median of 31. Non-addictive drugs showed a mean of 6.4 with a median of 0.125. Figure 6 shows the distribution for the addictive and non-addictive target interactions of the drugs.

**Figure 6.**
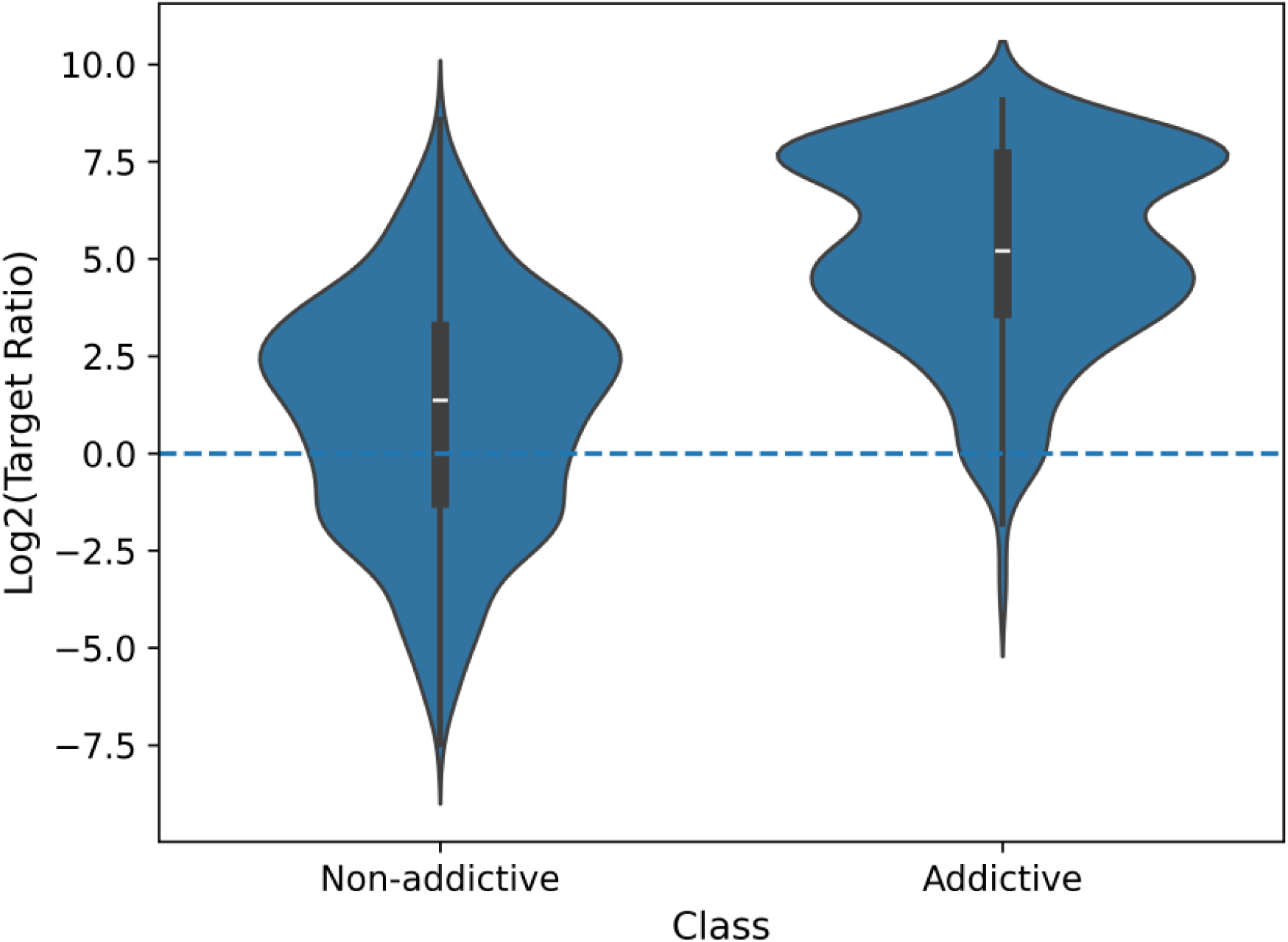
Violin plot comparing the distribution of log2-transformed target ratios (addictive/non-addictive) between non-addictive and addictive drugs. The dashed horizontal line indicates equal representation of targets. Distribution above the line indicates addictive target interaction, and distribution below the line indicates non-addictive target interaction.

This disparity reflects a fundamental asymmetry: non-addictive drugs engage both addictive and non-addictive targets concurrently, whereas addictive drugs interact predominantly with addiction-enriched targets, but what about the protein-protein interactions of these first-order targets?

To assess how these drugs perturb the PPI network, we examined their second-order interactions. The PPI network was constructed from the enriched addictive and non-addictive targets (Tier 2). This helped to identify the second-order perturbation pattern for the drugs. Critically, the second-order interactions of non-addictive enriched protein targets of addictive drugs show substantial overlap with first-order non-addictive enriched protein targets of non-addictive drugs. To characterize the relationship between addictive and non-addictive drug target spaces, we compared first- and second-order interaction neighborhoods across drug classes. Specifically, the non-addictive second-order targets of addictive drugs overlap 55% with the first-order non-addictive targets of non-addictive drugs. Second-order non-addictive targets of addictive drugs also show a 79% overlap with the second-order non-addictive targets of non-addictive drugs. Both drug classes engage the same molecular circuits but differ in the interaction order at which key nodes are reached. The overlap of non-addictive enriched protein targets was 726, comprising a diverse set of proteins that can be mapped to homeostatic, regulatory, and dampening states. A representative protein of interest is CDK5, a synaptic/neuro dampening protein known to act as a homeostatic brake, opposing synaptic potentiation associated with addiction-related plasticity (34). The list was also enriched for potassium, sodium, and calcium channel genes, including KCNQ1-5, KCNJ2-16, SCN1B-9A, and CACNA1S. Many of these channels regulate neuronal excitability and contribute to electrophysiological stability, which can influence reward circuit homeostasis relevant to addiction. Thus, the list of proteins potentially provides homeostatic, regulatory, and dampening nodes in the addictive and non-addictive interdigitated network. Combined with the PPI network, these results show that addictive-enriched and non-addictive-enriched proteins physically interact with each other, suggesting they are positioned to counter one another’s signaling. The 726 proteins of interest are listed in Supplementary File S9.

The above results suggest the following comprehensive mechanistic model. Non-addictive drugs coactivate the nearest neighbor regulatory targets, which results in multiscale damping across corticostriatal reinforcement circuits. Addictive drugs lack this additive suppression response. Addictive drugs classically enhance dopaminergic signaling from the ventral tegmental area (VTA) to the nucleus accumbens (NAc), potentiating corticostriatal synapses and driving long-term plasticity within prefrontal–striatal loops. When a compound selectively amplifies dopaminergic tone, particularly through D1 receptor–biased signaling, it promotes sustained cAMP/PKA activation, CREB phosphorylation, ΔFosB accumulation, and structural synaptic remodeling in medium spiny neurons. These molecular events consolidate reinforcement learning into persistent neuroadaptation. However, when the same compound simultaneously engages non-addictive targets enriched in regulatory, transport, or metabolic functions, these pathways may recruit inhibitory or homeostatic mechanisms that counterbalance pro-addictive signaling. Such engagement can enhance GABAergic tone, stabilize glutamate homeostasis, modulate dopamine reuptake kinetics, buffer intracellular calcium flux, or constrain AMPA receptor trafficking. The net effect is attenuation of synaptic potentiation within the NAc and prevention of reinforcement-driven plasticity from consolidating into a sustained addictive state.

From a systems perspective, addiction can be viewed as a transition of the reward network into a high-gain, self-reinforcing dynamical regime. Selective activation of pro-addictive targets increases network controllability in a direction that favors positive feedback amplification and synchronization of dopaminergic and glutamatergic signaling. In contrast, concurrent activation of regulatory and metabolic targets introduces negative feedback loops, signal dispersion, and energetic constraints, effectively reducing network gain and increasing damping. These mechanisms may prevent the circuit from crossing the critical threshold required to enter a stable addictive attractor state. In this framework, addiction emerges not merely from activation of reward circuitry, but from insufficient engagement of intrinsic regulatory constraints. Compounds that co-activate buffering and homeostatic pathways may therefore remain non-addictive because they impose early counter-regulatory control that limits amplification, block long-term plasticity, and prevent the coordinated physiological adaptations necessary for addiction.

## Discussion

This study represents a system-level framework for different engagement patterns of addictive and non-addictive drugs that integrates predicted protein-ligand interactions, brain-region-specific expression, pathway engagement, and protein-protein interaction topology. Our results reveal that addictive and non-addictive compounds largely converge on the same pathways and neural regions and yet differ in how they distribute their interactions within shared molecular programs. Rather than operating through distinct anatomical or pathway modules, differences in addiction emerge from differential engagement patterns embedded within the common biological circuits. As conjectured above, the pathways are highly interdigitated. When combined with Figure 5, this suggests that non-addictive drugs modulate or dampen addiction-related signaling within shared circuits. In addition to interacting with a protein that causes addiction, they interact with a nearest neighbor partner protein in their interaction complex that suppresses the addiction.

Our findings reveal that addictive and non-addictive drugs engage targets within a shared neural architecture spanning 120 brain regions and 1,864 pathways. Despite this circuit-level convergence, the two drug classes exhibit fundamentally different patterns of pathway-level engagement. Remarkably, even within shared pathways, addictive and non-addictive drugs distribute their molecular interactions asymmetrically; addictive drugs preferentially engage plasticity-associated protein targets, whereas non-addictive drugs, while also interacting with plasticity associated targets, preferentially engage regulatory and homeostatic targets within the same pathway structures. Rather than reflecting differences in pathway identity, these results indicate differences in functional engagement topology. Addictive drugs exhibit functional convergence, targeting a subset of signaling nodes linked to reinforcement and synaptic plasticity. In contrast, non-addictive drugs display distributed interactions, spreading interactions across diverse targets. We propose that addiction liability depends not merely on which pathways a drug interacts with, but on whether it preferentially targets proteins for amplification/plasticity or targets proteins that attenuate addiction response in that specific pathway.

This pattern supports a model based on the differential engagement of molecular targets within shared systems. The finding that the addictive and non-addictive drugs have different addictive/non-addictive target ratios further consolidates the model. The addictive signal is amplified before reaching regulatory checkpoints, as addictive drugs predominantly engage plasticity nodes. Non-addictive drugs, by contrast, interact with both target classes simultaneously, providing a buffering or regulatory signal that attenuates this amplification.

These findings have direct translational implications for developing safer psychiatric and analgesic medications. The presented mechanistic model in the results can also determine why some drugs are addictive and some are non-addictive. In subsequent work, we will develop a predictive addiction liability model grounded in the target engagement patterns identified here.

Although our findings are robust, this study relies on predicted protein-ligand interactions and computational network inference. The framework does not incorporate dose-response relationships, binding kinetics, or temporal dynamics of drug action, all of which may influence addiction liability in ways not captured by static interaction profiles. Experimental validation through biochemical binding assays, circuit-level perturbations, and behavioral addiction models will be required to establish the causal relationships implied by these computational findings.

## Methods

### Drug dataset assembly

Addictive compounds were obtained from the Drug Enforcement Administration (DEA) controlled substance list (13). Non-addictive compounds were derived from Phase IV-approved drugs in ChEMBL (14). Drug structures were represented using SMILES strings. Having assembled chemically diverse drug sets, we next predicted their molecular targets across the human proteome.

### Chemical Standardization and Deduplication

All compounds were standardized using RDKit, salts were removed, and the largest fragment was retained to define the parent structures. The InChIKey was used to identify the identical structures that had a conflict between the labels for the addictive and non-addictive classes. For those structures that had such a conflict, addictive ones were kept, and non-addictive ones were removed. Canonical SMILES were also generated for further downstream analysis.

### Structural Clustering

Chemical similarity was quantified using Morgan Fingerprints (radius = 2, 2048 bits). Compounds were clustered using the Butina algorithm at a Tanimoto similarity threshold of 0.95 and 0.80 (35). One representative compound per cluster was taken and retained. The 0.80 Tanimoto similarity threshold set was used for downstream analysis. To restrict the analysis to compounds with potential brain exposure, we applied Blood Brain Barrier filtering on these compounds.

### Proteome-wide target expression

Drug-target interactions were predicted using FINDSITE^comb2.0^ across the human proteome (∼19k proteins) (1). Protein interactions with predicted binding probability >= 0.70 were retained for downstream analysis. To ensure chemical diversity and eliminate redundancy in the drug sets, we performed systematic standardization and structural clustering.

### Blood brain barrier filtering

Brain exposure potential was estimated using the RDKit-derived physicochemical descriptors and heuristic Blood Brain Barrier (BBB) permeability (36, 37). The descriptors used are molecular weight (MW), topological polar surface area (TPSA), hydrogen bond donors (HBD), rotatable bond (RotB), and calculated logP (cLogP). A BBB-likeness score (0–5) was assigned by giving one point for each of the following criteria: MW <= 480, TPSA <= 90, HBD <= 2, RotB <=10, and 0 <= cLogP <=5. Compounds with a score >= 3 were taken as BBB-permeable (38–40). Only compounds predicted to be BBB permeable were retained for further analysis. We then filtered the targets present in the brain for minimal expression. This ensured that the predicted protein targets have at least one region where they are expressed.

### Brain region expression filtering

Protein targets in the brain were filtered for brain relevance using transcriptomic data from the Human Protein Atlas (2). Proteins with expression >= 1 pTPM in at least one brain region were retained. To identify targets preferentially targeted by each drug class, we calculated enrichment factors, as described below.

### Target Enrichment Analysis

For each protein, the enrichment factor was calculated with the formula:

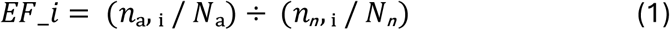

where, 𝐸𝐹 _𝑖 is the Enrichment Factor for target i, 𝑛ₐ, ᵢ is the number of addictive compounds predicted to bind protein target i, 𝑁ₐ is the total number of addictive compounds, 𝑛*ₙ*, ᵢ is the number of non-addictive compounds predicted to bind to protein target i, and 𝑁*ₙ* is the total number of non-addictive compounds. Targets with EF > 2 were considered preferentially bound by addictive drugs, while those with EF < 0.5 were preferentially bound by non-addictive drugs. Statistical significance was assessed at the target level using Fisher’s exact test with the Benjamini-Hochberg correction (FDR < 0.05).

### Target Tiers

Two complementary target sets were defined:

Tier 1 (high confidence): EF thresholds (Addictive EF > 2; Non-addictive EF < 0.5) and FDR <= 0.05

Tier 2 (high consensus): EF thresholds met with >= 10 supporting compounds, independent of FDR. To contextualize these computationally derived targets with existing biology, we performed literature-based validation.

### Literature-Based Target Validation

Predicted Tier 2 targets were taken for literature-based validation. A representative set of 100 targets from each class (addictive and non-addictive) was evaluated using an automated pipeline. Claims were formulated as “[gene] is implicated in addiction” for addictive targets and “[gene] has been associated with regulatory or homeostatic biological processes in neural systems” for non-addictive targets. Each claim was evaluated independently using GPT-5.4 with medium reasoning effort and web search enabled (search context size: high), querying peer-reviewed literature, systematic reviews, and authoritative biological databases.

Each claim was assigned a verdict from a fixed set: Highly Supported, Likely True, Mixed Evidence, Likely False, Contradicted, or No Evidence, along with a confidence label (Low, Medium, or High). Claims were evaluated independently with no crossclaim context. The resulting verdicts and confidence scores were aggregated by target class to assess overall validation rates.

### Brain region Prioritization

For each protein, the top five brain regions with the highest expression levels were selected based on pTPM values. Region-level specific and overlap analysis was performed. To resolve functional differences of these targets, we mapped them to their respective biological pathways.

### Pathway Annotation

Pathway annotation was performed using the Comparative Toxicogenomic Database for both addiction and non-addiction target drug sets (12). The pathway membership for overlapping regions was restricted to proteins expressed in those regions. For class-specific regions, pathway membership was restricted to expressed targets in those regions. To quantify how each drug class distributes its interactions within shared pathways, we developed normalized engagement metrics.

### Pathway Engagement Metrics

The number of addictive and non-addictive drugs differs substantially, and pathways often contain both target types; simple target counts fail to capture true engagement biases. To quantify the normalized difference between the interactions with addiction-enriched and non-addiction-enriched targets, we defined three pathway engagement metrics.

First to capture the balance of interactions for a specific drug within a given pathway, the per-drug engagement score is defined as:

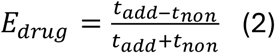

t_add and t_non are the number of addictive enriched and non-addictive enriched targets for the pathways that are engaged by the drug. E_drug_ ranges from -1 (non-addictive biased) to + 1 (addictive biased).

Second, to control potential biases arising from unequal target availability within pathways, we computed the Compositional engagement score, which normalizes for the total target capacity of the pathway:

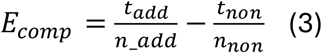

n_add and n_non are the total number of addictive enriched and non-addictive enriched targets present in the pathway. E_comp_ ranges from -1 (non-addictive biased) to + 1 (addictive biased).

Third, to measure the consistency of this engagement bias across individual drugs, we defined the Addiction tilt. While the mean engagement scores capture the average magnitude, the tilt quantifies the uniformity with which drugs interacting with a pathway exhibit an addictive bias. It is calculated as the fraction of drugs that yield a positive compositional engagement score:

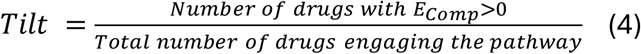

The tilt determines the fraction of drugs that engage more addictive enriched targets irrespective of the class.

Finally, to assess whether target classes form distinct or integrated functional modules, we constructed protein-protein interaction networks.

### Protein-Protein Interaction Network Analysis

Protein interaction networks were constructed using STRINGdb (confidence >= 0.70) in Cytoscape (32, 33). Network properties were calculated using the network analyzer in Cytoscape. To assess how each drug class perturbs the PPI network, first- and second-order interaction neighborhoods were computed for each drug by combining the STRING edge table with drug-target interaction data. For each drug, direct (first order) targets were classified by computing the ratio of addiction-enriched to non-addiction enriched hits among classified targets. Second order neighbors were defined as proteins reachable via one PPI hop from the direct target set, excluding the direct targets themselves. Each protein was assigned a label, addictive-dominant, non-addictive-dominant, or mixed, based on whether the ratio of addiction-enriched to non-addictive-enriched interactions met or exceeded a 0.7 threshold. Drug trajectories were then defined as the combination of first- and second-order neighborhood labels. Statistical enrichment of trajectories by drug class was assessed using Fisher’s exact test with the Benjamini-Hochberg correction (FDR < 0.05). To identify candidate suppression nodes, the second-order non-addictive neighborhood of addictive drugs was intersected with the first-order non-addictive targets of non-addictive drugs. This yielded 726 proteins reachable directly by non-addictive drugs but only indirectly by addictive drugs, representing candidate nodes at which non-addictive compounds may impose regulatory constraints on shared addiction circuitry.

## Supporting information

Folder for supplementary files

## Code and Data Availability

All code used for data processing, target enrichment analysis, pathway engagement scoring, and PPI network perturbation analysis is available at https://github.com/Harry-gatech/addiction_brain. The drug-target interaction dataset and supplementary files are provided with this manuscript.

## Acknowledgments

This work was supported in part by grant GM118039 of the Division of General Medical Sciences of NIH. A gift from the Ovarian Cancer Institute is also gratefully acknowledged. We thank Jessica Forness for helping prepare the manuscript and Bartosz Ilkowski for technical guidance and support.

## Author Contributions

HS: Designed the computational framework, performed the analysis, interpreted the results, and drafted the initial manuscript. HZ: Ran FINDSITE binding probability predictions, assisted with statistical modeling, and contributed to result interpretation. SS: Initialized and conceptualized the project and provided key interpretation of the results. JS: Led the overall direction of the project, identified which features were likely to drive addictive and non-addictive behavior, suggested which drug classes should be analyzed, provided critical guidance throughout the project, led the interpretation of results, and critically revised and edited the manuscript.

## Competing Interest Statement

The authors do not have any competing interests.

## Notes

https://doi.org/10.5281/zenodo.19698192

### Competing Interest Statement

The authors have declared no competing interest.

