## Supplementary material for "Understanding Substance Dependence: What Differentiates Addictive from Non-Addictive Drugs?": Folder for supplementary files: addictive-genes-claims-first100-001-avp-gene-protein-is-implicated-in-addiction-78c005df6210.md

# addictive-genes-claims-first100-001-78c005df6210

**Original claim:** AVP gene/protein is implicated in addiction.

## Claim

AVP (arginine vasopressin) gene/protein is implicated in addiction. ([pubmed.ncbi.nlm.nih.gov](https://pubmed.ncbi.nlm.nih.gov/21677651/)) [S1]

## Verdict

Verdict: Likely True

## Confidence

Medium. The case is strongest in **preclinical addiction models** and in **stress/relapse-related biology**; human evidence exists but is still limited, indirect, and more convincing for the broader **vasopressin signaling system** (especially AVPR1A/AVPR1B receptors) than for the **AVP gene itself** as a major causal addiction gene. ([pubmed.ncbi.nlm.nih.gov](https://pubmed.ncbi.nlm.nih.gov/21575018/)) [S2]

## Supporting evidence

Multiple rodent studies tie AVP biology to addiction-relevant behaviors. In a cocaine conditioning model, **AVP gene expression changed in the nucleus accumbens during all stages of cocaine-conditioned responding**, AVP protein in accumbal tissue also changed, and **local V1A receptor blockade in the nucleus accumbens interrupted acquisition/expression of the conditioned response**, while agonist stimulation enhanced it. That is direct evidence that AVP signaling can modulate cocaine-associated reward learning. ([pubmed.ncbi.nlm.nih.gov](https://pubmed.ncbi.nlm.nih.gov/19596360/)) [S3]

In cocaine withdrawal models, AVP also tracks stress-related adaptations that are highly relevant to addiction. After chronic escalating-dose “binge” cocaine, rats showed **persistent elevation of PVN AVP mRNA during withdrawal**, alongside increased pituitary **V1b** and **POMC** mRNA and elevated ACTH/corticosterone; a **selective V1b antagonist (SSR149415)** attenuated acute withdrawal-induced HPA activation. A separate study found **increased amygdalar AVP mRNA during acute cocaine withdrawal**, suggesting a role in the aversive, stress-related state of withdrawal. ([pubmed.ncbi.nlm.nih.gov](https://pubmed.ncbi.nlm.nih.gov/21677651/)) [S4]

Evidence also extends to opioids. In rats with heroin self-administration, animals that later showed **high stress-induced reinstatement** had **higher AVP mRNA in medial/basolateral amygdala**, and AVP mRNA levels correlated with stress-induced heroin-seeking. That links AVP not just to drug exposure, but to a core addiction phenotype: vulnerability to relapse under stress. ([pubmed.ncbi.nlm.nih.gov](https://pubmed.ncbi.nlm.nih.gov/25446223/)) [S5]

For alcohol, the preclinical signal is substantial. In selectively bred Sardinian alcohol-preferring rats, **basal AVP mRNA was higher** in medial/central amygdala and medial hypothalamus than in non-preferring rats, and an **AVP V1b antagonist reduced alcohol intake**. More recently, intranasal AVP in socially housed mice **reduced sweetened ethanol intake**, an effect blocked by a **V1a antagonist**. Taken together, these findings indicate that AVP signaling can modulate alcohol drinking, though the direction depends on dose, receptor subtype, and model. ([pubmed.ncbi.nlm.nih.gov](https://pubmed.ncbi.nlm.nih.gov/21575018/)) [S6]

There is also some human evidence, though it is weaker and more indirect. A case-control study of 111 patients in alcohol detoxification found **higher promoter methylation of the vasopressin gene** in alcohol-dependent patients than in controls, suggesting altered AVP regulation in human alcohol dependence, even though peripheral vasopressin mRNA itself was not significantly different. ([pubmed.ncbi.nlm.nih.gov](https://pubmed.ncbi.nlm.nih.gov/19046820/)) [S7]

Human genetics more strongly implicates the **vasopressin receptor system** than the AVP gene itself. In a candidate-gene study, SNPs in **AVPR1A** were associated with drug use disorder, with partial replication in independent samples and evidence that genotype influenced postmortem brain AVPR1A expression. That does not prove the AVP peptide gene causes addiction, but it supports involvement of the vasopressin pathway in addiction liability. ([pubmed.ncbi.nlm.nih.gov](https://pubmed.ncbi.nlm.nih.gov/21514569/)) [S8]

Human pharmacology is suggestive but not definitive. In a 150-participant phase 2 trial for alcohol dependence, the **V1b antagonist ABT-436** did **not** significantly improve the primary endpoint (heavy drinking days), but it **did** increase percent days abstinent and seemed more promising in participants with higher baseline stress. A small human laboratory study in cocaine use disorder also found a **distinct ACTH response pattern** to intranasal desmopressin/oxytocin challenge versus controls, consistent with altered vasopressin-related stress-axis regulation in addicted patients. ([pubmed.ncbi.nlm.nih.gov](https://pubmed.ncbi.nlm.nih.gov/27658483/)) [S9]

## Contradicting or limiting evidence

The evidence is not uniformly positive, and several findings argue against overstatement. In mice with targeted deletion of **Avpr1a** or **Avpr1b**, one study reported that **acute intoxicating effects of ethanol** and **voluntary ethanol consumption** were not altered by loss of either receptor. That means AVP signaling is probably **not a universal driver of all alcohol-related behaviors**. ([pubmed.ncbi.nlm.nih.gov](https://pubmed.ncbi.nlm.nih.gov/17049983/?utm_source=openai)) [S10]

Likewise, in vasopressin-deficient Brattleboro rats, AVP deficiency **did not change ethanol-withdrawal convulsions**, suggesting AVP is not required for every withdrawal phenotype. Older work on alcohol dependence/withdrawal therefore points to some important negative results. ([pubmed.ncbi.nlm.nih.gov](https://pubmed.ncbi.nlm.nih.gov/3834412/)) [S11]

The human literature remains limited. The alcohol-dependence methylation study used **peripheral blood**, not brain tissue, so it cannot establish central nervous system causality. The AVPR1A association paper involved a receptor gene, not the AVP peptide gene, and replication was not directionally uniform across all samples. The ABT-436 trial showed **mixed efficacy**, with a null primary outcome. Overall, the best-supported claim is that **vasopressin signaling is involved in addiction-related processes**, especially stress and relapse, rather than that AVP is a fully established addiction gene/protein in humans. ([pubmed.ncbi.nlm.nih.gov](https://pubmed.ncbi.nlm.nih.gov/19046820/)) [S12]

## Mechanistic / biological rationale

A biologically plausible mechanism exists. AVP signaling through the **V1b receptor** is a key regulator of the **hypothalamic-pituitary-adrenal (HPA) axis**: V1b-receptor knockout mice have lower basal ACTH/corticosterone and a blunted ACTH response to forced-swim stress. That makes AVP well positioned to influence the stress systems that are known to drive withdrawal distress and relapse vulnerability. ([pubmed.ncbi.nlm.nih.gov](https://pubmed.ncbi.nlm.nih.gov/14722621/?utm_source=openai)) [S13]

Addiction models fit that mechanism. During cocaine withdrawal, **PVN AVP expression rises**, pituitary **V1b** signaling is upregulated, and stress hormones stay elevated; blocking V1b dampens this response. In heroin models, higher amygdalar AVP marks animals more vulnerable to **stress-induced reinstatement**. These findings are consistent with AVP acting as a stress-amplifying signal that can promote negative reinforcement and relapse-like behavior. ([pubmed.ncbi.nlm.nih.gov](https://pubmed.ncbi.nlm.nih.gov/21677651/)) [S14]

AVP may also affect **reward learning and cue conditioning** outside the HPA axis. In the nucleus accumbens, AVP expression changes during cocaine conditioning, and local V1A manipulation changes conditioned responding. In alcohol-preferring rats, elevated AVP expression in amygdala/hypothalamus and sensitivity to V1b antagonism suggest AVP can contribute to high-drinking phenotypes. So the most coherent model is that AVP influences addiction via **stress integration, emotional valence, and conditioned cue/reinforcement processes**, rather than acting as a primary reward transmitter on its own. ([pubmed.ncbi.nlm.nih.gov](https://pubmed.ncbi.nlm.nih.gov/19596360/)) [S15]

## Bottom line

Yes—**the claim is likely true**, with an important qualifier: AVP is best supported as part of the **vasopressin stress-signaling network** that modulates **withdrawal, stress reactivity, relapse-like behavior, and some forms of alcohol/cocaine-conditioned responding**, mainly in animal models. Human evidence is **suggestive but not yet strong enough** to say AVP itself is a firmly established addiction gene/protein in clinical populations. A careful wording would be: **“AVP/vasopressin signaling is implicated in addiction-related neurobiology, especially stress-linked relapse and alcohol/cocaine models.”** ([pubmed.ncbi.nlm.nih.gov](https://pubmed.ncbi.nlm.nih.gov/21575018/)) [S16]

## Cited sources

- **[S1]** Persistent increase in hypothalamic arginine vasopressin gene expression during protracted withdrawal from chronic escalating-dose cocaine in rodents - PubMed — https://pubmed.ncbi.nlm.nih.gov/21677651/
- **[S2]** Involvement of arginine vasopressin and V1b receptor in alcohol drinking in Sardinian alcohol-preferring rats - PubMed — https://pubmed.ncbi.nlm.nih.gov/21575018/
- **[S3]** Arginine vasopressin gene expression changes within the nucleus accumbens during environment elicited cocaine-conditioned response in rats - PubMed — https://pubmed.ncbi.nlm.nih.gov/19596360/
- **[S4]** Persistent increase in hypothalamic arginine vasopressin gene expression during protracted withdrawal from chronic escalating-dose cocaine in rodents - PubMed — https://pubmed.ncbi.nlm.nih.gov/21677651/
- **[S5]** Individual differences in gene expression of vasopressin, D2 receptor, POMC and orexin: vulnerability to relapse to heroin-seeking in rats - PubMed — https://pubmed.ncbi.nlm.nih.gov/25446223/
- **[S6]** Involvement of arginine vasopressin and V1b receptor in alcohol drinking in Sardinian alcohol-preferring rats - PubMed — https://pubmed.ncbi.nlm.nih.gov/21575018/
- **[S7]** Epigenetic regulation and gene expression of vasopressin and atrial natriuretic peptide in alcohol withdrawal - PubMed — https://pubmed.ncbi.nlm.nih.gov/19046820/
- **[S8]** The AVPR1A gene and substance use disorders: association, replication, and functional evidence - PubMed — https://pubmed.ncbi.nlm.nih.gov/21514569/
- **[S9]** A Phase 2, Double-Blind, Placebo-Controlled Randomized Trial Assessing the Efficacy of ABT-436, a Novel V1b Receptor Antagonist, for Alcohol Dependence - PubMed — https://pubmed.ncbi.nlm.nih.gov/27658483/
- **[S10]** The acute intoxicating effects of ethanol are not dependent on the vasopressin 1a or 1b receptors - PubMed — https://pubmed.ncbi.nlm.nih.gov/17049983/?utm_source=openai
- **[S11]** The role of arginine vasopressin in alcohol dependence and withdrawal - PubMed — https://pubmed.ncbi.nlm.nih.gov/3834412/
- **[S12]** Epigenetic regulation and gene expression of vasopressin and atrial natriuretic peptide in alcohol withdrawal - PubMed — https://pubmed.ncbi.nlm.nih.gov/19046820/
- **[S13]** The vasopressin V1b receptor critically regulates hypothalamic-pituitary-adrenal axis activity under both stress and resting conditions - PubMed — https://pubmed.ncbi.nlm.nih.gov/14722621/?utm_source=openai
- **[S14]** Persistent increase in hypothalamic arginine vasopressin gene expression during protracted withdrawal from chronic escalating-dose cocaine in rodents - PubMed — https://pubmed.ncbi.nlm.nih.gov/21677651/
- **[S15]** Arginine vasopressin gene expression changes within the nucleus accumbens during environment elicited cocaine-conditioned response in rats - PubMed — https://pubmed.ncbi.nlm.nih.gov/19596360/
- **[S16]** Involvement of arginine vasopressin and V1b receptor in alcohol drinking in Sardinian alcohol-preferring rats - PubMed — https://pubmed.ncbi.nlm.nih.gov/21575018/
