## Supplementary material for "Understanding Substance Dependence: What Differentiates Addictive from Non-Addictive Drugs?": Folder for supplementary files: addictive-genes-claims-first100-002-actb-gene-protein-is-implicated-in-addiction-f037b7422ae8.md

# addictive-genes-claims-first100-002-f037b7422ae8

**Original claim:** ACTB gene/protein is implicated in addiction.

## Claim

ACTB gene/protein is implicated in addiction.

## Verdict

Verdict: Mixed Evidence

## Confidence

Medium

## Supporting evidence

ACTB encodes **β-actin**, a ubiquitous non-muscle actin isoform that forms part of the actin cytoskeleton and helps determine cell shape, motility, and intracellular signaling. That basic biology matters because addiction is widely understood as a disorder of maladaptive neuroplasticity, and actin dynamics are central to synaptic structural plasticity. A recent review on substance use disorders (SUDs) states that cytoskeletal dynamics are fundamental to the morphological and behavioral plasticity associated with drug exposure, and specifically highlights actin-based mechanisms across cocaine, methamphetamine, alcohol, opioids, and nicotine. ([medlineplus.gov](https://medlineplus.gov/genetics/gene/actb)) [S1]

For **cocaine**, there is solid preclinical evidence that actin remodeling is functionally involved in addiction-related behavior. In rats, repeated cocaine exposure and withdrawal increased filamentous actin (F-actin) in the nucleus accumbens, altered actin-binding proteins, and manipulating actin polymerization/depolymerization changed cocaine-induced reinstatement of drug seeking. That is not ACTB-specific genetics, but it does implicate β-actin-containing cytoskeletal machinery in a core addiction-relevant phenotype. ([pubmed.ncbi.nlm.nih.gov](https://pubmed.ncbi.nlm.nih.gov/16452681/)) [S2]

For **methamphetamine**, the evidence is somewhat closer to ACTB itself. A review summarizing the methamphetamine literature notes that, in dorsal striatal synaptosomes from rats after 30 days of methamphetamine self-administration and 14 days of abstinence, altered proteins included **actin cytoplasmic 1 (Actb)**. The same review also summarizes mouse studies showing that methamphetamine conditioning increased ADF, cofilin, G-actin, and F-actin, and that disrupting actin dynamics impaired methamphetamine conditioned place preference or methamphetamine seeking. These findings support involvement of actin/β-actin-related plasticity in methamphetamine-associated learning and motivation. ([pmc.ncbi.nlm.nih.gov](https://pmc.ncbi.nlm.nih.gov/articles/PMC11034636/)) [S3]

Additional methamphetamine work in nonhuman primates found hippocampal atrophy plus downregulation of cytoskeleton-related genes after acute or chronic exposure, with reduced profilin-2 expression as one validated change. Again, that is not direct proof about ACTB itself, but it supports a broader actin-cytoskeleton mechanism in stimulant-related neuroadaptation. ([pubmed.ncbi.nlm.nih.gov](https://pubmed.ncbi.nlm.nih.gov/29802913/)) [S4]

For **heroin/opioids**, a 2024 rat study found that inhibiting actin polymerization in the nucleus accumbens core during the reconsolidation window reduced cue-induced and heroin-primed reinstatement for at least 28 days. This supports a causal role for actin dynamics in drug-memory maintenance and relapse-like behavior. ([pubmed.ncbi.nlm.nih.gov](https://pubmed.ncbi.nlm.nih.gov/39716383/)) [S5]

For **alcohol**, preclinical studies summarized in the same cytoskeleton review report altered F-actin and actin regulators after excessive drinking, and manipulations of upstream actin regulators such as Prosapip1 and mTORC2 changed alcohol seeking, conditioned place preference, or binge-like consumption. So across multiple substances, actin-dependent structural plasticity is repeatedly linked to addiction-relevant behaviors. ([pmc.ncbi.nlm.nih.gov](https://pmc.ncbi.nlm.nih.gov/articles/PMC11034636/)) [S6]

## Contradicting or limiting evidence

The main limitation is **specificity**: most of the stronger evidence implicates the **actin cytoskeleton as a system**, not **ACTB as a reproducible addiction gene**. ACTB is a broadly expressed structural gene; official reference material describes it mainly as a ubiquitous cytoskeletal component involved in cell architecture, movement, and signaling, not as an addiction-focused gene. ([medlineplus.gov](https://medlineplus.gov/genetics/gene/actb)) [S7]

Large human genetic studies of SUDs have identified many loci and pathways, but ACTB does **not** emerge as a well-established recurrent risk gene. For example, a systematic integration of 75 GWAS/RNA-seq studies across substance use behaviors/disorders highlighted shared genes such as **CACNB2, GRIN2B, PLXDC2, and PKNOX2**, while a large multivariate study of 2.2 million individuals emphasized broad externalizing/SUD genetic architecture rather than ACTB specifically. Another shared-dependence GWAS identified **ANKS1B, AGBL4, and CTNNA2** as replicated loci for alcoholism, heroin dependence, and methamphetamine dependence. ([pubmed.ncbi.nlm.nih.gov](https://pubmed.ncbi.nlm.nih.gov/35491750/)) [S8]

There is also a cautionary literature issue: one later pathway paper listed **ACTB** as associated with amphetamine dependence, but the cited source was actually a **mouse study on actin depolymerizing factor (ADF/cofilin) in methamphetamine-induced place preference**, not a robust human ACTB association study. That means some apparent ACTB-addiction claims may reflect pathway-level actin biology rather than gene-specific evidence. ([pmc.ncbi.nlm.nih.gov](https://pmc.ncbi.nlm.nih.gov/articles/PMC4504822/)) [S9]

So, if the claim is interpreted narrowly as “**ACTB is a proven human addiction susceptibility gene/protein target**,” the evidence is weak. If interpreted more broadly as “**β-actin-related cytoskeletal dynamics participate in addiction-relevant neuroplasticity**,” the evidence is appreciably stronger. ([medlineplus.gov](https://medlineplus.gov/genetics/gene/actb)) [S10]

## Mechanistic / biological rationale

Mechanistically, the claim is biologically plausible. Actin is described as the primary component of the membrane cytoskeleton and a critical component of synaptic structural plasticity. In reward circuitry, changes in actin polymerization affect dendritic spine morphology, synaptic transmission, memory reconsolidation, and cue-associated seeking. Those processes are highly relevant to addiction, which depends on durable maladaptive learning in regions such as the nucleus accumbens, amygdala, prefrontal cortex, and hippocampus. ([pmc.ncbi.nlm.nih.gov](https://pmc.ncbi.nlm.nih.gov/articles/PMC11034636/)) [S11]

That said, β-actin is only one part of a larger machinery involving ADF/cofilin, Rac1, LIMK, cortactin, profilins, myosins, and other actin-binding proteins. Much of the most persuasive addiction biology points to these **regulators of actin turnover** or to **actin dynamics as a process**, rather than singling out ACTB expression or ACTB genetic variation as the primary driver. ([pmc.ncbi.nlm.nih.gov](https://pmc.ncbi.nlm.nih.gov/articles/PMC11034636/)) [S12]

## Bottom line

There is **credible preclinical evidence** that **β-actin/actin-cytoskeleton dynamics are involved in addiction-related neuroplasticity, drug-associated memory, and relapse-like behavior**, especially for cocaine, methamphetamine, heroin, and alcohol. However, the evidence that **ACTB itself** is a **direct, robust, and reproducible human addiction gene** is **limited**. So the claim is best judged **Mixed Evidence**: biologically plausible and partly supported at the pathway/protein-process level, but not strongly established as an ACTB-specific addiction association. ([pubmed.ncbi.nlm.nih.gov](https://pubmed.ncbi.nlm.nih.gov/16452681/)) [S13]

## Cited sources

- **[S1]** ACTB gene: MedlinePlus Genetics — https://medlineplus.gov/genetics/gene/actb
- **[S2]** Cocaine increases actin cycling: effects in the reinstatement model of drug seeking - PubMed — https://pubmed.ncbi.nlm.nih.gov/16452681/
- **[S3]** 
            Targeting the cytoskeleton as a therapeutic approach to substance use disorders - PMC
         — https://pmc.ncbi.nlm.nih.gov/articles/PMC11034636/
- **[S4]** Effects of acute and chronic methamphetamine administration on cynomolgus monkey hippocampus structure and cellular transcriptome - PubMed — https://pubmed.ncbi.nlm.nih.gov/29802913/
- **[S5]** Disrupting heroin-associated memory reconsolidation through actin polymerization inhibition in the nucleus accumbens core - PubMed — https://pubmed.ncbi.nlm.nih.gov/39716383/
- **[S6]** 
            Targeting the cytoskeleton as a therapeutic approach to substance use disorders - PMC
         — https://pmc.ncbi.nlm.nih.gov/articles/PMC11034636/
- **[S7]** ACTB gene: MedlinePlus Genetics — https://medlineplus.gov/genetics/gene/actb
- **[S8]** Common and distinguishing genetic factors for substance use behavior and disorder: an integrated analysis of genomic and transcriptomic studies from both human and animal studies - PubMed — https://pubmed.ncbi.nlm.nih.gov/35491750/
- **[S9]** 
            Multivariate Analysis of Subjective Responses to d-amphetamine In Healthy Volunteers Finds Novel Genetic Pathway Associations - PMC
         — https://pmc.ncbi.nlm.nih.gov/articles/PMC4504822/
- **[S10]** ACTB gene: MedlinePlus Genetics — https://medlineplus.gov/genetics/gene/actb
- **[S11]** 
            Targeting the cytoskeleton as a therapeutic approach to substance use disorders - PMC
         — https://pmc.ncbi.nlm.nih.gov/articles/PMC11034636/
- **[S12]** 
            Targeting the cytoskeleton as a therapeutic approach to substance use disorders - PMC
         — https://pmc.ncbi.nlm.nih.gov/articles/PMC11034636/
- **[S13]** Cocaine increases actin cycling: effects in the reinstatement model of drug seeking - PubMed — https://pubmed.ncbi.nlm.nih.gov/16452681/
