## Supplementary material for "Understanding Substance Dependence: What Differentiates Addictive from Non-Addictive Drugs?": Folder for supplementary files: addictive-genes-claims-first100-003-ptgds-gene-protein-is-implicated-in-addiction-412441c30216.md

# addictive-genes-claims-first100-003-412441c30216

**Original claim:** PTGDS gene/protein is implicated in addiction.

## Claim

PTGDS gene/protein is implicated in addiction.

## Verdict

Verdict: Mixed Evidence

There is **some evidence** linking PTGDS (the gene encoding lipocalin-type prostaglandin D synthase) to addiction-related biology, especially in **alcohol** and **nicotine** models, but the evidence is mostly **transcriptomic and associative**, not strong causal proof. PTGDS shows altered expression in alcohol-related human and mouse brain studies, and a nicotine self-administration study reported Ptgds among genes changing across addiction phases. However, PTGDS is **not currently a well-established, replicated core addiction gene** on the level of loci such as **ADH1B** for alcohol dependence or **OPRM1** for opioid addiction. ([pmc.ncbi.nlm.nih.gov](https://pmc.ncbi.nlm.nih.gov/articles/PMC10874299/)) [S1]

## Confidence

Medium.

## Supporting evidence

PTGDS is a biologically plausible brain gene for neuropsychiatric phenotypes. NCBI describes human **PTGDS** as a brain-preferential gene encoding a glutathione-independent prostaglandin D synthase that converts PGH2 to **PGD2**, with PGD2 acting as a **neuromodulator** and **trophic factor** in the CNS; mouse annotations also link Ptgds to regulation of the sleep/wake cycle. ([ncbi.nlm.nih.gov](https://www.ncbi.nlm.nih.gov/gene/5730?utm_source=openai)) [S2]

The strongest addiction-relevant evidence comes from **alcohol** studies. In a recent mouse model of alcohol dependence using chronic intermittent ethanol exposure with snRNA-seq and spatial transcriptomics in medial prefrontal cortex, **Ptgds was dysregulated in 17 of 19 identified cell types**, placing it among the broadly alcohol-responsive genes in that dataset. The same study also found strong region-specific alcohol-dependent transcriptional changes in Ptgds-enriched spatial clusters. ([pmc.ncbi.nlm.nih.gov](https://pmc.ncbi.nlm.nih.gov/articles/PMC10874299/)) [S3]

Human alcohol-related transcriptomic work also points in the same general direction, though less cleanly. A convergent functional genomics review listed **PTGDS as decreased in human alcoholic brain**, citing the classic frontal-cortex microarray study by Lewohl et al. Another systems-oriented paper likewise stated that PTGDS had already been linked to alcoholism in prior studies. This suggests PTGDS has appeared repeatedly in alcohol-related expression literature, even if it has not emerged as a top consensus locus across modern large-scale genetic studies. ([pmc.ncbi.nlm.nih.gov](https://pmc.ncbi.nlm.nih.gov/articles/PMC3309466/)) [S4]

There is also some evidence beyond alcohol. In a 2024 single-nucleus transcriptomics study of the ventral tegmental area during **nicotine self-administration**, the search result summary reports that **Ptgds** was among genes upregulated in both neurons and glia across addiction progression. The paper’s abstract frames these transcriptional changes as informative for mechanisms underlying **nicotine addiction**. ([sciencedirect.com](https://www.sciencedirect.com/science/article/pii/S1673852724002327?utm_source=openai)) [S5]

More broadly, alcohol-related single-cell studies in human brain show that **glial cells**—especially astrocytes, oligodendrocytes, and microglia—carry many of the strongest alcohol-associated transcriptomic changes. That matters because PTGDS/L-PGDS is heavily tied to non-neuronal and extracellular CNS biology, making its appearance in addiction-related datasets mechanistically coherent rather than random. ([pubmed.ncbi.nlm.nih.gov](https://pubmed.ncbi.nlm.nih.gov/32142123/?utm_source=openai)) [S6]

## Contradicting or limiting evidence

The main limitation is that the PTGDS-addiction link is **not yet strong causal evidence**. Most of the data are **expression changes** in postmortem tissue or animal models. Differential expression can reflect consequences of chronic drug exposure, compensatory responses, cell-state shifts, inflammation, or tissue composition differences; it does **not** by itself show that PTGDS drives addictive behavior. ([pmc.ncbi.nlm.nih.gov](https://pmc.ncbi.nlm.nih.gov/articles/PMC10874299/)) [S7]

Also, PTGDS is **not prominent in major replicated GWAS findings** for substance use disorders. For alcohol dependence, one of the clearest genome-wide significant signals in a large transancestral GWAS was **ADH1B**, not PTGDS. For opioid addiction, a large multi-trait GWAS highlighted **OPRM1** and gene-based associations with **PPP6C** and **FURIN**, again not PTGDS. This does not rule out PTGDS involvement, but it does argue against PTGDS being a leading, well-validated addiction-risk gene at present. ([pubmed.ncbi.nlm.nih.gov](https://pubmed.ncbi.nlm.nih.gov/30482948/)) [S8]

Evidence outside alcohol and nicotine is especially thin. A rat transcriptomic/metabolomic study on opioid tolerance explicitly noted that **little is known about Ptgds or prostaglandin D2 in opioid tolerance or chronic opioid use in general**. That is a useful reality check: the literature is not yet mature enough to say PTGDS is broadly established across addiction types. ([pmc.ncbi.nlm.nih.gov](https://pmc.ncbi.nlm.nih.gov/articles/PMC11201153/?utm_source=openai)) [S9]

There is also a specificity problem. PTGDS has been implicated in other CNS contexts—sleep regulation, oxidative stress responses, ischemia/neuroprotection, and some psychiatric conditions—so its appearance in addiction datasets may reflect **general brain homeostasis or glial response** rather than a uniquely addiction-specific function. ([pubmed.ncbi.nlm.nih.gov](https://pubmed.ncbi.nlm.nih.gov/22248185/?utm_source=openai)) [S10]

## Mechanistic / biological rationale

Mechanistically, PTGDS is plausible. PTGDS produces **PGD2**, and PGD2 signaling is deeply involved in CNS physiology. Authoritative gene summaries and a classic review describe PGD2/PGDS as important for **neuromodulation** and especially **sleep regulation**; inhibition of PGDS markedly reduces sleep in vivo. ([ncbi.nlm.nih.gov](https://www.ncbi.nlm.nih.gov/gene/5730?utm_source=openai)) [S11]

That matters because addiction is tightly intertwined with **sleep/circadian disruption**, **glial signaling**, and **neuroimmune adaptation**. Alcohol dependence studies increasingly highlight glial and immune pathways in affected brain regions, and nicotine addiction studies similarly emphasize glial transcriptional remodeling in reward circuitry. A PTGDS-centered prostaglandin pathway could therefore influence addiction indirectly by altering inflammatory tone, extracellular signaling, sleep pressure, and neuronal support functions. ([pubmed.ncbi.nlm.nih.gov](https://pubmed.ncbi.nlm.nih.gov/31358844/)) [S12]

PTGDS/L-PGDS also has reported **neuroprotective** properties. Experimental studies describe it as protecting against oxidative stress-induced neuronal death and as beneficial in cerebral ischemia models. Those functions fit a model in which PTGDS changes during chronic drug exposure could represent either a stress-response adaptation or a compensatory attempt to preserve neural homeostasis. But that remains a **plausible mechanism**, not proof that PTGDS is a causal addiction mediator. ([pubmed.ncbi.nlm.nih.gov](https://pubmed.ncbi.nlm.nih.gov/22248185/?utm_source=openai)) [S13]

## Bottom line

**PTGDS is plausibly involved in addiction-related neurobiology, but the current evidence is not definitive.** The case is strongest for **alcohol-related** brain transcriptomic changes, with additional suggestive evidence in **nicotine** models. Still, the literature is dominated by **association/expression** findings, and PTGDS has **not** emerged as a major replicated genetic risk locus for addiction in the way that ADH1B or OPRM1 have. So the fairest assessment is: **PTGDS is a credible but still preliminary addiction-implicated gene/protein, not a firmly established core addiction gene.** ([pmc.ncbi.nlm.nih.gov](https://pmc.ncbi.nlm.nih.gov/articles/PMC10874299/)) [S14]

## Cited sources

- **[S1]** 
            Cell-type brain-region specific changes in prefrontal cortex of a mouse model of alcohol dependence - PMC
         — https://pmc.ncbi.nlm.nih.gov/articles/PMC10874299/
- **[S2]** PTGDS prostaglandin D2 synthase [Homo sapiens (human)] - Gene - NCBI — https://www.ncbi.nlm.nih.gov/gene/5730?utm_source=openai
- **[S3]** 
            Cell-type brain-region specific changes in prefrontal cortex of a mouse model of alcohol dependence - PMC
         — https://pmc.ncbi.nlm.nih.gov/articles/PMC10874299/
- **[S4]** 
            Convergent functional genomic studies of omega-3 fatty acids in stress reactivity, bipolar disorder and alcoholism - PMC
         — https://pmc.ncbi.nlm.nih.gov/articles/PMC3309466/
- **[S5]** Nicotine-induced transcriptional changes and mitochondrial dysfunction in the ventral tegmental area revealed by single-nucleus transcriptomics - ScienceDirect — https://www.sciencedirect.com/science/article/pii/S1673852724002327?utm_source=openai
- **[S6]** Single cell transcriptome profiling of the human alcohol-dependent brain - PubMed — https://pubmed.ncbi.nlm.nih.gov/32142123/?utm_source=openai
- **[S7]** 
            Cell-type brain-region specific changes in prefrontal cortex of a mouse model of alcohol dependence - PMC
         — https://pmc.ncbi.nlm.nih.gov/articles/PMC10874299/
- **[S8]** Transancestral GWAS of alcohol dependence reveals common genetic underpinnings with psychiatric disorders - PubMed — https://pubmed.ncbi.nlm.nih.gov/30482948/
- **[S9]** Gene expression in the dorsal root ganglion and the cerebrospinal fluid metabolome in polyneuropathy and opioid tolerance in rats - PMC — https://pmc.ncbi.nlm.nih.gov/articles/PMC11201153/?utm_source=openai
- **[S10]** Lipocalin-type prostaglandin D synthase protects against oxidative stress-induced neuronal cell death - PubMed — https://pubmed.ncbi.nlm.nih.gov/22248185/?utm_source=openai
- **[S11]** PTGDS prostaglandin D2 synthase [Homo sapiens (human)] - Gene - NCBI — https://www.ncbi.nlm.nih.gov/gene/5730?utm_source=openai
- **[S12]** Glial gene networks associated with alcohol dependence - PubMed — https://pubmed.ncbi.nlm.nih.gov/31358844/
- **[S13]** Lipocalin-type prostaglandin D synthase protects against oxidative stress-induced neuronal cell death - PubMed — https://pubmed.ncbi.nlm.nih.gov/22248185/?utm_source=openai
- **[S14]** 
            Cell-type brain-region specific changes in prefrontal cortex of a mouse model of alcohol dependence - PMC
         — https://pmc.ncbi.nlm.nih.gov/articles/PMC10874299/
