## Supplementary material for "Understanding Substance Dependence: What Differentiates Addictive from Non-Addictive Drugs?": Folder for supplementary files: addictive-genes-claims-first100-004-apoe-gene-protein-is-implicated-in-addiction-10403b405567.md

# addictive-genes-claims-first100-004-10403b405567

**Original claim:** APOE gene/protein is implicated in addiction.

## Claim

**“APOE gene/protein is implicated in addiction.”**

## Verdict

Verdict: Mixed Evidence

## Confidence

Medium

## Supporting evidence

There **is some evidence** linking APOE to addiction-related phenotypes, especially for **alcohol** and, to a lesser extent, **smoking relapse/cessation**. In a preclinical study using human APOE-targeted replacement mice, APOE genotype affected alcohol-related behavior: during alcohol re-exposure, **male APOE4 mice showed increased alcohol preference and motivation** versus APOE3 mice, and APOE4 males also showed greater punishment-resistant responding for alcohol. That is a direct addiction-relevant signal, albeit in mice and in one experimental system. ([pubmed.ncbi.nlm.nih.gov](https://pubmed.ncbi.nlm.nih.gov/34585391/)) [S1]

Human evidence also suggests APOE may influence some **relapse-related** outcomes. In a pooled analysis of **917 adult smokers** from three randomized smoking-cessation trials, APOE ε4 carriage interacted with age: among smokers **older than 60 years**, ε4 carriers had lower quit rates and relapsed faster than noncarriers. This does not show APOE causes nicotine addiction generally, but it does support a possible role in **cessation failure/relapse vulnerability** in a specific subgroup. ([pubmed.ncbi.nlm.nih.gov](https://pubmed.ncbi.nlm.nih.gov/23247396/)) [S2]

For alcohol use disorder (AUD), APOE has also been linked to **clinical features** rather than clearly to disorder risk itself. In a 2023 study of abstinent middle-aged patients with AUD, APOE4 was associated with worse **memory/learning** in the AUD group and altered reelin-related signaling, suggesting APOE may shape cognitive consequences or recovery biology in AUD. A 2007 study in 194 alcohol-dependent patients reported a preliminary association between APOE alleles and **alcohol-withdrawal seizures** (positive for ε3, negative for ε2, not significant for ε4), again pointing to a possible role in some addiction-related complications. ([pubmed.ncbi.nlm.nih.gov](https://pubmed.ncbi.nlm.nih.gov/37350760/)) [S3]

Other substance-related studies are suggestive but indirect. In chronic heavy methamphetamine users, APOE ε4 did not define methamphetamine use disorder itself, but ε4 plus heavy methamphetamine exposure was associated with more adverse **brain morphometry/cognitive** findings, especially in frontal regions. That supports APOE as a modifier of substance-related brain effects, though not necessarily as a core addiction gene. ([pubmed.ncbi.nlm.nih.gov](https://pubmed.ncbi.nlm.nih.gov/37563863/)) [S4]

There is also mechanistic preclinical evidence that alcohol exposure engages APOE-relevant biology. In rats, chronic voluntary alcohol intake followed by abstinence altered expression of multiple **cholesterol-homeostasis genes** in the **prefrontal cortex, nucleus accumbens, mesencephalon, and amygdala**—regions central to addiction and relapse—with the authors concluding that alcohol produced durable dysregulation of cholesterol metabolism in key addiction circuits. Separately, chronic binge ethanol reduced cortical **APOE** expression in female AD-model mice, and apoE4 enhanced ethanol-induced neurotoxicity in APP-expressing neuronal cells. These studies are not proof of addiction causality, but they make APOE biologically plausible in addiction-related neuroadaptation. ([pubmed.ncbi.nlm.nih.gov](https://pubmed.ncbi.nlm.nih.gov/30618609/)) [S5]

## Contradicting or limiting evidence

The biggest limitation is that **large-scale human genetics has not established APOE as a major addiction-risk locus**. Recent reviews of substance use disorder genetics and alcohol use disorder genetics highlight replicated loci such as **ADH1B, ALDH2, DRD2, KLB, SLC39A8,** and **PDE4B**. A multivariate GWAS of **over 1 million people** for shared liability across substance use disorders identified multiple loci and genes, but APOE does not appear among the reported genome-wide signals or discussion of key genes. Likewise, major reviews and large AUD genetics papers do not identify APOE as a robust, replicated addiction gene. ([pmc.ncbi.nlm.nih.gov](https://pmc.ncbi.nlm.nih.gov/articles/PMC8477224/)) [S6]

The same pattern holds for smoking genetics. A very large meta-analysis of up to **622,409 individuals** identified many smoking-behavior loci and expanded the known genetic architecture of smoking traits, but APOE was not among the reported findings. So while one smoking-cessation study found an age-dependent APOE ε4 effect, that observation has not translated into APOE being a broadly accepted nicotine-addiction locus. ([nature.com](https://www.nature.com/articles/s41380-018-0313-0)) [S7]

Many APOE-positive studies are also **small, candidate-gene, or consequence-focused**. They often examine cognition, neurotoxicity, brain structure, withdrawal complications, or Alzheimer-related vulnerability in substance users, rather than **core addiction phenotypes** such as incident disorder risk, severity, cross-substance liability, or replicated genome-wide associations. That makes the evidence real but **indirect** for the specific claim that APOE is “implicated in addiction” itself. ([pubmed.ncbi.nlm.nih.gov](https://pubmed.ncbi.nlm.nih.gov/37350760/)) [S8]

Some studies are explicitly negative or only conditional. In alcohol-dependent patients, one MRI study found **no main effect of APOE4** on hippocampal volume loss; the effect emerged only in interaction with **hyperhomocysteinemia**. In the 2023 AUD cognition study, APOE4 carrier frequency was not elevated in AUD, and plasma APOE levels in ε4 carriers were similar regardless of AUD status. These findings argue against a simple, strong APOE→AUD-risk relationship. ([pubmed.ncbi.nlm.nih.gov](https://pubmed.ncbi.nlm.nih.gov/17420762/?utm_source=openai)) [S9]

## Mechanistic / biological rationale

APOE is a highly plausible **modifier** of addiction-related brain biology because it is a central regulator of **CNS lipid transport and cholesterol homeostasis**. In the brain, apoE is produced mainly by **astrocytes** and helps deliver cholesterol and other lipids to neurons through LDL-receptor-family pathways. This matters because cholesterol supply is important for **synapse formation, neuronal repair/remodeling, and synaptic communication**. ([pmc.ncbi.nlm.nih.gov](https://pmc.ncbi.nlm.nih.gov/articles/PMC4942259/)) [S10]

That biology intersects with addiction, because addiction is fundamentally a disorder of **lasting neuroplasticity** in cortico-striatal-limbic circuits. A recent Neuron study showed that astrocytic ApoE regulates neuronal cholesterol metabolism, histone acetylation, immediate-early gene expression, and **memory consolidation**; importantly, human **apoE4 was less effective than apoE3** in this neuronal metabolic/epigenetic regulation. Since addiction and relapse depend on learning, memory, and synaptic remodeling, this provides a credible pathway by which APOE isoforms could influence addiction-related adaptations. ([pubmed.ncbi.nlm.nih.gov](https://pubmed.ncbi.nlm.nih.gov/33503410/)) [S11]

APOE4 also appears less efficient than APOE3 in several neurobiological functions relevant to resilience after substance exposure, including cholesterol efflux/lipidation and aspects of neuronal support and repair. Preclinical alcohol studies further suggest that alcohol can durably disturb cholesterol-handling programs in the **PFC, NAc, mesencephalon, and amygdala**, which are classic relapse and reward regions. So the biological rationale is strong for **APOE as a modulator of neural homeostasis under substance exposure**. ([pmc.ncbi.nlm.nih.gov](https://pmc.ncbi.nlm.nih.gov/articles/PMC3022415/)) [S12]

Still, **biological plausibility is not the same as proof**. The mechanistic case mainly supports APOE as a factor that could influence how the brain responds to chronic substance use, abstinence, and neurotoxic stress—not yet as a firmly established primary addiction gene. ([pmc.ncbi.nlm.nih.gov](https://pmc.ncbi.nlm.nih.gov/articles/PMC8477224/)) [S13]

## Bottom line

APOE is **not** a well-established, replicated **core addiction-risk gene** in the way that ADH1B, ALDH2, CHRNA5/A3/B4, OPRM1, or DRD2-related signals are for specific substance phenotypes. However, there **is meaningful but uneven evidence** that APOE—especially **APOE4**—can influence **alcohol-related behavior in mice**, **smoking cessation/relapse in older adults**, and the **cognitive/neurobiological consequences** of substance exposure. Overall, APOE is best viewed as a **plausible modifier of addiction-related neurobiology and outcomes**, rather than a strongly proven general addiction gene. ([pubmed.ncbi.nlm.nih.gov](https://pubmed.ncbi.nlm.nih.gov/34585391/)) [S14]

## Cited sources

- **[S1]** Effects of sex and genotype in human APOE-targeted replacement mice on alcohol self-administration measured with the automated IntelliCage system before and after repeated mild traumatic brain injury - PubMed — https://pubmed.ncbi.nlm.nih.gov/34585391/
- **[S2]** APOE ɛ4, an Alzheimer's disease susceptibility allele, and smoking cessation - PubMed — https://pubmed.ncbi.nlm.nih.gov/23247396/
- **[S3]** Reelin Plasma Levels Identify Cognitive Decline in Alcohol Use Disorder Patients During Early Abstinence: The Influence of APOE4 Expression - PubMed — https://pubmed.ncbi.nlm.nih.gov/37350760/
- **[S4]** Independent and combined effects of methamphetamine use disorders and APOEε4 allele on cognitive performance and brain morphometry - PubMed — https://pubmed.ncbi.nlm.nih.gov/37563863/
- **[S5]** Persistent Neuroadaptations in the Expression of Genes Involved in Cholesterol Homeostasis Induced by Chronic, Voluntary Alcohol Intake in Rats - PubMed — https://pubmed.ncbi.nlm.nih.gov/30618609/
- **[S6]** 
            Genetics of substance use disorders: a review - PMC
         — https://pmc.ncbi.nlm.nih.gov/articles/PMC8477224/
- **[S7]** Meta-analysis of up to 622,409 individuals identifies 40 novel smoking behaviour associated genetic loci | Molecular Psychiatry — https://www.nature.com/articles/s41380-018-0313-0
- **[S8]** Reelin Plasma Levels Identify Cognitive Decline in Alcohol Use Disorder Patients During Early Abstinence: The Influence of APOE4 Expression - PubMed — https://pubmed.ncbi.nlm.nih.gov/37350760/
- **[S9]** Apolipoprotein E polymorphism, homocysteine serum levels and hippocampal volume in patients with alcoholism: an investigation of a gene-environment interaction - PubMed — https://pubmed.ncbi.nlm.nih.gov/17420762/?utm_source=openai
- **[S10]** 
            Central Nervous System Lipoproteins: ApoE and Regulation of Cholesterol Metabolism - PMC
         — https://pmc.ncbi.nlm.nih.gov/articles/PMC4942259/
- **[S11]** Astrocytic ApoE reprograms neuronal cholesterol metabolism and histone-acetylation-mediated memory - PubMed — https://pubmed.ncbi.nlm.nih.gov/33503410/
- **[S12]** 
            Apolipoprotein E: from lipid transport to neurobiology - PMC
         — https://pmc.ncbi.nlm.nih.gov/articles/PMC3022415/
- **[S13]** 
            Genetics of substance use disorders: a review - PMC
         — https://pmc.ncbi.nlm.nih.gov/articles/PMC8477224/
- **[S14]** Effects of sex and genotype in human APOE-targeted replacement mice on alcohol self-administration measured with the automated IntelliCage system before and after repeated mild traumatic brain injury - PubMed — https://pubmed.ncbi.nlm.nih.gov/34585391/
