## Supplementary material for "Understanding Substance Dependence: What Differentiates Addictive from Non-Addictive Drugs?": Folder for supplementary files: addictive-genes-claims-first100-005-gprc5b-gene-protein-is-implicated-in-addiction-2c8d05931975.md

# addictive-genes-claims-first100-005-2c8d05931975

**Original claim:** GPRC5B gene/protein is implicated in addiction.

## Claim

GPRC5B gene/protein is implicated in addiction.

## Verdict

Verdict: Mixed Evidence

## Confidence

Medium

## Supporting evidence

There is **some evidence linking GPRC5B to addiction-related biology, especially alcohol-related risk prioritization**, but it is not yet strong or direct. A cross-species prioritization study integrating human, mouse, fly, and worm ethanol/alcoholism datasets placed **GPRC5B among 47 top-ranked candidate genes**; in that study, higher-ranked genes were enriched for lower q-values in a COGA alcohol-dependence GWAS, and the authors explicitly interpreted the resulting set as relevant to “brain responses to ethanol and neural adaptations occurring with alcoholism.” GPRC5B was one of the genes in that top-priority list. ([bmcgenomics.biomedcentral.com](https://bmcgenomics.biomedcentral.com/articles/10.1186/1471-2164-13-S8-S16)) [S1]

A separate rat transcriptomic/genetic study of **lines selectively bred for high vs low ethanol consumption** reported **Gprc5b** among genes considered potentially interesting because they fell within a rat alcohol QTL and were significant in at least two line-pairs. That is not proof of causality, but it is the kind of convergent animal evidence often used to nominate addiction-relevant genes. ([pmc.ncbi.nlm.nih.gov](https://pmc.ncbi.nlm.nih.gov/articles/PMC3866700/?utm_source=openai)) [S2]

GPRC5B is also **well positioned anatomically** to matter for addiction circuitry. A detailed mouse-brain mapping study found expression in the **medial prefrontal cortex, nucleus accumbens, caudate-putamen, hippocampus, hypothalamus, medial and lateral habenula, amygdala, pons, and cerebellum**; the paper also noted that human GTEx data show GPRC5B is among the more highly expressed orphan GPCRs in the CNS. The same study found notable expression in **GABAergic neurons** across several regions and in **striatal cholinergic interneurons / GABAergic populations**, which are cell classes highly relevant to reward learning and substance-use phenotypes. ([pubmed.ncbi.nlm.nih.gov](https://pubmed.ncbi.nlm.nih.gov/35812210/)) [S3]

There is also broader neurobiological evidence that GPRC5B affects behaviors and neural functions that can be relevant to addiction vulnerability, even if not addiction-specific per se. Gprc5b knockout mice have been reported to show **behavioral abnormalities including reduced spontaneous locomotor activity and reduced response to novelty**, and loss of GPRC5B impairs **cerebellar synaptic plasticity and motor learning**. More recently, Gprc5b knockout mice were reported to be **resilient to retinoic-acid–induced depressive-like behavioral effects** and resistant to retinoic-acid suppression of adult hippocampal neurogenesis. These findings support a role in brain plasticity and behavior, which makes addiction involvement biologically plausible. ([frontiersin.org](https://www.frontiersin.org/journals/genetics/articles/10.3389/fgene.2013.00297/pdf)) [S4]

## Contradicting or limiting evidence

The biggest limitation is that **direct addiction evidence is sparse**. In the material reviewed, I did **not** find a strong human GWAS signal establishing GPRC5B as a replicated genome-wide significant addiction gene, nor did I find clear primary studies showing that manipulating GPRC5B directly changes classic addiction endpoints such as **drug self-administration, conditioned place preference, relapse/reinstatement, withdrawal severity, or compulsive intake**. The strongest human genetic signal I found was **candidate prioritization / enrichment**, not definitive association. ([bmcgenomics.biomedcentral.com](https://bmcgenomics.biomedcentral.com/articles/10.1186/1471-2164-13-S8-S16)) [S5]

Likewise, major reference material for the gene emphasizes other biology more than addiction. NCBI Gene lists GPRC5B literature tied to **body mass index, inflammatory signaling, diabetes-related biology, and rare leukodystrophy**, and the 2023 *Brain* review emphasizes **brain edema / water-homeostasis** links. Those are important functions, but they do not amount to established addiction involvement. ([ncbi.nlm.nih.gov](https://www.ncbi.nlm.nih.gov/gene/51704)) [S6]

Even the supportive alcohol-related studies are mostly **indirect**. The BMC Genomics study used a cross-species ranking strategy, which is valuable for hypothesis generation, but it does not itself prove that GPRC5B drives addiction in humans. The rat high/low-ethanol-consumption study similarly identifies Gprc5b as a **candidate within a QTL/module context**, not as a validated causal effector. ([bmcgenomics.biomedcentral.com](https://bmcgenomics.biomedcentral.com/articles/10.1186/1471-2164-13-S8-S16)) [S7]

## Mechanistic / biological rationale

Mechanistically, the claim is **plausible but still incomplete**. GPRC5B is an **orphan class C GPCR**, and its endogenous ligand is still not established in standard pharmacology references. That alone makes mechanistic interpretation harder than for better-characterized addiction targets. ([pharos.nih.gov](https://pharos.nih.gov/targets/GPRC5B?utm_source=openai)) [S8]

Still, several observations support possible relevance to addiction-related neuroadaptation. First, GPRC5B is enriched in **reward/control circuitry** such as mPFC, nucleus accumbens, and habenula. The habenula is particularly notable because it helps encode aversion, nicotine-related signaling, and negative motivational states; expression there does not prove addiction involvement, but it increases plausibility. ([pubmed.ncbi.nlm.nih.gov](https://pubmed.ncbi.nlm.nih.gov/35812210/)) [S9]

Second, GPRC5B has been linked to **synaptic development and plasticity**. Loss of GPRC5B disrupts Purkinje-cell synapse formation and cerebellar plasticity, while other work links it to adult hippocampal neurogenesis and behavioral responses to chronic retinoic acid. Addiction is fundamentally a disorder of maladaptive neuroplasticity, so a receptor that affects synapse formation, circuit responsiveness, and experience-dependent behavior is a credible candidate for involvement. That said, this is still an inference from broader neurobiology, not direct addiction proof. ([pubmed.ncbi.nlm.nih.gov](https://pubmed.ncbi.nlm.nih.gov/29481883/?utm_source=openai)) [S10]

Third, some non-neuronal signaling linked to GPRC5B may also be relevant in principle. Studies outside addiction show GPRC5B can modulate **NF-κB**, **Fyn-related signaling**, **cAMP/CREB-linked pathways**, and receptor interactions such as with prostaglandin signaling partners. Neuroimmune signaling, CREB-related transcription, and GPCR crosstalk are all important in substance-use disorders. But again, those links are mostly extrapolations; they have not yet been tied cleanly to addiction phenotypes for GPRC5B itself. ([pubmed.ncbi.nlm.nih.gov](https://pubmed.ncbi.nlm.nih.gov/31285284/?utm_source=openai)) [S11]

## Bottom line

**Yes, but only weakly to moderately.** GPRC5B has been **nominated as an alcohol/addiction-relevant candidate gene** by cross-species and animal-selection studies, and it is expressed in several **addiction-relevant brain regions** while influencing **neural plasticity and behavior**. However, the evidence is still **mostly indirect and hypothesis-generating**, not definitive. I would not call GPRC5B an established addiction gene at this point; I would call it a **plausible, understudied candidate with suggestive evidence**, strongest so far for **alcohol-related** relevance. ([bmcgenomics.biomedcentral.com](https://bmcgenomics.biomedcentral.com/articles/10.1186/1471-2164-13-S8-S16)) [S12]

## Cited sources

- **[S1]** Multi-species data integration and gene ranking enrich significant results in an alcoholism genome-wide association study | BMC Genomics | Springer Nature Link — https://bmcgenomics.biomedcentral.com/articles/10.1186/1471-2164-13-S8-S16
- **[S2]** Gene Expression within the Extended Amygdala of 5 Pairs of Rat Lines Selectively Bred for High or Low Ethanol Consumption - PMC — https://pmc.ncbi.nlm.nih.gov/articles/PMC3866700/?utm_source=openai
- **[S3]** Comprehensive Spatial Profile of the Orphan G Protein Coupled Receptor GPRC5B Expression in Mouse Brain - PubMed — https://pubmed.ncbi.nlm.nih.gov/35812210/
- **[S4]** G protein-linked signaling pathways in bipolar and major depressive disorders — https://www.frontiersin.org/journals/genetics/articles/10.3389/fgene.2013.00297/pdf
- **[S5]** Multi-species data integration and gene ranking enrich significant results in an alcoholism genome-wide association study | BMC Genomics | Springer Nature Link — https://bmcgenomics.biomedcentral.com/articles/10.1186/1471-2164-13-S8-S16
- **[S6]** GPRC5B G protein-coupled receptor class C group 5 member B [Homo sapiens (human)] - Gene - NCBI — https://www.ncbi.nlm.nih.gov/gene/51704
- **[S7]** Multi-species data integration and gene ranking enrich significant results in an alcoholism genome-wide association study | BMC Genomics | Springer Nature Link — https://bmcgenomics.biomedcentral.com/articles/10.1186/1471-2164-13-S8-S16
- **[S8]** Pharos : Target Details - GPRC5B — https://pharos.nih.gov/targets/GPRC5B?utm_source=openai
- **[S9]** Comprehensive Spatial Profile of the Orphan G Protein Coupled Receptor GPRC5B Expression in Mouse Brain - PubMed — https://pubmed.ncbi.nlm.nih.gov/35812210/
- **[S10]** Loss of GPRC5B impairs synapse formation of Purkinje cells with cerebellar nuclear neurons and disrupts cerebellar synaptic plasticity and motor learning - PubMed — https://pubmed.ncbi.nlm.nih.gov/29481883/?utm_source=openai
- **[S11]** GPRC5b Modulates Inflammatory Response in Glomerular Diseases via NF- κ B Pathway - PubMed — https://pubmed.ncbi.nlm.nih.gov/31285284/?utm_source=openai
- **[S12]** Multi-species data integration and gene ranking enrich significant results in an alcoholism genome-wide association study | BMC Genomics | Springer Nature Link — https://bmcgenomics.biomedcentral.com/articles/10.1186/1471-2164-13-S8-S16
