## Supplementary material for "Understanding Substance Dependence: What Differentiates Addictive from Non-Addictive Drugs?": Folder for supplementary files: addictive-genes-claims-first100-006-actg1-gene-protein-is-implicated-in-addiction-5244599d2ca9.md

# addictive-genes-claims-first100-006-5244599d2ca9

**Original claim:** ACTG1 gene/protein is implicated in addiction.

## Claim

ACTG1 gene/protein is implicated in addiction.

## Verdict

Verdict: Mixed Evidence

## Confidence

Medium

## Supporting evidence

ACTG1 encodes cytoplasmic γ-actin, a core component of the actin cytoskeleton that helps determine cell shape and motility. In neurons, actin dynamics are fundamental to dendritic spine remodeling and synaptic plasticity, which are central processes in addiction neurobiology. MedlinePlus describes ACTG1 as producing γ-actin, one of the ubiquitous non-muscle actins that help organize the intracellular actin network. In a mouse study, synaptic ACTG1 levels were shown to regulate hippocampal spine density, long-term potentiation, and contextual fear memory through TRIM3-dependent control of local ACTG1 turnover. ([medlineplus.gov](https://medlineplus.gov/download/genetics/gene/actg1.pdf)) [S1]

That mechanistic backdrop matters because addiction research broadly implicates actin-cytoskeleton remodeling in drug-induced neural plasticity. A review focused on addiction concluded that cytoskeletal dynamics are a necessary component of the synaptic and structural changes seen in addicted animals, and linked actin-regulating pathways to ethanol consumption and other drug-induced adaptations in reward-related circuits. ([pmc.ncbi.nlm.nih.gov](https://pmc.ncbi.nlm.nih.gov/articles/PMC3683094/)) [S2]

The strongest substance-related evidence for ACTG1 itself comes from alcohol-focused proteomics. A review of alcoholism proteomics found ACTG1 among the “well-consolidated” alcohol-sensitive proteins, reported as differentially expressed in **five** alcohol-related proteomic studies across human alcoholic brain and animal/cell models. In that same review, ACTG1 contributed to enriched pathways such as clathrin-mediated endocytosis and caveolar-mediated endocytosis, and the authors explicitly framed such proteomic signatures as relevant to biological processes underlying alcohol dependence. ([pmc.ncbi.nlm.nih.gov](https://pmc.ncbi.nlm.nih.gov/articles/PMC3857647/)) [S3]

There is also some opioid-related evidence. In rats examined after prolonged morphine withdrawal, hippocampal phosphoproteomics showed hypophosphorylated Actg1 among altered cytoskeletal proteins, and the authors interpreted the overall pattern as suggesting long-term homeostatic changes in hippocampal plasticity after morphine exposure/abstinence. That does not prove ACTG1 drives addictive behavior, but it does place the protein within neuroadaptations following chronic opioid exposure. ([pmc.ncbi.nlm.nih.gov](https://pmc.ncbi.nlm.nih.gov/articles/PMC8772819/)) [S4]

A mouse transcriptomic study of chronic ethanol self-administration further reported that striatal **Actg1** was downregulated in a “Heavy Drinkers” group, in a brain region the authors identified as critical for alcohol addiction. This is only one study and from an abstract/snippet rather than full-text evaluation here, but it is directionally consistent with ACTG1 being responsive to alcohol-related behavioral states. ([sciencedirect.com](https://www.sciencedirect.com/science/article/abs/pii/S0166432816304296?utm_source=openai)) [S5]

Finally, in humans, ACTG1 expression has been reported as increased in blood and buccal cells of smokers, where the authors proposed it as a tobacco smoking-responsive biomarker. This is weaker evidence for addiction per se because it addresses smoking exposure/biology rather than nicotine dependence or compulsive use, but it still shows substance-related responsiveness of ACTG1 in human samples. ([pubmed.ncbi.nlm.nih.gov](https://pubmed.ncbi.nlm.nih.gov/25447457/)) [S6]

## Contradicting or limiting evidence

The biggest limitation is that the evidence is mostly **indirect**. Much of it comes from proteomic or transcriptomic studies showing ACTG1 changes after alcohol, morphine, or smoking exposure, not from experiments demonstrating that ACTG1 manipulation changes drug self-administration, relapse, craving, or DSM-defined substance use disorder. The addiction-relevant ACTG1 literature located here is therefore largely correlative or mechanistic rather than causal. ([pmc.ncbi.nlm.nih.gov](https://pmc.ncbi.nlm.nih.gov/articles/PMC3857647/)) [S7]

Evidence is also not consistent across substances or models. In a rhesus monkey nucleus accumbens proteomics study after cocaine self-administration, ACTG1 was measured but **not** significantly different between cocaine and control groups; the study’s significant protein changes involved other targets instead. So ACTG1 is not a reproducibly altered addiction signal across all paradigms. ([pmc.ncbi.nlm.nih.gov](https://pmc.ncbi.nlm.nih.gov/articles/PMC3272768/)) [S8]

Direct human genetic evidence is especially sparse. In a GWAS Catalog-derived 2025 alcohol use disorder gene set, ACTG1 was not among the 12 listed associated genes; the genes highlighted instead included loci such as **ADH1B, ADH1C, DRD2,** and **SLC39A8**. That does not rule out a small effect or a role in other substance phenotypes, but it argues against ACTG1 being an established major susceptibility gene for AUD at present. ([maayanlab.cloud](https://maayanlab.cloud/Harmonizome/gene_set/Alcohol%2Buse%2Bdisorder/GWAS%2BCatalog%2BSNP-Phenotype%2BAssociations%2B2025)) [S9]

The tobacco study is another example of why precision matters: its conclusion was that ACTG1 may be a **smoking-responsive biomarker**, not that ACTG1 causes nicotine addiction. Likewise, the morphine and alcohol omics papers support involvement in substance-induced neuroadaptation, but they do not by themselves establish ACTG1 as an addiction gene in the stronger genetic/causal sense. ([pubmed.ncbi.nlm.nih.gov](https://pubmed.ncbi.nlm.nih.gov/25447457/)) [S10]

## Mechanistic / biological rationale

Biologically, the claim is plausible. ACTG1 encodes γ-actin, and actin remodeling is essential for dendritic spine structure, receptor trafficking, and synaptic plasticity. Those same processes are widely thought to underlie the transition from controlled drug use to compulsive use, especially in mesocorticolimbic circuits. ([medlineplus.gov](https://medlineplus.gov/download/genetics/gene/actg1.pdf)) [S11]

More specifically, the TRIM3-ACTG1 work showed that altering synaptic ACTG1 availability changes spine density and early LTP, and the authors proposed that ACTG1 helps control actin stability during synaptic plasticity. They further noted that changes in actin stability would be expected to affect spine morphogenesis as well as AMPA receptor recycling and anchoring. These are exactly the kinds of plasticity mechanisms often invoked in addiction models. ([pmc.ncbi.nlm.nih.gov](https://pmc.ncbi.nlm.nih.gov/articles/PMC4639863/)) [S12]

Alcohol proteomics also places ACTG1 in endocytosis-related pathways, which fits with the idea that substance exposure can alter trafficking and membrane protein organization at synapses. So, from a mechanistic standpoint, ACTG1 is a credible participant in neural homeostasis/plasticity responses to drugs. What remains uncertain is whether its role is merely downstream and nonspecific, or whether it is an important upstream determinant of addiction vulnerability or persistence. ([pmc.ncbi.nlm.nih.gov](https://pmc.ncbi.nlm.nih.gov/articles/PMC3857647/)) [S13]

## Bottom line

ACTG1 is **not** a well-established addiction gene in the way that classic alcohol- or nicotine-related loci are, and I did not find strong replicated human genetic or causal behavioral evidence tying ACTG1 directly to substance use disorder risk. However, there **is** meaningful indirect evidence: ACTG1 participates in synaptic actin biology, regulates plasticity-related neuronal phenotypes, and appears repeatedly in alcohol- and opioid-related omics studies, with weaker smoking-related exposure evidence as well. The fairest summary is that ACTG1 is a **plausible, indirectly supported plasticity-related candidate in addiction neurobiology**, but the current evidence is too limited to call it strongly established. ([pmc.ncbi.nlm.nih.gov](https://pmc.ncbi.nlm.nih.gov/articles/PMC4639863/)) [S14]

## Cited sources

- **[S1]** ACTG1 gene — https://medlineplus.gov/download/genetics/gene/actg1.pdf
- **[S2]** 
            Emerging roles of actin cytoskeleton regulating enzymes in drug addiction: Actin or reactin’? - PMC
         — https://pmc.ncbi.nlm.nih.gov/articles/PMC3683094/
- **[S3]** 
            Proteomic Approaches and Identification of Novel Therapeutic Targets for Alcoholism - PMC
         — https://pmc.ncbi.nlm.nih.gov/articles/PMC3857647/
- **[S4]** 
            Alterations in the Proteome and Phosphoproteome Profiles of Rat Hippocampus after Six Months of Morphine Withdrawal: Comparison with the Forebrain Cortex - PMC
         — https://pmc.ncbi.nlm.nih.gov/articles/PMC8772819/
- **[S5]** Inflexible ethanol intake: A putative link with the Lrrk2 pathway - ScienceDirect — https://www.sciencedirect.com/science/article/abs/pii/S0166432816304296?utm_source=openai
- **[S6]** Tobacco smoking-response genes in blood and buccal cells - PubMed — https://pubmed.ncbi.nlm.nih.gov/25447457/
- **[S7]** 
            Proteomic Approaches and Identification of Novel Therapeutic Targets for Alcoholism - PMC
         — https://pmc.ncbi.nlm.nih.gov/articles/PMC3857647/
- **[S8]** 
            Integrative proteomic analysis of the nucleus accumbens in rhesus monkeys following cocaine self-administration - PMC
         — https://pmc.ncbi.nlm.nih.gov/articles/PMC3272768/
- **[S9]** Gene Set - Alcohol use disorder — https://maayanlab.cloud/Harmonizome/gene_set/Alcohol%2Buse%2Bdisorder/GWAS%2BCatalog%2BSNP-Phenotype%2BAssociations%2B2025
- **[S10]** Tobacco smoking-response genes in blood and buccal cells - PubMed — https://pubmed.ncbi.nlm.nih.gov/25447457/
- **[S11]** ACTG1 gene — https://medlineplus.gov/download/genetics/gene/actg1.pdf
- **[S12]** 
            Ubiquitin ligase TRIM3 controls hippocampal plasticity and learning by regulating synaptic γ-actin levels - PMC
         — https://pmc.ncbi.nlm.nih.gov/articles/PMC4639863/
- **[S13]** 
            Proteomic Approaches and Identification of Novel Therapeutic Targets for Alcoholism - PMC
         — https://pmc.ncbi.nlm.nih.gov/articles/PMC3857647/
- **[S14]** 
            Ubiquitin ligase TRIM3 controls hippocampal plasticity and learning by regulating synaptic γ-actin levels - PMC
         — https://pmc.ncbi.nlm.nih.gov/articles/PMC4639863/
