## Supplementary material for "Understanding Substance Dependence: What Differentiates Addictive from Non-Addictive Drugs?": Folder for supplementary files: addictive-genes-claims-first100-007-hpca-gene-protein-is-implicated-in-addiction-bda2afee4f9b.md

# addictive-genes-claims-first100-007-bda2afee4f9b

**Original claim:** HPCA gene/protein is implicated in addiction.

## Claim

HPCA gene/protein is implicated in addiction.

## Verdict

Verdict: Mixed Evidence

There is **some evidence** linking HPCA/hippocalcin to addiction-relevant biology, especially in **opioid abstinence** and **alcohol-related brain gene networks**, but the evidence is **indirect and limited**. I did **not** find strong, replicated human genetic evidence that HPCA is a major addiction gene, nor direct causal studies showing that manipulating HPCA alone changes drug self-administration, relapse, or dependence phenotypes across substances. ([nature.com](https://www.nature.com/articles/s41386-020-00927-x)) [S1]

## Confidence

Medium

## Supporting evidence

HPCA encodes **hippocalcin**, a neuron-specific calcium sensor protein that is expressed almost exclusively in the brain and is particularly abundant in the **striatum/putamen**; Human Protein Atlas data also classify it as **brain-enriched** and relatively enriched in **basal ganglia** and **hippocampal formation**. Those are regions central to reward learning, habit formation, and neuroadaptations in substance use disorders, so the anatomic distribution is at least consistent with a possible role in addiction-related circuitry. ([pubmed.ncbi.nlm.nih.gov](https://pubmed.ncbi.nlm.nih.gov/25799108/)) [S2]

A notable **animal-study signal** comes from a 2021 mouse study comparing prolonged abstinence after morphine versus cocaine exposure. In that work, **Hpca** was one of seven medium spiny neuron marker genes showing **consistent down-regulation** in caudate-putamen, nucleus accumbens, and central amygdala during **morphine abstinence**, whereas this pattern was **not** seen during cocaine abstinence. In the nucleus accumbens specifically, Hpca down-regulation during morphine abstinence was reported and was “rescued” alongside behavioral improvement by chronic mGluR4 positive allosteric modulation. This does not prove HPCA causes addiction, but it does place Hpca within opioid-abstinence-related molecular adaptations in core addiction circuitry. ([nature.com](https://www.nature.com/articles/s41386-020-00927-x)) [S3]

There is also **human-plus-animal systems-level evidence** in alcohol research. A cross-species systems genomics study of binge drinking/alcohol dependence found that **Hpca** was among 13 genes differentially expressed in both **human alcoholic brain** and **ethanol-naive high-drinking mice**. The overlapping neuronal subnetworks were enriched for pathways involving **dopamine signaling, synaptic plasticity, glutamatergic signaling, long-term depression, and CREB signaling**, all highly relevant to addiction neurobiology. This again supports HPCA as part of an addiction-relevant molecular network, though not as a proven causal driver. ([pmc.ncbi.nlm.nih.gov](https://pmc.ncbi.nlm.nih.gov/articles/PMC6459809/)) [S4]

More broadly, addiction involves persistent molecular and circuit-level adaptations in cortico-striatal and accumbal systems, and the nucleus accumbens/striatal medium spiny neurons are a major substrate for these changes. Because HPCA is enriched in these neuronal populations and tracks with drug-related transcriptional changes in some models, it is reasonable to say HPCA is **plausibly implicated** in addiction biology at a network/mechanistic level. ([pubmed.ncbi.nlm.nih.gov](https://pubmed.ncbi.nlm.nih.gov/27475769/?utm_source=openai)) [S5]

## Contradicting or limiting evidence

The biggest limitation is the **lack of strong direct human genetic evidence**. A current GWAS Catalog search for **HPCA** returns **no cataloged GWAS results**, which argues against HPCA being a replicated, genome-wide-established addiction susceptibility gene at present. ([ebi.ac.uk](https://www.ebi.ac.uk/gwas/search?query=HPCA&utm_source=openai)) [S6]

The available positive evidence is also **substance- and context-specific** rather than broad. In the morphine-versus-cocaine abstinence study, Hpca changes were prominent in **morphine abstinence** but **not** in cocaine abstinence. That weakens any claim that HPCA has a general, cross-drug role in addiction. ([nature.com](https://www.nature.com/articles/s41386-020-00927-x)) [S7]

The alcohol study is important but still **indirect**. It used network overlap and differential expression across species to infer shared biology; it did **not** demonstrate that HPCA variants or HPCA-specific perturbation alter alcohol intake, craving, relapse, or dependence in humans. In fact, the authors framed the identified genes as **potential targets** and convergent network components rather than validated causal addiction genes. ([pmc.ncbi.nlm.nih.gov](https://pmc.ncbi.nlm.nih.gov/articles/PMC6459809/)) [S8]

Also, much of the best-established literature on HPCA is about **neuronal calcium signaling** and **dystonia**, not addiction. HPCA mutations are a recognized cause of autosomal-recessive dystonia, and functional work on hippocalcin has focused heavily on calcium-channel interactions, excitability, and synaptic physiology. That supports biological relevance in neurons, but it does **not** by itself establish an addiction-specific role. ([pubmed.ncbi.nlm.nih.gov](https://pubmed.ncbi.nlm.nih.gov/25799108/)) [S9]

## Mechanistic / biological rationale

Mechanistically, HPCA is a credible candidate for involvement in addiction-related plasticity because hippocalcin is a **neuronal calcium sensor**. Functional studies indicate it contributes to **voltage-gated calcium-channel regulation**, **slow afterhyperpolarization (sAHP)**, and **NMDAR-dependent long-term depression via AMPAR endocytosis**. These processes shape neuronal excitability, firing adaptation, and synaptic weakening/selection. ([pubmed.ncbi.nlm.nih.gov](https://pubmed.ncbi.nlm.nih.gov/25799108/)) [S10]

Those functions matter because addiction is driven by long-lasting changes in **synaptic plasticity and excitability** within reward-related circuits, especially the **nucleus accumbens and other striatal medium spiny neurons**. If HPCA changes alter calcium handling, sAHP, or LTD in these cells, that could influence how drug exposure or abstinence remodels reward learning, cue reactivity, or negative affect during withdrawal. This rationale is biologically coherent, and the morphine-abstinence data fit that model. ([pubmed.ncbi.nlm.nih.gov](https://pubmed.ncbi.nlm.nih.gov/27475769/?utm_source=openai)) [S11]

Still, the mechanistic case remains **plausible rather than proven** for addiction. The strongest direct mechanistic evidence for HPCA itself concerns neuronal signaling generally, not addiction phenotypes specifically. Put differently: HPCA has the **right kind of function in the right kind of cells**, but the step from “relevant neuronal regulator” to “established addiction gene/protein” has not yet been firmly demonstrated. ([v22.proteinatlas.org](https://v22.proteinatlas.org/ENSG00000121905-HPCA/brain)) [S12]

## Bottom line

HPCA/hippocalcin is **not well established as a canonical addiction gene**, but it is **not unsupported either**. The best current evidence suggests HPCA is an **indirect, mechanistically plausible participant** in addiction-related neurobiology—particularly in **opioid abstinence** and in **alcohol-related neuronal gene networks**—rather than a strongly validated, replicated addiction-risk gene in humans. So the claim is **too strong if meant as a proven human addiction gene**, but **reasonable if meant in the weaker sense of being involved in addiction-relevant neural adaptations**. ([nature.com](https://www.nature.com/articles/s41386-020-00927-x)) [S13]

## Cited sources

- **[S1]** Facilitating mGluR4 activity reverses the long-term deleterious consequences of chronic morphine exposure in male mice | Neuropsychopharmacology — https://www.nature.com/articles/s41386-020-00927-x
- **[S2]** Mutations in HPCA cause autosomal-recessive primary isolated dystonia - PubMed — https://pubmed.ncbi.nlm.nih.gov/25799108/
- **[S3]** Facilitating mGluR4 activity reverses the long-term deleterious consequences of chronic morphine exposure in male mice | Neuropsychopharmacology — https://www.nature.com/articles/s41386-020-00927-x
- **[S4]** 
            Dissecting Brain Networks Underlying Alcohol Binge Drinking Using a Systems Genomics Approach - PMC
         — https://pmc.ncbi.nlm.nih.gov/articles/PMC6459809/
- **[S5]** Neurobiology of addiction: a neurocircuitry analysis - PubMed — https://pubmed.ncbi.nlm.nih.gov/27475769/?utm_source=openai
- **[S6]** GWAS Catalog — https://www.ebi.ac.uk/gwas/search?query=HPCA&utm_source=openai
- **[S7]** Facilitating mGluR4 activity reverses the long-term deleterious consequences of chronic morphine exposure in male mice | Neuropsychopharmacology — https://www.nature.com/articles/s41386-020-00927-x
- **[S8]** 
            Dissecting Brain Networks Underlying Alcohol Binge Drinking Using a Systems Genomics Approach - PMC
         — https://pmc.ncbi.nlm.nih.gov/articles/PMC6459809/
- **[S9]** Mutations in HPCA cause autosomal-recessive primary isolated dystonia - PubMed — https://pubmed.ncbi.nlm.nih.gov/25799108/
- **[S10]** Mutations in HPCA cause autosomal-recessive primary isolated dystonia - PubMed — https://pubmed.ncbi.nlm.nih.gov/25799108/
- **[S11]** Neurobiology of addiction: a neurocircuitry analysis - PubMed — https://pubmed.ncbi.nlm.nih.gov/27475769/?utm_source=openai
- **[S12]** Brain tissue expression of HPCA - Summary - The Human Protein Atlas — https://v22.proteinatlas.org/ENSG00000121905-HPCA/brain
- **[S13]** Facilitating mGluR4 activity reverses the long-term deleterious consequences of chronic morphine exposure in male mice | Neuropsychopharmacology — https://www.nature.com/articles/s41386-020-00927-x
