## Supplementary material for "Understanding Substance Dependence: What Differentiates Addictive from Non-Addictive Drugs?": Folder for supplementary files: addictive-genes-claims-first100-008-atp1b2-gene-protein-is-implicated-in-addiction-5f52856df931.md

# addictive-genes-claims-first100-008-5f52856df931

**Original claim:** ATP1B2 gene/protein is implicated in addiction.

## Claim

ATP1B2 gene/protein is implicated in addiction.

## Verdict

Verdict: Mixed Evidence

There is **some indirect and low-to-moderate evidence** linking ATP1B2 to addiction-related biology, especially **alcohol use disorder (AUD)**, but I did **not** find strong evidence that ATP1B2 is a **replicated addiction gene/protein** in the way that term is usually used. The main supportive signal is a **small human postmortem AUD expression study** plus broader evidence that **astrocytes/glial biology** matter in addiction, while ATP1B2 is an established **astrocyte-enriched Na,K-ATPase β2 subunit**. However, I did **not** find a robust human GWAS signal, replicated clinical association, or a causal addiction-behavior study directly manipulating ATP1B2 itself. ([pmc.ncbi.nlm.nih.gov](https://pmc.ncbi.nlm.nih.gov/articles/PMC4221234/?utm_source=openai)) [S1]

## Confidence

Medium

## Supporting evidence

A human postmortem prefrontal cortex study in AUD reported ATP1B2 among genes showing altered expression: the paper’s table lists ATP1B2 with higher mean expression in AUD cases than controls and a permutation-based p-value of 0.029. That is a signal in the right direction for “implication,” but the same study explicitly stated that its differential-expression findings **did not survive multiple-testing correction**, which substantially weakens the claim. ([pmc.ncbi.nlm.nih.gov](https://pmc.ncbi.nlm.nih.gov/articles/PMC4221234/?utm_source=openai)) [S2]

Independent of ATP1B2 specifically, there is now fairly good evidence that **glial/astrocyte adaptations are relevant to alcohol addiction biology**. In human single-nucleus RNA-seq of prefrontal cortex from alcohol-dependent donors, the largest numbers of differentially expressed genes were found in **astrocytes, oligodendrocytes, and microglia**, not neurons. Mouse studies likewise show that chronic ethanol changes astrocyte-specific transcriptomes/translatomes in addiction-relevant regions including the **prefrontal cortex** and **nucleus accumbens**. This matters because ATP1B2 is a well-established astrocyte surface protein. ([pubmed.ncbi.nlm.nih.gov](https://pubmed.ncbi.nlm.nih.gov/32142123/?utm_source=openai)) [S3]

ATP1B2 itself is biologically positioned in those cell types. A JBC study showed that the widely used astrocyte antibody **ACSA-2 targets ATP1B2 on the plasma membrane of primary astrocytes**, making ATP1B2 a bona fide astrocyte-enriched marker rather than a generic housekeeping ATPase component. That supports the plausibility that ATP1B2 could participate in addiction-relevant glial adaptations, at least in alcohol-related settings. ([pubmed.ncbi.nlm.nih.gov](https://pubmed.ncbi.nlm.nih.gov/28373281/?utm_source=openai)) [S4]

More broadly, alcohol studies focused on astrocytes support the concept that **astrocyte-derived molecules can causally influence drinking behavior**. For example, a 2023 study reported that the astrocytic matricellular protein **hevin/SPARCL1** is increased in multiple brain regions of AUD subjects, and knockdown of hevin in **nucleus accumbens astrocytes** increased ethanol consumption in mice. This does not implicate ATP1B2 directly, but it strengthens the case that astrocyte-expressed proteins are legitimate addiction-relevant candidates. ([pubmed.ncbi.nlm.nih.gov](https://pubmed.ncbi.nlm.nih.gov/36830603/?utm_source=openai)) [S5]

## Contradicting or limiting evidence

The biggest limitation is the lack of **direct** evidence. I did **not** find a study showing that ATP1B2 variants are robustly associated with AUD or other substance use disorders in major human genetics datasets, nor did I find an experiment where ATP1B2 manipulation directly altered classic addiction phenotypes such as **drug self-administration, conditioned place preference, escalation, relapse, or withdrawal severity**. By contrast, major alcohol genetics studies retrieved in this search emphasized established loci such as **ADH1B** and, in rare-variant analyses, genes such as **ADH1C** and **PMM2**, rather than ATP1B2. ([bbrfoundation.org](https://bbrfoundation.org/sites/default/files/pdfs/jul-2023-webinar-slides.pdf?utm_source=openai)) [S6]

The only clearly ATP1B2-specific human addiction signal I found was that older microarray study in postmortem AUD cortex, and even there the authors cautioned that the expression findings **did not survive multiple-testing correction**. That makes the result **hypothesis-generating**, not confirmatory. ([pubmed.ncbi.nlm.nih.gov](https://pubmed.ncbi.nlm.nih.gov/25383235/?utm_source=openai)) [S7]

Also, a more recent human single-cell transcriptomic study of alcohol-dependent brain found substantial glial dysregulation overall, but ATP1B2 was **not highlighted** among the reported alcohol-related genes in the searchable article text. That does not rule out relevance, but it suggests ATP1B2 is **not currently a prominent or replicated signal** in the human AUD transcriptomics literature. ([pubmed.ncbi.nlm.nih.gov](https://pubmed.ncbi.nlm.nih.gov/32142123/?utm_source=openai)) [S8]

Finally, much of the strongest ATP1B2 literature is about **neurodevelopment, retinal degeneration, glioma biology, and astrocyte isolation/identity**, not addiction. So the evidence base is currently **indirectly supportive** rather than addiction-specific. ([pubmed.ncbi.nlm.nih.gov](https://pubmed.ncbi.nlm.nih.gov/9801359/?utm_source=openai)) [S9]

## Mechanistic / biological rationale

ATP1B2 encodes the **β2 subunit of the Na,K-ATPase**, also known as **AMOG**. Classic mouse work shows this subunit has dual relevance: it is both part of the ion pump and has **adhesion/neurite outgrowth-promoting** properties. Genetic replacement/rescue studies showed that loss of β2 has major CNS consequences, including juvenile lethality unless partially rescued and progressive photoreceptor degeneration, indicating that ATP1B2 is functionally important in nervous system homeostasis. ([pubmed.ncbi.nlm.nih.gov](https://pubmed.ncbi.nlm.nih.gov/9801359/?utm_source=openai)) [S10]

A reasonable **inference** is that ATP1B2 could influence addiction through **astrocyte-mediated ion and glutamate homeostasis**, rather than through a classic neurotransmitter receptor mechanism. Astrocytic glutamate uptake depends on the transmembrane sodium gradient maintained by Na,K-ATPase; experimental work on astrocytes showed that Na,K-ATPase α2 activity supports efficient restoration of sodium loads and glutamate/aspartate uptake. Because ATP1B2 is an astrocyte membrane protein and β subunits help assemble/stabilize Na,K-ATPase complexes, perturbing ATP1B2 could plausibly alter excitatory signaling, synaptic plasticity, or extracellular glutamate handling in reward circuits. That is mechanistically plausible, but it remains an **inference**, not direct proof for addiction. ([pubmed.ncbi.nlm.nih.gov](https://pubmed.ncbi.nlm.nih.gov/24901986/?utm_source=openai)) [S11]

This rationale fits the broader addiction literature: chronic ethanol exposure alters astrocyte-specific gene expression in prefrontal cortex and nucleus accumbens, and glial dysfunction is increasingly recognized in AUD. So ATP1B2 sits in a biologically credible pathway, even though the pathway-level evidence is stronger than the gene-specific evidence. ([nature.com](https://www.nature.com/articles/s41397-017-0012-2)) [S12]

## Bottom line

**Best current reading:** ATP1B2 is **not a well-established addiction gene/protein**, but it is a **plausible, weakly supported candidate**, especially in **alcohol-related** brain biology. The direct evidence is limited to a small, non-definitive human AUD expression signal, while the stronger case comes from ATP1B2’s role in **astrocytes**, **Na,K-ATPase function**, and **glutamate/ion homeostasis** in neural circuits that are relevant to addiction. So if “implicated” means **firmly established**, the claim is too strong; if it means **biologically plausible with some preliminary AUD-related evidence**, the claim is reasonable. ([pmc.ncbi.nlm.nih.gov](https://pmc.ncbi.nlm.nih.gov/articles/PMC4221234/?utm_source=openai)) [S13]

I found **suggestive but sparse** ATP1B2-addiction evidence, mostly around alcohol and astrocytes; what I did **not** find were replicated genetic associations or direct causal addiction-behavior studies focused on ATP1B2 itself. ([bbrfoundation.org](https://bbrfoundation.org/sites/default/files/pdfs/jul-2023-webinar-slides.pdf?utm_source=openai)) [S14]

## Cited sources

- **[S1]** Differential Expression of miR-130a in Postmortem Prefrontal Cortex of Subjects with Alcohol Use Disorders - PMC — https://pmc.ncbi.nlm.nih.gov/articles/PMC4221234/?utm_source=openai
- **[S2]** Differential Expression of miR-130a in Postmortem Prefrontal Cortex of Subjects with Alcohol Use Disorders - PMC — https://pmc.ncbi.nlm.nih.gov/articles/PMC4221234/?utm_source=openai
- **[S3]** Single cell transcriptome profiling of the human alcohol-dependent brain - PubMed — https://pubmed.ncbi.nlm.nih.gov/32142123/?utm_source=openai
- **[S4]** An immunoaffinity-based method for isolating ultrapure adult astrocytes based on ATP1B2 targeting by the ACSA-2 antibody - PubMed — https://pubmed.ncbi.nlm.nih.gov/28373281/?utm_source=openai
- **[S5]** The Matricellular Protein Hevin Is Involved in Alcohol Use Disorder - PubMed — https://pubmed.ncbi.nlm.nih.gov/36830603/?utm_source=openai
- **[S6]** What Genetics is Telling Us About — https://bbrfoundation.org/sites/default/files/pdfs/jul-2023-webinar-slides.pdf?utm_source=openai
- **[S7]** Differential Expression of miR-130a in Postmortem Prefrontal Cortex of Subjects with Alcohol Use Disorders - PubMed — https://pubmed.ncbi.nlm.nih.gov/25383235/?utm_source=openai
- **[S8]** Single cell transcriptome profiling of the human alcohol-dependent brain - PubMed — https://pubmed.ncbi.nlm.nih.gov/32142123/?utm_source=openai
- **[S9]** Na,K-ATPase subunit beta1 knock-in prevents lethality of beta2 deficiency in mice - PubMed — https://pubmed.ncbi.nlm.nih.gov/9801359/?utm_source=openai
- **[S10]** Na,K-ATPase subunit beta1 knock-in prevents lethality of beta2 deficiency in mice - PubMed — https://pubmed.ncbi.nlm.nih.gov/9801359/?utm_source=openai
- **[S11]** Role of Na,K-ATPase α1 and α2 isoforms in the support of astrocyte glutamate uptake - PubMed — https://pubmed.ncbi.nlm.nih.gov/24901986/?utm_source=openai
- **[S12]** Astrocyte-specific transcriptome responses to chronic ethanol consumption | The Pharmacogenomics Journal — https://www.nature.com/articles/s41397-017-0012-2
- **[S13]** Differential Expression of miR-130a in Postmortem Prefrontal Cortex of Subjects with Alcohol Use Disorders - PMC — https://pmc.ncbi.nlm.nih.gov/articles/PMC4221234/?utm_source=openai
- **[S14]** What Genetics is Telling Us About — https://bbrfoundation.org/sites/default/files/pdfs/jul-2023-webinar-slides.pdf?utm_source=openai
