## Supplementary material for "Understanding Substance Dependence: What Differentiates Addictive from Non-Addictive Drugs?": Folder for supplementary files: addictive-genes-claims-first100-009-atp6v0c-gene-protein-is-implicated-in-addiction-aca9c022ffec.md

# addictive-genes-claims-first100-009-aca9c022ffec

**Original claim:** ATP6V0C gene/protein is implicated in addiction.

## Claim

**ATP6V0C gene/protein is implicated in addiction.**

## Verdict

Verdict: Mixed Evidence

There is **some evidence consistent with involvement**, but it is **mostly indirect and preclinical**. ATP6V0C is a brain-expressed V-ATPase subunit with clear roles in neuronal vesicle acidification and neurotransmission, and it appears in several drug-exposure or addiction-model transcriptomic studies. However, I did **not** find strong evidence that ATP6V0C is an **established human addiction-risk gene** or a replicated causal addiction gene/protein in the way that loci such as **CHRNA5**, **DRD2**, or **PDE4B** are discussed in major addiction genetics papers. ([ncbi.nlm.nih.gov](https://www.ncbi.nlm.nih.gov/gene/527)) [S1]

## Confidence

**Medium.**

Confidence is moderate because the biological rationale is strong and multiple animal/omics studies point in the same direction, but the evidence base for **direct addiction-specific causality in humans** is limited. ([ncbi.nlm.nih.gov](https://www.ncbi.nlm.nih.gov/gene/527)) [S2]

## Supporting evidence

ATP6V0C encodes the **V0 subunit c** of the vacuolar ATPase (V-ATPase). NCBI states that V-ATPase-dependent organelle acidification is necessary for processes including **synaptic vesicle proton gradient generation**, and ATP6V0C is expressed broadly, including in **brain**. The Human Protein Atlas places ATP6V0C in a neuronal single-cell expression cluster annotated as **“Neurons – Neurotransmission & signaling”** and shows enhanced expression across multiple brain cell types. That gives ATP6V0C a plausible foothold in circuits relevant to reward and dependence. ([ncbi.nlm.nih.gov](https://www.ncbi.nlm.nih.gov/gene/527)) [S3]

In a 2024 single-nucleus transcriptomic study of the **ventral tegmental area (VTA)** during **nicotine self-administration** in mice, investigators reported that nicotine addiction progression was accompanied by significant transcriptional changes in energy-metabolism genes, and the search snippet for the paper specifically notes **Atp6v0c** among genes **upregulated** during this process. Because the VTA is central to nicotine reinforcement, this is one of the more relevant pieces of addiction-adjacent evidence for ATP6V0C. ([sciencedirect.com](https://www.sciencedirect.com/science/article/pii/S1673852724002327)) [S4]

In an alcohol-related mouse study, chronic ethanol exposure followed by **21 days of abstinence** altered prefrontal cortex transcription in selectively bred mouse lines differing in withdrawal severity. **Atp6v0c** was among the significantly changed genes in the dataset, and it also appeared in enriched GO categories such as **vacuolar membrane** and **monovalent inorganic cation transport**. Importantly, that study linked abstinence-associated pathway changes to **relapse drinking** biology, making ATP6V0C at least a candidate component of alcohol-related neuroadaptation. ([pmc.ncbi.nlm.nih.gov](https://pmc.ncbi.nlm.nih.gov/articles/PMC3014377/)) [S5]

A chronic **methamphetamine** mouse study also found **Atp6v0c** among lysosome-associated genes upregulated in oligodendrocytes after meth exposure. This does not prove a role in addiction behavior, but it supports the idea that ATP6V0C participates in brain cellular responses to addictive drugs, especially in lysosomal/autophagic pathways. ([pmc.ncbi.nlm.nih.gov](https://pmc.ncbi.nlm.nih.gov/articles/PMC11766323/)) [S6]

There is also older experimental literature, summarized in a review, suggesting ATP6V0C can affect **dopamine release**. The review describes earlier studies in which ATP6V0C overexpression increased **striatal dopamine overflow** in mice. Because mesolimbic dopamine signaling is central to reinforcement and drug reward, this is mechanistically relevant, though it is not addiction-specific evidence by itself. ([pmc.ncbi.nlm.nih.gov](https://pmc.ncbi.nlm.nih.gov/articles/PMC10717279/)) [S7]

## Contradicting or limiting evidence

The biggest limitation is that I did **not find strong, replicated human genetic evidence** placing ATP6V0C among the major addiction loci. In a recent multi-ancestral GWAS of **clinically defined nicotine dependence** involving **61,861 individuals**, the abstract highlights the well-known **CHRNA5** association and broader polygenic relationships with other substance-use disorders, but ATP6V0C is not highlighted. Likewise, a large multivariate GWAS of over **1 million** individuals identified a general addiction-risk factor with **17 genome-wide significant loci** and gene-based hits including **FTO, DRD2, and PDE4B**; ATP6V0C was not among the named top genes in the material reviewed. This is not definitive proof against involvement, but it argues against ATP6V0C being a currently well-established addiction gene. ([pubmed.ncbi.nlm.nih.gov](https://pubmed.ncbi.nlm.nih.gov/40831304/)) [S8]

Much of the positive evidence is also **indirect**. Drug-induced expression changes can reflect **toxicity, stress responses, mitochondrial remodeling, lysosomal adaptation, or withdrawal neurobiology**, rather than a specific role in reinforcement, compulsive use, craving, or relapse. The nicotine, alcohol, and methamphetamine studies support **association with drug-response biology**, but they do not by themselves show that manipulating ATP6V0C changes addiction-like behavior. ([sciencedirect.com](https://www.sciencedirect.com/science/article/pii/S1673852724002327)) [S9]

Another limitation is that ATP6V0C is best established in other neurologic contexts, especially **epilepsy/neurodevelopmental disease**, not substance-use disorder. Multiple human studies now support ATP6V0C variants as causes of neurodevelopmental disorders often associated with epilepsy, which strengthens its CNS relevance but does **not** specifically validate it as an addiction gene. ([pubmed.ncbi.nlm.nih.gov](https://pubmed.ncbi.nlm.nih.gov/37161035/?utm_source=openai)) [S10]

## Mechanistic / biological rationale

Mechanistically, ATP6V0C is highly plausible. V-ATPase acidifies intracellular vesicles, and that proton gradient is required for **loading neurotransmitters into synaptic vesicles** and maintaining neuronal organelle homeostasis. Addiction-related plasticity depends heavily on vesicle cycling, dopamine/glutamate/GABA signaling, and cellular stress adaptation, so a gene like ATP6V0C could reasonably modulate those processes. ([ncbi.nlm.nih.gov](https://www.ncbi.nlm.nih.gov/gene/527)) [S11]

ATP6V0C also sits at the intersection of several processes repeatedly implicated in addiction neurobiology: **synaptic transmission**, **lysosomal/autophagic function**, and **mitochondrial/energy metabolism**. The nicotine VTA study emphasized mitochondrial dysfunction and energy-metabolism shifts during self-administration, while the methamphetamine study linked Atp6v0c to lysosomal pathway changes. Those findings fit a broader model in which ATP6V0C contributes to neuronal and glial homeostasis under chronic drug exposure. ([sciencedirect.com](https://www.sciencedirect.com/science/article/pii/S1673852724002327?utm_source=openai)) [S12]

Human disease genetics further supports that ATP6V0C is functionally important in the nervous system. NCBI lists **early-onset epilepsy with or without developmental delay** as an associated condition, and published studies show ATP6V0C variants impair V-ATPase function and produce seizure/neurodevelopmental phenotypes. That does not equal addiction, but it indicates the gene is biologically potent in brain excitability and network stability—traits that can intersect with addiction vulnerability and drug responses. ([ncbi.nlm.nih.gov](https://www.ncbi.nlm.nih.gov/gene/527)) [S13]

## Bottom line

ATP6V0C is **biologically plausible** and has **some supportive preclinical evidence** linking it to drug-response pathways relevant to addiction, especially nicotine, alcohol-abstinence neuroadaptation, methamphetamine-related lysosomal changes, and possibly dopamine handling. But the current evidence is **not strong enough to call ATP6V0C a clearly established addiction gene/protein**, especially in humans. The most accurate summary is: **ATP6V0C is a candidate, indirectly implicated in addiction-related neurobiology, but not yet convincingly established as a core addiction gene.** ([ncbi.nlm.nih.gov](https://www.ncbi.nlm.nih.gov/gene/527)) [S14]

## Cited sources

- **[S1]** ATP6V0C ATPase H+ transporting V0 subunit c [Homo sapiens (human)] - Gene - NCBI — https://www.ncbi.nlm.nih.gov/gene/527
- **[S2]** ATP6V0C ATPase H+ transporting V0 subunit c [Homo sapiens (human)] - Gene - NCBI — https://www.ncbi.nlm.nih.gov/gene/527
- **[S3]** ATP6V0C ATPase H+ transporting V0 subunit c [Homo sapiens (human)] - Gene - NCBI — https://www.ncbi.nlm.nih.gov/gene/527
- **[S4]** Nicotine-induced transcriptional changes and mitochondrial dysfunction in the ventral tegmental area revealed by single-nucleus transcriptomics - ScienceDirect — https://www.sciencedirect.com/science/article/pii/S1673852724002327
- **[S5]** 
            Importance of genetic background for risk of relapse shown in altered prefrontal cortex gene expression during abstinence following chronic alcohol intoxication - PMC
         — https://pmc.ncbi.nlm.nih.gov/articles/PMC3014377/
- **[S6]** 
            Single-Cell RNA-Seq Uncovers Robust Glial Cell Transcriptional Changes in Methamphetamine-Administered Mice - PMC
         — https://pmc.ncbi.nlm.nih.gov/articles/PMC11766323/
- **[S7]** 
            Neurotransmitter release: vacuolar ATPase V0 sector c-subunits in possible gene or cell therapies for Parkinson’s, Alzheimer’s, and psychiatric diseases - PMC
         — https://pmc.ncbi.nlm.nih.gov/articles/PMC10717279/
- **[S8]** Multi-ancestral genome-wide association study of clinically defined nicotine dependence reveals strong genetic correlations with other substance use disorders and health-related traits - PubMed — https://pubmed.ncbi.nlm.nih.gov/40831304/
- **[S9]** Nicotine-induced transcriptional changes and mitochondrial dysfunction in the ventral tegmental area revealed by single-nucleus transcriptomics - ScienceDirect — https://www.sciencedirect.com/science/article/pii/S1673852724002327
- **[S10]** ATP6V0C gene variants were identified in individuals with epilepsy, with or without developmental delay - PubMed — https://pubmed.ncbi.nlm.nih.gov/37161035/?utm_source=openai
- **[S11]** ATP6V0C ATPase H+ transporting V0 subunit c [Homo sapiens (human)] - Gene - NCBI — https://www.ncbi.nlm.nih.gov/gene/527
- **[S12]** Nicotine-induced transcriptional changes and mitochondrial dysfunction in the ventral tegmental area revealed by single-nucleus transcriptomics - ScienceDirect — https://www.sciencedirect.com/science/article/pii/S1673852724002327?utm_source=openai
- **[S13]** ATP6V0C ATPase H+ transporting V0 subunit c [Homo sapiens (human)] - Gene - NCBI — https://www.ncbi.nlm.nih.gov/gene/527
- **[S14]** ATP6V0C ATPase H+ transporting V0 subunit c [Homo sapiens (human)] - Gene - NCBI — https://www.ncbi.nlm.nih.gov/gene/527
