## Supplementary material for "Understanding Substance Dependence: What Differentiates Addictive from Non-Addictive Drugs?": Folder for supplementary files: addictive-genes-claims-first100-010-tmem144-gene-protein-is-implicated-in-addiction-065940a44f53.md

# addictive-genes-claims-first100-010-065940a44f53

**Original claim:** TMEM144 gene/protein is implicated in addiction.

## Claim

TMEM144 gene/protein is implicated in addiction.

## Verdict

Verdict: No Evidence

Current evidence does **not** support TMEM144 as an established addiction gene/protein. In the literature retrieved, TMEM144-specific primary studies concerned **brain-enriched expression, hypothalamic/kisspeptin biology, and aging/Alzheimer’s-related metabolic effects**, not substance use disorder, drug reward, withdrawal, relapse, or other addiction phenotypes. ([ncbi.nlm.nih.gov](https://www.ncbi.nlm.nih.gov/gene/55314)) [S1]

## Confidence

Medium

TMEM144 appears to be a **poorly characterized, understudied** membrane protein. That makes the conclusion reasonably solid for **current direct evidence**, but with some uncertainty because lack of evidence may partly reflect lack of study rather than proof of absence. ([ncbi.nlm.nih.gov](https://www.ncbi.nlm.nih.gov/gene/55314)) [S2]

## Supporting evidence

There are a few **indirect** observations that could make TMEM144 worth future neuroscience study. NCBI currently annotates TMEM144 as a protein-coding gene predicted to enable **carbohydrate transmembrane transporter activity**, with **biased expression in brain**. The Human Protein Atlas likewise classifies TMEM144 as **brain-enriched** and places it in an **oligodendrocyte/myelination expression cluster**. ([ncbi.nlm.nih.gov](https://www.ncbi.nlm.nih.gov/gene/55314)) [S3]

TMEM144 also shows some CNS-related biology outside addiction. In a 2011 mouse hypothalamus study, TMEM144 was described as an “orphan 10-transmembrane transporter” implicated in **GPR54/kisspeptin function**, and its up-regulation in hypothalamus was confirmed by **RNA in situ hybridization and western blot**. ([pubmed.ncbi.nlm.nih.gov](https://pubmed.ncbi.nlm.nih.gov/21527035/)) [S4]

A 2025 study linked TMEM144 to aging/neurodegeneration-related pathways rather than addiction: in human samples and cell models, TMEM144 expression increased with **age** and in **Alzheimer’s disease**, and the authors suggested roles in **glucose transport** and **mitochondrial respiration** via TIMMDC1. ([pubmed.ncbi.nlm.nih.gov](https://pubmed.ncbi.nlm.nih.gov/41026247/)) [S5]

An additional rat study found **Tmem144** elevated across several models of hereditary **internalizing/externalizing behavioral temperament**, and the authors noted that its function remains **unknown**. They also described developmental expression in microglia. These are neurobehavioral observations, but they are **not addiction experiments**. ([pmc.ncbi.nlm.nih.gov](https://pmc.ncbi.nlm.nih.gov/articles/PMC11913853/)) [S6]

More broadly, addiction research increasingly recognizes roles for **glia, oligodendrocytes, microglia, and myelin plasticity**. Recent reviews note that omics studies implicate glial biology in addiction, and a 2024 study showed that **myelin plasticity in the ventral tegmental area is required for opioid reward** in mice. Because TMEM144 is enriched in brain and associated with an oligodendrocyte/myelination cluster, one can make a **plausibility argument** that it could someday matter in addiction-related circuit biology. But that is only an inference from adjacent biology, not direct evidence for this gene. ([pubmed.ncbi.nlm.nih.gov](https://pubmed.ncbi.nlm.nih.gov/38894643/)) [S7]

## Contradicting or limiting evidence

The key limitation is simple: I did **not identify direct evidence** that TMEM144 is associated with addiction in humans or causes addiction-relevant phenotypes in model systems. In the retrieved TMEM144-focused literature, the topics were **hypothalamic signaling**, **aging/Alzheimer’s-related metabolism**, and **non-addiction behavioral temperament**, not addiction itself. ([pubmed.ncbi.nlm.nih.gov](https://pubmed.ncbi.nlm.nih.gov/21527035/)) [S8]

Likewise, broad addiction reviews emphasize many transcriptomic and epigenetic changes in the addicted brain and growing interest in glial mechanisms, but the retrieved reviews did **not present TMEM144 as an established addiction-related molecule**. Instead, they discuss other pathways and genes involved in cocaine, alcohol, opioid, and glial biology. ([pubmed.ncbi.nlm.nih.gov](https://pubmed.ncbi.nlm.nih.gov/25172478/)) [S9]

A recent multi-omic opioid addiction resource curated hundreds of opioid addiction-associated genes from GWAS, RNA-seq, proteomics, and epigenomic data, but TMEM144 was **not found** in the displayed thresholded gene set page for that resource. This does not prove irrelevance, but it argues against TMEM144 being a prominent currently replicated addiction signal. ([geneweaver.org](https://geneweaver.org/viewgenesetdetails/408166)) [S10]

Also important: NCBI’s current summary for TMEM144 is still largely **predictive** rather than experimentally mature, describing predicted transporter activity and brain-biased expression rather than a well-established neural function. That weakens any strong claim that the gene is already “implicated in addiction.” ([ncbi.nlm.nih.gov](https://www.ncbi.nlm.nih.gov/gene/55314)) [S11]

## Mechanistic / biological rationale

If one were to hypothesize an addiction connection, the strongest rationale would be **indirect**. Addiction involves durable changes in reward circuitry, and modern work shows that not only neurons but also **glial cells** and **myelin plasticity** can shape drug reward, withdrawal, and relapse-related circuit adaptations. ([pubmed.ncbi.nlm.nih.gov](https://pubmed.ncbi.nlm.nih.gov/27475769/?utm_source=openai)) [S12]

TMEM144 is expressed in brain, enriched in an **oligodendrocyte/myelination-related cluster**, and has been proposed to participate in **membrane transport** and possibly **cellular energy/metabolic regulation**. Since oligodendrocytes, myelin remodeling, and metabolic support can influence neural circuit timing and reward learning, these features make TMEM144 **biologically conceivable** as a candidate for future addiction studies. ([ncbi.nlm.nih.gov](https://www.ncbi.nlm.nih.gov/gene/55314)) [S13]

However, that rationale remains speculative. There is currently **no direct chain of evidence** showing that TMEM144 alters dopamine signaling, drug self-administration, conditioned place preference, craving, relapse, withdrawal severity, or human substance use disorder risk. Without such evidence, the mechanistic story is hypothesis-generating rather than claim-supporting. ([pubmed.ncbi.nlm.nih.gov](https://pubmed.ncbi.nlm.nih.gov/27475769/?utm_source=openai)) [S14]

## Bottom line

TMEM144 is a **brain-enriched, still poorly characterized membrane protein** with some links to **hypothalamic signaling**, **oligodendrocyte/myelination-related expression**, and **aging/mitochondrial-metabolic biology**. Those features make an addiction role **plausible in principle**, especially because glia and myelin plasticity are relevant to addiction. But as of the evidence retrieved here, there is **no direct human genetic, transcriptomic, proteomic, or experimental behavioral evidence** that TMEM144 is implicated in addiction. The claim is therefore best rated **No Evidence**, not because addiction involvement is impossible, but because it is **currently unsubstantiated**. ([ncbi.nlm.nih.gov](https://www.ncbi.nlm.nih.gov/gene/55314)) [S15]

## Cited sources

- **[S1]** TMEM144 transmembrane protein 144 [Homo sapiens (human)] - Gene - NCBI — https://www.ncbi.nlm.nih.gov/gene/55314
- **[S2]** TMEM144 transmembrane protein 144 [Homo sapiens (human)] - Gene - NCBI — https://www.ncbi.nlm.nih.gov/gene/55314
- **[S3]** TMEM144 transmembrane protein 144 [Homo sapiens (human)] - Gene - NCBI — https://www.ncbi.nlm.nih.gov/gene/55314
- **[S4]** The testosterone-dependent and independent transcriptional networks in the hypothalamus of Gpr54 and Kiss1 knockout male mice are not fully equivalent - PubMed — https://pubmed.ncbi.nlm.nih.gov/21527035/
- **[S5]** Comprehensive study of W06A7.4 and TMEM144 mediated pathways in aging: insights from Caenorhabditis elegans to human - PubMed — https://pubmed.ncbi.nlm.nih.gov/41026247/
- **[S6]** 
            Bioenergetic-related gene expression in the hippocampus predicts internalizing vs. externalizing behavior in an animal model of temperament - PMC
         — https://pmc.ncbi.nlm.nih.gov/articles/PMC11913853/
- **[S7]** How omics is revealing new roles for glia in addiction - PubMed — https://pubmed.ncbi.nlm.nih.gov/38894643/
- **[S8]** The testosterone-dependent and independent transcriptional networks in the hypothalamus of Gpr54 and Kiss1 knockout male mice are not fully equivalent - PubMed — https://pubmed.ncbi.nlm.nih.gov/21527035/
- **[S9]** Gene expression in the addicted brain - PubMed — https://pubmed.ncbi.nlm.nih.gov/25172478/
- **[S10]** GeneSet Details :: GeneWeaver — https://geneweaver.org/viewgenesetdetails/408166
- **[S11]** TMEM144 transmembrane protein 144 [Homo sapiens (human)] - Gene - NCBI — https://www.ncbi.nlm.nih.gov/gene/55314
- **[S12]** Neurobiology of addiction: a neurocircuitry analysis - PubMed — https://pubmed.ncbi.nlm.nih.gov/27475769/?utm_source=openai
- **[S13]** TMEM144 transmembrane protein 144 [Homo sapiens (human)] - Gene - NCBI — https://www.ncbi.nlm.nih.gov/gene/55314
- **[S14]** Neurobiology of addiction: a neurocircuitry analysis - PubMed — https://pubmed.ncbi.nlm.nih.gov/27475769/?utm_source=openai
- **[S15]** TMEM144 transmembrane protein 144 [Homo sapiens (human)] - Gene - NCBI — https://www.ncbi.nlm.nih.gov/gene/55314
