## Supplementary material for "Understanding Substance Dependence: What Differentiates Addictive from Non-Addictive Drugs?": Folder for supplementary files: addictive-genes-claims-first100-011-timp1-gene-protein-is-implicated-in-addiction-86162b0c08e5.md

# addictive-genes-claims-first100-011-86162b0c08e5

**Original claim:** TIMP1 gene/protein is implicated in addiction.

## Claim

TIMP1 gene/protein is implicated in addiction.

## Verdict

Verdict: Likely True

## Confidence

Medium

## Supporting evidence

Multiple **human postmortem transcriptomic studies** link TIMP1 to substance-use disorders. In chronic **cocaine** users, TIMP1 was among the 91 robust differentially expressed genes in ventral midbrain, showing about **1.5-fold upregulation** versus matched controls in a study designed to profile dopaminergic-cell–enriched tissue. ([pmc.ncbi.nlm.nih.gov](https://pmc.ncbi.nlm.nih.gov/articles/PMC4104338/)) [S1]

In **opioid use disorder (OUD)**, RNA-seq of human postmortem **nucleus accumbens (NAc)** and dorsolateral prefrontal cortex found strong enrichment of **neuroinflammation and extracellular-matrix (ECM) remodeling** pathways, and specifically identified **TIMP1 as an OUD-specific hub gene** in NAc co-expression networks. The authors explicitly proposed TIMP1 as part of a mechanistic bridge between opioid-induced neuroinflammation, ECM remodeling, synaptic plasticity, and OUD-related behavioral change. ([pmc.ncbi.nlm.nih.gov](https://pmc.ncbi.nlm.nih.gov/articles/PMC8463497/)) [S2]

There is also evidence in **alcohol use disorder (AUD)**. In a transcriptomic study of withdrawal from chronic alcohol exposure, rat hippocampus showed induction of a STAT3-linked neuroimmune program; the investigators then examined human postmortem hippocampus and reported that **TIMP1 mRNA was significantly increased in AUD** compared with controls, with TIMP1 expression correlating with STAT3-pathway transcripts. ([pmc.ncbi.nlm.nih.gov](https://pmc.ncbi.nlm.nih.gov/articles/PMC8170676/?utm_source=openai)) [S3]

These converging studies matter because they are not just generic injury markers measured in blood: they come from **brain regions central to addiction circuitry**—ventral midbrain, NAc, prefrontal cortex, and hippocampus—using human tissue from individuals with cocaine, opioid, or alcohol use disorders. That pattern supports the narrower claim that TIMP1 is **implicated in addiction-related neurobiology**, not merely in peripheral consequences of substance use. ([pmc.ncbi.nlm.nih.gov](https://pmc.ncbi.nlm.nih.gov/articles/PMC4104338/)) [S4]

## Contradicting or limiting evidence

The main limitation is that the evidence is **mostly associative**, not causal. The cocaine, OUD, and AUD studies above show altered expression or network centrality, but they do **not** directly test whether selective manipulation of TIMP1 changes drug reward, escalation, craving, relapse, or withdrawal severity. In other words, TIMP1 is implicated by transcriptomic and network evidence, but I did not find strong primary evidence that TIMP1 itself has been experimentally validated as a driver of addiction behavior in the way some better-established addiction genes have been. ([pmc.ncbi.nlm.nih.gov](https://pmc.ncbi.nlm.nih.gov/articles/PMC4104338/)) [S5]

A second limitation is specificity. TIMP1 is a well-known **ECM-remodeling, injury-response, and inflammatory-response** gene. In OUD and AUD datasets, TIMP1 appears alongside broader neuroimmune and glial activation programs, so its altered expression could reflect a **reactive response** to chronic drug exposure rather than a primary addiction mechanism. The OUD paper itself frames the TIMP1 model partly as a hypothesis requiring direct future testing. ([pmc.ncbi.nlm.nih.gov](https://pmc.ncbi.nlm.nih.gov/articles/PMC8463497/)) [S6]

Finally, some of the broader mechanistic literature on TIMP1 comes from **non-addiction contexts** such as seizures, neurodegeneration, or injury. Those studies are useful for biological plausibility, but they do not by themselves prove a role in substance-use disorders. ([pubmed.ncbi.nlm.nih.gov](https://pubmed.ncbi.nlm.nih.gov/9151739/?utm_source=openai)) [S7]

## Mechanistic / biological rationale

TIMP1 is biologically plausible as an addiction-relevant molecule because it regulates **matrix metalloproteinases (MMPs)** and thus helps control **extracellular-matrix remodeling**, a process increasingly tied to synaptic plasticity. In the CNS, TIMP1 has been described as a **candidate plasticity gene**: it is induced in neurons and astrocytes after seizures and other strong activity-dependent states, consistent with responsiveness to neural activity. ([pubmed.ncbi.nlm.nih.gov](https://pubmed.ncbi.nlm.nih.gov/9151739/?utm_source=openai)) [S8]

Primary studies also show TIMP1 can directly affect neuronal structure and function. In cultured cortical neurons, TIMP1 altered **neurite outgrowth and morphology**, consistent with a role in structural plasticity. More recently, exogenous TIMP-1 in an Alzheimer’s mouse model enhanced **Akt/BDNF signaling** and promoted **postsynaptic long-term potentiation (LTP)**, showing that TIMP1 can modulate synaptic-function pathways highly relevant to learning-related adaptations. ([pmc.ncbi.nlm.nih.gov](https://pmc.ncbi.nlm.nih.gov/articles/PMC2788270/?utm_source=openai)) [S9]

That matters for addiction because persistent drug seeking depends on long-lasting synaptic and circuit remodeling in regions such as the **NAc, prefrontal cortex, ventral midbrain, and hippocampus**. If TIMP1 changes the balance of MMP activity and ECM structure, it could influence dendritic remodeling, synaptic stabilization, and neuroimmune-ECM interactions that support drug-associated memories and relapse vulnerability. The human OUD study specifically linked TIMP1-centered networks to **ECM remodeling and inflammatory signaling** in the NAc, which fits this mechanistic model. ([pmc.ncbi.nlm.nih.gov](https://pmc.ncbi.nlm.nih.gov/articles/PMC8463497/)) [S10]

There is also some behavioral relevance outside addiction per se. **Timp1 knockout mice** show increased anxiety-like behavior, indicating that endogenous TIMP1 can shape affect-related brain function. That does not establish addiction causality, but it supports the idea that TIMP1 is not merely a passive matrix inhibitor; it can influence behaviorally relevant neural states. ([pmc.ncbi.nlm.nih.gov](https://pmc.ncbi.nlm.nih.gov/articles/PMC11178759/?utm_source=openai)) [S11]

## Bottom line

The claim is **reasonable and supported at an association/mechanism level**: TIMP1 is repeatedly altered or network-central in **human brain studies of cocaine use, opioid use disorder, and alcohol use disorder**, and its known biology in **ECM remodeling, neuroinflammation, neurite structure, and synaptic plasticity** makes that connection plausible. ([pmc.ncbi.nlm.nih.gov](https://pmc.ncbi.nlm.nih.gov/articles/PMC4104338/)) [S12]

However, the evidence is **not yet strong enough to say TIMP1 is a proven causal addiction gene/protein**. The current literature more strongly supports: **“TIMP1 is implicated in addiction-related brain adaptations”** than **“TIMP1 directly causes addiction behavior.”** ([pmc.ncbi.nlm.nih.gov](https://pmc.ncbi.nlm.nih.gov/articles/PMC4104338/)) [S13]

## Cited sources

- **[S1]** 
            A Molecular Profile of Cocaine Abuse Includes the Differential Expression of Genes that Regulate Transcription, Chromatin, and Dopamine Cell Phenotype - PMC
         — https://pmc.ncbi.nlm.nih.gov/articles/PMC4104338/
- **[S2]** 
            Transcriptional alterations in dorsolateral prefrontal cortex and nucleus accumbens implicate neuroinflammation and synaptic remodeling in opioid use disorder - PMC
         — https://pmc.ncbi.nlm.nih.gov/articles/PMC8463497/
- **[S3]** Transcriptomics identifies STAT3 as a key regulator of hippocampal gene expression and anhedonia during withdrawal from chronic alcohol exposure - PMC — https://pmc.ncbi.nlm.nih.gov/articles/PMC8170676/?utm_source=openai
- **[S4]** 
            A Molecular Profile of Cocaine Abuse Includes the Differential Expression of Genes that Regulate Transcription, Chromatin, and Dopamine Cell Phenotype - PMC
         — https://pmc.ncbi.nlm.nih.gov/articles/PMC4104338/
- **[S5]** 
            A Molecular Profile of Cocaine Abuse Includes the Differential Expression of Genes that Regulate Transcription, Chromatin, and Dopamine Cell Phenotype - PMC
         — https://pmc.ncbi.nlm.nih.gov/articles/PMC4104338/
- **[S6]** 
            Transcriptional alterations in dorsolateral prefrontal cortex and nucleus accumbens implicate neuroinflammation and synaptic remodeling in opioid use disorder - PMC
         — https://pmc.ncbi.nlm.nih.gov/articles/PMC8463497/
- **[S7]** Tissue inhibitor of metalloproteinases-1 (TIMP-1) is differentially induced in neurons and astrocytes after seizures: evidence for developmental, immediate early gene, and lesion response - PubMed — https://pubmed.ncbi.nlm.nih.gov/9151739/?utm_source=openai
- **[S8]** Tissue inhibitor of metalloproteinases-1 (TIMP-1) is differentially induced in neurons and astrocytes after seizures: evidence for developmental, immediate early gene, and lesion response - PubMed — https://pubmed.ncbi.nlm.nih.gov/9151739/?utm_source=openai
- **[S9]** A New Role for TIMP-1 in Modulating Neurite Outgrowth and Morphology of Cortical Neurons - PMC — https://pmc.ncbi.nlm.nih.gov/articles/PMC2788270/?utm_source=openai
- **[S10]** 
            Transcriptional alterations in dorsolateral prefrontal cortex and nucleus accumbens implicate neuroinflammation and synaptic remodeling in opioid use disorder - PMC
         — https://pmc.ncbi.nlm.nih.gov/articles/PMC8463497/
- **[S11]** Timp1 Deletion Induces Anxiety-like Behavior in Mice - PMC — https://pmc.ncbi.nlm.nih.gov/articles/PMC11178759/?utm_source=openai
- **[S12]** 
            A Molecular Profile of Cocaine Abuse Includes the Differential Expression of Genes that Regulate Transcription, Chromatin, and Dopamine Cell Phenotype - PMC
         — https://pmc.ncbi.nlm.nih.gov/articles/PMC4104338/
- **[S13]** 
            A Molecular Profile of Cocaine Abuse Includes the Differential Expression of Genes that Regulate Transcription, Chromatin, and Dopamine Cell Phenotype - PMC
         — https://pmc.ncbi.nlm.nih.gov/articles/PMC4104338/
