## Supplementary material for "Understanding Substance Dependence: What Differentiates Addictive from Non-Addictive Drugs?": Folder for supplementary files: addictive-genes-claims-first100-012-cartpt-gene-protein-is-implicated-in-addiction-16b464cadd5a.md

# addictive-genes-claims-first100-012-16b464cadd5a

**Original claim:** CARTPT gene/protein is implicated in addiction.

## Claim

CARTPT gene/protein is implicated in addiction. ([pubmed.ncbi.nlm.nih.gov](https://pubmed.ncbi.nlm.nih.gov/8647455/)) [S1]

## Verdict

Verdict: Likely True

The claim is **likely true**, with the strongest support coming from **preclinical addiction neuroscience** rather than definitive human genetics or clinical data. CARTPT encodes CART peptide, a molecule originally identified because its transcript is induced by psychostimulants in striatum, and multiple animal studies show that changing CART signaling changes cocaine-, amphetamine-, and alcohol-related behaviors. Human evidence exists, but it is narrower and less consistent than the animal literature. ([pubmed.ncbi.nlm.nih.gov](https://pubmed.ncbi.nlm.nih.gov/8647455/)) [S2]

## Confidence

High

There is a substantial, coherent body of evidence that CART/CARTPT participates in drug-reward and drug-seeking circuitry, especially for **psychostimulants** and to a lesser extent **alcohol-related behaviors**. Confidence is not maximal because much of the evidence is from rodents, some findings are circuit- and sex-dependent, and human genetic evidence is mixed. ([pubmed.ncbi.nlm.nih.gov](https://pubmed.ncbi.nlm.nih.gov/16099925/)) [S3]

## Supporting evidence

CARTPT is biologically tied to addiction from its discovery onward. The human CART gene was characterized after a rat transcript, “cocaine- and amphetamine-regulated transcript,” had been identified as being transcriptionally regulated in striatum after acute psychomotor stimulant exposure. Later work in rat nucleus accumbens showed CART expression is regulated by cAMP/PKA/CREB signaling and can be increased under cocaine-related conditions, placing it squarely in a pathway long linked to reward-related neuroadaptation. ([pubmed.ncbi.nlm.nih.gov](https://pubmed.ncbi.nlm.nih.gov/8647455/)) [S4]

There is strong **animal behavioral evidence** that CART signaling modifies cocaine-related phenotypes. In rats, intra-accumbens CART peptide reduced cocaine-induced locomotor activity and also blunted dopamine-induced locomotion, suggesting CART acts within nucleus accumbens circuitry relevant to drug reward. In a separate self-administration study, intra-accumbens CART peptide reduced cocaine self-administration on a progressive-ratio schedule, consistent with lowered motivation to obtain cocaine. Local knockdown of CART in nucleus accumbens by shRNA had the opposite effect, increasing cocaine-induced locomotor activity. ([pubmed.ncbi.nlm.nih.gov](https://pubmed.ncbi.nlm.nih.gov/14551286/?utm_source=openai)) [S5]

**Genetic loss-of-function models** also support involvement. In CART knockout mice, acute amphetamine responses were attenuated, repeated amphetamine produced little sensitization, amphetamine conditioned place preference was weaker, and intravenous cocaine self-administration was reduced versus wild type. Those data indicate CART peptides influence the locomotor and motivational properties of psychostimulants, even though the direction of effect is not always simple. ([pubmed.ncbi.nlm.nih.gov](https://pubmed.ncbi.nlm.nih.gov/16099925/)) [S6]

More recent work strengthens the case that CARTPT is not just correlated with drug exposure, but can be **causally relevant to addiction-related plasticity**. After cocaine self-administration and abstinence, Cartpt becomes activated in mouse nucleus accumbens along with epigenetic changes at its promoter, and Nr4a1-related regulation of Cartpt was linked to suppression of cocaine-induced behavior. A 2025 CRISPR epigenetic editing study then directly manipulated repressive chromatin marks at the Cartpt promoter in vivo and showed that reducing Cartpt expression in D1 neurons increased cocaine conditioned place preference and altered extinction-related behavior. ([pubmed.ncbi.nlm.nih.gov](https://pubmed.ncbi.nlm.nih.gov/31980629/)) [S7]

Human evidence, while less extensive, points in the same general direction. A postmortem study of nucleus accumbens from human cocaine abusers reported **upregulated CART gene expression** by microarray and RT-PCR. For alcohol-related phenotypes, a Korean study found the CARTPT intronic variant rs2239670 associated with alcoholism in men. These are not definitive proof of causation, but they do support relevance of CARTPT to human addictive disorders. ([pmc.ncbi.nlm.nih.gov](https://pmc.ncbi.nlm.nih.gov/articles/PMC2215309/)) [S8]

The addiction link is not limited to stimulants. In alcohol models, CART knockout mice consumed and preferred less ethanol than wild-type mice, and this could not be explained by altered total intake or ethanol metabolism. In rats, CART signaling in the central amygdala was activated during yohimbine-induced reinstatement of alcohol seeking, and local CART antibody infusion into the central amygdala attenuated reinstatement. Newer work also found sex-dependent roles in binge drinking, with central amygdala CART neutralization reducing binge-like alcohol intake in females. ([pmc.ncbi.nlm.nih.gov](https://pmc.ncbi.nlm.nih.gov/articles/PMC4148419/)) [S9]

## Contradicting or limiting evidence

The main limitation is that **human evidence is modest and inconsistent**. A case-control study of individuals of African descent found no significant association between tested CARTPT SNPs/haplotypes and cocaine dependence, concluding that their data did not support CARTPT as a major susceptibility gene for cocaine dependence, while also noting sample-size and coverage limitations. So, CARTPT is not currently established as a robust, replicated human addiction-risk gene in the way some stronger GWAS loci are. ([pmc.ncbi.nlm.nih.gov](https://pmc.ncbi.nlm.nih.gov/articles/PMC2507865/)) [S10]

There is also **directional complexity** in the animal data. For example, local CART infusion into nucleus accumbens tends to suppress cocaine-related behaviors, yet global CART knockout can also reduce cocaine self-administration and ethanol intake. That apparent tension suggests CART’s role is likely **circuit-specific, state-dependent, and possibly compensatory** rather than simply “pro-addiction” or “anti-addiction.” Alcohol studies likewise show sex-dependent effects, including stronger reductions in ethanol intake in female CART knockout mice and female-specific effects of central amygdala CART manipulations on binge drinking. ([pubmed.ncbi.nlm.nih.gov](https://pubmed.ncbi.nlm.nih.gov/14551286/?utm_source=openai)) [S11]

Another limitation is mechanistic incompleteness. CART has long been treated as a neuropeptide system relevant to reward and addiction, but its cognate receptor biology remains unsettled. GPR160 was proposed as a CART receptor, yet later binding studies and reviews concluded that specific CART-GPR160 binding was not demonstrated and that the true CART receptor remains unresolved. That does not negate the addiction evidence, but it does limit mechanistic certainty and translational drug development. ([pmc.ncbi.nlm.nih.gov](https://pmc.ncbi.nlm.nih.gov/articles/PMC10953447/)) [S12]

## Mechanistic / biological rationale

A plausible biological rationale exists for CARTPT involvement in addiction because CART peptide is embedded in **mesolimbic reward circuitry** and responds to psychostimulant-relevant signaling. CART expression is regulated by cAMP/PKA/CREB in nucleus accumbens, a canonical addiction-related pathway, and cocaine can augment this signaling context. CART peptide in nucleus accumbens can blunt dopamine- and cocaine-driven locomotor effects, consistent with a modulatory or homeostatic role in excessive dopaminergic activation. ([pubmed.ncbi.nlm.nih.gov](https://pubmed.ncbi.nlm.nih.gov/16322355/)) [S13]

Mechanistically, several studies suggest CART can shape addiction-related plasticity by modifying intracellular signaling downstream of dopamine. In a cocaine sensitization model, CART peptide reduced cocaine-enhanced CaMKIIα-related signaling and downstream pCREB changes. In abstinence models, Cartpt is re-activated after prolonged cocaine withdrawal with promoter-level chromatin remodeling, and manipulating that chromatin state changes cocaine preference behavior. Together, these data fit a model in which CARTPT participates in **homeostatic adaptation within reward circuits** during both drug exposure and abstinence. ([pubmed.ncbi.nlm.nih.gov](https://pubmed.ncbi.nlm.nih.gov/29313985/)) [S14]

For alcohol-related behaviors, the rationale expands beyond dopamine alone to include **stress and relapse circuitry**, especially the central amygdala. CART neuron activation during stress-induced reinstatement of alcohol seeking and the effect of central amygdala CART blockade on reinstatement support a role in relapse-like behavior. Female-specific binge-drinking effects additionally suggest that CART can modulate addiction-related behavior through sensory/affective processes such as bitter-taste sensitivity and amygdala signaling, not just classic stimulant reward. ([pubmed.ncbi.nlm.nih.gov](https://pubmed.ncbi.nlm.nih.gov/32826981/)) [S15]

## Bottom line

Yes—**CARTPT/CART is implicated in addiction**, especially in **preclinical models of psychostimulant addiction** and, secondarily, in **alcohol-related behaviors**. The evidence includes drug-induced regulation of Cartpt expression, effects of CART peptide infusion or knockdown on cocaine reward/seeking phenotypes, knockout phenotypes affecting stimulant and alcohol behaviors, human postmortem expression changes, and a limited positive human alcoholism association. ([pubmed.ncbi.nlm.nih.gov](https://pubmed.ncbi.nlm.nih.gov/8647455/)) [S16]

The caveat is that CARTPT is **not yet established as a major, consistently replicated human addiction gene**, and its effects appear **context-, circuit-, and sex-dependent** rather than uniformly causal in one direction. So the most accurate interpretation is: **CARTPT/CART is very plausibly involved in addiction neurobiology, but the strongest evidence is mechanistic and animal-based, not definitive human clinical proof.** ([pmc.ncbi.nlm.nih.gov](https://pmc.ncbi.nlm.nih.gov/articles/PMC2507865/)) [S17]

## Cited sources

- **[S1]** Characterization of the human cDNA and genomic DNA encoding CART: a cocaine- and amphetamine-regulated transcript - PubMed — https://pubmed.ncbi.nlm.nih.gov/8647455/
- **[S2]** Characterization of the human cDNA and genomic DNA encoding CART: a cocaine- and amphetamine-regulated transcript - PubMed — https://pubmed.ncbi.nlm.nih.gov/8647455/
- **[S3]** Cocaine- and amphetamine-regulated transcript (CART) peptides modulate the locomotor and motivational properties of psychostimulants - PubMed — https://pubmed.ncbi.nlm.nih.gov/16099925/
- **[S4]** Characterization of the human cDNA and genomic DNA encoding CART: a cocaine- and amphetamine-regulated transcript - PubMed — https://pubmed.ncbi.nlm.nih.gov/8647455/
- **[S5]** Intra-accumbal injection of CART (cocaine-amphetamine regulated transcript) peptide reduces cocaine-induced locomotor activity - PubMed — https://pubmed.ncbi.nlm.nih.gov/14551286/?utm_source=openai
- **[S6]** Cocaine- and amphetamine-regulated transcript (CART) peptides modulate the locomotor and motivational properties of psychostimulants - PubMed — https://pubmed.ncbi.nlm.nih.gov/16099925/
- **[S7]** Nr4a1 suppresses cocaine-induced behavior via epigenetic regulation of homeostatic target genes - PubMed — https://pubmed.ncbi.nlm.nih.gov/31980629/
- **[S8]** 
            Gene expression profile of the nucleus accumbens of human cocaine abusers: evidence for dysregulation of myelin - PMC
         — https://pmc.ncbi.nlm.nih.gov/articles/PMC2215309/
- **[S9]** 
            Reduced ethanol consumption and preference in cocaine- and amphetamine-regulated transcript (CART) knockout mice - PMC
         — https://pmc.ncbi.nlm.nih.gov/articles/PMC4148419/
- **[S10]** 
            Genetic variants in the cocaine- and amphetamine-regulated transcript gene (CARTPT) and cocaine dependence - PMC
         — https://pmc.ncbi.nlm.nih.gov/articles/PMC2507865/
- **[S11]** Intra-accumbal injection of CART (cocaine-amphetamine regulated transcript) peptide reduces cocaine-induced locomotor activity - PubMed — https://pubmed.ncbi.nlm.nih.gov/14551286/?utm_source=openai
- **[S12]** 
            Orphan peptide and G protein‐coupled receptor signalling in alcohol use disorder - PMC
         — https://pmc.ncbi.nlm.nih.gov/articles/PMC10953447/
- **[S13]** Cocaine-amphetamine-regulated transcript expression in the rat nucleus accumbens is regulated by adenylyl cyclase and the cyclic adenosine 5'-monophosphate/protein kinase a second messenger system - PubMed — https://pubmed.ncbi.nlm.nih.gov/16322355/
- **[S14]** Cocaine- and amphetamine-regulated transcript peptide in the nucleus accumbens shell inhibits cocaine-induced locomotor sensitization to transient over-expression of α-Ca2+ /calmodulin-dependent protein kinase II - PubMed — https://pubmed.ncbi.nlm.nih.gov/29313985/
- **[S15]** Cocaine and amphetamine regulated transcript (CART) signalling in the central nucleus of the amygdala modulates stress-induced alcohol seeking - PubMed — https://pubmed.ncbi.nlm.nih.gov/32826981/
- **[S16]** Characterization of the human cDNA and genomic DNA encoding CART: a cocaine- and amphetamine-regulated transcript - PubMed — https://pubmed.ncbi.nlm.nih.gov/8647455/
- **[S17]** 
            Genetic variants in the cocaine- and amphetamine-regulated transcript gene (CARTPT) and cocaine dependence - PMC
         — https://pmc.ncbi.nlm.nih.gov/articles/PMC2507865/
