## Supplementary material for "Understanding Substance Dependence: What Differentiates Addictive from Non-Addictive Drugs?": Folder for supplementary files: addictive-genes-claims-first100-013-cox6a1-gene-protein-is-implicated-in-addiction-7aab99e0538a.md

# addictive-genes-claims-first100-013-7aab99e0538a

**Original claim:** COX6A1 gene/protein is implicated in addiction.

## Claim

COX6A1 gene/protein is implicated in addiction.

## Verdict

Verdict: Mixed Evidence

There is **some** evidence linking COX6A1 to addiction-relevant biology, but it is **limited, indirect, and mostly from alcohol-exposure animal omics studies** rather than direct causal addiction experiments or robust human genetic data. COX6A1 is therefore **not an established addiction gene**, though it may participate in mitochondrial/metabolic changes that accompany substance exposure, especially ethanol. ([ncbi.nlm.nih.gov](https://www.ncbi.nlm.nih.gov/gtr/genes/1337/?utm_source=openai)) [S1]

## Confidence

Medium

The confidence is medium because the literature does show repeated COX6A1 signal changes in some addiction-relevant contexts, especially alcohol-exposed rodent brain, but the evidence base is narrow and lacks strong human or mechanistic-causality studies specific to addiction. ([pmc.ncbi.nlm.nih.gov](https://pmc.ncbi.nlm.nih.gov/articles/PMC4516281/)) [S2]

## Supporting evidence

COX6A1 encodes a nuclear subunit of **mitochondrial complex IV (cytochrome c oxidase)**, the terminal enzyme of the respiratory chain. NCBI summarizes that this gene encodes the ubiquitous non-muscle isoform of subunit VIa, and an early brain study showed COX6A1 expression in brain tissue and downregulation after neuronal activity deprivation in monkey visual neurons, indicating that it is both neuronal and activity-responsive. That makes it biologically plausible that COX6A1 could change in addiction-relevant circuits, where neuronal activity and energy demand are heavily remodeled. ([ncbi.nlm.nih.gov](https://www.ncbi.nlm.nih.gov/gtr/genes/1337/?utm_source=openai)) [S3]

The strongest direct support comes from **ethanol-related rodent transcriptomics**. In inbred alcohol-preferring rats, ethanol self-administration altered nucleus accumbens gene expression, and **Cox6a1 was increased** (reported fold change about 1.13, p=0.002). Because the nucleus accumbens is a core reward/addiction region, this is a relevant association between COX6A1 expression and alcohol-taking behavior. ([pmc.ncbi.nlm.nih.gov](https://pmc.ncbi.nlm.nih.gov/articles/PMC4516281/)) [S4]

A second alcohol study in adolescent alcohol-preferring rats found **region-specific Cox6a1 changes** after binge-like drinking: decreased expression in ventral hippocampus and increased expression in medial prefrontal cortex. In the medial prefrontal cortex, Cox6a1 also appeared in significantly altered **oxidative phosphorylation** and **mitochondrial dysfunction** pathways. These findings support the idea that COX6A1 participates in alcohol-induced remodeling of brain energy metabolism in circuits tied to executive control and reinforcement. ([pmc.ncbi.nlm.nih.gov](https://pmc.ncbi.nlm.nih.gov/articles/PMC5851801/)) [S5]

Proteomic evidence points in the same direction. In rats exposed to chronic ethanol vapor, a brain-region proteomics study reported **Cox6a1 protein increased to 117% in prefrontal cortex**. That is still not proof of causation, but it shows that COX6A1 changes are detectable at the protein level after prolonged ethanol exposure, not only at the transcript level. ([pmc.ncbi.nlm.nih.gov](https://pmc.ncbi.nlm.nih.gov/articles/PMC3129429/?utm_source=openai)) [S6]

The broader addiction literature also strengthens plausibility. Reviews on addiction biology report that **mitochondria are altered by drugs of abuse**, including alcohol, cocaine, amphetamine, and opioids, and that cocaine exposure changes mitochondrial gene/protein networks in reward regions such as the nucleus accumbens, ventral tegmental area, and prefrontal cortex. Reviews focused on alcohol likewise describe mitochondrial changes as relevant to tolerance, dependence, withdrawal, and alcohol-related brain pathology. Since COX6A1 is a complex IV subunit in oxidative phosphorylation, it fits naturally within this mitochondria-centered framework. ([pubmed.ncbi.nlm.nih.gov](https://pubmed.ncbi.nlm.nih.gov/33626141/?utm_source=openai)) [S7]

## Contradicting or limiting evidence

The main limitation is that the evidence is **not addiction-specific enough**. The cited COX6A1 findings mostly come from transcriptomic or proteomic surveys after alcohol exposure. Such studies show correlation with exposure state, but they do **not** demonstrate that changing COX6A1 itself changes drug seeking, relapse, withdrawal severity, or reinforcement. I did not identify strong studies in which COX6A1 was genetically manipulated in addiction models to test behavior directly. ([pmc.ncbi.nlm.nih.gov](https://pmc.ncbi.nlm.nih.gov/articles/PMC4516281/)) [S8]

There is also a lack of convincing **human genetic evidence** for addiction. In the sources reviewed, COX6A1 is well established in **neuromuscular mitochondrial disease**—including recessive Charcot-Marie-Tooth phenotypes and broader mitochondrial dysfunction—but that is different from showing involvement in substance use disorder risk. The disease literature around COX6A1 is therefore real, but it does not by itself support addiction causality. ([pmc.ncbi.nlm.nih.gov](https://pmc.ncbi.nlm.nih.gov/articles/PMC4157141/?utm_source=openai)) [S9]

Another limitation is specificity across substances. I found the clearest evidence for **alcohol**, but not similarly strong direct COX6A1 evidence for cocaine, nicotine, opioids, or cannabis. Meanwhile, mitochondrial alterations are extremely common responses to toxic, metabolic, inflammatory, and neuronal stress states, so COX6A1 changes could simply reflect a **general bioenergetic response** rather than a specific addiction mechanism. Its known regulation by neuronal activity supports that concern. ([pubmed.ncbi.nlm.nih.gov](https://pubmed.ncbi.nlm.nih.gov/10773445/?utm_source=openai)) [S10]

## Mechanistic / biological rationale

Mechanistically, the case is plausible. COX6A1 is part of **complex IV**, which helps maintain the proton gradient that drives ATP synthesis. Reward-circuit neurons require tightly regulated mitochondrial ATP production for neurotransmission, ion pumping, synaptic plasticity, and structural remodeling. Because addictive drugs reshape firing patterns and plasticity in brain reward circuits, genes involved in oxidative phosphorylation are reasonable candidates for downstream adaptation. ([ncbi.nlm.nih.gov](https://www.ncbi.nlm.nih.gov/gtr/genes/1337/?utm_source=openai)) [S11]

Alcohol and other abused substances also promote **oxidative stress, altered mitochondrial trafficking/dynamics, and apoptotic signaling**. A review on alcohol and mitochondrial trafficking explicitly frames mitochondria as relevant to alcohol addiction and other drug addictions, while broader cocaine literature links mitochondrial changes to addiction-related neuroplasticity. If COX6A1 expression shifts, that could influence cellular energetic capacity or redox tone in these circuits. ([pubmed.ncbi.nlm.nih.gov](https://pubmed.ncbi.nlm.nih.gov/33626141/?utm_source=openai)) [S12]

There is also some direct mechanistic evidence that COX6A1 can affect cell survival pathways: one study identified COX6A1 as a suppressor of Bax-mediated cell death and reported reduced ROS generation and downstream apoptotic signaling when COX6A1 was expressed. That supports a potential role in stress resilience under toxic exposure. Still, this is **general mitochondrial/cell-death biology**, not addiction-specific proof. ([pubmed.ncbi.nlm.nih.gov](https://pubmed.ncbi.nlm.nih.gov/18549809/?utm_source=openai)) [S13]

So biologically, the best interpretation is: **COX6A1 is a plausible mediator of mitochondrial adaptations in addiction-relevant brain tissue**, but the current literature does not show it to be a validated driver of addictive behavior. ([pmc.ncbi.nlm.nih.gov](https://pmc.ncbi.nlm.nih.gov/articles/PMC4516281/)) [S14]

## Bottom line

COX6A1 is **not strongly established as an addiction gene/protein**, but there is **credible, limited evidence** that it participates in addiction-relevant mitochondrial changes—especially in **alcohol-exposed rodent brain**. The claim is therefore best rated **Mixed Evidence**: plausible and partially supported, but currently too indirect and sparse for a stronger conclusion. ([pmc.ncbi.nlm.nih.gov](https://pmc.ncbi.nlm.nih.gov/articles/PMC4516281/)) [S15]

## Cited sources

- **[S1]** COX6A1 cytochrome c oxidase subunit 6A1 - NIH Genetic Testing Registry (GTR) - NCBI — https://www.ncbi.nlm.nih.gov/gtr/genes/1337/?utm_source=openai
- **[S2]** 
            Differential gene expression in the nucleus accumbens with ethanol self-administration in inbred alcohol-preferring rats - PMC
         — https://pmc.ncbi.nlm.nih.gov/articles/PMC4516281/
- **[S3]** COX6A1 cytochrome c oxidase subunit 6A1 - NIH Genetic Testing Registry (GTR) - NCBI — https://www.ncbi.nlm.nih.gov/gtr/genes/1337/?utm_source=openai
- **[S4]** 
            Differential gene expression in the nucleus accumbens with ethanol self-administration in inbred alcohol-preferring rats - PMC
         — https://pmc.ncbi.nlm.nih.gov/articles/PMC4516281/
- **[S5]** 
            Gene expression changes in the ventral hippocampus and medial prefrontal cortex of adolescent alcohol-preferring (P) rats following binge-like-alcohol drinking - PMC
         — https://pmc.ncbi.nlm.nih.gov/articles/PMC5851801/
- **[S6]** Ethanol-Induced Changes in the Expression of Proteins Related to Neurotransmission and Metabolism in Different Regions of the Rat Brain - PMC — https://pmc.ncbi.nlm.nih.gov/articles/PMC3129429/?utm_source=openai
- **[S7]** The role of mitochondria in cocaine addiction - PubMed — https://pubmed.ncbi.nlm.nih.gov/33626141/?utm_source=openai
- **[S8]** 
            Differential gene expression in the nucleus accumbens with ethanol self-administration in inbred alcohol-preferring rats - PMC
         — https://pmc.ncbi.nlm.nih.gov/articles/PMC4516281/
- **[S9]** A Mutation of COX6A1 Causes a Recessive Axonal or Mixed Form of Charcot-Marie-Tooth Disease - PMC — https://pmc.ncbi.nlm.nih.gov/articles/PMC4157141/?utm_source=openai
- **[S10]** Human COX6A1 gene: promoter analysis, cDNA isolation and expression in the monkey brain - PubMed — https://pubmed.ncbi.nlm.nih.gov/10773445/?utm_source=openai
- **[S11]** COX6A1 cytochrome c oxidase subunit 6A1 - NIH Genetic Testing Registry (GTR) - NCBI — https://www.ncbi.nlm.nih.gov/gtr/genes/1337/?utm_source=openai
- **[S12]** The role of mitochondria in cocaine addiction - PubMed — https://pubmed.ncbi.nlm.nih.gov/33626141/?utm_source=openai
- **[S13]** Identification of cytochrome c oxidase subunit 6A1 as a suppressor of Bax-induced cell death by yeast-based functional screening - PubMed — https://pubmed.ncbi.nlm.nih.gov/18549809/?utm_source=openai
- **[S14]** 
            Differential gene expression in the nucleus accumbens with ethanol self-administration in inbred alcohol-preferring rats - PMC
         — https://pmc.ncbi.nlm.nih.gov/articles/PMC4516281/
- **[S15]** 
            Differential gene expression in the nucleus accumbens with ethanol self-administration in inbred alcohol-preferring rats - PMC
         — https://pmc.ncbi.nlm.nih.gov/articles/PMC4516281/
