## Supplementary material for "Understanding Substance Dependence: What Differentiates Addictive from Non-Addictive Drugs?": Folder for supplementary files: addictive-genes-claims-first100-014-tmem130-gene-protein-is-implicated-in-addiction-d72a155ecda5.md

# addictive-genes-claims-first100-014-d72a155ecda5

**Original claim:** TMEM130 gene/protein is implicated in addiction.

## Claim

TMEM130 gene/protein is implicated in addiction.

## Verdict

Verdict: No Evidence

Based on the scholarly and authoritative sources I could verify, there is **no direct evidence** that TMEM130 is implicated in addiction or substance use disorders. TMEM130 is a real, brain-expressed, neuron-associated gene/protein, but I did **not** find a human GWAS signal, replicated association study, addiction-focused postmortem transcriptomic paper, or causal animal-behavior study tying TMEM130 specifically to addiction phenotypes. Major current addiction-genetics papers instead highlight other genes/loci such as **OPRM1, FURIN, PPP6C, RABEPK, NCAM1, KCNN1,** and **PDE4B**. ([ncbi.nlm.nih.gov](https://www.ncbi.nlm.nih.gov/gene/222865?utm_source=openai)) [S1]

## Confidence

Medium

Confidence is **medium** because “no evidence” is always harder to prove than “some evidence”: a supplementary dataset, preprint, or niche omics paper could have mentioned TMEM130 without my searches surfacing it prominently. Still, the retrievable literature and major reference resources I checked were sparse and did not yield addiction-specific evidence for TMEM130. ([informatics.jax.org](https://informatics.jax.org/marker/MGI%3A3607706)) [S2]

## Supporting evidence

The strongest argument **for** the claim is only **indirect biological plausibility**. TMEM130 is a protein-coding gene with **biased/tissue-enhanced expression in brain**, and the Human Protein Atlas reports it is **detected across all examined human brain regions** and clusters with **neuronal/synaptic-function** genes. In mouse, Tmem130 was identified as a **neuron-specific** transcript with in situ expression in **hippocampus and cortex**. These findings make TMEM130 at least a plausible CNS-relevant molecule. ([ncbi.nlm.nih.gov](https://www.ncbi.nlm.nih.gov/gene/222865?utm_source=openai)) [S3]

There is also a weak mechanistic reason one might have suspected relevance: NCBI/Alliance and HPA place TMEM130 in the **Golgi apparatus**, and conserved-domain resources indicate it contains **PKD/KLD-related domains** within the **PKAT** family architecture. A neuronal Golgi/membrane protein could, in principle, affect trafficking or processing of synaptic proteins. But that is an inference, not addiction evidence. ([proteinatlas.org](https://www.proteinatlas.org/ENSG00000166448-TMEM130)) [S4]

## Contradicting or limiting evidence

The main problem for the claim is the absence of **direct addiction data**. When I checked the GWAS Catalog gene page for **TMEM130**, the page did not show populated associations/studies/traits entries, which argues against TMEM130 being a recognized mapped GWAS gene at present. Meanwhile, large contemporary addiction GWAS/meta-analyses identify other loci—such as **OPRM1, PPP6C, FURIN, RABEPK, NCAM1, KCNN1,** and **PDE4B**—rather than TMEM130. ([ebi.ac.uk](https://www.ebi.ac.uk/gwas/genes/TMEM130)) [S5]

The TMEM130 literature I could retrieve is also **not addiction-centered**. The most visible functional papers are about **cancer methylation/migration** in nasopharyngeal, gastric, breast, and cervical cancer. One 2022 paper explicitly framed itself as the **first** study to investigate TMEM130 expression/function in nasopharyngeal carcinoma, underscoring how limited the functional literature still is. That overall pattern is not what you would expect for a gene already implicated in addiction. ([pubmed.ncbi.nlm.nih.gov](https://pubmed.ncbi.nlm.nih.gov/34958002/?utm_source=openai)) [S6]

A further limitation is that even basic functional annotation remains thin. Mouse Genome Informatics states that, after literature review, there is **no experimental evidence** supporting a **molecular function** or **biological process** annotation for Tmem130. That does not rule out an addiction role, but it does mean the biology is too underdeveloped to support a confident implication in addiction. ([informatics.jax.org](https://informatics.jax.org/marker/MGI%3A3607706)) [S7]

## Mechanistic / biological rationale

Addiction-relevant genes usually gain support through one or more of the following: human genetic association, altered expression in addiction-relevant brain circuits, or causal animal studies affecting reward, reinforcement, withdrawal, relapse, or compulsive intake. For TMEM130, I found only the **background prerequisites** for possible relevance—brain expression, neuronal enrichment, Golgi localization, and membrane-protein/domain architecture—not the addiction-specific evidence that would elevate it from “plausible CNS gene” to “implicated in addiction.” ([proteinatlas.org](https://www.proteinatlas.org/ENSG00000166448-TMEM130/brain)) [S8]

So the best mechanistic summary is: **TMEM130 could conceivably participate in neuronal membrane trafficking or synaptic homeostasis**, because it is neuron-enriched and Golgi-associated. However, I found **no direct evidence** linking it to dopamine, opioid, GABA, glutamate, nicotine-receptor, or other canonical addiction pathways, and no data showing that manipulating TMEM130 changes addiction-related behavior. That gap is decisive here. ([proteinatlas.org](https://www.proteinatlas.org/ENSG00000166448-TMEM130)) [S9]

## Bottom line

TMEM130 is a **brain-expressed, neuron-associated, Golgi-localized** protein with **sparse functional characterization**, so a nervous-system role is plausible. But on the evidence currently retrievable from major scholarly and genetics resources, **TMEM130 is not established as an addiction gene/protein**, and I found **no direct evidence** supporting the claim that it is implicated in addiction. ([ncbi.nlm.nih.gov](https://www.ncbi.nlm.nih.gov/gene/222865?utm_source=openai)) [S10]

If you want, I can do a second-pass review specifically for **alcohol**, **opioid**, **nicotine**, or **cocaine** datasets/supplements to see whether TMEM130 ever appears as a low-confidence omics hit.

## Cited sources

- **[S1]** TMEM130 transmembrane protein 130 [Homo sapiens (human)] - Gene - NCBI — https://www.ncbi.nlm.nih.gov/gene/222865?utm_source=openai
- **[S2]** Tmem130 MGI Mouse Gene Detail - MGI:3607706 - transmembrane protein 130 — https://informatics.jax.org/marker/MGI%3A3607706
- **[S3]** TMEM130 transmembrane protein 130 [Homo sapiens (human)] - Gene - NCBI — https://www.ncbi.nlm.nih.gov/gene/222865?utm_source=openai
- **[S4]** TMEM130 protein expression summary - The Human Protein Atlas — https://www.proteinatlas.org/ENSG00000166448-TMEM130
- **[S5]** GWAS Catalog — https://www.ebi.ac.uk/gwas/genes/TMEM130
- **[S6]** TMEM130 regulates cell migration through DNA methylation in nasopharyngeal carcinoma - PubMed — https://pubmed.ncbi.nlm.nih.gov/34958002/?utm_source=openai
- **[S7]** Tmem130 MGI Mouse Gene Detail - MGI:3607706 - transmembrane protein 130 — https://informatics.jax.org/marker/MGI%3A3607706
- **[S8]** Brain tissue expression of TMEM130 - Summary - The Human Protein Atlas — https://www.proteinatlas.org/ENSG00000166448-TMEM130/brain
- **[S9]** TMEM130 protein expression summary - The Human Protein Atlas — https://www.proteinatlas.org/ENSG00000166448-TMEM130
- **[S10]** TMEM130 transmembrane protein 130 [Homo sapiens (human)] - Gene - NCBI — https://www.ncbi.nlm.nih.gov/gene/222865?utm_source=openai
