## Supplementary material for "Understanding Substance Dependence: What Differentiates Addictive from Non-Addictive Drugs?": Folder for supplementary files: addictive-genes-claims-first100-015-gng7-gene-protein-is-implicated-in-addiction-3c837df727e3.md

# addictive-genes-claims-first100-015-3c837df727e3

**Original claim:** GNG7 gene/protein is implicated in addiction.

## Claim

GNG7 gene/protein is implicated in addiction.

## Verdict

Verdict: Likely True

## Confidence

Medium

## Supporting evidence

GNG7 encodes the G-protein γ7 subunit, which is enriched in striatal neurons and dendrites and is predicted to participate in GPCR signaling and regulation of adenylate cyclase. The striatum is a core node for reward learning, psychostimulant responses, and habit formation, so a gene with a selective role in this circuitry is biologically plausible as an addiction-related factor. ([ncbi.nlm.nih.gov](https://www.ncbi.nlm.nih.gov/gene/58979?utm_source=openai)) [S1]

There is direct **preclinical** evidence that manipulating Gng7 changes behavioral responses to drugs of abuse. In a 2021 mouse study, conditional deletion of Gng7 in striatal medium spiny neuron subtypes disrupted assembly of the Gαolfβ2γ7 complex and altered in vivo responses to psychostimulants and opioids: loss in A2AR/D2R-expressing neurons produced hyperlocomotion and enhanced amphetamine response, while loss in D1R-expressing neurons selectively affected the acute locomotor-enhancing effects of morphine. The authors explicitly frame γ7 as a key determinant of striatal cAMP signaling in disorders including drug addiction. ([pubmed.ncbi.nlm.nih.gov](https://pubmed.ncbi.nlm.nih.gov/34544837/)) [S2]

A newer 2024 study strengthens the addiction link by testing cocaine-related phenotypes more directly. That study found that although **global** Gng7 deletion did not alter acute cocaine locomotion or overall cocaine sensitization, **cell-type-specific** loss mattered: D1R-pathway Gng7 loss impaired establishment of cocaine sensitization in high-responder male mice, while D2R/A2AR-pathway loss increased acute cocaine locomotor sensitivity in males. The same study also showed increased dendritic spine density in striatal medium spiny neurons after Gng7 deletion, connecting Gng7 to structural plasticity relevant to addiction-related neuroadaptations. ([frontiersin.org](https://www.frontiersin.org/journals/neuroanatomy/articles/10.3389/fnana.2024.1394659/full)) [S3]

Mechanistic papers preceding those behavioral studies support that this is not an incidental association. Gγ7 is required for proper assembly of a striatal Gαolfβ2γ7 heterotrimer that mediates D1 dopamine receptor and A2A adenosine receptor signaling, and this signaling module has been described as critical for pathways controlling locomotion and reward. Earlier work also showed that loss of Gng7 reduces striatal Gαolf levels, lowers cAMP production, and alters behavior. ([pubmed.ncbi.nlm.nih.gov](https://pubmed.ncbi.nlm.nih.gov/20639202/?utm_source=openai)) [S4]

There is also **alcohol-related animal evidence**. A transcriptomic study in alcohol-preferring versus non-preferring rats identified Gng7 among genes that “may constitute novel participants of brain ethanol response,” and broader expression studies across selectively bred high- vs low-ethanol-consuming rat lines also reported Gng7 differences in addiction-relevant regions. This does not prove causality, but it is consistent with a role in substance-use phenotypes beyond cocaine. ([sciencedirect.com](https://www.sciencedirect.com/science/article/abs/pii/S0091305715300757?utm_source=openai)) [S5]

Human evidence exists, but it is weaker and more indirect. A 2025 twin/Mendelian-randomization epigenetics study reported that hypermethylation at a GNG7-linked CpG (cg13078421) may increase smoking liability, with brain regulatory enrichment among loci showing reverse or bidirectional effects on smoking. The paper explicitly notes prior model-organism evidence linking GNG7 differential expression to addiction-related phenotypes. ([pmc.ncbi.nlm.nih.gov](https://pmc.ncbi.nlm.nih.gov/articles/PMC11213072/)) [S6]

## Contradicting or limiting evidence

The biggest limitation is that the strongest evidence is **from rodent models**, mostly using locomotor sensitization and acute drug-response paradigms rather than full addiction endpoints such as escalation, compulsive use, relapse, or self-administration. Those phenotypes are informative but are not equivalent to human substance use disorder. ([frontiersin.org](https://www.frontiersin.org/journals/neuroanatomy/articles/10.3389/fnana.2024.1394659/full)) [S7]

Even within the cocaine literature, the signal is nuanced rather than uniform. The 2024 study found **no effect of global Gng7 deletion** on acute cocaine locomotor response or overall cocaine-induced sensitization; the effect emerged mainly in specific neuronal subtypes and in a subset of male “high responder” mice. Females did not show the same genotype effect in that sensitization paradigm. That argues for a context-dependent role, not a broad universal addiction gene effect. ([frontiersin.org](https://www.frontiersin.org/journals/neuroanatomy/articles/10.3389/fnana.2024.1394659/full)) [S8]

The human data are still sparse. The smoking study was based on **peripheral blood DNA methylation**, not brain tissue, and the reverse-effect signal for GNG7 was modest and requires replication. I did not find strong, replicated evidence that common GNG7 variants are among the major genome-wide significant loci repeatedly established for alcohol dependence/AUD in large GWAS reviews, which instead emphasize other genes and pathways. ([pmc.ncbi.nlm.nih.gov](https://pmc.ncbi.nlm.nih.gov/articles/PMC11213072/)) [S9]

Some supporting findings are also inferential. For example, alcohol-preferring rat transcriptomics and ethanol-response candidate-gene studies nominate Gng7, but these are association-style animal data rather than direct intervention studies showing that changing Gng7 alters alcohol-seeking behavior. ([sciencedirect.com](https://www.sciencedirect.com/science/article/abs/pii/S0091305715300757?utm_source=openai)) [S10]

## Mechanistic / biological rationale

The mechanistic case is fairly coherent. GNG7 helps form the striatal Gαolfβ2γ7 complex, which couples key receptors involved in reward circuitry—especially D1 dopamine and A2A adenosine receptors—to adenylyl cyclase/cAMP signaling. Because dopamine- and adenosine-dependent cAMP signaling in direct and indirect pathway medium spiny neurons is central to drug-induced plasticity, altering GNG7 abundance or localization can plausibly reshape responses to addictive drugs. ([pmc.ncbi.nlm.nih.gov](https://pmc.ncbi.nlm.nih.gov/articles/PMC2943273/?utm_source=openai)) [S11]

The 2024 work adds a plausible structural-plasticity mechanism: Gng7 transcript 1 is enriched in dendritic neuropil, miRNA-sensitive, and linked to local γ7 protein production; Gng7 loss increased dendritic spine density in medium spiny neurons. Since repeated psychostimulant exposure is known to remodel spines in these neurons, GNG7 could influence addiction-relevant behavior by modulating synaptic organization and pathway balance rather than simply changing acute receptor signaling. ([frontiersin.org](https://www.frontiersin.org/journals/neuroanatomy/articles/10.3389/fnana.2024.1394659/full)) [S12]

This fits with the broader striatal literature showing that Gng7/Gαolf loss reduces cAMP signaling and can mute or redirect behavioral responses to psychostimulants and opioids. In other words, GNG7 is biologically positioned more as a **modulator of addiction-related neuroadaptation and drug responsiveness** than as a definitively proven primary human addiction-risk gene. ([pubmed.ncbi.nlm.nih.gov](https://pubmed.ncbi.nlm.nih.gov/12488442/?utm_source=openai)) [S13]

## Bottom line

The claim is **likely true**, but mainly on the basis of **preclinical neuroscience** rather than definitive human genetics. GNG7 has credible mechanistic and animal-model evidence linking it to addiction-relevant circuitry and to behavioral responses to cocaine, amphetamine, morphine, and possibly alcohol-related phenotypes. However, the effects are cell-type-, sex-, and paradigm-dependent, and strong replicated human evidence for GNG7 as a major addiction gene is still limited. ([frontiersin.org](https://www.frontiersin.org/journals/neuroanatomy/articles/10.3389/fnana.2024.1394659/full)) [S14]

## Cited sources

- **[S1]** Gng7 G protein subunit gamma 7 [Rattus norvegicus (Norway rat)] - Gene - NCBI — https://www.ncbi.nlm.nih.gov/gene/58979?utm_source=openai
- **[S2]** Selective Manipulation of G-Protein γ7 Subunit in Mice Provides New Insights into Striatal Control of Motor Behavior - PubMed — https://pubmed.ncbi.nlm.nih.gov/34544837/
- **[S3]** Frontiers | Post-transcriptional regulation and subcellular localization of G-protein γ7 subunit: implications for striatal function and behavioral responses to cocaine — https://www.frontiersin.org/journals/neuroanatomy/articles/10.3389/fnana.2024.1394659/full
- **[S4]** Adenosine A2A receptor signaling and golf assembly show a specific requirement for the gamma7 subtype in the striatum - PubMed — https://pubmed.ncbi.nlm.nih.gov/20639202/?utm_source=openai
- **[S5]** Novel candidate genes for alcoholism — transcriptomic analysis of prefrontal medial cortex, hippocampus and nucleus accumbens of Warsaw alcohol-preferring and non-preferring rats - ScienceDirect — https://www.sciencedirect.com/science/article/abs/pii/S0091305715300757?utm_source=openai
- **[S6]** 
            Unidirectional and Bidirectional Causation between Smoking and Blood DNA Methylation: Evidence from Twin-based Mendelian Randomisation - PMC
         — https://pmc.ncbi.nlm.nih.gov/articles/PMC11213072/
- **[S7]** Frontiers | Post-transcriptional regulation and subcellular localization of G-protein γ7 subunit: implications for striatal function and behavioral responses to cocaine — https://www.frontiersin.org/journals/neuroanatomy/articles/10.3389/fnana.2024.1394659/full
- **[S8]** Frontiers | Post-transcriptional regulation and subcellular localization of G-protein γ7 subunit: implications for striatal function and behavioral responses to cocaine — https://www.frontiersin.org/journals/neuroanatomy/articles/10.3389/fnana.2024.1394659/full
- **[S9]** 
            Unidirectional and Bidirectional Causation between Smoking and Blood DNA Methylation: Evidence from Twin-based Mendelian Randomisation - PMC
         — https://pmc.ncbi.nlm.nih.gov/articles/PMC11213072/
- **[S10]** Novel candidate genes for alcoholism — transcriptomic analysis of prefrontal medial cortex, hippocampus and nucleus accumbens of Warsaw alcohol-preferring and non-preferring rats - ScienceDirect — https://www.sciencedirect.com/science/article/abs/pii/S0091305715300757?utm_source=openai
- **[S11]** Adenosine A2A Receptor Signaling and Golf Assembly Show a Specific Requirement for the γ7 Subtype in the Striatum - PMC — https://pmc.ncbi.nlm.nih.gov/articles/PMC2943273/?utm_source=openai
- **[S12]** Frontiers | Post-transcriptional regulation and subcellular localization of G-protein γ7 subunit: implications for striatal function and behavioral responses to cocaine — https://www.frontiersin.org/journals/neuroanatomy/articles/10.3389/fnana.2024.1394659/full
- **[S13]** Loss of G protein gamma 7 alters behavior and reduces striatal alpha(olf) level and cAMP production - PubMed — https://pubmed.ncbi.nlm.nih.gov/12488442/?utm_source=openai
- **[S14]** Frontiers | Post-transcriptional regulation and subcellular localization of G-protein γ7 subunit: implications for striatal function and behavioral responses to cocaine — https://www.frontiersin.org/journals/neuroanatomy/articles/10.3389/fnana.2024.1394659/full
