## Supplementary material for "Understanding Substance Dependence: What Differentiates Addictive from Non-Addictive Drugs?": Folder for supplementary files: addictive-genes-claims-first100-016-slc6a4-gene-protein-is-implicated-in-addiction-6c854e4ce2cd.md

# addictive-genes-claims-first100-016-6c854e4ce2cd

**Original claim:** SLC6A4 gene/protein is implicated in addiction.

## Claim

SLC6A4, the gene encoding the serotonin transporter (SERT/5-HTT), is implicated in addiction. SLC6A4 encodes the membrane transporter that clears serotonin from the synaptic cleft; notably, the protein is also a known target of psychostimulants including cocaine and amphetamines. ([ncbi.nlm.nih.gov](https://www.ncbi.nlm.nih.gov/gene/6532)) [S1]

## Verdict

Verdict: Likely True

At a **protein/pathway level**, this claim is well grounded: SERT is directly engaged by cocaine and other psychostimulants, and serotonergic signaling is broadly involved in drug reinforcement, withdrawal, relapse vulnerability, and addiction-related behaviors across multiple substances. At a **human inherited-risk level**, however, the evidence is less clean: candidate-gene studies and some meta-analyses suggest association, but results are heterogeneous and not uniformly replicated, and stronger modern genome-wide evidence is limited. So the fairest reading is that **SLC6A4/SERT is implicated in addiction biology, but not established as a robust universal addiction-risk gene across all substances and populations**. ([ncbi.nlm.nih.gov](https://www.ncbi.nlm.nih.gov/gene/6532)) [S2]

## Confidence

Medium

Confidence is **medium** because the mechanistic and preclinical evidence is substantial, and some human association data support involvement, but the human genetic literature is inconsistent by phenotype, ancestry, and substance, with several null or weak findings for major outcomes such as alcohol dependence. ([pubmed.ncbi.nlm.nih.gov](https://pubmed.ncbi.nlm.nih.gov/24769172/)) [S3]

## Supporting evidence

First, the biological plausibility is strong. NCBI’s reviewed gene summary states that SLC6A4 encodes the serotonin transporter that terminates serotonergic signaling by reuptake, and that this transporter is a target of psychomotor stimulants such as amphetamines and cocaine. In a PNAS knock-in study, altering SERT to abolish high-affinity cocaine recognition left baseline serotonin transport intact but markedly reduced cocaine sensitivity, supporting a direct role for SERT in at least some cocaine effects. ([ncbi.nlm.nih.gov](https://www.ncbi.nlm.nih.gov/gene/6532)) [S4]

Second, broader addiction-neuroscience reviews place serotonin centrally in addiction-relevant processes. A review in *Behavioural Brain Research* concluded that the serotonergic system has a “crucial, but distinct involvement” in the establishment of drug-use behaviors and in the transition to and maintenance of addiction across cocaine, amphetamine, methamphetamine, MDMA, heroin, cannabis, alcohol, and nicotine. A separate review on serotonin and addiction vulnerability summarizes evidence that drug exposure and withdrawal alter serotonergic tone, and that these changes may contribute to dysphoria, relapse, and compulsive use. ([pubmed.ncbi.nlm.nih.gov](https://pubmed.ncbi.nlm.nih.gov/24769172/)) [S5]

Third, there is supportive—though not definitive—human genetic evidence. A 2021 umbrella review of more than 150 meta-analyses on substance use disorders reported that **SLC6A4** was among the genes associated with two or more substance-use disorders, and singled it out as one of the few with odds ratios above two in the reviewed candidate-gene literature. A multi-study meta-analysis focused on SLC6A4 concluded that association signals varied by substance but were seen across alcohol, heroin, cocaine, and methamphetamine dependence/abuse datasets. ([pubmed.ncbi.nlm.nih.gov](https://pubmed.ncbi.nlm.nih.gov/33556390/)) [S6]

Fourth, several human studies link SLC6A4 variation to addiction-related phenotypes, even when not proving overall disease susceptibility. In treatment-seeking alcohol-dependent individuals, the rs1042173 TT genotype was associated with greater drinking intensity and lower SLC6A4 mRNA/protein expression in vitro. Another study found preliminary evidence that this same functional variant modulated cue-induced alcohol craving. In a cohort of opioid users, the low-expression SLC6A4 promoter genotype was not associated with categorical opioid dependence, but it was associated with heavier cocaine exposure and more rapid escalation of cocaine use. ([pmc.ncbi.nlm.nih.gov](https://pmc.ncbi.nlm.nih.gov/articles/PMC2657929/)) [S7]

Fifth, animal data show that manipulating SERT changes addiction-relevant behaviors. In SERT knockout rats, cocaine produced stronger conditioned place preference and greater locomotor activation than in wild-type rats, suggesting enhanced sensitivity to cocaine reward. In another rat study, inherited serotonin-transporter down-regulation was described as associated with increased vulnerability to cocaine addiction and altered BDNF signaling in the amygdala after long-access cocaine self-administration. For alcohol, SERT knockout mice still learned ethanol self-administration, but showed lower breakpoints and lower unconstrained demand for ethanol than wild-type or heterozygous mice, indicating altered motivation for alcohol. ([link.springer.com](https://link.springer.com/article/10.1007/s00213-008-1212-x)) [S8]

## Contradicting or limiting evidence

The main limitation is that **human genetic association evidence is inconsistent**. The best direct counterexample here is a 2015 meta-analysis of 25 studies (8,885 participants) on the classic SLC6A4 promoter polymorphism 5-HTTLPR and alcohol dependence, which found **no overall association** under dominant, recessive, or additive models, and no significant modification by sex, ancestry, psychiatric comorbidity, publication year, or diagnostic criteria. ([pubmed.ncbi.nlm.nih.gov](https://pubmed.ncbi.nlm.nih.gov/25710844/)) [S9]

Even supportive meta-analytic literature emphasizes heterogeneity. The large multi-cultural SLC6A4 meta-analysis concluded that associations vary by substance and ancestry; it reported the short allele as the risk allele in European, Asian, and Mexican populations, but the long allele as the risk allele in African populations. That same paper also noted heterogeneity between alcohol, heroin, cocaine, and methamphetamine datasets, especially for cocaine. This makes any simple global statement about “the SLC6A4 addiction allele” untenable. ([pmc.ncbi.nlm.nih.gov](https://pmc.ncbi.nlm.nih.gov/articles/PMC3717550/)) [S10]

Modern psychiatric genetics also tempers enthusiasm for older candidate-gene claims. A recent review of substance-use-disorder genetics emphasizes that many pharmacogenomic and candidate-gene studies have had limited success, and that current genome-wide data for SUDs are still not sufficiently informative for clinical risk prediction. Notably, the strongest replicated GWAS signals for specific SUDs tend to involve other loci, such as nicotinic receptor genes for nicotine dependence, rather than SLC6A4. ([pmc.ncbi.nlm.nih.gov](https://pmc.ncbi.nlm.nih.gov/articles/PMC8477224/)) [S11]

Some positive SLC6A4 findings are also modest or preliminary. The opioid/cocaine cohort reported only a nominally significant association with heavy cocaine exposure, not categorical opioid dependence, and the cocaine methylation study used a small sample and explicitly called for replication. Those results support possible involvement, but not firm causality. ([pubmed.ncbi.nlm.nih.gov](https://pubmed.ncbi.nlm.nih.gov/33658787/)) [S12]

## Mechanistic / biological rationale

Mechanistically, SLC6A4 is a sensible addiction candidate because SERT regulates extracellular serotonin, and serotonin shapes reward processing, stress reactivity, impulsivity, and aversive withdrawal states. When SERT activity is reduced—by genotype, epigenetic change, or drug blockade—synaptic serotonin signaling changes, which can alter neural responses to drug cues and reinforcement. ([ncbi.nlm.nih.gov](https://www.ncbi.nlm.nih.gov/gene/6532)) [S13]

For psychostimulants, the rationale is especially direct: cocaine and related stimulants interact with monoamine transporters, including SERT. The PNAS knock-in model showing reduced cocaine sensitivity after selectively changing SERT’s cocaine recognition site provides unusually strong mechanistic evidence that SERT contributes to cocaine’s in vivo effects, even if dopamine transporters remain more central for many reinforcing effects. ([pubmed.ncbi.nlm.nih.gov](https://pubmed.ncbi.nlm.nih.gov/21282638/)) [S14]

More broadly, serotonin-system adaptations appear to track phases of addiction. Reviews describe acute drug exposure as often increasing extracellular serotonin, whereas chronic exposure and withdrawal can produce serotonergic dysregulation or reduced serotonergic output. That pattern could contribute to negative affect, stress sensitivity, relapse risk, and compulsive drug-seeking. Because SLC6A4 is the principal serotonin reuptake mechanism, it sits at a biologically plausible control point for these processes. ([pubmed.ncbi.nlm.nih.gov](https://pubmed.ncbi.nlm.nih.gov/24769172/)) [S15]

## Bottom line

**Yes—SLC6A4/SERT is reasonably described as implicated in addiction, especially in serotonergic modulation of addiction-related behavior and in psychostimulant pharmacology.** But the strongest support is **mechanistic, pharmacologic, and preclinical**, not a clean, universally replicated human genetic-risk signal. If the intended meaning is “SLC6A4 is part of addiction biology,” the claim is **likely true**. If the intended meaning is “SLC6A4 variants are a firmly established major cause of addiction risk in humans,” the evidence is **too mixed for that stronger claim**. ([ncbi.nlm.nih.gov](https://www.ncbi.nlm.nih.gov/gene/6532)) [S16]

## Cited sources

- **[S1]** SLC6A4 solute carrier family 6 member 4 [Homo sapiens (human)] - Gene - NCBI — https://www.ncbi.nlm.nih.gov/gene/6532
- **[S2]** SLC6A4 solute carrier family 6 member 4 [Homo sapiens (human)] - Gene - NCBI — https://www.ncbi.nlm.nih.gov/gene/6532
- **[S3]** The role of serotonin in drug use and addiction - PubMed — https://pubmed.ncbi.nlm.nih.gov/24769172/
- **[S4]** SLC6A4 solute carrier family 6 member 4 [Homo sapiens (human)] - Gene - NCBI — https://www.ncbi.nlm.nih.gov/gene/6532
- **[S5]** The role of serotonin in drug use and addiction - PubMed — https://pubmed.ncbi.nlm.nih.gov/24769172/
- **[S6]** Molecular genetics of substance use disorders: An umbrella review - PubMed — https://pubmed.ncbi.nlm.nih.gov/33556390/
- **[S7]** 
            Characterization of a functional polymorphism in the 3’UTR of SLC6A4 and its association with drinking intensity - PMC
         — https://pmc.ncbi.nlm.nih.gov/articles/PMC2657929/
- **[S8]** Adaptations in pre- and postsynaptic 5-HT1A receptor function and cocaine supersensitivity in serotonin transporter knockout rats | Psychopharmacology | Springer Nature Link — https://link.springer.com/article/10.1007/s00213-008-1212-x
- **[S9]** A meta-analysis of the associations between the SLC6A4 promoter polymorphism (5HTTLPR) and the risk for alcohol dependence - PubMed — https://pubmed.ncbi.nlm.nih.gov/25710844/
- **[S10]** 
            Multi-Cultural Association of the Serotonin Transporter Gene (SLC6A4) with Substance Use Disorder - PMC
         — https://pmc.ncbi.nlm.nih.gov/articles/PMC3717550/
- **[S11]** 
            Genetics of substance use disorders: a review - PMC
         — https://pmc.ncbi.nlm.nih.gov/articles/PMC8477224/
- **[S12]** Association of Serotonin Transporter (SERT) Polymorphisms with Opioid Dependence and Dimensional Aspects of Cocaine Use in a Caucasian Cohort of Opioid Users - PubMed — https://pubmed.ncbi.nlm.nih.gov/33658787/
- **[S13]** SLC6A4 solute carrier family 6 member 4 [Homo sapiens (human)] - Gene - NCBI — https://www.ncbi.nlm.nih.gov/gene/6532
- **[S14]** Transgenic elimination of high-affinity antidepressant and cocaine sensitivity in the presynaptic serotonin transporter - PubMed — https://pubmed.ncbi.nlm.nih.gov/21282638/
- **[S15]** The role of serotonin in drug use and addiction - PubMed — https://pubmed.ncbi.nlm.nih.gov/24769172/
- **[S16]** SLC6A4 solute carrier family 6 member 4 [Homo sapiens (human)] - Gene - NCBI — https://www.ncbi.nlm.nih.gov/gene/6532
