## Supplementary material for "Understanding Substance Dependence: What Differentiates Addictive from Non-Addictive Drugs?": Folder for supplementary files: addictive-genes-claims-first100-017-padi2-gene-protein-is-implicated-in-addiction-96b0e8d05c07.md

# addictive-genes-claims-first100-017-96b0e8d05c07

**Original claim:** PADI2 gene/protein is implicated in addiction.

## Claim

PADI2 gene/protein is implicated in addiction.

## Verdict

Verdict: No Evidence

At present, I did **not** find direct evidence that **PADI2** is an addiction gene/protein in the usual sense used in addiction biology: replicated human genetic association with substance use disorder, consistent differential expression in addiction-relevant human brain studies, or animal experiments showing that manipulating PADI2 changes drug reward, self-administration, dependence, withdrawal, craving, or relapse. ([pubmed.ncbi.nlm.nih.gov](https://pubmed.ncbi.nlm.nih.gov/36207451/)) [S1]

## Confidence

Medium. The indirect biology is plausible, but the **addiction-specific** evidence base appears sparse to absent. Large addiction GWAS and recent multiomic/transcriptomic studies point to other genes and pathways, and targeted PADI2 literature retrieval mainly surfaced non-addiction topics. ([nature.com](https://www.nature.com/articles/s41591-023-02653-5)) [S2]

## Supporting evidence

The strongest argument **for plausibility** is that PADI2 is a real CNS-expressed enzyme with functions relevant to neural homeostasis. NCBI describes PADI2 as a calcium-dependent peptidyl arginine deiminase that citrullinates proteins including **myelin basic protein (MBP)**, is expressed in brain, and has been discussed more in neurodegenerative/demyelinating conditions than in addiction. Independent CNS-focused work shows Padi2 is upregulated during oligodendrocyte differentiation, enriched in mature oligodendrocytes, and contributes to efficient myelination in mice. ([ncbi.nlm.nih.gov](https://www.ncbi.nlm.nih.gov/gene/11240/)) [S3]

PADI2 is also broadly present across CNS regions relevant to motivation and reward circuitry. A CNS expression study found detectable PADI2 transcripts in multiple human brain regions, including white matter, basal ganglia, midbrain, thalamus, and pons, with especially high expression in oligodendrocyte-lineage cells. That makes it biologically conceivable that altered PADI2 activity could affect brain circuits that are also involved in addiction. ([pmc.ncbi.nlm.nih.gov](https://pmc.ncbi.nlm.nih.gov/articles/PMC9207393/)) [S4]

More broadly, addiction research increasingly implicates **oligodendrocytes, myelin plasticity, and white matter integrity**. In mice, activity-driven myelin plasticity in the ventral tegmental area was reported to be required for opioid reward, with dopaminergic activity increasing oligodendrocyte precursor proliferation and later myelination in the VTA. In another mouse study, adolescent alcohol exposure impaired myelination and reduced oligodendrocyte-lineage cells in medial prefrontal cortex. Human imaging studies in cocaine and heroin use disorders also report widespread white-matter abnormalities consistent with demyelination/lower axonal packing. A review of cocaine-related neuroglial changes likewise summarizes altered myelin-gene expression and oligodendrocyte disturbances in substance use contexts. ([pubmed.ncbi.nlm.nih.gov](https://pubmed.ncbi.nlm.nih.gov/38839962/)) [S5]

Taken together, these findings support an **indirect** chain of reasoning: if PADI2 helps regulate oligodendrocyte differentiation and myelin biology, and myelin/oligodendrocyte plasticity contributes to reward-circuit adaptation, then PADI2 could in principle influence addiction-relevant neurobiology. But this remains a hypothesis, not direct evidence. ([pmc.ncbi.nlm.nih.gov](https://pmc.ncbi.nlm.nih.gov/articles/PMC6486480/)) [S6]

## Contradicting or limiting evidence

The main limitation is simple: major addiction genetics papers do **not** currently point to PADI2. In a very large problematic alcohol use GWAS, the reported significant signals included loci such as **ADH1B** and **BDNF**; a search within that article did not identify PADI2. ([nature.com](https://www.nature.com/articles/s41591-023-02653-5)) [S7]

For opioid addiction, a multi-trait GWAS highlighted **OPRM1** most strongly and reported gene-based associations with **PPP6C** and **FURIN**; PADI2 was not found in that paper either. Likewise, a cannabis use disorder GWAS implicated **CHRNA2**, not PADI2. ([pubmed.ncbi.nlm.nih.gov](https://pubmed.ncbi.nlm.nih.gov/36207451/)) [S8]

Recent human brain omics also do not provide a clear PADI2 signal. A multiomic network analysis of opioid addiction in dorsolateral prefrontal cortex emphasized pathways involving **BDNF/ERK/Akt**, glutamate transport, receptor subunits, and immediate-early genes; PADI2 was not identified in the paper text. A separate comparative transcriptomic analysis of alcohol and opioid use disorders in dorsolateral prefrontal cortex emphasized inflammatory, extracellular matrix, and synaptic pathways, again without a PADI2 signal in the article text. ([pmc.ncbi.nlm.nih.gov](https://pmc.ncbi.nlm.nih.gov/articles/PMC12119972/)) [S9]

Targeted retrieval of PADI2-focused literature also argues against an established addiction link. Searches surfaced PADI2 papers on cancer, smoking-related lung citrullination, and testis development rather than substance use disorder neurobiology. Notably, the smoking paper concerns **airway/lung PAD2 expression in smokers**, which is a peripheral exposure/inflammation observation and should not be conflated with addiction genetics or brain mechanisms. ([pubmed.ncbi.nlm.nih.gov](https://pubmed.ncbi.nlm.nih.gov/32629995/?utm_source=openai)) [S10]

## Mechanistic / biological rationale

Biologically, PADI2 is a plausible **modulator of neural homeostasis**, especially through citrullination of MBP and regulation of oligodendrocyte differentiation. Since adaptive myelination can tune circuit timing and reward-related learning, a myelin regulator like PADI2 could hypothetically affect susceptibility to drug-induced neuroadaptation. This is particularly plausible in regions where PADI2 is expressed and where myelin changes have been linked to substance exposure, such as midbrain and prefrontal systems. ([ncbi.nlm.nih.gov](https://www.ncbi.nlm.nih.gov/gene/11240/)) [S11]

However, that rationale remains **one step removed** from the claim. A mechanistic bridge is not the same as implication in addiction. To support the claim directly, one would want at least one of the following: replicated human association data, addiction-relevant postmortem brain dysregulation, or causal animal studies showing that PADI2 gain/loss alters drug-seeking or dependence phenotypes. I did not find that level of evidence. ([pubmed.ncbi.nlm.nih.gov](https://pubmed.ncbi.nlm.nih.gov/36207451/)) [S12]

## Bottom line

**PADI2 is a brain/myelin-related enzyme with a plausible indirect connection to addiction-relevant neural plasticity, but I found no direct evidence that it is specifically implicated in addiction.** The literature currently supports PADI2 far more strongly in **oligodendrocyte/myelin biology** and in **non-addiction diseases** than in substance use disorders. If you want, I can next do a stricter subtype check—e.g., **alcohol**, **opioid**, **nicotine**, **cocaine**, or **behavioral addiction** specifically. ([pmc.ncbi.nlm.nih.gov](https://pmc.ncbi.nlm.nih.gov/articles/PMC6486480/)) [S13]

## Cited sources

- **[S1]** Multi-trait genome-wide association study of opioid addiction: OPRM1 and beyond - PubMed — https://pubmed.ncbi.nlm.nih.gov/36207451/
- **[S2]** Multi-ancestry study of the genetics of problematic alcohol use in over 1 million individuals | Nature Medicine — https://www.nature.com/articles/s41591-023-02653-5
- **[S3]** PADI2 peptidyl arginine deiminase 2 [Homo sapiens (human)] - Gene - NCBI — https://www.ncbi.nlm.nih.gov/gene/11240/
- **[S4]** 
            Peptidylarginine Deiminase 2 Autoantibodies Are Linked to Less Severe Disease in Multiple Sclerosis and Post-treatment Lyme Disease - PMC
         — https://pmc.ncbi.nlm.nih.gov/articles/PMC9207393/
- **[S5]** Myelin plasticity in the ventral tegmental area is required for opioid reward - PubMed — https://pubmed.ncbi.nlm.nih.gov/38839962/
- **[S6]** 
            PAD2-Mediated Citrullination Contributes to Efficient Oligodendrocyte Differentiation and Myelination - PMC
         — https://pmc.ncbi.nlm.nih.gov/articles/PMC6486480/
- **[S7]** Multi-ancestry study of the genetics of problematic alcohol use in over 1 million individuals | Nature Medicine — https://www.nature.com/articles/s41591-023-02653-5
- **[S8]** Multi-trait genome-wide association study of opioid addiction: OPRM1 and beyond - PubMed — https://pubmed.ncbi.nlm.nih.gov/36207451/
- **[S9]** 
            Multiomic Network Analysis Identifies Dysregulated Neurobiological Pathways in Opioid Addiction - PMC
         — https://pmc.ncbi.nlm.nih.gov/articles/PMC12119972/
- **[S10]** Putative Roles for Peptidylarginine Deiminases in COVID-19 - PubMed — https://pubmed.ncbi.nlm.nih.gov/32629995/?utm_source=openai
- **[S11]** PADI2 peptidyl arginine deiminase 2 [Homo sapiens (human)] - Gene - NCBI — https://www.ncbi.nlm.nih.gov/gene/11240/
- **[S12]** Multi-trait genome-wide association study of opioid addiction: OPRM1 and beyond - PubMed — https://pubmed.ncbi.nlm.nih.gov/36207451/
- **[S13]** 
            PAD2-Mediated Citrullination Contributes to Efficient Oligodendrocyte Differentiation and Myelination - PMC
         — https://pmc.ncbi.nlm.nih.gov/articles/PMC6486480/
