## Supplementary material for "Understanding Substance Dependence: What Differentiates Addictive from Non-Addictive Drugs?": Folder for supplementary files: addictive-genes-claims-first100-018-gnb1-gene-protein-is-implicated-in-addiction-1f404ae089cf.md

# addictive-genes-claims-first100-018-1f404ae089cf

**Original claim:** GNB1 gene/protein is implicated in addiction.

## Claim

GNB1 gene/protein is implicated in addiction.

## Verdict

Verdict: Mixed Evidence

GNB1 is **implicated in addiction-related biology in preclinical models**, especially for psychostimulant sensitization and alcohol intake, and there is also some opioid-related evidence in mice. However, the evidence is **mostly rodent and mechanistic**, not robust human clinical or human-genetic evidence. So the claim is too broad if read as “established in human addiction,” but it is reasonable if read as “involved in addiction-relevant neurobiology.” ([pubmed.ncbi.nlm.nih.gov](https://pubmed.ncbi.nlm.nih.gov/9221795/)) [S1]

## Confidence

Medium

Confidence is moderate because multiple independent rodent studies point in the same direction, including a functional manipulation study in nucleus accumbens, but the human evidence base appears sparse and lacks clear genome-wide or clinical replication. ([pubmed.ncbi.nlm.nih.gov](https://pubmed.ncbi.nlm.nih.gov/9221795/)) [S2]

## Supporting evidence

The strongest evidence comes from **psychostimulant models**. A classic primary study identified rGβ1/GNB1 as a psychostimulant-regulated gene: cocaine or amphetamine increased its expression in neurons of the nucleus accumbens shell, and antisense suppression of rGβ1 during repeated cocaine exposure **abolished the development of behavioral sensitization** without blocking the acute cocaine response. That is important because sensitization is widely used as an addiction-relevant neuroadaptation model, even though it is not identical to human addiction itself. ([pubmed.ncbi.nlm.nih.gov](https://pubmed.ncbi.nlm.nih.gov/9221795/)) [S3]

There is also meaningful **alcohol-related evidence**. In selectively bred high- vs low-alcohol-preferring mice, Gnb1 was one of the candidate transcripts associated with alcohol preference. In that study, Gnb1 expression differed between lines, and targeted **Gnb1 shRNA knockdown in the nucleus accumbens increased ethanol consumption** in DBA/2 mice, providing functional evidence rather than correlation alone. ([pubmed.ncbi.nlm.nih.gov](https://pubmed.ncbi.nlm.nih.gov/25160899/)) [S4]

A separate mouse systems-genetics study likewise found that **Gnb1 expression was negatively correlated with alcohol consumption**, and in one model combination of C1qc and Gnb1 expression explained a large proportion of variance in strain-level alcohol intake. The authors interpreted lower Gnb1 expression as part of a predisposition network for higher alcohol preference. ([pmc.ncbi.nlm.nih.gov](https://pmc.ncbi.nlm.nih.gov/articles/PMC2583933/)) [S5]

For **opioid-related phenotypes**, a nucleus accumbens transcriptional profiling study in C57 vs DBA mice reported that Gnb1 expression correlated with **morphine consumption** in a two-bottle choice paradigm, was confirmed by qPCR, and showed modest morphine-induced upregulation in nucleus accumbens. The same paper also noted prior evidence linking Gnb1 to psychostimulant responses and cited downregulation during morphine withdrawal in locus coeruleus. ([pmc.ncbi.nlm.nih.gov](https://pmc.ncbi.nlm.nih.gov/articles/PMC1851712/)) [S6]

A broader review of heterotrimeric G-protein subunits and drugs of abuse summarized these findings and concluded that Gβ1/GNB1 changes are associated with psychostimulant sensitization, opioid exposure, withdrawal-related adaptations, and long-term ethanol exposure. The review specifically notes that acute cocaine/amphetamine elevate GNB1 in striatum/nucleus accumbens, antisense blockade prevents sensitization, chronic morphine can increase GNB1 in nucleus accumbens, and chronic ethanol decreases hippocampal GNB1. ([pmc.ncbi.nlm.nih.gov](https://pmc.ncbi.nlm.nih.gov/articles/PMC2644661/)) [S7]

## Contradicting or limiting evidence

The main limitation is that the evidence is **not primarily human**. The best-supported results are in rodents and focus on addiction-relevant endophenotypes such as sensitization, voluntary intake, preference, withdrawal-related expression changes, and strain correlations. These are informative, but they do not by themselves establish GNB1 as a human addiction-risk gene. ([pubmed.ncbi.nlm.nih.gov](https://pubmed.ncbi.nlm.nih.gov/9221795/)) [S8]

Also, some of the evidence is **indirect or associative**. For example, strain-expression correlations and QTL/eQTL studies can nominate plausible genes, but they do not prove that natural variation in GNB1 is a primary driver rather than part of a broader network. Even the review summarizing the field notes that some GNB1 expression changes after psychostimulants may be part of a sequence of adaptations rather than the sole causal factor. ([pmc.ncbi.nlm.nih.gov](https://pmc.ncbi.nlm.nih.gov/articles/PMC2583933/)) [S9]

Direct human-genetic support appears limited. The current **GWAS Catalog page for GNB1 shows no curated associations listed**, which argues against GNB1 being an established genome-wide susceptibility locus for substance use disorders at present. ([ebi.ac.uk](https://www.ebi.ac.uk/gwas/genes/GNB1?utm_source=openai)) [S10]

Likewise, GeneReviews for GNB1 states that **no germline phenotypes other than GNB1 encephalopathy are known** for pathogenic variants in this gene. That does not rule out subtle common-variant effects on addiction, but it underscores that addiction is not a recognized core human GNB1 clinical phenotype. ([ncbi.nlm.nih.gov](https://www.ncbi.nlm.nih.gov/books/NBK554743/)) [S11]

## Mechanistic / biological rationale

Mechanistically, the claim is plausible. GNB1 encodes the **Gβ1 subunit of heterotrimeric G proteins**. After GPCR activation, Gα dissociates from the Gβγ dimer, and both arms regulate downstream effectors. GeneReviews summarizes that these pathways include **ion channels, phospholipases, adenylyl cyclases, PI3K, and MAPK signaling**. Those signaling nodes are central to synaptic plasticity and neuromodulation. ([ncbi.nlm.nih.gov](https://www.ncbi.nlm.nih.gov/books/NBK554743/)) [S12]

That matters for addiction because many receptors central to reward, reinforcement, withdrawal, and relapse biology are GPCRs or converge on GPCR-linked cascades. In the addiction-focused G-protein review, drug exposure altered GNB1/Gβ1 in nucleus accumbens, locus coeruleus, and hippocampus, and the authors proposed that these shifts could modify intracellular signaling involved in sensitization, tolerance, dependence, and withdrawal. ([pmc.ncbi.nlm.nih.gov](https://pmc.ncbi.nlm.nih.gov/articles/PMC2644661/)) [S13]

The regional pattern also makes biological sense. The key positive studies implicate **nucleus accumbens**, a core reward region, and **locus coeruleus**, a classic opioid withdrawal-related site. The psychostimulant study placed GNB1 regulation in nucleus accumbens shell, while the alcohol study showed that nucleus accumbens knockdown changed drinking behavior, and the morphine study connected nucleus accumbens Gnb1 expression with morphine consumption. ([pubmed.ncbi.nlm.nih.gov](https://pubmed.ncbi.nlm.nih.gov/9221795/)) [S14]

So biologically, GNB1 sits in a position where altered Gβγ signaling could reasonably influence dopamine- and opioid-linked plasticity. The mechanistic case is therefore **stronger than the human disease-association case**. ([ncbi.nlm.nih.gov](https://www.ncbi.nlm.nih.gov/books/NBK554743/)) [S15]

## Bottom line

If the claim means **“GNB1 has evidence of involvement in addiction-related neural adaptations and drug-related behaviors”**, that is supported by several rodent studies, especially for **cocaine sensitization** and **alcohol intake**, with some additional opioid-related data. ([pubmed.ncbi.nlm.nih.gov](https://pubmed.ncbi.nlm.nih.gov/9221795/)) [S16]

If the claim means **“GNB1 is an established human addiction gene”**, the evidence is currently insufficient; I did not find strong replicated human-genetic or clinical evidence, and the GWAS Catalog does not show curated GNB1 associations. ([ebi.ac.uk](https://www.ebi.ac.uk/gwas/genes/GNB1?utm_source=openai)) [S17]

So the most accurate summary is: **GNB1 is plausibly and experimentally implicated in addiction-related biology in animals, but not yet firmly established as a human addiction gene.** ([pubmed.ncbi.nlm.nih.gov](https://pubmed.ncbi.nlm.nih.gov/9221795/)) [S18]

## Cited sources

- **[S1]** rGbeta1: a psychostimulant-regulated gene essential for establishing cocaine sensitization - PubMed — https://pubmed.ncbi.nlm.nih.gov/9221795/
- **[S2]** rGbeta1: a psychostimulant-regulated gene essential for establishing cocaine sensitization - PubMed — https://pubmed.ncbi.nlm.nih.gov/9221795/
- **[S3]** rGbeta1: a psychostimulant-regulated gene essential for establishing cocaine sensitization - PubMed — https://pubmed.ncbi.nlm.nih.gov/9221795/
- **[S4]** Genetics of gene expression characterizes response to selective breeding for alcohol preference - PubMed — https://pubmed.ncbi.nlm.nih.gov/25160899/
- **[S5]** 
            The genomic determinants of alcohol preference in mice - PMC
         — https://pmc.ncbi.nlm.nih.gov/articles/PMC2583933/
- **[S6]** 
            Transcriptional profiling of C57 and DBA strains of mice in the absence and presence of morphine - PMC
         — https://pmc.ncbi.nlm.nih.gov/articles/PMC1851712/
- **[S7]** 
            Alterations in the levels of heterotrimeric G protein subunits induced by psychostimulants, opiates, barbiturates, and ethanol: Implications for drug dependence, tolerance, and withdrawal - PMC
         — https://pmc.ncbi.nlm.nih.gov/articles/PMC2644661/
- **[S8]** rGbeta1: a psychostimulant-regulated gene essential for establishing cocaine sensitization - PubMed — https://pubmed.ncbi.nlm.nih.gov/9221795/
- **[S9]** 
            The genomic determinants of alcohol preference in mice - PMC
         — https://pmc.ncbi.nlm.nih.gov/articles/PMC2583933/
- **[S10]** GWAS Catalog — https://www.ebi.ac.uk/gwas/genes/GNB1?utm_source=openai
- **[S11]** GNB1 Encephalopathy - GeneReviews® - NCBI Bookshelf — https://www.ncbi.nlm.nih.gov/books/NBK554743/
- **[S12]** GNB1 Encephalopathy - GeneReviews® - NCBI Bookshelf — https://www.ncbi.nlm.nih.gov/books/NBK554743/
- **[S13]** 
            Alterations in the levels of heterotrimeric G protein subunits induced by psychostimulants, opiates, barbiturates, and ethanol: Implications for drug dependence, tolerance, and withdrawal - PMC
         — https://pmc.ncbi.nlm.nih.gov/articles/PMC2644661/
- **[S14]** rGbeta1: a psychostimulant-regulated gene essential for establishing cocaine sensitization - PubMed — https://pubmed.ncbi.nlm.nih.gov/9221795/
- **[S15]** GNB1 Encephalopathy - GeneReviews® - NCBI Bookshelf — https://www.ncbi.nlm.nih.gov/books/NBK554743/
- **[S16]** rGbeta1: a psychostimulant-regulated gene essential for establishing cocaine sensitization - PubMed — https://pubmed.ncbi.nlm.nih.gov/9221795/
- **[S17]** GWAS Catalog — https://www.ebi.ac.uk/gwas/genes/GNB1?utm_source=openai
- **[S18]** rGbeta1: a psychostimulant-regulated gene essential for establishing cocaine sensitization - PubMed — https://pubmed.ncbi.nlm.nih.gov/9221795/
