## Supplementary material for "Understanding Substance Dependence: What Differentiates Addictive from Non-Addictive Drugs?": Folder for supplementary files: addictive-genes-claims-first100-019-timp2-gene-protein-is-implicated-in-addiction-938ee05d6ff8.md

# addictive-genes-claims-first100-019-938ee05d6ff8

**Original claim:** TIMP2 gene/protein is implicated in addiction.

## Claim

**“TIMP2 gene/protein is implicated in addiction.”**

## Verdict

Verdict: Likely True

The best available evidence supports **some implication of TIMP2 in addiction-related neurobiology**, but the support is **mainly preclinical**. Across rodent studies, TIMP2 is altered during methamphetamine sensitization/reward, cocaine relapse, and heroin cue-induced seeking, and experimental manipulation of the broader MMP/TIMP system affects addiction-like behaviors. However, TIMP2 has **not emerged as a well-established human addiction gene** in the way major dopaminergic, opioid, nicotinic, or alcohol-metabolism loci have in large genomic studies. ([pubmed.ncbi.nlm.nih.gov](https://pubmed.ncbi.nlm.nih.gov/17472698/)) [S1]

## Confidence

**Medium.**

Confidence is limited because the most direct evidence is from **animal and cellular work**, not replicated human genetics or clinical biomarker studies specifically centered on TIMP2 and substance use disorder. Still, the preclinical signal is fairly consistent across several addiction-relevant paradigms. ([pubmed.ncbi.nlm.nih.gov](https://pubmed.ncbi.nlm.nih.gov/17472698/)) [S2]

## Supporting evidence

The strongest direct support comes from a **methamphetamine study** in rodents. Repeated methamphetamine increased **TIMP-2 expression** in brain, and antisense suppression of TIMP-2 **enhanced** methamphetamine behavioral sensitization and potentiated methamphetamine-induced dopamine release in the nucleus accumbens. In the same line of work, MMP-2/9 inhibition reduced methamphetamine sensitization and conditioned place preference, and the authors concluded that the MMP/TIMP system is involved in methamphetamine-induced sensitization and reward. That is not proof that TIMP2 alone “causes addiction,” but it is strong evidence that TIMP2 participates in addiction-relevant adaptations. ([pubmed.ncbi.nlm.nih.gov](https://pubmed.ncbi.nlm.nih.gov/17472698/)) [S3]

A complementary review from the same group summarized these rodent findings similarly: repeated methamphetamine exposure was accompanied by increased **MMP-2/-9/TIMP-2** activity, antisense TIMP-2 enhanced sensitization, and MMP inhibition or MMP knockout attenuated sensitization and reward-related behavior. This reinforces that TIMP2 is part of a functionally relevant extracellular-matrix remodeling system in stimulant dependence models. ([pubmed.ncbi.nlm.nih.gov](https://pubmed.ncbi.nlm.nih.gov/18198472/)) [S4]

Evidence also exists in **cocaine relapse models**. In rats, cocaine extinction and reinstatement were associated with dynamic extracellular protease changes in nucleus accumbens core, and **TIMP-2 protein increased during reinstatement**. In that same study, MMP-2 and MMP-9 inhibition reduced cue-induced reinstatement behavior, placing TIMP-2 within a molecular system that tracks and modulates relapse-associated synaptic plasticity. ([pubmed.ncbi.nlm.nih.gov](https://pubmed.ncbi.nlm.nih.gov/25326689/)) [S5]

In **heroin seeking**, a 2021 study reported that heroin-paired cues increased **TIMP-1/2** around accumbens medium spiny neurons and that these changes transiently inhibited MMP-2 activity around D2-MSNs during cue-induced heroin seeking. Importantly, combined local inhibition of TIMP-1/2 **potentiated** cue-induced heroin seeking. This does not isolate TIMP2 cleanly from TIMP1, but it places TIMP2 directly in the circuitry and molecular events that regulate opioid cue reactivity. ([pubmed.ncbi.nlm.nih.gov](https://pubmed.ncbi.nlm.nih.gov/33579535/)) [S6]

Broader addiction-neuroscience reviews also place TIMP2 within the extracellular-matrix remodeling framework now considered relevant to relapse and reward memory. Reviews of the “tetrapartite synapse” and extracellular matrix in substance use disorders note that TIMP-2 rises after cocaine-primed reinstatement and after repeated methamphetamine exposure, fitting a model in which MMP/TIMP balance helps shape plasticity underlying drug seeking. ([pmc.ncbi.nlm.nih.gov](https://pmc.ncbi.nlm.nih.gov/articles/PMC4589426/?utm_source=openai)) [S7]

## Contradicting or limiting evidence

The main limitation is that **human evidence is weak or indirect**. A systematic review integrating GWAS and RNA-seq across alcohol, nicotine, cannabis, opioid, cocaine, and methamphetamine-related traits identified many shared and substance-specific genes, but the highlighted replicated/shared signals were genes such as **CACNB2, GRIN2B, PLXDC2, PKNOX2, ADH1B, ALDH2, CHRNA3, and CHRNA4**—not TIMP2. Likewise, a large multivariate GWAS of more than 1 million participants implicated loci such as **DRD2** and **PDE4B**, again not establishing TIMP2 as a major replicated human SUD locus. ([pubmed.ncbi.nlm.nih.gov](https://pubmed.ncbi.nlm.nih.gov/35491750/)) [S8]

Even in the animal literature, TIMP2 often appears as part of the **MMP/TIMP axis**, not as a definitively independent addiction driver. In cocaine relapse, TIMP-2 rises during reinstatement, but the causal behavioral manipulations in that paper targeted **MMP-2 and MMP-9**, not TIMP2 itself. So for cocaine, the evidence is more “TIMP2 is associated with the relapse-related molecular state” than “TIMP2 is proven to drive relapse.” ([pubmed.ncbi.nlm.nih.gov](https://pubmed.ncbi.nlm.nih.gov/25326689/)) [S9]

There is also an interpretive nuance: in both methamphetamine and heroin paradigms, the observed TIMP2-related effects may be **compensatory or protective rather than pro-addictive**. In methamphetamine models, lowering TIMP-2 made sensitization worse; in heroin models, inhibiting TIMP-1/2 potentiated cue-induced seeking. That means TIMP2 may be implicated in addiction **as a regulator of the system**, possibly buffering maladaptive plasticity, rather than as a straightforward risk-promoting molecule. ([pubmed.ncbi.nlm.nih.gov](https://pubmed.ncbi.nlm.nih.gov/17472698/)) [S10]

## Mechanistic / biological rationale

Mechanistically, the claim is biologically plausible. TIMP2 is an endogenous regulator of **matrix metalloproteinases** and extracellular matrix turnover; NCBI describes TIMP family proteins as essential regulators of extracellular matrix remodeling. ([ncbi.nlm.nih.gov](https://www.ncbi.nlm.nih.gov/gene/7077)) [S11]

That matters because addiction is increasingly understood as a disorder of **activity-dependent synaptic plasticity** and reward-memory remodeling. Reviews of reward memory and substance use disorders describe extracellular-matrix proteases and their inhibitors as key modulators of synaptic remodeling, perineuronal nets, and relapse-related plasticity. ([frontiersin.org](https://www.frontiersin.org/articles/10.3389/fncel.2023.1208974/full)) [S12]

TIMP2 also has bona fide CNS roles beyond generic matrix inhibition. A recent neuronal study showed that neuronal TIMP2 is necessary for **ECM remodeling, dendritic spine regulation, adult neurogenesis, and hippocampus-dependent plasticity/memory**. Deleting neuronal TIMP2 increased extracellular-matrix density around synapses and impaired plasticity-related processes. Those are exactly the kinds of mechanisms that could intersect with addiction, where persistent cue-drug memories and relapse depend on maladaptive synaptic remodeling. ([pmc.ncbi.nlm.nih.gov](https://pmc.ncbi.nlm.nih.gov/articles/PMC10730400/)) [S13]

In addition, TIMP2 can regulate **neuroinflammatory signaling** in microglia. In cultured microglia, TIMP-2 overexpression reduced NO, TNF-α, IL-1β, ROS, MMP activity, MAPK phosphorylation, and NF-κB signaling, while increasing IL-10. Because neuroimmune and ECM pathways both contribute to substance-induced plasticity and relapse vulnerability, this gives TIMP2 a plausible route into addiction biology even if the direct causal evidence remains mostly preclinical. ([jneuroinflammation.biomedcentral.com](https://jneuroinflammation.biomedcentral.com/articles/10.1186/1742-2094-11-116)) [S14]

## Bottom line

**Yes—TIMP2 is reasonably described as “implicated in addiction,” but mainly in a preclinical, mechanistic sense.** Rodent studies tie TIMP2 to methamphetamine sensitization/reward, cocaine relapse-associated molecular remodeling, and heroin cue-induced seeking, all within the MMP/TIMP extracellular-matrix system. However, **direct human evidence is sparse**, and current large-scale human genetics has **not established TIMP2 as a major replicated addiction gene**. A careful formulation would be:

> **TIMP2 is implicated in addiction-related synaptic/extracellular-matrix plasticity in animal models, but its role in human substance use disorder remains unconfirmed and probably indirect.** ([pubmed.ncbi.nlm.nih.gov](https://pubmed.ncbi.nlm.nih.gov/17472698/)) [S15]

## Cited sources

- **[S1]** Role of matrix metalloproteinase and tissue inhibitor of MMP in methamphetamine-induced behavioral sensitization and reward: implications for dopamine receptor down-regulation and dopamine release - PubMed — https://pubmed.ncbi.nlm.nih.gov/17472698/
- **[S2]** Role of matrix metalloproteinase and tissue inhibitor of MMP in methamphetamine-induced behavioral sensitization and reward: implications for dopamine receptor down-regulation and dopamine release - PubMed — https://pubmed.ncbi.nlm.nih.gov/17472698/
- **[S3]** Role of matrix metalloproteinase and tissue inhibitor of MMP in methamphetamine-induced behavioral sensitization and reward: implications for dopamine receptor down-regulation and dopamine release - PubMed — https://pubmed.ncbi.nlm.nih.gov/17472698/
- **[S4]** Neuropsychotoxicity of abused drugs: involvement of matrix metalloproteinase-2 and -9 and tissue inhibitor of matrix metalloproteinase-2 in methamphetamine-induced behavioral sensitization and reward in rodents - PubMed — https://pubmed.ncbi.nlm.nih.gov/18198472/
- **[S5]** Synaptic plasticity mediating cocaine relapse requires matrix metalloproteinases - PubMed — https://pubmed.ncbi.nlm.nih.gov/25326689/
- **[S6]** Heroin Seeking and Extinction From Seeking Activate Matrix Metalloproteinases at Synapses on Distinct Subpopulations of Accumbens Cells - PubMed — https://pubmed.ncbi.nlm.nih.gov/33579535/
- **[S7]** The tetrapartite synapse: extracellular matrix remodeling contributes to corticoaccumbens plasticity underlying drug addiction - PMC — https://pmc.ncbi.nlm.nih.gov/articles/PMC4589426/?utm_source=openai
- **[S8]** Common and distinguishing genetic factors for substance use behavior and disorder: an integrated analysis of genomic and transcriptomic studies from both human and animal studies - PubMed — https://pubmed.ncbi.nlm.nih.gov/35491750/
- **[S9]** Synaptic plasticity mediating cocaine relapse requires matrix metalloproteinases - PubMed — https://pubmed.ncbi.nlm.nih.gov/25326689/
- **[S10]** Role of matrix metalloproteinase and tissue inhibitor of MMP in methamphetamine-induced behavioral sensitization and reward: implications for dopamine receptor down-regulation and dopamine release - PubMed — https://pubmed.ncbi.nlm.nih.gov/17472698/
- **[S11]** TIMP2 TIMP metallopeptidase inhibitor 2 [Homo sapiens (human)] - Gene - NCBI — https://www.ncbi.nlm.nih.gov/gene/7077
- **[S12]** Frontiers | Dynamic regulation of the extracellular matrix in reward memory processes: a question of time — https://www.frontiersin.org/articles/10.3389/fncel.2023.1208974/full
- **[S13]** 
            Neuronal TIMP2 regulates hippocampus-dependent plasticity and extracellular matrix complexity - PMC
         — https://pmc.ncbi.nlm.nih.gov/articles/PMC10730400/
- **[S14]** The anti-inflammatory role of tissue inhibitor of metalloproteinase-2 in lipopolysaccharide-stimulated microglia | Journal of Neuroinflammation | Springer Nature Link — https://jneuroinflammation.biomedcentral.com/articles/10.1186/1742-2094-11-116
- **[S15]** Role of matrix metalloproteinase and tissue inhibitor of MMP in methamphetamine-induced behavioral sensitization and reward: implications for dopamine receptor down-regulation and dopamine release - PubMed — https://pubmed.ncbi.nlm.nih.gov/17472698/
