## Supplementary material for "Understanding Substance Dependence: What Differentiates Addictive from Non-Addictive Drugs?": Folder for supplementary files: addictive-genes-claims-first100-020-atp1a1-gene-protein-is-implicated-in-addiction-3a7cea2784ce.md

# addictive-genes-claims-first100-020-3a7cea2784ce

**Original claim:** ATP1A1 gene/protein is implicated in addiction.

## Claim

ATP1A1 gene/protein is implicated in addiction.

## Verdict

Verdict: No Evidence

## Confidence

Medium.

## Supporting evidence

ATP1A1 encodes the α1 subunit of the Na+/K+-ATPase, a membrane pump that uses ATP to move sodium out of cells and potassium into cells, thereby helping maintain electrochemical gradients required for cell excitability and signaling. It is highly relevant to basic neurobiology and is expressed in the nervous system. ([medlineplus.gov](https://medlineplus.gov/genetics/gene/atp1a1/)) [S1]

There is also some **indirect, mechanistic plausibility** for sodium-pump biology to matter in addiction-relevant circuits. In astrocytes, Na,K-ATPase α1 and α2 physically interact with the glutamate transporters GLAST and GLT-1, and pump function supports glutamate clearance/homeostasis. Because glutamate dysregulation is central to many addiction models, this puts the broader Na/K-ATPase system in a biologically relevant neighborhood. ([pubmed.ncbi.nlm.nih.gov](https://pubmed.ncbi.nlm.nih.gov/24901986/)) [S2]

Likewise, pharmacologic inhibition of brain Na+,K+-ATPase with intracerebroventricular ouabain caused hyperactivity and impulsivity in mice, increased c-fos in nucleus accumbens and prefrontal cortex, and appeared to act through dopamine D2 receptor-related signaling. Those are addiction-adjacent behaviors/circuits, so they support a plausible pathway-level connection between Na/K-ATPase dysfunction and altered motivational/behavioral control. ([pubmed.ncbi.nlm.nih.gov](https://pubmed.ncbi.nlm.nih.gov/30296459/)) [S3]

However, these supportive points are **not direct evidence that ATP1A1 itself is implicated in addiction**. They show only that the protein’s normal biology overlaps with neural systems often involved in addiction. ([pubmed.ncbi.nlm.nih.gov](https://pubmed.ncbi.nlm.nih.gov/30296459/)) [S4]

## Contradicting or limiting evidence

The main limitation is straightforward: I could not identify convincing human genetic, human clinical, or animal addiction-model evidence specifically linking **ATP1A1** to addiction liability, diagnosis, drug reinforcement, relapse, or treatment response in the way that would usually justify saying a gene is “implicated in addiction.” The strongest current large-scale cross-disorder GWAS of substance use disorders highlighted loci such as **DRD2, PDE4B, ADH1B, CHRNA5, OPRM1,** and **CACNA1A** rather than ATP1A1. ([pubmed.ncbi.nlm.nih.gov](https://pubmed.ncbi.nlm.nih.gov/37250466/)) [S5]

Consistent with that, the GWAS Catalog gene page for **ATP1A1** currently shows empty sections for “Associations,” “Studies,” and “Traits,” which argues against ATP1A1 being a recognized GWAS-supported addiction gene at present. ([ebi.ac.uk](https://www.ebi.ac.uk/gwas/genes/ATP1A1)) [S6]

What *is* well established for ATP1A1 is a set of other disease links: aldosterone-producing adenoma, Charcot-Marie-Tooth neuropathy, spastic paraplegia, hypomagnesemia with seizures/intellectual disability, and related neurodevelopmental disorders. Authoritative summaries and disease papers describe these phenotypes, but not addiction as a defining ATP1A1-associated phenotype. ([medlineplus.gov](https://medlineplus.gov/genetics/gene/atp1a1/)) [S7]

That distinction matters. A gene can be crucial for neuronal ion homeostasis without having evidence for a specific role in addiction. ATP1A1 currently fits that pattern: important for neural physiology, yes; directly implicated in addiction, not yet demonstrated. ([medlineplus.gov](https://medlineplus.gov/genetics/gene/atp1a1/)) [S8]

A further limiting point is that the closest addiction-related evidence in this gene family often involves **other isoforms**, especially **ATP1A2**, not ATP1A1. For example, Atp1a2 has been reported to contribute modestly to alcohol-related behaviors in mice, and a 2025 mouse systems-genetics study identified **Atp1a2** as a candidate gene for oxycodone-induced locomotor activation and withdrawal-related anxiety-like behavior. That family-level signal does not transfer automatically to ATP1A1. ([pmc.ncbi.nlm.nih.gov](https://pmc.ncbi.nlm.nih.gov/articles/PMC5123668/)) [S9]

## Mechanistic / biological rationale

Biologically, ATP1A1 is a plausible **background regulator** rather than an established addiction gene. The α1 Na+/K+-ATPase is fundamental for maintaining membrane potential, ion gradients, and neuronal/glial homeostasis. Disturbing those gradients can alter excitability, transmitter handling, and network function. ([medlineplus.gov](https://medlineplus.gov/genetics/gene/atp1a1/)) [S10]

There are at least three reasons one might have hypothesized a role in addiction:

1. **Excitability and dopamine/glutamate balance.** Addiction involves long-lasting changes in corticostriatal and mesolimbic circuits. ATP1A1 contributes to the ionic gradients that underlie firing, synaptic transmission, and glial support functions. ([medlineplus.gov](https://medlineplus.gov/genetics/gene/atp1a1/)) [S11]

2. **Astrocyte glutamate homeostasis.** Na,K-ATPase α subunits interact with GLAST/GLT-1 and help restore intracellular sodium after glutamate uptake. Since impaired glutamate homeostasis is a recurring theme in relapse/reinstatement models, this is conceptually relevant. Still, the cited work is not addiction-specific and is not uniquely about ATP1A1. ([pubmed.ncbi.nlm.nih.gov](https://pubmed.ncbi.nlm.nih.gov/24901986/)) [S12]

3. **Behavioral effects of pump inhibition.** Broad Na+/K+-ATPase inhibition can produce dopamine-linked hyperactivity/impulsivity and alter activity in nucleus accumbens/prefrontal cortex. But this was a pharmacologic manipulation of the pump system, not evidence that inherited or acquired ATP1A1 variation drives addiction phenotypes. ([pubmed.ncbi.nlm.nih.gov](https://pubmed.ncbi.nlm.nih.gov/30296459/)) [S13]

So the mechanistic story is: **plausible indirect relevance**, but **no clear direct implication**. That is weaker than what is usually meant in genetics/neurobiology when a protein is said to be “implicated in addiction.” ([pubmed.ncbi.nlm.nih.gov](https://pubmed.ncbi.nlm.nih.gov/30296459/)) [S14]

## Bottom line

ATP1A1 is clearly important for neuronal and glial homeostasis, and the broader Na+/K+-ATPase system can influence behaviors and circuits that overlap with addiction biology. But based on currently visible scholarly and authoritative sources, there is **no solid direct evidence** that **ATP1A1 itself** is an established addiction gene/protein. Large substance-use GWAS do not currently highlight it, the GWAS Catalog does not list curated ATP1A1 associations, and the well-documented human phenotypes for ATP1A1 are neurological/endocrine rather than addiction-related. ([ebi.ac.uk](https://www.ebi.ac.uk/gwas/genes/ATP1A1)) [S15]

If you want, I can do a second-pass review focused narrowly on one addiction subtype—alcohol, nicotine, opioids, stimulants, or cannabis—to see whether any niche transcriptomic/proteomic study has reported ATP1A1 specifically.

## Cited sources

- **[S1]** ATP1A1 gene: MedlinePlus Genetics — https://medlineplus.gov/genetics/gene/atp1a1/
- **[S2]** Role of Na,K-ATPase α1 and α2 isoforms in the support of astrocyte glutamate uptake - PubMed — https://pubmed.ncbi.nlm.nih.gov/24901986/
- **[S3]** Na+, K+-ATPase inhibition causes hyperactivity and impulsivity in mice via dopamine D2 receptor-mediated mechanism - PubMed — https://pubmed.ncbi.nlm.nih.gov/30296459/
- **[S4]** Na+, K+-ATPase inhibition causes hyperactivity and impulsivity in mice via dopamine D2 receptor-mediated mechanism - PubMed — https://pubmed.ncbi.nlm.nih.gov/30296459/
- **[S5]** Multivariate genome-wide association meta-analysis of over 1 million subjects identifies loci underlying multiple substance use disorders - PubMed — https://pubmed.ncbi.nlm.nih.gov/37250466/
- **[S6]** GWAS Catalog — https://www.ebi.ac.uk/gwas/genes/ATP1A1
- **[S7]** ATP1A1 gene: MedlinePlus Genetics — https://medlineplus.gov/genetics/gene/atp1a1/
- **[S8]** ATP1A1 gene: MedlinePlus Genetics — https://medlineplus.gov/genetics/gene/atp1a1/
- **[S9]** 
            Atp1a2 Contributes Modestly to Alcohol-Related Behaviors - PMC
         — https://pmc.ncbi.nlm.nih.gov/articles/PMC5123668/
- **[S10]** ATP1A1 gene: MedlinePlus Genetics — https://medlineplus.gov/genetics/gene/atp1a1/
- **[S11]** ATP1A1 gene: MedlinePlus Genetics — https://medlineplus.gov/genetics/gene/atp1a1/
- **[S12]** Role of Na,K-ATPase α1 and α2 isoforms in the support of astrocyte glutamate uptake - PubMed — https://pubmed.ncbi.nlm.nih.gov/24901986/
- **[S13]** Na+, K+-ATPase inhibition causes hyperactivity and impulsivity in mice via dopamine D2 receptor-mediated mechanism - PubMed — https://pubmed.ncbi.nlm.nih.gov/30296459/
- **[S14]** Na+, K+-ATPase inhibition causes hyperactivity and impulsivity in mice via dopamine D2 receptor-mediated mechanism - PubMed — https://pubmed.ncbi.nlm.nih.gov/30296459/
- **[S15]** GWAS Catalog — https://www.ebi.ac.uk/gwas/genes/ATP1A1
