## Supplementary material for "Understanding Substance Dependence: What Differentiates Addictive from Non-Addictive Drugs?": Folder for supplementary files: addictive-genes-claims-first100-021-myl6-gene-protein-is-implicated-in-addiction-0dc2161d8973.md

# addictive-genes-claims-first100-021-0dc2161d8973

**Original claim:** MYL6 gene/protein is implicated in addiction.

## Claim

MYL6 gene/protein is implicated in addiction.

## Verdict

Verdict: Likely True

## Confidence

Medium

## Supporting evidence

The strongest direct evidence I found is a 2024 single-nucleus transcriptomics study of human dorsal striatum in opioid use disorder (OUD). In postmortem tissue from people with OUD versus unaffected comparison subjects, **MYL6 was significantly upregulated in striatal medium spiny neurons**, with reported **log2 fold-change >1.13 and FDR <0.045**. In that same analysis, the affected neuronal programs were enriched for stress-response, DNA replication/cell-cycle re-entry, and related injury-response pathways, placing MYL6 within a biologically relevant OUD-associated transcriptional signature in a core reward/motivation circuit. ([pmc.ncbi.nlm.nih.gov](https://pmc.ncbi.nlm.nih.gov/articles/PMC10831093/)) [S1]

There is also some cross-substance support from alcohol-related work. In a nonhuman primate model of heavy drinking, orbitofrontal cortex synaptomic profiling identified **MYL6 among alcohol-sensitive synaptic proteins** in heavy-drinking macaques. That does not prove causality, but it indicates that MYL6 protein abundance is altered in an addiction-relevant cortical region during chronic alcohol exposure. ([pmc.ncbi.nlm.nih.gov](https://pmc.ncbi.nlm.nih.gov/articles/PMC5373140/?utm_source=openai)) [S2]

Independent of addiction, MYL6 is clearly a plausible CNS-relevant molecule rather than a purely peripheral artifact. NCBI describes MYL6 as a **protein-coding myosin light chain** expressed in smooth muscle and non-muscle tissues, and the Human Protein Atlas reports that MYL6 RNA is **detected in all major human brain regions** with low regional specificity. A spatial transcriptomics study in mouse brain further found **region-specific isoform usage** of Myl6 across olfactory bulb and coronal brain sections, supporting a bona fide neural role. ([ncbi.nlm.nih.gov](https://www.ncbi.nlm.nih.gov/gene/4637?utm_source=openai)) [S3]

Taken together, these data support the statement that MYL6 is **implicated** in addiction if “implicated” is interpreted in the usual literature-review sense of **associated with addiction-related molecular alterations in relevant brain tissue**, especially for OUD. ([pmc.ncbi.nlm.nih.gov](https://pmc.ncbi.nlm.nih.gov/articles/PMC10831093/)) [S4]

## Contradicting or limiting evidence

The evidence is still **limited and mostly associative**. The clearest positive study is transcriptomic, not functional: it shows MYL6 dysregulation in human OUD brain, but it does **not** demonstrate that MYL6 drives drug seeking, tolerance, withdrawal, relapse, or other addiction behaviors. In other words, MYL6 could be a downstream consequence, compensatory response, or correlated marker rather than a causal mediator. ([pmc.ncbi.nlm.nih.gov](https://pmc.ncbi.nlm.nih.gov/articles/PMC10831093/)) [S5]

The main human OUD single-nucleus study was also modest in size, using **6 individuals with OUD and 6 unaffected comparison subjects** for the human striatal analysis. That is a valuable dataset, but still small for declaring a robust, generalized addiction gene across populations and substances. ([pmc.ncbi.nlm.nih.gov](https://pmc.ncbi.nlm.nih.gov/articles/PMC10831093/)) [S6]

The alcohol-related primate evidence is weaker than the OUD evidence. The macaque synaptomics study was partly exploratory and used a small proteomics cohort (**3 controls, 5 drinkers**). Moreover, while MYL6 appeared among alcohol-sensitive proteins, the authors’ discriminative biomarker analysis highlighted **MYL6B** and **NRCAM**, not MYL6 itself, as the top proteins separating control from heavy-drinking animals. That makes the alcohol signal for MYL6 suggestive rather than definitive. ([pubmed.ncbi.nlm.nih.gov](https://pubmed.ncbi.nlm.nih.gov/28270566/?utm_source=openai)) [S7]

More broadly, recent multiomic opioid-addiction network work has emphasized genes and pathways centered on **OPRM1, BDNF/ERK signaling, glutamatergic/GABAergic signaling, ion channels, and inflammatory/stress pathways**. MYL6 is not yet presented as a leading, replicated addiction gene in the way those pathways are. That does not negate the OUD finding, but it does limit how strongly one should generalize the claim. ([pubmed.ncbi.nlm.nih.gov](https://pubmed.ncbi.nlm.nih.gov/39615775/)) [S8]

## Mechanistic / biological rationale

MYL6 encodes **myosin light chain 6**, a component of the actomyosin motor machinery in non-muscle cells. Cytoskeletal and actomyosin systems are highly relevant to neuronal function because they help govern **spine structure, receptor trafficking, neurite dynamics, and synaptic remodeling**—all processes central to drug-induced plasticity. MYL6 is also broadly present in the brain, which strengthens the plausibility of a neural role. ([ncbi.nlm.nih.gov](https://www.ncbi.nlm.nih.gov/gene/4637?utm_source=openai)) [S9]

The OUD single-nucleus paper specifically interprets MYL6 as linked to **Rho-GTPase signaling**, with potential relevance to **glutamate receptor endocytosis** and **neuroplasticity**, and discusses it alongside neuronal stress/DNA-damage-associated programs in striatal medium spiny neurons. That is mechanistically coherent with addiction biology, since maladaptive corticostriatal plasticity is a core feature of substance use disorders. Still, this remains a **biological rationale**, not proof of mechanism in addiction. ([pmc.ncbi.nlm.nih.gov](https://pmc.ncbi.nlm.nih.gov/articles/PMC10831093/)) [S10]

So biologically, MYL6 is a plausible participant in addiction-related neural remodeling; empirically, however, direct tests are still missing. I did **not** find evidence in the sources reviewed showing that selective MYL6 manipulation in animals changes self-administration, conditioned place preference, reinstatement, craving-like behavior, or withdrawal severity. ([pmc.ncbi.nlm.nih.gov](https://pmc.ncbi.nlm.nih.gov/articles/PMC10831093/)) [S11]

## Bottom line

There **is** evidence that MYL6 is implicated in addiction-related biology, with the best support coming from **human OUD striatal single-nucleus transcriptomics** showing significant MYL6 upregulation in medium spiny neurons, plus weaker **alcohol-related primate proteomic** evidence. ([pmc.ncbi.nlm.nih.gov](https://pmc.ncbi.nlm.nih.gov/articles/PMC10831093/)) [S12]

However, the evidence is **not yet strong enough** to say MYL6 is an established causal addiction gene or a major addiction driver across substances. Right now, the most accurate reading is:

- **Yes, MYL6 is implicated**, especially in **opioid use disorder-related brain molecular changes**.
- **No, it is not yet firmly established** as a replicated, causal addiction mechanism or therapeutic target. ([pmc.ncbi.nlm.nih.gov](https://pmc.ncbi.nlm.nih.gov/articles/PMC10831093/)) [S13]

## Cited sources

- **[S1]** 
            Single nuclei transcriptomics in human and non-human primate striatum in opioid use disorder - PMC
         — https://pmc.ncbi.nlm.nih.gov/articles/PMC10831093/
- **[S2]** Orbitofrontal Neuroadaptations and Cross-Species Synaptic Biomarkers in Heavy-Drinking Macaques - PMC — https://pmc.ncbi.nlm.nih.gov/articles/PMC5373140/?utm_source=openai
- **[S3]** MYL6 myosin light chain 6 [Homo sapiens (human)] - Gene - NCBI — https://www.ncbi.nlm.nih.gov/gene/4637?utm_source=openai
- **[S4]** 
            Single nuclei transcriptomics in human and non-human primate striatum in opioid use disorder - PMC
         — https://pmc.ncbi.nlm.nih.gov/articles/PMC10831093/
- **[S5]** 
            Single nuclei transcriptomics in human and non-human primate striatum in opioid use disorder - PMC
         — https://pmc.ncbi.nlm.nih.gov/articles/PMC10831093/
- **[S6]** 
            Single nuclei transcriptomics in human and non-human primate striatum in opioid use disorder - PMC
         — https://pmc.ncbi.nlm.nih.gov/articles/PMC10831093/
- **[S7]** Orbitofrontal Neuroadaptations and Cross-Species Synaptic Biomarkers in Heavy-Drinking Macaques - PubMed — https://pubmed.ncbi.nlm.nih.gov/28270566/?utm_source=openai
- **[S8]** Multiomic Network Analysis Identifies Dysregulated Neurobiological Pathways in Opioid Addiction - PubMed — https://pubmed.ncbi.nlm.nih.gov/39615775/
- **[S9]** MYL6 myosin light chain 6 [Homo sapiens (human)] - Gene - NCBI — https://www.ncbi.nlm.nih.gov/gene/4637?utm_source=openai
- **[S10]** 
            Single nuclei transcriptomics in human and non-human primate striatum in opioid use disorder - PMC
         — https://pmc.ncbi.nlm.nih.gov/articles/PMC10831093/
- **[S11]** 
            Single nuclei transcriptomics in human and non-human primate striatum in opioid use disorder - PMC
         — https://pmc.ncbi.nlm.nih.gov/articles/PMC10831093/
- **[S12]** 
            Single nuclei transcriptomics in human and non-human primate striatum in opioid use disorder - PMC
         — https://pmc.ncbi.nlm.nih.gov/articles/PMC10831093/
- **[S13]** 
            Single nuclei transcriptomics in human and non-human primate striatum in opioid use disorder - PMC
         — https://pmc.ncbi.nlm.nih.gov/articles/PMC10831093/
