## Supplementary material for "Understanding Substance Dependence: What Differentiates Addictive from Non-Addictive Drugs?": Folder for supplementary files: addictive-genes-claims-first100-022-aplnr-gene-protein-is-implicated-in-addiction-96ceec0a6f86.md

# addictive-genes-claims-first100-022-96ceec0a6f86

**Original claim:** APLNR gene/protein is implicated in addiction.

## Claim

APLNR (the apelin receptor, also called APJ) gene/protein is implicated in addiction.

## Verdict

Verdict: Mixed Evidence

## Confidence

Medium

## Supporting evidence

APLNR has some **preclinical** links to addiction-relevant biology, especially in opioid-related models, but the evidence is mostly indirect. The strongest signal is that chronic mu-opioid receptor activation in mice altered lateral hypothalamic gene expression and **downregulated apelin**; the authors explicitly suggested this points to a possible role in opioid signaling, hedonic homeostasis, and reward-process changes relevant to addictive disease. Importantly, this was observed in a morphine-dependence paradigm and was mu-opioid-receptor-dependent. ([pubmed.ncbi.nlm.nih.gov](https://pubmed.ncbi.nlm.nih.gov/18591478/)) [S1]

A newer rat study of morphine dependence/withdrawal reported that **hypothalamic APLNR mRNA changed across morphine/naloxone conditions**, with reduced APLNR expression in the morphine-plus-naloxone withdrawal group versus controls. The authors interpreted these findings as suggesting involvement of the apelinergic system in opiate-related adaptations, though the experiment was framed around hypothalamic feeding-regulation peptides rather than addiction behavior per se. ([dergipark.org.tr](https://dergipark.org.tr/tr/download/article-file/4473656)) [S2]

There is also evidence of **functional cross-talk between the apelin/APLNR system and opioid systems**. In rats, chronic intrathecal morphine and chronic apelin each produced **cross-tolerance** to the other’s analgesic effect, and chronic morphine significantly **downregulated apelin receptor expression** in spinal tissue. This supports biological interaction between opioid and apelin/APLNR signaling, although it addresses tolerance/antinociception more than compulsive drug seeking. ([pubmed.ncbi.nlm.nih.gov](https://pubmed.ncbi.nlm.nih.gov/38517751/)) [S3]

At the receptor level, APLNR has been shown to **heterodimerize with kappa-opioid receptor (KOR)** in cell systems, changing downstream signaling toward increased PKC and altered cAMP/PKA signaling. Because KOR signaling is well known to influence dysphoria, aversion, stress, and relapse-related processes in addiction, this receptor–receptor interaction provides a plausible mechanistic route by which APLNR could affect addiction biology. ([pubmed.ncbi.nlm.nih.gov](https://pubmed.ncbi.nlm.nih.gov/22200678/)) [S4]

APLNR is also expressed in **addiction-relevant brain regions**. Rat apelin receptor mRNA is present not only in the hypothalamus but also in the **substantia nigra pars compacta, dorsal raphe, dentate gyrus, and hippocampus**. More recently, nigral dopaminergic neurons were shown to express APJ, and apelin modulated their firing and motor output in vivo. Since mesolimbic/mesostriatal dopamine systems are core components of reward and addiction neurocircuitry, this supports biological plausibility. ([pubmed.ncbi.nlm.nih.gov](https://pubmed.ncbi.nlm.nih.gov/11146423/)) [S5]

Finally, stress-related regulation may matter. In rats, **acute and repeated restraint stress upregulated APJ receptor mRNA in the hypothalamic paraventricular nucleus**, suggesting APLNR participates in stress-axis regulation. Because addiction involves withdrawal/negative-affect and stress-driven relapse circuitry, this is relevant, though still indirect. ([pubmed.ncbi.nlm.nih.gov](https://pubmed.ncbi.nlm.nih.gov/14622440/)) [S6]

## Contradicting or limiting evidence

The main limitation is that **direct addiction evidence is sparse**. I did not find strong human evidence that APLNR variants are reproducibly associated with substance use disorder, nor clear clinical studies showing altered APLNR as a biomarker or causal driver of addiction. The current literature is dominated by animal, cellular, and mechanistic studies rather than human addiction genetics or treatment-response studies. ([pmc.ncbi.nlm.nih.gov](https://pmc.ncbi.nlm.nih.gov/articles/PMC11491857/)) [S7]

Much of the literature is also **about apelin/APLNR in pain, neuroprotection, feeding, cardiovascular regulation, or stress**, not addiction behavior itself. For example, methamphetamine work showed apelin-13 protected PC12 cells from oxidative stress, autophagy, and apoptosis after meth exposure, but that is evidence for **drug-induced neurotoxicity modulation**, not evidence that APLNR drives reinforcement, craving, relapse, or compulsive use. ([pubmed.ncbi.nlm.nih.gov](https://pubmed.ncbi.nlm.nih.gov/31385138/)) [S8]

Similarly, the morphine cross-tolerance study addresses **analgesic tolerance** and receptor downregulation in spinal cord, not addiction-like endpoints such as self-administration, conditioned place preference, escalation, reinstatement, or relapse. Tolerance and addiction are related but not equivalent constructs. ([pubmed.ncbi.nlm.nih.gov](https://pubmed.ncbi.nlm.nih.gov/38517751/)) [S9]

A notable limiting study is the Aplnr knockout mouse behavioral paper: although the knockout altered **fear-related behavior** and acoustic startle, the abstract did **not report a clear reward-preference abnormality**, despite reward preference being included in the behavioral battery. This does not rule out an addiction role, but it weakens any claim that APLNR has an established, robust reward phenotype. ([pubmed.ncbi.nlm.nih.gov](https://pubmed.ncbi.nlm.nih.gov/33309737/)) [S10]

Also, some of the pro-addiction interpretations come from relatively **small or specialized preclinical studies**, including a conference abstract and a small gene-expression paper focused on hypothalamic feeding peptides. Those studies are hypothesis-generating, not definitive. ([pmc.ncbi.nlm.nih.gov](https://pmc.ncbi.nlm.nih.gov/articles/PMC11491857/)) [S11]

## Mechanistic / biological rationale

Biologically, the claim is plausible. Addiction depends on interactions among **reward circuitry, stress systems, and neuroadaptation**, especially within dopamine, opioid, and cortico-limbic networks. APLNR is positioned to intersect with several of these domains. ([pmc.ncbi.nlm.nih.gov](https://pmc.ncbi.nlm.nih.gov/articles/PMC6135092/)) [S12]

First, APLNR is a **Gi/o-coupled GPCR** that can regulate cAMP/PKA, PI3K/Akt, PKC, ERK, calcium-linked signaling, and β-arrestin pathways. These are classic signaling modules involved in synaptic plasticity and drug-induced neuroadaptation. ([pubmed.ncbi.nlm.nih.gov](https://pubmed.ncbi.nlm.nih.gov/25275559/)) [S13]

Second, APLNR is present in brain regions relevant to **dopamine, stress, memory, and interoception**, including hypothalamus, substantia nigra, dorsal raphe, and hippocampus. Expression in nigral dopamine neurons, plus evidence that apelin modifies their excitability, supports a potential influence on motivational or reinforcement processes. This is an inference from neuroanatomy and physiology, not direct proof of addiction causality. ([pubmed.ncbi.nlm.nih.gov](https://pubmed.ncbi.nlm.nih.gov/11146423/)) [S14]

Third, APLNR shows **molecular cross-talk with opioid systems**. Mu-opioid activation alters apelin/apelinergic expression; APLNR can heterodimerize with KOR and ORL1; and chronic morphine can downregulate apelin receptor expression in pain models. Since opioid receptors are central to reward, withdrawal, and relapse, this cross-talk could plausibly make APLNR a modulatory node in addiction-related adaptation. Again, this is mechanistic plausibility rather than settled demonstration. ([pubmed.ncbi.nlm.nih.gov](https://pubmed.ncbi.nlm.nih.gov/18591478/)) [S15]

## Bottom line

APLNR is **not an established major addiction gene/protein on the level of canonical targets** like OPRM1, DRD2, or CHRNA5. However, there is **credible preclinical evidence** that the apelin/APLNR system interacts with opioid signaling, stress circuitry, and dopaminergic neurons in ways that are relevant to addiction biology. The current evidence supports saying APLNR is **a plausible, emerging, but still weakly validated addiction-related candidate**, especially for **opioid-related neuroadaptation** rather than addiction broadly. ([pubmed.ncbi.nlm.nih.gov](https://pubmed.ncbi.nlm.nih.gov/18591478/)) [S16]

## Cited sources

- **[S1]** Gene expression is altered in the lateral hypothalamus upon activation of the mu opioid receptor - PubMed — https://pubmed.ncbi.nlm.nih.gov/18591478/
- **[S2]** Untitled source — https://dergipark.org.tr/tr/download/article-file/4473656
- **[S3]** Evidence for the Induction of Analgesic Cross-Tolerance Between Opioid and Apelin/APJ Systems in Male Rats - PubMed — https://pubmed.ncbi.nlm.nih.gov/38517751/
- **[S4]** Heterodimerization of human apelin and kappa opioid receptors: roles in signal transduction - PubMed — https://pubmed.ncbi.nlm.nih.gov/22200678/
- **[S5]** Cloning, pharmacological characterization and brain distribution of the rat apelin receptor - PubMed — https://pubmed.ncbi.nlm.nih.gov/11146423/
- **[S6]** APJ receptor mRNA expression in the rat hypothalamic paraventricular nucleus: regulation by stress and glucocorticoids - PubMed — https://pubmed.ncbi.nlm.nih.gov/14622440/
- **[S7]** 
            Potential Involvement of Apelin/APJ System in Addiction and Neuroprotection Against Drugs of Abuse - PMC
         — https://pmc.ncbi.nlm.nih.gov/articles/PMC11491857/
- **[S8]** Apelin-13 Protects PC12 Cells Against Methamphetamine-Induced Oxidative Stress, Autophagy and Apoptosis - PubMed — https://pubmed.ncbi.nlm.nih.gov/31385138/
- **[S9]** Evidence for the Induction of Analgesic Cross-Tolerance Between Opioid and Apelin/APJ Systems in Male Rats - PubMed — https://pubmed.ncbi.nlm.nih.gov/38517751/
- **[S10]** Aplnr knockout mice display sex-specific changes in conditioned fear - PubMed — https://pubmed.ncbi.nlm.nih.gov/33309737/
- **[S11]** 
            Potential Involvement of Apelin/APJ System in Addiction and Neuroprotection Against Drugs of Abuse - PMC
         — https://pmc.ncbi.nlm.nih.gov/articles/PMC11491857/
- **[S12]** 
            Neurobiology of addiction: a neurocircuitry analysis - PMC
         — https://pmc.ncbi.nlm.nih.gov/articles/PMC6135092/
- **[S13]** The apelin receptor: physiology, pathology, cell signalling, and ligand modulation of a peptide-activated class A GPCR - PubMed — https://pubmed.ncbi.nlm.nih.gov/25275559/
- **[S14]** Cloning, pharmacological characterization and brain distribution of the rat apelin receptor - PubMed — https://pubmed.ncbi.nlm.nih.gov/11146423/
- **[S15]** Gene expression is altered in the lateral hypothalamus upon activation of the mu opioid receptor - PubMed — https://pubmed.ncbi.nlm.nih.gov/18591478/
- **[S16]** Gene expression is altered in the lateral hypothalamus upon activation of the mu opioid receptor - PubMed — https://pubmed.ncbi.nlm.nih.gov/18591478/
