## Supplementary material for "Understanding Substance Dependence: What Differentiates Addictive from Non-Addictive Drugs?": Folder for supplementary files: addictive-genes-claims-first100-023-pmp2-gene-protein-is-implicated-in-addiction-198b7a0f9381.md

# addictive-genes-claims-first100-023-198b7a0f9381

**Original claim:** PMP2 gene/protein is implicated in addiction.

## Claim

PMP2 gene/protein is implicated in addiction.

## Verdict

Verdict: No Evidence

Based on the scholarly and authoritative sources I could identify, **there is currently no direct evidence that PMP2 is an established addiction gene/protein**. The literature consistently describes PMP2 as a **myelin-associated, lipid-binding protein** with roles in **Schwann-cell biology, myelination, remyelination, and Charcot-Marie-Tooth (CMT) neuropathy**, not as a replicated risk gene, biomarker, or mechanistic driver for substance use disorders. ([ncbi.nlm.nih.gov](https://www.ncbi.nlm.nih.gov/gene/5375)) [S1]

## Confidence

Medium

The confidence is **medium** because several large addiction-genetics and addiction-omics studies do **not** surface PMP2, but “absence from major studies” is not the same as a formal proof of no involvement. ([pmc.ncbi.nlm.nih.gov](https://pmc.ncbi.nlm.nih.gov/articles/PMC10217792/)) [S2]

## Supporting evidence

What **does** support the claim is only **indirect biological plausibility**, not addiction-specific evidence.

- **PMP2 encodes peripheral myelin protein 2**, a protein that localizes to **myelin sheaths of the peripheral nervous system**, binds lipids/membranes, and is thought to help stabilize myelin. NCBI also lists **dominant demyelinating CMT neuropathy** as the established disease link for defects in this gene. ([ncbi.nlm.nih.gov](https://www.ncbi.nlm.nih.gov/gene/5375)) [S3]
- A recent review states that P2/PMP2 is a **fatty-acid-binding protein** expressed in vertebrate peripheral myelin and **human astrocytes**, with suggested functions in **lipid transport** and **membrane stacking**. The same review frames the major human disease connection as **Charcot-Marie-Tooth disease**, not addiction. ([pubmed.ncbi.nlm.nih.gov](https://pubmed.ncbi.nlm.nih.gov/34138518/)) [S4]
- Recent mechanistic studies show PMP2 can enhance **fatty-acid uptake**, **mitochondrial ATP production**, and **myelination/remyelination** in Schwann cells, supporting a role in glial metabolic homeostasis. ([pubmed.ncbi.nlm.nih.gov](https://pubmed.ncbi.nlm.nih.gov/38311982/?utm_source=openai)) [S5]
- Human Protein Atlas data also show **substantial cortical RNA expression** of PMP2 in brain tissue datasets, so the gene is not purely irrelevant to the nervous system. In the cerebral cortex page, consensus RNA expression is reported around **162.6 nTPM**, with GTEx around **139.0 nTPM**. ([proteinatlas.org](https://www.proteinatlas.org/ENSG00000147588-PMP2/brain/cerebral%2Bcortex)) [S6]

Taken together, these findings mean PMP2 is a **real neural/glial biology gene** with plausible relevance to membrane lipid handling and energetic support. Since glia, myelin, and lipid metabolism can modulate neuronal signaling, one can imagine a hypothetical path from PMP2 biology to addiction-related neural function. But that is an **inference**, not direct evidence of involvement in addiction. ([pubmed.ncbi.nlm.nih.gov](https://pubmed.ncbi.nlm.nih.gov/34138518/)) [S7]

## Contradicting or limiting evidence

The strongest limiting evidence is that **major addiction-focused discovery studies do not identify PMP2**.

- In a multivariate GWAS meta-analysis of **over 1 million subjects** spanning problematic alcohol use, problematic tobacco use, cannabis use disorder, and opioid use disorder, the reported addiction-risk loci included genes such as **PDE4B**, but a full-text search of the paper does **not** find PMP2. ([pmc.ncbi.nlm.nih.gov](https://pmc.ncbi.nlm.nih.gov/articles/PMC10217792/)) [S8]
- In a human-brain **proteome-/transcriptome-wide association** analysis of smoking initiation, alcohol use disorder, cannabis use disorder, and opioid use disorder, the study identified significant proteins/genes for these traits, but PMP2 does **not** appear in the text. ([pmc.ncbi.nlm.nih.gov](https://pmc.ncbi.nlm.nih.gov/articles/PMC9630289/)) [S9]
- A recent re-analysis of **brain transcriptomes for substance-associated genes** emphasizes addiction-circuit enrichment and neurotransmission-related pathways, but PMP2 is **not** mentioned in the paper text. ([pmc.ncbi.nlm.nih.gov](https://pmc.ncbi.nlm.nih.gov/articles/PMC10539015/)) [S10]
- A recent human postmortem study of **gene expression differences in alcohol use disorder** identified hundreds of AUD-associated differentially expressed genes and some with GWAS support; PMP2 is **not** mentioned in the text. ([pubmed.ncbi.nlm.nih.gov](https://pubmed.ncbi.nlm.nih.gov/39394458/)) [S11]
- A 2025 multi-omics Mendelian-randomization study of **problematic alcohol use** prioritized genes/proteins such as **AMT, CCDC88B, PPP6C, SLC4A8, ADH1B, and GCKR** as potential causal targets; PMP2 was **not** found in the text. ([pmc.ncbi.nlm.nih.gov](https://pmc.ncbi.nlm.nih.gov/articles/PMC12478272/)) [S12]
- A large GWAS meta-analysis of **problematic alcohol use** identified **66 associated genes**, including **DRD2, BDNF, ADH4/5/7, and PDE4B**; PMP2 was again **not** found in the paper text. ([pmc.ncbi.nlm.nih.gov](https://pmc.ncbi.nlm.nih.gov/articles/PMC7485556/)) [S13]

Also important: the main disease literature around PMP2 is dominated by **peripheral neuropathy/CMT**, not by alcohol, nicotine, opioid, cocaine, or polysubstance phenotypes. Multiple PMP2 papers focus on neuropathy-causing mutations and myelin dysfunction. ([pubmed.ncbi.nlm.nih.gov](https://pubmed.ncbi.nlm.nih.gov/27009151/?utm_source=openai)) [S14]

So the evidence is not merely “weak”; it is that the gene’s **established literature footprint points somewhere else**. ([ncbi.nlm.nih.gov](https://www.ncbi.nlm.nih.gov/gene/5375)) [S15]

## Mechanistic / biological rationale

A biologically charitable interpretation would be:

1. **Addiction is a brain-circuit disorder** in which glia, lipid metabolism, and myelin integrity can influence synaptic timing, reward circuitry, and neural plasticity. Large addiction transcriptome analyses do highlight brain-region and neurotransmission biology as relevant. ([pmc.ncbi.nlm.nih.gov](https://pmc.ncbi.nlm.nih.gov/articles/PMC10539015/)) [S16]
2. **PMP2 participates in glial lipid handling and myelin support**, and in humans it is reported in astrocytes as well as peripheral myelin-related contexts. ([pubmed.ncbi.nlm.nih.gov](https://pubmed.ncbi.nlm.nih.gov/34138518/)) [S17]
3. Therefore, **if** altered glial lipid homeostasis or myelin remodeling contributes to addiction vulnerability, PMP2 could be considered a speculative upstream candidate. ([pubmed.ncbi.nlm.nih.gov](https://pubmed.ncbi.nlm.nih.gov/34138518/)) [S18]

However, that rationale has major weaknesses:

- Most strong PMP2 evidence is in **Schwann cells/peripheral myelin**, whereas addiction is usually studied via **central reward circuitry**. ([ncbi.nlm.nih.gov](https://www.ncbi.nlm.nih.gov/gene/5375)) [S19]
- Human cortical RNA expression or astrocyte expression does **not** by itself show involvement in drug reward, compulsive use, relapse, craving, or treatment response. ([pubmed.ncbi.nlm.nih.gov](https://pubmed.ncbi.nlm.nih.gov/34138518/)) [S20]
- No convincing human genetic, epigenetic, transcriptomic, proteomic, or behavioral-model study from the retrieved literature directly ties PMP2 to addiction phenotypes. ([pmc.ncbi.nlm.nih.gov](https://pmc.ncbi.nlm.nih.gov/articles/PMC10217792/)) [S21]

So the mechanistic case is **possible but highly speculative**.

## Bottom line

**PMP2 is not currently supported as an addiction-implicated gene/protein.** The literature supports PMP2 as a **myelin/lipid-handling protein** best known for roles in **Schwann-cell metabolism, myelination, remyelination, and Charcot-Marie-Tooth neuropathy**. I found **no direct addiction evidence** and no signal from major modern addiction GWAS or addiction multi-omics papers. At most, there is **indirect, hypothesis-generating plausibility** through glial/myelin biology. ([ncbi.nlm.nih.gov](https://www.ncbi.nlm.nih.gov/gene/5375)) [S22]

## Cited sources

- **[S1]** PMP2 peripheral myelin protein 2 [Homo sapiens (human)] - Gene - NCBI — https://www.ncbi.nlm.nih.gov/gene/5375
- **[S2]** 
            Multivariate genome-wide association meta-analysis of over 1 million subjects identifies loci underlying multiple substance use disorders - PMC
         — https://pmc.ncbi.nlm.nih.gov/articles/PMC10217792/
- **[S3]** PMP2 peripheral myelin protein 2 [Homo sapiens (human)] - Gene - NCBI — https://www.ncbi.nlm.nih.gov/gene/5375
- **[S4]** Human myelin protein P2: from crystallography to time-lapse membrane imaging and neuropathy-associated variants - PubMed — https://pubmed.ncbi.nlm.nih.gov/34138518/
- **[S5]** PMP2 regulates myelin thickening and ATP production during remyelination - PubMed — https://pubmed.ncbi.nlm.nih.gov/38311982/?utm_source=openai
- **[S6]** Brain tissue expression of PMP2 in cerebral cortex - The Human Protein Atlas — https://www.proteinatlas.org/ENSG00000147588-PMP2/brain/cerebral%2Bcortex
- **[S7]** Human myelin protein P2: from crystallography to time-lapse membrane imaging and neuropathy-associated variants - PubMed — https://pubmed.ncbi.nlm.nih.gov/34138518/
- **[S8]** 
            Multivariate genome-wide association meta-analysis of over 1 million subjects identifies loci underlying multiple substance use disorders - PMC
         — https://pmc.ncbi.nlm.nih.gov/articles/PMC10217792/
- **[S9]** 
            Integrating human brain proteomic data with genome-wide association study findings identifies novel brain proteins in substance use traits - PMC
         — https://pmc.ncbi.nlm.nih.gov/articles/PMC9630289/
- **[S10]** 
            Analysis of the brain transcriptome for substance-associated genes: An update on large-scale genome-wide association studies - PMC
         — https://pmc.ncbi.nlm.nih.gov/articles/PMC10539015/
- **[S11]** Gene expression differences associated with alcohol use disorder in human brain - PubMed — https://pubmed.ncbi.nlm.nih.gov/39394458/
- **[S12]** 
            Identification of potential therapeutic targets for problematic alcohol use using multi-omics data - PMC
         — https://pmc.ncbi.nlm.nih.gov/articles/PMC12478272/
- **[S13]** 
            Genome-wide meta-analysis of problematic alcohol use in 435,563 individuals yields insights into biology and relationships with other traits - PMC
         — https://pmc.ncbi.nlm.nih.gov/articles/PMC7485556/
- **[S14]** De novo PMP2 mutations in families with type 1 Charcot-Marie-Tooth disease - PubMed — https://pubmed.ncbi.nlm.nih.gov/27009151/?utm_source=openai
- **[S15]** PMP2 peripheral myelin protein 2 [Homo sapiens (human)] - Gene - NCBI — https://www.ncbi.nlm.nih.gov/gene/5375
- **[S16]** 
            Analysis of the brain transcriptome for substance-associated genes: An update on large-scale genome-wide association studies - PMC
         — https://pmc.ncbi.nlm.nih.gov/articles/PMC10539015/
- **[S17]** Human myelin protein P2: from crystallography to time-lapse membrane imaging and neuropathy-associated variants - PubMed — https://pubmed.ncbi.nlm.nih.gov/34138518/
- **[S18]** Human myelin protein P2: from crystallography to time-lapse membrane imaging and neuropathy-associated variants - PubMed — https://pubmed.ncbi.nlm.nih.gov/34138518/
- **[S19]** PMP2 peripheral myelin protein 2 [Homo sapiens (human)] - Gene - NCBI — https://www.ncbi.nlm.nih.gov/gene/5375
- **[S20]** Human myelin protein P2: from crystallography to time-lapse membrane imaging and neuropathy-associated variants - PubMed — https://pubmed.ncbi.nlm.nih.gov/34138518/
- **[S21]** 
            Multivariate genome-wide association meta-analysis of over 1 million subjects identifies loci underlying multiple substance use disorders - PMC
         — https://pmc.ncbi.nlm.nih.gov/articles/PMC10217792/
- **[S22]** PMP2 peripheral myelin protein 2 [Homo sapiens (human)] - Gene - NCBI — https://www.ncbi.nlm.nih.gov/gene/5375
