## Supplementary material for "Understanding Substance Dependence: What Differentiates Addictive from Non-Addictive Drugs?": Folder for supplementary files: addictive-genes-claims-first100-024-nrxn2-gene-protein-is-implicated-in-addiction-a3d18ba9d349.md

# addictive-genes-claims-first100-024-a3d18ba9d349

**Original claim:** NRXN2 gene/protein is implicated in addiction.

## Claim

NRXN2 gene/protein is implicated in addiction.

## Verdict

Verdict: Mixed Evidence

There is **some evidence** linking **NRXN2** to addiction-related phenotypes, but it is **limited, mostly indirect, and not strongly replicated across major human addiction genetics studies**. The clearest direct signal is a **candidate-gene rare-variant association with smoking status/nicotine dependence-related phenotype** in one human study; there is also **indirect preclinical evidence** from alcohol-vulnerability epigenetics in nonhuman primates. However, the broader addiction genetics literature has much stronger and more replicated support for other loci, and within the neurexin family, **NRXN3** is much more consistently tied to substance-use phenotypes than **NRXN2**. ([pubmed.ncbi.nlm.nih.gov](https://pubmed.ncbi.nlm.nih.gov/25450229/?utm_source=openai)) [S1]

## Confidence

Medium

## Supporting evidence

A human candidate-gene sequencing/genotyping study of nicotine dependence-related traits reported that **rare variants in NRXN2 were significantly associated with smoking status in African American participants** from the Mid-South Tobacco Case-Control sample. Importantly, the same paper also reported that **none of the common NRXN2 variants tested were significant after Bonferroni correction**, so the positive signal was restricted to a rare-variant burden analysis rather than a robust common-variant effect. ([pubmed.ncbi.nlm.nih.gov](https://pubmed.ncbi.nlm.nih.gov/25450229/?utm_source=openai)) [S2]

There is also **indirect preclinical support** from a longitudinal rhesus macaque study. In **alcohol-naïve** animals, differential DNA methylation patterns in dorsolateral prefrontal cortex that predicted **future risky ethanol consumption** included synaptic genes such as **NRXN2** and **NRXN3**. However, the authors explicitly stated that the **functional implications of these DMRs were not yet known**, and they did **not** report a corresponding differential-expression signal for the same animals at baseline. That makes this evidence suggestive of vulnerability biology, not proof that NRXN2 is a causal addiction gene. ([pmc.ncbi.nlm.nih.gov](https://pmc.ncbi.nlm.nih.gov/articles/PMC12044430/)) [S3]

At a broader family level, reviews of neurexin biology note that **NRXN-family variants have been linked to neuropsychiatric disorders including addiction**, which makes NRXN2 a biologically plausible candidate within a synaptic-adhesion family relevant to circuit plasticity. But those family-level statements do not by themselves establish a strong NRXN2-specific addiction effect. ([pubmed.ncbi.nlm.nih.gov](https://pubmed.ncbi.nlm.nih.gov/29221905/?utm_source=openai)) [S4]

## Contradicting or limiting evidence

The biggest limitation is **lack of strong replication in large-scale human addiction genomics**. Very large GWAS/meta-analyses of tobacco and alcohol use have identified many robust loci and genes, including well-known smoking loci such as **CHRNA5-CHRNA3-CHRNB4** and others, but **NRXN2 has not emerged as a prominent, repeatedly replicated lead locus in the available summaries of those studies**. That is an inference from the major published results, not proof of no effect. ([pubmed.ncbi.nlm.nih.gov](https://pubmed.ncbi.nlm.nih.gov/30643251/?utm_source=openai)) [S5]

Likewise, an integrated review of genomic and transcriptomic studies across alcohol, nicotine, and other drug-use phenotypes identified a small set of **shared genes** and pathways across substance-use traits, but **NRXN2 was not among the highlighted cross-substance shared genes**. This again suggests that, if NRXN2 contributes, its effect is likely **modest, context-specific, ancestry-specific, or otherwise not yet robustly established**. ([pubmed.ncbi.nlm.nih.gov](https://pubmed.ncbi.nlm.nih.gov/35491750/?utm_source=openai)) [S6]

A second major limitation is **specificity within the neurexin family**. Reviews focused on addiction-related cell-adhesion genes point much more strongly to **NRXN3** than to NRXN2, citing NRXN3 associations with nicotine, alcohol, opioid dependence, and cocaine-related expression changes. In other words, some literature that says “neurexins are implicated in addiction” is really driven more by **NRXN3** than by NRXN2. ([pmc.ncbi.nlm.nih.gov](https://pmc.ncbi.nlm.nih.gov/articles/PMC5838467/?utm_source=openai)) [S7]

Even neurexin-focused disease reviews emphasize that **NRXN1-related disease is much more common** than NRXN2- or NRXN3-related disease, and note **substance-abuse associations for NRXN3 SNPs** rather than clearly established NRXN2 addiction associations. ([rcastoragev2.blob.core.windows.net](https://rcastoragev2.blob.core.windows.net/b57401280c418fe5aff36e900754d009/PMC8491281.pdf?utm_source=openai)) [S8]

## Mechanistic / biological rationale

Mechanistically, **NRXN2 is a presynaptic neurexin**, part of a trans-synaptic adhesion system that helps organize synapses and shape neurotransmission. SynGO annotates NRXN2 to **presynaptic modulation of chemical synaptic transmission**, and primary studies show that Nrxn2 influences **excitatory synapse number, presynaptic release probability, and NMDAR-related synaptic physiology**. ([syngoportal.org](https://www.syngoportal.org/gene_hgnc8009.html?utm_source=openai)) [S9]

That matters because addiction is widely understood as a disorder of **maladaptive synaptic plasticity and circuit-level transcriptional remodeling** in reward-related brain regions. Cocaine self-administration studies, for example, show transcriptome-wide changes across reward circuitry that correlate with addiction-like behaviors. So, from a biological standpoint, a synaptic organizer such as NRXN2 is a **credible candidate** for modulating addiction vulnerability or neuroadaptation. ([pubmed.ncbi.nlm.nih.gov](https://pubmed.ncbi.nlm.nih.gov/29861096/)) [S10]

But plausibility is not the same as proof. In fact, NRXN2 biology is complex: newer work suggests Nrxn2 can **restrict excitatory synapse formation and restrain release probability** in some hippocampal contexts, rather than simply promoting synaptogenesis. That complexity supports a possible role in neural homeostasis and plasticity, but it also means the direction and behavioral consequences of NRXN2 perturbation may depend heavily on **cell type, splice isoform, circuit, and developmental stage**. ([pmc.ncbi.nlm.nih.gov](https://pmc.ncbi.nlm.nih.gov/articles/PMC10068831/)) [S11]

## Bottom line

**NRXN2 is not a strongly established addiction gene**, but saying it is **entirely unrelated** to addiction would also go too far. The current literature supports a **tentative, biologically plausible, but weakly substantiated** link: one notable human rare-variant signal for smoking status, some indirect alcohol-vulnerability epigenetic evidence, and strong synaptic-plasticity rationale. However, the evidence falls short of the level seen for **major replicated addiction loci** or even for **NRXN3** within the same gene family. Overall, the claim is best judged as **mixed evidence** rather than clearly true or clearly false. ([pubmed.ncbi.nlm.nih.gov](https://pubmed.ncbi.nlm.nih.gov/25450229/?utm_source=openai)) [S12]

## Cited sources

- **[S1]** The contribution of rare and common variants in 30 genes to risk nicotine dependence - PubMed — https://pubmed.ncbi.nlm.nih.gov/25450229/?utm_source=openai
- **[S2]** The contribution of rare and common variants in 30 genes to risk nicotine dependence - PubMed — https://pubmed.ncbi.nlm.nih.gov/25450229/?utm_source=openai
- **[S3]** 
            Pre-existing DNA methylation signatures in the prefrontal cortex of alcohol-naïve nonhuman primates define neural vulnerability for future risky ethanol consumption - PMC
         — https://pmc.ncbi.nlm.nih.gov/articles/PMC12044430/
- **[S4]** Neurexins and neuropsychiatric disorders - PubMed — https://pubmed.ncbi.nlm.nih.gov/29221905/?utm_source=openai
- **[S5]** Association studies of up to 1.2 million individuals yield new insights into the genetic etiology of tobacco and alcohol use - PubMed — https://pubmed.ncbi.nlm.nih.gov/30643251/?utm_source=openai
- **[S6]** Common and distinguishing genetic factors for substance use behavior and disorder: an integrated analysis of genomic and transcriptomic studies from both human and animal studies - PubMed — https://pubmed.ncbi.nlm.nih.gov/35491750/?utm_source=openai
- **[S7]** The Role of Cell Adhesion Molecule Genes Regulating Neuroplasticity in Addiction - PMC — https://pmc.ncbi.nlm.nih.gov/articles/PMC5838467/?utm_source=openai
- **[S8]** HHS Public Access — https://rcastoragev2.blob.core.windows.net/b57401280c418fe5aff36e900754d009/PMC8491281.pdf?utm_source=openai
- **[S9]** SynGO - gene details — https://www.syngoportal.org/gene_hgnc8009.html?utm_source=openai
- **[S10]** Cocaine Self-administration Alters Transcriptome-wide Responses in the Brain's Reward Circuitry - PubMed — https://pubmed.ncbi.nlm.nih.gov/29861096/
- **[S11]** 
            Neurexin-2 restricts synapse numbers and restrains the presynaptic release probability by an alternative splicing-dependent mechanism - PMC
         — https://pmc.ncbi.nlm.nih.gov/articles/PMC10068831/
- **[S12]** The contribution of rare and common variants in 30 genes to risk nicotine dependence - PubMed — https://pubmed.ncbi.nlm.nih.gov/25450229/?utm_source=openai
