## Supplementary material for "Understanding Substance Dependence: What Differentiates Addictive from Non-Addictive Drugs?": Folder for supplementary files: addictive-genes-claims-first100-025-skp1-gene-protein-is-implicated-in-addiction-4d43d91dcbd9.md

# addictive-genes-claims-first100-025-4d43d91dcbd9

**Original claim:** SKP1 gene/protein is implicated in addiction.

## Claim

SKP1 gene/protein is implicated in addiction.

## Verdict

Verdict: No Evidence

## Confidence

Medium

## Supporting evidence

SKP1 is a real **biologically plausible** candidate in the broadest sense: it encodes a core adaptor component of the **SCF (SKP1–CUL1–F-box) E3 ubiquitin ligase machinery**, a major protein-degradation system, and it is expressed in human brain, including the **nucleus accumbens**, a key reward/addiction region. That makes it reasonable to ask whether SKP1 might matter for addiction biology. ([ncbi.nlm.nih.gov](https://www.ncbi.nlm.nih.gov/gene/6500?utm_source=openai)) [S1]

There is also solid evidence that the **ubiquitin-proteasome system (UPS)** more broadly is involved in addiction-related neuroplasticity. A review of UPS in drug abuse describes drug-induced regulation of UPS components, UPS-dependent synaptic remodeling, and behavioral data showing that **proteasome inhibition in nucleus accumbens disrupted acquisition of opioid-conditioned behaviors and context-dependent sensitization**. So, the pathway SKP1 belongs to is relevant to addiction mechanisms at a systems level. ([frontiersin.org](https://www.frontiersin.org/journals/molecular-neuroscience/articles/10.3389/fnmol.2014.00099/pdf)) [S2]

I also found one **very indirect** signal involving SKP1 in an alcohol-related context: in lymphoblastoid cell lines derived from alcoholics versus controls, an enriched **protein ubiquitination pathway** included **SKP1** among genes differing by phenotype. However, this was not brain tissue, not a direct addiction-gene association test, and not evidence that SKP1 itself drives addictive behavior. At most, it is a weak omics hint that SKP1 can appear in alcohol-related molecular signatures. ([downstate.edu](https://www.downstate.edu/research/centers-departments/henri-begleiter-neurodynamics-laboratory/_documents/2014-McClintick-Ethanoltreatmentoflymphoblastoidcelllinesfromalcoholicsandnon-alcoholicscaus.pdf)) [S3]

## Contradicting or limiting evidence

The major limitation is that I did **not find direct scholarly evidence** showing that SKP1 itself is associated with addiction risk, differentially and reproducibly altered in addiction-relevant human brain tissue, or causally modulates addiction-like behavior when experimentally manipulated. In other words, the evidence supports the **pathway** more than the **specific gene**. ([frontiersin.org](https://www.frontiersin.org/journals/molecular-neuroscience/articles/10.3389/fnmol.2014.00099/pdf)) [S4]

Recent, higher-level genetics reviews of substance use disorders highlight other genes/loci instead. For example, a recent Molecular Psychiatry review summarizes robust signals such as **ADH1B/ADH1C** for alcohol use disorder, **CHRNA2/FOXP2** for cannabis use disorder, and **OPRM1** plus loci involving **FURIN** and **SCAI/PPP6C/RABEPK** for opioid use disorder. A JCI review of opioid addiction genetics similarly emphasizes **OPRM1, FURIN, and SCAI/PPP6C/RABEPK** as established GWAS-level findings. SKP1 is not presented as a replicated addiction risk gene in these reviews. ([nature.com](https://www.nature.com/articles/s41380-024-02547-z)) [S5]

A GWAS Catalog–derived **Alcohol use disorder** gene set lists 12 associated genes, including **ADH1B, ADH1C, SLC39A8, DRD2, MAP4K5, FTO, SIX3, ADH7, TSPAN5, PKHD1, MICB, and MYC**; a search of that set shows **no SKP1 entry**. This does not prove SKP1 has no role, but it argues against SKP1 being an established AUD gene by current GWAS evidence. ([maayanlab.cloud](https://maayanlab.cloud/Harmonizome/gene_set/Alcohol%2Buse%2Bdisorder/GWAS%2BCatalog%2BSNP-Phenotype%2BAssociations%2B2025)) [S6]

Similarly, a recent meta-analysis of **20,098 genes** in human prefrontal cortex from opioid overdose/opioid addiction-related cases found **335 significant DEGs** and highlighted genes/pathways such as **OPRK1, NPAS4, DUSP/EGR family members, NR4A2, SYT1, HCRTR2, BDNF**, and orexin/RTK/MAPK signaling. The paper explicitly states there was **little evidence of genetic drivers** for most of the observed differential expression. SKP1 was not among the genes emphasized in the study’s main findings. ([pmc.ncbi.nlm.nih.gov](https://pmc.ncbi.nlm.nih.gov/articles/PMC10802752/)) [S7]

## Mechanistic / biological rationale

Biologically, the claim is **plausible but unproven**. SKP1 is a central adaptor in SCF ubiquitin ligases, and SCF-mediated ubiquitination controls turnover of many proteins involved in signaling, transcription, and cellular homeostasis. Because addiction depends on long-lasting synaptic and transcriptional remodeling, any core UPS component could in principle influence addiction-related plasticity. ([ncbi.nlm.nih.gov](https://www.ncbi.nlm.nih.gov/gene/6500?utm_source=openai)) [S8]

That said, this rationale is still **generic**. SKP1 is a broadly used cellular adaptor, not a molecule currently known to be selectively tied to drug reward, craving, relapse, withdrawal, or reinforcement. High expression in the nucleus accumbens and participation in proteostasis are not enough, by themselves, to say that SKP1 is “implicated in addiction” in the same sense as genes like **OPRM1** or repeatedly identified pathway mediators in human and animal addiction studies. ([proteinatlas.org](https://www.proteinatlas.org/ENSG00000113558-SKP1/brain/nucleus%2Baccumbens?utm_source=openai)) [S9]

So the best mechanistic interpretation is: **SKP1 may be relevant to neural homeostasis and protein-turnover processes that addiction co-opts**, but current evidence does not show that SKP1 is a specific addiction gene/protein. That distinction matters. A housekeeping component of a pathway involved in addiction is not automatically an addiction-implicated gene unless there is direct association or intervention evidence. ([frontiersin.org](https://www.frontiersin.org/journals/molecular-neuroscience/articles/10.3389/fnmol.2014.00099/pdf)) [S10]

## Bottom line

The current literature does **not** support a strong statement that **SKP1 itself is implicated in addiction**. What is supported is a weaker claim: **SKP1 belongs to the ubiquitin-proteasome machinery, and that machinery is relevant to addiction-related synaptic plasticity and behavior**. I found only **indirect** SKP1-related signals, not direct human genetic, human brain, or causal animal evidence tying SKP1 specifically to addiction. On present evidence, SKP1 is better described as a **biologically plausible but unvalidated upstream/homeostatic protein**, not an established addiction gene/protein. ([frontiersin.org](https://www.frontiersin.org/journals/molecular-neuroscience/articles/10.3389/fnmol.2014.00099/pdf)) [S11]

## Cited sources

- **[S1]** SKP1 S-phase kinase associated protein 1 [Homo sapiens (human)] - Gene - NCBI — https://www.ncbi.nlm.nih.gov/gene/6500?utm_source=openai
- **[S2]** Roles of the ubiquitin proteasome system in the effects of drugs of abuse — https://www.frontiersin.org/journals/molecular-neuroscience/articles/10.3389/fnmol.2014.00099/pdf
- **[S3]** Ethanol treatment of lymphoblastoid cell lines from alcoholics and non-alcoholics causes many subtle changes in gene expression — https://www.downstate.edu/research/centers-departments/henri-begleiter-neurodynamics-laboratory/_documents/2014-McClintick-Ethanoltreatmentoflymphoblastoidcelllinesfromalcoholicsandnon-alcoholicscaus.pdf
- **[S4]** Roles of the ubiquitin proteasome system in the effects of drugs of abuse — https://www.frontiersin.org/journals/molecular-neuroscience/articles/10.3389/fnmol.2014.00099/pdf
- **[S5]** The genetic landscape of substance use disorders | Molecular Psychiatry — https://www.nature.com/articles/s41380-024-02547-z
- **[S6]** Gene Set - Alcohol use disorder — https://maayanlab.cloud/Harmonizome/gene_set/Alcohol%2Buse%2Bdisorder/GWAS%2BCatalog%2BSNP-Phenotype%2BAssociations%2B2025
- **[S7]** 
            Identifying novel gene dysregulation associated with opioid overdose death: A meta-analysis of differential gene expression in human prefrontal cortex - PMC
         — https://pmc.ncbi.nlm.nih.gov/articles/PMC10802752/
- **[S8]** SKP1 S-phase kinase associated protein 1 [Homo sapiens (human)] - Gene - NCBI — https://www.ncbi.nlm.nih.gov/gene/6500?utm_source=openai
- **[S9]** Brain tissue expression of SKP1 in nucleus accumbens - The Human Protein Atlas — https://www.proteinatlas.org/ENSG00000113558-SKP1/brain/nucleus%2Baccumbens?utm_source=openai
- **[S10]** Roles of the ubiquitin proteasome system in the effects of drugs of abuse — https://www.frontiersin.org/journals/molecular-neuroscience/articles/10.3389/fnmol.2014.00099/pdf
- **[S11]** Roles of the ubiquitin proteasome system in the effects of drugs of abuse — https://www.frontiersin.org/journals/molecular-neuroscience/articles/10.3389/fnmol.2014.00099/pdf
