## Supplementary material for "Understanding Substance Dependence: What Differentiates Addictive from Non-Addictive Drugs?": Folder for supplementary files: addictive-genes-claims-first100-026-hhatl-gene-protein-is-implicated-in-addiction-26e33a9705e5.md

# addictive-genes-claims-first100-026-26e33a9705e5

**Original claim:** HHATL gene/protein is implicated in addiction.

## Claim

HHATL gene/protein is implicated in addiction.

## Verdict

Verdict: No Evidence

## Confidence

Medium. The direct addiction literature for **HHATL** appears extremely sparse to absent in the sources I could find, but absence of retrieved evidence is not absolute proof of no role.

## Supporting evidence

HHATL is a real, biologically characterized gene/protein, but the literature I found links it to **Hedgehog pathway regulation, muscle/cardiac biology, and cellular stress responses** rather than addiction phenotypes. NCBI summarizes human **HHATL** as a protein-coding gene predicted to be involved in **negative regulation of N-terminal protein palmitoylation**, with expression reported as **restricted toward heart**. A foundational mechanistic paper identified mammalian Gup1/HHATL as a **negative regulator of Sonic hedgehog (Shh) N-terminal palmitoylation** in cells. ([ncbi.nlm.nih.gov](https://www.ncbi.nlm.nih.gov/gene/57467)) [S1]

Follow-up functional studies also point away from addiction as the primary known biology. In mice, **Hhatl/Mitsugumin-56** was described as a sarcoplasmic-reticulum resident protein essential for **postnatal muscle maturation**. In zebrafish, reduced **hhatla** promoted **cardiac hypertrophy-related defects**. More recently, cell biology work reported that Hhatl can reduce **ER stress** by promoting **autophagy via LC3 interaction**. These are meaningful biological findings, but they do not constitute evidence for a role in substance use disorders or reward-related behavior. ([pubmed.ncbi.nlm.nih.gov](https://pubmed.ncbi.nlm.nih.gov/25841338/?utm_source=openai)) [S2]

There is at least some evidence that **HHATL transcripts are present in brain tissue**, especially in parts of the brainstem, which means a neural role is not impossible. The Human Protein Atlas shows substantial **RNA expression in pons/medulla subregions**, although it reports **no protein-expression data** there on that page. NCBI, however, still summarizes overall expression as heart-restricted at the tissue level, underscoring that HHATL is not currently recognized as a prominent addiction-circuit gene. ([proteinatlas.org](https://www.proteinatlas.org/ENSG00000010282-HHATL/brain/pons%2Band%2Bmedulla)) [S3]

A broader, indirect point is that **Hedgehog-related signaling can intersect addiction-relevant circuitry**. For example, a 2022 PNAS study showed that **HHIP**—a different Hedgehog-pathway component, not HHATL—is highly enriched in **medial habenula** cholinergic neurons and that disrupting **Hhip** in these neurons enhanced nicotine’s motivational effects in mice. That establishes that **some Hedgehog-pathway biology** can affect nicotine-related behavior, but it does **not** specifically implicate HHATL. ([pubmed.ncbi.nlm.nih.gov](https://pubmed.ncbi.nlm.nih.gov/36346845/)) [S4]

## Contradicting or limiting evidence

The main limitation for the claim is simple: I did **not** find direct evidence that **HHATL** is associated with addiction in the usual evidentiary categories—human GWAS, candidate-gene studies, addiction transcriptomics, animal addiction models, or mechanistic addiction experiments. Instead, the HHATL literature retrieved was centered on **Shh palmitoylation**, **muscle/cardiac phenotypes**, and **ER-stress/autophagy biology**. That pattern argues that the statement “HHATL is implicated in addiction” is currently **unsubstantiated** in the mainstream literature I found. ([pubmed.ncbi.nlm.nih.gov](https://pubmed.ncbi.nlm.nih.gov/18081866/?utm_source=openai)) [S5]

Recent reviews of **substance use disorder genetics** emphasize a highly polygenic architecture and highlight established loci such as **ADH1B/ALDH2** for alcohol-related traits and **CHRNA5-CHRNA3-CHRNB4** for nicotine-related traits. These reviews discuss major advances across alcohol, tobacco, opioid, cannabis, and cocaine genetics, but the retrieved review material does not identify **HHATL** as an established or replicated addiction gene. That does not prove HHATL has no role, but it does weigh against the claim that it is already “implicated” in addiction in the same sense as recognized SUD loci. ([pubmed.ncbi.nlm.nih.gov](https://pubmed.ncbi.nlm.nih.gov/33879270/)) [S6]

Similarly, addiction-focused curated resources describe compilation of genes from GWAS, candidate-gene, transcriptomic, methylation, and literature-mining evidence for alcohol, nicotine, cocaine, and opioid dependence. In the materials I could retrieve about such resources, I found descriptions of the database framework, but no clear evidence placing HHATL among well-supported addiction genes. Again, this is **inferential** rather than definitive, because database querying through the browser was limited. ([pubmed.ncbi.nlm.nih.gov](https://pubmed.ncbi.nlm.nih.gov/30403795/?utm_source=openai)) [S7]

## Mechanistic / biological rationale

If one wanted to argue for a **possible** link, the best rationale would be **indirect**. HHATL modulates **Hedgehog signaling-related biochemistry** by interfering with Shh palmitoylation, and Hedgehog signaling is a major regulator of development and can remain relevant in adult tissues, including the nervous system. ([pubmed.ncbi.nlm.nih.gov](https://pubmed.ncbi.nlm.nih.gov/18081866/?utm_source=openai)) [S8]

That matters because addiction risk and progression involve long-term changes in neural circuits, synaptic signaling, and cellular homeostasis. A pathway that shapes neural development or plasticity could, in principle, affect addiction vulnerability. The HHIP-habenula-nicotine study adds a biologically plausible bridge: a Hedgehog-pathway regulator can modulate **nicotine aversion/motivation circuitry** in the habenula. However, that bridge is still **one step removed** from HHATL, and I found no study showing that manipulating HHATL changes drug taking, withdrawal, craving, reinforcement, or relapse-like behavior. ([pubmed.ncbi.nlm.nih.gov](https://pubmed.ncbi.nlm.nih.gov/36346845/)) [S9]

HHATL’s more recent connection to **ER stress and autophagy** also offers only a speculative route. ER stress, autophagy, and proteostasis are relevant to neuronal homeostasis broadly, and dysregulation of these processes has been discussed in many neuropsychiatric conditions. But the retrieved HHATL-autophagy study was not addiction-focused, and translating generic cellular-stress biology into a claim about addiction would be an overreach. ([pubmed.ncbi.nlm.nih.gov](https://pubmed.ncbi.nlm.nih.gov/38705394/?utm_source=openai)) [S10]

So mechanistically, the most defensible statement is: **HHATL has functions that make a neural role conceivable, but there is no direct evidence here that those functions translate into addiction relevance.** ([ncbi.nlm.nih.gov](https://www.ncbi.nlm.nih.gov/gene/57467)) [S11]

## Bottom line

Based on the best scholarly and authoritative sources I could find, **HHATL is not currently supported as an addiction-implicated gene/protein**. The literature supports roles in **Hedgehog-related palmitoylation control, muscle/cardiac biology, and ER-stress/autophagy**, with some transcript presence in brain tissue, but I found **no direct addiction evidence** for HHATL itself. The closest mechanistic support is indirect, via other Hedgehog-pathway components such as **HHIP** in nicotine-related habenular circuits. For the claim as stated, **“No Evidence”** is the most accurate verdict. ([ncbi.nlm.nih.gov](https://www.ncbi.nlm.nih.gov/gene/57467)) [S12]

## Cited sources

- **[S1]** HHATL hedgehog acyltransferase like [Homo sapiens (human)] - Gene - NCBI — https://www.ncbi.nlm.nih.gov/gene/57467
- **[S2]** Mitsugumin 56 (hedgehog acyltransferase-like) is a sarcoplasmic reticulum-resident protein essential for postnatal muscle maturation - PubMed — https://pubmed.ncbi.nlm.nih.gov/25841338/?utm_source=openai
- **[S3]** Brain tissue expression of HHATL in pons and medulla - The Human Protein Atlas — https://www.proteinatlas.org/ENSG00000010282-HHATL/brain/pons%2Band%2Bmedulla
- **[S4]** Hedgehog-interacting protein acts in the habenula to regulate nicotine intake - PubMed — https://pubmed.ncbi.nlm.nih.gov/36346845/
- **[S5]** Mammalian Gup1, a homolog of Saccharomyces cerevisiae glycerol uptake/transporter 1, acts as a negative regulator for N-terminal palmitoylation of Sonic hedgehog - PubMed — https://pubmed.ncbi.nlm.nih.gov/18081866/?utm_source=openai
- **[S6]** Genetics of substance use disorders: a review - PubMed — https://pubmed.ncbi.nlm.nih.gov/33879270/
- **[S7]** ANCO-GeneDB: annotations and comprehensive analysis of candidate genes for alcohol, nicotine, cocaine and opioid dependence - PubMed — https://pubmed.ncbi.nlm.nih.gov/30403795/?utm_source=openai
- **[S8]** Mammalian Gup1, a homolog of Saccharomyces cerevisiae glycerol uptake/transporter 1, acts as a negative regulator for N-terminal palmitoylation of Sonic hedgehog - PubMed — https://pubmed.ncbi.nlm.nih.gov/18081866/?utm_source=openai
- **[S9]** Hedgehog-interacting protein acts in the habenula to regulate nicotine intake - PubMed — https://pubmed.ncbi.nlm.nih.gov/36346845/
- **[S10]** Hhatl ameliorates endoplasmic reticulum stress through autophagy by associating with LC3 - PubMed — https://pubmed.ncbi.nlm.nih.gov/38705394/?utm_source=openai
- **[S11]** HHATL hedgehog acyltransferase like [Homo sapiens (human)] - Gene - NCBI — https://www.ncbi.nlm.nih.gov/gene/57467
- **[S12]** HHATL hedgehog acyltransferase like [Homo sapiens (human)] - Gene - NCBI — https://www.ncbi.nlm.nih.gov/gene/57467
