## Supplementary material for "Understanding Substance Dependence: What Differentiates Addictive from Non-Addictive Drugs?": Folder for supplementary files: addictive-genes-claims-first100-027-rps5-gene-protein-is-implicated-in-addiction-ea4199fecbe5.md

# addictive-genes-claims-first100-027-ea4199fecbe5

**Original claim:** RPS5 gene/protein is implicated in addiction.

## Claim

RPS5 gene/protein is implicated in addiction.

## Verdict

Verdict: Mixed Evidence

## Confidence

Medium

## Supporting evidence

RPS5 does show up in some **alcohol-related molecular studies**, but the signal is weak and indirect rather than addiction-specific. In a postmortem hippocampus study of chronic alcoholics, altered **EIF2 signaling** included **RPS5** among a broader set of ribosomal genes within larger changes involving stress, inflammation, glial/myelin biology, and glucocorticoid signaling. That places RPS5 inside an alcohol-associated molecular signature, but not as a singled-out driver gene. ([pmc.ncbi.nlm.nih.gov](https://pmc.ncbi.nlm.nih.gov/articles/PMC3836826/)) [S1]

There is also a **peripheral epigenetic association**: an epigenome-wide association study in saliva from 1,135 European American male veterans reported a promoter-proximal CpG annotated to **RPS5** (cg11701312) associated with alcohol consumption measured by AUDIT-C score. The same paper found that a subset of alcohol-consumption-associated CpGs also differed between hazardous and non-hazardous drinking groups, although the methylation differences were small and the tissue was peripheral rather than brain. ([pmc.ncbi.nlm.nih.gov](https://pmc.ncbi.nlm.nih.gov/articles/PMC9377208/)) [S2]

Additional support is even more indirect. In cultured oligodendrocytes, alcohol exposure decreased **Rps5** expression together with other translation-related genes, consistent with alcohol affecting ribosome/protein-synthesis programs. Again, this supports alcohol responsiveness of RPS5-linked biology, but not a demonstrated role in addiction behavior or vulnerability. ([pmc.ncbi.nlm.nih.gov](https://pmc.ncbi.nlm.nih.gov/articles/PMC10542481/?utm_source=openai)) [S3]

## Contradicting or limiting evidence

The strongest limitation is that I did **not find direct evidence** that RPS5 is a replicated **addiction gene/protein** in the usual sense used in addiction biology: no clear human genetic association repeatedly implicating RPS5 in substance use disorder risk, and no direct functional study showing that manipulating RPS5 changes drug taking, withdrawal, craving, relapse, or reinforcement. The available positive findings are mainly incidental appearances in broader alcohol-response omics datasets rather than targeted addiction studies. ([pmc.ncbi.nlm.nih.gov](https://pmc.ncbi.nlm.nih.gov/articles/PMC3836826/)) [S4]

Recent large, addiction-focused human brain studies also do **not** elevate RPS5 as a notable signal. A 2024 alcohol use disorder methylation study in the **nucleus accumbens** and **dorsolateral prefrontal cortex** identified 105 AUD-associated CpGs annotated to 120 genes and emphasized overlap with prior substance-use loci, but RPS5 was not among the reported genes on the source page. ([pubmed.ncbi.nlm.nih.gov](https://pubmed.ncbi.nlm.nih.gov/39399155/)) [S5]

Likewise, a 2025 multiomic network analysis of **opioid addiction** retained 211 interconnected genes and highlighted pathways centered on **Akt, BDNF, ERK, OPRM1, GABA/glutamate signaling, and ion-channel biology**; RPS5 was not among the reported implicated genes on that page. ([pubmed.ncbi.nlm.nih.gov](https://pubmed.ncbi.nlm.nih.gov/39615775/)) [S6]

A broader curated addiction resource, **ANCO-GeneDB**, integrated alcohol, nicotine, cocaine, and opioid dependence genetics/genomics and highlighted recurrent addiction genes such as **CHRNA5** and regionally enriched signals in anterior cingulate, frontal cortex, amygdala, caudate, putamen, and hypothalamus. In the article summary page, RPS5 was not highlighted among the recurrent addiction genes. ([academic.oup.com](https://academic.oup.com/database/article/doi/10.1093/database/bay121/5161354)) [S7]

So, while RPS5 can appear in alcohol-related signatures, the current literature does **not** support treating it as a well-established addiction locus or addiction effector comparable to canonical addiction-relevant genes/pathways. ([pmc.ncbi.nlm.nih.gov](https://pmc.ncbi.nlm.nih.gov/articles/PMC3836826/)) [S8]

## Mechanistic / biological rationale

Biologically, RPS5 is a **highly conserved 40S ribosomal protein** involved in core translation machinery. Review and mechanistic work describe RPS5 as a conserved ribosomal protein important for ribosome function, and one experimental paper specifically implicated ribosomal protein S5e in **translation initiation** through interaction with **eIF2α**. That means RPS5 could plausibly influence how neurons or glia adjust protein synthesis during chronic drug or alcohol exposure. ([pubmed.ncbi.nlm.nih.gov](https://pubmed.ncbi.nlm.nih.gov/36834797/?utm_source=openai)) [S9]

That said, addiction is usually framed around neuroadaptations in **basal ganglia/reward circuitry, extended amygdala/stress systems, and prefrontal executive-control circuits**, with key roles for dopamine, opioid peptides, glutamate, GABA, stress mediators, and downstream transcription/plasticity programs. Reviews by Koob and Volkow emphasize these systems rather than ubiquitous ribosomal structural proteins as primary addiction determinants. ([pubmed.ncbi.nlm.nih.gov](https://pubmed.ncbi.nlm.nih.gov/27475769/?utm_source=openai)) [S10]

A reasonable interpretation is that RPS5 may participate in **general cellular stress/translation responses** that occur in brains exposed to chronic alcohol or other insults, not necessarily in a pathway specific to addiction. This is consistent with the alcohol studies above, where RPS5 appears as one member of broad ribosomal/EIF2 signatures, not as a uniquely prioritized addiction mediator. ([pmc.ncbi.nlm.nih.gov](https://pmc.ncbi.nlm.nih.gov/articles/PMC3836826/)) [S11]

There are also non-neural studies showing RPS5 can modulate signaling pathways such as **Akt**, **p38**, and **NF-κB** in fibrosis/inflammation models. Those pathways overlap conceptually with mechanisms often discussed in addiction and neuroinflammation, but the evidence comes from liver, cardiac, and pulmonary contexts rather than addiction-relevant brain experiments. So this is mechanistic plausibility, not addiction evidence. ([pubmed.ncbi.nlm.nih.gov](https://pubmed.ncbi.nlm.nih.gov/24668691/?utm_source=openai)) [S12]

## Bottom line

The claim is **not strongly supported** as stated. RPS5 has **some indirect alcohol-related omics evidence** and a plausible role in general translation/stress biology, but I found **no convincing direct evidence** that RPS5 is a specific, replicated addiction gene/protein or that altering RPS5 changes core addiction phenotypes. The fairest summary is that RPS5 is, at most, a **weak secondary signal** in some alcohol-related datasets rather than a well-established addiction factor. ([pmc.ncbi.nlm.nih.gov](https://pmc.ncbi.nlm.nih.gov/articles/PMC3836826/)) [S13]

## Cited sources

- **[S1]** 
            Stress-response pathways are altered in the hippocampus of chronic alcoholics - PMC
         — https://pmc.ncbi.nlm.nih.gov/articles/PMC3836826/
- **[S2]** 
            Epigenome-wide DNA Methylation Association Analysis Identified Novel Loci in Peripheral Cells for Alcohol Consumption among European American Male Veterans - PMC
         — https://pmc.ncbi.nlm.nih.gov/articles/PMC9377208/
- **[S3]** Alcohol induces concentration-dependent transcriptomic changes in oligodendrocytes - PMC — https://pmc.ncbi.nlm.nih.gov/articles/PMC10542481/?utm_source=openai
- **[S4]** 
            Stress-response pathways are altered in the hippocampus of chronic alcoholics - PMC
         — https://pmc.ncbi.nlm.nih.gov/articles/PMC3836826/
- **[S5]** Alcohol Use Disorder-Associated DNA Methylation in the Nucleus Accumbens and Dorsolateral Prefrontal Cortex - PubMed — https://pubmed.ncbi.nlm.nih.gov/39399155/
- **[S6]** Multiomic Network Analysis Identifies Dysregulated Neurobiological Pathways in Opioid Addiction - PubMed — https://pubmed.ncbi.nlm.nih.gov/39615775/
- **[S7]** ANCO-GeneDB: annotations and comprehensive analysis of candidate genes for alcohol, nicotine, cocaine and opioid dependence | Database | Oxford Academic — https://academic.oup.com/database/article/doi/10.1093/database/bay121/5161354
- **[S8]** 
            Stress-response pathways are altered in the hippocampus of chronic alcoholics - PMC
         — https://pmc.ncbi.nlm.nih.gov/articles/PMC3836826/
- **[S9]** Eukaryotic Ribosomal Protein S5 of the 40S Subunit: Structure and Function - PubMed — https://pubmed.ncbi.nlm.nih.gov/36834797/?utm_source=openai
- **[S10]** Neurobiology of addiction: a neurocircuitry analysis - PubMed — https://pubmed.ncbi.nlm.nih.gov/27475769/?utm_source=openai
- **[S11]** 
            Stress-response pathways are altered in the hippocampus of chronic alcoholics - PMC
         — https://pmc.ncbi.nlm.nih.gov/articles/PMC3836826/
- **[S12]** Bioactive compound reveals a novel function for ribosomal protein S5 in hepatic stellate cell activation and hepatic fibrosis - PubMed — https://pubmed.ncbi.nlm.nih.gov/24668691/?utm_source=openai
- **[S13]** 
            Stress-response pathways are altered in the hippocampus of chronic alcoholics - PMC
         — https://pmc.ncbi.nlm.nih.gov/articles/PMC3836826/
