## Supplementary material for "Understanding Substance Dependence: What Differentiates Addictive from Non-Addictive Drugs?": Folder for supplementary files: addictive-genes-claims-first100-028-cox8a-gene-protein-is-implicated-in-addiction-56233a903c9f.md

# addictive-genes-claims-first100-028-56233a903c9f

**Original claim:** COX8A gene/protein is implicated in addiction.

## Claim

COX8A gene/protein is implicated in addiction.

## Verdict

Verdict: Mixed Evidence

## Confidence

Medium

## Supporting evidence

COX8A encodes a small nuclear-encoded subunit of cytochrome c oxidase (mitochondrial respiratory chain complex IV), so it is clearly relevant to cellular bioenergetics and neuronal metabolic function. The NCBI Gene record describes COX8A as part of the terminal respiratory enzyme that helps generate the proton gradient used for ATP production, and human loss-of-function causes a neurological mitochondrial disease (mitochondrial complex IV deficiency, Leigh-like syndrome/epilepsy), showing that the gene matters for brain biology. ([ncbi.nlm.nih.gov](https://www.ncbi.nlm.nih.gov/gene?Cmd=DetailsSearch&Db=gene&Term=1351)) [S1]

The most direct addiction-relevant signal I found was in an alcohol model. In adolescent alcohol-preferring rats exposed to binge-like alcohol drinking, **Cox8a** expression in medial prefrontal cortex was increased about 1.20-fold, with reported significance (p = 1.7×10^-3), and the gene was included in the study’s significantly altered “oxidative phosphorylation” and “mitochondrial dysfunction” pathways. Because the medial prefrontal cortex is a key circuit node for control, valuation, and compulsive drug-seeking, this is legitimate but still modest evidence that COX8A can change in an addiction-relevant brain context. ([pmc.ncbi.nlm.nih.gov](https://pmc.ncbi.nlm.nih.gov/articles/PMC5851801/)) [S2]

More broadly, curated addiction resources do contain many transcriptomic and genetic datasets for alcohol, nicotine, cocaine, and opioid phenotypes, indicating that mitochondrial/energy-metabolism pathways are repeatedly examined in addiction research. The ANCO-GeneDB documentation states that it aggregates addiction-related genes from genetics, transcriptomics, and epigenetics studies across alcohol, nicotine, cocaine, and opioid dependence, and its analyses page lists multiple human brain differential-expression datasets for alcohol dependence and cocaine addiction. That context makes it biologically plausible that a mitochondrial complex IV subunit like COX8A could appear as a secondary signal in some studies. ([bioinfo.uth.edu](https://bioinfo.uth.edu/ancogenedb/views/documents.php?utm_source=openai)) [S3]

## Contradicting or limiting evidence

The main limitation is that I did **not** find strong evidence that COX8A is an established addiction gene in humans. In a large integrative study of cocaine dependence/use, the genes highlighted by GWAS and follow-up transcriptomic analyses were **C1QL2, KCTD20, and STK38**; COX8A was not identified as a lead susceptibility gene. That does not exclude a small effect, but it argues against COX8A being a well-supported primary addiction locus. ([pubmed.ncbi.nlm.nih.gov](https://pubmed.ncbi.nlm.nih.gov/30734435/)) [S4]

Likewise, the curated addiction-expression summaries I reviewed do not elevate COX8A as a major human signal. ANCO-GeneDB lists human alcohol- and cocaine-related brain transcriptome datasets, including nucleus accumbens and hippocampus studies, but the displayed significant-gene lists for the alcohol datasets do not feature COX8A. For example, the GSE62699 alcohol-dependence nucleus accumbens list contains many significant genes, but COX8A is absent from the shown p-adjusted <0.05, |log2FC|>1 results; in the GSE44456 alcoholic hippocampus dataset, the displayed stringent-hit list contains only FKBP5. ([bioinfo.uth.edu](https://bioinfo.uth.edu/ancogenedb/views/analyses.php)) [S5]

Also importantly, authoritative gene-level clinical annotation does not link COX8A to substance use disorders. The NCBI COX8A record lists mitochondrial complex IV deficiency as the associated condition, not alcohol, nicotine, cocaine, opioid, or broader addiction phenotypes. That is not definitive disproof, but it does show that addiction is not an established disease association for this gene in standard reference resources. ([ncbi.nlm.nih.gov](https://www.ncbi.nlm.nih.gov/gene?Cmd=DetailsSearch&Db=gene&Term=1351)) [S6]

A further limitation is that the positive evidence is largely **indirect**. The rat alcohol study shows differential expression, but differential expression after drug exposure does not by itself mean a gene drives addiction vulnerability, compulsion, relapse, or reward learning. It may instead reflect downstream metabolic adaptation to repeated alcohol exposure. ([pmc.ncbi.nlm.nih.gov](https://pmc.ncbi.nlm.nih.gov/articles/PMC5851801/)) [S7]

## Mechanistic / biological rationale

There is a reasonable mechanistic story for why COX8A might matter indirectly. COX8A is part of complex IV, and complex IV supports oxidative phosphorylation, ATP generation, and mitochondrial redox balance. Neurons in reward circuitry have high energetic demand, and addiction-related exposures can perturb mitochondrial function, oxidative stress, and energy metabolism. Reviews and experimental work on alcohol and cocaine note mitochondrial dysfunction as a recurring feature of drug exposure and neurotoxicity. ([ncbi.nlm.nih.gov](https://www.ncbi.nlm.nih.gov/gene?Cmd=DetailsSearch&Db=gene&Term=1351)) [S8]

That makes the alcohol-rat finding biologically plausible: if binge alcohol alters oxidative phosphorylation genes in prefrontal cortex, Cox8a may be part of a compensatory or stress-response program. The same study reported coordinated increases across multiple mitochondrial respiratory genes, not an isolated COX8A-specific effect, which fits a pathway-level response more than a gene-specific addiction mechanism. ([pmc.ncbi.nlm.nih.gov](https://pmc.ncbi.nlm.nih.gov/articles/PMC5851801/)) [S9]

However, this rationale cuts both ways. Because COX8A is a core mitochondrial housekeeping component, altered expression may simply index disturbed neuronal homeostasis rather than addiction per se. In other words, COX8A has a stronger case for being relevant to **neural energy homeostasis** than for being specifically implicated in the behavioral syndrome of addiction. The distinction matters: a gene can be altered in addicted brains without being a validated addiction-risk gene or therapeutic target. ([ncbi.nlm.nih.gov](https://www.ncbi.nlm.nih.gov/gene?Cmd=DetailsSearch&Db=gene&Term=1351)) [S10]

## Bottom line

COX8A is **not strongly established as an addiction gene/protein**, but there is **some indirect evidence** connecting it to addiction-relevant biology. The best support I found is a rodent binge-alcohol transcriptomic study showing increased **Cox8a** expression in medial prefrontal cortex and enrichment of oxidative-phosphorylation/mitochondrial pathways. Against that, I found **no convincing human genetic evidence**, no clear functional studies showing that manipulating COX8A changes addictive behaviors, and no authoritative clinical annotation linking COX8A to substance use disorders. ([pmc.ncbi.nlm.nih.gov](https://pmc.ncbi.nlm.nih.gov/articles/PMC5851801/)) [S11]

So the fairest interpretation is: **COX8A may participate in mitochondrial adaptations seen during substance exposure, especially alcohol-related models, but current evidence is too limited to call it a well-supported addiction gene.** ([pmc.ncbi.nlm.nih.gov](https://pmc.ncbi.nlm.nih.gov/articles/PMC5851801/)) [S12]

## Cited sources

- **[S1]** COX8A cytochrome c oxidase subunit 8A [Homo sapiens (human)] - Gene - NCBI — https://www.ncbi.nlm.nih.gov/gene?Cmd=DetailsSearch&Db=gene&Term=1351
- **[S2]** 
            Gene expression changes in the ventral hippocampus and medial prefrontal cortex of adolescent alcohol-preferring (P) rats following binge-like-alcohol drinking - PMC
         — https://pmc.ncbi.nlm.nih.gov/articles/PMC5851801/
- **[S3]** ANCO-GeneDB — https://bioinfo.uth.edu/ancogenedb/views/documents.php?utm_source=openai
- **[S4]** Cocaine'omics: Genome-wide and transcriptome-wide analyses provide biological insight into cocaine use and dependence - PubMed — https://pubmed.ncbi.nlm.nih.gov/30734435/
- **[S5]** Analyses Results — https://bioinfo.uth.edu/ancogenedb/views/analyses.php
- **[S6]** COX8A cytochrome c oxidase subunit 8A [Homo sapiens (human)] - Gene - NCBI — https://www.ncbi.nlm.nih.gov/gene?Cmd=DetailsSearch&Db=gene&Term=1351
- **[S7]** 
            Gene expression changes in the ventral hippocampus and medial prefrontal cortex of adolescent alcohol-preferring (P) rats following binge-like-alcohol drinking - PMC
         — https://pmc.ncbi.nlm.nih.gov/articles/PMC5851801/
- **[S8]** COX8A cytochrome c oxidase subunit 8A [Homo sapiens (human)] - Gene - NCBI — https://www.ncbi.nlm.nih.gov/gene?Cmd=DetailsSearch&Db=gene&Term=1351
- **[S9]** 
            Gene expression changes in the ventral hippocampus and medial prefrontal cortex of adolescent alcohol-preferring (P) rats following binge-like-alcohol drinking - PMC
         — https://pmc.ncbi.nlm.nih.gov/articles/PMC5851801/
- **[S10]** COX8A cytochrome c oxidase subunit 8A [Homo sapiens (human)] - Gene - NCBI — https://www.ncbi.nlm.nih.gov/gene?Cmd=DetailsSearch&Db=gene&Term=1351
- **[S11]** 
            Gene expression changes in the ventral hippocampus and medial prefrontal cortex of adolescent alcohol-preferring (P) rats following binge-like-alcohol drinking - PMC
         — https://pmc.ncbi.nlm.nih.gov/articles/PMC5851801/
- **[S12]** 
            Gene expression changes in the ventral hippocampus and medial prefrontal cortex of adolescent alcohol-preferring (P) rats following binge-like-alcohol drinking - PMC
         — https://pmc.ncbi.nlm.nih.gov/articles/PMC5851801/
