## Supplementary material for "Understanding Substance Dependence: What Differentiates Addictive from Non-Addictive Drugs?": Folder for supplementary files: addictive-genes-claims-first100-029-hpcal1-gene-protein-is-implicated-in-addiction-6be478532013.md

# addictive-genes-claims-first100-029-6be478532013

**Original claim:** HPCAL1 gene/protein is implicated in addiction.

## Claim

HPCAL1 gene/protein is implicated in addiction.

## Verdict

Verdict: Mixed Evidence

HPCAL1 has **some suggestive links** to addiction-related biology, especially in **nicotine, alcohol, and opioid-related transcriptomic/genetic screens**, but the evidence is **indirect, sparse, and not yet robustly replicated**. I did **not** find strong human evidence placing HPCAL1 among the established, repeatedly replicated addiction-risk genes in the way that loci such as **CHRNA5-CHRNA3-CHRNB4, ADH1B, OPRM1, FURIN, DRD2, TENM2,** or **MAGI2/GNAI1** are discussed in modern substance-use-disorder genetics reviews. ([pmc.ncbi.nlm.nih.gov](https://pmc.ncbi.nlm.nih.gov/articles/PMC2581605/)) [S1]

## Confidence

Medium

Confidence is **medium** because there is enough literature to say the claim is **not baseless**, but the supporting evidence is mostly **candidate-gene/omics/network-level** rather than **direct causal or replicated human association evidence**. The current record supports “possibly implicated” better than “clearly established.” ([pmc.ncbi.nlm.nih.gov](https://pmc.ncbi.nlm.nih.gov/articles/PMC3193935/?utm_source=openai)) [S2]

## Supporting evidence

HPCAL1 encodes **hippocalcin-like 1 (also called VILIP-3)**, a **neuron-specific calcium-binding protein** expressed in the **brain and retina**. NCBI describes it as a protein likely relevant to **neuronal signaling in the central nervous system**, and the original cloning paper showed a brain-restricted transcript and a structure consistent with the **neuronal calcium sensor** family. That makes HPCAL1 biologically plausible as a modulator of neural circuits that are often altered in addiction. ([ncbi.nlm.nih.gov](https://www.ncbi.nlm.nih.gov/gene/3241?utm_source=openai)) [S3]

The most direct human genetic hint I found was an older pooled-DNA genome-wide study of **nicotine dependence and smoking cessation**. In that study, the authors looked for **clusters of nominally positive SNPs** rather than standard genome-wide significant loci, and **HPCAL1** appeared among the genes containing clustered nominal signals, with **5 clustered SNPs**. This is real signal-generation evidence, but it is exploratory and methodologically weaker than current individual-level GWAS standards. ([pmc.ncbi.nlm.nih.gov](https://pmc.ncbi.nlm.nih.gov/articles/PMC2581605/)) [S4]

There is also preclinical transcriptomic evidence. In a rat study comparing nucleus accumbens shell-to-ventral pallidum GABA neurons from **Lewis rats** versus **Fisher rats**—strains that differ in drug-taking vulnerability—the authors found **Hpcal1 upregulated** in Lewis rats. They interpreted Hpcal1, together with several other signaling/synaptic genes, as potentially contributing to altered accumbal signaling in a phenotype predisposed to nicotine self-administration. That supports relevance to **addiction-vulnerable circuitry**, though it does not establish Hpcal1 as a driver. ([pmc.ncbi.nlm.nih.gov](https://pmc.ncbi.nlm.nih.gov/articles/PMC3193935/?utm_source=openai)) [S5]

More recently, a human postmortem study of **opioid use disorder (OUD)** found that rhythmic transcripts in the **nucleus accumbens** of subjects with OUD formed OUD-specific coexpression modules enriched for synaptic functions. **HPCAL1** was listed among synaptic regulators in one of these OUD-specific modules. This is meaningful because it places HPCAL1 inside a molecular network altered in a core reward region in human OUD, but the result is **network membership**, not proof that HPCAL1 itself is a causal addiction gene. ([wickedneuro.com](https://wickedneuro.com/wp-content/uploads/2024/04/xue2022.pdf)) [S6]

An additional cross-species alcohol study integrated human AUD brain transcriptomics with primate and mouse alcohol-use models. It identified conserved coexpression networks associated with **human AUD** and **animal alcohol-use phenotypes**, and **HPCAL1** emerged as a **hub gene** in a nucleus accumbens network. The paper further linked HPCAL1, via GWAS catalog annotation, to **GGT levels from alcohol consumption**. This again supports an alcohol-related signal, but it is still indirect: the human GWAS trait was a **biomarker related to alcohol use**, not a definitive diagnosis of alcohol use disorder. ([pmc.ncbi.nlm.nih.gov](https://pmc.ncbi.nlm.nih.gov/articles/PMC11245012/)) [S7]

## Contradicting or limiting evidence

The clearest limitation is that even the rat accumbens transcriptomics paper that reported elevated **Hpcal1** explicitly noted that Hpcal1 had **“none”** of the literature-mined relationships with **addiction/nicotine** at that time, and only relatively few links to signaling and synaptic plasticity. In other words, the authors themselves treated Hpcal1 as a **novel/weakly connected candidate**, not an already established addiction gene. ([pmc.ncbi.nlm.nih.gov](https://pmc.ncbi.nlm.nih.gov/articles/PMC3193935/?utm_source=openai)) [S8]

Modern reviews of substance use disorder genetics emphasize other loci as the replicated core findings. For tobacco use disorder these include **CHRNA5-CHRNA3-CHRNB4, DNMT3B, MAGI2/GNAI1,** and **TENM2**; for opioid use disorder they emphasize **OPRM1, FURIN,** and **TSNARE1**; and for cross-disorder addiction risk they highlight loci near **DRD2**. **HPCAL1 is not presented in that review as an established replicated SUD gene**, which strongly suggests it is **not currently part of the mainstream replicated genetic architecture** of addiction. This last point is an inference from what the review does and does not highlight. ([nature.com](https://www.nature.com/articles/s41380-024-02547-z)) [S9]

The OUD circadian-transcriptome paper is also limiting in an important way: although it identified OUD-related rhythmic transcript changes and included HPCAL1 in an OUD-specific module, the authors reported **no significant enrichment** of those rhythmic transcripts for **opioid dependence GWAS** loci. That weakens any claim that HPCAL1 has direct human genetic support for OUD. ([wickedneuro.com](https://wickedneuro.com/wp-content/uploads/2024/04/xue2022.pdf)) [S10]

Likewise, the alcohol-related evidence is mostly **network-based** or tied to **alcohol-consumption biomarkers** rather than direct case-control AUD genetics or interventional functional studies. I did not find strong evidence that HPCAL1 variants have been reproducibly associated at genome-wide significance with **AUD, OUD, nicotine dependence, cocaine use disorder,** or other substance use disorders in contemporary large studies. This is a literature-search inference based on the sources reviewed here. ([pmc.ncbi.nlm.nih.gov](https://pmc.ncbi.nlm.nih.gov/articles/PMC11245012/)) [S11]

## Mechanistic / biological rationale

Mechanistically, the claim is **plausible**. HPCAL1 is a **neuronal calcium sensor** with EF-hand calcium-binding motifs, and neuronal calcium sensor proteins help translate intracellular **Ca²⁺ signals** into changes in membrane trafficking, signaling, and neuronal plasticity. Since addiction involves long-lasting plasticity in reward and stress circuits, a protein like HPCAL1 could reasonably influence those processes. ([pubmed.ncbi.nlm.nih.gov](https://pubmed.ncbi.nlm.nih.gov/8038222/?utm_source=openai)) [S12]

That plausibility fits the human transcriptomic studies. In OUD, HPCAL1 sits in a nucleus accumbens module enriched for **dopamine, GABAergic, and glutamatergic synaptic functions**; in the alcohol cross-species study, HPCAL1 is a hub in a conserved accumbens network enriched for **synaptic plasticity** and other neural processes relevant to addictive behavior. So there is a coherent biological story in which HPCAL1 could modulate addiction-related neuroadaptation. But this is still **mechanistic plausibility plus omics correlation**, not direct proof of a causal role. ([wickedneuro.com](https://wickedneuro.com/wp-content/uploads/2024/04/xue2022.pdf)) [S13]

## Bottom line

**HPCAL1 is not an established addiction gene, but it is a plausible and weakly supported candidate.** The best current evidence comes from **exploratory nicotine genetics**, **rodent nucleus accumbens expression studies**, and **human alcohol/OUD transcriptomic network analyses**. Those findings are enough to say HPCAL1 is **possibly implicated in addiction-related biology**, especially in reward-circuit molecular networks, but **not enough to claim a strong, replicated causal implication in addiction itself**. ([pmc.ncbi.nlm.nih.gov](https://pmc.ncbi.nlm.nih.gov/articles/PMC2581605/)) [S14]

## Cited sources

- **[S1]** 
            Genome wide association for nicotine dependence and smoking cessation success in NIH research volunteers - PMC
         — https://pmc.ncbi.nlm.nih.gov/articles/PMC2581605/
- **[S2]** Gene Expression in Accumbens GABA Neurons from Inbred Rats with Different Drug-Taking Behavior - PMC — https://pmc.ncbi.nlm.nih.gov/articles/PMC3193935/?utm_source=openai
- **[S3]** HPCAL1 hippocalcin like 1 [Homo sapiens (human)] - Gene - NCBI — https://www.ncbi.nlm.nih.gov/gene/3241?utm_source=openai
- **[S4]** 
            Genome wide association for nicotine dependence and smoking cessation success in NIH research volunteers - PMC
         — https://pmc.ncbi.nlm.nih.gov/articles/PMC2581605/
- **[S5]** Gene Expression in Accumbens GABA Neurons from Inbred Rats with Different Drug-Taking Behavior - PMC — https://pmc.ncbi.nlm.nih.gov/articles/PMC3193935/?utm_source=openai
- **[S6]** Molecular rhythm alterations in prefrontal cortex and nucleus accumbens associated with opioid use disorder — https://wickedneuro.com/wp-content/uploads/2024/04/xue2022.pdf
- **[S7]** 
            Bulk and Single-cell Transcriptomic Brain Data Identify Overlapping Processes and Cell-types with Human AUD and Mammalian Models of Alcohol Use - PMC
         — https://pmc.ncbi.nlm.nih.gov/articles/PMC11245012/
- **[S8]** Gene Expression in Accumbens GABA Neurons from Inbred Rats with Different Drug-Taking Behavior - PMC — https://pmc.ncbi.nlm.nih.gov/articles/PMC3193935/?utm_source=openai
- **[S9]** The genetic landscape of substance use disorders | Molecular Psychiatry — https://www.nature.com/articles/s41380-024-02547-z
- **[S10]** Molecular rhythm alterations in prefrontal cortex and nucleus accumbens associated with opioid use disorder — https://wickedneuro.com/wp-content/uploads/2024/04/xue2022.pdf
- **[S11]** 
            Bulk and Single-cell Transcriptomic Brain Data Identify Overlapping Processes and Cell-types with Human AUD and Mammalian Models of Alcohol Use - PMC
         — https://pmc.ncbi.nlm.nih.gov/articles/PMC11245012/
- **[S12]** Molecular cloning of a novel calcium-binding protein structurally related to hippocalcin from human brain and chromosomal mapping of its gene - PubMed — https://pubmed.ncbi.nlm.nih.gov/8038222/?utm_source=openai
- **[S13]** Molecular rhythm alterations in prefrontal cortex and nucleus accumbens associated with opioid use disorder — https://wickedneuro.com/wp-content/uploads/2024/04/xue2022.pdf
- **[S14]** 
            Genome wide association for nicotine dependence and smoking cessation success in NIH research volunteers - PMC
         — https://pmc.ncbi.nlm.nih.gov/articles/PMC2581605/
