## Supplementary material for "Understanding Substance Dependence: What Differentiates Addictive from Non-Addictive Drugs?": Folder for supplementary files: addictive-genes-claims-first100-030-slc39a14-gene-protein-is-implicated-in-addiction-b0392fa69d0d.md

# addictive-genes-claims-first100-030-b0392fa69d0d

**Original claim:** SLC39A14 gene/protein is implicated in addiction.

## Claim

SLC39A14 gene/protein is implicated in addiction.

## Verdict

Verdict: Likely False

The claim is **overstated based on current evidence**. SLC39A14 is a well-established **manganese/zinc/iron transporter** whose strongest human and animal evidence links it to **manganese homeostasis, hypermanganesemia, basal ganglia injury, dystonia/parkinsonism, and motor dysfunction**—not to addiction as a primary phenotype. I found only **indirect** support for an addiction link: one small, exploratory postmortem substance-use-disorder transcriptomic study placed SLC39A14 in enriched pathways in an alcohol-use subgroup, and SLC39A14-related manganese dysregulation can alter dopaminergic signaling, which is biologically relevant to addiction. But I did **not** find replicated human genetic associations, clinical addiction phenotypes, or direct animal addiction-behavior studies specifically implicating **SLC39A14** itself. ([ncbi.nlm.nih.gov](https://www.ncbi.nlm.nih.gov/gene/23516)) [S1]

## Confidence

Medium. The core biology of SLC39A14 is well established, and the absence of direct addiction evidence in the main authoritative sources is fairly clear. Confidence is not high because “implicated” is a broad term and there is some indirect mechanistic plausibility plus limited exploratory transcriptomic signal. ([ncbi.nlm.nih.gov](https://www.ncbi.nlm.nih.gov/gene/23516)) [S2]

## Supporting evidence

- A 2025 integrative postmortem brain transcriptomic study of substance use disorders reported that, within the **alcohol use disorder** subgroup, **SLC39A14** appeared in enriched Gene Ontology pathways such as **intracellular zinc ion homeostasis** and **positive regulation of G protein-coupled receptor signaling**. However, the authors explicitly stated that these subtype-specific findings were **exploratory** and limited by **small sample size/power**. This is the strongest addiction-adjacent evidence I found for SLC39A14, but it is not a replicated gene-level association with addiction risk or behavior. ([genomicpress.kglmeridian.com](https://genomicpress.kglmeridian.com/view/journals/genpsych/1/3/article-p44.xml)) [S3]

- SLC39A14 loss of function clearly affects brain systems relevant to reward circuitry. In mice, Slc39a14 deficiency causes **marked manganese accumulation in brain**, especially basal ganglia-related regions, along with locomotor impairment. In a more detailed neurochemical study, Slc39a14-knockout mice showed **85–90% inhibition of potassium-stimulated striatal dopamine release** despite preserved dopamine neuron number and striatal dopamine content. Because mesostriatal dopaminergic signaling is central to reinforcement and addiction biology, this creates a plausible indirect link. ([pubmed.ncbi.nlm.nih.gov](https://pubmed.ncbi.nlm.nih.gov/29437953/?utm_source=openai)) [S4]

- More broadly, manganese itself can interact with addiction-relevant circuitry. A PubMed-indexed review summarized prior work suggesting manganese influences the **brain reward system** and reported reduced morphine conditioned place preference/withdrawal intensity in rat experiments. Separate toxicology work showed chronic manganese exposure altered **amphetamine-evoked** brain responses in dopaminergic circuits. These findings support plausibility for a manganese–reward connection, though they do **not** specifically establish SLC39A14 as an addiction gene. ([pubmed.ncbi.nlm.nih.gov](https://pubmed.ncbi.nlm.nih.gov/24505901/)) [S5]

## Contradicting or limiting evidence

- Authoritative gene summaries do **not** describe addiction as an established SLC39A14 phenotype. NCBI summarizes SLC39A14 as a divalent metal transporter important for **manganese homeostasis**, with naturally occurring mutations associated with **neurodegeneration with brain iron accumulation** and **early-onset parkinsonism-dystonia with hypermanganesemia**. GeneReviews likewise describes SLC39A14 deficiency as a disorder of **developmental delay, dystonia, spasticity, dysarthria, and parkinsonian signs**, and states that no other phenotypes beyond those discussed there are known for germline pathogenic variants. ([ncbi.nlm.nih.gov](https://www.ncbi.nlm.nih.gov/gene/23516)) [S6]

- The strongest human genetic literature on metal transporters and addiction points to a **different gene, SLC39A8**, not SLC39A14. A JCI review on manganese homeostasis specifically notes that the common **SLC39A8** Ala391Thr variant is associated with **alcohol use disorder** and alcohol-related traits, whereas **SLC39A14** is discussed in the context of **HMNDYT2/dystonia-parkinsonism** from manganese excess. That distinction matters because the two genes are related transporters and can easily be conflated. ([jci.org](https://www.jci.org/articles/view/133120)) [S7]

- In the primary SLC39A14 animal literature, reported phenotypes are overwhelmingly **motor** and **movement-disorder-like**, not addiction-like. Slc39a14 knockout models show manganese dyshomeostasis, locomotor deficits, dystonia-like behavior, and impaired dopamine release, but the cited studies do not test core addiction phenotypes such as drug self-administration, escalation, reinstatement, compulsive intake, or withdrawal-related reinforcement. ([pubmed.ncbi.nlm.nih.gov](https://pubmed.ncbi.nlm.nih.gov/29437953/?utm_source=openai)) [S8]

- The one SUD transcriptomic signal is weak evidence. In that 2025 study, SLC39A14 appeared in **pathway enrichment**, not as a replicated causal locus; subgroup analyses were explicitly called **exploratory**. That is hypothesis-generating, not proof of implication in addiction. ([genomicpress.kglmeridian.com](https://genomicpress.kglmeridian.com/view/journals/genpsych/1/3/article-p44.xml)) [S9]

## Mechanistic / biological rationale

SLC39A14 encodes **ZIP14**, a transporter for **manganese, zinc, iron, and cadmium**. It is highly expressed in peripheral tissues such as **jejunum and liver**, where it helps regulate systemic manganese trafficking and excretion. Loss of ZIP14 function impairs manganese handling, causing **elevated blood and brain manganese**, especially in basal ganglia-linked regions, and produces a syndrome resembling **manganism/dystonia-parkinsonism**. ([ncbi.nlm.nih.gov](https://www.ncbi.nlm.nih.gov/gene/23516)) [S10]

That biology gives a **credible indirect route** to addiction relevance: manganese homeostasis can shape dopaminergic neurotransmission, and Slc39a14-deficient mice show a profound presynaptic defect in **stimulated striatal dopamine release**. Since dopamine signaling in striatal circuits is central to reward learning, incentive salience, and drug reinforcement, one can reasonably hypothesize that severe SLC39A14 dysfunction might influence addiction-related processes. ([pubmed.ncbi.nlm.nih.gov](https://pubmed.ncbi.nlm.nih.gov/34358615/)) [S11]

But the existing data point more strongly toward **neurotoxicity/movement disorder** than toward addiction. SLC39A14-associated disease is not characterized clinically as a substance use disorder, and current experiments do not demonstrate that manipulating SLC39A14 changes addictive behavior in the way expected for a bona fide addiction gene. So the mechanistic rationale is **plausible but indirect**. ([ncbi.nlm.nih.gov](https://www.ncbi.nlm.nih.gov/books/NBK431123/)) [S12]

## Bottom line

**SLC39A14 is not currently well supported as an addiction gene/protein.** The best evidence ties it to **manganese homeostasis** and **dystonia/parkinsonism**, with only **indirect** addiction relevance through dopaminergic/manganese biology and one **exploratory** AUD-adjacent transcriptomic signal. If the intended claim is that SLC39A14 is **firmly implicated in addiction**, the evidence does **not** support that. If the intended claim is that it has a **possible indirect connection** to addiction biology via metal homeostasis and dopamine signaling, that is more defensible—but still unproven. ([ncbi.nlm.nih.gov](https://www.ncbi.nlm.nih.gov/gene/23516)) [S13]

## Cited sources

- **[S1]** SLC39A14 solute carrier family 39 member 14 [Homo sapiens (human)] - Gene - NCBI — https://www.ncbi.nlm.nih.gov/gene/23516
- **[S2]** SLC39A14 solute carrier family 39 member 14 [Homo sapiens (human)] - Gene - NCBI — https://www.ncbi.nlm.nih.gov/gene/23516
- **[S3]** Deciphering the molecular basis of accelerated biological aging in substance use disorder: Integrative transcriptomic analysis in: Genomic Psychiatry Volume 1: Issue 3 | Genomic Press — https://genomicpress.kglmeridian.com/view/journals/genpsych/1/3/article-p44.xml
- **[S4]** SLC39A14 deficiency alters manganese homeostasis and excretion resulting in brain manganese accumulation and motor deficits in mice - PubMed — https://pubmed.ncbi.nlm.nih.gov/29437953/?utm_source=openai
- **[S5]** Manganese interactions with dopaminergic system - PubMed — https://pubmed.ncbi.nlm.nih.gov/24505901/
- **[S6]** SLC39A14 solute carrier family 39 member 14 [Homo sapiens (human)] - Gene - NCBI — https://www.ncbi.nlm.nih.gov/gene/23516
- **[S7]** 
JCI -
Manganese homeostasis: from rare single-gene disorders to complex phenotypes and diseases
 — https://www.jci.org/articles/view/133120
- **[S8]** SLC39A14 deficiency alters manganese homeostasis and excretion resulting in brain manganese accumulation and motor deficits in mice - PubMed — https://pubmed.ncbi.nlm.nih.gov/29437953/?utm_source=openai
- **[S9]** Deciphering the molecular basis of accelerated biological aging in substance use disorder: Integrative transcriptomic analysis in: Genomic Psychiatry Volume 1: Issue 3 | Genomic Press — https://genomicpress.kglmeridian.com/view/journals/genpsych/1/3/article-p44.xml
- **[S10]** SLC39A14 solute carrier family 39 member 14 [Homo sapiens (human)] - Gene - NCBI — https://www.ncbi.nlm.nih.gov/gene/23516
- **[S11]** Behavioral and neurochemical studies of inherited manganese-induced dystonia-parkinsonism in Slc39a14-knockout mice - PubMed — https://pubmed.ncbi.nlm.nih.gov/34358615/
- **[S12]** SLC39A14 Deficiency - GeneReviews® - NCBI Bookshelf — https://www.ncbi.nlm.nih.gov/books/NBK431123/
- **[S13]** SLC39A14 solute carrier family 39 member 14 [Homo sapiens (human)] - Gene - NCBI — https://www.ncbi.nlm.nih.gov/gene/23516
