## Supplementary material for "Understanding Substance Dependence: What Differentiates Addictive from Non-Addictive Drugs?": Folder for supplementary files: addictive-genes-claims-first100-031-fabp5-gene-protein-is-implicated-in-addiction-336eb7cad01a.md

# addictive-genes-claims-first100-031-336eb7cad01a

**Original claim:** FABP5 gene/protein is implicated in addiction.

## Claim

FABP5 gene/protein is implicated in addiction.

## Verdict

Verdict: Likely True

## Confidence

Medium

## Supporting evidence

There is **direct preclinical evidence** that manipulating FABP5 alters addiction-relevant behaviors. In a rat cocaine model, a transcriptomic screen identified **FABP5 as highly enriched in the nucleus accumbens shell (NAcSh)**, and **AAV-mediated knockdown of FABP5 in the NAcSh decreased cocaine self-administration** while altering medium spiny neuron excitability. The authors explicitly framed FABP5/retinoic-acid signaling as a candidate pathway for cocaine use disorder. ([pubmed.ncbi.nlm.nih.gov](https://pubmed.ncbi.nlm.nih.gov/33181146/?utm_source=openai)) [S1]

There is also **substance-specific evidence in nicotine reward**. In mice, **global FABP5 deletion increased nicotine-conditioned place preference (CPP)** at **0.1 mg/kg nicotine**, indicating enhanced nicotine reward sensitivity in the knockout animals. That is a standard addiction-relevant behavioral assay, even though it is not the same as compulsive drug self-administration. ([pubmed.ncbi.nlm.nih.gov](https://pubmed.ncbi.nlm.nih.gov/36864947/)) [S2]

Beyond those direct behavioral studies, FABP5 has been linked to **reward-circuit neurochemistry**. FABP5 knockout mice show **altered dopamine signaling**, including **higher D1 receptor levels in several striatal subregions**, and FABP5 deletion also alters **brain endocannabinoid tone**, including elevated anandamide (AEA) in the striatum, medial prefrontal cortex, midbrain, and thalamus. These systems are strongly implicated in reinforcement and drug reward, so these findings support biological relevance to addiction. ([pubmed.ncbi.nlm.nih.gov](https://pubmed.ncbi.nlm.nih.gov/37664218/)) [S3]

Mechanistically, FABP5 is not just correlated with these phenotypes; it has a plausible causal role because it is a **carrier/chaperone for endocannabinoid-related lipids**. Primary studies show FABP5 is important for **AEA handling**, **2-AG retrograde signaling**, and even intracellular handling of **THC/CBD**. Since endocannabinoid signaling modulates reward, motivation, and drug responses, this strengthens the inference that FABP5 can influence addiction-related processes. ([pubmed.ncbi.nlm.nih.gov](https://pubmed.ncbi.nlm.nih.gov/29531087/?utm_source=openai)) [S4]

## Contradicting or limiting evidence

The evidence is **not yet strong enough to call FABP5 a well-established human addiction gene**. The most convincing data I found are **rodent and mechanistic**, not replicated human genetic or clinical studies. The cocaine and nicotine findings are important, but they do not by themselves prove that FABP5 is a major determinant of human substance use disorder. ([pubmed.ncbi.nlm.nih.gov](https://pubmed.ncbi.nlm.nih.gov/33181146/?utm_source=openai)) [S5]

Some findings are also **mixed across paradigms and manipulations**. A pharmacology study of **SBFI26**, a FABP5/7 inhibitor, found that it **did not produce CPP or conditioned place aversion** and concluded that, under the tested conditions, it **did not appear to pose a dependence risk**. That argues against a simplistic claim that reducing FABP5 signaling is intrinsically addictive. ([pubmed.ncbi.nlm.nih.gov](https://pubmed.ncbi.nlm.nih.gov/27092087/)) [S6]

Likewise, a study of **FABP5/7 double-deficient mice** found **increased sucrose consumption** and reduced forced-swim immobility, but **acute pharmacologic inhibition** did **not** reproduce the sucrose effect. That raises the possibility of **developmental compensation** or other indirect effects in constitutive knockout models. Also, that experiment was not specific to FABP5 alone and used a natural reward rather than an addictive drug. ([pubmed.ncbi.nlm.nih.gov](https://pubmed.ncbi.nlm.nih.gov/29570114/)) [S7]

The nicotine study is also narrower than the claim sounds: the effect was seen in **CPP**, not self-administration, and apparently at **one nicotine dose (0.1 mg/kg) but not 0.5 mg/kg**. The cocaine study was stronger behaviorally, but it was **region-specific**, done in **male rats**, and the authors note that they **did not show an effect on reinstatement** in the paradigm they used. ([pubmed.ncbi.nlm.nih.gov](https://pubmed.ncbi.nlm.nih.gov/36864947/)) [S8]

## Mechanistic / biological rationale

FABP5 is biologically plausible as an addiction-related modulator because it sits at the intersection of **endocannabinoid**, **dopamine**, and **reward-circuit** signaling. FABP5 promotes intracellular trafficking of **AEA** and can influence its hydrolysis and downstream signaling; FABP5 deletion elevates brain AEA and disrupts PPARβ/δ-linked signaling. ([pubmed.ncbi.nlm.nih.gov](https://pubmed.ncbi.nlm.nih.gov/24644281/?utm_source=openai)) [S9]

In reward-relevant brain regions, FABP5 regulates synaptic endocannabinoid signaling. In the **dorsal raphe**, pharmacologic inhibition or genetic deletion of FABP5 **abolished phasic and tonic retrograde endocannabinoid control of excitatory transmission**. In the **striatum**, FABP5 deletion elevated AEA and impaired tonic 2-AG/AEA signaling at GABA synapses. More recently, **astrocytic FABP5** was shown to be necessary for normal **2-AG-mediated depolarization-induced suppression of inhibition** in the hippocampus. Together, these data make FABP5 a genuine regulator of synaptic eCB signaling rather than a distant bystander. ([pubmed.ncbi.nlm.nih.gov](https://pubmed.ncbi.nlm.nih.gov/29531087/)) [S10]

That matters because endocannabinoid signaling is deeply intertwined with addiction biology. In the FABP5 literature itself, FABP5 deletion was associated with **greater striatal D1 receptor levels**, suggesting downstream effects on dopaminergic signaling. Since dopamine signaling in the striatum and nucleus accumbens is central to reinforcement learning and drug seeking, this provides a reasonable path by which FABP5 could alter vulnerability to drug reward. ([pubmed.ncbi.nlm.nih.gov](https://pubmed.ncbi.nlm.nih.gov/37664218/)) [S11]

FABP5 also interacts with **cannabinoid drugs directly**. Human FABPs, including FABP5, bind **THC and CBD**, and these cannabinoids can compete with AEA for FABP-mediated transport. In mice, FABP5 also altered the **behavioral effects and pharmacokinetics of acute THC inhalation**. This does not prove cannabis addiction involvement, but it shows FABP5 can influence the brain handling and effects of an addictive psychoactive drug. ([pubmed.ncbi.nlm.nih.gov](https://pubmed.ncbi.nlm.nih.gov/25666611/)) [S12]

A key nuance is that the direction of effect is **not uniform across substances or circuits**. **Global FABP5 loss enhanced nicotine CPP**, whereas **NAcSh-specific FABP5 knockdown reduced cocaine self-administration**. That pattern suggests FABP5 is best viewed as a **modulator of addiction-relevant neurobiology**, not a simple “pro-addiction” or “anti-addiction” switch. ([pubmed.ncbi.nlm.nih.gov](https://pubmed.ncbi.nlm.nih.gov/36864947/)) [S13]

## Bottom line

The claim is **likely true**, but mainly on the basis of **preclinical** evidence. FABP5 has been **causally linked** to addiction-relevant phenotypes in rodents—most notably **reduced cocaine self-administration after NAc shell FABP5 knockdown** and **enhanced nicotine CPP after FABP5 deletion**—and there is strong mechanistic support because FABP5 regulates **endocannabinoid and dopamine-related signaling** in reward circuits. ([pubmed.ncbi.nlm.nih.gov](https://pubmed.ncbi.nlm.nih.gov/33181146/?utm_source=openai)) [S14]

What is **not** established is that FABP5 is a confirmed **human addiction gene/protein target** with robust clinical or GWAS-level evidence. So the fairest summary is: **FABP5 is credibly implicated in addiction biology, especially in animal models and mechanistic studies, but human evidence remains limited.** ([pubmed.ncbi.nlm.nih.gov](https://pubmed.ncbi.nlm.nih.gov/27092087/)) [S15]

## Cited sources

- **[S1]** Topographic transcriptomics of the nucleus accumbens shell: Identification and validation of fatty acid binding protein 5 as target for cocaine addiction - PubMed — https://pubmed.ncbi.nlm.nih.gov/33181146/?utm_source=openai
- **[S2]** Fatty Acid-Binding Protein 5 Gene Deletion Enhances Nicotine-Conditioned Place Preference: Illuminating the Putative Gateway Mechanisms - PubMed — https://pubmed.ncbi.nlm.nih.gov/36864947/
- **[S3]** Fatty acid-binding protein 5 differentially impacts dopamine signaling independent of sex and environment - PubMed — https://pubmed.ncbi.nlm.nih.gov/37664218/
- **[S4]** Fatty-acid-binding protein 5 controls retrograde endocannabinoid signaling at central glutamate synapses - PubMed — https://pubmed.ncbi.nlm.nih.gov/29531087/?utm_source=openai
- **[S5]** Topographic transcriptomics of the nucleus accumbens shell: Identification and validation of fatty acid binding protein 5 as target for cocaine addiction - PubMed — https://pubmed.ncbi.nlm.nih.gov/33181146/?utm_source=openai
- **[S6]** Examination of the Addictive and Behavioral Properties of Fatty Acid-Binding Protein Inhibitor SBFI26 - PubMed — https://pubmed.ncbi.nlm.nih.gov/27092087/
- **[S7]** Fatty acid-binding proteins 5 and 7 gene deletion increases sucrose consumption and diminishes forced swim immobility time - PubMed — https://pubmed.ncbi.nlm.nih.gov/29570114/
- **[S8]** Fatty Acid-Binding Protein 5 Gene Deletion Enhances Nicotine-Conditioned Place Preference: Illuminating the Putative Gateway Mechanisms - PubMed — https://pubmed.ncbi.nlm.nih.gov/36864947/
- **[S9]** Fatty acid-binding protein 5 (FABP5) regulates cognitive function both by decreasing anandamide levels and by activating the nuclear receptor peroxisome proliferator-activated receptor β/δ (PPARβ/δ) in the brain - PubMed — https://pubmed.ncbi.nlm.nih.gov/24644281/?utm_source=openai
- **[S10]** Fatty-acid-binding protein 5 controls retrograde endocannabinoid signaling at central glutamate synapses - PubMed — https://pubmed.ncbi.nlm.nih.gov/29531087/
- **[S11]** Fatty acid-binding protein 5 differentially impacts dopamine signaling independent of sex and environment - PubMed — https://pubmed.ncbi.nlm.nih.gov/37664218/
- **[S12]** Fatty acid-binding proteins (FABPs) are intracellular carriers for Δ9-tetrahydrocannabinol (THC) and cannabidiol (CBD) - PubMed — https://pubmed.ncbi.nlm.nih.gov/25666611/
- **[S13]** Fatty Acid-Binding Protein 5 Gene Deletion Enhances Nicotine-Conditioned Place Preference: Illuminating the Putative Gateway Mechanisms - PubMed — https://pubmed.ncbi.nlm.nih.gov/36864947/
- **[S14]** Topographic transcriptomics of the nucleus accumbens shell: Identification and validation of fatty acid binding protein 5 as target for cocaine addiction - PubMed — https://pubmed.ncbi.nlm.nih.gov/33181146/?utm_source=openai
- **[S15]** Examination of the Addictive and Behavioral Properties of Fatty Acid-Binding Protein Inhibitor SBFI26 - PubMed — https://pubmed.ncbi.nlm.nih.gov/27092087/
