## Supplementary material for "Understanding Substance Dependence: What Differentiates Addictive from Non-Addictive Drugs?": Folder for supplementary files: addictive-genes-claims-first100-032-nampt-gene-protein-is-implicated-in-addiction-2e4db1ab96b3.md

# addictive-genes-claims-first100-032-2e4db1ab96b3

**Original claim:** NAMPT gene/protein is implicated in addiction.

## Claim

**Claim:** *NAMPT gene/protein is implicated in addiction.*

## Verdict

Verdict: Likely True

## Confidence

**Medium.** There is credible **causal preclinical evidence** linking NAMPT to cocaine reward circuitry, but the evidence base is still **narrow**, centered mostly on mouse studies of cocaine-conditioned reward rather than broad human addiction data. ([pubmed.ncbi.nlm.nih.gov](https://pubmed.ncbi.nlm.nih.gov/29753648/?utm_source=openai)) [S1]

## Supporting evidence

The strongest direct evidence comes from a 2018 mouse study showing that **NAMPT is upregulated in the ventral tegmental area (VTA)** after cocaine conditioning, and that **pharmacologic inhibition of NAMPT with FK866 attenuates cocaine reward**. In that same study, the anti-reward effect of FK866 was weakened by **restoring NAMPT signaling** via NAMPT overexpression or **NMN supplementation**, and cocaine-conditioned mice showed increased **VTA NAD and NMN** along with increased **SIRT1**. Importantly, the behavioral effect of NAMPT inhibition was substantially reduced in **Sirt1 conditional knockout** mice, supporting a NAMPT→NAD→SIRT1 mechanism rather than a nonspecific drug effect. ([pubmed.ncbi.nlm.nih.gov](https://pubmed.ncbi.nlm.nih.gov/29753648/?utm_source=openai)) [S2]

A subsequent commentary summarizing that paper adds useful detail: NAMPT protein was reported as increased not only in the **VTA** but also in the **nucleus accumbens (NAc)** of cocaine-conditioned mice, while other regions tested did not show the same change. The commentary also notes that **VTA NAMPT overexpression enhanced cocaine conditioned place preference (CPP)**, whereas **VTA NAMPT inhibition reduced CPP**, and that this behavioral dependence was largely **SIRT1-dependent**. Together, that is reasonably strong evidence that NAMPT is not just passively altered by cocaine exposure, but functionally contributes to at least one addiction-relevant phenotype. ([pmc.ncbi.nlm.nih.gov](https://pmc.ncbi.nlm.nih.gov/articles/PMC6971506/)) [S3]

There is also broader support from adjacent reward-pathway work showing that the **NAD/SIRT1 axis** is relevant to addiction biology. In the nucleus accumbens, **SIRT1 regulates cocaine-associated synaptic plasticity and reward**, and both SIRT1 overexpression and pharmacologic activation have been reported to increase cocaine preference, whereas antagonism decreases it. A separate study found that **NAD+ cellular redox and SIRT1 regulate conditioned cocaine reward** and dopaminergic transcriptional control. These studies do not prove NAMPT’s role by themselves, but they make the NAMPT finding more biologically coherent because NAMPT is the rate-limiting enzyme feeding this pathway. ([pmc.ncbi.nlm.nih.gov](https://pmc.ncbi.nlm.nih.gov/articles/PMC9355311/)) [S4]

More recent work in the NAc further supports this general framework: cocaine exposure altered the diurnal regulation of **Nampt expression, Sirt1 expression, and NAD+ levels** in the nucleus accumbens, and the authors explicitly note that prior VTA work showed **NAMPT-mediated NAD biosynthesis directly promotes cocaine reward in a SIRT1-dependent manner**. This is still not independent proof of NAMPT as a pan-addiction gene, but it reinforces that NAMPT sits in a molecular system already tied to reward regulation in multiple brain regions. ([pmc.ncbi.nlm.nih.gov](https://pmc.ncbi.nlm.nih.gov/articles/PMC9355311/)) [S5]

## Contradicting or limiting evidence

The biggest limitation is **scope**. The direct literature connecting NAMPT itself to addiction is **sparse**. The commentary on the 2018 study explicitly describes that paper as **uniquely linking NAMPT and SIRT1 to cocaine addiction**, which implies the field is still early rather than mature. ([pmc.ncbi.nlm.nih.gov](https://pmc.ncbi.nlm.nih.gov/articles/PMC6971506/)) [S6]

Also, the best direct evidence is in **mouse cocaine CPP**, which is an addiction-relevant reward model but **not the same as the full syndrome of substance use disorder**. The same commentary specifically notes that **self-administration**, reinforcement, and withdrawal-related models still need to be tested for NAMPT, and that whether NAMPT contributes to withdrawal symptoms remained unknown. So the current evidence is stronger for **cocaine reward-related behavior** than for addiction broadly defined. ([pmc.ncbi.nlm.nih.gov](https://pmc.ncbi.nlm.nih.gov/articles/PMC6971506/)) [S7]

Human evidence is notably thin. I did **not** find convincing human clinical, postmortem, or genetic studies showing that NAMPT is an established addiction-risk gene. By contrast, large human GWAS of substance use disorders have highlighted other loci—such as **ADH1B** in alcohol dependence and **PDE4B** plus other substance-specific receptor/metabolic genes in multivariate SUD analyses. That does **not** prove NAMPT is uninvolved, but it does suggest NAMPT is **not yet a major replicated human signal** in addiction genetics. ([pubmed.ncbi.nlm.nih.gov](https://pubmed.ncbi.nlm.nih.gov/30482948/)) [S8]

So the claim is better supported if interpreted narrowly as **“NAMPT is implicated in addiction-relevant neurobiology, especially cocaine reward”** than if interpreted broadly as **“NAMPT is a well-established addiction gene/protein across human substance use disorders.”** ([pubmed.ncbi.nlm.nih.gov](https://pubmed.ncbi.nlm.nih.gov/29753648/?utm_source=openai)) [S9]

## Mechanistic / biological rationale

Mechanistically, the claim is plausible. NAMPT is the **rate-limiting enzyme in the nicotinamide salvage pathway for NAD+ biosynthesis**, and reviews describe NAMPT and SIRT1 as part of a coupled **NAMPT/NAD/SIRT1 regulatory network** that links metabolism, stress responses, and gene regulation. Because SIRT1 is NAD-dependent, changes in NAMPT can alter downstream transcriptional and epigenetic programs relevant to neuronal plasticity. ([pubmed.ncbi.nlm.nih.gov](https://pubmed.ncbi.nlm.nih.gov/26215259/?utm_source=openai)) [S10]

NAMPT also has well-established roles in **neural homeostasis**, which strengthens the biological rationale for an addiction link. In neurons, Nampt is required for **long-term depression (LTD)** and normal function of **GluN2B-containing NMDA receptors**. Separate mouse work shows that **NAMPT depletion disturbs mitochondrial homeostasis and causes neuronal degeneration**, and inducible deletion of Nampt in adult projection neurons causes **mitochondrial dysfunction, synaptic abnormalities, motor deficits, and death**. These findings show NAMPT is not a peripheral bystander protein; it is deeply involved in neuronal energy balance, synaptic function, and survival. ([pubmed.ncbi.nlm.nih.gov](https://pubmed.ncbi.nlm.nih.gov/26481044/)) [S11]

That matters because addiction involves durable changes in **synaptic plasticity, redox state, circadian regulation, and mitochondrial function** within reward circuits such as the VTA and NAc. Reviews on NAD biology in addiction note that impaired NAD/SIRT1 signaling can alter **brain energy homeostasis and behavioral plasticity** during substance exposure, and NAc studies show cocaine disrupts the diurnal regulation of **Nampt, Sirt1, and NAD+**. So while the mechanistic chain is still incomplete, the pathway is biologically credible: **NAMPT could influence addiction-related behavior by modulating NAD availability, SIRT1 activity, dopaminergic regulation, and plasticity in reward circuits.** ([mdpi.com](https://www.mdpi.com/2076-3921/9/5/425/html)) [S12]

## Bottom line

The best available evidence supports **NAMPT as an addiction-relevant molecule in preclinical cocaine models**, especially through a **VTA NAMPT/NAD/SIRT1 pathway** that modulates cocaine reward. That is enough to say the claim is **likely true** in a **narrow, mechanistic/preclinical sense**. ([pubmed.ncbi.nlm.nih.gov](https://pubmed.ncbi.nlm.nih.gov/29753648/?utm_source=openai)) [S13]

However, the evidence is **not yet broad or definitive**: it relies heavily on a limited set of mouse studies, emphasizes **conditioned reward rather than full addiction phenotypes**, and lacks strong **human genetic or clinical confirmation**. So NAMPT is best described as a **promising but not yet firmly established addiction gene/protein**. ([pmc.ncbi.nlm.nih.gov](https://pmc.ncbi.nlm.nih.gov/articles/PMC6971506/)) [S14]

## Cited sources

- **[S1]** Nicotinamide phosphoribosyltransferase regulates cocaine reward through Sirtuin 1 - PubMed — https://pubmed.ncbi.nlm.nih.gov/29753648/?utm_source=openai
- **[S2]** Nicotinamide phosphoribosyltransferase regulates cocaine reward through Sirtuin 1 - PubMed — https://pubmed.ncbi.nlm.nih.gov/29753648/?utm_source=openai
- **[S3]** 
            Nicotinamide phosphoribosyltransferase contributes to cocaine addiction through sirtuin 1 - PMC
         — https://pmc.ncbi.nlm.nih.gov/articles/PMC6971506/
- **[S4]** 
            Circadian Transcription Factor NPAS2 and the NAD+-Dependent Deacetylase SIRT1 Interact in the Mouse Nucleus Accumbens and Regulate Reward - PMC
         — https://pmc.ncbi.nlm.nih.gov/articles/PMC9355311/
- **[S5]** 
            Circadian Transcription Factor NPAS2 and the NAD+-Dependent Deacetylase SIRT1 Interact in the Mouse Nucleus Accumbens and Regulate Reward - PMC
         — https://pmc.ncbi.nlm.nih.gov/articles/PMC9355311/
- **[S6]** 
            Nicotinamide phosphoribosyltransferase contributes to cocaine addiction through sirtuin 1 - PMC
         — https://pmc.ncbi.nlm.nih.gov/articles/PMC6971506/
- **[S7]** 
            Nicotinamide phosphoribosyltransferase contributes to cocaine addiction through sirtuin 1 - PMC
         — https://pmc.ncbi.nlm.nih.gov/articles/PMC6971506/
- **[S8]** Transancestral GWAS of alcohol dependence reveals common genetic underpinnings with psychiatric disorders - PubMed — https://pubmed.ncbi.nlm.nih.gov/30482948/
- **[S9]** Nicotinamide phosphoribosyltransferase regulates cocaine reward through Sirtuin 1 - PubMed — https://pubmed.ncbi.nlm.nih.gov/29753648/?utm_source=openai
- **[S10]** Physiological and pathophysiological roles of NAMPT and NAD metabolism - PubMed — https://pubmed.ncbi.nlm.nih.gov/26215259/?utm_source=openai
- **[S11]** Nampt is required for long-term depression and the function of GluN2B subunit-containing NMDA receptors - PubMed — https://pubmed.ncbi.nlm.nih.gov/26481044/
- **[S12]** Sobriety and Satiety: Is NAD+ the Answer? — https://www.mdpi.com/2076-3921/9/5/425/html
- **[S13]** Nicotinamide phosphoribosyltransferase regulates cocaine reward through Sirtuin 1 - PubMed — https://pubmed.ncbi.nlm.nih.gov/29753648/?utm_source=openai
- **[S14]** 
            Nicotinamide phosphoribosyltransferase contributes to cocaine addiction through sirtuin 1 - PMC
         — https://pmc.ncbi.nlm.nih.gov/articles/PMC6971506/
