## Supplementary material for "Understanding Substance Dependence: What Differentiates Addictive from Non-Addictive Drugs?": Folder for supplementary files: addictive-genes-claims-first100-033-pcsk6-gene-protein-is-implicated-in-addiction-d08bb50df07f.md

# addictive-genes-claims-first100-033-d08bb50df07f

**Original claim:** PCSK6 gene/protein is implicated in addiction.

## Claim

**PCSK6 gene/protein is implicated in addiction.**

## Verdict

Verdict: Likely False

## Confidence

**Medium**

## Supporting evidence

There is **very limited, preliminary** evidence that even touches this claim. A 2010 conference abstract describing a GWAS of **DSM-IV nicotine withdrawal** reported that two of the top three SNPs were located within **TMEM62 and PCSK6**. However, this signal was only among SNPs with **p < 1×10⁻⁴**, and the authors explicitly stated that the findings **required expansion and replication**. This is not the same as a replicated, peer-reviewed association with addiction broadly, or even with nicotine dependence specifically. ([srnt.org](https://srnt.org/wp-content/uploads/2024/11/2010_Annual_Meeting_Abstract.pdf)) [S1]

PCSK6 is also **biologically relevant to the nervous system** in a general sense. NCBI describes PCSK6 as a subtilisin-like proprotein convertase expressed in many tissues, including **neuroendocrine tissues and brain**, and the Human Protein Atlas reports **brain-enhanced** expression with detection across all major sampled brain regions. ([ncbi.nlm.nih.gov](https://www.ncbi.nlm.nih.gov/gene/5046?utm_source=openai)) [S2]

Human genetic studies have linked PCSK6 to **brain asymmetry/handedness-related traits**, not addiction. For example, PCSK6 was associated with handedness in individuals with dyslexia, and later work linked PCSK6 variants to handedness, structural brain asymmetry, and some dimensional traits such as autism-spectrum score or magical ideation in non-clinical samples. These findings support a **neurodevelopmental** role for PCSK6, which gives some indirect plausibility that it could matter in brain-related phenotypes. ([pmc.ncbi.nlm.nih.gov](https://pmc.ncbi.nlm.nih.gov/articles/PMC3016905/?utm_source=openai)) [S3]

More recently, a mouse/cellular Alzheimer’s disease study found that PCSK6 can activate **MT5-MMP**, influence APP processing, and affect **hippocampal long-term potentiation and memory**. Again, this shows CNS relevance, but it does **not** show addiction relevance. ([pubmed.ncbi.nlm.nih.gov](https://pubmed.ncbi.nlm.nih.gov/38216110/?utm_source=openai)) [S4]

## Contradicting or limiting evidence

The biggest limitation is that I could not find **robust, replicated, peer-reviewed evidence** directly connecting PCSK6 to addiction, substance use disorder, craving, relapse, reinforcement, or drug self-administration. The one direct signal I found was the **conference abstract** above, which is preliminary and not sufficient on its own to establish implication in addiction. ([srnt.org](https://srnt.org/wp-content/uploads/2024/11/2010_Annual_Meeting_Abstract.pdf)) [S5]

Importantly, later **peer-reviewed** nicotine-withdrawal GWAS work pointed to **other loci**, not PCSK6. A Finnish twin GWAS highlighted associations for nicotine withdrawal at loci including **TMEM163, AP2A2, and MUC6**. ([pmc.ncbi.nlm.nih.gov](https://pmc.ncbi.nlm.nih.gov/articles/PMC6519128/?utm_source=openai)) [S6] Another peer-reviewed GWAS of DSM-IV nicotine withdrawal identified a **protocadherin gene cluster regulatory variant** associated with nicotine withdrawal and urge to smoke. ([pmc.ncbi.nlm.nih.gov](https://pmc.ncbi.nlm.nih.gov/articles/PMC5390815/?utm_source=openai)) [S7]

Likewise, modern reviews of substance-use-disorder genetics emphasize other genes/loci as established or emerging findings. For nicotine-related phenotypes, the strongest replicated signals involve loci such as **CHRNA5–CHRNA3–CHRNB4**, **CYP2A6**, and **CHRNA4**; for other SUDs, reviews discuss genes such as **FOXP2**, **EPHX2**, and others depending on the phenotype. PCSK6 is **not** presented as an established addiction gene in these reviews. ([pmc.ncbi.nlm.nih.gov](https://pmc.ncbi.nlm.nih.gov/articles/PMC9210391/?utm_source=openai)) [S8]

The broader PCSK6 literature is also dominated by **cardiovascular**, **developmental asymmetry**, **fibrosis**, and some **neurodegeneration** topics rather than addiction. NCBI Gene’s summary and bibliography, along with recent reviews, highlight roles in precursor-protein processing, left-right patterning, vascular biology, hypertension, and idiopathic pulmonary fibrosis. ([ncbi.nlm.nih.gov](https://www.ncbi.nlm.nih.gov/gene/5046?utm_source=openai)) [S9]

So the direct evidence base is not just weak; it is also **outcompeted by other genes** in later addiction GWAS and reviews.

## Mechanistic / biological rationale

Mechanistically, PCSK6 is a **secreted serine endoprotease** that cleaves precursor proteins at basic residues and helps activate or inactivate multiple extracellular substrates. It participates in the constitutive secretory pathway and has known roles in processing signaling molecules such as **NODAL/TGF-β-related proteins**, as well as other substrates in vascular and developmental biology. ([ncbi.nlm.nih.gov](https://www.ncbi.nlm.nih.gov/gene/5046?utm_source=openai)) [S10]

That biology makes it conceivable that PCSK6 could influence neural development or neural homeostasis indirectly. Its links to **left-right patterning**, **brain asymmetry**, and **hippocampal plasticity in an Alzheimer’s model** show that it can matter in brain structure and function. ([pmc.ncbi.nlm.nih.gov](https://pmc.ncbi.nlm.nih.gov/articles/PMC3016905/?utm_source=openai)) [S11]

But addiction claims need a tighter chain of evidence: for example, replicated association with substance-related phenotypes, altered expression in addiction-relevant circuits, or behavioral effects in animal models of drug reward, withdrawal, relapse, or self-administration. I did **not** find convincing evidence that PCSK6 has been shown to regulate the canonical addiction systems most often implicated in substance use disorders, such as **mesolimbic dopamine signaling, cue-reward learning, reinforcement, or relapse circuitry**. The available PCSK6 brain evidence is better described as **neurodevelopmental or neurodegenerative**, not addiction-specific. ([pubmed.ncbi.nlm.nih.gov](https://pubmed.ncbi.nlm.nih.gov/38216110/?utm_source=openai)) [S12]

So the mechanistic case is **indirect and speculative**: a brain-expressed protease with some neural relevance is not enough to conclude addiction implication.

## Bottom line

PCSK6 is a **brain-expressed protease with credible roles in neurodevelopment and some CNS biology**, but that is **not the same** as being implicated in addiction. The only direct addiction-adjacent signal I found was a **single preliminary conference abstract** on nicotine withdrawal that specifically called for replication. Later peer-reviewed GWAS and reviews of addiction genetics point instead to **other loci**, and I found no solid human, animal, or clinical evidence establishing PCSK6 as an addiction gene/protein. ([srnt.org](https://srnt.org/wp-content/uploads/2024/11/2010_Annual_Meeting_Abstract.pdf)) [S13]

**Best current interpretation:** PCSK6 is **not an established addiction-related gene**; at most, it has **weak, unreplicated, indirect** relevance.

## Cited sources

- **[S1]** 2010 SRNT Proceedings-F — https://srnt.org/wp-content/uploads/2024/11/2010_Annual_Meeting_Abstract.pdf
- **[S2]** PCSK6 proprotein convertase subtilisin/kexin type 6 [Homo sapiens (human)] - Gene - NCBI — https://www.ncbi.nlm.nih.gov/gene/5046?utm_source=openai
- **[S3]** PCSK6 is associated with handedness in individuals with dyslexia - PMC — https://pmc.ncbi.nlm.nih.gov/articles/PMC3016905/?utm_source=openai
- **[S4]** PCSK6 exacerbates Alzheimer's disease pathogenesis by promoting MT5-MMP maturation - PubMed — https://pubmed.ncbi.nlm.nih.gov/38216110/?utm_source=openai
- **[S5]** 2010 SRNT Proceedings-F — https://srnt.org/wp-content/uploads/2024/11/2010_Annual_Meeting_Abstract.pdf
- **[S6]** Genome‐wide association study in Finnish twins highlights the connection between nicotine addiction and neurotrophin signaling pathway - PMC — https://pmc.ncbi.nlm.nih.gov/articles/PMC6519128/?utm_source=openai
- **[S7]** A protocadherin gene cluster regulatory variant is associated with nicotine withdrawal and the urge to smoke - PMC — https://pmc.ncbi.nlm.nih.gov/articles/PMC5390815/?utm_source=openai
- **[S8]** Genetics of substance use disorders in the era of big data - PMC — https://pmc.ncbi.nlm.nih.gov/articles/PMC9210391/?utm_source=openai
- **[S9]** PCSK6 proprotein convertase subtilisin/kexin type 6 [Homo sapiens (human)] - Gene - NCBI — https://www.ncbi.nlm.nih.gov/gene/5046?utm_source=openai
- **[S10]** PCSK6 proprotein convertase subtilisin/kexin type 6 [Homo sapiens (human)] - Gene - NCBI — https://www.ncbi.nlm.nih.gov/gene/5046?utm_source=openai
- **[S11]** PCSK6 is associated with handedness in individuals with dyslexia - PMC — https://pmc.ncbi.nlm.nih.gov/articles/PMC3016905/?utm_source=openai
- **[S12]** PCSK6 exacerbates Alzheimer's disease pathogenesis by promoting MT5-MMP maturation - PubMed — https://pubmed.ncbi.nlm.nih.gov/38216110/?utm_source=openai
- **[S13]** 2010 SRNT Proceedings-F — https://srnt.org/wp-content/uploads/2024/11/2010_Annual_Meeting_Abstract.pdf
