## Supplementary material for "Understanding Substance Dependence: What Differentiates Addictive from Non-Addictive Drugs?": Folder for supplementary files: addictive-genes-claims-first100-034-pmp22-gene-protein-is-implicated-in-addiction-10218677e87d.md

# addictive-genes-claims-first100-034-10218677e87d

**Original claim:** PMP22 gene/protein is implicated in addiction.

## Claim

PMP22 gene/protein is implicated in addiction.

## Verdict

Verdict: Mixed Evidence

There is **some** evidence linking **PMP22** to **alcohol-related addiction phenotypes**, but it is **indirect, limited, and not sufficient to make PMP22 an established addiction gene/protein**. The strongest positive evidence comes from **postmortem human brain transcriptomic studies in alcoholism/AUD** and a **rat alcohol-dependence model**, where PMP22/Pmp22 appears among differentially expressed or alcohol-responsive genes. However, I did **not** find convincing evidence of **replicated human genetic association**, **genome-wide significant addiction risk signals**, or **direct mechanistic experiments** showing that altering PMP22 changes addictive behavior. ([genomebiology.biomedcentral.com](https://genomebiology.biomedcentral.com/articles/10.1186/gb-2012-13-2-239/tables/1)) [S1]

## Confidence

Medium

The literature is enough to say the claim is **not baseless**, especially for **alcohol dependence/AUD**, but the evidence is mostly **expression-level association** rather than **causal addiction biology**. The gap between “altered in addicted/alcohol-exposed brains” and “implicated in addiction mechanisms” is substantial here. ([pmc.ncbi.nlm.nih.gov](https://pmc.ncbi.nlm.nih.gov/articles/PMC4047192/)) [S2]

## Supporting evidence

A review of alcohol-related candidate genes reported **PMP22** as a replicated finding from **postmortem brain microarray studies** in alcoholism, specifically in **prefrontal cortex** and **frontal cortex/nucleus accumbens** samples. Importantly, the same review lists PMP22 under **microarray** evidence rather than **GWAS**, meaning the support is transcriptomic rather than inherited-risk based. ([genomebiology.biomedcentral.com](https://genomebiology.biomedcentral.com/articles/10.1186/gb-2012-13-2-239/tables/1)) [S3]

A broader review of **human substance-use transcriptomics** concluded that myelination-related genes are repeatedly altered in substance-use disorders and stated that, among reviewed substance-use datasets, **PMP22 changes were identified exclusively in alcoholic cases**. That places PMP22 within the alcohol-related molecular signature observed in human postmortem brain, even if it does not prove causality. ([pmc.ncbi.nlm.nih.gov](https://pmc.ncbi.nlm.nih.gov/articles/PMC2593736/)) [S4]

The original human postmortem study by Flatscher-Bader et al. compared **prefrontal cortex** and **nucleus accumbens** from alcoholics and controls and found region-specific **alcohol-responsive genes** in mesocorticolimbic circuitry, supporting the idea that chronic alcohol dependence is associated with broad transcriptional remodeling in addiction-relevant brain regions. The later review cited this study as one of the sources linking PMP22 to alcoholic brain transcriptomes. ([pubmed.ncbi.nlm.nih.gov](https://pubmed.ncbi.nlm.nih.gov/15816859/?utm_source=openai)) [S5]

In a **rat medial prefrontal cortex** study after a **history of alcohol dependence**, 165 mRNAs were significantly altered; **Pmp22** appeared in enriched gene ontology categories including **transmission of nerve impulse**, **cellular ion homeostasis**, **cellular homeostasis**, and **myelin sheath**. The paper also framed this as a model of **protracted abstinence**, a stage associated with **craving and relapse risk**, which makes the finding relevant to addiction biology rather than merely acute intoxication. ([pmc.ncbi.nlm.nih.gov](https://pmc.ncbi.nlm.nih.gov/articles/PMC3546132/)) [S6]

A later human dorsolateral prefrontal cortex exon-microarray study in alcoholism likewise summarized prior work as showing disturbances in **myelination-related genes**, specifically naming **PMP22** among genes previously reported as altered in alcoholic frontal cortex. This gives some cross-study consistency within alcohol-related brain transcriptomics. ([pmc.ncbi.nlm.nih.gov](https://pmc.ncbi.nlm.nih.gov/articles/PMC4047192/)) [S7]

## Contradicting or limiting evidence

The main limitation is that **PMP22’s established biology is not addiction-specific**. PMP22 encodes **peripheral myelin protein 22**, is found mainly in the **peripheral nervous system**, is produced primarily by **Schwann cells**, and has a central known role in **myelin development/maintenance** and **dosage-sensitive peripheral neuropathies** such as **Charcot-Marie-Tooth disease** and **hereditary neuropathy with liability to pressure palsies**. That is a very different evidence base from the classic neurotransmission, reward, stress, or synaptic-plasticity genes usually implicated in addiction. ([medlineplus.gov](https://medlineplus.gov/genetics/gene/pmp22)) [S8]

There is also a notable lack of **human genetics evidence**. A direct search of the **NHGRI-EBI GWAS Catalog** for **PMP22** returned **0 studies, 0 publications, 0 variants, 0 traits, and 0 genes**, which argues against PMP22 being an established genome-wide addiction-risk locus at present. ([ebi.ac.uk](https://www.ebi.ac.uk/gwas/search?query=PMP22)) [S9]

Similarly, the alcoholism review that includes PMP22 explicitly categorizes it as supported by **microarray/postmortem expression** data, **not** by GWAS. So even within alcohol research, the evidence points more to a **state marker or downstream consequence** of chronic alcohol exposure than to a robust inherited susceptibility gene. ([genomebiology.biomedcentral.com](https://genomebiology.biomedcentral.com/articles/10.1186/gb-2012-13-2-239/tables/1)) [S10]

Postmortem transcriptomic studies also have important confounders. Reviews of human substance-use brain expression datasets note that **agonal state/brain pH**, **smoking history**, and other clinical variables can substantially affect expression profiles. The alcoholism exon-array study also notes that **co-morbid tobacco use**, **hepatic disease/cirrhosis**, and other medical complications may influence brain gene expression. ([pmc.ncbi.nlm.nih.gov](https://pmc.ncbi.nlm.nih.gov/articles/PMC2593736/)) [S11]

Finally, one of the later alcoholism transcriptomic papers explicitly states that although white-matter and developmental gene disturbances are seen in alcoholic cortex, **the specific impact of these disturbances on abuse behavior itself remains to be elucidated**. That is a direct warning against overinterpreting PMP22 as a mechanistic addiction driver. ([pmc.ncbi.nlm.nih.gov](https://pmc.ncbi.nlm.nih.gov/articles/PMC4047192/)) [S12]

## Mechanistic / biological rationale

A biologically plausible, but still **indirect**, rationale exists. Addiction involves **mesolimbic/prefrontal circuitry**, and chronic alcohol exposure is associated with transcriptional changes in **prefrontal cortex** and **nucleus accumbens**, the same general systems that regulate reward, executive control, craving, and relapse. White-matter and glial abnormalities can alter signal propagation and network coordination, so genes tied to myelin biology can plausibly modulate neural function in addiction-relevant circuits. ([pmc.ncbi.nlm.nih.gov](https://pmc.ncbi.nlm.nih.gov/articles/PMC4047192/)) [S13]

PMP22 itself is a **dosage-sensitive myelin gene** whose expression must be tightly controlled for normal myelin homeostasis. It is also regulated post-transcriptionally by **miR-29a** in Schwann cells, illustrating that PMP22 sits in regulatory pathways that can respond to cellular state changes. In principle, long-term alcohol-related dysregulation of myelin-associated genes could contribute to altered circuit function. ([pubmed.ncbi.nlm.nih.gov](https://pubmed.ncbi.nlm.nih.gov/31586623/)) [S14]

But the mechanistic case weakens when one asks whether this is **core addiction biology** or **secondary tissue pathology**. PMP22 is primarily a **peripheral Schwann-cell/myelin** protein, not a canonical synaptic, dopaminergic, glutamatergic, GABAergic, or transcription-factor hub of addiction. So the most plausible interpretation is that PMP22 changes reflect **alcohol-related myelin/glial remodeling or white-matter injury**, rather than a directly demonstrated role in reward, reinforcement, or compulsive drug seeking. ([medlineplus.gov](https://medlineplus.gov/genetics/gene/pmp22)) [S15]

## Bottom line

**PMP22 is not a well-established addiction gene/protein.** There is **real but limited evidence** that it is **associated with alcohol dependence/AUD-related brain transcriptomic changes**, and that makes the claim **partly supportable** in a narrow, indirect sense. But the current literature does **not** show strong human genetic evidence, direct behavioral causality, or a clearly defined addiction-specific mechanism. The fairest summary is: **PMP22 is an indirect, alcohol-linked transcriptomic correlate of addiction-related states, not a firmly validated addiction gene.** ([genomebiology.biomedcentral.com](https://genomebiology.biomedcentral.com/articles/10.1186/gb-2012-13-2-239/tables/1)) [S16]

## Cited sources

- **[S1]** The genetic basis of alcoholism: multiple phenotypes, many genes, complex networks | Genome Biology | Full Text — https://genomebiology.biomedcentral.com/articles/10.1186/gb-2012-13-2-239/tables/1
- **[S2]** 
            Exon Microarray Analysis of Human Dorsolateral Prefrontal Cortex in Alcoholism - PMC
         — https://pmc.ncbi.nlm.nih.gov/articles/PMC4047192/
- **[S3]** The genetic basis of alcoholism: multiple phenotypes, many genes, complex networks | Genome Biology | Full Text — https://genomebiology.biomedcentral.com/articles/10.1186/gb-2012-13-2-239/tables/1
- **[S4]** 
            Transcriptional correlates of human substance use - PMC
         — https://pmc.ncbi.nlm.nih.gov/articles/PMC2593736/
- **[S5]** Alcohol-responsive genes in the frontal cortex and nucleus accumbens of human alcoholics - PubMed — https://pubmed.ncbi.nlm.nih.gov/15816859/?utm_source=openai
- **[S6]** 
            Coordinated dysregulation of mRNAs and microRNAs in the rat medial prefrontal cortex following a history of alcohol dependence - PMC
         — https://pmc.ncbi.nlm.nih.gov/articles/PMC3546132/
- **[S7]** 
            Exon Microarray Analysis of Human Dorsolateral Prefrontal Cortex in Alcoholism - PMC
         — https://pmc.ncbi.nlm.nih.gov/articles/PMC4047192/
- **[S8]** PMP22 gene: MedlinePlus Genetics — https://medlineplus.gov/genetics/gene/pmp22
- **[S9]** GWAS Catalog — https://www.ebi.ac.uk/gwas/search?query=PMP22
- **[S10]** The genetic basis of alcoholism: multiple phenotypes, many genes, complex networks | Genome Biology | Full Text — https://genomebiology.biomedcentral.com/articles/10.1186/gb-2012-13-2-239/tables/1
- **[S11]** 
            Transcriptional correlates of human substance use - PMC
         — https://pmc.ncbi.nlm.nih.gov/articles/PMC2593736/
- **[S12]** 
            Exon Microarray Analysis of Human Dorsolateral Prefrontal Cortex in Alcoholism - PMC
         — https://pmc.ncbi.nlm.nih.gov/articles/PMC4047192/
- **[S13]** 
            Exon Microarray Analysis of Human Dorsolateral Prefrontal Cortex in Alcoholism - PMC
         — https://pmc.ncbi.nlm.nih.gov/articles/PMC4047192/
- **[S14]** Regulating PMP22 expression as a dosage sensitive neuropathy gene - PubMed — https://pubmed.ncbi.nlm.nih.gov/31586623/
- **[S15]** PMP22 gene: MedlinePlus Genetics — https://medlineplus.gov/genetics/gene/pmp22
- **[S16]** The genetic basis of alcoholism: multiple phenotypes, many genes, complex networks | Genome Biology | Full Text — https://genomebiology.biomedcentral.com/articles/10.1186/gb-2012-13-2-239/tables/1
