## Supplementary material for "Understanding Substance Dependence: What Differentiates Addictive from Non-Addictive Drugs?": Folder for supplementary files: addictive-genes-claims-first100-035-ppp2r1a-gene-protein-is-implicated-in-addiction-53989adbeef9.md

# addictive-genes-claims-first100-035-53989adbeef9

**Original claim:** PPP2R1A gene/protein is implicated in addiction.

## Claim

**PPP2R1A gene/protein is implicated in addiction.**

## Verdict

Verdict: No Evidence

I did **not find direct, specific evidence** that **PPP2R1A** itself is a replicated addiction gene/protein in humans or in addiction-focused animal models. What I did find is that **PP2A biology more broadly** is involved in addiction-related phenotypes, and PPP2R1A is the **scaffolding Aα subunit** of the PP2A holoenzyme. That creates **biological plausibility**, but it is **not the same as evidence that PPP2R1A itself is an established addiction factor**. ([ncbi.nlm.nih.gov](https://www.ncbi.nlm.nih.gov/books/NBK580243/)) [S1]

## Confidence

**Medium**

Confidence is medium because the literature supports a role for the **PP2A pathway/complex** in several addiction-relevant paradigms, but the evidence I found does **not isolate PPP2R1A** as the implicated component. Large modern human genetics / multi-omics studies of substance use disorders that I checked did not identify or even mention PPP2R1A in the article text. ([pmc.ncbi.nlm.nih.gov](https://pmc.ncbi.nlm.nih.gov/articles/PMC10217792/)) [S2]

## Supporting evidence

PPP2R1A encodes the **Aα scaffolding subunit of protein phosphatase 2A (PP2A)**, which assembles PP2A catalytic and regulatory subunits. GeneReviews describes PPP2R1A as the PP2A scaffolding subunit and emphasizes its importance in brain biology; pathogenic variants cause a neurodevelopmental disorder, not an addiction syndrome. ([ncbi.nlm.nih.gov](https://www.ncbi.nlm.nih.gov/books/NBK580243/)) [S3]

PPP2R1A is also broadly expressed in the brain. Human Protein Atlas data show expression across addiction-relevant regions such as **amygdala, basal ganglia, cerebral cortex, hippocampal formation, hypothalamus, midbrain, and thalamus**. A recent mouse study similarly found widespread brain expression, with relatively high expression in cortex/hippocampus and notable expression in **glutamatergic neurons**. ([v20.proteinatlas.org](https://v20.proteinatlas.org/ENSG00000105568-PPP2R1A/tissue)) [S4]

That same 2025 mouse study provides an important **indirect** link to addiction-relevant neurobiology: Ppp2r1a haploinsufficiency increased excitatory synaptic transmission in medial prefrontal cortex neurons and impaired **endocannabinoid (2-AG) signaling** via increased MAGL expression. The authors concluded that impaired 2-AG signaling contributed to altered synaptic transmission and behavioral deficits. Because endocannabinoid signaling is a recognized modulator of reward, habit formation, and drug reinforcement, these findings make PPP2R1A **mechanistically plausible** as a participant in circuits relevant to addiction. ([pmc.ncbi.nlm.nih.gov](https://pmc.ncbi.nlm.nih.gov/articles/PMC12404745/)) [S5]

More generally, **PP2A** has been implicated in addiction-related animal models. In mice, PP2A signaling in the caudate-putamen contributed to **acute methamphetamine-induced hyperlocomotion**. In another study, hippocampal PP2A deficiency reduced **morphine priming** in conditioned place preference. A 2024 study further showed that disrupting the **PP2A–GABA\_B receptor interaction** in the ventral tegmental area reduced methamphetamine sensitization and drug-seeking behavior. These studies support the idea that **PP2A-dependent phosphatase signaling** matters in addiction biology. ([pubmed.ncbi.nlm.nih.gov](https://pubmed.ncbi.nlm.nih.gov/34237414/)) [S6]

## Contradicting or limiting evidence

The major limitation is simple: the evidence above is about **PP2A**, **PP2A activity**, or **PP2A interactions**, **not PPP2R1A specifically**. None of the addiction studies I located manipulated PPP2R1A directly or showed that PPP2R1A variants/expression changes predict addiction risk, severity, or relapse. ([academic.oup.com](https://academic.oup.com/cercor/article/33/11/6594/6982733)) [S7]

Likewise, I did **not identify PPP2R1A as a highlighted signal** in major contemporary human studies of substance use disorders. In a large multivariate GWAS/meta-analysis of over 1 million subjects for multiple SUDs, a text search of the article did not find PPP2R1A. The same was true for a large cross-SUD meta-analysis in diverse populations and for recent human cocaine-use-disorder multi-omics / transcriptomic papers I checked. That does not prove PPP2R1A has no role, but it argues against it being an established or replicated addiction gene at present. ([pmc.ncbi.nlm.nih.gov](https://pmc.ncbi.nlm.nih.gov/articles/PMC10217792/)) [S8]

A broader umbrella review of molecular genetics of substance use disorders summarized repeatedly implicated genes such as **OPRM1, DRD2, DRD4, BDNF, SLC6A4, CHRNA5, FOXP2, PEX, and AUTS2**; PPP2R1A was not among the genes highlighted. ([pubmed.ncbi.nlm.nih.gov](https://pubmed.ncbi.nlm.nih.gov/33556390/)) [S9]

Finally, the best-established disease association for PPP2R1A is **PPP2R1A-related neurodevelopmental disorder**, with developmental delay, intellectual disability, hypotonia, epilepsy, and brain malformations in some patients. That phenotype is neurologic/developmental rather than addiction-related. ([ncbi.nlm.nih.gov](https://www.ncbi.nlm.nih.gov/books/NBK580243/)) [S10]

## Mechanistic / biological rationale

A biologic rationale **can** be made, but it remains indirect. PPP2R1A is the structural subunit that helps assemble PP2A holoenzymes, and PP2A regulates phosphorylation states of proteins involved in synaptic plasticity, inhibitory signaling, and reward-circuit function. ([ncbi.nlm.nih.gov](https://www.ncbi.nlm.nih.gov/books/NBK580243/)) [S11]

In addiction models, PP2A has been tied to at least three relevant processes:  
1. **dopamine/CaMKII-linked methamphetamine responses**,  
2. **morphine-associated memory/priming**, and  
3. **GABA\_B receptor trafficking/signaling** affecting methamphetamine sensitization and seeking. ([pubmed.ncbi.nlm.nih.gov](https://pubmed.ncbi.nlm.nih.gov/34237414/)) [S12]

PPP2R1A-specific mouse work also suggests that reduced PPP2R1A can shift the excitation/inhibition balance and weaken **2-AG endocannabinoid signaling**, partly through MAGL upregulation. Since endocannabinoid signaling has a well-described role in reward and addiction, one could infer that PPP2R1A perturbation **might** influence addiction vulnerability or drug responses. But that is still an inference from circuit biology, **not direct addiction evidence**. ([pmc.ncbi.nlm.nih.gov](https://pmc.ncbi.nlm.nih.gov/articles/PMC12404745/)) [S13]

## Bottom line

The claim is **not currently supported directly**. The literature supports a role for **PP2A-related signaling** in addiction-relevant behaviors, and PPP2R1A is an essential PP2A scaffolding subunit expressed in the brain, so there is **indirect mechanistic plausibility**. But I did **not find direct human genetic, transcriptomic, or causal animal evidence** establishing **PPP2R1A itself** as an addiction-implicated gene/protein. The most accurate summary is:

**PPP2R1A is biologically relevant to neural signaling pathways that overlap with addiction biology, but it is not presently an established addiction gene based on the available evidence I found.** ([ncbi.nlm.nih.gov](https://www.ncbi.nlm.nih.gov/books/NBK580243/)) [S14]

## Cited sources

- **[S1]** PPP2R1A-Related Neurodevelopmental Disorder - GeneReviews® - NCBI Bookshelf — https://www.ncbi.nlm.nih.gov/books/NBK580243/
- **[S2]** 
            Multivariate genome-wide association meta-analysis of over 1 million subjects identifies loci underlying multiple substance use disorders - PMC
         — https://pmc.ncbi.nlm.nih.gov/articles/PMC10217792/
- **[S3]** PPP2R1A-Related Neurodevelopmental Disorder - GeneReviews® - NCBI Bookshelf — https://www.ncbi.nlm.nih.gov/books/NBK580243/
- **[S4]** Tissue expression of PPP2R1A - Summary - The Human Protein Atlas — https://v20.proteinatlas.org/ENSG00000105568-PPP2R1A/tissue
- **[S5]** 
            Ppp2r1a haploinsufficiency increases excitatory synaptic transmission and decreases spatial learning by impairing endocannabinoid signaling - PMC
         — https://pmc.ncbi.nlm.nih.gov/articles/PMC12404745/
- **[S6]** D1R/PP2A/p-CaMKIIα signaling in the caudate putamen is involved in acute methamphetamine-induced hyperlocomotion - PubMed — https://pubmed.ncbi.nlm.nih.gov/34237414/
- **[S7]** Untitled source — https://academic.oup.com/cercor/article/33/11/6594/6982733
- **[S8]** 
            Multivariate genome-wide association meta-analysis of over 1 million subjects identifies loci underlying multiple substance use disorders - PMC
         — https://pmc.ncbi.nlm.nih.gov/articles/PMC10217792/
- **[S9]** Molecular genetics of substance use disorders: An umbrella review - PubMed — https://pubmed.ncbi.nlm.nih.gov/33556390/
- **[S10]** PPP2R1A-Related Neurodevelopmental Disorder - GeneReviews® - NCBI Bookshelf — https://www.ncbi.nlm.nih.gov/books/NBK580243/
- **[S11]** PPP2R1A-Related Neurodevelopmental Disorder - GeneReviews® - NCBI Bookshelf — https://www.ncbi.nlm.nih.gov/books/NBK580243/
- **[S12]** D1R/PP2A/p-CaMKIIα signaling in the caudate putamen is involved in acute methamphetamine-induced hyperlocomotion - PubMed — https://pubmed.ncbi.nlm.nih.gov/34237414/
- **[S13]** 
            Ppp2r1a haploinsufficiency increases excitatory synaptic transmission and decreases spatial learning by impairing endocannabinoid signaling - PMC
         — https://pmc.ncbi.nlm.nih.gov/articles/PMC12404745/
- **[S14]** PPP2R1A-Related Neurodevelopmental Disorder - GeneReviews® - NCBI Bookshelf — https://www.ncbi.nlm.nih.gov/books/NBK580243/
