## Supplementary material for "Understanding Substance Dependence: What Differentiates Addictive from Non-Addictive Drugs?": Folder for supplementary files: addictive-genes-claims-first100-036-ppp3r1-gene-protein-is-implicated-in-addiction-4fa10dcfa52a.md

# addictive-genes-claims-first100-036-4fa10dcfa52a

**Original claim:** PPP3R1 gene/protein is implicated in addiction.

## Claim

PPP3R1 gene/protein is implicated in addiction.

## Verdict

Verdict: Mixed Evidence

PPP3R1 is **plausibly involved** in addiction-related biology, and there is **suggestive preclinical evidence** linking it to alcohol and broader drug-response pathways. However, the strongest evidence is mostly for **calcineurin signaling as a pathway** rather than for **PPP3R1 itself as a directly established addiction gene/protein**, especially in humans. ([proteinatlas.org](https://www.proteinatlas.org/ENSG00000221823-PPP3R1/tissue)) [S1]

## Confidence

Medium

## Supporting evidence

PPP3R1 encodes the **regulatory B subunit of calcineurin (protein phosphatase 3/PP2B)**, a calcium/calmodulin-dependent phosphatase with clear neuronal relevance. Human Protein Atlas data place PPP3R1 in a **brain/retina neuronal signaling expression cluster**, with enhanced expression in brain and retina, which makes it biologically credible as a modulator of addiction-relevant neural plasticity. ([ncbi.nlm.nih.gov](https://www.ncbi.nlm.nih.gov/gene/5534?utm_source=openai)) [S2]

In **alcohol-related models**, PPP3R1 shows up repeatedly in drug-responsive datasets. In a mouse nucleus accumbens proteomics study after chronic alcohol drinking, PPP3R1 was among the proteins significantly altered and grouped in a functional cluster containing the Gene Ontology category **“response to alcohol.”** In a rhesus macaque ethanol self-administration transcriptomic study, **Ppp3r1** emerged as an **intake-correlated hub** in cortical Area 32 after more than 12 months of alcohol self-administration. These are not proof of causality, but they do place PPP3R1 within alcohol-responsive molecular networks in addiction-relevant brain regions. ([pubmed.ncbi.nlm.nih.gov](https://pubmed.ncbi.nlm.nih.gov/29502276/)) [S3]

There is also stronger **pathway-level** evidence in alcohol models implicating calcineurin. A review focused on alcohol use disorder concluded that calcineurin signaling is a relevant target in AUD, and a mouse study found that the calcineurin inhibitor **tacrolimus** reduced binge-like ethanol intake and blood alcohol levels in High Drinking in the Dark mice. That does not isolate PPP3R1 specifically, but because PPP3R1 is part of the calcineurin holoenzyme, it supports the claim that the **PPP3R1-containing pathway** is addiction-relevant. ([pubmed.ncbi.nlm.nih.gov](https://pubmed.ncbi.nlm.nih.gov/31601401/)) [S4]

For **cocaine**, several studies support calcineurin involvement in addiction-related neuroadaptations. Repeated cocaine administration decreased calcineurin in rat nucleus accumbens neurons and altered downstream excitability-related signaling. Another study showed that systemic **cyclosporine**, a calcineurin inhibitor, changed cocaine-induced locomotor responses and mimicked cocaine-related phosphorylation changes in synapsin I. A third study found that cocaine regulates **MEF2-dependent synaptic and behavioral plasticity** partly through a mechanism involving **calcineurin**. Together, these data strongly implicate calcineurin signaling in cocaine-induced plasticity, even if they do not prove PPP3R1 is the unique driver. ([nature.com](https://www.nature.com/articles/1300654)) [S5]

For **opioid-related biology**, calcineurin again appears relevant. In cultured hippocampal neurons, morphine increased calcineurin activity and drove **GluR1 dephosphorylation and AMPA receptor internalization**, effects blocked by FK506. In rats, opiate exposure state altered **cdk5/calcineurin signaling** in the basolateral amygdala during reward and withdrawal-aversion memory formation, with calcineurin inhibition switching the memory mechanism. These findings fit well with addiction as a disorder of maladaptive synaptic plasticity and associative memory. ([pubmed.ncbi.nlm.nih.gov](https://pubmed.ncbi.nlm.nih.gov/21068335/)) [S6]

## Contradicting or limiting evidence

The biggest limitation is that most of the evidence is **not PPP3R1-specific**. Much of the literature manipulates or measures **calcineurin as a whole pathway** rather than PPP3R1 itself, and the addiction-related behavioral studies generally do **not** use PPP3R1-selective genetic deletion, overexpression, or rescue to show that PPP3R1 alone is necessary or sufficient for addiction phenotypes. ([pubmed.ncbi.nlm.nih.gov](https://pubmed.ncbi.nlm.nih.gov/33294845/)) [S7]

Even the alcohol proteomics paper that identified PPP3R1 did **not functionally validate PPP3R1**; instead, it experimentally followed up **GSTP1** as the actionable target. So PPP3R1 was a signal in the omics data, but not the demonstrated causal mediator in that study. ([pubmed.ncbi.nlm.nih.gov](https://pubmed.ncbi.nlm.nih.gov/29502276/)) [S8]

The human evidence is also limited. A recent systematic review of functional genomic mechanisms in opioid action/OUD found that, across 73 studies, **replicable findings were surprisingly limited**, with only one gene consistently identified across many studies and substantial heterogeneity by species, brain region, exposure duration, and other factors. That review supports caution about elevating any single candidate gene—especially one like PPP3R1 that has not emerged as a major replicated human signal. ([nature.com](https://www.nature.com/articles/s41380-023-02238-1)) [S9]

Likewise, large human addiction GWAS have highlighted other genes more clearly than PPP3R1. For opioid addiction, large studies emphasized loci such as **OPRM1, FURIN, and PPP6C**. For alcohol use disorder, very large GWAS emphasized numerous loci, especially established alcohol-related and psychiatric/shared-risk loci, rather than PPP3R1 as a leading replicated finding. This does **not** rule PPP3R1 out, but it argues against calling it a firmly established human addiction gene at present. ([pmc.ncbi.nlm.nih.gov](https://pmc.ncbi.nlm.nih.gov/articles/PMC9546890/?utm_source=openai)) [S10]

## Mechanistic / biological rationale

Mechanistically, the claim is quite plausible. Calcineurin is a **Ca2+/calmodulin-activated serine/threonine phosphatase** composed of catalytic and regulatory subunits; PPP3R1 encodes the major **regulatory B subunit**. Because addiction involves repeated activity-dependent remodeling of synapses in regions such as the **nucleus accumbens, amygdala, cortex, and hippocampus**, a calcium-sensitive phosphatase that shapes phosphorylation state, receptor trafficking, and transcriptional responses is well-positioned to influence addiction-related plasticity. ([ncbi.nlm.nih.gov](https://www.ncbi.nlm.nih.gov/gene/5534?utm_source=openai)) [S11]

The available mechanistic data are consistent with that idea. Calcineurin participates in **AMPA receptor trafficking** after morphine exposure, regulates **MEF2-linked structural and behavioral plasticity** after cocaine, and is discussed in reviews as relevant to **AUD-related neuroimmune and synaptic processes**. Because PPP3R1 controls calcineurin complex function, altered PPP3R1 expression or regulation could reasonably affect reward learning, sensitization, withdrawal memory, or relapse-related circuitry. Still, that is a **mechanistic inference**, not direct proof that PPP3R1 is itself a primary addiction gene. ([pubmed.ncbi.nlm.nih.gov](https://pubmed.ncbi.nlm.nih.gov/21068335/)) [S12]

## Bottom line

PPP3R1 is best described as a **candidate addiction-related gene/protein with biologically credible, but still limited, direct evidence**. The direct PPP3R1 data are mainly **preclinical and omics-based**, while the stronger body of evidence points to **calcineurin signaling more broadly** as relevant to alcohol, cocaine, and opioid-induced neuroadaptations. ([pubmed.ncbi.nlm.nih.gov](https://pubmed.ncbi.nlm.nih.gov/28247455/)) [S13]

So if the claim means **“PPP3R1 has some evidence linking it to addiction biology”**, that is fair. If it means **“PPP3R1 is an established, replicated addiction gene in humans”**, the current literature does **not** support that stronger statement. ([nature.com](https://www.nature.com/articles/s41380-023-02238-1)) [S14]

## Cited sources

- **[S1]** Tissue expression of PPP3R1 - Summary - The Human Protein Atlas — https://www.proteinatlas.org/ENSG00000221823-PPP3R1/tissue
- **[S2]** PPP3R1 protein phosphatase 3 regulatory subunit B, alpha [Homo sapiens (human)] - Gene - NCBI — https://www.ncbi.nlm.nih.gov/gene/5534?utm_source=openai
- **[S3]** Mining the nucleus accumbens proteome for novel targets of alcohol self-administration in male C57BL/6J mice - PubMed — https://pubmed.ncbi.nlm.nih.gov/29502276/
- **[S4]** Calcineurin signaling as a target for the treatment of alcohol abuse and neuroinflammatory disorders - PubMed — https://pubmed.ncbi.nlm.nih.gov/31601401/
- **[S5]** Repeated Cocaine Administration Decreases Calcineurin (PP2B) but Enhances DARPP-32 Modulation of Sodium Currents in Rat Nucleus Accumbens Neurons | Neuropsychopharmacology — https://www.nature.com/articles/1300654
- **[S6]** Morphine induces AMPA receptor internalization in primary hippocampal neurons via calcineurin-dependent dephosphorylation of GluR1 subunits - PubMed — https://pubmed.ncbi.nlm.nih.gov/21068335/
- **[S7]** Effects of Tacrolimus and Other Immune Targeting Compounds on Binge-Like Ethanol Drinking in High Drinking in the Dark Mice - PubMed — https://pubmed.ncbi.nlm.nih.gov/33294845/
- **[S8]** Mining the nucleus accumbens proteome for novel targets of alcohol self-administration in male C57BL/6J mice - PubMed — https://pubmed.ncbi.nlm.nih.gov/29502276/
- **[S9]** Functional genomic mechanisms of opioid action and opioid use disorder: a systematic review of animal models and human studies | Molecular Psychiatry — https://www.nature.com/articles/s41380-023-02238-1
- **[S10]** Multi-trait genome-wide association study of opioid addiction: OPRM1 and beyond - PMC — https://pmc.ncbi.nlm.nih.gov/articles/PMC9546890/?utm_source=openai
- **[S11]** PPP3R1 protein phosphatase 3 regulatory subunit B, alpha [Homo sapiens (human)] - Gene - NCBI — https://www.ncbi.nlm.nih.gov/gene/5534?utm_source=openai
- **[S12]** Morphine induces AMPA receptor internalization in primary hippocampal neurons via calcineurin-dependent dephosphorylation of GluR1 subunits - PubMed — https://pubmed.ncbi.nlm.nih.gov/21068335/
- **[S13]** On the relationships in rhesus macaques between chronic ethanol consumption and the brain transcriptome - PubMed — https://pubmed.ncbi.nlm.nih.gov/28247455/
- **[S14]** Functional genomic mechanisms of opioid action and opioid use disorder: a systematic review of animal models and human studies | Molecular Psychiatry — https://www.nature.com/articles/s41380-023-02238-1
