## Supplementary material for "Understanding Substance Dependence: What Differentiates Addictive from Non-Addictive Drugs?": Folder for supplementary files: addictive-genes-claims-first100-037-rgs4-gene-protein-is-implicated-in-addiction-ac29dfa9c05f.md

# addictive-genes-claims-first100-037-ac29dfa9c05f

**Original claim:** RGS4 gene/protein is implicated in addiction.

## Claim

RGS4 gene/protein is implicated in addiction.

## Verdict

Verdict: Likely True

## Confidence

Medium.

## Supporting evidence

RGS4 is a biologically plausible addiction-related gene/protein because it is a regulator of G-protein signaling that modulates GPCR pathways central to reward and dependence, including opioid and dopamine-linked signaling. Reviews on RGS proteins and addictive drugs specifically identify RGS-family members, including RGS4, as relevant to the actions of abused substances and to addiction-related intracellular signaling. ([pubmed.ncbi.nlm.nih.gov](https://pubmed.ncbi.nlm.nih.gov/20201861/?utm_source=openai)) [S1]

The strongest evidence is in **opioid-related preclinical models**. In rats, morphine, cocaine, and amphetamine all altered RGS4 mRNA in multiple brain regions implicated in reward and dependence, suggesting that RGS4 is a common drug-responsive signaling node rather than a drug-irrelevant bystander. ([pubmed.ncbi.nlm.nih.gov](https://pubmed.ncbi.nlm.nih.gov/12270694/)) [S2]

More importantly, **direct manipulation of RGS4 changes addiction-relevant behaviors** in vivo. In mice, inducible deletion of RGS4 in the nucleus accumbens increased sensitivity to morphine reward in conditioned place preference assays, while overexpression of RGS4 in the nucleus accumbens blocked morphine CPP. The same study also found that RGS4 in the locus coeruleus opposes morphine physical dependence. ([pubmed.ncbi.nlm.nih.gov](https://pubmed.ncbi.nlm.nih.gov/19914603/)) [S3]

A later mechanistic mouse study reinforced this opioid link: nucleus accumbens knockdown of RGS4 significantly increased morphine-associated behaviors and enhanced phosphorylation of GluR1 and NR2A glutamate receptor subunits, pointing to a defined synaptic mechanism by which lower RGS4 can potentiate opioid reward/withdrawal-related signaling. ([pubmed.ncbi.nlm.nih.gov](https://pubmed.ncbi.nlm.nih.gov/29754475/?utm_source=openai)) [S4]

Other opioid-related data also support involvement. In rat locus coeruleus, chronic morphine elevated RGS4 protein, and intracellular application of wild-type RGS4 diminished electrophysiological responses to morphine; the authors concluded that RGS4 induction could contribute to opiate tolerance and dependence in this circuit. In cell models, opioid agonists reduced RGS4 protein via ubiquitin-proteasome/lysosomal pathways, and this RGS4 loss enhanced signaling cross-talk at selected receptors. ([pubmed.ncbi.nlm.nih.gov](https://pubmed.ncbi.nlm.nih.gov/12653973/)) [S5]

There is also **cocaine-related evidence**. In RGS4 knockout mice, cocaine CPP was attenuated in males, indicating that RGS4 can influence cocaine reward, although the direction differs from the opioid findings. Separately, rats with a history of cocaine self-administration followed by abstinence showed reduced dorsal striatal RGS4, and re-exposure to the cocaine-paired context triggered cocaine-seeking together with downstream PKC/ERK/CREB-related signaling changes. ([pubmed.ncbi.nlm.nih.gov](https://pubmed.ncbi.nlm.nih.gov/30036561/)) [S6]

Evidence also extends, though less strongly, to **alcohol-related phenotypes**. A human postmortem study reported lower RGS4 mRNA in superior frontal gyrus/primary motor cortex of cirrhotic alcoholics versus controls and suggested a possible protective RGS4 haplotype in European Caucasians, while a rat chronic intermittent ethanol model found increased RGS4 mRNA and protein in prefrontal cortex after exposure/withdrawal. These findings are consistent with RGS4 being altered in alcohol-related neuroadaptation. ([pmc.ncbi.nlm.nih.gov](https://pmc.ncbi.nlm.nih.gov/articles/PMC2902459/?utm_source=openai)) [S7]

## Contradicting or limiting evidence

The biggest limitation is that **most of the evidence is preclinical**. The clearest data come from rodent knockout/knockdown, overexpression, self-administration, CPP, and withdrawal models, plus cell-signaling experiments. Those models are useful and mechanistically informative, but they do not by themselves prove that RGS4 is a major causal determinant of human substance use disorders. ([pubmed.ncbi.nlm.nih.gov](https://pubmed.ncbi.nlm.nih.gov/19914603/)) [S8]

The effect of RGS4 is also **not directionally uniform across drugs, circuits, or behaviors**. In opioids, reduced RGS4 in nucleus accumbens tends to enhance morphine reward, whereas for cocaine, RGS4 deletion attenuated CPP in male mice. That pattern means RGS4 is better described as a context-dependent modulator of addiction-related neuroplasticity than as a simple “pro-addiction” factor. ([pubmed.ncbi.nlm.nih.gov](https://pubmed.ncbi.nlm.nih.gov/19914603/)) [S9]

Human evidence is limited and somewhat indirect. The alcohol postmortem/genetic study was relatively small, the haplotype finding was presented cautiously and stated to need confirmation, and the lower RGS4 expression was especially evident in **cirrhotic** alcoholics, leaving room for confounding by severe medical comorbidity, nutrition, liver disease, or other consequences of chronic alcoholism rather than addiction biology per se. ([pmc.ncbi.nlm.nih.gov](https://pmc.ncbi.nlm.nih.gov/articles/PMC2902459/?utm_source=openai)) [S10]

Also, large modern GWAS of alcohol dependence and opioid addiction have highlighted other loci—such as **ADH1B** for alcohol dependence and **OPRM1**, **PPP6C**, and **FURIN** for opioid addiction—rather than establishing RGS4 as a leading replicated genome-wide risk gene. That does not rule out RGS4 involvement, but it does weaken any claim that RGS4 is a firmly established major human addiction-susceptibility locus. ([pubmed.ncbi.nlm.nih.gov](https://pubmed.ncbi.nlm.nih.gov/30482948/)) [S11]

## Mechanistic / biological rationale

Mechanistically, the claim makes sense. RGS4 is a GTPase-accelerating regulator that shortens signaling downstream of heterotrimeric G proteins. Because many addiction-relevant receptors are GPCRs, changing RGS4 levels can reshape the intensity, duration, and bias of signaling in reward circuits. Reviews of opioid receptor signaling specifically emphasize RGS4 as a prominent modulator of MOR-family signaling. ([pubmed.ncbi.nlm.nih.gov](https://pubmed.ncbi.nlm.nih.gov/32038168/)) [S12]

RGS4 also modulates non-opioid pathways relevant to addictive neuroplasticity. It inhibits signaling by group I metabotropic glutamate receptors and can inhibit dopamine D2/D3 receptor signaling, placing it at the intersection of glutamatergic and dopaminergic control of reward learning, sensitization, withdrawal, and relapse-related plasticity. ([pubmed.ncbi.nlm.nih.gov](https://pubmed.ncbi.nlm.nih.gov/9437012/?utm_source=openai)) [S13]

The opioid studies suggest a concrete circuit model: drug exposure changes RGS4 abundance; altered RGS4 then changes GPCR signaling tone, receptor cross-talk, and downstream phosphorylation events. In nucleus accumbens this can influence glutamate receptor phosphorylation and morphine reward; in locus coeruleus it can influence dependence-related neuronal adaptations; in dorsal striatum after cocaine abstinence it may contribute to an altered Gαq/Gαi balance and relapse-associated PKC/ERK/CREB signaling. ([pubmed.ncbi.nlm.nih.gov](https://pubmed.ncbi.nlm.nih.gov/29754475/)) [S14]

## Bottom line

The claim is **likely true**, but mainly because of **convergent animal and mechanistic evidence**, especially for **opioid reward/dependence** and to a lesser extent **cocaine** and **alcohol-related neuroadaptation**. RGS4 clearly influences addiction-relevant signaling and behaviors in preclinical systems. ([pubmed.ncbi.nlm.nih.gov](https://pubmed.ncbi.nlm.nih.gov/19914603/)) [S15]

However, the evidence is **not yet strong enough to say RGS4 is a definitive major human addiction gene**. Human data are limited, drug- and brain-region-specific effects are mixed, and large GWAS have not made RGS4 a top replicated locus. So the safest summary is: **RGS4 is credibly implicated in addiction biology, especially in preclinical opioid models, but its importance in human addiction susceptibility remains only partially established.** ([pmc.ncbi.nlm.nih.gov](https://pmc.ncbi.nlm.nih.gov/articles/PMC2902459/?utm_source=openai)) [S16]

## Cited sources

- **[S1]** Regulator of G protein-signaling proteins and addictive drugs - PubMed — https://pubmed.ncbi.nlm.nih.gov/20201861/?utm_source=openai
- **[S2]** Abused drugs modulate RGS4 mRNA levels in rat brain: comparison between acute drug treatment and a drug challenge after chronic treatment - PubMed — https://pubmed.ncbi.nlm.nih.gov/12270694/
- **[S3]** Brain region specific actions of regulator of G protein signaling 4 oppose morphine reward and dependence but promote analgesia - PubMed — https://pubmed.ncbi.nlm.nih.gov/19914603/
- **[S4]** Regulator of G-Protein Signaling 4 (RGS4) Controls Morphine Reward by Glutamate Receptor Activation in the Nucleus Accumbens of Mouse Brain - PubMed — https://pubmed.ncbi.nlm.nih.gov/29754475/?utm_source=openai
- **[S5]** Regulation of RGS proteins by chronic morphine in rat locus coeruleus - PubMed — https://pubmed.ncbi.nlm.nih.gov/12653973/
- **[S6]** Regulators of G-protein signaling 2 and 4 differentially regulate cocaine-induced rewarding effects - PubMed — https://pubmed.ncbi.nlm.nih.gov/30036561/
- **[S7]** Association of Polymorphisms in RGS4 and Expression of RGS transcripts in the Brains of Human Alcoholics - PMC — https://pmc.ncbi.nlm.nih.gov/articles/PMC2902459/?utm_source=openai
- **[S8]** Brain region specific actions of regulator of G protein signaling 4 oppose morphine reward and dependence but promote analgesia - PubMed — https://pubmed.ncbi.nlm.nih.gov/19914603/
- **[S9]** Brain region specific actions of regulator of G protein signaling 4 oppose morphine reward and dependence but promote analgesia - PubMed — https://pubmed.ncbi.nlm.nih.gov/19914603/
- **[S10]** Association of Polymorphisms in RGS4 and Expression of RGS transcripts in the Brains of Human Alcoholics - PMC — https://pmc.ncbi.nlm.nih.gov/articles/PMC2902459/?utm_source=openai
- **[S11]** Transancestral GWAS of alcohol dependence reveals common genetic underpinnings with psychiatric disorders - PubMed — https://pubmed.ncbi.nlm.nih.gov/30482948/
- **[S12]** Regulator of G-Protein Signaling (RGS) Protein Modulation of Opioid Receptor Signaling as a Potential Target for Pain Management - PubMed — https://pubmed.ncbi.nlm.nih.gov/32038168/
- **[S13]** RGS4 inhibits signaling by group I metabotropic glutamate receptors - PubMed — https://pubmed.ncbi.nlm.nih.gov/9437012/?utm_source=openai
- **[S14]** Regulator of G-Protein Signaling 4 (RGS4) Controls Morphine Reward by Glutamate Receptor Activation in the Nucleus Accumbens of Mouse Brain - PubMed — https://pubmed.ncbi.nlm.nih.gov/29754475/
- **[S15]** Brain region specific actions of regulator of G protein signaling 4 oppose morphine reward and dependence but promote analgesia - PubMed — https://pubmed.ncbi.nlm.nih.gov/19914603/
- **[S16]** Association of Polymorphisms in RGS4 and Expression of RGS transcripts in the Brains of Human Alcoholics - PMC — https://pmc.ncbi.nlm.nih.gov/articles/PMC2902459/?utm_source=openai
