## Supplementary material for "Understanding Substance Dependence: What Differentiates Addictive from Non-Addictive Drugs?": Folder for supplementary files: addictive-genes-claims-first100-038-kiaa1755-gene-protein-is-implicated-in-addiction-bef6e0c44f52.md

# addictive-genes-claims-first100-038-bef6e0c44f52

**Original claim:** KIAA1755 gene/protein is implicated in addiction.

## Claim

KIAA1755 gene/protein is implicated in addiction.

## Verdict

Verdict: Likely False

## Confidence

Medium.

## Supporting evidence

There is a **very limited, indirect signal** connecting KIAA1755 to a smoking-related phenotype. In NCBI Gene, KIAA1755 has a GeneRIF link to a 2010 smoking-cessation trial titled *Personalized smoking cessation: interactions between nicotine dose, dependence and quit-success genotype score*. That study reported that a **polygenic** “quit-success genotype score” interacted with nicotine dose and dependence level to predict quitting outcomes, and the score was built from **12,058 SNPs** showing nominal signals in prior datasets. In other words, the paper supports a broad polygenic smoking-cessation signal, not a specific causal role for KIAA1755 itself. NCBI’s inclusion of the paper under KIAA1755 shows the gene appeared in that context, but the study does not establish KIAA1755 as an addiction gene. ([ncbi.nlm.nih.gov](https://www.ncbi.nlm.nih.gov/gene/85449)) [S1]

KIAA1755 also has some **general neurobiological plausibility**. NCBI currently describes it as a protein-coding gene with predicted guanyl-nucleotide exchange factor activity and predicted involvement in axon guidance, with broad expression in brain and other tissues. Human Protein Atlas data further indicate low regional specificity across the brain, with expression patterns clustering toward white matter/signal-transduction and oligodendrocyte/myelination-associated cell types. Those features mean KIAA1755 is not biologically absurd as a brain-relevant gene, but they are still far from showing a role in addiction specifically. ([ncbi.nlm.nih.gov](https://www.ncbi.nlm.nih.gov/gene/85449)) [S2]

## Contradicting or limiting evidence

The strongest evidence argues **against** KIAA1755 being an established addiction-implicated gene. In curated human genetics resources, KIAA1755 is mainly linked to **cardiac rhythm traits**, not substance use disorders. NCBI’s phenotype section for KIAA1755 points to GWAS findings on heart rate loci, and multiple primary studies have associated KIAA1755 variants with heart rate and heart-rate variability (HRV). One exome-chip study identified a low-frequency loss-of-function variant at the KIAA1755 locus for heart rate, and another large HRV GWAS found KIAA1755-tagging variants among genome-wide significant HRV loci. ([ncbi.nlm.nih.gov](https://www.ncbi.nlm.nih.gov/gene/85449?utm_source=openai)) [S3]

Functional follow-up work also supports a **cardiac/autonomic**, not addiction-specific, interpretation. In zebrafish CRISPR work translating GWAS loci, KIAA1755 ortholog disruption tended to increase HRV without clearly affecting heart rate, and the authors interpreted the locus as potentially influencing heart-rhythm regulation. They explicitly called KIAA1755 a previously uncharacterized gene and said that future mechanistic studies are needed to determine how it affects heart rhythm. That is evidence of incomplete functional annotation, but the phenotype being pursued was cardiovascular/autonomic control rather than reward, dependence, craving, withdrawal, or compulsive drug use. ([pmc.ncbi.nlm.nih.gov](https://pmc.ncbi.nlm.nih.gov/articles/PMC7367351/)) [S4]

More importantly, I did **not** find KIAA1755 in the full text of several major modern addiction-GWAS papers. In the 2023 multivariate GWAS/meta-analysis of over 1 million subjects across problematic alcohol use, problematic tobacco use, cannabis use disorder, and opioid use disorder, gene-based signals included genes such as **FTO, DRD2, and PDE4B**, and KIAA1755 was not mentioned in the PMC full text. Likewise, in a 2024 multi-ancestry GWAS of alcohol use disorder involving over 1 million individuals, KIAA1755 was not mentioned in the PMC full text. In a 2025 large GWAS of DSM-defined nicotine dependence, KIAA1755 was again not mentioned in the PMC full text. ([pmc.ncbi.nlm.nih.gov](https://pmc.ncbi.nlm.nih.gov/articles/PMC10217792/)) [S5]

The same pattern extends to broader tobacco genetics. A recent large multi-ancestry study of tobacco use disorder identified hundreds of potential risk genes, but I did not find KIAA1755 mentioned in the available full text. That absence does not prove the gene has no effect, but it does argue strongly against calling it a recognized addiction-implicated gene on current evidence. ([nature.com](https://www.nature.com/articles/s41562-024-01851-6)) [S6]

Finally, the overall direction of current addiction genetics does not center on KIAA1755. NIH’s summary of the large cross-substance study emphasized that shared addiction-risk signals mapped most strongly to regions involved in **dopamine signaling regulation**, and the paper’s reported leading gene-based results were not KIAA1755. ([nih.gov](https://www.nih.gov/news-events/news-releases/nih-study-reveals-shared-genetic-markers-underlying-substance-use-disorders)) [S7]

## Mechanistic / biological rationale

From a mechanistic standpoint, the case for KIAA1755 in addiction is **weak and speculative**. Addiction-related genes are usually implicated through one or more of the following: replicated human associations with substance use disorder phenotypes, expression or functional effects in reward/salience circuits, involvement in known transmitter systems such as dopamine, glutamate, GABA, nicotinic acetylcholine, opioid, or endocannabinoid signaling, or convergent evidence from animal and cellular models. The major contemporary cross-SUD genetics literature emphasizes dopamine-regulatory biology and other well-characterized neurobehavioral pathways; KIAA1755 is not among the highlighted genes in those analyses. ([nih.gov](https://www.nih.gov/news-events/news-releases/nih-study-reveals-shared-genetic-markers-underlying-substance-use-disorders)) [S8]

KIAA1755’s predicted roles—guanyl-nucleotide exchange factor activity and axon guidance—could, in principle, matter for neural circuit assembly or signaling. Its expression in brain, white matter-associated clusters, and oligodendrocyte-related cell types could also fit a broad “nervous system homeostasis” narrative. But this is still only **generic neural plausibility**, not addiction-specific evidence. There is no strong human evidence here tying KIAA1755 to craving, reinforcement learning, withdrawal, relapse, treatment response, or substance-specific liability. ([ncbi.nlm.nih.gov](https://www.ncbi.nlm.nih.gov/gene/85449)) [S9]

In fact, the better-developed mechanistic literature around KIAA1755 currently points elsewhere: autonomic/cardiac physiology and possibly broader cellular signaling, not reward circuitry. The HRV and heart-rate associations, plus zebrafish follow-up suggesting effects on HRV, are much more concrete than anything in addiction biology. ([nature.com](https://www.nature.com/articles/ncomms15805)) [S10]

## Bottom line

At present, **KIAA1755 is not a well-supported addiction gene/protein**. There is a thin, indirect smoking-cessation link through an old **polygenic** quit-success score, and the gene is brain-expressed with predicted neural-related functions. But the best available human evidence links KIAA1755 mainly to **heart rate/heart-rate variability**, and I did not find it implicated in the full text of several large, modern GWAS studies of alcohol use disorder, nicotine dependence, tobacco use disorder, or cross-substance addiction liability. On current evidence, the claim is best judged **Likely False** rather than established or even clearly mixed. ([pmc.ncbi.nlm.nih.gov](https://pmc.ncbi.nlm.nih.gov/articles/PMC2896464/)) [S11]

## Cited sources

- **[S1]** KIAA1755 KIAA1755 [Homo sapiens (human)] - Gene - NCBI — https://www.ncbi.nlm.nih.gov/gene/85449
- **[S2]** KIAA1755 KIAA1755 [Homo sapiens (human)] - Gene - NCBI — https://www.ncbi.nlm.nih.gov/gene/85449
- **[S3]** KIAA1755 KIAA1755 [Homo sapiens (human)] - Gene - NCBI — https://www.ncbi.nlm.nih.gov/gene/85449?utm_source=openai
- **[S4]** 
            Translating GWAS-identified loci for cardiac rhythm and rate using an in vivo image- and CRISPR/Cas9-based approach - PMC
         — https://pmc.ncbi.nlm.nih.gov/articles/PMC7367351/
- **[S5]** 
            Multivariate genome-wide association meta-analysis of over 1 million subjects identifies loci underlying multiple substance use disorders - PMC
         — https://pmc.ncbi.nlm.nih.gov/articles/PMC10217792/
- **[S6]** Multi-ancestry meta-analysis of tobacco use disorder identifies 461 potential risk genes and reveals associations with multiple health outcomes | Nature Human Behaviour — https://www.nature.com/articles/s41562-024-01851-6
- **[S7]** NIH study reveals shared genetic markers underlying substance use disorders | National Institutes of Health (NIH) — https://www.nih.gov/news-events/news-releases/nih-study-reveals-shared-genetic-markers-underlying-substance-use-disorders
- **[S8]** NIH study reveals shared genetic markers underlying substance use disorders | National Institutes of Health (NIH) — https://www.nih.gov/news-events/news-releases/nih-study-reveals-shared-genetic-markers-underlying-substance-use-disorders
- **[S9]** KIAA1755 KIAA1755 [Homo sapiens (human)] - Gene - NCBI — https://www.ncbi.nlm.nih.gov/gene/85449
- **[S10]** Genetic loci associated with heart rate variability and their effects on cardiac disease risk | Nature Communications — https://www.nature.com/articles/ncomms15805
- **[S11]** 
            Personalized Smoking Cessation: Interactions between Nicotine Dose, Dependence and Quit-Success Genotype Score - PMC
         — https://pmc.ncbi.nlm.nih.gov/articles/PMC2896464/
