## Supplementary material for "Understanding Substance Dependence: What Differentiates Addictive from Non-Addictive Drugs?": Folder for supplementary files: addictive-genes-claims-first100-039-maob-gene-protein-is-implicated-in-addiction-9f0c98bd6bac.md

# addictive-genes-claims-first100-039-9f0c98bd6bac

**Original claim:** MAOB gene/protein is implicated in addiction.

## Claim

**MAOB gene/protein is implicated in addiction.**

## Verdict

Verdict: Likely True

## Confidence

**Medium.**

## Supporting evidence

MAOB encodes monoamine oxidase B, a mitochondrial outer-membrane enzyme involved in oxidative deamination of monoamines; it preferentially degrades phenethylamine and also participates in metabolism of other neuroactive amines, placing it in pathways relevant to reward signaling. Because dopamine-related reinforcement is central to addiction biology, MAO-B is biologically plausible as a contributor to addictive phenotypes. ([ncbi.nlm.nih.gov](https://www.ncbi.nlm.nih.gov/gtr/genes/4129)) [S1]

The strongest evidence concerns **tobacco/nicotine addiction at the protein/activity level**. In a classic PET study of living humans, smokers had about a **40% reduction in brain MAO-B** relative to nonsmokers or former smokers, and the authors proposed that reduced MAO-B could synergize with nicotine to shape smoking behavior. Subsequent reviews concluded there is a **clear association** between smoking and reduced MAO-A/MAO-B binding in smokers’ brains, with strong evidence that tobacco smoke contains one or more MAO-inhibiting substances. ([pubmed.ncbi.nlm.nih.gov](https://pubmed.ncbi.nlm.nih.gov/8602220/)) [S2]

Authoritative public-health summaries are aligned with that interpretation. NIDA states that smoking is linked to a marked decrease in monoamine oxidase levels and reduced MAO binding sites in the brain, that this is likely due to a tobacco-smoke constituent other than nicotine, and that animal work suggests MAO inhibition can make nicotine more reinforcing and may contribute to tobacco addiction. ([nida.nih.gov](https://nida.nih.gov/publications/research-reports/tobacco-nicotine-e-cigarettes/nicotine-addictive?utm_source=openai)) [S3]

Preclinical evidence also supports involvement in nicotine reinforcement. A 2022 review summarizing rodent work reported that MAO activity is reduced in smokers and that MAO inhibition can increase nicotine self-administration, especially at low nicotine doses, potentially lowering the threshold at which nicotine becomes reinforcing. Earlier reviews similarly concluded that MAO inhibition potentiates nicotine’s reinforcing effects in rodents. ([pubmed.ncbi.nlm.nih.gov](https://pubmed.ncbi.nlm.nih.gov/36051642/)) [S4]

There is also **some human genetic/epigenetic evidence**, though it is much weaker than the tobacco imaging/activity evidence. In a population study of 1,230 Russian adults, a MAOB intron 13 variant was associated with smoking status in women, and MAOB methylation patterns were associated with smoking history. A smaller 2021 study in 302 polysubstance users found a MAOB VNTR association with cocaine and opiate addiction. These findings support the idea that MAOB may be implicated in susceptibility or phenotype variation, but they are not yet definitive. ([pubmed.ncbi.nlm.nih.gov](https://pubmed.ncbi.nlm.nih.gov/28858992/)) [S5]

More broadly, a review on MAO in personality and drug use concluded that MAO-A and MAO-B have been frequent targets in studies of personality and substance-related behaviors, and that associations have been reported across gene variants, peripheral enzyme activity, brain imaging, and methylation measures. ([sciencedirect.com](https://www.sciencedirect.com/science/article/abs/pii/S0278584616300203)) [S6]

## Contradicting or limiting evidence

The main limitation is that the evidence is **not equally strong for “MAOB gene” and “MAO-B protein,” nor for all addictions**. For tobacco dependence, reviews emphasize that although MAO inhibition is clearly associated with smoking, there is **little evidence that low MAO is a pre-existing predictive factor** for becoming a smoker, and no convincing human proof that MAO-inhibiting smoke constituents directly drive dependence severity. ([pubmed.ncbi.nlm.nih.gov](https://pubmed.ncbi.nlm.nih.gov/26508396/)) [S7]

Human causality also remains unsettled because nicotine itself does not seem to be the direct MAO-B inhibitor. In vivo baboon PET work found that an acute nicotine dose did **not** inhibit brain MAO-B, implying that other smoke constituents or longer-term adaptations must explain the smoker/non-smoker difference. Reviews likewise note that no single absorbed tobacco constituent has been definitively shown to account for MAO inhibition in human smokers. ([pubmed.ncbi.nlm.nih.gov](https://pubmed.ncbi.nlm.nih.gov/9674950/?utm_source=openai)) [S8]

Clinical intervention data are also sobering. If selective MAO-B inhibition were a strong standalone driver of tobacco addiction, MAO-B inhibitors might be expected to help smoking cessation; however, a Cochrane review found **no significant benefit** of MAO inhibitors overall for quitting smoking, and an RCT of the selective MAO-B inhibitor EVT302 found it was **not superior to placebo**, either alone or with nicotine patch. That does not rule out MAO-B involvement, but it argues against a simple “MAO-B inhibition alone explains addiction” model. ([pubmed.ncbi.nlm.nih.gov](https://pubmed.ncbi.nlm.nih.gov/24402784/)) [S9]

For **alcohol dependence**, older platelet-MAO literature is especially vulnerable to confounding by smoking. In a twin-registry study of 1,551 subjects, current smoking strongly lowered platelet MAO activity, but alcohol use and lifetime alcohol dependence were **not associated** with MAO activity after smoking was accounted for; the authors concluded platelet MAO activity could not be sustained as a direct genetic marker of vulnerability to alcohol dependence. ([pubmed.ncbi.nlm.nih.gov](https://pubmed.ncbi.nlm.nih.gov/10824664/)) [S10]

Gene-level evidence across addictions is also inconsistent. The MAO review notes that what is known about MAOA/MAOB variants does **not explain enzyme activities well in vivo**, and that single-gene associations in this area have generally been hard to unify into a mechanistic model. The positive cocaine/opiate genetic study was relatively small and compared addiction subgroups within polyconsumers rather than establishing a robust genome-wide signal in the general population. ([sciencedirect.com](https://www.sciencedirect.com/science/article/abs/pii/S0278584616300203)) [S11]

## Mechanistic / biological rationale

Mechanistically, MAO-B is relevant because addiction circuits depend heavily on monoamine signaling. MAO-B contributes to catabolism of dopamine-related amines; reduced MAO-B activity would be expected to increase availability of dopamine or related trace amines in reward-related circuits, potentially amplifying reinforcement from nicotine or other drugs. That mechanistic logic underlies both the PET findings in smokers and the rodent nicotine self-administration data. ([ncbi.nlm.nih.gov](https://www.ncbi.nlm.nih.gov/gtr/genes/4129)) [S12]

There is also indirect support from endophenotype studies. MAO-B knockout mice show behavioral disinhibition, increased novelty seeking, and risk-taking-like behavior—traits often discussed as addiction-relevant vulnerability phenotypes. But the same paper explicitly cautions that rodent-to-human translation is imperfect, including species differences in MAO isoform expression and dopamine metabolism, so this is supportive rather than decisive evidence. ([pmc.ncbi.nlm.nih.gov](https://pmc.ncbi.nlm.nih.gov/articles/PMC2783894/)) [S13]

Overall, the most coherent model is: **tobacco smoke reduces MAO activity (including MAO-B), this can enhance nicotine reinforcement in animals, and in humans it plausibly contributes to the addictive properties of cigarettes**. That is a meaningful form of implication, even though it falls short of proving MAOB is a major causal gene for addiction broadly defined. ([pubmed.ncbi.nlm.nih.gov](https://pubmed.ncbi.nlm.nih.gov/36051642/)) [S14]

## Bottom line

The claim is **reasonable but should be stated narrowly**. **MAO-B/MAOB is most clearly implicated in tobacco addiction**, especially at the **protein/activity** level: smokers show reduced brain MAO-B, tobacco smoke inhibits MAO, and animal studies show MAO inhibition can enhance nicotine reinforcement. However, **gene-level evidence is limited and inconsistent**, and evidence for **non-tobacco addictions** such as alcohol, cocaine, or opioids is much thinner and less replicated. So: **yes, MAOB is implicated in addiction, but mainly in tobacco dependence and mainly as a modulatory factor rather than a conclusively established universal addiction gene.** ([pubmed.ncbi.nlm.nih.gov](https://pubmed.ncbi.nlm.nih.gov/8602220/)) [S15]

## Cited sources

- **[S1]** MAOB monoamine oxidase B - NIH Genetic Testing Registry (GTR) - NCBI — https://www.ncbi.nlm.nih.gov/gtr/genes/4129
- **[S2]** Inhibition of monoamine oxidase B in the brains of smokers - PubMed — https://pubmed.ncbi.nlm.nih.gov/8602220/
- **[S3]** Is nicotine addictive? | National Institute on Drug Abuse (NIDA) — https://nida.nih.gov/publications/research-reports/tobacco-nicotine-e-cigarettes/nicotine-addictive?utm_source=openai
- **[S4]** Monoamine oxidase inhibition in cigarette smokers: From preclinical studies to tobacco product regulation - PubMed — https://pubmed.ncbi.nlm.nih.gov/36051642/
- **[S5]** The genotypes and methylation of MAO genes as factors behind smoking behavior - PubMed — https://pubmed.ncbi.nlm.nih.gov/28858992/
- **[S6]** The role of MAO in personality and drug use - ScienceDirect — https://www.sciencedirect.com/science/article/abs/pii/S0278584616300203
- **[S7]** Contribution of Monoamine Oxidase Inhibition to Tobacco Dependence: A Review of the Evidence - PubMed — https://pubmed.ncbi.nlm.nih.gov/26508396/
- **[S8]** An acute dose of nicotine does not inhibit MAO B in baboon brain in vivo - PubMed — https://pubmed.ncbi.nlm.nih.gov/9674950/?utm_source=openai
- **[S9]** Antidepressants for smoking cessation - PubMed — https://pubmed.ncbi.nlm.nih.gov/24402784/
- **[S10]** Monoamine oxidase: associations with alcohol dependence, smoking and other measures of psychopathology - PubMed — https://pubmed.ncbi.nlm.nih.gov/10824664/
- **[S11]** The role of MAO in personality and drug use - ScienceDirect — https://www.sciencedirect.com/science/article/abs/pii/S0278584616300203
- **[S12]** MAOB monoamine oxidase B - NIH Genetic Testing Registry (GTR) - NCBI — https://www.ncbi.nlm.nih.gov/gtr/genes/4129
- **[S13]** 
            Behavioral disinhibition and reduced anxiety-like behaviors in monoamine oxidase B deficient mice - PMC
         — https://pmc.ncbi.nlm.nih.gov/articles/PMC2783894/
- **[S14]** Monoamine oxidase inhibition in cigarette smokers: From preclinical studies to tobacco product regulation - PubMed — https://pubmed.ncbi.nlm.nih.gov/36051642/
- **[S15]** Inhibition of monoamine oxidase B in the brains of smokers - PubMed — https://pubmed.ncbi.nlm.nih.gov/8602220/
