## Supplementary material for "Understanding Substance Dependence: What Differentiates Addictive from Non-Addictive Drugs?": Folder for supplementary files: addictive-genes-claims-first100-040-elovl5-gene-protein-is-implicated-in-addiction-c9c9a7c5fed6.md

# addictive-genes-claims-first100-040-c9c9a7c5fed6

**Original claim:** ELOVL5 gene/protein is implicated in addiction.

## Claim

ELOVL5 gene/protein is implicated in addiction.

## Verdict

Verdict: Mixed Evidence

## Confidence

Medium

## Supporting evidence

ELOVL5 is a biologically credible brain gene rather than a purely peripheral lipid-metabolism gene. In mice, Elovl5 is expressed in a region- and cell-type-specific way across the CNS, with notable expression in cerebellum, brainstem, olfactory regions, hippocampal CA2/CA3 pyramidal cells, oligodendroglia, and microglia. ELOVL5 encodes a fatty-acid elongase important for omega-3 and omega-6 long-chain PUFA synthesis, so it plausibly affects neuronal membrane composition, glial biology, and signaling relevant to reward circuitry and plasticity. ([pubmed.ncbi.nlm.nih.gov](https://pubmed.ncbi.nlm.nih.gov/33994961/)) [S1]

There are also a few addiction-related omics hints involving ELOVL5, especially for alcohol. In a human postmortem frontal-cortex study of alcoholics, ELOVL5 appeared among predicted targets within enriched lipid/fatty-acid biosynthetic pathways linked to upregulated microRNAs. That does not prove causality, but it does place ELOVL5 inside an alcohol-associated human brain regulatory signature. ([pmc.ncbi.nlm.nih.gov](https://pmc.ncbi.nlm.nih.gov/articles/PMC3170679/)) [S2]

A separate exploratory alcohol use disorder (AUD) DNA-methylation study also listed ELOVL5 among genes in one of the top functional networks derived from associated CpG findings. Again, this is hypothesis-generating rather than definitive, but it is another human AUD-related signal touching ELOVL5. ([pmc.ncbi.nlm.nih.gov](https://pmc.ncbi.nlm.nih.gov/articles/PMC5108727/?utm_source=openai)) [S3]

More broadly, PUFA biology has been linked to addiction vulnerability and drug-related behaviors. A review of omega-3 fatty acids and addiction concluded that preclinical literature generally supports effects of n-3 fatty-acid status on drug sensitivity and addiction-relevant behavior, and a mouse study reported that dietary omega-3 supplementation reduced opioid-seeking behaviors. Because ELOVL5 is one of the enzymes that helps generate long-chain PUFAs, these findings provide an indirect mechanistic bridge between ELOVL5 biology and addiction phenotypes. ([pubmed.ncbi.nlm.nih.gov](https://pubmed.ncbi.nlm.nih.gov/32348211/?utm_source=openai)) [S4]

## Contradicting or limiting evidence

The main limitation is that I did not find strong, replicated, direct evidence that ELOVL5 itself is a bona fide addiction gene in the way that CHRNA5, OPRM1, ADH1B, or related loci are. Large modern addiction-genetics papers and reviews emphasize many replicated loci for alcohol, nicotine, opioid, and stimulant phenotypes, but ELOVL5 is not highlighted among the established signals in the sources reviewed. For example, a 2023 review reanalyzing large-scale GWAS-derived substance-associated genes discusses broad addiction-circuit transcriptomic enrichment without identifying ELOVL5 as a prominent replicated addiction gene, while large nicotine-dependence GWAS continue to foreground CHRNA5-region findings. ([pubmed.ncbi.nlm.nih.gov](https://pubmed.ncbi.nlm.nih.gov/37753566/)) [S5]

Likewise, major alcohol-dependence/AUD genetics papers cited in the reviewed material identified other loci or pathways as the strongest signals, not ELOVL5. Examples include chromosome 11 loci in older alcohol-dependence GWAS and newer cross-substance/common-risk loci such as ANKS1B, AGBL4, and CTNNA2 in alcoholism/heroin/methamphetamine analyses. ([pubmed.ncbi.nlm.nih.gov](https://pubmed.ncbi.nlm.nih.gov/20201924/?utm_source=openai)) [S6]

The positive ELOVL5 findings are also weak in evidentiary hierarchy. The alcohol microRNA study implicates ELOVL5 only as part of an enriched target/pathway set, not as a directly measured causal driver. The AUD methylation study was explicitly exploratory. I did not find a human genetic association replicated across cohorts, a transcriptome-wide association result, or an animal addiction experiment where ELOVL5 manipulation itself altered drug self-administration, conditioned place preference, relapse, or withdrawal severity. ([pmc.ncbi.nlm.nih.gov](https://pmc.ncbi.nlm.nih.gov/articles/PMC3170679/)) [S7]

Finally, some ELOVL-family evidence in addiction points to other family members rather than ELOVL5. For instance, one study on RNA splicing and AUD implicated ELOVL7, not ELOVL5. That does not count against ELOVL5 directly, but it underscores that evidence for one elongase-family gene should not be transferred automatically to another. ([pubmed.ncbi.nlm.nih.gov](https://pubmed.ncbi.nlm.nih.gov/37217680/?utm_source=openai)) [S8]

## Mechanistic / biological rationale

A plausible biological rationale exists. ELOVL5 helps synthesize long-chain polyunsaturated fatty acids, which are important components of neuronal membranes and influence receptor function, synaptic vesicle dynamics, inflammatory tone, and glial signaling. In the CNS, its expression in hippocampal neurons, oligodendroglia, and microglia suggests roles in neural homeostasis rather than only liver lipid metabolism. ([pubmed.ncbi.nlm.nih.gov](https://pubmed.ncbi.nlm.nih.gov/33994961/)) [S9]

That matters for addiction because addictive drugs produce long-lasting neuroplastic and neuroimmune changes in cortico-striatal-limbic circuits, and PUFA-related pathways have been proposed to modulate those adaptations. Reviews of omega-3/addiction biology argue that n-3 PUFA status can affect drug sensitivity, neuroinflammation, stress responsivity, and relapse-related behaviors; the opioid-seeking mouse study is one concrete example of that broader framework. Thus, if ELOVL5 meaningfully alters local PUFA availability in relevant circuits or glia, it could in principle modulate addiction vulnerability or severity. ([pubmed.ncbi.nlm.nih.gov](https://pubmed.ncbi.nlm.nih.gov/32348211/?utm_source=openai)) [S10]

But this remains an inference, not a demonstrated ELOVL5-specific addiction mechanism. Existing direct evidence for ELOVL5 is much stronger for neural lipid homeostasis and certain neurologic disorders than for substance use disorders. Mutations in ELOVL5 cause spinocerebellar ataxia type 38, and disease-associated variants can mislocalize the protein away from the endoplasmic reticulum, showing that disrupted ELOVL5 function can have meaningful neural consequences. That supports neurobiological relevance, but it is not addiction evidence per se. ([pubmed.ncbi.nlm.nih.gov](https://pubmed.ncbi.nlm.nih.gov/25065913/)) [S11]

## Bottom line

ELOVL5 is **not currently a well-established addiction gene/protein** based on the evidence I found. There are **some indirect and exploratory signals** connecting it to addiction-related biology—especially alcohol-related human omics findings and a plausible PUFA-based mechanism—but there is **little direct, replicated evidence** that ELOVL5 itself drives addiction risk or addiction behaviors. Overall, the best characterization is **suggestive but weak/indirect evidence**, not strong confirmation. ([pmc.ncbi.nlm.nih.gov](https://pmc.ncbi.nlm.nih.gov/articles/PMC3170679/)) [S12]

## Cited sources

- **[S1]** Elovl5 Expression in the Central Nervous System of the Adult Mouse - PubMed — https://pubmed.ncbi.nlm.nih.gov/33994961/
- **[S2]** 
            Up-regulation of MicroRNAs in Brain of Human Alcoholics - PMC
         — https://pmc.ncbi.nlm.nih.gov/articles/PMC3170679/
- **[S3]** An Exploratory Association Study of Alcohol Use Disorder and DNA Methylation - PMC — https://pmc.ncbi.nlm.nih.gov/articles/PMC5108727/?utm_source=openai
- **[S4]** Omega-3 Fatty Acids and Vulnerability to Addiction: Reviewing Preclinical and Clinical Evidence - PubMed — https://pubmed.ncbi.nlm.nih.gov/32348211/?utm_source=openai
- **[S5]** Analysis of the brain transcriptome for substance-associated genes: An update on large-scale genome-wide association studies - PubMed — https://pubmed.ncbi.nlm.nih.gov/37753566/
- **[S6]** Genome-wide association study of alcohol dependence implicates a region on chromosome 11 - PubMed — https://pubmed.ncbi.nlm.nih.gov/20201924/?utm_source=openai
- **[S7]** 
            Up-regulation of MicroRNAs in Brain of Human Alcoholics - PMC
         — https://pmc.ncbi.nlm.nih.gov/articles/PMC3170679/
- **[S8]** RNA alternative splicing impacts the risk for alcohol use disorder - PubMed — https://pubmed.ncbi.nlm.nih.gov/37217680/?utm_source=openai
- **[S9]** Elovl5 Expression in the Central Nervous System of the Adult Mouse - PubMed — https://pubmed.ncbi.nlm.nih.gov/33994961/
- **[S10]** Omega-3 Fatty Acids and Vulnerability to Addiction: Reviewing Preclinical and Clinical Evidence - PubMed — https://pubmed.ncbi.nlm.nih.gov/32348211/?utm_source=openai
- **[S11]** ELOVL5 mutations cause spinocerebellar ataxia 38 - PubMed — https://pubmed.ncbi.nlm.nih.gov/25065913/
- **[S12]** 
            Up-regulation of MicroRNAs in Brain of Human Alcoholics - PMC
         — https://pmc.ncbi.nlm.nih.gov/articles/PMC3170679/
