## Supplementary material for "Understanding Substance Dependence: What Differentiates Addictive from Non-Addictive Drugs?": Folder for supplementary files: addictive-genes-claims-first100-041-slc12a5-gene-protein-is-implicated-in-addiction-952b153216a4.md

# addictive-genes-claims-first100-041-952b153216a4

**Original claim:** SLC12A5 gene/protein is implicated in addiction.

## Claim

SLC12A5, which encodes the neuronal K\(^+\)-Cl\(^-\) cotransporter KCC2, is implicated in addiction. KCC2 is a major neuronal chloride extruder needed for mature inhibitory GABA/glycine signaling, so the claim is really about whether altered SLC12A5/KCC2 function contributes to addiction-relevant neurobiology or behavior. ([pubmed.ncbi.nlm.nih.gov](https://pubmed.ncbi.nlm.nih.gov/30576625/)) [S1]

## Verdict

Verdict: Likely True

## Confidence

Medium

## Supporting evidence

The strongest evidence is **preclinical**, especially in rodent ventral tegmental area (VTA) circuits central to reward, withdrawal, and drug seeking. In opioid models, chronic morphine withdrawal activates VTA microglia, increases BDNF, and reduces KCC2 expression/function in VTA GABA neurons. In that study, blocking microglial activation or BDNF signaling prevented loss of chloride extrusion and restored cocaine conditioned place preference in opioid-dependent animals; conversely, intra-VTA BDNF or a KCC2 inhibitor in opioid-naive animals reproduced a blunted reward phenotype. That is a fairly direct demonstration that KCC2 dysfunction can causally alter addiction-relevant reward processing. ([pubmed.ncbi.nlm.nih.gov](https://pubmed.ncbi.nlm.nih.gov/26202104/)) [S2]

Older work on opiate motivation points in the same direction. In rats, intra-VTA furosemide, used there as a chloride cotransporter inhibitor, shifted morphine motivation into the dopamine-dependent state usually seen after dependence/withdrawal, and electrophysiology showed reduced muscimol inhibition or even excitation of some VTA GABA neurons under furosemide. Those results support the idea that perturbing KCC2-related chloride handling in the VTA can change the motivational circuitry engaged by opioids. ([pubmed.ncbi.nlm.nih.gov](https://pubmed.ncbi.nlm.nih.gov/23392354/)) [S3]

Evidence also extends beyond opioids. A 2025 study reported that **both cocaine and morphine** caused a depolarizing shift in \(E_{GABA}\) in VTA GABA neurons, and that this effect depended on dopamine D1/D5 receptor signaling. The authors concluded that acute cocaine impairs chloride extrusion through KCC2 dephosphorylation, while morphine produces convergent disruption of chloride homeostasis; morphine self-administration also produced a lasting disturbance in VTA GABA chloride homeostasis. This places KCC2 dysfunction within a shared drug-induced plasticity mechanism rather than an opioid-only phenomenon. ([pmc.ncbi.nlm.nih.gov](https://pmc.ncbi.nlm.nih.gov/articles/PMC12678199/)) [S4]

For alcohol-related behaviors, the evidence is also fairly direct. In male rats exposed to intermittent ethanol, withdrawal reduced phosphorylated KCC2 in the VTA, and a KCC2 activator (CLP290) reduced excessive alcohol consumption, whereas a KCC2 inhibitor increased intake. The same paper showed that the alcohol-limiting effects of BDNF/TrkB activation were reversed by KCC2 inhibition, arguing that KCC2 is not merely correlated with alcohol behavior but functionally involved in it. ([pubmed.ncbi.nlm.nih.gov](https://pubmed.ncbi.nlm.nih.gov/36401060/)) [S5]

Stress studies are relevant because stress is a major addiction vulnerability factor and relapse driver. In a Neuron paper, acute stress reduced VTA KCC2 function/phosphorylation, shifted GABA signaling toward excitation, and increased ethanol self-administration; intra-VTA CLP290 normalized chloride homeostasis and reduced that stress-driven drinking. A later mechanistic study showed that 5-HT\(_{2A}\) receptor activation could restore KCC2 S940 phosphorylation and normalize stress-induced VTA chloride dysregulation. Together, these studies support KCC2 as a modulator of addiction-relevant stress-drug interactions. ([pubmed.ncbi.nlm.nih.gov](https://pubmed.ncbi.nlm.nih.gov/27720487/)) [S6]

Nicotine exposure has likewise been linked to KCC2-mediated vulnerability to other drug use. After acute nicotine, rats showed reduced VTA KCC2 expression, a depolarizing shift in GABA signaling, and increased diazepam intake; CLP290 normalized diazepam-related VTA signaling and reduced diazepam consumption back toward control levels. That is another causal manipulation consistent with KCC2 involvement in polysubstance-use vulnerability. ([pmc.ncbi.nlm.nih.gov](https://pmc.ncbi.nlm.nih.gov/articles/PMC7082131/)) [S7]

Recent synthesis papers now explicitly frame midbrain KCC2 downregulation as relevant to substance-use behavior. A 2024 review summarized evidence that stress and drugs including nicotine, alcohol, and morphine downregulate KCC2 in VTA GABA neurons and that pharmacologic KCC2 activation reduces alcohol or diazepam consumption in preclinical models. ([pmc.ncbi.nlm.nih.gov](https://pmc.ncbi.nlm.nih.gov/articles/PMC11392611/)) [S8]

## Contradicting or limiting evidence

The main limitation is that the case is **not yet strong in humans**. The literature I found is dominated by rodent electrophysiology, place preference, and self-administration experiments, with little direct evidence that inherited human SLC12A5 variation meaningfully alters addiction risk. By contrast, authoritative human genetics reviews of SLC12A5 focus on epilepsy and neurodevelopmental disease, where pathogenic variants are clearly established. ([pubmed.ncbi.nlm.nih.gov](https://pubmed.ncbi.nlm.nih.gov/30576625/)) [S9]

There is also an important interpretive complication: KCC2 downregulation does **not always map onto a simple “more reward” phenotype**. Reviews of the field note that reduced KCC2 can produce blunted cocaine or morphine place preference and decreased reward sensitivity in some paradigms, yet increase alcohol or diazepam intake, enhance early drug-taking, or alter learning in others. So KCC2 seems better viewed as a regulator of reward/allostasis/learning circuitry than as a straightforward “pro-addiction” switch. ([pmc.ncbi.nlm.nih.gov](https://pmc.ncbi.nlm.nih.gov/articles/PMC11392611/)) [S10]

Some of the newer work also remains incomplete mechanistically. For example, the 2025 cocaine/morphine study notes that the precise site of D1/D5 action is unresolved and that self-administration experiments were conducted only in male rats. More broadly, it is still uncertain how much of the behavioral effect arises from VTA interneurons versus long-range VTA GABA projections, and whether the same KCC2 mechanism generalizes across sexes, substances, and stages of addiction. ([pmc.ncbi.nlm.nih.gov](https://pmc.ncbi.nlm.nih.gov/articles/PMC12678199/)) [S11]

## Mechanistic / biological rationale

The biological rationale is strong. SLC12A5 encodes KCC2, the main neuronal chloride extruder, which is essential for keeping intracellular chloride low enough for GABA\(_A\) signaling to remain inhibitory. When KCC2 function falls, \(E_{GABA}\) depolarizes, GABAergic inhibition weakens, and in some circumstances GABA responses can become paradoxically excitatory. ([pubmed.ncbi.nlm.nih.gov](https://pubmed.ncbi.nlm.nih.gov/30576625/)) [S12]

In addiction-relevant midbrain circuitry, KCC2 is especially important in **VTA GABA neurons**. Drug exposure, withdrawal, or stress can reduce KCC2 expression or phosphorylation, impair chloride extrusion, and alter how VTA GABA neurons respond to GABAergic input. That can reshape inhibition onto dopamine neurons and modify tonic versus phasic dopamine signaling, cue learning, reward sensitivity, and drug intake. Recent work further suggests that KCC2-dependent chloride dynamics help regulate synchronization of VTA GABA networks during reward learning and may be “hijacked” in addiction. ([pmc.ncbi.nlm.nih.gov](https://pmc.ncbi.nlm.nih.gov/articles/PMC11392611/)) [S13]

Several upstream regulators fit known addiction biology. BDNF can reduce KCC2 and switch VTA inhibitory physiology in opioid-related states; stress-related signaling alters KCC2 phosphorylation at S940; and cocaine/morphine can engage dopamine D1/D5-dependent mechanisms that converge on KCC2 dysfunction. This makes KCC2 a plausible integrator of stress, withdrawal, and drug-evoked plasticity rather than an isolated finding. ([pubmed.ncbi.nlm.nih.gov](https://pubmed.ncbi.nlm.nih.gov/26202104/)) [S14]

## Bottom line

Yes—**SLC12A5/KCC2 is plausibly implicated in addiction**, but mainly at the level of **preclinical circuit and behavioral neuroscience**, not yet as an established human addiction gene. The best-supported interpretation is that KCC2 is an addiction-relevant regulator of mesolimbic inhibitory plasticity, especially in VTA GABA neurons, influencing withdrawal-related reward deficits, alcohol and diazepam intake, and drug-induced reward learning. Because direct human genetic and clinical evidence is still limited, “Likely True” is more appropriate than “Highly Supported.” ([pubmed.ncbi.nlm.nih.gov](https://pubmed.ncbi.nlm.nih.gov/26202104/)) [S15]

## Cited sources

- **[S1]** Pathogenic potential of human SLC12A5 variants causing KCC2 dysfunction - PubMed — https://pubmed.ncbi.nlm.nih.gov/30576625/
- **[S2]** Neuroimmune Regulation of GABAergic Neurons Within the Ventral Tegmental Area During Withdrawal from Chronic Morphine - PubMed — https://pubmed.ncbi.nlm.nih.gov/26202104/
- **[S3]** Ventral tegmental area GABA neurons and opiate motivation - PubMed — https://pubmed.ncbi.nlm.nih.gov/23392354/
- **[S4]** 
            Cocaine and Morphine Converge to Disrupt Chloride Homeostasis in Ventral Tegmental Area GABA Neurons - PMC
         — https://pmc.ncbi.nlm.nih.gov/articles/PMC12678199/
- **[S5]** Role of KCC2 in the Regulation of Brain-Derived Neurotrophic Factor on Ethanol Consumption in Rats - PubMed — https://pubmed.ncbi.nlm.nih.gov/36401060/
- **[S6]** Stress Increases Ethanol Self-Administration via a Shift toward Excitatory GABA Signaling in the Ventral Tegmental Area - PubMed — https://pubmed.ncbi.nlm.nih.gov/27720487/
- **[S7]** 
            Acute Nicotine Exposure Alters Ventral Tegmental Area Inhibitory Transmission and Promotes Diazepam Consumption - PMC
         — https://pmc.ncbi.nlm.nih.gov/articles/PMC7082131/
- **[S8]** 
            Midbrain KCC2 Downregulation: Implications for stress-related and substance use behaviors - PMC
         — https://pmc.ncbi.nlm.nih.gov/articles/PMC11392611/
- **[S9]** Pathogenic potential of human SLC12A5 variants causing KCC2 dysfunction - PubMed — https://pubmed.ncbi.nlm.nih.gov/30576625/
- **[S10]** 
            Midbrain KCC2 Downregulation: Implications for stress-related and substance use behaviors - PMC
         — https://pmc.ncbi.nlm.nih.gov/articles/PMC11392611/
- **[S11]** 
            Cocaine and Morphine Converge to Disrupt Chloride Homeostasis in Ventral Tegmental Area GABA Neurons - PMC
         — https://pmc.ncbi.nlm.nih.gov/articles/PMC12678199/
- **[S12]** Pathogenic potential of human SLC12A5 variants causing KCC2 dysfunction - PubMed — https://pubmed.ncbi.nlm.nih.gov/30576625/
- **[S13]** 
            Midbrain KCC2 Downregulation: Implications for stress-related and substance use behaviors - PMC
         — https://pmc.ncbi.nlm.nih.gov/articles/PMC11392611/
- **[S14]** Neuroimmune Regulation of GABAergic Neurons Within the Ventral Tegmental Area During Withdrawal from Chronic Morphine - PubMed — https://pubmed.ncbi.nlm.nih.gov/26202104/
- **[S15]** Neuroimmune Regulation of GABAergic Neurons Within the Ventral Tegmental Area During Withdrawal from Chronic Morphine - PubMed — https://pubmed.ncbi.nlm.nih.gov/26202104/
