## Supplementary material for "Understanding Substance Dependence: What Differentiates Addictive from Non-Addictive Drugs?": Folder for supplementary files: addictive-genes-claims-first100-042-dbi-gene-protein-is-implicated-in-addiction-b42c8956cc8f.md

# addictive-genes-claims-first100-042-b42c8956cc8f

**Original claim:** DBI gene/protein is implicated in addiction.

## Claim

DBI gene/protein is implicated in addiction.

## Verdict

Verdict: Mixed Evidence

DBI (diazepam binding inhibitor, also called ACBP) is **plausibly implicated in addiction-related biology**, especially in **preclinical models of opioid and alcohol dependence/withdrawal**, and there is **some human alcohol-dependence genetic signal** from small candidate-gene studies. However, the strongest evidence is **animal/mechanistic rather than robust human clinical-genetic evidence**, and at least one human biomarker study was negative. ([pmc.ncbi.nlm.nih.gov](https://pmc.ncbi.nlm.nih.gov/articles/PMC3987987/)) [S1]

## Confidence

Medium

The claim is not unsupported, but the evidence base is uneven: there are multiple relevant preclinical studies, while the human evidence is limited, older, and not clearly replicated at modern large-study standards. ([pubmed.ncbi.nlm.nih.gov](https://pubmed.ncbi.nlm.nih.gov/15950760/?utm_source=openai)) [S2]

## Supporting evidence

DBI is not a random metabolic protein with no brain role: it is an endogenous ligand/modulator at benzodiazepine-sensitive GABA\(_A\) receptor sites, and DBI deletion in mice abolished endogenous potentiation of GABAergic transmission in the thalamic reticular nucleus; re-expression rescued the effect. That establishes DBI as a bona fide regulator of inhibitory neurotransmission in the CNS, which is highly relevant to addiction circuitry. ([pmc.ncbi.nlm.nih.gov](https://pmc.ncbi.nlm.nih.gov/articles/PMC3987987/)) [S3]

In a rat morphine conditioned-place-preference model, chronic morphine increased **DBI mRNA** in several classic addiction-related regions, including the **ventral tegmental area, nucleus accumbens, amygdala, and hippocampal CA1**. During withdrawal, DBI mRNA remained elevated, and expression in those regions positively correlated with conditioned place preference. That is direct preclinical evidence linking DBI expression to drug reward/withdrawal-associated behavior. ([pubmed.ncbi.nlm.nih.gov](https://pubmed.ncbi.nlm.nih.gov/15950760/?utm_source=openai)) [S4]

A later opioid study went beyond correlation. In mice, DBI and DBI-derived peptides modulated morphine responses, and blocking DBI fragments reduced **naloxone-precipitated withdrawal jumping**; anti-DBI antiserum also reversed morphine analgesic tolerance. The authors concluded that DBI-related signaling could be a biological target for **opioid tolerance and dependence**, which is stronger evidence for functional involvement than expression data alone. ([pubmed.ncbi.nlm.nih.gov](https://pubmed.ncbi.nlm.nih.gov/33259865/)) [S5]

For alcohol, older preclinical work reported that chronic ethanol exposure and withdrawal increase **DBI mRNA** in mouse brain, and another study specifically described “ethanol-induced increases of diazepam binding inhibitor (DBI) and its mRNA in the mouse brain.” These findings support DBI responsiveness to chronic alcohol exposure in dependence-like states. ([sciencedirect.com](https://www.sciencedirect.com/science/article/abs/pii/S0169328X97003306?utm_source=openai)) [S6]

There is also limited human genetic evidence. A 2014 Japanese case-control study reported a significant difference in **DBI rs2276596** allele frequency between alcohol-dependent patients and controls, concluding that a DBI variant may be a risk factor for alcohol dependence in that population. ([pubmed.ncbi.nlm.nih.gov](https://pubmed.ncbi.nlm.nih.gov/24818357/)) [S7]

## Contradicting or limiting evidence

Human evidence is much weaker than the animal evidence. In a cerebrospinal-fluid study of abstinent alcoholics versus controls, investigators found **no significant difference** in CSF DBI concentration and no significant correlation with anxiety measures, although DBI correlated with CRH in both groups. So at least one human biomarker study did **not** support a clear case-control difference. ([pubmed.ncbi.nlm.nih.gov](https://pubmed.ncbi.nlm.nih.gov/2333358/?utm_source=openai)) [S8]

The positive human genetic evidence also appears limited to relatively small candidate-gene studies in Japanese samples. By contrast, a large community-based study examining **GABA system genes**, which included **DBI**, found that **no method produced results indicative of an effect** of GABA-system variants on alcohol use or misuse in that sample. That does not disprove a DBI effect specifically, but it weakens the argument that DBI is a broadly established human addiction gene. ([deepblue.lib.umich.edu](https://deepblue.lib.umich.edu/bitstream/handle/2027.42/106921/acer12352.pdf?sequence=1&utm_source=openai)) [S9]

The substance coverage is also uneven. The clearest experimental evidence is for **opioids** and to a lesser extent **alcohol**; I did not find comparably strong primary evidence in humans for DBI in nicotine, stimulant, or cannabis addiction. So the broad phrase “implicated in addiction” is more defensible if interpreted as **some addiction-related contexts**, not as a general, firmly established addiction gene across disorders. ([pubmed.ncbi.nlm.nih.gov](https://pubmed.ncbi.nlm.nih.gov/15950760/?utm_source=openai)) [S10]

## Mechanistic / biological rationale

A biologically plausible mechanism exists. DBI is encoded as a protein that can modulate **GABA\(_A\)** receptor signaling at benzodiazepine-sensitive sites in brain synapses, and experimental deletion/rescue studies show that endogenous DBI can tune inhibitory synaptic strength. Because addiction involves maladaptive plasticity in reward and stress circuits that depend heavily on inhibitory control, DBI is well positioned to influence those processes. ([pmc.ncbi.nlm.nih.gov](https://pmc.ncbi.nlm.nih.gov/articles/PMC3987987/)) [S11]

The opioid studies fit that logic well. DBI expression rose in the **VTA, nucleus accumbens, amygdala, and hippocampus** during morphine dependence/withdrawal, and another study linked DBI’s anti-opioid effects to **lateral habenula–periaqueductal gray** circuitry plus **NMDA receptor / nitric oxide** signaling. Taken together, these data suggest DBI may shape withdrawal, tolerance, and reinforcement by altering inhibitory/excitatory balance in addiction-relevant networks. This is still an inference from animal work, but it is a biologically coherent one. ([pubmed.ncbi.nlm.nih.gov](https://pubmed.ncbi.nlm.nih.gov/15950760/?utm_source=openai)) [S12]

For alcohol, the rationale is similarly plausible: ethanol strongly affects GABAergic signaling, and chronic alcohol exposure/withdrawal was associated with increased DBI expression in preclinical studies. That makes DBI a credible participant in neural adaptation to repeated alcohol exposure, even though the human validation remains limited. ([sciencedirect.com](https://www.sciencedirect.com/science/article/abs/pii/S0169328X97003306?utm_source=openai)) [S13]

## Bottom line

DBI is **reasonably implicated in addiction-related neurobiology**, but the support is **mostly preclinical**. The best evidence shows DBI changes in opioid and alcohol dependence models and can influence opioid tolerance/withdrawal behaviors. Human evidence exists but is limited and mixed, with a small positive alcohol-genetics study and at least one negative CSF study. So the most accurate overall judgment is **Mixed Evidence**, not “Highly Supported.” ([pubmed.ncbi.nlm.nih.gov](https://pubmed.ncbi.nlm.nih.gov/15950760/?utm_source=openai)) [S14]

## Cited sources

- **[S1]** 
            Endogenous Positive Allosteric Modulation of GABAA Receptors by Diazepam binding inhibitor - PMC
         — https://pmc.ncbi.nlm.nih.gov/articles/PMC3987987/
- **[S2]** Conditioned place preference associates with the mRNA expression of diazepam binding inhibitor in brain regions of the addicted rat during withdrawal - PubMed — https://pubmed.ncbi.nlm.nih.gov/15950760/?utm_source=openai
- **[S3]** 
            Endogenous Positive Allosteric Modulation of GABAA Receptors by Diazepam binding inhibitor - PMC
         — https://pmc.ncbi.nlm.nih.gov/articles/PMC3987987/
- **[S4]** Conditioned place preference associates with the mRNA expression of diazepam binding inhibitor in brain regions of the addicted rat during withdrawal - PubMed — https://pubmed.ncbi.nlm.nih.gov/15950760/?utm_source=openai
- **[S5]** Possible mechanism and potential application of anti-opioid effect of diazepam-binding inhibitor - PubMed — https://pubmed.ncbi.nlm.nih.gov/33259865/
- **[S6]** Functional involvement of benzodiazepine receptors in ethanol-induced increases of diazepam binding inhibitor (DBI) and its mRNA in the mouse brain - ScienceDirect — https://www.sciencedirect.com/science/article/abs/pii/S0169328X97003306?utm_source=openai
- **[S7]** Relationship between alcohol dependence and the DBI rs2276596 (C/A) polymorphism in Japanese - PubMed — https://pubmed.ncbi.nlm.nih.gov/24818357/
- **[S8]** CSF diazepam-binding inhibitor in alcoholics and normal controls - PubMed — https://pubmed.ncbi.nlm.nih.gov/2333358/?utm_source=openai
- **[S9]** ALCOHOLISM: CLINICAL AND EXPERIMENTAL RESEARCH — https://deepblue.lib.umich.edu/bitstream/handle/2027.42/106921/acer12352.pdf?sequence=1&utm_source=openai
- **[S10]** Conditioned place preference associates with the mRNA expression of diazepam binding inhibitor in brain regions of the addicted rat during withdrawal - PubMed — https://pubmed.ncbi.nlm.nih.gov/15950760/?utm_source=openai
- **[S11]** 
            Endogenous Positive Allosteric Modulation of GABAA Receptors by Diazepam binding inhibitor - PMC
         — https://pmc.ncbi.nlm.nih.gov/articles/PMC3987987/
- **[S12]** Conditioned place preference associates with the mRNA expression of diazepam binding inhibitor in brain regions of the addicted rat during withdrawal - PubMed — https://pubmed.ncbi.nlm.nih.gov/15950760/?utm_source=openai
- **[S13]** Functional involvement of benzodiazepine receptors in ethanol-induced increases of diazepam binding inhibitor (DBI) and its mRNA in the mouse brain - ScienceDirect — https://www.sciencedirect.com/science/article/abs/pii/S0169328X97003306?utm_source=openai
- **[S14]** Conditioned place preference associates with the mRNA expression of diazepam binding inhibitor in brain regions of the addicted rat during withdrawal - PubMed — https://pubmed.ncbi.nlm.nih.gov/15950760/?utm_source=openai
