## Supplementary material for "Understanding Substance Dependence: What Differentiates Addictive from Non-Addictive Drugs?": Folder for supplementary files: addictive-genes-claims-first100-043-tmbim1-gene-protein-is-implicated-in-addiction-9d8d830ca7a7.md

# addictive-genes-claims-first100-043-9d8d830ca7a7

**Original claim:** TMBIM1 gene/protein is implicated in addiction.

## Claim

TMBIM1 (also called **RECS1** or **LFG3**) gene/protein is implicated in addiction. TMBIM1 encodes a transmembrane BAX inhibitor motif-containing protein with reported calcium-channel activity, death-receptor binding, and localization to endosomal/lysosomal/Golgi membranes. ([ncbi.nlm.nih.gov](https://www.ncbi.nlm.nih.gov/gene/64114)) [S1]

## Verdict

Verdict: No Evidence

I did **not** find direct evidence that TMBIM1 is an addiction gene/protein in the usual sense used in addiction biology: i.e., replicated association with substance use disorder risk, altered expression in addiction-relevant human or animal datasets, or functional experiments showing effects on drug reward, self-administration, withdrawal, craving, or relapse. The literature I found instead centers on TMBIM1/RECS1 as a lysosomal/endosomal calcium-homeostasis and cell-death regulator, with disease links mainly in steatohepatitis, cardiovascular remodeling, vascular biology, and cancer. ([pmc.ncbi.nlm.nih.gov](https://pmc.ncbi.nlm.nih.gov/articles/PMC8589314/)) [S2]

## Confidence

Medium

Confidence is **medium** because the searched evidence base is fairly informative: representative addiction-genetics reviews, integrated GWAS/transcriptomic reviews, a large cross-substance GWAS paper, and a nicotine TWAS paper did not identify TMBIM1, and targeted scholarly searches for “TMBIM1/RECS1 + addiction/substance use disorder” did not surface direct addiction studies. Still, absence from these sources is not absolute proof that no niche or unpublished signal exists. ([pubmed.ncbi.nlm.nih.gov](https://pubmed.ncbi.nlm.nih.gov/35491750/)) [S3]

## Supporting evidence

The strongest case **for plausibility** is indirect, not addiction-specific. TMBIM1/RECS1 is a member of the TMBIM family involved in intracellular calcium homeostasis and cell-death regulation. In mechanistic work, RECS1 localized to lysosomes, promoted lysosomal acidification, functioned as a pH-dependent calcium/sodium channel, and modulated lysosomal membrane permeabilization and cell death under stress. Broader TMBIM-family reviews also place TMBIM1 in endosomal/lysosomal/Golgi compartments and link it to death-receptor/Fas signaling and calcium handling. ([pmc.ncbi.nlm.nih.gov](https://pmc.ncbi.nlm.nih.gov/articles/PMC8589314/)) [S4]

TMBIM1 is also not absent from the nervous system. Human Protein Atlas brain data show that TMBIM1 RNA is **detected in all major brain regions**, has **low regional specificity**, and clusters with an **astrocyte-associated** expression module (“Astrocytes – Mixed function”). That means the gene is at least biologically positioned to matter in CNS homeostasis. ([proteinatlas.org](https://www.proteinatlas.org/ENSG00000135926-TMBIM1/brain)) [S5]

This matters because astrocytes and glia are increasingly recognized as participants in addiction biology. Reviews describe astrocytes as contributors to synaptic plasticity and drug-induced adaptations, and broader glial/neuroimmune literature links astrocytic and microglial signaling to addiction-like states. ([pubmed.ncbi.nlm.nih.gov](https://pubmed.ncbi.nlm.nih.gov/37724324/)) [S6]

A second indirect argument is through **innate immune/TLR4 signaling**. TMBIM1 has been shown, in non-addiction contexts, to promote lysosomal degradation of TLR4 and suppress inflammatory pathology. Separately, addiction/neuroimmune reviews report that TLR4 and related innate immune pathways can influence alcohol- and drug-related behaviors in some models, including craving/reinstatement-related phenomena and aspects of reward learning. ([pubmed.ncbi.nlm.nih.gov](https://pubmed.ncbi.nlm.nih.gov/28481357/?utm_source=openai)) [S7]

Taken together, these points give TMBIM1 some **biological plausibility** as a potential modifier of neural or neuroimmune processes that could, in theory, intersect with addiction. But plausibility is not the same as implication. ([proteinatlas.org](https://www.proteinatlas.org/ENSG00000135926-TMBIM1/brain)) [S8]

## Contradicting or limiting evidence

The main limitation for the claim is simple: **direct addiction evidence is missing**. A 2022 integrated review of genomic and transcriptomic studies across alcohol, nicotine, cannabis, opioid, cocaine, and methamphetamine phenotypes identified shared and distinct addiction-related genes/pathways, but a text search of that paper did not find TMBIM1. ([pubmed.ncbi.nlm.nih.gov](https://pubmed.ncbi.nlm.nih.gov/35491750/)) [S9]

Likewise, a 2021 review of substance-use-disorder genetics summarizes robust findings such as alcohol-metabolizing loci (**ADH1B**, **ALDH2**) and the nicotine-related **CHRNA5-CHRNA3-CHRNB4** cluster; TMBIM1 is not discussed there. ([cambridge.org](https://www.cambridge.org/core/services/aop-cambridge-core/content/view/B3BAE9D2DCF78C7C4833A8DB4420F5B2/S0033291721000969a.pdf/genetics-of-substance-use-disorders-a-review.pdf)) [S10]

A 2024 JCI review of generalized genetic liability to substance use disorders emphasizes cross-substance shared risk, dopaminergic regulation, and neuronal development mechanisms; again, TMBIM1 was not found in the article text. The same is true for a large multivariate GWAS of over 1 million participants and for a brain-tissue TWAS meta-analysis of nicotine addiction. ([jci.org](https://www.jci.org/articles/view/172881)) [S11]

The gene-centric annotation sources I found also do not point toward addiction. On the human NCBI Gene page, the mapped phenotype note references inflammatory bowel disease, and GeneRIFs center on steatohepatitis, colorectal cancer, calcium homeostasis, and Fas/apoptosis-related biology rather than substance use disorder. ([ncbi.nlm.nih.gov](https://www.ncbi.nlm.nih.gov/gene/64114)) [S12]

So the problem is not that there is strong evidence *against* TMBIM1 in addiction; it is that the current literature does not supply the kind of evidence needed to say it is actually implicated. ([pubmed.ncbi.nlm.nih.gov](https://pubmed.ncbi.nlm.nih.gov/35491750/)) [S13]

## Mechanistic / biological rationale

If TMBIM1 were eventually linked to addiction, the most plausible routes would be **indirect**: regulation of lysosomal calcium homeostasis, astrocyte biology, cell-stress responses, or neuroimmune/TLR4 signaling. Each of those processes has some relevance to addiction-related plasticity, especially in glial modulation of glutamate signaling and inflammatory tone in reward circuits. ([pmc.ncbi.nlm.nih.gov](https://pmc.ncbi.nlm.nih.gov/articles/PMC8589314/)) [S14]

However, the key mechanistic bridge is missing. I did not find evidence that TMBIM1 variants, expression changes, or perturbation specifically alter mesolimbic dopamine signaling, nucleus accumbens plasticity, drug reinforcement, sensitization, dependence, withdrawal, or relapse behavior. Nor did I find direct human postmortem, GWAS, TWAS, or animal-behavior studies making that connection. ([pubmed.ncbi.nlm.nih.gov](https://pubmed.ncbi.nlm.nih.gov/35491750/)) [S15]

## Bottom line

At present, **TMBIM1 should not be described as an established addiction-implicated gene/protein**. The available literature supports TMBIM1 as a brain-expressed, astrocyte-associated lysosomal/endosomal regulator of calcium homeostasis, stress responses, and inflammatory signaling, but I found **no direct addiction-specific human genetic, transcriptomic, or functional behavioral evidence** tying it to substance use disorders. The most accurate summary is: **biologically plausible, but currently unsupported for addiction.** ([proteinatlas.org](https://www.proteinatlas.org/ENSG00000135926-TMBIM1/brain)) [S16]

## Cited sources

- **[S1]** TMBIM1 transmembrane BAX inhibitor motif containing 1 [Homo sapiens (human)] - Gene - NCBI — https://www.ncbi.nlm.nih.gov/gene/64114
- **[S2]** 
            Control of lysosomal-mediated cell death by the pH-dependent calcium channel RECS1 - PMC
         — https://pmc.ncbi.nlm.nih.gov/articles/PMC8589314/
- **[S3]** Common and distinguishing genetic factors for substance use behavior and disorder: an integrated analysis of genomic and transcriptomic studies from both human and animal studies - PubMed — https://pubmed.ncbi.nlm.nih.gov/35491750/
- **[S4]** 
            Control of lysosomal-mediated cell death by the pH-dependent calcium channel RECS1 - PMC
         — https://pmc.ncbi.nlm.nih.gov/articles/PMC8589314/
- **[S5]** Brain tissue expression of TMBIM1 - Summary - The Human Protein Atlas — https://www.proteinatlas.org/ENSG00000135926-TMBIM1/brain
- **[S6]** Astrocytes: the neglected stars in the central nervous system and drug addiction - PubMed — https://pubmed.ncbi.nlm.nih.gov/37724324/
- **[S7]** Tmbim1 is a multivesicular body regulator that protects against non-alcoholic fatty liver disease in mice and monkeys by targeting the lysosomal degradation of Tlr4 - PubMed — https://pubmed.ncbi.nlm.nih.gov/28481357/?utm_source=openai
- **[S8]** Brain tissue expression of TMBIM1 - Summary - The Human Protein Atlas — https://www.proteinatlas.org/ENSG00000135926-TMBIM1/brain
- **[S9]** Common and distinguishing genetic factors for substance use behavior and disorder: an integrated analysis of genomic and transcriptomic studies from both human and animal studies - PubMed — https://pubmed.ncbi.nlm.nih.gov/35491750/
- **[S10]** S0033291721000969jrv 2189..2200 — https://www.cambridge.org/core/services/aop-cambridge-core/content/view/B3BAE9D2DCF78C7C4833A8DB4420F5B2/S0033291721000969a.pdf/genetics-of-substance-use-disorders-a-review.pdf
- **[S11]** 
JCI -
Generalized genetic liability to substance use disorders
 — https://www.jci.org/articles/view/172881
- **[S12]** TMBIM1 transmembrane BAX inhibitor motif containing 1 [Homo sapiens (human)] - Gene - NCBI — https://www.ncbi.nlm.nih.gov/gene/64114
- **[S13]** Common and distinguishing genetic factors for substance use behavior and disorder: an integrated analysis of genomic and transcriptomic studies from both human and animal studies - PubMed — https://pubmed.ncbi.nlm.nih.gov/35491750/
- **[S14]** 
            Control of lysosomal-mediated cell death by the pH-dependent calcium channel RECS1 - PMC
         — https://pmc.ncbi.nlm.nih.gov/articles/PMC8589314/
- **[S15]** Common and distinguishing genetic factors for substance use behavior and disorder: an integrated analysis of genomic and transcriptomic studies from both human and animal studies - PubMed — https://pubmed.ncbi.nlm.nih.gov/35491750/
- **[S16]** Brain tissue expression of TMBIM1 - Summary - The Human Protein Atlas — https://www.proteinatlas.org/ENSG00000135926-TMBIM1/brain
