## Supplementary material for "Understanding Substance Dependence: What Differentiates Addictive from Non-Addictive Drugs?": Folder for supplementary files: addictive-genes-claims-first100-044-cox7a2-gene-protein-is-implicated-in-addiction-872225ee655a.md

# addictive-genes-claims-first100-044-872225ee655a

**Original claim:** COX7A2 gene/protein is implicated in addiction.

## Claim

COX7A2 gene/protein is implicated in addiction.

## Verdict

Verdict: Mixed Evidence

## Confidence

Medium

## Supporting evidence

COX7A2 is a real, biologically relevant brain-energy gene: it encodes a nuclear-encoded subunit of cytochrome c oxidase/mitochondrial complex IV, the terminal enzyme of the respiratory chain, and is broadly expressed in human tissues including brain. NCBI describes it as part of oxidative phosphorylation, and Human Protein Atlas reports low regional specificity across brain regions rather than absence from brain. ([ncbi.nlm.nih.gov](https://www.ncbi.nlm.nih.gov/gene/1347)) [S1]

The strongest addiction-relevant signal I found is in alcohol use disorder (AUD), but it is **pathway/transcriptomic** rather than **gene-causal** evidence. In a pathway-level analysis of 17 postmortem human brain-expression datasets from alcohol-dependent patients, oxidative phosphorylation was the most consistently downregulated pathway, appearing in 6 of 17 datasets; the authors specifically listed **COX7A2** among the recurrent genes in that pathway. The affected regions included prefrontal cortex, nucleus accumbens, hippocampus, and amygdala-related samples. ([pmc.ncbi.nlm.nih.gov](https://pmc.ncbi.nlm.nih.gov/articles/PMC6956180/)) [S2]

That finding fits older human brain transcriptomic work showing that chronic cocaine exposure, and to a lesser degree alcohol exposure, is associated with suppression of mitochondrial inner-membrane genes in hippocampus, suggesting a recurring bioenergetic signature in addiction-related brain tissue. This does **not** prove COX7A2 specifically drives addiction, but it makes its appearance in the AUD pathway analysis biologically plausible rather than random. ([pmc.ncbi.nlm.nih.gov](https://pmc.ncbi.nlm.nih.gov/articles/PMC3081016/)) [S3]

There is also **cellular exposure evidence** for cannabinoids. In human iPSC-derived neurons, acute and chronic THC exposure altered many genes; the paper explicitly names **COX7A2** among affected mitochondrial genes and reports that oxidative phosphorylation was strongly altered after THC exposure. Again, this is not addiction-behavior evidence, but it places COX7A2 within a drug-responsive neuronal mitochondrial program. ([pubmed.ncbi.nlm.nih.gov](https://pubmed.ncbi.nlm.nih.gov/29691375/)) [S4]

More broadly, mitochondrial dysfunction is a recognized theme in addiction biology. A review focused on cocaine concluded that cocaine affects brain metabolism and impairs oxidative phosphorylation, and animal/human studies in other substances report mitochondrial abnormalities as well, including decreased mitochondrial DNA copy number in opiate addiction and reduced respiratory capacity after chronic alcohol exposure in mouse medial prefrontal cortex. These findings support a plausible context in which a complex IV subunit such as COX7A2 could matter. ([pubmed.ncbi.nlm.nih.gov](https://pubmed.ncbi.nlm.nih.gov/33626141/)) [S5]

## Contradicting or limiting evidence

Despite those signals, I did **not** find strong evidence that COX7A2 is an established addiction gene in the way genes such as **OPRM1**, **ADH1B/ADH1C**, **CHRNA3/CHRNA4**, or **FURIN/PPP6C** are discussed in the substance-use literature. In a 2022 integrated analysis of GWAS and RNA-seq studies across alcohol, nicotine, and other drugs, the shared cross-substance genes highlighted were **CACNB2, GRIN2B, PLXDC2, and PKNOX2**—not COX7A2. ([pubmed.ncbi.nlm.nih.gov](https://pubmed.ncbi.nlm.nih.gov/35491750/)) [S6]

Likewise, modern human genetic studies of opioid addiction have prioritized other loci. A multi-trait GWAS of opioid addiction emphasized **OPRM1** and gene-level associations such as **PPP6C** and **FURIN**, with no COX7A2 signal reported in the paper text. ([nature.com](https://www.nature.com/articles/s41598-022-21003-y)) [S7]

For alcohol dependence, review-level summaries of GWAS findings emphasize the **ADH cluster** as the most robust locus, plus genes such as **SERINC2, KIAA0040, NRD1, and HTR7**; COX7A2 is not among the repeatedly prioritized risk genes in those reviews. ([pmc.ncbi.nlm.nih.gov](https://pmc.ncbi.nlm.nih.gov/articles/PMC4187224/)) [S8]

Evidence is also limited by poor reproducibility across transcriptomic studies. A 2023 systematic review of opioid functional genomics found that across 44 transcriptomic studies in major brain regions, only **Cdkn1a** was consistently identified in eleven studies, and overall cross-study convergence was surprisingly low. That makes isolated omics appearances—especially for metabolic genes—hard to interpret as proof of addiction-specific causality. ([nature.com](https://www.nature.com/articles/s41380-023-02238-1)) [S9]

Most importantly, I did not find direct evidence that **genetic variation in COX7A2 increases addiction risk**, nor experiments showing that selective manipulation of COX7A2 in addiction-relevant neurons changes drug taking, reward, craving, withdrawal, or relapse. The available evidence is therefore largely **indirect**: postmortem expression/pathway changes, cellular drug-exposure signatures, and general mitochondrial rationale. ([pmc.ncbi.nlm.nih.gov](https://pmc.ncbi.nlm.nih.gov/articles/PMC6956180/)) [S10]

## Mechanistic / biological rationale

Mechanistically, the claim is plausible. Addiction involves durable neuroadaptations in reward and control circuits, and those processes are energetically demanding. Mitochondria regulate ATP production, calcium buffering, redox balance, apoptosis vulnerability, and synaptic plasticity; reviews of cocaine and other substance exposures repeatedly point to oxidative phosphorylation and mitochondrial dysfunction as part of the pathology. Because COX7A2 is a complex IV subunit, altered COX7A2 expression could in principle affect neuronal bioenergetics and ROS handling in addiction-relevant circuits. ([ncbi.nlm.nih.gov](https://www.ncbi.nlm.nih.gov/gene/1347)) [S11]

There is also precedent that COX7A2 matters in high-energy tissues and redox biology outside addiction. NCBI classifies it as a core complex IV gene, and a review of knockout mouse models notes human COX7A2 association with a rare fatal infantile cardioencephalomyopathy, underscoring that this is not a trivial bystander gene. In a non-neural cell model, Cox7a2 overexpression increased reactive oxygen species and altered steroidogenesis, which at least shows the protein can influence cellular redox state. ([ncbi.nlm.nih.gov](https://www.ncbi.nlm.nih.gov/gene/1347)) [S12]

But that mechanistic plausibility cuts both ways: many oxidative-phosphorylation genes change in stressed or diseased tissue, and such changes may reflect downstream metabolic adaptation rather than a primary addiction driver. The AUD pathway paper itself cautions that one should not automatically infer lower pathway function from transcript changes alone—only that many downregulated genes belong to oxidative phosphorylation. ([pmc.ncbi.nlm.nih.gov](https://pmc.ncbi.nlm.nih.gov/articles/PMC6956180/)) [S13]

## Bottom line

There **is some evidence** linking COX7A2 to addiction-related biology, mainly because it appears within **downregulated oxidative-phosphorylation signatures in postmortem AUD brain** and within **drug-responsive mitochondrial gene programs** such as THC-exposed human neurons. ([pmc.ncbi.nlm.nih.gov](https://pmc.ncbi.nlm.nih.gov/articles/PMC6956180/)) [S14]

However, the evidence is **indirect and not yet gene-specific enough** to call COX7A2 an established addiction gene. I found **no robust replicated GWAS signal, no clear causal animal-behavior data, and no major review that prioritizes COX7A2 as a core addiction susceptibility gene**. ([pubmed.ncbi.nlm.nih.gov](https://pubmed.ncbi.nlm.nih.gov/35491750/)) [S15]

So the fairest summary is: **COX7A2 is plausibly involved in addiction-associated mitochondrial/homeostatic changes, especially in alcohol-related brain transcriptomics, but current evidence is too limited to say it is firmly implicated as a direct addiction risk gene or therapeutic target.** ([pmc.ncbi.nlm.nih.gov](https://pmc.ncbi.nlm.nih.gov/articles/PMC6956180/)) [S16]

## Cited sources

- **[S1]** https://www.ncbi.nlm.nih.gov/gene/1347 — https://www.ncbi.nlm.nih.gov/gene/1347
- **[S2]** https://pmc.ncbi.nlm.nih.gov/articles/PMC6956180/ — https://pmc.ncbi.nlm.nih.gov/articles/PMC6956180/
- **[S3]** https://pmc.ncbi.nlm.nih.gov/articles/PMC3081016/ — https://pmc.ncbi.nlm.nih.gov/articles/PMC3081016/
- **[S4]** https://pubmed.ncbi.nlm.nih.gov/29691375/ — https://pubmed.ncbi.nlm.nih.gov/29691375/
- **[S5]** https://pubmed.ncbi.nlm.nih.gov/33626141/ — https://pubmed.ncbi.nlm.nih.gov/33626141/
- **[S6]** https://pubmed.ncbi.nlm.nih.gov/35491750/ — https://pubmed.ncbi.nlm.nih.gov/35491750/
- **[S7]** https://www.nature.com/articles/s41598-022-21003-y — https://www.nature.com/articles/s41598-022-21003-y
- **[S8]** https://pmc.ncbi.nlm.nih.gov/articles/PMC4187224/ — https://pmc.ncbi.nlm.nih.gov/articles/PMC4187224/
- **[S9]** https://www.nature.com/articles/s41380-023-02238-1 — https://www.nature.com/articles/s41380-023-02238-1
- **[S10]** https://pmc.ncbi.nlm.nih.gov/articles/PMC6956180/ — https://pmc.ncbi.nlm.nih.gov/articles/PMC6956180/
- **[S11]** https://www.ncbi.nlm.nih.gov/gene/1347 — https://www.ncbi.nlm.nih.gov/gene/1347
- **[S12]** https://www.ncbi.nlm.nih.gov/gene/1347 — https://www.ncbi.nlm.nih.gov/gene/1347
- **[S13]** https://pmc.ncbi.nlm.nih.gov/articles/PMC6956180/ — https://pmc.ncbi.nlm.nih.gov/articles/PMC6956180/
- **[S14]** https://pmc.ncbi.nlm.nih.gov/articles/PMC6956180/ — https://pmc.ncbi.nlm.nih.gov/articles/PMC6956180/
- **[S15]** https://pubmed.ncbi.nlm.nih.gov/35491750/ — https://pubmed.ncbi.nlm.nih.gov/35491750/
- **[S16]** https://pmc.ncbi.nlm.nih.gov/articles/PMC6956180/ — https://pmc.ncbi.nlm.nih.gov/articles/PMC6956180/
