## Supplementary material for "Understanding Substance Dependence: What Differentiates Addictive from Non-Addictive Drugs?": Folder for supplementary files: addictive-genes-claims-first100-045-timp3-gene-protein-is-implicated-in-addiction-d4a19bf2678f.md

# addictive-genes-claims-first100-045-d4a19bf2678f

**Original claim:** TIMP3 gene/protein is implicated in addiction.

## Claim

TIMP3 gene/protein is implicated in addiction.

## Verdict

Verdict: No Evidence

Based on the literature I could find, there is **not currently convincing direct evidence** that **TIMP3 itself** is an addiction-implicated gene/protein in the usual sense of having reproducible human genetic association, addiction-relevant differential regulation, or causal behavioral evidence from manipulation studies. What **is** supported is a broader role for **extracellular matrix (ECM) remodeling and matrix metalloproteinase (MMP)/TIMP signaling** in addiction-related plasticity; however, the addiction literature points much more clearly to **MMP-2/MMP-9 and TIMP-1/TIMP-2** than to TIMP3. ([pmc.ncbi.nlm.nih.gov](https://pmc.ncbi.nlm.nih.gov/articles/PMC4589426/)) [S1]

## Confidence

High

The confidence is high because addiction-focused reviews specifically discussing MMP/TIMP biology either do **not** identify TIMP3 as an established addiction factor or explicitly state that relatively little is known about TIMP-3 in brain, while the directly discussed addiction-related TIMP findings center on TIMP2 rather than TIMP3. ([pmc.ncbi.nlm.nih.gov](https://pmc.ncbi.nlm.nih.gov/articles/PMC4589426/)) [S2]

## Supporting evidence

There **is** a biologically relevant framework in which TIMP3 could matter. Addiction-related neuroplasticity involves ECM remodeling at cortico-striatal/accumbal synapses, and reviews conclude that MMP-regulated extracellular signaling is important for developing and expressing drug-seeking behaviors. An earlier review likewise argued that MMP/TIMP-regulated ECM remodeling may contribute to neural plasticity underlying addiction and relapse. So, at the pathway level, TIMP-family biology is clearly addiction-adjacent. ([pmc.ncbi.nlm.nih.gov](https://pmc.ncbi.nlm.nih.gov/articles/PMC4589426/)) [S3]

The strongest TIMP3-specific addiction-adjacent paper I found was a **2025 rat methamphetamine conditioned place preference (CPP)** study. In that work, TIMP3 was identified by network pharmacology as one of 12 putative targets of exosome-associated **miR-181a-5p**; the authors also showed miR-181a-5p could bind the **TIMP3 3′UTR** in a luciferase assay and speculated that these targets may participate in methamphetamine dependence. This is **hypothesis-generating** evidence that places TIMP3 in an addiction-relevant molecular network. ([pmc.ncbi.nlm.nih.gov](https://pmc.ncbi.nlm.nih.gov/articles/PMC12117265/)) [S4]

That said, even this “supporting” evidence is indirect: it links TIMP3 to a **miRNA network in an addiction model**, not to addiction behavior through direct TIMP3 manipulation. So it suggests plausibility, not implication in a strong evidentiary sense. ([pmc.ncbi.nlm.nih.gov](https://pmc.ncbi.nlm.nih.gov/articles/PMC12117265/)) [S5]

## Contradicting or limiting evidence

A key limitation is that the addiction-focused ECM/MMP review from 2015 explicitly notes that **“relatively little is known about TIMP-3 and TIMP-4.”** The same review states that TIMP-3 is expressed at **very low levels** in brain regions such as cortex, cerebellum, olfactory bulb, and brain stem, but when it discusses addiction-related TIMP findings, the examples are **TIMP-1 and especially TIMP-2**—including methamphetamine-induced TIMP2 upregulation and evidence that blocking TIMP2 upregulation increases methamphetamine locomotor sensitization, plus TIMP-2 increases after cocaine-primed reinstatement. No analogous direct addiction evidence is presented for TIMP3. ([pmc.ncbi.nlm.nih.gov](https://pmc.ncbi.nlm.nih.gov/articles/PMC4589426/)) [S6]

The older review on MMPs/TIMPs and addiction similarly treats TIMPs largely at the family level and does not provide TIMP3-specific addiction findings. It lists TIMP3 among TIMP family members that inhibit MMPs, but the paper’s addiction discussion remains broad and does not establish TIMP3 as a causal or associated addiction molecule. ([pmc.ncbi.nlm.nih.gov](https://pmc.ncbi.nlm.nih.gov/articles/PMC2821634/)) [S7]

The 2025 methamphetamine paper is also limited methodologically. It used **CPP**, network pharmacology, molecular docking, and luciferase assays, but did **not** manipulate TIMP3 expression or protein function in vivo and did not show that TIMP3 changes alter methamphetamine reward, relapse, or drug seeking. The authors also acknowledged that CPP does not assess relapse/drug-seeking, and they noted other mechanistic gaps. So this study does not establish TIMP3 as an addiction gene/protein. ([pmc.ncbi.nlm.nih.gov](https://pmc.ncbi.nlm.nih.gov/articles/PMC12117265/)) [S8]

Meanwhile, the direct TIMP3 literature that does exist is largely about **brain injury, BBB regulation, neuroinflammation, and neuronal survival**, not addiction. Recombinant TIMP3 preserves blood-brain barrier integrity after traumatic brain injury, promotes neuronal survival and neurite/dendritic outgrowth, and improves neurocognitive outcomes in TBI models. In transient global ischemia, **Timp3 knockout** reduces astrocytosis, microglial reactivity, TNFR1 expression, and neuronal death. Those are meaningful CNS roles, but they do **not** demonstrate addiction involvement. ([pubmed.ncbi.nlm.nih.gov](https://pubmed.ncbi.nlm.nih.gov/23175708/?utm_source=openai)) [S9]

## Mechanistic / biological rationale

Mechanistically, TIMP3 is still interesting. TIMP3 is an **ECM-bound inhibitor** of multiple metalloproteinases and an important endogenous inhibitor of **ADAM17/TACE** and other proteases, placing it in pathways that regulate ECM turnover, ectodomain shedding, inflammatory signaling, and barrier integrity. Those processes are relevant to addiction because addiction involves persistent synaptic and circuit remodeling, and ECM remodeling has been implicated in drug-seeking plasticity. ([pubmed.ncbi.nlm.nih.gov](https://pubmed.ncbi.nlm.nih.gov/19581416/?utm_source=openai)) [S10]

There is also broader addiction evidence for ECM/perineuronal-net biology. For example, repeated binge drinking increases perineuronal nets in the insular cortex, and enzymatic disruption of insular perineuronal nets reduces aversion-resistant alcohol drinking. This strengthens the idea that ECM regulators could, in principle, influence addiction phenotypes. ([pubmed.ncbi.nlm.nih.gov](https://pubmed.ncbi.nlm.nih.gov/26332441/?utm_source=openai)) [S11]

But moving from **“ECM biology matters in addiction”** to **“TIMP3 is implicated in addiction”** requires TIMP3-specific evidence—e.g., human association data, region-specific expression changes after drug exposure, or causal behavioral effects of Timp3/TIMP3 manipulation in nucleus accumbens, insula, prefrontal cortex, or related circuits. I did not find such evidence. At present, the mechanistic case is therefore **plausible but unproven**. ([pmc.ncbi.nlm.nih.gov](https://pmc.ncbi.nlm.nih.gov/articles/PMC4589426/)) [S12]

## Bottom line

Current evidence does **not** substantiate **TIMP3** as an addiction-implicated gene/protein. The field supports a role for **ECM/MMP/TIMP systems broadly** in addiction-related plasticity, and TIMP3 clearly has CNS roles in neuroprotection, BBB regulation, and neuroinflammatory signaling. But the available TIMP3-addiction link is presently **indirect and insufficient**, with stronger direct addiction evidence existing for other family members such as **TIMP2** rather than TIMP3. ([pmc.ncbi.nlm.nih.gov](https://pmc.ncbi.nlm.nih.gov/articles/PMC4589426/)) [S13]

If you want, I can next do a **gene-by-gene comparison of TIMP1, TIMP2, TIMP3, and MMP9 in addiction** and rank which are actually supported.

## Cited sources

- **[S1]** 
            The tetrapartite synapse: extracellular matrix remodeling contributes to corticoaccumbens plasticity underlying drug addiction - PMC
         — https://pmc.ncbi.nlm.nih.gov/articles/PMC4589426/
- **[S2]** 
            The tetrapartite synapse: extracellular matrix remodeling contributes to corticoaccumbens plasticity underlying drug addiction - PMC
         — https://pmc.ncbi.nlm.nih.gov/articles/PMC4589426/
- **[S3]** 
            The tetrapartite synapse: extracellular matrix remodeling contributes to corticoaccumbens plasticity underlying drug addiction - PMC
         — https://pmc.ncbi.nlm.nih.gov/articles/PMC4589426/
- **[S4]** 
            Cardiomyocyte-derived exosomes carrying miR-181a-5p facilitate heart-brain crosstalk and exacerbate methamphetamine dependence in rats - PMC
         — https://pmc.ncbi.nlm.nih.gov/articles/PMC12117265/
- **[S5]** 
            Cardiomyocyte-derived exosomes carrying miR-181a-5p facilitate heart-brain crosstalk and exacerbate methamphetamine dependence in rats - PMC
         — https://pmc.ncbi.nlm.nih.gov/articles/PMC12117265/
- **[S6]** 
            The tetrapartite synapse: extracellular matrix remodeling contributes to corticoaccumbens plasticity underlying drug addiction - PMC
         — https://pmc.ncbi.nlm.nih.gov/articles/PMC4589426/
- **[S7]** 
            Contributions of Matrix Metalloproteinases to Neural Plasticity, Habituation, Associative Learning and Drug Addiction - PMC
         — https://pmc.ncbi.nlm.nih.gov/articles/PMC2821634/
- **[S8]** 
            Cardiomyocyte-derived exosomes carrying miR-181a-5p facilitate heart-brain crosstalk and exacerbate methamphetamine dependence in rats - PMC
         — https://pmc.ncbi.nlm.nih.gov/articles/PMC12117265/
- **[S9]** Mesenchymal stem cells regulate blood-brain barrier integrity through TIMP3 release after traumatic brain injury - PubMed — https://pubmed.ncbi.nlm.nih.gov/23175708/?utm_source=openai
- **[S10]** TIMP3 is reduced in atherosclerotic plaques from subjects with type 2 diabetes and increased by SirT1 - PubMed — https://pubmed.ncbi.nlm.nih.gov/19581416/?utm_source=openai
- **[S11]** Repeated Binge Drinking Increases Perineuronal Nets in the Insular Cortex - PubMed — https://pubmed.ncbi.nlm.nih.gov/26332441/?utm_source=openai
- **[S12]** 
            The tetrapartite synapse: extracellular matrix remodeling contributes to corticoaccumbens plasticity underlying drug addiction - PMC
         — https://pmc.ncbi.nlm.nih.gov/articles/PMC4589426/
- **[S13]** 
            The tetrapartite synapse: extracellular matrix remodeling contributes to corticoaccumbens plasticity underlying drug addiction - PMC
         — https://pmc.ncbi.nlm.nih.gov/articles/PMC4589426/
