## Supplementary material for "Understanding Substance Dependence: What Differentiates Addictive from Non-Addictive Drugs?": Folder for supplementary files: addictive-genes-claims-first100-046-slc6a8-gene-protein-is-implicated-in-addiction-e0fa60b93587.md

# addictive-genes-claims-first100-046-e0fa60b93587

**Original claim:** SLC6A8 gene/protein is implicated in addiction.

## Claim

SLC6A8 gene/protein is implicated in addiction. ([medlineplus.gov](https://medlineplus.gov/download/genetics/gene/slc6a8.pdf)) [S1]

## Verdict

Verdict: Likely False

Current evidence does **not** support SLC6A8 as an established addiction gene/protein. SLC6A8 encodes the creatine transporter (CRT1), and the best-supported biology links it to **creatine uptake, cellular/brain energy homeostasis, and cerebral creatine deficiency syndromes**, not to substance-use disorders themselves. ([medlineplus.gov](https://medlineplus.gov/download/genetics/gene/slc6a8.pdf)) [S2]

## Confidence

Medium. The negative conclusion is based on the **absence of direct human genetic or clinical addiction evidence** plus the presence of only **indirect mechanistic/preclinical hints**. Those hints are interesting, but they fall short of showing that SLC6A8 is meaningfully involved in addiction pathogenesis. ([ebi.ac.uk](https://www.ebi.ac.uk/gwas/genes/SLC6A8)) [S3]

## Supporting evidence

SLC6A8 is expressed in the brain, including midbrain and basal ganglia-related regions, and is broadly detected across brain regions rather than being absent from reward circuitry. The Human Protein Atlas classifies it as having **low regional specificity** and being **detected in all** surveyed brain regions. ([proteinatlas.org](https://www.proteinatlas.org/ENSG00000130821-SLC6A8/brain)) [S4]

Mechanistically, that matters because addiction-relevant circuits rely heavily on dopamine signaling and neuronal energetics. A 2019 mouse study deleted **Slc6a8 specifically in dopamine-transporter-expressing neurons** and found persistent hyperactivity, showing that the transporter can influence dopaminergic-neuron-related behavior. However, the study did **not** test drug reward, self-administration, relapse, withdrawal, or dependence phenotypes. ([pubmed.ncbi.nlm.nih.gov](https://pubmed.ncbi.nlm.nih.gov/31520365/)) [S5]

There is also broader indirect support for creatine-related biology touching addiction-relevant systems. A review on creatine and drug abuse noted that altered brain creatine levels have been observed in drug-exposed individuals and argued that brain energetics may matter in addiction biology, while explicitly stating that the significance of those changes and the role of creatine supplementation in drug abuse remained unresolved. ([pubmed.ncbi.nlm.nih.gov](https://pubmed.ncbi.nlm.nih.gov/21399936/)) [S6]

More recent systems-level work also points to drug-related changes in **creatine metabolism**, not necessarily SLC6A8 itself. For example, a 2024 spatial multiomic study of acute cocaine exposure reported that creatine was downregulated across reward-circuit regions, suggesting that cocaine can perturb energetic/metabolic pathways involving creatine. ([academic.oup.com](https://academic.oup.com/pnasnexus/article/doi/10.1093/pnasnexus/pgae458/7816872?utm_source=openai)) [S7]

Finally, SLC6A8 belongs to the SLC6 transporter family, several members of which are central to addiction biology. Reviews of SLC6 transporters highlight that **DAT/SLC6A3, NET/SLC6A2, and SERT/SLC6A4** regulate reward-related monoamine signaling and are direct targets of addictive drugs, whereas SLC6A8 transports creatine and is discussed mainly in relation to creatine deficiency and cognition. ([pmc.ncbi.nlm.nih.gov](https://pmc.ncbi.nlm.nih.gov/articles/PMC3602807/)) [S8]

## Contradicting or limiting evidence

The biggest limitation for the claim is that direct evidence is missing. Authoritative disease summaries describe SLC6A8 as the creatine transporter whose pathogenic variants cause **X-linked creatine deficiency / cerebral creatine deficiency syndrome**, with manifestations such as intellectual disability, behavioral problems, seizures, autism-like features, and muscle weakness—not addiction. ([medlineplus.gov](https://medlineplus.gov/download/genetics/gene/slc6a8.pdf)) [S9]

Likewise, transporter-focused brain-disease reviews list SLC6A8 under **cognitive deficits / cerebral creatine deficiency syndrome**, while addiction-related transporter associations in the same review are assigned to genes such as **SLC6A4** and **SLC18A2**, not SLC6A8. ([pmc.ncbi.nlm.nih.gov](https://pmc.ncbi.nlm.nih.gov/articles/PMC7193445/)) [S10]

Human genetics also does not currently support the addiction claim. The GWAS Catalog gene page for **SLC6A8** shows the framework for associations/studies/traits but no listed entries on the accessible page, which is consistent with a lack of established GWAS associations for addiction or related substance-use traits. ([ebi.ac.uk](https://www.ebi.ac.uk/gwas/genes/SLC6A8)) [S11]

That contrasts with addiction-GWAS reviews, which repeatedly identify other loci—such as **CNIH3, FAM53B, KAT2B, CREB1-adjacent signals,** and others—for opioid, stimulant, or general substance dependence, without highlighting SLC6A8 as a replicated finding. ([pmc.ncbi.nlm.nih.gov](https://pmc.ncbi.nlm.nih.gov/articles/PMC4996024/)) [S12]

Even the most favorable creatine-focused review is cautious: it frames creatine as a **possible** area of interest in drug abuse but explicitly says the role of creatine supplementation and altered brain creatine in drug abuse “has yet to be explored,” which is not evidence that SLC6A8 is already implicated. ([pubmed.ncbi.nlm.nih.gov](https://pubmed.ncbi.nlm.nih.gov/21399936/)) [S13]

## Mechanistic / biological rationale

Biologically, SLC6A8 is plausible as an **indirect modulator** of addiction-relevant neurobiology because it imports creatine, a molecule important for ATP buffering and cellular energy handling. MedlinePlus states that SLC6A8 transports creatine into cells and that creatine is needed for proper energy storage and use. ([medlineplus.gov](https://medlineplus.gov/download/genetics/gene/slc6a8.pdf)) [S14]

In the CNS, creatine biology appears relevant to neuronal function. A recent experimental paper even proposed creatine as a possible neurotransmitter-like signaling molecule and showed that loss of **Slc6a8** reduced synaptic-vesicle creatine and creatine uptake into synaptosomes. That supports a role in neural homeostasis and signaling. ([pmc.ncbi.nlm.nih.gov](https://pmc.ncbi.nlm.nih.gov/articles/PMC10735228/)) [S15]

Because addiction heavily involves mesolimbic dopamine circuits, and because drugs of abuse reinforce behavior largely through dopamine-related signaling in reward pathways, any protein that affects dopaminergic neuron energetics could, in theory, influence addiction vulnerability or expression. ([nida.nih.gov](https://nida.nih.gov/publications/drugs-brains-behavior-science-addiction/drugs-brain?utm_source=openai)) [S16]

But that is still an **inference**, not a demonstrated addiction link. The preclinical SLC6A8 evidence so far shows effects on **dopaminergic-neuron-associated behavior** and brain energetics, not on addiction-defining endpoints such as drug reinforcement, escalation, compulsive use, cue-induced seeking, withdrawal, or relapse. ([pubmed.ncbi.nlm.nih.gov](https://pubmed.ncbi.nlm.nih.gov/31520365/)) [S17]

So the mechanistic rationale is: **possible indirect relevance via neuronal energy homeostasis and dopamine-circuit physiology**. What is missing is the crucial next step—evidence that perturbing SLC6A8 actually changes addiction-like phenotypes in animals or addiction risk/outcomes in humans. ([pubmed.ncbi.nlm.nih.gov](https://pubmed.ncbi.nlm.nih.gov/31520365/)) [S18]

## Bottom line

SLC6A8 is **well supported** as a creatine transporter important for brain energy homeostasis and neurodevelopmental disease, but it is **not currently well supported as an addiction gene/protein**. The available literature provides only **indirect, mechanistically plausible hints**—for example, expression in brain reward-related regions, effects in dopaminergic neurons, and drug-related changes in creatine metabolism—without direct human genetic, clinical, or addiction-behavior evidence. On present evidence, the claim is best judged **Likely False**. ([medlineplus.gov](https://medlineplus.gov/download/genetics/gene/slc6a8.pdf)) [S19]

## Cited sources

- **[S1]** SLC6A8 gene — https://medlineplus.gov/download/genetics/gene/slc6a8.pdf
- **[S2]** SLC6A8 gene — https://medlineplus.gov/download/genetics/gene/slc6a8.pdf
- **[S3]** GWAS Catalog — https://www.ebi.ac.uk/gwas/genes/SLC6A8
- **[S4]** Brain tissue expression of SLC6A8 - Summary - The Human Protein Atlas — https://www.proteinatlas.org/ENSG00000130821-SLC6A8/brain
- **[S5]** Deletion of the Creatine Transporter (Slc6a8) in Dopaminergic Neurons Leads to Hyperactivity in Mice - PubMed — https://pubmed.ncbi.nlm.nih.gov/31520365/
- **[S6]** A potential role for creatine in drug abuse? - PubMed — https://pubmed.ncbi.nlm.nih.gov/21399936/
- **[S7]** Spatial multiomic insights into acute cocaine exposure | PNAS Nexus | Oxford Academic — https://academic.oup.com/pnasnexus/article/doi/10.1093/pnasnexus/pgae458/7816872?utm_source=openai
- **[S8]** 
            SLC6 Transporters: Structure, Function, Regulation, Disease Association and Therapeutics - PMC
         — https://pmc.ncbi.nlm.nih.gov/articles/PMC3602807/
- **[S9]** SLC6A8 gene — https://medlineplus.gov/download/genetics/gene/slc6a8.pdf
- **[S10]** 
            The solute carrier transporters and the brain: Physiological and pharmacological implications - PMC
         — https://pmc.ncbi.nlm.nih.gov/articles/PMC7193445/
- **[S11]** GWAS Catalog — https://www.ebi.ac.uk/gwas/genes/SLC6A8
- **[S12]** 
            A Review of Genome-Wide Association Studies of Stimulant and Opioid Use Disorders - PMC
         — https://pmc.ncbi.nlm.nih.gov/articles/PMC4996024/
- **[S13]** A potential role for creatine in drug abuse? - PubMed — https://pubmed.ncbi.nlm.nih.gov/21399936/
- **[S14]** SLC6A8 gene — https://medlineplus.gov/download/genetics/gene/slc6a8.pdf
- **[S15]** 
            Suggestion of creatine as a new neurotransmitter by approaches ranging from chemical analysis and biochemistry to electrophysiology - PMC
         — https://pmc.ncbi.nlm.nih.gov/articles/PMC10735228/
- **[S16]** Drugs, Brains, and Behavior: The Science of Addiction: Drugs and the Brain | NIDA — https://nida.nih.gov/publications/drugs-brains-behavior-science-addiction/drugs-brain?utm_source=openai
- **[S17]** Deletion of the Creatine Transporter (Slc6a8) in Dopaminergic Neurons Leads to Hyperactivity in Mice - PubMed — https://pubmed.ncbi.nlm.nih.gov/31520365/
- **[S18]** Deletion of the Creatine Transporter (Slc6a8) in Dopaminergic Neurons Leads to Hyperactivity in Mice - PubMed — https://pubmed.ncbi.nlm.nih.gov/31520365/
- **[S19]** SLC6A8 gene — https://medlineplus.gov/download/genetics/gene/slc6a8.pdf
