## Supplementary material for "Understanding Substance Dependence: What Differentiates Addictive from Non-Addictive Drugs?": Folder for supplementary files: addictive-genes-claims-first100-047-mylk-gene-protein-is-implicated-in-addiction-667bc3257807.md

# addictive-genes-claims-first100-047-667bc3257807

**Original claim:** MYLK gene/protein is implicated in addiction.

## Claim

**MYLK gene/protein is implicated in addiction.**

## Verdict

Verdict: Mixed Evidence

## Confidence

**Medium**

## Supporting evidence

MYLK encodes myosin light chain kinase (MLCK), a calcium/calmodulin-dependent kinase that phosphorylates myosin regulatory light chains and regulates actomyosin contractility. NCBI notes that it produces smooth-muscle and nonmuscle isoforms, and the Human Protein Atlas shows that MYLK is expressed in brain with low regional specificity and falls into a brain RNA-expression cluster annotated as **“White matter - Myelination (mainly)”**, indicating CNS expression but not a classic reward-circuit–specific pattern. ([ncbi.nlm.nih.gov](https://www.ncbi.nlm.nih.gov/gene/4638?utm_source=openai)) [S1]

There is at least some **preclinical addiction-related transcriptomic evidence**. In a mouse nucleus accumbens RNA-seq study using a methamphetamine regimen that produced locomotor sensitization, 2,171 mRNAs were differentially expressed after methamphetamine exposure, and the authors interpreted these changes as relevant to methamphetamine addiction biology. In the study’s miRNA–mRNA network table, **Mylk** appears among the downregulated genes linked to altered miR-204-3p signaling. This does not prove causality, but it does place Mylk within an addiction-relevant brain-region transcriptomic response to an abused drug. ([pmc.ncbi.nlm.nih.gov](https://pmc.ncbi.nlm.nih.gov/articles/PMC5116666/)) [S2]

There is also **mechanistic neuronal evidence** that MLCK can participate in processes often relevant to addiction neuroadaptation. In hippocampal neurons, BDNF increased phosphorylation of myosin light chain 2 and nonmuscle myosin II activity through a LYN-kinase/MLCK-dependent pathway; the study also confirmed this effect after BDNF injection into mouse dorsal hippocampus. Because addiction involves experience-dependent synaptic plasticity, this supports biological plausibility that MLCK-related signaling could matter in drug-induced neural remodeling, even though the paper was not an addiction study. ([pmc.ncbi.nlm.nih.gov](https://pmc.ncbi.nlm.nih.gov/articles/PMC9194867/?utm_source=openai)) [S3]

A separate brain study showed that **nonmuscle MYLK mediates microglial migration** induced by HIV Tat, including reduced migration of nmMYLK-knockout microglia toward Tat injected into hippocampus. This again is not addiction evidence per se, but it supports the idea that MYLK can influence neuroimmune processes inside the brain, and neuroimmune signaling is relevant to many substance-use disorders. ([pubmed.ncbi.nlm.nih.gov](https://pubmed.ncbi.nlm.nih.gov/23292072/?utm_source=openai)) [S4]

## Contradicting or limiting evidence

The main limitation is that the evidence above is **indirect and sparse**. The core functional annotations and disease associations for MYLK emphasize smooth/nonmuscle contractility, vascular permeability, inflammation, and certain Mendelian or cardiopulmonary phenotypes—not substance-use disorders. NCBI’s gene summary describes MYLK primarily in contractile biology, and MYLK-focused primary papers retrieved in this search center on acute lung injury/ARDS, asthma, vascular barrier dysfunction, and related inflammatory phenotypes rather than addiction. ([ncbi.nlm.nih.gov](https://www.ncbi.nlm.nih.gov/gene/4638?utm_source=openai)) [S5]

Large human addiction-genetics studies located here do **not** point to MYLK as a leading signal. For example, a gene/pathway analysis of smoking behaviors in 74,053 individuals reported 21 implicated genes and highlighted loci such as **IREB2**, **BDNF-AS**, and **SLC25A21**, plus strong nicotinic receptor pathway signals; MYLK was not one of the named findings in the main results. ([pmc.ncbi.nlm.nih.gov](https://pmc.ncbi.nlm.nih.gov/articles/PMC5777181/)) [S6]

Likewise, a recent **multiomic opioid-addiction** analysis integrating GWAS and postmortem dorsolateral prefrontal cortex omics prioritized pathways such as **BDNF**, **ERK**, and **Akt/PI3K**, and highlighted genes such as **BDNF, NTRK2, GSK3B, PIK3R1, OPRM1, SLC1A2, GABBR2,** and **KCNMA1**. MYLK was not among the study’s emphasized retained genes, conceptual model genes, or drug-target discussion. ([pmc.ncbi.nlm.nih.gov](https://pmc.ncbi.nlm.nih.gov/articles/PMC12119972/)) [S7]

So the present literature is not consistent with MYLK being an **established addiction gene** in the way that genes such as **OPRM1**, nicotinic receptor subunits, or alcohol-metabolism genes are. At most, the current evidence suggests MYLK may sit near downstream cellular processes that drugs can perturb. ([pmc.ncbi.nlm.nih.gov](https://pmc.ncbi.nlm.nih.gov/articles/PMC5777181/)) [S8]

## Mechanistic / biological rationale

Biologically, the claim is not implausible. MYLK/MLCK regulates cytoskeletal contraction by phosphorylating myosin light chains, with important roles in cell motility, barrier regulation, and cytoskeletal organization. Nonmuscle MYLK variants and isoforms have been studied especially in endothelial and inflammatory contexts. ([ncbi.nlm.nih.gov](https://www.ncbi.nlm.nih.gov/gene/4638?utm_source=openai)) [S9]

That biology could intersect addiction **indirectly** in several ways. First, cytoskeletal remodeling is part of synaptic plasticity, and the BDNF–LYN–MLCK–myosin II pathway in hippocampal neurons provides a plausible route by which MYLK-related signaling could influence structural or functional plasticity relevant to learning and reward. Second, MYLK’s role in microglial migration suggests it could affect neuroimmune responses, which are increasingly recognized in addiction and relapse biology. Third, its broader roles in endothelial permeability and cell migration suggest possible links to blood-brain barrier or inflammatory adaptations after drug exposure. These are mechanistic inferences from MYLK biology, not direct demonstrations that MYLK drives addictive behavior. ([pmc.ncbi.nlm.nih.gov](https://pmc.ncbi.nlm.nih.gov/articles/PMC9194867/?utm_source=openai)) [S10]

Crucially, there is a difference between:
1. a gene being **changed by drug exposure** or sitting in a downstream pathway, and  
2. a gene being **causally implicated in addiction susceptibility, maintenance, or relapse**.

For MYLK, the evidence found here is much stronger for the first category than the second. The methamphetamine nucleus accumbens study shows association with a drug-induced transcriptional state, but not necessity or sufficiency for addiction-like behavior. ([pmc.ncbi.nlm.nih.gov](https://pmc.ncbi.nlm.nih.gov/articles/PMC5116666/)) [S11]

## Bottom line

The claim is **not clearly wrong**, but it is **too strong if stated without qualification**.

A more accurate version would be:

**“MYLK has limited, mainly indirect preclinical evidence connecting it to addiction-related neurobiology, but it is not currently a well-established addiction gene/protein.”** ([pmc.ncbi.nlm.nih.gov](https://pmc.ncbi.nlm.nih.gov/articles/PMC5116666/)) [S12]

So overall, the evidence is **mixed**: there is some brain expression, mechanistic plausibility, and at least one addiction-relevant animal transcriptomic signal, but there is **little direct, replicated human genetic or causal experimental evidence** showing that MYLK itself is a major driver of addiction.

## Cited sources

- **[S1]** MYLK myosin light chain kinase [Homo sapiens (human)] - Gene - NCBI — https://www.ncbi.nlm.nih.gov/gene/4638?utm_source=openai
- **[S2]** 
            mRNA changes in nucleus accumbens related to methamphetamine addiction in mice - PMC
         — https://pmc.ncbi.nlm.nih.gov/articles/PMC5116666/
- **[S3]** Brain-derived neurotrophic factor regulates LYN kinase–mediated myosin light chain kinase activation to modulate nonmuscle myosin II activity in hippocampal neurons - PMC — https://pmc.ncbi.nlm.nih.gov/articles/PMC9194867/?utm_source=openai
- **[S4]** Nonmuscle myosin light-chain kinase mediates microglial migration induced by HIV Tat: involvement of β1 integrins - PubMed — https://pubmed.ncbi.nlm.nih.gov/23292072/?utm_source=openai
- **[S5]** MYLK myosin light chain kinase [Homo sapiens (human)] - Gene - NCBI — https://www.ncbi.nlm.nih.gov/gene/4638?utm_source=openai
- **[S6]** 
            Pathways to smoking behaviors: biological insights from the Tobacco and Genetics Consortium meta-analysis - PMC
         — https://pmc.ncbi.nlm.nih.gov/articles/PMC5777181/
- **[S7]** 
            Multiomic Network Analysis Identifies Dysregulated Neurobiological Pathways in Opioid Addiction - PMC
         — https://pmc.ncbi.nlm.nih.gov/articles/PMC12119972/
- **[S8]** 
            Pathways to smoking behaviors: biological insights from the Tobacco and Genetics Consortium meta-analysis - PMC
         — https://pmc.ncbi.nlm.nih.gov/articles/PMC5777181/
- **[S9]** MYLK myosin light chain kinase [Homo sapiens (human)] - Gene - NCBI — https://www.ncbi.nlm.nih.gov/gene/4638?utm_source=openai
- **[S10]** Brain-derived neurotrophic factor regulates LYN kinase–mediated myosin light chain kinase activation to modulate nonmuscle myosin II activity in hippocampal neurons - PMC — https://pmc.ncbi.nlm.nih.gov/articles/PMC9194867/?utm_source=openai
- **[S11]** 
            mRNA changes in nucleus accumbens related to methamphetamine addiction in mice - PMC
         — https://pmc.ncbi.nlm.nih.gov/articles/PMC5116666/
- **[S12]** 
            mRNA changes in nucleus accumbens related to methamphetamine addiction in mice - PMC
         — https://pmc.ncbi.nlm.nih.gov/articles/PMC5116666/
