## Supplementary material for "Understanding Substance Dependence: What Differentiates Addictive from Non-Addictive Drugs?": Folder for supplementary files: addictive-genes-claims-first100-048-asph-gene-protein-is-implicated-in-addiction-973937f2b4bf.md

# addictive-genes-claims-first100-048-973937f2b4bf

**Original claim:** ASPH gene/protein is implicated in addiction.

## Claim

**“ASPH gene/protein is implicated in addiction.”**

## Verdict

Verdict: Mixed Evidence

## Confidence

**Medium**

## Supporting evidence

There is **some evidence linking ASPH to alcohol-related brain changes**, particularly in models and tissues relevant to alcohol use disorder (AUD), but this evidence is mostly **indirect** and does not firmly establish ASPH as an addiction gene in the usual sense of affecting reward, reinforcement, craving, relapse, or substance-seeking behavior. ([pmc.ncbi.nlm.nih.gov](https://pmc.ncbi.nlm.nih.gov/articles/PMC12620835/)) [S1]

ASPH encodes **aspartate beta-hydroxylase**, a protein-coding gene with multiple isoforms; NCBI notes it is thought to have an important role in **calcium homeostasis**, and the longest isoforms hydroxylate Asp/Asn residues in EGF-like domains of target proteins. ASPH biology is therefore relevant to signaling systems that could, in principle, affect neural function. ([ncbi.nlm.nih.gov](https://www.ncbi.nlm.nih.gov/gene/444?utm_source=openai)) [S2]

In the nervous system, ASPH has been connected to **Notch signaling**. In a rat developmental model, experimental suppression of AAH/ASPH in the cerebellum impaired motor performance and reduced expression of **Notch-1, Jagged-1, and HES-1**, whereas overexpression had opposite effects. The authors concluded that AAH/ASPH is an important mediator of cerebellar development and function. This does not show addiction, but it supports a bona fide neurobiological role for ASPH in brain signaling. ([pubmed.ncbi.nlm.nih.gov](https://pubmed.ncbi.nlm.nih.gov/21050474/)) [S3]

More specifically for alcohol, a rat study found that chronic-plus-binge ethanol exposure caused **progressive white-matter atrophy** and was associated with reduced **ASPH, Jagged1, HES-1, and HIF-1α** expression in frontal white matter. The paper interprets ASPH suppression as part of ethanol-related white-matter injury via Notch pathway impairment. ([pubmed.ncbi.nlm.nih.gov](https://pubmed.ncbi.nlm.nih.gov/29204305/?utm_source=openai)) [S4]

Human postmortem evidence also exists. In cerebellar tissue from men with AUD, investigators reported **reduced ASPH and HES1 mRNA** relative to controls and argued that AUD-related cerebellar pathology involves downregulated ASPH/Notch signaling and oligodendrocyte-white-matter dysfunction. This is the strongest direct human evidence tying ASPH to a substance-use disorder context. ([pmc.ncbi.nlm.nih.gov](https://pmc.ncbi.nlm.nih.gov/articles/PMC12620835/)) [S5]

A secondary integrative resource also places ASPH in a broad **“substance-related disorder”** gene set derived from GWASdb associations. That is supportive only at a very low level, because it is a downstream aggregation/database signal rather than a primary replicated study. ([maayanlab.cloud](https://maayanlab.cloud/Harmonizome/gene_set/substance-related%2Bdisorder/GWASdb%2BSNP-Disease%2BAssociations)) [S6]

## Contradicting or limiting evidence

The biggest limitation is that the available ASPH literature is largely about **alcohol-induced neurodevelopmental injury or alcohol-related white-matter degeneration**, not about the core behavioral neurobiology of addiction. The fetal alcohol and white-matter studies ask whether ethanol alters ASPH expression and downstream signaling; they do **not** test whether ASPH variation or manipulation changes alcohol self-administration, drug reward, compulsive use, withdrawal severity, cue reactivity, or relapse. ([pubmed.ncbi.nlm.nih.gov](https://pubmed.ncbi.nlm.nih.gov/19393862/?utm_source=openai)) [S7]

The main human AUD paper is also **small and cross-sectional**: only **6 AUD brains and 6 controls**, males only, using postmortem tissue. The authors themselves note these limitations and that the work cannot resolve earlier disease stages or reversibility well. That makes the human signal interesting but far from definitive. ([pmc.ncbi.nlm.nih.gov](https://pmc.ncbi.nlm.nih.gov/articles/PMC12620835/)) [S8]

By contrast, large modern human genetics studies of substance use disorders have highlighted other loci and pathways. A large trans-ancestral GWAS of alcohol dependence found genome-wide significant effects in **ADH1B**, not ASPH. A large multi-ancestral GWAS of DSM-defined nicotine dependence replicated the well-known **CHRNA5** locus. Recent reviews of generalized SUD liability emphasize loci and pathways such as **DRD2, OPRM1, FURIN**, dopaminergic regulation, neuronal development, and immune processes. ([pubmed.ncbi.nlm.nih.gov](https://pubmed.ncbi.nlm.nih.gov/30482948/)) [S9]

Consistent with that, searches within major recent addiction-genetics review/GWAS sources did **not** find ASPH mentioned. This does not prove ASPH is irrelevant, but it strongly suggests ASPH is **not currently an established or replicated addiction gene** in the mainstream human genetics literature. ([nature.com](https://www.nature.com/articles/s41380-024-02547-z)) [S10]

So the most accurate reading is: ASPH is implicated in **alcohol-associated brain pathology**, and perhaps in neural processes that could intersect with addiction, but evidence for a **direct addiction role** is sparse. ([pmc.ncbi.nlm.nih.gov](https://pmc.ncbi.nlm.nih.gov/articles/PMC12620835/)) [S11]

## Mechanistic / biological rationale

A plausible mechanism exists. ASPH hydroxylates EGF-like domains and promotes **Notch-Jagged-HES/HEY** signaling; NCBI also notes roles related to calcium homeostasis. In the adult brain, Notch signaling is not just developmental: it has recognized roles in **migration, morphology, synaptic plasticity, and neuronal survival**. Those processes are highly relevant to long-term substance-induced neuroadaptation. ([ncbi.nlm.nih.gov](https://www.ncbi.nlm.nih.gov/gtr/genes/444/?utm_source=openai)) [S12]

There is also some addiction-adjacent support for the pathway. A review on post-developmental Notch functions includes a section on **reward and addiction**, and a Drosophila study reported that alcohol activates **Scabrous/Notch** signaling to influence alcohol-associated memories. That means an ASPH→Notch connection is biologically plausible as part of alcohol-related neural plasticity. ([mdpi.com](https://www.mdpi.com/2218-273X/10/7/985?utm_source=openai)) [S13]

However, this remains a **mechanistic inference**, not a demonstrated ASPH-addiction pathway in humans. The current evidence better supports ASPH as a mediator of **ethanol-responsive neural homeostasis / white-matter biology / developmental signaling** than as a proven determinant of addiction vulnerability or addictive behavior itself. ([pubmed.ncbi.nlm.nih.gov](https://pubmed.ncbi.nlm.nih.gov/21050474/)) [S14]

## Bottom line

ASPH is **not well established as an addiction gene/protein** by current human genetics or behavioral-neuroscience standards. Still, it is **not unsupported either**: animal and postmortem human studies indicate that ASPH is altered in alcohol-exposed brain tissue and may participate in **AUD-related white-matter and Notch-signaling abnormalities**. Overall, the claim is best judged **partly supported but too broad as stated**. A more accurate claim would be: **“ASPH is implicated in alcohol-related brain pathology, with indirect but not yet strong direct evidence for a role in addiction.”** ([pmc.ncbi.nlm.nih.gov](https://pmc.ncbi.nlm.nih.gov/articles/PMC12620835/)) [S15]

## Cited sources

- **[S1]** 
            Molecular and biochemical pathologies in human alcohol-related cerebellar white matter degeneration - PMC
         — https://pmc.ncbi.nlm.nih.gov/articles/PMC12620835/
- **[S2]** ASPH aspartate beta-hydroxylase [Homo sapiens (human)] - Gene - NCBI — https://www.ncbi.nlm.nih.gov/gene/444?utm_source=openai
- **[S3]** Role of aspartyl-(asparaginyl)-β-hydroxylase mediated notch signaling in cerebellar development and function - PubMed — https://pubmed.ncbi.nlm.nih.gov/21050474/
- **[S4]** Ethanol-Induced White Matter Atrophy Is Associated with Impaired Expression of Aspartyl-Asparaginyl- β-Hydroxylase (ASPH) and Notch Signaling in an Experimental Rat Model - PubMed — https://pubmed.ncbi.nlm.nih.gov/29204305/?utm_source=openai
- **[S5]** 
            Molecular and biochemical pathologies in human alcohol-related cerebellar white matter degeneration - PMC
         — https://pmc.ncbi.nlm.nih.gov/articles/PMC12620835/
- **[S6]** Gene Set - substance-related disorder — https://maayanlab.cloud/Harmonizome/gene_set/substance-related%2Bdisorder/GWASdb%2BSNP-Disease%2BAssociations
- **[S7]** Ethanol inhibition of aspartyl-asparaginyl-beta-hydroxylase in fetal alcohol spectrum disorder: potential link to the impairments in central nervous system neuronal migration - PubMed — https://pubmed.ncbi.nlm.nih.gov/19393862/?utm_source=openai
- **[S8]** 
            Molecular and biochemical pathologies in human alcohol-related cerebellar white matter degeneration - PMC
         — https://pmc.ncbi.nlm.nih.gov/articles/PMC12620835/
- **[S9]** Transancestral GWAS of alcohol dependence reveals common genetic underpinnings with psychiatric disorders - PubMed — https://pubmed.ncbi.nlm.nih.gov/30482948/
- **[S10]** The genetic landscape of substance use disorders | Molecular Psychiatry — https://www.nature.com/articles/s41380-024-02547-z
- **[S11]** 
            Molecular and biochemical pathologies in human alcohol-related cerebellar white matter degeneration - PMC
         — https://pmc.ncbi.nlm.nih.gov/articles/PMC12620835/
- **[S12]** ASPH aspartate beta-hydroxylase - NIH Genetic Testing Registry (GTR) - NCBI — https://www.ncbi.nlm.nih.gov/gtr/genes/444/?utm_source=openai
- **[S13]** Post-Developmental Roles of Notch Signaling in the Nervous System — https://www.mdpi.com/2218-273X/10/7/985?utm_source=openai
- **[S14]** Role of aspartyl-(asparaginyl)-β-hydroxylase mediated notch signaling in cerebellar development and function - PubMed — https://pubmed.ncbi.nlm.nih.gov/21050474/
- **[S15]** 
            Molecular and biochemical pathologies in human alcohol-related cerebellar white matter degeneration - PMC
         — https://pmc.ncbi.nlm.nih.gov/articles/PMC12620835/
