## Supplementary material for "Understanding Substance Dependence: What Differentiates Addictive from Non-Addictive Drugs?": Folder for supplementary files: addictive-genes-claims-first100-049-cmtm5-gene-protein-is-implicated-in-addiction-f5650d8e1e1e.md

# addictive-genes-claims-first100-049-f5650d8e1e1e

**Original claim:** CMTM5 gene/protein is implicated in addiction.

## Claim

CMTM5 gene/protein is implicated in addiction.

## Verdict

Verdict: No Evidence

I did not find direct peer-reviewed evidence that **CMTM5** is implicated in addiction, substance use disorder, craving, withdrawal, relapse, or drug self-administration. The literature retrieved for CMTM5 is dominated by **CNS myelin/oligodendrocyte biology** and **cancer biology**, not addiction research. In targeted PubMed searches using addiction-related terms, the CMTM5 papers surfaced were about oligodendrocyte/myelin function, demyelination, and tumor suppression rather than addiction phenotypes. ([pubmed.ncbi.nlm.nih.gov](https://pubmed.ncbi.nlm.nih.gov/35274615/?utm_source=openai)) [S1]

## Confidence

Medium

Confidence is **medium** because CMTM5 is a relatively understudied gene, so small omics signals could exist in supplementary datasets that are not easy to retrieve by title/abstract search alone. But based on the accessible scholarly literature I found, there is **no direct human genetic, human clinical, postmortem, or animal behavioral evidence** tying CMTM5 specifically to addiction. ([pubmed.ncbi.nlm.nih.gov](https://pubmed.ncbi.nlm.nih.gov/35274615/?utm_source=openai)) [S2]

## Supporting evidence

The strongest evidence *potentially supportive of plausibility* is indirect: **CMTM5 is a bona fide CNS myelin/oligodendrocyte protein**. In the eLife mouse study, CMTM5 expression was highly enriched in mature oligodendrocytes and CNS myelin, and oligodendrocyte-specific loss of Cmtm5 caused a progressive axonopathy, indicating a role in maintaining axonal integrity rather than in initial myelin formation. Human Protein Atlas data likewise place CMTM5 in an oligodendrocyte/myelination expression cluster. ([pmc.ncbi.nlm.nih.gov](https://pmc.ncbi.nlm.nih.gov/articles/PMC8916772/)) [S3]

There is also broader addiction literature showing that **myelin and oligodendrocyte biology can matter for reward and substance-use phenotypes**. A Nature study reported that myelin plasticity in the ventral tegmental area is required for opioid reward in mice, and human neuroimaging work in cocaine and heroin use disorders found widespread white-matter abnormalities that correlated with craving and duration of use. These findings do **not** implicate CMTM5 specifically, but they show that a myelin-associated gene could, in principle, be relevant to addiction circuitry. ([pubmed.ncbi.nlm.nih.gov](https://pubmed.ncbi.nlm.nih.gov/38839962/)) [S4]

So, if one wants to build the best case *for* the claim, it would be this: CMTM5 participates in neural homeostasis at the level of oligodendrocytes/myelin, and myelin plasticity can influence reward-related circuitry. That is a **biological plausibility argument**, not direct evidence of addiction involvement. ([pmc.ncbi.nlm.nih.gov](https://pmc.ncbi.nlm.nih.gov/articles/PMC8916772/)) [S5]

## Contradicting or limiting evidence

A major limitation for the claim is the **absence of direct CMTM5-addiction literature**. When CMTM5 was searched together with addiction-related terms, the returned PubMed records were not addiction studies; instead they were on myelin biology, cancer, glioma, and autoimmune disease. That pattern argues that CMTM5 is **not currently an established addiction candidate gene** in the peer-reviewed literature. ([pubmed.ncbi.nlm.nih.gov](https://pubmed.ncbi.nlm.nih.gov/35274615/?utm_source=openai)) [S6]

The available functional work on CMTM5 also points elsewhere. The best mechanistic studies show roles in **axonal integrity**, **oligodendrocyte-lineage expression**, and **demyelinating disease contexts** such as multiple sclerosis lesions, but they do not test addiction-relevant endpoints such as reinforcement, conditioned place preference for drugs, self-administration, escalation, withdrawal, or relapse. ([pmc.ncbi.nlm.nih.gov](https://pmc.ncbi.nlm.nih.gov/articles/PMC8916772/)) [S7]

It is also important not to over-interpret the broader myelin/addiction literature. White-matter abnormalities in substance use disorder are **nonspecific** and do not identify CMTM5 as a causal gene. Likewise, the opioid-reward myelin-plasticity study implicates oligodendrogenesis in reward learning, but it did not examine CMTM5 expression or manipulate Cmtm5. ([pubmed.ncbi.nlm.nih.gov](https://pubmed.ncbi.nlm.nih.gov/38839962/)) [S8]

Some CMTM5 papers report effects on signaling pathways such as **EGFR/PI3K/AKT**, but those data come from cancer systems, not addiction models or neural circuit studies. Such findings are too indirect to count as evidence that CMTM5 is implicated in addiction. ([pubmed.ncbi.nlm.nih.gov](https://pubmed.ncbi.nlm.nih.gov/29213215/?utm_source=openai)) [S9]

## Mechanistic / biological rationale

CMTM5 belongs to the CKLF-like MARVEL transmembrane family and encodes a small multi-pass membrane protein. Early work showed broad normal-tissue expression and emphasized tumor-suppressor roles in carcinoma. More recent neuroscience work substantially sharpened its CNS assignment: CMTM5 is enriched in mature oligodendrocytes and CNS myelin, and loss of Cmtm5 compromises long-term axonal integrity. ([pubmed.ncbi.nlm.nih.gov](https://pubmed.ncbi.nlm.nih.gov/17908965/)) [S10]

That biology gives a **reasonable but indirect** route by which CMTM5 *could* matter to addiction. Addiction involves durable circuit adaptations, and myelin plasticity can alter conduction timing, network synchrony, and dopaminergic reward-circuit function. If a myelin protein helps maintain axon-glia support, then altered CMTM5 could hypothetically influence reward learning or vulnerability to substance-induced circuit remodeling. ([pubmed.ncbi.nlm.nih.gov](https://pubmed.ncbi.nlm.nih.gov/38839962/)) [S11]

However, the currently available CMTM5 data do **not** bridge that hypothesis to addiction. In fact, the mouse and MS-related studies suggest CMTM5 is more tightly linked to **axonal support/homeostasis** than to classic neurotransmitter receptor signaling or reward behavior, and one MS-focused study found that Cmtm5 knockdown did **not** alter oligodendroglial ER-stress responses in vitro. So the mechanistic rationale is plausible only at a high level and remains untested for addiction specifically. ([pubmed.ncbi.nlm.nih.gov](https://pubmed.ncbi.nlm.nih.gov/37626895/)) [S12]

## Bottom line

At present, **CMTM5 is supported as a myelin/oligodendrocyte/axonal-integrity gene, not as an addiction gene**. The most defensible assessment is that the claim is **unsubstantiated**: there is some **indirect biological plausibility** through myelin biology, but I found **no direct evidence** that CMTM5 is implicated in addiction itself. ([pmc.ncbi.nlm.nih.gov](https://pmc.ncbi.nlm.nih.gov/articles/PMC8916772/)) [S13]

## Cited sources

- **[S1]** Progressive axonopathy when oligodendrocytes lack the myelin protein CMTM5 - PubMed — https://pubmed.ncbi.nlm.nih.gov/35274615/?utm_source=openai
- **[S2]** Progressive axonopathy when oligodendrocytes lack the myelin protein CMTM5 - PubMed — https://pubmed.ncbi.nlm.nih.gov/35274615/?utm_source=openai
- **[S3]** 
            Progressive axonopathy when oligodendrocytes lack the myelin protein CMTM5 - PMC
         — https://pmc.ncbi.nlm.nih.gov/articles/PMC8916772/
- **[S4]** Myelin plasticity in the ventral tegmental area is required for opioid reward - PubMed — https://pubmed.ncbi.nlm.nih.gov/38839962/
- **[S5]** 
            Progressive axonopathy when oligodendrocytes lack the myelin protein CMTM5 - PMC
         — https://pmc.ncbi.nlm.nih.gov/articles/PMC8916772/
- **[S6]** Progressive axonopathy when oligodendrocytes lack the myelin protein CMTM5 - PubMed — https://pubmed.ncbi.nlm.nih.gov/35274615/?utm_source=openai
- **[S7]** 
            Progressive axonopathy when oligodendrocytes lack the myelin protein CMTM5 - PMC
         — https://pmc.ncbi.nlm.nih.gov/articles/PMC8916772/
- **[S8]** Myelin plasticity in the ventral tegmental area is required for opioid reward - PubMed — https://pubmed.ncbi.nlm.nih.gov/38839962/
- **[S9]** CMTM5 is downregulated and suppresses tumour growth in hepatocellular carcinoma through regulating PI3K-AKT signalling - PubMed — https://pubmed.ncbi.nlm.nih.gov/29213215/?utm_source=openai
- **[S10]** CMTM5 exhibits tumor suppressor activities and is frequently silenced by methylation in carcinoma cell lines - PubMed — https://pubmed.ncbi.nlm.nih.gov/17908965/
- **[S11]** Myelin plasticity in the ventral tegmental area is required for opioid reward - PubMed — https://pubmed.ncbi.nlm.nih.gov/38839962/
- **[S12]** Loss of the Novel Myelin Protein CMTM5 in Multiple Sclerosis Lesions and Its Involvement in Oligodendroglial Stress Responses - PubMed — https://pubmed.ncbi.nlm.nih.gov/37626895/
- **[S13]** 
            Progressive axonopathy when oligodendrocytes lack the myelin protein CMTM5 - PMC
         — https://pmc.ncbi.nlm.nih.gov/articles/PMC8916772/
