## Supplementary material for "Understanding Substance Dependence: What Differentiates Addictive from Non-Addictive Drugs?": Folder for supplementary files: addictive-genes-claims-first100-050-adcyap1r1-gene-protein-is-implicated-in-addiction-92dedd4b83b8.md

# addictive-genes-claims-first100-050-92dedd4b83b8

**Original claim:** ADCYAP1R1 gene/protein is implicated in addiction.

## Claim

ADCYAP1R1 gene/protein is implicated in addiction.

## Verdict

Verdict: Likely True

ADCYAP1R1 encodes the PAC1 receptor, and the strongest available evidence does place this receptor in addiction-relevant biology—particularly excessive alcohol intake, alcohol seeking, withdrawal-associated negative affect, and stress-related relapse circuitry in rodents. Human evidence exists, but it is much weaker than the animal literature and is not yet the kind of large-scale, replicated genetic signal that would make ADCYAP1R1 a firmly established human addiction gene. ([pubmed.ncbi.nlm.nih.gov](https://pubmed.ncbi.nlm.nih.gov/31284862/)) [S1]

## Confidence

Medium

Confidence is **medium** because multiple independent rodent studies support PAC1/ADCYAP1R1 involvement in addiction-related behaviors, especially for alcohol, but human evidence is limited mostly to small candidate-gene work rather than robust replication in large genome-wide studies. ([pubmed.ncbi.nlm.nih.gov](https://pubmed.ncbi.nlm.nih.gov/28652748/?utm_source=openai)) [S2]

## Supporting evidence

The most direct support comes from **alcohol models**. In alcohol-preferring Scr:sP rats, central PAC1 blockade with PACAP(6-38) reduced ethanol intake, motivation, and alcohol-seeking behavior, and these effects were selective relative to inactive lever pressing and sweet reinforcers. In the same line, PAC1R-positive cells were increased in the nucleus accumbens core after alcohol drinking, and both intra-core PAC1 antagonism and PAC1R knockdown reduced excessive alcohol drinking. ([pubmed.ncbi.nlm.nih.gov](https://pubmed.ncbi.nlm.nih.gov/35460713/)) [S3]

A complementary study in the **nucleus accumbens shell** found the opposite direction of manipulation produced the opposite phenotype: shRNA knockdown of PAC1R in shell increased ethanol self-administration, ethanol preference, and motivation to drink, without a similar rise in saccharin self-administration. That suggests PAC1 signaling is not just correlated with alcohol behavior, but functionally regulates it in a circuit-dependent way. ([pubmed.ncbi.nlm.nih.gov](https://pubmed.ncbi.nlm.nih.gov/34924973/)) [S4]

Evidence also links the PACAP/PAC1 system to the **dependence/withdrawal side** of alcohol addiction. In ethanol-dependent male rats, chronic intermittent ethanol exposure increased PACAP immunoreactivity in the BNST, and intra-BNST PACAP(6-38) blocked excessive drinking and reversed withdrawal-associated anxiety-like behavior, while sparing basal drinking in nondependent controls. This pattern fits a role in the “dark side” of addiction—negative reinforcement during withdrawal—rather than only acute reward. ([pubmed.ncbi.nlm.nih.gov](https://pubmed.ncbi.nlm.nih.gov/33191400/)) [S5]

For **human data**, a 2017 study of 491 women aged 18–28 reported that an intronic ADCYAP1R1 polymorphism (rs2302475) was significantly associated with problematic alcohol use measured by AUDIT, with supporting haplotype evidence. That is not definitive proof, but it is direct human genetic support for the claim. ([pubmed.ncbi.nlm.nih.gov](https://pubmed.ncbi.nlm.nih.gov/28652748/?utm_source=openai)) [S6]

There is also **cocaine-relapse evidence**, although it is less specifically tied to ADCYAP1R1 than the alcohol work. In a rat cocaine self-administration model, cocaine increased BNST PACAP transcript levels, intra-BNST PACAP(6-38) prevented footshock-induced reinstatement, and intra-BNST PACAP was sufficient to reinstate cocaine seeking without footshock. A newer rat study further showed that PACAP signaling in the nucleus accumbens core can suppress D1-like agonist-primed reinstatement of cocaine seeking. Together, these studies place the PACAP/PAC1 axis in relapse circuitry, though the direction of effect depends on brain region and paradigm. ([pubmed.ncbi.nlm.nih.gov](https://pubmed.ncbi.nlm.nih.gov/28656976/)) [S7]

For **opioids**, PAC1-deficient mice showed a dramatic increase in physical withdrawal signs after naloxone-precipitated morphine withdrawal. This is addiction-relevant, although notably the same study found no change in morphine reward or locomotor sensitization, so the receptor seems more involved in dependence/withdrawal than in every dimension of opioid reinforcement. ([pubmed.ncbi.nlm.nih.gov](https://pubmed.ncbi.nlm.nih.gov/12573539/)) [S8]

## Contradicting or limiting evidence

The main limitation is that the **best evidence is preclinical**, not clinical. The human genetic signal for ADCYAP1R1 comes from a relatively small, sex-limited candidate-gene study, whereas large GWAS of alcohol dependence have most robustly highlighted loci such as **ADH1B** and a small number of other regions. So, by inference from current large-scale human genetics, ADCYAP1R1 is **not yet a strongly established, replicated genome-wide addiction locus**. ([pubmed.ncbi.nlm.nih.gov](https://pubmed.ncbi.nlm.nih.gov/28652748/?utm_source=openai)) [S9]

A second limitation is that the phenotype appears **domain-specific rather than universal**. In the opioid study, PAC1 deficiency changed withdrawal severity but not morphine reward or sensitization. That means “implicated in addiction” is more accurate than “a core universal addiction gene/protein across all drugs and stages.” ([pubmed.ncbi.nlm.nih.gov](https://pubmed.ncbi.nlm.nih.gov/12573539/)) [S10]

A third limitation is **strong circuit specificity and sometimes opposite effects depending on brain region**. In alcohol models, reducing PAC1R signaling in nucleus accumbens core suppresses excessive drinking, whereas reducing PAC1R in shell increases drinking. In cocaine models, BNST PACAP/PAC1 signaling promotes stress-induced reinstatement, while accumbens PACAP signaling can suppress D1-driven reinstatement. So the literature supports involvement, but not a simple “more PAC1 equals more addiction” rule. ([pubmed.ncbi.nlm.nih.gov](https://pubmed.ncbi.nlm.nih.gov/35460713/)) [S11]

Finally, not every study isolates **ADCYAP1R1 alone** with perfect specificity: some experiments manipulate PACAP ligand levels or use pharmacologic tools within the broader PACAP receptor system. That still supports pathway involvement, but it slightly weakens the specificity of the claim when compared with the strongest receptor-knockdown data in alcohol-preferring rats. ([pubmed.ncbi.nlm.nih.gov](https://pubmed.ncbi.nlm.nih.gov/34924973/)) [S12]

## Mechanistic / biological rationale

ADCYAP1R1 encodes **PAC1**, a class B GPCR and the principal high-affinity receptor for PACAP. PAC1 is widely expressed in the nervous system, and receptor isoforms can engage multiple signaling modes, including G-protein and β-arrestin-linked pathways. Reviews describe maladaptive PACAP/PAC1 signaling as relevant to stress-related psychopathology, and PAC1 has been discussed as a therapeutic target in brain disorders. ([pubmed.ncbi.nlm.nih.gov](https://pubmed.ncbi.nlm.nih.gov/31284862/)) [S13]

At the cellular level, PAC1 activation can directly increase neuronal excitability. In dentate gyrus granule cells, PACAP increased excitability through PAC1-dependent **MEK/ERK** and endosomal signaling, with dependence on persistent sodium current rather than a simple cAMP-only mechanism in that preparation. This gives a plausible biophysical route by which PAC1 signaling could reshape activity in reward-, stress-, and relapse-related circuits. ([pubmed.ncbi.nlm.nih.gov](https://pubmed.ncbi.nlm.nih.gov/32186931/)) [S14]

That mechanism maps well onto addiction biology. The nucleus accumbens and BNST are key nodes for reinforcement, negative affect, and relapse. Alcohol exposure increased PAC1R in accumbens core in alcohol-preferring rats, and chronic ethanol or cocaine paradigms recruited PACAP/PAC1-related signaling in BNST/accumbens circuits that changed seeking, intake, or reinstatement behavior. The overall picture is that ADCYAP1R1 is most plausibly involved in **stress-coupled neuroadaptation, withdrawal/negative reinforcement, and relapse vulnerability**, with especially consistent evidence in alcohol models. ([pubmed.ncbi.nlm.nih.gov](https://pubmed.ncbi.nlm.nih.gov/35460713/)) [S15]

## Bottom line

Yes—**ADCYAP1R1/PAC1 is plausibly implicated in addiction**, especially in **alcohol-use phenotypes** and in **stress/withdrawal/relapse-related circuitry**. The claim is best supported by rodent functional studies and modestly supported by a human candidate-gene association, but it is **not yet a top-tier, broadly replicated human addiction gene**. So the fairest summary is: **likely true, but strongest for preclinical alcohol and relapse biology, with important limits on generalization.** ([pubmed.ncbi.nlm.nih.gov](https://pubmed.ncbi.nlm.nih.gov/28652748/?utm_source=openai)) [S16]

## Cited sources

- **[S1]** Targeting the PAC1 Receptor for Neurological and Metabolic Disorders - PubMed — https://pubmed.ncbi.nlm.nih.gov/31284862/
- **[S2]** PAC1 receptor (ADCYAP1R1) genotype and problematic alcohol use in a sample of young women - PubMed — https://pubmed.ncbi.nlm.nih.gov/28652748/?utm_source=openai
- **[S3]** Pituitary adenylate cyclase-activating polypeptide type 1 receptor within the nucleus accumbens core mediates excessive alcohol drinking in alcohol-preferring rats - PubMed — https://pubmed.ncbi.nlm.nih.gov/35460713/
- **[S4]** Viral-Mediated Knockdown of Nucleus Accumbens Shell PAC1 Receptor Promotes Excessive Alcohol Drinking in Alcohol-Preferring Rats - PubMed — https://pubmed.ncbi.nlm.nih.gov/34924973/
- **[S5]** Pituitary adenylate cyclase-activating polypeptide (PACAP) modulates dependence-induced alcohol drinking and anxiety-like behavior in male rats - PubMed — https://pubmed.ncbi.nlm.nih.gov/33191400/
- **[S6]** PAC1 receptor (ADCYAP1R1) genotype and problematic alcohol use in a sample of young women - PubMed — https://pubmed.ncbi.nlm.nih.gov/28652748/?utm_source=openai
- **[S7]** Pituitary Adenylate Cyclase-Activating Peptide in the Bed Nucleus of the Stria Terminalis Mediates Stress-Induced Reinstatement of Cocaine Seeking in Rats - PubMed — https://pubmed.ncbi.nlm.nih.gov/28656976/
- **[S8]** Morphine withdrawal is modified in pituitary adenylate cyclase-activating polypeptide type I-receptor-deficient mice - PubMed — https://pubmed.ncbi.nlm.nih.gov/12573539/
- **[S9]** PAC1 receptor (ADCYAP1R1) genotype and problematic alcohol use in a sample of young women - PubMed — https://pubmed.ncbi.nlm.nih.gov/28652748/?utm_source=openai
- **[S10]** Morphine withdrawal is modified in pituitary adenylate cyclase-activating polypeptide type I-receptor-deficient mice - PubMed — https://pubmed.ncbi.nlm.nih.gov/12573539/
- **[S11]** Pituitary adenylate cyclase-activating polypeptide type 1 receptor within the nucleus accumbens core mediates excessive alcohol drinking in alcohol-preferring rats - PubMed — https://pubmed.ncbi.nlm.nih.gov/35460713/
- **[S12]** Viral-Mediated Knockdown of Nucleus Accumbens Shell PAC1 Receptor Promotes Excessive Alcohol Drinking in Alcohol-Preferring Rats - PubMed — https://pubmed.ncbi.nlm.nih.gov/34924973/
- **[S13]** Targeting the PAC1 Receptor for Neurological and Metabolic Disorders - PubMed — https://pubmed.ncbi.nlm.nih.gov/31284862/
- **[S14]** Pituitary adenylate cyclase-activating polypeptide-induced PAC1 receptor internalization and recruitment of MEK/ERK signaling enhance excitability of dentate gyrus granule cells - PubMed — https://pubmed.ncbi.nlm.nih.gov/32186931/
- **[S15]** Pituitary adenylate cyclase-activating polypeptide type 1 receptor within the nucleus accumbens core mediates excessive alcohol drinking in alcohol-preferring rats - PubMed — https://pubmed.ncbi.nlm.nih.gov/35460713/
- **[S16]** PAC1 receptor (ADCYAP1R1) genotype and problematic alcohol use in a sample of young women - PubMed — https://pubmed.ncbi.nlm.nih.gov/28652748/?utm_source=openai
