## Supplementary material for "Understanding Substance Dependence: What Differentiates Addictive from Non-Addictive Drugs?": Folder for supplementary files: addictive-genes-claims-first100-051-rbp1-gene-protein-is-implicated-in-addiction-f7c3d57ba6ca.md

# addictive-genes-claims-first100-051-f7c3d57ba6ca

**Original claim:** RBP1 gene/protein is implicated in addiction.

## Claim

**RBP1** (here, the official human gene symbol for **retinol binding protein 1 / cellular retinol-binding protein 1**) is implicated in addiction. RBP1 encodes an intracellular retinol-binding protein involved in retinoid metabolism rather than one of the classic dopamine/opioid/nicotinic receptor genes most strongly established in substance use disorders. ([ncbi.nlm.nih.gov](https://www.ncbi.nlm.nih.gov/gene/5947?utm_source=openai)) [S1]

## Verdict

Verdict: Mixed Evidence

## Confidence

**Medium**

## Supporting evidence

There **is** preclinical evidence linking the **retinoic acid (RA) signaling pathway**—a pathway in which RBP1 participates—to addiction-related neurobiology. In a rat nucleus accumbens (NAc) transcriptomic study comparing environmentally enriched versus isolated animals after cocaine or saline exposure, RA signaling emerged as a prominent pathway. In that same study, **Rbp1** was identified as one of several RA-pathway genes with selective/enhanced expression in the **NAc shell**, a key reward region. Importantly, experimentally increasing RA signaling in NAc shell by knocking down the RA-degrading enzyme **Cyp26b1** increased cocaine self-administration/drug-seeking measures, supporting a role for this pathway in addiction-related behavior. ([pmc.ncbi.nlm.nih.gov](https://pmc.ncbi.nlm.nih.gov/articles/PMC5110542/)) [S2]

More recent work strengthens the case for **RA signaling** in cocaine-related plasticity. In rats, increasing RA signaling in nucleus accumbens core during abstinence by **Cyp26b1** knockdown accelerated the “incubation” of cocaine craving and accelerated synaptic insertion of Ca²⁺-permeable AMPA receptors. A follow-up study reported that RA signaling in D1-type accumbens neurons contributes to maintenance of incubation-associated synaptic changes and cocaine seeking. These studies do **not** manipulate RBP1 directly, but they make the broader RA pathway biologically relevant to addiction phenotypes. ([pubmed.ncbi.nlm.nih.gov](https://pubmed.ncbi.nlm.nih.gov/38935096/?utm_source=openai)) [S3]

There is also older **genetic/transcriptomic candidate-gene evidence** from alcohol-preference models. A mouse transcriptome meta-analysis of high- versus low-alcohol-consuming strains identified **Rbp1** among cis-regulated candidate genes within an alcohol preference QTL. That finding suggests Rbp1 may track with inherited liability to alcohol-drinking phenotypes in rodents, at least as a positional/candidate signal. ([pubmed.ncbi.nlm.nih.gov](https://pubmed.ncbi.nlm.nih.gov/16618939/?utm_source=openai)) [S4]

Taken together, these data support a **plausible, indirect implication**: RBP1 sits inside a pathway—retinoid/RA signaling—that has experimental links to cocaine-related neuroplasticity, and it has appeared as a candidate in rodent alcohol-preference genetics. That is enough to say the claim is **not baseless**. ([pmc.ncbi.nlm.nih.gov](https://pmc.ncbi.nlm.nih.gov/articles/PMC5110542/)) [S5]

## Contradicting or limiting evidence

The main limitation is that the evidence is **mostly indirect and preclinical**. In the cocaine/NAc study, the causal manipulation was of **Cyp26b1**, not **Rbp1** itself; the authors explicitly framed several RA-pathway components—including **Rbp1**—as candidate targets for future work rather than demonstrating that Rbp1 specifically drives the behavioral effect. ([pmc.ncbi.nlm.nih.gov](https://pmc.ncbi.nlm.nih.gov/articles/PMC5110542/)) [S6]

Likewise, the alcohol evidence is **candidate-gene/QTL** evidence, not strong causal validation. In a later, more tightly filtered systems-genetics analysis of alcohol preference in mice, the final candidate list emphasized genes such as **Gnb1, C1qb, C1qc, Scn4b, Cep63,** and **Ube2j2** rather than Rbp1. The paper also noted that an earlier study had identified many more correlated transcripts because it did **not** systematically apply the eQTL/bQTL overlap filter. That does not refute Rbp1, but it does weaken confidence that Rbp1 is a robust, repeatedly prioritized alcohol gene. ([pmc.ncbi.nlm.nih.gov](https://pmc.ncbi.nlm.nih.gov/articles/PMC2583933/)) [S7]

Human evidence is also weak relative to established addiction genes. A recent review of substance use disorder genetics highlights replicated loci such as **ADH1B/ADH1C/DRD2** for problematic alcohol use, **CHRNA2** for cannabis use disorder, **CHRNA5-CHRNA3-CHRNB4** for nicotine dependence, and **OPRM1/FURIN/TSNARE1** for opioid use disorder. RBP1 is not among the major robust loci emphasized there, which argues against it being an established human addiction gene at present. ([nature.com](https://www.nature.com/articles/s41380-024-02547-z)) [S8]

So, if the claim is interpreted strongly—meaning **RBP1 itself has direct, replicated causal or human genetic evidence in addiction**—the current literature does **not** support that level of confidence. The best-supported statement is narrower: **RBP1 is a plausible pathway-level candidate within retinoid signaling, not a firmly established addiction gene/protein.** ([pmc.ncbi.nlm.nih.gov](https://pmc.ncbi.nlm.nih.gov/articles/PMC5110542/)) [S9]

## Mechanistic / biological rationale

Biologically, the idea is plausible. RBP1 binds intracellular retinol and helps regulate the availability of substrates for **retinoic acid** production. Retinoid signaling is active in the adult brain, and reviews describe roles for RA in neuronal gene regulation, synaptic function, and homeostatic plasticity. ([ncbi.nlm.nih.gov](https://www.ncbi.nlm.nih.gov/gene/5947?utm_source=openai)) [S10]

That matters because addiction involves persistent plasticity in reward circuitry, especially in the **nucleus accumbens**. The cocaine studies above argue that RA signaling in accumbal neurons can regulate AMPA-receptor remodeling and craving-related plasticity during abstinence. In principle, a protein like RBP1—by shaping local retinoid handling—could influence that process. ([pubmed.ncbi.nlm.nih.gov](https://pubmed.ncbi.nlm.nih.gov/38935096/?utm_source=openai)) [S11]

However, this remains a **pathway inference**, not a demonstrated RBP1-specific mechanism. RA signaling depends on multiple components, including retinol uptake/binding proteins, synthetic enzymes (for example **Aldh1a1/Aldh1a3**), degradative enzymes (**Cyp26b1**), and receptors (**Rarb/Rxrg**). The available addiction studies manipulate other RA-pathway nodes, not RBP1 itself. So the mechanistic rationale is biologically coherent but still **one step removed** from proving that RBP1 is a causal addiction factor. ([pmc.ncbi.nlm.nih.gov](https://pmc.ncbi.nlm.nih.gov/articles/PMC5110542/)) [S12]

## Bottom line

There is **some real scientific basis** for connecting **RBP1** to addiction, but the support is **indirect and mostly rodent-based**. The strongest evidence is that **retinoic acid signaling in nucleus accumbens is involved in cocaine-related plasticity and craving**, and **Rbp1** is one of the RA-pathway genes enriched in that circuitry; an older mouse alcohol-genetics study also flagged **Rbp1** as a candidate gene. But I did **not** find strong direct evidence that **RBP1 itself** is a replicated human addiction gene or that manipulating RBP1 alone changes addiction behavior. So the fairest assessment is: **RBP1 is a plausible candidate within an addiction-relevant retinoid pathway, but it is not yet firmly established as an addiction gene/protein.** ([pmc.ncbi.nlm.nih.gov](https://pmc.ncbi.nlm.nih.gov/articles/PMC5110542/)) [S13]

## Cited sources

- **[S1]** RBP1 retinol binding protein 1 [Homo sapiens (human)] - Gene - NCBI — https://www.ncbi.nlm.nih.gov/gene/5947?utm_source=openai
- **[S2]** 
            Transcriptomics of Environmental Enrichment Reveals a Role for Retinoic Acid Signaling in Addiction - PMC
         — https://pmc.ncbi.nlm.nih.gov/articles/PMC5110542/
- **[S3]** Retinoic acid-mediated homeostatic plasticity in the nucleus accumbens core contributes to incubation of cocaine craving - PubMed — https://pubmed.ncbi.nlm.nih.gov/38935096/?utm_source=openai
- **[S4]** Toward understanding the genetics of alcohol drinking through transcriptome meta-analysis - PubMed — https://pubmed.ncbi.nlm.nih.gov/16618939/?utm_source=openai
- **[S5]** 
            Transcriptomics of Environmental Enrichment Reveals a Role for Retinoic Acid Signaling in Addiction - PMC
         — https://pmc.ncbi.nlm.nih.gov/articles/PMC5110542/
- **[S6]** 
            Transcriptomics of Environmental Enrichment Reveals a Role for Retinoic Acid Signaling in Addiction - PMC
         — https://pmc.ncbi.nlm.nih.gov/articles/PMC5110542/
- **[S7]** 
            The genomic determinants of alcohol preference in mice - PMC
         — https://pmc.ncbi.nlm.nih.gov/articles/PMC2583933/
- **[S8]** The genetic landscape of substance use disorders | Molecular Psychiatry — https://www.nature.com/articles/s41380-024-02547-z
- **[S9]** 
            Transcriptomics of Environmental Enrichment Reveals a Role for Retinoic Acid Signaling in Addiction - PMC
         — https://pmc.ncbi.nlm.nih.gov/articles/PMC5110542/
- **[S10]** RBP1 retinol binding protein 1 [Homo sapiens (human)] - Gene - NCBI — https://www.ncbi.nlm.nih.gov/gene/5947?utm_source=openai
- **[S11]** Retinoic acid-mediated homeostatic plasticity in the nucleus accumbens core contributes to incubation of cocaine craving - PubMed — https://pubmed.ncbi.nlm.nih.gov/38935096/?utm_source=openai
- **[S12]** 
            Transcriptomics of Environmental Enrichment Reveals a Role for Retinoic Acid Signaling in Addiction - PMC
         — https://pmc.ncbi.nlm.nih.gov/articles/PMC5110542/
- **[S13]** 
            Transcriptomics of Environmental Enrichment Reveals a Role for Retinoic Acid Signaling in Addiction - PMC
         — https://pmc.ncbi.nlm.nih.gov/articles/PMC5110542/
