## Supplementary material for "Understanding Substance Dependence: What Differentiates Addictive from Non-Addictive Drugs?": Folder for supplementary files: addictive-genes-claims-first100-052-hcn2-gene-protein-is-implicated-in-addiction-f85a0385292b.md

# addictive-genes-claims-first100-052-f85a0385292b

**Original claim:** HCN2 gene/protein is implicated in addiction.

## Claim

HCN2 gene/protein is implicated in addiction.

## Verdict

Verdict: Likely True

Multiple **rodent** studies link HCN2 to addiction-relevant phenotypes, especially **alcohol reward/intake** and **morphine conditioned place preference (CPP)**, and cocaine exposure alters HCN2-related signaling in reward circuitry. However, the evidence is still **mostly preclinical**, not strongly established in humans, and some of the cocaine data are mechanistically mixed or not clearly **HCN2-specific**. ([pubmed.ncbi.nlm.nih.gov](https://pubmed.ncbi.nlm.nih.gov/35377277/)) [S1]

## Confidence

Medium

The conclusion rests on **multiple independent animal studies** with behavioral and electrophysiologic readouts, but there is **limited direct human genetic or clinical evidence** identifying HCN2 as a confirmed addiction-risk gene or therapeutic target. ([pubmed.ncbi.nlm.nih.gov](https://pubmed.ncbi.nlm.nih.gov/35377277/)) [S2]

## Supporting evidence

The strongest evidence is for **alcohol-related behavior**. In alcohol-preferring UChB rats, **VTA overexpression of HCN2** increased voluntary ethanol intake by about twofold, increased ethanol CPP by about threefold, increased ethanol-induced locomotion, and increased ethanol-evoked dopamine release in nucleus accumbens. In a complementary study from the same line of work, **VTA knockdown of HCN2** reduced ethanol preference and intake by about 70% without reducing total fluid intake, supporting a causal role in alcohol reinforcement rather than a nonspecific sickness effect. ([pubmed.ncbi.nlm.nih.gov](https://pubmed.ncbi.nlm.nih.gov/24460767/)) [S3]

There is also supportive evidence for **opioid reward**. In a rat chronic morphine model, investigators found **upregulated HCN2 mRNA and protein in the VTA**, stronger HCN2 signal in **TH-positive dopaminergic neurons**, and increased **Ih** current magnitude/density after morphine exposure. In the same study, **intra-VTA ZD7288** (an HCN-channel blocker) reduced morphine CPP, consistent with HCN-channel involvement in morphine reward learning. ([pubmed.ncbi.nlm.nih.gov](https://pubmed.ncbi.nlm.nih.gov/39267158/)) [S4]

For **cocaine**, the evidence is more indirect but still supportive. A 2012 rat study found that cocaine sensitization increased HCN2 protein expression across several mesocorticolimbic regions: about **55% in VTA**, **35% in prefrontal cortex**, **36% in nucleus accumbens**, and **22% in hippocampus**. The authors explicitly interpreted these HCN2 changes as potentially relevant to addiction-related neuroadaptations. ([pubmed.ncbi.nlm.nih.gov](https://pubmed.ncbi.nlm.nih.gov/23203153/)) [S5]

More broadly, a 2023 *Molecular Psychiatry* study showed that chronic **cocaine self-administration** increased **Ih** in VTA dopamine neurons and that systemic or intra-VTA **ivabradine** reduced cocaine self-administration under a progressive-ratio schedule and shifted the cocaine dose-response curve downward. That study strengthens the case for **HCN-channel involvement** in cocaine reinforcement, though it does **not** pin the effect specifically on HCN2. ([pubmed.ncbi.nlm.nih.gov](https://pubmed.ncbi.nlm.nih.gov/37845497/)) [S6]

Transcriptomic evidence also points in the same direction, though again mostly preclinically. A 2024 systematic review/meta-analysis of alcohol use disorder transcriptomics reported that in **rodent prefrontal cortex**, **Hcn2** emerged among “previously well-characterized alcohol-sensitive genes.” This is not definitive on its own, but it supports the idea that Hcn2 is part of alcohol-responsive molecular adaptation in addiction-relevant circuitry. ([nature.com](https://www.nature.com/articles/s41380-024-02719-x)) [S7]

## Contradicting or limiting evidence

The main limitation is that **human evidence is weak**. A 2021 umbrella review covering more than 150 meta-analyses of substance use disorder genetics highlighted recurrent genes such as **OPRM1, DRD2, DRD4, BDNF, SLC6A4**, and GWAS signals such as **CHRNA5, FOXP2, PEX, AUTS2**; **HCN2 was not identified as an established cross-study signal**. That does not prove HCN2 is irrelevant, but it argues against calling it a well-established **human addiction gene** at present. ([pubmed.ncbi.nlm.nih.gov](https://pubmed.ncbi.nlm.nih.gov/33556390/)) [S8]

The cocaine literature is also not fully internally consistent. One study found that cocaine sensitization **reduced Ih amplitude** in VTA dopamine neurons and reduced cell capacitance/cell size, even though related work found increased HCN2 protein expression after cocaine sensitization. The authors suggested this might reflect **homeostatic scaling**, altered channel trafficking, or changes in surface/intracellular ratios rather than a simple monotonic increase in HCN2 function. ([pmc.ncbi.nlm.nih.gov](https://pmc.ncbi.nlm.nih.gov/articles/PMC3331609/)) [S9]

Specificity is another issue. Some key pharmacology uses **HCN-channel blockers** such as **ZD7288** or **ivabradine**, which affect the HCN family rather than HCN2 alone. Also, the 2023 cocaine study reported that in rat VTA, **Hcn3 and Hcn4** were the predominant HCN mRNAs in dopamine neurons, whereas **Hcn2 was barely detectable in Slc6a3-positive dopamine neurons** and was more prominent in VTA GABA neurons. So for cocaine in particular, “HCN channels” may be a stronger statement than “HCN2 specifically.” ([pubmed.ncbi.nlm.nih.gov](https://pubmed.ncbi.nlm.nih.gov/39267158/)) [S10]

## Mechanistic / biological rationale

HCN2 encodes a **hyperpolarization-activated cyclic nucleotide-gated cation channel** that contributes to **Ih**, a current important for neuronal membrane properties and excitability. HCN2 is **brain-enriched** and shows enhanced expression across several brain neuronal cell types. More generally, HCN channels in midbrain dopamine systems regulate resting membrane behavior, afterhyperpolarization, dendritic integration, and the balance between tonic and burst firing—features tightly tied to dopamine signaling in reward circuits. ([ncbi.nlm.nih.gov](https://www.ncbi.nlm.nih.gov/gene/610)) [S11]

That biology fits addiction plausibly. Reviews of VTA dopamine physiology note that HCN/Ih signaling can shape dopamine neuron firing patterns and thereby alter dopamine release to nucleus accumbens and related targets. Ethanol has been reported to augment **Ih** in VTA dopamine neurons, and chronic exposure can alter that current, making HCN-family channels a credible substrate for reinforcement-related plasticity. ([pmc.ncbi.nlm.nih.gov](https://pmc.ncbi.nlm.nih.gov/articles/PMC4002296/)) [S12]

Still, the mechanistic picture may differ by drug and cell type. For alcohol and morphine, the data more directly support HCN2-related reward effects in VTA-centered circuitry. For cocaine, HCN-related adaptations clearly occur, but the available evidence suggests they may involve **multiple HCN subunits**, altered trafficking, and compensatory/homeostatic processes rather than a simple “more HCN2 = more addiction” model. ([pubmed.ncbi.nlm.nih.gov](https://pubmed.ncbi.nlm.nih.gov/24460767/)) [S13]

## Bottom line

The claim is **likely true**, with an important qualifier: **HCN2 is implicated mainly by preclinical evidence**, not yet as a firmly validated human addiction gene. The best support is for **alcohol-related behaviors** and **morphine reward** in rodent VTA circuitry, with additional but less clean evidence in **cocaine-related** neuroadaptations. So it is fair to say HCN2 is **implicated in addiction-related biology and behavior**, but not fair to say it is already a **well-established human addiction determinant**. ([pubmed.ncbi.nlm.nih.gov](https://pubmed.ncbi.nlm.nih.gov/35377277/)) [S14]

## Cited sources

- **[S1]** https://pubmed.ncbi.nlm.nih.gov/35377277/ — https://pubmed.ncbi.nlm.nih.gov/35377277/
- **[S2]** https://pubmed.ncbi.nlm.nih.gov/35377277/ — https://pubmed.ncbi.nlm.nih.gov/35377277/
- **[S3]** https://pubmed.ncbi.nlm.nih.gov/24460767/ — https://pubmed.ncbi.nlm.nih.gov/24460767/
- **[S4]** https://pubmed.ncbi.nlm.nih.gov/39267158/ — https://pubmed.ncbi.nlm.nih.gov/39267158/
- **[S5]** https://pubmed.ncbi.nlm.nih.gov/23203153/ — https://pubmed.ncbi.nlm.nih.gov/23203153/
- **[S6]** https://pubmed.ncbi.nlm.nih.gov/37845497/ — https://pubmed.ncbi.nlm.nih.gov/37845497/
- **[S7]** https://www.nature.com/articles/s41380-024-02719-x — https://www.nature.com/articles/s41380-024-02719-x
- **[S8]** https://pubmed.ncbi.nlm.nih.gov/33556390/ — https://pubmed.ncbi.nlm.nih.gov/33556390/
- **[S9]** https://pmc.ncbi.nlm.nih.gov/articles/PMC3331609/ — https://pmc.ncbi.nlm.nih.gov/articles/PMC3331609/
- **[S10]** https://pubmed.ncbi.nlm.nih.gov/39267158/ — https://pubmed.ncbi.nlm.nih.gov/39267158/
- **[S11]** https://www.ncbi.nlm.nih.gov/gene/610 — https://www.ncbi.nlm.nih.gov/gene/610
- **[S12]** https://pmc.ncbi.nlm.nih.gov/articles/PMC4002296/ — https://pmc.ncbi.nlm.nih.gov/articles/PMC4002296/
- **[S13]** https://pubmed.ncbi.nlm.nih.gov/24460767/ — https://pubmed.ncbi.nlm.nih.gov/24460767/
- **[S14]** https://pubmed.ncbi.nlm.nih.gov/35377277/ — https://pubmed.ncbi.nlm.nih.gov/35377277/
