## Supplementary material for "Understanding Substance Dependence: What Differentiates Addictive from Non-Addictive Drugs?": Folder for supplementary files: addictive-genes-claims-first100-053-cacna1a-gene-protein-is-implicated-in-addiction-f2bd1988370a.md

# addictive-genes-claims-first100-053-f2bd1988370a

**Original claim:** CACNA1A gene/protein is implicated in addiction.

## Claim

CACNA1A gene/protein is implicated in addiction.

## Verdict

Verdict: Likely True

## Confidence

Medium

## Supporting evidence

CACNA1A encodes the pore-forming α1A subunit of the neuronal P/Q-type voltage-gated calcium channel CaV2.1, which is highly expressed in brain and is central to calcium-dependent neurotransmitter release. That function makes it biologically plausible as a modulator of reward-circuit signaling and drug-induced synaptic plasticity. ([ncbi.nlm.nih.gov](https://www.ncbi.nlm.nih.gov/gene/773)) [S1]

The strongest direct human evidence I found is for **cannabis use disorder (CUD)** rather than “addiction” in general. In a 2023 multivariate GWAS/meta-analysis of more than 1 million participants across problematic alcohol use, problematic tobacco use, CUD, and opioid use disorder, the authors reported a **CUD-specific genome-wide significant signal at rs8104317 near CACNA1A** with **P = 1.17 × 10⁻13**. In the same paper, CACNA1A was not presented as a general shared addiction-risk locus; it appeared as a substance-specific association for CUD. ([icuddr.org](https://icuddr.org/wp-content/uploads/2024/12/s44220-023-00034-y.pdf)) [S2]

There is also **addiction-relevant animal/circuit evidence** for CaV2.1/P/Q-type channels. In rat central amygdala slices, acute ethanol increased GABA release through a mechanism requiring **presynaptic CRF1 receptors and P/Q-type voltage-gated calcium channels**; importantly, this mechanism remained engaged after chronic intermittent ethanol exposure, a dependence model. The authors explicitly framed these channels as “key regulators” of alcohol effects on CeA GABAergic signaling during the transition to dependence. ([pubmed.ncbi.nlm.nih.gov](https://pubmed.ncbi.nlm.nih.gov/28734867/)) [S3]

Another rodent alcohol-withdrawal study found that **Cacna1a mRNA increased 1.9-fold at 3 h and 2.1-fold at 24 h** in the inferior colliculus during ethanol withdrawal, before peak withdrawal-seizure susceptibility, suggesting involvement in withdrawal neuroadaptation. ([pubmed.ncbi.nlm.nih.gov](https://pubmed.ncbi.nlm.nih.gov/29277284/)) [S4]

A methamphetamine study in mouse medial prefrontal cortex likewise reported altered **Cacna1a** expression after repeated METH exposure/withdrawal, alongside changes in calcium currents and synaptic physiology that the authors linked to prefrontal dysfunction in METH addiction. ([pubmed.ncbi.nlm.nih.gov](https://pubmed.ncbi.nlm.nih.gov/25871318/)) [S5]

Together, these findings support the narrower statement that **CACNA1A/CaV2.1 is implicated in at least some substance use disorder biology, especially cannabis use disorder genetics and alcohol-related neuroadaptation in animal models**. ([icuddr.org](https://icuddr.org/wp-content/uploads/2024/12/s44220-023-00034-y.pdf)) [S6]

## Contradicting or limiting evidence

The main limitation is **scope**. CACNA1A is not currently established as a **broad, consistently replicated pan-addiction gene** on the level of canonical loci such as **ADH1B** for alcohol, **CHRNA5/CHRNA3/CHRNB4** for nicotine/tobacco, **CHRNA2** for CUD, or **OPRM1** for opioid use disorder. Large GWASs of alcohol dependence, nicotine dependence, cannabis use disorder, and opioid use disorder have highlighted those loci as robust signals. ([pubmed.ncbi.nlm.nih.gov](https://pubmed.ncbi.nlm.nih.gov/30482948/?utm_source=openai)) [S7]

A 2022 integrated review of genomic and transcriptomic studies across alcohol, nicotine, and drug use/disorders identified shared genes such as **CACNB2, GRIN2B, PLXDC2, and PKNOX2**, but did **not** single out CACNA1A as one of the main shared cross-substance genes. ([pubmed.ncbi.nlm.nih.gov](https://pubmed.ncbi.nlm.nih.gov/35491750/)) [S8]

Similarly, a 2025 large cross-SUD meta-analysis reported **785 prioritized shared genes** across alcohol, cannabis, opioid, and tobacco-related disorders, but CACNA1A was not highlighted as one of the prioritized shared loci in the paper text. That does not disprove a role in addiction, but it argues against CACNA1A being a major broadly shared addiction gene at present. ([pmc.ncbi.nlm.nih.gov](https://pmc.ncbi.nlm.nih.gov/articles/PMC12916498/)) [S9]

Older human association evidence for alcohol-related phenotypes is also weak. In one community-based study of GABA-system-related genes, several **CACNA1A SNPs showed only nominal associations** with alcohol phenotypes, but **none survived appropriate multiple-testing correction**, and the paper’s overall conclusion was “no evidence for a role” of the tested GABA-system markers in alcohol use/abuse in that sample. ([pmc.ncbi.nlm.nih.gov](https://pmc.ncbi.nlm.nih.gov/articles/PMC3984361/)) [S10]

The animal alcohol-withdrawal paper also found a disconnect between transcript and protein: **Cacna1a mRNA rose, but CaV2.1 protein did not significantly change** in the measured tissue. That weakens any claim that altered CACNA1A expression is already proven to drive addiction phenotypes directly. ([pubmed.ncbi.nlm.nih.gov](https://pubmed.ncbi.nlm.nih.gov/29277284/)) [S11]

Finally, CACNA1A is best known clinically for **episodic ataxia type 2, familial hemiplegic migraine, spinocerebellar ataxia type 6, and developmental/epileptic encephalopathy**, not addiction. So addiction is not its dominant established disease association. ([medlineplus.gov](https://medlineplus.gov/genetics/gene/cacna1a/)) [S12]

## Mechanistic / biological rationale

Mechanistically, CACNA1A is plausible because CaV2.1 channels sit at presynaptic terminals and control calcium entry that triggers vesicle fusion and neurotransmitter release. Drug exposure and withdrawal often remodel synaptic release probability, inhibitory/excitatory balance, and stress-circuit signaling; a presynaptic calcium channel can therefore influence addiction-relevant plasticity even if it is not a top GWAS hit across every substance. ([ncbi.nlm.nih.gov](https://www.ncbi.nlm.nih.gov/gene/773)) [S13]

In alcohol models, P/Q-type channels appear to participate in **central amygdala GABA release**, interacting with **CRF signaling**, a pathway strongly tied to dependence, stress, and negative affect during addiction. ([pubmed.ncbi.nlm.nih.gov](https://pubmed.ncbi.nlm.nih.gov/28734867/)) [S14]

In psychostimulant models, repeated methamphetamine altered **Cacna1a** expression together with calcium currents and excitatory transmission in medial prefrontal cortex, consistent with a role in drug-induced cortical plasticity. ([pubmed.ncbi.nlm.nih.gov](https://pubmed.ncbi.nlm.nih.gov/25871318/)) [S15]

That said, the mechanistic evidence remains mostly **indirect and preclinical**. It supports biological plausibility and involvement in certain drug responses, but it does not yet prove that CACNA1A is a central driver of addiction across substances or that targeting CaV2.1 would treat addiction in humans. ([pubmed.ncbi.nlm.nih.gov](https://pubmed.ncbi.nlm.nih.gov/25871318/)) [S16]

## Bottom line

The claim is **reasonable but should be stated narrowly**. Current evidence supports that **CACNA1A/CaV2.1 is implicated in some addiction-related phenotypes**, with the clearest direct human evidence being a **genome-wide significant association with cannabis use disorder**, plus animal evidence linking P/Q-type channels to **alcohol-related dependence/withdrawal mechanisms** and methamphetamine-induced cortical adaptations. ([icuddr.org](https://icuddr.org/wp-content/uploads/2024/12/s44220-023-00034-y.pdf)) [S17]

However, CACNA1A is **not yet a well-established, broadly replicated general addiction gene** across alcohol, nicotine, opioid, and cannabis disorders. So the best summary is: **likely true in a substance-specific/biological-systems sense, but overstated if meant as a major universal addiction gene.** ([pubmed.ncbi.nlm.nih.gov](https://pubmed.ncbi.nlm.nih.gov/35491750/)) [S18]

## Cited sources

- **[S1]** CACNA1A calcium voltage-gated channel subunit alpha1 A [Homo sapiens (human)] - Gene - NCBI — https://www.ncbi.nlm.nih.gov/gene/773
- **[S2]** Multivariate genome-wide association meta-analysis of over 1 million subjects identifies loci underlying multiple substance use disorders — https://icuddr.org/wp-content/uploads/2024/12/s44220-023-00034-y.pdf
- **[S3]** P/Q-type voltage-gated calcium channels mediate the ethanol and CRF sensitivity of central amygdala GABAergic synapses - PubMed — https://pubmed.ncbi.nlm.nih.gov/28734867/
- **[S4]** Alcohol withdrawal upregulates mRNA encoding for CaV2.1-α1 subunit in the rat inferior colliculus - PubMed — https://pubmed.ncbi.nlm.nih.gov/29277284/
- **[S5]** Methamphetamine blunts Ca(2+) currents and excitatory synaptic transmission through D1/5 receptor-mediated mechanisms in the mouse medial prefrontal cortex - PubMed — https://pubmed.ncbi.nlm.nih.gov/25871318/
- **[S6]** Multivariate genome-wide association meta-analysis of over 1 million subjects identifies loci underlying multiple substance use disorders — https://icuddr.org/wp-content/uploads/2024/12/s44220-023-00034-y.pdf
- **[S7]** Transancestral GWAS of alcohol dependence reveals common genetic underpinnings with psychiatric disorders - PubMed — https://pubmed.ncbi.nlm.nih.gov/30482948/?utm_source=openai
- **[S8]** Common and distinguishing genetic factors for substance use behavior and disorder: an integrated analysis of genomic and transcriptomic studies from both human and animal studies - PubMed — https://pubmed.ncbi.nlm.nih.gov/35491750/
- **[S9]** 
            Genome-wide meta-analyses of cross substance use disorders in diverse populations - PMC
         — https://pmc.ncbi.nlm.nih.gov/articles/PMC12916498/
- **[S10]** 
            GABA System Genes – No Evidence for a Role in Alcohol Use and Abuse in a Community-based Sample - PMC
         — https://pmc.ncbi.nlm.nih.gov/articles/PMC3984361/
- **[S11]** Alcohol withdrawal upregulates mRNA encoding for CaV2.1-α1 subunit in the rat inferior colliculus - PubMed — https://pubmed.ncbi.nlm.nih.gov/29277284/
- **[S12]** CACNA1A gene: MedlinePlus Genetics — https://medlineplus.gov/genetics/gene/cacna1a/
- **[S13]** CACNA1A calcium voltage-gated channel subunit alpha1 A [Homo sapiens (human)] - Gene - NCBI — https://www.ncbi.nlm.nih.gov/gene/773
- **[S14]** P/Q-type voltage-gated calcium channels mediate the ethanol and CRF sensitivity of central amygdala GABAergic synapses - PubMed — https://pubmed.ncbi.nlm.nih.gov/28734867/
- **[S15]** Methamphetamine blunts Ca(2+) currents and excitatory synaptic transmission through D1/5 receptor-mediated mechanisms in the mouse medial prefrontal cortex - PubMed — https://pubmed.ncbi.nlm.nih.gov/25871318/
- **[S16]** Methamphetamine blunts Ca(2+) currents and excitatory synaptic transmission through D1/5 receptor-mediated mechanisms in the mouse medial prefrontal cortex - PubMed — https://pubmed.ncbi.nlm.nih.gov/25871318/
- **[S17]** Multivariate genome-wide association meta-analysis of over 1 million subjects identifies loci underlying multiple substance use disorders — https://icuddr.org/wp-content/uploads/2024/12/s44220-023-00034-y.pdf
- **[S18]** Common and distinguishing genetic factors for substance use behavior and disorder: an integrated analysis of genomic and transcriptomic studies from both human and animal studies - PubMed — https://pubmed.ncbi.nlm.nih.gov/35491750/
