## Supplementary material for "Understanding Substance Dependence: What Differentiates Addictive from Non-Addictive Drugs?": Folder for supplementary files: addictive-genes-claims-first100-054-gltp-gene-protein-is-implicated-in-addiction-bf3c4e7a601b.md

# addictive-genes-claims-first100-054-bf3c4e7a601b

**Original claim:** GLTP gene/protein is implicated in addiction.

## Claim

GLTP gene/protein is implicated in addiction.

## Verdict

Verdict: Mixed Evidence

There is **some preclinical, indirect evidence** that GLTP could be relevant to addiction-related neurobiology, mainly through its roles in oligodendrocytes, myelin growth, glycosphingolipid handling, and an observed expression change in a mouse alcohol-use-disorder model. However, I did **not** find strong evidence that GLTP is an established addiction gene/protein in humans, nor direct demonstrations that GLTP manipulation changes core addiction phenotypes such as drug reward, self-administration, relapse, or withdrawal. ([mdpi.com](https://www.mdpi.com/2073-4409/12/5/745)) [S1]

## Confidence

Medium

The verdict is moderately confident because the relevant literature is limited and uneven: there is a real biological signal around GLTP in myelin/oligodendrocyte biology and one direct alcohol-model transcriptomic association, but the usual stronger forms of evidence for “implication in addiction” (human genetics, replicated human brain data, causal animal-behavior studies) were not evident in the retrieved literature. ([mdpi.com](https://www.mdpi.com/2073-4409/12/5/745)) [S2]

## Supporting evidence

GLTP encodes **glycolipid transfer protein**, a reviewed human protein-coding gene on chromosome 12. NCBI describes it as a cytoplasmic protein that accelerates transfer of certain glycosphingolipids/glyceroglycolipids between membranes, and lists broad tissue expression. Independent mechanistic studies further show that GLTP regulates glycosphingolipid homeostasis and can affect vesicular trafficking. Those are potentially relevant functions for neural signaling, because membrane lipid composition strongly influences cell identity and communication. ([ncbi.nlm.nih.gov](https://www.ncbi.nlm.nih.gov/gene/51228)) [S3]

The strongest addiction-adjacent evidence I found is from a **mouse chronic-plus-binge alcohol model**: cerebellar RNA-seq identified **Gltp** among ethanol-dysregulated oligodendrocyte-lineage genes, with reported downregulation of about **logFC −0.48, adjusted p = 0.0006**. This does not prove causality, but it is a direct observation tying Gltp expression to an alcohol-use-disorder experimental paradigm. ([pubmed.ncbi.nlm.nih.gov](https://pubmed.ncbi.nlm.nih.gov/36899881/?utm_source=openai)) [S4]

Separate work strengthens the biological relevance of GLTP in the CNS. In 2023, Zhang et al. reported that **Gltp is selectively expressed in differentiated oligodendrocytes**, is **essential for oligodendrocyte differentiation**, and promotes oligodendrocyte membrane outgrowth. In 2024, a CNS myelination study found GLTP **highly enriched in the growing myelin sheath**; oligodendrocyte-specific Gltp knockout mice showed **ER pathology, hypomyelination, and reduced myelin glycolipid content**. These findings place GLTP squarely in pathways that shape white matter and circuit function. ([pubmed.ncbi.nlm.nih.gov](https://pubmed.ncbi.nlm.nih.gov/37191854/)) [S5]

That matters because addiction research increasingly implicates **oligodendrocytes and myelin plasticity** in drug-induced circuit remodeling. A recent Nature study reported that **myelin plasticity in the ventral tegmental area is required for opioid reward**, arguing that maladaptive myelination can contribute to addiction-related learning. In alcohol biology, additional work shows that alcohol exposure alters oligodendrocyte transcriptomes and myelination-related pathways, providing a context in which GLTP-dependent oligodendrocyte biology could matter. ([nature.com](https://www.nature.com/articles/s41586-024-07525-7?utm_source=openai)) [S6]

## Contradicting or limiting evidence

The main limitation is the **absence of strong direct addiction evidence for GLTP**. In major human substance-use GWAS, the leading genes highlighted are other loci: for example, a cross-substance meta-analysis of more than 1 million subjects emphasized loci including **PDE4B** for general addiction liability, and a cannabis-use-disorder GWAS implicated **CHRNA2**. These papers do not present GLTP as a headline addiction locus. ([pubmed.ncbi.nlm.nih.gov](https://pubmed.ncbi.nlm.nih.gov/37250466/)) [S7]

Likewise, the NCBI Gene page for GLTP does not point to an addiction phenotype in its phenotype section; the visible GWAS-linked phenotype there is **metabolic syndrome**, not a substance-use disorder. That does not rule out future addiction relevance, but it argues against GLTP currently being a well-established addiction gene in human genetics. ([ncbi.nlm.nih.gov](https://www.ncbi.nlm.nih.gov/gene/51228)) [S8]

Also, the alcohol-model result is **expression-level association in one preclinical context**, not evidence that GLTP drives addictive behavior. The retrieved GLTP papers are focused mainly on lipid transfer, cell biology, oligodendrocyte maturation, myelin growth, autophagy/inflammation, and non-neural peripheral phenotypes such as epidermal differentiation disorder. Based on the literature retrieved, I did not identify studies showing that GLTP perturbation changes drug self-administration, conditioned place preference, craving, relapse, or withdrawal severity. ([pubmed.ncbi.nlm.nih.gov](https://pubmed.ncbi.nlm.nih.gov/37191854/)) [S9]

## Mechanistic / biological rationale

A biologically plausible case can be made, but it remains **indirect**. GLTP transfers glycosphingolipids and participates in glycosphingolipid homeostasis; it has also been linked to vesicular trafficking. Glycosphingolipids and myelin lipids influence membrane organization, axonal conduction, and glia-neuron interactions. If GLTP is important for oligodendrocyte maturation and myelin growth, then altered GLTP could in principle modify neural circuit timing, plasticity, and homeostatic support in reward-related pathways. ([ncbi.nlm.nih.gov](https://www.ncbi.nlm.nih.gov/gene/51228)) [S10]

That rationale is strengthened by modern addiction work showing that **myelin plasticity itself can participate in drug reward learning**, at least for opioids in mice. It is therefore reasonable to hypothesize that a myelin-relevant lipid-transfer protein like GLTP could modulate addiction vulnerability or adaptation. But that is still a **hypothesis**, not a demonstrated causal chain. No retrieved study directly showed that GLTP-specific dysfunction alters dopaminergic signaling, reward learning, compulsive drug taking, or relapse behavior. ([nature.com](https://www.nature.com/articles/s41586-024-07525-7?utm_source=openai)) [S11]

One more nuance: the available direct signal is **strongest for alcohol exposure**, not for addiction broadly across nicotine, stimulants, opioids, and cannabis. Even there, the evidence is a transcriptomic association in cerebellar oligodendrocyte-lineage genes, which is meaningfully suggestive but still far from establishing GLTP as a core alcohol-addiction gene. ([mdpi.com](https://www.mdpi.com/2073-4409/12/5/745)) [S12]

## Bottom line

GLTP is **not currently well established as an addiction gene/protein** by the standards usually used in addiction neuroscience. The best available evidence supports a narrower statement: **GLTP has plausible indirect relevance to addiction-related biology because it is important for oligodendrocyte/myelin lipid handling, and Gltp expression is altered in a mouse alcohol-use model**. That is enough to call the claim **partly plausible but weakly supported overall**, hence **Mixed Evidence** rather than Likely True or Highly Supported. ([mdpi.com](https://www.mdpi.com/2073-4409/12/5/745)) [S13]

## Cited sources

- **[S1]** Cerebellar Transcriptomic Analysis in a Chronic plus Binge Mouse Model of Alcohol Use Disorder Demonstrates Ethanol-Induced Neuroinflammation and Altered Glial Gene Expression | MDPI — https://www.mdpi.com/2073-4409/12/5/745
- **[S2]** Cerebellar Transcriptomic Analysis in a Chronic plus Binge Mouse Model of Alcohol Use Disorder Demonstrates Ethanol-Induced Neuroinflammation and Altered Glial Gene Expression | MDPI — https://www.mdpi.com/2073-4409/12/5/745
- **[S3]** GLTP glycolipid transfer protein [Homo sapiens (human)] - Gene - NCBI — https://www.ncbi.nlm.nih.gov/gene/51228
- **[S4]** Cerebellar Transcriptomic Analysis in a Chronic plus Binge Mouse Model of Alcohol Use Disorder Demonstrates Ethanol-Induced Neuroinflammation and Altered Glial Gene Expression - PubMed — https://pubmed.ncbi.nlm.nih.gov/36899881/?utm_source=openai
- **[S5]** Role of Gltp in Maturation of Oligodendrocytes Under the Regulation of Nkx2.2 - PubMed — https://pubmed.ncbi.nlm.nih.gov/37191854/
- **[S6]** Myelin plasticity in the ventral tegmental area is required for opioid reward | Nature — https://www.nature.com/articles/s41586-024-07525-7?utm_source=openai
- **[S7]** Multivariate genome-wide association meta-analysis of over 1 million subjects identifies loci underlying multiple substance use disorders - PubMed — https://pubmed.ncbi.nlm.nih.gov/37250466/
- **[S8]** GLTP glycolipid transfer protein [Homo sapiens (human)] - Gene - NCBI — https://www.ncbi.nlm.nih.gov/gene/51228
- **[S9]** Role of Gltp in Maturation of Oligodendrocytes Under the Regulation of Nkx2.2 - PubMed — https://pubmed.ncbi.nlm.nih.gov/37191854/
- **[S10]** GLTP glycolipid transfer protein [Homo sapiens (human)] - Gene - NCBI — https://www.ncbi.nlm.nih.gov/gene/51228
- **[S11]** Myelin plasticity in the ventral tegmental area is required for opioid reward | Nature — https://www.nature.com/articles/s41586-024-07525-7?utm_source=openai
- **[S12]** Cerebellar Transcriptomic Analysis in a Chronic plus Binge Mouse Model of Alcohol Use Disorder Demonstrates Ethanol-Induced Neuroinflammation and Altered Glial Gene Expression | MDPI — https://www.mdpi.com/2073-4409/12/5/745
- **[S13]** Cerebellar Transcriptomic Analysis in a Chronic plus Binge Mouse Model of Alcohol Use Disorder Demonstrates Ethanol-Induced Neuroinflammation and Altered Glial Gene Expression | MDPI — https://www.mdpi.com/2073-4409/12/5/745
