## Supplementary material for "Understanding Substance Dependence: What Differentiates Addictive from Non-Addictive Drugs?": Folder for supplementary files: addictive-genes-claims-first100-055-gabrb2-gene-protein-is-implicated-in-addiction-07f928524993.md

# addictive-genes-claims-first100-055-07f928524993

**Original claim:** GABRB2 gene/protein is implicated in addiction.

## Claim

**GABRB2 gene/protein is implicated in addiction.**

## Verdict

Verdict: Mixed Evidence

## Confidence

**Medium.**

## Supporting evidence

GABRB2 encodes the **β2 subunit of the GABA\_A receptor**, a ligand-gated chloride channel that mediates fast inhibitory synaptic transmission in the CNS; the gene lies in the chromosome **5q34 GABA\_A receptor cluster** with **GABRA1** and **GABRG2**. That makes it biologically plausible as an addiction candidate, because GABAergic signaling is central to alcohol and other drug responses. ([ncbi.nlm.nih.gov](https://www.ncbi.nlm.nih.gov/gene/2561)) [S1]

For **alcohol dependence**, an early study in two population isolates reported that the **GABRB2 1412T allele** was associated with alcohol dependence in both **Finns** and **Southwestern Native Americans**; the reported allele-level P values were **0.01** and **0.008**, respectively. ([jamanetwork.com](https://jamanetwork.com/journals/jamapsychiatry/fullarticle/208168)) [S2]

A separate **Scottish** candidate-gene study also investigated the chromosome 5 GABA\_A cluster and concluded that variants in **GABRB2/GABRA6/GABRG2** were associated with alcohol dependence in that population, supporting at least some replication of a chromosome 5 cluster signal. ([pubmed.ncbi.nlm.nih.gov](https://pubmed.ncbi.nlm.nih.gov/10578235/?utm_source=openai)) [S3]

For **opioid/heroin-related phenotypes**, a Han Chinese study found that while **single-marker allele/genotype tests were not significant**, several **GABRB2 haplotypes** across the S1–S29 segment showed strong association with **heroin dependence**, and the authors concluded that their results support a possible role of GABRB2 in heroin dependence but require replication. ([pubmed.ncbi.nlm.nih.gov](https://pubmed.ncbi.nlm.nih.gov/26561861/?utm_source=openai)) [S4]

Another study in subjects of **African and European ancestry** examining glutamatergic and GABAergic loci in heroin/cocaine addiction reported **GABRB2** among the **potential susceptibility loci/markers**, again suggesting possible involvement, though not definitive proof. ([pmc.ncbi.nlm.nih.gov](https://pmc.ncbi.nlm.nih.gov/articles/PMC4564302/?utm_source=openai)) [S5]

There is also **epigenetic evidence**. In a study of **alcohol-nicotine codependence**, one **GABRB2 promoter CpG** (**cg02095443**) was among the loci that survived Bonferroni correction in European Americans. The authors interpreted these methylation changes as potential biomarkers of addiction-related states, while also noting that causality and directionality were unresolved. ([nature.com](https://www.nature.com/articles/srep41816)) [S6]

Finally, a 2021 review focused on GABRB2 summarized that the gene has been associated with **substance dependence** among several neuropsychiatric phenotypes. That review is not primary evidence, but it reflects that the signal has appeared repeatedly enough in the literature to be noticed. ([pubmed.ncbi.nlm.nih.gov](https://pubmed.ncbi.nlm.nih.gov/34673206/)) [S7]

## Contradicting or limiting evidence

The strongest limitation is that **more robust pooled and family-based analyses have not established GABRB2 as a major addiction gene**. In a 2014 meta-analysis of five GABA\_A receptor genes across **4,739 cases** and **4,924 controls**, the significant pooled signals were for **GABRA2**, **GABRG2**, and **GABRA6**. **GABRB2 was included in the analysis but was not highlighted among the significant pooled associations**; that is a reasonable inference that the available evidence for GABRB2 was weaker and not robust in that meta-analysis. ([pubmed.ncbi.nlm.nih.gov](https://pubmed.ncbi.nlm.nih.gov/24136292/)) [S8]

Likewise, the large **COGA** family-based study of the chromosome 5 GABA\_A cluster found **“no compelling evidence”** for association of **GABRA1, GABRA6, GABRB2, or GABRG2** with alcohol dependence or alcohol dependence plus antisocial personality disorder. The authors specifically argued that sporadic GABRB2 SNP findings were likely **false positives**, citing inconsistency across methods, local LD structure, and multiple-testing concerns. ([downstate.edu](https://www.downstate.edu/research/centers-departments/henri-begleiter-neurodynamics-laboratory/_documents/dick_2005-NoAssociationGABAAReceptorGenesonChromosome5withAlcoholismintheCollaborativeStudyo.pdf)) [S9]

Negative studies also exist at the individual-study level. In **349 German alcohol-dependent subjects** and **182 controls**, investigators found **no evidence** that the tested **GABRB2/GABRA6/GABRG2** variants conferred vulnerability to alcohol dependence, withdrawal severity, or familial alcoholism. ([pubmed.ncbi.nlm.nih.gov](https://pubmed.ncbi.nlm.nih.gov/10195814/?utm_source=openai)) [S10]

Even the positive heroin study in Han Chinese is limited: the association was **haplotype-level only**, not single-marker; the odds ratios for rare recombinants were extremely large and therefore potentially unstable; and the authors themselves explicitly called for **replication**. ([pubmed.ncbi.nlm.nih.gov](https://pubmed.ncbi.nlm.nih.gov/26561861/?utm_source=openai)) [S11]

The epigenetic evidence is also **non-causal**. In the alcohol-nicotine codependence methylation study, the authors emphasized that methylation differences could reflect **predisposition, consequence of chronic exposure, or both**. Peripheral blood methylation is also not the same as demonstrating causal brain mechanisms. ([nature.com](https://www.nature.com/articles/srep41816)) [S12]

Overall, the literature around GABRB2 and addiction is dominated by **older candidate-gene**, **haplotype**, and **targeted methylation** studies rather than strong modern **genome-wide replicated** evidence. Reviews of substance-use genetics increasingly emphasize that many older candidate-gene findings did not replicate consistently, and in the GABA\_A family the best-supported addiction associations are more clearly with **GABRA2** than with **GABRB2**. ([pubmed.ncbi.nlm.nih.gov](https://pubmed.ncbi.nlm.nih.gov/24136292/)) [S13]

## Mechanistic / biological rationale

The mechanistic case is plausible but mostly **indirect**. GABRB2 encodes a core subunit of the major **GABA\_A receptor** population, and GABA\_A receptors are major mediators of inhibitory tone in circuits relevant to **reward, withdrawal, anxiety, impulsivity, and relapse**. Alcohol dependence research broadly supports GABAergic involvement, and pharmacologic modulation of GABA systems is relevant to treatment and withdrawal biology. ([ncbi.nlm.nih.gov](https://www.ncbi.nlm.nih.gov/gene/2561)) [S14]

Because alcohol and other addictive drugs drive neuroadaptations in inhibitory signaling, variation in a receptor subunit gene such as **GABRB2** could, in principle, alter receptor assembly, trafficking, stoichiometry, or network inhibition. That said, the literature linking addiction phenotypes to **specific functional GABRB2 variants** is thin: the 2014 meta-analysis explicitly noted that no commonly studied GABA receptor variants associated with substance dependence had yet been shown to be clearly **functional or pathogenic**. ([ncbi.nlm.nih.gov](https://www.ncbi.nlm.nih.gov/gene/2561)) [S15]

There is some supportive systems-level evidence that addiction-related states are accompanied by altered GABAergic regulation, and one alcohol-nicotine codependence study found differential **GABRB2 promoter methylation**. But direct causal experiments showing that altering **GABRB2 itself** changes addiction-like behavior are much less developed than for some other GABA receptor subunits. ([nature.com](https://www.nature.com/articles/srep41816)) [S16]

## Bottom line

**Yes, GABRB2 has been implicated in addiction in the literature—but only tentatively.** There are **positive candidate-gene, haplotype, and methylation studies** linking GABRB2 to alcohol dependence, heroin dependence, or alcohol-nicotine codependence. However, **larger and more rigorous evidence is inconsistent**, including a major family-based negative study and a meta-analysis in which stronger signals emerged for other GABA receptor genes, not GABRB2. ([jamanetwork.com](https://jamanetwork.com/journals/jamapsychiatry/fullarticle/208168)) [S17]

So the most accurate summary is: **GABRB2 is a biologically plausible and repeatedly suggested candidate in addiction, but it is not among the most robustly established addiction genes.** ([ncbi.nlm.nih.gov](https://www.ncbi.nlm.nih.gov/gene/2561)) [S18]

## Cited sources

- **[S1]** GABRB2 gamma-aminobutyric acid type A receptor subunit beta2 [Homo sapiens (human)] - Gene - NCBI — https://www.ncbi.nlm.nih.gov/gene/2561
- **[S2]** Haplotype-Based Localization of an Alcohol Dependence Gene to the 5q34 γ-Aminobutyric Acid Type A Gene Cluster | Genetics and Genomics | JAMA Psychiatry | JAMA Network — https://jamanetwork.com/journals/jamapsychiatry/fullarticle/208168
- **[S3]** Association between variants at the GABAAbeta2, GABAAalpha6 and GABAAgamma2 gene cluster and alcohol dependence in a Scottish population - PubMed — https://pubmed.ncbi.nlm.nih.gov/10578235/?utm_source=openai
- **[S4]** GABRB2 Haplotype Association with Heroin Dependence in Chinese Population - PubMed — https://pubmed.ncbi.nlm.nih.gov/26561861/?utm_source=openai
- **[S5]** Glutamatergic and GABAergic susceptibility loci for heroin and cocaine addiction in subjects of African and European ancestry - PMC — https://pmc.ncbi.nlm.nih.gov/articles/PMC4564302/?utm_source=openai
- **[S6]** Alcohol and nicotine codependence-associated DNA methylation changes in promoter regions of addiction-related genes | Scientific Reports — https://www.nature.com/articles/srep41816
- **[S7]** GABRB2, a key player in neuropsychiatric disorders and beyond - PubMed — https://pubmed.ncbi.nlm.nih.gov/34673206/
- **[S8]** Association of gamma-aminobutyric acid A receptor α2 gene (GABRA2) with alcohol use disorder - PubMed — https://pubmed.ncbi.nlm.nih.gov/24136292/
- **[S9]** No association of the GABAA receptor genes on chromosome 5 with alcoholism in the collaborative study on the genetics of alcoholism sample — https://www.downstate.edu/research/centers-departments/henri-begleiter-neurodynamics-laboratory/_documents/dick_2005-NoAssociationGABAAReceptorGenesonChromosome5withAlcoholismintheCollaborativeStudyo.pdf
- **[S10]** Association analysis of sequence variants of GABA(A) alpha6, beta2, and gamma2 gene cluster and alcohol dependence - PubMed — https://pubmed.ncbi.nlm.nih.gov/10195814/?utm_source=openai
- **[S11]** GABRB2 Haplotype Association with Heroin Dependence in Chinese Population - PubMed — https://pubmed.ncbi.nlm.nih.gov/26561861/?utm_source=openai
- **[S12]** Alcohol and nicotine codependence-associated DNA methylation changes in promoter regions of addiction-related genes | Scientific Reports — https://www.nature.com/articles/srep41816
- **[S13]** Association of gamma-aminobutyric acid A receptor α2 gene (GABRA2) with alcohol use disorder - PubMed — https://pubmed.ncbi.nlm.nih.gov/24136292/
- **[S14]** GABRB2 gamma-aminobutyric acid type A receptor subunit beta2 [Homo sapiens (human)] - Gene - NCBI — https://www.ncbi.nlm.nih.gov/gene/2561
- **[S15]** GABRB2 gamma-aminobutyric acid type A receptor subunit beta2 [Homo sapiens (human)] - Gene - NCBI — https://www.ncbi.nlm.nih.gov/gene/2561
- **[S16]** Alcohol and nicotine codependence-associated DNA methylation changes in promoter regions of addiction-related genes | Scientific Reports — https://www.nature.com/articles/srep41816
- **[S17]** Haplotype-Based Localization of an Alcohol Dependence Gene to the 5q34 γ-Aminobutyric Acid Type A Gene Cluster | Genetics and Genomics | JAMA Psychiatry | JAMA Network — https://jamanetwork.com/journals/jamapsychiatry/fullarticle/208168
- **[S18]** GABRB2 gamma-aminobutyric acid type A receptor subunit beta2 [Homo sapiens (human)] - Gene - NCBI — https://www.ncbi.nlm.nih.gov/gene/2561
