## Supplementary material for "Understanding Substance Dependence: What Differentiates Addictive from Non-Addictive Drugs?": Folder for supplementary files: addictive-genes-claims-first100-056-slc6a11-gene-protein-is-implicated-in-addiction-314315fa126f.md

# addictive-genes-claims-first100-056-314315fa126f

**Original claim:** SLC6A11 gene/protein is implicated in addiction.

## Claim

**SLC6A11 gene/protein is implicated in addiction.**

## Verdict

Verdict: Likely True

## Confidence

**Medium.**

## Supporting evidence

SLC6A11 encodes **GABA transporter 3 (GAT-3)**, a largely astrocyte-enriched transporter that helps clear extracellular GABA and regulate tonic/extrasynaptic inhibition. That function is biologically relevant to addiction circuits because altered inhibitory tone in the amygdala and related regions is strongly implicated in alcohol use disorder (AUD). ([ncbi.nlm.nih.gov](https://www.ncbi.nlm.nih.gov/gene/6538)) [S1]

The strongest direct evidence links **SLC6A11/GAT-3 to alcohol addiction-like behavior**, rather than to addiction broadly across all substances. In a 2018 *Science* study, rats given a mutually exclusive choice between alcohol and a high-value non-drug reward showed that the alcohol-preferring subgroup had **selectively decreased GAT-3 expression in the amygdala**. The same paper reported that **viral knockdown of Slc6a11** in rats that originally preferred saccharin shifted behavior toward alcohol choice, supporting a **causal** role in this alcohol-choice phenotype. The authors also found **downregulated GAT-3 in the central amygdala of postmortem brains from people with alcohol dependence**. ([pubmed.ncbi.nlm.nih.gov](https://pubmed.ncbi.nlm.nih.gov/29930131/)) [S2]

That causal animal evidence was followed by additional preclinical support. In a 2021 rat study, animals that persisted in drinking alcohol despite quinine adulteration or the availability of saccharin—behaviors often used as models of compulsive alcohol use—had **decreased GAT-3 mRNA in the central amygdala**, and baclofen preferentially reduced compulsive drinking in this subgroup. The paper explicitly proposed low GAT-3 as an **endophenotype of vulnerability** to compulsive alcohol drinking. ([pubmed.ncbi.nlm.nih.gov](https://pubmed.ncbi.nlm.nih.gov/33527681/)) [S3]

Human genetic evidence is weaker but not absent. A 2015 genome-wide meta-analysis of alcohol dependence reported an **African-ancestry-specific top-ranked SNP at SLC6A11** with p = 2.7 × 10^-7, which is suggestive and hypothesis-generating, though not definitive by modern GWAS standards. The same study concluded that SLC6A11 was among the genes appropriate for follow-up in alcohol dependence. ([pubmed.ncbi.nlm.nih.gov](https://pubmed.ncbi.nlm.nih.gov/26173551/?utm_source=openai)) [S4]

A recent review focused on astrocytic GABA regulation in alcohol-related disorders summarized this literature similarly: decreased amygdala GAT3/GAT-3 expression has been observed in alcohol-preferring animals, GAT3 knockdown can shift preference toward alcohol over saccharin, and low amygdala GAT3 has been linked to persistent alcohol seeking despite aversive adulteration. That review does not replace the primary studies, but it shows that the field now treats GAT-3 as a credible alcohol-relevant candidate mechanism. ([mdpi.com](https://www.mdpi.com/2073-4409/13/4/318)) [S5]

## Contradicting or limiting evidence

The main limitation is **scope**: the evidence is strongest for **alcohol addiction/AUD**, not for addiction in general. I did not find comparably strong primary evidence tying SLC6A11 to nicotine, opioid, stimulant, or polysubstance addiction in humans. So the claim becomes more accurate if narrowed to **alcohol-related addiction phenotypes**. ([pubmed.ncbi.nlm.nih.gov](https://pubmed.ncbi.nlm.nih.gov/29930131/)) [S6]

Also, not all alcohol models point in the same direction. A more recent rat study on ethanol dependence found that chronic ethanol exposure produced **increased astrocytic GAT3 expression during withdrawal**, and that **GAT3 knockdown or overexpression in central amygdala astrocytes did not change** withdrawal severity, dependence-escalated drinking, quinine-resistant drinking, or post-dependent drinking. The authors concluded that astrocytic GAT3, although altered by dependence, appeared **dispensable** for several dependence-related drinking phenotypes in that model. ([pmc.ncbi.nlm.nih.gov](https://pmc.ncbi.nlm.nih.gov/articles/PMC11195260/)) [S7]

That discrepancy suggests the role of SLC6A11/GAT-3 is probably **context-dependent**: it may matter more for **alcohol choice vulnerability or compulsive preference over alternative rewards** than for every measure of ethanol dependence. It may also depend on **cell type, circuit, brain subregion, withdrawal state, and task design**. This is an inference from the contrasting animal studies, rather than a single paper’s conclusion. ([ww2.uthscsa.edu](https://ww2.uthscsa.edu/artt/AddictionJC/2018-09-11-Augier.pdf)) [S8]

The human genetics case is also limited. The 2015 SLC6A11 signal was **suggestive**, not a universally replicated, genome-wide-significant finding across large studies. In contrast, a 2023 multi-ancestry AUD GWAS with over **1 million participants** identified **105 genome-wide significant variants from 37 loci**, emphasizing many other loci; taken together with the older report, this suggests SLC6A11 is **not yet an established major common-variant AUD locus**. That last point is an inference from comparing the older suggestive result with the newer very large GWAS. ([pmc.ncbi.nlm.nih.gov](https://pmc.ncbi.nlm.nih.gov/articles/PMC10775504/)) [S9]

## Mechanistic / biological rationale

Mechanistically, the claim is plausible. GAT-3 is an astrocytic GABA transporter that helps set extracellular GABA levels and tonic inhibition. Experimental work in hippocampus shows that GAT-3 contributes to control of **extracellular GABA** and becomes especially relevant when GABA spillover is high; inhibiting GAT-3 can augment extracellular GABA and tonic conductance. ([pmc.ncbi.nlm.nih.gov](https://pmc.ncbi.nlm.nih.gov/articles/PMC3678035/)) [S10]

In the alcohol-addiction studies, reduced amygdala GAT-3 was interpreted as causing **impaired GABA clearance**, thereby increasing local GABA tone. In the 2018 *Science* paper, alcohol-preferring rats showed decreased GAT-3 along with electrophysiologic evidence consistent with increased inhibitory tone in central amygdala, and knockdown of Slc6a11 reproduced synaptic effects and shifted behavior toward alcohol choice. ([ww2.uthscsa.edu](https://ww2.uthscsa.edu/artt/AddictionJC/2018-09-11-Augier.pdf)) [S11]

This fits broader addiction neurobiology because the **central amygdala** is a key hub for negative reinforcement, stress-related drinking, and compulsive alcohol seeking. If SLC6A11/GAT-3 alters inhibitory homeostasis there, it could change the salience of alcohol relative to non-drug rewards or alter vulnerability to compulsive use. However, the newer dependence study indicates that GAT3 changes may also represent a **homeostatic adaptation** rather than a universal driver of alcohol-related behavior. ([ww2.uthscsa.edu](https://ww2.uthscsa.edu/artt/AddictionJC/2018-09-11-Augier.pdf)) [S12]

## Bottom line

The claim is **best judged as likely true**, with an important qualifier: **SLC6A11/GAT-3 is implicated mainly in alcohol addiction-related phenotypes**, especially in preclinical studies of alcohol choice and compulsive drinking, and there is some translational support from human postmortem brain data. ([pubmed.ncbi.nlm.nih.gov](https://pubmed.ncbi.nlm.nih.gov/29930131/)) [S13]

What is **not** yet strongly established is that SLC6A11 is a broadly replicated **human addiction gene across substances** or a consistently validated common-variant risk locus in large modern GWAS. So: **implicated, yes; definitively established across addiction as a whole, no.** ([pubmed.ncbi.nlm.nih.gov](https://pubmed.ncbi.nlm.nih.gov/26173551/?utm_source=openai)) [S14]

## Cited sources

- **[S1]** SLC6A11 solute carrier family 6 member 11 [Homo sapiens (human)] - Gene - NCBI — https://www.ncbi.nlm.nih.gov/gene/6538
- **[S2]** A molecular mechanism for choosing alcohol over an alternative reward - PubMed — https://pubmed.ncbi.nlm.nih.gov/29930131/
- **[S3]** Baclofen decreases compulsive alcohol drinking in rats characterized by reduced levels of GAT-3 in the central amygdala - PubMed — https://pubmed.ncbi.nlm.nih.gov/33527681/
- **[S4]** A New Genomewide Association Meta-Analysis of Alcohol Dependence - PubMed — https://pubmed.ncbi.nlm.nih.gov/26173551/?utm_source=openai
- **[S5]** Astrocytic GABAergic Regulation in Alcohol Use and Major Depressive Disorders — https://www.mdpi.com/2073-4409/13/4/318
- **[S6]** A molecular mechanism for choosing alcohol over an alternative reward - PubMed — https://pubmed.ncbi.nlm.nih.gov/29930131/
- **[S7]** 
            Central Amygdala Astrocyte Plasticity Underlies GABAergic Dysregulation in Ethanol Dependence - PMC
         — https://pmc.ncbi.nlm.nih.gov/articles/PMC11195260/
- **[S8]** Science Journals — AAAS — https://ww2.uthscsa.edu/artt/AddictionJC/2018-09-11-Augier.pdf
- **[S9]** 
            Identification of Novel Loci and Cross-Disorder Pleiotropy Through Multi-Ancestry Genome-Wide Analysis of Alcohol Use Disorder in Over One Million Individuals - PMC
         — https://pmc.ncbi.nlm.nih.gov/articles/PMC10775504/
- **[S10]** 
            A functional role for both γ-aminobutyric acid (GABA) transporter-1 and GABA transporter-3 in the modulation of extracellular GABA and GABAergic tonic conductances in the rat hippocampus - PMC
         — https://pmc.ncbi.nlm.nih.gov/articles/PMC3678035/
- **[S11]** Science Journals — AAAS — https://ww2.uthscsa.edu/artt/AddictionJC/2018-09-11-Augier.pdf
- **[S12]** Science Journals — AAAS — https://ww2.uthscsa.edu/artt/AddictionJC/2018-09-11-Augier.pdf
- **[S13]** A molecular mechanism for choosing alcohol over an alternative reward - PubMed — https://pubmed.ncbi.nlm.nih.gov/29930131/
- **[S14]** A New Genomewide Association Meta-Analysis of Alcohol Dependence - PubMed — https://pubmed.ncbi.nlm.nih.gov/26173551/?utm_source=openai
