## Supplementary material for "Understanding Substance Dependence: What Differentiates Addictive from Non-Addictive Drugs?": Folder for supplementary files: addictive-genes-claims-first100-057-gabra5-gene-protein-is-implicated-in-addiction-3e93a86ebd69.md

# addictive-genes-claims-first100-057-3e93a86ebd69

**Original claim:** GABRA5 gene/protein is implicated in addiction.

## Claim

**“GABRA5 gene/protein is implicated in addiction.”**

## Verdict

Verdict: Likely True

The claim is **best supported for alcohol-related addiction phenotypes**, where multiple animal studies show that manipulating **GABRA5 / α5-containing GABA\_A receptors** changes alcohol reinforcement, alcohol seeking, and relapse-like drinking. Human evidence exists but is **weaker and less consistent**: there are suggestive candidate-gene and postmortem findings, but not strong, reproducible genome-wide human genetic evidence. So, as a broad statement that GABRA5 is “implicated” in addiction, the claim is reasonable; as a claim of **firmly established causal importance across all substance addictions**, it would overstate the evidence. ([ncbi.nlm.nih.gov](https://www.ncbi.nlm.nih.gov/gene/2558?utm_source=openai)) [S1]

## Confidence

**Medium.**

Confidence is medium because the preclinical evidence is fairly convergent, including mouse knockout, rat pharmacology, and nonhuman primate self-administration studies, but the **human evidence is mixed and mostly alcohol-specific** rather than broad across all addictive disorders. ([pubmed.ncbi.nlm.nih.gov](https://pubmed.ncbi.nlm.nih.gov/16253225/?utm_source=openai)) [S2]

## Supporting evidence

GABRA5 encodes the **α5 subunit of the GABA\_A receptor**, an inhibitory ligand-gated chloride channel subunit expressed in the brain. α5-containing GABA\_A receptors are especially noted for roles in hippocampal inhibition and cognition, making them biologically plausible candidates for drug-reward and cue-learning circuitry. ([ncbi.nlm.nih.gov](https://www.ncbi.nlm.nih.gov/gene/2558?utm_source=openai)) [S3]

A strong line of evidence comes from **mouse genetics**. In a targeted deletion study, **α5 knockout mice** showed reduced ethanol consumption at higher ethanol concentrations, and the α5-preferring inverse agonist **α5IA-II** dose-dependently reduced operant responding for ethanol without reducing responding for sucrose. That pattern argues against a merely nonspecific suppression of behavior and supports a role for α5-containing receptors in ethanol reward/reinforcement. ([pubmed.ncbi.nlm.nih.gov](https://pubmed.ncbi.nlm.nih.gov/16253225/?utm_source=openai)) [S4]

The evidence extends across species. In **rhesus monkeys**, the α5-selective agonist **QH-ii-066** increased alcohol drinking, whereas the α5 inverse agonist **L-655,708** decreased alcohol drinking; importantly, these effects were selective for alcohol versus sucrose and were reversible with an α5 antagonist. That is unusually strong pharmacologic evidence that α5-containing GABA\_A receptors modulate alcohol’s reinforcing effects in a primate model. ([pmc.ncbi.nlm.nih.gov](https://pmc.ncbi.nlm.nih.gov/articles/PMC3951841/?utm_source=openai)) [S5]

Rodent relapse and cue-reactivity models point in the same direction. In male rats, **L-655,708 attenuated cue-induced reinstatement** of alcohol seeking, while α5 agonist treatment could enhance alcohol-seeking behavior. The same study also found that α5 inverse agonists selectively reduced alcohol+sucrose self-administration more than sucrose alone, again suggesting a specific link to alcohol reinforcement rather than generalized motor suppression or reward blunting. ([rti.org](https://www.rti.org/publication/alpha-5gaba-subunit-containing-receptors-sweetened-alcohol-cue-induced-reinstatement-active-sweetene?utm_source=openai)) [S6]

A later rat study using the **alcohol deprivation effect**, a common relapse-like model, found that low doses of the α5 agonist **enhanced** relapse-like drinking, whereas **L-655,708 dose-dependently inhibited** it. This strengthens the case that α5 receptor signaling is involved not only in ongoing alcohol intake, but also in **relapse-related behavior**. ([pmc.ncbi.nlm.nih.gov](https://pmc.ncbi.nlm.nih.gov/articles/PMC8511242/?utm_source=openai)) [S7]

There is also **suggestive human evidence**. A 2003 family-based study from COGA reported association between alcoholism and the chromosome 15 GABA\_A cluster, with the strongest signal in paternal transmission analyses for **GABRA5** under a more severe ICD-10 alcohol dependence phenotype. Separately, a postmortem human brain study found **lower GABA\_A α5 mRNA expression in frontal cortex** of alcoholics compared with controls, consistent with long-term alcohol-related neuroadaptation involving this receptor subunit. ([downstate.edu](https://www.downstate.edu/research/centers-departments/henri-begleiter-neurodynamics-laboratory/_documents/pdf/2003-Song-Association%20of%20GABAA%20Receptors%20and%20Alcohol%20Dependence%20and%20the%20Effects%20of%20Genetic%20Imprinting.pdf)) [S8]

Additional molecular studies are compatible with that interpretation. In a rat ethanol-withdrawal model, **Gabra5 mRNA increased in the ventral tegmental area** during withdrawal, suggesting dynamic regulation of this subunit during dependence/withdrawal states in a core reward-region. ([pubmed.ncbi.nlm.nih.gov](https://pubmed.ncbi.nlm.nih.gov/30103280/?utm_source=openai)) [S9]

## Contradicting or limiting evidence

The clearest limitation is that **human genetics is inconsistent**. A subsequent COGA family-based study tested GABRA5, GABRB3, and GABRG3 and found **no consistent evidence of association for GABRA5**, instead finding the more reproducible signal in **GABRG3** and no parent-of-origin effect. That directly tempers the earlier positive candidate-gene result. ([pubmed.ncbi.nlm.nih.gov](https://pubmed.ncbi.nlm.nih.gov/14745296/?utm_source=openai)) [S10]

Also, large GWAS of alcohol dependence/AUD have reliably highlighted loci such as **ADH1B** and some other non-GABRA5 signals; **GABRA5 is not prominently reported in the genome-wide significant findings in these abstracts**. That does not prove no effect, but it suggests any human genetic effect of GABRA5 is likely **modest, context-dependent, or hard to detect reproducibly**. This is an inference from the published GWAS summaries, not a direct negative test of every GABRA5 variant. ([nature.com](https://www.nature.com/articles/mp2013145?utm_source=openai)) [S11]

The claim is also too broad if interpreted as applying equally to **all addictions**. In this search, the strongest direct primary evidence was for **alcohol-related** phenotypes. I did **not** find comparably strong, direct human or preclinical evidence tying GABRA5 specifically to nicotine, opioid, stimulant, or cannabis addiction. So “implicated in addiction” is defensible mainly if addiction includes **alcohol use disorder**, not as a well-established pan-addiction gene. ([pmc.ncbi.nlm.nih.gov](https://pmc.ncbi.nlm.nih.gov/articles/PMC3951841/?utm_source=openai)) [S12]

## Mechanistic / biological rationale

α5-containing GABA\_A receptors are concentrated in the hippocampus and are important mediators of **tonic inhibition**, with both extrasynaptic and some synaptic localization. Because tonic inhibition shapes neuronal excitability, these receptors are positioned to influence learning, memory, contextual processing, and inhibitory tone in circuits relevant to substance reinforcement and relapse. ([pubmed.ncbi.nlm.nih.gov](https://pubmed.ncbi.nlm.nih.gov/14993607/?utm_source=openai)) [S13]

That matters for addiction biology because alcohol-related behaviors are driven not only by acute reward, but also by **contextual memory, cue reactivity, and relapse circuitry**. Background sections of the rat alcohol-seeking literature note α5-rich ventral hippocampal CA1/CA3 pathways projecting to the nucleus accumbens, ventral tegmental area, amygdala, and hypothalamus—regions central to reward, motivation, and cue-triggered drug seeking. Pharmacologic modulation of α5 receptors changes alcohol seeking, self-administration, and relapse-like drinking in exactly those kinds of behavioral paradigms. ([pmc.ncbi.nlm.nih.gov](https://pmc.ncbi.nlm.nih.gov/articles/PMC6606346/?utm_source=openai)) [S14]

A plausible biological model, therefore, is that **GABRA5-dependent inhibitory signaling helps tune the salience of alcohol-associated cues and the reinforcing effects of alcohol**, while chronic alcohol exposure induces compensatory changes in GABRA5 expression/function that may contribute to dependence and withdrawal-related states. The withdrawal-associated VTA changes in rats and altered cortical expression in human alcoholics are consistent with that model, though they do not by themselves establish causality. ([pubmed.ncbi.nlm.nih.gov](https://pubmed.ncbi.nlm.nih.gov/30103280/?utm_source=openai)) [S15]

## Bottom line

**GABRA5 is reasonably implicated in addiction, but mainly in alcohol-related addiction.** The most convincing evidence comes from **preclinical studies**: knockout mice, rat self-administration/reinstatement/relapse models, and rhesus monkey alcohol self-administration all support a role for **α5-containing GABA\_A receptors** in alcohol reinforcement and relapse-like behavior. Human evidence is **suggestive but mixed**, with inconsistent candidate-gene associations and limited direct clinical validation. So the fairest summary is: **likely true for alcohol addiction biology; not yet firmly established as a broad, general addiction gene across substances.** ([pubmed.ncbi.nlm.nih.gov](https://pubmed.ncbi.nlm.nih.gov/16253225/?utm_source=openai)) [S16]

## Cited sources

- **[S1]** GABRA5 gamma-aminobutyric acid type A receptor subunit alpha5 [Homo sapiens (human)] - Gene - NCBI — https://www.ncbi.nlm.nih.gov/gene/2558?utm_source=openai
- **[S2]** Role of GABAA alpha5-containing receptors in ethanol reward: the effects of targeted gene deletion, and a selective inverse agonist - PubMed — https://pubmed.ncbi.nlm.nih.gov/16253225/?utm_source=openai
- **[S3]** GABRA5 gamma-aminobutyric acid type A receptor subunit alpha5 [Homo sapiens (human)] - Gene - NCBI — https://www.ncbi.nlm.nih.gov/gene/2558?utm_source=openai
- **[S4]** Role of GABAA alpha5-containing receptors in ethanol reward: the effects of targeted gene deletion, and a selective inverse agonist - PubMed — https://pubmed.ncbi.nlm.nih.gov/16253225/?utm_source=openai
- **[S5]** Modulation of α5 subunit-containing GABAA receptors alters alcohol drinking by rhesus monkeys - PMC — https://pmc.ncbi.nlm.nih.gov/articles/PMC3951841/?utm_source=openai
- **[S6]** alpha 5GABA(A) subunit-containing receptors and sweetened alcohol cue-induced reinstatement and active sweetened alcohol self-administration in male rats | RTI — https://www.rti.org/publication/alpha-5gaba-subunit-containing-receptors-sweetened-alcohol-cue-induced-reinstatement-active-sweetene?utm_source=openai
- **[S7]** Modulation of relapse-like drinking in male Sprague-Dawley rats by ligands targeting the α5GABAA receptor - PMC — https://pmc.ncbi.nlm.nih.gov/articles/PMC8511242/?utm_source=openai
- **[S8]** Untitled source — https://www.downstate.edu/research/centers-departments/henri-begleiter-neurodynamics-laboratory/_documents/pdf/2003-Song-Association%20of%20GABAA%20Receptors%20and%20Alcohol%20Dependence%20and%20the%20Effects%20of%20Genetic%20Imprinting.pdf
- **[S9]** Histone Deacetylase Inhibitor Suberanilohydroxamic Acid Treatment Reverses Hyposensitivity to γ-Aminobutyric Acid in the Ventral Tegmental Area During Ethanol Withdrawal - PubMed — https://pubmed.ncbi.nlm.nih.gov/30103280/?utm_source=openai
- **[S10]** Association of GABRG3 with alcohol dependence - PubMed — https://pubmed.ncbi.nlm.nih.gov/14745296/?utm_source=openai
- **[S11]** Genome-wide association study of alcohol dependence:significant findings in African- and European-Americans including novel risk loci | Molecular Psychiatry — https://www.nature.com/articles/mp2013145?utm_source=openai
- **[S12]** Modulation of α5 subunit-containing GABAA receptors alters alcohol drinking by rhesus monkeys - PMC — https://pmc.ncbi.nlm.nih.gov/articles/PMC3951841/?utm_source=openai
- **[S13]** Tonic inhibition in mouse hippocampal CA1 pyramidal neurons is mediated by alpha5 subunit-containing gamma-aminobutyric acid type A receptors - PubMed — https://pubmed.ncbi.nlm.nih.gov/14993607/?utm_source=openai
- **[S14]** α5GABAA subunit-containing receptors and sweetened alcohol cue-induced reinstatement and active sweetened alcohol self-administration in male rats - PMC — https://pmc.ncbi.nlm.nih.gov/articles/PMC6606346/?utm_source=openai
- **[S15]** Histone Deacetylase Inhibitor Suberanilohydroxamic Acid Treatment Reverses Hyposensitivity to γ-Aminobutyric Acid in the Ventral Tegmental Area During Ethanol Withdrawal - PubMed — https://pubmed.ncbi.nlm.nih.gov/30103280/?utm_source=openai
- **[S16]** Role of GABAA alpha5-containing receptors in ethanol reward: the effects of targeted gene deletion, and a selective inverse agonist - PubMed — https://pubmed.ncbi.nlm.nih.gov/16253225/?utm_source=openai
