## Supplementary material for "Understanding Substance Dependence: What Differentiates Addictive from Non-Addictive Drugs?": Folder for supplementary files: addictive-genes-claims-first100-058-ppp2r2c-gene-protein-is-implicated-in-addiction-15791bce5e76.md

# addictive-genes-claims-first100-058-15791bce5e76

**Original claim:** PPP2R2C gene/protein is implicated in addiction.

## Claim

PPP2R2C gene/protein is implicated in addiction.

## Verdict

Verdict: No Evidence

## Confidence

Medium

## Supporting evidence

PPP2R2C is a **biologically plausible brain gene**, but that is not the same as being implicated in addiction. It encodes a PP2A regulatory B55γ subunit that was originally described as **highly brain-enriched** and nearly undetectable in most other tissues. Human Protein Atlas data also classify it as **brain-enhanced**, with enrichment in neuronal cell-type clusters related to synaptic function/neurogenesis. ([pubmed.ncbi.nlm.nih.gov](https://pubmed.ncbi.nlm.nih.gov/10945473/)) [S1]

There is also evidence that PPP2R2C has **important neural functions**. A familial chromosomal rearrangement disrupting PPP2R2C was reported in autosomal dominant intellectual disability with epilepsy/behavioral problems, and the authors noted a mouse-brain expression pattern consistent with a role in **synaptic plasticity, learning, and memory**. More recently, mouse work showed that Ppp2r2c is enriched at the **axon initial segment (AIS)** and can modulate neuronal firing and Kv1.2 phosphorylation, supporting a role in neuronal excitability/homeostasis. ([pubmed.ncbi.nlm.nih.gov](https://pubmed.ncbi.nlm.nih.gov/20601260/)) [S2]

PPP2R2C also has an **indirect signaling link** to dopamine biology: in a non-neural system, PPP2R2C co-immunoprecipitated with the dopamine D2 receptor and regulated PP2A/Akt/GSK3β signaling downstream of D2R. Because D2 receptor signaling is central to addiction neurobiology, this gives PPP2R2C some **mechanistic plausibility** as a candidate worth studying. However, this is still indirect and not addiction evidence per se. ([pubmed.ncbi.nlm.nih.gov](https://pubmed.ncbi.nlm.nih.gov/26290374/)) [S3]

## Contradicting or limiting evidence

I did **not find direct peer-reviewed evidence** that PPP2R2C is a replicated addiction gene/protein in humans or a validated causal mediator in animal addiction models. In several representative, high-profile addiction-genetics sources reviewed here, searching within the papers found **no mention of PPP2R2C**: a 2024 JCI paper on generalized genetic liability to substance use disorders, a 2025 multi-ancestral GWAS of clinically defined nicotine dependence, and a review of alcohol-dependence GWAS discoveries. That does not prove PPP2R2C has no role, but it argues against it being a currently established addiction locus. ([jci.org](https://www.jci.org/articles/view/172881)) [S4]

The PPP2R2C literature that surfaced most readily was centered on **other phenotypes**—for example Alzheimer-related tau dephosphorylation, brain aging/neural senescence, Parkinson/LRRK2 signaling, cancer biology, and neurodevelopmental phenotypes—rather than addiction, substance use disorder, or drug reward/withdrawal models. ([pubmed.ncbi.nlm.nih.gov](https://pubmed.ncbi.nlm.nih.gov/32291379/?utm_source=openai)) [S5]

Even some of the more relevant mechanistic studies do **not** support a simple dopamine/addiction interpretation. In the zebrafish ppp2r2c-loss model, the mutant animals had behavioral abnormalities, but measured brain catecholamines—including **dopamine**—were reportedly unchanged. That weakens any claim that PPP2R2C has already been shown to alter addiction-relevant dopamine tone directly. ([pmc.ncbi.nlm.nih.gov](https://pmc.ncbi.nlm.nih.gov/articles/PMC10014060/)) [S6]

So, the main limitation is not contradictory positive data; it is the **absence of direct addiction-focused evidence**. Right now, PPP2R2C looks more like a **neurobiologically interesting candidate** than an addiction-implicated gene established by genetics, transcriptomics, or functional addiction experiments. ([jci.org](https://www.jci.org/articles/view/172881)) [S7]

## Mechanistic / biological rationale

There is a reasonable **biological rationale** for why PPP2R2C could matter in addiction, even though the direct evidence is missing. Addiction-related plasticity depends heavily on phosphorylation state, dopaminergic signaling, and changes in neuronal excitability within cortico-striatal and mesolimbic circuits. PPP2R2C encodes a **brain-enriched PP2A regulatory subunit**, and PP2A complexes are major controllers of phospho-signaling. ([pubmed.ncbi.nlm.nih.gov](https://pubmed.ncbi.nlm.nih.gov/10945473/)) [S8]

Recent work places Ppp2r2c at the **AIS**, where it can alter phosphorylation of Kv1.2 and increase neuronal network activity when overexpressed. In principle, proteins that shape excitability and activity homeostasis can influence reward learning, reinforcement, and drug-induced plasticity. That is a legitimate mechanistic bridge—but still an **inference**, not direct addiction evidence. ([pmc.ncbi.nlm.nih.gov](https://pmc.ncbi.nlm.nih.gov/articles/PMC12675759/)) [S9]

Likewise, the reported PPP2R2C-D2R-PP2A/Akt/GSK3β interaction is interesting because D2 signaling is strongly tied to reward dysfunction and compulsive behavior. But the published PPP2R2C-D2R study was performed in **renal proximal tubule cells**, not in nucleus accumbens/striatum or an addiction model, so extrapolation to addiction remains speculative. ([pubmed.ncbi.nlm.nih.gov](https://pubmed.ncbi.nlm.nih.gov/26290374/)) [S10]

Overall, the mechanism says **“possible and worth testing”**, not **“already implicated.”** ([pmc.ncbi.nlm.nih.gov](https://pmc.ncbi.nlm.nih.gov/articles/PMC12675759/)) [S11]

## Bottom line

At present, the claim that **PPP2R2C is implicated in addiction** is **not supported by direct evidence** from the main human addiction-genetics and addiction-biology literature I reviewed. PPP2R2C is clearly a **brain-relevant phosphatase regulatory gene** with roles in synaptic plasticity, neuronal excitability, and signaling pathways that are broadly relevant to addiction, so it is a **plausible candidate**. But plausibility is not the same as implication. Based on current accessible evidence, PPP2R2C should be described as an **indirectly plausible but unestablished addiction candidate**, not as a gene/protein already implicated in addiction. ([pubmed.ncbi.nlm.nih.gov](https://pubmed.ncbi.nlm.nih.gov/10945473/)) [S12]

## Cited sources

- **[S1]** Molecular cloning and mapping of the brain-abundant B1gamma subunit of protein phosphatase 2A, PPP2R2C, to human chromosome 4p16 - PubMed — https://pubmed.ncbi.nlm.nih.gov/10945473/
- **[S2]** PPP2R2C, a gene disrupted in autosomal dominant intellectual disability - PubMed — https://pubmed.ncbi.nlm.nih.gov/20601260/
- **[S3]** Dopamine D2 receptors' effects on renal inflammation are mediated by regulation of PP2A function - PubMed — https://pubmed.ncbi.nlm.nih.gov/26290374/
- **[S4]** 
JCI -
Generalized genetic liability to substance use disorders
 — https://www.jci.org/articles/view/172881
- **[S5]** PP2A subunit PPP2R2C is downregulated in the brains of Alzheimer's transgenic mice - PubMed — https://pubmed.ncbi.nlm.nih.gov/32291379/?utm_source=openai
- **[S6]** 
            Protein phosphatase 2A activators reverse age‐related behavioral changes by targeting neural cell senescence - PMC
         — https://pmc.ncbi.nlm.nih.gov/articles/PMC10014060/
- **[S7]** 
JCI -
Generalized genetic liability to substance use disorders
 — https://www.jci.org/articles/view/172881
- **[S8]** Molecular cloning and mapping of the brain-abundant B1gamma subunit of protein phosphatase 2A, PPP2R2C, to human chromosome 4p16 - PubMed — https://pubmed.ncbi.nlm.nih.gov/10945473/
- **[S9]** 
            A distinct PP2A subunit regulates local protein phosphorylation at the axon initial segment - PMC
         — https://pmc.ncbi.nlm.nih.gov/articles/PMC12675759/
- **[S10]** Dopamine D2 receptors' effects on renal inflammation are mediated by regulation of PP2A function - PubMed — https://pubmed.ncbi.nlm.nih.gov/26290374/
- **[S11]** 
            A distinct PP2A subunit regulates local protein phosphorylation at the axon initial segment - PMC
         — https://pmc.ncbi.nlm.nih.gov/articles/PMC12675759/
- **[S12]** Molecular cloning and mapping of the brain-abundant B1gamma subunit of protein phosphatase 2A, PPP2R2C, to human chromosome 4p16 - PubMed — https://pubmed.ncbi.nlm.nih.gov/10945473/
