## Supplementary material for "Understanding Substance Dependence: What Differentiates Addictive from Non-Addictive Drugs?": Folder for supplementary files: addictive-genes-claims-first100-059-prkcz-gene-protein-is-implicated-in-addiction-b5beb0f402b3.md

# addictive-genes-claims-first100-059-b5beb0f402b3

**Original claim:** PRKCZ gene/protein is implicated in addiction.

## Claim

PRKCZ gene/protein is implicated in addiction.

## Verdict

Verdict: Likely True

## Confidence

Medium

## Supporting evidence

**What PRKCZ is.** PRKCZ encodes atypical protein kinase C-ζ, and the same locus also gives rise to the brain-enriched catalytic isoform **PKMζ**, which has been studied extensively in synaptic plasticity and memory. For addiction-related claims, much of the literature is really about **PKMζ**, a product of the **Prkcz/PRKCZ** locus, rather than full-length PKCζ alone. ([ncbi.nlm.nih.gov](https://www.ncbi.nlm.nih.gov/gene/5590?utm_source=openai)) [S1]

**Rodent cocaine evidence is fairly consistent.** In mice, chronic cocaine self-administration increased PKMζ mRNA and protein in the **nucleus accumbens**, a core reward region, and constitutive deletion of PKMζ increased cocaine self-administration and cue-induced reinstatement; nucleus-accumbens-specific deletion recapitulated the increase in cocaine taking/seeking in males. The authors concluded that PKMζ acts to **dampen** cocaine taking and seeking rather than promote it. That still counts as implication in addiction biology, because altering the gene changes addiction-relevant behavior. ([pubmed.ncbi.nlm.nih.gov](https://pubmed.ncbi.nlm.nih.gov/30111812/?utm_source=openai)) [S2]

**Additional psychostimulant work supports a role in drug-induced neuroadaptation.** In rats, inhibition of PKMζ signaling in the mesolimbic system altered AMPA currents, AMPA/NMDA ratio, and the expression of cocaine locomotor sensitization, with region- and time-specific effects in the VTA and nucleus accumbens. Sensitization is not identical to addiction, but it is a widely used preclinical model of drug-induced plasticity relevant to addiction mechanisms. ([pmc.ncbi.nlm.nih.gov](https://pmc.ncbi.nlm.nih.gov/articles/PMC3980506/?utm_source=openai)) [S3]

**Alcohol-related evidence also supports involvement.** In mice, deletion of Prkcz increased intermittent ethanol consumption, and intermittent ethanol exposure increased PKMζ protein in ventral striatum. The authors interpreted this as a negative-feedback role in limiting binge-like ethanol intake. Earlier expression work also identified Prkcz as positively correlated with ethanol consumption and alcohol-responsive in mouse models. ([pubmed.ncbi.nlm.nih.gov](https://pubmed.ncbi.nlm.nih.gov/23905844/?utm_source=openai)) [S4]

**Opioid-related evidence extends the signal beyond stimulants and alcohol.** In rats, intra-accumbens ZIP inhibition of PKCζ/PKMζ disrupted retrieval, consolidation, and reconsolidation of **remifentanil-associated drug memory** and drug seeking. More recently, constitutive PKMζ deletion altered **oxycodone** self-administration, motivation, and sensitization in dose- and sex-dependent ways, with the abstract concluding that PKMζ dampens oxycodone taking. ([pubmed.ncbi.nlm.nih.gov](https://pubmed.ncbi.nlm.nih.gov/22348011/?utm_source=openai)) [S5]

**Addiction-related memory studies are especially notable.** A frequently cited rat study found that inhibiting PKMζ activity in nucleus accumbens core abolished long-term conditioned place preference memories for **morphine** and **cocaine**, suggesting a role in maintaining drug-cue reward memories. Another study found PKMζ contributes to morphine reward/aversion and extinction memory in amygdala/infralimbic cortex circuits. These are addiction-relevant processes because persistent drug-cue memories are central to relapse vulnerability. ([pmc.ncbi.nlm.nih.gov](https://pmc.ncbi.nlm.nih.gov/articles/PMC3150199/?utm_source=openai)) [S6]

**Human evidence exists but is much weaker and more indirect.** I did not find strong, replicated evidence that PRKCZ is a major direct human addiction-risk gene. The clearest human signal I found was an integrative multi-omics study nominating **PRKCZ** among genes shared between **schizophrenia and smoking behaviors**, based on SNP, methylation, and expression analyses. That supports possible relevance to smoking-related phenotypes, but it is indirect and not the same as demonstrating PRKCZ as a robust standalone addiction gene in humans. ([pubmed.ncbi.nlm.nih.gov](https://pubmed.ncbi.nlm.nih.gov/32012119/?utm_source=openai)) [S7]

## Contradicting or limiting evidence

**Most evidence is preclinical, not clinical.** The case for PRKCZ in addiction rests mainly on rodent knockout, expression, and intracranial inhibition studies. That is enough to say the locus is *implicated* in addiction biology, but not enough to call it a firmly established human addiction gene or therapeutic target. ([pubmed.ncbi.nlm.nih.gov](https://pubmed.ncbi.nlm.nih.gov/23905844/?utm_source=openai)) [S8]

**A major caveat is that many classic studies used ZIP, which is not fully specific for PKMζ/PRKCZ products.** Later work showed that ZIP can still reverse LTP or memory-related phenomena in PKMζ/Prkcz knockout mice, and biochemical studies indicate it can affect other PKC-family isoforms or act through mechanisms beyond selective PKMζ inhibition. Therefore, older “PKMζ is necessary” claims from ZIP-only addiction-memory studies should be interpreted cautiously. ([pubmed.ncbi.nlm.nih.gov](https://pubmed.ncbi.nlm.nih.gov/23283174/?utm_source=openai)) [S9]

**The broader PKMζ memory field has been controversial.** Nature papers reported that Prkcz-null mice show normal learning and memory, challenging the idea that PKMζ is uniquely required for long-term memory. Follow-up work suggested compensation by the related atypical kinase **PKCι/λ** in knockout animals. This does not erase the addiction findings, but it does weaken overly simple claims that PRKCZ alone is the indispensable molecular substrate of drug memory. ([pubmed.ncbi.nlm.nih.gov](https://pubmed.ncbi.nlm.nih.gov/23283171/)) [S10]

**Directionality is complex.** Several of the stronger studies suggest PKMζ may act as a **brake** or homeostatic counterweight on drug taking/seeking rather than a pro-addiction driver. So “implicated in addiction” is fair, but “causes addiction” or “promotes addiction” would overstate the evidence. ([pubmed.ncbi.nlm.nih.gov](https://pubmed.ncbi.nlm.nih.gov/30111812/?utm_source=openai)) [S11]

## Mechanistic / biological rationale

A biologically plausible rationale exists. Addiction-related behavior is strongly linked to **glutamatergic plasticity** in the **nucleus accumbens**, including AMPA receptor trafficking and shifts in GluA2-containing vs calcium-permeable AMPA receptors. Reviews of addiction neurobiology emphasize that these glutamate/AMPAR processes are central to cocaine seeking and relapse-like behavior. ([pmc.ncbi.nlm.nih.gov](https://pmc.ncbi.nlm.nih.gov/articles/PMC4931870/)) [S12]

PKMζ is mechanistically positioned to affect exactly these processes. In neuronal systems, PKMζ promotes synaptic clustering of PSD-95 and increases **synaptic GluA2-containing AMPA receptors**. Because cocaine- and cue-induced seeking are sensitive to manipulations of AMPAR and GluA2 trafficking in the accumbens, it is biologically plausible that PRKCZ-derived PKMζ modulates addiction-related plasticity and cue memory through regulation of postsynaptic AMPAR composition. ([pubmed.ncbi.nlm.nih.gov](https://pubmed.ncbi.nlm.nih.gov/22378468/)) [S13]

Putting those findings together, the most defensible interpretation is that PRKCZ/PKMζ is implicated mainly in **addiction-related synaptic plasticity, drug-cue memory, and homeostatic regulation of reward seeking**, rather than being proven as a primary human susceptibility gene. That last step is an inference from the combined animal behavioral and synaptic evidence, not a direct human clinical demonstration. ([pubmed.ncbi.nlm.nih.gov](https://pubmed.ncbi.nlm.nih.gov/30111812/?utm_source=openai)) [S14]

## Bottom line

If **PRKCZ** is interpreted to include its brain product **PKMζ**, then the claim is **likely true**: multiple rodent studies across **cocaine, alcohol, and opioids** show that manipulating this locus changes addiction-relevant behaviors and drug-associated memory, especially in mesolimbic circuits such as the **nucleus accumbens**. ([pubmed.ncbi.nlm.nih.gov](https://pubmed.ncbi.nlm.nih.gov/23905844/?utm_source=openai)) [S15]

However, the evidence is **mostly animal/preclinical**, some influential experiments relied on the imperfect inhibitor **ZIP**, and direct **human genetic/clinical evidence is sparse and indirect**. So PRKCZ is best described as **implicated in addiction-related neuroplasticity**, not as a definitively established human addiction gene. ([pubmed.ncbi.nlm.nih.gov](https://pubmed.ncbi.nlm.nih.gov/23283174/?utm_source=openai)) [S16]

## Cited sources

- **[S1]** PRKCZ protein kinase C zeta [Homo sapiens (human)] - Gene - NCBI — https://www.ncbi.nlm.nih.gov/gene/5590?utm_source=openai
- **[S2]** PKMζ in the nucleus accumbens acts to dampen cocaine seeking - PubMed — https://pubmed.ncbi.nlm.nih.gov/30111812/?utm_source=openai
- **[S3]** Inhibition of Protein kinase Mzeta (PKMζ) in the mesolimbic system alters cocaine sensitization in rats - PMC — https://pmc.ncbi.nlm.nih.gov/articles/PMC3980506/?utm_source=openai
- **[S4]** Deletion of Prkcz increases intermittent ethanol consumption in mice - PubMed — https://pubmed.ncbi.nlm.nih.gov/23905844/?utm_source=openai
- **[S5]** Activation of PKCzeta and PKMzeta in the nucleus accumbens core is necessary for the retrieval, consolidation and reconsolidation of drug memory - PubMed — https://pubmed.ncbi.nlm.nih.gov/22348011/?utm_source=openai
- **[S6]** Inhibition of PKMζ in Nucleus Accumbens Core Abolishes Long-Term Drug Reward Memory - PMC — https://pmc.ncbi.nlm.nih.gov/articles/PMC3150199/?utm_source=openai
- **[S7]** Identification of 34 genes conferring genetic and pharmacological risk for the comorbidity of schizophrenia and smoking behaviors - PubMed — https://pubmed.ncbi.nlm.nih.gov/32012119/?utm_source=openai
- **[S8]** Deletion of Prkcz increases intermittent ethanol consumption in mice - PubMed — https://pubmed.ncbi.nlm.nih.gov/23905844/?utm_source=openai
- **[S9]** PKM-ζ is not required for hippocampal synaptic plasticity, learning and memory - PubMed — https://pubmed.ncbi.nlm.nih.gov/23283174/?utm_source=openai
- **[S10]** Prkcz null mice show normal learning and memory - PubMed — https://pubmed.ncbi.nlm.nih.gov/23283171/
- **[S11]** PKMζ in the nucleus accumbens acts to dampen cocaine seeking - PubMed — https://pubmed.ncbi.nlm.nih.gov/30111812/?utm_source=openai
- **[S12]** 
            The Nucleus Accumbens: Mechanisms of Addiction across Drug Classes Reflect the Importance of Glutamate Homeostasis - PMC
         — https://pmc.ncbi.nlm.nih.gov/articles/PMC4931870/
- **[S13]** PKMζ is necessary and sufficient for synaptic clustering of PSD-95 - PubMed — https://pubmed.ncbi.nlm.nih.gov/22378468/
- **[S14]** PKMζ in the nucleus accumbens acts to dampen cocaine seeking - PubMed — https://pubmed.ncbi.nlm.nih.gov/30111812/?utm_source=openai
- **[S15]** Deletion of Prkcz increases intermittent ethanol consumption in mice - PubMed — https://pubmed.ncbi.nlm.nih.gov/23905844/?utm_source=openai
- **[S16]** PKM-ζ is not required for hippocampal synaptic plasticity, learning and memory - PubMed — https://pubmed.ncbi.nlm.nih.gov/23283174/?utm_source=openai
