## Supplementary material for "Understanding Substance Dependence: What Differentiates Addictive from Non-Addictive Drugs?": Folder for supplementary files: addictive-genes-claims-first100-060-tecr-gene-protein-is-implicated-in-addiction-c80ad298704f.md

# addictive-genes-claims-first100-060-c80ad298704f

**Original claim:** TECR gene/protein is implicated in addiction.

## Claim

**TECR gene/protein is implicated in addiction.**

## Verdict

Verdict: No Evidence

## Confidence

**Medium**

## Supporting evidence

TECR encodes **trans-2,3-enoyl-CoA reductase**, an **endoplasmic-reticulum membrane enzyme** involved in the final reduction step of **long- and very-long-chain fatty-acid elongation** and in sphingolipid-related lipid metabolism. Authoritative gene resources describe it primarily as a **lipid-metabolic enzyme**, not as an established addiction gene. ([ncbi.nlm.nih.gov](https://www.ncbi.nlm.nih.gov/gene/9524?utm_source=openai)) [S1]

There is also some bona fide **neurobiological relevance** for TECR outside addiction. In human/cell studies, TECR directly interacts with **SERCA2b** and acts as a **negative regulator of ER Ca²⁺ accumulation**, linking lipid metabolism to intracellular calcium signaling. ClinGen’s TECR curation notes this as potentially relevant to **synaptic potentiation, learning, and memory**. ([pubmed.ncbi.nlm.nih.gov](https://pubmed.ncbi.nlm.nih.gov/33482198/?utm_source=openai)) [S2]

A separate mouse/endothelial study found that Tecr supports **blood-brain barrier homeostasis**: endothelial Tecr was enriched during barriergenesis, and endothelial knockout increased **BBB permeability/transcytosis** while altering omega-3–related lipid profiles. That shows TECR can affect CNS-relevant biology, but it is still not addiction-specific evidence. ([pubmed.ncbi.nlm.nih.gov](https://pubmed.ncbi.nlm.nih.gov/35465346/)) [S3]

Human disease evidence for TECR is currently centered on **autosomal recessive intellectual disability**, not substance use disorder. ClinGen lists TECR–intellectual disability as a curated gene-disease relationship with **limited** evidence, and the original family-based report linked TECR mutation to nonsyndromic intellectual disability in a Hutterite family. ([search.clinicalgenome.org](https://search.clinicalgenome.org/kb/gene-validity/CGGV%3Aassertion_8c7adee2-6a14-4348-a018-9097954bd39d-2022-03-10T070000.000Z)) [S4]

## Contradicting or limiting evidence

I did **not** find direct primary evidence that TECR is associated with addiction in humans or animal models. In a large cross-substance GWAS/meta-analysis covering problematic alcohol use, problematic tobacco use, cannabis use disorder, and opioid use disorder in over **1 million** individuals, the reported addiction-risk loci included genes such as **PDE4B** and substance-specific receptor/metabolic genes; a search of the article text did **not** identify **TECR**. ([nature.com](https://www.nature.com/articles/s44220-023-00034-y)) [S5]

Likewise, searches of major disorder-specific genomic studies showed **no TECR mention** in a large GWAS of **problematic alcohol use**, a large GWAS of **nicotine dependence**, a large GWAS/meta-analysis of **cannabis use disorder**, or a GWAS of **opioid use disorder**. ([discovery.ucl.ac.uk](https://discovery.ucl.ac.uk/id/eprint/10099547/1/PAU_meta_NN_merged%20%281%29.pdf)) [S6]

Recent scholarly reviews of substance-use-disorder genetics summarize the field around repeatedly implicated loci and pathways, but the accessible review material I found did not provide evidence elevating TECR as an addiction gene. Instead, the literature emphasizes established signals such as **alcohol-metabolizing genes**, **nicotinic receptor genes**, and **OPRM1** or cross-disorder polygenic risk. ([pubmed.ncbi.nlm.nih.gov](https://pubmed.ncbi.nlm.nih.gov/33879270/?utm_source=openai)) [S7]

A possible source of confusion is **PECR**, a different gene (**peroxisomal** trans-2-enoyl-CoA reductase). Some older alcohol-dependence GWAS/reviews discussed **PECR**, not TECR, and even there the evidence was described as weaker or less consistently important than top alcohol loci. That does **not** transfer to TECR. ([pmc.ncbi.nlm.nih.gov](https://pmc.ncbi.nlm.nih.gov/articles/PMC4187224/?utm_source=openai)) [S8]

## Mechanistic / biological rationale

Biologically, one can make only a **speculative** case that TECR could matter to addiction. Lipid composition, sphingolipid metabolism, ER calcium handling, synaptic plasticity, and BBB integrity are all broad processes that can influence brain function. TECR participates in several of those processes through **fatty-acid elongation**, **sphingolipid-related metabolism**, **SERCA2b/ER calcium regulation**, and **BBB maintenance**. ([ncbi.nlm.nih.gov](https://www.ncbi.nlm.nih.gov/gene/9524?utm_source=openai)) [S9]

However, addiction claims usually require more specific evidence: for example, replicated **GWAS associations**, differential expression in addiction-relevant brain regions from human cases or validated animal models, functional perturbation changing **drug reward/reinforcement/withdrawal**, or consistent inclusion in convergent multi-omic analyses of substance use disorders. I did not find that level of evidence for TECR. ([nature.com](https://www.nature.com/articles/s44220-023-00034-y)) [S10]

So the best current interpretation is: **TECR has plausible CNS-related biology, but plausibility is not implication in addiction**. Its known roles are too general and indirect to support the claim as stated. ([pubmed.ncbi.nlm.nih.gov](https://pubmed.ncbi.nlm.nih.gov/33482198/?utm_source=openai)) [S11]

## Bottom line

At present, **TECR is not an established addiction gene/protein**. The published evidence I found supports TECR’s roles in **lipid metabolism**, **ER calcium regulation**, **BBB homeostasis**, and **rare intellectual-disability phenotypes**, but I found **no direct, replicated human or animal evidence** linking TECR itself to addiction or substance use disorders. If the intended gene was **PECR** rather than TECR, there is some older, still limited alcohol-genetics literature on that separate gene. ([ncbi.nlm.nih.gov](https://www.ncbi.nlm.nih.gov/gene/9524?utm_source=openai)) [S12]

## Cited sources

- **[S1]** TECR trans-2,3-enoyl-CoA reductase [Homo sapiens (human)] - Gene - NCBI — https://www.ncbi.nlm.nih.gov/gene/9524?utm_source=openai
- **[S2]** Trans-2-enoyl-CoA reductase limits Ca2+ accumulation in the endoplasmic reticulum by inhibiting the Ca2+ pump SERCA2b - PubMed — https://pubmed.ncbi.nlm.nih.gov/33482198/?utm_source=openai
- **[S3]** trans-2-Enoyl-CoA Reductase Tecr-Driven Lipid Metabolism in Endothelial Cells Protects against Transcytosis to Maintain Blood-Brain Barrier Homeostasis - PubMed — https://pubmed.ncbi.nlm.nih.gov/35465346/
- **[S4]**   curation results for Gene-Disease Validity   — https://search.clinicalgenome.org/kb/gene-validity/CGGV%3Aassertion_8c7adee2-6a14-4348-a018-9097954bd39d-2022-03-10T070000.000Z
- **[S5]** Multivariate genome-wide association meta-analysis of over 1 million subjects identifies loci underlying multiple substance use disorders | Nature Mental Health — https://www.nature.com/articles/s44220-023-00034-y
- **[S6]** Untitled source — https://discovery.ucl.ac.uk/id/eprint/10099547/1/PAU_meta_NN_merged%20%281%29.pdf
- **[S7]** Genetics of substance use disorders: a review - PubMed — https://pubmed.ncbi.nlm.nih.gov/33879270/?utm_source=openai
- **[S8]** Genome-wide association discoveries of alcohol dependence - PMC — https://pmc.ncbi.nlm.nih.gov/articles/PMC4187224/?utm_source=openai
- **[S9]** TECR trans-2,3-enoyl-CoA reductase [Homo sapiens (human)] - Gene - NCBI — https://www.ncbi.nlm.nih.gov/gene/9524?utm_source=openai
- **[S10]** Multivariate genome-wide association meta-analysis of over 1 million subjects identifies loci underlying multiple substance use disorders | Nature Mental Health — https://www.nature.com/articles/s44220-023-00034-y
- **[S11]** Trans-2-enoyl-CoA reductase limits Ca2+ accumulation in the endoplasmic reticulum by inhibiting the Ca2+ pump SERCA2b - PubMed — https://pubmed.ncbi.nlm.nih.gov/33482198/?utm_source=openai
- **[S12]** TECR trans-2,3-enoyl-CoA reductase [Homo sapiens (human)] - Gene - NCBI — https://www.ncbi.nlm.nih.gov/gene/9524?utm_source=openai
