## Supplementary material for "Understanding Substance Dependence: What Differentiates Addictive from Non-Addictive Drugs?": Folder for supplementary files: addictive-genes-claims-first100-061-kalrn-gene-protein-is-implicated-in-addiction-00dcbe002f78.md

# addictive-genes-claims-first100-061-00dcbe002f78

**Original claim:** KALRN gene/protein is implicated in addiction.

## Claim

KALRN gene/protein is implicated in addiction.

## Verdict

Verdict: Likely True

KALRN (encoding kalirin, especially the Kalirin-7 isoform) is supported by **substantial preclinical evidence** as a regulator of addiction-relevant neuroplasticity and cocaine-related behaviors, particularly in the nucleus accumbens. However, the **human evidence is still limited and mostly indirect**, so the claim is better framed as “KALRN is implicated in addiction-related biology, especially cocaine-related plasticity,” rather than “KALRN is a definitively established human addiction-risk gene.” ([pubmed.ncbi.nlm.nih.gov](https://pubmed.ncbi.nlm.nih.gov/20452575/)) [S1]

## Confidence

Medium

Confidence is **medium** because multiple independent animal studies converge on KALRN/kalirin in cocaine-related phenotypes and synaptic adaptations, but human evidence is narrower, with less replication and more endophenotypic than diagnostic support. ([pubmed.ncbi.nlm.nih.gov](https://pubmed.ncbi.nlm.nih.gov/20452575/)) [S2]

## Supporting evidence

Several rodent studies directly connect kalirin to **cocaine-induced behavioral and structural plasticity**. In mice, chronic cocaine increased Kalirin-7 mRNA and protein in the nucleus accumbens, while Kal7 knockout prevented the usual cocaine-induced rise in dendritic spine density. The same knockout also altered cocaine behaviors: locomotor sensitization was enhanced, while cocaine conditioned place preference was reduced, despite normal food place preference. This is direct evidence that Kal7 participates in drug-specific reward/plasticity mechanisms rather than general learning or reward. ([pubmed.ncbi.nlm.nih.gov](https://pubmed.ncbi.nlm.nih.gov/20452575/)) [S3]

The evidence extends beyond passive drug exposure to more addiction-relevant reinforcement paradigms. In a mouse intravenous self-administration study, Kal7 knockout animals took **more cocaine at lower doses** than wild-type mice. The authors also found altered nucleus accumbens transcriptional responses to self-administered cocaine, including failure to upregulate the NMDA receptor subunit NR2B transcript in the knockout animals. That supports involvement in reinforcing effects of cocaine, not just locomotor activation. ([pubmed.ncbi.nlm.nih.gov](https://pubmed.ncbi.nlm.nih.gov/23894151/)) [S4]

A rat study provided stronger mechanistic resolution. After repeated cocaine and withdrawal, Kal-7 protein, Rac1/PAK signaling, AMPA receptor surface expression, and dendritic spine density all increased in nucleus accumbens. Knocking down Kal-7 in the accumbens core blocked the cocaine-induced AMPAR and spine changes and **severely impaired incentive sensitization**, measured as faster learning to self-administer a threshold cocaine dose, even though locomotor sensitization still developed. This is important because it places kalirin in a pathway that affects the “wanting” or incentive-motivational properties of cocaine rather than nonspecific activation alone. ([pubmed.ncbi.nlm.nih.gov](https://pubmed.ncbi.nlm.nih.gov/23825406/)) [S5]

Additional work showed that repeated cocaine changes **Kalrn promoter usage and isoform expression** in the nucleus accumbens, favoring Kalirin-7/ΔKalirin-7-related transcripts. The authors concluded these cocaine-driven splicing/promoter shifts may contribute to the behavioral, morphological, and biochemical responses to cocaine. Another study in rat medium spiny neurons found that repeated cocaine increased Kal7 and Kal12 expression, that Kal7 overexpression increased spine density, and that reducing endogenous kalirin decreased dendritic length and spine density, supporting kalirin as an obligatory intermediate in MSN responses to repeated cocaine. ([pubmed.ncbi.nlm.nih.gov](https://pubmed.ncbi.nlm.nih.gov/21329509/)) [S6]

There is also **human translational evidence**, though less definitive. In the IMAGEN adolescent cohort (n=1,423), intronic KALRN variation was associated with greater ventral striatal activation during reward anticipation, and another KALRN intronic variant in the same haplotype block was associated with increased binge-drinking frequency. That does not prove KALRN causes addiction, but it does connect the gene to reward-circuit responsivity and alcohol misuse in humans—both relevant addiction phenotypes. ([pubmed.ncbi.nlm.nih.gov](https://pubmed.ncbi.nlm.nih.gov/27092175/)) [S7]

## Contradicting or limiting evidence

The main limitation is that the **strongest evidence is preclinical and cocaine-focused**. Most direct causal data come from mice and rats, particularly from nucleus accumbens studies of cocaine exposure, cocaine sensitization, place preference, and cocaine self-administration. That is enough to implicate KALRN in addiction-related neurobiology, but it does **not** establish a broad role across all addictive substances or all stages of addiction in humans. ([pubmed.ncbi.nlm.nih.gov](https://pubmed.ncbi.nlm.nih.gov/20452575/)) [S8]

The human data are also **limited and indirect**. The best positive human paper linked KALRN variants to ventral striatal activation and adolescent binge drinking, but those are risk-related or intermediate phenotypes, not replicated genome-wide-significant associations with diagnosed substance use disorders. ([pubmed.ncbi.nlm.nih.gov](https://pubmed.ncbi.nlm.nih.gov/27092175/)) [S9]

Phenotypically, KALRN manipulation does **not** produce a simple one-directional addiction effect. Kal7 loss can increase cocaine locomotor sensitization and increase low-dose cocaine self-administration, yet reduce cocaine conditioned place preference. That pattern argues for a **complex modulatory role** in reward learning, motivation, and glutamatergic plasticity, rather than a straightforward “more KALRN = more addiction” model. ([pubmed.ncbi.nlm.nih.gov](https://pubmed.ncbi.nlm.nih.gov/20452575/)) [S10]

Finally, larger human GWAS of alcohol and multi-substance addiction have highlighted other loci—such as **ADH1B, CRHR1, FGF14, and PDE4B**—as more robust genome-wide signals. That does not refute KALRN involvement, but it shows KALRN is **not currently a top-tier, consistently replicated human addiction locus** on the basis of available large-scale genetics. ([pubmed.ncbi.nlm.nih.gov](https://pubmed.ncbi.nlm.nih.gov/22004471/?utm_source=openai)) [S11]

## Mechanistic / biological rationale

The biological rationale is strong. Kalirin is a **brain-enriched Rho guanine nucleotide exchange factor** that regulates Rac-family signaling, actin remodeling, dendritic spine morphogenesis, and glutamatergic synapse structure/function. Independent work showed KALRN knockout reduces cortical Rac1 signaling, spine density, and glutamatergic transmission, consistent with kalirin being a major organizer of excitatory synaptic architecture. Because addiction depends heavily on long-lasting plasticity in mesocorticolimbic circuits, this makes KALRN mechanistically plausible from the outset. ([pubmed.ncbi.nlm.nih.gov](https://pubmed.ncbi.nlm.nih.gov/19625617/)) [S12]

More specifically, Kalirin-7 interacts with the **GluN2B/NR2B subunit of NMDA receptors** and regulates its synaptic localization and function. Kal7 knockout mice show reduced NR2B-dependent currents and reduced surface expression of NR2B. In the cocaine literature, pretreatment with the GluN2B antagonist ifenprodil abolished the exaggerated cocaine locomotor response in Kal7-deficient mice, linking the behavioral phenotype to glutamatergic/NMDA signaling rather than to a vague developmental abnormality. ([pubmed.ncbi.nlm.nih.gov](https://pubmed.ncbi.nlm.nih.gov/21880917/)) [S13]

Kalirin also appears to couple drug exposure to **structural and AMPAR-mediated plasticity** in the nucleus accumbens. Repeated cocaine increases Kal-7 and activates downstream Rac1/PAK signaling; Kal-7 knockdown prevents cocaine-induced increases in AMPAR surface expression and dendritic spine density. This offers a biologically coherent mechanism by which KALRN could influence incentive sensitization, cue responsiveness, and persistent drug-related learning. ([pubmed.ncbi.nlm.nih.gov](https://pubmed.ncbi.nlm.nih.gov/23825406/)) [S14]

So mechanistically, KALRN is best viewed as a **plasticity-regulating node in reward circuitry**, especially at glutamatergic synapses of the nucleus accumbens and related circuits, rather than as a classic drug-metabolism gene or a universally proven addiction susceptibility gene. ([pubmed.ncbi.nlm.nih.gov](https://pubmed.ncbi.nlm.nih.gov/21329509/)) [S15]

## Bottom line

Yes—**KALRN/kalirin is reasonably described as implicated in addiction**, especially in **cocaine-related addiction neurobiology**. The support is strongest from rodent studies showing that cocaine changes kalirin expression/isoforms and that disrupting Kalirin-7 alters cocaine-induced spine plasticity, incentive sensitization, and self-administration. But the human evidence is still limited, so the most accurate formulation is:

**KALRN is implicated in addiction-related neural plasticity and addiction-relevant behaviors, with strongest evidence in preclinical cocaine models and more limited, indirect evidence in humans.** ([pubmed.ncbi.nlm.nih.gov](https://pubmed.ncbi.nlm.nih.gov/20452575/)) [S16]

## Cited sources

- **[S1]** Behavioral and morphological responses to cocaine require kalirin7 - PubMed — https://pubmed.ncbi.nlm.nih.gov/20452575/
- **[S2]** Behavioral and morphological responses to cocaine require kalirin7 - PubMed — https://pubmed.ncbi.nlm.nih.gov/20452575/
- **[S3]** Behavioral and morphological responses to cocaine require kalirin7 - PubMed — https://pubmed.ncbi.nlm.nih.gov/20452575/
- **[S4]** Constitutive knockout of kalirin-7 leads to increased rates of cocaine self-administration - PubMed — https://pubmed.ncbi.nlm.nih.gov/23894151/
- **[S5]** Kalirin-7 mediates cocaine-induced AMPA receptor and spine plasticity, enabling incentive sensitization - PubMed — https://pubmed.ncbi.nlm.nih.gov/23825406/
- **[S6]** Kalrn promoter usage and isoform expression respond to chronic cocaine exposure - PubMed — https://pubmed.ncbi.nlm.nih.gov/21329509/
- **[S7]** Mouse and Human Genetic Analyses Associate Kalirin with Ventral Striatal Activation during Impulsivity and with Alcohol Misuse - PubMed — https://pubmed.ncbi.nlm.nih.gov/27092175/
- **[S8]** Behavioral and morphological responses to cocaine require kalirin7 - PubMed — https://pubmed.ncbi.nlm.nih.gov/20452575/
- **[S9]** Mouse and Human Genetic Analyses Associate Kalirin with Ventral Striatal Activation during Impulsivity and with Alcohol Misuse - PubMed — https://pubmed.ncbi.nlm.nih.gov/27092175/
- **[S10]** Behavioral and morphological responses to cocaine require kalirin7 - PubMed — https://pubmed.ncbi.nlm.nih.gov/20452575/
- **[S11]** Genome-wide significant association between alcohol dependence and a variant in the ADH gene cluster - PubMed — https://pubmed.ncbi.nlm.nih.gov/22004471/?utm_source=openai
- **[S12]** Kalirin regulates cortical spine morphogenesis and disease-related behavioral phenotypes - PubMed — https://pubmed.ncbi.nlm.nih.gov/19625617/
- **[S13]** Kalirin binds the NR2B subunit of the NMDA receptor, altering its synaptic localization and function - PubMed — https://pubmed.ncbi.nlm.nih.gov/21880917/
- **[S14]** Kalirin-7 mediates cocaine-induced AMPA receptor and spine plasticity, enabling incentive sensitization - PubMed — https://pubmed.ncbi.nlm.nih.gov/23825406/
- **[S15]** Kalrn promoter usage and isoform expression respond to chronic cocaine exposure - PubMed — https://pubmed.ncbi.nlm.nih.gov/21329509/
- **[S16]** Behavioral and morphological responses to cocaine require kalirin7 - PubMed — https://pubmed.ncbi.nlm.nih.gov/20452575/
