## Supplementary material for "Understanding Substance Dependence: What Differentiates Addictive from Non-Addictive Drugs?": Folder for supplementary files: addictive-genes-claims-first100-062-ednrb-gene-protein-is-implicated-in-addiction-c483ed422700.md

# addictive-genes-claims-first100-062-c483ed422700

**Original claim:** EDNRB gene/protein is implicated in addiction.

## Claim

EDNRB gene/protein is implicated in addiction.

## Verdict

Verdict: Likely False

## Confidence

Medium — large human genetics resources and major addiction-genetics reviews do not identify **EDNRB** as a replicated addiction locus, and the closest addiction-related endothelin studies point more toward **ETA/EDNRA-system** involvement than a clear **EDNRB-specific** signal. However, this is still mostly a conclusion from the pattern of available evidence rather than a definitive falsification experiment. ([ebi.ac.uk](https://www.ebi.ac.uk/gwas/genes/EDNRB)) [S1]

## Supporting evidence

The best case *for* the claim is indirect, not direct. **EDNRB** encodes endothelin receptor type B, a GPCR with recognized roles in cell signaling, vascular biology, and development; it is also expressed in the nervous system and can affect CNS biology. MedlinePlus and NCBI describe EDNRB as a receptor involved in signaling, blood-vessel function, hormone-related processes, and neural crest/enteric nervous system development. ([medlineplus.gov](https://medlineplus.gov/genetics/gene/ednrb/?utm_source=openai)) [S2]

There is also evidence that **EDNRB has bona fide CNS plasticity/homeostasis functions**. In a mouse/zebrafish study, reduced endothelin signaling decreased oligodendrocyte myelin sheath number, oligodendroglial **EDNRB** loss phenocopied part of the effect of social isolation in medial prefrontal cortex, and intranasal **EDNRB agonist** treatment rescued isolation-associated myelination deficits. That does not demonstrate addiction, but it shows EDNRB can influence adaptive brain responses in circuits relevant to behavior. ([pmc.ncbi.nlm.nih.gov](https://pmc.ncbi.nlm.nih.gov/articles/PMC6831104/)) [S3]

A second indirect point is that the broader **endothelin system** has been investigated in opioid withdrawal biology. In neonatal morphine-withdrawal experiments, investigators tested endothelin receptor pharmacology including an **ET(B) agonist** and concluded that central endothelin receptors are relevant enough to examine in withdrawal paradigms. A later review likewise discussed the endothelin system as a potential target in opioid withdrawal. ([pubmed.ncbi.nlm.nih.gov](https://pubmed.ncbi.nlm.nih.gov/16741068/)) [S4]

Taken together, these findings give **biological plausibility** that EDNRB could, in principle, affect neural adaptation. But plausibility is not the same as evidence that it is actually implicated in addiction.

## Contradicting or limiting evidence

The strongest limitation is the **lack of direct human genetic evidence**. When accessed, the GWAS Catalog entry for **EDNRB** showed the framework for “Associations,” “Studies,” and “Traits,” but no listed entries, which argues against any currently curated genome-wide significant trait associations for this gene at that source. ([ebi.ac.uk](https://www.ebi.ac.uk/gwas/genes/EDNRB)) [S5]

Large alcohol-dependence genetics studies also do **not** point to EDNRB. A review of 45 alcohol-dependence GWASs identified robust or replicated signals at loci such as the **ADH cluster**, **SERINC2**, **KIAA0040**, **PKNOX2**, **HTR7**, and others, while discussing broader candidate systems such as dopaminergic, GABAergic, cholinergic, and opioidergic pathways; **EDNRB was not among the highlighted replicated loci**. ([pmc.ncbi.nlm.nih.gov](https://pmc.ncbi.nlm.nih.gov/articles/PMC4187224/)) [S6]

Similarly, the largest trans-ancestral GWAS of DSM-IV alcohol dependence reported genome-wide significant effects in **ADH1B** and emphasized shared genetics with other psychiatric and substance-use traits, but not EDNRB. ([pubmed.ncbi.nlm.nih.gov](https://pubmed.ncbi.nlm.nih.gov/30482948/)) [S7]

For opioid use disorder, a 2025 review concluded that the most repeatable genome-wide findings concentrate at **OPRM1** and **FURIN**, and it cautioned that many older candidate-gene findings have not held up in large GWAS. EDNRB was not presented as a robust OUD locus. ([cambridge.org](https://www.cambridge.org/core/product/8605BFBA52432F57F98A918E6CD64E00)) [S8]

The closest **directly addiction-adjacent** endothelin experiment is also not supportive of an EDNRB-specific role. In neonatal morphine withdrawal, the **ET(B) agonist IRL1620** produced similar G-protein stimulation in placebo and withdrawal groups, whereas the paper’s conclusion emphasized **ET(A)** involvement. The later review on opioid withdrawal likewise centered on **ETA receptor antagonism** as the relevant therapeutic signal. ([pubmed.ncbi.nlm.nih.gov](https://pubmed.ncbi.nlm.nih.gov/16741068/)) [S9]

Finally, authoritative reference descriptions of **EDNRB** focus on Hirschsprung disease, Waardenburg syndrome type IV, melanocyte biology, and vascular/developmental functions, not addiction phenotypes. That does not prove absence of a role, but it is consistent with EDNRB **not** being an established addiction gene/protein. ([medlineplus.gov](https://medlineplus.gov/genetics/gene/ednrb/?utm_source=openai)) [S10]

## Mechanistic / biological rationale

Biologically, one can make a **speculative** case for EDNRB having some relevance to addiction-related neuroadaptation. EDNRB is a GPCR, and GPCR signaling often intersects with pathways involved in synaptic plasticity, glial biology, stress responses, and circuit remodeling. The myelination study suggests EDNRB participates in **experience-dependent oligodendrocyte responses** in medial prefrontal cortex, a region deeply involved in executive control and relapse vulnerability. ([pmc.ncbi.nlm.nih.gov](https://pmc.ncbi.nlm.nih.gov/articles/PMC6831104/)) [S11]

But addiction claims require more than generic CNS relevance. For a gene/protein to be considered “implicated in addiction,” one would usually want at least one of the following: replicated human genetic association, reproducible differential expression in addicted brains, or animal experiments showing altered drug reward, self-administration, withdrawal, craving, or relapse when the gene/protein is manipulated. Across the major sources reviewed here, **that level of evidence is missing for EDNRB**. ([cambridge.org](https://www.cambridge.org/core/product/8605BFBA52432F57F98A918E6CD64E00)) [S12]

The broader endothelin literature actually suggests a more specific interpretation: if endothelin signaling matters in addiction-related states, the evidence so far leans more toward **ETA-mediated opioid withdrawal mechanisms** than toward **EDNRB/ETB** as a recurring addiction target. ([pubmed.ncbi.nlm.nih.gov](https://pubmed.ncbi.nlm.nih.gov/16741068/)) [S13]

## Bottom line

Current evidence does **not** support **EDNRB** as an established or recurring addiction gene/protein. There is **indirect plausibility** because EDNRB participates in CNS signaling and adaptive myelination, and the broader endothelin system has been examined in opioid withdrawal. But direct addiction evidence for **EDNRB specifically**—human genetics, replicated omics, or addiction-behavior experiments—is lacking, and the limited opioid-withdrawal literature points more to **ETA** than **ETB/EDNRB**. On present evidence, the claim is **overstated**. ([pmc.ncbi.nlm.nih.gov](https://pmc.ncbi.nlm.nih.gov/articles/PMC6831104/)) [S14]

## Cited sources

- **[S1]** GWAS Catalog — https://www.ebi.ac.uk/gwas/genes/EDNRB
- **[S2]** EDNRB gene: MedlinePlus Genetics — https://medlineplus.gov/genetics/gene/ednrb/?utm_source=openai
- **[S3]** 
            Endothelin signalling mediates experience-dependent myelination in the CNS - PMC
         — https://pmc.ncbi.nlm.nih.gov/articles/PMC6831104/
- **[S4]** Involvement of central endothelin receptors in neonatal morphine withdrawal - PubMed — https://pubmed.ncbi.nlm.nih.gov/16741068/
- **[S5]** GWAS Catalog — https://www.ebi.ac.uk/gwas/genes/EDNRB
- **[S6]** 
            Genome-wide association discoveries of alcohol dependence - PMC
         — https://pmc.ncbi.nlm.nih.gov/articles/PMC4187224/
- **[S7]** Transancestral GWAS of alcohol dependence reveals common genetic underpinnings with psychiatric disorders - PubMed — https://pubmed.ncbi.nlm.nih.gov/30482948/
- **[S8]** Genetic and Epigenetic Approaches to Opioid Use Disorder | Expert Reviews in Molecular Medicine | Cambridge Core — https://www.cambridge.org/core/product/8605BFBA52432F57F98A918E6CD64E00
- **[S9]** Involvement of central endothelin receptors in neonatal morphine withdrawal - PubMed — https://pubmed.ncbi.nlm.nih.gov/16741068/
- **[S10]** EDNRB gene: MedlinePlus Genetics — https://medlineplus.gov/genetics/gene/ednrb/?utm_source=openai
- **[S11]** 
            Endothelin signalling mediates experience-dependent myelination in the CNS - PMC
         — https://pmc.ncbi.nlm.nih.gov/articles/PMC6831104/
- **[S12]** Genetic and Epigenetic Approaches to Opioid Use Disorder | Expert Reviews in Molecular Medicine | Cambridge Core — https://www.cambridge.org/core/product/8605BFBA52432F57F98A918E6CD64E00
- **[S13]** Involvement of central endothelin receptors in neonatal morphine withdrawal - PubMed — https://pubmed.ncbi.nlm.nih.gov/16741068/
- **[S14]** 
            Endothelin signalling mediates experience-dependent myelination in the CNS - PMC
         — https://pmc.ncbi.nlm.nih.gov/articles/PMC6831104/
