## Supplementary material for "Understanding Substance Dependence: What Differentiates Addictive from Non-Addictive Drugs?": Folder for supplementary files: addictive-genes-claims-first100-063-slc6a1-gene-protein-is-implicated-in-addiction-b326d43a1e7a.md

# addictive-genes-claims-first100-063-b326d43a1e7a

**Original claim:** SLC6A1 gene/protein is implicated in addiction.

## Claim

SLC6A1 gene/protein is implicated in addiction. ([ncbi.nlm.nih.gov](https://www.ncbi.nlm.nih.gov/sites/books/NBK589173/)) [S1]

## Verdict

Verdict: Mixed Evidence

SLC6A1 encodes GAT-1, the major neuronal GABA transporter, so it is biologically plausible that it could influence addiction-relevant processes. There is some supporting evidence from human postmortem brain studies, a candidate-gene alcohol study, human pharmacology with the GAT-1 inhibitor tiagabine, and several animal studies linking GAT-1 to alcohol preference or opioid withdrawal. However, the evidence is not robust enough to say SLC6A1 is an established addiction gene in the same sense as the strongest replicated substance-use loci. In particular, the human genetic evidence is limited and does not appear to be supported by large modern AUD GWAS, and clinical trials of tiagabine for cocaine dependence have been mixed to negative. ([pmc.ncbi.nlm.nih.gov](https://pmc.ncbi.nlm.nih.gov/articles/PMC3258238/)) [S2]

## Confidence

Medium. Confidence is limited because the signal comes from heterogeneous evidence types—transcriptomic, candidate-gene, pharmacologic, and animal-model data—rather than from strong replicated human genetic associations or clearly effective human intervention studies. ([pmc.ncbi.nlm.nih.gov](https://pmc.ncbi.nlm.nih.gov/articles/PMC3258238/)) [S3]

## Supporting evidence

**1) Human postmortem brain data support altered SLC6A1 expression in addiction.**  
In postmortem hippocampus, SLC6A1/GAT1 expression was reported as downregulated in both alcoholics and cocaine addicts versus controls. A later review of human alcoholic brain transcriptomics summarized SLC6A1 among GABA-related genes repeatedly found to be downregulated in alcoholics, and noted overlap with cocaine-addicted brains. That is not proof of causation, but it does place SLC6A1 among molecular changes observed in human addicted brain tissue. ([pmc.ncbi.nlm.nih.gov](https://pmc.ncbi.nlm.nih.gov/articles/PMC3258238/)) [S4]

**2) There is at least some human genetic association evidence for alcohol dependence.**  
A 2015 study in three ethnically distinct populations found a protective signal for a functional SLC6A1 promoter insertion polymorphism in the African American sample, with the insertion allele more common in controls than in alcohol-dependent cases; combined analysis with GABBR1 suggested genotype combinations associated with resilience versus risk. Importantly, though, the SLC6A1 finding was ancestry-specific and partly exploratory rather than a broad, replicated cross-population hit. ([pmc.ncbi.nlm.nih.gov](https://pmc.ncbi.nlm.nih.gov/articles/PMC4700558/)) [S5]

**3) Human pharmacology suggests GAT-1 modulation can affect cocaine-related subjective responses.**  
In a small human laboratory study, tiagabine pretreatment attenuated some cocaine-induced subjective ratings, including feeling “stimulated” and craving cocaine, without changing cardiovascular effects. This does not show SLC6A1 causes addiction, but it suggests the protein can modulate at least some acute cocaine-related effects in humans. ([pubmed.ncbi.nlm.nih.gov](https://pubmed.ncbi.nlm.nih.gov/16332385/)) [S6]

**4) Preclinical work supports a role in addiction-related behaviors and withdrawal.**  
A Nature Neuroscience study found that GAT-1 activity in periaqueductal gray contributed to opioid withdrawal behaviors, and that GAT-1 inhibition reduced withdrawal-associated GABAergic changes. This is strong mechanistic evidence that the SLC6A1 protein can participate in a core addiction-related process: dependence/withdrawal. ([nature.com](https://www.nature.com/articles/nn.2940)) [S7]

**5) Newer mouse work links Slc6a1/GAT1 to alcohol preference.**  
A 2025 mouse study reported reduced Slc6a1/GAT1 expression in amygdala of high-alcohol-drinking mice, found that a GAT1 inhibitor blocked a protective menin overexpression effect, and showed that restoring GAT1 in specific interneurons reduced alcohol preference in genetically manipulated mice. This is still animal evidence, but it directly connects Slc6a1 manipulation to alcohol-drinking behavior. ([pubmed.ncbi.nlm.nih.gov](https://pubmed.ncbi.nlm.nih.gov/40425852/)) [S8]

## Contradicting or limiting evidence

**1) Large human GWAS do not currently make SLC6A1 a robustly established AUD locus.**  
The largest recent multi-ancestry AUD GWAS identified 105 genome-wide significant variants across 37 loci, with mapped signals in multiple brain-relevant pathways, but SLC6A1 is not identified in that report. Likewise, an earlier large trans-ancestral alcohol dependence GWAS highlighted ADH1B, not SLC6A1. I am inferring from the lack of reported signal that SLC6A1 is not presently a top-tier replicated common-variant AUD locus. ([pmc.ncbi.nlm.nih.gov](https://pmc.ncbi.nlm.nih.gov/articles/PMC10775504/)) [S9]

**2) The candidate-gene evidence is limited.**  
The main positive SLC6A1 association study relied on a promoter insertion polymorphism unique to African ancestry and exploratory genotype-combination analyses; the paper itself notes some results were nominal or trend-level. That is much weaker than consistent replication across large independent cohorts or genome-wide significance. ([pmc.ncbi.nlm.nih.gov](https://pmc.ncbi.nlm.nih.gov/articles/PMC4700558/)) [S10]

**3) Clinical evidence for GAT-1–targeting treatment in addiction is mixed to negative.**  
Although tiagabine reduced some acute subjective cocaine effects in a small lab study, a larger randomized controlled trial in cocaine dependence found no significant differences in craving/global functioning between groups and no robust reduction in cocaine use. A review of psychostimulant medication development concluded that tiagabine’s findings in both animals and humans were conflicting and suggested limited therapeutic potential. ([pubmed.ncbi.nlm.nih.gov](https://pubmed.ncbi.nlm.nih.gov/16332385/)) [S11]

**4) SLC6A1’s best-established human disease role is not addiction.**  
GeneReviews states that SLC6A1 encodes GAT-1 and that known germline pathogenic-variant phenotypes are SLC6A1-related neurodevelopmental disorder/epilepsy; it specifically notes no other germline phenotypes are currently known. So, unlike some genes with strong syndromic or highly replicated addiction associations, SLC6A1 is primarily established in neurodevelopmental disease, not substance-use disorder. ([ncbi.nlm.nih.gov](https://www.ncbi.nlm.nih.gov/sites/books/NBK589173/)) [S12]

## Mechanistic / biological rationale

SLC6A1 encodes the sodium- and chloride-dependent GABA transporter GAT-1, which clears GABA from the synaptic cleft into neurons and glia. Because GABA is the principal inhibitory neurotransmitter in the brain, SLC6A1 helps regulate excitation/inhibition balance and therefore can influence circuits relevant to reward, stress, craving, and withdrawal. ([ncbi.nlm.nih.gov](https://www.ncbi.nlm.nih.gov/sites/books/NBK589173/)) [S13]

Experimental evidence supports this mechanism. GAT1-deficient mice have elevated extracellular GABA and increased tonic GABAergic signaling, showing that changing SLC6A1 function materially alters inhibitory tone. In addiction-relevant terms, altered inhibitory tone can affect drug reward, tolerance, and withdrawal circuitry. The opioid-withdrawal study is especially informative because it ties GAT-1 activity directly to expression of withdrawal signs in a defined brain region. ([pubmed.ncbi.nlm.nih.gov](https://pubmed.ncbi.nlm.nih.gov/12815026/?utm_source=openai)) [S14]

That said, mechanistic plausibility is not the same as disease-level proof. Many genes in GABA signaling are biologically relevant to addiction circuitry, but only some show strong, replicated human association. For SLC6A1, the mechanistic case is stronger than the human causal-genetic case. ([pubmed.ncbi.nlm.nih.gov](https://pubmed.ncbi.nlm.nih.gov/27475769/?utm_source=openai)) [S15]

## Bottom line

SLC6A1/GAT-1 is **plausibly and partially implicated** in addiction biology, especially through GABAergic regulation of withdrawal, alcohol-related behaviors, and drug-response phenotypes. But the evidence is **not yet strong enough to call it a firmly established human addiction gene**. The best summary is: **mechanistically relevant, supported by some human and animal evidence, but not robustly confirmed by large-scale human genetics or clinical treatment data**. ([pmc.ncbi.nlm.nih.gov](https://pmc.ncbi.nlm.nih.gov/articles/PMC3258238/)) [S16]

## Cited sources

- **[S1]** SLC6A1-Related Neurodevelopmental Disorder - GeneReviews® - NCBI Bookshelf — https://www.ncbi.nlm.nih.gov/sites/books/NBK589173/
- **[S2]** 
            GABAergic Gene Expression in Postmortem Hippocampus from Alcoholics and Cocaine Addicts; Corresponding Findings in Alcohol-Naïve P and NP Rats - PMC
         — https://pmc.ncbi.nlm.nih.gov/articles/PMC3258238/
- **[S3]** 
            GABAergic Gene Expression in Postmortem Hippocampus from Alcoholics and Cocaine Addicts; Corresponding Findings in Alcohol-Naïve P and NP Rats - PMC
         — https://pmc.ncbi.nlm.nih.gov/articles/PMC3258238/
- **[S4]** 
            GABAergic Gene Expression in Postmortem Hippocampus from Alcoholics and Cocaine Addicts; Corresponding Findings in Alcohol-Naïve P and NP Rats - PMC
         — https://pmc.ncbi.nlm.nih.gov/articles/PMC3258238/
- **[S5]** 
            GABBR1 and SLC6A1, two genes involved in modulation of GABA synaptic transmission influence risk for alcoholism; results from three ethnically diverse populations - PMC
         — https://pmc.ncbi.nlm.nih.gov/articles/PMC4700558/
- **[S6]** Tiagabine affects the subjective responses to cocaine in humans - PubMed — https://pubmed.ncbi.nlm.nih.gov/16332385/
- **[S7]** Drug-induced GABA transporter currents enhance GABA release to induce opioid withdrawal behaviors | Nature Neuroscience — https://www.nature.com/articles/nn.2940
- **[S8]** A neural substrate of comorbid depressive symptoms in alcohol use - PubMed — https://pubmed.ncbi.nlm.nih.gov/40425852/
- **[S9]** 
            Identification of Novel Loci and Cross-Disorder Pleiotropy Through Multi-Ancestry Genome-Wide Analysis of Alcohol Use Disorder in Over One Million Individuals - PMC
         — https://pmc.ncbi.nlm.nih.gov/articles/PMC10775504/
- **[S10]** 
            GABBR1 and SLC6A1, two genes involved in modulation of GABA synaptic transmission influence risk for alcoholism; results from three ethnically diverse populations - PMC
         — https://pmc.ncbi.nlm.nih.gov/articles/PMC4700558/
- **[S11]** Tiagabine affects the subjective responses to cocaine in humans - PubMed — https://pubmed.ncbi.nlm.nih.gov/16332385/
- **[S12]** SLC6A1-Related Neurodevelopmental Disorder - GeneReviews® - NCBI Bookshelf — https://www.ncbi.nlm.nih.gov/sites/books/NBK589173/
- **[S13]** SLC6A1-Related Neurodevelopmental Disorder - GeneReviews® - NCBI Bookshelf — https://www.ncbi.nlm.nih.gov/sites/books/NBK589173/
- **[S14]** GABA transporter-1 (GAT1)-deficient mice: differential tonic activation of GABAA versus GABAB receptors in the hippocampus - PubMed — https://pubmed.ncbi.nlm.nih.gov/12815026/?utm_source=openai
- **[S15]** Neurobiology of addiction: a neurocircuitry analysis - PubMed — https://pubmed.ncbi.nlm.nih.gov/27475769/?utm_source=openai
- **[S16]** 
            GABAergic Gene Expression in Postmortem Hippocampus from Alcoholics and Cocaine Addicts; Corresponding Findings in Alcohol-Naïve P and NP Rats - PMC
         — https://pmc.ncbi.nlm.nih.gov/articles/PMC3258238/
