## Supplementary material for "Understanding Substance Dependence: What Differentiates Addictive from Non-Addictive Drugs?": Folder for supplementary files: addictive-genes-claims-first100-064-gabra1-gene-protein-is-implicated-in-addiction-9bf753461568.md

# addictive-genes-claims-first100-064-9bf753461568

**Original claim:** GABRA1 gene/protein is implicated in addiction.

## Claim

**“GABRA1 gene/protein is implicated in addiction.”**

## Verdict

Verdict: Mixed Evidence

## Confidence

**Medium.**

## Supporting evidence

GABRA1 encodes the **α1 subunit of the GABA\_A receptor**, a major component of fast inhibitory neurotransmission in the brain; GABA\_A receptors are chloride channels modulated by benzodiazepines and other sedative-hypnotic drugs, making GABRA1 biologically plausible as a candidate in substance-related phenotypes. ([ncbi.nlm.nih.gov](https://www.ncbi.nlm.nih.gov/gene/2554)) [S1]

**Human genetic evidence exists, but is limited and substance-specific.** In a family-based study from the Collaborative Study on the Genetics of Alcoholism (COGA), investigators found **association between GABRA1 variants and several alcohol-related phenotypes**, including a broader alcohol dependence definition, blackouts, age at first drunkenness, and level of response to alcohol. Importantly, the same paper notes that earlier family-based analyses had **not** found association with DSM-IV alcohol dependence, so even this positive human signal is nuanced rather than definitive. ([pubmed.ncbi.nlm.nih.gov](https://pubmed.ncbi.nlm.nih.gov/16792556/)) [S2]

**Rodent alcohol-dependence models provide stronger mechanistic support.** Multiple studies report that chronic ethanol exposure reduces α1/Gabra1 expression or surface availability. In rat cortex, chronic ethanol increased internalization of α1-containing GABA\_A receptors and decreased α1 peptide in the synaptic fraction, consistent with receptor trafficking changes during dependence-related neuroadaptation. ([pubmed.ncbi.nlm.nih.gov](https://pubmed.ncbi.nlm.nih.gov/12859683/)) [S3] In cultured cortical neurons, ethanol decreased Gabra1 transcription via histone deacetylation at the Gabra1 promoter, and class I HDAC inhibition prevented that decrease. ([pubmed.ncbi.nlm.nih.gov](https://pubmed.ncbi.nlm.nih.gov/28798030/)) [S4] In rat dependence models, chronic ethanol exposure plus withdrawal decreased Gabra1 expression in cortex and medial prefrontal cortex, and HDAC inhibition prevented both the molecular change and several dependence-like behaviors, while also reducing voluntary ethanol consumption. ([pubmed.ncbi.nlm.nih.gov](https://pubmed.ncbi.nlm.nih.gov/29520058/)) [S5]

A review of the chronic intermittent ethanol model concluded that ethanol-induced GABA\_A receptor plasticity contributes critically to alcohol dependence and increased drinking. That review specifically notes **downregulation of α1-containing receptors** during adaptation to ethanol, though it argues the most important receptors for maintained positive reinforcement may be α2- and α4-containing subtypes rather than α1 itself. ([pubmed.ncbi.nlm.nih.gov](https://pubmed.ncbi.nlm.nih.gov/28931433/)) [S6]

There is also **some support in sedative-hypnotic/benzodiazepine abuse-liability literature**. A review focused on zolpidem concluded that α1-containing GABA\_A receptors play a prominent role in several zolpidem and benzodiazepine effects, and in primates appear to contribute more strongly to **discriminative stimulus, reinforcing, and withdrawal effects** than in rodents. This does not prove that GABRA1 is a major “addiction gene,” but it does support involvement in some abuse-related phenotypes for α1-preferring GABAergic drugs. ([pubmed.ncbi.nlm.nih.gov](https://pubmed.ncbi.nlm.nih.gov/24563183/)) [S7]

## Contradicting or limiting evidence

The biggest limitation is that **direct human evidence is sparse and inconsistent**, and most positive findings are concentrated in **alcohol-related phenotypes**, not addiction broadly across substances. The COGA study itself reported that prior family-based analyses showed **no association with DSM-IV alcohol dependence**; the positive findings emerged when using broader or intermediate phenotypes. ([pubmed.ncbi.nlm.nih.gov](https://pubmed.ncbi.nlm.nih.gov/16792556/)) [S8]

Replication is also weak. A case-control study in Italian men found **no significant association** between GABRA1 polymorphisms and alcohol dependence after correction for multiple testing. ([pubmed.ncbi.nlm.nih.gov](https://pubmed.ncbi.nlm.nih.gov/25191505/)) [S9] Human postmortem data are also mixed: in frontal cortex from alcoholics, one study found **no significant difference** in α1 mRNA or peptide expression versus controls. ([pubmed.ncbi.nlm.nih.gov](https://pubmed.ncbi.nlm.nih.gov/9660306/)) [S10]

For **cross-substance addiction genetics**, GABRA1 does not currently stand out as a leading signal. In a 2023 multivariate GWAS/meta-analysis of over 1 million individuals examining general and substance-specific liability across alcohol, tobacco, cannabis, and opioid use disorders, the abstract highlights other significant loci such as **PDE4B**, not GABRA1. This is an inference from what the paper reports prominently—not proof of no effect—but it argues against GABRA1 being a major common-variant driver of broad addiction risk. ([pubmed.ncbi.nlm.nih.gov](https://pubmed.ncbi.nlm.nih.gov/37250466/)) [S11]

Animal pharmacology also tempers the claim. In α1-subunit knockout mice, α1-containing receptors were **not necessary for most acute ethanol effects** tested: anxiolysis, ataxia, anticonvulsant effects, hypnosis, and acute functional tolerance were largely unchanged, though locomotor stimulation increased. ([pubmed.ncbi.nlm.nih.gov](https://pubmed.ncbi.nlm.nih.gov/12606632/)) [S12] In rhesus macaques, α1-preferring compounds did **not** support a strong contribution of α1-containing GABA\_A receptors to alcohol reinforcement. ([pubmed.ncbi.nlm.nih.gov](https://pubmed.ncbi.nlm.nih.gov/24330519/)) [S13]

Likewise, for benzodiazepine reward, α1 may be less central than sometimes assumed. In a mouse intracranial self-stimulation study, diazepam’s reward-enhancing effects were preserved in **α1 point-mutant mice** but abolished in **α2 and α3 point-mutant mice**, suggesting α2/α3-containing receptors—not α1—were necessary for that reward-related effect. ([pubmed.ncbi.nlm.nih.gov](https://pubmed.ncbi.nlm.nih.gov/22763624/)) [S14]

## Mechanistic / biological rationale

The biological rationale is still reasonable. Because GABRA1 encodes a major synaptic GABA\_A receptor subunit in brain, changing its abundance or function could shift inhibitory tone in circuits relevant to **reward, withdrawal, tolerance, and drug cue processing**. GABA\_A receptors are direct pharmacologic targets of benzodiazepines and are functionally altered by ethanol exposure, so GABRA1 sits in a pathway that is unquestionably relevant to addiction neurobiology. ([ncbi.nlm.nih.gov](https://www.ncbi.nlm.nih.gov/gene/2554)) [S15]

The strongest mechanistic thread is in **alcohol-induced neuroadaptation**. Chronic ethanol can drive α1 receptor internalization and suppress Gabra1 transcription through HDAC-linked chromatin changes, potentially weakening normal inhibitory signaling and contributing to compensatory receptor-subtype remodeling, tolerance, anxiety-like withdrawal states, and escalated drinking in animal models. ([pubmed.ncbi.nlm.nih.gov](https://pubmed.ncbi.nlm.nih.gov/12859683/)) [S16]

For **benzodiazepine-type drugs**, α1-containing receptors appear relevant to some abuse-related properties—especially zolpidem-like discriminative and reinforcing effects in primates—but even here the circuitry is not simple, and some reward-related outcomes depend more on α2/α3-containing receptors than α1. ([pubmed.ncbi.nlm.nih.gov](https://pubmed.ncbi.nlm.nih.gov/24563183/)) [S17]

So, biologically, GABRA1 is plausibly involved in addiction-related processes, but the evidence suggests it is **one component of a broader GABAergic adaptation network**, not a uniquely established addiction determinant.

## Bottom line

GABRA1 is **plausibly and partially implicated** in addiction-related biology, with the best support coming from **preclinical alcohol dependence/neuroadaptation studies** and some **benzodiazepine/zolpidem abuse-liability literature**. However, the **human genetic evidence is limited and inconsistent**, and stronger evidence for addiction risk often points to other GABA receptor subunits or other genes altogether. Overall, the claim is **too broad to call clearly true**, but it is also **not unsupported**—hence **Mixed Evidence**. ([pubmed.ncbi.nlm.nih.gov](https://pubmed.ncbi.nlm.nih.gov/16792556/)) [S18]

## Cited sources

- **[S1]** GABRA1 gamma-aminobutyric acid type A receptor subunit alpha1 [Homo sapiens (human)] - Gene - NCBI — https://www.ncbi.nlm.nih.gov/gene/2554
- **[S2]** Association between GABRA1 and drinking behaviors in the collaborative study on the genetics of alcoholism sample - PubMed — https://pubmed.ncbi.nlm.nih.gov/16792556/
- **[S3]** Chronic ethanol consumption enhances internalization of alpha1 subunit-containing GABAA receptors in cerebral cortex - PubMed — https://pubmed.ncbi.nlm.nih.gov/12859683/
- **[S4]** Ethanol Exposure Regulates Gabra1 Expression via Histone Deacetylation at the Promoter in Cultured Cortical Neurons - PubMed — https://pubmed.ncbi.nlm.nih.gov/28798030/
- **[S5]** Histone deacetylases mediate GABAA receptor expression, physiology, and behavioral maladaptations in rat models of alcohol dependence - PubMed — https://pubmed.ncbi.nlm.nih.gov/29520058/
- **[S6]** Role of GABAA receptors in alcohol use disorders suggested by chronic intermittent ethanol (CIE) rodent model - PubMed — https://pubmed.ncbi.nlm.nih.gov/28931433/
- **[S7]** The behavioral pharmacology of zolpidem: evidence for the functional significance of α1-containing GABA(A) receptors - PubMed — https://pubmed.ncbi.nlm.nih.gov/24563183/
- **[S8]** Association between GABRA1 and drinking behaviors in the collaborative study on the genetics of alcoholism sample - PubMed — https://pubmed.ncbi.nlm.nih.gov/16792556/
- **[S9]** GABA Receptors Genes Polymorphisms and Alcohol Dependence: No Evidence of an Association in an Italian Male Population - PubMed — https://pubmed.ncbi.nlm.nih.gov/25191505/
- **[S10]** GABA(A) receptor alpha1, alpha4, and beta3 subunit mRNA and protein expression in the frontal cortex of human alcoholics - PubMed — https://pubmed.ncbi.nlm.nih.gov/9660306/
- **[S11]** Multivariate genome-wide association meta-analysis of over 1 million subjects identifies loci underlying multiple substance use disorders - PubMed — https://pubmed.ncbi.nlm.nih.gov/37250466/
- **[S12]** Deletion of GABAA receptor alpha 1 subunit-containing receptors alters responses to ethanol and other anesthetics - PubMed — https://pubmed.ncbi.nlm.nih.gov/12606632/
- **[S13]** Little evidence of a role for the α1GABAA subunit-containing receptor in a rhesus monkey model of alcohol drinking - PubMed — https://pubmed.ncbi.nlm.nih.gov/24330519/
- **[S14]** Differential roles of GABA(A) receptor subtypes in benzodiazepine-induced enhancement of brain-stimulation reward - PubMed — https://pubmed.ncbi.nlm.nih.gov/22763624/
- **[S15]** GABRA1 gamma-aminobutyric acid type A receptor subunit alpha1 [Homo sapiens (human)] - Gene - NCBI — https://www.ncbi.nlm.nih.gov/gene/2554
- **[S16]** Chronic ethanol consumption enhances internalization of alpha1 subunit-containing GABAA receptors in cerebral cortex - PubMed — https://pubmed.ncbi.nlm.nih.gov/12859683/
- **[S17]** The behavioral pharmacology of zolpidem: evidence for the functional significance of α1-containing GABA(A) receptors - PubMed — https://pubmed.ncbi.nlm.nih.gov/24563183/
- **[S18]** Association between GABRA1 and drinking behaviors in the collaborative study on the genetics of alcoholism sample - PubMed — https://pubmed.ncbi.nlm.nih.gov/16792556/
