## Supplementary material for "Understanding Substance Dependence: What Differentiates Addictive from Non-Addictive Drugs?": Folder for supplementary files: addictive-genes-claims-first100-065-elovl1-gene-protein-is-implicated-in-addiction-6f22bdf869a9.md

# addictive-genes-claims-first100-065-6f22bdf869a9

**Original claim:** ELOVL1 gene/protein is implicated in addiction.

## Claim

ELOVL1 gene/protein is implicated in addiction.

## Verdict

Verdict: Mixed Evidence

## Confidence

Medium

## Supporting evidence

The strongest human evidence I found is **indirect rather than addiction-specific**. In a 2022 human genetics study of nine genes involved in regulating ω-3 long-chain polyunsaturated fatty acid levels, the **9-gene set** was associated with **alcohol consumption, alcohol use disorder (AUD), and broad externalizing**. However, when genes were tested individually, **ELOVL1 showed association with broad externalizing**, while it **did not show association with alcohol consumption or AUD** in the reported table. That matters because externalizing traits overlap with substance-use liability, but they are not the same as addiction itself. ([pmc.ncbi.nlm.nih.gov](https://pmc.ncbi.nlm.nih.gov/articles/PMC9509483/?utm_source=openai)) [S1]

A second human clue comes from psychiatric genomics rather than SUD genetics directly. A transcriptome-wide association study (TWAS) of ADHD found that conditioning on predicted expression of **ELOVL1, CCDC24, and ARTN** explained **77.4% of the GWAS signal** at one ADHD locus. The paper also notes that ADHD-associated eQTL/phenotype links included risk-related traits such as smoking-related phenotypes. This again places ELOVL1 in an **externalizing/neurodevelopmental risk neighborhood** relevant to later substance misuse, but still not as direct proof of addiction involvement. ([nature.com](https://www.nature.com/articles/s41467-019-12450-9)) [S2]

There is also a modest preclinical transcriptomic signal. In a rat model of cocaine relapse vulnerability, **Elovl1** was among 28 transcripts upregulated in dorsomedial prefrontal cortex in the “exercise throughout withdrawal” condition that reduced cue-induced cocaine seeking. That suggests Elovl1 expression can vary in an addiction-relevant brain state, but the study did **not** show that Elovl1 itself drove the behavioral effect, nor that manipulating Elovl1 alters cocaine reinforcement or relapse. ([pmc.ncbi.nlm.nih.gov](https://pmc.ncbi.nlm.nih.gov/articles/PMC10593899/)) [S3]

## Contradicting or limiting evidence

The main limitation is that **direct SUD evidence is weak**. In a multivariate GWAS/meta-analysis of **over 1 million subjects** spanning problematic alcohol use, problematic tobacco use, cannabis use disorder, and opioid use disorder, the main genome-wide results emphasized loci/genes such as **DRD2** and **PDE4B**; **ELOVL1 was not reported among the paper’s main loci/results**. That does not prove ELOVL1 is irrelevant, but it argues against ELOVL1 being a currently established major addiction gene. ([pmc.ncbi.nlm.nih.gov](https://pmc.ncbi.nlm.nih.gov/articles/PMC10217792/)) [S4]

Even the best positive paper here stops short of supporting the exact claim. In the ω-3 pathway study, **ELOVL1’s individual signal was for broad externalizing**, not for alcohol consumption or AUD. And the same large addiction GWAS explicitly states that addiction liability and externalizing are **overlapping but distinct** constructs. So using those data to claim “ELOVL1 is implicated in addiction” is a stretch unless the statement is qualified as indirect or cross-trait. ([pmc.ncbi.nlm.nih.gov](https://pmc.ncbi.nlm.nih.gov/articles/PMC9509483/?utm_source=openai)) [S5]

Also, the core ELOVL1 literature is centered on **lipid metabolism, myelin biology, and neurological disease**, not addiction. Pathogenic **ELOVL1** variants are linked to hypomyelinating leukodystrophy-like syndromes, ichthyosis, movement disorder, spasticity, and deafness. In mice, Elovl1 deficiency shortens myelin sphingolipids, lowers galactosylceramides, causes hypomyelination, and worsens motor coordination. Those findings strongly support a CNS role, but they are **not addiction-specific evidence**. ([pubmed.ncbi.nlm.nih.gov](https://pubmed.ncbi.nlm.nih.gov/29496980/)) [S6]

## Mechanistic / biological rationale

Biologically, ELOVL1 is very plausible as a **modifier of neural function**. It is a fatty-acid elongase required for synthesis of **saturated and monounsaturated very-long-chain fatty acids**, and especially for production of **C24 sphingolipids**. These lipids are important structural components of membranes and myelin. ([pubmed.ncbi.nlm.nih.gov](https://pubmed.ncbi.nlm.nih.gov/20937905/?utm_source=openai)) [S7]

That matters because addiction is not only about neurotransmitters; it also involves **persistent membrane, glial, inflammatory, and white-matter changes**. More broadly, sphingolipid pathways have been implicated in reward- and drug-related biology, and studies show that **ethanol alters brain ceramide metabolism** and that **morphine withdrawal enhances sphingolipid metabolism**. So an enzyme upstream of very-long-chain sphingolipid production is mechanistically relevant in principle. Still, this is a **pathway-level rationale**, not direct proof that ELOVL1 specifically mediates addiction. ([pmc.ncbi.nlm.nih.gov](https://pmc.ncbi.nlm.nih.gov/articles/PMC4405906/?utm_source=openai)) [S8]

There are also ELOVL1-specific CNS mechanistic data that support plausibility. In reactive astrocytes, deleting **ELOVL1** reduced saturated-lipid–mediated neurotoxicity. In separate model systems, excess glial ELOVL1/very-long-chain fatty acids increased sphingosine-1-phosphate signaling, neuroinflammation, dopaminergic neuron loss, motor impairment, and shortened lifespan. These phenomena intersect with processes often discussed in addiction neuroscience—glial activation, lipid signaling, and dopaminergic vulnerability—but they still fall short of demonstrating effects on reinforcement, compulsive use, craving, or relapse. ([pubmed.ncbi.nlm.nih.gov](https://pubmed.ncbi.nlm.nih.gov/34616039/)) [S9]

## Bottom line

The claim is **too strong if read literally**. Current evidence suggests that **ELOVL1 may be indirectly relevant to addiction-related biology**, mainly through **externalizing/ADHD-associated genetics** and through **very-long-chain lipid/sphingolipid pathways** important for CNS homeostasis. But I did **not** find strong evidence that ELOVL1 is a **directly established addiction gene/protein** in human SUD genetics or in causal addiction-model experiments. A more accurate wording would be: **ELOVL1 is plausibly involved in neurobiological pathways that may influence addiction risk, but direct addiction-specific evidence is limited.** ([pmc.ncbi.nlm.nih.gov](https://pmc.ncbi.nlm.nih.gov/articles/PMC9509483/?utm_source=openai)) [S10]

## Cited sources

- **[S1]** Genes regulating levels of ω‐3 long‐chain polyunsaturated fatty acids are associated with alcohol use disorder and consumption, and broader externalizing behavior in humans - PMC — https://pmc.ncbi.nlm.nih.gov/articles/PMC9509483/?utm_source=openai
- **[S2]** Transcriptome-wide association study of attention deficit hyperactivity disorder identifies associated genes and phenotypes | Nature Communications — https://www.nature.com/articles/s41467-019-12450-9
- **[S3]** 
            Transcriptional Profile of Exercise-Induced Protection Against Relapse to Cocaine Seeking in a Rat Model - PMC
         — https://pmc.ncbi.nlm.nih.gov/articles/PMC10593899/
- **[S4]** 
            Multivariate genome-wide association meta-analysis of over 1 million subjects identifies loci underlying multiple substance use disorders - PMC
         — https://pmc.ncbi.nlm.nih.gov/articles/PMC10217792/
- **[S5]** Genes regulating levels of ω‐3 long‐chain polyunsaturated fatty acids are associated with alcohol use disorder and consumption, and broader externalizing behavior in humans - PMC — https://pmc.ncbi.nlm.nih.gov/articles/PMC9509483/?utm_source=openai
- **[S6]** Dominant ELOVL1 mutation causes neurological disorder with ichthyotic keratoderma, spasticity, hypomyelination and dysmorphic features - PubMed — https://pubmed.ncbi.nlm.nih.gov/29496980/
- **[S7]** ELOVL1 production of C24 acyl-CoAs is linked to C24 sphingolipid synthesis - PubMed — https://pubmed.ncbi.nlm.nih.gov/20937905/?utm_source=openai
- **[S8]** Ceramide metabolism analysis in a model of binge drinking reveals both neuroprotective and toxic effects of ethanol - PMC — https://pmc.ncbi.nlm.nih.gov/articles/PMC4405906/?utm_source=openai
- **[S9]** Neurotoxic reactive astrocytes induce cell death via saturated lipids - PubMed — https://pubmed.ncbi.nlm.nih.gov/34616039/
- **[S10]** Genes regulating levels of ω‐3 long‐chain polyunsaturated fatty acids are associated with alcohol use disorder and consumption, and broader externalizing behavior in humans - PMC — https://pmc.ncbi.nlm.nih.gov/articles/PMC9509483/?utm_source=openai
