## Supplementary material for "Understanding Substance Dependence: What Differentiates Addictive from Non-Addictive Drugs?": Folder for supplementary files: addictive-genes-claims-first100-066-cox7a1-gene-protein-is-implicated-in-addiction-6e2e2c41febb.md

# addictive-genes-claims-first100-066-6e2e2c41febb

**Original claim:** COX7A1 gene/protein is implicated in addiction.

## Claim

COX7A1 gene/protein is implicated in addiction.

## Verdict

Verdict: Likely False

The best available literature does **not** support COX7A1 as an established addiction gene/protein. What exists is mostly **indirect, low-level evidence**: a cocaine-responsive transcriptomic signal in one mouse ventral tegmental area study, plus a general rationale that mitochondrial/oxidative phosphorylation biology matters in substance-related neuroadaptations. By contrast, I did **not** find replicated human genetic, transcriptome-wide, or functional addiction studies that specifically identify COX7A1 as a risk gene or causal mediator. ([nature.com](https://www.nature.com/articles/s41386-021-01031-4)) [S1]

## Confidence

Medium

Confidence is **medium** because the negative case here is mainly based on the **absence of strong direct evidence** rather than a definitive falsifying experiment. However, the addiction-genetics literature is large enough that if COX7A1 were a robust addiction gene, one would expect it to appear in major reviews, replicated GWAS/TWAS findings, or functional validation studies; it does not. ([pmc.ncbi.nlm.nih.gov](https://pmc.ncbi.nlm.nih.gov/articles/PMC8960411/)) [S2]

## Supporting evidence

There is a biologically plausible **indirect** path from COX7A1 to addiction. COX7A1 encodes a nuclear-encoded subunit of cytochrome c oxidase (mitochondrial complex IV), and broader addiction literature supports roles for mitochondrial function, reactive oxygen species, and oxidative homeostasis in substance-related brain adaptations. Reviews describe mitochondria as participating in neuronal energy production, calcium handling, ROS generation, and drug-induced epigenetic/transcriptional changes, while an alcohol-withdrawal systems study implicated an oxidative-phosphorylation/oxidative-homeostasis brain network in addiction-relevant phenotypes. ([ncbi.nlm.nih.gov](https://www.ncbi.nlm.nih.gov/gene/1346)) [S3]

The most direct addiction-relevant signal I found is a **mouse cocaine transcriptomics** paper. In that study, cocaine exposure across several paradigms altered biological processes in the ventral tegmental area (VTA), and the authors specifically noted **Cox7a1** among genes appearing in a shared mitochondrial GO-term analysis across paradigms. That does put Cox7a1 inside an addiction-relevant brain region and molecular process. ([pmc.ncbi.nlm.nih.gov](https://pmc.ncbi.nlm.nih.gov/articles/PMC8357835/)) [S4]

COX7A1 also appears to be at least **detectable in human brain RNA datasets**, so a CNS role is not impossible. The Human Protein Atlas classifies it as **group enriched** overall in heart muscle, skeletal muscle, and tongue, but also reports brain RNA detection with **low regional specificity** and detection across brain regions. ([proteinatlas.org](https://www.proteinatlas.org/ENSG00000161281-COX7A1)) [S5]

## Contradicting or limiting evidence

The main problem with the claim is that the evidence is **too sparse and too indirect**. In the cocaine VTA paper, Cox7a1 was mentioned only as one gene contributing to a broader mitochondrial GO-term overlap; it was **not** one of the genes highlighted as correlating with cocaine intake. The genes the authors reported as correlating with intake were **Tnfrsf12a, Hip1, Idnk, Zbtb16, Mef2a,** and **Rn45s**, not Cox7a1. That makes the COX7A1 signal exploratory rather than a strong implication in addiction behavior. ([pmc.ncbi.nlm.nih.gov](https://pmc.ncbi.nlm.nih.gov/articles/PMC8357835/)) [S6]

Major human genetics reviews of substance use disorders do **not** identify COX7A1 as a replicated addiction locus. A broad 2021 review of SUD genetics highlights robust loci such as **ADH1B/ALDH2** for alcohol-related traits and the **CHRNA5-CHRNA3-CHRNB4** cluster for nicotine-related traits. A cocaine-focused human molecular genetics review discusses genes such as **KCTD20, PLCB1, NFAT5,** and cocaine-responsive miRNAs, and also notes that a TWAS found **no significant associations in any tissue** for cocaine dependence. COX7A1 is not presented as a validated human addiction gene in these syntheses. ([cambridge.org](https://www.cambridge.org/core/services/aop-cambridge-core/content/view/B3BAE9D2DCF78C7C4833A8DB4420F5B2/S0033291721000969a.pdf/genetics-of-substance-use-disorders-a-review.pdf)) [S7]

COX7A1’s best-established biology is also **not addiction-centered**. RefSeq/NCBI describes it as the **muscle isoform** of cytochrome c oxidase subunit VIIa. A detailed review of COX isoforms reports strongest COX7A1 expression in heart and skeletal muscle, while mouse knockout studies show reduced COX activity, impaired oxidative phosphorylation, cardiomyopathy, and exercise intolerance. Those phenotypes support a role in **bioenergetics**, especially in muscle and heart, but they do not constitute evidence for a specific addiction function. ([ncbi.nlm.nih.gov](https://www.ncbi.nlm.nih.gov/gene/1346)) [S8]

There is also some nuance on tissue expression. Older reference descriptions emphasize muscle specificity, whereas Human Protein Atlas RNA data show broader detectability including brain. Even with that broader detectability, HPA still places its strongest enrichment outside the brain and reports **low brain regional specificity**, which argues against a clearly defined addiction-circuit-specific role at present. ([ncbi.nlm.nih.gov](https://www.ncbi.nlm.nih.gov/gene/1346)) [S9]

## Mechanistic / biological rationale

From a mechanistic standpoint, the claim is **plausible in principle but unproven in practice**. Addiction involves persistent remodeling of reward circuitry, and those neurons are metabolically demanding. Because complex IV helps drive oxidative phosphorylation, any subunit that materially alters mitochondrial respiration could theoretically affect neuronal excitability, synaptic plasticity, ROS balance, and transcriptional adaptation after drug exposure. That broader logic is consistent with reviews linking mitochondrial function and oxidative stress to drug addiction biology. ([pubmed.ncbi.nlm.nih.gov](https://pubmed.ncbi.nlm.nih.gov/24956109/)) [S10]

But plausibility is not the same as implication. COX7A1 is one of several COX isoforms, and isoform biology matters: the isoform review describes **COX7A1** as the heart/skeletal-muscle-type form, while **COX7A2** is the more broadly expressed form across adult and fetal tissues. That weakens any assumption that COX7A1 would be a primary driver of brain addiction phenotypes specifically. ([pmc.ncbi.nlm.nih.gov](https://pmc.ncbi.nlm.nih.gov/articles/PMC5448071/)) [S11]

So the most defensible mechanistic statement is: **mitochondrial respiration is relevant to addiction, and COX7A1 is a mitochondrial respiration gene, therefore a role is conceivable**. What is missing are the key next steps that would elevate it from conceivable to implicated: replicated human association evidence, cell-type-specific brain expression/function in reward circuitry, and perturbation studies showing altered drug seeking, reinforcement, withdrawal, or relapse when COX7A1 is manipulated. ([pmc.ncbi.nlm.nih.gov](https://pmc.ncbi.nlm.nih.gov/articles/PMC8960411/)) [S12]

## Bottom line

COX7A1 is **not currently well supported as an addiction gene/protein**. The literature I found provides only **weak, indirect support**: one mouse cocaine transcriptomic observation plus a general mitochondrial rationale. Against that, there is **no clear replicated human genetic evidence**, no strong addiction-focused functional validation, and the gene’s established biology is much more centered on mitochondrial respiration in heart/skeletal muscle than on addiction circuitry. On current evidence, saying COX7A1 is “implicated in addiction” is **an overstatement**. ([pmc.ncbi.nlm.nih.gov](https://pmc.ncbi.nlm.nih.gov/articles/PMC8357835/)) [S13]

## Cited sources

- **[S1]** Cocaine induces paradigm-specific changes to the transcriptome within the ventral tegmental area | Neuropsychopharmacology — https://www.nature.com/articles/s41386-021-01031-4
- **[S2]** 
            Molecular genetics of cocaine use disorders in humans - PMC
         — https://pmc.ncbi.nlm.nih.gov/articles/PMC8960411/
- **[S3]** COX7A1 cytochrome c oxidase subunit 7A1 [Homo sapiens (human)] - Gene - NCBI — https://www.ncbi.nlm.nih.gov/gene/1346
- **[S4]** 
            Cocaine induces paradigm-specific changes to the transcriptome within the ventral tegmental area - PMC
         — https://pmc.ncbi.nlm.nih.gov/articles/PMC8357835/
- **[S5]** COX7A1 protein expression summary - The Human Protein Atlas — https://www.proteinatlas.org/ENSG00000161281-COX7A1
- **[S6]** 
            Cocaine induces paradigm-specific changes to the transcriptome within the ventral tegmental area - PMC
         — https://pmc.ncbi.nlm.nih.gov/articles/PMC8357835/
- **[S7]** S0033291721000969jrv 2189..2200 — https://www.cambridge.org/core/services/aop-cambridge-core/content/view/B3BAE9D2DCF78C7C4833A8DB4420F5B2/S0033291721000969a.pdf/genetics-of-substance-use-disorders-a-review.pdf
- **[S8]** COX7A1 cytochrome c oxidase subunit 7A1 [Homo sapiens (human)] - Gene - NCBI — https://www.ncbi.nlm.nih.gov/gene/1346
- **[S9]** COX7A1 cytochrome c oxidase subunit 7A1 [Homo sapiens (human)] - Gene - NCBI — https://www.ncbi.nlm.nih.gov/gene/1346
- **[S10]** Mitoepigenetics and drug addiction - PubMed — https://pubmed.ncbi.nlm.nih.gov/24956109/
- **[S11]** 
            Tissue- and Condition-Specific Isoforms of Mammalian Cytochrome c Oxidase Subunits: From Function to Human Disease - PMC
         — https://pmc.ncbi.nlm.nih.gov/articles/PMC5448071/
- **[S12]** 
            Molecular genetics of cocaine use disorders in humans - PMC
         — https://pmc.ncbi.nlm.nih.gov/articles/PMC8960411/
- **[S13]** 
            Cocaine induces paradigm-specific changes to the transcriptome within the ventral tegmental area - PMC
         — https://pmc.ncbi.nlm.nih.gov/articles/PMC8357835/
