## Supplementary material for "Understanding Substance Dependence: What Differentiates Addictive from Non-Addictive Drugs?": Folder for supplementary files: addictive-genes-claims-first100-067-larp1-gene-protein-is-implicated-in-addiction-ffc77a7cd0be.md

# addictive-genes-claims-first100-067-ffc77a7cd0be

**Original claim:** LARP1 gene/protein is implicated in addiction.

## Claim

LARP1 gene/protein is implicated in addiction.

## Verdict

Verdict: No Evidence

Current scholarly evidence does **not** establish LARP1 as an addiction gene/protein. I did **not** find peer-reviewed human genetic, clinical, or animal addiction studies directly showing that LARP1 alters risk for substance use disorder, is differentially implicated by addiction GWAS, or causally changes addiction-relevant behaviors such as self-administration, conditioned place preference, relapse, craving, or withdrawal. In addiction-focused reviews of mTOR signaling and systems-omics of substance use disorders, searches for “LARP1” returned no match. ([pubmed.ncbi.nlm.nih.gov](https://pubmed.ncbi.nlm.nih.gov/24666346/)) [S1]

## Confidence

Medium

The confidence is **medium** because the addiction literature is large and negative findings are hard to prove exhaustively, but the absence of direct evidence across the sources I reviewed was fairly consistent. ([pubmed.ncbi.nlm.nih.gov](https://pubmed.ncbi.nlm.nih.gov/24666346/)) [S2]

## Supporting evidence

What **is** supported is that LARP1 is a real, biologically important **brain-expressed RNA-binding protein**. Human data show LARP1 is expressed in developing cortex from early fetal stages onward and is present across major inhibitory and excitatory neuronal subclasses, with more modest expression in glia. Rare damaging variants in LARP1 have been linked to neurodevelopmental disorder phenotypes, including autism spectrum disorder. ([pmc.ncbi.nlm.nih.gov](https://pmc.ncbi.nlm.nih.gov/articles/PMC11418108/)) [S3]

There is also emerging evidence that Larp1 matters for brain function in mice. In a recent **preprint** indexed in PubMed, brain-specific Larp1 knockout reduced brain mass and neuronal density, depleted synaptic TOP mRNAs, and caused severe impairment in spatial learning and memory. That supports a role in neuronal development/plasticity, but it is **not** addiction evidence. ([pubmed.ncbi.nlm.nih.gov](https://pubmed.ncbi.nlm.nih.gov/41278816/)) [S4]

At the pathway level, LARP1 is a well-established downstream effector of **mTORC1**, a pathway that has been implicated in neuroadaptations to drugs of abuse. Reviews of addiction biology describe mTORC1 as promoting drug-related neuroadaptations and behaviors such as reward seeking, excessive intake, drug seeking, and relapse. ([pubmed.ncbi.nlm.nih.gov](https://pubmed.ncbi.nlm.nih.gov/24666346/)) [S5]

However, the fact that **mTORC1** is implicated in addiction does **not** automatically mean each mTOR effector is implicated. Indeed, the addiction-focused mTOR reviews I checked did not identify LARP1 specifically. ([pubmed.ncbi.nlm.nih.gov](https://pubmed.ncbi.nlm.nih.gov/24666346/)) [S6]

As a contrast, large addiction genetics studies currently highlight other genes instead. For opioid addiction/use disorder, major signals include **OPRM1**, **FURIN**, and **PPP6C**; for alcohol dependence, **ADH1B** is a replicated genome-wide signal; and for smoking behavior, strong human genetic evidence includes nicotinic-pathway genes such as **CHRNB2** and **CHRNA5**. LARP1 was not reported among the headline loci in these large studies. ([pmc.ncbi.nlm.nih.gov](https://pmc.ncbi.nlm.nih.gov/articles/PMC9682545/)) [S7]

## Contradicting or limiting evidence

The main limitation is that “implicated in addiction” can be interpreted broadly. If one means **indirectly plausible because it regulates translation downstream of mTOR in neurons**, then LARP1 has some mechanistic plausibility. But if one means **directly supported by addiction-specific human genetics, transcriptomics, proteomics, or behavioral experiments**, I did not find that evidence. ([pubmed.ncbi.nlm.nih.gov](https://pubmed.ncbi.nlm.nih.gov/24666346/)) [S8]

Another limitation is that pathway-level reviews themselves caution that, although mTOR signaling is involved in drug reward and neuroadaptation, its precise role in addictive disorders still needs more work. So even the broader pathway inference is not fully settled, much less the narrower claim about LARP1 specifically. ([pubmed.ncbi.nlm.nih.gov](https://pubmed.ncbi.nlm.nih.gov/32854585/)) [S9]

Also, some of the most brain-specific functional evidence for Larp1 comes from a **preprint**, not yet a fully peer-reviewed journal article, so those data should be treated as provisional. ([pubmed.ncbi.nlm.nih.gov](https://pubmed.ncbi.nlm.nih.gov/41278816/)) [S10]

## Mechanistic / biological rationale

Mechanistically, LARP1 binds mTORC1 via **RAPTOR**, interacts with **5′TOP mRNAs**, and represses their translation downstream of mTORC1. Structural work further shows that LARP1 directly binds the mRNA cap and adjacent TOP motif, preventing eIF4F assembly and thereby controlling ribosome-biogenesis/translation-related transcripts. ([pubmed.ncbi.nlm.nih.gov](https://pubmed.ncbi.nlm.nih.gov/25940091/)) [S11]

Other work shows that LARP1, together with **PABPC1**, shapes selective mRNA binding and translational repression, including transcripts related to mitochondrial functions. This is relevant because addiction involves durable changes in neuronal protein synthesis, metabolism, and synaptic plasticity. ([pmc.ncbi.nlm.nih.gov](https://pmc.ncbi.nlm.nih.gov/articles/PMC7797073/)) [S12]

So a reasonable **biological hypothesis** is that LARP1 could influence addiction-related neuroplasticity by modulating translation in reward-circuit neurons. But that remains a hypothesis. I did not find studies demonstrating that drugs of abuse change LARP1 in nucleus accumbens/VTA/PFC, or that LARP1 manipulation alters addiction phenotypes in vivo. ([pubmed.ncbi.nlm.nih.gov](https://pubmed.ncbi.nlm.nih.gov/25940091/)) [S13]

## Bottom line

LARP1 is a **brain-relevant, mTOR-regulated translational regulator**, and that gives it a **plausible but unproven** connection to addiction biology. Based on currently accessible scholarly evidence, it should **not** be described as an established addiction gene/protein. A more accurate statement would be: **“LARP1 has indirect mechanistic plausibility for addiction-related neuroplasticity, but direct addiction evidence is currently lacking.”** ([pubmed.ncbi.nlm.nih.gov](https://pubmed.ncbi.nlm.nih.gov/24666346/)) [S14]

I found LARP1 studies on **mTOR/TOP-mRNA regulation, neurodevelopment, and memory**, but not direct peer-reviewed studies showing that LARP1 is itself implicated in addiction. ([pmc.ncbi.nlm.nih.gov](https://pmc.ncbi.nlm.nih.gov/articles/PMC11418108/)) [S15]

## Cited sources

- **[S1]** mTOR complex 1: a key player in neuroadaptations induced by drugs of abuse - PubMed — https://pubmed.ncbi.nlm.nih.gov/24666346/
- **[S2]** mTOR complex 1: a key player in neuroadaptations induced by drugs of abuse - PubMed — https://pubmed.ncbi.nlm.nih.gov/24666346/
- **[S3]** 
            LARP1 haploinsufficiency is associated with an autosomal dominant neurodevelopmental disorder - PMC
         — https://pmc.ncbi.nlm.nih.gov/articles/PMC11418108/
- **[S4]** Larp1 supports brain growth and spatial memory via post-transcriptional control of the translation machinery - PubMed — https://pubmed.ncbi.nlm.nih.gov/41278816/
- **[S5]** mTOR complex 1: a key player in neuroadaptations induced by drugs of abuse - PubMed — https://pubmed.ncbi.nlm.nih.gov/24666346/
- **[S6]** mTOR complex 1: a key player in neuroadaptations induced by drugs of abuse - PubMed — https://pubmed.ncbi.nlm.nih.gov/24666346/
- **[S7]** 
            Cross-ancestry meta-analysis of opioid use disorder uncovers novel loci with predominant effects in brain regions associated with addiction - PMC
         — https://pmc.ncbi.nlm.nih.gov/articles/PMC9682545/
- **[S8]** mTOR complex 1: a key player in neuroadaptations induced by drugs of abuse - PubMed — https://pubmed.ncbi.nlm.nih.gov/24666346/
- **[S9]** The role of the mTOR pathway in models of drug-induced reward and the behavioural constituents of addiction - PubMed — https://pubmed.ncbi.nlm.nih.gov/32854585/
- **[S10]** Larp1 supports brain growth and spatial memory via post-transcriptional control of the translation machinery - PubMed — https://pubmed.ncbi.nlm.nih.gov/41278816/
- **[S11]** La-related Protein 1 (LARP1) Represses Terminal Oligopyrimidine (TOP) mRNA Translation Downstream of mTOR Complex 1 (mTORC1) - PubMed — https://pubmed.ncbi.nlm.nih.gov/25940091/
- **[S12]** 
            The mTOR regulated RNA-binding protein LARP1 requires PABPC1 for guided mRNA interaction - PMC
         — https://pmc.ncbi.nlm.nih.gov/articles/PMC7797073/
- **[S13]** La-related Protein 1 (LARP1) Represses Terminal Oligopyrimidine (TOP) mRNA Translation Downstream of mTOR Complex 1 (mTORC1) - PubMed — https://pubmed.ncbi.nlm.nih.gov/25940091/
- **[S14]** mTOR complex 1: a key player in neuroadaptations induced by drugs of abuse - PubMed — https://pubmed.ncbi.nlm.nih.gov/24666346/
- **[S15]** 
            LARP1 haploinsufficiency is associated with an autosomal dominant neurodevelopmental disorder - PMC
         — https://pmc.ncbi.nlm.nih.gov/articles/PMC11418108/
