## Supplementary material for "Understanding Substance Dependence: What Differentiates Addictive from Non-Addictive Drugs?": Folder for supplementary files: addictive-genes-claims-first100-068-slc18a2-gene-protein-is-implicated-in-addiction-7a648e35d086.md

# addictive-genes-claims-first100-068-7a648e35d086

**Original claim:** SLC18A2 gene/protein is implicated in addiction.

## Claim

SLC18A2 gene/protein is implicated in addiction. ([ncbi.nlm.nih.gov](https://www.ncbi.nlm.nih.gov/gene/6571)) [S1]

## Verdict

Verdict: Likely True

At a broad biological level, this claim is supported. SLC18A2 encodes VMAT2, the vesicular monoamine transporter that packages dopamine, serotonin, norepinephrine, and other monoamines into synaptic vesicles. Because addictive drugs—especially amphetamine-class psychostimulants—directly exploit monoamine storage and release, VMAT2 is mechanistically relevant to addiction. Human studies also report SLC18A2/VMAT2 associations with several substance-use phenotypes and altered VMAT2 measures in addicted populations. However, the **human genetic evidence is not definitive**: much of it comes from candidate-gene studies with modest sample sizes, some findings are nominal rather than genome-wide robust, and replication is incomplete. ([ncbi.nlm.nih.gov](https://www.ncbi.nlm.nih.gov/gene/6571)) [S2]

## Confidence

Medium. The mechanistic and preclinical evidence is strong, and there is meaningful human imaging/genetic support, but the claim is stronger for **VMAT2’s role in addiction biology**, particularly psychostimulants, than for **SLC18A2 as a conclusively established major human addiction-risk gene**. ([pubmed.ncbi.nlm.nih.gov](https://pubmed.ncbi.nlm.nih.gov/41166311/)) [S3]

## Supporting evidence

SLC18A2 is the gene for VMAT2, a proton-driven vesicular monoamine transporter that loads dopamine and other monoamines into synaptic vesicles for regulated release. NCBI and structural studies describe VMAT2 as central to monoamine storage/release, and recent cryo-EM work explicitly notes that VMAT2 contributes to amphetamine-induced psychostimulation and informs substance-abuse biology. This alone makes SLC18A2 highly plausible as an addiction-relevant gene/protein, because dopamine signaling in mesolimbic reward circuits is a core substrate of many addictive drugs. ([ncbi.nlm.nih.gov](https://www.ncbi.nlm.nih.gov/gene/6571)) [S4]

Human genetic studies provide direct, though not definitive, support. In a family-based study, polymorphisms in SLC18A2 were associated with **alcohol dependence**, and haplotypes also showed association in a combined alcohol/nicotine-dependent sample; the same study did **not** find association in the nicotine-only families. The authors concluded that SLC18A2 variants “might contribute” to substance dependence. ([pubmed.ncbi.nlm.nih.gov](https://pubmed.ncbi.nlm.nih.gov/15988470/)) [S5]

Additional candidate-gene studies link SLC18A2 to opioid and stimulant-related phenotypes. A study of opioid dependence reported that three intronic SLC18A2 SNPs were associated with opioid dependence, and noted prior evidence from a separate sample that other SLC18A2 SNPs were protective against opioid dependence with or without cocaine comorbidity. A systems-omics review of substance use disorders summarized SLC18A2 as showing **nominal associations** with heroin and cocaine addiction in human cohorts. ([pmc.ncbi.nlm.nih.gov](https://pmc.ncbi.nlm.nih.gov/articles/PMC6566135/)) [S6]

More recently, a 2024 study in methamphetamine use disorder found that lower **blood SLC18A2 mRNA** levels were present in people who used methamphetamine versus controls, and certain SLC18A2 variants related to methamphetamine-use severity or inhibitory-control performance. That does not prove causality, but it is another human line of evidence connecting SLC18A2 to addiction-related phenotypes. ([pubmed.ncbi.nlm.nih.gov](https://pubmed.ncbi.nlm.nih.gov/39394974/)) [S7]

Human neuroimaging also implicates VMAT2 in addiction. In cocaine dependence, PET imaging with [¹¹C]DTBZ found **lower striatal VMAT2 availability** across limbic, associative, and sensorimotor striatal subdivisions, consistent with altered presynaptic dopamine storage capacity in cocaine abusers. A human review of psychostimulant addiction likewise summarizes reduced VMAT2 availability in chronic cocaine abusers. ([pubmed.ncbi.nlm.nih.gov](https://pubmed.ncbi.nlm.nih.gov/22193525/)) [S8]

For methamphetamine users, the imaging literature is more complicated but still addiction-relevant. A PET study of people in **early abstinence** from methamphetamine found **increased**, not decreased, striatal VMAT2 binding, especially close to last use, and concluded that VMAT2 binding is strongly influenced by recent abstinence status. Even though the direction differs from cocaine studies, the finding still supports involvement of VMAT2 in methamphetamine-related brain adaptation. ([pmc.ncbi.nlm.nih.gov](https://pmc.ncbi.nlm.nih.gov/articles/PMC6671215/)) [S9]

Preclinical studies are especially supportive. In VMAT2 heterozygous knockout mice, reduced VMAT2 levels led to **reduced amphetamine-conditioned reward**, indicating that intact vesicular monoamine function contributes to amphetamine reward-related behavior. Separate work also showed that full knockout causes neonatal lethality and that partial deficiency can alter sensitivity to psychostimulants. ([pubmed.ncbi.nlm.nih.gov](https://pubmed.ncbi.nlm.nih.gov/9275230/)) [S10]

Pharmacology studies further reinforce this. Reviews of methamphetamine abuse describe methamphetamine as acting in part through VMAT2 to inhibit vesicular dopamine uptake and promote vesicular dopamine release, increasing cytosolic dopamine available for reverse transport via DAT. In rats, several VMAT2-targeting compounds reduce methamphetamine self-administration, and a 2024 paper reported that the VMAT2 inhibitor JPC-141 specifically decreased methamphetamine self-administration without reducing food-maintained responding. ([pubmed.ncbi.nlm.nih.gov](https://pubmed.ncbi.nlm.nih.gov/24484975/)) [S11]

## Contradicting or limiting evidence

The biggest limitation is that much of the human genetic literature for SLC18A2 in addiction comes from the **candidate-gene era**. Major reviews of addiction genetics note that many candidate-gene findings across substance dependence did not replicate consistently in later GWAS-era research, implying a substantial false-positive risk for small association studies in general. That caution applies to SLC18A2 as well. ([pmc.ncbi.nlm.nih.gov](https://pmc.ncbi.nlm.nih.gov/articles/PMC3095110/)) [S12]

Some specific results are mixed. The 2005 family study found association for alcohol dependence but **not nicotine dependence alone**. The heroin/cocaine evidence summarized in the systems-omics review was described as **nominal**, not definitive. The 2024 methamphetamine study was small and used **peripheral blood mRNA**, which is not the same as brain expression. ([pubmed.ncbi.nlm.nih.gov](https://pubmed.ncbi.nlm.nih.gov/15988470/)) [S13]

Imaging findings are also not uniform across substances or abstinence states. Cocaine dependence is associated with lower striatal VMAT2 availability, whereas methamphetamine studies have reported increased VMAT2 binding in early abstinence and other patterns after longer abstinence. That means VMAT2 is clearly involved in drug-related neuroadaptation, but its signal may reflect a mix of state effects, compensation, terminal integrity, and recent exposure rather than a simple one-direction biomarker of “addiction.” ([pubmed.ncbi.nlm.nih.gov](https://pubmed.ncbi.nlm.nih.gov/22193525/)) [S14]

Finally, not every manipulation of VMAT2 maps cleanly onto reward. In VMAT2-overexpressing mice, increased VMAT2 protected against methamphetamine neurotoxicity but did **not** increase methamphetamine-conditioned place preference. So VMAT2 affects toxicity and dopamine handling strongly, but its effect on reward-related behavior may depend on context, drug class, and direction of perturbation. ([pmc.ncbi.nlm.nih.gov](https://pmc.ncbi.nlm.nih.gov/articles/PMC4489556/)) [S15]

## Mechanistic / biological rationale

Biologically, the rationale is straightforward. VMAT2 packages monoamines into synaptic vesicles, thereby controlling how much dopamine is available for regulated exocytotic release and how much remains in the cytosol. Since addictive psychostimulants derive much of their reinforcing effect from boosting extracellular dopamine, a transporter that determines vesicular dopamine storage is intrinsically relevant to addiction biology. ([ncbi.nlm.nih.gov](https://www.ncbi.nlm.nih.gov/gene/6571)) [S16]

Recent structural studies show that amphetamine binds VMAT2 and can drive vesicular monoamine release through VMAT2-dependent exchange-like mechanisms. Earlier pharmacology reviews similarly describe methamphetamine as interacting at VMAT2 to inhibit vesicular uptake and promote dopamine redistribution/release. This places VMAT2 not at the periphery, but near the core of psychostimulant action. ([pubmed.ncbi.nlm.nih.gov](https://pubmed.ncbi.nlm.nih.gov/41166311/)) [S17]

VMAT2 also influences neuronal resilience. By sequestering dopamine into vesicles, it limits cytosolic dopamine oxidation and toxic byproduct formation. In animal models, higher VMAT2 protects against methamphetamine-induced dopaminergic damage, whereas reduced VMAT2 increases vulnerability. This matters because addiction is not only about acute reward; chronic drug exposure also produces neuroadaptations and sometimes neurotoxic stress in dopamine systems. ([pubmed.ncbi.nlm.nih.gov](https://pubmed.ncbi.nlm.nih.gov/20181938/)) [S18]

## Bottom line

Yes—**SLC18A2/VMAT2 is reasonably described as implicated in addiction**, especially in the biology of **psychostimulant addiction**. The strongest support comes from mechanism, animal models, drug-target studies, and human PET findings, with additional but less definitive support from human candidate-gene studies in alcohol, opioid, heroin/cocaine, and methamphetamine-related phenotypes. Still, the evidence does **not** yet justify calling SLC18A2 a conclusively proven major addiction gene in humans. The most accurate summary is: **implicated, particularly functionally and mechanistically; genetically suggestive, but not fully settled.** ([pubmed.ncbi.nlm.nih.gov](https://pubmed.ncbi.nlm.nih.gov/41166311/)) [S19]

## Cited sources

- **[S1]** SLC18A2 solute carrier family 18 member A2 [Homo sapiens (human)] - Gene - NCBI — https://www.ncbi.nlm.nih.gov/gene/6571
- **[S2]** SLC18A2 solute carrier family 18 member A2 [Homo sapiens (human)] - Gene - NCBI — https://www.ncbi.nlm.nih.gov/gene/6571
- **[S3]** Molecular basis of vesicular monoamine transport and neurological drug interactions - PubMed — https://pubmed.ncbi.nlm.nih.gov/41166311/
- **[S4]** SLC18A2 solute carrier family 18 member A2 [Homo sapiens (human)] - Gene - NCBI — https://www.ncbi.nlm.nih.gov/gene/6571
- **[S5]** Association of DNA polymorphisms in the synaptic vesicular amine transporter gene (SLC18A2) with alcohol and nicotine dependence - PubMed — https://pubmed.ncbi.nlm.nih.gov/15988470/
- **[S6]** 
            VMAT2 gene (SLC18A2) variants associated with a greater risk for developing opioid dependence - PMC
         — https://pmc.ncbi.nlm.nih.gov/articles/PMC6566135/
- **[S7]** Associations between methamphetamine use disorder and SLC18A1, SLC18A2, BDNF, and FAAH gene sequence variants and expression levels - PubMed — https://pubmed.ncbi.nlm.nih.gov/39394974/
- **[S8]** In vivo evidence for low striatal vesicular monoamine transporter 2 (VMAT2) availability in cocaine abusers - PubMed — https://pubmed.ncbi.nlm.nih.gov/22193525/
- **[S9]** 
            Increased Vesicular Monoamine Transporter Binding during Early Abstinence In Human Methamphetamine Users: Is VMAT2 a Stable Dopamine Neuron Biomarker? - PMC
         — https://pmc.ncbi.nlm.nih.gov/articles/PMC6671215/
- **[S10]** VMAT2 knockout mice: heterozygotes display reduced amphetamine-conditioned reward, enhanced amphetamine locomotion, and enhanced MPTP toxicity - PubMed — https://pubmed.ncbi.nlm.nih.gov/9275230/
- **[S11]** The vesicular monoamine transporter-2: an important pharmacological target for the discovery of novel therapeutics to treat methamphetamine abuse - PubMed — https://pubmed.ncbi.nlm.nih.gov/24484975/
- **[S12]** 
            Genetic Vulnerability and Susceptibility to Substance Dependence - PMC
         — https://pmc.ncbi.nlm.nih.gov/articles/PMC3095110/
- **[S13]** Association of DNA polymorphisms in the synaptic vesicular amine transporter gene (SLC18A2) with alcohol and nicotine dependence - PubMed — https://pubmed.ncbi.nlm.nih.gov/15988470/
- **[S14]** In vivo evidence for low striatal vesicular monoamine transporter 2 (VMAT2) availability in cocaine abusers - PubMed — https://pubmed.ncbi.nlm.nih.gov/22193525/
- **[S15]** 
            Increased Vesicular Monoamine Transporter 2 (VMAT2; Slc18a2) Protects against Methamphetamine Toxicity - PMC
         — https://pmc.ncbi.nlm.nih.gov/articles/PMC4489556/
- **[S16]** SLC18A2 solute carrier family 18 member A2 [Homo sapiens (human)] - Gene - NCBI — https://www.ncbi.nlm.nih.gov/gene/6571
- **[S17]** Molecular basis of vesicular monoamine transport and neurological drug interactions - PubMed — https://pubmed.ncbi.nlm.nih.gov/41166311/
- **[S18]** High regulatability favors genetic selection in SLC18A2, a vesicular monoamine transporter essential for life - PubMed — https://pubmed.ncbi.nlm.nih.gov/20181938/
- **[S19]** Molecular basis of vesicular monoamine transport and neurological drug interactions - PubMed — https://pubmed.ncbi.nlm.nih.gov/41166311/
