## Supplementary material for "Understanding Substance Dependence: What Differentiates Addictive from Non-Addictive Drugs?": Folder for supplementary files: addictive-genes-claims-first100-069-dync1i2-gene-protein-is-implicated-in-addiction-3df7ee59e7cf.md

# addictive-genes-claims-first100-069-3df7ee59e7cf

**Original claim:** DYNC1I2 gene/protein is implicated in addiction.

## Claim

DYNC1I2 gene/protein is implicated in addiction.

## Verdict

Verdict: Likely False

Current evidence does **not** support DYNC1I2 as an established addiction gene/protein. The strongest authoritative annotations for DYNC1I2 concern **neurodevelopmental disease**, not substance use or addiction, and recent large-scale genetics papers on substance use disorders emphasize other loci instead. What exists for DYNC1I2 is mostly **indirect biological plausibility** plus a few **weak secondary-database signals**, not convincing human or animal addiction evidence. ([ncbi.nlm.nih.gov](https://www.ncbi.nlm.nih.gov/gene/1781)) [S1]

## Confidence

Medium

## Supporting evidence

DYNC1I2 encodes a **cytoplasmic dynein 1 intermediate chain**, a non-catalytic component of the dynein motor complex involved in **retrograde transport of organelles and vesicles**. NCBI lists it as brain-expressed, and dynein biology more broadly is central to neuronal trafficking, axonal transport, and synaptic function. That gives DYNC1I2 a plausible **indirect** connection to neural processes that addiction depends on. ([ncbi.nlm.nih.gov](https://www.ncbi.nlm.nih.gov/gene/1781)) [S2]

There is also a small amount of **indirect addiction-related signal**. In a mouse study of voluntary alcohol drinking, **Dync1i2** appeared within an overrepresented pathway (“phagosome maturation”) among genes correlated with recent alcohol intake in **ventral tegmental area dopaminergic neurons**. However, the study did **not** identify Dync1i2 as a validated causal driver of alcohol-seeking or dependence; it was one gene inside a pathway-level enrichment result. ([pmc.ncbi.nlm.nih.gov](https://pmc.ncbi.nlm.nih.gov/articles/PMC4785873/)) [S3]

A secondary aggregation resource, Harmonizome/GWASdb, includes **DYNC1I2** in a broad **nicotine dependence** gene set, but with a low standardized value and without demonstrating a replicated, gene-specific causal association. Importantly, the ANCO-GeneDB methods show that such resources often include genes mapped from SNP-based sources using relatively loose thresholds, including GWASdb and PheWAS data, so inclusion alone is weak evidence. ([maayanlab.cloud](https://maayanlab.cloud/Harmonizome/gene_set/nicotine%2Bdependence/GWASdb%2BSNP-Disease%2BAssociations)) [S4]

## Contradicting or limiting evidence

Authoritative gene-disease resources do **not** present addiction as an established DYNC1I2-associated phenotype. NCBI Gene lists DYNC1I2 as encoding a dynein subunit and identifies its associated condition as **“neurodevelopmental disorder with microcephaly and structural brain anomalies.”** GenCC likewise classifies DYNC1I2 for that neurodevelopmental disorder, not for addiction. ([ncbi.nlm.nih.gov](https://www.ncbi.nlm.nih.gov/gene/1781)) [S5]

The strongest primary human evidence for DYNC1I2 concerns **recessive neurodevelopmental disease**. In the key human genetics paper, bi-allelic DYNC1I2 variants were linked to **syndromic microcephaly, intellectual disability, and cerebral malformations**, and zebrafish modeling supported a neurodevelopmental role. That is meaningful biology, but it is **not addiction evidence**. ([pubmed.ncbi.nlm.nih.gov](https://pubmed.ncbi.nlm.nih.gov/31079899/?utm_source=openai)) [S6]

Recent high-level reviews of substance use disorder genetics and large GWAS emphasize other replicated loci: for alcohol-related phenotypes, genes such as **ADH1B/ADH1C**; for cannabis use disorder, **CHRNA2**; for opioid use disorder, **OPRM1**, **FURIN**, and loci near **TSNARE1**; and for nicotine dependence, the **CHRNA5** region remains a major signal. These sources do not elevate DYNC1I2 as a recognized addiction locus. ([nature.com](https://www.nature.com/articles/s41380-024-02547-z)) [S7]

Even for nicotine dependence specifically, a recent multi-ancestral GWAS of clinically defined nicotine dependence reported replication at the **CHRNA5** locus and strong cross-trait genetic correlations, but did not present DYNC1I2 as a lead replicated finding. That does not prove DYNC1I2 has no effect, but it argues against calling it an established addiction gene. ([pubmed.ncbi.nlm.nih.gov](https://pubmed.ncbi.nlm.nih.gov/39974067/)) [S8]

## Mechanistic / biological rationale

A mechanistic case for DYNC1I2 is easy to imagine, but remains mostly **generic rather than specific**. Cytoplasmic dynein is the principal minus-end-directed microtubule motor in neurons and is crucial for retrograde movement of cargoes, signaling endosomes, and other organelles. Dynein dysfunction can disrupt neuronal trafficking, synaptic organization, and neurodevelopment. ([journals.biologists.com](https://journals.biologists.com/jcs/article/126/11/2319/32725/Microtubule-based-transport-basic-mechanisms?utm_source=openai)) [S9]

Because addiction involves long-term plasticity in mesolimbic circuitry, especially dopaminergic neurons and their targets, any gene affecting intracellular transport could in principle influence neural homeostasis relevant to reward learning. The alcohol-transcriptomics study in VTA dopamine neurons is consistent with that broad idea. ([pmc.ncbi.nlm.nih.gov](https://pmc.ncbi.nlm.nih.gov/articles/PMC4785873/)) [S10]

But that rationale is still **one step removed** from the claim. Many genes are important for neuronal transport or synaptic maintenance; only a subset have direct evidence linking them to addiction risk, reinforcement, withdrawal, relapse, or treatment response. For DYNC1I2, the retrieved literature is centered on **cell biology and neurodevelopment**, not addiction phenotypes. Its known human disease association is neurodevelopmental, and the available functional data point to brain development and dynein-complex biology rather than drug-related behavior. ([ncbi.nlm.nih.gov](https://www.ncbi.nlm.nih.gov/gene/1781)) [S11]

## Bottom line

DYNC1I2 has a **credible neuronal role** as part of the dynein transport machinery, and there are a few **weak, indirect** hints that it may sit in pathways touched by substance exposure. However, the present scholarly and authoritative evidence does **not** support saying that DYNC1I2 is meaningfully “implicated in addiction.” A more accurate statement would be: **DYNC1I2 is a neurodevelopmental/dynein transport gene with indirect relevance to neuronal biology, but it is not currently an established addiction gene/protein.** ([ncbi.nlm.nih.gov](https://www.ncbi.nlm.nih.gov/gene/1781)) [S12]

## Cited sources

- **[S1]** DYNC1I2 dynein cytoplasmic 1 intermediate chain 2 [Homo sapiens (human)] - Gene - NCBI — https://www.ncbi.nlm.nih.gov/gene/1781
- **[S2]** DYNC1I2 dynein cytoplasmic 1 intermediate chain 2 [Homo sapiens (human)] - Gene - NCBI — https://www.ncbi.nlm.nih.gov/gene/1781
- **[S3]** 
            Alcohol consumption induces global gene expression changes in VTA dopaminergic neurons - PMC
         — https://pmc.ncbi.nlm.nih.gov/articles/PMC4785873/
- **[S4]** Gene Set - nicotine dependence — https://maayanlab.cloud/Harmonizome/gene_set/nicotine%2Bdependence/GWASdb%2BSNP-Disease%2BAssociations
- **[S5]** DYNC1I2 dynein cytoplasmic 1 intermediate chain 2 [Homo sapiens (human)] - Gene - NCBI — https://www.ncbi.nlm.nih.gov/gene/1781
- **[S6]** Bi-allelic Variants in DYNC1I2 Cause Syndromic Microcephaly with Intellectual Disability, Cerebral Malformations, and Dysmorphic Facial Features - PubMed — https://pubmed.ncbi.nlm.nih.gov/31079899/?utm_source=openai
- **[S7]** The genetic landscape of substance use disorders | Molecular Psychiatry — https://www.nature.com/articles/s41380-024-02547-z
- **[S8]** Multi-ancestral genome-wide association study of clinically defined nicotine dependence reveals strong genetic correlations with other substance use disorders and health-related traits - PubMed — https://pubmed.ncbi.nlm.nih.gov/39974067/
- **[S9]** Microtubule-based transport – basic mechanisms, traffic rules and role in neurological pathogenesis | Journal of Cell Science | The Company of Biologists — https://journals.biologists.com/jcs/article/126/11/2319/32725/Microtubule-based-transport-basic-mechanisms?utm_source=openai
- **[S10]** 
            Alcohol consumption induces global gene expression changes in VTA dopaminergic neurons - PMC
         — https://pmc.ncbi.nlm.nih.gov/articles/PMC4785873/
- **[S11]** DYNC1I2 dynein cytoplasmic 1 intermediate chain 2 [Homo sapiens (human)] - Gene - NCBI — https://www.ncbi.nlm.nih.gov/gene/1781
- **[S12]** DYNC1I2 dynein cytoplasmic 1 intermediate chain 2 [Homo sapiens (human)] - Gene - NCBI — https://www.ncbi.nlm.nih.gov/gene/1781
