## Supplementary material for "Understanding Substance Dependence: What Differentiates Addictive from Non-Addictive Drugs?": Folder for supplementary files: addictive-genes-claims-first100-070-atp6v0b-gene-protein-is-implicated-in-addiction-8b6cc1221c87.md

# addictive-genes-claims-first100-070-8b6cc1221c87

**Original claim:** ATP6V0B gene/protein is implicated in addiction.

## Claim

ATP6V0B gene/protein is implicated in addiction.

## Verdict

Verdict: Mixed Evidence

There is **some indirect evidence** connecting ATP6V0B to addiction-related biology, but it is **not strong enough to treat ATP6V0B as an established addiction gene/protein**. The main supportive findings I found are: (1) a male-mouse nicotine self-administration single-nucleus RNA-seq study reporting **Atp6v0b** among downregulated energy-metabolism genes in ventral tegmental area cells, and (2) a human GWAS of problematic opioid prescription use in which the lead variant was an eQTL for **ATP6V0B** among several nearby genes. However, that GWAS itself favored **KDM4A** rather than ATP6V0B as the more likely causal gene, and the GWAS Catalog currently shows **no direct ATP6V0B entries**. ([sciencedirect.com](https://www.sciencedirect.com/science/article/pii/S1673852724002327?utm_source=openai)) [S1]

## Confidence

Low

## Supporting evidence

A 2024 single-nucleus transcriptomics study of the mouse ventral tegmental area during nicotine self-administration found addiction-phase transcriptional remodeling and specifically reported that **Atp6v0b** was among the downregulated energy-metabolism genes. That is a relevant signal because the VTA is a core reward-region for nicotine dependence, but it is still an observational expression change in one animal model, not a causal demonstration. ([sciencedirect.com](https://www.sciencedirect.com/science/article/pii/S1673852724002327)) [S2]

A 2021 human GWAS of **problematic opioid prescription use** reported that the lead signal in one locus was an eQTL for several nearby genes, including **ATP6V0B**. This means common human variation associated with an opioid-related phenotype may influence ATP6V0B expression at that locus. That said, the same paper noted that credible-set analysis pointed more strongly toward **KDM4A** as the probable causal gene in the region, so ATP6V0B was a plausible nearby effector, not the clearly established driver. ([pmc.ncbi.nlm.nih.gov](https://pmc.ncbi.nlm.nih.gov/articles/PMC8562028/)) [S3]

ATP6V0B is biologically positioned to matter for neuronal signaling. It encodes the **c'' subunit** of the membrane V0 domain of the vacuolar H\+-ATPase, and structural work describes the V-ATPase c-ring as containing nine ATP6V0C-encoded c subunits plus one **ATP6V0B-encoded c'' subunit**. Because addiction phenotypes depend heavily on synaptic transmission and plasticity, genes in this machinery are at least mechanistically plausible candidates for involvement. ([pubmed.ncbi.nlm.nih.gov](https://pubmed.ncbi.nlm.nih.gov/36074901/)) [S4]

## Contradicting or limiting evidence

The strongest limitation is the **lack of direct, replicated human genetic evidence**. A current search of the NHGRI-EBI GWAS Catalog for **ATP6V0B** returns **no mapped studies, variants, traits, or gene entries**, which is not what you would expect for a well-supported addiction gene. ([ebi.ac.uk](https://www.ebi.ac.uk/gwas/search?query=ATP6V0B)) [S5]

The human opioid GWAS signal is **ambiguous at the gene level**. The associated SNP was an eQTL for ATP6V0B, but also for multiple neighboring genes, and the authors explicitly stated that credible-set analysis favored **KDM4A**. So the study supports, at most, **indirect locus-level involvement**, not a clean ATP6V0B-specific association. ([pmc.ncbi.nlm.nih.gov](https://pmc.ncbi.nlm.nih.gov/articles/PMC8562028/)) [S6]

ATP6V0B also looks more like a **broad cellular/homeostatic gene** than an addiction-specific gene. Human Protein Atlas data classify it as having **low tissue specificity**, **low regional brain specificity**, and being **detected in all** major brain regions. That does not rule out involvement in addiction, but it weakens the idea that ATP6V0B is selectively tied to addiction biology rather than to general vesicular/lysosomal function. ([proteinatlas.org](https://www.proteinatlas.org/ENSG00000117410-ATP6V0B/brain)) [S7]

I also did **not identify** convincing studies showing that selective manipulation of ATP6V0B itself alters drug self-administration, withdrawal, craving, relapse, or other canonical addiction behaviors in animals or humans. The available evidence is mostly associative, transcriptomic, or mechanistic rather than causal.

## Mechanistic / biological rationale

ATP6V0B is part of the **V-ATPase proton pump**, which acidifies intracellular organelles. In neurons, V-ATPases generate the proton electrochemical gradient that powers neurotransmitter loading into synaptic vesicles; one study notes that loading of neurotransmitters into synaptic vesicles is energized by about **one V-ATPase molecule per vesicle**, and another review states that synaptic-vesicle acidification by V-ATPase is required for neurotransmitter loading and contributes to transmitter release machinery. ([pubmed.ncbi.nlm.nih.gov](https://pubmed.ncbi.nlm.nih.gov/36418452/)) [S8]

That matters for addiction because addictive drugs ultimately act through altered synaptic signaling within reward circuits. If ATP6V0B function changed enough to affect V-ATPase assembly or performance, it could in principle perturb vesicular filling, release probability, or broader neuronal homeostasis. This is a **reasonable biological inference**, not direct proof of addiction causality. ([pubmed.ncbi.nlm.nih.gov](https://pubmed.ncbi.nlm.nih.gov/36074901/)) [S9]

ATP6V0B is indeed expressed in addiction-relevant midbrain regions. Human brain expression datasets show ATP6V0B expression in **substantia nigra** and **ventral tegmental area**, among many other regions, consistent with a role in general neuronal function within dopaminergic circuitry. ([v24.proteinatlas.org](https://v24.proteinatlas.org/ENSG00000117410-ATP6V0B/tissue/midbrain)) [S10]

There is also evidence that ATP6V0B can modulate broader signaling pathways tied to cell homeostasis: one study identified ATP6V0B as an **E2F1 transcriptional target** and found that ectopic ATP6V0B expression increased **V-ATPase** and **mTORC1** activity. Since lysosomal signaling, autophagy, and mTOR-related plasticity are often discussed in neuroadaptation, this adds mechanistic plausibility. But again, this is **pathway relevance**, not addiction-specific evidence. ([pubmed.ncbi.nlm.nih.gov](https://pubmed.ncbi.nlm.nih.gov/26356814/)) [S11]

## Bottom line

ATP6V0B is **plausibly relevant** to addiction biology because it is a neuronal V-ATPase component involved in synaptic vesicle acidification/neurotransmitter loading, is expressed in reward-related midbrain regions, and appears in a small amount of addiction-related transcriptomic/genetic evidence. ([pubmed.ncbi.nlm.nih.gov](https://pubmed.ncbi.nlm.nih.gov/36074901/)) [S12]

But the current literature does **not** support calling ATP6V0B a **validated addiction gene/protein**. The evidence is best described as **limited and indirect**: a suggestive nicotine transcriptomic signal, an ambiguous opioid GWAS locus, and mechanistic plausibility grounded in synaptic homeostasis. ([sciencedirect.com](https://www.sciencedirect.com/science/article/pii/S1673852724002327?utm_source=openai)) [S13]

## Cited sources

- **[S1]** Nicotine-induced transcriptional changes and mitochondrial dysfunction in the ventral tegmental area revealed by single-nucleus transcriptomics - ScienceDirect — https://www.sciencedirect.com/science/article/pii/S1673852724002327?utm_source=openai
- **[S2]** Nicotine-induced transcriptional changes and mitochondrial dysfunction in the ventral tegmental area revealed by single-nucleus transcriptomics - ScienceDirect — https://www.sciencedirect.com/science/article/pii/S1673852724002327
- **[S3]** 
            Genome-wide association study of problematic opioid prescription use in 132,113 23andMe research participants of European ancestry - PMC
         — https://pmc.ncbi.nlm.nih.gov/articles/PMC8562028/
- **[S4]** ATP6V0C variants impair V-ATPase function causing a neurodevelopmental disorder often associated with epilepsy - PubMed — https://pubmed.ncbi.nlm.nih.gov/36074901/
- **[S5]** GWAS Catalog — https://www.ebi.ac.uk/gwas/search?query=ATP6V0B
- **[S6]** 
            Genome-wide association study of problematic opioid prescription use in 132,113 23andMe research participants of European ancestry - PMC
         — https://pmc.ncbi.nlm.nih.gov/articles/PMC8562028/
- **[S7]** Brain tissue expression of ATP6V0B - Summary - The Human Protein Atlas — https://www.proteinatlas.org/ENSG00000117410-ATP6V0B/brain
- **[S8]** Regulation of the mammalian-brain V-ATPase through ultraslow mode-switching - PubMed — https://pubmed.ncbi.nlm.nih.gov/36418452/
- **[S9]** ATP6V0C variants impair V-ATPase function causing a neurodevelopmental disorder often associated with epilepsy - PubMed — https://pubmed.ncbi.nlm.nih.gov/36074901/
- **[S10]** Tissue expression of ATP6V0B - Staining in midbrain - The Human Protein Atlas — https://v24.proteinatlas.org/ENSG00000117410-ATP6V0B/tissue/midbrain
- **[S11]** V-ATPase: a master effector of E2F1-mediated lysosomal trafficking, mTORC1 activation and autophagy - PubMed — https://pubmed.ncbi.nlm.nih.gov/26356814/
- **[S12]** ATP6V0C variants impair V-ATPase function causing a neurodevelopmental disorder often associated with epilepsy - PubMed — https://pubmed.ncbi.nlm.nih.gov/36074901/
- **[S13]** Nicotine-induced transcriptional changes and mitochondrial dysfunction in the ventral tegmental area revealed by single-nucleus transcriptomics - ScienceDirect — https://www.sciencedirect.com/science/article/pii/S1673852724002327?utm_source=openai
