## Supplementary material for "Understanding Substance Dependence: What Differentiates Addictive from Non-Addictive Drugs?": Folder for supplementary files: addictive-genes-claims-first100-071-iqgap1-gene-protein-is-implicated-in-addiction-2ae1b3b0d756.md

# addictive-genes-claims-first100-071-2ae1b3b0d756

**Original claim:** IQGAP1 gene/protein is implicated in addiction.

## Claim

IQGAP1 gene/protein is implicated in addiction. ([pubmed.ncbi.nlm.nih.gov](https://pubmed.ncbi.nlm.nih.gov/23018238/?utm_source=openai)) [S1]

## Verdict

Verdict: Mixed Evidence

## Confidence

Medium

## Supporting evidence

The strongest **direct** evidence is a 2022 human case-control study of **morphine dependence**. It enrolled **40 patients with morphine dependence and 40 healthy controls**, measured **IQGAP1 mRNA in whole blood**, and found **lower IQGAP1 levels** in the morphine-dependent group. Within the dependent group, lower IQGAP1 correlated with higher opioid addiction severity (OASI), and IQGAP1 remained an independent factor in their regression model. The same paper also used a dual-luciferase assay to show that **miR-124 directly targets IQGAP1**, supporting a plausible regulatory pathway rather than a purely descriptive association. This does count as evidence that IQGAP1 is implicated in **opioid/morphine dependence**, at least preliminarily. ([frontiersin.org](https://www.frontiersin.org/articles/10.3389/fpsyt.2022.845357/full)) [S2]

There is also substantial **biological plausibility** from neuroscience studies. In mice, IQGAP1 is abundant in brain tissue and functions in **NMDA receptor/ERK signaling**, **dendritic spine maintenance**, and **synaptic plasticity**. IQGAP1 knockout reduced surface **NR2A/GluN2A**, impaired ERK activation, reduced spine density in hippocampus and lateral amygdala, impaired a weak form of hippocampal LTP, and caused deficits in fear conditioning and object/context memory tasks. A later study further linked **IQGAP1/ERK signaling** to histone modifications involved in fear-memory formation. These are not addiction studies per se, but they place IQGAP1 in pathways that are highly relevant to maladaptive learning and drug-induced neuroplasticity. ([pubmed.ncbi.nlm.nih.gov](https://pubmed.ncbi.nlm.nih.gov/21653857/)) [S3]

That matters because addiction research consistently shows that **glutamatergic/NMDAR-dependent synaptic plasticity** in the **VTA, nucleus accumbens, and prefrontal cortex** contributes to sensitization, drug seeking, craving, and relapse-related behaviors. Reviews of addiction neurobiology emphasize that altered NMDAR-dependent plasticity and dendritic remodeling in mesocorticolimbic circuits are core features of the addicted brain. Since IQGAP1 regulates NMDAR trafficking, ERK signaling, and spine structure, it is mechanistically credible that it could influence addiction-related adaptations. ([pmc.ncbi.nlm.nih.gov](https://pmc.ncbi.nlm.nih.gov/articles/PMC4299443/)) [S4]

## Contradicting or limiting evidence

The direct addiction evidence is still **thin**. The morphine-dependence paper is a **single**, **small**, **cross-sectional** study using **peripheral blood** rather than brain tissue. The authors explicitly note the small sample size, inability to infer temporal change or causality, and possible confounding from smoking, drinking, and other substance-related factors common in this population. So the study supports “possible involvement,” but not that IQGAP1 is an established addiction gene. ([frontiersin.org](https://www.frontiersin.org/articles/10.3389/fpsyt.2022.845357/full)) [S5]

Large-scale human genetics do **not** currently place IQGAP1 among the better-supported opioid addiction genes. A large **cross-ancestry GWAS of opioid use disorder** reported genome-wide significant loci near genes such as **OPRM1, FURIN, NCAM1, KCNN1, RNF114, TSNARE1, FBXW4, BTNL2, and CDKAL1**, but **not IQGAP1**. That does not disprove a role for IQGAP1, but it argues against IQGAP1 being a replicated, high-confidence OUD locus at present. ([pmc.ncbi.nlm.nih.gov](https://pmc.ncbi.nlm.nih.gov/articles/PMC9682545/)) [S6]

Similarly, a 2023 **systematic review** of opioid functional-genomic studies screened **2,709 articles**, included **73** for qualitative analysis, and concluded that cross-study consistency was generally poor; **Cdkn1a** was the only gene consistently identified across many studies. IQGAP1 was **not highlighted** as a reproducible opioid genomic signal in the review text. Again, absence from such a review is not proof of no role, but it is important limiting evidence. ([nature.com](https://www.nature.com/articles/s41380-023-02238-1)) [S7]

An additional nuance is that the best mouse mechanistic paper on IQGAP1 concluded that IQGAP1 had a specific role in **cognitive** processes, while **anxiety-, depression-like, and motivational** behaviors were largely intact in their assays. That does not exclude addiction relevance, since addiction overlaps with learning and plasticity, but it weakens any claim that IQGAP1 is already clearly established as a core reward/motivation gene. ([pubmed.ncbi.nlm.nih.gov](https://pubmed.ncbi.nlm.nih.gov/21653857/)) [S8]

## Mechanistic / biological rationale

IQGAP1 is a **multifunctional scaffold protein** that links **Rho-family GTPases, the actin cytoskeleton, MAPK/ERK signaling, and receptor signaling complexes**. Reviews describe it as important in **neuronal development**, **spine regulation**, and **synaptic plasticity**. In other words, IQGAP1 sits at exactly the kind of signaling/cytoskeletal interface that often matters for long-term experience-dependent circuit remodeling. ([pubmed.ncbi.nlm.nih.gov](https://pubmed.ncbi.nlm.nih.gov/23018238/?utm_source=openai)) [S9]

In neurons specifically, IQGAP1 helps couple **GluN2A-containing NMDARs** to downstream **ERK** responses and supports normal dendritic spine density and plasticity. Addiction is widely modeled as a form of **pathological learning**, where repeated drug exposure reshapes synapses in reward circuits. On mechanistic grounds, an IQGAP1 perturbation could therefore affect how opioid or other drug exposure is translated into persistent synaptic and behavioral change. That is a reasonable inference from current biology, though still an inference. ([pubmed.ncbi.nlm.nih.gov](https://pubmed.ncbi.nlm.nih.gov/21653857/)) [S10]

The most specific proposed addiction-related mechanism so far is the **miR-124 → IQGAP1** axis in morphine dependence. In the human morphine study, miR-124 was elevated, IQGAP1 was reduced, and reporter assays supported direct binding of miR-124 to the IQGAP1 3′UTR. That provides a plausible molecular route by which opioid exposure could alter plasticity-related signaling, but it still falls short of proving that IQGAP1 drives drug taking, relapse, or compulsive use. ([frontiersin.org](https://www.frontiersin.org/articles/10.3389/fpsyt.2022.845357/full)) [S11]

## Bottom line

There is **some real evidence** that IQGAP1 is implicated in **morphine dependence**, mainly from one small human study plus a strong mechanistic fit with synaptic-plasticity pathways known to matter in addiction. But the evidence is **preliminary**, **not yet well replicated**, and does **not** currently justify calling IQGAP1 a broadly established addiction gene across substances. Overall, the claim is best judged as **Mixed Evidence**: plausible and partially supported for opioid-related dependence, but not strongly established for addiction in general. ([frontiersin.org](https://www.frontiersin.org/articles/10.3389/fpsyt.2022.845357/full)) [S12]

## Cited sources

- **[S1]** IQGAP1: A microtubule-microfilament scaffolding protein with multiple roles in nerve cell development and synaptic plasticity - PubMed — https://pubmed.ncbi.nlm.nih.gov/23018238/?utm_source=openai
- **[S2]** Frontiers | MiR-124 Regulates IQGAP1 and Participates in the Relationship Between Morphine Dependence Susceptibility and Cognition — https://www.frontiersin.org/articles/10.3389/fpsyt.2022.845357/full
- **[S3]** IQGAP1 regulates NR2A signaling, spine density, and cognitive processes - PubMed — https://pubmed.ncbi.nlm.nih.gov/21653857/
- **[S4]** 
            Glutamatergic synaptic plasticity in the mesocorticolimbic system in addiction - PMC
         — https://pmc.ncbi.nlm.nih.gov/articles/PMC4299443/
- **[S5]** Frontiers | MiR-124 Regulates IQGAP1 and Participates in the Relationship Between Morphine Dependence Susceptibility and Cognition — https://www.frontiersin.org/articles/10.3389/fpsyt.2022.845357/full
- **[S6]** 
            Cross-ancestry meta-analysis of opioid use disorder uncovers novel loci with predominant effects in brain regions associated with addiction - PMC
         — https://pmc.ncbi.nlm.nih.gov/articles/PMC9682545/
- **[S7]** Functional genomic mechanisms of opioid action and opioid use disorder: a systematic review of animal models and human studies | Molecular Psychiatry — https://www.nature.com/articles/s41380-023-02238-1
- **[S8]** IQGAP1 regulates NR2A signaling, spine density, and cognitive processes - PubMed — https://pubmed.ncbi.nlm.nih.gov/21653857/
- **[S9]** IQGAP1: A microtubule-microfilament scaffolding protein with multiple roles in nerve cell development and synaptic plasticity - PubMed — https://pubmed.ncbi.nlm.nih.gov/23018238/?utm_source=openai
- **[S10]** IQGAP1 regulates NR2A signaling, spine density, and cognitive processes - PubMed — https://pubmed.ncbi.nlm.nih.gov/21653857/
- **[S11]** Frontiers | MiR-124 Regulates IQGAP1 and Participates in the Relationship Between Morphine Dependence Susceptibility and Cognition — https://www.frontiersin.org/articles/10.3389/fpsyt.2022.845357/full
- **[S12]** Frontiers | MiR-124 Regulates IQGAP1 and Participates in the Relationship Between Morphine Dependence Susceptibility and Cognition — https://www.frontiersin.org/articles/10.3389/fpsyt.2022.845357/full
