## Supplementary material for "Understanding Substance Dependence: What Differentiates Addictive from Non-Addictive Drugs?": Folder for supplementary files: addictive-genes-claims-first100-072-gnb2-gene-protein-is-implicated-in-addiction-d6532b105c81.md

# addictive-genes-claims-first100-072-d6532b105c81

**Original claim:** GNB2 gene/protein is implicated in addiction.

## Claim

GNB2 gene/protein is implicated in addiction.

## Verdict

Verdict: Mixed Evidence

There is a **plausible mechanistic basis** and **some direct preclinical evidence** linking GNB2 to addiction-relevant neurobiology, especially opioid-related adaptations in the striatum/nucleus accumbens. However, the evidence is **not strong enough to place GNB2 among the established human addiction genes**. Most support comes from pathway membership and animal omics studies after drug exposure, not from replicated human genetic associations or decisive causal experiments focused on GNB2 itself. ([ncbi.nlm.nih.gov](https://www.ncbi.nlm.nih.gov/gene/2783)) [S1]

## Confidence

Medium

## Supporting evidence

GNB2 encodes **G protein subunit beta-2 (Gβ2)**, a component of heterotrimeric G proteins that transduce signals from many GPCRs. NCBI summarizes GNB2 as a beta subunit important for regulating alpha subunits and other signal-transduction effectors, and Human Protein Atlas indicates it is expressed in human tissues including brain, albeit without strong regional brain specificity. That makes it biologically credible as a modulator of reward-circuit signaling. ([ncbi.nlm.nih.gov](https://www.ncbi.nlm.nih.gov/gene/2783)) [S2]

Authoritative pathway resources place GNB2 squarely in **addiction-relevant signaling pathways**. KEGG lists human GNB2 in **morphine addiction** and **alcoholism**, and also in dopaminergic, glutamatergic, GABAergic, serotonergic, cholinergic, and endocannabinoid synapse pathways. Reactome places it within **GPCR downstream signaling**, specifically **G alpha(i) signaling events** and **opioid signaling**. Pathway inclusion is not proof of disease causality, but it does show that GNB2 participates in canonical molecular machinery used by addictive drugs and reward circuits. ([kegg.jp](https://www.kegg.jp/entry/hsa%3A10681%2Bhsa%3A2782%2Bhsa%3A2783%2Bhsa%3A2784%2Bhsa%3A2785%2Bhsa%3A2786%2Bhsa%3A2787%2Bhsa%3A2788%2Bhsa%3A2790%2Bhsa%3A2791%2Bhsa%3A2792%2Bhsa%3A2793%2Bhsa%3A51764%2Bhsa%3A54331%2Bhsa%3A55970%2Bhsa%3A59345%2Bhsa%3A94235%2BH00264%2BH00729%2BH00773%2BH00787%2BH01481%2BH01633%2BH02705%2BH02885%2BH02895)) [S3]

There is also **direct preclinical drug-exposure evidence**. In a rat striatal presynaptic proteomics study, chronic morphine altered GNB2 protein levels; the study’s seed list specifically included **downregulated GNB2** in the presynaptic fraction after morphine treatment. ([pmc.ncbi.nlm.nih.gov](https://pmc.ncbi.nlm.nih.gov/articles/PMC3197197/?utm_source=openai)) [S4]

A second proteomics study found that prolonged morphine exposure altered the striatal postsynaptic density proteome, with **GNB2 among the upregulated proteins** in the morphine-regulated network. The authors framed these synaptic changes as relevant to morphine side effects including tolerance, dependence, and addiction. Taken together with the presynaptic study, this suggests compartment-specific morphine-related remodeling of GNB2-containing signaling complexes rather than a simple global increase or decrease. ([pubmed.ncbi.nlm.nih.gov](https://pubmed.ncbi.nlm.nih.gov/26149443/?utm_source=openai)) [S5]

A third opioid-related study used RNA-seq after continuous versus interrupted morphine exposure and found significant regulation of **Gnb2** among downstream signaling molecules in the **nucleus accumbens** and **dorsal striatum** following interrupted morphine. Those are core brain regions in reward, reinforcement, and relapse biology. ([pmc.ncbi.nlm.nih.gov](https://pmc.ncbi.nlm.nih.gov/articles/PMC7608117/?utm_source=openai)) [S6]

More broadly, addiction literature supports the importance of **G-protein signaling changes** in drug dependence. A review on heterotrimeric G-protein subunits concluded that abused drugs alter G-protein subunit expression and coupling in brain regions important for dependence, tolerance, and withdrawal. Although that review emphasized GNB1 more than GNB2, it supports the general framework in which beta-subunit remodeling is addiction-relevant. ([pmc.ncbi.nlm.nih.gov](https://pmc.ncbi.nlm.nih.gov/articles/PMC2644661/)) [S7]

## Contradicting or limiting evidence

The biggest limitation is that **human evidence specific to GNB2 is sparse**. In a major umbrella review of molecular genetics of substance use disorders covering more than 150 meta-analyses, the repeatedly implicated genes were things like **OPRM1, DRD2, DRD4, BDNF, SLC6A4**, and **CHRNA5**; GNB2 was not highlighted as a recurring established signal. That strongly suggests GNB2 is **not currently a leading replicated human addiction gene**. ([sciencedirect.com](https://www.sciencedirect.com/science/article/abs/pii/S0149763421000336?utm_source=openai)) [S8]

Likewise, a very large multivariate GWAS/meta-analysis of over **1 million** individuals across problematic alcohol use, problematic tobacco use, cannabis use disorder, and opioid use disorder emphasized loci such as **PDE4B** and other dopamine-regulation-related genes as cross-substance signals. Based on the high-level results available from that study and the NIDA summary, GNB2 does not appear to be among the best-supported cross-disorder loci. ([pubmed.ncbi.nlm.nih.gov](https://pubmed.ncbi.nlm.nih.gov/37250466/?utm_source=openai)) [S9]

Another limitation is that much of the direct GNB2 evidence comes from **omics correlation after drug exposure**. Changes in mRNA or protein abundance after morphine do not prove that GNB2 is a driver of addictive behavior; they may reflect downstream adaptation, cell-type shifts, or synaptic redistribution. The morphine proteomics studies themselves support neuroadaptation, but they do not establish that manipulating GNB2 alone changes self-administration, craving, relapse, or compulsive use. ([pmc.ncbi.nlm.nih.gov](https://pmc.ncbi.nlm.nih.gov/articles/PMC3197197/?utm_source=openai)) [S10]

Finally, when GNB2 has been firmly linked to human disease, the strongest evidence is for **cardiac conduction disease** and **neurodevelopmental disorders**, not addiction. That does not argue against a role in addiction, but it shows where the current evidence base is strongest. ([pubmed.ncbi.nlm.nih.gov](https://pubmed.ncbi.nlm.nih.gov/28219978/?utm_source=openai)) [S11]

## Mechanistic / biological rationale

The mechanistic rationale is real. Addictive drugs heavily engage **GPCR signaling** in reward circuits, especially opioid, dopamine, GABA, and related neuromodulatory pathways. GNB2 encodes a beta subunit of heterotrimeric G proteins, placing it near the center of that signaling architecture. KEGG and Reactome both map GNB2 to opioid/morphine addiction-related signaling and multiple neurotransmitter synapse pathways. ([ncbi.nlm.nih.gov](https://www.ncbi.nlm.nih.gov/gene/2783)) [S12]

One especially relevant mechanism is **Gβγ-mediated GIRK channel regulation**. A functional study showed that Gβ2γ2 can activate GIRK channels, and a review of μ-opioid receptor biased signaling notes that the **G_i/o βγ** branch activates GIRK channels and inhibits voltage-gated calcium channels, suppressing neuronal excitability. GIRK channels themselves have been reviewed as relevant to **learning, reward, and addiction**, including drug-induced adaptations in mesolimbic circuits. So even without addiction-specific GNB2 genetics, there is a biologically coherent path from GNB2-containing Gβγ dimers to opioid and reward-circuit plasticity. ([pubmed.ncbi.nlm.nih.gov](https://pubmed.ncbi.nlm.nih.gov/10944236/?utm_source=openai)) [S13]

That said, this remains mostly a **mechanistic plausibility argument** plus some morphine-responsive omics data. It is not the same as showing that GNB2 variants or GNB2-targeted interventions materially alter addiction risk in humans. ([pmc.ncbi.nlm.nih.gov](https://pmc.ncbi.nlm.nih.gov/articles/PMC7608117/?utm_source=openai)) [S14]

## Bottom line

GNB2 is **reasonably implicated in addiction-related biology**, especially through **GPCR/Gβγ signaling** and **opioid-induced synaptic adaptations** in preclinical models. But the evidence is **not yet strong, replicated, or human-centered enough** to call GNB2 an established addiction gene on par with better-supported loci like **OPRM1, DRD2, or CHRNA5**. The fairest summary is: **mechanistically plausible and supported by some animal/drug-response data, but not firmly established in human addiction genetics or causal addiction experiments.** ([pubmed.ncbi.nlm.nih.gov](https://pubmed.ncbi.nlm.nih.gov/26149443/?utm_source=openai)) [S15]

## Cited sources

- **[S1]** GNB2 G protein subunit beta 2 [Homo sapiens (human)] - Gene - NCBI — https://www.ncbi.nlm.nih.gov/gene/2783
- **[S2]** GNB2 G protein subunit beta 2 [Homo sapiens (human)] - Gene - NCBI — https://www.ncbi.nlm.nih.gov/gene/2783
- **[S3]** KEGG T01001: 10681 — https://www.kegg.jp/entry/hsa%3A10681%2Bhsa%3A2782%2Bhsa%3A2783%2Bhsa%3A2784%2Bhsa%3A2785%2Bhsa%3A2786%2Bhsa%3A2787%2Bhsa%3A2788%2Bhsa%3A2790%2Bhsa%3A2791%2Bhsa%3A2792%2Bhsa%3A2793%2Bhsa%3A51764%2Bhsa%3A54331%2Bhsa%3A55970%2Bhsa%3A59345%2Bhsa%3A94235%2BH00264%2BH00729%2BH00773%2BH00787%2BH01481%2BH01633%2BH02705%2BH02885%2BH02895
- **[S4]** Chronic Morphine Alters the Presynaptic Protein Profile: Identification of Novel Molecular Targets Using Proteomics and Network Analysis - PMC — https://pmc.ncbi.nlm.nih.gov/articles/PMC3197197/?utm_source=openai
- **[S5]** Morphine Regulated Synaptic Networks Revealed by Integrated Proteomics and Network Analysis - PubMed — https://pubmed.ncbi.nlm.nih.gov/26149443/?utm_source=openai
- **[S6]** Interruption of continuous opioid exposure exacerbates drug-evoked adaptations in the mesolimbic dopamine system - PMC — https://pmc.ncbi.nlm.nih.gov/articles/PMC7608117/?utm_source=openai
- **[S7]** 
            Alterations in the levels of heterotrimeric G protein subunits induced by psychostimulants, opiates, barbiturates, and ethanol: Implications for drug dependence, tolerance, and withdrawal - PMC
         — https://pmc.ncbi.nlm.nih.gov/articles/PMC2644661/
- **[S8]** Molecular genetics of substance use disorders: An umbrella review - ScienceDirect — https://www.sciencedirect.com/science/article/abs/pii/S0149763421000336?utm_source=openai
- **[S9]** Multivariate genome-wide association meta-analysis of over 1 million subjects identifies loci underlying multiple substance use disorders - PubMed — https://pubmed.ncbi.nlm.nih.gov/37250466/?utm_source=openai
- **[S10]** Chronic Morphine Alters the Presynaptic Protein Profile: Identification of Novel Molecular Targets Using Proteomics and Network Analysis - PMC — https://pmc.ncbi.nlm.nih.gov/articles/PMC3197197/?utm_source=openai
- **[S11]** A Mutation in the G-Protein Gene GNB2 Causes Familial Sinus Node and Atrioventricular Conduction Dysfunction - PubMed — https://pubmed.ncbi.nlm.nih.gov/28219978/?utm_source=openai
- **[S12]** GNB2 G protein subunit beta 2 [Homo sapiens (human)] - Gene - NCBI — https://www.ncbi.nlm.nih.gov/gene/2783
- **[S13]** Activation and inhibition of G protein-coupled inwardly rectifying potassium (Kir3) channels by G protein beta gamma subunits - PubMed — https://pubmed.ncbi.nlm.nih.gov/10944236/?utm_source=openai
- **[S14]** Interruption of continuous opioid exposure exacerbates drug-evoked adaptations in the mesolimbic dopamine system - PMC — https://pmc.ncbi.nlm.nih.gov/articles/PMC7608117/?utm_source=openai
- **[S15]** Morphine Regulated Synaptic Networks Revealed by Integrated Proteomics and Network Analysis - PubMed — https://pubmed.ncbi.nlm.nih.gov/26149443/?utm_source=openai
