## Supplementary material for "Understanding Substance Dependence: What Differentiates Addictive from Non-Addictive Drugs?": Folder for supplementary files: addictive-genes-claims-first100-073-cldn11-gene-protein-is-implicated-in-addiction-d2ffef0a9ad7.md

# addictive-genes-claims-first100-073-d2ffef0a9ad7

**Original claim:** CLDN11 gene/protein is implicated in addiction.

## Claim

CLDN11 gene/protein is implicated in addiction.

## Verdict

Verdict: Mixed Evidence

There is **some evidence** linking **CLDN11 (claudin-11)** to addiction-related biology, but it is **limited, indirect, and not well replicated**. The strongest direct evidence is from **small transcriptomic/postmortem studies** showing altered CLDN11 expression in cocaine abuse and from **animal alcohol studies** showing Cldn11 downregulation in reward circuitry. However, I did **not** find strong evidence that CLDN11 is a **major, replicated human addiction-risk gene** in large modern GWAS, nor strong causal evidence that manipulating CLDN11 specifically changes addictive behavior. Overall, the claim is **plausible but not firmly established**. ([pubmed.ncbi.nlm.nih.gov](https://pubmed.ncbi.nlm.nih.gov/18357522/?utm_source=openai)) [S1]

## Confidence

Medium

Confidence is **medium** because the direction of the available evidence is coherent—CLDN11 is a CNS myelin/oligodendrocyte gene, and myelin biology is increasingly implicated in substance use disorders—but the **direct CLDN11-specific evidence base is sparse**, based largely on small expression studies and indirect mechanistic inference rather than replicated human genetics or intervention studies. ([ncbi.nlm.nih.gov](https://www.ncbi.nlm.nih.gov/gene/5010/ortholog/?utm_source=openai)) [S2]

## Supporting evidence

**1) Human postmortem cocaine data show altered CLDN11 expression.** A postmortem study of brains from cocaine abusers measured multiple myelin-related transcripts and reported that **claudin-11 expression was decreased in the caudate**, alongside broader evidence of altered myelin-related gene expression after chronic cocaine abuse. This is direct human evidence that CLDN11 is altered in at least one substance-use context, although it is expression evidence rather than genetic causation. ([pubmed.ncbi.nlm.nih.gov](https://pubmed.ncbi.nlm.nih.gov/18357522/?utm_source=openai)) [S3]

**2) Animal alcohol data also implicate Cldn11 in reward-related circuitry.** In alcohol-preferring rats exposed to repeated excessive binge-like drinking, **Cldn11 was downregulated in the ventral tegmental area (VTA)**, a core reward-region heavily involved in addiction neurobiology. That does not prove CLDN11 causes addiction, but it places the gene within a neuroanatomically relevant alcohol-responsive program. ([pmc.ncbi.nlm.nih.gov](https://pmc.ncbi.nlm.nih.gov/articles/PMC3704142/)) [S4]

**3) Addiction broadly involves white matter/myelin abnormalities, which makes CLDN11 biologically relevant.** CLDN11 is a **major CNS myelin component** expressed in the oligodendrocyte lineage and contributes to oligodendrocyte proliferation/migration and myelin integrity. Independent human imaging work shows **widespread white-matter abnormalities** in cocaine and heroin use disorders, and these abnormalities correlate with craving and duration of use. This does not single out CLDN11, but it strengthens the plausibility that a core myelin gene could matter in addiction-related neuroadaptation. ([ncbi.nlm.nih.gov](https://www.ncbi.nlm.nih.gov/gene/5010/ortholog/?utm_source=openai)) [S5]

**4) Modern addiction neuroscience supports a causal role for myelin plasticity in reward learning.** A 2024 Nature study showed that **myelin plasticity/oligodendrogenesis in the VTA is required for opioid reward learning**; blocking oligodendrogenesis dampened nucleus accumbens dopamine dynamics and impaired morphine-conditioned behavior. Again, this is not CLDN11-specific, but it provides an important mechanistic bridge: genes essential for myelin structure, like CLDN11, sit in a pathway now known to influence reward-related plasticity. ([pubmed.ncbi.nlm.nih.gov](https://pubmed.ncbi.nlm.nih.gov/38839962/)) [S6]

## Contradicting or limiting evidence

**1) There is little evidence that CLDN11 is a replicated human addiction susceptibility gene.** Recent very large OUD GWAS identified loci such as **OPRM1, FURIN, RABEPK, FBXW4, NCAM1, KCNN1, and TSNARE1**, not CLDN11, among their lead signals. That does not exclude a modest role for CLDN11, but it argues against calling it a strongly established addiction gene on current human genetic evidence. ([pubmed.ncbi.nlm.nih.gov](https://pubmed.ncbi.nlm.nih.gov/36171425/)) [S7]

**2) Cross-study convergence in addiction functional genomics is generally weak, and CLDN11 is not a standout recurring signal.** A systematic review of functional genomic mechanisms in opioid action/OUD concluded that published omics results show **surprisingly low consistency**, with only one gene recurring across eleven studies in their synthesis. That broader lack of reproducibility lowers confidence in any single CLDN11 expression hit unless independently replicated. ([nature.com](https://www.nature.com/articles/s41380-023-02238-1)) [S8]

**3) The direct CLDN11 studies are small and mostly associative.** The cocaine postmortem study involved a relatively small matched sample and reported **region-specific** rather than uniform changes; notably, it did **not** find changes in nucleus accumbens in that analysis. Such studies can be influenced by chronic illness burden, polysubstance exposure, agonal factors, and postmortem confounding. ([pubmed.ncbi.nlm.nih.gov](https://pubmed.ncbi.nlm.nih.gov/18357522/?utm_source=openai)) [S9]

**4) CLDN11 knockout phenotypes are neurobehavioral, but not addiction-specific.** In mice, loss of Cldn11 perturbs behavior and neurotransmitter levels, including reduced anxiety-like behavior and altered auditory processing, consistent with disrupted myelin-dependent circuit timing. But these models have **not** established that CLDN11 deficiency itself produces drug seeking, sensitization, self-administration, relapse, or other core addiction phenotypes. ([pubmed.ncbi.nlm.nih.gov](https://pubmed.ncbi.nlm.nih.gov/29491447/)) [S10]

## Mechanistic / biological rationale

CLDN11 encodes **claudin-11**, a tight-junction protein that is a **major structural component of CNS myelin** and is largely restricted in the CNS to the oligodendrocyte lineage. Loss of claudin-11 disrupts radial tight junctions in myelin, slows conduction along myelinated axons, and changes neurotransmitter balance and behavior in mice. That makes CLDN11 a biologically credible contributor to disorders in which **timing, synchrony, and plasticity of distributed circuits** are altered. ([pmc.ncbi.nlm.nih.gov](https://pmc.ncbi.nlm.nih.gov/articles/PMC10827939/)) [S11]

This rationale fits addiction biology reasonably well. Substance use disorders are increasingly associated with **white-matter disruption**, oligodendrocyte/myelin changes, and reward-circuit myelin plasticity. Alcohol studies review a strong link between ethanol exposure and myelin dysfunction, and newer experimental work indicates that oligodendrogenesis in the VTA is functionally involved in opioid reward learning. If addiction recruits or damages myelin-regulated circuit timing, then CLDN11 is a plausible participant because it helps maintain the electrical properties of myelinated axons. ([pmc.ncbi.nlm.nih.gov](https://pmc.ncbi.nlm.nih.gov/articles/PMC6597285/)) [S12]

Still, that is a **pathway-level inference**, not proof of CLDN11-specific causation. Many myelin genes could contribute similarly, and available evidence currently supports **myelin/oligodendrocyte biology in addiction** more strongly than it supports **CLDN11 as a specific addiction gene**. ([pmc.ncbi.nlm.nih.gov](https://pmc.ncbi.nlm.nih.gov/articles/PMC12633287/)) [S13]

## Bottom line

CLDN11 is **not strongly established as a canonical addiction gene**, but there is **credible, limited evidence** that it is involved in **addiction-relevant biology**, mainly through **myelin/oligodendrocyte pathways**. The best direct support comes from **altered CLDN11 expression in human cocaine brain tissue** and **Cldn11 downregulation in alcohol-exposed rat VTA**, while broader myelin plasticity studies make the link biologically plausible. The main weakness is the absence of **robust replicated human genetic or causal behavioral evidence specific to CLDN11**. So the most accurate summary is: **possible/indirect implication, not yet firmly proven.** ([pubmed.ncbi.nlm.nih.gov](https://pubmed.ncbi.nlm.nih.gov/18357522/?utm_source=openai)) [S14]

## Cited sources

- **[S1]** Expression of transcripts for myelin related genes in postmortem brain from cocaine abusers - PubMed — https://pubmed.ncbi.nlm.nih.gov/18357522/?utm_source=openai
- **[S2]** CLDN11 orthologs - NCBI — https://www.ncbi.nlm.nih.gov/gene/5010/ortholog/?utm_source=openai
- **[S3]** Expression of transcripts for myelin related genes in postmortem brain from cocaine abusers - PubMed — https://pubmed.ncbi.nlm.nih.gov/18357522/?utm_source=openai
- **[S4]** 
            Changes in Gene Expression within the Ventral Tegmental Area Following Repeated Excessive Binge-Like Alcohol Drinking by Alcohol-Preferring (P) Rats - PMC
         — https://pmc.ncbi.nlm.nih.gov/articles/PMC3704142/
- **[S5]** CLDN11 orthologs - NCBI — https://www.ncbi.nlm.nih.gov/gene/5010/ortholog/?utm_source=openai
- **[S6]** Myelin plasticity in the ventral tegmental area is required for opioid reward - PubMed — https://pubmed.ncbi.nlm.nih.gov/38839962/
- **[S7]** Cross-ancestry meta-analysis of opioid use disorder uncovers novel loci with predominant effects in brain regions associated with addiction - PubMed — https://pubmed.ncbi.nlm.nih.gov/36171425/
- **[S8]** Functional genomic mechanisms of opioid action and opioid use disorder: a systematic review of animal models and human studies | Molecular Psychiatry — https://www.nature.com/articles/s41380-023-02238-1
- **[S9]** Expression of transcripts for myelin related genes in postmortem brain from cocaine abusers - PubMed — https://pubmed.ncbi.nlm.nih.gov/18357522/?utm_source=openai
- **[S10]** Absence of Claudin 11 in CNS Myelin Perturbs Behavior and Neurotransmitter Levels in Mice - PubMed — https://pubmed.ncbi.nlm.nih.gov/29491447/
- **[S11]** 
            Claudin-11 in health and disease: implications for myelin disorders, hearing, and fertility - PMC
         — https://pmc.ncbi.nlm.nih.gov/articles/PMC10827939/
- **[S12]** 
            Function and Mechanism of Myelin Regulation in Alcohol Abuse and Alcoholism - PMC
         — https://pmc.ncbi.nlm.nih.gov/articles/PMC6597285/
- **[S13]** 
            Variation of Myelin-associated Gene Expression and Ndrg1 within the Prefrontal Cortex as Determinants for Initial Level of Response to Alcohol - PMC
         — https://pmc.ncbi.nlm.nih.gov/articles/PMC12633287/
- **[S14]** Expression of transcripts for myelin related genes in postmortem brain from cocaine abusers - PubMed — https://pubmed.ncbi.nlm.nih.gov/18357522/?utm_source=openai
