## Supplementary material for "Understanding Substance Dependence: What Differentiates Addictive from Non-Addictive Drugs?": Folder for supplementary files: addictive-genes-claims-first100-074-chchd10-gene-protein-is-implicated-in-addiction-da0c8ecb5921.md

# addictive-genes-claims-first100-074-da0c8ecb5921

**Original claim:** CHCHD10 gene/protein is implicated in addiction.

## Claim

**“CHCHD10 gene/protein is implicated in addiction.”**

## Verdict

Verdict: No Evidence

By the usual biomedical standard for saying a gene is “implicated in addiction” — i.e., replicated human genetic association, gene-level prioritization in addiction GWAS, addiction-focused transcriptomic/functional studies, or animal experiments directly linking the gene to drug reward, dependence, withdrawal, relapse, or substance-use phenotypes — I did **not** find credible direct evidence for **CHCHD10**. Instead, CHCHD10 is primarily established as a **mitochondrial/neurodegeneration-related gene**, with recognized links to disorders such as mitochondrial myopathy, ALS/FTD-spectrum disease, spinal motor neuronopathy, and related neuropathies. ([ncbi.nlm.nih.gov](https://www.ncbi.nlm.nih.gov/books/NBK304142/)) [S1]

## Confidence

**Medium**

Confidence is **medium** rather than high because absence-of-evidence judgments are inherently limited by what is indexed, reported in main text versus supplements, and how broadly one defines “implicated.” Still, multiple addiction-focused resources and large-scale genomic studies reviewed here do **not** identify CHCHD10 as an addiction gene, while CHCHD10’s documented literature is centered elsewhere. ([academic.oup.com](https://academic.oup.com/database/article/doi/10.1093/database/bay121/5161354)) [S2]

## Supporting evidence

A strong negative indicator is that **GeneReviews** and **NCBI Gene** describe CHCHD10 as a mitochondrial protein involved in cristae/intermembrane-space biology and link pathogenic variants to **mitochondrial myopathy, amyotrophic lateral sclerosis, frontotemporal dementia, late-onset spinal motor neuronopathy, and axonal Charcot-Marie-Tooth neuropathy** — not addiction or substance use disorders. ([ncbi.nlm.nih.gov](https://www.ncbi.nlm.nih.gov/books/NBK304142/)) [S3]

A curated addiction genetics resource also weighs against the claim. **ANCO-GeneDB**, a database built from genetic/genomic evidence on **alcohol, nicotine, cocaine, and opioid dependence**, reports **5,633 unique ANCO-associated genes** from its collection/analyses; a direct search within the article text for **CHCHD10** returns **no match**. That does not prove impossibility, but it argues against CHCHD10 being an established addiction gene in the curated literature covered by that resource. ([academic.oup.com](https://academic.oup.com/database/article/doi/10.1093/database/bay121/5161354)) [S4]

Recent large-scale substance-use-disorder genomics point the same way. In a **multivariate GWAS/meta-analysis of over 1 million subjects**, the authors reported a shared addiction-risk factor and highlighted other genes/pathways — for example **DRD2**, **PDE4B**, and substance-specific signals such as **ADH1B**, **CHRNA5**, and **OPRM1-related biology** — while **CHCHD10 does not appear in the accessible main text**. ([pmc.ncbi.nlm.nih.gov](https://pmc.ncbi.nlm.nih.gov/articles/PMC10217792/)) [S5]

Likewise, a **2025 cross-population meta-analysis of cross-substance use disorders** identified **220 shared loci**, **785 prioritized genes**, and strong neuronal/brain-region enrichment, yet **CHCHD10 does not appear in the accessible main text** there either. Again, absence from main text is not absolute proof, but it is inconsistent with the claim that CHCHD10 is currently recognized as an addiction-implicated gene. ([nature.com](https://www.nature.com/articles/s41380-025-03294-5)) [S6]

Overall, the evidence reviewed places CHCHD10 in the domain of **mitochondrial stress, oxidative phosphorylation, synaptic dysfunction in neurodegeneration, and motor-neuron/neuromuscular disease**, not addiction genetics or addiction neurobiology specifically. ([ncbi.nlm.nih.gov](https://www.ncbi.nlm.nih.gov/books/NBK304142/)) [S7]

## Contradicting or limiting evidence

There **is** indirect biological overlap that could make CHCHD10 *plausibly relevant* to addiction biology in a broad sense. CHCHD10 regulates mitochondrial respiration/cellular stress responses, and addiction is widely understood to involve durable synaptic plasticity in reward circuitry. A review of addiction neurobiology describes addiction as partly driven by drug-induced hijacking of **synaptic plasticity mechanisms** in mesolimbic dopamine circuits. Separately, CHCHD10-mutant mouse work shows altered **hippocampal synaptic plasticity**, and CHCHD10 loss impairs a peripheral synapse (the neuromuscular junction) through ATP-dependent effects on acetylcholine-receptor clustering. ([pubmed.ncbi.nlm.nih.gov](https://pubmed.ncbi.nlm.nih.gov/17948030/)) [S8]

But this is still **indirect**. Synaptic plasticity defects in CHCHD10 models were studied mainly in **neurodegeneration** contexts, not in drug reward, dependence, cue reactivity, withdrawal, or relapse models. Likewise, neuromuscular-junction findings concern **peripheral cholinergic synapses**, not the central reward circuitry typically used to infer addiction mechanisms. ([pubmed.ncbi.nlm.nih.gov](https://pubmed.ncbi.nlm.nih.gov/35787294/)) [S9]

Another limitation is definitional: if “implicated” is used very loosely to mean “a gene affecting mitochondrial and synaptic homeostasis that could conceivably modulate addiction-relevant neuronal states,” then CHCHD10 has some theoretical relevance. But that is **not the same** as being supported as an addiction gene by addiction-specific evidence. ([pubmed.ncbi.nlm.nih.gov](https://pubmed.ncbi.nlm.nih.gov/29540477/)) [S10]

## Mechanistic / biological rationale

CHCHD10 encodes a **mitochondrial intermembrane-space/cristae-junction protein**. Authoritative gene resources describe it as enriched at mitochondrial cristae junctions and potentially involved in **cristae morphology maintenance** and **oxidative phosphorylation**. Experimental work further supports roles in **cellular stress responses**, oxygen-sensitive regulation, and mitochondrial respiration. ([ncbi.nlm.nih.gov](https://www.ncbi.nlm.nih.gov/gene/400916)) [S11]

That biology is relevant to neurons in general. CHCHD10 is expressed in brain tissue, including cerebral cortex, and mutant CHCHD10 models show abnormalities in synaptic markers and long-term potentiation-like physiology. Since addiction involves long-lasting adaptations in neural circuits, especially reward-related synaptic plasticity, mitochondrial homeostasis genes are biologically plausible modifiers of addiction vulnerability. ([proteinatlas.org](https://www.proteinatlas.org/ENSG00000250479-CHCHD10/brain/cerebral%2Bcortex)) [S12]

However, plausibility is not evidence of implication. Many mitochondrial or synaptic genes could, in theory, influence addiction-related neuroadaptation, but the field generally reserves “implicated in addiction” for genes with **direct addiction-linked evidence** — e.g., replicated association, expression change in addiction models/human tissues, or manipulation altering addictive behavior. For CHCHD10, I found evidence for **neurodegenerative and mitochondrial disease mechanisms**, not that kind of addiction-specific support. ([ncbi.nlm.nih.gov](https://www.ncbi.nlm.nih.gov/books/NBK304142/)) [S13]

## Bottom line

At present, **CHCHD10 should not be described as an addiction-implicated gene/protein based on available scholarly evidence**. The literature supports CHCHD10 as a **mitochondrial and neurodegeneration-related gene** with possible indirect relevance to neuronal homeostasis and synaptic plasticity, but I found **no convincing direct evidence** tying it to addiction phenotypes specifically. A more accurate statement would be: **CHCHD10 is implicated in mitochondrial and neurodegenerative biology, with only speculative indirect relevance to addiction.** ([ncbi.nlm.nih.gov](https://www.ncbi.nlm.nih.gov/books/NBK304142/)) [S14]

## Cited sources

- **[S1]** CHCHD10-Related Disorders - GeneReviews® - NCBI Bookshelf — https://www.ncbi.nlm.nih.gov/books/NBK304142/
- **[S2]** ANCO-GeneDB: annotations and comprehensive analysis of candidate genes for alcohol, nicotine, cocaine and opioid dependence | Database | Oxford Academic — https://academic.oup.com/database/article/doi/10.1093/database/bay121/5161354
- **[S3]** CHCHD10-Related Disorders - GeneReviews® - NCBI Bookshelf — https://www.ncbi.nlm.nih.gov/books/NBK304142/
- **[S4]** ANCO-GeneDB: annotations and comprehensive analysis of candidate genes for alcohol, nicotine, cocaine and opioid dependence | Database | Oxford Academic — https://academic.oup.com/database/article/doi/10.1093/database/bay121/5161354
- **[S5]** 
            Multivariate genome-wide association meta-analysis of over 1 million subjects identifies loci underlying multiple substance use disorders - PMC
         — https://pmc.ncbi.nlm.nih.gov/articles/PMC10217792/
- **[S6]** Genome-wide meta-analyses of cross substance use disorders in diverse populations | Molecular Psychiatry — https://www.nature.com/articles/s41380-025-03294-5
- **[S7]** CHCHD10-Related Disorders - GeneReviews® - NCBI Bookshelf — https://www.ncbi.nlm.nih.gov/books/NBK304142/
- **[S8]** Synaptic plasticity and addiction - PubMed — https://pubmed.ncbi.nlm.nih.gov/17948030/
- **[S9]** Modulation of synaptic plasticity, motor unit physiology, and TDP-43 pathology by CHCHD10 - PubMed — https://pubmed.ncbi.nlm.nih.gov/35787294/
- **[S10]** The cellular stress proteins CHCHD10 and MNRR1 (CHCHD2): Partners in mitochondrial and nuclear function and dysfunction - PubMed — https://pubmed.ncbi.nlm.nih.gov/29540477/
- **[S11]** CHCHD10 coiled-coil-helix-coiled-coil-helix domain containing 10 [Homo sapiens (human)] - Gene - NCBI — https://www.ncbi.nlm.nih.gov/gene/400916
- **[S12]** Brain tissue expression of CHCHD10 in cerebral cortex - The Human Protein Atlas — https://www.proteinatlas.org/ENSG00000250479-CHCHD10/brain/cerebral%2Bcortex
- **[S13]** CHCHD10-Related Disorders - GeneReviews® - NCBI Bookshelf — https://www.ncbi.nlm.nih.gov/books/NBK304142/
- **[S14]** CHCHD10-Related Disorders - GeneReviews® - NCBI Bookshelf — https://www.ncbi.nlm.nih.gov/books/NBK304142/
