## Supplementary material for "Understanding Substance Dependence: What Differentiates Addictive from Non-Addictive Drugs?": Folder for supplementary files: addictive-genes-claims-first100-075-slc48a1-gene-protein-is-implicated-in-addiction-36b0eccd20d5.md

# addictive-genes-claims-first100-075-36b0eccd20d5

**Original claim:** SLC48A1 gene/protein is implicated in addiction.

## Claim

SLC48A1 gene/protein is implicated in addiction.

## Verdict

Verdict: No Evidence

I did not find direct evidence that **SLC48A1** (also called **HRG1**) is implicated in addiction or substance use disorders. The literature and major gene resources I found instead describe SLC48A1 primarily as a **heme transporter** involved in **lysosomal/endosomal heme handling**, **systemic iron recycling**, and, more recently, **oligodendrocyte/myelin biology** in the central nervous system. ([pmc.ncbi.nlm.nih.gov](https://pmc.ncbi.nlm.nih.gov/articles/PMC8525403/)) [S1]

## Confidence

Medium

## Supporting evidence

SLC48A1’s best-supported function is **heme transport**, not addiction biology. NCBI Gene annotates the human gene with heme binding, heme transmembrane transporter activity, and heme transport, and experimental papers show that SLC48A1/HRG1 transports heme from lysosomes/phagolysosomes to the cytoplasm. In SLC48A1-deficient mice, heme accumulates in reticuloendothelial macrophages as hemozoin, supporting a role in mammalian heme-iron recycling. ([ncbi.nlm.nih.gov](https://www.ncbi.nlm.nih.gov/gene/55652)) [S2]

There is also a plausible **CNS-relevant** role for SLC48A1, but it is still not addiction-specific. A 2025 Glia study reported that **Hrg1/Slc48a1 is enriched in mature myelinating oligodendrocytes** in mouse and human datasets, localizes within myelin sheaths, and that knockout mice show **reduced myelin iron**, lower **MAG** expression, and ultrastructural myelin defects. The Human Protein Atlas likewise places SLC48A1 in oligodendrocyte-associated brain expression clusters. ([pmc.ncbi.nlm.nih.gov](https://pmc.ncbi.nlm.nih.gov/articles/PMC11662986/)) [S3]

Those observations matter because addiction neuroscience has increasingly implicated **myelin plasticity** and **iron homeostasis** more broadly. For example, a 2024 Nature study found that **myelin plasticity in the ventral tegmental area is required for opioid reward**, and human imaging studies in cocaine use disorder reported **abnormal brain/peripheral iron regulation** and **elevated pallidal iron**. These findings show that biological domains connected to SLC48A1—myelin and iron—can be relevant to addiction, even though SLC48A1 itself has not been directly tied to addiction in the studies I found. ([nature.com](https://www.nature.com/articles/s41586-024-07525-7)) [S4]

## Contradicting or limiting evidence

The main limitation is simple: **the direct evidence is missing**. The indexed SLC48A1 literature retrieved here centers on heme transport, iron recycling, and oligodendrocyte/myelin function rather than addiction phenotypes, drug reward, self-administration, withdrawal, craving, or relapse. Even the most CNS-focused SLC48A1 paper is about myelin integrity, not addictive behavior. ([pmc.ncbi.nlm.nih.gov](https://pmc.ncbi.nlm.nih.gov/articles/PMC8525403/)) [S5]

NCBI Gene’s phenotype/GWAS section for human **SLC48A1** lists an association with **serum urate concentrations**; I did not find addiction-related phenotypes there. That does not prove no association exists anywhere, but it is notable that a major curated resource does **not** flag addiction as a phenotype for this gene. ([ncbi.nlm.nih.gov](https://www.ncbi.nlm.nih.gov/gene/55652)) [S6]

The 2025 oligodendrocyte/myelin paper is important for brain biology, but it should not be overinterpreted. Its reported phenotypes are **myelin iron loss, reduced MAG, and myelin structural abnormalities**; it does not report addiction-relevant behavioral assays. So, at present, it supports a neural homeostasis/myelin role, not an addiction role. ([pmc.ncbi.nlm.nih.gov](https://pmc.ncbi.nlm.nih.gov/articles/PMC11662986/)) [S7]

## Mechanistic / biological rationale

A **hypothesis-generating** rationale does exist. SLC48A1 controls intracellular heme availability and contributes to iron liberation from heme after lysosomal export. Because iron is important for oligodendrocyte maturation and myelin maintenance, and because SLC48A1 is enriched in mature oligodendrocytes/myelin, altered SLC48A1 function could in principle affect white-matter or circuit plasticity. ([pmc.ncbi.nlm.nih.gov](https://pmc.ncbi.nlm.nih.gov/articles/PMC8525403/)) [S8]

That matters because addiction-related circuit remodeling is not purely synaptic. The opioid-reward paper argues that **adaptive myelination** can shape reward circuitry, and cocaine studies suggest **iron dysregulation** may accompany addictive illness. By inference, a gene that influences heme/iron handling in oligodendrocytes could someday prove relevant to addiction vulnerability or consequences. But that is still an **indirect mechanistic inference**, not a demonstrated gene-addiction link. ([nature.com](https://www.nature.com/articles/s41586-024-07525-7)) [S9]

A second speculative route is via **endosomal pH and trafficking**. Earlier work showed HRG1/SLC48A1 can associate with the **V-ATPase** and influence endosomal acidification and receptor trafficking. Since vesicular trafficking and receptor recycling are important in neural plasticity, one could imagine downstream relevance to addiction biology. However, I found no study showing that this SLC48A1-dependent trafficking mechanism has been connected specifically to addiction in humans or animal models. ([pubmed.ncbi.nlm.nih.gov](https://pubmed.ncbi.nlm.nih.gov/19875448/?utm_source=openai)) [S10]

## Bottom line

Based on the available scholarly and authoritative sources I found, the claim that **“SLC48A1 gene/protein is implicated in addiction”** is **not currently supported by direct evidence**. SLC48A1 is well supported as a **heme transporter** with roles in **iron recycling** and emerging roles in **oligodendrocyte/myelin integrity**, and those broader biological systems are relevant to addiction research. But that is not the same as showing that **SLC48A1 itself** is implicated in addiction. Right now, the strongest fair summary is: **biologically plausible as an indirect candidate, but not evidenced as an addiction gene/protein.** ([pmc.ncbi.nlm.nih.gov](https://pmc.ncbi.nlm.nih.gov/articles/PMC8525403/)) [S11]

## Cited sources

- **[S1]** 
            Normal Iron Homeostasis Requires the Transporter SLC48A1 for Efficient Heme-Iron Recycling in Mammals - PMC
         — https://pmc.ncbi.nlm.nih.gov/articles/PMC8525403/
- **[S2]** SLC48A1 solute carrier family 48 member 1 [Homo sapiens (human)] - Gene - NCBI — https://www.ncbi.nlm.nih.gov/gene/55652
- **[S3]** 
            Oligodendrocyte Slc48a1 (Hrg1) encodes a functional heme transporter required for myelin integrity - PMC
         — https://pmc.ncbi.nlm.nih.gov/articles/PMC11662986/
- **[S4]** Myelin plasticity in the ventral tegmental area is required for opioid reward | Nature — https://www.nature.com/articles/s41586-024-07525-7
- **[S5]** 
            Normal Iron Homeostasis Requires the Transporter SLC48A1 for Efficient Heme-Iron Recycling in Mammals - PMC
         — https://pmc.ncbi.nlm.nih.gov/articles/PMC8525403/
- **[S6]** SLC48A1 solute carrier family 48 member 1 [Homo sapiens (human)] - Gene - NCBI — https://www.ncbi.nlm.nih.gov/gene/55652
- **[S7]** 
            Oligodendrocyte Slc48a1 (Hrg1) encodes a functional heme transporter required for myelin integrity - PMC
         — https://pmc.ncbi.nlm.nih.gov/articles/PMC11662986/
- **[S8]** 
            Normal Iron Homeostasis Requires the Transporter SLC48A1 for Efficient Heme-Iron Recycling in Mammals - PMC
         — https://pmc.ncbi.nlm.nih.gov/articles/PMC8525403/
- **[S9]** Myelin plasticity in the ventral tegmental area is required for opioid reward | Nature — https://www.nature.com/articles/s41586-024-07525-7
- **[S10]** Heme-binding protein HRG-1 is induced by insulin-like growth factor I and associates with the vacuolar H+-ATPase to control endosomal pH and receptor trafficking - PubMed — https://pubmed.ncbi.nlm.nih.gov/19875448/?utm_source=openai
- **[S11]** 
            Normal Iron Homeostasis Requires the Transporter SLC48A1 for Efficient Heme-Iron Recycling in Mammals - PMC
         — https://pmc.ncbi.nlm.nih.gov/articles/PMC8525403/
