## Supplementary material for "Understanding Substance Dependence: What Differentiates Addictive from Non-Addictive Drugs?": Folder for supplementary files: addictive-genes-claims-first100-076-dync1i1-gene-protein-is-implicated-in-addiction-0530dd54dfda.md

# addictive-genes-claims-first100-076-0530dd54dfda

**Original claim:** DYNC1I1 gene/protein is implicated in addiction.

## Claim

DYNC1I1 gene/protein is implicated in addiction.

## Verdict

Verdict: Mixed Evidence

## Confidence

Medium

## Supporting evidence

DYNC1I1 encodes cytoplasmic dynein 1 intermediate chain 1, a neuronal dynein subunit involved in intracellular transport. Independent sources indicate that the neuronal IC-1 isoform is specifically expressed from the endogenous Dync1i1 locus in mouse neurons, and dynein/DYNC1I1-related complexes help stabilize microtubules and maintain synaptic density in cortical neurons. Those are biologically relevant processes for addiction-related plasticity. ([pubmed.ncbi.nlm.nih.gov](https://pubmed.ncbi.nlm.nih.gov/23475693/?utm_source=openai)) [S1]

There is also some drug-exposure-linked molecular evidence. A conference abstract in rats reported that acute ethanol exposure altered m6A methylation of Dync1i1 transcripts in amygdala, and that MeRIP-qPCR changes correlated with Dync1i1 mRNA expression. The authors framed this as a possible way alcohol exposure could “prime” the amygdala for alcohol use disorder, but this was an acute-exposure, preclinical, abstract-level result rather than a full addiction-behavior study. ([pmc.ncbi.nlm.nih.gov](https://pmc.ncbi.nlm.nih.gov/articles/PMC10729598/)) [S2]

A separate secondary annotation, linked to a human postmortem cocaine-abuse transcriptional profiling study, lists DYNC1I1 mRNA as downregulated after cocaine exposure/abuse. However, in the paper text accessible through web search, I could verify the overall cocaine-related transcriptional profiling study but not independently confirm the DYNC1I1-specific line item from the primary article text itself. That makes this supportive signal suggestive, but weak. ([go.drugbank.com](https://go.drugbank.com/articles/A21701)) [S3]

Finally, systems-genomics work on substance-use traits shows that addiction risk is enriched in neuronal chromatin architecture and reward-circuit cell types, especially dopaminergic, GABAergic, serotonergic neurons, and nucleus accumbens medium spiny neurons. This does not prove a role for DYNC1I1 specifically, but it supports the plausibility that genes involved in neuronal transport/synaptic maintenance could matter in addiction biology. ([pubmed.ncbi.nlm.nih.gov](https://pubmed.ncbi.nlm.nih.gov/35422469/)) [S4]

## Contradicting or limiting evidence

The biggest limitation is that I did **not** find strong, direct, replicated evidence that DYNC1I1 is an established addiction gene. In the sources reviewed, there was no clear peer-reviewed human study showing a robust, replicated association of DYNC1I1 variants with substance use disorder diagnosis, no Mendelian evidence tying DYNC1I1 dysfunction to addictive phenotypes, and no animal paper directly demonstrating that Dync1i1 manipulation changes alcohol, nicotine, cocaine, or opioid self-administration, relapse, or sensitization. That absence of direct evidence matters. ([pubmed.ncbi.nlm.nih.gov](https://pubmed.ncbi.nlm.nih.gov/25555482/)) [S5]

Large human genetic studies of nicotine dependence and related tobacco/alcohol traits have highlighted other loci—such as CHRNA5-CHRNA3-CHRNB4 and DNMT3B for nicotine dependence—rather than DYNC1I1 as a prominent, established signal in the abstracts and summaries reviewed. Likewise, the chromatin-architecture paper on smoking/alcohol traits discusses many mapped risk genes and pathways, but the accessible main text does not single out DYNC1I1 as a validated core addiction gene. ([pubmed.ncbi.nlm.nih.gov](https://pubmed.ncbi.nlm.nih.gov/25555482/)) [S6]

The ethanol-related Dync1i1 result is also preliminary: it comes from a conference poster abstract, concerns acute ethanol exposure in rat amygdala, and does not demonstrate addiction-like behavior or causality. Similarly, the cocaine-related DYNC1I1 signal is presently best treated as a database-level annotation unless independently confirmed from the original primary dataset/report. ([pmc.ncbi.nlm.nih.gov](https://pmc.ncbi.nlm.nih.gov/articles/PMC10729598/)) [S7]

## Mechanistic / biological rationale

Mechanistically, the claim is plausible. Addiction depends on long-lasting synaptic and circuit plasticity in reward-related regions, and dynein-mediated intracellular transport is central to neuronal growth, dendritic differentiation, spine development, microtubule organization, and synaptic maintenance. Dync1i1 knockout mice show impaired parvalbumin interneuron activity and gamma oscillations, and dynein-NCAM interactions are required for normal synaptic density in cortical neurons. Those functions are relevant to neural homeostasis and could, in principle, influence vulnerability to substance-induced plasticity. ([pubmed.ncbi.nlm.nih.gov](https://pubmed.ncbi.nlm.nih.gov/37257662/?utm_source=openai)) [S8]

That said, biological plausibility is not the same as implication in addiction. Many neuronal housekeeping or plasticity-related genes could theoretically modulate addiction circuits, but a gene is usually considered “implicated” only when there is direct evidence from human genetics, reproducible transcriptomic/proteomic data, or causal perturbation studies in addiction models. For DYNC1I1, the rationale is stronger than the direct addiction evidence. ([pubmed.ncbi.nlm.nih.gov](https://pubmed.ncbi.nlm.nih.gov/23960070/)) [S9]

## Bottom line

DYNC1I1 has a **credible neurobiological connection** to processes that matter for addiction—neuronal transport, synaptic stability, interneuron function, and drug-responsive molecular regulation—but the current evidence is **too sparse and indirect** to call it an established addiction gene/protein. The best characterization is: **preliminary/indirect implication, not strong confirmation**. ([pubmed.ncbi.nlm.nih.gov](https://pubmed.ncbi.nlm.nih.gov/23960070/)) [S10]

If you want, I can next do a **deeper paper-by-paper review** focused on one substance class only—alcohol, nicotine, cocaine, or opioids—to see whether DYNC1I1 has stronger evidence in a narrower addiction context.

## Cited sources

- **[S1]** Establishing a novel knock-in mouse line for studying neuronal cytoplasmic dynein under normal and pathologic conditions - PubMed — https://pubmed.ncbi.nlm.nih.gov/23475693/?utm_source=openai
- **[S2]** https://pmc.ncbi.nlm.nih.gov/articles/PMC10729598/ — https://pmc.ncbi.nlm.nih.gov/articles/PMC10729598/
- **[S3]** https://go.drugbank.com/articles/A21701 — https://go.drugbank.com/articles/A21701
- **[S4]** https://pubmed.ncbi.nlm.nih.gov/35422469/ — https://pubmed.ncbi.nlm.nih.gov/35422469/
- **[S5]** https://pubmed.ncbi.nlm.nih.gov/25555482/ — https://pubmed.ncbi.nlm.nih.gov/25555482/
- **[S6]** https://pubmed.ncbi.nlm.nih.gov/25555482/ — https://pubmed.ncbi.nlm.nih.gov/25555482/
- **[S7]** https://pmc.ncbi.nlm.nih.gov/articles/PMC10729598/ — https://pmc.ncbi.nlm.nih.gov/articles/PMC10729598/
- **[S8]** Pain sensitivity related to gamma oscillation of parvalbumin interneuron in primary somatosensory cortex in Dync1i1-/- mice - PubMed — https://pubmed.ncbi.nlm.nih.gov/37257662/?utm_source=openai
- **[S9]** Dynein interacts with the neural cell adhesion molecule (NCAM180) to tether dynamic microtubules and maintain synaptic density in cortical neurons - PubMed — https://pubmed.ncbi.nlm.nih.gov/23960070/
- **[S10]** Dynein interacts with the neural cell adhesion molecule (NCAM180) to tether dynamic microtubules and maintain synaptic density in cortical neurons - PubMed — https://pubmed.ncbi.nlm.nih.gov/23960070/
