## Supplementary material for "Understanding Substance Dependence: What Differentiates Addictive from Non-Addictive Drugs?": Folder for supplementary files: addictive-genes-claims-first100-077-pafah1b1-gene-protein-is-implicated-in-addiction-eea2278335ee.md

# addictive-genes-claims-first100-077-eea2278335ee

**Original claim:** PAFAH1B1 gene/protein is implicated in addiction.

## Claim

PAFAH1B1 gene/protein is implicated in addiction.

## Verdict

Verdict: Mixed Evidence

## Confidence

Medium

## Supporting evidence

There **is** some evidence linking **PAFAH1B1 (LIS1/Pafah1b1)** to **alcohol-related phenotypes**, but it is mostly **preclinical** and not yet strong enough to establish it as a well-supported human addiction gene. The strongest direct paper I found is a cross-species integrative genomics study that prioritized **Pafah1b1** across 32 alcohol-related datasets and then tested a heterozygous knockout mouse. In that study, **Pafah1b1-mutant mice showed altered acute alcohol responses**, including reduced hypothermic response at higher ethanol doses, altered alcohol-related anxiety-like behavior, and a **trend** toward higher ethanol intake in the two-bottle choice assay. The authors explicitly described Pafah1b1 as a **novel alcohol-related candidate** emerging from convergent genomics and then “validated” it in mice. ([pmc.ncbi.nlm.nih.gov](https://pmc.ncbi.nlm.nih.gov/articles/PMC4720795/)) [S1]

Additional animal/omics studies are directionally consistent with an alcohol link. In a mouse nucleus accumbens transcriptomics study after repeated binge ethanol drinking, **Pafah1b1** appeared among the **top 30 significantly regulated genes in females**. ([frontiersin.org](https://www.frontiersin.org/journals/genetics/articles/10.3389/fgene.2018.00325/pdf)) [S2] In an older rat study of **ethanol self-administration**, **Pafah1b1** expression in nucleus accumbens was **downregulated** relative to controls. ([pmc.ncbi.nlm.nih.gov](https://pmc.ncbi.nlm.nih.gov/articles/PMC4516281/)) [S3] In a mouse amygdala proteomics study after moderate alcohol drinking, **PAFAH1B1 protein was decreased** within alcohol-sensitive protein networks related to cellular development, microtubule dynamics, and synaptic plasticity. ([pmc.ncbi.nlm.nih.gov](https://pmc.ncbi.nlm.nih.gov/articles/PMC4417085/)) [S4]

Taken together, those findings make it reasonable to say PAFAH1B1 is **a plausible preclinical alcohol-response candidate**. However, that is narrower than saying it is firmly implicated in “addiction” generally. Most of the evidence is about **alcohol exposure/response** rather than human substance use disorder itself. ([pmc.ncbi.nlm.nih.gov](https://pmc.ncbi.nlm.nih.gov/articles/PMC4720795/)) [S5]

## Contradicting or limiting evidence

The biggest limitation is that I found **little convincing human genetic or clinical evidence** tying **PAFAH1B1** directly to addiction. In GeneReviews, the established phenotype spectrum for intragenic germline **PAFAH1B1** variants is essentially **lissencephaly / subcortical band heterotopia and related neurodevelopmental manifestations**, and it explicitly states that **no phenotypes other than those discussed there are known** for intragenic pathogenic variants. Addiction is not part of that recognized Mendelian phenotype. ([ncbi.nlm.nih.gov](https://www.ncbi.nlm.nih.gov/books/NBK5189/)) [S6]

Large human addiction GWAS also do **not** provide strong support for PAFAH1B1 as a leading locus. For alcohol dependence, a large trans-ancestral GWAS identified **ADH1B** as the genome-wide significant signal, not PAFAH1B1. ([pubmed.ncbi.nlm.nih.gov](https://pubmed.ncbi.nlm.nih.gov/30482948/)) [S7] For clinically defined nicotine dependence, a large multi-ancestral GWAS replicated the well-known **CHRNA5** signal and modeled broader addiction liability, again without highlighting PAFAH1B1 as a key locus in the abstracted findings. ([pubmed.ncbi.nlm.nih.gov](https://pubmed.ncbi.nlm.nih.gov/39974067/)) [S8] For cocaine dependence/use disorder, an integrative human genomic/transcriptomic analysis identified other candidates such as **NDUFB9** and **C1QL2** in associated networks rather than PAFAH1B1. ([pubmed.ncbi.nlm.nih.gov](https://pubmed.ncbi.nlm.nih.gov/32457073/)) [S9]

Even the main supportive mouse paper is not definitive. Its drinking result was a **trend** rather than a conventional statistically significant increase in alcohol intake, and the paper itself noted that **Pafah1b1 had not previously been associated with alcohol-use disorder** in annotated human or mouse resources before their integrative analysis. ([pmc.ncbi.nlm.nih.gov](https://pmc.ncbi.nlm.nih.gov/articles/PMC4720795/)) [S10] Also, one cited proteomic study examined **moderate, nondependent** alcohol drinking and specifically reported **absence of dependence**, so those findings are better interpreted as early alcohol-response biology than addiction per se. ([pmc.ncbi.nlm.nih.gov](https://pmc.ncbi.nlm.nih.gov/articles/PMC4417085/)) [S11]

So the limiting picture is: **preclinical alcohol-related signals exist**, but **human addiction genetics/clinical evidence is sparse** and the current evidence base does not put PAFAH1B1 among established addiction genes. ([ncbi.nlm.nih.gov](https://www.ncbi.nlm.nih.gov/books/NBK5189/)) [S12]

## Mechanistic / biological rationale

Biologically, PAFAH1B1 is a **credible neural systems gene**. NCBI and GeneReviews describe it as a highly conserved protein involved in the **PAFAH1B complex**, with major roles in **microtubule organization, dynein-related transport, mitotic spindle orientation, nucleokinesis, and radial/tangential neuronal migration**. ([ncbi.nlm.nih.gov](https://www.ncbi.nlm.nih.gov/gene/5048?utm_source=openai)) [S13]

That matters because addiction vulnerability and drug-induced plasticity depend on intact **reward-circuit development, synaptic organization, excitation/inhibition balance, and plasticity**. Independent experimental work shows that **Pafah1b1 haploinsufficiency disrupts GABAergic interneuron migration**, reduces parvalbumin- and somatostatin-positive interneurons in dentate gyrus, shifts circuits toward **greater excitation and less inhibition**, and produces seizures/cognitive deficits in mice. ([nature.com](https://www.nature.com/articles/s41598-017-08809-x)) [S14] Other work shows LIS1 influences **synaptogenesis, dendritic spine dynamics, synaptic integrity, and synaptic homeostasis** in post-migrational neurons. ([pubmed.ncbi.nlm.nih.gov](https://pubmed.ncbi.nlm.nih.gov/23483716/?utm_source=openai)) [S15]

The alcohol-candidate paper also noted that **Pafah1b1 is co-expressed with GABA-A receptor subunits** previously associated with alcohol-related phenotypes and discussed possible roles in **synaptic integration** and **excitatory neurotransmission**. ([pmc.ncbi.nlm.nih.gov](https://pmc.ncbi.nlm.nih.gov/articles/PMC4720795/)) [S16] That makes mechanistic sense: a gene that shapes inhibitory interneuron development and synaptic plasticity could plausibly modulate **drug sensitivity, reinforcement, or vulnerability**. But mechanistic plausibility is **not the same as proof of addiction involvement**, especially without robust human association or causal clinical data. ([nature.com](https://www.nature.com/articles/s41598-017-08809-x)) [S17]

## Bottom line

If the claim means **“PAFAH1B1 has some experimental evidence suggesting relevance to alcohol-related behaviors or neural processes relevant to addiction”**, that is **fair**. There are several **rodent and omics** signals, plus one notable **mouse functional study**, supporting a possible role. ([pmc.ncbi.nlm.nih.gov](https://pmc.ncbi.nlm.nih.gov/articles/PMC4720795/)) [S18]

If the claim means **“PAFAH1B1 is an established human addiction gene/protein”**, the current evidence does **not** support that. Human clinical phenotype resources and major addiction GWAS do not presently make PAFAH1B1 a well-established addiction locus. ([ncbi.nlm.nih.gov](https://www.ncbi.nlm.nih.gov/books/NBK5189/)) [S19]

So overall: **PAFAH1B1 is a plausible, mostly alcohol-related preclinical candidate, but not a strongly established addiction gene in humans at this time.** ([pmc.ncbi.nlm.nih.gov](https://pmc.ncbi.nlm.nih.gov/articles/PMC4720795/)) [S20]

## Cited sources

- **[S1]** 
            Cross-Species Integrative Functional Genomics in GeneWeaver Reveals a Role for Pafah1b1 in Altered Response to Alcohol - PMC
         — https://pmc.ncbi.nlm.nih.gov/articles/PMC4720795/
- **[S2]** Binge Ethanol Drinking Produces Sexually Divergent and Distinct Changes in Nucleus Accumbens Signaling Cascades and Pathways in Adult C57BL/6J Mice — https://www.frontiersin.org/journals/genetics/articles/10.3389/fgene.2018.00325/pdf
- **[S3]** 
            Differential gene expression in the nucleus accumbens with ethanol self-administration in inbred alcohol-preferring rats - PMC
         — https://pmc.ncbi.nlm.nih.gov/articles/PMC4516281/
- **[S4]** 
            Moderate Alcohol Drinking and the Amygdala Proteome: Identification and Validation of Calcium/Calmodulin Dependent Kinase II and AMPA Receptor Activity as Novel Molecular Mechanisms of the Positive Reinforcing Effects of Alcohol - PMC
         — https://pmc.ncbi.nlm.nih.gov/articles/PMC4417085/
- **[S5]** 
            Cross-Species Integrative Functional Genomics in GeneWeaver Reveals a Role for Pafah1b1 in Altered Response to Alcohol - PMC
         — https://pmc.ncbi.nlm.nih.gov/articles/PMC4720795/
- **[S6]** PAFAH1B1-Related Lissencephaly / Subcortical Band Heterotopia - GeneReviews® - NCBI Bookshelf — https://www.ncbi.nlm.nih.gov/books/NBK5189/
- **[S7]** Transancestral GWAS of alcohol dependence reveals common genetic underpinnings with psychiatric disorders - PubMed — https://pubmed.ncbi.nlm.nih.gov/30482948/
- **[S8]** Multi-ancestral genome-wide association study of clinically defined nicotine dependence reveals strong genetic correlations with other substance use disorders and health-related traits - PubMed — https://pubmed.ncbi.nlm.nih.gov/39974067/
- **[S9]** Genetic Architecture and Molecular Neuropathology of Human Cocaine Addiction - PubMed — https://pubmed.ncbi.nlm.nih.gov/32457073/
- **[S10]** 
            Cross-Species Integrative Functional Genomics in GeneWeaver Reveals a Role for Pafah1b1 in Altered Response to Alcohol - PMC
         — https://pmc.ncbi.nlm.nih.gov/articles/PMC4720795/
- **[S11]** 
            Moderate Alcohol Drinking and the Amygdala Proteome: Identification and Validation of Calcium/Calmodulin Dependent Kinase II and AMPA Receptor Activity as Novel Molecular Mechanisms of the Positive Reinforcing Effects of Alcohol - PMC
         — https://pmc.ncbi.nlm.nih.gov/articles/PMC4417085/
- **[S12]** PAFAH1B1-Related Lissencephaly / Subcortical Band Heterotopia - GeneReviews® - NCBI Bookshelf — https://www.ncbi.nlm.nih.gov/books/NBK5189/
- **[S13]** PAFAH1B1 platelet activating factor acetylhydrolase 1b regulatory subunit 1 [Homo sapiens (human)] - Gene - NCBI — https://www.ncbi.nlm.nih.gov/gene/5048?utm_source=openai
- **[S14]** PAFAH1B1 haploinsufficiency disrupts GABA neurons and synaptic E/I balance in the dentate gyrus | Scientific Reports — https://www.nature.com/articles/s41598-017-08809-x
- **[S15]** Lis1 controls dynamics of neuronal filopodia and spines to impact synaptogenesis and social behaviour - PubMed — https://pubmed.ncbi.nlm.nih.gov/23483716/?utm_source=openai
- **[S16]** 
            Cross-Species Integrative Functional Genomics in GeneWeaver Reveals a Role for Pafah1b1 in Altered Response to Alcohol - PMC
         — https://pmc.ncbi.nlm.nih.gov/articles/PMC4720795/
- **[S17]** PAFAH1B1 haploinsufficiency disrupts GABA neurons and synaptic E/I balance in the dentate gyrus | Scientific Reports — https://www.nature.com/articles/s41598-017-08809-x
- **[S18]** 
            Cross-Species Integrative Functional Genomics in GeneWeaver Reveals a Role for Pafah1b1 in Altered Response to Alcohol - PMC
         — https://pmc.ncbi.nlm.nih.gov/articles/PMC4720795/
- **[S19]** PAFAH1B1-Related Lissencephaly / Subcortical Band Heterotopia - GeneReviews® - NCBI Bookshelf — https://www.ncbi.nlm.nih.gov/books/NBK5189/
- **[S20]** 
            Cross-Species Integrative Functional Genomics in GeneWeaver Reveals a Role for Pafah1b1 in Altered Response to Alcohol - PMC
         — https://pmc.ncbi.nlm.nih.gov/articles/PMC4720795/
