## Supplementary material for "Understanding Substance Dependence: What Differentiates Addictive from Non-Addictive Drugs?": Folder for supplementary files: addictive-genes-claims-first100-078-slc6a2-gene-protein-is-implicated-in-addiction-a18c07fef48d.md

# addictive-genes-claims-first100-078-a18c07fef48d

**Original claim:** SLC6A2 gene/protein is implicated in addiction.

## Claim

**Claim:** “SLC6A2 gene/protein is implicated in addiction.”

## Verdict

Verdict: Likely True

Broadly, this claim is supported **at the protein/pathway level**: **SLC6A2 encodes the norepinephrine transporter (NET)**, a key regulator of norepinephrine homeostasis and a **direct pharmacologic target of cocaine**. Preclinical work also links NET/noradrenergic signaling to **drug seeking, cue reactivity, and relapse-like behavior**, especially for stimulants and opioids. However, if the claim is interpreted more narrowly to mean that **SLC6A2 is a firmly established inherited addiction-risk gene in humans**, the evidence is much less consistent. ([ncbi.nlm.nih.gov](https://www.ncbi.nlm.nih.gov/gene/6530)) [S1]

## Confidence

**Medium.** Confidence is not high because the strongest evidence is **mechanistic, pharmacologic, and animal-based**, whereas **human genetic association data are mixed** and I did not find convincing evidence that SLC6A2 is among the most robustly replicated genome-wide addiction loci. ([pubmed.ncbi.nlm.nih.gov](https://pubmed.ncbi.nlm.nih.gov/26773688/)) [S2]

## Supporting evidence

SLC6A2 is the human gene for the **sodium-dependent norepinephrine transporter (NET)**, which removes norepinephrine from the synaptic cleft and thereby regulates **norepinephrine homeostasis**. That alone makes it biologically plausible in disorders driven by altered arousal, stress responsiveness, and reward-related signaling. ([ncbi.nlm.nih.gov](https://www.ncbi.nlm.nih.gov/gene/6530)) [S3]

The most direct addiction relevance is that **cocaine interacts with NET**. Classic transporter work showed that cocaine acts as a **competitive inhibitor** at the human norepinephrine transporter, and later work in relapse models explicitly describes NET blockade as a mechanism by which cocaine raises extracellular norepinephrine. NET is also recognized as one of cocaine’s three major high-affinity monoamine transporter targets, alongside DAT and SERT. ([pubmed.ncbi.nlm.nih.gov](https://pubmed.ncbi.nlm.nih.gov/9852563/)) [S4]

A substantial preclinical literature implicates **noradrenergic signaling in relapse-related behaviors**. A review focused on norepinephrine in arousal/motivation concluded that norepinephrine plays an important role in **relapse**, and that noradrenergic signaling can be sufficient to reinstate drug seeking under some conditions. ([pubmed.ncbi.nlm.nih.gov](https://pubmed.ncbi.nlm.nih.gov/26773688/)) [S5]

More specifically, **selective NET inhibition** altered addiction-like behavior in animals. In rats, **atomoxetine**—a selective norepinephrine reuptake inhibitor acting at NET—reduced **cue-controlled cocaine seeking**, **cue-controlled heroin seeking**, and **cue-induced relapse to cocaine seeking** after abstinence. That is fairly direct evidence that the NET/SLC6A2 axis participates in addiction-relevant behavior, particularly relapse and cue-driven motivation. ([pubmed.ncbi.nlm.nih.gov](https://pubmed.ncbi.nlm.nih.gov/21109233/)) [S6]

Additional circuit-level evidence points the same way. In a rat reinstatement model, **α1-adrenergic receptor signaling in medial prefrontal cortex** was required for **cocaine-primed reinstatement**, and the authors’ model explicitly links this to cocaine-induced elevation of norepinephrine via **NET blockade** in the mPFC. This supports a mechanistic chain from cocaine → NET inhibition → increased norepinephrine signaling → relapse-like behavior. ([pubmed.ncbi.nlm.nih.gov](https://pubmed.ncbi.nlm.nih.gov/28392265/)) [S7]

Genetic manipulation studies are also relevant, though nuanced. NET knockout mice show **altered reward circuitry** on imaging-based analyses, consistent with a role for NET in reward-network organization. Meanwhile, older cocaine studies showed that **NET knockout mice can still show some cocaine reward-related behaviors**, implying that NET is **involved but not singularly required**; addiction biology is distributed across multiple systems rather than controlled by one transporter alone. ([pubmed.ncbi.nlm.nih.gov](https://pubmed.ncbi.nlm.nih.gov/23469209/)) [S8]

Human evidence is suggestive but less definitive. A 2023 study found that the **SLC6A2 rs36024 genotype interacted with cocaine use disorder diagnosis** to influence **inhibitory control**, with the CC genotype showing worse stop-signal performance in people with cocaine use disorder. That does not prove causation, but it supports the idea that SLC6A2 variation may modulate an addiction-relevant endophenotype. ([pubmed.ncbi.nlm.nih.gov](https://pubmed.ncbi.nlm.nih.gov/36774338/)) [S9]

There are also candidate-gene signals in some substance-use phenotypes. One study reported associations between **SLC6A2 SNPs and alcoholism**, and another found a heroin-dependence study with largely negative single-marker analyses but a significant haplotype difference. Separate work linked SLC6A2 variants to **acute subjective response to D-amphetamine**, a phenotype plausibly related to abuse liability. Taken together, these studies support “possible involvement,” but not a settled human genetic effect. ([pubmed.ncbi.nlm.nih.gov](https://pubmed.ncbi.nlm.nih.gov/21070505/)) [S10]

## Contradicting or limiting evidence

The strongest limitation is that **human genetic evidence is inconsistent**. A study of alcohol dependence found that two functional SLC6A2 polymorphisms were **not major risk factors** for alcohol dependence or its clinical subtypes. The heroin-dependence study likewise reported **no significant single-marker or genotype effects** after logistic regression, despite one haplotype signal. ([pubmed.ncbi.nlm.nih.gov](https://pubmed.ncbi.nlm.nih.gov/17630229/)) [S11]

Some apparently supportive candidate-gene findings also failed replication in broader psychostimulant-response work. A replication-focused study of candidate genes for acute amphetamine response reported that previous findings, including those involving **SLC6A2**, were **not replicated** in an additional sample, underscoring the fragility of small candidate-gene associations. ([pubmed.ncbi.nlm.nih.gov](https://pubmed.ncbi.nlm.nih.gov/23303064/)) [S12]

Clinical translation is also limited. Although atomoxetine reduced cocaine-related attentional bias in one human laboratory study, a randomized 12-week outpatient trial found that atomoxetine **did not reduce cocaine use** relative to placebo in treatment-seeking cocaine-dependent individuals. So NET modulation clearly affects some addiction-relevant processes, but it has **not translated into a clearly effective anti-addiction therapy**. ([pubmed.ncbi.nlm.nih.gov](https://pubmed.ncbi.nlm.nih.gov/28551713/)) [S13]

At the broader genomics level, modern large-scale substance-use GWASs identify many loci for alcohol, tobacco, cannabis, opioid, and cross-substance phenotypes, but I did **not** find SLC6A2 presented as one of the most reproducible genome-wide significant addiction loci in the large studies/reviews I examined. That is an inference from what these large studies highlight, and it argues against overstating SLC6A2 as a major established addiction gene. ([pubmed.ncbi.nlm.nih.gov](https://pubmed.ncbi.nlm.nih.gov/37250466/)) [S14]

## Mechanistic / biological rationale

Biologically, the rationale is strong. NET/SLC6A2 regulates the duration and magnitude of **norepinephrine signaling**, which shapes **arousal, stress responses, attention, impulse control, and prefrontal cortical function**—all processes relevant to addiction vulnerability and relapse. ([ncbi.nlm.nih.gov](https://www.ncbi.nlm.nih.gov/gene/6530)) [S15]

This is particularly relevant for stimulants. Cocaine blocks NET, increasing extracellular norepinephrine, and preclinical work suggests that this can amplify **prefrontal output** and facilitate **reinstatement of drug seeking** through α1-adrenergic signaling. Thus, even if dopamine remains central to reward, NET can influence the **motivational, cue-reactive, and relapse-related** components of addiction. ([pubmed.ncbi.nlm.nih.gov](https://pubmed.ncbi.nlm.nih.gov/28392265/)) [S16]

There is also evidence of drug-induced plasticity involving NET itself. In postmortem human cortex from cocaine users, chronic cocaine exposure was associated with **upregulated NET protein and binding sites**, and mechanistic cell work has shown **cocaine-induced NET up-regulation** via p38 MAPK-dependent phosphorylation. These findings fit the idea that SLC6A2/NET is not just a passive target, but part of the adaptive biology of chronic cocaine exposure. ([pubmed.ncbi.nlm.nih.gov](https://pubmed.ncbi.nlm.nih.gov/15763139/)) [S17]

## Bottom line

**Yes—SLC6A2/NET is reasonably considered implicated in addiction**, especially in the biology of **psychostimulant action, cue-driven drug seeking, and relapse-related noradrenergic signaling**. But the implication is strongest at the **protein/pathway and animal-mechanistic level**, not as a **firmly established major human addiction-risk gene**. So the fairest summary is: **likely true in a broad biological sense, but only modestly supported as a direct, reproducibly inherited human addiction gene.** ([ncbi.nlm.nih.gov](https://www.ncbi.nlm.nih.gov/gene/6530)) [S18]

## Cited sources

- **[S1]** https://www.ncbi.nlm.nih.gov/gene/6530 — https://www.ncbi.nlm.nih.gov/gene/6530
- **[S2]** https://pubmed.ncbi.nlm.nih.gov/26773688/ — https://pubmed.ncbi.nlm.nih.gov/26773688/
- **[S3]** https://www.ncbi.nlm.nih.gov/gene/6530 — https://www.ncbi.nlm.nih.gov/gene/6530
- **[S4]** https://pubmed.ncbi.nlm.nih.gov/9852563/ — https://pubmed.ncbi.nlm.nih.gov/9852563/
- **[S5]** https://pubmed.ncbi.nlm.nih.gov/26773688/ — https://pubmed.ncbi.nlm.nih.gov/26773688/
- **[S6]** https://pubmed.ncbi.nlm.nih.gov/21109233/ — https://pubmed.ncbi.nlm.nih.gov/21109233/
- **[S7]** https://pubmed.ncbi.nlm.nih.gov/28392265/ — https://pubmed.ncbi.nlm.nih.gov/28392265/
- **[S8]** https://pubmed.ncbi.nlm.nih.gov/23469209/ — https://pubmed.ncbi.nlm.nih.gov/23469209/
- **[S9]** https://pubmed.ncbi.nlm.nih.gov/36774338/ — https://pubmed.ncbi.nlm.nih.gov/36774338/
- **[S10]** https://pubmed.ncbi.nlm.nih.gov/21070505/ — https://pubmed.ncbi.nlm.nih.gov/21070505/
- **[S11]** https://pubmed.ncbi.nlm.nih.gov/17630229/ — https://pubmed.ncbi.nlm.nih.gov/17630229/
- **[S12]** https://pubmed.ncbi.nlm.nih.gov/23303064/ — https://pubmed.ncbi.nlm.nih.gov/23303064/
- **[S13]** https://pubmed.ncbi.nlm.nih.gov/28551713/ — https://pubmed.ncbi.nlm.nih.gov/28551713/
- **[S14]** https://pubmed.ncbi.nlm.nih.gov/37250466/ — https://pubmed.ncbi.nlm.nih.gov/37250466/
- **[S15]** https://www.ncbi.nlm.nih.gov/gene/6530 — https://www.ncbi.nlm.nih.gov/gene/6530
- **[S16]** https://pubmed.ncbi.nlm.nih.gov/28392265/ — https://pubmed.ncbi.nlm.nih.gov/28392265/
- **[S17]** https://pubmed.ncbi.nlm.nih.gov/15763139/ — https://pubmed.ncbi.nlm.nih.gov/15763139/
- **[S18]** https://www.ncbi.nlm.nih.gov/gene/6530 — https://www.ncbi.nlm.nih.gov/gene/6530
