## Supplementary material for "Understanding Substance Dependence: What Differentiates Addictive from Non-Addictive Drugs?": Folder for supplementary files: addictive-genes-claims-first100-079-slc8a1-gene-protein-is-implicated-in-addiction-293d15530574.md

# addictive-genes-claims-first100-079-293d15530574

**Original claim:** SLC8A1 gene/protein is implicated in addiction.

## Claim

SLC8A1 gene/protein is implicated in addiction.

## Verdict

Verdict: Mixed Evidence

## Confidence

Medium

## Supporting evidence

SLC8A1 encodes NCX1, a plasma-membrane Na\+/Ca\(2+\) exchanger. More broadly, the SLC8/NCX family regulates cellular Ca\(2+\) fluxes, contributes to neurotransmitter secretion, and has recognized roles in long-term potentiation and learning-related signaling. That makes SLC8A1 biologically plausible as a modifier of addiction-related neuroplasticity, even before looking at disorder-specific data. ([pubmed.ncbi.nlm.nih.gov](https://pubmed.ncbi.nlm.nih.gov/23506867/)) [S1]

There is preclinical evidence linking NCX1/SLC8A1 to alcohol-related phenotypes. In a 2024 rat study, the NCX1 reverse-mode inhibitor SN-6 significantly reduced voluntary ethanol intake, and in some analyses reduced ethanol preference, without materially changing water intake. The authors interpreted this as evidence that NCX1 activity can influence alcohol consumption behavior. ([pubmed.ncbi.nlm.nih.gov](https://pubmed.ncbi.nlm.nih.gov/38612942/?utm_source=openai)) [S2]

Related alcohol-withdrawal work strengthens that signal. In rat inferior colliculus, NCX1 protein levels were reported to be elevated at 3 and 24 hours after alcohol withdrawal, and local inhibition of NCX1 reverse-mode activity with SN-6 markedly suppressed the occurrence and severity of alcohol-withdrawal seizures. That does not prove SLC8A1 causes addiction, but it does implicate NCX1 in a dependence/withdrawal-relevant neuroadaptation. ([pmc.ncbi.nlm.nih.gov](https://pmc.ncbi.nlm.nih.gov/articles/PMC10577795/?utm_source=openai)) [S3]

There is also methamphetamine-related transcriptomic evidence. In a long-access methamphetamine self-administration rat model, Slc8a1 was listed among the genes strongly upregulated shortly after withdrawal in striatum, with a reported 7.56-fold increase at the 2-hour time point versus the 1-month withdrawal group. This is an addiction-model expression change, albeit in animals and without direct causal manipulation of Slc8a1 itself. ([pmc.ncbi.nlm.nih.gov](https://pmc.ncbi.nlm.nih.gov/articles/PMC4359351/)) [S4]

A separate mouse genetics study of methamphetamine sensitivity found Slc8a1 among differentially expressed genes in striatum within a locus affecting acute methamphetamine response. Importantly, however, that study ultimately validated **Hnrnph1**, not Slc8a1, as the causal quantitative trait gene. So Slc8a1 appeared in an addiction-relevant molecular signature, but was not established as the driver. ([pmc.ncbi.nlm.nih.gov](https://pmc.ncbi.nlm.nih.gov/articles/PMC4675533/)) [S5]

## Contradicting or limiting evidence

The biggest limitation is the lack of strong human genetic evidence. The current GWAS Catalog page for **SLC8A1** shows empty association/study/trait tables, which argues against SLC8A1 being an established genome-wide significant human addiction-risk gene at present. ([ebi.ac.uk](https://www.ebi.ac.uk/gwas/genes/SLC8A1?utm_source=openai)) [S6]

Recent reviews of substance-use-disorder genetics describe large GWAS now identifying many loci across alcohol, cannabis, opioid, and tobacco use disorders, but SLC8A1 is not highlighted there as a replicated, established locus. That is not proof of absence, but it suggests any human association is at most weak, indirect, or not yet reproducibly demonstrated. ([pubmed.ncbi.nlm.nih.gov](https://pubmed.ncbi.nlm.nih.gov/38811691/)) [S7]

The positive evidence is also uneven in scope. The clearest functional data concern **alcohol intake and alcohol withdrawal** in rodents, whereas direct evidence for SLC8A1 in cocaine, opioid, nicotine, or cannabis addiction is sparse. Methamphetamine-related evidence is mainly transcriptomic/correlative, not a demonstration that altering SLC8A1 changes compulsive drug seeking, relapse, or dependence in a validated causal way. ([pubmed.ncbi.nlm.nih.gov](https://pubmed.ncbi.nlm.nih.gov/38612942/?utm_source=openai)) [S8]

Even within the alcohol literature, the data are preclinical and rely heavily on pharmacologic inhibition of NCX reverse-mode activity. That supports involvement of the **protein function** in addiction-relevant phenotypes, but it is still a step short of showing that inherited or acquired changes in the **human SLC8A1 gene** materially alter addiction risk. ([pubmed.ncbi.nlm.nih.gov](https://pubmed.ncbi.nlm.nih.gov/38612942/?utm_source=openai)) [S9]

## Mechanistic / biological rationale

Mechanistically, the claim is plausible. Addiction is driven in part by maladaptive synaptic plasticity, altered excitability, and calcium-dependent signaling in reward-related circuits. NCX proteins regulate transmembrane Ca\(2+\) exchange and thereby influence Ca\(2+\)-dependent cellular events, neurotransmitter secretion, and plasticity-related processes such as long-term potentiation and learning. A molecule with that role could reasonably modulate addiction-related adaptations if its expression or operating mode changes in relevant neurons. ([pubmed.ncbi.nlm.nih.gov](https://pubmed.ncbi.nlm.nih.gov/23506867/)) [S10]

The alcohol-withdrawal studies offer a concrete proposed mechanism: when NCX1 operates in reverse mode, it can facilitate Ca\(2+\) entry rather than extrusion. In the withdrawal model, investigators proposed that increased NCX1 expression/activity contributed to disturbed Ca\(2+\) homeostasis, hyperexcitability, and withdrawal seizures; blocking NCX1 reverse mode reduced those phenotypes. That rationale fits known addiction biology, where calcium signaling and excitability are central to withdrawal and circuit remodeling. ([pmc.ncbi.nlm.nih.gov](https://pmc.ncbi.nlm.nih.gov/articles/PMC10577795/?utm_source=openai)) [S11]

Still, “biological plausibility” is not the same as “established addiction gene.” Many calcium-handling proteins can affect excitability and plasticity. To move SLC8A1 from plausible to firmly implicated, stronger evidence would be needed: replicated human genetic associations, human brain transcriptomic links in addiction, or animal studies showing that selective genetic manipulation of Slc8a1 changes drug self-administration, relapse, or dependence-related behavior. ([pubmed.ncbi.nlm.nih.gov](https://pubmed.ncbi.nlm.nih.gov/38811691/)) [S12]

## Bottom line

SLC8A1/NCX1 is **plausibly involved** in addiction biology and has **meaningful preclinical support**, especially for alcohol-related consumption and withdrawal phenotypes in rodents, plus some methamphetamine-associated expression changes in addiction models. However, the evidence is **not yet strong enough to call SLC8A1 an established human addiction gene**, because direct, replicated human genetic evidence appears limited or absent. Overall, the fairest assessment is **Mixed Evidence**: promising mechanistic and animal data, but insufficient human confirmation. ([pubmed.ncbi.nlm.nih.gov](https://pubmed.ncbi.nlm.nih.gov/38612942/?utm_source=openai)) [S13]

## Cited sources

- **[S1]** The SLC8 gene family of sodium-calcium exchangers (NCX) - structure, function, and regulation in health and disease - PubMed — https://pubmed.ncbi.nlm.nih.gov/23506867/
- **[S2]** Inhibition of the Sodium-Calcium Exchanger Reverse Mode Activity Reduces Alcohol Consumption in Rats - PubMed — https://pubmed.ncbi.nlm.nih.gov/38612942/?utm_source=openai
- **[S3]** Targeted inhibition of upregulated sodium-calcium exchanger in rat inferior colliculus suppresses alcohol withdrawal seizures - PMC — https://pmc.ncbi.nlm.nih.gov/articles/PMC10577795/?utm_source=openai
- **[S4]** 
            Transcriptional and Epigenetic Substrates of Methamphetamine Addiction and Withdrawal: Evidence from a Long-Access Self-Administration Model in the Rat - PMC
         — https://pmc.ncbi.nlm.nih.gov/articles/PMC4359351/
- **[S5]** 
            Hnrnph1 Is A Quantitative Trait Gene for Methamphetamine Sensitivity - PMC
         — https://pmc.ncbi.nlm.nih.gov/articles/PMC4675533/
- **[S6]** GWAS Catalog — https://www.ebi.ac.uk/gwas/genes/SLC8A1?utm_source=openai
- **[S7]** The genetic landscape of substance use disorders - PubMed — https://pubmed.ncbi.nlm.nih.gov/38811691/
- **[S8]** Inhibition of the Sodium-Calcium Exchanger Reverse Mode Activity Reduces Alcohol Consumption in Rats - PubMed — https://pubmed.ncbi.nlm.nih.gov/38612942/?utm_source=openai
- **[S9]** Inhibition of the Sodium-Calcium Exchanger Reverse Mode Activity Reduces Alcohol Consumption in Rats - PubMed — https://pubmed.ncbi.nlm.nih.gov/38612942/?utm_source=openai
- **[S10]** The SLC8 gene family of sodium-calcium exchangers (NCX) - structure, function, and regulation in health and disease - PubMed — https://pubmed.ncbi.nlm.nih.gov/23506867/
- **[S11]** Targeted inhibition of upregulated sodium-calcium exchanger in rat inferior colliculus suppresses alcohol withdrawal seizures - PMC — https://pmc.ncbi.nlm.nih.gov/articles/PMC10577795/?utm_source=openai
- **[S12]** The genetic landscape of substance use disorders - PubMed — https://pubmed.ncbi.nlm.nih.gov/38811691/
- **[S13]** Inhibition of the Sodium-Calcium Exchanger Reverse Mode Activity Reduces Alcohol Consumption in Rats - PubMed — https://pubmed.ncbi.nlm.nih.gov/38612942/?utm_source=openai
