## Supplementary material for "Understanding Substance Dependence: What Differentiates Addictive from Non-Addictive Drugs?": Folder for supplementary files: addictive-genes-claims-first100-080-ddah1-gene-protein-is-implicated-in-addiction-9a9120c51d4f.md

# addictive-genes-claims-first100-080-9a9120c51d4f

**Original claim:** DDAH1 gene/protein is implicated in addiction.

## Claim

DDAH1 gene/protein is implicated in addiction.

## Verdict

Verdict: Mixed Evidence

## Confidence

Medium. ([pubmed.ncbi.nlm.nih.gov](https://pubmed.ncbi.nlm.nih.gov/41317783/)) [S1]

## Supporting evidence

A recent dedicated review focused specifically on DDAH1 and addiction concluded that proteomic, metabolomic, and functional studies **suggest DDAH1 may play a role** in addiction-related phenotypes, especially for psychostimulants, while also emphasizing that the literature is still sparse. The authors summarized evidence that psychostimulants can increase brain DDAH1 levels, that opioids may also increase DDAH1, and that existing data are insufficient for firm mechanistic conclusions. ([pubmed.ncbi.nlm.nih.gov](https://pubmed.ncbi.nlm.nih.gov/41317783/)) [S2]

The strongest direct preclinical support comes from **methamphetamine models**. In a rat proteomics study, DDAH1 expression increased in the striatum and frontal cortex after methamphetamine exposure, placing DDAH1 among proteins altered by methamphetamine neurotoxicity. ([pubmed.ncbi.nlm.nih.gov](https://pubmed.ncbi.nlm.nih.gov/17904249/)) [S3] A follow-up study reported that acute methamphetamine activated the **DDAH1/ADMA/NOS** pathway, increased nitric oxide production and protein tyrosine nitration, and that a DDAH1 inhibitor suppressed these changes and reduced apoptosis/cell death in rat brain and PC12 cells. That is stronger-than-associative evidence that DDAH1 can modulate at least one biologically relevant consequence of psychostimulant exposure. ([pubmed.ncbi.nlm.nih.gov](https://pubmed.ncbi.nlm.nih.gov/23583342/)) [S4]

There is also **genetic/functional animal evidence** linking DDAH1 to stimulant-related behavioral responses. In global Ddah1 knockout mice, loss of DDAH1 altered amphetamine response kinetics and reduced DOPAC, a dopamine metabolite, in striatal/caudoputamen-related regions. The authors interpreted this as support for involvement of the DDAH/ADMA pathway in CNS processes shaping behavioral output. Although this is not an addiction model per se, amphetamine sensitivity is a relevant addiction-related endophenotype. ([pubmed.ncbi.nlm.nih.gov](https://pubmed.ncbi.nlm.nih.gov/36792833/)) [S5]

More broadly, DDAH1 is biologically positioned to matter in addiction-relevant circuits because it is **widely expressed in adult mouse and human brain**, including cortex, caudoputamen/striatum, dorsal raphe, hippocampal formation, and vascular/endothelial compartments. This distribution overlaps regions central to reward, salience, and drug-induced plasticity. ([pubmed.ncbi.nlm.nih.gov](https://pubmed.ncbi.nlm.nih.gov/34014421/)) [S6]

Human evidence is limited but not absent. A 2025 psychiatric review summarized **GWAS/gene-based associations involving DDAH1** with alcohol dependence, alcohol consumption, and amphetamine response, while also noting broader DDAH1/DDAH2 links to psychiatric phenotypes. That does not prove causality, but it supports the statement that DDAH1 has appeared in human addiction-relevant association datasets. ([pmc.ncbi.nlm.nih.gov](https://pmc.ncbi.nlm.nih.gov/articles/PMC12466234/)) [S7]

Finally, the mechanistic background is plausible: nitric oxide signaling has long been implicated in substance dependence across opioids, ethanol, psychostimulants, and nicotine, with NOS inhibitors reported to reduce withdrawal signs in several drug classes. Since DDAH1 is a major regulator of ADMA, an endogenous NOS inhibitor, DDAH1 sits upstream of a pathway already linked to addiction biology. ([pubmed.ncbi.nlm.nih.gov](https://pubmed.ncbi.nlm.nih.gov/11166077/)) [S8]

## Contradicting or limiting evidence

The main limitation is that the evidence base is **small and uneven**. Even the recent addiction-focused review states that mechanistic evidence is limited and that available data are sparse, especially outside psychostimulants. Its own conclusion is cautious: DDAH1 **may** play a role, but further work is needed to define its specific function in addiction. ([pubmed.ncbi.nlm.nih.gov](https://pubmed.ncbi.nlm.nih.gov/41317783/)) [S9]

Human evidence is particularly weak. Much of it is **indirect**, often relying on peripheral ADMA measurements rather than direct brain DDAH1 assays, and even those peripheral studies are inconsistent. The psychiatric review noted that alcohol-dependence studies of ADMA have yielded conflicting directions, with some reporting increases and others decreases; it also noted that withdrawal tends to change ADMA levels regardless of baseline direction. That means the human biomarker signal is not robust or specific. ([pmc.ncbi.nlm.nih.gov](https://pmc.ncbi.nlm.nih.gov/articles/PMC12466234/)) [S10]

Some addiction-relevant preclinical findings are negative or non-supportive. In mice made tolerant/dependent to morphine, plasma ADMA, SDMA, homoarginine, and arginine did **not** differ from controls, and the authors concluded these measures were not associated with development of morphine tolerance/dependence. This does not rule out a brain-specific DDAH1 role, but it weakens claims of a generalizable addiction signal. ([pubmed.ncbi.nlm.nih.gov](https://pubmed.ncbi.nlm.nih.gov/20661755/)) [S11]

Even the more positive Ddah1 knockout study did **not** show a classic addiction phenotype such as self-administration, escalation, relapse, or conditioned place preference. It showed altered amphetamine sensitivity and dopamine metabolism, which are relevant but still indirect. ([pubmed.ncbi.nlm.nih.gov](https://pubmed.ncbi.nlm.nih.gov/36792833/)) [S12]

Likewise, the methamphetamine studies mainly address **neurotoxicity/oxidative injury**, not addiction behavior itself. Addiction and neurotoxicity overlap biologically but are not the same claim. So evidence that DDAH1 contributes to methamphetamine-induced cellular damage is not by itself proof that DDAH1 drives compulsive drug seeking or dependence. ([pubmed.ncbi.nlm.nih.gov](https://pubmed.ncbi.nlm.nih.gov/23583342/)) [S13]

## Mechanistic / biological rationale

DDAH1 degrades **asymmetric dimethylarginine (ADMA)**, a major endogenous inhibitor of nitric oxide synthase. By lowering ADMA, DDAH1 can increase nitric oxide signaling. In the CNS, nitric oxide functions as a signaling molecule involved in neurotransmission and plasticity, and DDAH1 is the predominant, broadly distributed DDAH isoform in the postnatal brain. ([pubmed.ncbi.nlm.nih.gov](https://pubmed.ncbi.nlm.nih.gov/34014421/)) [S14]

That matters because nitric oxide signaling has been implicated in multiple addiction-relevant processes, including withdrawal, glutamatergic signaling, and drug-induced neural adaptations. Reviews of NO in substance dependence describe NOS inhibition as attenuating withdrawal signs for opioids, ethanol, and nicotine, suggesting that NO is part of a shared dependence-related signaling framework. ([pubmed.ncbi.nlm.nih.gov](https://pubmed.ncbi.nlm.nih.gov/11166077/)) [S15]

Within stimulant models, the proposed DDAH1 mechanism is especially concrete: increased DDAH1 lowers ADMA, disinhibits NOS, increases NO/peroxynitrite-related stress and protein nitration, and thereby contributes to methamphetamine-associated neuronal injury. Separately, Ddah1 knockout alters dopamine metabolite levels and amphetamine responsiveness, suggesting a link between the DDAH1/ADMA/NO axis and dopaminergic tone. Since addiction depends heavily on dopamine- and glutamate-dependent plasticity in striatal and cortical circuits, this is a biologically credible route by which DDAH1 could influence addiction phenotypes. ([pubmed.ncbi.nlm.nih.gov](https://pubmed.ncbi.nlm.nih.gov/23583342/)) [S16]

Still, this rationale is stronger for **modulating addiction-related neurobiology** than for proving DDAH1 is a core addiction gene in humans. The mechanistic bridge is plausible, but not yet fully established. ([pubmed.ncbi.nlm.nih.gov](https://pubmed.ncbi.nlm.nih.gov/41317783/)) [S17]

## Bottom line

The claim is **not unsupported**, but it is also **not firmly established**. DDAH1 has credible preclinical links to stimulant-related neurobiology, including methamphetamine-induced DDAH1 upregulation and pharmacologic rescue, and Ddah1 loss alters amphetamine response and dopamine metabolism. There are also some human association signals summarized in recent reviews. However, direct human evidence is limited, biomarker findings are inconsistent, and many studies address neurotoxicity or endophenotypes rather than addiction diagnosis/behavior itself. Overall, DDAH1 is **reasonably viewed as a candidate or emerging addiction-related gene/protein**, especially in psychostimulant contexts, but the current evidence is best described as **mixed rather than definitive**. ([pubmed.ncbi.nlm.nih.gov](https://pubmed.ncbi.nlm.nih.gov/41317783/)) [S18]

## Cited sources

- **[S1]** Involvement of dimethylarginine dimethylaminohydrolase type 1 (DDAH1) in addiction and social stress phenotypes: Insights from proteomic, metabolomic and functional studies - PubMed — https://pubmed.ncbi.nlm.nih.gov/41317783/
- **[S2]** Involvement of dimethylarginine dimethylaminohydrolase type 1 (DDAH1) in addiction and social stress phenotypes: Insights from proteomic, metabolomic and functional studies - PubMed — https://pubmed.ncbi.nlm.nih.gov/41317783/
- **[S3]** Proteomic profiling of proteins associated with methamphetamine-induced neurotoxicity in different regions of rat brain - PubMed — https://pubmed.ncbi.nlm.nih.gov/17904249/
- **[S4]** Up-regulation of protein tyrosine nitration in methamphetamine-induced neurotoxicity through DDAH/ADMA/NOS pathway - PubMed — https://pubmed.ncbi.nlm.nih.gov/23583342/
- **[S5]** Knock-out of the critical nitric oxide synthase regulator DDAH1 in mice impacts amphetamine sensitivity and dopamine metabolism - PubMed — https://pubmed.ncbi.nlm.nih.gov/36792833/
- **[S6]** Divergent Dimethylarginine Dimethylaminohydrolase Isoenzyme Expression in the Central Nervous System - PubMed — https://pubmed.ncbi.nlm.nih.gov/34014421/
- **[S7]** 
            The Emerging Role of the DDAH Proteins in Psychiatric Disorders - PMC
         — https://pmc.ncbi.nlm.nih.gov/articles/PMC12466234/
- **[S8]** Nitric oxide and substance dependence - PubMed — https://pubmed.ncbi.nlm.nih.gov/11166077/
- **[S9]** Involvement of dimethylarginine dimethylaminohydrolase type 1 (DDAH1) in addiction and social stress phenotypes: Insights from proteomic, metabolomic and functional studies - PubMed — https://pubmed.ncbi.nlm.nih.gov/41317783/
- **[S10]** 
            The Emerging Role of the DDAH Proteins in Psychiatric Disorders - PMC
         — https://pmc.ncbi.nlm.nih.gov/articles/PMC12466234/
- **[S11]** Modulatory role of the endogenous nitric oxide synthase inhibitor, asymmetric dimethylarginine (ADMA), in morphine tolerance and dependence in mice - PubMed — https://pubmed.ncbi.nlm.nih.gov/20661755/
- **[S12]** Knock-out of the critical nitric oxide synthase regulator DDAH1 in mice impacts amphetamine sensitivity and dopamine metabolism - PubMed — https://pubmed.ncbi.nlm.nih.gov/36792833/
- **[S13]** Up-regulation of protein tyrosine nitration in methamphetamine-induced neurotoxicity through DDAH/ADMA/NOS pathway - PubMed — https://pubmed.ncbi.nlm.nih.gov/23583342/
- **[S14]** Divergent Dimethylarginine Dimethylaminohydrolase Isoenzyme Expression in the Central Nervous System - PubMed — https://pubmed.ncbi.nlm.nih.gov/34014421/
- **[S15]** Nitric oxide and substance dependence - PubMed — https://pubmed.ncbi.nlm.nih.gov/11166077/
- **[S16]** Up-regulation of protein tyrosine nitration in methamphetamine-induced neurotoxicity through DDAH/ADMA/NOS pathway - PubMed — https://pubmed.ncbi.nlm.nih.gov/23583342/
- **[S17]** Involvement of dimethylarginine dimethylaminohydrolase type 1 (DDAH1) in addiction and social stress phenotypes: Insights from proteomic, metabolomic and functional studies - PubMed — https://pubmed.ncbi.nlm.nih.gov/41317783/
- **[S18]** Involvement of dimethylarginine dimethylaminohydrolase type 1 (DDAH1) in addiction and social stress phenotypes: Insights from proteomic, metabolomic and functional studies - PubMed — https://pubmed.ncbi.nlm.nih.gov/41317783/
