## Supplementary material for "Understanding Substance Dependence: What Differentiates Addictive from Non-Addictive Drugs?": Folder for supplementary files: addictive-genes-claims-first100-081-sf3b1-gene-protein-is-implicated-in-addiction-1aa91d114e96.md

# addictive-genes-claims-first100-081-1aa91d114e96

**Original claim:** SF3B1 gene/protein is implicated in addiction.

## Claim

SF3B1 gene/protein is implicated in addiction.

## Verdict

Verdict: Mixed Evidence

## Confidence

Medium

## Supporting evidence

SF3B1 is a biologically plausible nervous-system gene. It encodes a core component of the U2 snRNP spliceosome, which helps recognize intron branch points and 3′ splice sites, and NCBI summarizes it as a ubiquitously expressed splicing factor with expression across many tissues. Human Protein Atlas data also indicate brain-region expression rather than brain absence. ([ncbi.nlm.nih.gov](https://www.ncbi.nlm.nih.gov/gene/23451)) [S1]

There is also some **indirect human evidence** connecting SF3B1 to neuropsychiatric phenotypes relevant to addiction biology. NCBI’s gene page lists an EBI GWAS Catalog association for schizophrenia at the SF3B1 locus, and multiple psychiatric-genetics papers discuss SF3B1 as lying within or near schizophrenia-associated loci. That does **not** prove addiction involvement, but it does show that SF3B1 has prior evidence linking it to brain-related disease rather than being purely a hematology/cancer gene. ([ncbi.nlm.nih.gov](https://www.ncbi.nlm.nih.gov/gene/23451)) [S2]

A 2026 UK Biobank study on clonal hematopoiesis reported that CHIP was associated prospectively with later psychiatric disorders, including substance use disorders, and that gene-specific CHIP signals included SF3B1 among the seven most frequent CHIP driver genes with increased psychiatric risk. This is not a standard addiction-genetics result, because it concerns **somatic blood-cell mutations** rather than inherited brain risk variants, but it is still a published human association placing SF3B1 in the orbit of substance-use-disorder risk. ([pubmed.ncbi.nlm.nih.gov](https://pubmed.ncbi.nlm.nih.gov/41709299/)) [S3]

There is also **molecularly relevant but indirect** evidence from opioid biology. In a study of an OPRM1 intronic SNP associated with heroin addiction severity, RNA affinity purification identified SF3B1 among U2 snRNP-related proteins preferentially recruited to the addiction-associated RNA element. That does not show that SF3B1 variants cause addiction, but it does place the SF3B1-containing spliceosomal machinery in a mechanistic context linked to heroin-relevant OPRM1 splicing. ([pmc.ncbi.nlm.nih.gov](https://pmc.ncbi.nlm.nih.gov/articles/PMC4131016/)) [S4]

Finally, addiction research broadly supports the idea that RNA splicing matters in reward circuits. Cocaine self-administration in mice caused widespread alternative splicing changes across reward-related brain regions, and heroin exposure in humans and rats was associated with altered expression of alternatively spliced OPRM1 transcripts in medial prefrontal cortex. This broader literature makes SF3B1 mechanistically plausible as a contributor to addiction-related biology, even though these studies did not establish SF3B1 itself as the driver. ([pubmed.ncbi.nlm.nih.gov](https://pubmed.ncbi.nlm.nih.gov/34480866/)) [S5]

## Contradicting or limiting evidence

The main limitation is that **direct evidence is sparse**. I did not find strong human studies showing that inherited SF3B1 variants reproducibly increase risk for alcohol, nicotine, opioid, stimulant, or general substance use disorder in the way that established addiction genes do. By contrast, large GWAS for opioid use disorder, alcohol dependence, and nicotine dependence highlight loci such as **OPRM1/DRD2/FURIN**, **ADH1B**, and **CHRNA5**, respectively, rather than SF3B1. Absence from a few GWAS is not definitive disproof, but it argues against SF3B1 being an established addiction gene. ([pubmed.ncbi.nlm.nih.gov](https://pubmed.ncbi.nlm.nih.gov/37252880/)) [S6]

The strongest “supporting” human addiction-related signal I found for SF3B1 comes from the CHIP study, but that evidence is hard to interpret as proof of addiction biology per se. CHIP involves age-related somatic mutations in hematopoietic cells, not germline variation or direct neuronal manipulation, so the finding could reflect systemic inflammation, vascular effects, or other indirect pathways rather than a primary addiction mechanism in reward circuits. ([pubmed.ncbi.nlm.nih.gov](https://pubmed.ncbi.nlm.nih.gov/41709299/)) [S7]

Likewise, the OPRM1 splicing paper does not identify SF3B1 as a causal addiction gene. SF3B1 appeared as one member of a larger spliceosomal protein network associated with an OPRM1 RNA element; the study’s main mechanistic driver was hnRNPH and the functional phenotype centered on an OPRM1 SNP. So this is best viewed as **contextual involvement of spliceosomal machinery**, not proof that SF3B1 itself is implicated in addiction at the organism level. ([pmc.ncbi.nlm.nih.gov](https://pmc.ncbi.nlm.nih.gov/articles/PMC4131016/)) [S8]

More generally, the best-established disease roles of SF3B1 remain in **myelodysplastic syndromes, hematologic malignancy, solid tumors, and more recently neurodevelopmental disorders**, not addiction. NCBI and recent primary papers emphasize those domains far more strongly than substance-use phenotypes. ([ncbi.nlm.nih.gov](https://www.ncbi.nlm.nih.gov/gene/23451)) [S9]

## Mechanistic / biological rationale

A plausible mechanistic case exists, but it is mostly **indirect**. SF3B1 is a core splicing factor, and neurons rely heavily on alternative splicing to shape receptor isoforms, signaling proteins, synaptic components, and activity-dependent plasticity. Because addiction is fundamentally a disorder of durable neuroplasticity, any major splicing regulator could in principle influence vulnerability or adaptation. ([ncbi.nlm.nih.gov](https://www.ncbi.nlm.nih.gov/gene/23451)) [S10]

That rationale becomes more concrete for opioid biology. OPRM1 undergoes extensive alternative splicing, and human heroin abusers as well as heroin self-administering rats show dysregulated OPRM1 splice-variant expression in medial prefrontal cortex. A separate mechanistic study linked an addiction-severity-associated OPRM1 intronic variant to recruitment of hnRNPH and spliceosomal proteins including SF3B1. Together, these data make it biologically reasonable to hypothesize that SF3B1-containing spliceosomal complexes could modulate addiction-relevant receptor isoforms. Still, this remains a hypothesis more than a demonstrated SF3B1-specific addiction mechanism. ([pubmed.ncbi.nlm.nih.gov](https://pubmed.ncbi.nlm.nih.gov/32506472/)) [S11]

There is also evidence that SF3B1 has bona fide neuronal functions outside cancer. Experimental work shows roles in neurogenesis, neurite/dendrite maintenance, and mitochondrial homeostasis in neurons, and recent human genetics links de novo SF3B1 variants to neurodevelopmental disorders. Those findings strengthen the idea that perturbing SF3B1 could affect brain circuitry, but they stop short of showing addiction-specific effects. ([pmc.ncbi.nlm.nih.gov](https://pmc.ncbi.nlm.nih.gov/articles/PMC12504175/)) [S12]

## Bottom line

SF3B1 is **not an established addiction gene** in the same sense as OPRM1, ADH1B, or CHRNA5. However, there is **some indirect evidence** putting it near addiction-related biology: it is a neuronal splicing factor, splice regulation is clearly relevant to addiction, SF3B1-containing spliceosomal machinery appears in an OPRM1/heroin-severity splicing study, and a recent CHIP study linked SF3B1-mutant hematopoietic clones with later substance use disorders. Overall, the claim is **not strongly proven**, but it is **not baseless either**—best summarized as **mixed and mostly indirect evidence**. ([pmc.ncbi.nlm.nih.gov](https://pmc.ncbi.nlm.nih.gov/articles/PMC4131016/)) [S13]

## Cited sources

- **[S1]** SF3B1 splicing factor 3b subunit 1 [Homo sapiens (human)] - Gene - NCBI — https://www.ncbi.nlm.nih.gov/gene/23451
- **[S2]** SF3B1 splicing factor 3b subunit 1 [Homo sapiens (human)] - Gene - NCBI — https://www.ncbi.nlm.nih.gov/gene/23451
- **[S3]** Clonal haematopoiesis of indeterminate potential and risk of incident psychiatric disorders - PubMed — https://pubmed.ncbi.nlm.nih.gov/41709299/
- **[S4]** 
            A Heroin Addiction Severity-Associated Intronic Single Nucleotide Polymorphism Modulates Alternative Pre-mRNA Splicing of the μ Opioid Receptor Gene OPRM1 via hnRNPH Interactions - PMC
         — https://pmc.ncbi.nlm.nih.gov/articles/PMC4131016/
- **[S5]** Chromatin-mediated alternative splicing regulates cocaine-reward behavior - PubMed — https://pubmed.ncbi.nlm.nih.gov/34480866/
- **[S6]** Genome-wide analyses reveal novel opioid use disorder loci and genetic overlap with schizophrenia, bipolar disorder, and major depression - PubMed — https://pubmed.ncbi.nlm.nih.gov/37252880/
- **[S7]** Clonal haematopoiesis of indeterminate potential and risk of incident psychiatric disorders - PubMed — https://pubmed.ncbi.nlm.nih.gov/41709299/
- **[S8]** 
            A Heroin Addiction Severity-Associated Intronic Single Nucleotide Polymorphism Modulates Alternative Pre-mRNA Splicing of the μ Opioid Receptor Gene OPRM1 via hnRNPH Interactions - PMC
         — https://pmc.ncbi.nlm.nih.gov/articles/PMC4131016/
- **[S9]** SF3B1 splicing factor 3b subunit 1 [Homo sapiens (human)] - Gene - NCBI — https://www.ncbi.nlm.nih.gov/gene/23451
- **[S10]** SF3B1 splicing factor 3b subunit 1 [Homo sapiens (human)] - Gene - NCBI — https://www.ncbi.nlm.nih.gov/gene/23451
- **[S11]** Dysregulated expression of the alternatively spliced variant mRNAs of the mu opioid receptor gene, OPRM1, in the medial prefrontal cortex of male human heroin abusers and heroin self-administering male rats - PubMed — https://pubmed.ncbi.nlm.nih.gov/32506472/
- **[S12]** 
            Splicing factor Sf3b1 facilitates maintenance of neuronal dendrites by modulating mitochondrial health - PMC
         — https://pmc.ncbi.nlm.nih.gov/articles/PMC12504175/
- **[S13]** 
            A Heroin Addiction Severity-Associated Intronic Single Nucleotide Polymorphism Modulates Alternative Pre-mRNA Splicing of the μ Opioid Receptor Gene OPRM1 via hnRNPH Interactions - PMC
         — https://pmc.ncbi.nlm.nih.gov/articles/PMC4131016/
