## Supplementary material for "Understanding Substance Dependence: What Differentiates Addictive from Non-Addictive Drugs?": Folder for supplementary files: addictive-genes-claims-first100-082-crabp1-gene-protein-is-implicated-in-addiction-5026d06a7aa8.md

# addictive-genes-claims-first100-082-5026d06a7aa8

**Original claim:** CRABP1 gene/protein is implicated in addiction.

## Claim

CRABP1 gene/protein is implicated in addiction.

## Verdict

Verdict: Mixed Evidence

Direct evidence that **CRABP1 itself** contributes to addiction is limited. One pathway-level analysis of smoking-quantity GWAS found enrichment of retinoid-binding genes and included **CRABP1** among contributing genes, with CRABP1 noted as significant in the ARIC dataset. However, larger and more direct addiction-genetics studies repeatedly localize the strongest nicotine-risk signals at **15q25** to **CHRNA5-CHRNA3-CHRNB4** and related expression effects—especially **CHRNA5**—rather than to CRABP1. ([genepi.qimr.edu.au](https://genepi.qimr.edu.au/contents/p/staff/Harari_PLOSONE_e50913_.pdf)) [S1]

## Confidence

Medium

CRABP1 is biologically plausible because it binds retinoic acid and modulates intracellular signaling, while retinoid signaling in nucleus accumbens circuits has been shown to affect drug reward sensitivity, fentanyl taking, and cocaine-craving-related plasticity. But the step from “retinoid signaling matters in addiction” to “CRABP1 specifically matters” is still mostly inferential. ([ncbi.nlm.nih.gov](https://www.ncbi.nlm.nih.gov/gene/1381)) [S2]

## Supporting evidence

**CRABP1** encodes cellular retinoic acid-binding protein 1 and is located at **15q25.1**. In a pathway analysis of smoking quantity across the OZALC-NAG, SAGE, and ARIC datasets, retinoid-binding/retinoic-acid-binding gene sets were enriched, and the authors specifically note that **CRABP1** contributed to the ARIC signal. This is the clearest human genetic evidence I found linking CRABP1 to an addiction-related phenotype, although it is pathway-level rather than a definitive gene-level hit. ([ncbi.nlm.nih.gov](https://www.ncbi.nlm.nih.gov/gene/1381)) [S3]

CRABP1 also has neurobehavioral functions that are relevant to addiction biology. In mice, **Crabp1 knockout** reduced anxiety-like behavior, lowered stress-induced corticosterone, and altered HPA-axis feedback, with reduced hypothalamic **FKBP5** expression. That does not prove an addiction role, but it places CRABP1 in stress-responsive neuroendocrine biology that often intersects with substance-use vulnerability and relapse. ([pubmed.ncbi.nlm.nih.gov](https://pubmed.ncbi.nlm.nih.gov/34830120/)) [S4]

Mechanistically, CRABP1 directly interacts with and dampens **CaMKII** signaling. Separate studies showed CRABP1 suppresses CaMKII activity and, in motor neurons, CRABP1 loss increased phospho-CaMKII and reduced **Agrin** expression. Because CaMKII-linked plasticity is relevant to reward learning and long-term neural adaptation, this gives CRABP1 a plausible route into addiction-related mechanisms. ([pubmed.ncbi.nlm.nih.gov](https://pubmed.ncbi.nlm.nih.gov/29371236/?utm_source=openai)) [S5]

Recent primary studies also strengthen the broader retinoid-addiction connection. In nucleus accumbens, **RXRα** controls addiction-relevant transcriptional programs and drug reward sensitivity; reducing retinoic acid synthesis in accumbens shell produced a more “protective” phenotype and altered fentanyl-taking behavior; and manipulating retinoic acid signaling in accumbens core changed CP-AMPAR accumulation and incubation of cocaine craving. These studies did **not** test CRABP1 directly, but they make a CRABP1 role biologically plausible by inference. ([pubmed.ncbi.nlm.nih.gov](https://pubmed.ncbi.nlm.nih.gov/36889314/)) [S6]

## Contradicting or limiting evidence

The strongest limitation is that the best-replicated addiction genetics in the same general **15q25** region do **not** point to CRABP1. Large nicotine-dependence GWAS/meta-analyses consistently replicate **CHRNA5-CHRNA3-CHRNB4**, and a 15q25 brain eQTL study linked shared smoking/schizophrenia risk variants to expression of **CHRNA5, CHRNA3, CHRNB4, PSMA4,** and **HYKK**—not CRABP1. That makes CRABP1 a weaker candidate than the nicotinic receptor genes for direct involvement in tobacco addiction. ([pubmed.ncbi.nlm.nih.gov](https://pubmed.ncbi.nlm.nih.gov/28972577/)) [S7]

Even the smoking-quantity paper that is most favorable to CRABP1 is not decisive. Its support came from **gene-set/pathway enrichment**, not from a replicated CRABP1-specific genome-wide significant association. The same paper reports that only the **cholinergic receptor** GO terms were consistently significant across both pathway methods used, indicating that the retinoid/CRABP1 signal was weaker and more method-dependent. ([genepi.qimr.edu.au](https://genepi.qimr.edu.au/contents/p/staff/Harari_PLOSONE_e50913_.pdf)) [S8]

Across other addiction domains, recent high-powered primary studies emphasize other genes. For opioid addiction/OUD, replicated signals center on **OPRM1**, with additional gene-based hits such as **PPP6C** and **FURIN**. I did not find comparable primary evidence elevating CRABP1 as a replicated locus for opioid addiction. ([pubmed.ncbi.nlm.nih.gov](https://pubmed.ncbi.nlm.nih.gov/36207451/)) [S9]

There is also a phenotype mismatch. Most direct CRABP1 experiments so far concern **stress/HPA regulation**, **energy homeostasis**, **cardiac signaling**, or **motor-neuron maintenance**, rather than drug reward, compulsive intake, withdrawal, or relapse. So the current case for CRABP1 in addiction is still largely associative and mechanistic rather than demonstration-based. ([pubmed.ncbi.nlm.nih.gov](https://pubmed.ncbi.nlm.nih.gov/34830120/)) [S10]

## Mechanistic / biological rationale

CRABP1 is a cytosolic retinoic-acid-binding protein with signaling effects that extend beyond classical nuclear retinoid receptor transcription. It can modulate intracellular pathways including **CaMKII**, which is relevant because addiction involves durable signaling and plasticity changes in reward circuits. ([ncbi.nlm.nih.gov](https://www.ncbi.nlm.nih.gov/gene/1381)) [S11]

A cautious mechanistic model would be: **CRABP1 alters local RA signaling and CaMKII-linked plasticity; retinoid signaling in nucleus accumbens influences reward sensitivity, emotional reactivity, fentanyl taking, and cocaine-craving-related adaptations; therefore CRABP1 could modulate addiction vulnerability or expression of addiction-related behaviors.** But that remains an inference, because the key reward-circuit studies manipulated **RXRα, Aldh1a1,** or **Cyp26b1/RA tone**, not CRABP1 itself. ([pubmed.ncbi.nlm.nih.gov](https://pubmed.ncbi.nlm.nih.gov/36889314/)) [S12]

CRABP1’s effects on HPA-axis feedback and anxiety-like behavior provide a second plausible bridge, especially for stress-triggered escalation or relapse. Still, those data more directly support a role in **stress biology** than in addiction biology. ([pubmed.ncbi.nlm.nih.gov](https://pubmed.ncbi.nlm.nih.gov/34830120/)) [S13]

## Bottom line

CRABP1 is **not currently a strongly established addiction gene/protein**, but it is **not an implausible candidate**. The evidence is best described as **indirect and limited**: a pathway-level smoking association plus mechanistic plausibility from CRABP1’s roles in retinoid/CaMKII signaling and stress regulation, set against stronger replicated addiction evidence for neighboring or alternative genes such as **CHRNA5-CHRNA3-CHRNB4** and **OPRM1**. So the claim that **“CRABP1 is implicated in addiction”** should be stated cautiously. ([genepi.qimr.edu.au](https://genepi.qimr.edu.au/contents/p/staff/Harari_PLOSONE_e50913_.pdf)) [S14]

## Cited sources

- **[S1]** pone.0050913 1..9 — https://genepi.qimr.edu.au/contents/p/staff/Harari_PLOSONE_e50913_.pdf
- **[S2]** CRABP1 cellular retinoic acid binding protein 1 [Homo sapiens (human)] - Gene - NCBI — https://www.ncbi.nlm.nih.gov/gene/1381
- **[S3]** CRABP1 cellular retinoic acid binding protein 1 [Homo sapiens (human)] - Gene - NCBI — https://www.ncbi.nlm.nih.gov/gene/1381
- **[S4]** Crabp1 Modulates HPA Axis Homeostasis and Anxiety-like Behaviors by Altering FKBP5 Expression - PubMed — https://pubmed.ncbi.nlm.nih.gov/34830120/
- **[S5]** CRABP1 protects the heart from isoproterenol-induced acute and chronic remodeling - PubMed — https://pubmed.ncbi.nlm.nih.gov/29371236/?utm_source=openai
- **[S6]** Transcriptional control of nucleus accumbens neuronal excitability by retinoid X receptor alpha tunes sensitivity to drug rewards - PubMed — https://pubmed.ncbi.nlm.nih.gov/36889314/
- **[S7]** Genome-wide association study across European and African American ancestries identifies a SNP in DNMT3B contributing to nicotine dependence - PubMed — https://pubmed.ncbi.nlm.nih.gov/28972577/
- **[S8]** pone.0050913 1..9 — https://genepi.qimr.edu.au/contents/p/staff/Harari_PLOSONE_e50913_.pdf
- **[S9]** Multi-trait genome-wide association study of opioid addiction: OPRM1 and beyond - PubMed — https://pubmed.ncbi.nlm.nih.gov/36207451/
- **[S10]** Crabp1 Modulates HPA Axis Homeostasis and Anxiety-like Behaviors by Altering FKBP5 Expression - PubMed — https://pubmed.ncbi.nlm.nih.gov/34830120/
- **[S11]** CRABP1 cellular retinoic acid binding protein 1 [Homo sapiens (human)] - Gene - NCBI — https://www.ncbi.nlm.nih.gov/gene/1381
- **[S12]** Transcriptional control of nucleus accumbens neuronal excitability by retinoid X receptor alpha tunes sensitivity to drug rewards - PubMed — https://pubmed.ncbi.nlm.nih.gov/36889314/
- **[S13]** Crabp1 Modulates HPA Axis Homeostasis and Anxiety-like Behaviors by Altering FKBP5 Expression - PubMed — https://pubmed.ncbi.nlm.nih.gov/34830120/
- **[S14]** pone.0050913 1..9 — https://genepi.qimr.edu.au/contents/p/staff/Harari_PLOSONE_e50913_.pdf
