## Supplementary material for "Understanding Substance Dependence: What Differentiates Addictive from Non-Addictive Drugs?": Folder for supplementary files: addictive-genes-claims-first100-083-wsb1-gene-protein-is-implicated-in-addiction-353c38d03d51.md

# addictive-genes-claims-first100-083-353c38d03d51

**Original claim:** WSB1 gene/protein is implicated in addiction.

## Claim

WSB1 gene/protein is implicated in addiction.

## Verdict

Verdict: No Evidence

I did **not** find convincing direct evidence that **WSB1** is an established addiction gene/protein in humans or animal models. In large addiction-genetics reviews and multi-omics studies, other genes and pathways recur, but WSB1 does not emerge as a replicated signal. ([pmc.ncbi.nlm.nih.gov](https://pmc.ncbi.nlm.nih.gov/articles/PMC11541208/)) [S1]

## Confidence

Medium

The confidence is not high only because “implicated” is a broad claim and a very small or obscure study could exist. But across recent high-level reviews and major SUD genomic papers, I found no direct WSB1-addiction evidence. ([pmc.ncbi.nlm.nih.gov](https://pmc.ncbi.nlm.nih.gov/articles/PMC11541208/)) [S2]

## Supporting evidence

The best arguments **for plausibility** are indirect and neurobiological rather than addiction-specific:

- **WSB1 is expressed in the brain.** A review of WSB1 biology notes that its transcripts are highly expressed in brain among other tissues, and Human Protein Atlas data show WSB1 RNA is detected broadly across major human brain regions, with low regional specificity. ([pmc.ncbi.nlm.nih.gov](https://pmc.ncbi.nlm.nih.gov/articles/PMC4992216/)) [S3]
- **WSB1 has bona fide neural functions.** In brain, WSB1 participates in ubiquitination of **type 2 iodothyronine deiodinase (D2)**, a regulator of local thyroid hormone activation; astrocyte-specific Wsb1 deletion in mice altered hypothalamic D2 ubiquitination and thyroid-hormone sensitivity. ([pmc.ncbi.nlm.nih.gov](https://pmc.ncbi.nlm.nih.gov/articles/PMC4992216/)) [S4]
- **WSB1 can affect neuronal survival/proteostasis in a disease model.** In a Parkinson’s disease model, WSB1 interacted with LRRK2, promoted its K27/K29-linked ubiquitination and aggregation, and reduced mutant-LRRK2 neuronal toxicity in primary neurons and Drosophila; the paper also reported effects on dopaminergic neuron loss and motor phenotype in flies. ([pubmed.ncbi.nlm.nih.gov](https://pubmed.ncbi.nlm.nih.gov/27273569/)) [S5]
- **Some WSB1-linked pathways overlap conceptually with addiction biology.** WSB1 has been linked in non-addiction literature to NF-κB termination and GSK3β ubiquitination, while addiction literature does implicate NF-κB and GSK3β-related signaling in drug-related plasticity and drug-seeking behaviors. That creates a mechanistic rationale for investigation, but not evidence of actual addiction involvement by WSB1 itself. ([academic.oup.com](https://academic.oup.com/nar/article-abstract/52/9/4969/7624079?utm_source=openai)) [S6]

So, WSB1 is **neurobiologically relevant**, but that is not the same as being implicated in addiction.

## Contradicting or limiting evidence

Several broad, addiction-focused sources argue against the claim being currently supported:

- A 2024 expert review, **“The genetic landscape of substance use disorders,”** summarizes robust genetic findings across AUD, OUD, CUD, and TUD and discusses replicated loci and polygenic findings; **WSB1 is not mentioned** in the text. ([pmc.ncbi.nlm.nih.gov](https://pmc.ncbi.nlm.nih.gov/articles/PMC11541208/)) [S7]
- A 2022 systematic review integrating **75 GWAS studies and 17 brain RNA-seq studies** for substance use behavior/disorder identified shared genes such as **CACNB2, GRIN2B, PLXDC2, and PKNOX2**; **WSB1 is not found in the article text**. ([pubmed.ncbi.nlm.nih.gov](https://pubmed.ncbi.nlm.nih.gov/35491750/)) [S8]
- A 2023 multivariate GWAS meta-analysis of **over 1 million individuals** identified 19 independent SNPs for a general addiction-risk factor and highlighted genes such as **PDE4B** and dopamine-regulation biology; WSB1 was not a highlighted locus. ([pubmed.ncbi.nlm.nih.gov](https://pubmed.ncbi.nlm.nih.gov/37250466/?utm_source=openai)) [S9]
- A 2025 multi-omic opioid-addiction analysis integrating GWAS, RNA-seq, proteomic, methylation, and chromatin data identified **404 opioid-addiction genes** and a core network of **211 interrelated genes** involving Akt, BDNF, and ERK pathways; **WSB1 is not found in the article text**. ([sciencedirect.com](https://www.sciencedirect.com/science/article/pii/S0006322324017815)) [S10]
- A 2021 systematic review of **methamphetamine use disorder genetics** evaluated 76 studies covering 75 genes and highlighted signals such as **FAAH** and **BDNF**; **WSB1 is not found in the article text**. ([pubmed.ncbi.nlm.nih.gov](https://pubmed.ncbi.nlm.nih.gov/33217458/)) [S11]
- Likewise, an older review of proteomics across multiple drugs of abuse compiled **497 proteins** associated with exposure to addictive drugs, and **WSB1 is not found in that review text**. ([pubmed.ncbi.nlm.nih.gov](https://pubmed.ncbi.nlm.nih.gov/21922273/)) [S12]

This does **not** prove WSB1 has no role. But it does mean there is no clear replicated human-genetic, brain-omics, or functional-addiction signal placing WSB1 among recognized addiction genes at present. ([pmc.ncbi.nlm.nih.gov](https://pmc.ncbi.nlm.nih.gov/articles/PMC11541208/)) [S13]

## Mechanistic / biological rationale

If one wanted to hypothesize a WSB1-addiction link, the rationale would be **indirect**:

1. **Protein homeostasis / ubiquitination:** addiction involves durable synaptic and transcriptional remodeling, and ubiquitin-mediated proteostasis is relevant to neural plasticity. WSB1 is an E3-ligase adaptor with established substrate-directed ubiquitination roles. ([pmc.ncbi.nlm.nih.gov](https://pmc.ncbi.nlm.nih.gov/articles/PMC4992216/)) [S14]  
2. **Brain hormone regulation:** WSB1 controls ubiquitination of D2 deiodinase, affecting local T3 generation in brain regions such as the hypothalamus. Thyroid hormone signaling can influence neural development and function, though a direct addiction-specific WSB1 pathway has not been shown. ([pmc.ncbi.nlm.nih.gov](https://pmc.ncbi.nlm.nih.gov/articles/PMC4992216/)) [S15]  
3. **Neuroimmune / kinase pathways:** WSB1 has been implicated in NF-κB regulation and GSK3β biology outside addiction models, while NF-κB and GSK3β are implicated in addiction-related neuroplasticity and drug-seeking in separate literature. This is a **pathway-overlap argument**, not direct evidence. ([academic.oup.com](https://academic.oup.com/nar/article-abstract/52/9/4969/7624079?utm_source=openai)) [S16]  
4. **Dopaminergic neuron relevance:** WSB1 modulates LRRK2 toxicity in dopaminergic systems in Parkinson models, which makes it neuronally relevant, but Parkinson dopaminergic neurodegeneration is not the same phenomenon as addiction circuitry remodeling. ([pubmed.ncbi.nlm.nih.gov](https://pubmed.ncbi.nlm.nih.gov/27273569/)) [S17]

Overall, the biology makes WSB1 **worth exploring**, but current evidence does not justify saying it is already implicated in addiction.

## Bottom line

At present, the claim that **WSB1 gene/protein is implicated in addiction** is **not supported by direct evidence**. WSB1 is a real brain-expressed ubiquitination-related protein with plausible links to neuronal signaling and proteostasis, but I could not find replicated human genetic associations, addiction-focused transcriptomic/proteomic signals, or causal addiction-model experiments that specifically implicate WSB1 in substance use disorder. ([proteinatlas.org](https://www.proteinatlas.org/ENSG00000109046-WSB1/brain)) [S18]

If you want, I can next do a **stricter deep-dive by evidence type**—for example:
- human GWAS only,
- postmortem brain omics only,
- animal addiction models only,
- or compare WSB1 against a known addiction gene to show the difference in evidence strength.

## Cited sources

- **[S1]** 
            The genetic landscape of substance use disorders - PMC
         — https://pmc.ncbi.nlm.nih.gov/articles/PMC11541208/
- **[S2]** 
            The genetic landscape of substance use disorders - PMC
         — https://pmc.ncbi.nlm.nih.gov/articles/PMC11541208/
- **[S3]** 
            WSB1: from homeostasis to hypoxia - PMC
         — https://pmc.ncbi.nlm.nih.gov/articles/PMC4992216/
- **[S4]** 
            WSB1: from homeostasis to hypoxia - PMC
         — https://pmc.ncbi.nlm.nih.gov/articles/PMC4992216/
- **[S5]** Ubiqutination via K27 and K29 chains signals aggregation and neuronal protection of LRRK2 by WSB1 - PubMed — https://pubmed.ncbi.nlm.nih.gov/27273569/
- **[S6]** WSB1/2 target chromatin-bound lysine-methylated RelA for proteasomal degradation and NF-κB termination | Nucleic Acids Research | Oxford Academic — https://academic.oup.com/nar/article-abstract/52/9/4969/7624079?utm_source=openai
- **[S7]** 
            The genetic landscape of substance use disorders - PMC
         — https://pmc.ncbi.nlm.nih.gov/articles/PMC11541208/
- **[S8]** Common and distinguishing genetic factors for substance use behavior and disorder: an integrated analysis of genomic and transcriptomic studies from both human and animal studies - PubMed — https://pubmed.ncbi.nlm.nih.gov/35491750/
- **[S9]** Multivariate genome-wide association meta-analysis of over 1 million subjects identifies loci underlying multiple substance use disorders - PubMed — https://pubmed.ncbi.nlm.nih.gov/37250466/?utm_source=openai
- **[S10]** Multiomic Network Analysis Identifies Dysregulated Neurobiological Pathways in Opioid Addiction - ScienceDirect — https://www.sciencedirect.com/science/article/pii/S0006322324017815
- **[S11]** Genetics of methamphetamine use disorder: A systematic review and meta-analyses of gene association studies - PubMed — https://pubmed.ncbi.nlm.nih.gov/33217458/
- **[S12]** Genes and pathways co-associated with the exposure to multiple drugs of abuse, including alcohol, amphetamine/methamphetamine, cocaine, marijuana, morphine, and/or nicotine: a review of proteomics analyses - PubMed — https://pubmed.ncbi.nlm.nih.gov/21922273/
- **[S13]** 
            The genetic landscape of substance use disorders - PMC
         — https://pmc.ncbi.nlm.nih.gov/articles/PMC11541208/
- **[S14]** 
            WSB1: from homeostasis to hypoxia - PMC
         — https://pmc.ncbi.nlm.nih.gov/articles/PMC4992216/
- **[S15]** 
            WSB1: from homeostasis to hypoxia - PMC
         — https://pmc.ncbi.nlm.nih.gov/articles/PMC4992216/
- **[S16]** WSB1/2 target chromatin-bound lysine-methylated RelA for proteasomal degradation and NF-κB termination | Nucleic Acids Research | Oxford Academic — https://academic.oup.com/nar/article-abstract/52/9/4969/7624079?utm_source=openai
- **[S17]** Ubiqutination via K27 and K29 chains signals aggregation and neuronal protection of LRRK2 by WSB1 - PubMed — https://pubmed.ncbi.nlm.nih.gov/27273569/
- **[S18]** Brain tissue expression of WSB1 - Summary - The Human Protein Atlas — https://www.proteinatlas.org/ENSG00000109046-WSB1/brain
