## Supplementary material for "Understanding Substance Dependence: What Differentiates Addictive from Non-Addictive Drugs?": Folder for supplementary files: addictive-genes-claims-first100-084-far1-gene-protein-is-implicated-in-addiction-13130fbe53e0.md

# addictive-genes-claims-first100-084-13130fbe53e0

**Original claim:** FAR1 gene/protein is implicated in addiction.

## Claim

FAR1 gene/protein is implicated in addiction.

## Verdict

Verdict: No Evidence

Current published evidence does **not** establish FAR1 as an addiction gene/protein. FAR1 is clearly important for ether lipid/plasmalogen biology and for brain development/homeostasis, but I did not find convincing human genetic, human transcriptomic, animal behavioral, or mechanistic studies directly linking **FAR1 itself** to substance addiction phenotypes such as drug self-administration, craving, relapse, cue-reactivity, or diagnosed substance use disorder. ([pubmed.ncbi.nlm.nih.gov](https://pubmed.ncbi.nlm.nih.gov/24108123/)) [S1]

## Confidence

Medium

The confidence is **medium** because the FAR1 literature is substantial for lipid biology and neurological disease, and the addiction literature is substantial for other genes, yet proving a total absence of evidence is inherently harder than confirming a positive association. Still, based on the major peer-reviewed sources I found, FAR1 is **not an established addiction-linked gene**. ([pubmed.ncbi.nlm.nih.gov](https://pubmed.ncbi.nlm.nih.gov/36781603/)) [S2]

## Supporting evidence

FAR1 encodes fatty acyl-CoA reductase 1, a peroxisomal membrane protein that generates fatty alcohols required for ether lipid/plasmalogen synthesis; experiments in mammalian cells indicate FAR1 is a **rate-limiting enzyme** in this pathway and is regulated by plasmalogen-dependent feedback. ([pubmed.ncbi.nlm.nih.gov](https://pubmed.ncbi.nlm.nih.gov/24108123/)) [S3]

That biology matters to the nervous system. FAR1 deficiency in humans causes a peroxisomal disorder with severe neurological manifestations, including intellectual disability, epilepsy, cataracts, spasticity, and white-matter abnormalities; newer case series broaden the phenotype to additional neurological syndromes from FAR1 variants. In mice, Far1 knockout reduces brain ether lipids/plasmalogens and causes hypomyelination, supporting a real role for FAR1 in brain structure and function. ([pubmed.ncbi.nlm.nih.gov](https://pubmed.ncbi.nlm.nih.gov/25439727/)) [S4]

There is also indirect pathway-level evidence that ether lipid deficiency can alter neural systems relevant to psychiatric phenotypes. In a mouse model of **global ether lipid deficiency** (Gnpat knockout, not FAR1 knockout), investigators reported disturbed neurotransmitter homeostasis, altered neurotransmitter turnover, hyperactivity, impaired social interaction, and broader behavioral abnormalities. A review on plasmalogens in the brain likewise summarizes links between plasmalogen deficiency and disturbed synaptic neurotransmission, intracellular signaling, myelin abnormalities, neuroinflammation, cognitive changes, and aberrant behaviors. ([pubmed.ncbi.nlm.nih.gov](https://pubmed.ncbi.nlm.nih.gov/30759250/?utm_source=openai)) [S5]

A recent human psychiatry paper found that **peripheral blood FAR1 expression** was positively associated with internalizing symptoms in youths, but not with externalizing symptoms or addiction phenotypes. This suggests FAR1 may be relevant to some aspects of psychopathology, but it is still far from evidence for addiction. ([pubmed.ncbi.nlm.nih.gov](https://pubmed.ncbi.nlm.nih.gov/40018685/)) [S6]

## Contradicting or limiting evidence

The strongest limitation is the absence of **direct addiction-specific evidence**. Recent reviews of substance use disorder genetics summarize replicated loci and pathways across alcohol, tobacco, cannabis, and opioid use disorders, but FAR1 is not presented as an established signal. In opioid addiction specifically, recent multi-omic reviews emphasize loci such as **OPRM1, FURIN, and PPP6C/SCAI/RABEPK**, not FAR1. From this omission in current state-of-the-art summaries, I infer that FAR1 is **not currently a replicated or widely recognized addiction gene**. ([pubmed.ncbi.nlm.nih.gov](https://pubmed.ncbi.nlm.nih.gov/38811691/)) [S7]

Likewise, very large association studies of tobacco and alcohol use have identified many loci, but the prominent signals discussed are not FAR1. This does not prove FAR1 has no effect whatsoever, but it argues against FAR1 being a major or well-supported addiction locus at present. ([pubmed.ncbi.nlm.nih.gov](https://pubmed.ncbi.nlm.nih.gov/30643251/?utm_source=openai)) [S8]

The FAR1 disease literature is centered on **peroxisomal lipid metabolism and neurological developmental disorders**, not addiction biology. Human FAR1 papers focus on plasmalogen synthesis defects, epilepsy, cataracts, spasticity, white-matter disease, and movement/neurologic phenotypes. ([pubmed.ncbi.nlm.nih.gov](https://pubmed.ncbi.nlm.nih.gov/25439727/)) [S9]

Even the more suggestive pathway studies are indirect: the neurotransmitter and behavior findings were observed in **ether lipid-deficient models upstream of addiction**, not in FAR1-specific addiction paradigms. Hyperactivity, impaired social behavior, or altered monoamines are not sufficient to claim implication in addiction unless they are tied to addiction-relevant outcomes. ([pubmed.ncbi.nlm.nih.gov](https://pubmed.ncbi.nlm.nih.gov/30759250/?utm_source=openai)) [S10]

## Mechanistic / biological rationale

A cautious biological rationale **can** be made. FAR1 controls production of fatty alcohols used for ether lipids/plasmalogens, and plasmalogens are abundant brain phospholipids with roles in membrane organization, vesicle fusion, myelination, oxidative balance, and synaptic function. Because addiction depends on neural homeostasis and synaptic plasticity in reward-related circuits, disruption of FAR1-dependent lipid homeostasis could, in principle, influence processes relevant to addiction vulnerability. ([pubmed.ncbi.nlm.nih.gov](https://pubmed.ncbi.nlm.nih.gov/24108123/)) [S11]

That plausibility is strengthened by pathway data showing that ether lipid deficiency can disturb monoamine-related neurochemistry and behavior. Since dopamine and related signaling are central to addiction, one can reasonably hypothesize that severe disturbances in plasmalogen biology might modulate reward circuitry. But this remains a **hypothesis**, not demonstrated evidence for FAR1 in addiction. The available studies do not show that FAR1 variants or FAR1 manipulation alter drug reward, reinforcement, escalation, withdrawal, relapse, or human SUD risk. ([pubmed.ncbi.nlm.nih.gov](https://pubmed.ncbi.nlm.nih.gov/30759250/?utm_source=openai)) [S12]

So biologically, FAR1 looks more like a **general neural lipid-homeostasis gene** with possible indirect relevance to psychiatric/neurobehavioral function than a gene specifically implicated in addiction. ([pubmed.ncbi.nlm.nih.gov](https://pubmed.ncbi.nlm.nih.gov/24108123/)) [S13]

## Bottom line

FAR1 is well supported as a key enzyme in ether lipid/plasmalogen synthesis and is clearly relevant to brain development, myelination, and neural homeostasis. However, based on current peer-reviewed and authoritative sources, there is **no direct published evidence** that FAR1 is implicated in addiction per se. At most, there is an **indirect, biologically plausible pathway-level link** through lipid-dependent neurotransmission and brain membrane biology. ([pubmed.ncbi.nlm.nih.gov](https://pubmed.ncbi.nlm.nih.gov/24108123/)) [S14]

## Cited sources

- **[S1]** Topogenesis and homeostasis of fatty acyl-CoA reductase 1 - PubMed — https://pubmed.ncbi.nlm.nih.gov/24108123/
- **[S2]** Spectrum of FAR1 (Fatty Acyl-CoA Reductase 1) Variants and Related Neurological Conditions - PubMed — https://pubmed.ncbi.nlm.nih.gov/36781603/
- **[S3]** Topogenesis and homeostasis of fatty acyl-CoA reductase 1 - PubMed — https://pubmed.ncbi.nlm.nih.gov/24108123/
- **[S4]** A peroxisomal disorder of severe intellectual disability, epilepsy, and cataracts due to fatty acyl-CoA reductase 1 deficiency - PubMed — https://pubmed.ncbi.nlm.nih.gov/25439727/
- **[S5]** Disturbed neurotransmitter homeostasis in ether lipid deficiency - PubMed — https://pubmed.ncbi.nlm.nih.gov/30759250/?utm_source=openai
- **[S6]** Impact of life adversity and gene expression on psychiatric symptoms in children and adolescents: findings from the Brazilian high risk cohort study - PubMed — https://pubmed.ncbi.nlm.nih.gov/40018685/
- **[S7]** The genetic landscape of substance use disorders - PubMed — https://pubmed.ncbi.nlm.nih.gov/38811691/
- **[S8]** Association studies of up to 1.2 million individuals yield new insights into the genetic etiology of tobacco and alcohol use - PubMed — https://pubmed.ncbi.nlm.nih.gov/30643251/?utm_source=openai
- **[S9]** A peroxisomal disorder of severe intellectual disability, epilepsy, and cataracts due to fatty acyl-CoA reductase 1 deficiency - PubMed — https://pubmed.ncbi.nlm.nih.gov/25439727/
- **[S10]** Disturbed neurotransmitter homeostasis in ether lipid deficiency - PubMed — https://pubmed.ncbi.nlm.nih.gov/30759250/?utm_source=openai
- **[S11]** Topogenesis and homeostasis of fatty acyl-CoA reductase 1 - PubMed — https://pubmed.ncbi.nlm.nih.gov/24108123/
- **[S12]** Disturbed neurotransmitter homeostasis in ether lipid deficiency - PubMed — https://pubmed.ncbi.nlm.nih.gov/30759250/?utm_source=openai
- **[S13]** Topogenesis and homeostasis of fatty acyl-CoA reductase 1 - PubMed — https://pubmed.ncbi.nlm.nih.gov/24108123/
- **[S14]** Topogenesis and homeostasis of fatty acyl-CoA reductase 1 - PubMed — https://pubmed.ncbi.nlm.nih.gov/24108123/
