## Supplementary material for "Understanding Substance Dependence: What Differentiates Addictive from Non-Addictive Drugs?": Folder for supplementary files: addictive-genes-claims-first100-085-gabrq-gene-protein-is-implicated-in-addiction-f6c345d91bcb.md

# addictive-genes-claims-first100-085-f6c345d91bcb

**Original claim:** GABRQ gene/protein is implicated in addiction.

## Claim

**“GABRQ gene/protein is implicated in addiction.”**

## Verdict

Verdict: Mixed Evidence

## Confidence

**Medium**

## Supporting evidence

GABRQ encodes the **theta (θ) subunit of the GABA\_A receptor**, a ligand-gated chloride channel involved in fast inhibitory neurotransmission. Because GABAergic signaling is centrally involved in alcohol and other substance-use phenotypes, a priori it is biologically plausible that specific GABA\_A subunits could influence addiction-related processes. GABRQ is a bona fide brain-expressed GABA\_A receptor gene, and experimental work shows that θ-containing receptors have distinct pharmacology from more common receptor assemblies. ([ncbi.nlm.nih.gov](https://www.ncbi.nlm.nih.gov/gene/55879?utm_source=openai)) [S1]

A notable mechanistic clue is that **GABRQ/the θ subunit is enriched in the locus coeruleus (LC)** in rat brain, alongside α3/β1-containing GABA\_A receptor assemblies. The LC is a brainstem noradrenergic nucleus with established roles in arousal, stress responses, and especially opioid withdrawal physiology. Recombinant studies indicate that θ-containing α3β1 receptors respond differently to anesthetics and modulators than canonical γ2-containing receptors, suggesting that GABRQ can meaningfully alter inhibitory signaling in a circuit that is relevant to dependence/withdrawal states. ([pmc.ncbi.nlm.nih.gov](https://pmc.ncbi.nlm.nih.gov/articles/PMC1363348/)) [S2]

There is also **some preclinical transcriptomic evidence** linking Gabrq expression to addiction-related states. In a mouse fentanyl-exposure model, Gabrq was among the differentially expressed genes in prefrontal cortex and was listed within addiction-related pathway analyses, with expression reduced in fentanyl-exposed animals relative to controls. The authors interpreted these broader transcriptional changes as potentially contributing to fentanyl addiction-like behavior and withdrawal-related neuroadaptation. ([pmc.ncbi.nlm.nih.gov](https://pmc.ncbi.nlm.nih.gov/articles/PMC11630728/)) [S3]

Similarly, in a rat model of **alcohol withdrawal**, one study reported that Gabrq showed delayed expression dynamics during the withdrawal time course in the dorsal vagal complex, a homeostatic brainstem region. That does not establish causality for addiction, but it does suggest that Gabrq can be transcriptionally responsive during dependence/withdrawal-related neuroadaptation. ([pmc.ncbi.nlm.nih.gov](https://pmc.ncbi.nlm.nih.gov/articles/PMC4419739/?utm_source=openai)) [S4]

Taken together, these data support a **limited, mostly indirect implication**: GABRQ sits in the right neurotransmitter system, in at least some relevant brain circuits, and shows state-related expression changes in a few animal models of drug exposure or withdrawal. ([pmc.ncbi.nlm.nih.gov](https://pmc.ncbi.nlm.nih.gov/articles/PMC1363348/)) [S5]

## Contradicting or limiting evidence

The main limitation is that **direct human evidence is weak**. I did not find robust, replicated human genetic association studies showing that GABRQ variants materially increase risk for alcohol, nicotine, opioid, cocaine, or other substance use disorders. By contrast, reviews of GABAergic genetics in nicotine/alcohol dependence emphasize other genes—especially **GABRA2, GABRG1, GABBR2, and related loci**—rather than GABRQ. ([pubmed.ncbi.nlm.nih.gov](https://pubmed.ncbi.nlm.nih.gov/22048727/)) [S6]

A particularly important negative result comes from a **large community-based study of GABA-system genes** and alcohol-related phenotypes. That analysis explicitly included **GABRQ** (with 2 SNPs on chromosome X) among 31 GABA-related genes and found **no significant association** of GABA-system polymorphisms with alcohol use or alcohol abuse/dependence symptom measures by single-SNP, polygenic, variance-component, or gene-based analyses. ([pmc.ncbi.nlm.nih.gov](https://pmc.ncbi.nlm.nih.gov/articles/PMC3984361/)) [S7]

Postmortem human brain data are also not very supportive for a strong role. In a study of hippocampal GABAergic gene expression from **alcoholics, cocaine addicts, and controls**, GABRQ expression did **not** differ significantly by diagnosis. A follow-up factor-analysis paper similarly grouped GABRQ with low-cerebellar-expression genes that appeared **unresponsive to chronic alcohol/cocaine exposure** in that dataset. ([pmc.ncbi.nlm.nih.gov](https://pmc.ncbi.nlm.nih.gov/articles/PMC3661725/)) [S8]

More broadly, the addiction literature strongly implicates the **GABA system**, but that is not the same as implicating **GABRQ specifically**. For alcohol and nicotine dependence genetics, the strongest and most replicated GABA\_A findings have centered on other receptor subunits, especially **GABRA2/GABRG1**. That weakens any attempt to generalize from “GABA is involved in addiction” to “GABRQ is implicated in addiction.” ([pubmed.ncbi.nlm.nih.gov](https://pubmed.ncbi.nlm.nih.gov/22048727/)) [S9]

Finally, some of the “supporting” evidence is **indirect or low-resolution**: transcriptomic shifts after drug exposure can reflect downstream adaptation rather than causal contribution, and pathway enrichment tables that include Gabrq do not prove that the gene is a driver of addictive behavior. ([pmc.ncbi.nlm.nih.gov](https://pmc.ncbi.nlm.nih.gov/articles/PMC11630728/)) [S10]

## Mechanistic / biological rationale

Biologically, the claim is **plausible but not proven**. GABRQ encodes a GABA\_A receptor subunit, and GABA\_A receptors are major regulators of neuronal excitability, reward-circuit inhibition, stress responsiveness, and withdrawal physiology. θ-containing receptors are enriched in specialized nuclei such as the **locus coeruleus**, where altered inhibitory tone could in principle influence noradrenergic output, autonomic arousal, anxiety, and withdrawal-related symptoms—all relevant to relapse vulnerability. ([pmc.ncbi.nlm.nih.gov](https://pmc.ncbi.nlm.nih.gov/articles/PMC1363348/)) [S11]

The pharmacology is also suggestive. θ-containing α3β1 receptors differ from standard GABA\_A receptor assemblies in sensitivity to modulators such as **etomidate, propofol, pregnanolone, and pentobarbital-related modulation patterns**, meaning that incorporation of θ can change how inhibitory synapses respond to endogenous neurosteroids and drugs. Because alcohol and several drugs of abuse interact with GABAergic signaling directly or indirectly, altered GABRQ expression could theoretically shift circuit homeostasis during intoxication, tolerance, or withdrawal. ([pmc.ncbi.nlm.nih.gov](https://pmc.ncbi.nlm.nih.gov/articles/PMC1363348/)) [S12]

Still, that rationale remains mostly **inferential**. The strongest mechanistic models in addiction currently involve other GABA receptor subunits and other neurotransmitter systems with better human and causal experimental support. For GABRQ, the mechanistic story is best viewed as a **hypothesis-generating framework**, not established evidence of disease involvement. ([pubmed.ncbi.nlm.nih.gov](https://pubmed.ncbi.nlm.nih.gov/22048727/)) [S13]

## Bottom line

GABRQ is **not a well-established addiction gene/protein** in the way that some other GABA-related genes are. There is **some indirect preclinical evidence**—brain-region expression in withdrawal-relevant circuitry, transcriptomic changes in alcohol withdrawal or fentanyl exposure, and a plausible role in inhibitory homeostasis—but **human genetic and postmortem evidence is sparse and largely non-supportive**, with at least one sizable study finding no alcohol-related association. ([pmc.ncbi.nlm.nih.gov](https://pmc.ncbi.nlm.nih.gov/articles/PMC1363348/)) [S14]

So the fairest reading is: **GABRQ may be a plausible, understudied candidate in addiction biology, but current evidence does not show a strong or replicated implication in addiction.** ([pubmed.ncbi.nlm.nih.gov](https://pubmed.ncbi.nlm.nih.gov/22048727/)) [S15]

## Cited sources

- **[S1]** GABRQ gamma-aminobutyric acid type A receptor subunit theta [Homo sapiens (human)] - Gene - NCBI — https://www.ncbi.nlm.nih.gov/gene/55879?utm_source=openai
- **[S2]** 
            Impact of ε and θ subunits on pharmacological properties of α3β1 GABAA receptors expressed in Xenopus oocytes - PMC
         — https://pmc.ncbi.nlm.nih.gov/articles/PMC1363348/
- **[S3]** 
            Synaptic Structure and Transcriptomic Profiling of Reward and Sensory Brain Areas in Male Mice of Fentanyl Addiction - PMC
         — https://pmc.ncbi.nlm.nih.gov/articles/PMC11630728/
- **[S4]** Rapid temporal changes in the expression of a set of neuromodulatory genes during alcohol withdrawal in the dorsal vagal complex: molecular evidence of homeostatic disturbance - PMC — https://pmc.ncbi.nlm.nih.gov/articles/PMC4419739/?utm_source=openai
- **[S5]** 
            Impact of ε and θ subunits on pharmacological properties of α3β1 GABAA receptors expressed in Xenopus oocytes - PMC
         — https://pmc.ncbi.nlm.nih.gov/articles/PMC1363348/
- **[S6]** Genetics of GABAergic signaling in nicotine and alcohol dependence - PubMed — https://pubmed.ncbi.nlm.nih.gov/22048727/
- **[S7]** 
            GABA System Genes – No Evidence for a Role in Alcohol Use and Abuse in a Community-based Sample - PMC
         — https://pmc.ncbi.nlm.nih.gov/articles/PMC3984361/
- **[S8]** 
            A Factor Analysis of Global GABAergic Gene Expression in Human Brain Identifies Specificity in Response to Chronic Alcohol and Cocaine Exposure - PMC
         — https://pmc.ncbi.nlm.nih.gov/articles/PMC3661725/
- **[S9]** Genetics of GABAergic signaling in nicotine and alcohol dependence - PubMed — https://pubmed.ncbi.nlm.nih.gov/22048727/
- **[S10]** 
            Synaptic Structure and Transcriptomic Profiling of Reward and Sensory Brain Areas in Male Mice of Fentanyl Addiction - PMC
         — https://pmc.ncbi.nlm.nih.gov/articles/PMC11630728/
- **[S11]** 
            Impact of ε and θ subunits on pharmacological properties of α3β1 GABAA receptors expressed in Xenopus oocytes - PMC
         — https://pmc.ncbi.nlm.nih.gov/articles/PMC1363348/
- **[S12]** 
            Impact of ε and θ subunits on pharmacological properties of α3β1 GABAA receptors expressed in Xenopus oocytes - PMC
         — https://pmc.ncbi.nlm.nih.gov/articles/PMC1363348/
- **[S13]** Genetics of GABAergic signaling in nicotine and alcohol dependence - PubMed — https://pubmed.ncbi.nlm.nih.gov/22048727/
- **[S14]** 
            Impact of ε and θ subunits on pharmacological properties of α3β1 GABAA receptors expressed in Xenopus oocytes - PMC
         — https://pmc.ncbi.nlm.nih.gov/articles/PMC1363348/
- **[S15]** Genetics of GABAergic signaling in nicotine and alcohol dependence - PubMed — https://pubmed.ncbi.nlm.nih.gov/22048727/
