## Supplementary material for "Understanding Substance Dependence: What Differentiates Addictive from Non-Addictive Drugs?": Folder for supplementary files: addictive-genes-claims-first100-086-gabra6-gene-protein-is-implicated-in-addiction-4596ae9d91dd.md

# addictive-genes-claims-first100-086-4596ae9d91dd

**Original claim:** GABRA6 gene/protein is implicated in addiction.

## Claim

**“GABRA6 gene/protein is implicated in addiction.”**

## Verdict

Verdict: Mixed Evidence

## Confidence

**Medium.**

## Supporting evidence

GABRA6 encodes the **GABA\_A receptor α6 subunit**, which is highly enriched in **cerebellar granule cells** and is a key component of certain **extrasynaptic GABA\_A receptors** involved in tonic inhibition. This makes it biologically plausible that GABRA6 variation could influence alcohol-related phenotypes, especially intoxication sensitivity and perhaps downstream addiction risk. ([pubmed.ncbi.nlm.nih.gov](https://pubmed.ncbi.nlm.nih.gov/35017178/)) [S1]

Several **candidate-gene association studies** reported links between GABRA6 variants and **alcohol dependence/alcoholism**. A Scottish case-control study examined markers in the chromosome 5 GABA\_A receptor cluster, including GABRA6, in 108 alcohol-dependent subjects and 54 controls. ([pubmed.ncbi.nlm.nih.gov](https://pubmed.ncbi.nlm.nih.gov/10578235/)) [S2] A Korean study of 162 hospitalized male alcoholics and 172 controls reported a significant association between polymorphisms in **GABRA1 and GABRA6** and alcoholism. ([pubmed.ncbi.nlm.nih.gov](https://pubmed.ncbi.nlm.nih.gov/16778401/)) [S3] A Taiwanese Han study also tested GABRA6 variants within the same receptor-gene cluster for association with alcohol dependence. ([nature.com](https://www.nature.com/articles/4001110?utm_source=openai)) [S4]

A more informative source is a **2014 meta-analysis** of candidate-gene studies across GABA receptor genes. For **GABRA6 rs3219151**, the authors reported that, in the subset of **alcohol dependence** studies, the combined evidence supported association with **OR 1.34 (95% CI 1.17–1.54), P = 4×10⁻⁵**, without significant heterogeneity. In the paper’s summary, they described **significant association between GABRA6 and alcohol dependence** based on the combined candidate-gene literature. ([pmc.ncbi.nlm.nih.gov](https://pmc.ncbi.nlm.nih.gov/articles/PMC3924525/)) [S5]

There is also evidence that GABRA6 may influence **alcohol-response endophenotypes**, which are relevant because lower sensitivity to alcohol’s sedative/impairing effects is a recognized risk marker for later alcohol problems. A critical review of alcohol-sensitivity genetics summarized that oral alcohol challenge studies linked **GABRA6 Pro385Ser** to a **lower level of response** to alcohol. ([pmc.ncbi.nlm.nih.gov](https://pmc.ncbi.nlm.nih.gov/articles/PMC5916326/)) [S6] Reviews focused on GABAergic genetics in dependence similarly state that GABRA6 Pro385Ser has been associated with both **alcohol dependence** and **reduced sedative response to alcohol**. ([pmc.ncbi.nlm.nih.gov](https://pmc.ncbi.nlm.nih.gov/articles/PMC3746562/)) [S7]

Human molecular data are limited but not absent. In postmortem cortex from people with alcohol use disorder (AUD), one study found that the **rs3219151 genotype mediated GABRA6 transcript expression**, with sex-dependent effects, supporting possible functional consequences of this variant in AUD-related biology. ([pmc.ncbi.nlm.nih.gov](https://pmc.ncbi.nlm.nih.gov/articles/PMC9744570/)) [S8]

## Contradicting or limiting evidence

The evidence is **not consistently replicated**, and it is notably weaker than for some other addiction-related GABA genes such as **GABRA2**. In a family-based COGA study, **GABRA6 did not provide evidence for association with alcoholism**. ([pubmed.ncbi.nlm.nih.gov](https://pubmed.ncbi.nlm.nih.gov/12555233/)) [S9] A later community-based study examining many GABA-system markers likewise found **no association** between GABA-system variation and alcohol use/abuse phenotypes in that more general-population sample. ([pmc.ncbi.nlm.nih.gov](https://pmc.ncbi.nlm.nih.gov/articles/PMC3984361/)) [S10]

Even within the 2014 meta-analysis, the GABRA6 signal looks **fragile rather than definitive**. When all studies across substance-dependence phenotypes were combined, there was **heterogeneity**, and no significant association under the random-effects model; the stronger signal was mainly in the **alcohol dependence subset**, not in addiction broadly. The same paper emphasized stronger and more reproducible evidence for **GABRA2** and **GABRG2** than for GABRA6. ([pmc.ncbi.nlm.nih.gov](https://pmc.ncbi.nlm.nih.gov/articles/PMC3924525/)) [S11]

Crucially, **large modern GWAS of AUD** have not highlighted GABRA6 as a major common-risk locus. For example, a Han Chinese alcohol-dependence GWAS found significant signals linked to **ADH1B/ALDH2-related loci**, not GABRA6. An East Asian AUD meta-analysis likewise detected only two risk loci, again at **ADH1B** and near **ALDH2**, without GABRA6 emerging as a genome-wide hit. ([pubmed.ncbi.nlm.nih.gov](https://pubmed.ncbi.nlm.nih.gov/31591379/)) [S12] This does not prove GABRA6 is irrelevant, but it argues against GABRA6 being a **strong, broadly replicated common variant contributor** to addiction risk. ([pubmed.ncbi.nlm.nih.gov](https://pubmed.ncbi.nlm.nih.gov/35094024/)) [S13]

The claim is also too broad if “addiction” means **multiple substance-use disorders**. The best human evidence for GABRA6 is concentrated in **alcohol-related phenotypes**. By contrast, in the same meta-analysis, broader cross-substance evidence was stronger for other genes; for GABRA6, the main reproducible signal was **AUD/alcohol dependence**, not heroin, methamphetamine, nicotine, or cocaine dependence in general. ([pmc.ncbi.nlm.nih.gov](https://pmc.ncbi.nlm.nih.gov/articles/PMC3924525/)) [S14]

Finally, expression/pathology data do not show a simple consistent picture. In human cerebellum from AUD subjects, one epigenetic study found changes in the **δ subunit (GABRD)** but reported **no changes in α6-containing GABA\_A receptor subunits**, limiting the case that GABRA6 itself is a robustly altered downstream effector in chronic AUD brains. ([pubmed.ncbi.nlm.nih.gov](https://pubmed.ncbi.nlm.nih.gov/29020412/)) [S15]

## Mechanistic / biological rationale

Mechanistically, GABRA6 is relevant mainly because α6-containing GABA\_A receptors participate in **tonic inhibitory signaling** and are unusually important in the **cerebellum**, a region strongly affected by ethanol. Human brain-expression analyses show GABRA6 is **highly expressed in cerebellar cortex** and much less elsewhere. ([pmc.ncbi.nlm.nih.gov](https://pmc.ncbi.nlm.nih.gov/articles/PMC3661725/)) [S16]

Preclinical work gives a plausible alcohol-related mechanism. In rats, a naturally occurring **Gabra6 R100Q** variant altered the sensitivity of **α6β3δ extrasynaptic GABA\_A receptors** to ethanol, and animals carrying the more sensitive form showed greater **alcohol-induced motor impairment**. That supports a role for α6-containing receptors in acute behavioral responses to alcohol. ([pubmed.ncbi.nlm.nih.gov](https://pubmed.ncbi.nlm.nih.gov/15696164/)) [S17]

This is relevant to addiction biology because **initial response to alcohol** is an established intermediate phenotype: people with lower sensitivity to some alcohol effects may drink more to achieve the same subjective state, increasing risk over time. Reviews of human genetics of alcohol sensitivity specifically note GABRA6 as one of the loci linked to **lower alcohol response**. ([pmc.ncbi.nlm.nih.gov](https://pmc.ncbi.nlm.nih.gov/articles/PMC5916326/)) [S18]

Another indirect pathway is **stress responsivity**. A human experimental study found that a GABRA6 polymorphism was associated with altered **ACTH, cortisol, and blood-pressure responses** to psychosocial stress. Because stress responsivity is a recognized modifier of substance-use vulnerability and relapse risk, this offers another biologically plausible route by which GABRA6 could influence AUD risk. Still, this is **indirect evidence**, not proof of a direct causal role in addiction. ([pubmed.ncbi.nlm.nih.gov](https://pubmed.ncbi.nlm.nih.gov/15197399/)) [S19]

## Bottom line

GABRA6 is **plausibly involved in addiction-related biology**, but the evidence is **strongest for alcohol dependence/alcohol sensitivity**, not addiction broadly. There are multiple supportive candidate-gene studies and some mechanistic plausibility, including alcohol-response and stress-response effects, but the findings are **inconsistent**, often modest, and **not strongly reinforced by modern large-scale GWAS**. ([pmc.ncbi.nlm.nih.gov](https://pmc.ncbi.nlm.nih.gov/articles/PMC3924525/)) [S20]

So the most accurate summary is: **GABRA6 is a plausible and somewhat supported candidate in alcohol-use disorder biology, but it is not firmly established as a major addiction gene overall.** ([pmc.ncbi.nlm.nih.gov](https://pmc.ncbi.nlm.nih.gov/articles/PMC3924525/)) [S21]

## Cited sources

- **[S1]** α 6-Containing GABAA Receptors: Functional Roles and Therapeutic Potentials - PubMed — https://pubmed.ncbi.nlm.nih.gov/35017178/
- **[S2]** Association between variants at the GABAAbeta2, GABAAalpha6 and GABAAgamma2 gene cluster and alcohol dependence in a Scottish population - PubMed — https://pubmed.ncbi.nlm.nih.gov/10578235/
- **[S3]** Association between alcoholism and the genetic polymorphisms of the GABAA receptor genes on chromosome 5q33-34 in Korean population - PubMed — https://pubmed.ncbi.nlm.nih.gov/16778401/
- **[S4]** Association of the gamma-aminobutyric acid A receptor gene cluster with alcohol dependence in Taiwanese Han | Molecular Psychiatry — https://www.nature.com/articles/4001110?utm_source=openai
- **[S5]** 
            Association of Gamma-Aminobutyric Acid A Receptor α2 Gene (GABRA2) with Alcohol Use Disorder - PMC
         — https://pmc.ncbi.nlm.nih.gov/articles/PMC3924525/
- **[S6]** 
            A Critical Review of Methods and Results in the Search for Genetic Contributors to Alcohol Sensitivity - PMC
         — https://pmc.ncbi.nlm.nih.gov/articles/PMC5916326/
- **[S7]** 
            Genetics of GABAergic signaling in nicotine and alcohol dependence - PMC
         — https://pmc.ncbi.nlm.nih.gov/articles/PMC3746562/
- **[S8]** 
            Sex differences in GABAA receptor subunit transcript expression are mediated by genotype in subjects with alcohol‐related cirrhosis of the liver - PMC
         — https://pmc.ncbi.nlm.nih.gov/articles/PMC9744570/
- **[S9]** Association of GABA(A) receptors and alcohol dependence and the effects of genetic imprinting - PubMed — https://pubmed.ncbi.nlm.nih.gov/12555233/
- **[S10]** 
            GABA System Genes – No Evidence for a Role in Alcohol Use and Abuse in a Community-based Sample - PMC
         — https://pmc.ncbi.nlm.nih.gov/articles/PMC3984361/
- **[S11]** 
            Association of Gamma-Aminobutyric Acid A Receptor α2 Gene (GABRA2) with Alcohol Use Disorder - PMC
         — https://pmc.ncbi.nlm.nih.gov/articles/PMC3924525/
- **[S12]** Genome-wide association study of alcohol dependence in male Han Chinese and cross-ethnic polygenic risk score comparison - PubMed — https://pubmed.ncbi.nlm.nih.gov/31591379/
- **[S13]** Genome-wide meta-analysis of alcohol use disorder in East Asians - PubMed — https://pubmed.ncbi.nlm.nih.gov/35094024/
- **[S14]** 
            Association of Gamma-Aminobutyric Acid A Receptor α2 Gene (GABRA2) with Alcohol Use Disorder - PMC
         — https://pmc.ncbi.nlm.nih.gov/articles/PMC3924525/
- **[S15]** Emerging Role of One-Carbon Metabolism and DNA Methylation Enrichment on δ-Containing GABAA Receptor Expression in the Cerebellum of Subjects with Alcohol Use Disorders (AUD) - PubMed — https://pubmed.ncbi.nlm.nih.gov/29020412/
- **[S16]** 
            A Factor Analysis of Global GABAergic Gene Expression in Human Brain Identifies Specificity in Response to Chronic Alcohol and Cocaine Exposure - PMC
         — https://pmc.ncbi.nlm.nih.gov/articles/PMC3661725/
- **[S17]** Alcohol-induced motor impairment caused by increased extrasynaptic GABA(A) receptor activity - PubMed — https://pubmed.ncbi.nlm.nih.gov/15696164/
- **[S18]** 
            A Critical Review of Methods and Results in the Search for Genetic Contributors to Alcohol Sensitivity - PMC
         — https://pmc.ncbi.nlm.nih.gov/articles/PMC5916326/
- **[S19]** GABRA6 gene polymorphism and an attenuated stress response - PubMed — https://pubmed.ncbi.nlm.nih.gov/15197399/
- **[S20]** 
            Association of Gamma-Aminobutyric Acid A Receptor α2 Gene (GABRA2) with Alcohol Use Disorder - PMC
         — https://pmc.ncbi.nlm.nih.gov/articles/PMC3924525/
- **[S21]** 
            Association of Gamma-Aminobutyric Acid A Receptor α2 Gene (GABRA2) with Alcohol Use Disorder - PMC
         — https://pmc.ncbi.nlm.nih.gov/articles/PMC3924525/
