## Supplementary material for "Understanding Substance Dependence: What Differentiates Addictive from Non-Addictive Drugs?": Folder for supplementary files: addictive-genes-claims-first100-087-dpp6-gene-protein-is-implicated-in-addiction-6dbff2ef4083.md

# addictive-genes-claims-first100-087-6dbff2ef4083

**Original claim:** DPP6 gene/protein is implicated in addiction.

## Claim

DPP6 gene/protein is implicated in addiction.

## Verdict

Verdict: Likely True

The best current evidence supports **DPP6 as implicated in alcohol-related addiction phenotypes**, rather than as a broadly established addiction gene across all substances. Multiple large human genetic studies have linked the DPP6 locus to alcohol consumption/problem-use traits, postmortem human AUD tissue shows altered DPP6 expression, and a recent mouse loss-of-function study found enhanced ethanol reward-related behavior. However, the evidence is still largely associative, and direct evidence across non-alcohol addictions is limited. ([pmc.ncbi.nlm.nih.gov](https://pmc.ncbi.nlm.nih.gov/articles/PMC7711277/)) [S1]

## Confidence

Medium

## Supporting evidence

Large human GWAS provide the strongest support. In a large meta-analysis of alcohol consumption, a DPP6-region variant (rs6969458) was associated with drinks per week at genome-wide significance (P = 6.4 × 10^-14). ([pmc.ncbi.nlm.nih.gov](https://pmc.ncbi.nlm.nih.gov/articles/PMC7711277/)) [S2]

The signal is not isolated to simple consumption. In a GWAS meta-analysis of **problematic alcohol use** involving 435,563 individuals, the DPP6 locus again reached genome-wide significance (rs2533200; P = 1.79 × 10^-8). ([pmc.ncbi.nlm.nih.gov](https://pmc.ncbi.nlm.nih.gov/articles/PMC7485556/)) [S3]

A separate study of **maximum habitual alcohol intake** in U.S. veterans also identified DPP6 in multitrait analysis (rs2098112; P = 8.99 × 10^-10), linking the locus to a phenotype that genetically loads more strongly on alcohol-problem traits than on nondependent consumption. ([pmc.ncbi.nlm.nih.gov](https://pmc.ncbi.nlm.nih.gov/articles/PMC9614582/)) [S4]

More recently, a multi-ancestry GWAS of **alcohol use disorder (AUD)** in over one million individuals reported a genome-wide significant DPP6 locus in the European-ancestry analysis (rs2533199; P = 3.27 × 10^-8), with DPP6 listed among EUR-specific associated loci. ([pmc.ncbi.nlm.nih.gov](https://pmc.ncbi.nlm.nih.gov/articles/PMC10775504/)) [S5]

Human brain expression data are also consistent with involvement. In postmortem mediodorsal thalamus from people with alcohol-related disorders/AUD, DPP6 was among the differentially expressed genes; the authors noted that DPP6 was one of the genes showing both altered expression in AUD brain tissue and strong prior GWAS associations with AUD/alcohol phenotypes. ([pubmed.ncbi.nlm.nih.gov](https://pubmed.ncbi.nlm.nih.gov/33805312/)) [S6]

There is also supporting animal evidence. A recent Dpp6 knockout mouse study reported that **male homozygous knockouts showed increased ethanol conditioned place preference**, and knockout mice also showed altered ethanol-induced anxiolytic and sedative responses. That pattern is consistent with Dpp6 affecting sensitivity to alcohol’s rewarding effects. ([pubmed.ncbi.nlm.nih.gov](https://pubmed.ncbi.nlm.nih.gov/40894538/?utm_source=openai)) [S7]

## Contradicting or limiting evidence

The main limitation is **scope**: most direct evidence is about **alcohol-related phenotypes**, not addiction in general. The claim “implicated in addiction” is therefore better supported if interpreted as “implicated in alcohol use disorder/alcohol-related addiction biology” than if interpreted as a pan-addiction gene for nicotine, opioids, stimulants, and other substances. The strongest cited human studies above are all alcohol-focused. ([pmc.ncbi.nlm.nih.gov](https://pmc.ncbi.nlm.nih.gov/articles/PMC7711277/)) [S8]

The human evidence is also mostly **association-based**, not causal. GWAS signals map a locus to a phenotype, but they do not by themselves prove that DPP6 is the sole causal gene at that locus or define the direction of effect in specific cell types. The postmortem AUD study is informative but observational, and altered expression in diseased tissue could reflect consequences of chronic alcohol exposure as well as pre-existing vulnerability. ([pubmed.ncbi.nlm.nih.gov](https://pubmed.ncbi.nlm.nih.gov/33805312/)) [S9]

The animal evidence is promising but still limited. The recent ethanol conditioned place preference study is a **preprint indexed in PubMed/PMC pilot**, not a fully peer-reviewed journal article, and its strongest reward-related effect was sex-specific (seen mainly in male homozygous knockouts). It also did not show a clear intake difference in the limited drinking assay reported by the authors. ([pubmed.ncbi.nlm.nih.gov](https://pubmed.ncbi.nlm.nih.gov/40894538/?utm_source=openai)) [S10]

Finally, I did not find comparably strong primary evidence tying DPP6 directly to cocaine, opioid, or nicotine addiction outcomes in humans. That does not rule out such a role, but it means the broader wording of the claim should be treated cautiously.

## Mechanistic / biological rationale

DPP6 is biologically plausible as an addiction-relevant gene because it is a brain-enriched auxiliary subunit for Kv4-family A-type potassium channels and also has Kv4-independent roles in synaptic development. DPP6 helps regulate dendritic excitability, firing properties, synaptic maturation, cell adhesion, and learning-related plasticity. ([pmc.ncbi.nlm.nih.gov](https://pmc.ncbi.nlm.nih.gov/articles/PMC9409431/)) [S11]

Primary work in hippocampal neurons shows that DPP6 is critical for the normal dendritic A-type K^+ current gradient in CA1 neurons; DPP6 deletion disrupts Kv4.2 distribution and slows channel recovery/inactivation, altering dendritic excitability. ([pubmed.ncbi.nlm.nih.gov](https://pubmed.ncbi.nlm.nih.gov/21943606/)) [S12]

DPP6 loss also impairs dendritic filopodia/spine development and hippocampus-dependent learning and memory. DPP6-knockout mice show deficits in Morris water maze, T-maze, and object-related memory tasks, alongside altered synaptic structure and motility. ([pmc.ncbi.nlm.nih.gov](https://pmc.ncbi.nlm.nih.gov/articles/PMC5884885/)) [S13]

That matters for addiction because reward learning and relapse are heavily dependent on **contextual and associative circuitry**, especially hippocampal projections to nucleus accumbens and related forebrain networks. Independent studies show that hippocampus-to-NAc synaptic strength regulates reward behavior, repeated cocaine potentiates hippocampal inputs to NAc, and ventral hippocampus contributes to context-induced reinstatement of cocaine seeking. ([pmc.ncbi.nlm.nih.gov](https://pmc.ncbi.nlm.nih.gov/articles/PMC6292781/?utm_source=openai)) [S14]

DPP6 is also expressed in regions relevant to addiction circuitry, including the hippocampus, striatum, and substantia nigra, which further strengthens biological plausibility. ([pmc.ncbi.nlm.nih.gov](https://pmc.ncbi.nlm.nih.gov/articles/PMC9409431/)) [S15]

So, a reasonable mechanistic interpretation is: **variation or dysregulation of DPP6 could alter neural excitability and synaptic plasticity in circuits that encode reward, context, and reinforcement learning**, thereby influencing vulnerability to alcohol-related addictive behavior. That rationale is plausible, but still not fully proven in humans. ([pmc.ncbi.nlm.nih.gov](https://pmc.ncbi.nlm.nih.gov/articles/PMC9409431/)) [S16]

## Bottom line

DPP6 is **likely implicated in addiction if the claim is meant mainly in the context of alcohol-related addiction phenotypes**. The evidence includes repeated human GWAS signals for alcohol use/problem traits, altered DPP6 expression in human AUD brain tissue, and supportive mouse behavioral data. But the case is **not yet strong enough to call DPP6 a broadly established addiction gene across substances**, and much of the evidence remains associative rather than causal. ([pmc.ncbi.nlm.nih.gov](https://pmc.ncbi.nlm.nih.gov/articles/PMC7711277/)) [S17]

## Cited sources

- **[S1]** 
            Novel alcohol-related genes suggest shared genetic mechanisms with neuropsychiatric disorders - PMC
         — https://pmc.ncbi.nlm.nih.gov/articles/PMC7711277/
- **[S2]** 
            Novel alcohol-related genes suggest shared genetic mechanisms with neuropsychiatric disorders - PMC
         — https://pmc.ncbi.nlm.nih.gov/articles/PMC7711277/
- **[S3]** 
            Genome-wide meta-analysis of problematic alcohol use in 435,563 individuals yields insights into biology and relationships with other traits - PMC
         — https://pmc.ncbi.nlm.nih.gov/articles/PMC7485556/
- **[S4]** 
            Genome-Wide Investigation of Maximum Habitual Alcohol Intake in US Veterans in Relation to Alcohol Consumption Traits and Alcohol Use Disorder - PMC
         — https://pmc.ncbi.nlm.nih.gov/articles/PMC9614582/
- **[S5]** 
            Identification of Novel Loci and Cross-Disorder Pleiotropy Through Multi-Ancestry Genome-Wide Analysis of Alcohol Use Disorder in Over One Million Individuals - PMC
         — https://pmc.ncbi.nlm.nih.gov/articles/PMC10775504/
- **[S6]** Chronic Alcohol Use Induces Molecular Genetic Changes in the Dorsomedial Thalamus of People with Alcohol-Related Disorders - PubMed — https://pubmed.ncbi.nlm.nih.gov/33805312/
- **[S7]** Dpp6 Homozygous Knockout Mice Exhibit Increased Ethanol Conditioned Place Preference and Acute Ethanol-Induced Anxiolytic Behavior - PubMed — https://pubmed.ncbi.nlm.nih.gov/40894538/?utm_source=openai
- **[S8]** 
            Novel alcohol-related genes suggest shared genetic mechanisms with neuropsychiatric disorders - PMC
         — https://pmc.ncbi.nlm.nih.gov/articles/PMC7711277/
- **[S9]** Chronic Alcohol Use Induces Molecular Genetic Changes in the Dorsomedial Thalamus of People with Alcohol-Related Disorders - PubMed — https://pubmed.ncbi.nlm.nih.gov/33805312/
- **[S10]** Dpp6 Homozygous Knockout Mice Exhibit Increased Ethanol Conditioned Place Preference and Acute Ethanol-Induced Anxiolytic Behavior - PubMed — https://pubmed.ncbi.nlm.nih.gov/40894538/?utm_source=openai
- **[S11]** 
            Neuronal Roles of the Multifunctional Protein Dipeptidyl Peptidase-like 6 (DPP6) - PMC
         — https://pmc.ncbi.nlm.nih.gov/articles/PMC9409431/
- **[S12]** DPP6 establishes the A-type K(+) current gradient critical for the regulation of dendritic excitability in CA1 hippocampal neurons - PubMed — https://pubmed.ncbi.nlm.nih.gov/21943606/
- **[S13]** 
            DPP6 Loss Impacts Hippocampal Synaptic Development and Induces Behavioral Impairments in Recognition, Learning and Memory - PMC
         — https://pmc.ncbi.nlm.nih.gov/articles/PMC5884885/
- **[S14]** Reward behavior is regulated by the strength of hippocampus-nucleus accumbens synapses - PMC — https://pmc.ncbi.nlm.nih.gov/articles/PMC6292781/?utm_source=openai
- **[S15]** 
            Neuronal Roles of the Multifunctional Protein Dipeptidyl Peptidase-like 6 (DPP6) - PMC
         — https://pmc.ncbi.nlm.nih.gov/articles/PMC9409431/
- **[S16]** 
            Neuronal Roles of the Multifunctional Protein Dipeptidyl Peptidase-like 6 (DPP6) - PMC
         — https://pmc.ncbi.nlm.nih.gov/articles/PMC9409431/
- **[S17]** 
            Novel alcohol-related genes suggest shared genetic mechanisms with neuropsychiatric disorders - PMC
         — https://pmc.ncbi.nlm.nih.gov/articles/PMC7711277/
