## Supplementary material for "Understanding Substance Dependence: What Differentiates Addictive from Non-Addictive Drugs?": Folder for supplementary files: addictive-genes-claims-first100-088-exph5-gene-protein-is-implicated-in-addiction-b33e2418f390.md

# addictive-genes-claims-first100-088-b33e2418f390

**Original claim:** EXPH5 gene/protein is implicated in addiction.

## Claim

EXPH5 gene/protein is implicated in addiction.

## Verdict

Verdict: No Evidence

## Confidence

Medium

## Supporting evidence

The strongest current human genetics resources do **not** support EXPH5 as an addiction gene. In the NHGRI-EBI GWAS Catalog, the EXPH5 gene page currently shows no listed associations, no studies, and no traits mapped to EXPH5, which argues against a replicated genome-wide signal for addiction or other traits being curated to this gene. ([ebi.ac.uk](https://www.ebi.ac.uk/gwas/genes/EXPH5)) [S1]

Recent high-level reviews of substance use disorder genetics also do not identify EXPH5 among the loci most strongly implicated in shared or substance-specific addiction risk. A 2024 *Journal of Clinical Investigation* review of generalized genetic liability to substance use disorders highlighted dopaminergic regulation and neuronal development mechanisms, with examples including **DRD2, PDE4B, BDNF, FTO**, and substance-specific loci such as **ADH1B, CHRNA5, CHRNB2,** and **OPRM1**. EXPH5 was not among the emphasized genes or loci. ([jci.org](https://www.jci.org/articles/view/172881)) [S2]

The curated functional annotation of EXPH5 points elsewhere biologically. NCBI describes EXPH5 as encoding a synaptotagmin-like family protein that acts as a **Rab27B effector** involved in **exosome secretion** and **intracellular vesicle trafficking**; reduced expression causes **keratin filament defects**, and pathogenic variants are associated with **epidermolysis bullosa simplex**, a skin fragility disorder. That disease-focused annotation is also what appears in the gene’s phenotype literature. ([ncbi.nlm.nih.gov](https://www.ncbi.nlm.nih.gov/gene/23086)) [S3]

Primary human disease papers on EXPH5 likewise center on skin disease, not neuropsychiatric or addictive phenotypes. In the original human report linking EXPH5 loss-of-function to inherited skin fragility, affected individuals had skin findings but **no neurologic abnormalities** reported. ([pmc.ncbi.nlm.nih.gov](https://pmc.ncbi.nlm.nih.gov/articles/PMC3516608/)) [S4]

A review of Rab27 effectors is also informative: it summarizes exophilin-5/EXPH5 as causing epidermolysis bullosa simplex and affecting airway/Th2 inflammatory biology, while noting that a **different family member**, exophilin-6, has reported neuronal migration and presynaptic neurotransmitter-release effects. That pattern suggests Rab27-family membership alone is not enough to infer an addiction role for EXPH5 specifically. ([pmc.ncbi.nlm.nih.gov](https://pmc.ncbi.nlm.nih.gov/articles/PMC10511049/)) [S5]

## Contradicting or limiting evidence

There is some **indirect biological plausibility**, but it falls short of evidence for addiction. Human Protein Atlas data label EXPH5 as **tissue enhanced in brain and skin**, and the original EXPH5 mutation paper detected EXPH5 mRNA amplification from **brain** as well as skin, muscle, and heart. So EXPH5 is not purely skin-restricted, and a CNS-related role is not impossible. ([proteinatlas.org](https://www.proteinatlas.org/ENSG00000110723-EXPH5/brain)) [S6]

Also, EXPH5 participates in vesicle/exosome trafficking, and vesicle trafficking processes can matter for neuronal signaling broadly. However, that is a generic mechanistic possibility, not addiction-specific evidence. Neither the curated GWAS resource nor the major recent addiction-genetics reviews cited above provide direct support for EXPH5 as an addiction locus. ([ncbi.nlm.nih.gov](https://www.ncbi.nlm.nih.gov/gene/23086)) [S7]

A further limitation is that absence from current large studies does **not** prove EXPH5 has zero effect. Addiction genetics remains polygenic and incomplete, and reviews note that larger and more granular studies are still needed to resolve modest, context-specific, or ancestry-specific signals. So the appropriate conclusion is “no evidence found,” not “definitively excluded.” ([jci.org](https://www.jci.org/articles/view/172881)) [S8]

## Mechanistic / biological rationale

For a gene to be credibly “implicated in addiction,” one usually wants at least one of the following: replicated human genetic association, reproducible differential regulation in addiction-relevant brain tissue, or functional evidence connecting the gene to reward circuitry, drug reinforcement, withdrawal, relapse, or core neuroadaptations. Current addiction literature instead emphasizes pathways involving **dopamine signaling, synaptic plasticity, glutamatergic regulation, neuronal development, and stress/inflammatory responses** in defined reward-circuit cell types. ([jci.org](https://www.jci.org/articles/view/172881)) [S9]

EXPH5’s known biology does not map cleanly onto that evidence base. The gene encodes exophilin-5/Slac2-b, a **Rab27B effector** involved in vesicle trafficking and exosome secretion. In humans, the clearest phenotype is **recessive epidermolysis bullosa simplex**. In mice, exophilin-5 has also been linked to **allergic airway inflammation** and regulation of IL-33/Th2 responses. These are real biological functions, but they are not the canonical addiction mechanisms established for stronger addiction genes. ([ncbi.nlm.nih.gov](https://www.ncbi.nlm.nih.gov/gene/23086)) [S10]

The most one can say mechanistically is that vesicle-trafficking proteins can, in principle, influence neuronal homeostasis or signaling, and EXPH5 is expressed in brain tissue. But that remains speculative for addiction unless tied to specific data in reward-related circuits or substance-related phenotypes. I did not find such direct evidence in the scholarly sources reviewed here. ([proteinatlas.org](https://www.proteinatlas.org/ENSG00000110723-EXPH5/brain)) [S11]

## Bottom line

As of **March 16, 2026**, the claim that **EXPH5 is implicated in addiction** is **not supported by current curated human genetics or mainstream addiction-biology literature**. The best-supported roles for EXPH5 are in **Rab27B-mediated vesicle/exosome trafficking**, **skin integrity/epidermolysis bullosa simplex**, and some **airway immune biology**. Brain expression gives mild mechanistic plausibility, but I found **no convincing direct human genetic, transcriptomic, or functional evidence** linking EXPH5 specifically to addiction. ([ebi.ac.uk](https://www.ebi.ac.uk/gwas/genes/EXPH5)) [S12]

## Cited sources

- **[S1]** GWAS Catalog — https://www.ebi.ac.uk/gwas/genes/EXPH5
- **[S2]** 
JCI -
Generalized genetic liability to substance use disorders
 — https://www.jci.org/articles/view/172881
- **[S3]** EXPH5 exophilin 5 [Homo sapiens (human)] - Gene - NCBI — https://www.ncbi.nlm.nih.gov/gene/23086
- **[S4]** 
            Germline Mutation in EXPH5 Implicates the Rab27B Effector Protein Slac2-b in Inherited Skin Fragility - PMC
         — https://pmc.ncbi.nlm.nih.gov/articles/PMC3516608/
- **[S5]** 
            In vivo Roles of Rab27 and Its Effectors in Exocytosis - PMC
         — https://pmc.ncbi.nlm.nih.gov/articles/PMC10511049/
- **[S6]** Brain tissue expression of EXPH5 - Summary - The Human Protein Atlas — https://www.proteinatlas.org/ENSG00000110723-EXPH5/brain
- **[S7]** EXPH5 exophilin 5 [Homo sapiens (human)] - Gene - NCBI — https://www.ncbi.nlm.nih.gov/gene/23086
- **[S8]** 
JCI -
Generalized genetic liability to substance use disorders
 — https://www.jci.org/articles/view/172881
- **[S9]** 
JCI -
Generalized genetic liability to substance use disorders
 — https://www.jci.org/articles/view/172881
- **[S10]** EXPH5 exophilin 5 [Homo sapiens (human)] - Gene - NCBI — https://www.ncbi.nlm.nih.gov/gene/23086
- **[S11]** Brain tissue expression of EXPH5 - Summary - The Human Protein Atlas — https://www.proteinatlas.org/ENSG00000110723-EXPH5/brain
- **[S12]** GWAS Catalog — https://www.ebi.ac.uk/gwas/genes/EXPH5
