## Supplementary material for "Understanding Substance Dependence: What Differentiates Addictive from Non-Addictive Drugs?": Folder for supplementary files: addictive-genes-claims-first100-089-slc44a2-gene-protein-is-implicated-in-addiction-08f6b3b403df.md

# addictive-genes-claims-first100-089-08f6b3b403df

**Original claim:** SLC44A2 gene/protein is implicated in addiction.

## Claim

**Claim:** *SLC44A2 gene/protein is implicated in addiction.*

## Verdict

Verdict: No Evidence

I did **not** find convincing direct evidence that **SLC44A2** is a replicated addiction gene/protein in humans or a validated addiction-relevant target in animal models. The current literature around SLC44A2 is dominated by **thrombosis/platelet biology, blood-group antigen biology, hearing loss, and inner-ear disease**, not substance use disorders. ([ddd.uab.cat](https://ddd.uab.cat/pub/artpub/2020/283367/283367.pdf)) [S1]

## Confidence

**Medium**

Confidence is **medium** because absence of evidence is harder to prove than the presence of a positive association. However, targeted searches of major addiction-genetics reviews and landmark GWAS papers did not identify SLC44A2, while the SLC44A2-specific literature consistently points to other phenotypes. ([nature.com](https://www.nature.com/articles/s41380-024-02547-z)) [S2]

## Supporting evidence

There is a **plausible indirect biological rationale** for why someone might wonder about SLC44A2. SLC44A2 encodes **CTL2**, a choline transporter-like protein. A review of choline transport at the blood-brain barrier reports that CTL2 is expressed in **brain microvascular endothelial cells**, helps move choline across the blood-brain barrier, and is also present in **mitochondria**; choline itself is essential for **acetylcholine synthesis**, membrane phospholipids, and methyl-donor metabolism through betaine/SAM pathways. Those processes are broadly relevant to brain function. ([pubmed.ncbi.nlm.nih.gov](https://pubmed.ncbi.nlm.nih.gov/31547050/)) [S3]

There is also functional biology showing SLC44A2 can affect **cellular energetics**. In mice, Slc44a2 was shown to mediate **mitochondrial choline transport**, increasing mitochondrial oxygen consumption and ATP production in platelets; knockout mice had impaired platelet activation and delayed thrombosis. That shows SLC44A2 is a biologically active transporter/scaffold with real physiological effects, not an uncharacterized gene. ([ddd.uab.cat](https://ddd.uab.cat/pub/artpub/2020/283367/283367.pdf)) [S4]

SLC44A2 is not purely “non-neural.” Older family-level review work described **SLC44A2-4 as mainly peripheral**, in contrast to **SLC44A1**, which is widely expressed in the nervous system, but later work identified CTL2 in the **blood-brain barrier** and in multiple tissues. So an indirect role in neural homeostasis is not impossible. ([pubmed.ncbi.nlm.nih.gov](https://pubmed.ncbi.nlm.nih.gov/23506897/)) [S5]

## Contradicting or limiting evidence

The biggest limitation for the claim is that **direct addiction evidence is missing**. In a 2024 review of the genetics of substance use disorders, the replicated loci discussed include genes such as **ADH1B/ADH1C**, **DRD2**, **CHRNA2**, and **OPRM1/FURIN/TSNARE1** for different substance use disorders and shared addiction risk. A text search of that review did **not** find SLC44A2. ([nature.com](https://www.nature.com/articles/s41380-024-02547-z)) [S6]

Likewise, text searches of landmark large-scale addiction GWAS papers available here did **not** identify SLC44A2 in a major tobacco/alcohol genetics paper, a cannabis use disorder GWAS, or a transancestral alcohol dependence GWAS. That does **not** prove SLC44A2 has zero effect, but it strongly argues against it being a currently established or replicated addiction locus. ([gwern.net](https://gwern.net/doc/genetics/heritable/correlation/2019-liu.pdf)) [S7]

The **SLC44A2-centered literature** that is easy to retrieve is focused elsewhere. A 2020 *Nature Communications* paper links SLC44A2 to **thrombosis and platelet mitochondrial function**. Mouse knockout work links Slc44a2 to **progressive sensorineural hearing loss, hair-cell loss, and spiral ganglion degeneration**. Human SLC44A2-null cases were reported with **hearing impairment and a rare red blood phenotype**, not addiction phenotypes. ([ddd.uab.cat](https://ddd.uab.cat/pub/artpub/2020/283367/283367.pdf)) [S8]

Even within cholinergic biology, SLC44A2 is **not** the best-established neuronal addiction candidate. The blood-brain-barrier review states that **high-affinity choline transporter 1 (CHT1/SLC5A7)** is the classic transporter involved in acetylcholine synthesis in neurons, while **CTL1/SLC44A1** is also functionally expressed in neurons/astrocytes and is more strongly tied to nervous-system phospholipid biology. CTL2/SLC44A2 appears more prominent at the **BBB** and in non-neuronal contexts. ([pubmed.ncbi.nlm.nih.gov](https://pubmed.ncbi.nlm.nih.gov/31547050/)) [S9]

## Mechanistic / biological rationale

If one wanted to argue for a possible role, the argument would have to be **indirect and speculative**. Because CTL2 can affect **brain choline availability** at the blood-brain barrier and because choline contributes to **acetylcholine synthesis, membrane phospholipid turnover, mitochondrial metabolism, and methyl-donor/epigenetic pathways**, SLC44A2 could in principle modulate aspects of neural homeostasis that are upstream of reward circuitry or drug responses. ([pubmed.ncbi.nlm.nih.gov](https://pubmed.ncbi.nlm.nih.gov/31547050/)) [S10]

But that is still far from saying it is “implicated in addiction.” To justify that stronger claim, one would normally want at least one of the following: replicated human genetic associations with substance use disorder, brain-expression/TWAS evidence in addiction cohorts, differential expression in addiction-relevant tissue, or animal studies showing effects on **drug reward, reinforcement, withdrawal, craving, sensitization, or relapse**. I did not identify such evidence for SLC44A2 in the materials retrieved. Instead, the strongest SLC44A2 biology points to **platelets/thrombosis**, **hearing/inner ear**, and **blood-cell phenotypes**. ([ddd.uab.cat](https://ddd.uab.cat/pub/artpub/2020/283367/283367.pdf)) [S11]

So biologically, the fairest summary is: **possible indirect relevance to choline handling and cellular energetics**, but **no demonstrated addiction-specific role** at present. ([pubmed.ncbi.nlm.nih.gov](https://pubmed.ncbi.nlm.nih.gov/31547050/)) [S12]

## Bottom line

Based on the currently retrievable scholarly and authoritative sources, the claim that **SLC44A2 is implicated in addiction** is **not supported by direct evidence**. The gene/protein has real biology and some plausible indirect links to brain homeostasis through **choline transport** and **mitochondrial function**, but the published evidence I found ties it primarily to **thrombosis/platelet biology, hearing loss, and blood antigen phenotypes**, not addiction. ([ddd.uab.cat](https://ddd.uab.cat/pub/artpub/2020/283367/283367.pdf)) [S13]

If you want the strongest cholinergic/addiction-related alternatives, genes with much better evidence include **CHRNA2**, **nicotinic receptor loci**, **DRD2**, **OPRM1**, and alcohol-metabolism genes such as **ADH1B**. ([nature.com](https://www.nature.com/articles/s41380-024-02547-z)) [S14]

## Cited sources

- **[S1]** The choline transporter Slc44a2 controls platelet activation and thrombosis by regulating mitochondrial function — https://ddd.uab.cat/pub/artpub/2020/283367/283367.pdf
- **[S2]** The genetic landscape of substance use disorders | Molecular Psychiatry — https://www.nature.com/articles/s41380-024-02547-z
- **[S3]** Functional Expression of Choline Transporters in the Blood-Brain Barrier - PubMed — https://pubmed.ncbi.nlm.nih.gov/31547050/
- **[S4]** The choline transporter Slc44a2 controls platelet activation and thrombosis by regulating mitochondrial function — https://ddd.uab.cat/pub/artpub/2020/283367/283367.pdf
- **[S5]** The choline transporter-like family SLC44: properties and roles in human diseases - PubMed — https://pubmed.ncbi.nlm.nih.gov/23506897/
- **[S6]** The genetic landscape of substance use disorders | Molecular Psychiatry — https://www.nature.com/articles/s41380-024-02547-z
- **[S7]** Association studies of up to 1.2 million individuals yield new insights into the genetic etiology of tobacco and alcohol use — https://gwern.net/doc/genetics/heritable/correlation/2019-liu.pdf
- **[S8]** The choline transporter Slc44a2 controls platelet activation and thrombosis by regulating mitochondrial function — https://ddd.uab.cat/pub/artpub/2020/283367/283367.pdf
- **[S9]** Functional Expression of Choline Transporters in the Blood-Brain Barrier - PubMed — https://pubmed.ncbi.nlm.nih.gov/31547050/
- **[S10]** Functional Expression of Choline Transporters in the Blood-Brain Barrier - PubMed — https://pubmed.ncbi.nlm.nih.gov/31547050/
- **[S11]** The choline transporter Slc44a2 controls platelet activation and thrombosis by regulating mitochondrial function — https://ddd.uab.cat/pub/artpub/2020/283367/283367.pdf
- **[S12]** Functional Expression of Choline Transporters in the Blood-Brain Barrier - PubMed — https://pubmed.ncbi.nlm.nih.gov/31547050/
- **[S13]** The choline transporter Slc44a2 controls platelet activation and thrombosis by regulating mitochondrial function — https://ddd.uab.cat/pub/artpub/2020/283367/283367.pdf
- **[S14]** The genetic landscape of substance use disorders | Molecular Psychiatry — https://www.nature.com/articles/s41380-024-02547-z
