## Supplementary material for "Understanding Substance Dependence: What Differentiates Addictive from Non-Addictive Drugs?": Folder for supplementary files: addictive-genes-claims-first100-090-apbb2-gene-protein-is-implicated-in-addiction-23d64ad8bf4d.md

# addictive-genes-claims-first100-090-23d64ad8bf4d

**Original claim:** APBB2 gene/protein is implicated in addiction.

## Claim

APBB2 gene/protein is implicated in addiction.

## Verdict

Verdict: Mixed Evidence

APBB2 can reasonably be described as a **tentative candidate gene** for some **opioid-related addiction phenotypes**, because human genetic studies have reported associations with opioid dependence and with amphetamine co-use in heroin-dependent patients on methadone. However, those signals are limited, ancestry/context-specific, and have **not emerged as robust, consistently replicated leading findings** in larger later opioid-use-disorder GWAS. ([pmc.ncbi.nlm.nih.gov](https://pmc.ncbi.nlm.nih.gov/articles/PMC3992201/)) [S1]

## Confidence

**Medium.** The claim is not baseless: there are published human association studies directly naming APBB2 in opioid-related phenotypes. But the evidence base is small, mostly genetic/associational rather than mechanistic, and later larger studies have shifted emphasis toward other loci such as **OPRM1**, **PPP6C**, and **FURIN**. ([pmc.ncbi.nlm.nih.gov](https://pmc.ncbi.nlm.nih.gov/articles/PMC3992201/)) [S2]

## Supporting evidence

The strongest direct support comes from a 2014 three-phase GWAS of opioid dependence by Gelernter et al. In African American participants, the study reported **genome-wide significant** APBB2-region associations in the symptom-count model after phase 1+2 meta-analysis: rs115368721 in **APBB2** reached **p = 1.92 × 10^-8**, and the linked proxy rs114070671 reached **p = 2.26 × 10^-9**. The authors explicitly described APBB2 as an additional genome-wide-significant association of biological interest in opioid dependence. ([pmc.ncbi.nlm.nih.gov](https://pmc.ncbi.nlm.nih.gov/articles/PMC3992201/)) [S3]

A second supportive study came from Liu et al. (2018), who examined **344 heroin-dependent patients undergoing methadone maintenance treatment**. They reported that APBB2 intronic variants **rs3935357** and **rs4861075** were significantly associated with **amphetamine use** within this opioid-dependent cohort, and that the rs4861075 TT genotype was associated with higher APBB2 expression in EBV-transformed lymphoblastoid cells. This does not show APBB2 causes addiction, but it does connect APBB2 variation to a clinically relevant addiction-related phenotype: polysubstance stimulant use among people with opioid dependence. ([pubmed.ncbi.nlm.nih.gov](https://pubmed.ncbi.nlm.nih.gov/29330135/)) [S4]

Review-level summaries of opioid addiction genetics have also listed APBB2 among GWAS-identified opioid-use-disorder loci. For example, a focused review of opioid addiction genetics tabulated **APBB2 rs114070671** among significant GWAS variants for opioid use disorder. That review did not elevate APBB2 to the level of the most robust loci, but it supports the narrower statement that APBB2 has been implicated in human addiction genetics literature. ([pmc.ncbi.nlm.nih.gov](https://pmc.ncbi.nlm.nih.gov/articles/PMC6368898/)) [S5]

## Contradicting or limiting evidence

The main limitation is **replication**. In the original 2014 GWAS, the APBB2 signal weakened in the independent phase 3 sample: the proxy APBB2 SNP **rs114070671** had **p = 0.55** in phase 3, and the combined phase 1+2+3 signal attenuated to **p = 1.04 × 10^-7**, which is below conventional genome-wide significance. The paper itself described APBB2 as a “not-yet-replicated” association of interest rather than a firmly established locus. ([pmc.ncbi.nlm.nih.gov](https://pmc.ncbi.nlm.nih.gov/articles/PMC3992201/)) [S6]

Later, much larger GWAS did **not** confirm APBB2 as one of the top replicated opioid-risk genes. A large JAMA Psychiatry meta-analysis of **10,544 European-ancestry OUD cases and 72,163 opioid-exposed controls** found **OPRM1 rs1799971** as the significant replicated association, while stating that **most previously reported variants associated with opioid-related traits were not significant in MVP**. The paper specifically referenced earlier GWAS hits mapped to APBB2, but APBB2 did not emerge as a replicated lead signal in that larger analysis. ([jamanetwork.com](https://jamanetwork.com/journals/jamapsychiatry/fullarticle/2766708)) [S7]

Similarly, a 2022 large-scale multi-trait GWAS of opioid addiction involving **23,367 cases** and effective sample size **88,114** emphasized **OPRM1**, and gene-based signals at **PPP6C** and **FURIN**. The authors noted that, prior to their study, only **rs1799971** had been independently replicated; APBB2 was not among the highlighted genome-wide significant loci. ([pmc.ncbi.nlm.nih.gov](https://pmc.ncbi.nlm.nih.gov/articles/PMC9546890/)) [S8]

A 2024 JCI review reached a similar overall conclusion: as GWAS sample sizes have grown, the strongest current variant-level signals are in loci such as **OPRM1** and **FURIN**, while many other omics findings remain insufficiently robust or hard to compare across studies. That broader field-level assessment argues against treating APBB2 as an established addiction gene at present. ([pmc.ncbi.nlm.nih.gov](https://pmc.ncbi.nlm.nih.gov/articles/PMC11473141/)) [S9]

Also important: the APBB2 literature here is mostly about **opioid dependence/OUD** and **amphetamine co-use in opioid-dependent patients**. I did not identify strong primary evidence from large human studies tying APBB2 to addiction across alcohol, nicotine, cocaine, or cannabis in the way seen for better-established addiction genes. The current implication is therefore **narrow**, not general across all addictions. ([pmc.ncbi.nlm.nih.gov](https://pmc.ncbi.nlm.nih.gov/articles/PMC3992201/)) [S10]

## Mechanistic / biological rationale

APBB2 encodes **amyloid beta precursor protein-binding family B member 2** (also called **FE65L1**), an adaptor protein that binds the cytoplasmic domains of APP-family proteins and contains **two phosphotyrosine-binding domains**, consistent with a role in intracellular signaling rather than direct neurotransmitter reception. That basic biology makes it plausible as a neural modifier gene, but it is not an obvious canonical addiction receptor or transporter. ([ncbi.nlm.nih.gov](https://www.ncbi.nlm.nih.gov/gtr/genes/323?utm_source=openai)) [S11]

Expression data support neural relevance: Human Protein Atlas data show APBB2 expression in the brain, including the **basal ganglia** and **nucleus accumbens**, regions deeply involved in reward and addiction circuitry. ([proteinatlas.org](https://www.proteinatlas.org/ENSG00000163697-APBB2/brain/basal%2Bganglia)) [S12]

There is also indirect family-level functional support from mouse work on **FE65/FE65L1** proteins. FE65/FE65L1 genetic disruption has been linked to altered **synaptic plasticity/LTP**, neuromuscular junction abnormalities, abnormal open-field habituation, and reduced anxiety-like behavior. Those phenotypes indicate that FE65-family proteins can influence neural circuit function and plasticity, which are biologically relevant to addiction. But these experiments do **not** directly show that APBB2 itself drives drug reward, self-administration, withdrawal, or relapse. ([pmc.ncbi.nlm.nih.gov](https://pmc.ncbi.nlm.nih.gov/articles/PMC4899880/)) [S13]

So the mechanistic case is best described as **plausible but indirect**: APBB2 is a brain-expressed adaptor involved in neuronal signaling/synaptic biology, and human association studies hint at opioid-related relevance, but a direct addiction mechanism has not been established. ([ncbi.nlm.nih.gov](https://www.ncbi.nlm.nih.gov/gtr/genes/323?utm_source=openai)) [S14]

## Bottom line

APBB2 is **not a definitive, strongly validated addiction gene**, but it **has been implicated** in addiction-related phenotypes—mainly **opioid dependence/OUD** and **amphetamine co-use within opioid-dependent cohorts**—by human genetic association studies. Because those findings are limited and not robustly replicated in later larger GWAS, the most accurate overall assessment is **Mixed Evidence**, not “highly supported.” ([pmc.ncbi.nlm.nih.gov](https://pmc.ncbi.nlm.nih.gov/articles/PMC3992201/)) [S15]

## Cited sources

- **[S1]** 
            Genomewide association study of opioid dependence: multiple associations mapped to calcium and potassium pathways - PMC
         — https://pmc.ncbi.nlm.nih.gov/articles/PMC3992201/
- **[S2]** 
            Genomewide association study of opioid dependence: multiple associations mapped to calcium and potassium pathways - PMC
         — https://pmc.ncbi.nlm.nih.gov/articles/PMC3992201/
- **[S3]** 
            Genomewide association study of opioid dependence: multiple associations mapped to calcium and potassium pathways - PMC
         — https://pmc.ncbi.nlm.nih.gov/articles/PMC3992201/
- **[S4]** APBB2 is associated with amphetamine use and plasma beta-amyloids in patients receiving methadone maintenance treatment - PubMed — https://pubmed.ncbi.nlm.nih.gov/29330135/
- **[S5]** 
            A Review of Opioid Addiction Genetics - PMC
         — https://pmc.ncbi.nlm.nih.gov/articles/PMC6368898/
- **[S6]** 
            Genomewide association study of opioid dependence: multiple associations mapped to calcium and potassium pathways - PMC
         — https://pmc.ncbi.nlm.nih.gov/articles/PMC3992201/
- **[S7]** Association of OPRM1 Functional Coding Variant With Opioid Use Disorder: A Genome-Wide Association Study | Genetics and Genomics | JAMA Psychiatry | JAMA Network — https://jamanetwork.com/journals/jamapsychiatry/fullarticle/2766708
- **[S8]** 
            Multi-trait genome-wide association study of opioid addiction: OPRM1 and beyond - PMC
         — https://pmc.ncbi.nlm.nih.gov/articles/PMC9546890/
- **[S9]** 
            An emerging multi-omic understanding of the genetics of opioid addiction - PMC
         — https://pmc.ncbi.nlm.nih.gov/articles/PMC11473141/
- **[S10]** 
            Genomewide association study of opioid dependence: multiple associations mapped to calcium and potassium pathways - PMC
         — https://pmc.ncbi.nlm.nih.gov/articles/PMC3992201/
- **[S11]** APBB2 amyloid beta precursor protein binding family B member 2 - NIH Genetic Testing Registry (GTR) - NCBI — https://www.ncbi.nlm.nih.gov/gtr/genes/323?utm_source=openai
- **[S12]** Brain tissue expression of APBB2 in basal ganglia - The Human Protein Atlas — https://www.proteinatlas.org/ENSG00000163697-APBB2/brain/basal%2Bganglia
- **[S13]** 
            FE65 and FE65L1 share common synaptic functions and genetically interact with the APP family in neuromuscular junction formation - PMC
         — https://pmc.ncbi.nlm.nih.gov/articles/PMC4899880/
- **[S14]** APBB2 amyloid beta precursor protein binding family B member 2 - NIH Genetic Testing Registry (GTR) - NCBI — https://www.ncbi.nlm.nih.gov/gtr/genes/323?utm_source=openai
- **[S15]** 
            Genomewide association study of opioid dependence: multiple associations mapped to calcium and potassium pathways - PMC
         — https://pmc.ncbi.nlm.nih.gov/articles/PMC3992201/
