## Supplementary material for "Understanding Substance Dependence: What Differentiates Addictive from Non-Addictive Drugs?": Folder for supplementary files: addictive-genes-claims-first100-091-pdyn-gene-protein-is-implicated-in-addiction-b81f2121f08d.md

# addictive-genes-claims-first100-091-b81f2121f08d

**Original claim:** PDYN gene/protein is implicated in addiction.

## Claim

**“PDYN gene/protein is implicated in addiction.”**

## Verdict

Verdict: Likely True

## Confidence

**Medium**

## Supporting evidence

PDYN encodes **prodynorphin**, a precursor protein that is processed into **dynorphin peptides**, endogenous ligands for the **kappa opioid receptor (KOR)**. NCBI’s gene record specifically notes that dynorphin is involved in modulating responses to psychoactive substances, including cocaine, making PDYN biologically relevant to addiction from the outset. ([ncbi.nlm.nih.gov](https://www.ncbi.nlm.nih.gov/gene/5173)) [S1]

**Human genetic association studies** support an addiction link, though not uniformly. In a family-based study of **1,860 European American individuals from 219 multiplex alcohol-dependent families**, multiple SNPs in **PDYN** were associated with alcohol dependence, and haplotype analyses strengthened the signal. Another study found a PDYN haplotype associated both with **alcohol dependence** and with a tendency to drink in **negative emotional states** (“negative craving”), which is clinically relevant to relapse-prone addiction phenotypes. ([pubmed.ncbi.nlm.nih.gov](https://pubmed.ncbi.nlm.nih.gov/16924269/)) [S2]

Evidence also extends to **other substances**. A 2009 study reported that a **3′UTR PDYN haplotype** was associated with **cocaine dependence/cocaine-alcohol codependence** in Caucasians, and the same risk haplotype was linked to **lower PDYN mRNA expression in human dorsal and ventral striatum**, adding functional plausibility. A separate study found **PDYN polymorphisms associated with opioid addiction** in European Americans, with stronger odds ratios in women in sex-stratified analyses. A Japanese case-control study reported that a higher-expression **PDYN promoter VNTR allele** was more frequent in **methamphetamine dependence**. ([pubmed.ncbi.nlm.nih.gov](https://pubmed.ncbi.nlm.nih.gov/18923396/)) [S3]

A meta-analysis of PDYN polymorphisms and **opioid dependence** concluded that **rs1022563** showed a significant association in **Asian populations**, although the authors emphasized that the included evidence base was too limited for a robust final conclusion. That pattern—signal present but not definitive—is consistent with a gene that is implicated but not yet settled as a major universal risk locus. ([pubmed.ncbi.nlm.nih.gov](https://pubmed.ncbi.nlm.nih.gov/31510971/)) [S4]

**Human brain studies** also support involvement. In postmortem tissue, **heroin abusers** showed a selective reduction of **PDYN mRNA in the periamygdaloid cortex (PAC)** across two independent cohorts; rats with chronic heroin self-administration showed a similar PAC Pdyn reduction during withdrawal. The authors concluded that impaired PAC-Pdyn function likely contributes to stress/negative-affect circuitry relevant to opiate addiction vulnerability. In alcohol-related human brain studies, PDYN expression and PDYN/OPRK1 coordination are altered in reward- and control-related regions, including the nucleus accumbens and dorsolateral prefrontal cortex. ([jci.org](https://www.jci.org/articles/view/70395)) [S5]

**Animal and circuit-level causal evidence** is stronger than the human genetics. PDYN knockout mice show **higher ethanol-conditioned place preference** and **greater ethanol consumption/preference**, consistent with increased vulnerability to alcohol reinforcement when prodynorphin is lost. Region-specific work likewise found that **knocking out Pdyn in the central amygdala reduced alcohol intake** in male mice. During opioid withdrawal, blockade of dynorphin release from **Pdyn-expressing dorsal raphe neurons** prevented withdrawal-related social deficits in mice, implicating dynorphin signaling in withdrawal states that can promote relapse. ([pubmed.ncbi.nlm.nih.gov](https://pubmed.ncbi.nlm.nih.gov/21966993/)) [S6]

Finally, in humans with **cocaine use disorder**, PET imaging of the dynorphin/KOR system found that **greater KOR availability** was associated with **more cocaine choice** under stress, and a 3-day cocaine binge reduced tracer binding, interpreted in context as consistent with altered endogenous dynorphin/KOR signaling in cocaine addiction. This does not measure PDYN directly, but it supports involvement of the **PDYN-dynorphin-KOR axis** in addictive behavior. ([pubmed.ncbi.nlm.nih.gov](https://pubmed.ncbi.nlm.nih.gov/31026862/)) [S7]

## Contradicting or limiting evidence

The evidence is **not uniformly positive**, especially in human association studies. A 2023 case-control study in Turkish males found that **OPRK1**, but **not PDYN**, was associated with alcohol use disorder risk; the authors concluded PDYN did not appear associated with AUD, depression, anxiety, or craving in that sample. Another study found **no association between a PDYN VNTR and categorical alcohol dependence diagnosis**, although it did find an association with **disinhibited behavior**, a related trait. ([pubmed.ncbi.nlm.nih.gov](https://pubmed.ncbi.nlm.nih.gov/37177778/)) [S8]

The **human genetic literature is heterogeneous** by ancestry, sex, substance, and phenotype definition. Even the opioid-dependence meta-analysis that found a significant PDYN signal stressed that the number of studies was **insufficient for a robust and convincing conclusion**. More broadly, a recent review of genetic/epigenetic work in opioid use disorder lists **PDYN** among implicated candidates but explicitly calls for **large, ancestrally diverse GWAS and replication studies**, indicating that current evidence is still short of firm genomic consensus. ([pubmed.ncbi.nlm.nih.gov](https://pubmed.ncbi.nlm.nih.gov/31510971/)) [S9]

There is also an important conceptual limitation: much of the strongest evidence concerns the **dynorphin/KOR system** or **Pdyn-expressing neurons** rather than inherited PDYN variation per se. That still supports the user’s broad claim that “PDYN gene/protein is implicated,” but it is weaker support for narrower claims such as “PDYN is a major replicated human addiction-risk gene” or “PDYN variants are established clinical biomarkers.” ([pmc.ncbi.nlm.nih.gov](https://pmc.ncbi.nlm.nih.gov/articles/PMC9789200/)) [S10]

## Mechanistic / biological rationale

The mechanistic logic is strong. PDYN-derived dynorphins are the major endogenous ligands at **KOR**, a system widely linked to **aversion, dysphoria, stress responses, and anti-reward signaling**. Reviews of the dynorphin/KOR literature describe this system as particularly relevant to the **withdrawal/negative-affect** side of addiction rather than simply the acute pleasurable effects of drugs. ([ncbi.nlm.nih.gov](https://www.ncbi.nlm.nih.gov/gene/5173)) [S11]

In mesolimbic circuitry, repeated drug exposure can increase dopamine signaling, which in turn can drive **prodynorphin expression** in nucleus accumbens pathways; dynorphin then feeds back through KOR to **dampen dopamine-related reward signaling**. This is often interpreted as a **homeostatic/anti-reward adaptation**: initially counter-regulatory, but over time capable of contributing to dysphoria, stress sensitivity, compulsive use, and relapse vulnerability. ([pmc.ncbi.nlm.nih.gov](https://pmc.ncbi.nlm.nih.gov/articles/PMC2879894/?utm_source=openai)) [S12]

The withdrawal data fit that model. In mice, dynorphin release from **Pdyn neurons** contributes to opioid-withdrawal-related social dysfunction via KOR effects on serotonin signaling in the nucleus accumbens. In heroin-related human and rat PAC studies, reduced PDYN expression tracked with negative affect/stress circuitry, and experimental inhibition of PAC Pdyn neurons produced behavioral and physiological changes consistent with negative affect. Together, these findings make biological sense for addiction, especially the **maintenance and relapse** phases. ([pmc.ncbi.nlm.nih.gov](https://pmc.ncbi.nlm.nih.gov/articles/PMC9789200/)) [S13]

## Bottom line

The claim is **likely true**. PDYN is implicated in addiction by **multiple converging lines of evidence**: biologically it encodes dynorphin peptides central to KOR signaling; in animals, manipulating Pdyn alters alcohol intake and withdrawal-related behaviors; in humans, PDYN variation and PDYN expression changes have been associated with alcohol, opioid, cocaine, and methamphetamine-related phenotypes. ([ncbi.nlm.nih.gov](https://www.ncbi.nlm.nih.gov/gene/5173)) [S14]

However, the implication is **stronger mechanistically and preclinically than it is as a replicated human genetic risk factor**. Human association findings are mixed, often modest, and sometimes population- or phenotype-specific. So PDYN should be viewed as a **credible addiction-related gene/protein within the dynorphin-KOR stress/anti-reward system**, not as a definitively established universal addiction gene. ([pubmed.ncbi.nlm.nih.gov](https://pubmed.ncbi.nlm.nih.gov/31510971/)) [S15]

## Cited sources

- **[S1]** PDYN prodynorphin [Homo sapiens (human)] - Gene - NCBI — https://www.ncbi.nlm.nih.gov/gene/5173
- **[S2]** Association of the kappa-opioid system with alcohol dependence - PubMed — https://pubmed.ncbi.nlm.nih.gov/16924269/
- **[S3]** A functional haplotype implicated in vulnerability to develop cocaine dependence is associated with reduced PDYN expression in human brain - PubMed — https://pubmed.ncbi.nlm.nih.gov/18923396/
- **[S4]** Association between prodynorphin gene polymorphisms and opioid dependence susceptibility: a meta-analysis - PubMed — https://pubmed.ncbi.nlm.nih.gov/31510971/
- **[S5]** 
JCI -
Impaired periamygdaloid-cortex prodynorphin is characteristic of opiate addiction and depression
 — https://www.jci.org/articles/view/70395
- **[S6]** Increased ethanol intake in prodynorphin knockout mice is associated to changes in opioid receptor function and dopamine transmission - PubMed — https://pubmed.ncbi.nlm.nih.gov/21966993/
- **[S7]** Kappa-opioid receptors, dynorphin, and cocaine addiction: a positron emission tomography study - PubMed — https://pubmed.ncbi.nlm.nih.gov/31026862/
- **[S8]** Effect of PDYN and OPRK1 polymorphisms on the risk of alcohol use disorder and the intensity of depressive symptoms - PubMed — https://pubmed.ncbi.nlm.nih.gov/37177778/
- **[S9]** Association between prodynorphin gene polymorphisms and opioid dependence susceptibility: a meta-analysis - PubMed — https://pubmed.ncbi.nlm.nih.gov/31510971/
- **[S10]** 
            Modulation of 5-HT release by dynorphin mediates social deficits during opioid withdrawal - PMC
         — https://pmc.ncbi.nlm.nih.gov/articles/PMC9789200/
- **[S11]** PDYN prodynorphin [Homo sapiens (human)] - Gene - NCBI — https://www.ncbi.nlm.nih.gov/gene/5173
- **[S12]** The role of the dynorphin–κ opioid system in the reinforcing effects of drugs of abuse - PMC — https://pmc.ncbi.nlm.nih.gov/articles/PMC2879894/?utm_source=openai
- **[S13]** 
            Modulation of 5-HT release by dynorphin mediates social deficits during opioid withdrawal - PMC
         — https://pmc.ncbi.nlm.nih.gov/articles/PMC9789200/
- **[S14]** PDYN prodynorphin [Homo sapiens (human)] - Gene - NCBI — https://www.ncbi.nlm.nih.gov/gene/5173
- **[S15]** Association between prodynorphin gene polymorphisms and opioid dependence susceptibility: a meta-analysis - PubMed — https://pubmed.ncbi.nlm.nih.gov/31510971/
