## Supplementary material for "Understanding Substance Dependence: What Differentiates Addictive from Non-Addictive Drugs?": Folder for supplementary files: addictive-genes-claims-first100-092-tle4-gene-protein-is-implicated-in-addiction-cdc523831a17.md

# addictive-genes-claims-first100-092-cdc523831a17

**Original claim:** TLE4 gene/protein is implicated in addiction.

## Claim

TLE4 gene/protein is implicated in addiction.

## Verdict

Verdict: Mixed Evidence

There is **some evidence** linking TLE4 to addiction-related biology, especially in **opioid** models and human **opioid use disorder (OUD)** transcriptomics, but it is **not yet a well-established addiction gene/protein**. The current support is mostly **observational**: gene-expression correlations in rat nucleus accumbens after oxycodone self-administration, cell-type-specific differential expression in human/primate striatum in OUD, and weaker alcohol-related omics findings. I did **not** find strong evidence from **causal perturbation studies** showing that changing TLE4 itself alters drug reward, intake, or relapse, nor did I find major replicated human GWAS signals placing TLE4 among the leading addiction-risk loci. ([pubmed.ncbi.nlm.nih.gov](https://pubmed.ncbi.nlm.nih.gov/38710443/)) [S1]

## Confidence

Medium

## Supporting evidence

The strongest addiction-relevant signal I found is a **2024 rat oxycodone self-administration study**. Male rats given long-access oxycodone developed addiction-like behaviors, and **nucleus accumbens tle4 expression was strongly negatively correlated with reinstatement behavior** after abstinence; tle4 also negatively correlated with total oxycodone intake, and miR-182 negatively correlated with tle4. The authors explicitly interpreted this as suggesting a possible role for tle4 in maintaining an addiction-like brain state, while noting the need for direct manipulation studies. ([pubmed.ncbi.nlm.nih.gov](https://pubmed.ncbi.nlm.nih.gov/38710443/)) [S2]

There is also **human tissue evidence**. In a **single-nucleus transcriptomic study of human and non-human primate striatum in OUD**, TLE4 was among the **top downregulated differentially expressed genes** in specific neuronal subtypes, including interneurons, within pathway clusters related to cellular stress/DNA-damage biology. This does not prove causality, but it does place TLE4 inside an OUD-associated transcriptional signature in a core addiction-relevant brain region. ([pmc.ncbi.nlm.nih.gov](https://pmc.ncbi.nlm.nih.gov/articles/PMC10831093/)) [S3]

A weaker, older line of support comes from **alcohol-related omics**. In alcohol-preferring rats, **Tle4** appeared among genes with altered nucleus accumbens expression in the **ethanol vs. saccharin** comparison. Separately, a chronic ethanol proteomics study reported **TLE4** among top altered proteins in mouse striatum. These findings are compatible with substance-related regulation of TLE4, though they are much less persuasive than the opioid data. ([pmc.ncbi.nlm.nih.gov](https://pmc.ncbi.nlm.nih.gov/articles/PMC4516281/)) [S4]

Biological plausibility is also present. TLE4 is a **transcriptional corepressor** involved in **negative regulation of canonical Wnt signaling/transcription**, and brain studies show that **Tle4 is expressed in striatal medium spiny neurons** as well as corticothalamic cortical neurons. Because addiction involves durable transcriptional and synaptic remodeling in cortico-striatal circuits, a transcriptional corepressor expressed in those neurons is a plausible participant in the process. ([ncbi.nlm.nih.gov](https://www.ncbi.nlm.nih.gov/gene/7091?utm_source=openai)) [S5]

## Contradicting or limiting evidence

The main limitation is that the evidence is **mostly correlational**. In the oxycodone rat study, tle4 was **measured**, not manipulated; the paper states that it is **unclear** whether the observed expression pattern reflects a **compensatory response** to oxycodone exposure or a factor contributing to susceptibility/severity. That is an important distinction for the claim “is implicated in addiction.” ([pmc.ncbi.nlm.nih.gov](https://pmc.ncbi.nlm.nih.gov/articles/PMC11164234/?utm_source=openai)) [S6]

The alcohol transcriptomic support is also **weak/exploratory**. In the alcohol-preferring rat study, the **ethanol vs. water** comparison had an **FDR of 1.0**, and the **Tle4 signal was in the ethanol vs. saccharin comparison**, which had **FDR 0.5–0.8** overall; the authors selected other genes, not Tle4, for qRT-PCR confirmation. So this is not robust validation of TLE4 as an alcohol-addiction driver. ([pmc.ncbi.nlm.nih.gov](https://pmc.ncbi.nlm.nih.gov/articles/PMC4516281/)) [S7]

The human OUD transcriptomic study is valuable, but it is still **postmortem observational biology**. Differential expression can reflect many things besides causal disease mechanisms, including chronic drug exposure, overdose-related stress, comorbidities, or downstream adaptations. In that study, TLE4 appears as one of several genes within a broader stress-related signature rather than as a singular standout mechanistic node. ([pmc.ncbi.nlm.nih.gov](https://pmc.ncbi.nlm.nih.gov/articles/PMC10831093/)) [S8]

Large human genetic studies also **do not currently elevate TLE4 as a leading addiction-risk locus**. A large OUD GWAS/meta-analysis identified genome-wide significant signals at **OPRM1** and **FURIN** and reported 19 independent significant loci overall; a large trans-ancestral alcohol dependence GWAS highlighted **ADH1B**. These studies do not support TLE4 as a major, replicated common-variant risk gene for OUD or alcohol dependence. ([pubmed.ncbi.nlm.nih.gov](https://pubmed.ncbi.nlm.nih.gov/35879402/)) [S9]

## Mechanistic / biological rationale

Mechanistically, TLE4 is a reasonable candidate for influencing addiction-related neuroadaptation. It encodes a **non-DNA-binding transcriptional corepressor** and is predicted to participate in **Wnt-related transcriptional repression**. In the nervous system, Tle4 is required for **acquisition and postnatal stability of corticothalamic neuron identity**, and broader cell-atlas work shows Tle4 expression in **striatal medium spiny neurons**. Developmental striatal studies also identify **Grg4/Tle4** among **striatopallidal neuron-specific genes**, alongside markers such as **Drd2**, **Penk**, and **Adora2a**. Those are all highly relevant circuits/cell classes for reward learning, behavioral inhibition, withdrawal-related plasticity, and relapse. ([pmc.ncbi.nlm.nih.gov](https://pmc.ncbi.nlm.nih.gov/articles/PMC10542749/?utm_source=openai)) [S10]

This means a plausible hypothesis is that **reduced TLE4 could alter transcriptional programs that stabilize neuronal identity or synaptic homeostasis in cortico-striatal pathways**, thereby affecting addiction vulnerability or persistence. The convergence between the rat oxycodone study—where **lower tle4 tracked worse reinstatement-related phenotypes**—and the human OUD striatal study—where **TLE4 was downregulated in certain neuronal populations**—is suggestive, though still not definitive. ([pubmed.ncbi.nlm.nih.gov](https://pubmed.ncbi.nlm.nih.gov/38710443/)) [S11]

There is also an **indirect neuroimmune rationale**. Outside the brain, TLE4 represses inflammatory cytokine programs such as **IL-6** and **IL-12** in macrophages, while addiction research increasingly implicates **microglia/neuroimmune signaling** in the progression of drug addiction. That does **not** show TLE4 is an addiction protein, but it provides a biologically coherent route by which TLE4-related transcriptional control could intersect with addiction-relevant inflammatory states. This remains speculative until shown directly in addiction models. ([pmc.ncbi.nlm.nih.gov](https://pmc.ncbi.nlm.nih.gov/articles/PMC6418302/?utm_source=openai)) [S12]

## Bottom line

If “implicated” means **there are credible data associating TLE4 with addiction-related phenotypes**, then the claim is **partly supported**—most clearly for **opioid-related** phenotypes. If it means **TLE4 is an established causal addiction gene/protein**, the evidence is **not there yet**. Overall, TLE4 looks like a **plausible, emerging candidate** in addiction biology rather than a confirmed core addiction gene. ([pubmed.ncbi.nlm.nih.gov](https://pubmed.ncbi.nlm.nih.gov/38710443/)) [S13]

## Cited sources

- **[S1]** Neuroplasticity-related genes correlate with individual differences in distinct phases of oxycodone self-administration in male rats - PubMed — https://pubmed.ncbi.nlm.nih.gov/38710443/
- **[S2]** Neuroplasticity-related genes correlate with individual differences in distinct phases of oxycodone self-administration in male rats - PubMed — https://pubmed.ncbi.nlm.nih.gov/38710443/
- **[S3]** 
            Single nuclei transcriptomics in human and non-human primate striatum in opioid use disorder - PMC
         — https://pmc.ncbi.nlm.nih.gov/articles/PMC10831093/
- **[S4]** 
            Differential gene expression in the nucleus accumbens with ethanol self-administration in inbred alcohol-preferring rats - PMC
         — https://pmc.ncbi.nlm.nih.gov/articles/PMC4516281/
- **[S5]** TLE4 TLE family member 4, transcriptional corepressor [Homo sapiens (human)] - Gene - NCBI — https://www.ncbi.nlm.nih.gov/gene/7091?utm_source=openai
- **[S6]** Neuroplasticity-Related Genes Correlate with Individual Differences in Distinct Phases of Oxycodone Self-Administration in Male Rats - PMC — https://pmc.ncbi.nlm.nih.gov/articles/PMC11164234/?utm_source=openai
- **[S7]** 
            Differential gene expression in the nucleus accumbens with ethanol self-administration in inbred alcohol-preferring rats - PMC
         — https://pmc.ncbi.nlm.nih.gov/articles/PMC4516281/
- **[S8]** 
            Single nuclei transcriptomics in human and non-human primate striatum in opioid use disorder - PMC
         — https://pmc.ncbi.nlm.nih.gov/articles/PMC10831093/
- **[S9]** Genome-wide association study in individuals of European and African ancestry and multi-trait analysis of opioid use disorder identifies 19 independent genome-wide significant risk loci - PubMed — https://pubmed.ncbi.nlm.nih.gov/35879402/
- **[S10]** Tle4 controls both developmental acquisition and early post-natal maturation of corticothalamic projection neuron identity - PMC — https://pmc.ncbi.nlm.nih.gov/articles/PMC10542749/?utm_source=openai
- **[S11]** Neuroplasticity-related genes correlate with individual differences in distinct phases of oxycodone self-administration in male rats - PubMed — https://pubmed.ncbi.nlm.nih.gov/38710443/
- **[S12]** TLE4 acts as a corepressor of Hes1 to inhibit inflammatory responses in macrophages - PMC — https://pmc.ncbi.nlm.nih.gov/articles/PMC6418302/?utm_source=openai
- **[S13]** Neuroplasticity-related genes correlate with individual differences in distinct phases of oxycodone self-administration in male rats - PubMed — https://pubmed.ncbi.nlm.nih.gov/38710443/
