## Supplementary material for "Understanding Substance Dependence: What Differentiates Addictive from Non-Addictive Drugs?": Folder for supplementary files: addictive-genes-claims-first100-093-wsb2-gene-protein-is-implicated-in-addiction-e36f13d9fc21.md

# addictive-genes-claims-first100-093-e36f13d9fc21

**Original claim:** WSB2 gene/protein is implicated in addiction.

## Claim

WSB2 gene/protein is implicated in addiction.

## Verdict

Verdict: No Evidence

## Confidence

Medium. The gene is biologically plausible in a very broad sense because it is brain-expressed and participates in ubiquitin-mediated protein regulation, but I did not find direct peer-reviewed evidence tying WSB2 itself to addiction phenotypes, substance use disorder risk, or addiction-relevant neural adaptations. ([ncbi.nlm.nih.gov](https://www.ncbi.nlm.nih.gov/gene/55884)) [S1]

## Supporting evidence

The strongest points **in favor** of the claim are indirect rather than addiction-specific. Human NCBI and Human Protein Atlas resources indicate that WSB2 is broadly expressed, including in brain/CNS, and mouse annotations likewise note expression in the central nervous system. That means the gene is at least present in tissues relevant to addiction biology. ([ncbi.nlm.nih.gov](https://www.ncbi.nlm.nih.gov/gene/55884)) [S2]

WSB2 also has bona fide molecular functions that could, in principle, matter for neural plasticity: it encodes a WD-repeat/SOCS-box protein that serves as a substrate-recognition component for Cullin-RING E3 ubiquitin ligase machinery, and curated annotations link it to protein ubiquitination/polyubiquitination. Ubiquitin-dependent protein turnover is broadly important in synaptic remodeling and long-term cellular adaptation, both of which are relevant to addiction neuroscience at a systems level. However, this is mechanistic plausibility, not direct addiction evidence. ([ncbi.nlm.nih.gov](https://www.ncbi.nlm.nih.gov/gene/55884)) [S3]

A recent human genetics paper showed that recessive WSB2 variants can cause a neurodevelopmental syndrome with developmental delay, seizures, hypotonia, abnormal brain morphology, and failure to thrive, and the authors noted broad brain expression with neocortical enrichment in human single-cell datasets. This supports the idea that WSB2 has genuine neural importance. But again, neurodevelopmental relevance is not the same as implication in addiction. ([nature.com](https://www.nature.com/articles/s41431-025-01863-4)) [S4]

There is also at least one peripheral epigenetic exposure signal: a study of sperm DNA methylation in smokers reported a small methylation difference at a WSB2-annotated CpG. That finding shows WSB2 can appear in smoking-associated omics data, but it does **not** establish a role in nicotine dependence, reward circuitry, craving, withdrawal, or substance use disorder pathogenesis. It is best interpreted as a weak, exposure-associated peripheral biomarker signal. ([pmc.ncbi.nlm.nih.gov](https://pmc.ncbi.nlm.nih.gov/articles/PMC5679018/)) [S5]

## Contradicting or limiting evidence

The main limitation is simple: I did **not** find a direct body of literature connecting WSB2 to addiction. In curated NCBI Gene references, the named functional links for WSB2 focus on NF-κB termination, p53 degradation, breast cancer migration, melanoma biology, and receptor trafficking rather than addiction phenotypes. Likewise, the recent European Journal of Human Genetics paper on WSB2 states that prior work on WSB2 had mainly focused on cancer tissues/cells and a limited set of substrates, not substance use disorders. ([ncbi.nlm.nih.gov](https://www.ncbi.nlm.nih.gov/gene/55884)) [S6]

I also did not identify a peer-reviewed human genetic association study showing WSB2 as a replicated risk gene for alcohol use disorder, nicotine dependence, opioid use disorder, cannabis use disorder, or a cross-substance addiction factor. Recent reviews and large GWAS summaries of substance use disorders emphasize other loci/genes—such as DRD2, PDE4B, CHRNB2, OPRM1, and FURIN—and discuss general addiction-risk loci without highlighting WSB2 as an established signal. That does not prove WSB2 has no role, but it strongly suggests it is not currently recognized as a replicated addiction gene. ([nature.com](https://www.nature.com/articles/s41380-024-02547-z)) [S7]

I likewise did not find an animal study manipulating Wsb2 and measuring canonical addiction-related behaviors such as drug self-administration, conditioned place preference for abused drugs, escalation of intake, cue-induced reinstatement, withdrawal, or compulsive-like use. Available mouse annotations instead emphasize expression patterns and general molecular roles, not addiction phenotyping. ([ncbi.nlm.nih.gov](https://www.ncbi.nlm.nih.gov/gene/59043)) [S8]

Finally, the few hints that touch substance exposure are weak and non-specific. The smoking-associated sperm methylation signal is peripheral and modest, and an in silico study linked WSB2 to smoking-related phenotypes in cancer datasets rather than to addiction biology itself. These are far below the evidentiary bar needed to say a gene is “implicated in addiction.” ([pmc.ncbi.nlm.nih.gov](https://pmc.ncbi.nlm.nih.gov/articles/PMC5679018/)) [S9]

## Mechanistic / biological rationale

If one wanted to make a **hypothesis** for future study, there is a plausible route. WSB2 is a substrate receptor in ubiquitin-ligase biology, and curated/experimental work links it to degradation of specific proteins, including methylated RelA/NF-κB pathway components; other work links WSB2 to p53 and NOXA regulation in non-neural, mostly cancer contexts. Because ubiquitin-proteasome signaling, inflammatory transcriptional control, and protein-stability programs can shape neuronal homeostasis and plasticity, WSB2 could conceivably influence addiction-relevant adaptations someday if tested directly. ([pubmed.ncbi.nlm.nih.gov](https://pubmed.ncbi.nlm.nih.gov/38452206/)) [S10]

But that remains speculative. The currently documented WSB2 substrates and phenotypes are mostly derived from cancer, immune, or developmental studies, not from reward circuitry, dopaminergic signaling, synaptic transmission, or human SUD cohorts. Even the neurodevelopmental syndrome data support a role in brain development/function broadly, not addiction specifically. So the mechanistic rationale is “possible in theory,” not “demonstrated in addiction.” ([nature.com](https://www.nature.com/articles/s41431-025-01863-4)) [S11]

## Bottom line

At present, the claim that **WSB2 is implicated in addiction** is **not supported by direct evidence**. WSB2 is a real brain-expressed ubiquitin-pathway gene with emerging importance in neurodevelopment and established roles in protein turnover, NF-κB signaling, and cancer biology, so it is not biologically absurd as a future candidate. But based on the literature I found, it is **not currently an established addiction gene/protein**, and the available signals are indirect, peripheral, or from non-addiction contexts. ([ncbi.nlm.nih.gov](https://www.ncbi.nlm.nih.gov/gene/55884)) [S12]

## Cited sources

- **[S1]** https://www.ncbi.nlm.nih.gov/gene/55884 — https://www.ncbi.nlm.nih.gov/gene/55884
- **[S2]** https://www.ncbi.nlm.nih.gov/gene/55884 — https://www.ncbi.nlm.nih.gov/gene/55884
- **[S3]** https://www.ncbi.nlm.nih.gov/gene/55884 — https://www.ncbi.nlm.nih.gov/gene/55884
- **[S4]** https://www.nature.com/articles/s41431-025-01863-4 — https://www.nature.com/articles/s41431-025-01863-4
- **[S5]** https://pmc.ncbi.nlm.nih.gov/articles/PMC5679018/ — https://pmc.ncbi.nlm.nih.gov/articles/PMC5679018/
- **[S6]** https://www.ncbi.nlm.nih.gov/gene/55884 — https://www.ncbi.nlm.nih.gov/gene/55884
- **[S7]** https://www.nature.com/articles/s41380-024-02547-z — https://www.nature.com/articles/s41380-024-02547-z
- **[S8]** https://www.ncbi.nlm.nih.gov/gene/59043 — https://www.ncbi.nlm.nih.gov/gene/59043
- **[S9]** https://pmc.ncbi.nlm.nih.gov/articles/PMC5679018/ — https://pmc.ncbi.nlm.nih.gov/articles/PMC5679018/
- **[S10]** https://pubmed.ncbi.nlm.nih.gov/38452206/ — https://pubmed.ncbi.nlm.nih.gov/38452206/
- **[S11]** https://www.nature.com/articles/s41431-025-01863-4 — https://www.nature.com/articles/s41431-025-01863-4
- **[S12]** https://www.ncbi.nlm.nih.gov/gene/55884 — https://www.ncbi.nlm.nih.gov/gene/55884
