## Supplementary material for "Understanding Substance Dependence: What Differentiates Addictive from Non-Addictive Drugs?": Folder for supplementary files: addictive-genes-claims-first100-094-fabp7-gene-protein-is-implicated-in-addiction-a52683607b44.md

# addictive-genes-claims-first100-094-a52683607b44

**Original claim:** FABP7 gene/protein is implicated in addiction.

## Claim

FABP7 gene/protein is implicated in addiction.

## Verdict

Verdict: Likely True

## Confidence

Medium

## Supporting evidence

FABP7 is a brain-enriched, largely astrocyte-associated fatty-acid binding protein involved in lipid trafficking, endocannabinoid-related signaling, stress responses, sleep/circadian regulation, and synaptic/metabolic homeostasis rather than being a classic “reward gene” like *FOSB* or *OPRM1*. That matters because addiction biology is not only neuronal reward signaling; it also includes glial, stress, immune, and homeostatic processes. FABP7 is expressed in astrocytes and neural progenitors, and human/mouse studies show that altered FABP7 function changes sleep architecture and stress-related biology. ([pmc.ncbi.nlm.nih.gov](https://pmc.ncbi.nlm.nih.gov/articles/PMC5381954/)) [S1]

The strongest direct addiction-relevant evidence is in opioids. In a 2023 *Cell* study, blood from people with opioid use disorder showed expansion of fragile-like regulatory T cells, and the abundance of these cells correlated positively with withdrawal severity. In the corresponding mouse experiments, opioid exposure reduced astrocyte-derived Fabp7, increased blood-brain barrier permeability, promoted Treg infiltration into the nucleus accumbens, altered synaptic structure via IFN-γ signaling, and worsened withdrawal symptoms. The paper’s summary explicitly states that opioids disrupted BBB integrity through downregulation of astrocyte-derived Fabp7, and follow-up commentary notes that rescuing astrocyte-derived Fabp7 improved synaptic stability and withdrawal phenotypes. This is not merely correlation; it places Fabp7 in a causal pathway for an addiction-relevant outcome: opioid withdrawal. ([sciencedirect.com](https://www.sciencedirect.com/science/article/pii/S0092867422015732)) [S2]

There is also preclinical evidence linking FABP7 to cocaine-related vulnerability. In a mouse model where early-life social stress increased adult cocaine-seeking and reinstatement after withdrawal, nucleus accumbens RNA-seq identified *Fabp7* among a small set of candidate transcripts associated with this heightened relapse vulnerability, and the authors specifically highlighted FABP7 as a candidate target for vulnerability to cocaine addiction. This supports implication, although it is still associative rather than definitive proof of causality. ([pubmed.ncbi.nlm.nih.gov](https://pubmed.ncbi.nlm.nih.gov/26706499/?utm_source=openai)) [S3]

Additional support comes from studies manipulating brain FABPs that include FABP7. In FABP5/7 double-knockout mice, baseline acquisition of cocaine conditioned place preference was preserved, but stress-induced reinstatement of cocaine preference was abolished, and the knockout mice showed blunted corticosterone responses to stress. In a separate study, pharmacologic inhibition of FABP5/7 reduced ethanol consumption in mice. These findings suggest that brain FABPs, including FABP7, can influence stress-linked drug seeking and alcohol intake. However, because these experiments were not FABP7-selective, they support involvement of the pathway more strongly than they prove a unique role for FABP7 itself. ([pubmed.ncbi.nlm.nih.gov](https://pubmed.ncbi.nlm.nih.gov/29457656/?utm_source=openai)) [S4]

More recent FABP7-specific mechanistic work strengthens biological plausibility. A 2025 mouse study found that genetic deletion of *Fabp7* reduced CB1 receptor binding in several brain regions and described FABP7 as participating in endocannabinoid transport and stress regulation. Since CB1/endocannabinoid signaling is broadly involved in reward, stress responsivity, withdrawal, and relapse-related behavior, this places FABP7 upstream of a major addiction-relevant signaling system. ([pubmed.ncbi.nlm.nih.gov](https://pubmed.ncbi.nlm.nih.gov/39542341/)) [S5]

## Contradicting or limiting evidence

The main limitation is that most of the evidence is preclinical or mechanistic, not robustly clinical. I did not find strong, replicated human genetic evidence showing that common or rare *FABP7* variants are a major risk factor for substance use disorders across populations. Nor did I find large human studies establishing FABP7 as a diagnostic biomarker or treatment target for addiction itself. The human evidence is strongest for opioid withdrawal biology, not for addiction broadly defined across substances. ([sciencedirect.com](https://www.sciencedirect.com/science/article/pii/S0092867422015732)) [S6]

Also, several “positive” studies involve combined FABP5/7 deletion or inhibition rather than isolated FABP7 manipulation. That means some behavioral effects on cocaine stress-reinstatement or ethanol intake could be driven partly, or even mainly, by FABP5 or by shared endocannabinoid transport effects rather than FABP7 specifically. ([pubmed.ncbi.nlm.nih.gov](https://pubmed.ncbi.nlm.nih.gov/29457656/?utm_source=openai)) [S7]

A recent narrative review of FABPs in neuropsychiatric disease explicitly states that the FABP–SUD link is “not sufficiently elucidated.” In that review’s summary, FABP3—not FABP7—was the subtype more clearly tied to dopamine receptor regulation linked to ADHD and substance use disorders, whereas elevated FABP7 was emphasized more for major depression and autism spectrum disorder. That does not negate an addiction role for FABP7, but it argues against overstating the case. ([pubmed.ncbi.nlm.nih.gov](https://pubmed.ncbi.nlm.nih.gov/40613283/)) [S8]

## Mechanistic / biological rationale

A biologically coherent case can be made for FABP7 in addiction because it sits at the intersection of several systems already known to matter in substance use disorders. First, FABP7 participates in intracellular lipid trafficking and endocannabinoid handling. The endocannabinoid system is well established as a modulator of reward processing, emotional homeostasis, drug reinforcement, withdrawal-related states, and relapse-like behavior. If FABP7 changes endocannabinoid availability or CB1 receptor biology, it could plausibly alter addiction-related phenotypes even without being a “primary addiction gene.” ([pubmed.ncbi.nlm.nih.gov](https://pubmed.ncbi.nlm.nih.gov/39542341/)) [S9]

Second, FABP7 is an astrocytic homeostasis gene tied to sleep and circadian regulation. Human carriers of a FABP7 missense variant had more fragmented sleep, and Fabp7-deficient mice likewise showed fragmented sleep. Independent reviews of addiction biology show that circadian disruption and sleep disturbance can increase vulnerability to substance use disorders and relapse. So FABP7 could contribute to addiction risk indirectly by altering sleep/circadian stability and stress responsiveness. ([pmc.ncbi.nlm.nih.gov](https://pmc.ncbi.nlm.nih.gov/articles/PMC5381954/)) [S10]

Third, the opioid data suggest a distinctly glial/neuroimmune mechanism: astrocytic Fabp7 helps maintain BBB integrity, and loss of that protection permits immune-cell entry into reward circuitry, leading to synaptic remodeling and worse withdrawal. That is especially notable because addiction research increasingly recognizes BBB dysfunction, neuroinflammation, and glial signaling as part of the disease process, particularly in negative affect and withdrawal states. ([sciencedirect.com](https://www.sciencedirect.com/science/article/pii/S0092867422015732)) [S11]

## Bottom line

The claim is **likely true**, but with an important qualifier: **FABP7 is implicated in addiction-related biology more convincingly than it is established as a broad, primary human addiction gene**. The best evidence is for **opioid withdrawal/neuroimmune mechanisms**, with additional **preclinical links to cocaine relapse vulnerability, stress-induced drug seeking, alcohol intake, and endocannabinoid/CB1 signaling**. What is still missing are large, replicated human genetic or clinical studies showing that FABP7 has a major, substance-general effect on addiction risk or treatment response. ([sciencedirect.com](https://www.sciencedirect.com/science/article/pii/S0092867422015732)) [S12]

## Cited sources

- **[S1]** 
            Normal sleep requires the astrocyte brain-type fatty acid binding protein FABP7 - PMC
         — https://pmc.ncbi.nlm.nih.gov/articles/PMC5381954/
- **[S2]** Opioid-induced fragile-like regulatory T cells contribute to withdrawal - ScienceDirect — https://www.sciencedirect.com/science/article/pii/S0092867422015732
- **[S3]** Regulation of nucleus accumbens transcript levels in mice by early-life social stress and cocaine - PubMed — https://pubmed.ncbi.nlm.nih.gov/26706499/?utm_source=openai
- **[S4]** Fatty acid binding protein deletion prevents stress-induced preference for cocaine and dampens stress-induced corticosterone levels - PubMed — https://pubmed.ncbi.nlm.nih.gov/29457656/?utm_source=openai
- **[S5]** Fatty acid-binding protein 7 gene deletion promotes decreases in brain cannabinoid type 1 receptor binding - PubMed — https://pubmed.ncbi.nlm.nih.gov/39542341/
- **[S6]** Opioid-induced fragile-like regulatory T cells contribute to withdrawal - ScienceDirect — https://www.sciencedirect.com/science/article/pii/S0092867422015732
- **[S7]** Fatty acid binding protein deletion prevents stress-induced preference for cocaine and dampens stress-induced corticosterone levels - PubMed — https://pubmed.ncbi.nlm.nih.gov/29457656/?utm_source=openai
- **[S8]** The Role of Fatty Acid Binding Proteins in Neuropsychiatric Diseases: A Narrative Review - PubMed — https://pubmed.ncbi.nlm.nih.gov/40613283/
- **[S9]** Fatty acid-binding protein 7 gene deletion promotes decreases in brain cannabinoid type 1 receptor binding - PubMed — https://pubmed.ncbi.nlm.nih.gov/39542341/
- **[S10]** 
            Normal sleep requires the astrocyte brain-type fatty acid binding protein FABP7 - PMC
         — https://pmc.ncbi.nlm.nih.gov/articles/PMC5381954/
- **[S11]** Opioid-induced fragile-like regulatory T cells contribute to withdrawal - ScienceDirect — https://www.sciencedirect.com/science/article/pii/S0092867422015732
- **[S12]** Opioid-induced fragile-like regulatory T cells contribute to withdrawal - ScienceDirect — https://www.sciencedirect.com/science/article/pii/S0092867422015732
