## Supplementary material for "Understanding Substance Dependence: What Differentiates Addictive from Non-Addictive Drugs?": Folder for supplementary files: addictive-genes-claims-first100-095-gpr37-gene-protein-is-implicated-in-addiction-9d4e5ea99fa6.md

# addictive-genes-claims-first100-095-9d4e5ea99fa6

**Original claim:** GPR37 gene/protein is implicated in addiction.

## Claim

GPR37 gene/protein is implicated in addiction.

## Verdict

Verdict: Likely True

The best available evidence supports **a role for GPR37 in addiction-related neurobiology, especially for psychostimulants such as cocaine and amphetamine, in rodent models**. Multiple mouse studies link GPR37 loss to altered dopamine transporter function, changed behavioral responses to psychostimulants, absent cocaine/amphetamine conditioned place preference, impaired cocaine-induced ΔFosB induction, and altered corticostriatal synaptic effects of cocaine. However, this evidence is **mostly preclinical**, and I did **not** find strong human genetic, clinical, or treatment data establishing GPR37 as a confirmed addiction gene in people. ([pubmed.ncbi.nlm.nih.gov](https://pubmed.ncbi.nlm.nih.gov/17519329/)) [S1]

## Confidence

Medium

There are several converging rodent studies pointing in the same general direction, which is stronger than a single isolated finding. But the literature is still narrow: it is centered on psychostimulants, uses mostly knockout mice and related mechanistic experiments, and lacks comparable human evidence. ([pubmed.ncbi.nlm.nih.gov](https://pubmed.ncbi.nlm.nih.gov/17519329/)) [S2]

## Supporting evidence

A foundational line of evidence is that **GPR37 directly interfaces with dopamine signaling**, which is central to psychostimulant reward. In a 2007 PNAS study, GPR37 was found to colocalize with the dopamine transporter (DAT) in mouse striatal presynaptic membranes and to physically interact with DAT. Gpr37-null mice had increased plasma-membrane DAT and enhanced DAT-mediated dopamine uptake, along with reduced cocaine-induced locomotor activity. An earlier 2004 PNAS paper also found that Gpr37-knockout mice had altered nigrostriatal dopaminergic signaling, reduced striatal dopamine content, and enhanced amphetamine sensitivity. Together, these findings place GPR37 squarely in a dopamine-regulatory pathway highly relevant to stimulant reward and reinforcement. ([pubmed.ncbi.nlm.nih.gov](https://pubmed.ncbi.nlm.nih.gov/17519329/)) [S3]

The most direct addiction-relevant behavioral evidence comes from a 2011 FASEB Journal study. In that work, **GPR37-knockout mice failed to show conditioned place preference (CPP) to either amphetamine or cocaine**, and acute psychostimulant treatment no longer produced the expected ERK2 response seen in wild-type mice. Chronic cocaine also failed to increase ΔFosB in the knockout animals. Because CPP is a well-established model for drug reward/addiction-like behavior, and ΔFosB is a classic molecular mediator of long-lasting drug-induced adaptations, this is strong preclinical evidence that GPR37 contributes to psychostimulant reward signaling. ([pubmed.ncbi.nlm.nih.gov](https://pubmed.ncbi.nlm.nih.gov/21372109/)) [S4]

A third, independent line of support comes from synaptic physiology. A 2017 Neuroscience Letters study found that cocaine decreased corticostriatal synaptic transmission in both wild-type and GPR37-knockout mouse slices, but the effect was **larger in the knockout**. The authors concluded that GPR37 is involved in cocaine-induced modification of basal synaptic transmission. Since addiction involves drug-driven remodeling of corticostriatal circuits, this strengthens the case that GPR37 participates in psychostimulant-induced neuroplasticity rather than only affecting general locomotion. ([pubmed.ncbi.nlm.nih.gov](https://pubmed.ncbi.nlm.nih.gov/28007645/)) [S5]

There is also **more indirect, newer evidence** extending the signal to methamphetamine-related adaptations. A 2024 study of methamphetamine sensitization in mouse nucleus accumbens identified **Gpr37 as a candidate target** in a miR-29c-3p-regulated network, and reported aberrant Gpr37 expression in the nucleus accumbens during methamphetamine sensitization. This does not prove causality for GPR37 itself, but it is directionally consistent with a broader role in stimulant-related neuroadaptation. ([pubmed.ncbi.nlm.nih.gov](https://pubmed.ncbi.nlm.nih.gov/38256016/?utm_source=openai)) [S6]

## Contradicting or limiting evidence

The biggest limitation is that **the evidence is overwhelmingly from animal and mechanistic studies**, not humans. In a very large 2023 multivariate GWAS/meta-analysis of substance use disorders involving over 1 million participants, the main reported signals included genes such as **PDE4B, ADH1B, CHRNA5, OPRM1, and CACNA1A**; I did not find GPR37 mentioned in the article text. That does not prove GPR37 is irrelevant in humans, but it does mean it is **not currently established as a major replicated human SUD locus** on the level of better-known addiction genes. ([pmc.ncbi.nlm.nih.gov](https://pmc.ncbi.nlm.nih.gov/articles/PMC10217792/)) [S7]

A second limitation is that the phenotype is **complex rather than uniformly “pro-addiction” or “anti-addiction.”** Gpr37 loss increases amphetamine sensitivity in one study, reduces cocaine-induced locomotor activity in another, and abolishes cocaine/amphetamine CPP in a third. Those findings are compatible with an important modulatory role, but they do not map cleanly onto a simple one-direction narrative such as “more GPR37 always increases addiction risk” or “less GPR37 is always protective.” ([pubmed.ncbi.nlm.nih.gov](https://pubmed.ncbi.nlm.nih.gov/15218106/)) [S8]

A third limitation is scope. The direct evidence is strongest for **cocaine and amphetamine**, with only indirect support for methamphetamine, and I did not find comparably direct evidence for opioid, alcohol, nicotine, or cannabis addiction. Recent GPR37 reviews still emphasize Parkinson’s disease, pain, inflammation, autism, and other CNS roles much more than addiction, underscoring that addiction is not yet one of the receptor’s most firmly established disease domains. ([pmc.ncbi.nlm.nih.gov](https://pmc.ncbi.nlm.nih.gov/articles/PMC11144446/)) [S9]

Even within cocaine-related physiology, effects are selective. The 2017 synaptic study found altered **basal synaptic transmission** responses to cocaine but **no genotype difference in paired-pulse ratio**, suggesting that GPR37 does not broadly control every form of cocaine-related synaptic plasticity. ([pubmed.ncbi.nlm.nih.gov](https://pubmed.ncbi.nlm.nih.gov/28007645/)) [S10]

## Mechanistic / biological rationale

Biologically, the claim is plausible. In humans, GPR37 is expressed in the **basal ganglia, including nucleus accumbens**, and in mice it is enriched in the **striatal presynaptic fraction**, where it colocalizes with DAT. Those are exactly the kinds of regions and compartments expected for a protein influencing stimulant reward circuitry. ([proteinatlas.org](https://www.proteinatlas.org/ENSG00000170775-GPR37/brain/basal%2Bganglia)) [S11]

Mechanistically, GPR37 appears to sit at a crossroads of several addiction-relevant signaling systems. It modulates **DAT trafficking and dopamine uptake**, interacts with **dopamine D2 receptors**, and also forms functionally relevant associations with **adenosine A2A receptors** in striatum. In GPR37-deficient mice, A2A receptor surface expression/function is enhanced. These interactions matter because DAT is central to the reinforcing actions of cocaine and related psychostimulants, while A2A signaling has documented roles in drug-seeking and addiction-related striatal signaling. ([pubmed.ncbi.nlm.nih.gov](https://pubmed.ncbi.nlm.nih.gov/17519329/)) [S12]

So, as an inference from these data, **GPR37 likely influences addiction-related behavior by tuning dopamine clearance, dopaminergic receptor signaling, and striatal synaptic plasticity**. The 2011 findings that GPR37 loss blocks psychostimulant-induced ERK2 activation and ΔFosB accumulation provide a particularly compelling bridge from receptor biology to known molecular pathways of long-term drug adaptation. That mechanistic logic is strongest for **psychostimulant addiction-like phenotypes**, not yet for addiction in general. ([pubmed.ncbi.nlm.nih.gov](https://pubmed.ncbi.nlm.nih.gov/21372109/)) [S13]

## Bottom line

Yes—**GPR37 is fairly described as being implicated in addiction-related biology**, but with an important qualifier: the evidence is **mainly preclinical and most convincing for psychostimulant reward/addiction-like phenotypes in rodents**. If the intended meaning is “GPR37 has been shown to influence cocaine/amphetamine reward circuitry and related molecular adaptations,” the claim is well supported. If the intended meaning is “GPR37 is an established human addiction gene across substance use disorders,” the evidence is not there yet. ([pubmed.ncbi.nlm.nih.gov](https://pubmed.ncbi.nlm.nih.gov/17519329/)) [S14]

## Cited sources

- **[S1]** GPR37 associates with the dopamine transporter to modulate dopamine uptake and behavioral responses to dopaminergic drugs - PubMed — https://pubmed.ncbi.nlm.nih.gov/17519329/
- **[S2]** GPR37 associates with the dopamine transporter to modulate dopamine uptake and behavioral responses to dopaminergic drugs - PubMed — https://pubmed.ncbi.nlm.nih.gov/17519329/
- **[S3]** GPR37 associates with the dopamine transporter to modulate dopamine uptake and behavioral responses to dopaminergic drugs - PubMed — https://pubmed.ncbi.nlm.nih.gov/17519329/
- **[S4]** Absence of the GPR37/PAEL receptor impairs striatal Akt and ERK2 phosphorylation, DeltaFosB expression, and conditioned place preference to amphetamine and cocaine - PubMed — https://pubmed.ncbi.nlm.nih.gov/21372109/
- **[S5]** Parkinson's disease-associated GPR37 receptor regulates cocaine-mediated synaptic depression in corticostriatal synapses - PubMed — https://pubmed.ncbi.nlm.nih.gov/28007645/
- **[S6]** Impact of miR-29c-3p in the Nucleus Accumbens on Methamphetamine-Induced Behavioral Sensitization and Neuroplasticity-Related Proteins - PubMed — https://pubmed.ncbi.nlm.nih.gov/38256016/?utm_source=openai
- **[S7]** 
            Multivariate genome-wide association meta-analysis of over 1 million subjects identifies loci underlying multiple substance use disorders - PMC
         — https://pmc.ncbi.nlm.nih.gov/articles/PMC10217792/
- **[S8]** Altered dopamine signaling and MPTP resistance in mice lacking the Parkinson's disease-associated GPR37/parkin-associated endothelin-like receptor - PubMed — https://pubmed.ncbi.nlm.nih.gov/15218106/
- **[S9]** 
            Orphan G Protein-Coupled Receptor GPR37 as an Emerging Therapeutic Target - PMC
         — https://pmc.ncbi.nlm.nih.gov/articles/PMC11144446/
- **[S10]** Parkinson's disease-associated GPR37 receptor regulates cocaine-mediated synaptic depression in corticostriatal synapses - PubMed — https://pubmed.ncbi.nlm.nih.gov/28007645/
- **[S11]** Brain tissue expression of GPR37 in basal ganglia - The Human Protein Atlas — https://www.proteinatlas.org/ENSG00000170775-GPR37/brain/basal%2Bganglia
- **[S12]** GPR37 associates with the dopamine transporter to modulate dopamine uptake and behavioral responses to dopaminergic drugs - PubMed — https://pubmed.ncbi.nlm.nih.gov/17519329/
- **[S13]** Absence of the GPR37/PAEL receptor impairs striatal Akt and ERK2 phosphorylation, DeltaFosB expression, and conditioned place preference to amphetamine and cocaine - PubMed — https://pubmed.ncbi.nlm.nih.gov/21372109/
- **[S14]** GPR37 associates with the dopamine transporter to modulate dopamine uptake and behavioral responses to dopaminergic drugs - PubMed — https://pubmed.ncbi.nlm.nih.gov/17519329/
