## Supplementary material for "Understanding Substance Dependence: What Differentiates Addictive from Non-Addictive Drugs?": Folder for supplementary files: addictive-genes-claims-first100-096-znf106-gene-protein-is-implicated-in-addiction-2f2d499c4596.md

# addictive-genes-claims-first100-096-2f2d499c4596

**Original claim:** ZNF106 gene/protein is implicated in addiction.

## Claim

**ZNF106 gene/protein is implicated in addiction.**

## Verdict

Verdict: No Evidence

I did **not** find convincing direct evidence that **ZNF106** is an established addiction gene/protein in humans or animal models. In the addiction literature I reviewed, ZNF106 was **not** identified as a replicated locus, core pathway gene, or experimentally validated mediator of substance use disorder (SUD), opioid use disorder (OUD), alcohol use disorder (AUD), nicotine dependence, or other addictions. ([pubmed.ncbi.nlm.nih.gov](https://pubmed.ncbi.nlm.nih.gov/33556390/)) [S1]

## Confidence

**Medium**

Confidence is **medium** rather than high because absence-of-evidence judgments are inherently limited: a small, unreplicated, or very recent signal could exist. But across major reviews and large-scale human genetic studies of SUDs/OUD, I did not see ZNF106 emerge as a supported addiction-relevant gene. ([pubmed.ncbi.nlm.nih.gov](https://pubmed.ncbi.nlm.nih.gov/33556390/)) [S2]

## Supporting evidence

The only support I found is **indirect biological plausibility**, not addiction-specific evidence. ZNF106 is a protein-coding gene with **RNA-binding activity** and predicted localization to the **nucleolus** and **nuclear speck**, which makes it a plausible regulator of neuronal gene expression or RNA handling. NCBI also notes predicted involvement in insulin receptor signaling. ([ncbi.nlm.nih.gov](https://www.ncbi.nlm.nih.gov/gene/64397?utm_source=openai)) [S3]

ZNF106/ZFP106 also has bona fide **neural relevance**. In mice, loss of Zfp106 causes **motor and sensory neurodegeneration**, and the phenotype is linked to altered transcriptional programs and mitochondrial dysfunction. Separate work identified Zfp106 as a **GGGGCC repeat RNA-binding protein** that interacts with other RNA-binding proteins such as **TDP-43** and **FUS**, again supporting a role in neuronal RNA biology. These findings show that ZNF106 matters for neuronal maintenance/homeostasis broadly. ([pubmed.ncbi.nlm.nih.gov](https://pubmed.ncbi.nlm.nih.gov/26604141/?utm_source=openai)) [S4]

That said, none of these studies tested addiction-like behaviors, drug reward, relapse, withdrawal, sensitization, self-administration, or substance-specific clinical phenotypes. So they support only a **general neurobiological relevance**, not implication in addiction per se. ([pubmed.ncbi.nlm.nih.gov](https://pubmed.ncbi.nlm.nih.gov/26604141/?utm_source=openai)) [S5]

## Contradicting or limiting evidence

A major **umbrella review** of molecular genetics of SUDs, summarizing more than 150 meta-analyses, highlighted genes such as **OPRM1, DRD2, DRD4, BDNF, SLC6A4, ADH1B, CHRNA5, FOXP2, PEX, and AUTS2** as implicated across different addictions or substances; **ZNF106 was not among the genes highlighted**. This does not prove ZNF106 is uninvolved, but it strongly suggests it is **not an established or replicated addiction gene**. ([pubmed.ncbi.nlm.nih.gov](https://pubmed.ncbi.nlm.nih.gov/33556390/)) [S6]

Likewise, a very large multivariate GWAS meta-analysis across more than 1 million individuals found cross-substance and substance-specific loci for alcohol, tobacco, cannabis, and opioid disorders, with signals including **PDE4B** and other addiction-relevant loci; **ZNF106 was not reported among the lead findings** in the abstracted results. ([pubmed.ncbi.nlm.nih.gov](https://pubmed.ncbi.nlm.nih.gov/37250466/)) [S7]

A large **cross-ancestry OUD GWAS** identified loci including **OPRM1, FURIN, RABEPK, FBXW4, NCAM1, KCNN1, and TSNARE1**, with CNS enrichment in addiction-related brain regions. Again, **ZNF106 was not identified as an OUD locus** in the reported lead results. ([pubmed.ncbi.nlm.nih.gov](https://pubmed.ncbi.nlm.nih.gov/36171425/)) [S8]

A recent **multiomic network analysis of opioid addiction** integrating GWAS, transcriptomics, proteomics, epigenetics, and chromatin data from dorsolateral prefrontal cortex implicated pathways centered on **Akt, BDNF, ERK, OPRM1, GABA/glutamate signaling, and related genes**. ZNF106 was not among the named opioid addiction genes/pathways highlighted in the paper’s abstract and figures. ([pubmed.ncbi.nlm.nih.gov](https://pubmed.ncbi.nlm.nih.gov/39615775/)) [S9]

Importantly, some addiction-associated studies identify **epigenetic** or **expression** changes in many genes secondary to drug exposure, but that is not the same as showing that a gene is a causal or replicated addiction susceptibility factor. I did not find a replicated human genetic association, Mendelian evidence, animal addiction phenotype, or mechanistic addiction study specifically centered on ZNF106. ([pubmed.ncbi.nlm.nih.gov](https://pubmed.ncbi.nlm.nih.gov/33556390/)) [S10]

## Mechanistic / biological rationale

From a mechanistic standpoint, ZNF106 looks more like a **general neuronal RNA-regulatory/homeostatic factor** than a known addiction effector. NCBI describes it as an **RNA-binding** gene product, and mouse/cellular studies place it in RNA metabolism and neuronal survival pathways. ([ncbi.nlm.nih.gov](https://www.ncbi.nlm.nih.gov/gene/64397?utm_source=openai)) [S11]

The mouse knockout data are especially informative: Zfp106 deficiency leads to progressive **motor and sensory neuron degeneration**, with evidence of **mitochondrial dysfunction**, altered metabolic/inflammatory gene-expression programs, and neuromuscular pathology. Those findings fit a role in **neural homeostasis** and neuronal maintenance. ([pubmed.ncbi.nlm.nih.gov](https://pubmed.ncbi.nlm.nih.gov/26604141/?utm_source=openai)) [S12]

However, the addiction field’s most replicated biological themes involve reward and reinforcement circuitry, including **dopamine regulation**, **opioid receptor biology**, **glutamatergic plasticity**, and stress/externalizing pathways. Large human studies consistently nominate genes/pathways in those domains, whereas ZNF106 has not emerged as a recurrent signal. So even if ZNF106 contributes in some subtle way to neuronal resilience or transcriptional control, current evidence does **not** place it in the core mechanistic architecture of addiction. ([pubmed.ncbi.nlm.nih.gov](https://pubmed.ncbi.nlm.nih.gov/37250466/)) [S13]

In other words: **neural relevance ≠ addiction implication**. Many genes are important for neurons; far fewer are specifically implicated in addictive phenotypes. For ZNF106, the evidence currently fits the former, not the latter. ([pubmed.ncbi.nlm.nih.gov](https://pubmed.ncbi.nlm.nih.gov/26604141/?utm_source=openai)) [S14]

## Bottom line

Based on the scholarly and authoritative sources I reviewed, **ZNF106 is not currently supported as an addiction-implicated gene/protein**. It is a **neuronal RNA-binding protein with evidence for roles in neural maintenance and neurodegeneration**, which gives it some indirect biological plausibility, but I found **no replicated direct human or experimental evidence** tying it to addiction itself. The most accurate present-day assessment is therefore **No Evidence**. ([ncbi.nlm.nih.gov](https://www.ncbi.nlm.nih.gov/gene/64397?utm_source=openai)) [S15]

## Cited sources

- **[S1]** Molecular genetics of substance use disorders: An umbrella review - PubMed — https://pubmed.ncbi.nlm.nih.gov/33556390/
- **[S2]** Molecular genetics of substance use disorders: An umbrella review - PubMed — https://pubmed.ncbi.nlm.nih.gov/33556390/
- **[S3]** ZNF106 zinc finger protein 106 [Homo sapiens (human)] - Gene - NCBI — https://www.ncbi.nlm.nih.gov/gene/64397?utm_source=openai
- **[S4]** Deficiency of the zinc finger protein ZFP106 causes motor and sensory neurodegeneration - PubMed — https://pubmed.ncbi.nlm.nih.gov/26604141/?utm_source=openai
- **[S5]** Deficiency of the zinc finger protein ZFP106 causes motor and sensory neurodegeneration - PubMed — https://pubmed.ncbi.nlm.nih.gov/26604141/?utm_source=openai
- **[S6]** Molecular genetics of substance use disorders: An umbrella review - PubMed — https://pubmed.ncbi.nlm.nih.gov/33556390/
- **[S7]** Multivariate genome-wide association meta-analysis of over 1 million subjects identifies loci underlying multiple substance use disorders - PubMed — https://pubmed.ncbi.nlm.nih.gov/37250466/
- **[S8]** Cross-ancestry meta-analysis of opioid use disorder uncovers novel loci with predominant effects in brain regions associated with addiction - PubMed — https://pubmed.ncbi.nlm.nih.gov/36171425/
- **[S9]** Multiomic Network Analysis Identifies Dysregulated Neurobiological Pathways in Opioid Addiction - PubMed — https://pubmed.ncbi.nlm.nih.gov/39615775/
- **[S10]** Molecular genetics of substance use disorders: An umbrella review - PubMed — https://pubmed.ncbi.nlm.nih.gov/33556390/
- **[S11]** ZNF106 zinc finger protein 106 [Homo sapiens (human)] - Gene - NCBI — https://www.ncbi.nlm.nih.gov/gene/64397?utm_source=openai
- **[S12]** Deficiency of the zinc finger protein ZFP106 causes motor and sensory neurodegeneration - PubMed — https://pubmed.ncbi.nlm.nih.gov/26604141/?utm_source=openai
- **[S13]** Multivariate genome-wide association meta-analysis of over 1 million subjects identifies loci underlying multiple substance use disorders - PubMed — https://pubmed.ncbi.nlm.nih.gov/37250466/
- **[S14]** Deficiency of the zinc finger protein ZFP106 causes motor and sensory neurodegeneration - PubMed — https://pubmed.ncbi.nlm.nih.gov/26604141/?utm_source=openai
- **[S15]** ZNF106 zinc finger protein 106 [Homo sapiens (human)] - Gene - NCBI — https://www.ncbi.nlm.nih.gov/gene/64397?utm_source=openai
