## Supplementary material for "Understanding Substance Dependence: What Differentiates Addictive from Non-Addictive Drugs?": Folder for supplementary files: addictive-genes-claims-first100-097-sfrp5-gene-protein-is-implicated-in-addiction-9103492e475c.md

# addictive-genes-claims-first100-097-9103492e475c

**Original claim:** SFRP5 gene/protein is implicated in addiction.

## Claim

**“SFRP5 gene/protein is implicated in addiction.”**

## Verdict

Verdict: No Evidence

Current evidence does **not** support SFRP5 as an established addiction gene/protein. I found **indirect biological plausibility** through its role as a Wnt-signaling modulator and through broader evidence that Wnt/β-catenin pathways participate in addiction-related neuroadaptations, but I did **not** find convincing direct human genetic, transcriptomic, postmortem, or animal-behavioral evidence specifically implicating **SFRP5** in substance use disorder or addiction phenotypes. ([proteinatlas.org](https://www.proteinatlas.org/ENSG00000120057-SFRP5)) [S1]

## Confidence

**Medium.**

Confidence is not high because “absence of evidence” is always provisional, and niche omics supplements can be missed by keyword search. But confidence is above low because targeted searches for **SFRP5** with nicotine, alcohol, opioid, and substance-use-disorder terms did not yield addiction-focused SFRP5 papers, and several large addiction omics/genetics papers did not mention SFRP5 in their searchable full text. ([pubmed.ncbi.nlm.nih.gov](https://pubmed.ncbi.nlm.nih.gov/33573682/?utm_source=openai)) [S2]

## Supporting evidence

The strongest **supporting** evidence is only **indirect**. SFRP5 is a secreted frizzled-related protein that modulates Wnt signaling; the Human Protein Atlas lists it as **brain regional enriched in choroid plexus**, and NCBI/primary literature indicate central Sfrp5 signaling in rats can act through a **hypothalamic–brain-hepatic vagus** circuit to regulate glucose and lipid metabolism. That shows SFRP5 can participate in CNS homeostatic signaling, so a brain-related role is biologically plausible. ([proteinatlas.org](https://www.proteinatlas.org/ENSG00000120057-SFRP5)) [S3]

Separately, **Wnt signaling itself** has been implicated in addiction-relevant processes. In rodents, cocaine sensitization involves Wnt/β-catenin changes in prefrontal cortex, and pharmacologic manipulation of that pathway altered cocaine-induced neuroadaptations. Adolescent social isolation also altered prefrontal Wnt/β-catenin signaling and increased later cocaine vulnerability. A review further argued that Wnt/β-catenin signaling in nucleus accumbens may contribute to long-term cocaine neuroplasticity and could be a treatment target. ([pubmed.ncbi.nlm.nih.gov](https://pubmed.ncbi.nlm.nih.gov/26910786/?utm_source=openai)) [S4]

Opioid-related work points in the same pathway-level direction: morphine withdrawal has been linked to Wnt signaling, with inhibition of Wnt synthesis/signaling suppressing naloxone-precipitated withdrawal-related behavioral and neurochemical alterations in preclinical models. This supports the idea that **Wnt-family biology** can matter in addiction/withdrawal states. ([pubmed.ncbi.nlm.nih.gov](https://pubmed.ncbi.nlm.nih.gov/31738230/?utm_source=openai)) [S5]

Taken together, these studies mean that saying **“Wnt signaling is implicated in addiction-related neurobiology”** is reasonable. Because SFRP5 is a Wnt modulator and has some CNS expression/signaling data, one can make an **inference** that SFRP5 is *biologically plausible* as a candidate regulator. But that is still only an inference, not direct evidence. ([proteinatlas.org](https://www.proteinatlas.org/ENSG00000120057-SFRP5)) [S6]

## Contradicting or limiting evidence

The central limitation is simple: I did **not** find a direct paper showing that **SFRP5** variation, expression, protein level, or manipulation is associated with addiction risk, substance use disorder diagnosis, craving, relapse, self-administration, or withdrawal severity in humans or standard animal addiction models. Searches combining SFRP5 with nicotine dependence, alcohol dependence, opioid-related disorders, or substance-related disorders mainly returned non-addiction papers on inflammation, cancer/drug resistance, or other biology. ([pubmed.ncbi.nlm.nih.gov](https://pubmed.ncbi.nlm.nih.gov/33573682/?utm_source=openai)) [S7]

This absence also shows up in large addiction datasets. In a recent multiomic opioid addiction study integrating GWAS, transcriptomic, proteomic, and epigenetic data from human dorsolateral prefrontal cortex, searchable full text did **not** mention SFRP5. ([pubmed.ncbi.nlm.nih.gov](https://pubmed.ncbi.nlm.nih.gov/39615775/)) [S8]

Likewise, a transcriptome-wide association meta-analysis across 13 brain tissues for nicotine addiction had **no searchable mention of SFRP5**, and a large multi-ancestry tobacco use disorder paper whose full text discusses many implicated genes also had **no searchable SFRP5 mention**. That does **not** prove SFRP5 is uninvolved, but it argues against SFRP5 being a currently recognized, replicated addiction gene. ([pmc.ncbi.nlm.nih.gov](https://pmc.ncbi.nlm.nih.gov/articles/PMC8775257/)) [S9]

An additional caution is that evidence for **other SFRP family members** should not be transferred automatically to SFRP5. For example, **Sfrp2** has direct preclinical evidence in morphine withdrawal, but that is not evidence for **SFRP5**. Family membership and shared pathway biology are not enough to establish gene-specific implication. ([pubmed.ncbi.nlm.nih.gov](https://pubmed.ncbi.nlm.nih.gov/30291843/?utm_source=openai)) [S10]

Finally, the broader SFRP5 literature is dominated by roles in **adipose biology, inflammation, metabolism, and Wnt regulation**, not reward circuitry or substance-use phenotypes. In mouse and human reference resources, SFRP5 is especially tied to adipose/metabolic contexts, and human brain expression appears regionally enriched in **choroid plexus**, not prominently in classic mesolimbic reward nodes in the sources reviewed here. ([pubmed.ncbi.nlm.nih.gov](https://pubmed.ncbi.nlm.nih.gov/20558665/?utm_source=openai)) [S11]

## Mechanistic / biological rationale

Mechanistically, SFRP5 is a **secreted extracellular modulator of Wnt signaling**. It can antagonize Wnt signaling and has been linked in multiple non-addiction systems to regulation of inflammation, differentiation, and JNK-related signaling. Those functions are relevant in principle because addiction involves synaptic plasticity, stress signaling, neuroimmune changes, and persistent circuit remodeling. ([proteinatlas.org](https://www.proteinatlas.org/ENSG00000120057-SFRP5)) [S12]

That said, a plausible mechanism is not the same as demonstrated involvement. The addiction literature more directly implicates **Wnt/β-catenin pathway components broadly** rather than SFRP5 specifically. So the most defensible mechanistic statement is: **SFRP5 could theoretically influence addiction-related biology as an upstream Wnt regulator, but this remains unproven.** ([pubmed.ncbi.nlm.nih.gov](https://pubmed.ncbi.nlm.nih.gov/26910786/?utm_source=openai)) [S13]

A nuanced interpretation is that SFRP5 may fit better, based on current evidence, into **neural/metabolic homeostasis** than into addiction per se. The hypothalamic Sfrp5 study supports a CNS role in metabolic regulation, and the Human Protein Atlas signal in choroid plexus suggests specialized brain expression, but neither finding demonstrates a role in reward, reinforcement, or compulsive drug use. ([pubmed.ncbi.nlm.nih.gov](https://pubmed.ncbi.nlm.nih.gov/31770545/?utm_source=openai)) [S14]

## Bottom line

At present, the claim **“SFRP5 is implicated in addiction”** is **not supported by direct evidence**. The best available reading is:

- **Direct evidence for SFRP5 in addiction:** not found. ([pubmed.ncbi.nlm.nih.gov](https://pubmed.ncbi.nlm.nih.gov/33573682/?utm_source=openai)) [S15]
- **Indirect pathway plausibility via Wnt signaling:** yes. ([pubmed.ncbi.nlm.nih.gov](https://pubmed.ncbi.nlm.nih.gov/26910786/?utm_source=openai)) [S16]
- **Best overall judgment:** SFRP5 is a **plausible but currently unvalidated candidate**, not an established addiction-related gene/protein. ([proteinatlas.org](https://www.proteinatlas.org/ENSG00000120057-SFRP5)) [S17]

If you want, I can also do a **deeper gene-centric review** limited to:
1. human GWAS/TWAS evidence,  
2. postmortem brain expression data, or  
3. animal addiction-model studies only.

## Cited sources

- **[S1]** SFRP5 protein expression summary - The Human Protein Atlas — https://www.proteinatlas.org/ENSG00000120057-SFRP5
- **[S2]** Secreted frizzled-related protein 5 (SFRP5) protects ATDC5 cells against LPS-induced inflammation and apoptosis via inhibiting Wnt5a/JNK pathway - PubMed — https://pubmed.ncbi.nlm.nih.gov/33573682/?utm_source=openai
- **[S3]** SFRP5 protein expression summary - The Human Protein Atlas — https://www.proteinatlas.org/ENSG00000120057-SFRP5
- **[S4]** Wnt/β-catenin pathway in the prefrontal cortex is required for cocaine-induced neuroadaptations - PubMed — https://pubmed.ncbi.nlm.nih.gov/26910786/?utm_source=openai
- **[S5]** Wnt signaling contributes to withdrawal symptoms from opioid receptor activation induced by morphine exposure or chronic inflammation - PubMed — https://pubmed.ncbi.nlm.nih.gov/31738230/?utm_source=openai
- **[S6]** SFRP5 protein expression summary - The Human Protein Atlas — https://www.proteinatlas.org/ENSG00000120057-SFRP5
- **[S7]** Secreted frizzled-related protein 5 (SFRP5) protects ATDC5 cells against LPS-induced inflammation and apoptosis via inhibiting Wnt5a/JNK pathway - PubMed — https://pubmed.ncbi.nlm.nih.gov/33573682/?utm_source=openai
- **[S8]** Multiomic Network Analysis Identifies Dysregulated Neurobiological Pathways in Opioid Addiction - PubMed — https://pubmed.ncbi.nlm.nih.gov/39615775/
- **[S9]** 
            Meta-Analysis of Transcriptome-Wide Association Studies across 13 Brain Tissues Identified Novel Clusters of Genes Associated with Nicotine Addiction - PMC
         — https://pmc.ncbi.nlm.nih.gov/articles/PMC8775257/
- **[S10]** Loss of Sfrp2 contributes to the neurological disorders related with morphine withdrawal via Wnt/β-catenin signaling - PubMed — https://pubmed.ncbi.nlm.nih.gov/30291843/?utm_source=openai
- **[S11]** Sfrp5 is an anti-inflammatory adipokine that modulates metabolic dysfunction in obesity - PubMed — https://pubmed.ncbi.nlm.nih.gov/20558665/?utm_source=openai
- **[S12]** SFRP5 protein expression summary - The Human Protein Atlas — https://www.proteinatlas.org/ENSG00000120057-SFRP5
- **[S13]** Wnt/β-catenin pathway in the prefrontal cortex is required for cocaine-induced neuroadaptations - PubMed — https://pubmed.ncbi.nlm.nih.gov/26910786/?utm_source=openai
- **[S14]** Central Sfrp5 regulates hepatic glucose flux and VLDL-triglyceride secretion - PubMed — https://pubmed.ncbi.nlm.nih.gov/31770545/?utm_source=openai
- **[S15]** Secreted frizzled-related protein 5 (SFRP5) protects ATDC5 cells against LPS-induced inflammation and apoptosis via inhibiting Wnt5a/JNK pathway - PubMed — https://pubmed.ncbi.nlm.nih.gov/33573682/?utm_source=openai
- **[S16]** Wnt/β-catenin pathway in the prefrontal cortex is required for cocaine-induced neuroadaptations - PubMed — https://pubmed.ncbi.nlm.nih.gov/26910786/?utm_source=openai
- **[S17]** SFRP5 protein expression summary - The Human Protein Atlas — https://www.proteinatlas.org/ENSG00000120057-SFRP5
