## Supplementary material for "Understanding Substance Dependence: What Differentiates Addictive from Non-Addictive Drugs?": Folder for supplementary files: addictive-genes-claims-first100-098-rora-gene-protein-is-implicated-in-addiction-b11a1e4bc9b5.md

# addictive-genes-claims-first100-098-b11a1e4bc9b5

**Original claim:** RORA gene/protein is implicated in addiction.

## Claim

RORA gene/protein is implicated in addiction.

## Verdict

Verdict: Mixed Evidence

There is **some evidence** linking **RORA (RORα)** to addiction-related phenotypes, especially through **human genetic association studies of self-reported addiction/substance-specific dependence**, **drug-responsive brain transcriptomics**, and **biological plausibility via circadian/stress/inflammatory pathways**. However, the evidence is **not strong or consistently replicated enough** to place RORA among the best-established addiction genes. Much of the positive evidence is **candidate-gene, nominal, indirect, or pathway-level**, rather than definitive causal evidence from large unbiased GWAS or intervention studies. ([ncbi.nlm.nih.gov](https://www.ncbi.nlm.nih.gov/gene/6095)) [S1]

## Confidence

Medium

## Supporting evidence

**RORA is biologically relevant to brain function and circadian control.** The NCBI Gene summary describes RORA as a **nuclear hormone receptor transcription factor** that regulates transcription and “aids in the transcriptional regulation of some genes involved in circadian rhythm.” A classic mechanistic study further showed that **RORα regulates transcription of the core clock gene BMAL1**, placing it directly in circadian machinery. Because circadian disruption is strongly tied to reward processing and substance-use vulnerability, this makes RORA a plausible addiction-relevant regulator. ([ncbi.nlm.nih.gov](https://www.ncbi.nlm.nih.gov/gene/6095)) [S2]

**Human association data provide some direct, though not definitive, support.** In a recent UK Biobank analysis of **98,800 participants**, variants in circadian-related genes including **RORA** were associated with **self-reported addiction**, with additional sex-specific associations for **narcotic** and **prescription/opioid addiction**. The authors explicitly reported that RORA variants were among the influential signals in both overall and substance-specific analyses. That said, the study relied on **self-report** and tested a **preselected circadian-gene panel**, so it is supportive but not decisive. ([pmc.ncbi.nlm.nih.gov](https://pmc.ncbi.nlm.nih.gov/articles/PMC11674644/)) [S3]

**Earlier nicotine-dependence GWAS work also pointed to RORA, albeit weakly.** In NIH research volunteers, a genome-wide study reported **clusters of nominally positive SNPs** in **RORA** among genes associated with **nicotine dependence**; this was not a clean genome-wide-significant single-locus discovery, but it does count as prior human genetic signal in an addiction phenotype. ([pmc.ncbi.nlm.nih.gov](https://pmc.ncbi.nlm.nih.gov/articles/PMC2581605/)) [S4]

**Preclinical transcriptomic data suggest Rora is drug-responsive in reward-related brain tissue.** In mouse striatum, **acute amphetamine exposure** increased **Rora** expression modestly but significantly across lines. The striatum is central to reward learning and addiction-related neuroplasticity, so drug-induced regulation of Rora is consistent with involvement in stimulant-response biology. Still, this study examined **acute exposure/transcriptomics**, not addiction behavior per se. ([pmc.ncbi.nlm.nih.gov](https://pmc.ncbi.nlm.nih.gov/articles/PMC6291885/)) [S5]

**Systems-level opioid studies also place RORA in addiction-relevant pathways.** A recent multiomic network analysis integrating **GWAS, transcriptomic, proteomic, and epigenetic** data from opioid-related datasets identified interconnected pathways involving ERK, Akt, BDNF, immediate-early genes, and neuroinflammatory processes; in the model, **ERK signaling could activate RORA**, which then promotes transcription of genes tied to microglial inflammatory biology. This is best interpreted as **pathway implication**, not proof that RORA itself is a primary opioid-risk gene. ([pubmed.ncbi.nlm.nih.gov](https://pubmed.ncbi.nlm.nih.gov/39615775/)) [S6]

## Contradicting or limiting evidence

**The most direct negative/limiting human evidence I found comes from a RORA-focused psychopathology study that included an addiction-relevant externalizing dimension.** In that analysis, RORA variants showed a corrected association with **fear-related internalizing**, but for **externalizing**—the latent factor that included substance-related pathology—**none of the RORA SNPs survived multiple-testing correction**. This argues against a robust, broad association of RORA with the externalizing/addiction spectrum. ([pmc.ncbi.nlm.nih.gov](https://pmc.ncbi.nlm.nih.gov/articles/PMC3797180/)) [S7]

**Recent large, high-powered GWAS of substance use disorders have highlighted other loci rather than RORA.** For example, a cross-ancestry OUD meta-analysis in **425,944** individuals reported lead loci such as **OPRM1, FURIN, RABEPK, FBXW4, NCAM1, KCNN1,** and **TSNARE1**. A large cannabis use disorder GWAS highlighted **CHRNA2**. These results do not disprove a small RORA effect, but they suggest that **RORA is not currently among the most robust replicated genome-wide signals** for major substance use disorders. That is an inference from what these high-powered studies reported. ([pubmed.ncbi.nlm.nih.gov](https://pubmed.ncbi.nlm.nih.gov/36171425/)) [S8]

**Much of the positive RORA-addiction evidence is indirect.** The UK Biobank study used **self-reported addiction**, the older nicotine study was **nominal/cluster-based**, the amphetamine study measured **expression changes rather than addiction behavior**, and the opioid multiomic paper used **network inference**. I did **not identify strong primary studies** showing that direct manipulation of RORA itself reliably alters classic addiction endpoints such as **drug self-administration, escalation, reinstatement/relapse, or withdrawal severity** in standard animal models. That absence lowers confidence in a strong causal claim. ([pmc.ncbi.nlm.nih.gov](https://pmc.ncbi.nlm.nih.gov/articles/PMC11674644/)) [S9]

## Mechanistic / biological rationale

A plausible rationale exists. **RORA encodes a DNA-binding nuclear receptor transcription factor** with a recognized role in **circadian regulation**, including control of **BMAL1** and broader circadian gene-expression programs. Circadian mechanisms influence **dopamine signaling, sleep-wake timing, stress responsivity, and reward-related behavior**, all of which are relevant to addiction vulnerability and relapse. ([ncbi.nlm.nih.gov](https://www.ncbi.nlm.nih.gov/gene/6095)) [S10]

RORA also appears relevant to **neuroprotection and oxidative/inflammatory stress**. In the fear-psychopathology paper, the authors summarize prior evidence that RORA is expressed in brain regions such as cortex, hippocampus, and hypothalamus, is **activated during oxidative stress**, and can promote expression of genes involved in **reactive oxygen species clearance**. Addiction-related drugs can perturb oxidative stress, neuroimmune tone, and circadian homeostasis, so a transcription factor sitting at that intersection is mechanistically credible. ([pmc.ncbi.nlm.nih.gov](https://pmc.ncbi.nlm.nih.gov/articles/PMC3797180/)) [S11]

The opioid multiomic study adds a further plausible link: **ERK/Akt/BDNF-linked addiction circuitry** may converge on transcriptional regulators including **RORA**, potentially coupling reward-circuit signaling to **microglial/inflammatory** and plasticity-related downstream programs. This is still a model, not proof, but it gives a biologically coherent reason why RORA could matter in addiction. ([pubmed.ncbi.nlm.nih.gov](https://pubmed.ncbi.nlm.nih.gov/39615775/)) [S12]

## Bottom line

**RORA is plausibly involved in addiction-related biology, but the case is not yet strong enough to call it an established addiction gene.** The best reading of the literature is:

- **Yes, there are positive signals**: human association findings, stimulant-responsive expression changes, and credible circadian/neuroimmune mechanisms. ([pmc.ncbi.nlm.nih.gov](https://pmc.ncbi.nlm.nih.gov/articles/PMC11674644/)) [S13]
- **But the evidence is still limited and uneven**: some negative/nonnull findings for externalizing, lack of standout replication in large unbiased SUD GWAS, and little direct causal manipulation evidence. ([pmc.ncbi.nlm.nih.gov](https://pmc.ncbi.nlm.nih.gov/articles/PMC3797180/)) [S14]

So the claim is **reasonable as a tentative statement**—“RORA has been implicated in addiction-related pathways and some addiction phenotypes”—but **too strong if meant as firmly established or broadly replicated across addictions**.

## Cited sources

- **[S1]** RORA RAR related orphan receptor A [Homo sapiens (human)] - Gene - NCBI — https://www.ncbi.nlm.nih.gov/gene/6095
- **[S2]** RORA RAR related orphan receptor A [Homo sapiens (human)] - Gene - NCBI — https://www.ncbi.nlm.nih.gov/gene/6095
- **[S3]** 
            Sex- and Substance-Specific Associations of Circadian-Related Genes with Addiction in the UK Biobank Cohort Implicate Neuroplasticity Pathways - PMC
         — https://pmc.ncbi.nlm.nih.gov/articles/PMC11674644/
- **[S4]** 
            Genome wide association for nicotine dependence and smoking cessation success in NIH research volunteers - PMC
         — https://pmc.ncbi.nlm.nih.gov/articles/PMC2581605/
- **[S5]** 
            Interplay Between Amphetamine and Activity Level in Gene Networks of the Mouse Striatum - PMC
         — https://pmc.ncbi.nlm.nih.gov/articles/PMC6291885/
- **[S6]** Multiomic Network Analysis Identifies Dysregulated Neurobiological Pathways in Opioid Addiction - PubMed — https://pubmed.ncbi.nlm.nih.gov/39615775/
- **[S7]** 
            The retinoid-related orphan receptor alpha (RORA) gene and fear-related psychopathology - PMC
         — https://pmc.ncbi.nlm.nih.gov/articles/PMC3797180/
- **[S8]** Cross-ancestry meta-analysis of opioid use disorder uncovers novel loci with predominant effects in brain regions associated with addiction - PubMed — https://pubmed.ncbi.nlm.nih.gov/36171425/
- **[S9]** 
            Sex- and Substance-Specific Associations of Circadian-Related Genes with Addiction in the UK Biobank Cohort Implicate Neuroplasticity Pathways - PMC
         — https://pmc.ncbi.nlm.nih.gov/articles/PMC11674644/
- **[S10]** RORA RAR related orphan receptor A [Homo sapiens (human)] - Gene - NCBI — https://www.ncbi.nlm.nih.gov/gene/6095
- **[S11]** 
            The retinoid-related orphan receptor alpha (RORA) gene and fear-related psychopathology - PMC
         — https://pmc.ncbi.nlm.nih.gov/articles/PMC3797180/
- **[S12]** Multiomic Network Analysis Identifies Dysregulated Neurobiological Pathways in Opioid Addiction - PubMed — https://pubmed.ncbi.nlm.nih.gov/39615775/
- **[S13]** 
            Sex- and Substance-Specific Associations of Circadian-Related Genes with Addiction in the UK Biobank Cohort Implicate Neuroplasticity Pathways - PMC
         — https://pmc.ncbi.nlm.nih.gov/articles/PMC11674644/
- **[S14]** 
            The retinoid-related orphan receptor alpha (RORA) gene and fear-related psychopathology - PMC
         — https://pmc.ncbi.nlm.nih.gov/articles/PMC3797180/
