## Supplementary material for "Understanding Substance Dependence: What Differentiates Addictive from Non-Addictive Drugs?": Folder for supplementary files: addictive-genes-claims-first100-099-kcnk1-gene-protein-is-implicated-in-addiction-e897048ea762.md

# addictive-genes-claims-first100-099-e897048ea762

**Original claim:** KCNK1 gene/protein is implicated in addiction.

## Claim

KCNK1 gene/protein is implicated in addiction.

## Verdict

Verdict: Mixed Evidence

There is **some preclinical and transcriptomic evidence** connecting **KCNK1/TWIK-1** to substance-related neurobiology, particularly nicotine- and alcohol-associated gene-expression changes, and a **very preliminary conference abstract** reported that nucleus accumbens KCNK1 knockdown reduced stress-potentiated alcohol drinking in mice. However, I did **not** find strong, replicated **human genetic**, **clinical**, or widely validated **causal** evidence placing KCNK1 among established addiction genes. ([frontiersin.org](https://www.frontiersin.org/journals/psychiatry/articles/10.3389/fpsyt.2011.00005/full)) [S1]

## Confidence

Medium

## Supporting evidence

**1) KCNK1 is biologically relevant to brain excitability/homeostasis.**  
KCNK1 encodes the two-pore-domain potassium channel **TWIK-1/K2P1.1**, a potassium leak/background channel. NCBI Gene annotates it with potassium channel and potassium leak channel activity, and brain-focused resources describe it as contributing to resting membrane potential regulation, especially in astrocytes. Experimental work shows TWIK-1 and TREK-1 contribute substantially to astrocyte passive conductance, and later work supports TWIK1/TREK1 heterodimers as regulators of astrocytic passive conductance. ([ncbi.nlm.nih.gov](https://www.ncbi.nlm.nih.gov/gene/3775)) [S2]

**2) Nicotine exposure alters KCNK1 expression in addiction-relevant settings.**  
A rat study of chronic nicotine self-administration across addiction-relevant brain regions reported effects of nicotine on **TWIK-1 mRNA** in the **amygdala** and **ventral tegmental area**. Separately, acute nicotine exposure in SH-SY5Y cells upregulated **KCNK1** among transport-related genes after 1 hour. These findings suggest nicotine can modulate KCNK1 expression, although they do not by themselves show that KCNK1 drives addiction behavior. ([pubmed.ncbi.nlm.nih.gov](https://pubmed.ncbi.nlm.nih.gov/11478936/?utm_source=openai)) [S3]

**3) Kcnk1 has shown an indirect association with a nicotine-withdrawal phenotype in mice.**  
In a systems-genetics study of **nicotine withdrawal deficits in contextual fear learning** across BXD mouse strains, hippocampal **Kcnk1** expression significantly correlated with the withdrawal-related phenotype. This is relevant because withdrawal is part of addiction biology, though the paper’s prioritized positional candidate genes for the mapped QTL were other genes, not Kcnk1. ([pmc.ncbi.nlm.nih.gov](https://pmc.ncbi.nlm.nih.gov/articles/PMC12666787/)) [S4]

**4) Alcohol exposure studies have reported Kcnk1 transcript changes.**  
Older ethanol-expression studies reported **Twik-1/Kcnk1 downregulation** in cerebellum after chronic ethanol exposure in wild-type mice, and another study found **Kcnk1** altered in the dorsal raphe after adolescent binge-like alcohol drinking in alcohol-preferring rats. These are again supportive of responsiveness to drug exposure, but they are expression studies rather than proof of causal involvement in addiction. ([pmc.ncbi.nlm.nih.gov](https://pmc.ncbi.nlm.nih.gov/articles/PMC1931504/?utm_source=openai)) [S5]

**5) There is one direct but preliminary behavioral claim from a conference abstract.**  
A 2025 LSU symposium abstract reported that transcriptomic analysis downstream of accumbal G9a manipulation highlighted **KCNK1**, and that selectively targeting **NAc KCNK1 in dynorphin-positive neurons** blocked **stress-potentiated alcohol drinking** and reproduced excitability changes. If this result holds up in a peer-reviewed full paper, it would be meaningful direct evidence. At present, though, it appears to be **conference-level, unpublished/preliminary** evidence. ([faculty.lsu.edu](https://faculty.lsu.edu/sfnbr/files/sfnbtr_annual_symposium_program_2025.pdf?utm_source=openai)) [S6]

## Contradicting or limiting evidence

**1) Human genetics support appears weak or absent.**  
A current **GWAS Catalog** search for **KCNK1** returned **no results**, which argues against KCNK1 being a well-established genome-wide significant addiction locus in curated human GWAS literature. Consistent with that, the NCBI Gene page lists GWAS-related entries for non-addiction traits such as periodontal pathogen colonization and severe diabetic retinopathy, not substance use disorders. ([ebi.ac.uk](https://www.ebi.ac.uk/gwas/search?query=KCNK1)) [S7]

**2) Large-scale cross-substance genomic reviews do not highlight KCNK1.**  
A systematic integration of **75 GWAS studies** plus brain RNA-seq studies spanning alcohol, nicotine, cannabis, opioid, cocaine, and methamphetamine traits identified thousands of genes, but the highlighted shared/specific genes were things like **CACNB2, GRIN2B, PLXDC2, PKNOX2, ADH1B, ALDH2, CHRNA3, and CHRNA4**—not KCNK1. That does not prove absence of effect, but it suggests KCNK1 is not a robust signal in current large-scale addiction genomics. ([pubmed.ncbi.nlm.nih.gov](https://pubmed.ncbi.nlm.nih.gov/35491750/)) [S8]

**3) Addiction genetics reviews implicate other potassium-channel genes more clearly than KCNK1.**  
For example, an opioid genetics review specifically notes GWAS signals in **KCNC1** and **KCNG2**, not KCNK1. Likewise, a NIDA-authored review on potassium channels in substance use disorders discusses stronger evidence for families such as **GIRK/KCNJ**, **KCNQ**, **KCNN**, **KCNMA1**, and **KCNK13**, whereas KCNK1 appears mainly in the background classification of potassium channels or as an indirect expression-change observation. ([pubmed.ncbi.nlm.nih.gov](https://pubmed.ncbi.nlm.nih.gov/29302220/)) [S9]

**4) Most positive evidence is indirect.**  
The majority of KCNK1-addiction links I found were:  
- drug-induced expression changes,  
- correlation with a withdrawal-related cognitive phenotype, or  
- a conference abstract not yet established in the peer-reviewed literature.  
That is a much lower evidentiary standard than replicated human association, Mendelian/randomization support, validated causal animal manipulations across labs, or therapeutic modulation data. ([frontiersin.org](https://www.frontiersin.org/journals/psychiatry/articles/10.3389/fpsyt.2011.00005/full)) [S10]

## Mechanistic / biological rationale

There is a **plausible mechanism**, but plausibility is not the same as proof. KCNK1/TWIK-1 helps set background potassium conductance and influences **resting membrane potential** and **cellular excitability**. In astrocytes, TWIK-1—alone or in heterodimeric partnership with TREK-1—contributes to **passive conductance** and therefore to aspects of **potassium homeostasis** in brain tissue. Because addiction involves long-lasting adaptations in excitability and synaptic/network function in circuits such as the **nucleus accumbens**, **amygdala**, and **VTA**, a channel like KCNK1 is mechanistically credible as a modulator of vulnerability or drug adaptation. ([ncbi.nlm.nih.gov](https://www.ncbi.nlm.nih.gov/gene/3775)) [S11]

That said, KCNK1’s best-established biology is currently broader **neural homeostasis/excitability regulation**, not addiction per se. In newer knockout work, TWIK-1 deficiency increased susceptibility to **kainic acid-induced seizures**, supporting a role in network stability and astrocyte-mediated neuromodulation. This strengthens the case that KCNK1 could influence addiction-relevant circuitry, but it remains **indirect evidence** for addiction. ([pmc.ncbi.nlm.nih.gov](https://pmc.ncbi.nlm.nih.gov/articles/PMC11732521/)) [S12]

## Bottom line

**KCNK1 should not currently be described as a well-established addiction gene/protein.** The best characterization is that **limited, mostly preclinical and indirect evidence** suggests KCNK1 **may participate in addiction-related neuroadaptation**, especially in nicotine/alcohol contexts and possibly via neuronal/astrocytic excitability in reward-stress circuits. But the claim is **not strongly established**, because robust human genetics, replicated causal studies, and peer-reviewed mechanistic addiction papers specifically centered on KCNK1 are still sparse. ([pubmed.ncbi.nlm.nih.gov](https://pubmed.ncbi.nlm.nih.gov/35491750/)) [S13]

## Cited sources

- **[S1]** Frontiers | Genome-Wide Expression Analysis Reveals Diverse Effects of Acute Nicotine Exposure on Neuronal Function-Related Genes and Pathways — https://www.frontiersin.org/journals/psychiatry/articles/10.3389/fpsyt.2011.00005/full
- **[S2]** KCNK1 potassium two pore domain channel subfamily K member 1 [Homo sapiens (human)] - Gene - NCBI — https://www.ncbi.nlm.nih.gov/gene/3775
- **[S3]** Region-specific transcriptional response to chronic nicotine in rat brain - PubMed — https://pubmed.ncbi.nlm.nih.gov/11478936/?utm_source=openai
- **[S4]** 
            Systems genetic analysis of nicotine withdrawal deficits in hippocampus-dependent learning - PMC
         — https://pmc.ncbi.nlm.nih.gov/articles/PMC12666787/
- **[S5]** Microarray Analysis Identifies Cerebellar Genes Sensitive to Chronic Ethanol Treatment in PKCγ Mice - PMC — https://pmc.ncbi.nlm.nih.gov/articles/PMC1931504/?utm_source=openai
- **[S6]** Baton Rouge Chapter — https://faculty.lsu.edu/sfnbr/files/sfnbtr_annual_symposium_program_2025.pdf?utm_source=openai
- **[S7]** GWAS Catalog — https://www.ebi.ac.uk/gwas/search?query=KCNK1
- **[S8]** Common and distinguishing genetic factors for substance use behavior and disorder: an integrated analysis of genomic and transcriptomic studies from both human and animal studies - PubMed — https://pubmed.ncbi.nlm.nih.gov/35491750/
- **[S9]** A brief review of the genetics and pharmacogenetics of opioid use disorders - PubMed — https://pubmed.ncbi.nlm.nih.gov/29302220/
- **[S10]** Frontiers | Genome-Wide Expression Analysis Reveals Diverse Effects of Acute Nicotine Exposure on Neuronal Function-Related Genes and Pathways — https://www.frontiersin.org/journals/psychiatry/articles/10.3389/fpsyt.2011.00005/full
- **[S11]** KCNK1 potassium two pore domain channel subfamily K member 1 [Homo sapiens (human)] - Gene - NCBI — https://www.ncbi.nlm.nih.gov/gene/3775
- **[S12]** 
            Mice deficient in TWIK-1 are more susceptible to kainic acid-induced seizures - PMC
         — https://pmc.ncbi.nlm.nih.gov/articles/PMC11732521/
- **[S13]** Common and distinguishing genetic factors for substance use behavior and disorder: an integrated analysis of genomic and transcriptomic studies from both human and animal studies - PubMed — https://pubmed.ncbi.nlm.nih.gov/35491750/
