## Supplementary material for "Understanding Substance Dependence: What Differentiates Addictive from Non-Addictive Drugs?": Folder for supplementary files: addictive-genes-claims-first100-100-thoc3-gene-protein-is-implicated-in-addiction-a230bbf5f804.md

# addictive-genes-claims-first100-100-a230bbf5f804

**Original claim:** THOC3 gene/protein is implicated in addiction.

## Claim

THOC3 gene/protein is implicated in addiction.

## Verdict

Verdict: No Evidence

I did not find direct evidence that **THOC3 itself** is implicated in addiction or substance use disorders. THOC3 is a real, conserved component of the **THO/TREX mRNA-processing/export machinery**, is expressed in brain tissue, and appears in neuronal transcript-processing complexes, so the idea is **biologically plausible**. But the addiction-relevant evidence I found concerns the **broader THO complex**—especially other subunits such as **THOC5**—not THOC3 specifically. ([ncbi.nlm.nih.gov](https://www.ncbi.nlm.nih.gov/gene/84321)) [S1]

## Confidence

Medium

Confidence is **medium** because the literature search found substantial information on THOC3’s molecular function and on THO/TREX neurobiology, but not THOC3-specific addiction findings. That is not proof of no role, but it does argue against THOC3 being an established addiction gene at present. ([ncbi.nlm.nih.gov](https://www.ncbi.nlm.nih.gov/gene/84321)) [S2]

## Supporting evidence

THOC3 encodes **THO complex subunit 3**, a core part of the nuclear **THO/TREX complex** that couples mRNA processing with nuclear export. NCBI describes THOC3 as a component of the transcription export complex recruited to mRNAs in a splicing- and cap-dependent manner. Structural resources also place THOC3 in the human THO core complex. ([ncbi.nlm.nih.gov](https://www.ncbi.nlm.nih.gov/gene/84321)) [S3]

THOC3 is present in the nervous system. Human Protein Atlas data show THOC3 expression in brain tissue, including **hippocampal formation**; the hippocampal consensus mRNA expression shown there is **40.2 nTPM**. At the same time, THOC3 is classified as having **low tissue specificity**, meaning it is broadly expressed rather than selectively enriched in addiction circuitry. ([v21.proteinatlas.org](https://v21.proteinatlas.org/ENSG00000051596-THOC3/brain/Hippocampal%2Bformation)) [S4]

There is also indirect neuronal evidence. In a study of neuronal RNA processing, all core THO members—including **THOC3**—were detected in a THO complex associated with **ZC3H14**, and the authors proposed that this machinery helps coordinate processing/export of neuronal transcripts. That supports a role for THOC3 in neuronal RNA handling, though not specifically in addiction. ([pubmed.ncbi.nlm.nih.gov](https://pubmed.ncbi.nlm.nih.gov/29912477/)) [S5]

A separate THO-complex study makes the addiction link more plausible at the **complex** level. In mice, conditional deletion of **Thoc5** in dopaminergic neurons caused substantial **loss of dopamine synapses**, later **dopaminergic neurodegeneration**, and severely impaired dopamine release. Because addiction depends heavily on dopamine-linked reinforcement and plasticity, this shows that THO-complex dysfunction can affect addiction-relevant neural systems. But again, those data are about **THOC5/THO-complex biology**, not THOC3 itself. ([pmc.ncbi.nlm.nih.gov](https://pmc.ncbi.nlm.nih.gov/articles/PMC6542560/)) [S6]

## Contradicting or limiting evidence

The biggest limitation is simple: I found **no THOC3-specific addiction study**. In targeted searches using THOC3 together with addiction-related terms, the returned scholarly results were unrelated THOC3 biology papers rather than substance-use papers; for example, a search intended to retrieve THOC3/addiction literature surfaced a 2026 THOC3 paper on **intestinal stem-cell self-renewal**, not addiction. ([pubmed.ncbi.nlm.nih.gov](https://pubmed.ncbi.nlm.nih.gov/41486589/?utm_source=openai)) [S7]

Likewise, I found no mention of THOC3 in the article text of recent large-scale substance-use genetics papers. A **multivariate GWAS/meta-analysis of over 1 million subjects** and a later **MTAG study** on substance-use traits were both searchable in full text, and direct text searches returned **no “THOC3” hit** in either article. That does not rule out a buried supplementary or sub-threshold association, but it argues against THOC3 being among the more clearly supported current SUD loci. ([pmc.ncbi.nlm.nih.gov](https://pmc.ncbi.nlm.nih.gov/articles/PMC10217792/?utm_source=openai)) [S8]

I also did not find human association, causal-variant, or addiction-model papers specifically assigning THOC3 a role in alcohol, nicotine, opioid, stimulant, or cannabis addiction in the sources consulted. By contrast, when THO/TREX subunits are discussed in disease contexts, the literature more often points to **THOC2** and **THOC6** in neurodevelopmental disorders, or **THOC5** in dopaminergic-neuron maintenance. ([pmc.ncbi.nlm.nih.gov](https://pmc.ncbi.nlm.nih.gov/articles/PMC5095910/)) [S9]

## Mechanistic / biological rationale

A cautious mechanistic rationale does exist. Addiction involves persistent neuroadaptations in reward and control circuits, and dopamine signaling is a major part of reinforcement-related brain function. So, in principle, genes that shape **activity-dependent RNA processing/export** could influence vulnerability to addictive behavior. ([nida.nih.gov](https://nida.nih.gov/publications/drugs-brains-behavior-science-addiction/drugs-brain?utm_source=openai)) [S10]

THO/TREX biology fits that broad framework: THOC proteins help regulate export/processing of selected transcripts, and THO dysfunction in neurons can disrupt **synapse-related gene expression** and **dopaminergic neuron integrity**. Since THOC3 is a core THO subunit, it is reasonable to hypothesize that THOC3 perturbation might affect neuronal homeostasis or plasticity. But that remains an **inference from complex-level biology**, not direct evidence that THOC3 is implicated in addiction. ([ncbi.nlm.nih.gov](https://www.ncbi.nlm.nih.gov/gene/84321)) [S11]

## Bottom line

THOC3 is **biologically plausible but not demonstrated** as an addiction gene. The evidence supports THOC3 as a core mRNA-export factor present in brain and as part of a neuronal THO/TREX complex, while separate work on another subunit (**THOC5**) shows that THO-complex dysfunction can damage dopaminergic circuits relevant to addiction. But I found **no direct THOC3-specific human genetic, transcriptomic, or experimental evidence** linking it to addiction or substance use disorder. On current evidence, the claim is best rated **No Evidence**. ([ncbi.nlm.nih.gov](https://www.ncbi.nlm.nih.gov/gene/84321)) [S12]

## Cited sources

- **[S1]** THOC3 THO complex subunit 3 [Homo sapiens (human)] - Gene - NCBI — https://www.ncbi.nlm.nih.gov/gene/84321
- **[S2]** THOC3 THO complex subunit 3 [Homo sapiens (human)] - Gene - NCBI — https://www.ncbi.nlm.nih.gov/gene/84321
- **[S3]** THOC3 THO complex subunit 3 [Homo sapiens (human)] - Gene - NCBI — https://www.ncbi.nlm.nih.gov/gene/84321
- **[S4]** Brain tissue expression of THOC3 in Hippocampal formation - The Human Protein Atlas — https://v21.proteinatlas.org/ENSG00000051596-THOC3/brain/Hippocampal%2Bformation
- **[S5]** The polyadenosine RNA-binding protein ZC3H14 interacts with the THO complex and coordinately regulates the processing of neuronal transcripts - PubMed — https://pubmed.ncbi.nlm.nih.gov/29912477/
- **[S6]** 
            The THO complex coordinates transcripts for synapse development and dopamine neuron survival - PMC
         — https://pmc.ncbi.nlm.nih.gov/articles/PMC6542560/
- **[S7]** Small Nucleolar RNA Snord17 Promotes Self-Renewal of Intestinal Stem Cells through Yy2 mRNA Export and Tead4 Activation - PubMed — https://pubmed.ncbi.nlm.nih.gov/41486589/?utm_source=openai
- **[S8]** Multivariate genome-wide association meta-analysis of over 1 million subjects identifies loci underlying multiple substance use disorders - PMC — https://pmc.ncbi.nlm.nih.gov/articles/PMC10217792/?utm_source=openai
- **[S9]** 
            The role of TREX in gene expression and disease - PMC
         — https://pmc.ncbi.nlm.nih.gov/articles/PMC5095910/
- **[S10]** Drugs, Brains, and Behavior: The Science of Addiction: Drugs and the Brain | NIDA — https://nida.nih.gov/publications/drugs-brains-behavior-science-addiction/drugs-brain?utm_source=openai
- **[S11]** THOC3 THO complex subunit 3 [Homo sapiens (human)] - Gene - NCBI — https://www.ncbi.nlm.nih.gov/gene/84321
- **[S12]** THOC3 THO complex subunit 3 [Homo sapiens (human)] - Gene - NCBI — https://www.ncbi.nlm.nih.gov/gene/84321
