## Supplementary material for "Understanding Substance Dependence: What Differentiates Addictive from Non-Addictive Drugs?": Folder for supplementary files: non-addictive-genes-claims-first100-001-plekhb1-gene-protein-has-been-associated-with-regu-6c190dda5f2d.md

# non-addictive-genes-claims-first100-001-6c190dda5f2d

**Original claim:** PLEKHB1 gene/protein has been associated with regulatory or homeostatic biological processes in neural systems.

## Claim

PLEKHB1 gene/protein has been associated with regulatory or homeostatic biological processes in neural systems.

## Verdict

Verdict: Mixed Evidence

PLEKHB1 is clearly associated with neural and sensory cell types, especially retina and other primary sensory systems, and several studies link it to membrane trafficking, phototransduction-related signaling, and mechanotransduction-related protein complexes. However, the strongest evidence is largely from expression/localization and protein-interaction studies rather than decisive functional loss-of-function phenotypes, and direct evidence for a bona fide **homeostatic** role is limited. ([ncbi.nlm.nih.gov](https://www.ncbi.nlm.nih.gov/gene/58473)) [S1]

## Confidence

Medium

## Supporting evidence

PLEKHB1 is not a generic housekeeping gene; it is enriched in neural/sensory contexts. NCBI summarizes it as having biased expression in brain, and the Human Protein Atlas classifies it as tissue-enhanced in **brain and retina**. Single-cell data further show cell-type-enhanced expression in **cone and rod photoreceptors**, **Müller glia**, and **oligodendrocytes**, which supports a bona fide association with neural and neuroglial systems rather than only non-neural tissues. ([ncbi.nlm.nih.gov](https://www.ncbi.nlm.nih.gov/gene/58473)) [S2]

The best direct retinal evidence comes from the original PHR1/PLEKHB1 studies. Xu et al. reported that PHR1 is highly expressed in photoreceptors, localizes to photoreceptor outer segments, and binds **transducin βγ** rather than inositol phosphates. On that basis, the authors proposed that it could modulate the phototransduction pathway. NCBI/GO also lists PLEKHB1 as “involved in phototransduction,” although that annotation is based on a non-traceable author statement rather than a definitive causal experiment. ([pubmed.ncbi.nlm.nih.gov](https://pubmed.ncbi.nlm.nih.gov/10585447/)) [S3]

A second foundational study identified PLEKHB1 as **evectin-1**, a vesicular protein localized to **rhodopsin-bearing post-Golgi membranes** in photoreceptors. The authors concluded that its structure and subcellular distribution strongly implicate it as a mediator of **post-Golgi trafficking** in cells producing large membrane-rich organelles. Because membrane trafficking in photoreceptors is a key regulatory process for outer-segment maintenance and signaling compartmentalization, this is relevant evidence for a neural regulatory association, even if it is not a direct demonstration of “homeostasis” per se. ([pubmed.ncbi.nlm.nih.gov](https://pubmed.ncbi.nlm.nih.gov/10200314/)) [S4]

PLEKHB1 also appears in other sensory neural systems. In a mouse knock-in/knockout study, Phr1 expression was detected in retina, olfactory epithelium, olfactory bulb, inner ear sensory cells, and other primary sensory neurons. The authors noted that many of these sensory systems use specialized signaling architectures and suggested that PHR1 may mark a distinct sensory-neuronal program. This broad sensory-neural distribution strengthens the claim that PLEKHB1 is associated with neural regulatory biology rather than being retina-specific only. ([pmc.ncbi.nlm.nih.gov](https://pmc.ncbi.nlm.nih.gov/articles/PMC517893/)) [S5]

In inner-ear sensory cells, PHR1/PLEKHB1 directly interacted with **myosin 1c** and **myosin VIIa**. The authors proposed that these interactions could help anchor the actin cytoskeleton to the plasma membrane and participate in the **slow adaptation** phase of mechanotransduction in hair bundles. That is a plausible regulatory role in a neural sensory system, though it remains more mechanistic/proposed than conclusively established in vivo. ([pubmed.ncbi.nlm.nih.gov](https://pubmed.ncbi.nlm.nih.gov/15976448/)) [S6]

More recent disease-oriented work is compatible with a neural role. In TDP-43 ALS mouse models, PLEKHB1 protein was downregulated in motor neurons, and the authors interpreted older retinal, sensory-neuron, and trafficking studies as suggesting a possible role in membrane/G-protein/myosin-related neuronal processes. This is only associative and not disease-defining, but it shows that PLEKHB1 continues to be considered relevant to neuronal regulatory biology. ([pmc.ncbi.nlm.nih.gov](https://pmc.ncbi.nlm.nih.gov/articles/PMC7530531/)) [S7]

## Contradicting or limiting evidence

The main limitation is that **functional necessity is weakly established**. In the 2004 mouse knockout-knockin study, despite clear expression in primary sensory neurons, the investigators did **not** detect an obvious phenotype in Phr1-null mice up to 1 year of age. They explicitly suggested possible redundancy with the homolog PHR2/EVT2. That weakens any claim that PLEKHB1 is a proven, indispensable homeostatic regulator in neural systems. ([pmc.ncbi.nlm.nih.gov](https://pmc.ncbi.nlm.nih.gov/articles/PMC517893/)) [S8]

The strongest explicit test of a homeostasis/adaptation-type function in a neural system was negative. In olfactory neurons, PHR1 was abundant and vesicle-associated, but knockout mice showed no detectable differences in olfactory morphology, regeneration after methyl bromide injury, odor function, or short-term olfactory adaptation. The authors concluded that the exact function remained unknown. This directly limits the “homeostatic” part of the claim. ([pure.johnshopkins.edu](https://pure.johnshopkins.edu/en/publications/phr1-is-a-vesicle-bound-protein-abundantly-expressed-in-mature-ol-3/)) [S9]

Much of the positive case rests on **expression, localization, binding, and author inference**, not on robust perturbation data. For example, transducin βγ binding suggests a signaling role in photoreceptors, and myosin interactions suggest a mechanotransduction-related role in hair cells, but these are not the same as showing that PLEKHB1 measurably regulates neural homeostasis in vivo. ([pubmed.ncbi.nlm.nih.gov](https://pubmed.ncbi.nlm.nih.gov/10585447/)) [S10]

The ALS literature is also preliminary. The motor-neuron translatome study explicitly stated that PLEKHB1 had not previously been implicated in neurodegenerative disease and that future work would be needed to characterize its healthy and pathological functions in motor neurons. So disease association should not be overread as proof of a homeostatic neural role. ([pmc.ncbi.nlm.nih.gov](https://pmc.ncbi.nlm.nih.gov/articles/PMC7530531/)) [S11]

There is also an important nomenclature hazard: **PLEKHB1 uses the alias “PHR1,”** but in a large neurodevelopment/axon-degeneration literature, **Phr1** often refers instead to **MYCBP2**, an unrelated E3 ubiquitin ligase involved in axon growth and degeneration. This alias collision can falsely inflate the apparent neural evidence if sources are not carefully disambiguated. ([ncbi.nlm.nih.gov](https://www.ncbi.nlm.nih.gov/gene/58473)) [S12]

## Mechanistic / biological rationale

Biologically, PLEKHB1 is a plausible regulator of specialized neural membrane processes. It is an unusual **integral membrane protein** with an **N-terminal pleckstrin homology domain** and a **C-terminal transmembrane anchor**. In photoreceptors, it localizes to outer segments and post-Golgi membranes, fitting a role in membrane sorting/trafficking. Its binding to **transducin βγ** makes a signaling-modulatory role in phototransduction plausible. In auditory/vestibular hair cells, its direct interaction with **myosin 1c** and **myosin VIIa** provides a credible route by which it could influence actin–membrane coupling and sensory mechanotransduction. These are all regulatory-type mechanisms in neural sensory systems, but mostly mechanistic and inferential rather than firmly demonstrated as essential homeostatic functions. ([pubmed.ncbi.nlm.nih.gov](https://pubmed.ncbi.nlm.nih.gov/10585447/)) [S13]

## Bottom line

A careful reading of the literature supports saying that **PLEKHB1 has been associated with regulatory processes in neural sensory systems**, especially retinal membrane trafficking/signaling and possibly inner-ear mechanotransduction. But the evidence for a broader or clearly established **homeostatic** role is sparse, and knockout studies show mild or absent phenotypes in at least some sensory contexts. So the claim is **partly supported, but overstated if interpreted as a well-established neural homeostasis gene/protein**. ([proteinatlas.org](https://www.proteinatlas.org/ENSG00000021300-PLEKHB1/brain)) [S14]

## Cited sources

- **[S1]** PLEKHB1 pleckstrin homology domain containing B1 [Homo sapiens (human)] - Gene - NCBI — https://www.ncbi.nlm.nih.gov/gene/58473
- **[S2]** PLEKHB1 pleckstrin homology domain containing B1 [Homo sapiens (human)] - Gene - NCBI — https://www.ncbi.nlm.nih.gov/gene/58473
- **[S3]** PHR1 encodes an abundant, pleckstrin homology domain-containing integral membrane protein in the photoreceptor outer segments - PubMed — https://pubmed.ncbi.nlm.nih.gov/10585447/
- **[S4]** Evectins: vesicular proteins that carry a pleckstrin homology domain and localize to post-Golgi membranes - PubMed — https://pubmed.ncbi.nlm.nih.gov/10200314/
- **[S5]** 
            PHR1, a PH Domain-Containing Protein Expressed in Primary Sensory Neurons - PMC
         — https://pmc.ncbi.nlm.nih.gov/articles/PMC517893/
- **[S6]** PHR1, an integral membrane protein of the inner ear sensory cells, directly interacts with myosin 1c and myosin VIIa - PubMed — https://pubmed.ncbi.nlm.nih.gov/15976448/
- **[S7]** 
            Motor neuron translatome reveals deregulation of SYNGR4 and PLEKHB1 in mutant TDP-43 amyotrophic lateral sclerosis models - PMC
         — https://pmc.ncbi.nlm.nih.gov/articles/PMC7530531/
- **[S8]** 
            PHR1, a PH Domain-Containing Protein Expressed in Primary Sensory Neurons - PMC
         — https://pmc.ncbi.nlm.nih.gov/articles/PMC517893/
- **[S9]** 
        PHR1 is a vesicle-bound protein abundantly expressed in mature olfactory neurons
      -  Johns Hopkins University — https://pure.johnshopkins.edu/en/publications/phr1-is-a-vesicle-bound-protein-abundantly-expressed-in-mature-ol-3/
- **[S10]** PHR1 encodes an abundant, pleckstrin homology domain-containing integral membrane protein in the photoreceptor outer segments - PubMed — https://pubmed.ncbi.nlm.nih.gov/10585447/
- **[S11]** 
            Motor neuron translatome reveals deregulation of SYNGR4 and PLEKHB1 in mutant TDP-43 amyotrophic lateral sclerosis models - PMC
         — https://pmc.ncbi.nlm.nih.gov/articles/PMC7530531/
- **[S12]** PLEKHB1 pleckstrin homology domain containing B1 [Homo sapiens (human)] - Gene - NCBI — https://www.ncbi.nlm.nih.gov/gene/58473
- **[S13]** PHR1 encodes an abundant, pleckstrin homology domain-containing integral membrane protein in the photoreceptor outer segments - PubMed — https://pubmed.ncbi.nlm.nih.gov/10585447/
- **[S14]** Brain tissue expression of PLEKHB1 - Summary - The Human Protein Atlas — https://www.proteinatlas.org/ENSG00000021300-PLEKHB1/brain
