## Supplementary material for "Understanding Substance Dependence: What Differentiates Addictive from Non-Addictive Drugs?": Folder for supplementary files: non-addictive-genes-claims-first100-002-dbndd2-gene-protein-has-been-associated-with-regul-c8cf2d7a6b50.md

# non-addictive-genes-claims-first100-002-c8cf2d7a6b50

**Original claim:** DBNDD2 gene/protein has been associated with regulatory or homeostatic biological processes in neural systems.

## Claim

DBNDD2 gene/protein has been associated with regulatory or homeostatic biological processes in neural systems.

## Verdict

Verdict: Likely True

## Confidence

Medium

## Supporting evidence

DBNDD2 is expressed in the nervous system, which makes a neural role biologically plausible. NCBI lists **broad human brain expression** for DBNDD2, and the mouse ortholog **Dbndd2** is also broadly expressed in adult cerebellum and cortex. Human Protein Atlas additionally places DBNDD2 in a **brain white-matter/signal-transduction** expression cluster and in a **single-cell oligodendrocyte/myelination** cluster. ([ncbi.nlm.nih.gov](https://www.ncbi.nlm.nih.gov/gene/55861)) [S1]

There is direct evidence that DBNDD2/CK1BP has **regulatory biochemical activity**. In a human study, DBNDD2 (then studied as **CK1BP**) was identified from a **human brain cDNA library** as an isoform-selective binding partner of casein kinase-1, binding CK1δ/CK1ε and **inhibiting kinase activity** toward substrates including tau and α-synuclein. NCBI therefore annotates human DBNDD2 as being involved in **negative regulation of protein kinase activity**. That is a bona fide regulatory process, even if not by itself proof of a dedicated neural homeostasis role. ([pubmed.ncbi.nlm.nih.gov](https://pubmed.ncbi.nlm.nih.gov/16618118/?utm_source=openai)) [S2]

Model-organism curation links the mouse ortholog more specifically to neural and homeostatic biology. The current NCBI/MGI record for **Dbndd2** annotates it as enabling **ATPase binding** and acting upstream of or within **monoatomic ion homeostasis**, **neurogenesis**, **neuron projection development**, and the **ERK1/ERK2 cascade**; several of those annotations are marked as based on **direct assay** or **mutant phenotype** evidence. Because Dbndd2 is the mouse ortholog of human DBNDD2, this is meaningful support for the claim of association. ([ncbi.nlm.nih.gov](https://www.ncbi.nlm.nih.gov/gene/52840)) [S3]

The strongest neural-functional evidence comes from recent rodent work on axon growth. In a 2025 rat study, DBNDD2 expression in dorsal root ganglia decreased after **regenerative peripheral axon injury** but increased after **non-regenerative central axon injury**; DBNDD2 also rose with development as regenerative capacity declined. Experimentally, **DBNDD2 knockdown promoted neurite outgrowth** in neonatal and adult DRG neurons and improved **axon regeneration** after sciatic nerve crush. That is a direct association with neural growth-regulatory processes. ([pubmed.ncbi.nlm.nih.gov](https://pubmed.ncbi.nlm.nih.gov/40658351/?utm_source=openai)) [S4]

There is also suggestive evidence for a role in neural **proteostasis**, a key homeostatic process. In a 2021 cell-model study, CK1BP/DBNDD2 overexpression reduced total α-synuclein levels, **reduced α-synuclein oligomerization and insoluble accumulation**, and also lowered aggregation of a polyglutamine protein. The authors interpreted this as potentially beneficial in neurodegenerative disease contexts. Although these were HEK293T-cell experiments rather than neurons or animals, they still support association with protein homeostasis-relevant regulation. ([pmc.ncbi.nlm.nih.gov](https://pmc.ncbi.nlm.nih.gov/articles/PMC8616157/)) [S5]

Finally, the historical mouse literature and curation connect Dbndd2/NKIP to ion-balance biology that is highly relevant to neural homeostasis. NCBI’s GeneRIF summary for the mouse protein states that NKIP/Dbndd2 interacts with the **Na+/K+-ATPase complex**, influencing **cellular ion balance**, **apoptosis**, and **neuronal differentiation**. Since ion gradients and Na+/K+-ATPase activity are foundational for neural excitability and stability, this is a meaningful mechanistic association with neural homeostatic processes. ([ncbi.nlm.nih.gov](https://www.ncbi.nlm.nih.gov/gene/52840)) [S6]

## Contradicting or limiting evidence

The main limitation is that the evidence base is **small and uneven**, and much of it is **not human in vivo evidence**. The clearest neural-functional data come from **mouse/rat** studies and curated model-organism annotations, while the best human biochemical evidence is mainly that DBNDD2 can bind and inhibit CK1-family kinases. That supports a regulatory role, but it does not establish DBNDD2 as a well-validated human neural-homeostasis gene. ([pubmed.ncbi.nlm.nih.gov](https://pubmed.ncbi.nlm.nih.gov/16618118/?utm_source=openai)) [S7]

Some of the “homeostatic” evidence is also indirect. The mouse annotations include **monoatomic ion homeostasis** and **neurogenesis/neuron projection development**, but curated annotations can combine primary experimental data with inference frameworks; they are useful evidence of association, not the same thing as a large independent body of confirmatory studies. ([ncbi.nlm.nih.gov](https://www.ncbi.nlm.nih.gov/gene/52840)) [S8]

The Na+/K+-ATPase angle is suggestive rather than fully resolved mechanistically. A 2021 review of Na,K-ATPase maturation describes NKIP as an **endogenous suppressor** of Na,K-ATPase activity that interacts with the β-subunit in the ER and lysosomes, but explicitly says its role in ER processing is **currently unknown**. That means the homeostatic mechanism is plausible but not fully nailed down. ([link.springer.com](https://link.springer.com/article/10.1007/s00232-021-00184-z)) [S9]

The proteostasis/neurodegeneration evidence is also limited by model choice. The 2021 α-synuclein paper used **HEK293T cells**, not neurons or animal nervous tissue, so it should be treated as mechanistic support rather than definitive evidence of a physiological neural homeostasis role. ([pubmed.ncbi.nlm.nih.gov](https://pubmed.ncbi.nlm.nih.gov/34831053/)) [S10]

## Mechanistic / biological rationale

A plausible mechanistic picture is that DBNDD2 participates in **regulatory control of signaling and membrane-protein function**, with downstream relevance to neural homeostasis. One branch of evidence shows DBNDD2/CK1BP binding and inhibiting **CK1 isoforms**, which can alter phosphorylation-dependent signaling and protein handling. Another branch places the mouse ortholog at the **Na+/K+-ATPase complex**, linked to ion balance, ERK signaling, apoptosis, and neuronal differentiation. Together, these findings make it biologically credible that DBNDD2 could modulate neuronal or glial stability, growth state, and stress responses. ([pubmed.ncbi.nlm.nih.gov](https://pubmed.ncbi.nlm.nih.gov/16618118/?utm_source=openai)) [S11]

Its annotated localization also fits that idea. Human DBNDD2 is predicted at the **cytoplasm**, **endoplasmic reticulum**, and **lysosome**, while the mouse ortholog is annotated to the **ER** and **lysosome**. Those compartments are central to protein quality control, trafficking, and membrane-protein maturation—classic cellular homeostasis functions. ([ncbi.nlm.nih.gov](https://www.ncbi.nlm.nih.gov/gene/55861)) [S12]

The expression data strengthen the neural relevance, especially on the glial/white-matter side. Human Protein Atlas clustering associates DBNDD2 with **white matter/signal transduction** and **oligodendrocytes/myelination**, suggesting it may be more related to neural support-cell biology than to a classic synaptic neurotransmission gene. That is still consistent with a role in neural-system regulation or homeostasis. ([proteinatlas.org](https://www.proteinatlas.org/ENSG00000244274-DBNDD2/brain)) [S13]

## Bottom line

The claim is **probably correct** if “associated with” is interpreted modestly. DBNDD2 has documented links to **negative regulation of kinase activity**, **ion-homeostasis-related biology**, **neurogenesis/neuron projection development**, and **axon growth/regeneration**, with additional suggestive evidence for **proteostasis** in neurodegeneration models. However, the evidence is still **limited**, relies heavily on **model-organism and cell studies**, and does **not yet establish DBNDD2 as a firmly characterized core neural homeostasis gene in humans**. ([ncbi.nlm.nih.gov](https://www.ncbi.nlm.nih.gov/gene/55861)) [S14]

## Cited sources

- **[S1]** DBNDD2 dysbindin domain containing 2 [Homo sapiens (human)] - Gene - NCBI — https://www.ncbi.nlm.nih.gov/gene/55861
- **[S2]** Dysbindin structural homologue CK1BP is an isoform-selective binding partner of human casein kinase-1 - PubMed — https://pubmed.ncbi.nlm.nih.gov/16618118/?utm_source=openai
- **[S3]** Dbndd2 dysbindin domain containing 2 [Mus musculus (house mouse)] - Gene - NCBI — https://www.ncbi.nlm.nih.gov/gene/52840
- **[S4]** Suppressing DBNDD2 promotes neuron growth and axon regeneration in adult mammals - PubMed — https://pubmed.ncbi.nlm.nih.gov/40658351/?utm_source=openai
- **[S5]** 
            CK1BP Reduces α-Synuclein Oligomerization and Aggregation Independent of Serine 129 Phosphorylation - PMC
         — https://pmc.ncbi.nlm.nih.gov/articles/PMC8616157/
- **[S6]** Dbndd2 dysbindin domain containing 2 [Mus musculus (house mouse)] - Gene - NCBI — https://www.ncbi.nlm.nih.gov/gene/52840
- **[S7]** Dysbindin structural homologue CK1BP is an isoform-selective binding partner of human casein kinase-1 - PubMed — https://pubmed.ncbi.nlm.nih.gov/16618118/?utm_source=openai
- **[S8]** Dbndd2 dysbindin domain containing 2 [Mus musculus (house mouse)] - Gene - NCBI — https://www.ncbi.nlm.nih.gov/gene/52840
- **[S9]** Maturation of the Na,K-ATPase in the Endoplasmic Reticulum in Health and Disease | The Journal of Membrane Biology | Springer Nature Link — https://link.springer.com/article/10.1007/s00232-021-00184-z
- **[S10]** CK1BP Reduces α-Synuclein Oligomerization and Aggregation Independent of Serine 129 Phosphorylation - PubMed — https://pubmed.ncbi.nlm.nih.gov/34831053/
- **[S11]** Dysbindin structural homologue CK1BP is an isoform-selective binding partner of human casein kinase-1 - PubMed — https://pubmed.ncbi.nlm.nih.gov/16618118/?utm_source=openai
- **[S12]** DBNDD2 dysbindin domain containing 2 [Homo sapiens (human)] - Gene - NCBI — https://www.ncbi.nlm.nih.gov/gene/55861
- **[S13]** Brain tissue expression of DBNDD2 - Summary - The Human Protein Atlas — https://www.proteinatlas.org/ENSG00000244274-DBNDD2/brain
- **[S14]** DBNDD2 dysbindin domain containing 2 [Homo sapiens (human)] - Gene - NCBI — https://www.ncbi.nlm.nih.gov/gene/55861
