## Supplementary material for "Understanding Substance Dependence: What Differentiates Addictive from Non-Addictive Drugs?": Folder for supplementary files: non-addictive-genes-claims-first100-003-cldnd1-gene-protein-has-been-associated-with-regul-72b8054567bc.md

# non-addictive-genes-claims-first100-003-72b8054567bc

**Original claim:** CLDND1 gene/protein has been associated with regulatory or homeostatic biological processes in neural systems.

## Claim

CLDND1 gene/protein has been associated with regulatory or homeostatic biological processes in neural systems.

## Verdict

Verdict: Likely True

The best available evidence supports an association, but mainly in the **brain endothelium / blood-brain barrier (BBB)** rather than in neurons themselves. CLDND1 (also called claudin-25) is described in reviews as a BBB tight-junction protein with high mRNA expression in brain endothelial cells, localization at tight junctions, and effects on small-molecule permeability when knocked down. Because BBB tight junctions are central to maintaining the brain’s internal milieu, this places CLDND1 within a plausible **neural homeostasis** framework. However, the evidence base is still relatively small, functionally narrower than for major BBB claudins such as CLDN5, and much weaker for direct roles in neuronal or synaptic regulation. ([pmc.ncbi.nlm.nih.gov](https://pmc.ncbi.nlm.nih.gov/articles/PMC11172262/)) [S1]

## Confidence

Medium. The signal is consistent across a few primary studies and recent BBB reviews, but much of the mechanistic literature appears to come from a limited set of groups, and direct evidence for broader neural regulatory/homeostatic roles outside the BBB is sparse. ([pmc.ncbi.nlm.nih.gov](https://pmc.ncbi.nlm.nih.gov/articles/PMC11172262/)) [S2]

## Supporting evidence

A 2024 review of BBB tight-junction proteins lists **claudin-25/CLDND1** as having **very high mRNA expression in vivo in brain endothelial cells**, localization to tight junctions in human brain endothelial cells, and a knockdown phenotype in which paracellular permeability for small molecules rises. The same review explicitly states that the BBB maintains a constant cerebral milieu and is decisive for CNS homeostasis, which makes CLDND1’s BBB role relevant to neural homeostasis even if it is not neuron-intrinsic. ([pmc.ncbi.nlm.nih.gov](https://pmc.ncbi.nlm.nih.gov/articles/PMC11172262/)) [S3]

Primary experimental work supports that interpretation. In adult murine brain, CLDND1 was reported to be abundant in the cerebellum, expressed on endothelial cells, and strongly localized at tight junctions in human brain endothelial cells. In that study, CLDND1 knockdown increased permeability to **small** molecules across confluent human brain endothelial cells, and CLDND1 levels fell transiently after cerebellar hemorrhage, consistent with a barrier-stabilizing role. ([pubmed.ncbi.nlm.nih.gov](https://pubmed.ncbi.nlm.nih.gov/28244141/)) [S4]

Additional mechanistic papers show that CLDND1 is not just present but **regulated** in brain endothelial cells. One study found that the transcription factor **RORα** directly enhances CLDND1 transcription in human brain endothelial cells. Another found that **MZF1** positively regulates CLDND1, and that MZF1 knockdown reduces CLDND1 and increases endothelial permeability. A further paper implicated **EGF/ELK1** signaling in upregulating CLDND1 expression. Together, these studies support the idea that CLDND1 participates in a regulated barrier program rather than being a passive structural bystander. ([pubmed.ncbi.nlm.nih.gov](https://pubmed.ncbi.nlm.nih.gov/28130419/)) [S5]

Post-transcriptional regulation has also been described. In stroke-prone hypertensive rats, a conserved **miR-124** site in Cldnd1 was identified; miR-124 reduced CLDND1 reporter activity and CLDND1 expression in human brain endothelial cells, while lower brain miR-124 levels were associated with higher Cldnd1 expression in vivo. That is direct evidence linking CLDND1 to a regulated cerebrovascular program in the brain. ([pubmed.ncbi.nlm.nih.gov](https://pubmed.ncbi.nlm.nih.gov/29530526/)) [S6]

Expression data add a second, more indirect line of support for “neural system” association. The Human Protein Atlas classifies **CLDND1 as brain-enriched at the RNA level** and places it in expression clusters labeled **“Oligodendrocytes - Myelination”** and **“White matter - Myelination.”** Its nearest transcriptomic neighbors include canonical myelin/oligodendrocyte genes such as **PLP1, MOG, MAG, MOBP, and MBP**. This does not prove function, but it does support a reproducible association with CNS white-matter biology. ([proteinatlas.org](https://www.proteinatlas.org/ENSG00000080822-CLDND1/tissue)) [S7]

## Contradicting or limiting evidence

The strongest limitation is that the evidence is **mostly BBB/endothelial**, not neuronal. The claim says “neural systems,” which can reasonably include the neurovascular unit and BBB, but if interpreted as a claim about **neuronal homeostasis, synaptic regulation, or neural circuit regulation**, the current evidence is much thinner. ([pmc.ncbi.nlm.nih.gov](https://pmc.ncbi.nlm.nih.gov/articles/PMC11172262/)) [S8]

CLDND1 also does **not** appear to be a dominant BBB tight-junction determinant on the level of CLDN5. In the 2019 BBB paper, Cldn25 knockdown altered tight-junction strand morphology, but unlike Cldn5 knockdown, it did **not** significantly reduce transendothelial electrical resistance in the tested brain endothelial cell line. That suggests CLDND1 contributes to junction architecture or selective permeability, but may not be essential for gross barrier tightness in the way major claudins are. ([pmc.ncbi.nlm.nih.gov](https://pmc.ncbi.nlm.nih.gov/articles/PMC11105330/)) [S9]

For oligodendrocyte/white-matter links, the evidence is even more limited. A recent human brain transcriptomics study noted that CLDND1 is highly expressed in oligodendrocytes and showed disease-associated splicing changes, but the authors explicitly stated that its **precise function in the brain remains poorly understood** and framed BBB-homeostasis implications as a hypothesis rather than a demonstrated mechanism. ([pmc.ncbi.nlm.nih.gov](https://pmc.ncbi.nlm.nih.gov/articles/PMC12336083/)) [S10]

There are also data-quality caveats. In the Human Protein Atlas tissue resource, CLDND1 is brain-enriched by RNA and placed in oligodendrocyte/myelination clusters, but the Atlas also flags **low consistency between antibody staining and RNA expression** and says external verification is pending for protein-level tissue interpretation. So the transcriptomic association is useful, but protein localization outside endothelial studies should be treated cautiously. ([proteinatlas.org](https://www.proteinatlas.org/ENSG00000080822-CLDND1/tissue)) [S11]

## Mechanistic / biological rationale

Biologically, the association makes sense. Tight junction proteins in brain endothelial cells help form the BBB, and BBB function is fundamental for controlling paracellular flux, excluding toxins and blood-borne signals, and maintaining the stable extracellular conditions required for normal CNS function. Reviews explicitly describe the BBB as a decisive factor for CNS homeostasis. If CLDND1 contributes to BBB junction organization and selective permeability, then it is mechanistically linked to a homeostatic process in neural systems. ([pmc.ncbi.nlm.nih.gov](https://pmc.ncbi.nlm.nih.gov/articles/PMC11172262/)) [S12]

The CLDND1-specific studies fit that model. CLDND1 localizes to tight junctions in human brain endothelial cells; lowering it increases small-molecule permeability; and its expression is modulated by RORα, MZF1, ELK1/EGF signaling, and miR-124. That pattern is what one would expect for a gene participating in regulated maintenance of barrier properties. ([pubmed.ncbi.nlm.nih.gov](https://pubmed.ncbi.nlm.nih.gov/28244141/)) [S13]

At the same time, the available data suggest CLDND1 may be more of a **supporting or modulatory tight-junction component** than a master barrier determinant. The morphology/permeability effects are real, but they are subtler than those seen for canonical BBB claudins in many systems. So the mechanistic rationale supports **association**, but not yet a claim that CLDND1 is a major central controller of neural homeostasis. ([pmc.ncbi.nlm.nih.gov](https://pmc.ncbi.nlm.nih.gov/articles/PMC11105330/)) [S14]

## Bottom line

The claim is **likely true**, provided “neural systems” is interpreted broadly to include the **brain vasculature / blood-brain barrier**. CLDND1 has credible evidence linking it to **regulated tight-junction biology in brain endothelial cells** and, by extension, to **BBB-dependent CNS homeostasis**. What is **not** well established is a direct role in neurons, synapses, or general neural homeostatic signaling. The strongest conclusion is therefore: **CLDND1 is associated with neurovascular barrier regulation relevant to CNS homeostasis, but evidence for broader neural regulatory roles remains limited.** ([pmc.ncbi.nlm.nih.gov](https://pmc.ncbi.nlm.nih.gov/articles/PMC11172262/)) [S15]

## Cited sources

- **[S1]** 
            The Basic Requirement of Tight Junction Proteins in Blood-Brain Barrier Function and Their Role in Pathologies - PMC
         — https://pmc.ncbi.nlm.nih.gov/articles/PMC11172262/
- **[S2]** 
            The Basic Requirement of Tight Junction Proteins in Blood-Brain Barrier Function and Their Role in Pathologies - PMC
         — https://pmc.ncbi.nlm.nih.gov/articles/PMC11172262/
- **[S3]** 
            The Basic Requirement of Tight Junction Proteins in Blood-Brain Barrier Function and Their Role in Pathologies - PMC
         — https://pmc.ncbi.nlm.nih.gov/articles/PMC11172262/
- **[S4]** Claudin domain containing 1 contributing to endothelial cell adhesion decreases in presence of cerebellar hemorrhage - PubMed — https://pubmed.ncbi.nlm.nih.gov/28244141/
- **[S5]** The retinoic acid receptor-related orphan receptor α positively regulates tight junction protein claudin domain-containing 1 mRNA expression in human brain endothelial cells - PubMed — https://pubmed.ncbi.nlm.nih.gov/28130419/
- **[S6]** Levels of tight junction protein CLDND1 are regulated by microRNA-124 in the cerebellum of stroke-prone spontaneously hypertensive rats - PubMed — https://pubmed.ncbi.nlm.nih.gov/29530526/
- **[S7]** Tissue expression of CLDND1 - Summary - The Human Protein Atlas — https://www.proteinatlas.org/ENSG00000080822-CLDND1/tissue
- **[S8]** 
            The Basic Requirement of Tight Junction Proteins in Blood-Brain Barrier Function and Their Role in Pathologies - PMC
         — https://pmc.ncbi.nlm.nih.gov/articles/PMC11172262/
- **[S9]** 
            Tight junction proteins at the blood–brain barrier: far more than claudin-5 - PMC
         — https://pmc.ncbi.nlm.nih.gov/articles/PMC11105330/
- **[S10]** 
            Brain transcriptomics highlight abundant gene expression and splicing alterations in non-neuronal cells in aFTLD-U - PMC
         — https://pmc.ncbi.nlm.nih.gov/articles/PMC12336083/
- **[S11]** Tissue expression of CLDND1 - Summary - The Human Protein Atlas — https://www.proteinatlas.org/ENSG00000080822-CLDND1/tissue
- **[S12]** 
            The Basic Requirement of Tight Junction Proteins in Blood-Brain Barrier Function and Their Role in Pathologies - PMC
         — https://pmc.ncbi.nlm.nih.gov/articles/PMC11172262/
- **[S13]** Claudin domain containing 1 contributing to endothelial cell adhesion decreases in presence of cerebellar hemorrhage - PubMed — https://pubmed.ncbi.nlm.nih.gov/28244141/
- **[S14]** 
            Tight junction proteins at the blood–brain barrier: far more than claudin-5 - PMC
         — https://pmc.ncbi.nlm.nih.gov/articles/PMC11105330/
- **[S15]** 
            The Basic Requirement of Tight Junction Proteins in Blood-Brain Barrier Function and Their Role in Pathologies - PMC
         — https://pmc.ncbi.nlm.nih.gov/articles/PMC11172262/
