## Supplementary material for "Understanding Substance Dependence: What Differentiates Addictive from Non-Addictive Drugs?": Folder for supplementary files: non-addictive-genes-claims-first100-004-peg10-gene-protein-has-been-associated-with-regula-be960a02d4b8.md

# non-addictive-genes-claims-first100-004-be960a02d4b8

**Original claim:** PEG10 gene/protein has been associated with regulatory or homeostatic biological processes in neural systems.

## Claim

PEG10 gene/protein has been associated with regulatory or homeostatic biological processes in neural systems. ([pmc.ncbi.nlm.nih.gov](https://pmc.ncbi.nlm.nih.gov/articles/PMC8385294/)) [S1]

## Verdict

Verdict: Likely True

Multiple peer-reviewed studies link PEG10 to neural regulatory biology, especially in **human neuronal models** and **neurologic disease contexts**. The strongest evidence is that PEG10 is regulated by neural proteostasis factors such as **UBE3A** and **UBQLN2**, can alter the **neuronal transcriptome**, localizes to **stress granules** and **extracellular vesicles** in neurons, and affects **neuronal migration** during development. However, evidence that PEG10 serves a normal, well-established **physiologic homeostatic role in healthy neural tissue** is still limited, so the claim is better supported as an **association** than as a definitively proven core homeostatic function. ([pmc.ncbi.nlm.nih.gov](https://pmc.ncbi.nlm.nih.gov/articles/PMC8385294/)) [S2]

## Confidence

Medium. The association is supported by more than one direct neural study, but much of the evidence comes from **disease models, perturbation experiments, and human cell culture**, not from definitive demonstration of a normal endogenous homeostatic role in intact neural circuits. ([pmc.ncbi.nlm.nih.gov](https://pmc.ncbi.nlm.nih.gov/articles/PMC8385294/)) [S3]

## Supporting evidence

A 2021 study in human iPSC-derived neurons from Angelman syndrome found that PEG10 is a **UBE3A-regulated protein** in neurons and characterized PEG10 as an RNA-binding, retrovirus-like protein that localizes to **stress granules** and is secreted in **extracellular vesicles**. In that study, PEG10 knockdown and UBE3A reinstatement produced strongly similar changes in the neuronal transcriptome, supporting PEG10 as a downstream regulator of neuronal molecular state. The same paper also reported that PEG10 overexpression altered **neuronal migration** during mouse cortical development, and that PEG10 expression is detectable in human excitatory neurons, inhibitory neurons, and astrocytes, with relatively high expression in human fetal brain. ([pmc.ncbi.nlm.nih.gov](https://pmc.ncbi.nlm.nih.gov/articles/PMC8385294/)) [S4]

That same Angelman study specifically linked PEG10-dependent transcriptional effects to pathways involving **cell adhesion**, **cell-matrix interactions**, and genes relevant to **neuronal migration and development**, including laminin-related programs. Those are classic regulatory processes in neural development and tissue organization, making the claim of neural regulatory association substantially supported. ([pmc.ncbi.nlm.nih.gov](https://pmc.ncbi.nlm.nih.gov/articles/PMC8385294/)) [S5]

A 2023 eLife study provided a second, independent neural association: **UBQLN2 restrains PEG10 to maintain neuronal health in ALS**. Mechanistically, the authors showed that UBQLN2 is necessary and sufficient to limit PEG10 gag-pol abundance, and that loss of UBQLN function leads to PEG10 accumulation through impaired proteasomal control. Because UBQLN2 is a neuronal proteostasis factor implicated in ALS, this places PEG10 within a neural protein-homeostasis network rather than merely a generic developmental pathway. ([pmc.ncbi.nlm.nih.gov](https://pmc.ncbi.nlm.nih.gov/articles/PMC10076021/)) [S6]

The ALS study also found that PEG10 accumulates in human spinal cord disease tissue and that PEG10-derived products can drive transcriptional changes affecting **neuronal genes**, including genes involved in **axon remodeling**. The authors reported that PEG10 pol was among the most significantly upregulated proteins in ALS spinal cord proteomics and that PEG10 signal was present in regions containing somatosensory and apparent motor neurons. This is strong evidence that PEG10 dysregulation is associated with disturbed neuronal regulatory programs in a neurodegenerative setting. ([pmc.ncbi.nlm.nih.gov](https://pmc.ncbi.nlm.nih.gov/articles/PMC10076021/)) [S7]

There is also more indirect evidence from neural progenitor biology. A 2019 study on **TET3 in neural progenitor cells** found altered methylation at the **Peg10 imprinting control region** when TET3 was knocked down, in a context where TET3 was required to maintain **neural stem cell identity**. This does not prove PEG10 itself is the causal effector of that identity program, but it does place the PEG10 locus within an epigenetic regulatory network operating in neural precursor cells. ([pubmed.ncbi.nlm.nih.gov](https://pubmed.ncbi.nlm.nih.gov/31646359/)) [S8]

## Contradicting or limiting evidence

The biggest limitation is that the field has **not yet established a definitive, normal physiologic homeostatic role** for PEG10 in healthy neural tissue. The Angelman paper explicitly states that “the precise role of PEG10 in human neurons remains unknown,” despite showing strong disease-related and transcriptomic effects. That means the current literature supports association and plausible function, but not a settled textbook role in neural homeostasis. ([pmc.ncbi.nlm.nih.gov](https://pmc.ncbi.nlm.nih.gov/articles/PMC8385294/)) [S9]

Species consistency is also limited. In the Angelman study, the authors reported that PEG10 was **highly expressed in humans but almost undetectable in mice**, and they noted failure to detect PEG10 protein in mouse brain in their own proteomic experiments. This suggests that at least some neural roles of PEG10 may be **human-enriched or species-specific**, which weakens broad generalization across mammalian neural biology. ([pmc.ncbi.nlm.nih.gov](https://pmc.ncbi.nlm.nih.gov/articles/PMC8385294/)) [S10]

Much of the mechanistic evidence comes from **overexpression, knockdown, disease tissue, or cell-culture systems** rather than from endogenous manipulation in healthy adult neural circuits. For example, the ALS study inferred neuronal dysfunction from PEG10 accumulation and transcriptional effects, but also noted that further work is needed to determine PEG10’s precise contribution to in vivo neurodegeneration. ([pmc.ncbi.nlm.nih.gov](https://pmc.ncbi.nlm.nih.gov/articles/PMC10076021/)) [S11]

Even within ALS, some results are nuanced. The eLife study reported no dramatic difference in PEG10 immunofluorescence intensity/localization between a cohort of sporadic ALS cases and controls, despite proteomic evidence that PEG10 pol was significantly elevated in ALS spinal cord. That discrepancy suggests the neural disease association is real but method-dependent and not yet fully resolved. ([pmc.ncbi.nlm.nih.gov](https://pmc.ncbi.nlm.nih.gov/articles/PMC10076021/)) [S12]

Finally, the neural progenitor/TET3 evidence is **indirect** with respect to PEG10 function. It shows Peg10 locus regulation in a neural stem-cell maintenance context, but not that PEG10 protein itself executes homeostatic neural control. ([pubmed.ncbi.nlm.nih.gov](https://pubmed.ncbi.nlm.nih.gov/31646359/)) [S13]

## Mechanistic / biological rationale

The available data make biological sense. PEG10 is a **retrotransposon-derived RNA-binding protein** with gag/pol-like features, and in neurons it has been observed in **stress granules** and **extracellular vesicles**. Both compartments are highly relevant to neural regulation: stress granules influence RNA handling during cellular stress, and extracellular vesicles can alter intercellular signaling and cargo exchange. A protein that binds RNA and modulates vesicle content could plausibly influence neuronal state and adaptation. ([pmc.ncbi.nlm.nih.gov](https://pmc.ncbi.nlm.nih.gov/articles/PMC8385294/)) [S14]

PEG10 is also under control of two protein-quality-control systems with known neural importance: **UBE3A** and **UBQLN2**. In neurons, UBE3A regulates PEG10 abundance in the Angelman context, while UBQLN2 restricts PEG10 gag-pol abundance through proteasome-linked degradation in the ALS context. That pattern is consistent with PEG10 being a substrate whose levels must be restrained to preserve neural molecular balance. ([pmc.ncbi.nlm.nih.gov](https://pmc.ncbi.nlm.nih.gov/articles/PMC8385294/)) [S15]

A further mechanistic clue is that PEG10 gag-pol can **self-cleave**, releasing a **nucleocapsid-like fragment** that moves to the nucleus. That fragment contains a zinc-finger region reported to bind nucleic acids, and its expression was sufficient to induce transcriptional changes, with neuronal genes among the most affected. This offers a plausible route by which PEG10 dysregulation could perturb neural regulatory programs, including oxidative-stress-related and axon-remodeling pathways. ([pmc.ncbi.nlm.nih.gov](https://pmc.ncbi.nlm.nih.gov/articles/PMC10076021/)) [S16]

## Bottom line

The claim is **likely true**. PEG10 has been associated with **regulatory processes in neural systems**, including neuronal transcriptome control, stress-granule biology, extracellular-vesicle composition, neuronal migration, and proteostatic restraint linked to neuronal health in ALS. What is **not yet firmly established** is that PEG10 has a broadly accepted, endogenous **homeostatic role in normal healthy neural tissue** independent of disease or experimental perturbation. So the best evidence supports **association and plausible functional involvement**, but not yet a fully settled canonical neural-homeostasis role. ([pmc.ncbi.nlm.nih.gov](https://pmc.ncbi.nlm.nih.gov/articles/PMC8385294/)) [S17]

## Cited sources

- **[S1]** 
            Secreted retrovirus-like GAG-domain-containing protein PEG10 is regulated by UBE3A and is involved in Angelman syndrome pathophysiology - PMC
         — https://pmc.ncbi.nlm.nih.gov/articles/PMC8385294/
- **[S2]** 
            Secreted retrovirus-like GAG-domain-containing protein PEG10 is regulated by UBE3A and is involved in Angelman syndrome pathophysiology - PMC
         — https://pmc.ncbi.nlm.nih.gov/articles/PMC8385294/
- **[S3]** 
            Secreted retrovirus-like GAG-domain-containing protein PEG10 is regulated by UBE3A and is involved in Angelman syndrome pathophysiology - PMC
         — https://pmc.ncbi.nlm.nih.gov/articles/PMC8385294/
- **[S4]** 
            Secreted retrovirus-like GAG-domain-containing protein PEG10 is regulated by UBE3A and is involved in Angelman syndrome pathophysiology - PMC
         — https://pmc.ncbi.nlm.nih.gov/articles/PMC8385294/
- **[S5]** 
            Secreted retrovirus-like GAG-domain-containing protein PEG10 is regulated by UBE3A and is involved in Angelman syndrome pathophysiology - PMC
         — https://pmc.ncbi.nlm.nih.gov/articles/PMC8385294/
- **[S6]** 
            UBQLN2 restrains the domesticated retrotransposon PEG10 to maintain neuronal health in ALS - PMC
         — https://pmc.ncbi.nlm.nih.gov/articles/PMC10076021/
- **[S7]** 
            UBQLN2 restrains the domesticated retrotransposon PEG10 to maintain neuronal health in ALS - PMC
         — https://pmc.ncbi.nlm.nih.gov/articles/PMC10076021/
- **[S8]** Tet3 regulates cellular identity and DNA methylation in neural progenitor cells - PubMed — https://pubmed.ncbi.nlm.nih.gov/31646359/
- **[S9]** 
            Secreted retrovirus-like GAG-domain-containing protein PEG10 is regulated by UBE3A and is involved in Angelman syndrome pathophysiology - PMC
         — https://pmc.ncbi.nlm.nih.gov/articles/PMC8385294/
- **[S10]** 
            Secreted retrovirus-like GAG-domain-containing protein PEG10 is regulated by UBE3A and is involved in Angelman syndrome pathophysiology - PMC
         — https://pmc.ncbi.nlm.nih.gov/articles/PMC8385294/
- **[S11]** 
            UBQLN2 restrains the domesticated retrotransposon PEG10 to maintain neuronal health in ALS - PMC
         — https://pmc.ncbi.nlm.nih.gov/articles/PMC10076021/
- **[S12]** 
            UBQLN2 restrains the domesticated retrotransposon PEG10 to maintain neuronal health in ALS - PMC
         — https://pmc.ncbi.nlm.nih.gov/articles/PMC10076021/
- **[S13]** Tet3 regulates cellular identity and DNA methylation in neural progenitor cells - PubMed — https://pubmed.ncbi.nlm.nih.gov/31646359/
- **[S14]** 
            Secreted retrovirus-like GAG-domain-containing protein PEG10 is regulated by UBE3A and is involved in Angelman syndrome pathophysiology - PMC
         — https://pmc.ncbi.nlm.nih.gov/articles/PMC8385294/
- **[S15]** 
            Secreted retrovirus-like GAG-domain-containing protein PEG10 is regulated by UBE3A and is involved in Angelman syndrome pathophysiology - PMC
         — https://pmc.ncbi.nlm.nih.gov/articles/PMC8385294/
- **[S16]** 
            UBQLN2 restrains the domesticated retrotransposon PEG10 to maintain neuronal health in ALS - PMC
         — https://pmc.ncbi.nlm.nih.gov/articles/PMC10076021/
- **[S17]** 
            Secreted retrovirus-like GAG-domain-containing protein PEG10 is regulated by UBE3A and is involved in Angelman syndrome pathophysiology - PMC
         — https://pmc.ncbi.nlm.nih.gov/articles/PMC8385294/
