## Supplementary material for "Understanding Substance Dependence: What Differentiates Addictive from Non-Addictive Drugs?": Folder for supplementary files: non-addictive-genes-claims-first100-005-slc22a17-gene-protein-has-been-associated-with-reg-0baf9c28909a.md

# non-addictive-genes-claims-first100-005-0baf9c28909a

**Original claim:** SLC22A17 gene/protein has been associated with regulatory or homeostatic biological processes in neural systems.

## Claim

SLC22A17 gene/protein has been associated with regulatory or homeostatic biological processes in neural systems. ([pubmed.ncbi.nlm.nih.gov](https://pubmed.ncbi.nlm.nih.gov/37589051/)) [S1]

## Verdict

Verdict: Likely True

The claim is supported by multiple **preclinical** studies showing SLC22A17 involvement in neural-system regulation, especially **brain iron/redox homeostasis, hippocampal neurogenesis, astroglial state regulation, blood-brain barrier integrity, and stress-related neuronal signaling**. However, the evidence base is still dominated by **mouse and cell-model work**, with limited direct causal evidence in humans. ([pubmed.ncbi.nlm.nih.gov](https://pubmed.ncbi.nlm.nih.gov/41397957)) [S2]

## Confidence

Medium

## Supporting evidence

SLC22A17 is an atypical member of the SLC22 family and is better characterized as a **lipocalin-2 (LCN2/24p3) receptor** that mediates **receptor-mediated endocytosis**, rather than as a conventional organic cation transporter. That biology makes it plausible as a regulator of intracellular iron handling and stress responses. Human transcriptomic resources also show brain expression, with Human Protein Atlas clustering SLC22A17 in a **brain-expression cluster centered on choroid plexus / mixed function**, consistent with neural-system relevance. ([pubmed.ncbi.nlm.nih.gov](https://pubmed.ncbi.nlm.nih.gov/37589051/)) [S3]

The strongest direct support for a **homeostatic neural role** comes from a 2025 mouse study showing that **brain-specific Slc22a17 deletion disrupted hippocampal iron homeostasis and postnatal neurogenesis**. In that study, conditional knockout caused early postnatal mortality, neural stem cell apoptosis, oxidative stress from iron overload, impaired hippocampal development, and cognitive deficits. The authors concluded that Slc22a17 is critical for postnatal neurogenesis through regulation of iron homeostasis in the hippocampus. That is a direct association with both a **homeostatic** process and a **neural** system. ([pubmed.ncbi.nlm.nih.gov](https://pubmed.ncbi.nlm.nih.gov/41397957)) [S4]

That same study also provided a mechanistic link between Slc22a17 and intracellular redox regulation: loss of Slc22a17 altered p62/Nrf2/HO-1 signaling, increased reactive oxygen species, and impaired neural stem-cell survival. In other words, SLC22A17 was not merely expressed in neural tissue; perturbing it changed a core regulatory axis governing **iron balance, oxidative stress, and neural stem-cell maintenance**. ([pubmed.ncbi.nlm.nih.gov](https://pubmed.ncbi.nlm.nih.gov/41397957)) [S5]

A second supporting line of evidence comes from an Alzheimer’s disease-related neurogenesis study. In that work, reactive astroglial **Lcn2/Slc22a17 signaling** was implicated in regulating the balance between reactive and pro-neurogenic astroglial states. The paper reported that **astrocyte-specific Slc22a17 knockdown mimicked the pro-neurogenic effect** seen with Ngfr induction, supporting the idea that Slc22a17 participates in regulating neurogenic plasticity within the hippocampal niche. This is again a regulatory neural-system role, though in a disease-model context. ([pubmed.ncbi.nlm.nih.gov](https://pubmed.ncbi.nlm.nih.gov/37429840/)) [S6]

A third line of evidence concerns **barrier homeostasis** in the brain. In a 2024 stroke study, SLC22A17 was identified as a brain-enriched mediator linked to cell-death pathways; its endothelial expression increased in both **human and mouse brain after cerebral ischemia**. In cultured human brain endothelial cells, SLC22A17 knockdown prevented TNF-α-induced ferroptosis, tight-junction loss, and increased permeability, and in mice it reduced blood-brain barrier leakage after focal ischemia. Since BBB integrity is a canonical homeostatic process in neural systems, this strongly supports association with neural regulation. ([pubmed.ncbi.nlm.nih.gov](https://pubmed.ncbi.nlm.nih.gov/38738428/)) [S7]

There is also evidence connecting SLC22A17 to **neuronal signaling and behavioral regulation**. A 2024 study of chronic stress reported that circulating LCN2 acted in medial prefrontal cortex neurons expressing SLC22A17, and that **Slc22a17 knockdown in mPFC reversed stress-associated calcium-signaling deficits and anxiety-like behavior** in mice. This does not prove a broad physiological role under all conditions, but it does associate SLC22A17 with regulatory neural processes affecting circuit activity and behavior. ([pubmed.ncbi.nlm.nih.gov](https://pubmed.ncbi.nlm.nih.gov/38589429/)) [S8]

Taken together, these studies show SLC22A17 associated with several kinds of neural regulation: **iron homeostasis in neural stem cells, astroglial state control, BBB integrity, and stress-responsive neuronal function**. That breadth makes the claim more credible than if it rested on only one disease model. ([pubmed.ncbi.nlm.nih.gov](https://pubmed.ncbi.nlm.nih.gov/41397957)) [S9]

## Contradicting or limiting evidence

The main limitation is that the best evidence is still **preclinical**. The strongest causal data come from mouse knockouts, viral knockdown, organotypic slices, and cultured human brain endothelial cells, not from human intervention studies or robust human genetics. So while the association is real, its **human physiological importance** is not yet established with high certainty. ([pubmed.ncbi.nlm.nih.gov](https://pubmed.ncbi.nlm.nih.gov/41397957)) [S10]

Also, some of the neural literature is really about the **LCN2–SLC22A17 axis**, where the receptor is one component of a broader inflammatory/iron-signaling pathway. In several papers, the strongest perturbation is on **LCN2**, with SLC22A17 involvement inferred from receptor expression or supported by targeted knockdown in selected cell types. That still supports association, but it weakens any claim that SLC22A17’s independent neural role is already fully defined. ([pubmed.ncbi.nlm.nih.gov](https://pubmed.ncbi.nlm.nih.gov/37429840/)) [S11]

Importantly, even the recent hippocampal paper notes that the physiological and molecular functions of SLC22A17 in the brain had remained **poorly understood**, and the renal-focused review describes several SLC22A17 functions as still **assumed** or incompletely resolved. That argues against calling the claim “highly supported.” ([pmc.ncbi.nlm.nih.gov](https://pmc.ncbi.nlm.nih.gov/articles/PMC12705704/?utm_source=openai)) [S12]

Finally, expression in brain datasets alone does not establish homeostatic function. Human Protein Atlas supports neural-system expression, but expression is only supportive context; the functional claim depends mostly on the experimental papers above. ([proteinatlas.org](https://www.proteinatlas.org/ENSG00000092096-SLC22A17/brain)) [S13]

## Mechanistic / biological rationale

A biologically coherent mechanism exists. SLC22A17 binds LCN2 and mediates endocytic trafficking of iron-associated cargo, positioning it to influence **cellular iron availability, redox balance, and stress signaling**. Because neural stem cells, neurons, astrocytes, and brain endothelial cells are all sensitive to iron-driven oxidative injury, a receptor that modulates iron handling is well placed to affect neural homeostasis. ([pubmed.ncbi.nlm.nih.gov](https://pubmed.ncbi.nlm.nih.gov/37589051/)) [S14]

In hippocampal neural stem cells, Slc22a17 loss caused iron overload, oxidative stress, and apoptosis, with evidence implicating the **p62–Nrf2–HO-1** pathway. That gives a concrete mechanism for how SLC22A17 can preserve neurogenesis: by keeping intracellular iron/redox status within a tolerable range. ([pubmed.ncbi.nlm.nih.gov](https://pubmed.ncbi.nlm.nih.gov/41397957)) [S15]

In brain endothelial cells after ischemia, SLC22A17 appears to couple inflammatory stress to **ferroptosis and tight-junction repression**, thereby influencing BBB permeability. In astroglial and neuronal settings, the LCN2/SLC22A17 axis appears to modulate **reactive-state programs, neurogenic plasticity, and stress-related neuronal activity**. These are all recognizable regulatory processes in neural systems, even if they occur partly in pathological settings. ([pubmed.ncbi.nlm.nih.gov](https://pubmed.ncbi.nlm.nih.gov/38738428/)) [S16]

## Bottom line

Overall, the claim is **likely true**. SLC22A17 has been associated with regulatory and homeostatic processes in neural systems, most convincingly through **hippocampal iron homeostasis/neurogenesis** and **blood-brain barrier regulation**, with additional evidence in **astroglial plasticity** and **stress-related neuronal signaling**. The caution is that this evidence is **mostly preclinical**, so the breadth and importance of SLC22A17’s role in human neural homeostasis are not yet fully settled. ([pubmed.ncbi.nlm.nih.gov](https://pubmed.ncbi.nlm.nih.gov/41397957)) [S17]

## Cited sources

- **[S1]** Role of the SLC22A17/lipocalin-2 receptor in renal endocytosis of proteins/metalloproteins: a focus on iron- and cadmium-binding proteins - PubMed — https://pubmed.ncbi.nlm.nih.gov/37589051/
- **[S2]** Slc22a17 governs postnatal neurogenesis by maintaining the iron homeostasis in hippocampus - PubMed — https://pubmed.ncbi.nlm.nih.gov/41397957
- **[S3]** Role of the SLC22A17/lipocalin-2 receptor in renal endocytosis of proteins/metalloproteins: a focus on iron- and cadmium-binding proteins - PubMed — https://pubmed.ncbi.nlm.nih.gov/37589051/
- **[S4]** Slc22a17 governs postnatal neurogenesis by maintaining the iron homeostasis in hippocampus - PubMed — https://pubmed.ncbi.nlm.nih.gov/41397957
- **[S5]** Slc22a17 governs postnatal neurogenesis by maintaining the iron homeostasis in hippocampus - PubMed — https://pubmed.ncbi.nlm.nih.gov/41397957
- **[S6]** Nerve growth factor receptor (Ngfr) induces neurogenic plasticity by suppressing reactive astroglial Lcn2/Slc22a17 signaling in Alzheimer's disease - PubMed — https://pubmed.ncbi.nlm.nih.gov/37429840/
- **[S7]** SLC22A17 as a Cell Death-Linked Regulator of Tight Junctions in Cerebral Ischemia - PubMed — https://pubmed.ncbi.nlm.nih.gov/38738428/
- **[S8]** Stress increases hepatic release of lipocalin 2 which contributes to anxiety-like behavior in mice - PubMed — https://pubmed.ncbi.nlm.nih.gov/38589429/
- **[S9]** Slc22a17 governs postnatal neurogenesis by maintaining the iron homeostasis in hippocampus - PubMed — https://pubmed.ncbi.nlm.nih.gov/41397957
- **[S10]** Slc22a17 governs postnatal neurogenesis by maintaining the iron homeostasis in hippocampus - PubMed — https://pubmed.ncbi.nlm.nih.gov/41397957
- **[S11]** Nerve growth factor receptor (Ngfr) induces neurogenic plasticity by suppressing reactive astroglial Lcn2/Slc22a17 signaling in Alzheimer's disease - PubMed — https://pubmed.ncbi.nlm.nih.gov/37429840/
- **[S12]** Slc22a17 governs postnatal neurogenesis by maintaining the iron homeostasis in hippocampus - PMC — https://pmc.ncbi.nlm.nih.gov/articles/PMC12705704/?utm_source=openai
- **[S13]** Brain tissue expression of SLC22A17 - Summary - The Human Protein Atlas — https://www.proteinatlas.org/ENSG00000092096-SLC22A17/brain
- **[S14]** Role of the SLC22A17/lipocalin-2 receptor in renal endocytosis of proteins/metalloproteins: a focus on iron- and cadmium-binding proteins - PubMed — https://pubmed.ncbi.nlm.nih.gov/37589051/
- **[S15]** Slc22a17 governs postnatal neurogenesis by maintaining the iron homeostasis in hippocampus - PubMed — https://pubmed.ncbi.nlm.nih.gov/41397957
- **[S16]** SLC22A17 as a Cell Death-Linked Regulator of Tight Junctions in Cerebral Ischemia - PubMed — https://pubmed.ncbi.nlm.nih.gov/38738428/
- **[S17]** Slc22a17 governs postnatal neurogenesis by maintaining the iron homeostasis in hippocampus - PubMed — https://pubmed.ncbi.nlm.nih.gov/41397957
