## Supplementary material for "Understanding Substance Dependence: What Differentiates Addictive from Non-Addictive Drugs?": Folder for supplementary files: non-addictive-genes-claims-first100-006-tph2-gene-protein-has-been-associated-with-regulat-588f5064fa75.md

# non-addictive-genes-claims-first100-006-588f5064fa75

**Original claim:** TPH2 gene/protein has been associated with regulatory or homeostatic biological processes in neural systems.

## Claim

TPH2 gene/protein has been associated with regulatory or homeostatic biological processes in neural systems.

## Verdict

Verdict: Likely True

TPH2 is well established as the predominant neuronal tryptophan hydroxylase isoform and the rate-limiting enzyme for brain serotonin synthesis. Because central serotonin is deeply involved in autonomic, circadian, sleep-wake, stress-response, developmental, and broader neural regulatory functions, and because TPH2 loss or altered expression changes several of those processes in animal models, the claim is supported overall. The main caveat is that the strongest evidence is mechanistic and animal-based; direct human evidence for “homeostatic” roles is more indirect and mostly genetic, imaging, or postmortem. ([ncbi.nlm.nih.gov](https://www.ncbi.nlm.nih.gov/gene/121278)) [S1]

## Confidence

High

## Supporting evidence

Authoritative gene resources and foundational papers agree that TPH2 encodes the brain-enriched tryptophan hydroxylase isoform that catalyzes the first, rate-limiting step in serotonin biosynthesis. NCBI describes TPH2 as catalyzing the rate-limiting step in serotonin production, and early molecular studies showed that TPH2 is predominantly expressed in brainstem/raphe serotonergic neurons, distinguishing it from the mainly peripheral isoform TPH1. ([ncbi.nlm.nih.gov](https://www.ncbi.nlm.nih.gov/gene/121278)) [S2]

That biochemical placement matters biologically because serotonin is a major neuromodulator for neural regulation. Reviews of TPH2 and serotonergic circuitry describe raphe-derived serotonin as participating in control of circadian rhythms, food intake, sleep, mood, cognition, reproduction, motor/autonomic function, and stress-related regulation. In that sense, TPH2 is not just “expressed in neurons”; it sits at a bottleneck for a transmitter system already known to support multiple homeostatic and regulatory functions. ([sciencedirect.com](https://www.sciencedirect.com/science/article/abs/pii/S0306453010003215)) [S3]

Direct functional evidence comes from Tph2-deficient mice. In a widely cited knockout study, loss of brain serotonin caused by Tph2 deletion led to impaired early postnatal growth and altered autonomic control of sleep, breathing, thermoregulation, heart rate, and blood pressure. The same paper concluded that TPH2-derived serotonin is involved in regulation of behavior and autonomic pathways. These are classic homeostatic or regulatory phenotypes in neural systems, not merely disease associations. ([pmc.ncbi.nlm.nih.gov](https://pmc.ncbi.nlm.nih.gov/articles/PMC2700938/)) [S4]

Independent work using adult, conditional depletion of brain serotonin by targeting Tph2 found disrupted daily activity timing, loss of the normal nocturnal “siesta,” and evidence that adult serotonin is required for sleep-wake homeostasis and normal circadian behavioral control. This strengthens the case that TPH2 is linked not only to development but also to ongoing homeostatic regulation in the mature nervous system. ([pubmed.ncbi.nlm.nih.gov](https://pubmed.ncbi.nlm.nih.gov/27656022/)) [S5]

Sleep-related evidence also extends across species. In zebrafish and mice, serotonergic raphe signaling and tph2 deficiency were linked to increased arousal, lighter sleep, and attenuated sleep homeostasis; in that study, tph2 mutant animals had difficulty initiating and maintaining normal sleep states. Cross-species convergence makes the association more credible than a single-model finding. ([pmc.ncbi.nlm.nih.gov](https://pmc.ncbi.nlm.nih.gov/articles/PMC6706304/)) [S6]

Developmental and circuit-level evidence also supports a regulatory role. In a knock-in model replacing Tph2 with a reporter, loss of brain serotonin altered serotonergic circuitry formation, changed innervation patterns in several brain regions, and was accompanied by increased hippocampal BDNF expression, interpreted as a possible feedback mechanism tuning serotonergic circuit formation. A related study of brain-specific Tph2 inactivation reported effects on growth, body-weight regulation, receptor adaptations, and noradrenergic markers while emphasizing serotonin’s role in developmental processes, synaptic plasticity, metabolic homeostasis, neuroendocrine function, appetite, energy expenditure, respiratory rate, and sleep. ([nature.com](https://www.nature.com/articles/mp2012128)) [S7]

There is also evidence that TPH2 itself is regulated in ways characteristic of homeostatic systems. Reviews describe daily rhythmic tph2 expression in raphe/retina, correlation with circadian changes in TPH protein and serotonin synthesis in projections to circadian centers such as the suprachiasmatic nucleus, and regulation by glucocorticoids and environmental stressors. Those findings place TPH2 within feedback-linked circadian and stress-response control loops rather than as a static housekeeping enzyme. ([pmc.ncbi.nlm.nih.gov](https://pmc.ncbi.nlm.nih.gov/articles/PMC3587664/)) [S8]

Human evidence is less direct but still consistent with a regulatory role. Reviews of TPH2 in cognitive control and emotion regulation summarize associations of TPH2 variants with affective traits, stress responsivity, and disorders involving dysregulation of cognitive and emotional control. Since those are neural regulatory phenotypes, they support the claim at least at the level of association, though not as strongly as the animal intervention studies. ([sciencedirect.com](https://www.sciencedirect.com/science/article/abs/pii/S0306453010003215)) [S9]

## Contradicting or limiting evidence

The biggest limitation is definitional: TPH2 is most directly a serotonin-synthesis enzyme, not a canonical “homeostatic regulator” in the narrow sense used for mechanisms like synaptic scaling or ion-channel set-point control. So the claim is best understood as “TPH2 is associated with neural regulatory/homeostatic processes through serotonergic signaling,” not that TPH2 itself is a specialized homeostasis gene in every usage of that term. ([ncbi.nlm.nih.gov](https://www.ncbi.nlm.nih.gov/gene/121278)) [S10]

Much of the strongest evidence comes from animal genetics, especially knockout or conditional-deletion models. Those models are informative for causality, but they do not automatically prove identical effect sizes or phenotypic breadth in humans. Human literature is dominated by associations, imaging correlates, and postmortem findings rather than direct experimental perturbation of TPH2. ([sciencedirect.com](https://www.sciencedirect.com/science/article/abs/pii/S0306453010003215)) [S11]

Some data also suggest that TPH2 deficiency does not broadly reorganize all neural homeostatic systems. In one knock-in model, major regulatory components of serotonergic neurotransmission such as SERT, MAO, and several 5-HT1A receptor measures did not show strong compensatory changes, and basal extracellular levels of several other neurotransmitters were largely unchanged. The authors explicitly suggested that Tph2 loss-of-function may not be substantially offset by homeostatic mechanisms. That does not refute a role in homeostatic outputs like sleep or autonomic control, but it does limit stronger claims about global neurotransmitter homeostasis. ([pmc.ncbi.nlm.nih.gov](https://pmc.ncbi.nlm.nih.gov/articles/PMC3536482/)) [S12]

There are also mild contradictions in the regulatory literature depending on species and context. For example, glucocorticoid effects on tph2 expression were reported in opposite directions in mice versus rats in one review, implying that TPH2 regulation can be species-, tissue-, and stress-paradigm-dependent. ([pmc.ncbi.nlm.nih.gov](https://pmc.ncbi.nlm.nih.gov/articles/PMC3587664/)) [S13]

## Mechanistic / biological rationale

Mechanistically, the rationale is straightforward. TPH2 controls the rate-limiting step of serotonin synthesis in raphe-centered serotonergic neurons, so changes in TPH2 expression or function alter central serotonin availability. Because serotonin projections innervate hypothalamic, brainstem autonomic, limbic, cortical, and circadian-control regions, altered TPH2 can plausibly shift neural set points for sleep-wake cycling, thermoregulation, respiratory control, stress responsivity, and aspects of developmental circuit tuning. ([pubmed.ncbi.nlm.nih.gov](https://pubmed.ncbi.nlm.nih.gov/14563478/)) [S14]

A second mechanistic point is that TPH2 is itself dynamically regulated. Circadian rhythmicity of tph2 expression, glucocorticoid sensitivity, and stress-responsive transcription all imply that the enzyme is embedded in feedback systems that help organisms adapt brain serotonin output to day-night cycles and environmental demands. That is biologically consistent with involvement in regulatory and homeostatic processes. ([pmc.ncbi.nlm.nih.gov](https://pmc.ncbi.nlm.nih.gov/articles/PMC3587664/)) [S15]

Finally, developmental studies suggest that brain serotonin made via TPH2 influences wiring and plasticity of serotonergic circuits and downstream targets. If a molecule shapes both acute transmitter availability and longer-term circuit formation, it can affect neural homeostasis at multiple timescales: immediate physiological regulation and developmental stabilization of network architecture. ([nature.com](https://www.nature.com/articles/mp2012128)) [S16]

## Bottom line

The claim is **likely true**. TPH2 is not merely loosely correlated with neural regulation; it is the central serotonin-synthesis bottleneck, and perturbing it changes several neural regulatory/homeostatic domains, especially **sleep-wake control, circadian timing, autonomic regulation, thermoregulation, breathing, stress-axis interactions, and aspects of circuit development**. The evidence is strongest in **animal and mechanistic studies**, while **human evidence is more indirect**, so the safest formulation is that TPH2 is **credibly associated with**—and in animal models often causally involved in—regulatory and homeostatic biological processes in neural systems. ([pmc.ncbi.nlm.nih.gov](https://pmc.ncbi.nlm.nih.gov/articles/PMC2700938/)) [S17]

## Cited sources

- **[S1]** TPH2 tryptophan hydroxylase 2 [Homo sapiens (human)] - Gene - NCBI — https://www.ncbi.nlm.nih.gov/gene/121278
- **[S2]** TPH2 tryptophan hydroxylase 2 [Homo sapiens (human)] - Gene - NCBI — https://www.ncbi.nlm.nih.gov/gene/121278
- **[S3]** Tryptophan hydroxylase-2 (TPH2) in disorders of cognitive control and emotion regulation: A perspective - ScienceDirect — https://www.sciencedirect.com/science/article/abs/pii/S0306453010003215
- **[S4]** 
            Growth retardation and altered autonomic control in mice lacking brain serotonin - PMC
         — https://pmc.ncbi.nlm.nih.gov/articles/PMC2700938/
- **[S5]** Adult Brain Serotonin Deficiency Causes Hyperactivity, Circadian Disruption, and Elimination of Siestas - PubMed — https://pubmed.ncbi.nlm.nih.gov/27656022/
- **[S6]** 
            The Serotonergic Raphe Promote Sleep in Zebrafish and Mice - PMC
         — https://pmc.ncbi.nlm.nih.gov/articles/PMC6706304/
- **[S7]** Lack of brain serotonin affects postnatal development and serotonergic neuronal circuitry formation | Molecular Psychiatry — https://www.nature.com/articles/mp2012128
- **[S8]** 
            Advances in Tryptophan Hydroxylase-2 Gene Expression Regulation: New Insights into Serotonin-Stress Interaction and Clinical Implications - PMC
         — https://pmc.ncbi.nlm.nih.gov/articles/PMC3587664/
- **[S9]** Tryptophan hydroxylase-2 (TPH2) in disorders of cognitive control and emotion regulation: A perspective - ScienceDirect — https://www.sciencedirect.com/science/article/abs/pii/S0306453010003215
- **[S10]** TPH2 tryptophan hydroxylase 2 [Homo sapiens (human)] - Gene - NCBI — https://www.ncbi.nlm.nih.gov/gene/121278
- **[S11]** Tryptophan hydroxylase-2 (TPH2) in disorders of cognitive control and emotion regulation: A perspective - ScienceDirect — https://www.sciencedirect.com/science/article/abs/pii/S0306453010003215
- **[S12]** 
            Deficient serotonin neurotransmission and depression-like serotonin biomarker alterations in tryptophan hydroxylase 2 (Tph2) loss-of-function mice - PMC
         — https://pmc.ncbi.nlm.nih.gov/articles/PMC3536482/
- **[S13]** 
            Advances in Tryptophan Hydroxylase-2 Gene Expression Regulation: New Insights into Serotonin-Stress Interaction and Clinical Implications - PMC
         — https://pmc.ncbi.nlm.nih.gov/articles/PMC3587664/
- **[S14]** A unique central tryptophan hydroxylase isoform - PubMed — https://pubmed.ncbi.nlm.nih.gov/14563478/
- **[S15]** 
            Advances in Tryptophan Hydroxylase-2 Gene Expression Regulation: New Insights into Serotonin-Stress Interaction and Clinical Implications - PMC
         — https://pmc.ncbi.nlm.nih.gov/articles/PMC3587664/
- **[S16]** Lack of brain serotonin affects postnatal development and serotonergic neuronal circuitry formation | Molecular Psychiatry — https://www.nature.com/articles/mp2012128
- **[S17]** 
            Growth retardation and altered autonomic control in mice lacking brain serotonin - PMC
         — https://pmc.ncbi.nlm.nih.gov/articles/PMC2700938/
