## Supplementary material for "Understanding Substance Dependence: What Differentiates Addictive from Non-Addictive Drugs?": Folder for supplementary files: non-addictive-genes-claims-first100-007-rps19-gene-protein-has-been-associated-with-regula-3e6e1f8a11f3.md

# non-addictive-genes-claims-first100-007-3e6e1f8a11f3

**Original claim:** RPS19 gene/protein has been associated with regulatory or homeostatic biological processes in neural systems.

## Claim

**RPS19 gene/protein has been associated with regulatory or homeostatic biological processes in neural systems.**

## Verdict

Verdict: Mixed Evidence

RPS19 **has some association** with neural regulatory contexts, but the evidence is **indirect and not yet strong enough** to say it has a well-established, specific homeostatic role in neural systems. The strongest support comes from curated gene annotations placing mouse **Rps19** at the **synapse** and upstream of **Notch signaling**, from animal models showing **brain/behavioral abnormalities** when Rps19 is altered, and from extraribosomal studies showing RPS19 can regulate **immune/inflammatory signaling** pathways that are relevant to brain homeostasis. However, the core, best-established function of RPS19 remains **ribosome biogenesis/translation**, and direct evidence that RPS19 normally serves a dedicated neural homeostatic regulator in humans or mammals is limited. ([ncbi.nlm.nih.gov](https://www.ncbi.nlm.nih.gov/gene/20085)) [S1]

## Confidence

**Medium.** The claim is broad enough that some supporting associations exist, but most neural links are from **model organisms, phenotypic studies, database curation, or mechanistic inference**, not from definitive human neurobiology experiments. ([ncbi.nlm.nih.gov](https://www.ncbi.nlm.nih.gov/gene/20085)) [S2]

## Supporting evidence

RPS19 is not just a generic housekeeping gene in databases: the mouse NCBI/Alliance entry states that **Rps19 is active in synapse** and acts upstream of or within the **Notch signaling pathway**. That is a direct curated association of this gene with a neural compartment and a canonical regulatory pathway that is important in nervous-system development and cell-state control. ([ncbi.nlm.nih.gov](https://www.ncbi.nlm.nih.gov/gene/20085)) [S3]

Independent annotation resources also connect human RPS19 to neural subcellular structures. The Human Protein Atlas/UniProt-linked annotations list **postsynaptic density** among associated cellular components, alongside molecular functions such as **fibroblast growth factor binding** and **protein kinase binding**. These do not prove a dedicated neural function, but they do place RPS19 in a molecular neighborhood that is relevant to synaptic regulation. ([proteinatlas.org](https://www.proteinatlas.org/ENSG00000105372-RPS19/summary/gene)) [S4]

Functional animal data also point toward nervous-system involvement. In a mouse **Rps19(Arg67del)** model, investigators reported **brain abnormalities**, **moderate cognitive impairment**, and **neuromuscular dysfunction with gait abnormalities**. That study does not prove a homeostatic role, but it does show that perturbing Rps19 can affect neural structure and behavior. ([pubmed.ncbi.nlm.nih.gov](https://pubmed.ncbi.nlm.nih.gov/27643579/)) [S5]

A separate mouse study targeting the **extraribosomal** function of RP S19 found altered behavior: the knock-in mice showed **increased grooming** and **enhanced contextual freezing**, which the authors interpreted as evidence that RP S19 oligomers may contribute to anxiety-like behavior. Again, this is not a direct demonstration of neural homeostasis, but it is a neural/behavioral association beyond erythropoiesis. ([lup.lub.lu.se](https://lup.lub.lu.se/record/0c191942-2da0-48c1-8a01-4869a6c02d7a)) [S6]

Developmental vertebrate data also support a nervous-system association. The zebrafish rps19 entry states that the gene acts upstream of or within **brain segmentation**, and experimental work on rps19 deficiency reported developmental abnormalities with **suppression of the neural field** that could affect the **eye, brain, and neural crest derivatives**. That puts rps19 squarely in a developmental neural-regulatory context, though still not specifically “homeostasis” in the mature nervous system. ([zfin.org](https://zfin.org/ZDB-GENE-040426-1716)) [S7]

There is also evidence for **regulatory** activity outside the ribosome. Human RPS19 can bind its own mRNA 5′UTR, and the authors proposed this as a **regulatory mechanism** controlling RPS19 expression. While that study was not neural, it supports the idea that RPS19 can have extraribosomal regulatory behavior rather than being only a static structural ribosomal component. ([pubmed.ncbi.nlm.nih.gov](https://pubmed.ncbi.nlm.nih.gov/20395159/)) [S8]

Finally, extracellular RPS19 has immunoregulatory properties that are plausibly relevant to neural homeostasis. RPS19 was shown to bind **macrophage migration inhibitory factor (MIF)** and attenuate its pro-inflammatory function, and another study showed extracellular RPS19 can signal through **C5aR1**. Since complement/C5a signaling is implicated in brain development, glial signaling, and synapse pruning, these findings provide a biologically plausible route by which RPS19 could influence neural homeostatic processes. ([pubmed.ncbi.nlm.nih.gov](https://pubmed.ncbi.nlm.nih.gov/19155217/)) [S9]

## Contradicting or limiting evidence

The main limitation is that RPS19’s **best-established function** is still classic ribosome biology: it is a **40S ribosomal protein** required for **pre-rRNA processing** and **maturation of 40S ribosomal subunits**, and pathogenic variants are best known for causing **Diamond-Blackfan anemia**. That is the dominant, high-confidence biology for this gene. ([ncbi.nlm.nih.gov](https://www.ncbi.nlm.nih.gov/gene/6223)) [S10]

Also, the neural links are not all equally strong. Some come from **curated database annotations** or **orthology-based inferences**, which are useful but weaker than direct neural functional experiments. For example, synapse/postsynaptic-density associations appear in curated resources, but there is not a large body of primary literature showing endogenous RPS19 acting as a canonical synaptic homeostasis factor in neurons or glia. ([ncbi.nlm.nih.gov](https://www.ncbi.nlm.nih.gov/gene/20085)) [S11]

The animal-model studies show that altering Rps19 can produce **neurological phenotypes**, but that does not necessarily mean RPS19 normally performs a specific homeostatic control function in neural tissue. Disrupting an essential ribosomal protein can cause broad developmental or translational stress effects that secondarily impact the brain. ([pubmed.ncbi.nlm.nih.gov](https://pubmed.ncbi.nlm.nih.gov/27643579/)) [S12]

Likewise, the immune-regulatory studies involving **MIF** and **C5aR1** were not primarily performed in normal neural tissue. Their relevance to brain homeostasis is therefore partly **inferred**, based on what is known about those pathways in microglia, astrocytes, and complement biology. That inference is reasonable, but still indirect. ([pubmed.ncbi.nlm.nih.gov](https://pubmed.ncbi.nlm.nih.gov/19155217/)) [S13]

## Mechanistic / biological rationale

A plausible rationale exists for why RPS19 could matter in neural regulation. Neurons depend heavily on tightly controlled **protein synthesis**, local ribosomal function, and proteostasis at synapses. Because RPS19 is required for small-subunit biogenesis and translation-related processes, perturbing it could alter synaptic maintenance, neuronal development, or activity-dependent protein production. ([proteinatlas.org](https://www.proteinatlas.org/ENSG00000105372-RPS19/summary/gene)) [S14]

RPS19 also appears capable of **extraribosomal regulation**. It can bind its own mRNA, interact with **MIF**, and signal through **C5aR1** in non-neural contexts. Those interactions matter because inflammatory/complement pathways are now recognized as part of normal and pathological neural homeostasis, including glial surveillance, synapse remodeling, and neurodevelopmental signaling. So, it is mechanistically plausible that extracellular or extraribosomal RPS19 could influence neural regulatory states. This is, however, still partly an inference from adjacent evidence rather than a settled neural mechanism. ([pubmed.ncbi.nlm.nih.gov](https://pubmed.ncbi.nlm.nih.gov/20395159/)) [S15]

Notch signaling provides another plausible bridge. The mouse gene annotation linking Rps19 to **Notch signaling** is notable because Notch is central to neural progenitor maintenance, differentiation, and cell-state regulation. But at present, the specific Rps19→Notch connection in neural tissue is not well developed in the primary literature. ([ncbi.nlm.nih.gov](https://www.ncbi.nlm.nih.gov/gene/20085)) [S16]

## Bottom line

RPS19 is **not well established as a dedicated neural homeostasis gene**, but it **has been associated** with neural regulatory contexts through curated synaptic annotations, vertebrate developmental data, neurological phenotypes in mutant animals, and extraribosomal immune-regulatory interactions that could plausibly affect brain homeostasis. So the claim is **not unsupported**, but the evidence is **too indirect and heterogeneous** to rate as likely true in a strong, specific sense. The fairest assessment is **Mixed Evidence**. ([ncbi.nlm.nih.gov](https://www.ncbi.nlm.nih.gov/gene/20085)) [S17]

## Cited sources

- **[S1]** https://www.ncbi.nlm.nih.gov/gene/20085 — https://www.ncbi.nlm.nih.gov/gene/20085
- **[S2]** https://www.ncbi.nlm.nih.gov/gene/20085 — https://www.ncbi.nlm.nih.gov/gene/20085
- **[S3]** https://www.ncbi.nlm.nih.gov/gene/20085 — https://www.ncbi.nlm.nih.gov/gene/20085
- **[S4]** https://www.proteinatlas.org/ENSG00000105372-RPS19/summary/gene — https://www.proteinatlas.org/ENSG00000105372-RPS19/summary/gene
- **[S5]** https://pubmed.ncbi.nlm.nih.gov/27643579/ — https://pubmed.ncbi.nlm.nih.gov/27643579/
- **[S6]** https://lup.lub.lu.se/record/0c191942-2da0-48c1-8a01-4869a6c02d7a — https://lup.lub.lu.se/record/0c191942-2da0-48c1-8a01-4869a6c02d7a
- **[S7]** https://zfin.org/ZDB-GENE-040426-1716 — https://zfin.org/ZDB-GENE-040426-1716
- **[S8]** https://pubmed.ncbi.nlm.nih.gov/20395159/ — https://pubmed.ncbi.nlm.nih.gov/20395159/
- **[S9]** https://pubmed.ncbi.nlm.nih.gov/19155217/ — https://pubmed.ncbi.nlm.nih.gov/19155217/
- **[S10]** https://www.ncbi.nlm.nih.gov/gene/6223 — https://www.ncbi.nlm.nih.gov/gene/6223
- **[S11]** https://www.ncbi.nlm.nih.gov/gene/20085 — https://www.ncbi.nlm.nih.gov/gene/20085
- **[S12]** https://pubmed.ncbi.nlm.nih.gov/27643579/ — https://pubmed.ncbi.nlm.nih.gov/27643579/
- **[S13]** https://pubmed.ncbi.nlm.nih.gov/19155217/ — https://pubmed.ncbi.nlm.nih.gov/19155217/
- **[S14]** https://www.proteinatlas.org/ENSG00000105372-RPS19/summary/gene — https://www.proteinatlas.org/ENSG00000105372-RPS19/summary/gene
- **[S15]** https://pubmed.ncbi.nlm.nih.gov/20395159/ — https://pubmed.ncbi.nlm.nih.gov/20395159/
- **[S16]** https://www.ncbi.nlm.nih.gov/gene/20085 — https://www.ncbi.nlm.nih.gov/gene/20085
- **[S17]** https://www.ncbi.nlm.nih.gov/gene/20085 — https://www.ncbi.nlm.nih.gov/gene/20085
