## Supplementary material for "Understanding Substance Dependence: What Differentiates Addictive from Non-Addictive Drugs?": Folder for supplementary files: non-addictive-genes-claims-first100-008-gpx3-gene-protein-has-been-associated-with-regulat-7675760c0d38.md

# non-addictive-genes-claims-first100-008-7675760c0d38

**Original claim:** GPX3 gene/protein has been associated with regulatory or homeostatic biological processes in neural systems.

## Claim

GPX3 gene/protein has been associated with regulatory or homeostatic biological processes in neural systems.

## Verdict

Verdict: Likely True

## Confidence

Medium

## Supporting evidence

GPX3 encodes a **secreted extracellular glutathione peroxidase** that reduces hydrogen peroxide and lipid/organic hydroperoxides using glutathione, placing it squarely in antioxidant and redox-control biology rather than in a purely structural role. Authoritative gene/protein resources also indicate that GPX3 is present in the **extracellular space** and is expressed in the **brain**, making a neural homeostatic role biologically plausible from the outset. ([ncbi.nlm.nih.gov](https://www.ncbi.nlm.nih.gov/gene/2878)) [S1]

There is direct literature linking **Gpx3 expression to developmental neural-supportive programs**. In a neonatal mouse study, a distinct **CD11c+ microglial subset** was described as a major source of **Igf1, Spp1, Lgals1, and Gpx3**, and these genes were reported as being associated with **neurogenic and myelinogenic processes** in the developing brain. The paper’s framing is important: it does not just say Gpx3 is present in brain tissue; it places Gpx3 inside a microglial program tied to **neurodevelopment and myelination**, both core regulatory/homeostatic neural processes. ([pmc.ncbi.nlm.nih.gov](https://pmc.ncbi.nlm.nih.gov/articles/PMC5686552/?utm_source=openai)) [S2]

Astrocyte-focused literature also supports an association between GPX3 and **brain redox homeostasis**. A review of brain glutathione metabolism notes that **GPX1 and GPX3 expression has been described in astrocytes in culture**, within the broader context that astrocytes supply glutathione precursors and antioxidant support to neurons. That same review emphasizes that disruption of astrocytic glutathione-related support compromises the homeostasis of other brain cell types and can contribute to neurodegeneration. ([pmc.ncbi.nlm.nih.gov](https://pmc.ncbi.nlm.nih.gov/articles/PMC10179008/)) [S3]

More specifically, an experimental study in **human astroglial cells** found that amyloid-β\(_{25-35}\) activated an **Nrf2-dependent antioxidant response** that included induction of **GPX3** along with other antioxidant genes. The authors’ model explicitly places astrocytic GPX3 within a stress-response program that is protective for astrocytes but can interact with neuron-facing biology through altered glutamate handling. This is not proof of normal physiology, but it is direct evidence that GPX3 participates in a neural-glial antioxidant regulatory response. ([pubmed.ncbi.nlm.nih.gov](https://pubmed.ncbi.nlm.nih.gov/34829555/)) [S4]

Several reviews of neuron–astrocyte redox coupling go further and synthesize prior work as showing that **neuronal activity regulates astrocytic antioxidant programs**, including **GPX3** and SOD3, as part of **non-cell-autonomous antioxidant support** for neurons. That places GPX3 conceptually in a recognized framework of neural homeostasis: neuron–glia signaling that stabilizes extracellular oxidative conditions and supports neuronal resilience. ([pmc.ncbi.nlm.nih.gov](https://pmc.ncbi.nlm.nih.gov/articles/PMC10011399/?utm_source=openai)) [S5]

Human disease-omics studies provide additional association-level support. In Alzheimer’s disease cohorts, **CSF GPX3 levels decreased across worsening amyloid/tau categories** and were associated with biomarkers of **neurodegeneration and neuroinflammation**. A separate ALS risk-locus study reported **lower GPX3 in brain**, lower GPX3 with more advanced disease, and a **zebrafish gpx3 knockdown motor phenotype** that was rescued by gpx3 expression. These data do not by themselves prove a homeostatic role, but they do show that GPX3 tracks with neural-system integrity and motor-neural function. ([pubmed.ncbi.nlm.nih.gov](https://pubmed.ncbi.nlm.nih.gov/38809917/)) [S6]

## Contradicting or limiting evidence

The main limitation is that much of the neural evidence is **associational, transcriptomic, or biomarker-based**, not a direct demonstration that GPX3 is a master regulator of neural homeostasis in vivo. For example, the neonatal microglia/myelination paper linked Gpx3 to a neurodevelopmental gene program, but its strongest causal validation within that program focused on **IGF1**, not GPX3 itself. So the study supports association more strongly than GPX3-specific necessity. ([pmc.ncbi.nlm.nih.gov](https://pmc.ncbi.nlm.nih.gov/articles/PMC5686552/?utm_source=openai)) [S7]

Similarly, the Alzheimer’s and ALS studies show that GPX3 levels change with disease state and genetic risk, but they do **not establish causality**. A falling CSF or brain GPX3 signal could reflect a downstream consequence of neurodegeneration, inflammation, altered cell composition, or oxidative stress rather than a primary homeostatic driver. ([pubmed.ncbi.nlm.nih.gov](https://pubmed.ncbi.nlm.nih.gov/38809917/)) [S8]

The astroglial amyloid-β study is also limited by model choice: it used **U373 astroglial cells** and an **Aβ fragment** rather than intact mammalian brain tissue under physiological conditions. That makes it useful mechanistically, but weaker as evidence for normal in vivo neural homeostasis. ([pubmed.ncbi.nlm.nih.gov](https://pubmed.ncbi.nlm.nih.gov/34829555/)) [S9]

Finally, some of the literature placing GPX3 in activity-dependent astrocyte antioxidant support comes from **reviews synthesizing prior work**, rather than from a large body of GPX3-specific functional experiments in intact neural circuits. So while the overall direction of evidence is supportive, the field is not yet at the level where one could say GPX3’s neural homeostatic role is definitively established. ([pmc.ncbi.nlm.nih.gov](https://pmc.ncbi.nlm.nih.gov/articles/PMC10011399/?utm_source=openai)) [S10]

## Mechanistic / biological rationale

Mechanistically, the claim makes sense. GPX3 is an **extracellular oxidoreductase/peroxidase**, and the extracellular compartment is crucial in neural tissue because synapses, axons, myelin, glial processes, and vascular interfaces are all vulnerable to oxidative stress. A secreted enzyme that detoxifies **H\(_2\)O\(_2\)** and **lipid hydroperoxides** is well positioned to contribute to **redox homeostasis** around neural cells. ([ncbi.nlm.nih.gov](https://www.ncbi.nlm.nih.gov/gene/2878)) [S11]

This is especially plausible for **astrocyte–neuron** and **microglia–oligodendrocyte/neuron** interactions. Astrocytes are central providers of antioxidant support and glutathione-related metabolites in the CNS, and reviews of brain glutathione biology place GPX3 within astrocytic antioxidant machinery. In parallel, developmental microglia expressing Gpx3 in myelinogenic/neurogenic contexts suggest that GPX3 could help buffer oxidative conditions during periods of intense membrane synthesis, pruning, and maturation. That is a biologically coherent homeostatic role, even if the degree of GPX3-specific causal importance remains only partly resolved. ([pmc.ncbi.nlm.nih.gov](https://pmc.ncbi.nlm.nih.gov/articles/PMC10179008/)) [S12]

The disease associations are also mechanistically consistent. In AD and ALS, oxidative stress, neuroinflammation, and neuron–glia dysfunction are prominent features; therefore, altered GPX3 in CSF, brain, or genetically influenced risk loci is consistent with a role in **maintenance of neural redox balance** rather than incidental expression alone. Still, that step from plausibility to proof is where the evidence remains incomplete. ([pubmed.ncbi.nlm.nih.gov](https://pubmed.ncbi.nlm.nih.gov/38809917/)) [S13]

## Bottom line

The claim is **likely true** because GPX3 has been linked in the literature to **brain redox homeostasis, astrocytic antioxidant support, developmental microglial programs involved in neurogenesis/myelination, and neural disease states such as AD and ALS**. However, the evidence is stronger for **association** than for **definitive GPX3-specific causal control** of neural homeostasis. In short: GPX3 is reasonably supported as a **neural-system-associated antioxidant/homeostatic factor**, but not yet as a fully established central regulator. ([pharos.nih.gov](https://pharos.nih.gov/targets/GPX3)) [S14]

## Cited sources

- **[S1]** GPX3 glutathione peroxidase 3 [Homo sapiens (human)] - Gene - NCBI — https://www.ncbi.nlm.nih.gov/gene/2878
- **[S2]** A novel microglial subset plays a key role in myelinogenesis in developing brain - PMC — https://pmc.ncbi.nlm.nih.gov/articles/PMC5686552/?utm_source=openai
- **[S3]** 
            Appraising the Role of Astrocytes as Suppliers of Neuronal Glutathione Precursors - PMC
         — https://pmc.ncbi.nlm.nih.gov/articles/PMC10179008/
- **[S4]** Amyloid-β 25-35 Induces Neurotoxicity through the Up-Regulation of Astrocytic System Xc - PubMed — https://pubmed.ncbi.nlm.nih.gov/34829555/
- **[S5]** Long-term plasticity of astrocytic phenotypes and their control by neurons in health and disease - PMC — https://pmc.ncbi.nlm.nih.gov/articles/PMC10011399/?utm_source=openai
- **[S6]** Post-GWAS multiomic functional investigation of the TNIP1 locus in Alzheimer's disease highlights a potential role for GPX3 - PubMed — https://pubmed.ncbi.nlm.nih.gov/38809917/
- **[S7]** A novel microglial subset plays a key role in myelinogenesis in developing brain - PMC — https://pmc.ncbi.nlm.nih.gov/articles/PMC5686552/?utm_source=openai
- **[S8]** Post-GWAS multiomic functional investigation of the TNIP1 locus in Alzheimer's disease highlights a potential role for GPX3 - PubMed — https://pubmed.ncbi.nlm.nih.gov/38809917/
- **[S9]** Amyloid-β 25-35 Induces Neurotoxicity through the Up-Regulation of Astrocytic System Xc - PubMed — https://pubmed.ncbi.nlm.nih.gov/34829555/
- **[S10]** Long-term plasticity of astrocytic phenotypes and their control by neurons in health and disease - PMC — https://pmc.ncbi.nlm.nih.gov/articles/PMC10011399/?utm_source=openai
- **[S11]** GPX3 glutathione peroxidase 3 [Homo sapiens (human)] - Gene - NCBI — https://www.ncbi.nlm.nih.gov/gene/2878
- **[S12]** 
            Appraising the Role of Astrocytes as Suppliers of Neuronal Glutathione Precursors - PMC
         — https://pmc.ncbi.nlm.nih.gov/articles/PMC10179008/
- **[S13]** Post-GWAS multiomic functional investigation of the TNIP1 locus in Alzheimer's disease highlights a potential role for GPX3 - PubMed — https://pubmed.ncbi.nlm.nih.gov/38809917/
- **[S14]** Pharos : Target Details - GPX3 — https://pharos.nih.gov/targets/GPX3
