## Supplementary material for "Understanding Substance Dependence: What Differentiates Addictive from Non-Addictive Drugs?": Folder for supplementary files: non-addictive-genes-claims-first100-009-cfl1-gene-protein-has-been-associated-with-regulat-2f096329a597.md

# non-addictive-genes-claims-first100-009-2f096329a597

**Original claim:** CFL1 gene/protein has been associated with regulatory or homeostatic biological processes in neural systems.

## Claim

CFL1 gene/protein has been associated with regulatory or homeostatic biological processes in neural systems.

## Verdict

Verdict: Likely True

CFL1 (cofilin-1, non-muscle cofilin) is strongly associated with **regulatory** processes in neural systems, especially actin-dependent control of neural development, neuronal migration, dendritic spine remodeling, glutamatergic synapse function, and synaptic plasticity. Authoritative gene annotations for mouse Cfl1 place it in the nervous system, postsynaptic density, and glutamatergic synapse, and indicate roles in neural crest migration and neural fold formation. ([ncbi.nlm.nih.gov](https://www.ncbi.nlm.nih.gov/gene/12631)) [S1]

The weaker part of the claim is **homeostatic**: the literature I found links CFL1 much more clearly to structural/synaptic regulation and plasticity than to classic neural homeostasis such as firing-rate homeostasis or synaptic scaling. So the overall claim is better supported if read as “associated with regulatory processes in neural systems” than as “established mediator of neural homeostasis.” ([pubmed.ncbi.nlm.nih.gov](https://pubmed.ncbi.nlm.nih.gov/26037722/)) [S2]

## Confidence

High

## Supporting evidence

An authoritative NCBI/Alliance summary for mouse **Cfl1** states that it is involved in cytoskeleton organization and embryonic development, acts upstream of or within **neural crest cell migration** and **neural fold formation**, is active in the **glutamatergic synapse** and **postsynaptic density**, and is expressed in the **nervous system**. That alone places CFL1 squarely within neural regulatory biology. ([ncbi.nlm.nih.gov](https://www.ncbi.nlm.nih.gov/gene/12631)) [S3]

Developmental evidence is strong. In mouse embryos, loss of **Cfl1** disrupts neural plate morphogenesis and blocks neural tube closure. The defect is mechanistically linked to failure of apical actomyosin activation, indicating that cofilin-1 is required for the regulated cytoskeletal changes needed for neurulation. A related study likewise concluded that normal spinal neural tube closure requires ROCK/LIMK/cofilin-dependent control of actin turnover and actomyosin disassembly. ([pmc.ncbi.nlm.nih.gov](https://pmc.ncbi.nlm.nih.gov/articles/PMC4378245/)) [S4]

There is also direct evidence for a role in **neuronal migration**. In a mouse hippocampal study, deletion of the C-terminal region of Reelin reduced cofilin phosphorylation, impaired migration of late-born CA1 neurons, and this positioning defect was rescued by expression of pseudo-phosphorylated cofilin. That is strong evidence that regulated cofilin signaling participates in neural circuit assembly during development. ([pubmed.ncbi.nlm.nih.gov](https://pubmed.ncbi.nlm.nih.gov/36435394/)) [S5]

At mature excitatory synapses, CFL1/cofilin-1 is repeatedly implicated in regulating dendritic spine structure and synaptic efficacy. A forebrain excitatory-neuron conditional **n-cofilin** knockout showed enlarged spines, increased axo-spinous synapse density, impaired late-phase LTP, abnormal LTD, altered AMPA receptor behavior, and impaired spatial and fear-associated learning. These are classic phenotypes of a molecule that regulates synaptic architecture and plasticity. ([pubmed.ncbi.nlm.nih.gov](https://pubmed.ncbi.nlm.nih.gov/20407421/)) [S6]

Cellular studies reinforce this. In hippocampal neurons, activity-dependent spine shrinkage and growth both required changes in cofilin regulation; the authors concluded cofilin is essential for normal spine maintenance and for activity-dependent structural plasticity. Another study showed that phosphorylated cofilin accumulation is a key local regulator of spine enlargement, while dephosphorylated cofilin diffusion promotes spine shrinkage and reduced PSD95. ([pubmed.ncbi.nlm.nih.gov](https://pubmed.ncbi.nlm.nih.gov/24740405/)) [S7]

Reviews of the field are consistent with these primary data. One review describes ADF/cofilin as a “crucial regulator of synapse physiology and behavior,” emphasizing roles in neurotransmitter release, AMPA receptor availability, LTP/LTD, and excitatory synapse function in the mammalian CNS. Another review states that ADF/cofilin regulate neuronal structure and function, including long-term potentiation and depression, through effects on dendritic spines and glutamate-receptor trafficking. ([pubmed.ncbi.nlm.nih.gov](https://pubmed.ncbi.nlm.nih.gov/26037722/)) [S8]

There is also some evidence in neural precursor systems. In neural stem cell and Neuro-2a experiments, cofilin-1 expression/phosphorylation changed during brain and NSC development, and manipulating cofilin-1 altered differentiation, neurite outgrowth, and neuronal maturation. This is not as definitive as the in vivo mouse work, but it supports a regulatory role in neural lineage biology. ([pubmed.ncbi.nlm.nih.gov](https://pubmed.ncbi.nlm.nih.gov/34655709/)) [S9]

## Contradicting or limiting evidence

The main limitation is that much of the evidence is from **mouse, rat, or cultured-cell systems**, not direct human physiology. For the narrow question of “association,” that is acceptable, but it weakens any claim that CFL1 is a proven human neural-homeostasis gene in the strict clinical sense. ([pubmed.ncbi.nlm.nih.gov](https://pubmed.ncbi.nlm.nih.gov/20407421/)) [S10]

Also, not every neural function depends uniquely on CFL1. A presynaptic study found that synapse physiology was preserved after genetic ablation of **ADF** alone, and noted that **n-cofilin mutant mice display postsynaptic plasticity defects but not presynaptic defects**; presynaptic abnormalities became clearer only when both ADF and cofilin family functions were disrupted, implying partial redundancy within the ADF/cofilin family. ([pubmed.ncbi.nlm.nih.gov](https://pubmed.ncbi.nlm.nih.gov/24770705/)) [S11]

Most importantly, the retrieved literature supports **regulation/plasticity** much more strongly than **canonical homeostasis**. The major reviews and primary studies focus on neurulation, migration, dendritic spine remodeling, LTP/LTD, receptor trafficking, and disease-related synaptic dysfunction. I did not find comparably strong direct evidence that CFL1 is a central mediator of classic firing-rate homeostasis or synaptic scaling in central neurons. That does not contradict the claim, but it limits how strongly the “homeostatic” part should be interpreted. ([pubmed.ncbi.nlm.nih.gov](https://pubmed.ncbi.nlm.nih.gov/26037722/)) [S12]

## Mechanistic / biological rationale

CFL1 encodes **cofilin-1**, an actin-binding protein that severs and depolymerizes actin filaments. Neural systems depend heavily on rapid, spatially precise actin remodeling: growth cones migrate, neural folds bend and close, dendritic spines enlarge or shrink, and glutamate receptors traffic into and out of synapses. A molecule that controls actin filament turnover is therefore biologically well positioned to regulate these neural processes. ([pubmed.ncbi.nlm.nih.gov](https://pubmed.ncbi.nlm.nih.gov/33282872/)) [S13]

Mechanistically, cofilin activity is switched by phosphorylation state and upstream pathways such as **LIMK** and **SSH/calcineurin**. In synapses, active vs. inactive cofilin changes where actin is severed or stabilized, which in turn affects spine morphology, AMPA receptor mobility, and the persistence of LTP or LTD. During development, the same core actin-remodeling logic helps explain why Cfl1 loss disrupts neural tube closure and why altered cofilin phosphorylation perturbs neuronal migration. ([pubmed.ncbi.nlm.nih.gov](https://pubmed.ncbi.nlm.nih.gov/33282872/)) [S14]

## Bottom line

The claim is **likely true**. CFL1/cofilin-1 is well supported as a regulator of neural-system biology, particularly **neural development**, **neuronal migration**, **dendritic spine dynamics**, **glutamatergic synapse organization**, and **synaptic plasticity**. The evidence is strongest in animal and cellular neuroscience, with authoritative database annotations aligning with the literature. The only reason not to score the claim higher is that the evidence more clearly supports **regulatory neural processes** than **classical neural homeostasis** specifically. ([ncbi.nlm.nih.gov](https://www.ncbi.nlm.nih.gov/gene/12631)) [S15]

## Cited sources

- **[S1]** Cfl1 cofilin 1, non-muscle [Mus musculus (house mouse)] - Gene - NCBI — https://www.ncbi.nlm.nih.gov/gene/12631
- **[S2]** ADF/cofilin: a crucial regulator of synapse physiology and behavior - PubMed — https://pubmed.ncbi.nlm.nih.gov/26037722/
- **[S3]** Cfl1 cofilin 1, non-muscle [Mus musculus (house mouse)] - Gene - NCBI — https://www.ncbi.nlm.nih.gov/gene/12631
- **[S4]** 
            Morphogenesis of the mouse neural plate depends on distinct roles of cofilin 1 in apical and basal epithelial domains - PMC
         — https://pmc.ncbi.nlm.nih.gov/articles/PMC4378245/
- **[S5]** Reelin regulates the migration of late-born hippocampal CA1 neurons via cofilin phosphorylation - PubMed — https://pubmed.ncbi.nlm.nih.gov/36435394/
- **[S6]** Learning, AMPA receptor mobility and synaptic plasticity depend on n-cofilin-mediated actin dynamics - PubMed — https://pubmed.ncbi.nlm.nih.gov/20407421/
- **[S7]** Activity-dependent dendritic spine shrinkage and growth involve downregulation of cofilin via distinct mechanisms - PubMed — https://pubmed.ncbi.nlm.nih.gov/24740405/
- **[S8]** ADF/cofilin: a crucial regulator of synapse physiology and behavior - PubMed — https://pubmed.ncbi.nlm.nih.gov/26037722/
- **[S9]** The effect of cofilin 1 expression and phosphorylation dynamics on cell fate determination and neuron maturation in neural stem cells - PubMed — https://pubmed.ncbi.nlm.nih.gov/34655709/
- **[S10]** Learning, AMPA receptor mobility and synaptic plasticity depend on n-cofilin-mediated actin dynamics - PubMed — https://pubmed.ncbi.nlm.nih.gov/20407421/
- **[S11]** ADF/Cofilin Controls Synaptic Actin Dynamics and Regulates Synaptic Vesicle Mobilization and Exocytosis - PubMed — https://pubmed.ncbi.nlm.nih.gov/24770705/
- **[S12]** ADF/cofilin: a crucial regulator of synapse physiology and behavior - PubMed — https://pubmed.ncbi.nlm.nih.gov/26037722/
- **[S13]** The Role of ADF/Cofilin in Synaptic Physiology and Alzheimer's Disease - PubMed — https://pubmed.ncbi.nlm.nih.gov/33282872/
- **[S14]** The Role of ADF/Cofilin in Synaptic Physiology and Alzheimer's Disease - PubMed — https://pubmed.ncbi.nlm.nih.gov/33282872/
- **[S15]** Cfl1 cofilin 1, non-muscle [Mus musculus (house mouse)] - Gene - NCBI — https://www.ncbi.nlm.nih.gov/gene/12631
