## Supplementary material for "Understanding Substance Dependence: What Differentiates Addictive from Non-Addictive Drugs?": Folder for supplementary files: non-addictive-genes-claims-first100-010-gadd45b-gene-protein-has-been-associated-with-regu-ca90ef3e7650.md

# non-addictive-genes-claims-first100-010-ca90ef3e7650

**Original claim:** GADD45B gene/protein has been associated with regulatory or homeostatic biological processes in neural systems.

## Claim

GADD45B gene/protein has been associated with regulatory or homeostatic biological processes in neural systems.

## Verdict

Verdict: Highly Supported

## Confidence

High

## Supporting evidence

A substantial preclinical literature links **GADD45B/Gadd45b** to **regulatory processes in neural systems**, especially activity-dependent epigenetic regulation, neural plasticity, neurogenesis, and injury responses. A recent review focused on neuronal Gadd45 proteins summarizes that Gadd45β is induced by neuronal activity and regulates DNA methylation/gene expression programs involved in memory formation, visual cortex plasticity, and adult neurogenesis. The same review notes that Gadd45β is expressed across brain regions and that both baseline and activity-induced pools contribute to neuronal function. ([pmc.ncbi.nlm.nih.gov](https://pmc.ncbi.nlm.nih.gov/articles/PMC9846468/)) [S1]

One of the strongest lines of evidence comes from adult hippocampus studies. In a landmark mouse study, Gadd45b was identified as an **activity-induced immediate early gene** in mature hippocampal neurons. Loss of Gadd45b impaired activity-dependent DNA demethylation of promoters for neurotrophic genes including **Bdnf** and **Fgf1**, reduced progenitor proliferation, and impaired dendritic growth of newborn neurons, supporting a role in regulating adult neurogenesis. Electroconvulsive stimulation/exercise-related work extended this by showing that transient Gadd45b induction can produce longer-lasting changes in hippocampal neurogenesis via neurotrophic gene regulation. ([pmc.ncbi.nlm.nih.gov](https://pmc.ncbi.nlm.nih.gov/articles/PMC2726986/)) [S2]

There is also direct evidence tying Gadd45b to **learning-related transcriptional regulation**. In hippocampal fear-conditioning experiments, learning increased Gadd45β expression in CA1 together with **BDNF promoter demethylation**; both effects were blocked when NF-κB/c-Rel signaling was inhibited or genetically disrupted. Independent work found that Gadd45b is rapidly induced after contextual fear conditioning and that knockout mice show impaired long-term hippocampus-dependent memory, consistent with a regulatory role during memory consolidation. ([pubmed.ncbi.nlm.nih.gov](https://pubmed.ncbi.nlm.nih.gov/26441517/)) [S3]

Beyond hippocampus, Gadd45b has been linked to **visual cortical critical-period plasticity**. In juvenile and adult mouse visual cortex, OTX2 regulates Gadd45b/g transcription, and viral Gadd45b expression in adult visual cortex was sufficient to induce ocular-dominance plasticity with accompanying epigenetic changes in plasticity-gene promoters. That is strong evidence that Gadd45b participates in regulatory programs controlling circuit plasticity in a defined neural system. ([pubmed.ncbi.nlm.nih.gov](https://pubmed.ncbi.nlm.nih.gov/29771284/)) [S4]

Gadd45b is also associated with **dopamine-dependent transcriptional regulation** in striatal reward circuitry. In rat striatal neurons, knockdown of Gadd45b blocked many DRD1-responsive transcriptional changes and disrupted dopamine-driven DNA methylation changes. The authors concluded that Gadd45b helps translate neuronal activity into programmed transcriptional and epigenetic responses, extending its regulatory role beyond hippocampal models. ([pmc.ncbi.nlm.nih.gov](https://pmc.ncbi.nlm.nih.gov/articles/PMC8027017/)) [S5]

Evidence also connects Gadd45b to broader neural maintenance/protective processes. In global ischemia models, Gadd45b increased in vulnerable hippocampal CA1 neurons, and knockdown accelerated neuronal death and mitochondrial dysfunction; the study proposed a protective mechanism involving BDNF promoter demethylation. Earlier retinal work likewise found Gadd45b protects retinal ganglion cells from oxidative stress, TNF-α toxicity, and glutamate excitotoxicity. These findings support association with neural stress-response and survival pathways, which are plausibly homeostasis-related even if not always labeled that way. ([pubmed.ncbi.nlm.nih.gov](https://pubmed.ncbi.nlm.nih.gov/30832463/)) [S6]

Additional animal studies broaden the picture: Gadd45b knockdown in neonatal amygdala altered juvenile social behavior, changed expression of genes such as **MeCP2, Reelin, BDNF**, and modified local cytokine expression; after experimental stroke, Gadd45b promoted axonal plasticity and functional recovery. Together, these findings repeatedly place Gadd45b in neural regulatory programs rather than purely generic cell-stress biology. ([pubmed.ncbi.nlm.nih.gov](https://pubmed.ncbi.nlm.nih.gov/25728234/)) [S7]

## Contradicting or limiting evidence

The main limitation is that the literature is **not fully consistent about the direction of effect**. One mouse study reported that Gadd45b knockout causes **deficits** in long-term hippocampus-dependent memory, whereas another found **enhanced persistent memory and facilitated hippocampal LTP** in knockout mice. A later review explicitly highlights these apparently contradictory findings and suggests they may reflect region-, cell-type-, developmental-, or baseline-vs-induced-pool differences. So the association with neural regulation is strong, but the exact functional consequence of increasing vs decreasing Gadd45b is context dependent. ([pmc.ncbi.nlm.nih.gov](https://pmc.ncbi.nlm.nih.gov/articles/PMC3407936/)) [S8]

A second limitation is that most evidence is from **rodents, injury models, or cultured neurons**. Direct human evidence is much thinner. Human/postmortem work in major psychosis found altered GADD45B expression/binding at the **BDNF** promoter and altered methylation marks, which is compatible with a neural regulatory role, but this is associative and disease-contextual rather than definitive proof of normal physiological function in humans. ([pmc.ncbi.nlm.nih.gov](https://pmc.ncbi.nlm.nih.gov/articles/PMC3242315/?utm_source=openai)) [S9]

A third limitation is semantic: the claim includes **“homeostatic”** processes. I found strong direct evidence for **regulatory** roles in neural systems, but less direct evidence for classic **neural homeostasis** in the narrow sense of set-point stabilization or canonical homeostatic synaptic plasticity. Most papers study activity-dependent epigenetic regulation, neurogenesis, memory, stress susceptibility, or neuroprotection instead. So the claim is best supported through the “regulatory” half of the wording, with “homeostatic” being a more cautious inference. ([pmc.ncbi.nlm.nih.gov](https://pmc.ncbi.nlm.nih.gov/articles/PMC9846468/)) [S10]

## Mechanistic / biological rationale

GADD45B is a member of the GADD45 stress-response family. The NCBI gene summary describes the family as responsive to environmental stress and involved in growth/apoptosis regulation and MAPK-related signaling. In the nervous system, however, the best-supported mechanism is not generic cell-cycle control but **activity-coupled epigenetic regulation**. ([ncbi.nlm.nih.gov](https://www.ncbi.nlm.nih.gov/gene/4616?utm_source=openai)) [S11]

Mechanistically, neuronal activity can induce Gadd45β through pathways involving **nuclear calcium, glutamatergic signaling, TRKB, DRD1, and c-Rel/NF-κB**. Once induced, Gadd45β is linked to promoter-specific DNA demethylation or methylation remodeling at plasticity-relevant genes such as **Bdnf, Fgf1, Reelin, Gad67**, and dopamine-responsive targets. In cortical neurons, Gadd45b knockdown abolishes NMDA-induced Bdnf IXa upregulation and prevents the corresponding reductions in 5mC/5hmC at that promoter, giving a plausible molecular explanation for effects on neurogenesis, memory-related transcription, and stress responses. ([pmc.ncbi.nlm.nih.gov](https://pmc.ncbi.nlm.nih.gov/articles/PMC9846468/)) [S12]

This mechanism fits a biologically coherent model: **transient neuronal activity** induces Gadd45b, which then helps convert short-lived signaling into **longer-lasting transcriptional states** that influence neural plasticity, progenitor proliferation, dendritic growth, circuit remodeling, and sometimes survival after injury. In ischemia and retinal injury models, Gadd45b also behaves as an anti-apoptotic/neuroprotective factor, suggesting a broader role in preserving neural integrity under stress. ([pmc.ncbi.nlm.nih.gov](https://pmc.ncbi.nlm.nih.gov/articles/PMC2726986/)) [S13]

## Bottom line

Yes. The claim is **well supported**, especially if interpreted as saying that **GADD45B is associated with regulatory neural processes**. Across multiple animal and cell-based studies, GADD45B/Gadd45b is linked to activity-dependent epigenetic regulation, adult neurogenesis, memory-related transcription, visual-cortex critical-period plasticity, dopaminergic transcriptional programs, and neuroprotection after injury. The main caveats are that the literature is mostly preclinical, some behavioral phenotypes are directionally inconsistent, and evidence for **classic neural homeostasis** is more indirect than evidence for **neural regulation/plasticity**. ([pmc.ncbi.nlm.nih.gov](https://pmc.ncbi.nlm.nih.gov/articles/PMC9846468/)) [S14]

## Cited sources

- **[S1]** 
            Regulation of neuronal plasticity by the DNA repair associated Gadd45 proteins - PMC
         — https://pmc.ncbi.nlm.nih.gov/articles/PMC9846468/
- **[S2]** 
            Neuronal Activity–Induced Gadd45b Promotes Epigenetic DNA Demethylation and Adult Neurogenesis - PMC
         — https://pmc.ncbi.nlm.nih.gov/articles/PMC2726986/
- **[S3]** NF-κB mediates Gadd45β expression and DNA demethylation in the hippocampus during fear memory formation - PubMed — https://pubmed.ncbi.nlm.nih.gov/26441517/
- **[S4]** Non-cell Autonomous OTX2 Homeoprotein Regulates Visual Cortex Plasticity Through Gadd45b/g - PubMed — https://pubmed.ncbi.nlm.nih.gov/29771284/
- **[S5]** 
            Regulation of dopamine-dependent transcription and cocaine action by Gadd45b - PMC
         — https://pmc.ncbi.nlm.nih.gov/articles/PMC8027017/
- **[S6]** Gadd45b Acts as Neuroprotective Effector in Global Ischemia-Induced Neuronal Death - PubMed — https://pubmed.ncbi.nlm.nih.gov/30832463/
- **[S7]** Gadd45b is an epigenetic regulator of juvenile social behavior and alters local pro-inflammatory cytokine production in the rodent amygdala - PubMed — https://pubmed.ncbi.nlm.nih.gov/25728234/
- **[S8]** 
            Gadd45b knockout mice exhibit selective deficits in hippocampus-dependent long-term memory - PMC
         — https://pmc.ncbi.nlm.nih.gov/articles/PMC3407936/
- **[S9]** Growth Arrest and DNA-Damage-Inducible, Beta (GADD45b)-Mediated DNA Demethylation in Major Psychosis - PMC — https://pmc.ncbi.nlm.nih.gov/articles/PMC3242315/?utm_source=openai
- **[S10]** 
            Regulation of neuronal plasticity by the DNA repair associated Gadd45 proteins - PMC
         — https://pmc.ncbi.nlm.nih.gov/articles/PMC9846468/
- **[S11]** GADD45B growth arrest and DNA damage inducible beta [Homo sapiens (human)] - Gene - NCBI — https://www.ncbi.nlm.nih.gov/gene/4616?utm_source=openai
- **[S12]** 
            Regulation of neuronal plasticity by the DNA repair associated Gadd45 proteins - PMC
         — https://pmc.ncbi.nlm.nih.gov/articles/PMC9846468/
- **[S13]** 
            Neuronal Activity–Induced Gadd45b Promotes Epigenetic DNA Demethylation and Adult Neurogenesis - PMC
         — https://pmc.ncbi.nlm.nih.gov/articles/PMC2726986/
- **[S14]** 
            Regulation of neuronal plasticity by the DNA repair associated Gadd45 proteins - PMC
         — https://pmc.ncbi.nlm.nih.gov/articles/PMC9846468/
