## Supplementary material for "Understanding Substance Dependence: What Differentiates Addictive from Non-Addictive Drugs?": Folder for supplementary files: non-addictive-genes-claims-first100-011-eef2-gene-protein-has-been-associated-with-regulat-e910419ad9db.md

# non-addictive-genes-claims-first100-011-e910419ad9db

**Original claim:** EEF2 gene/protein has been associated with regulatory or homeostatic biological processes in neural systems.

## Claim

EEF2 gene/protein has been associated with regulatory or homeostatic biological processes in neural systems. ([ncbi.nlm.nih.gov](https://www.ncbi.nlm.nih.gov/gene/1938?utm_source=openai)) [S1]

## Verdict

Verdict: Highly Supported

Across multiple neuronal cell, slice, and animal studies, eEF2 is not just a generic translation factor; its regulated activity is repeatedly implicated in activity-dependent control of dendritic protein synthesis, synaptic plasticity, excitation/inhibition balance, neurogenesis, and rapid homeostatic synaptic scaling. Direct human evidence is more limited and disease-oriented, but human EEF2 variants also produce neurodevelopmental and cerebellar phenotypes, supporting neural relevance. ([sciencedirect.com](https://www.sciencedirect.com/science/article/pii/S0896627307005752)) [S2]

## Confidence

High

The literature is internally consistent that the eEF2/eEF2K axis regulates neural translational responses and synaptic state. Confidence is slightly tempered only because the strongest “homeostatic” evidence is mostly from rodent and in vitro systems, and many experiments manipulate eEF2 phosphorylation via eEF2K rather than EEF2 alone. ([frontiersin.org](https://www.frontiersin.org/journals/cellular-neuroscience/articles/10.3389/fncel.2014.00035/full)) [S3]

## Supporting evidence

A foundational Neuron study identified eEF2 as a **biochemical sensor in dendrites** that couples miniature synaptic transmission to local protein synthesis. In cultured hippocampal neurons, spontaneous neurotransmission promoted eEF2 phosphorylation, and local control of eEF2 phosphorylation tuned dendritic translation. That is a direct regulatory role in neuronal information processing, not merely a housekeeping function. ([sciencedirect.com](https://www.sciencedirect.com/science/article/pii/S0896627307005752)) [S4]

A focused review of neuronal dendritic translation concluded that the eEF2K/eEF2 pathway is engaged by neuronal activity and glutamatergic signaling and is associated with multiple forms of synaptic plasticity. The review specifically notes well-established dendritic targets such as **BDNF, Arc, αCaMKII, and MAP1B**, molecules central to synaptic regulation and plasticity. ([frontiersin.org](https://www.frontiersin.org/journals/cellular-neuroscience/articles/10.3389/fncel.2014.00035/full)) [S5]

Primary studies support those review-level conclusions. In hippocampal neurons, mGluR activation requires eEF2K/eEF2 signaling for rapid Arc/Arg3.1 translation and mGluR-LTD, linking the pathway to regulated weakening of synapses. Another study showed that activity-dependent spine maturation and dendritic BDNF synthesis depend on eEF2-pathway signaling, tying eEF2 to structural synaptic regulation. ([neuroscience.jhu.edu](https://neuroscience.jhu.edu/files2/publications_Worley_P_park_Neuron.pdf)) [S6]

Evidence also extends to **homeostatic** processes specifically. A 2021 Cell Reports study found that **acute inhibition of eEF2K induces rapid synaptic scaling in the hippocampus**, and the authors explicitly frame eEF2K/eEF2 as regulators of protein translation and synaptic homeostasis. This is strong direct support for association with neural homeostatic plasticity. ([pubmed.ncbi.nlm.nih.gov](https://pubmed.ncbi.nlm.nih.gov/34731624/)) [S7]

Related work on ketamine’s mechanism further supports that interpretation: blockade of resting NMDAR activity deactivates eEF2K, reduces eEF2 phosphorylation, and de-suppresses BDNF translation; this mechanism has been tied to rapid antidepressant responses and, in later work, to homeostatic synaptic scaling. ([pubmed.ncbi.nlm.nih.gov](https://pubmed.ncbi.nlm.nih.gov/21677641/?utm_source=openai)) [S8]

The pathway also regulates broader neural stability variables. In mouse hippocampus, loss of eEF2K increased GABAergic transmission by upregulating Synapsin-2b and α5-containing GABA\(_A\) receptors, thereby altering the **excitation/inhibition balance** and seizure susceptibility. Because E/I balance is a core homeostatic property of neural circuits, this is important evidence for involvement in neural homeostasis. ([pubmed.ncbi.nlm.nih.gov](https://pubmed.ncbi.nlm.nih.gov/27005990/)) [S9]

In the adult dentate gyrus, physiological manipulations that enhance neurogenesis were reported to converge on the eEF2/eEF2K pathway. Both global and dentate-gyrus-targeted eEF2K knockout enhanced neurogenesis-related proteins, increased adult neurogenesis, and improved dentate-gyrus-dependent learning. This places the pathway in regulation of ongoing neural tissue maintenance and adaptive plasticity. ([pubmed.ncbi.nlm.nih.gov](https://pubmed.ncbi.nlm.nih.gov/32707059/)) [S10]

Direct perturbation of **Eef2 itself** also affects neural function. In Eef2 heterozygous mice, reduced eEF2 in excitatory neurons of the prefrontal cortex lowered protein synthesis, spine density, neuronal excitability, and AMPAR-mediated synaptic transmission, and impaired social novelty behavior. This shows the neural regulatory role is not limited to manipulating the upstream kinase. ([embopress.org](https://www.embopress.org/doi/10.15252/embr.202154543)) [S11]

Human genetics supports neural relevance as well. De novo EEF2 variants have been associated with a neurodevelopmental disorder featuring motor delay, macrocephaly/ventriculomegaly, and other brain abnormalities, and earlier work linked impaired EEF2 to spinocerebellar ataxia type 26. These findings do not prove “homeostasis” directly, but they are consistent with EEF2 being important for normal neural regulatory biology. ([pubmed.ncbi.nlm.nih.gov](https://pubmed.ncbi.nlm.nih.gov/33355653/)) [S12]

## Contradicting or limiting evidence

The main limitation is that much of the strongest mechanistic evidence concerns the **eEF2K/eEF2 phosphorylation pathway**, not EEF2 gene dosage or protein abundance by itself. In other words, the neural regulatory role is best established for the regulated state of eEF2 within its signaling axis. ([frontiersin.org](https://www.frontiersin.org/journals/cellular-neuroscience/articles/10.3389/fncel.2014.00035/full)) [S13]

Also, eEF2 is a ubiquitous translation elongation factor used in many tissues, so it is not a neuron-specific homeostasis protein. Its neural effects likely reflect specialized use of a general translational control mechanism rather than a dedicated neural-only homeostatic factor. ([ncbi.nlm.nih.gov](https://www.ncbi.nlm.nih.gov/gene/1938?utm_source=openai)) [S14]

Some mechanistic details remain context-dependent. The literature shows that eEF2 phosphorylation usually suppresses global elongation, yet in some neuronal paradigms it selectively increases translation of certain plasticity-related mRNAs such as Arc or BDNF. Reviews note that these effects are not fully understood and may differ by stimulus, compartment, or timescale. ([frontiersin.org](https://www.frontiersin.org/journals/cellular-neuroscience/articles/10.3389/fncel.2014.00035/full)) [S15]

Finally, direct human experimental evidence for homeostatic neural functions is sparse. Human data are mainly genetic associations and clinical phenotypes, whereas the strongest causal evidence comes from rodents, cultured neurons, and ex vivo slices. ([pubmed.ncbi.nlm.nih.gov](https://pubmed.ncbi.nlm.nih.gov/33355653/)) [S16]

## Mechanistic / biological rationale

EEF2 encodes eukaryotic elongation factor 2, the factor that drives GTP-dependent ribosomal translocation during translation elongation. In neurons, its activity is dynamically controlled by phosphorylation at Thr56 by eEF2K; phosphorylation inhibits eEF2 and slows elongation. ([ncbi.nlm.nih.gov](https://www.ncbi.nlm.nih.gov/gene/1938?utm_source=openai)) [S17]

Because neuronal signaling pathways strongly regulate intracellular calcium, mTOR, and AMPK, synaptic activity can rapidly modulate eEF2K and thereby eEF2. NMDAR and mGluR signaling alter eEF2 phosphorylation in dendrites, changing local protein synthesis near synapses. ([frontiersin.org](https://www.frontiersin.org/journals/cellular-neuroscience/articles/10.3389/fncel.2014.00035/full)) [S18]

This makes biological sense as a homeostatic/regulatory mechanism: changing eEF2 activity can suppress bulk translation while selectively favoring translation of specific synaptic effectors such as Arc, BDNF, αCaMKII, and MAP1B. Those downstream proteins control AMPAR trafficking, dendritic spine structure, synaptic efficacy, and circuit excitability. ([frontiersin.org](https://www.frontiersin.org/journals/cellular-neuroscience/articles/10.3389/fncel.2014.00035/full)) [S19]

At the systems level, that translational switch can contribute to rapid synaptic scaling, maintenance of excitation/inhibition balance, experience-dependent cortical plasticity during sleep, adult dentate-gyrus neurogenesis, and behavioral outputs such as learning, social novelty, and antidepressant-like responses. ([pubmed.ncbi.nlm.nih.gov](https://pubmed.ncbi.nlm.nih.gov/34731624/)) [S20]

## Bottom line

The claim is **well supported**. EEF2—especially as part of the regulated eEF2K/eEF2 signaling axis—has strong evidence linking it to **regulatory** neural processes and meaningful evidence linking it to **homeostatic** ones, including synaptic scaling and circuit E/I balance. The clearest caution is that the literature is stronger for **activity-dependent translational regulation in neurons** than for EEF2 as a standalone, neural-specific homeostasis gene. ([sciencedirect.com](https://www.sciencedirect.com/science/article/pii/S0896627307005752)) [S21]

## Cited sources

- **[S1]** EEF2 eukaryotic translation elongation factor 2 [Homo sapiens (human)] - Gene - NCBI — https://www.ncbi.nlm.nih.gov/gene/1938?utm_source=openai
- **[S2]** Postsynaptic Decoding of Neural Activity: eEF2 as a Biochemical Sensor Coupling Miniature Synaptic Transmission to Local Protein Synthesis - ScienceDirect — https://www.sciencedirect.com/science/article/pii/S0896627307005752
- **[S3]** Frontiers | Elongation factor-2 phosphorylation in dendrites and the regulation of dendritic mRNA translation in neurons — https://www.frontiersin.org/journals/cellular-neuroscience/articles/10.3389/fncel.2014.00035/full
- **[S4]** Postsynaptic Decoding of Neural Activity: eEF2 as a Biochemical Sensor Coupling Miniature Synaptic Transmission to Local Protein Synthesis - ScienceDirect — https://www.sciencedirect.com/science/article/pii/S0896627307005752
- **[S5]** Frontiers | Elongation factor-2 phosphorylation in dendrites and the regulation of dendritic mRNA translation in neurons — https://www.frontiersin.org/journals/cellular-neuroscience/articles/10.3389/fncel.2014.00035/full
- **[S6]** doi:10.1016/j.neuron.2008.05.023 — https://neuroscience.jhu.edu/files2/publications_Worley_P_park_Neuron.pdf
- **[S7]** Convergence of distinct signaling pathways on synaptic scaling to trigger rapid antidepressant action - PubMed — https://pubmed.ncbi.nlm.nih.gov/34731624/
- **[S8]** NMDA receptor blockade at rest triggers rapid behavioural antidepressant responses - PubMed — https://pubmed.ncbi.nlm.nih.gov/21677641/?utm_source=openai
- **[S9]** eEF2K/eEF2 Pathway Controls the Excitation/Inhibition Balance and Susceptibility to Epileptic Seizures - PubMed — https://pubmed.ncbi.nlm.nih.gov/27005990/
- **[S10]** eEF2/eEF2K Pathway in the Mature Dentate Gyrus Determines Neurogenesis Level and Cognition - PubMed — https://pubmed.ncbi.nlm.nih.gov/32707059/
- **[S11]** eEF2 in the prefrontal cortex promotes excitatory synaptic transmission and social novelty behavior | EMBO Reports — https://www.embopress.org/doi/10.15252/embr.202154543
- **[S12]** De Novo variants in EEF2 cause a neurodevelopmental disorder with benign external hydrocephalus - PubMed — https://pubmed.ncbi.nlm.nih.gov/33355653/
- **[S13]** Frontiers | Elongation factor-2 phosphorylation in dendrites and the regulation of dendritic mRNA translation in neurons — https://www.frontiersin.org/journals/cellular-neuroscience/articles/10.3389/fncel.2014.00035/full
- **[S14]** EEF2 eukaryotic translation elongation factor 2 [Homo sapiens (human)] - Gene - NCBI — https://www.ncbi.nlm.nih.gov/gene/1938?utm_source=openai
- **[S15]** Frontiers | Elongation factor-2 phosphorylation in dendrites and the regulation of dendritic mRNA translation in neurons — https://www.frontiersin.org/journals/cellular-neuroscience/articles/10.3389/fncel.2014.00035/full
- **[S16]** De Novo variants in EEF2 cause a neurodevelopmental disorder with benign external hydrocephalus - PubMed — https://pubmed.ncbi.nlm.nih.gov/33355653/
- **[S17]** EEF2 eukaryotic translation elongation factor 2 [Homo sapiens (human)] - Gene - NCBI — https://www.ncbi.nlm.nih.gov/gene/1938?utm_source=openai
- **[S18]** Frontiers | Elongation factor-2 phosphorylation in dendrites and the regulation of dendritic mRNA translation in neurons — https://www.frontiersin.org/journals/cellular-neuroscience/articles/10.3389/fncel.2014.00035/full
- **[S19]** Frontiers | Elongation factor-2 phosphorylation in dendrites and the regulation of dendritic mRNA translation in neurons — https://www.frontiersin.org/journals/cellular-neuroscience/articles/10.3389/fncel.2014.00035/full
- **[S20]** Convergence of distinct signaling pathways on synaptic scaling to trigger rapid antidepressant action - PubMed — https://pubmed.ncbi.nlm.nih.gov/34731624/
- **[S21]** Postsynaptic Decoding of Neural Activity: eEF2 as a Biochemical Sensor Coupling Miniature Synaptic Transmission to Local Protein Synthesis - ScienceDirect — https://www.sciencedirect.com/science/article/pii/S0896627307005752
