## Supplementary material for "Understanding Substance Dependence: What Differentiates Addictive from Non-Addictive Drugs?": Folder for supplementary files: non-addictive-genes-claims-first100-012-nr4a1-gene-protein-has-been-associated-with-regula-33a5cae05878.md

# non-addictive-genes-claims-first100-012-33a5cae05878

**Original claim:** NR4A1 gene/protein has been associated with regulatory or homeostatic biological processes in neural systems.

## Claim

NR4A1 gene/protein has been associated with regulatory or homeostatic biological processes in neural systems.

## Verdict

Verdict: Highly Supported

Across multiple lines of neuroscience literature, NR4A1/Nr4a1 is linked to **regulatory functions in neural systems**, including activity-dependent transcription, synaptic wiring, synaptic plasticity, mitochondrial–synaptic coupling, microglial activation thresholds, and compensatory/homeostatic responses to drug exposure and neural stress. The strongest causal evidence comes from **mouse and cell-based studies**; human evidence exists but is more **correlative** than interventional. ([ncbi.nlm.nih.gov](https://www.ncbi.nlm.nih.gov/gene/3164?utm_source=openai)) [S1]

## Confidence

High

The claim is broad and is supported by **multiple independent studies**, including a review, several mechanistic mouse studies in neurons and microglia, and some human correlative data. Confidence is high for the statement that NR4A1 is **associated with regulatory/homeostatic neural processes**, but lower for any stronger claim that it is always protective or uniformly homeostatic in every neural context. ([pmc.ncbi.nlm.nih.gov](https://pmc.ncbi.nlm.nih.gov/articles/PMC3078984/)) [S2]

## Supporting evidence

NR4A1 is a **nuclear transcription factor** in the NR4A orphan nuclear receptor family, and NR4A-family genes are described as **immediate-early genes** induced by diverse stimuli, including signaling pathways relevant to neural activity and stress. Review literature also notes hippocampal induction of Nr4a genes under memory-forming conditions, placing NR4A1 squarely in stimulus-responsive regulatory biology rather than a passive structural role. ([ncbi.nlm.nih.gov](https://www.ncbi.nlm.nih.gov/gene/3164?utm_source=openai)) [S3]

In **neurons**, recent work shows a direct regulatory role for Nr4a1 in circuit organization. In mouse forebrain inhibitory interneurons, Nr4a1 expression was transiently induced by sensory experience/associative learning and was required for normal learning. Loss of Nr4a1 caused cell-type-specific transcriptional switches in gene families involved in **surface adhesion, repulsion, synaptic differentiation, and interneuron wiring**, indicating a bona fide role in regulating neural circuit architecture. ([pubmed.ncbi.nlm.nih.gov](https://pubmed.ncbi.nlm.nih.gov/38754414/)) [S4]

A separate hippocampal study linked Nr4a1 to **excitatory synaptic function and cognition**. In mice, deleting Nr4a1 in CA1 pyramidal neurons impaired dendritic spine density, glutamate receptor expression, excitatory synaptic transmission, long-term potentiation, and cognitive performance; overexpressing Nr4a1 improved memory in aged animals. Mechanistically, Nr4a1 bound the **TrkB/Ntrk2 promoter** and enhanced its transcription, supporting a direct role in maintaining synaptic plasticity-related signaling. The same paper also reported that **NR4A1 declines with aging in the human hippocampus**, which is consistent with a role in neural maintenance, though still correlative in humans. ([pmc.ncbi.nlm.nih.gov](https://pmc.ncbi.nlm.nih.gov/articles/PMC11095431/)) [S5]

NR4A1 has also been implicated in **mitochondrial–synaptic homeostasis** under stress. In prefrontal cortex models, NR4A1 was reported to control the energetic competence of mitochondria relative to synapse number, effectively acting as a checkpoint that adjusts mitochondrial and synaptic responses to chronic stress. In that study, Nr4a1 knockdown or knockout protected pyramidal neurons from chronic-stress-induced adverse effects, while high transcriptionally active NR4A1 in human PFC correlated with synaptic loss and cognitive impairment. This supports a regulatory/homeostatic role, even though prolonged activation may become maladaptive. ([pubmed.ncbi.nlm.nih.gov](https://pubmed.ncbi.nlm.nih.gov/29295823/)) [S6]

Evidence is also strong in **microglia**, the CNS resident immune/homeostatic cells. A 2017 study identified Nr4a1 as a **“rheostat”** controlling the activation threshold and polarization of microglia. ATP-induced Nr4a1 expression in steady-state microglia helped maintain a **resting, noninflammatory phenotype**; microglia-specific or global deletion caused spontaneous overactivation, increased cytokine/NO production, and worse autoimmune CNS inflammation. That is close to a textbook example of a homeostatic regulator in a neural cell type. ([pubmed.ncbi.nlm.nih.gov](https://pubmed.ncbi.nlm.nih.gov/28411187/)) [S7]

Additional neuroimmune work is consistent with that picture. In CNS-recruited macrophages during experimental autoimmune encephalomyelitis, Nr4a1 limited neuroinflammation by restraining macrophage tyrosine hydroxylase expression and catecholamine production; Nr4a1-deficient mice developed accelerated, more severe disease. In ischemic stroke models, another study found microglial NR4A1 can act noncanonically as an **RNA-binding protein** that destabilizes **Tnf mRNA**, functioning as a post-transcriptional brake on inflammation and improving recovery. ([pmc.ncbi.nlm.nih.gov](https://pmc.ncbi.nlm.nih.gov/articles/PMC4833087/)) [S8]

The addiction literature provides direct support for the word **“homeostatic.”** In mouse reward circuitry during cocaine abstinence, investigators identified Nr4a1 as a regulator of **homeostatic target gene expression** across nucleus accumbens, ventral tegmental area, and prefrontal cortex. The paper explicitly states that Nr4a1 regulates genes involved in **neuronal homeostasis**, including **Cartpt**, and that increasing Nr4a1 reduced cocaine-reinforced behaviors. That does not prove NR4A1 is universally homeostatic in all neural settings, but it clearly supports an association with homeostatic processes in neural systems. ([nature.com](https://www.nature.com/articles/s41467-020-14331-y)) [S9]

## Contradicting or limiting evidence

The main limitation is **context dependence**. NR4A1 is not uniformly protective or homeostatic across all neural paradigms. For example, some ischemia studies reported the opposite of the anti-inflammatory microglial papers: NR4A1 deletion reduced infarct volume, neuron injury, M1 polarization, and cognitive deficits after ischemic stroke, implying that in those models NR4A1 promoted injury-associated neuroinflammation. Another cerebral ischemia-reperfusion study concluded that increased NR4A1 worsened injury by repressing protective mitophagy and promoting mitochondrial dysfunction and apoptosis. ([pubmed.ncbi.nlm.nih.gov](https://pubmed.ncbi.nlm.nih.gov/31445702/)) [S10]

Likewise, a preterm-birth/perinatal neuroinflammation model implicated Nr4a1 in the **fetal cerebral vasculature** as a mediator of inflammatory brain injury, again showing that NR4A1 can participate in damaging as well as stabilizing responses depending on cell type, developmental stage, and insult. ([pubmed.ncbi.nlm.nih.gov](https://pubmed.ncbi.nlm.nih.gov/31907354/)) [S11]

A second limitation is that **most mechanistic evidence is from rodents and cultured cells**. Human data in this area are mainly observational: e.g., associations of active NR4A1 with synaptic loss in postmortem PFC or age-related reduction in human hippocampus/peripheral blood. Those observations strengthen translational relevance, but they do not establish the full physiologic role of NR4A1 in the living human brain. ([pubmed.ncbi.nlm.nih.gov](https://pubmed.ncbi.nlm.nih.gov/29295823/)) [S12]

## Mechanistic / biological rationale

Biologically, the claim makes sense because NR4A1 is an **activity- and stress-responsive transcription factor**. NR4A-family genes are rapidly induced by upstream pathways such as cAMP/PKA, MAPK, CREB, and other stimulus-coupled cascades. That positioning allows NR4A1 to convert transient neural activity or stress signals into downstream gene-expression programs. ([pmc.ncbi.nlm.nih.gov](https://pmc.ncbi.nlm.nih.gov/articles/PMC3078984/)) [S13]

In neurons, those downstream programs appear to affect **synaptic differentiation, circuit wiring, spine structure, glutamate receptor expression, TrkB signaling, and memory-related plasticity**. In stressed cortex, NR4A1 also links **mitochondrial state to synapse number**, which is conceptually a homeostatic function because network stability depends on matching energetic supply to synaptic demand. ([pubmed.ncbi.nlm.nih.gov](https://pubmed.ncbi.nlm.nih.gov/38754414/)) [S14]

In glial cells, the mechanism is complementary: NR4A1 sets inflammatory thresholds. In microglia it helps maintain a **resting/noninflammatory state**, and in one stroke model it directly destabilized **Tnf mRNA**, acting as a molecular brake on excessive cytokine output. These are classic regulatory/homeostatic mechanisms in CNS innate immunity. ([pubmed.ncbi.nlm.nih.gov](https://pubmed.ncbi.nlm.nih.gov/28411187/)) [S15]

## Bottom line

Yes. The literature strongly supports that **NR4A1 is associated with regulatory—and in several cases explicitly homeostatic—processes in neural systems**. The evidence spans neuronal plasticity/circuit regulation, mitochondrial–synaptic coupling, microglial inflammatory set-point control, and homeostatic adaptations in reward circuitry. The main caveat is that NR4A1’s effects are **context-dependent**: in some injury models it appears protective, while in others it appears pathogenic. So the broad association is well supported, but any narrower claim that NR4A1 is uniformly “homeostatic” or always beneficial in the nervous system would be too strong. ([pubmed.ncbi.nlm.nih.gov](https://pubmed.ncbi.nlm.nih.gov/38754414/)) [S16]

## Cited sources

- **[S1]** NR4A1 nuclear receptor subfamily 4 group A member 1 [Homo sapiens (human)] - Gene - NCBI — https://www.ncbi.nlm.nih.gov/gene/3164?utm_source=openai
- **[S2]** 
            The role of NR4A transcription factors in memory formation - PMC
         — https://pmc.ncbi.nlm.nih.gov/articles/PMC3078984/
- **[S3]** NR4A1 nuclear receptor subfamily 4 group A member 1 [Homo sapiens (human)] - Gene - NCBI — https://www.ncbi.nlm.nih.gov/gene/3164?utm_source=openai
- **[S4]** Nr4a1 regulates cell-specific transcriptional programs in inhibitory GABAergic interneurons - PubMed — https://pubmed.ncbi.nlm.nih.gov/38754414/
- **[S5]** 
            Progressive reduction of nuclear receptor Nr4a1 mediates age‐dependent cognitive decline - PMC
         — https://pmc.ncbi.nlm.nih.gov/articles/PMC11095431/
- **[S6]** The Stress-Induced Transcription Factor NR4A1 Adjusts Mitochondrial Function and Synapse Number in Prefrontal Cortex - PubMed — https://pubmed.ncbi.nlm.nih.gov/29295823/
- **[S7]** The Nuclear Receptor Nr4a1 Acts as a Microglia Rheostat and Serves as a Therapeutic Target in Autoimmune-Driven Central Nervous System Inflammation - PubMed — https://pubmed.ncbi.nlm.nih.gov/28411187/
- **[S8]** 
            The orphan nuclear receptor Nr4a1 couples sympathetic and inflammatory cues in CNS-recruited macrophages to limit neuroinflammation - PMC
         — https://pmc.ncbi.nlm.nih.gov/articles/PMC4833087/
- **[S9]** Nr4a1 suppresses cocaine-induced behavior via epigenetic regulation of homeostatic target genes | Nature Communications — https://www.nature.com/articles/s41467-020-14331-y
- **[S10]** NR4A1 regulates cerebral ischemia-induced brain injury by regulating neuroinflammation through interaction with NF-κB/p65 - PubMed — https://pubmed.ncbi.nlm.nih.gov/31445702/
- **[S11]** The orphan nuclear receptor Nr4a1 mediates perinatal neuroinflammation in a murine model of preterm labor - PubMed — https://pubmed.ncbi.nlm.nih.gov/31907354/
- **[S12]** The Stress-Induced Transcription Factor NR4A1 Adjusts Mitochondrial Function and Synapse Number in Prefrontal Cortex - PubMed — https://pubmed.ncbi.nlm.nih.gov/29295823/
- **[S13]** 
            The role of NR4A transcription factors in memory formation - PMC
         — https://pmc.ncbi.nlm.nih.gov/articles/PMC3078984/
- **[S14]** Nr4a1 regulates cell-specific transcriptional programs in inhibitory GABAergic interneurons - PubMed — https://pubmed.ncbi.nlm.nih.gov/38754414/
- **[S15]** The Nuclear Receptor Nr4a1 Acts as a Microglia Rheostat and Serves as a Therapeutic Target in Autoimmune-Driven Central Nervous System Inflammation - PubMed — https://pubmed.ncbi.nlm.nih.gov/28411187/
- **[S16]** Nr4a1 regulates cell-specific transcriptional programs in inhibitory GABAergic interneurons - PubMed — https://pubmed.ncbi.nlm.nih.gov/38754414/
