## Supplementary material for "Understanding Substance Dependence: What Differentiates Addictive from Non-Addictive Drugs?": Folder for supplementary files: non-addictive-genes-claims-first100-013-s100a1-gene-protein-has-been-associated-with-regul-df048f2a3b8b.md

# non-addictive-genes-claims-first100-013-df048f2a3b8b

**Original claim:** S100A1 gene/protein has been associated with regulatory or homeostatic biological processes in neural systems.

## Claim

S100A1 gene/protein has been associated with regulatory or homeostatic biological processes in neural systems.

## Verdict

Verdict: Likely True

## Confidence

Medium

## Supporting evidence

S100A1 is clearly present in neural tissue, which is a prerequisite for any neural regulatory role. Human Protein Atlas data classify **S100A1 as group-enriched in brain, heart muscle, skeletal muscle, and tongue**, with brain RNA clustering most consistent with **astrocytes**, and curated human brain protein localization describing it as **“detected in glial cells, most abundant in astrocytes.”** Older experimental work also found S100A1 expression in **neuronal and glial cell lines**, and in mouse brain localized it to the **hippocampus, cerebral cortex, and amygdala**, with partial colocalization with GFAP-positive astrocytes. ([proteinatlas.org](https://www.proteinatlas.org/ENSG00000160678-S100A1/brain)) [S1]

There is also direct evidence that S100A1 participates in **neuronal differentiation-associated regulation**. In PC12 neuronal cells, S100A1 protein levels rose with differentiation, its subcellular distribution was linked to the **Golgi/secretory vesicle compartment**, and its binding-protein profile resembled that seen in brain tissue. A separate PC12 study reported that manipulating S100A1 altered **neurite organization, tubulin levels, and proliferation**, consistent with a role in regulating cytoskeletal and differentiation-related processes in neuron-like cells. ([pubmed.ncbi.nlm.nih.gov](https://pubmed.ncbi.nlm.nih.gov/7760053/)) [S2]

Evidence also connects S100A1 to **presynaptic regulatory machinery**. In brain extracts, synapsins Ia/Ib/IIa/IIb were identified as major **Ca²⁺/S100A1-interacting proteins**, and S100A1 bound synapsin IIa in a **Ca²⁺-dependent** manner with submicromolar affinity. The same study showed synapsins and S100A1 in the soma and neurites of PC12 cells, supporting the possibility of interaction in neurons in vivo. Because synapsins are established regulators of **synaptic vesicle dynamics and neurotransmitter release**, this is strong association-level evidence that S100A1 participates in neural regulatory biology. ([pmc.ncbi.nlm.nih.gov](https://pmc.ncbi.nlm.nih.gov/articles/PMC1220678/)) [S3]

More direct functional neural evidence comes from sympathetic neurons. In cultured superior cervical ganglion neurons, exogenous S100A1 was taken up into neurons, transported along axons, and then **enhanced CaV1 channel currents**, **amplified action-potential–evoked Ca²⁺ transients**, and **prolonged action potentials** in a **PKA-dependent** manner. The authors interpreted this as evidence that S100A1 can modulate neuronal excitability and calcium signaling. Even though this is peripheral rather than central nervous system work, it is still evidence for association with **homeostatic/regulatory neural processes**, especially calcium handling and excitability control. ([pmc.ncbi.nlm.nih.gov](https://pmc.ncbi.nlm.nih.gov/articles/PMC2770745/)) [S4]

S100A1 has also been tied to neural calcium regulation in disease-relevant brain tissue. In an Alzheimer’s disease mouse model, **genetic ablation of S100A1 reduced astrocytosis, microgliosis, and plaque burden**, and the study identified **S100A1–ryanodine receptor (RyR) complexes** in human and mouse AD brain, calling RyR a primary S100A1 target in brain. The authors concluded that S100A1 inhibition might help normalize **aberrant Ca²⁺ signaling** in AD. Reviews of AD-related S100 biology similarly describe S100A1 as linked to **tau/GSK3 pathways, APP-related biology, and Ca²⁺-dependent RyR interactions** in brain tissue. ([pubmed.ncbi.nlm.nih.gov](https://pubmed.ncbi.nlm.nih.gov/24931125/)) [S5]

At a broader mechanistic level, S100A1 is an **EF-hand Ca²⁺-binding protein** with experimentally measured Ca²⁺ and Zn²⁺ binding that would allow it to function as a **Ca²⁺/Zn²⁺ sensor** under physiological conditions. NCBI’s gene summary notes functions including **stimulation of Ca²⁺-induced Ca²⁺ release**, inhibition of microtubule assembly, and inhibition of PKC-mediated phosphorylation. Those biochemical properties fit well with the neural findings above, especially for roles in calcium-dependent regulation of excitability, vesicle trafficking, and cytoskeletal organization. ([pubmed.ncbi.nlm.nih.gov](https://pubmed.ncbi.nlm.nih.gov/34944467/)) [S6]

## Contradicting or limiting evidence

The main limitation is that **S100A1 is much better established in heart and skeletal muscle than in nervous system biology**. Many high-confidence mechanistic papers focus on cardiomyocytes or muscle excitation-contraction coupling, while the neural literature is comparatively small and scattered. The claim is therefore supported mainly at the level of **association and plausible function**, not by a deep, consensus body of neural-specific physiology studies. ([pubmed.ncbi.nlm.nih.gov](https://pubmed.ncbi.nlm.nih.gov/25157660/?utm_source=openai)) [S7]

Evidence for a major indispensable role in normal CNS homeostasis is also limited. A knockout-mouse study explicitly stated that S100A1’s CNS role was **“largely unknown,”** and found **normal brain morphology**, no obvious differences in astrocyte or neuron density/shape, and **no impairment in spatial learning and memory**. The behavioral phenotype was narrower: mainly **increased exploration and reduced anxiety-like behavior** in male mice. This argues against S100A1 being a universally critical master regulator of baseline neural homeostasis. ([pubmed.ncbi.nlm.nih.gov](https://pubmed.ncbi.nlm.nih.gov/17045663/)) [S8]

Several supportive studies are also **indirect**: PC12 cells are neuron-like rather than bona fide CNS neurons; synapsin binding demonstrates a plausible presynaptic regulatory interaction but not, by itself, a measured change in neurotransmitter release; and the Alzheimer’s work is largely in a **pathological context**, which may not reflect normal homeostatic function. Thus, the strongest statement supported by the literature is that S100A1 is **associated with** neural regulatory/homeostatic processes, not that it is a canonical or central controller of neural homeostasis. ([pubmed.ncbi.nlm.nih.gov](https://pubmed.ncbi.nlm.nih.gov/7760053/)) [S9]

## Mechanistic / biological rationale

Biologically, the claim is plausible. S100A1 is a small **Ca²⁺/Zn²⁺-binding sensor** that changes conformation upon ion binding and then engages target proteins. In neural cells, that kind of molecule would be well positioned to couple fluctuations in intracellular calcium to downstream changes in **channel activity, vesicle behavior, kinase signaling, and cytoskeletal organization**. ([pubmed.ncbi.nlm.nih.gov](https://pubmed.ncbi.nlm.nih.gov/34944467/)) [S10]

The most coherent neural mechanism emerging from the literature is **calcium-signal tuning**. In neurons, S100A1 has been linked to **CaV1 channel enhancement** and larger AP-evoked Ca²⁺ transients; in AD brain it forms complexes with **RyR**, a major intracellular Ca²⁺ release channel; and in neuron-like cells it localizes near **secretory vesicles** and binds **synapsins**, which organize vesicle pools and release competence. Taken together, this supports an inference that S100A1 can help shape **neuronal excitability and presynaptic calcium-dependent signaling**, both of which are homeostatic/regulatory processes. ([pmc.ncbi.nlm.nih.gov](https://pmc.ncbi.nlm.nih.gov/articles/PMC2770745/)) [S11]

Its glial expression also matters. Human brain atlas data place S100A1 prominently in **astrocytes/glial cells**, and older rodent work found expression in both **neurons and glia**. Because astrocytes are major regulators of extracellular ion balance, metabolic support, and synaptic environment, S100A1 expression in this compartment strengthens the argument that it is at least associated with neural homeostatic biology, even if direct astrocyte-specific functional studies remain limited. That last point is partly an inference from expression plus known astrocyte physiology, rather than a direct experimental demonstration for S100A1 itself. ([proteinatlas.org](https://www.proteinatlas.org/ENSG00000160678-S100A1/brain)) [S12]

## Bottom line

The literature supports saying that **S100A1 has been associated with regulatory/homeostatic processes in neural systems**, especially **calcium signaling, neuronal excitability, vesicle/cytoskeletal regulation, and glial/neuroinflammatory responses**. However, the evidence is **not as extensive or definitive** as it is for S100A1 in heart and muscle, and much of the neural evidence comes from **cell models, peripheral neurons, or neurodegenerative disease contexts** rather than normal CNS homeostasis. So the best overall judgment is **Likely True**, with **medium confidence**. ([proteinatlas.org](https://www.proteinatlas.org/ENSG00000160678-S100A1/brain)) [S13]

## Cited sources

- **[S1]** Brain tissue expression of S100A1 - Summary - The Human Protein Atlas — https://www.proteinatlas.org/ENSG00000160678-S100A1/brain
- **[S2]** Analysis of S100A1 expression during skeletal muscle and neuronal cell differentiation - PubMed — https://pubmed.ncbi.nlm.nih.gov/7760053/
- **[S3]** 
            Synapsins as major neuronal Ca2+/S100A1-interacting proteins - PMC
         — https://pmc.ncbi.nlm.nih.gov/articles/PMC1220678/
- **[S4]** 
            Augmentation of Cav1 channel current and action potential duration after uptake of S100A1 in sympathetic ganglion neurons - PMC
         — https://pmc.ncbi.nlm.nih.gov/articles/PMC2770745/
- **[S5]** The Ca2+ sensor S100A1 modulates neuroinflammation, histopathology and Akt activity in the PSAPP Alzheimer's disease mouse model - PubMed — https://pubmed.ncbi.nlm.nih.gov/24931125/
- **[S6]** Mechanism of Zn2+ and Ca2+ Binding to Human S100A1 - PubMed — https://pubmed.ncbi.nlm.nih.gov/34944467/
- **[S7]** S100A1: a major player in cardiovascular performance - PubMed — https://pubmed.ncbi.nlm.nih.gov/25157660/?utm_source=openai
- **[S8]** S100A1-deficient male mice exhibit increased exploratory activity and reduced anxiety-related responses - PubMed — https://pubmed.ncbi.nlm.nih.gov/17045663/
- **[S9]** Analysis of S100A1 expression during skeletal muscle and neuronal cell differentiation - PubMed — https://pubmed.ncbi.nlm.nih.gov/7760053/
- **[S10]** Mechanism of Zn2+ and Ca2+ Binding to Human S100A1 - PubMed — https://pubmed.ncbi.nlm.nih.gov/34944467/
- **[S11]** 
            Augmentation of Cav1 channel current and action potential duration after uptake of S100A1 in sympathetic ganglion neurons - PMC
         — https://pmc.ncbi.nlm.nih.gov/articles/PMC2770745/
- **[S12]** Brain tissue expression of S100A1 - Summary - The Human Protein Atlas — https://www.proteinatlas.org/ENSG00000160678-S100A1/brain
- **[S13]** Brain tissue expression of S100A1 - Summary - The Human Protein Atlas — https://www.proteinatlas.org/ENSG00000160678-S100A1/brain
