## Supplementary material for "Understanding Substance Dependence: What Differentiates Addictive from Non-Addictive Drugs?": Folder for supplementary files: non-addictive-genes-claims-first100-014-th-gene-protein-has-been-associated-with-regulator-cc0e005463a9.md

# non-addictive-genes-claims-first100-014-cc0e005463a9

**Original claim:** TH gene/protein has been associated with regulatory or homeostatic biological processes in neural systems.

## Claim

TH gene/protein has been associated with regulatory or homeostatic biological processes in neural systems.

## Verdict

Verdict: Likely True

## Confidence

High

## Supporting evidence

TH encodes tyrosine hydroxylase, the rate-limiting enzyme for catecholamine synthesis, converting tyrosine to L-DOPA and thereby controlling production of dopamine, norepinephrine, and epinephrine. NCBI states that TH “plays a key role in the physiology of adrenergic neurons,” and a major review describes TH as the rate-limiting step in catecholamine biosynthesis and a central point for regulation of dopamine synthesis. ([ncbi.nlm.nih.gov](https://www.ncbi.nlm.nih.gov/gene/7054)) [S1]

This is relevant to neural regulation and homeostasis because the catecholamines made downstream of TH are core neural regulators. MedlinePlus notes that dopamine helps the brain control movement and mood, while norepinephrine and epinephrine participate in the autonomic nervous system and regulate involuntary processes such as blood pressure and body temperature; TH deficiency reduces these catecholamines and disrupts normal nervous system function. ([medlineplus.gov](https://medlineplus.gov/genetics/condition/tyrosine-hydroxylase-deficiency)) [S2]

The evidence is not only that TH participates in neurotransmitter synthesis, but that TH itself is a regulated node in neural systems. A review of TH regulation reports that its N-terminal region controls enzyme activity, with multiple phosphorylation sites and dephosphorylation mechanisms, and that catecholamine neurotransmitters feed back to inhibit TH. Another review specifically on TH phosphorylation concludes that phosphorylation at Ser40 increases TH activity in vitro, in situ, and in vivo, and that hierarchical phosphorylation provides a form of control over TH activation. ([pubmed.ncbi.nlm.nih.gov](https://pubmed.ncbi.nlm.nih.gov/21176768/)) [S3]

More recent structural work strengthens the “homeostatic/regulatory” interpretation. A 2022 cryo-EM study found that dopamine binding inhibits TH, while Ser40 phosphorylation reactivates it, and explicitly describes these as key regulatory mechanisms for dopamine and TH homeostasis. That same paper notes that maintaining catecholamine homeostasis requires TH regulation at transcriptional, translational, and post-translational levels. ([pmc.ncbi.nlm.nih.gov](https://pmc.ncbi.nlm.nih.gov/articles/PMC8748767/)) [S4]

A systems/control paper also places TH within dopamine homeostasis: TH, DDC, and VMAT2 are discussed as a functional dopamine-synthesis module, and the paper describes TH as being inhibited by dopamine, tyrosine, and extracellular dopamine acting through D2 autoreceptor signaling that reduces PKA-dependent TH phosphorylation. That is a direct neural feedback mechanism linking TH to stabilization of dopaminergic tone. ([pubmed.ncbi.nlm.nih.gov](https://pubmed.ncbi.nlm.nih.gov/34884667/)) [S5]

Human genetic evidence also supports association with regulatory neural physiology. In a study of common TH variation, promoter polymorphisms were associated with catecholamine secretion, stress-induced blood pressure responses, hypertension, and broader autonomic function; the authors concluded that proximal TH promoter variation is functional and alters autonomic physiology. That is strong evidence that TH affects regulatory/homeostatic neural output in humans, especially through sympathetic/autonomic pathways. ([pubmed.ncbi.nlm.nih.gov](https://pubmed.ncbi.nlm.nih.gov/20571875/)) [S6]

There is also direct experimental evidence from neural circuits involved in homeostasis. In mice, arcuate nucleus TH neurons were shown to influence food intake and body weight; optogenetic stimulation increased feeding, food deprivation activated these cells, and the authors concluded that ARC TH cells play an influential positive role in energy homeostasis. ([pubmed.ncbi.nlm.nih.gov](https://pubmed.ncbi.nlm.nih.gov/27548245/)) [S7]

Similarly, a 2024 mouse study reported that signaling in hypothalamic TH neurons contributes to dysregulation of peripheral glucose homeostasis through neural autonomic and/or neuroendocrine mechanisms. A Nature review on appetite circuits also states that Th-expressing catecholaminergic neurons in the nucleus of the solitary tract have been implicated in promoting homeostatic appetite, including feeding responses to low blood glucose. ([pubmed.ncbi.nlm.nih.gov](https://pubmed.ncbi.nlm.nih.gov/38349753/)) [S8]

Finally, curated functional annotations on the NCBI Gene page support broader neural-process association. TH is annotated there as involved in catecholamine biosynthetic process, dopamine biosynthetic process, dopaminergic synaptic transmission, cognition, learning, and memory. While these annotations vary in evidence strength, they are consistent with TH being associated with regulatory neural functions rather than only a generic metabolic reaction. ([ncbi.nlm.nih.gov](https://www.ncbi.nlm.nih.gov/gene/7054)) [S9]

## Contradicting or limiting evidence

The main limitation is conceptual rather than contradictory: TH is best established as a biosynthetic enzyme, not as a master “homeostasis gene” in the same sense as canonical signaling receptors, ion channels, or transcriptional homeostatic regulators. Much of the strongest evidence concerns regulation of catecholamine synthesis and neural circuit function upstream of homeostatic outcomes, rather than TH being directly annotated as a generic neural homeostasis effector. ([pubmed.ncbi.nlm.nih.gov](https://pubmed.ncbi.nlm.nih.gov/21176768/)) [S10]

Also, some of the broader nervous-system associations on the NCBI/GO page, such as cognition, learning, memory, and dopaminergic synaptic transmission, include inference-based evidence codes rather than only direct human experimental data. Those annotations are useful but should be interpreted as supportive, not definitive proof of a primary TH-specific role in each listed process. ([ncbi.nlm.nih.gov](https://www.ncbi.nlm.nih.gov/gene/7054)) [S11]

In addition, some of the clearest “homeostasis” studies involve TH-expressing neuronal populations in mice, especially hypothalamic neurons regulating energy balance or glucose control. Those studies show a real association between TH-marked catecholaminergic circuits and homeostatic control, but they do not always isolate whether the phenotype depends specifically on TH enzymatic activity versus other properties of those neurons. ([pubmed.ncbi.nlm.nih.gov](https://pubmed.ncbi.nlm.nih.gov/27548245/)) [S12]

## Mechanistic / biological rationale

Mechanistically, the claim makes sense. TH sits at the rate-limiting step of catecholamine biosynthesis, so changing TH expression or activity can alter dopamine, norepinephrine, and epinephrine availability across central and peripheral neural systems. Because those transmitters regulate movement, reward, mood, arousal, autonomic output, cardiovascular responses, temperature regulation, feeding, and glucose control, TH is positioned upstream of many regulatory and homeostatic neural processes. ([medlineplus.gov](https://medlineplus.gov/genetics/condition/tyrosine-hydroxylase-deficiency)) [S13]

TH is also embedded in feedback control loops. Dopamine can inhibit TH directly, and extracellular dopamine can reduce TH activation through D2 autoreceptor-linked suppression of PKA signaling; conversely, phosphorylation, especially at Ser40, reactivates TH. This is a classic control architecture for maintaining transmitter levels within functional bounds, which is why recent literature explicitly discusses TH in the context of dopamine and catecholamine homeostasis. ([pubmed.ncbi.nlm.nih.gov](https://pubmed.ncbi.nlm.nih.gov/34884667/)) [S14]

At the neural-circuit level, TH-expressing catecholaminergic neurons are repeatedly implicated in circuits that sense internal state and adjust behavior or physiology accordingly, including appetite, body weight, and glucose regulation. That does not mean TH alone “causes” homeostasis, but it strongly supports association with homeostatic biological processes in neural systems. ([pubmed.ncbi.nlm.nih.gov](https://pubmed.ncbi.nlm.nih.gov/27548245/)) [S15]

## Bottom line

The claim is **likely true**. TH is not merely a generic metabolic enzyme; it is a tightly regulated, rate-limiting determinant of catecholamine synthesis in neural tissues, and human, mechanistic, and animal-circuit studies all connect it to neural regulatory functions and to homeostasis-related processes such as autonomic control, appetite, and glucose balance. The strongest evidence is for **regulation of catecholamine/dopamine synthesis and downstream neural homeostatic circuits**, while evidence for TH as a direct, standalone “homeostasis regulator” is somewhat more indirect. ([ncbi.nlm.nih.gov](https://www.ncbi.nlm.nih.gov/gene/7054)) [S16]

## Cited sources

- **[S1]** TH tyrosine hydroxylase [Homo sapiens (human)] - Gene - NCBI — https://www.ncbi.nlm.nih.gov/gene/7054
- **[S2]** Tyrosine hydroxylase deficiency: MedlinePlus Genetics — https://medlineplus.gov/genetics/condition/tyrosine-hydroxylase-deficiency
- **[S3]** Tyrosine hydroxylase and regulation of dopamine synthesis - PubMed — https://pubmed.ncbi.nlm.nih.gov/21176768/
- **[S4]** 
            Structural mechanism for tyrosine hydroxylase inhibition by dopamine and reactivation by Ser40 phosphorylation - PMC
         — https://pmc.ncbi.nlm.nih.gov/articles/PMC8748767/
- **[S5]** DOPA Homeostasis by Dopamine: A Control-Theoretic View - PubMed — https://pubmed.ncbi.nlm.nih.gov/34884667/
- **[S6]** Human tyrosine hydroxylase natural allelic variation: influence on autonomic function and hypertension - PubMed — https://pubmed.ncbi.nlm.nih.gov/20571875/
- **[S7]** Hypothalamic arcuate nucleus tyrosine hydroxylase neurons play orexigenic role in energy homeostasis - PubMed — https://pubmed.ncbi.nlm.nih.gov/27548245/
- **[S8]** (Pro)renin receptor signaling in hypothalamic tyrosine hydroxylase neurons is required for obesity-associated glucose metabolic impairment - PubMed — https://pubmed.ncbi.nlm.nih.gov/38349753/
- **[S9]** TH tyrosine hydroxylase [Homo sapiens (human)] - Gene - NCBI — https://www.ncbi.nlm.nih.gov/gene/7054
- **[S10]** Tyrosine hydroxylase and regulation of dopamine synthesis - PubMed — https://pubmed.ncbi.nlm.nih.gov/21176768/
- **[S11]** TH tyrosine hydroxylase [Homo sapiens (human)] - Gene - NCBI — https://www.ncbi.nlm.nih.gov/gene/7054
- **[S12]** Hypothalamic arcuate nucleus tyrosine hydroxylase neurons play orexigenic role in energy homeostasis - PubMed — https://pubmed.ncbi.nlm.nih.gov/27548245/
- **[S13]** Tyrosine hydroxylase deficiency: MedlinePlus Genetics — https://medlineplus.gov/genetics/condition/tyrosine-hydroxylase-deficiency
- **[S14]** DOPA Homeostasis by Dopamine: A Control-Theoretic View - PubMed — https://pubmed.ncbi.nlm.nih.gov/34884667/
- **[S15]** Hypothalamic arcuate nucleus tyrosine hydroxylase neurons play orexigenic role in energy homeostasis - PubMed — https://pubmed.ncbi.nlm.nih.gov/27548245/
- **[S16]** TH tyrosine hydroxylase [Homo sapiens (human)] - Gene - NCBI — https://www.ncbi.nlm.nih.gov/gene/7054
