## Supplementary material for "Understanding Substance Dependence: What Differentiates Addictive from Non-Addictive Drugs?": Folder for supplementary files: non-addictive-genes-claims-first100-015-c3-gene-protein-has-been-associated-with-regulator-3979c3eefe06.md

# non-addictive-genes-claims-first100-015-3979c3eefe06

**Original claim:** C3 gene/protein has been associated with regulatory or homeostatic biological processes in neural systems.

## Claim

C3 gene/protein has been associated with regulatory or homeostatic biological processes in neural systems.

## Verdict

Verdict: Highly Supported

Multiple independent studies support a role for complement C3 in **normal neural regulation**, especially in **developmental synapse elimination/circuit refinement**. Additional evidence links C3 or its cleavage products/receptors to **adult neurogenesis**, **neuronal migration**, and **developmental organization of retinal astrocyte/vascular networks**. The strongest causal data are from animal and cellular studies; human evidence is mainly transcriptomic and observational, but it is consistent with local complement activity in healthy brain tissue. ([pubmed.ncbi.nlm.nih.gov](https://pubmed.ncbi.nlm.nih.gov/18083105/)) [S1]

## Confidence

High

Confidence is high because the literature includes several well-cited primary studies with genetic loss-of-function or rescue experiments showing that C3-related pathways affect neural developmental and maintenance processes. The main limitation is that most causal evidence comes from mice rather than direct human intervention studies. ([pubmed.ncbi.nlm.nih.gov](https://pubmed.ncbi.nlm.nih.gov/18083105/)) [S2]

## Supporting evidence

The clearest evidence is for **developmental synaptic pruning**. In a seminal Cell paper, mice lacking **C3** showed sustained defects in CNS synapse elimination in the retinogeniculate system, indicating that C3 is required for normal anatomical refinement of neural circuits during development. A follow-up study showed that microglia engulf presynaptic inputs during the peak pruning period and that this engulfment depends on the **CR3/C3 pathway**; disrupting this signaling caused lasting wiring abnormalities. These are classic examples of a regulatory/homeostatic process in neural circuit maturation rather than a purely inflammatory role. ([pubmed.ncbi.nlm.nih.gov](https://pubmed.ncbi.nlm.nih.gov/18083105/)) [S3]

Evidence also supports a role for C3 in **ongoing physiological regulation of synapses in the healthy adult brain**. In adult C3-knockout mice, investigators found altered hippocampal synaptic function, enhanced spatial learning, and evidence that C3 negatively regulates the number of functional CA3-CA1 glutamatergic synapses. That does not prove C3 is always beneficial, but it does show that C3 is associated with normal regulation of synapse number/function outside overt disease states. ([sciencedirect.com](https://www.sciencedirect.com/science/article/pii/S0014488613003798)) [S4]

C3-related signaling has also been implicated in **adult neurogenesis**, another homeostatic neural process. In the EMBO Journal study by Rahpeymai et al., neural stem/progenitor populations expressed **C3aR**, and loss of C3aR signaling reduced basal neurogenesis in mice. The same study also linked complement signaling to ischemia-induced neurogenesis, suggesting that C3-derived signals can influence both steady-state and reparative neural progenitor behavior. Because C3a is generated from C3, these data support a functional connection between C3 biology and neural homeostasis, even if the most direct manipulation in that paper was at the receptor/signaling level. ([pmc.ncbi.nlm.nih.gov](https://pmc.ncbi.nlm.nih.gov/articles/PMC1422160/)) [S5]

Another line of evidence concerns **neuronal migration** in the developing cortex. Gorelik et al. showed that **C3, MASP1, and MASP2** are expressed in the embryonic mouse brain, and that C3 reduction impaired neuronal migration. Importantly, migration defects were rescued by molecular mimics of C3 cleavage products or by activating downstream receptors, which strengthens the causal inference that complement activity is part of normal developmental regulation rather than a bystander marker. ([pmc.ncbi.nlm.nih.gov](https://pmc.ncbi.nlm.nih.gov/articles/PMC5418580/)) [S6]

C3 has also been linked to **organization of non-neuronal elements that support neural homeostasis**, particularly in the retina. In a developmental retina study, loss of **C3** or **C3aR** impaired microglial phagocytic removal of astrocytes and altered astrocyte density/spatial patterning, which in turn affected vascular network development. That is a bona fide regulatory process within a neural tissue, expanding the evidence beyond synapses alone. ([pmc.ncbi.nlm.nih.gov](https://pmc.ncbi.nlm.nih.gov/articles/PMC10110418/)) [S7]

Human data are less causal but still supportive. A recent single-nucleus transcriptomic analysis of the healthy human hippocampus found that **C3** is predominantly expressed in **microglia**, with lower expression in astrocytes and other cell types, while **C3AR1** and several C3-fragment receptors are also mainly expressed in microglia. This pattern is consistent with a locally organized complement system operating in normal brain tissue rather than C3 being only a peripheral inflammatory molecule. Reviews of complement biology in the brain likewise describe complement as contributing to **tissue homeostasis**, including synaptic pruning and debris clearance in the healthy CNS. ([orca.cardiff.ac.uk](https://orca.cardiff.ac.uk/id/eprint/180307/1/Complement%20biosynthesis%20in%20human%20brain.pdf)) [S8]

## Contradicting or limiting evidence

The main limitation is that **most direct functional evidence is from rodents**, especially developmental mouse models. Human evidence largely consists of expression atlases, postmortem observations, and disease association studies rather than experiments proving that endogenous C3 regulates homeostatic neural processes in living humans. ([orca.cardiff.ac.uk](https://orca.cardiff.ac.uk/id/eprint/180307/1/Complement%20biosynthesis%20in%20human%20brain.pdf)) [S9]

Not every study shows a large baseline neural phenotype when C3-related signaling is disrupted. For example, one recent mouse study reported that **C3aR1 deficiency did not alter gross brain structure or functional connectivity across early life development**, suggesting that some complement-dependent roles may be subtle, region-specific, developmentally restricted, or buffered by compensatory pathways. That does not negate the pruning/migration/neurogenesis data, but it argues against overgeneralizing C3 as a universal master regulator of all neural homeostasis. ([pmc.ncbi.nlm.nih.gov](https://pmc.ncbi.nlm.nih.gov/articles/PMC12646079/)) [S10]

The literature also shows that C3 is **context-dependent**. In aging mice, C3 deficiency protected against age-related synapse loss, LTP decline, and cognitive impairment, implying that endogenous C3 can contribute to maladaptive synapse loss in later life. Conversely, the same study noted prior AD-model work in which C3 deficiency or inhibition increased amyloid deposition and neurodegeneration, indicating that C3 can also support clearance and tissue protection in some settings. Reviews likewise emphasize that complement-mediated pruning can be beneficial during development but harmful when excessive in the adult or injured brain. ([pmc.ncbi.nlm.nih.gov](https://pmc.ncbi.nlm.nih.gov/articles/PMC6605437/)) [S11]

## Mechanistic / biological rationale

The biology is plausible and fairly well worked out. **C3 is the central amplification node of the complement cascade**. When cleaved, it produces **C3b/iC3b**, which can opsonize synapses or cellular material and promote phagocytosis through microglial complement receptors such as **CR3**; it also produces **C3a**, an anaphylatoxin that signals through **C3aR**. In neural tissue, these routes provide mechanisms for selective synapse removal, glial activation, progenitor regulation, and other remodeling functions. ([pubmed.ncbi.nlm.nih.gov](https://pubmed.ncbi.nlm.nih.gov/33239010/)) [S12]

That framework fits the major experimental findings. In developmental pruning, C3-tagged synaptic elements are removed by microglia through CR3-dependent engulfment. In neurogenesis and migration studies, C3-derived signaling through C3aR was linked to progenitor behavior and neuroblast migration. In the retina, C3/C3aR signaling helped microglia eliminate excess astrocytes to shape tissue architecture. Together, these mechanisms are exactly the kind of localized regulatory processes that fall under neural homeostasis. ([pmc.ncbi.nlm.nih.gov](https://pmc.ncbi.nlm.nih.gov/articles/PMC3528177/)) [S13]

Human transcriptomic data strengthen the plausibility of these mechanisms in people: in healthy hippocampus, microglia are the dominant C3-expressing cells and also strongly express **C3AR1** and phagocytic receptors for C3 fragments, while other brain cell types express complement regulators. That architecture is consistent with a controlled local complement network capable of supporting surveillance, remodeling, and cleanup in the healthy brain. ([orca.cardiff.ac.uk](https://orca.cardiff.ac.uk/id/eprint/180307/1/Complement%20biosynthesis%20in%20human%20brain.pdf)) [S14]

## Bottom line

Yes—this claim is well supported.

The **best-established** neural regulatory/homeostatic role for C3 is in **developmental synapse pruning and circuit refinement**. There is also good evidence connecting C3-related signaling to **adult neurogenesis**, **neuronal migration**, and **developmental organization of glial/vascular networks in neural tissue**. The main caveat is that the strongest causal evidence comes from animal models, and C3’s effects are **highly context-dependent**: the same pathway can support normal homeostasis in one setting and drive maladaptive synapse loss or neuroinflammation in another. ([pubmed.ncbi.nlm.nih.gov](https://pubmed.ncbi.nlm.nih.gov/18083105/)) [S15]

## Cited sources

- **[S1]** The classical complement cascade mediates CNS synapse elimination - PubMed — https://pubmed.ncbi.nlm.nih.gov/18083105/
- **[S2]** The classical complement cascade mediates CNS synapse elimination - PubMed — https://pubmed.ncbi.nlm.nih.gov/18083105/
- **[S3]** The classical complement cascade mediates CNS synapse elimination - PubMed — https://pubmed.ncbi.nlm.nih.gov/18083105/
- **[S4]** Altered cognitive performance and synaptic function in the hippocampus of mice lacking C3 - ScienceDirect — https://www.sciencedirect.com/science/article/pii/S0014488613003798
- **[S5]** 
            Complement: a novel factor in basal and ischemia-induced neurogenesis - PMC
         — https://pmc.ncbi.nlm.nih.gov/articles/PMC1422160/
- **[S6]** 
            Developmental activities of the complement pathway in migrating neurons - PMC
         — https://pmc.ncbi.nlm.nih.gov/articles/PMC5418580/
- **[S7]** 
            Microglia refine developing retinal astrocytic and vascular networks through the complement C3/C3aR axis - PMC
         — https://pmc.ncbi.nlm.nih.gov/articles/PMC10110418/
- **[S8]** Untitled source — https://orca.cardiff.ac.uk/id/eprint/180307/1/Complement%20biosynthesis%20in%20human%20brain.pdf
- **[S9]** Untitled source — https://orca.cardiff.ac.uk/id/eprint/180307/1/Complement%20biosynthesis%20in%20human%20brain.pdf
- **[S10]** 
            Complement receptor C3ar1 deficiency does not alter brain structure or functional connectivity across early life development - PMC
         — https://pmc.ncbi.nlm.nih.gov/articles/PMC12646079/
- **[S11]** 
            Complement C3-Deficient Mice Fail to Display Age-Related Hippocampal Decline - PMC
         — https://pmc.ncbi.nlm.nih.gov/articles/PMC6605437/
- **[S12]** The good, the bad, and the opportunities of the complement system in neurodegenerative disease - PubMed — https://pubmed.ncbi.nlm.nih.gov/33239010/
- **[S13]** 
            Microglia Sculpt Postnatal Neural Circuits in an Activity and Complement-Dependent Manner - PMC
         — https://pmc.ncbi.nlm.nih.gov/articles/PMC3528177/
- **[S14]** Untitled source — https://orca.cardiff.ac.uk/id/eprint/180307/1/Complement%20biosynthesis%20in%20human%20brain.pdf
- **[S15]** The classical complement cascade mediates CNS synapse elimination - PubMed — https://pubmed.ncbi.nlm.nih.gov/18083105/
