## Supplementary material for "Understanding Substance Dependence: What Differentiates Addictive from Non-Addictive Drugs?": Folder for supplementary files: non-addictive-genes-claims-first100-016-ntrk2-gene-protein-has-been-associated-with-regula-7d99b6d54902.md

# non-addictive-genes-claims-first100-016-7d99b6d54902

**Original claim:** NTRK2 gene/protein has been associated with regulatory or homeostatic biological processes in neural systems.

## Claim

NTRK2 gene/protein has been associated with regulatory or homeostatic biological processes in neural systems.

## Verdict

Verdict: Highly Supported

## Confidence

High

## Supporting evidence

NTRK2 encodes TrkB, the high-affinity receptor for brain-derived neurotrophic factor (BDNF), and authoritative gene resources annotate it to neural regulatory processes including cellular response to BDNF, central nervous system neuron development, learning, long-term synaptic potentiation, and BDNF-mediated trans-synaptic signaling. That already places NTRK2 squarely within neural regulatory biology rather than being a peripheral or incidental marker. ([ncbi.nlm.nih.gov](https://www.ncbi.nlm.nih.gov/gene/4915)) [S1]

The strongest general evidence comes from the neurobiology literature on BDNF-TrkB signaling. A major review describes BDNF-TrkB as regulating neural progenitor selection, dendritic differentiation, neuronal connectivity, protein transcription/translation/trafficking during synaptic development, and multiple forms of synaptic plasticity via MAPK, PI3K, and PLCγ signaling. A second review specifically emphasizes that neuronal activity modulates TrkB surface expression, synaptic localization, and endocytosis, i.e. NTRK2 is itself under regulated control and mediates activity-dependent regulation at synapses. ([pubmed.ncbi.nlm.nih.gov](https://pubmed.ncbi.nlm.nih.gov/20186705/)) [S2]

There is also direct evidence for **homeostatic** neural roles, not just broad “regulatory” roles. In cultured hippocampal neurons, activity deprivation reduced inhibitory synaptic strength, and exogenous BDNF reversed this effect through TrkB; the authors concluded that activity-dependent scaling of GABAergic synapse strength is modulated by BDNF/TrkB signaling. This is classic homeostatic plasticity biology. ([pubmed.ncbi.nlm.nih.gov](https://pubmed.ncbi.nlm.nih.gov/16330218/)) [S3]

A second homeostatic-plasticity study found that prolonged activity blockade in hippocampal neurons reduced ERK1/2 activity, then reduced TrkB activation, and linked this pathway to decreased K_v7.3 expression at the axon initial segment during homeostatic scaling of action-potential firing. In other words, NTRK2/TrkB is not merely correlated with plasticity; it participates in mechanisms that stabilize neuronal excitability after chronic perturbation. ([pubmed.ncbi.nlm.nih.gov](https://pubmed.ncbi.nlm.nih.gov/35966206/)) [S4]

In vivo evidence extends beyond synapses to organism-level neural homeostasis. In the dorsomedial hypothalamus, TrkB-expressing neurons were activated by fasting/refeeding, chemogenetic activation suppressed feeding, inhibition increased feeding when animals were physiologically sated, and adult Ntrk2 deletion in this region caused hyperphagia, reduced energy expenditure, and obesity. The paper explicitly concludes these neurons are necessary and sufficient to suppress appetite and maintain physiological satiation. That is a direct neural homeostatic function. ([pubmed.ncbi.nlm.nih.gov](https://pubmed.ncbi.nlm.nih.gov/30718415/)) [S5]

Developmental circuit studies also support a regulatory/homeostatic role. Conditional deletion of Ntrk2/TrkB from immature dentate granule cells disrupted their integration and maturation, altered the developmental GABA shift by reducing Nkcc1 expression, disturbed early synchronized network activity, and led to impaired adult hippocampal plasticity and cognition. This shows NTRK2 is involved in regulating circuit maturation and maintaining appropriate developmental signaling balance in neural systems. ([pubmed.ncbi.nlm.nih.gov](https://pubmed.ncbi.nlm.nih.gov/32361506/?utm_source=openai)) [S6]

Human data are also consistent. A 2025 cohort study of 44 affected individuals concluded that pathogenic NTRK2 variants cause two neurodevelopmental syndromes: one associated with developmental delay, epilepsy, visual impairment, and feeding difficulties, and another loss-of-function group associated with developmental delay, obesity, and hyperphagia. Separately, functional characterization of an obesity-associated Y722C NTRK2 variant showed impaired MAPK/PLCγ/Akt signaling plus reduced BDNF-induced neurite outgrowth and cell survival. These observations do not prove every homeostatic role in humans directly, but they strongly support that normal NTRK2 function is required for regulated neural development and neuroendocrine control. ([pubmed.ncbi.nlm.nih.gov](https://pubmed.ncbi.nlm.nih.gov/39540377/)) [S7]

Additional human translational evidence comes from a study showing rare human TrkB variants impaired hippocampal synaptogenesis and were associated with neurobehavioral abnormalities; the paper summarizes TrkB as regulating neuronal development, synapse formation, and plasticity. ([pubmed.ncbi.nlm.nih.gov](https://pubmed.ncbi.nlm.nih.gov/32493978/?utm_source=openai)) [S8]

## Contradicting or limiting evidence

The main limitation is that the word **homeostatic** is narrower than **regulatory**. For NTRK2, broad neural regulatory roles are extremely well established, but direct evidence for “homeostasis” is strongest in selected contexts such as hypothalamic energy balance and homeostatic synaptic/intrinsic plasticity, rather than across every neural subsystem. ([pubmed.ncbi.nlm.nih.gov](https://pubmed.ncbi.nlm.nih.gov/30718415/)) [S9]

Also, much of the most mechanistically precise evidence comes from rodent or cultured-neuron experiments rather than direct interventional human studies. Human evidence is robust for disease association and functional variant effects, but usually indirect with respect to specific homeostatic mechanisms. ([pubmed.ncbi.nlm.nih.gov](https://pubmed.ncbi.nlm.nih.gov/39540377/)) [S10]

Finally, NTRK2 does not appear to mediate every hypothesized neural homeostatic process. In one mouse chemical-genetic study, acute NtrkB inhibition altered REM sleep and state transitions but had **no effect on sleep homeostasis**, suggesting that NTRK2’s homeostatic importance is domain-specific rather than universal across all neural regulation. ([pubmed.ncbi.nlm.nih.gov](https://pubmed.ncbi.nlm.nih.gov/34537852/?utm_source=openai)) [S11]

## Mechanistic / biological rationale

The mechanistic case is straightforward. TrkB is a membrane receptor that, upon BDNF binding, activates MAPK, PI3K-Akt, and PLCγ pathways. Those cascades regulate transcription, translation, protein trafficking, calcium signaling, dendritic growth, synaptic maturation, and synaptic strength. Reviews further show that neuronal activity changes TrkB surface availability, localization to synaptic/lipid-raft compartments, and receptor endocytosis, providing a plausible mechanism for adaptive regulation in active neural circuits. ([pubmed.ncbi.nlm.nih.gov](https://pubmed.ncbi.nlm.nih.gov/20186705/)) [S12]

For bona fide homeostatic plasticity, the logic is also biologically coherent: chronic activity reduction lowers ERK and TrkB signaling, which then changes expression/localization of excitability regulators such as K_v7.3 at the axon initial segment, helping reset firing properties. At inhibitory synapses, BDNF/TrkB can counter activity-deprivation-induced weakening, consistent with a role in stabilizing network function. ([pubmed.ncbi.nlm.nih.gov](https://pubmed.ncbi.nlm.nih.gov/35966206/)) [S13]

At the systems level, TrkB-expressing hypothalamic neurons respond to fasting and refeeding and regulate food intake and energy expenditure, providing a neural substrate for physiologic homeostasis. In development, Ntrk2 helps coordinate the timing of GABAergic maturation and early network synchronization, which is another form of regulatory stabilization during circuit assembly. ([pubmed.ncbi.nlm.nih.gov](https://pubmed.ncbi.nlm.nih.gov/30718415/)) [S14]

## Bottom line

Yes. The claim is well supported. NTRK2/TrkB is extensively associated with **regulatory** processes in neural systems—development, synaptic signaling, plasticity, and circuit maturation—and there is also direct evidence for **homeostatic** roles, especially in hypothalamic energy balance and homeostatic synaptic/excitability plasticity. The main caveat is that “homeostatic” evidence is strongest in specific neural contexts and is derived largely from animal/cellular work, while human evidence is more often genetic and phenotypic than mechanistic. ([ncbi.nlm.nih.gov](https://www.ncbi.nlm.nih.gov/gene/4915)) [S15]

## Cited sources

- **[S1]** NTRK2 neurotrophic receptor tyrosine kinase 2 [Homo sapiens (human)] - Gene - NCBI — https://www.ncbi.nlm.nih.gov/gene/4915
- **[S2]** Postsynaptic BDNF-TrkB signaling in synapse maturation, plasticity, and disease - PubMed — https://pubmed.ncbi.nlm.nih.gov/20186705/
- **[S3]** Activity-dependent scaling of GABAergic synapse strength is regulated by brain-derived neurotrophic factor - PubMed — https://pubmed.ncbi.nlm.nih.gov/16330218/
- **[S4]** Homeostatic regulation of extracellular signal-regulated kinase 1/2 activity and axonal Kv7.3 expression by prolonged blockade of hippocampal neuronal activity - PubMed — https://pubmed.ncbi.nlm.nih.gov/35966206/
- **[S5]** TrkB-expressing neurons in the dorsomedial hypothalamus are necessary and sufficient to suppress homeostatic feeding - PubMed — https://pubmed.ncbi.nlm.nih.gov/30718415/
- **[S6]** Immature Dentate Granule Cells Require Ntrk2/Trkb for the Formation of Functional Hippocampal Circuitry - PubMed — https://pubmed.ncbi.nlm.nih.gov/32361506/?utm_source=openai
- **[S7]** Upregulation versus loss of function of NTRK2 in 44 affected individuals leads to 2 distinct neurodevelopmental disorders - PubMed — https://pubmed.ncbi.nlm.nih.gov/39540377/
- **[S8]** Human BDNF/TrkB variants impair hippocampal synaptogenesis and associate with neurobehavioural abnormalities - PubMed — https://pubmed.ncbi.nlm.nih.gov/32493978/?utm_source=openai
- **[S9]** TrkB-expressing neurons in the dorsomedial hypothalamus are necessary and sufficient to suppress homeostatic feeding - PubMed — https://pubmed.ncbi.nlm.nih.gov/30718415/
- **[S10]** Upregulation versus loss of function of NTRK2 in 44 affected individuals leads to 2 distinct neurodevelopmental disorders - PubMed — https://pubmed.ncbi.nlm.nih.gov/39540377/
- **[S11]** A chemical-genetic investigation of BDNF-NtrkB signaling in mammalian sleep - PubMed — https://pubmed.ncbi.nlm.nih.gov/34537852/?utm_source=openai
- **[S12]** Postsynaptic BDNF-TrkB signaling in synapse maturation, plasticity, and disease - PubMed — https://pubmed.ncbi.nlm.nih.gov/20186705/
- **[S13]** Homeostatic regulation of extracellular signal-regulated kinase 1/2 activity and axonal Kv7.3 expression by prolonged blockade of hippocampal neuronal activity - PubMed — https://pubmed.ncbi.nlm.nih.gov/35966206/
- **[S14]** TrkB-expressing neurons in the dorsomedial hypothalamus are necessary and sufficient to suppress homeostatic feeding - PubMed — https://pubmed.ncbi.nlm.nih.gov/30718415/
- **[S15]** NTRK2 neurotrophic receptor tyrosine kinase 2 [Homo sapiens (human)] - Gene - NCBI — https://www.ncbi.nlm.nih.gov/gene/4915
