## Supplementary material for "Understanding Substance Dependence: What Differentiates Addictive from Non-Addictive Drugs?": Folder for supplementary files: non-addictive-genes-claims-first100-017-mat2a-gene-protein-has-been-associated-with-regula-9c06a71928d8.md

# non-addictive-genes-claims-first100-017-9c06a71928d8

**Original claim:** MAT2A gene/protein has been associated with regulatory or homeostatic biological processes in neural systems.

## Claim

MAT2A gene/protein has been associated with regulatory or homeostatic biological processes in neural systems.

## Verdict

Verdict: Likely True

MAT2A is not a classic “neural homeostasis gene” in the same sense as ion channels or neurotransmitter transporters, but the literature does support its association with regulatory processes in neural systems. The strongest evidence links MAT2A to astrocyte inflammatory-state regulation in CNS disease and to hippocampal synaptic plasticity, learning, and memory in mouse models. Its role is biologically plausible because MAT2A makes S-adenosylmethionine (SAM), the main methyl donor for cellular methylation reactions. ([nature.com](https://www.nature.com/articles/s41586-020-1999-0)) [S1]

## Confidence

Medium

Confidence is **Medium** because multiple peer-reviewed studies support neural relevance, including a high-impact Nature paper in astrocytes and targeted hippocampal studies in mice, and MAT2A is expressed in brain cells with astrocyte-enhanced single-nucleus expression in Human Protein Atlas data. However, most evidence is preclinical, disease-contextual, or mechanistic rather than large-scale human clinical evidence for a core homeostatic role under normal physiology. ([nature.com](https://www.nature.com/articles/s41586-020-1999-0)) [S2]

## Supporting evidence

**1) MAT2A has the right molecular function to regulate neural biology.**  
NCBI describes MAT2A as the enzyme that synthesizes S-adenosylmethionine from methionine and ATP, and SAM is “the key methyl donor in cellular processes.” Gene Ontology annotations on the same page place MAT2A in the SAM biosynthetic process and one-carbon metabolism. A recent review likewise states that SAM is the primary methyl donor for numerous methylation reactions and that mammalian MAT2A expression is tightly controlled as part of SAM homeostasis. ([ncbi.nlm.nih.gov](https://www.ncbi.nlm.nih.gov/gene/4144)) [S3]

**2) MAT2A is expressed in brain-relevant cell types, especially astrocytes.**  
Human Protein Atlas single-nucleus brain data classify MAT2A as “cell type enhanced (Astrocyte),” indicating that its expression is not merely absent from neural tissue but is enriched in a major homeostatic glial population. This does not by itself prove function, but it strengthens the plausibility of a neural regulatory role. ([proteinatlas.org](https://www.proteinatlas.org/ENSG00000168906-MAT2A/single%2Bcell)) [S4]

**3) Strongest direct neural regulatory evidence: astrocyte state regulation in CNS inflammation.**  
In a Nature study of multiple sclerosis and experimental autoimmune encephalomyelitis (EAE), astrocytes with reduced NRF2 and increased MAFG showed a program in which MAFG cooperated with MAT2α to promote DNA methylation and repress antioxidant and anti-inflammatory transcriptional programs. The paper further reported that GM-CSF signaling in astrocytes drives MAFG and MAT2α expression and pro-inflammatory transcriptional modules that contribute to CNS pathology. This is direct evidence that MAT2A participates in regulation of glial transcriptional state in the CNS. ([nature.com](https://www.nature.com/articles/s41586-020-1999-0)) [S5]

**4) Strongest direct neural plasticity evidence: hippocampal learning and memory.**  
A 2022 mouse study found that hippocampal METTL16 promoted MAT2A expression by m6A-dependent stabilization of MAT2A mRNA, and that this pathway was involved in improved dendritic spine plasticity, learning, and memory. The same report also stated that MAT2A overexpression rescued learning/memory and dendritic-spine defects caused by METTL16 knockdown. That is good evidence that MAT2A is functionally upstream of a neural regulatory phenotype, not just a passive marker. ([pubmed.ncbi.nlm.nih.gov](https://pubmed.ncbi.nlm.nih.gov/36307396/)) [S6]

**5) Disease-model extension: Alzheimer-like pathology.**  
In 5×FAD mice, overexpression of MAT2A in the hippocampus improved learning and memory, increased PSD95 and synaptophysin, increased dendritic spine density, and reduced Aβ1-42 accumulation. This places MAT2A within a neural regulatory pathway affecting synaptic structure/function and disease-relevant pathology, though still in a model system rather than humans. ([pubmed.ncbi.nlm.nih.gov](https://pubmed.ncbi.nlm.nih.gov/40308894/)) [S7]

**6) Additional, lower-strength supportive associations in neural tissue.**  
Human postmortem cerebellum studies in alcohol use disorder reported altered one-carbon-metabolism enzymes, including MAT2A, and noted that MAT2 is highly expressed in cerebellar Purkinje neurons and may help determine SAM availability for promoter methylation. A repetitive mild traumatic brain injury study also found dysregulation of hippocampal MAT2A/MAT2B and showed that pharmacologic MAT2A inhibition in hippocampal neuronal cells reduced PRMT7-related methylation output. These studies are more contextual and indirect, but they still support neural-system association. ([pmc.ncbi.nlm.nih.gov](https://pmc.ncbi.nlm.nih.gov/articles/PMC5716183/)) [S8]

## Contradicting or limiting evidence

The main limitation is **scope and directness**. Most of the evidence I found comes from mouse studies, disease models, cell culture, postmortem tissue, or injury/addiction paradigms rather than demonstrations that MAT2A is a central regulator of normal neural homeostasis across healthy human brain circuits. That means the claim is supported as an **association** with neural regulatory processes, but not yet as a universally established homeostatic controller in the nervous system. ([nature.com](https://www.nature.com/articles/s41586-020-1999-0)) [S9]

The evidence is also **context-dependent and region-specific**. For example, chronic ethanol exposure increased cerebellar Mat2a expression while leaving cerebellar SAM unchanged, whereas prenatal alcohol exposure lowered fetal whole-brain Mat2a and later showed sex- and region-specific adult hippocampal/hypothalamic changes. These mixed regional responses argue against a simple, uniform neural effect. ([pubmed.ncbi.nlm.nih.gov](https://pubmed.ncbi.nlm.nih.gov/28369960/)) [S10]

Finally, much of MAT2A’s neural relevance may be **upstream and metabolic-epigenetic**, rather than specific to a dedicated neural signaling pathway. In other words, MAT2A may influence neural regulation because it controls methyl-donor supply for many downstream pathways, not because it is a neuron- or glia-specific homeostatic effector. That still counts as regulatory association, but it narrows how strongly the claim should be interpreted. ([ncbi.nlm.nih.gov](https://www.ncbi.nlm.nih.gov/gene/4144)) [S11]

## Mechanistic / biological rationale

The mechanistic rationale is strong. MAT2A synthesizes SAM, and SAM is the major methyl donor for DNA, RNA, histone, and other methylation reactions. Because neural cells rely heavily on epigenetic and epitranscriptomic regulation for cell-state control, inflammatory programming, and synaptic plasticity, shifts in MAT2A activity can plausibly alter neural-system regulation through methylation capacity. ([ncbi.nlm.nih.gov](https://www.ncbi.nlm.nih.gov/gene/4144)) [S12]

This is not just abstract theory: mammalian cells regulate SAM homeostasis by controlling MAT2A expression/splicing, including METTL16-dependent regulation of MAT2A transcripts. That creates a feedback architecture in which nutrient/metabolite state influences MAT2A, which then influences methylation-dependent cellular programs. ([pubmed.ncbi.nlm.nih.gov](https://pubmed.ncbi.nlm.nih.gov/33949310/)) [S13]

In neural contexts, the downstream readouts match that framework. In astrocytes, MAT2A cooperates with MAFG to promote DNA methylation and suppress antioxidant/anti-inflammatory programs during CNS inflammation. In hippocampus, the METTL16–MAT2A axis affects global m6A, dendritic spine plasticity, synaptic proteins, and memory-related behavior. Together, these findings fit a model in which MAT2A serves as a metabolic-epigenetic regulator of neural cell state rather than a classic neurotransmission protein. ([nature.com](https://www.nature.com/articles/s41586-020-1999-0)) [S14]

## Bottom line

The claim is **likely true**. MAT2A has credible published associations with **regulatory processes in neural systems**, especially **astrocyte inflammatory programming** and **hippocampal synaptic plasticity/memory-related regulation**. The evidence is strongest in **preclinical and disease-related settings**, and weaker for a broad, canonical role in normal human neural homeostasis. So the most accurate wording is: **MAT2A is a plausible and increasingly supported metabolic-epigenetic regulator in neural systems, but the evidence is not yet broad enough to call it a firmly established master neural homeostasis gene.** ([nature.com](https://www.nature.com/articles/s41586-020-1999-0)) [S15]

## Cited sources

- **[S1]** MAFG-driven astrocytes promote CNS inflammation | Nature — https://www.nature.com/articles/s41586-020-1999-0
- **[S2]** MAFG-driven astrocytes promote CNS inflammation | Nature — https://www.nature.com/articles/s41586-020-1999-0
- **[S3]** MAT2A methionine adenosyltransferase 2A [Homo sapiens (human)] - Gene - NCBI — https://www.ncbi.nlm.nih.gov/gene/4144
- **[S4]** Single cell type - MAT2A - The Human Protein Atlas — https://www.proteinatlas.org/ENSG00000168906-MAT2A/single%2Bcell
- **[S5]** MAFG-driven astrocytes promote CNS inflammation | Nature — https://www.nature.com/articles/s41586-020-1999-0
- **[S6]** Knockdown of METTL16 disrupts learning and memory by reducing the stability of MAT2A mRNA - PubMed — https://pubmed.ncbi.nlm.nih.gov/36307396/
- **[S7]** Effects of m6A methylation of MAT2A mRNA regulated by METTL16 on learning and memory, hippocampal synaptic plasticity and Aβ1-42 in 5 × FAD mice - PubMed — https://pubmed.ncbi.nlm.nih.gov/40308894/
- **[S8]** 
            Emerging Role of One-Carbon Metabolism and DNA Methylation Enrichment on δ-Containing GABAA Receptor Expression in the Cerebellum of Subjects with Alcohol Use Disorders (AUD) - PMC
         — https://pmc.ncbi.nlm.nih.gov/articles/PMC5716183/
- **[S9]** MAFG-driven astrocytes promote CNS inflammation | Nature — https://www.nature.com/articles/s41586-020-1999-0
- **[S10]** Chronic Alcohol Exposure Differentially Alters One-Carbon Metabolism in Rat Liver and Brain - PubMed — https://pubmed.ncbi.nlm.nih.gov/28369960/
- **[S11]** MAT2A methionine adenosyltransferase 2A [Homo sapiens (human)] - Gene - NCBI — https://www.ncbi.nlm.nih.gov/gene/4144
- **[S12]** MAT2A methionine adenosyltransferase 2A [Homo sapiens (human)] - Gene - NCBI — https://www.ncbi.nlm.nih.gov/gene/4144
- **[S13]** SAM homeostasis is regulated by CFIm-mediated splicing of MAT2A - PubMed — https://pubmed.ncbi.nlm.nih.gov/33949310/
- **[S14]** MAFG-driven astrocytes promote CNS inflammation | Nature — https://www.nature.com/articles/s41586-020-1999-0
- **[S15]** MAFG-driven astrocytes promote CNS inflammation | Nature — https://www.nature.com/articles/s41586-020-1999-0
