## Supplementary material for "Understanding Substance Dependence: What Differentiates Addictive from Non-Addictive Drugs?": Folder for supplementary files: non-addictive-genes-claims-first100-018-gja1-gene-protein-has-been-associated-with-regulat-4b60b439e441.md

# non-addictive-genes-claims-first100-018-4b60b439e441

**Original claim:** GJA1 gene/protein has been associated with regulatory or homeostatic biological processes in neural systems.

## Claim

GJA1 gene/protein has been associated with regulatory or homeostatic biological processes in neural systems. ([ncbi.nlm.nih.gov](https://www.ncbi.nlm.nih.gov/gene/2697?utm_source=openai)) [S1]

## Verdict

Verdict: Highly Supported

GJA1 encodes connexin-43 (Cx43), a major astrocytic gap-junction protein. Multiple primary studies in the CNS show that astroglial Cx43 participates in classic neural homeostatic/regulatory functions, including extracellular potassium handling, glutamate/glutamine support of synaptic transmission, metabolic substrate trafficking, blood-brain barrier maintenance, and sleep-wake regulation. The strongest evidence is from rodent genetic and physiological studies rather than direct human intervention studies, but the broad claim of association is well supported. ([ncbi.nlm.nih.gov](https://www.ncbi.nlm.nih.gov/gene/2697?utm_source=openai)) [S2]

## Confidence

High

## Supporting evidence

- **GJA1 is the gene for connexin-43, a gap-junction protein with strong relevance to astrocyte communication.** NCBI identifies human **GJA1** as a protein-coding gene whose product is a component of gap junctions that mediate diffusion of small molecules between cells. In the brain, Cx43 is described as the major constituent of astrocytic gap junctions and is highly expressed throughout the CNS. ([ncbi.nlm.nih.gov](https://www.ncbi.nlm.nih.gov/gene/2697?utm_source=openai)) [S3]

- **Extracellular potassium homeostasis:** In a classic hippocampal study using mice with coupling-deficient astrocytes (conditional Cx43 loss plus Cx30 deletion), astrocytic gap junction disruption **slowed K+ clearance**, **increased K+ accumulation during synchronized firing**, impaired radial K+ relocation in part of the hippocampus, and lowered the threshold for epileptiform activity. The authors concluded that astrocytic gap junctions accelerate potassium clearance and contribute to extracellular K+ homeostasis. ([pmc.ncbi.nlm.nih.gov](https://pmc.ncbi.nlm.nih.gov/articles/PMC6675300/)) [S4]

- **Synaptic glutamate and excitability regulation:** In hippocampal Cx30/Cx43 double-knockout mice, altered astroglial physiology led to **increased extracellular potassium and glutamate levels during evoked activity**, directly linking connexin-mediated astroglial networks to regulation of basal synaptic transmission and plasticity. ([pmc.ncbi.nlm.nih.gov](https://pmc.ncbi.nlm.nih.gov/articles/PMC3100942/)) [S5]

- **Cx43-specific control of synaptic efficacy:** In an astrocyte-specific **Cx43** knockout model, Cx43 loss reduced astroglial coupling by about 60% and caused an approximately 30% reduction in excitatory synaptic transmission in hippocampal CA1, with the effect attributed to altered synaptically released glutamate rather than neuronal intrinsic excitability. That is a direct demonstration that **GJA1/Cx43** itself, not only astrocyte networks in general, regulates synaptic function. ([pmc.ncbi.nlm.nih.gov](https://pmc.ncbi.nlm.nih.gov/articles/PMC4173282/)) [S6]

- **Glutamine homeostasis at synapses:** A 2022 study showed that astroglial Cx43 is enriched in **perisynaptic astroglial processes**, and that its hemichannels open in an activity-dependent manner regulated by glutamate/K+ signaling via astroglial Kir4.1. The study linked this Cx43 activity to glutamine supply required for physiological synaptic activity and recognition memory, placing Cx43 squarely in neurotransmitter-cycle homeostasis. ([nature.com](https://www.nature.com/articles/s41467-022-28331-7)) [S7]

- **Presynaptic vesicle release support via Cx43-dependent glutamine supply:** A follow-up study in glial Cx43 knockout mice found slower synaptic vesicle release and lower release probability at hippocampal synapses, with the phenotype depending on glutamine supply through Cx43 hemichannel function. This further supports a homeostatic role in sustaining normal neurotransmission. ([pubmed.ncbi.nlm.nih.gov](https://pubmed.ncbi.nlm.nih.gov/37190042/)) [S8]

- **Neurometabolic homeostasis:** In Science, Rouach et al. showed that connexin 43 and 30 permit **intercellular trafficking of glucose and its metabolites through astroglial networks**. Under extracellular glucose deprivation, supplying glucose or lactate to astrocytes sustained glutamatergic synaptic transmission only when astrocytes remained gap-junction connected, demonstrating a homeostatic energy-delivery role. ([pubmed.ncbi.nlm.nih.gov](https://pubmed.ncbi.nlm.nih.gov/19056987/)) [S9]

- **Cx43-dependent metabolic support for plasticity:** In cortex, experimental disruption of astrocytic Cx43-based coupling reduced astrocyte network size and impaired long-term potentiation; intracellular lactate rescue restored LTP. This strongly supports the idea that Cx43-mediated astrocytic networks act as an energy reservoir for neural plasticity. ([nature.com](https://www.nature.com/articles/s41467-020-15778-9)) [S10]

- **Neurovascular/BBB homeostasis:** Mice lacking astroglial connexins Cx43/Cx30 developed astrocyte endfoot edema, reduced aquaporin-4 and β-dystroglycan at perivascular endfeet, and a blood-brain barrier that became abnormally permeable under stress. The authors concluded that astroglial connexins are necessary to maintain BBB integrity. ([pmc.ncbi.nlm.nih.gov](https://pmc.ncbi.nlm.nih.gov/articles/PMC3421093/)) [S11]

- **Behavioral-state regulation:** In astrocyte-specific Cx43 knockout mice, investigators observed excessive sleepiness and fragmented wakefulness, reduced orexin-neuron excitability, and rescue of the phenotype with lactate delivery to the lateral hypothalamus. This is direct evidence that Cx43-mediated astroglial metabolic networks regulate a major neural homeostatic function: the sleep-wake cycle. ([pmc.ncbi.nlm.nih.gov](https://pmc.ncbi.nlm.nih.gov/articles/PMC5617118/)) [S12]

## Contradicting or limiting evidence

- **Most strong evidence is preclinical.** The clearest demonstrations come from mouse knockouts, brain slices, and cellular physiology. That is enough to support the claim of association in neural systems, but it means direct causal inference to human neural physiology is more limited. Human data are mostly expression/pathology studies rather than mechanistic intervention studies. ([pmc.ncbi.nlm.nih.gov](https://pmc.ncbi.nlm.nih.gov/articles/PMC5617118/)) [S13]

- **Some functions are partly redundant with other connexins, especially Cx30.** Several important studies used **Cx43/Cx30 double knockout** models rather than isolated GJA1 loss, implying that some homeostatic effects reflect astrocyte network coupling more broadly and that Cx30 can compensate in some settings. In the potassium-buffering work, substantial K+ clearance was still preserved, indicating that gap-junction-dependent mechanisms only partially account for hippocampal buffering. ([pmc.ncbi.nlm.nih.gov](https://pmc.ncbi.nlm.nih.gov/articles/PMC6675300/)) [S14]

- **Cx43 is not uniformly “protective” in all neural contexts.** In inflammatory demyelination/EAE, brain gray matter astroglia-specific Cx43 ablation reduced pro-inflammatory astroglial and microglial signatures and attenuated spinal cord inflammatory demyelination. So Cx43 participates in neural regulation, but its net effect can become maladaptive in disease. ([jneuroinflammation.biomedcentral.com](https://jneuroinflammation.biomedcentral.com/articles/10.1186/s12974-021-02176-1)) [S15]

- **Not all observed effects are necessarily due only to canonical gap-junction channel function.** The BBB study explicitly notes that whether the barrier phenotype is solely due to loss of Cx-based gap-junction channels remains unresolved; Cx43 also has hemichannel and scaffolding/cytoskeletal roles. ([pmc.ncbi.nlm.nih.gov](https://pmc.ncbi.nlm.nih.gov/articles/PMC3421093/)) [S16]

- **Transcriptomic consequences can vary by context and genetic background.** Conditional Gja1 deletion altered numerous brain transcripts, but the specific gene-expression changes differed substantially by mouse strain, suggesting context-sensitive downstream biology. ([pubmed.ncbi.nlm.nih.gov](https://pubmed.ncbi.nlm.nih.gov/22771707/)) [S17]

## Mechanistic / biological rationale

Cx43 forms astrocytic **gap junction channels** and **hemichannels**, allowing passage of ions and small metabolites/signaling molecules between astrocytes and, in some contexts, between astrocytes and extracellular space. That architecture gives a plausible mechanism for several homeostatic functions: spatial redistribution of extracellular K+, diffusion of glucose/lactate/glutamine across astrocyte networks, and modulation of synaptic transmitter availability. ([ncbi.nlm.nih.gov](https://www.ncbi.nlm.nih.gov/gene/2697?utm_source=openai)) [S18]

This rationale is borne out experimentally. Loss of astroglial connexin coupling slows K+ clearance and changes extracellular K+ dynamics; loss of Cx43 alters astrocyte volume and synaptic glutamate handling; and activity-dependent Cx43 hemichannel opening near synapses supports glutamine availability for neurotransmission. Together, these findings fit a model in which GJA1/Cx43 helps astrocytes stabilize the extracellular ionic, metabolic, and transmitter environment required for normal neuronal signaling. ([pmc.ncbi.nlm.nih.gov](https://pmc.ncbi.nlm.nih.gov/articles/PMC6675300/)) [S19]

At the neurovascular interface, Cx43-enriched astrocyte endfeet are also positioned to support BBB and water/ion homeostasis. Deleting astroglial connexins disrupts perivascular endfoot structure, lowers AQP4/β-dystroglycan, and weakens BBB integrity under stress, which is mechanistically consistent with a role in maintaining neural tissue homeostasis beyond synapses alone. ([pmc.ncbi.nlm.nih.gov](https://pmc.ncbi.nlm.nih.gov/articles/PMC3421093/)) [S20]

## Bottom line

Yes. The claim is **strongly supported**: **GJA1/Cx43 is repeatedly implicated in regulatory and homeostatic processes in neural systems**, especially through astrocyte-mediated control of ionic balance, neurotransmitter support, metabolic coupling, neurovascular integrity, and vigilance-state regulation. The evidence is strongest in rodent CNS models, and some effects are context-dependent or partly redundant with other connexins, but the overall association is well established. ([pmc.ncbi.nlm.nih.gov](https://pmc.ncbi.nlm.nih.gov/articles/PMC6675300/)) [S21]

## Cited sources

- **[S1]** GJA1 gap junction protein alpha 1 [Homo sapiens (human)] - Gene - NCBI — https://www.ncbi.nlm.nih.gov/gene/2697?utm_source=openai
- **[S2]** GJA1 gap junction protein alpha 1 [Homo sapiens (human)] - Gene - NCBI — https://www.ncbi.nlm.nih.gov/gene/2697?utm_source=openai
- **[S3]** GJA1 gap junction protein alpha 1 [Homo sapiens (human)] - Gene - NCBI — https://www.ncbi.nlm.nih.gov/gene/2697?utm_source=openai
- **[S4]** 
            The Impact of Astrocytic Gap Junctional Coupling on Potassium Buffering in the Hippocampus - PMC
         — https://pmc.ncbi.nlm.nih.gov/articles/PMC6675300/
- **[S5]** 
            Astroglial networks scale synaptic activity and plasticity - PMC
         — https://pmc.ncbi.nlm.nih.gov/articles/PMC3100942/
- **[S6]** 
            Astroglial connexin 43 sustains glutamatergic synaptic efficacy - PMC
         — https://pmc.ncbi.nlm.nih.gov/articles/PMC4173282/
- **[S7]** Physiological synaptic activity and recognition memory require astroglial glutamine | Nature Communications — https://www.nature.com/articles/s41467-022-28331-7
- **[S8]** Astroglial Connexin 43 Regulates Synaptic Vesicle Release at Hippocampal Synapses - PubMed — https://pubmed.ncbi.nlm.nih.gov/37190042/
- **[S9]** Astroglial metabolic networks sustain hippocampal synaptic transmission - PubMed — https://pubmed.ncbi.nlm.nih.gov/19056987/
- **[S10]** Stress gates an astrocytic energy reservoir to impair synaptic plasticity | Nature Communications — https://www.nature.com/articles/s41467-020-15778-9
- **[S11]** 
            Deletion of astroglial connexins weakens the blood–brain barrier - PMC
         — https://pmc.ncbi.nlm.nih.gov/articles/PMC3421093/
- **[S12]** 
            Connexin 43-mediated astroglial metabolic networks contribute to the regulation of the sleep-wake cycle - PMC
         — https://pmc.ncbi.nlm.nih.gov/articles/PMC5617118/
- **[S13]** 
            Connexin 43-mediated astroglial metabolic networks contribute to the regulation of the sleep-wake cycle - PMC
         — https://pmc.ncbi.nlm.nih.gov/articles/PMC5617118/
- **[S14]** 
            The Impact of Astrocytic Gap Junctional Coupling on Potassium Buffering in the Hippocampus - PMC
         — https://pmc.ncbi.nlm.nih.gov/articles/PMC6675300/
- **[S15]** Brain gray matter astroglia-specific connexin 43 ablation attenuates spinal cord inflammatory demyelination | Journal of Neuroinflammation | Springer Nature Link — https://jneuroinflammation.biomedcentral.com/articles/10.1186/s12974-021-02176-1
- **[S16]** 
            Deletion of astroglial connexins weakens the blood–brain barrier - PMC
         — https://pmc.ncbi.nlm.nih.gov/articles/PMC3421093/
- **[S17]** The connexin43-dependent transcriptome during brain development: importance of genetic background - PubMed — https://pubmed.ncbi.nlm.nih.gov/22771707/
- **[S18]** GJA1 gap junction protein alpha 1 [Homo sapiens (human)] - Gene - NCBI — https://www.ncbi.nlm.nih.gov/gene/2697?utm_source=openai
- **[S19]** 
            The Impact of Astrocytic Gap Junctional Coupling on Potassium Buffering in the Hippocampus - PMC
         — https://pmc.ncbi.nlm.nih.gov/articles/PMC6675300/
- **[S20]** 
            Deletion of astroglial connexins weakens the blood–brain barrier - PMC
         — https://pmc.ncbi.nlm.nih.gov/articles/PMC3421093/
- **[S21]** 
            The Impact of Astrocytic Gap Junctional Coupling on Potassium Buffering in the Hippocampus - PMC
         — https://pmc.ncbi.nlm.nih.gov/articles/PMC6675300/
