## Supplementary material for "Understanding Substance Dependence: What Differentiates Addictive from Non-Addictive Drugs?": Folder for supplementary files: non-addictive-genes-claims-first100-019-nrxn1-gene-protein-has-been-associated-with-regula-737a663fa46e.md

# non-addictive-genes-claims-first100-019-737a663fa46e

**Original claim:** NRXN1 gene/protein has been associated with regulatory or homeostatic biological processes in neural systems.

## Claim

NRXN1 gene/protein has been associated with regulatory or homeostatic biological processes in neural systems.

## Verdict

Verdict: Highly Supported

## Confidence

High

## Supporting evidence

NRXN1 is directly annotated in expert synaptic ontologies and reference databases as participating in **regulatory synaptic processes**, not just structural presence at synapses. SynGO lists NRXN1 under presynapse assembly, regulation of presynapse assembly, regulation of postsynaptic density/specialization assembly, regulation of the synaptic vesicle cycle, and **regulation of trans-synaptic signaling by endocannabinoid, modulating synaptic transmission**. UniProt/Reactome likewise describe NRXN1 as a cell-surface protein involved in cell-cell interactions, exocytosis, and **regulation of signal transmission**, including calcium channel activity, Ca²⁺-triggered neurotransmitter release, and postsynaptic NMDA receptor regulation. ([syngoportal.org](https://www.syngoportal.org/gene_hgnc8008.html)) [S1]

The broader literature strongly supports NRXN1 as a regulator of synapse properties. A review focused specifically on NRXN1 concluded that available neurobiological evidence supports roles in **synaptic formation, organization, transmission, and plasticity**. A major neurexin review similarly describes neurexins as molecular codes that shape neuronal synapses and instruct synapse-specific functional properties through regulated isoform expression and alternative splicing. ([pubmed.ncbi.nlm.nih.gov](https://pubmed.ncbi.nlm.nih.gov/31138894/)) [S2]

Primary experimental work shows that neurexin-family proteins, including Nrxn1 isoforms, regulate core presynaptic functions. In triple α-neurexin knockout mice, α-neurexins were found to be **essential for Ca²⁺-triggered neurotransmitter release**, because synaptic Ca²⁺ channel function was markedly reduced even though synapse formation was preserved. A later study specifically showed that **Nrxn1α facilitates presynaptic CaV2.1-mediated Ca²⁺ influx**, providing a direct mechanism for regulation of release probability and synaptic output. ([pubmed.ncbi.nlm.nih.gov](https://pubmed.ncbi.nlm.nih.gov/12827191/)) [S3]

NRXN1 also regulates **trans-synaptic control of postsynaptic receptors**. In hippocampal synapses, alternative splicing of presynaptic Nrxn1 at splice site 4 strongly altered **postsynaptic NMDA receptor-mediated responses** without changing synapse density, demonstrating a clear regulatory role rather than a merely structural one. Related work showed that Nrxn1-containing neurexin-cerebellin complexes can regulate postsynaptic glutamate receptor responses in a circuit-dependent way. ([pubmed.ncbi.nlm.nih.gov](https://pubmed.ncbi.nlm.nih.gov/31005376/)) [S4]

Human neuronal models support the same interpretation. In isogenic human neurons, heterozygous NRXN1 loss-of-function mutations selectively **impaired neurotransmitter release** without altering neuronal differentiation or synapse formation, indicating a direct role in regulating synaptic function. In patient-derived cortical neurons with NRXN1α haploinsufficiency, investigators reported **increased excitability**, altered calcium signaling, and transcriptomic evidence for upregulated voltage-gated ion channel and glutamatergic signaling pathways. ([pmc.ncbi.nlm.nih.gov](https://pmc.ncbi.nlm.nih.gov/articles/PMC4560990/)) [S5]

There is also evidence that NRXN1 is subject to dynamic regulatory mechanisms at synapses. Super-resolution imaging showed that synaptic Nrxn1 forms presynaptic nanoclusters whose abundance and size change during development and are **dynamically regulated by ADAM10-mediated ectodomain cleavage**. This is consistent with an actively regulated synaptic organizer rather than a static adhesion molecule. ([pmc.ncbi.nlm.nih.gov](https://pmc.ncbi.nlm.nih.gov/articles/PMC6683742/)) [S6]

Finally, there is meaningful evidence for a **homeostatic** connection, although it is narrower than the evidence for general regulation. At the mouse neuromuscular junction, α-neurexins were reported to be required for efficient transmitter release and to contribute to the **molecular mechanism of synaptic homeostasis** during compensatory presynaptic upregulation after postsynaptic receptor reduction. In central excitatory circuits, β-neurexins regulated tonic endocannabinoid signaling and presynaptic LTP, showing that neurexin isoforms help stabilize and tune synaptic strength through retrograde signaling pathways. Because NRXN1 produces both α and β isoforms, these findings support the claim that NRXN1/neurexin-1 is associated with regulatory and, to a degree, homeostatic neural processes. ([pubmed.ncbi.nlm.nih.gov](https://pubmed.ncbi.nlm.nih.gov/16406382/)) [S7]

## Contradicting or limiting evidence

The strongest caveat is that **“homeostatic” evidence is less direct and less extensive than “regulatory” evidence**. Some of the clearest homeostasis language comes from α-neurexin studies at the peripheral neuromuscular junction, not from human NRXN1-specific experiments in central brain circuits. Likewise, the endocannabinoid work was performed on β-neurexins as a class rather than isolating NRXN1 alone in every experiment. So the claim is very well supported if interpreted as “regulatory,” but only moderately direct if interpreted as “NRXN1 is a core homeostatic gene” in the narrow technical sense. ([pubmed.ncbi.nlm.nih.gov](https://pubmed.ncbi.nlm.nih.gov/16406382/)) [S8]

There is also substantial **context dependence**. Some studies found that Nrxn1 is important for shaping synaptic properties but **not essential for basic synapse formation or maintenance** in certain hippocampal culture settings. Mouse circuit studies likewise show synapse-class specificity: Nrxn1α deletion affects some projections and cell types strongly while sparing others. That means NRXN1 is not a universal regulator of all neural homeostasis everywhere; its effects depend on isoform, splice state, ligand repertoire, and circuit context. ([pmc.ncbi.nlm.nih.gov](https://pmc.ncbi.nlm.nih.gov/articles/PMC6683742/)) [S9]

## Mechanistic / biological rationale

Mechanistically, NRXN1 is well positioned to mediate regulatory and stabilizing functions in neural systems. It is a **presynaptic adhesion receptor** that bridges pre- and postsynaptic compartments through interactions with neuroligins, LRRTMs, cerebellins/GluD complexes, dystroglycan, and intracellular partners such as CASK. These interactions let NRXN1 influence presynaptic Ca²⁺ entry, vesicle release, and postsynaptic receptor composition. ([reactome.org](https://www.reactome.org/content/schema/instance/browser/uniprot%3AQ9ULB1)) [S10]

Its regulatory capacity is amplified by **extensive alternative splicing** and post-translational modification. Splice variation in Nrxn1 changes ligand binding and selectively tunes NMDA receptor signaling; heparan-sulfate modification of neurexin-1 is required for normal ligand interactions and for proper synapse development and transmission; and ADAM10-mediated cleavage dynamically alters synaptic Nrxn1 nanoclusters. Together, these mechanisms provide a biologically plausible basis for how NRXN1 can contribute to synaptic tuning, stability, and adaptive regulation over time. ([pubmed.ncbi.nlm.nih.gov](https://pubmed.ncbi.nlm.nih.gov/31005376/)) [S11]

The heparan-sulfate study is especially important because it ties neurexin-1 to a broader signaling framework: mice lacking HS on neurexin-1 had structural and functional deficits at central synapses, and HS-dependent interactions expanded the set of potential extracellular ligands. That supports a model in which NRXN1 acts as a **modifiable signaling hub** for organizing and regulating synaptic communication. ([pubmed.ncbi.nlm.nih.gov](https://pubmed.ncbi.nlm.nih.gov/30100184/)) [S12]

## Bottom line

Yes. The claim is well supported. NRXN1/neurexin-1 is strongly associated with **regulatory biological processes in neural systems**, including presynaptic release control, Ca²⁺ channel function, trans-synaptic signaling, receptor regulation, synapse assembly/organization, and plasticity. Evidence for a specifically **homeostatic** role also exists, but it is narrower and often comes from isoform-family or non-central-synapse studies rather than exclusively from NRXN1-specific human CNS data. Overall, the safest summary is: **NRXN1 is clearly a synaptic regulatory gene/protein, with plausible and partly demonstrated links to synaptic homeostatic mechanisms.** ([syngoportal.org](https://www.syngoportal.org/gene_hgnc8008.html)) [S13]

## Cited sources

- **[S1]** SynGO - gene details — https://www.syngoportal.org/gene_hgnc8008.html
- **[S2]** Genetic insights and neurobiological implications from NRXN1 in neuropsychiatric disorders - PubMed — https://pubmed.ncbi.nlm.nih.gov/31138894/
- **[S3]** Alpha-neurexins couple Ca2+ channels to synaptic vesicle exocytosis - PubMed — https://pubmed.ncbi.nlm.nih.gov/12827191/
- **[S4]** Alternative Splicing of Presynaptic Neurexins Differentially Controls Postsynaptic NMDA and AMPA Receptor Responses - PubMed — https://pubmed.ncbi.nlm.nih.gov/31005376/
- **[S5]** 
            Human Neuropsychiatric Disease Modeling Using Conditional Deletion Reveals Synaptic Transmission Defects Caused By Heterozygous Mutations in NRXN1 - PMC
         — https://pmc.ncbi.nlm.nih.gov/articles/PMC4560990/
- **[S6]** 
            Synaptic neurexin-1 assembles into dynamically regulated active zone nanoclusters - PMC
         — https://pmc.ncbi.nlm.nih.gov/articles/PMC6683742/
- **[S7]** alpha-Neurexins are required for efficient transmitter release and synaptic homeostasis at the mouse neuromuscular junction - PubMed — https://pubmed.ncbi.nlm.nih.gov/16406382/
- **[S8]** alpha-Neurexins are required for efficient transmitter release and synaptic homeostasis at the mouse neuromuscular junction - PubMed — https://pubmed.ncbi.nlm.nih.gov/16406382/
- **[S9]** 
            Synaptic neurexin-1 assembles into dynamically regulated active zone nanoclusters - PMC
         — https://pmc.ncbi.nlm.nih.gov/articles/PMC6683742/
- **[S10]** Reactome | UniProt:Q9ULB1 NRXN1 — https://www.reactome.org/content/schema/instance/browser/uniprot%3AQ9ULB1
- **[S11]** Alternative Splicing of Presynaptic Neurexins Differentially Controls Postsynaptic NMDA and AMPA Receptor Responses - PubMed — https://pubmed.ncbi.nlm.nih.gov/31005376/
- **[S12]** Heparan Sulfate Organizes Neuronal Synapses through Neurexin Partnerships - PubMed — https://pubmed.ncbi.nlm.nih.gov/30100184/
- **[S13]** SynGO - gene details — https://www.syngoportal.org/gene_hgnc8008.html
