## Supplementary material for "Understanding Substance Dependence: What Differentiates Addictive from Non-Addictive Drugs?": Folder for supplementary files: non-addictive-genes-claims-first100-020-prdx1-gene-protein-has-been-associated-with-regula-3774df2b63a1.md

# non-addictive-genes-claims-first100-020-3774df2b63a1

**Original claim:** PRDX1 gene/protein has been associated with regulatory or homeostatic biological processes in neural systems.

## Claim

PRDX1 gene/protein has been associated with regulatory or homeostatic biological processes in neural systems.

## Verdict

Verdict: Highly Supported

## Confidence

High

## Supporting evidence

PRDX1 is not just a generic oxidative-stress marker. Authoritative gene resources describe PRDX1 as a thioredoxin-dependent peroxidase involved in hydrogen peroxide breakdown and **cell redox homeostasis**, with hydrogen peroxide explicitly noted as a signaling molecule whose levels influence growth, maturation, survival, and other cell functions. In other words, PRDX1 has an established biochemical role in regulating intracellular redox tone, which is a classic homeostatic process. ([medlineplus.gov](https://medlineplus.gov/download/genetics/gene/prdx1.pdf)) [S1]

That general role is linked to the nervous system by both expression and function. Reviews of CNS peroxiredoxins report distinct neural cell-type distribution of Prdx isoforms and note PRDX1/PrxI expression in glial and oligodendrocyte-related compartments, with region- and cell-type-specific patterns in murine and human CNS tissue. A recent review further states that peroxiredoxins in the CNS regulate processes including oxidative stress, neuroinflammation, apoptosis, mitochondrial dynamics, ER stress, and synaptic plasticity, all of which are regulatory/homeostatic neural processes. ([pubmed.ncbi.nlm.nih.gov](https://pubmed.ncbi.nlm.nih.gov/18084903/)) [S2]

There is also strong **direct mechanistic evidence** that Prdx1 regulates neural developmental programs. In a Cell paper, Prdx1 was shown to interact with GDE2, and loss of Prdx1 caused motor-neuron differentiation deficits analogous to GDE2 loss. The study concluded that Prdx1 is a “pivotal regulator” of GDE2 activity and that coupled thiol-redox cascades involving Prdx1 control neuronal differentiation in the spinal cord. This is clear evidence for a regulatory role in a neural system, not merely passive association. ([pubmed.ncbi.nlm.nih.gov](https://pubmed.ncbi.nlm.nih.gov/19766572/)) [S3]

Evidence is similarly strong for neural homeostatic roles in injury-response states, especially in glia. In acute ischemic stroke models, Prdx1 was required for a stroke-associated microglial population with enhanced antioxidant function; Prdx1 knockout reduced this microglial state, lowered resistance to ROS damage, increased inflammation, and worsened infarction. That directly supports a role for PRDX1 in maintaining redox and inflammatory balance in a neural immune cell population. ([pubmed.ncbi.nlm.nih.gov](https://pubmed.ncbi.nlm.nih.gov/35688114/)) [S4]

Another neural injury study found that Prdx1 was upregulated in astrocytes and microglia after intracerebral hemorrhage, and that overexpression reduced inflammation, apoptosis, symptoms, and mortality. Mechanistically, the paper reported that Prdx1 affected the stability of inflammation- and apoptosis-related mRNAs, showing a post-transcriptional regulatory function in neural tissue rather than only an antioxidant “cleanup” role. ([pubmed.ncbi.nlm.nih.gov](https://pubmed.ncbi.nlm.nih.gov/32210752/)) [S5]

Neuronal data also support a homeostatic role. In spiral ganglion neurons, PRDX1 deficiency suppressed autophagy and worsened cisplatin-induced neuronal loss, whereas PRDX1 upregulation activated autophagy, reduced ROS accumulation and apoptosis, and protected hearing-related neurons via PTEN-AKT signaling. Since autophagy and ROS buffering are core cellular homeostatic programs, this is directly relevant to the claim. ([pubmed.ncbi.nlm.nih.gov](https://pubmed.ncbi.nlm.nih.gov/33749526/)) [S6]

Likewise, in a hippocampal neuronal model of Alzheimer-like injury, Prx1 overexpression reduced neuronal apoptosis, synaptic loss, tau phosphorylation, mitochondrial dysfunction, and Ca²⁺/calpain/Cdk5-linked pathology. Separately, a neurodegeneration review summarizes human AD-brain studies showing altered PRDX1 levels and associations between reduced neuronal PRDX1 and increased oxidative damage/AMPK-mTOR-tau abnormalities. Together, these findings support association with neuronal stress-regulation and homeostatic maintenance pathways. ([pubmed.ncbi.nlm.nih.gov](https://pubmed.ncbi.nlm.nih.gov/38977865/)) [S7]

## Contradicting or limiting evidence

The biggest limitation is that much of the strongest evidence is **preclinical**: rodent models, chick spinal cord, primary glia, or neuronal cell lines. Direct evidence that PRDX1 governs baseline homeostasis in the healthy adult human brain is less extensive than evidence in development or injury/disease models. ([pubmed.ncbi.nlm.nih.gov](https://pubmed.ncbi.nlm.nih.gov/19766572/)) [S8]

Neural expression is also cell-type and context dependent. An older CNS review described basal PrxI/PRDX1 expression as predominantly glial rather than neuronal, whereas newer reviews note broader but still differential CNS expression. So PRDX1 should not be framed as a universal neuron-specific homeostatic factor across all neural compartments. ([pubmed.ncbi.nlm.nih.gov](https://pubmed.ncbi.nlm.nih.gov/18084903/)) [S9]

Importantly, PRDX1’s effects are **not uniformly protective**. In one ischemia-reperfusion study, PRDX1 knockout reduced infarct size and inflammation, and the authors concluded that PRDX1 promoted injury via TLR4/NF-κB-linked inflammation. Broader inflammation reviews also describe extracellular PRDX1/peroxiredoxins released from damaged brain cells as DAMP-like signals that can activate TLR2/TLR4 pathways and drive inflammatory cytokine production. So PRDX1 can participate in maladaptive signaling after neural injury, depending on localization and context. ([pubmed.ncbi.nlm.nih.gov](https://pubmed.ncbi.nlm.nih.gov/31526567/)) [S10]

## Mechanistic / biological rationale

The biological rationale is strong. PRDX1 catalyzes peroxide reduction and is annotated to hydrogen peroxide catabolism and cell redox homeostasis. Because low-level H₂O₂ acts as a signaling molecule, enzymes like PRDX1 are positioned to regulate signaling thresholds rather than merely detoxify damage. Reviews of PRDX1 specifically describe it as a modulator of peroxide signaling and redox-dependent protein interactions. ([medlineplus.gov](https://medlineplus.gov/download/genetics/gene/prdx1.pdf)) [S11]

In neural systems, that chemistry maps onto recognizable regulatory circuits. During spinal cord development, Prdx1 redox activity activates GDE2 to control neuronal differentiation. In stroke-associated microglia, Prdx1 supports an antioxidant transcriptional program and cell survival. In hemorrhagic injury, Prdx1 influences inflammatory/apoptotic mRNA stability in astrocytes and microglia. In auditory neurons, it promotes autophagy through PTEN-AKT signaling, reducing ROS and apoptosis. These are all mechanistically plausible ways a redox regulator would contribute to neural homeostasis. ([pubmed.ncbi.nlm.nih.gov](https://pubmed.ncbi.nlm.nih.gov/19766572/)) [S12]

At the same time, the same protein can become pro-inflammatory when released extracellularly from injured cells, consistent with the broader concept that redox/homeostatic proteins can switch from intracellular housekeeping to extracellular danger signaling after tissue damage. ([pubmed.ncbi.nlm.nih.gov](https://pubmed.ncbi.nlm.nih.gov/31526567/)) [S13]

## Bottom line

Yes. The literature strongly supports that PRDX1 is associated with **regulatory and homeostatic processes in neural systems**, especially redox control, neuronal differentiation, microglial antioxidant programs, inflammatory regulation, autophagy, and neuronal stress responses. The main caveat is that this evidence is mostly preclinical and context dependent: intracellular PRDX1 often supports neural homeostasis, whereas extracellular PRDX1 after injury can promote neuroinflammation. ([pubmed.ncbi.nlm.nih.gov](https://pubmed.ncbi.nlm.nih.gov/19766572/)) [S14]

## Cited sources

- **[S1]** PRDX1 gene — https://medlineplus.gov/download/genetics/gene/prdx1.pdf
- **[S2]** Peroxiredoxins in the central nervous system - PubMed — https://pubmed.ncbi.nlm.nih.gov/18084903/
- **[S3]** The antioxidant enzyme Prdx1 controls neuronal differentiation by thiol-redox-dependent activation of GDE2 - PubMed — https://pubmed.ncbi.nlm.nih.gov/19766572/
- **[S4]** The antioxidant enzyme Peroxiredoxin-1 controls stroke-associated microglia against acute ischemic stroke - PubMed — https://pubmed.ncbi.nlm.nih.gov/35688114/
- **[S5]** Prdx1 Reduces Intracerebral Hemorrhage-Induced Brain Injury via Targeting Inflammation- and Apoptosis-Related mRNA Stability - PubMed — https://pubmed.ncbi.nlm.nih.gov/32210752/
- **[S6]** PRDX1 activates autophagy via the PTEN-AKT signaling pathway to protect against cisplatin-induced spiral ganglion neuron damage - PubMed — https://pubmed.ncbi.nlm.nih.gov/33749526/
- **[S7]** Peroxiredoxin 1 inhibits streptozotocin-induced Alzheimer's disease-like pathology in hippocampal neuronal cells via the blocking of Ca2+/Calpain/Cdk5-mediated mitochondrial fragmentation - PubMed — https://pubmed.ncbi.nlm.nih.gov/38977865/
- **[S8]** The antioxidant enzyme Prdx1 controls neuronal differentiation by thiol-redox-dependent activation of GDE2 - PubMed — https://pubmed.ncbi.nlm.nih.gov/19766572/
- **[S9]** Peroxiredoxins in the central nervous system - PubMed — https://pubmed.ncbi.nlm.nih.gov/18084903/
- **[S10]** PRDX1 enhances cerebral ischemia-reperfusion injury through activation of TLR4-regulated inflammation and apoptosis - PubMed — https://pubmed.ncbi.nlm.nih.gov/31526567/
- **[S11]** PRDX1 gene — https://medlineplus.gov/download/genetics/gene/prdx1.pdf
- **[S12]** The antioxidant enzyme Prdx1 controls neuronal differentiation by thiol-redox-dependent activation of GDE2 - PubMed — https://pubmed.ncbi.nlm.nih.gov/19766572/
- **[S13]** PRDX1 enhances cerebral ischemia-reperfusion injury through activation of TLR4-regulated inflammation and apoptosis - PubMed — https://pubmed.ncbi.nlm.nih.gov/31526567/
- **[S14]** The antioxidant enzyme Prdx1 controls neuronal differentiation by thiol-redox-dependent activation of GDE2 - PubMed — https://pubmed.ncbi.nlm.nih.gov/19766572/
