## Supplementary material for "Understanding Substance Dependence: What Differentiates Addictive from Non-Addictive Drugs?": Folder for supplementary files: non-addictive-genes-claims-first100-021-phyhip-gene-protein-has-been-associated-with-regul-d13c43c0590b.md

# non-addictive-genes-claims-first100-021-d13c43c0590b

**Original claim:** PHYHIP gene/protein has been associated with regulatory or homeostatic biological processes in neural systems.

## Claim

PHYHIP gene/protein has been associated with regulatory or homeostatic biological processes in neural systems.

## Verdict

Verdict: Likely True

## Confidence

Medium

## Supporting evidence

PHYHIP is not a ubiquitous “housekeeping” gene with obvious non-neural dominance; rather, multiple authoritative resources describe it as brain-enriched and neuron-associated. NCBI summarizes human **PHYHIP** as a protein-coding gene that enables **protein tyrosine kinase binding**, is **involved in protein localization**, and shows **biased expression in brain**. The Human Protein Atlas likewise classifies PHYHIP as **tissue enriched in brain**, places it in a **“Brain - Synaptic signal transduction”** tissue-expression cluster and a **“Neurons - Mixed function”** brain-expression cluster, and reports enhancement in **excitatory and inhibitory neurons**. ([ncbi.nlm.nih.gov](https://www.ncbi.nlm.nih.gov/gene/9796)) [S1]

The earliest primary literature also supports a neural association. The original characterization identified PAHX-AP1/PHYHIP as a **brain-specific protein** interacting with **PHYH** (phytanoyl-CoA alpha-hydroxylase), a gene implicated in Refsum disease, a disorder with major neurologic manifestations. The authors explicitly linked the protein’s brain-specific expression pattern to neural deficits seen in that condition. This does not by itself prove function, but it establishes an early neural-system association. ([sciencedirect.com](https://www.sciencedirect.com/science/article/abs/pii/S0169328X99003046?utm_source=openai)) [S2]

More directly relevant to “regulatory” biology, a 2002 developmental neuroscience study examined PAHX-AP1/PHYHIP expression in rat retina and visual cortex during postnatal development and after **binocular visual deprivation**. Expression rose after eye opening, was maintained into adulthood, and **decreased after visual deprivation** in both retina and visual cortex. The authors concluded that visual stimulation is required to maintain expression and suggested involvement in the **developmental regulation** of photoreceptor function and visual-cortex plasticity. That is a clear association with an activity-dependent regulatory process in neural tissue, albeit in an animal model. ([sciencedirect.com](https://www.sciencedirect.com/science/article/abs/pii/S0736574802000175?utm_source=openai)) [S3]

A separate mouse study found that PAHX-AP1/PHYHIP binds **BAI1** (brain-specific angiogenesis inhibitor 1), with overlapping localization in neurons of the cerebral cortex, hippocampus, and selected cranial nuclei. In that paper, the authors linked the BAI1 system to **neuronal differentiation** and showed correlated developmental expression in brain. Because PHYHIP was a BAI1-binding partner with shared localization, this further associates PHYHIP with developmental regulatory programs in neural tissue. ([pubmed.ncbi.nlm.nih.gov](https://pubmed.ncbi.nlm.nih.gov/11245925/)) [S4]

The strongest mechanistic hint comes from the DYRK1A paper. In PC12 cells, PHYHIP was shown to bind the neural-development–relevant kinase **DYRK1A**, relocalize DYRK1A from the **nucleus to the cytoplasm**, and prevent its interaction with the nuclear transcription factor **CREB**. Since DYRK1A is well established as a regulator of neurodevelopment and cognition-related pathways, PHYHIP’s ability to alter DYRK1A localization and partner access is a plausible regulatory function in a neural context. ([pubmed.ncbi.nlm.nih.gov](https://pubmed.ncbi.nlm.nih.gov/15694837/)) [S5]

There are also more recent disease-context associations in neural tissue. An epigenome-wide association study of **human frontal cortex** identified a replicated differentially methylated locus near **PHYHIP** associated with **Braak Lewy body stage**. This is only correlative, but it supports the idea that PHYHIP sits in molecular neighborhoods relevant to neurodegenerative brain pathology. ([pubmed.ncbi.nlm.nih.gov](https://pubmed.ncbi.nlm.nih.gov/35995800/?utm_source=openai)) [S6]

Finally, a very recent preclinical paper reported **Phyhip-mediated active transport** across the injured blood-brain barrier in traumatic brain injury, with a Phyhip-targeted bFGF construct improving **neuronal survival**, **BBB integrity**, **neovascularization**, and **neuroinflammation** in rodents. This is emerging evidence and not yet a canonical function of endogenous PHYHIP, but it is consistent with possible involvement in neurovascular and homeostasis-related processes. ([pubmed.ncbi.nlm.nih.gov](https://pubmed.ncbi.nlm.nih.gov/40785555/)) [S7]

## Contradicting or limiting evidence

The main limitation is that PHYHIP is still **poorly characterized**. Even authoritative summaries remain conservative: NCBI’s current functional summary for human PHYHIP is essentially that it enables **protein tyrosine kinase binding** and is **involved in protein localization**; it does **not** describe PHYHIP as an established master regulator of neuronal homeostasis, synaptic homeostasis, excitability homeostasis, or glial homeostatic maintenance. ([ncbi.nlm.nih.gov](https://www.ncbi.nlm.nih.gov/gene/9796)) [S8]

Most of the functionally suggestive evidence is also **older and indirect**. The developmental retina/visual-cortex study, the BAI1-binding study, and the DYRK1A relocalization study are informative, but they are largely based on **rodent tissues, yeast-two-hybrid interactions, co-immunoprecipitation, overexpression, and cell-based localization assays** rather than definitive in vivo loss-of-function demonstrations that endogenous PHYHIP is required for a specific neural homeostatic program. ([sciencedirect.com](https://www.sciencedirect.com/science/article/abs/pii/S0736574802000175?utm_source=openai)) [S9]

The more recent human-brain disease links are mostly **associational**, not causal. For example, differential methylation near PHYHIP in Lewy body pathology shows relevance to diseased brain tissue, but does not establish whether PHYHIP drives, protects against, or merely tracks neuropathology. ([pubmed.ncbi.nlm.nih.gov](https://pubmed.ncbi.nlm.nih.gov/35995800/?utm_source=openai)) [S10]

Likewise, review-level discussion of a possible **PHYH/PHYHIP/DYRK1A** axis in Alzheimer disease is explicitly speculative. A recent review states that this pathway **“merits further investigation,”** which is essentially an acknowledgment that current evidence is hypothesis-generating rather than settled. ([pmc.ncbi.nlm.nih.gov](https://pmc.ncbi.nlm.nih.gov/articles/PMC10083964/?utm_source=openai)) [S11]

## Mechanistic / biological rationale

A plausible model is that PHYHIP acts less like a classic enzyme and more like an **intracellular interaction/localization protein**. That fits its formal annotations for kinase binding and protein localization, as well as experimental data showing interaction with **PHYH**, **BAI1**, and **DYRK1A**. If a brain-enriched protein alters where a signaling kinase such as DYRK1A resides inside the cell, it can change which substrates and transcriptional programs are accessible, which is a bona fide regulatory mechanism. ([ncbi.nlm.nih.gov](https://www.ncbi.nlm.nih.gov/gene/9796)) [S12]

Its brain enrichment and neuron-associated expression clusters make a neural role biologically plausible, and the Human Protein Atlas’ localization signal to **mitochondria** and **nucleoplasm** is consistent with participation in compartment-specific regulation. That said, inferring a direct role in “homeostasis” from localization alone would go too far; this is suggestive, not conclusive. ([proteinatlas.org](https://www.proteinatlas.org/ENSG00000168490-PHYHIP?utm_source=openai)) [S13]

The developmental visual-system data are especially relevant because they connect PHYHIP expression to **sensory experience** and **plasticity-related maintenance** of expression after eye opening. In other words, PHYHIP appears responsive to neural activity context, which is exactly the kind of pattern often seen in proteins involved in neural regulatory adaptation. Still, the literature has not yet shown that PHYHIP is a core homeostatic controller in the way that, for example, well-known synaptic-scaling or proteostasis factors are. ([sciencedirect.com](https://www.sciencedirect.com/science/article/abs/pii/S0736574802000175?utm_source=openai)) [S14]

## Bottom line

The claim is **supportable at the level of association**. PHYHIP is brain-enriched, neuron-associated, linked experimentally to **developmental regulation/plasticity** in retina and visual cortex, and able to modulate localization of the neural kinase **DYRK1A**, which supports a **regulatory role in neural systems**. ([proteinatlas.org](https://www.proteinatlas.org/ENSG00000168490-PHYHIP?utm_source=openai)) [S15]

However, evidence for a **well-established neural homeostatic function** is still limited. The field has more **expression, interaction, and correlative disease data** than definitive mechanistic proof from rigorous in vivo loss-of-function studies. So the most accurate summary is: **PHYHIP is likely associated with neural regulatory biology, but a specific, established role in neural homeostasis remains unproven.** ([ncbi.nlm.nih.gov](https://www.ncbi.nlm.nih.gov/gene/9796)) [S16]

## Cited sources

- **[S1]** PHYHIP phytanoyl-CoA 2-hydroxylase interacting protein [Homo sapiens (human)] - Gene - NCBI — https://www.ncbi.nlm.nih.gov/gene/9796
- **[S2]** Identification of a brain specific protein that associates with a Refsum disease gene product, phytanoyl-CoA alpha-hydroxylase - ScienceDirect — https://www.sciencedirect.com/science/article/abs/pii/S0169328X99003046?utm_source=openai
- **[S3]** Postnatal expression and distribution of Refsum disease gene associated protein in the rat retina and visual cortex: effect of binocular visual deprivation - ScienceDirect — https://www.sciencedirect.com/science/article/abs/pii/S0736574802000175?utm_source=openai
- **[S4]** Characterization of mouse brain-specific angiogenesis inhibitor 1 (BAI1) and phytanoyl-CoA alpha-hydroxylase-associated protein 1, a novel BAI1-binding protein - PubMed — https://pubmed.ncbi.nlm.nih.gov/11245925/
- **[S5]** Dual-specificity tyrosine-phosphorylated and regulated kinase 1A (DYRK1A) interacts with the phytanoyl-CoA alpha-hydroxylase associated protein 1 (PAHX-AP1), a brain specific protein - PubMed — https://pubmed.ncbi.nlm.nih.gov/15694837/
- **[S6]** Epigenome-wide association study of human frontal cortex identifies differential methylation in Lewy body pathology - PubMed — https://pubmed.ncbi.nlm.nih.gov/35995800/?utm_source=openai
- **[S7]** Phyhip-Targeted Delivery of bFGF Platform with Dual Modulation for Enhancing Neurovascular Repair in Traumatic Brain Injury - PubMed — https://pubmed.ncbi.nlm.nih.gov/40785555/
- **[S8]** PHYHIP phytanoyl-CoA 2-hydroxylase interacting protein [Homo sapiens (human)] - Gene - NCBI — https://www.ncbi.nlm.nih.gov/gene/9796
- **[S9]** Postnatal expression and distribution of Refsum disease gene associated protein in the rat retina and visual cortex: effect of binocular visual deprivation - ScienceDirect — https://www.sciencedirect.com/science/article/abs/pii/S0736574802000175?utm_source=openai
- **[S10]** Epigenome-wide association study of human frontal cortex identifies differential methylation in Lewy body pathology - PubMed — https://pubmed.ncbi.nlm.nih.gov/35995800/?utm_source=openai
- **[S11]** Exploring links between 2‐oxoglutarate‐dependent oxygenases and Alzheimer's disease - PMC — https://pmc.ncbi.nlm.nih.gov/articles/PMC10083964/?utm_source=openai
- **[S12]** PHYHIP phytanoyl-CoA 2-hydroxylase interacting protein [Homo sapiens (human)] - Gene - NCBI — https://www.ncbi.nlm.nih.gov/gene/9796
- **[S13]** PHYHIP protein expression summary - The Human Protein Atlas — https://www.proteinatlas.org/ENSG00000168490-PHYHIP?utm_source=openai
- **[S14]** Postnatal expression and distribution of Refsum disease gene associated protein in the rat retina and visual cortex: effect of binocular visual deprivation - ScienceDirect — https://www.sciencedirect.com/science/article/abs/pii/S0736574802000175?utm_source=openai
- **[S15]** PHYHIP protein expression summary - The Human Protein Atlas — https://www.proteinatlas.org/ENSG00000168490-PHYHIP?utm_source=openai
- **[S16]** PHYHIP phytanoyl-CoA 2-hydroxylase interacting protein [Homo sapiens (human)] - Gene - NCBI — https://www.ncbi.nlm.nih.gov/gene/9796
