## Supplementary material for "Understanding Substance Dependence: What Differentiates Addictive from Non-Addictive Drugs?": Folder for supplementary files: non-addictive-genes-claims-first100-022-prdx2-gene-protein-has-been-associated-with-regula-878e79a66a88.md

# non-addictive-genes-claims-first100-022-878e79a66a88

**Original claim:** PRDX2 gene/protein has been associated with regulatory or homeostatic biological processes in neural systems.

## Claim

PRDX2 gene/protein has been associated with regulatory or homeostatic biological processes in neural systems.

## Verdict

Verdict: Highly Supported

## Confidence

High

## Supporting evidence

PRDX2 is a well-established peroxide-detoxifying and redox-signaling protein, and curated reference resources describe it as a thiol-specific peroxidase that protects cells from oxidative stress and serves as a sensor of hydrogen-peroxide-mediated signaling. In the Human Protein Atlas, PRDX2 shows broad brain expression and is assigned to a brain RNA cluster labeled “Neurons & Synapses - Synaptic function,” supporting relevance to neural tissue rather than only non-neural biology. ([proteinatlas.org](https://www.proteinatlas.org/ENSG00000167815-PRDX2)) [S1]

Human brain histology also supports a neural association. In a classic immunohistochemical study of human brain, Prx-II/PRDX2 was reported as selectively neuronal across multiple regions including cerebral cortex, cerebellum, basal ganglia, substantia nigra, and spinal cord, with especially strong staining in large neurons such as hippocampal pyramidal neurons and Purkinje cells. The authors concluded that this restricted distribution likely reflects distinct functional roles in brain cell types. ([pubmed.ncbi.nlm.nih.gov](https://pubmed.ncbi.nlm.nih.gov/10494109/)) [S2]

A review focused on neurological disorders summarizes the field by stating that Prdx2 is an abundant antioxidant enzyme in the central nervous system, reduces reactive oxygen species, and participates in neuronal signaling-pathway regulation; importantly, the review also notes its dual nature in the injured brain. While reviews are secondary evidence, this one is consistent with the primary studies below. ([pubmed.ncbi.nlm.nih.gov](https://pubmed.ncbi.nlm.nih.gov/32002772/)) [S3]

There is strong mechanistic evidence from neuronal ischemia models that PRDX2 contributes to intracellular homeostasis and signaling control. In transgenic mice overexpressing PRX2, ischemic brain injury was reduced and neurological recovery improved after focal cerebral ischemia. In primary cortical neurons, PRX2 overexpression restored PRX and thioredoxin activity after oxygen-glucose deprivation, preserved Trx-ASK1 association, inhibited ASK1 activation, and reduced neuronal death. These findings directly tie PRDX2 to neuronal redox homeostasis and regulation of stress-response signaling. ([pubmed.ncbi.nlm.nih.gov](https://pubmed.ncbi.nlm.nih.gov/22356734/)) [S4]

A separate Parkinson-related study reached a similar conclusion in dopaminergic neurons. In vitro and in vivo 6-hydroxydopamine models showed that endogenous PRX2 was the most protective among tested 2-Cys peroxiredoxins, and PRX2 overexpression protected dopaminergic neurons while preserving dopamine-related motor function in mice. Mechanistically, PRX2 maintained thioredoxin in a reduced state and suppressed ASK1, JNK/c-Jun, and p38 pro-death pathways. That is a clear regulatory role in neuronal stress signaling, not merely a passive marker. ([pubmed.ncbi.nlm.nih.gov](https://pubmed.ncbi.nlm.nih.gov/21209210/)) [S5]

Human-relevant neuronal evidence also exists at the level of post-translational regulation. In dopaminergic neural cells and human iPSC-derived neurons, S-nitrosylation of Prx2 impaired its enzymatic and protective activity, while denitrosylation via sulfiredoxin reduced oxidative-stress vulnerability. The same study found markedly increased SNO-Prx2 in murine and human Parkinson’s disease brains, linking PRDX2 dysregulation to disturbed neuronal redox homeostasis in human disease tissue. ([pubmed.ncbi.nlm.nih.gov](https://pubmed.ncbi.nlm.nih.gov/27821734/)) [S6]

PRDX2 is also associated with glial homeostatic responses. In human multiple-sclerosis autopsy tissue, PRDX2 was upregulated mainly in astrocytes within white-matter lesions, and expression correlated positively with inflammation and oxidative stress. The authors interpreted this as suggesting a role in astrocyte resistance to oxidative damage, which fits a protective/homeostatic function in the CNS microenvironment. ([pubmed.ncbi.nlm.nih.gov](https://pubmed.ncbi.nlm.nih.gov/28375164/)) [S7]

More recent mechanistic work extends the association from redox balance to organelle homeostasis. In HT22 neuronal cells, Prx II preserved endoplasmic-reticulum/mitochondria contact sites, supported mitochondrial integrity, and helped maintain mitochondrial Ca2+ homeostasis, thereby reducing apoptosis under oxidative stress. This is directly relevant to neural homeostasis because mitochondrial Ca2+ control is central to neuronal survival and signaling. ([pubmed.ncbi.nlm.nih.gov](https://pubmed.ncbi.nlm.nih.gov/38637880/)) [S8]

## Contradicting or limiting evidence

The main limitation is context dependence. PRDX2 is not uniformly “homeostatic” in every neural context. When released extracellularly after brain injury, it can act as a damage-associated molecular pattern rather than a protective intracellular enzyme. After subarachnoid hemorrhage, extracellular Prx2 activated microglia through TLR4/MyD88/NF-κB signaling and promoted neuronal apoptosis; in a 2024 intracerebral hemorrhage model, Prx2 similarly drove brain swelling, microglial activation, neutrophil infiltration, neuronal death, and neurological deficits via TLR4. So PRDX2 can be protective inside neural cells yet harmful when spilled into the extracellular injury milieu. ([pubmed.ncbi.nlm.nih.gov](https://pubmed.ncbi.nlm.nih.gov/29554978/)) [S9]

Another limitation is that most causal evidence comes from cell culture and rodent injury models rather than human interventional studies. Human evidence is strongest for expression patterns, postmortem tissue changes, and lesion associations, not for direct clinical proof that PRDX2 is a master regulator of normal neural homeostasis in living humans. Thus, the evidence strongly supports association and plausible functional involvement, but it is less definitive for broad claims about indispensable physiologic control in the intact human nervous system. ([pubmed.ncbi.nlm.nih.gov](https://pubmed.ncbi.nlm.nih.gov/10494109/)) [S10]

## Mechanistic / biological rationale

Biologically, the claim makes sense. PRDX2 detoxifies hydrogen peroxide and organic hydroperoxides, and curated resources describe it not just as an antioxidant enzyme but also as a sensor/regulator of H2O2-mediated signaling. In neurons, where oxidative metabolism is high and redox imbalance can quickly trigger dysfunction or death, a protein that both removes peroxides and modulates peroxide-sensitive signaling is inherently positioned to support homeostatic regulation. ([proteinatlas.org](https://www.proteinatlas.org/ENSG00000167815-PRDX2)) [S11]

The best-supported neural mechanism is the PRDX2-thioredoxin-ASK1 axis. Under oxidative stress, PRDX2 helps preserve reduced thioredoxin, which keeps ASK1 restrained and limits downstream JNK/p38 stress signaling and apoptosis in neurons. That is a classic homeostatic buffering role: PRDX2 prevents redox perturbations from propagating into maladaptive cell-death signaling. ([pubmed.ncbi.nlm.nih.gov](https://pubmed.ncbi.nlm.nih.gov/21209210/)) [S12]

Additional studies indicate that PRDX2 function is regulated by post-translational modification and extends to organelle-level control. S-nitrosylation disables protective PRDX2 activity in neurons, while sulfiredoxin-mediated reversal restores resistance to oxidative stress; separately, Prx II supports ER-mitochondria contact sites and mitochondrial Ca2+ handling, linking redox control to mitochondrial homeostasis. Together, these findings place PRDX2 within a broader network that stabilizes neuronal physiology under stress. ([pubmed.ncbi.nlm.nih.gov](https://pubmed.ncbi.nlm.nih.gov/27821734/)) [S13]

At the same time, the extracellular DAMP behavior of PRDX2 after hemorrhagic injury shows that its biological effect depends on compartment and context: intracellular PRDX2 is largely homeostatic/protective, whereas extracellular PRDX2 can amplify neuroinflammation. ([pubmed.ncbi.nlm.nih.gov](https://pubmed.ncbi.nlm.nih.gov/29554978/)) [S14]

## Bottom line

Yes—this claim is **highly supported**. PRDX2 is strongly associated with **neural redox homeostasis**, **stress-response signaling regulation**, and, in newer work, **mitochondrial/Ca2+ homeostasis** in neuronal systems. Evidence comes from curated protein-function resources, human brain expression studies, human disease tissue, and multiple mechanistic neuron/brain models. The important nuance is that PRDX2 is **context dependent**: intracellularly it is usually protective and homeostatic, but extracellular PRDX2 released after CNS injury can become pro-inflammatory and damaging. ([proteinatlas.org](https://www.proteinatlas.org/ENSG00000167815-PRDX2)) [S15]

## Cited sources

- **[S1]** PRDX2 protein expression summary - The Human Protein Atlas — https://www.proteinatlas.org/ENSG00000167815-PRDX2
- **[S2]** Differential expression of peroxiredoxin subtypes in human brain cell types - PubMed — https://pubmed.ncbi.nlm.nih.gov/10494109/
- **[S3]** Effects of Peroxiredoxin 2 in Neurological Disorders: A Review of its Molecular Mechanisms - PubMed — https://pubmed.ncbi.nlm.nih.gov/32002772/
- **[S4]** Transgenic overexpression of peroxiredoxin-2 attenuates ischemic neuronal injury via suppression of a redox-sensitive pro-death signaling pathway - PubMed — https://pubmed.ncbi.nlm.nih.gov/22356734/
- **[S5]** Peroxiredoxin-2 protects against 6-hydroxydopamine-induced dopaminergic neurodegeneration via attenuation of the apoptosis signal-regulating kinase (ASK1) signaling cascade - PubMed — https://pubmed.ncbi.nlm.nih.gov/21209210/
- **[S6]** Role of sulfiredoxin as a peroxiredoxin-2 denitrosylase in human iPSC-derived dopaminergic neurons - PubMed — https://pubmed.ncbi.nlm.nih.gov/27821734/
- **[S7]** Expression of the Antioxidative Enzyme Peroxiredoxin 2 in Multiple Sclerosis Lesions in Relation to Inflammation - PubMed — https://pubmed.ncbi.nlm.nih.gov/28375164/
- **[S8]** Exploring the role of Prx II in mitigating endoplasmic reticulum stress and mitochondrial dysfunction in neurodegeneration - PubMed — https://pubmed.ncbi.nlm.nih.gov/38637880/
- **[S9]** Peroxiredoxin 2 activates microglia by interacting with Toll-like receptor 4 after subarachnoid hemorrhage - PubMed — https://pubmed.ncbi.nlm.nih.gov/29554978/
- **[S10]** Differential expression of peroxiredoxin subtypes in human brain cell types - PubMed — https://pubmed.ncbi.nlm.nih.gov/10494109/
- **[S11]** PRDX2 protein expression summary - The Human Protein Atlas — https://www.proteinatlas.org/ENSG00000167815-PRDX2
- **[S12]** Peroxiredoxin-2 protects against 6-hydroxydopamine-induced dopaminergic neurodegeneration via attenuation of the apoptosis signal-regulating kinase (ASK1) signaling cascade - PubMed — https://pubmed.ncbi.nlm.nih.gov/21209210/
- **[S13]** Role of sulfiredoxin as a peroxiredoxin-2 denitrosylase in human iPSC-derived dopaminergic neurons - PubMed — https://pubmed.ncbi.nlm.nih.gov/27821734/
- **[S14]** Peroxiredoxin 2 activates microglia by interacting with Toll-like receptor 4 after subarachnoid hemorrhage - PubMed — https://pubmed.ncbi.nlm.nih.gov/29554978/
- **[S15]** PRDX2 protein expression summary - The Human Protein Atlas — https://www.proteinatlas.org/ENSG00000167815-PRDX2
