## Supplementary material for "Understanding Substance Dependence: What Differentiates Addictive from Non-Addictive Drugs?": Folder for supplementary files: non-addictive-genes-claims-first100-023-s100a6-gene-protein-has-been-associated-with-regul-a9a2fef63910.md

# non-addictive-genes-claims-first100-023-a9a2fef63910

**Original claim:** S100A6 gene/protein has been associated with regulatory or homeostatic biological processes in neural systems.

## Claim

S100A6 gene/protein has been associated with regulatory or homeostatic biological processes in neural systems. ([pmc.ncbi.nlm.nih.gov](https://pmc.ncbi.nlm.nih.gov/articles/PMC7313082/)) [S1]

## Verdict

Verdict: Likely True

The literature supports an association between S100A6 and neural regulatory/homeostatic functions, especially in **astrocyte biology, neural stem-cell states, neuronal morphogenesis/proteostasis, and metal-ion stress responses**. However, much of the evidence is **preclinical or pathology-linked**, and direct proof that S100A6 is a core homeostatic regulator in healthy adult human neural circuits remains limited. ([pmc.ncbi.nlm.nih.gov](https://pmc.ncbi.nlm.nih.gov/articles/PMC12518605/)) [S2]

## Confidence

Medium. Multiple independent studies and reviews place S100A6 in neural cell populations and regulatory pathways, but the strongest causal evidence is still mainly from **mouse, cell-culture, and disease-model** work rather than definitive human functional studies. ([pmc.ncbi.nlm.nih.gov](https://pmc.ncbi.nlm.nih.gov/articles/PMC7313082/)) [S3]

## Supporting evidence

S100A6 is clearly **present in neural systems**, including neurons, astrocytes, ependymal cells, Schwann cells, hypothalamic tanycytes, and neurogenic niches. Reviews summarizing immunohistochemical and transcriptomic work report S100A6 expression in hippocampus, cortex, thalamus, brainstem, cerebellum, and ventricular border regions; human brain staining has also been reported in cortex and hippocampus. This establishes biological plausibility for neural roles rather than a purely peripheral function. ([pmc.ncbi.nlm.nih.gov](https://pmc.ncbi.nlm.nih.gov/articles/PMC7313082/)) [S4]

Evidence for a **regulatory role in neural development and neuron–glia signaling** became substantially stronger with a 2025 study showing that astrocytes secrete S100A6 during corticogenesis, that neurons contain the partner protein CaCyBp, and that extracellular S100A6 reduces neurite complexity by altering neuronal protein-turnover pathways. The authors concluded that S100A6-CaCyBp signaling modulates neuronal morphogenesis and proteostasis, effectively placing S100A6 in a bona fide intercellular regulatory pathway in the developing mouse brain. ([pmc.ncbi.nlm.nih.gov](https://pmc.ncbi.nlm.nih.gov/articles/PMC12518605/)) [S5]

S100A6 is also associated with **neural stem-cell quiescence and gliogenic regulation**. In the adult mouse hippocampal subgranular zone, S100A6 was highly expressed in neural stem cells and astrocyte precursors, and pharmacologic inhibition of S100 proteins reduced production of S100β-positive astrocyte-lineage cells without affecting neurogenesis. Reviews further note that single-cell RNA-seq linked S100A6 to the most quiescent adult stem-cell subpopulations, including neural stem/progenitor compartments. That pattern is consistent with a role in regulating stem-cell state and local tissue maintenance. ([pubmed.ncbi.nlm.nih.gov](https://pubmed.ncbi.nlm.nih.gov/24115312/)) [S6]

Several studies connect S100A6 to **stress-response and injury-response homeostasis** in neural tissue. In mouse models, astrocytic S100A6 rises with glutamate-related injury paradigms and epilepsy-like insults, and its increase can be attenuated by interventions that reduce excitotoxic signaling. In a rat traumatic brain injury model, early S100A6 reduction was followed by recovery toward baseline, with the authors linking later increases to cognitive improvement and possible regeneration/repair. These data do not prove a protective role, but they do support involvement in neural responses that attempt to restore tissue equilibrium after insult. ([pubmed.ncbi.nlm.nih.gov](https://pubmed.ncbi.nlm.nih.gov/22982625/)) [S7]

There is also evidence for **metal-ion homeostasis**, especially zinc-related regulation in neurodegenerative contexts. In Alzheimer-like mouse models, S100A6 is upregulated in astrocytes around amyloid plaques; zinc loading increases both plaque burden and S100A6 expression, while chelation reverses those changes. Recombinant S100A6 reduced plaque burden and lowered plaque-associated zinc in brain sections, supporting the idea that S100A6 can act as a zinc-sequestering factor in the diseased brain microenvironment. Because Zn²⁺ dyshomeostasis is a recognized feature of amyloid pathology, this is a fairly direct link to a homeostatic process in neural tissue. ([pubmed.ncbi.nlm.nih.gov](https://pubmed.ncbi.nlm.nih.gov/31440382/)) [S8]

Human relevance is supported, though less strongly, by neuropathology studies. S100A6 overexpression has been reported in astrocytes near impaired axons in both ALS-model mice and human sporadic ALS tissue, and reviews summarize altered S100A6 expression in AD, ALS, epilepsy, and aging-related brain changes. This supports association with neural regulatory/stress pathways, though it does not by itself establish whether S100A6 is compensatory, pathogenic, or both depending on context. ([pubmed.ncbi.nlm.nih.gov](https://pubmed.ncbi.nlm.nih.gov/12152788/)) [S9]

## Contradicting or limiting evidence

The biggest limitation is that **association is stronger than proof of indispensable function**. Much of the literature shows where S100A6 is expressed, or that it changes with injury, aging, epilepsy, AD, or ALS, but such data do not automatically establish a homeostatic role. Reviews explicitly state that the field is still far from defining S100A6’s exact role in the brain. ([pmc.ncbi.nlm.nih.gov](https://pmc.ncbi.nlm.nih.gov/articles/PMC7313082/)) [S10]

Directionality is also uncertain in several disease settings. For example, in ALS tissue the original authors observed selective astrocytic induction of S100A6 but stated that their findings did not determine whether its role was beneficial or detrimental. Likewise, glutamate-toxicity and epilepsy studies show S100A6 upregulation in activated astrocytes, but that could reflect a downstream stress marker rather than a primary regulator. ([pubmed.ncbi.nlm.nih.gov](https://pubmed.ncbi.nlm.nih.gov/12152788/)) [S11]

Finally, direct **human functional** evidence is sparse. The strongest mechanistic studies are in mouse development, rodent injury models, and ex vivo or in vitro systems. So while the claim of “associated with” neural regulatory/homeostatic processes is supported, a stronger claim such as “established key homeostatic regulator in human neural systems” would overstate the evidence. ([pmc.ncbi.nlm.nih.gov](https://pmc.ncbi.nlm.nih.gov/articles/PMC12518605/)) [S12]

## Mechanistic / biological rationale

S100A6 is an **EF-hand Ca²⁺/Zn²⁺-binding protein**, which already makes a regulatory role biologically plausible because calcium and zinc are central to neural signaling, stress responses, and proteostasis. Reviews describe S100A6 as interacting with cytoskeletal proteins, annexins, CacyBP/SIP, and Sgt1, implying roles in membrane dynamics, cytoskeletal organization, signaling, and protein quality-control pathways. ([pmc.ncbi.nlm.nih.gov](https://pmc.ncbi.nlm.nih.gov/articles/PMC9866648/)) [S13]

In neural systems, one plausible mechanism is **astrocyte-to-neuron signaling**. The 2025 developmental study showed that astrocyte-derived S100A6 can act on neuronal CaCyBp-linked pathways, downregulating unfolded-protein-response/ER-stress components and shifting ubiquitin-proteasome activity, which slows protein turnover and limits neuritogenesis. That is a classic regulatory/homeostatic type of mechanism: a glial signal constrains neuronal growth programs and protein-handling state. ([pmc.ncbi.nlm.nih.gov](https://pmc.ncbi.nlm.nih.gov/articles/PMC12518605/)) [S14]

A second plausible mechanism is **local ion buffering/stress adaptation**. In AD-like models, S100A6 appears capable of competing for zinc and reducing zinc-associated amyloid aggregation, suggesting a homeostatic role under conditions of metal-ion imbalance. More broadly, S100A6 changes with chronic stress, excitotoxicity, and injury, which fits a model in which it participates in cellular attempts to re-establish ionic and proteostatic balance. ([pubmed.ncbi.nlm.nih.gov](https://pubmed.ncbi.nlm.nih.gov/31440382/)) [S15]

A third is **stem-cell-state regulation**. Its enrichment in quiescent neural stem cells and astrocyte precursors suggests S100A6 may help govern the balance between quiescence, activation, and gliogenic differentiation in neurogenic niches. That is not definitive proof of a homeostatic function, but it is closely aligned with tissue-maintenance biology. ([pubmed.ncbi.nlm.nih.gov](https://pubmed.ncbi.nlm.nih.gov/24115312/)) [S16]

## Bottom line

The claim is **likely true**. S100A6 is not just incidentally present in the nervous system; it has been linked by multiple studies to **astrocyte-mediated regulation, neural stem-cell quiescence/gliogenesis, neuronal proteostasis and morphogenesis, and zinc-related stress responses in neurodegeneration**. The caveat is that much of this evidence is **preclinical and context-dependent**, so S100A6 should be described as **associated with** neural regulatory/homeostatic processes rather than definitively established as a central homeostatic controller in humans. ([pmc.ncbi.nlm.nih.gov](https://pmc.ncbi.nlm.nih.gov/articles/PMC12518605/)) [S17]

## Cited sources

- **[S1]** 
            S100A6 and Its Brain Ligands in Neurodegenerative Disorders - PMC
         — https://pmc.ncbi.nlm.nih.gov/articles/PMC7313082/
- **[S2]** 
            Astrocytes modulate neuronal development by S100A6 signaling - PMC
         — https://pmc.ncbi.nlm.nih.gov/articles/PMC12518605/
- **[S3]** 
            S100A6 and Its Brain Ligands in Neurodegenerative Disorders - PMC
         — https://pmc.ncbi.nlm.nih.gov/articles/PMC7313082/
- **[S4]** 
            S100A6 and Its Brain Ligands in Neurodegenerative Disorders - PMC
         — https://pmc.ncbi.nlm.nih.gov/articles/PMC7313082/
- **[S5]** 
            Astrocytes modulate neuronal development by S100A6 signaling - PMC
         — https://pmc.ncbi.nlm.nih.gov/articles/PMC12518605/
- **[S6]** S100A6 (calcyclin) is a novel marker of neural stem cells and astrocyte precursors in the subgranular zone of the adult mouse hippocampus - PubMed — https://pubmed.ncbi.nlm.nih.gov/24115312/
- **[S7]** Upregulation of calcium binding protein, S100A6, in activated astrocytes is linked to glutamate toxicity - PubMed — https://pubmed.ncbi.nlm.nih.gov/22982625/
- **[S8]** Glial S100A6 Degrades β-amyloid Aggregation through Targeting Competition with Zinc Ions - PubMed — https://pubmed.ncbi.nlm.nih.gov/31440382/
- **[S9]** S100A6 overexpression within astrocytes associated with impaired axons from both ALS mouse model and human patients - PubMed — https://pubmed.ncbi.nlm.nih.gov/12152788/
- **[S10]** 
            S100A6 and Its Brain Ligands in Neurodegenerative Disorders - PMC
         — https://pmc.ncbi.nlm.nih.gov/articles/PMC7313082/
- **[S11]** S100A6 overexpression within astrocytes associated with impaired axons from both ALS mouse model and human patients - PubMed — https://pubmed.ncbi.nlm.nih.gov/12152788/
- **[S12]** 
            Astrocytes modulate neuronal development by S100A6 signaling - PMC
         — https://pmc.ncbi.nlm.nih.gov/articles/PMC12518605/
- **[S13]** 
            S100A6 Protein—Expression and Function in Norm and Pathology - PMC
         — https://pmc.ncbi.nlm.nih.gov/articles/PMC9866648/
- **[S14]** 
            Astrocytes modulate neuronal development by S100A6 signaling - PMC
         — https://pmc.ncbi.nlm.nih.gov/articles/PMC12518605/
- **[S15]** Glial S100A6 Degrades β-amyloid Aggregation through Targeting Competition with Zinc Ions - PubMed — https://pubmed.ncbi.nlm.nih.gov/31440382/
- **[S16]** S100A6 (calcyclin) is a novel marker of neural stem cells and astrocyte precursors in the subgranular zone of the adult mouse hippocampus - PubMed — https://pubmed.ncbi.nlm.nih.gov/24115312/
- **[S17]** 
            Astrocytes modulate neuronal development by S100A6 signaling - PMC
         — https://pmc.ncbi.nlm.nih.gov/articles/PMC12518605/
