## Supplementary material for "Understanding Substance Dependence: What Differentiates Addictive from Non-Addictive Drugs?": Folder for supplementary files: non-addictive-genes-claims-first100-024-prkacb-gene-protein-has-been-associated-with-regul-cc8f16c93f71.md

# non-addictive-genes-claims-first100-024-cc8f16c93f71

**Original claim:** PRKACB gene/protein has been associated with regulatory or homeostatic biological processes in neural systems.

## Claim

PRKACB gene/protein has been associated with regulatory or homeostatic biological processes in neural systems.

## Verdict

Verdict: Likely True

The claim is supported for **neural regulatory processes** and only **partially/directly supported** for **neural homeostatic processes**. PRKACB encodes the catalytic β subunit of PKA, is enriched in brain tissue, shows regionally patterned expression in the CNS, and has direct experimental links to synaptic plasticity, neurotransmitter-release control, and neural development. Evidence that PRKACB specifically mediates bona fide **homeostatic** neural regulation exists, but is more indirect and less isoform-specific than the evidence for general neural regulatory roles. ([proteinatlas.org](https://www.proteinatlas.org/ENSG00000142875-PRKACB/tissue)) [S1]

## Confidence

Medium

## Supporting evidence

PRKACB is clearly relevant to the nervous system at the expression level. In the Human Protein Atlas, PRKACB is annotated as **tissue enhanced in brain**, and its RNA-expression cluster is labeled **“Brain - Synaptic signal transduction (mainly)”**. In mouse brain, classic in situ work found that **Cβ mRNA has a distinct regional pattern**, including relatively higher expression in the dentate gyrus than in CA1-3, implying neural-tissue specialization rather than merely ubiquitous housekeeping expression. ([proteinatlas.org](https://www.proteinatlas.org/ENSG00000142875-PRKACB/tissue)) [S2]

There is strong **direct genetic evidence** that Prkacb participates in neural regulatory processes governing synaptic plasticity. In mice lacking the **Cβ1** catalytic isoform, high-frequency stimulation still induced initial potentiation at Schaffer collateral-CA1 synapses, but the mutants failed to maintain normal late-phase LTP; the late potentiated response was reported as only about **30% of control**. The same mutants also failed to show lasting LTD and depotentiation. That is a direct link between Prkacb and regulation of long-term synaptic efficacy in the hippocampus. ([pmc.ncbi.nlm.nih.gov](https://pmc.ncbi.nlm.nih.gov/articles/PMC39982/?utm_source=openai)) [S3]

PRKACB is also linked to **activity-dependent regulation of neurotransmitter release**. In adult rat neuromuscular junction, a 2023 study reported that the **synaptic PKA Cβ subunit** controls activity-dependent phosphorylation of **SNAP-25** and **Synapsin-1**, and concluded that this retrograde signaling helps **balance acetylcholine release** during nerve-muscle communication. Although the neuromuscular junction is peripheral rather than CNS, it is still a neural synapse, and the paper provides isoform-specific evidence for PRKACB/Cβ in synaptic regulation. ([pubmed.ncbi.nlm.nih.gov](https://pubmed.ncbi.nlm.nih.gov/36869288/)) [S4]

There is also evidence for PRKACB in **developmental neural regulation**. Alliance/MGI experimental GO annotations link mouse Prkacb to **negative regulation of smoothened signaling involved in dorsal/ventral neural tube patterning** and to **neural tube closure**. The NCBI mouse gene summary similarly states that Prkacb acts upstream of or within **neural tube development** and is expressed in the **nervous system**. These are regulatory biological processes squarely within neural development. ([informatics.jax.org](https://www.informatics.jax.org/homology/GOGraph/Prkacb)) [S5]

For the **homeostatic** part of the claim, the evidence is suggestive rather than definitive. In a proteomic study of **homeostatic scaling at central synapses**, Prkacb abundance changed bidirectionally with chronic activity manipulations: it increased after **24 h TTX** and decreased after **2 h bicuculline**, alongside proteins important for synaptic transmission and plasticity. A complementary phosphoproteomic study in cortical neurons showed that thousands of phosphorylation events are regulated during **homeostatic up- and down-scaling**, with persistent reciprocal regulation pointing to common signaling axes in synaptic homeostasis. Taken together, these papers place Prkacb/PKA-associated signaling within the molecular landscape of neural homeostatic plasticity, even if they do not prove PRKACB alone is the essential driver. ([pmc.ncbi.nlm.nih.gov](https://pmc.ncbi.nlm.nih.gov/articles/PMC5814146/)) [S6]

## Contradicting or limiting evidence

The main limitation is that **homeostatic evidence is not as PRKACB-specific** as the synaptic-plasticity evidence. The strongest causal paper is the 1996 Cβ1 knockout study, but it addresses **LTP/LTD/depotentiation**, i.e. plasticity and regulation, not classical homeostatic set-point control. The homeostatic-scaling evidence comes mainly from proteomics and pathway-level inference, where PRKACB is one changing component among many. ([pmc.ncbi.nlm.nih.gov](https://pmc.ncbi.nlm.nih.gov/articles/PMC39982/?utm_source=openai)) [S7]

A second limitation is that much of the neural literature is on **PKA broadly**, not PRKACB specifically. For example, studies of neuronal excitability and synaptic downscaling often implicate **PKA-dependent signaling** without resolving which catalytic isoform is responsible. So the inference from “PKA is involved” to “PRKACB specifically is involved” is reasonable but not always directly demonstrated. ([embopress.org](https://www.embopress.org/doi/abs/10.15252/embj.2022110963)) [S8]

Human genetics also does not strongly anchor PRKACB as a major neural-homeostasis gene. Reported human PRKACB variant papers have focused mainly on **skeletal disease** and **adrenocortical hyperplasia/endocrine phenotypes**, not primary neurological syndromes. That does not argue against a neural role, but it limits confidence that PRKACB has a well-established, clinically validated homeostatic role in the human nervous system. ([pubmed.ncbi.nlm.nih.gov](https://pubmed.ncbi.nlm.nih.gov/33055300/?utm_source=openai)) [S9]

## Mechanistic / biological rationale

Mechanistically, the claim is biologically plausible. PRKACB encodes a **cAMP-dependent protein kinase catalytic subunit (Cβ)**, and PKA is a core transducer of cAMP signals in neurons. In brain tissue, PRKACB is one of the catalytic isoforms contributing to PKA signaling, and its patterned neural expression suggests isoform-specific signaling niches. ([informatics.jax.org](https://www.informatics.jax.org/homology/GOGraph/Prkacb)) [S10]

That fits well with the observed phenotypes. PKA signaling is well positioned to regulate **synaptic strength, vesicle release, and activity-dependent phosphorylation** of neuronal substrates. The NMJ study directly ties Cβ to phosphorylation of **SNAP-25** and **Synapsin-1**, which are presynaptic regulators of vesicle handling and transmitter output. The hippocampal knockout study shows that losing a Prkacb isoform disrupts the maintenance and bidirectional adjustment of synaptic efficacy, which is exactly the kind of regulatory architecture expected from a cAMP/PKA catalytic subunit. ([pubmed.ncbi.nlm.nih.gov](https://pubmed.ncbi.nlm.nih.gov/36869288/)) [S11]

For homeostasis, the rationale is that cAMP/PKA signaling can participate in negative-feedback stabilization of synapses and neuronal excitability. The homeostatic-scaling studies do not prove PRKACB is the unique mediator, but the fact that Prkacb abundance changes with chronic TTX/bicuculline manipulations is consistent with a role in adaptive set-point regulation. ([pmc.ncbi.nlm.nih.gov](https://pmc.ncbi.nlm.nih.gov/articles/PMC5814146/)) [S12]

## Bottom line

Overall, the statement is **better supported than not**. PRKACB has **clear associations with neural regulatory processes**—especially synaptic plasticity, neurotransmitter-release control, patterned brain expression, and neural development. The evidence for **neural homeostatic** roles is **present but less direct**, relying more on proteomic association and broader PKA-pathway biology than on definitive PRKACB-specific causal experiments. So the most accurate overall judgment is **Likely True**. ([pmc.ncbi.nlm.nih.gov](https://pmc.ncbi.nlm.nih.gov/articles/PMC39982/?utm_source=openai)) [S13]

## Cited sources

- **[S1]** Tissue expression of PRKACB - Summary - The Human Protein Atlas — https://www.proteinatlas.org/ENSG00000142875-PRKACB/tissue
- **[S2]** Tissue expression of PRKACB - Summary - The Human Protein Atlas — https://www.proteinatlas.org/ENSG00000142875-PRKACB/tissue
- **[S3]** Impaired hippocampal plasticity in mice lacking the Cbeta1 catalytic subunit of cAMP-dependent protein kinase - PMC — https://pmc.ncbi.nlm.nih.gov/articles/PMC39982/?utm_source=openai
- **[S4]** Synaptic retrograde regulation of the PKA-induced SNAP-25 and Synapsin-1 phosphorylation - PubMed — https://pubmed.ncbi.nlm.nih.gov/36869288/
- **[S5]** Comparative GO Graph (mouse, human, rat, zebrafish) — https://www.informatics.jax.org/homology/GOGraph/Prkacb
- **[S6]** 
            Time- and polarity-dependent proteomic changes associated with homeostatic scaling at central synapses - PMC
         — https://pmc.ncbi.nlm.nih.gov/articles/PMC5814146/
- **[S7]** Impaired hippocampal plasticity in mice lacking the Cbeta1 catalytic subunit of cAMP-dependent protein kinase - PMC — https://pmc.ncbi.nlm.nih.gov/articles/PMC39982/?utm_source=openai
- **[S8]** Autophagy regulates neuronal excitability by controlling cAMP/protein kinase A signaling at the synapse | The EMBO Journal | Springer Nature Link — https://www.embopress.org/doi/abs/10.15252/embj.2022110963
- **[S9]** PRKACB variants in skeletal disease or adrenocortical hyperplasia: effects on protein kinase A - PubMed — https://pubmed.ncbi.nlm.nih.gov/33055300/?utm_source=openai
- **[S10]** Comparative GO Graph (mouse, human, rat, zebrafish) — https://www.informatics.jax.org/homology/GOGraph/Prkacb
- **[S11]** Synaptic retrograde regulation of the PKA-induced SNAP-25 and Synapsin-1 phosphorylation - PubMed — https://pubmed.ncbi.nlm.nih.gov/36869288/
- **[S12]** 
            Time- and polarity-dependent proteomic changes associated with homeostatic scaling at central synapses - PMC
         — https://pmc.ncbi.nlm.nih.gov/articles/PMC5814146/
- **[S13]** Impaired hippocampal plasticity in mice lacking the Cbeta1 catalytic subunit of cAMP-dependent protein kinase - PMC — https://pmc.ncbi.nlm.nih.gov/articles/PMC39982/?utm_source=openai
