## Supplementary material for "Understanding Substance Dependence: What Differentiates Addictive from Non-Addictive Drugs?": Folder for supplementary files: non-addictive-genes-claims-first100-025-caly-gene-protein-has-been-associated-with-regulat-0eb7d70eedcf.md

# non-addictive-genes-claims-first100-025-0eb7d70eedcf

**Original claim:** CALY gene/protein has been associated with regulatory or homeostatic biological processes in neural systems.

## Claim

CALY gene/protein has been associated with regulatory or homeostatic biological processes in neural systems.

## Verdict

Verdict: Likely True

The claim is well supported **for neural regulatory processes** such as receptor trafficking, endocytosis, vesicle sorting, activity-dependent synaptic plasticity, and synaptic reorganization. The evidence is **less direct for “homeostasis” in the narrow sense** of canonical neural homeostatic plasticity or baseline network set-point control, but some reviews and mechanistic data make that interpretation plausible. ([ncbi.nlm.nih.gov](https://www.ncbi.nlm.nih.gov/gene/50632)) [S1]

## Confidence

High

## Supporting evidence

CALY encodes **calcyon**, a neuron-enriched transmembrane vesicular/endosomal protein with clear localization to neural compartments relevant to regulation of synaptic signaling. NCBI curates CALY as a brain-biased gene whose protein is required for maximal stimulated calcium release after purinergic or muscarinic receptor activation and interacts with the D1 dopamine receptor. The same record includes Gene Ontology annotations placing CALY in the **postsynaptic endocytic zone**, **glutamatergic synapse**, and processes including **positive regulation of endocytosis** and **postsynaptic neurotransmitter receptor internalization**. Independent expression data also show neuronal staining in human brain tissue, and macaque in situ hybridization found calcyon mRNA broadly distributed throughout the primate brain. ([ncbi.nlm.nih.gov](https://www.ncbi.nlm.nih.gov/gene/50632)) [S2]

A strong mechanistic foundation comes from the original biochemical/cell-biological work showing that calcyon is a **clathrin light chain-binding protein**. In that study, calcyon stimulated clathrin self-assembly, localized to pre- and postsynaptic vesicular compartments, and enhanced transferrin uptake when overexpressed; conversely, transferrin uptake was largely abolished in neurons from calcyon-null mice while recycling was preserved. Those findings directly support a role in **clathrin-mediated endocytosis**, which is a core regulatory process controlling receptor availability and synaptic signaling. ([pubmed.ncbi.nlm.nih.gov](https://pubmed.ncbi.nlm.nih.gov/16595675/)) [S3]

The most direct neural functional evidence comes from hippocampal studies of AMPA receptor trafficking and LTD. In calcyon knockout mice, deletion of the gene impaired agonist-stimulated internalization of GluR1/GluR2-containing AMPA receptors without changing basal surface levels, and it abolished or markedly impaired chemical and electrically induced LTD in CA1 neurons; re-expression of calcyon rescued LTD. The authors concluded that calcyon has a specialized role in **regulating activity-dependent removal of synaptic AMPA receptors**. Because AMPAR trafficking is a central regulator of synaptic strength, this is strong evidence that CALY is associated with regulatory processes in neural systems. ([pmc.ncbi.nlm.nih.gov](https://pmc.ncbi.nlm.nih.gov/articles/PMC2771427/)) [S4]

Additional work extends calcyon’s regulatory role beyond postsynaptic AMPAR internalization. A Journal of Neurochemistry study showed that calcyon interacts with AP-3 adaptor-complex subunits and regulates targeting of AP-3 cargoes such as **ZnT3** and **PI4KIIα** in hippocampal mossy fibers, indicating a role in **vesicle/cargo sorting** in neurons. Another study identified calcyon as a dynein-associated endosomal protein that regulates vesicle sorting and axonal motility, again pointing to active regulation of neuronal membrane traffic rather than passive expression alone. ([pubmed.ncbi.nlm.nih.gov](https://pubmed.ncbi.nlm.nih.gov/22650988/)) [S5]

Calcyon also appears to regulate signaling systems important for neural development and circuit function. One study found that calcyon is a key determinant of **neuregulin-1 ectodomain cleavage and signaling**, and that calcyon overexpression enhanced **GABA transmission** in mice. Another showed that phosphorylation-dependent association of calcyon with PSD-95 promotes **dopamine D1 receptor internalization**. These findings place calcyon within regulatory pathways affecting dopaminergic and glutamatergic/GABAergic signaling balance. ([pubmed.ncbi.nlm.nih.gov](https://pubmed.ncbi.nlm.nih.gov/25349163/)) [S6]

Evidence also exists for a role in structural/systems-level neural adaptation. In a brain-injury model, the hevin-calcyon interaction was shown to be crucial for **synaptic and functional recovery** in sensory-motor cortex, and the authors described direct evidence for a causal role in **synaptic reorganization after brain injury**. Separately, inducible calcyon overexpression during adolescence altered adult working memory and response inhibition, implying that abnormal calcyon levels can perturb maturation of neural circuits. Together these data support association with broader regulatory processes governing circuit remodeling and recovery. ([pubmed.ncbi.nlm.nih.gov](https://pubmed.ncbi.nlm.nih.gov/33753902/)) [S7]

## Contradicting or limiting evidence

The main limitation is that **“homeostatic” is only partially established**. The strongest evidence for CALY concerns receptor internalization, endocytosis, cargo sorting, synaptic plasticity, and post-injury remodeling. Those are unquestionably regulatory neural processes, but they are not identical to classic **homeostatic plasticity** or direct maintenance of firing-rate/network set points. A review of NEEP21/calcyon argues that balancing their expression or activity **could** be important for signaling-pathway homeostasis, but that statement is a synthesis/proposal rather than definitive proof from a dedicated homeostasis experiment. ([pubmed.ncbi.nlm.nih.gov](https://pubmed.ncbi.nlm.nih.gov/25376768/)) [S8]

Another limitation is that much of the causal evidence comes from **cell systems and rodent models**, not direct human physiological experiments. Human evidence is more indirect, consisting mainly of expression data and candidate-gene association studies. Those human genetic associations are mixed: one family-based study reported association with ADHD-related traits, whereas another study found that tested CALCYON variants did **not** play a major role in schizophrenia or cocaine dependence. These findings suggest clinical relevance is plausible but not settled, and disease association does not itself prove a normal homeostatic role. ([pubmed.ncbi.nlm.nih.gov](https://pubmed.ncbi.nlm.nih.gov/16172615/)) [S9]

## Mechanistic / biological rationale

Biologically, the claim makes sense because calcyon sits at the intersection of several processes that neurons use to regulate and stabilize signaling: **endocytosis, receptor internalization, endosomal sorting, vesicle transport, and activity-dependent synaptic remodeling**. Calcyon binds clathrin light chain, is annotated to the postsynaptic endocytic zone, regulates AP-3-dependent cargo targeting, associates with dynein-based trafficking machinery, and modulates internalization of AMPARs and D1 receptors. These are classic mechanisms by which neurons tune receptor surface abundance, neurotransmission, and synaptic efficacy. ([pubmed.ncbi.nlm.nih.gov](https://pubmed.ncbi.nlm.nih.gov/16595675/)) [S10]

That said, the **homeostatic** interpretation is mostly an inference from these trafficking roles. If a protein regulates receptor removal, cargo routing, and recovery-associated synaptic reorganization, it is reasonable to infer that it could contribute to neural homeostasis. But the literature located here supports that more strongly as a **plausible mechanistic extension** than as a directly demonstrated canonical homeostatic program. ([pubmed.ncbi.nlm.nih.gov](https://pubmed.ncbi.nlm.nih.gov/25376768/)) [S11]

## Bottom line

Overall, the claim is **likely true**. CALY/calcyon is convincingly associated with **regulatory biological processes in neural systems**, especially clathrin-mediated endocytosis, postsynaptic receptor internalization, synaptic plasticity, vesicle sorting, and synaptic reorganization. ([pubmed.ncbi.nlm.nih.gov](https://pubmed.ncbi.nlm.nih.gov/16595675/)) [S12]

If the intended meaning is specifically that CALY is a proven mediator of **neural homeostasis** in the strict sense, the evidence is weaker and more indirect. But for the broader wording you used—“regulatory or homeostatic biological processes in neural systems”—the literature supports the statement, with the strongest support on the **regulatory** side. ([pubmed.ncbi.nlm.nih.gov](https://pubmed.ncbi.nlm.nih.gov/25376768/)) [S13]

## Cited sources

- **[S1]** CALY calcyon neuron specific vesicular protein [Homo sapiens (human)] - Gene - NCBI — https://www.ncbi.nlm.nih.gov/gene/50632
- **[S2]** CALY calcyon neuron specific vesicular protein [Homo sapiens (human)] - Gene - NCBI — https://www.ncbi.nlm.nih.gov/gene/50632
- **[S3]** Calcyon, a novel partner of clathrin light chain, stimulates clathrin-mediated endocytosis - PubMed — https://pubmed.ncbi.nlm.nih.gov/16595675/
- **[S4]** 
            Calcyon is Necessary for Activity Dependent AMPA Receptor Internalization and LTD in CA1 Neurons of Hippocampus - PMC
         — https://pmc.ncbi.nlm.nih.gov/articles/PMC2771427/
- **[S5]** Calcyon, a mammalian specific NEEP21 family member, interacts with adaptor protein complex 3 (AP-3) and regulates targeting of AP-3 cargoes - PubMed — https://pubmed.ncbi.nlm.nih.gov/22650988/
- **[S6]** Calcyon stimulates neuregulin 1 maturation and signaling - PubMed — https://pubmed.ncbi.nlm.nih.gov/25349163/
- **[S7]** Hevin-calcyon interaction promotes synaptic reorganization after brain injury - PubMed — https://pubmed.ncbi.nlm.nih.gov/33753902/
- **[S8]** Complementary roles of the neuron-enriched endosomal proteins NEEP21 and calcyon in neuronal vesicle trafficking - PubMed — https://pubmed.ncbi.nlm.nih.gov/25376768/
- **[S9]** Association of the calcyon gene (DRD1IP) with attention deficit/hyperactivity disorder - PubMed — https://pubmed.ncbi.nlm.nih.gov/16172615/
- **[S10]** Calcyon, a novel partner of clathrin light chain, stimulates clathrin-mediated endocytosis - PubMed — https://pubmed.ncbi.nlm.nih.gov/16595675/
- **[S11]** Complementary roles of the neuron-enriched endosomal proteins NEEP21 and calcyon in neuronal vesicle trafficking - PubMed — https://pubmed.ncbi.nlm.nih.gov/25376768/
- **[S12]** Calcyon, a novel partner of clathrin light chain, stimulates clathrin-mediated endocytosis - PubMed — https://pubmed.ncbi.nlm.nih.gov/16595675/
- **[S13]** Complementary roles of the neuron-enriched endosomal proteins NEEP21 and calcyon in neuronal vesicle trafficking - PubMed — https://pubmed.ncbi.nlm.nih.gov/25376768/
