## Supplementary material for "Understanding Substance Dependence: What Differentiates Addictive from Non-Addictive Drugs?": Folder for supplementary files: non-addictive-genes-claims-first100-026-arf3-gene-protein-has-been-associated-with-regulat-3d5c054f76c6.md

# non-addictive-genes-claims-first100-026-3d5c054f76c6

**Original claim:** ARF3 gene/protein has been associated with regulatory or homeostatic biological processes in neural systems.

## Claim

ARF3 gene/protein has been associated with regulatory or homeostatic biological processes in neural systems.

## Verdict

Verdict: Likely True

ARF3 has credible evidence linking it to **regulatory processes in neural systems**, especially **neurodevelopmental regulation** rather than well-established adult neural homeostasis. The strongest support comes from human genetic and animal-model studies showing that pathogenic ARF3 variants disrupt brain development and affect early neural regulatory processes such as **neural precursor proliferation** and **planar cell polarity-dependent cell movements**. Evidence for a direct role in **mature neural homeostasis**, synaptic homeostatic plasticity, or circuit-level stabilization is much thinner. ([pubmed.ncbi.nlm.nih.gov](https://pubmed.ncbi.nlm.nih.gov/36369169/)) [S1]

## Confidence

Medium

## Supporting evidence

ARF3 is a bona fide trafficking GTPase with brain expression, making neural relevance biologically plausible. NCBI lists ARF3 as a reviewed protein-coding gene with **ubiquitous expression in brain**, and the Human Protein Atlas places it in a tissue-expression cluster annotated as **“Brain - Neuronal signaling (mainly)”**. Those resources do not prove neural function by themselves, but they support that ARF3 is present in neural tissues and expressed in a pattern compatible with neuronal biology. ([ncbi.nlm.nih.gov](https://www.ncbi.nlm.nih.gov/gene/377)) [S2]

The strongest direct evidence is from **human rare-disease genetics**. A 2021 Human Molecular Genetics report identified **de novo pathogenic ARF3 variants** in two unrelated children with **developmental delay, epilepsy, and brain abnormalities**. The same paper notes that ARF3 mainly works at the **trans-Golgi network (TGN)**, consistent with a role in regulated intracellular trafficking. This is clear evidence that ARF3 dysfunction can affect nervous-system development and brain structure in humans. ([pubmed.ncbi.nlm.nih.gov](https://pubmed.ncbi.nlm.nih.gov/34346499/)) [S3]

A 2022 Nature Communications study substantially strengthened that association. It reported additional de novo ARF3 missense variants causing a developmental disorder that impairs **nervous system formation**, and functional work showed disrupted **Golgi morphology**, **vesicle assembly/trafficking**, and—importantly for the present claim—early defects in **neural precursors’ proliferation** and **planar cell polarity-dependent cell movements** in vivo. Those are explicitly regulatory developmental processes in neural systems. ([pubmed.ncbi.nlm.nih.gov](https://pubmed.ncbi.nlm.nih.gov/36369169/)) [S4]

A 2022 Nature Communications review on “golgipathies” interpreted these findings in a broader developmental context. It states that ARF3 has been identified as a **Golgi-resident factor essential to brain and skeletal development**, and specifically summarizes evidence that ARF3 contributes to **mitotic spindle orientation in neural progenitor division during neurogenesis** and to **planar cell polarity**. That review also argues that Golgi-based trafficking mechanisms are important for development and homeostasis of differentiated tissues, placing ARF3 within a pathway with clear relevance to neural regulation. ([nature.com](https://www.nature.com/articles/s41467-022-35101-y)) [S5]

At the cell-biological level, ARF3 is not just a generic ARF-family member. A Molecular Biology of the Cell study found that ARF3 is activated **uniquely at the TGN**, depends specifically on **BIG-family guanine nucleotide exchange factors**, and likely serves a **unique function at the TGN**. Because the TGN is central to cargo sorting and membrane trafficking, this provides a mechanistic basis for why altered ARF3 activity could disturb developmental signaling and polarity in neural progenitors. ([pubmed.ncbi.nlm.nih.gov](https://pubmed.ncbi.nlm.nih.gov/20357002/)) [S6]

## Contradicting or limiting evidence

The main limitation is that the evidence is much stronger for **neural development** than for **neural homeostasis** in the narrower neuroscience sense. I did not find strong primary evidence showing that normal ARF3 directly governs adult neuronal homeostatic plasticity, neurotransmitter set-point regulation, firing-rate homeostasis, or glial homeostatic programs. The best-supported neural association is with **developmental regulation**, not established mature-system homeostasis. ([pubmed.ncbi.nlm.nih.gov](https://pubmed.ncbi.nlm.nih.gov/36369169/)) [S7]

Also, much of the mechanistic evidence comes from **rare pathogenic variants**, cultured non-neural cells, and zebrafish modeling. Those data are highly informative, but they do not fully establish how wild-type ARF3 functions in normal human neural physiology across the lifespan. ([pubmed.ncbi.nlm.nih.gov](https://pubmed.ncbi.nlm.nih.gov/36369169/)) [S8]

More broadly, synaptic-trafficking reviews discuss the **Arf family** as important in membrane traffic, but the most specific presynaptic evidence is centered on **ARF6**, while class I Arfs (including ARF3) are treated more generally. That pattern suggests direct ARF3-specific evidence in mature synapses remains sparse. This is an inference from the literature landscape, not proof that ARF3 lacks such roles. ([frontiersin.org](https://www.frontiersin.org/journals/synaptic-neuroscience/articles/10.3389/fnsyn.2016.00010/pdf)) [S9]

## Mechanistic / biological rationale

ARF3 encodes a **small GTPase** involved in **protein trafficking, vesicle budding/uncoating, and Golgi/TGN function**. NCBI and UniProt/HPA describe it as participating in intracellular protein transport and Golgi-associated trafficking, while the primary cell-biology literature shows that ARF3 has specialized TGN localization and regulation. ([ncbi.nlm.nih.gov](https://www.ncbi.nlm.nih.gov/gene/377)) [S10]

That biology fits well with neural regulatory roles. Neural progenitors and differentiating neurons rely heavily on polarized membrane trafficking to control **cell division orientation**, **cell polarity**, **cargo delivery**, receptor localization, and tissue morphogenesis. The ARF3 variant studies directly connect disturbed ARF3-dependent Golgi trafficking to **impaired neural precursor proliferation** and **planar cell polarity-related movements**, offering a plausible causal route from molecular defect to neural developmental dysregulation. ([pubmed.ncbi.nlm.nih.gov](https://pubmed.ncbi.nlm.nih.gov/36369169/)) [S11]

A more cautious extension toward “homeostasis” is also biologically reasonable but less directly proven. Reviews of Golgi biology emphasize that intracellular trafficking supports tissue **development and homeostasis**, and ARF3-related Golgi dysfunction can delay recycling and perturb cargo handling. It is therefore plausible that ARF3 contributes to maintenance of neural cellular equilibrium, but the current literature supports this more as a **mechanistic inference** than as a directly demonstrated neural-homeostasis function. ([nature.com](https://www.nature.com/articles/s41467-022-35101-y)) [S12]

## Bottom line

The claim is **reasonable overall** if “regulatory” is interpreted broadly to include **neurodevelopmental regulation**. ARF3 is associated with brain expression, human neurodevelopmental disorders, and experimentally supported effects on **neural precursor proliferation**, **cell polarity**, and Golgi-mediated trafficking relevant to nervous-system development. But if the claim is meant to assert a well-established role in **adult neural homeostasis**, the evidence is currently **limited and indirect**, not definitive. ([pubmed.ncbi.nlm.nih.gov](https://pubmed.ncbi.nlm.nih.gov/36369169/)) [S13]

## Cited sources

- **[S1]** Dominant ARF3 variants disrupt Golgi integrity and cause a neurodevelopmental disorder recapitulated in zebrafish - PubMed — https://pubmed.ncbi.nlm.nih.gov/36369169/
- **[S2]** ARF3 ARF GTPase 3 [Homo sapiens (human)] - Gene - NCBI — https://www.ncbi.nlm.nih.gov/gene/377
- **[S3]** De novo ARF3 variants cause neurodevelopmental disorder with brain abnormality - PubMed — https://pubmed.ncbi.nlm.nih.gov/34346499/
- **[S4]** Dominant ARF3 variants disrupt Golgi integrity and cause a neurodevelopmental disorder recapitulated in zebrafish - PubMed — https://pubmed.ncbi.nlm.nih.gov/36369169/
- **[S5]** Golgipathies reveal the critical role of the sorting machinery in brain and skeletal development | Nature Communications — https://www.nature.com/articles/s41467-022-35101-y
- **[S6]** Arf3 is activated uniquely at the trans-Golgi network by brefeldin A-inhibited guanine nucleotide exchange factors - PubMed — https://pubmed.ncbi.nlm.nih.gov/20357002/
- **[S7]** Dominant ARF3 variants disrupt Golgi integrity and cause a neurodevelopmental disorder recapitulated in zebrafish - PubMed — https://pubmed.ncbi.nlm.nih.gov/36369169/
- **[S8]** Dominant ARF3 variants disrupt Golgi integrity and cause a neurodevelopmental disorder recapitulated in zebrafish - PubMed — https://pubmed.ncbi.nlm.nih.gov/36369169/
- **[S9]** Molecular Machines Determining the Fate of Endocytosed Synaptic Vesicles in Nerve Terminals — https://www.frontiersin.org/journals/synaptic-neuroscience/articles/10.3389/fnsyn.2016.00010/pdf
- **[S10]** ARF3 ARF GTPase 3 [Homo sapiens (human)] - Gene - NCBI — https://www.ncbi.nlm.nih.gov/gene/377
- **[S11]** Dominant ARF3 variants disrupt Golgi integrity and cause a neurodevelopmental disorder recapitulated in zebrafish - PubMed — https://pubmed.ncbi.nlm.nih.gov/36369169/
- **[S12]** Golgipathies reveal the critical role of the sorting machinery in brain and skeletal development | Nature Communications — https://www.nature.com/articles/s41467-022-35101-y
- **[S13]** Dominant ARF3 variants disrupt Golgi integrity and cause a neurodevelopmental disorder recapitulated in zebrafish - PubMed — https://pubmed.ncbi.nlm.nih.gov/36369169/
