## Supplementary material for "Understanding Substance Dependence: What Differentiates Addictive from Non-Addictive Drugs?": Folder for supplementary files: non-addictive-genes-claims-first100-027-aatk-gene-protein-has-been-associated-with-regulat-9b354343aece.md

# non-addictive-genes-claims-first100-027-9b354343aece

**Original claim:** AATK gene/protein has been associated with regulatory or homeostatic biological processes in neural systems.

## Claim

AATK gene/protein has been associated with regulatory or homeostatic biological processes in neural systems.

## Verdict

Verdict: Likely True

AATK (also called AATYK or LMTK1 in much of the neuroscience literature) has substantial evidence linking it to **regulatory neural processes**: neuronal differentiation, neurite extension, axon outgrowth, dendrite/spine formation, endosomal trafficking in neurons, and regulation of neuronal apoptosis/survival signaling. However, I found **little direct evidence** that AATK has been studied as a mediator of **neural homeostasis in the strict sense** used for synaptic scaling, firing-rate homeostasis, or classic homeostatic plasticity. So the claim is well supported if “regulatory” is the key term, but only weakly supported if interpreted narrowly as “homeostatic.” ([pubmed.ncbi.nlm.nih.gov](https://pubmed.ncbi.nlm.nih.gov/10837911/)) [S1]

## Confidence

Confidence: Medium

The confidence is not low because multiple peer-reviewed studies, plus authoritative gene/expression resources, consistently place AATK in neural regulatory pathways. It is not high because much of the mechanistic work is in cultured cells or mouse models, and because “homeostatic” is broader than what the direct AATK literature usually tests. ([pubmed.ncbi.nlm.nih.gov](https://pubmed.ncbi.nlm.nih.gov/12670708/)) [S2]

## Supporting evidence

AATK is strongly enriched in neural tissue and broadly present across the brain, which is a basic prerequisite for a neural regulatory role. Early studies found widespread neuronal expression of AATYK mRNA/protein in adult brain and multiple brain regions, and later resources show human AATK is brain-biased or brain-enhanced, with RNA detected across all major sampled brain regions. ([nature.com](https://www.nature.com/articles/1204210)) [S3]

There is direct experimental evidence that AATK promotes **neuronal differentiation**. In SH-SY5Y neuroblastoma cells, AATYK overexpression increased the proportion of differentiated cells and neurite length, and also enhanced differentiation induced by retinoic acid, TPA, and IGF-I. In developing mouse brain/cerebellar granule cells, AATYK expression rose during postnatal development and wild-type AATYK promoted neurite outgrowth, whereas a kinase-defective mutant inhibited it. These are classic regulatory developmental processes in neural systems. ([pubmed.ncbi.nlm.nih.gov](https://pubmed.ncbi.nlm.nih.gov/10837911/)) [S4]

AATK has also been associated with regulation of **neuronal apoptosis/survival-related signaling**. In mature cerebellar granule neurons, AATYK expression increased during low-potassium apoptosis, and overexpression of wild-type AATYK increased apoptosis more than kinase-dead AATYK. A later study showed that AATYK phosphorylation states tracked survival versus apoptotic conditions in cerebellar granule cells, with distinct Ca2+-dependent regulation under high- vs low-potassium conditions. That supports involvement in neural-state regulation, especially survival/death responses. ([nature.com](https://www.nature.com/articles/1204210)) [S5]

The strongest mechanistic neuroscience literature places AATK/LMTK1 in **neuronal membrane trafficking and morphogenesis**. In mouse cortical neurons, LMTK1/AATYK1 was highly expressed in neurons, phosphorylated during axonal outgrowth, colocalized with Rab11A-positive recycling endosomes, and acted upstream of Rab11A in a Cdk5-dependent pathway. Knockdown or knockout enhanced axonal elongation and increased anterograde movement of Rab11A-positive vesicles, while rescue with wild-type or phosphomimetic LMTK1 reversed the phenotype. That is clear evidence of a regulatory role in neuronal structure formation. ([pubmed.ncbi.nlm.nih.gov](https://pubmed.ncbi.nlm.nih.gov/22573681/)) [S6]

AATK/LMTK1 also regulates **dendritic spines and synapse-related structure**. Knockdown of LMTK1 increased dendritic spine density in cultured hippocampal neurons and after in utero electroporation in developing mouse brain; the study further linked this to a LMTK1-TBC1D9B-Rab11A cascade regulating spine formation via endosome trafficking. Separately, LMTK1 knockout mice had increased synapse numbers despite grossly normal overall brain structure, along with behavioral abnormalities such as hyperactivity and impulsiveness, suggesting that AATK normally restrains aspects of neuronal circuit growth/synaptic organization rather than simply serving as a passive marker. ([pubmed.ncbi.nlm.nih.gov](https://pubmed.ncbi.nlm.nih.gov/31628178/)) [S7]

Authoritative annotation resources are consistent with these experiments. NCBI/Alliance annotate AATK/Aatk as involved in **brain development** and **neuron apoptotic process**, and the human NCBI entry summarizes that the gene has been shown to induce neuronal differentiation in a neuroblastoma cell line. These database summaries are not primary proof by themselves, but they align with the published experimental literature. ([ncbi.nlm.nih.gov](https://www.ncbi.nlm.nih.gov/gene/11302?utm_source=openai)) [S8]

## Contradicting or limiting evidence

The main limitation is that I did **not** find direct AATK studies on canonical **neural homeostasis** endpoints such as firing-rate homeostasis, synaptic scaling, excitatory/inhibitory set-point maintenance, or textbook “homeostatic plasticity.” The review literature instead centers on endosomal trafficking, neurite/axon/dendrite/spine growth, and prevention of neuronal overgrowth. So the evidence strongly supports “regulatory,” but only indirectly supports “homeostatic.” ([frontiersin.org](https://www.frontiersin.org/articles/10.3389/fnmol.2020.00112/full)) [S9]

Another limitation is that much of the functional evidence comes from **mouse neurons, mouse knockout models, cultured cerebellar granule neurons, or cell lines**. Human evidence is stronger for expression and association than for direct in vivo causality. For example, the human NCBI record emphasizes brain-biased expression and neuronal differentiation in a neuroblastoma line, whereas the most detailed mechanistic work on axons, Rab11 trafficking, and spine regulation is from rodent systems. ([ncbi.nlm.nih.gov](https://www.ncbi.nlm.nih.gov/gene/9625)) [S10]

There is also some unresolved biology around the protein itself. AATK was originally classified as a tyrosine kinase, but later literature treats LMTK1/AATK as a serine/threonine kinase; the 2020 review notes that LMTK1 kinase activity itself has not yet been clearly shown, even though kinase-negative mutants have strong phenotypic effects. That does not negate a regulatory neural role, but it does mean some molecular details remain unsettled. ([frontiersin.org](https://www.frontiersin.org/articles/10.3389/fnmol.2020.00112/full)) [S11]

## Mechanistic / biological rationale

A coherent mechanistic model has emerged in which AATK/LMTK1 acts as a **brake on neuronal membrane trafficking and structural growth**. AATK binds the neuronal kinase regulator p35/Cdk5, is phosphorylated by Cdk5, and localizes to Rab11-positive recycling endosomes. Phosphorylation at Ser34 alters its recycling-endosome localization and links Cdk5 activity to Rab11-dependent endosomal traffic. Because Rab11-mediated recycling supplies membrane and cargo needed for axon elongation, dendrite arborization, and spine formation, AATK is well positioned to regulate these processes by modulating endosomal flow. ([pubmed.ncbi.nlm.nih.gov](https://pubmed.ncbi.nlm.nih.gov/14521924/)) [S12]

This explains why loss or inhibition of AATK/LMTK1 tends to produce **overgrowth phenotypes**: longer axons, greater dendritic branching, and more spines/synapses. The spine paper further places TBC1D9B, a Rab11 GAP, downstream of LMTK1, strengthening the idea that AATK is not just correlated with neuronal morphology but participates in a specific trafficking-control cascade. Biologically, such a pathway is plausibly relevant to maintaining orderly neural architecture and preventing excess membrane/cargo delivery, which is “homeostasis-like” in a broad sense, even if not directly tested in classic homeostatic-plasticity paradigms. ([pubmed.ncbi.nlm.nih.gov](https://pubmed.ncbi.nlm.nih.gov/31628178/)) [S13]

A second mechanistic axis is its link to **neuronal survival/apoptosis regulation** in cerebellar granule cells. AATK abundance and phosphorylation state change with survival versus apoptotic conditions, and kinase-defective mutants blunt the apoptosis-associated phenotype. That places AATK in intracellular signaling networks that regulate neuronal state transitions, another bona fide regulatory role in neural biology. ([nature.com](https://www.nature.com/articles/1204210)) [S14]

## Bottom line

The claim is **likely true**. AATK has repeatedly been associated with **regulatory biological processes in neural systems**, including neuronal differentiation, neurite extension, axon/dendrite/spine regulation, endosomal trafficking, and neuronal apoptosis. What is **not** well established is a direct role in **neural homeostasis** in the narrow, classical neuroscience sense. So the best evidence-based reading is: **AATK is clearly linked to neural regulatory processes; evidence for explicit homeostatic roles is presently limited and indirect.** ([pubmed.ncbi.nlm.nih.gov](https://pubmed.ncbi.nlm.nih.gov/10837911/)) [S15]

## Cited sources

- **[S1]** A novel kinase, AATYK induces and promotes neuronal differentiation in a human neuroblastoma (SH-SY5Y) cell line - PubMed — https://pubmed.ncbi.nlm.nih.gov/10837911/
- **[S2]** Differential expression and function of apoptosis-associated tyrosine kinase (AATYK) in the developing mouse brain - PubMed — https://pubmed.ncbi.nlm.nih.gov/12670708/
- **[S3]** Characterization of the apoptosis-associated tyrosine kinase (AATYK) expressed in the CNS | Oncogene — https://www.nature.com/articles/1204210
- **[S4]** A novel kinase, AATYK induces and promotes neuronal differentiation in a human neuroblastoma (SH-SY5Y) cell line - PubMed — https://pubmed.ncbi.nlm.nih.gov/10837911/
- **[S5]** Characterization of the apoptosis-associated tyrosine kinase (AATYK) expressed in the CNS | Oncogene — https://www.nature.com/articles/1204210
- **[S6]** LMTK1/AATYK1 is a novel regulator of axonal outgrowth that acts via Rab11 in a Cdk5-dependent manner - PubMed — https://pubmed.ncbi.nlm.nih.gov/22573681/
- **[S7]** The LMTK1-TBC1D9B-Rab11A Cascade Regulates Dendritic Spine Formation via Endosome Trafficking - PubMed — https://pubmed.ncbi.nlm.nih.gov/31628178/
- **[S8]** Aatk apoptosis-associated tyrosine kinase [Mus musculus (house mouse)] - Gene - NCBI — https://www.ncbi.nlm.nih.gov/gene/11302?utm_source=openai
- **[S9]** Frontiers | LMTK1, a Novel Modulator of Endosomal Trafficking in Neurons — https://www.frontiersin.org/articles/10.3389/fnmol.2020.00112/full
- **[S10]** AATK apoptosis associated tyrosine kinase [Homo sapiens (human)] - Gene - NCBI — https://www.ncbi.nlm.nih.gov/gene/9625
- **[S11]** Frontiers | LMTK1, a Novel Modulator of Endosomal Trafficking in Neurons — https://www.frontiersin.org/articles/10.3389/fnmol.2020.00112/full
- **[S12]** Apoptosis-associated tyrosine kinase is a Cdk5 activator p35 binding protein - PubMed — https://pubmed.ncbi.nlm.nih.gov/14521924/
- **[S13]** The LMTK1-TBC1D9B-Rab11A Cascade Regulates Dendritic Spine Formation via Endosome Trafficking - PubMed — https://pubmed.ncbi.nlm.nih.gov/31628178/
- **[S14]** Characterization of the apoptosis-associated tyrosine kinase (AATYK) expressed in the CNS | Oncogene — https://www.nature.com/articles/1204210
- **[S15]** A novel kinase, AATYK induces and promotes neuronal differentiation in a human neuroblastoma (SH-SY5Y) cell line - PubMed — https://pubmed.ncbi.nlm.nih.gov/10837911/
