## Supplementary material for "Understanding Substance Dependence: What Differentiates Addictive from Non-Addictive Drugs?": Folder for supplementary files: non-addictive-genes-claims-first100-028-gpx4-gene-protein-has-been-associated-with-regulat-086307a891f1.md

# non-addictive-genes-claims-first100-028-086307a891f1

**Original claim:** GPX4 gene/protein has been associated with regulatory or homeostatic biological processes in neural systems.

## Claim

GPX4 gene/protein has been associated with regulatory or homeostatic biological processes in neural systems.

## Verdict

Verdict: Highly Supported

The claim is strongly supported, especially if “regulatory or homeostatic” is interpreted to include **neuronal lipid redox homeostasis, membrane protection, resistance to ferroptotic cell death, developmental maintenance, and injury-response programs in the CNS**. Multiple lines of evidence—from neural expression studies, neuron-specific loss-of-function models, human disease genetics, iPSC/organoid work, and disease-model rescue experiments—show that GPX4 is not just present in neural tissue, but functionally important for maintaining neural integrity. ([pubmed.ncbi.nlm.nih.gov](https://pubmed.ncbi.nlm.nih.gov/17603929/)) [S1]

## Confidence

High

The evidence base includes mechanistic biochemistry, developmental studies, conditional mouse genetics, disease-model intervention studies, and recent human mutation/iPSC/organoid data. The main limitation is that much of the strongest causal work is preclinical, but the overall conclusion is still robust. ([pubmed.ncbi.nlm.nih.gov](https://pubmed.ncbi.nlm.nih.gov/30888116/)) [S2]

## Supporting evidence

- **Neural expression and developmental patterning:** GPX4 is expressed in the developing brain, with large amounts of mRNA detected in neuronal layers during perinatal development; in adult brain, mitochondrial and cytosolic GPX4 isoforms were detected in neurons of cortex, hippocampus, and cerebellum. A review of the field similarly concluded that cytosolic and mitochondrial GPX4 are major isoforms expressed by neurons in the developing brain, supporting a role in neural development and function. ([pubmed.ncbi.nlm.nih.gov](https://pubmed.ncbi.nlm.nih.gov/17603929/)) [S3]

- **Direct evidence that GPX4 supports neural homeostasis:** In the 2007 brain study, selective GPX4 knockdown caused depletion of plasma-membrane phosphatidylinositol-(4,5)-bisphosphate and then apoptosis, indicating that GPX4 helps maintain membrane lipid/signaling homeostasis rather than merely serving as a passive antioxidant marker. The same study found GPX4 induction in reactive astrocytes after brain injury, consistent with a CNS protective/homeostatic response. ([pubmed.ncbi.nlm.nih.gov](https://pubmed.ncbi.nlm.nih.gov/17603929/)) [S4]

- **Adult nervous system requirement:** Systemic inducible ablation of Gpx4 in adult mice caused death within about 2 weeks and was accompanied by hippocampal neuronal loss and increased astrogliosis. The authors concluded that Gpx4 is essential for mitochondrial integrity and neuron survival in adult animals. ([pubmed.ncbi.nlm.nih.gov](https://pubmed.ncbi.nlm.nih.gov/22401858/)) [S5]

- **Neuron-specific causal evidence:** Forebrain neuron-specific ablation of Gpx4 caused cognitive impairment, hippocampal neurodegeneration, elevated lipid peroxidation, ERK activation, and neuroinflammation, with features consistent with ferroptotic rather than apoptotic death. Vitamin E deficiency accelerated this degeneration, reinforcing the interpretation that GPX4 participates in neural antioxidant/homeostatic defense. ([pmc.ncbi.nlm.nih.gov](https://pmc.ncbi.nlm.nih.gov/articles/PMC5312549/)) [S6]

- **Interneuron development and maintenance:** In neuron-specific Gpx4 knockout mice, hippocampal cell death appeared early and parvalbumin-positive interneurons were selectively reduced, while calretinin-positive interneurons were less affected. This supports a role for GPX4 in the maintenance of specific neuronal subtypes and network-relevant cell populations. ([pubmed.ncbi.nlm.nih.gov](https://pubmed.ncbi.nlm.nih.gov/19890015/)) [S7]

- **Human genetic evidence:** Truncating GPX4 mutations were identified in Sedaghatian-type spondylometaphyseal dysplasia, a neonatal lethal disorder that includes central nervous system abnormalities. The genetic report explicitly notes GPX4’s role in protecting against membrane lipid peroxidation and its importance for embryonic development. ([pubmed.ncbi.nlm.nih.gov](https://pubmed.ncbi.nlm.nih.gov/24706940/)) [S8]

- **Recent human-neuron and organoid evidence:** A 2026 Cell paper showed that a disease-causing GPX4 missense variant (R152H) disrupts membrane anchoring and causes early-onset neurodegeneration. In that study, spatiotemporal Gpx4 deletion or neuron-specific GPX4^R152H expression caused degeneration of cortical and cerebellar neurons with progressive neuroinflammation; patient iPSC-derived cortical neurons and forebrain organoids showed increased ferroptotic vulnerability that was sensitive to ferroptosis inhibition. This is especially strong evidence that GPX4 is functionally involved in neural homeostasis, not merely correlated with it. ([pubmed.ncbi.nlm.nih.gov](https://pubmed.ncbi.nlm.nih.gov/41349546/)) [S9]

- **Disease-model rescue studies support a homeostatic role:** In ALS models, reduced GPX4 in spinal cord was linked to motor-neuron loss and phospholipid peroxide accumulation; neuron-targeted GPX4 delivery or GPX4 transgenic overexpression improved motor performance, delayed disease onset, reduced lipid peroxidation, and ameliorated motor-neuron degeneration. One study explicitly concluded that GPX4 is important for maintaining phospholipid redox homeostasis in the spinal cord. ([pubmed.ncbi.nlm.nih.gov](https://pubmed.ncbi.nlm.nih.gov/36585109/)) [S10]

- **Broader CNS disease association:** In multiple sclerosis and experimental autoimmune encephalomyelitis, all three GPX4 isoforms declined in affected CNS tissue, and the study emphasizes GPX4 as the major inhibitor of ferroptosis in neurological disorders with oxidative stress. This does not by itself prove causality, but it is consistent with GPX4 participating in CNS protective homeostasis. ([pmc.ncbi.nlm.nih.gov](https://pmc.ncbi.nlm.nih.gov/articles/PMC6347488/)) [S11]

## Contradicting or limiting evidence

There is **little evidence contradicting the core claim**, but there are important limitations on how strongly it should be framed. First, GPX4 is a **general cellular lipid-peroxide defense enzyme**, not a neural-specific regulator in the way that neurotransmitter receptors, ion channels, or developmental transcription factors are. So the best-supported wording is that GPX4 is involved in **neural redox/lipid homeostasis and regulated cell-death control**, rather than being a dedicated neural signaling regulator. ([pubmed.ncbi.nlm.nih.gov](https://pubmed.ncbi.nlm.nih.gov/30888116/)) [S12]

Second, much of the strongest causal evidence comes from **mouse genetics, cultured neurons, and organoids**, often under injury or disease conditions. Human evidence exists, but it is still relatively limited and concentrated in rare disorders such as GPX4-related developmental disease, plus recent patient-derived cellular models. ([pubmed.ncbi.nlm.nih.gov](https://pubmed.ncbi.nlm.nih.gov/24706940/)) [S13]

Third, “homeostatic biological processes in neural systems” is a broad phrase. GPX4’s strongest demonstrated homeostatic role is in **membrane lipid redox balance and protection from ferroptosis**. Evidence for broader roles in neural circuit regulation, synaptic plasticity, or neurotransmission is more indirect. ([pubmed.ncbi.nlm.nih.gov](https://pubmed.ncbi.nlm.nih.gov/17603929/)) [S14]

## Mechanistic / biological rationale

GPX4 is unusual among glutathione peroxidases because it can directly reduce **phospholipid hydroperoxides in biological membranes**, thereby preventing propagation of lipid peroxidation and suppressing ferroptosis. Reviews describe GPX4 as central at the intersection of lipid homeostasis and ferroptosis, and CNS-focused reviews note that neuronal survival under steady-state and stress conditions depends on the cystine/GSH/GPX4 axis. ([pubmed.ncbi.nlm.nih.gov](https://pubmed.ncbi.nlm.nih.gov/30888116/)) [S15]

This mechanism is especially relevant in neural tissue because neurons are rich in **oxidation-prone polyunsaturated membrane lipids**, are metabolically active, and are vulnerable to iron- and oxidative-stress-linked damage. When GPX4 levels fall, phospholipid peroxides accumulate, membrane integrity and signaling lipids are disrupted, and neurons become susceptible to ferroptotic death and neuroinflammatory secondary responses. That pattern is seen in forebrain-neuron knockouts, ALS models, and the recent human R152H mutation study. ([pmc.ncbi.nlm.nih.gov](https://pmc.ncbi.nlm.nih.gov/articles/PMC5312549/)) [S16]

Additional mechanistic support comes from intervention studies: pharmacologic selenium can drive neuronal GPX4 expression and protect neurons in stroke models, and ferroptosis inhibitors rescue GPX4-deficient human neuronal models. These findings fit a coherent biology in which GPX4 is a key component of neural homeostatic defense against lipid oxidative damage. ([pubmed.ncbi.nlm.nih.gov](https://pubmed.ncbi.nlm.nih.gov/31056284/?utm_source=openai)) [S17]

## Bottom line

Yes—**GPX4 is strongly associated with, and in many models causally required for, homeostatic processes in neural systems**, particularly **neuronal membrane lipid redox homeostasis, resistance to ferroptosis, developmental maintenance, and injury/stress protection**. The evidence is strongest for a **protective/homeostatic antioxidant-lipid repair role in neurons and some glial injury responses**, rather than for a classic neurotransmission-regulatory role. Overall, the claim is **Highly Supported**. ([pubmed.ncbi.nlm.nih.gov](https://pubmed.ncbi.nlm.nih.gov/17603929/)) [S18]

## Cited sources

- **[S1]** Role for glutathione peroxidase-4 in brain development and neuronal apoptosis: specific induction of enzyme expression in reactive astrocytes following brain injury - PubMed — https://pubmed.ncbi.nlm.nih.gov/17603929/
- **[S2]** GPX4 at the Crossroads of Lipid Homeostasis and Ferroptosis - PubMed — https://pubmed.ncbi.nlm.nih.gov/30888116/
- **[S3]** Role for glutathione peroxidase-4 in brain development and neuronal apoptosis: specific induction of enzyme expression in reactive astrocytes following brain injury - PubMed — https://pubmed.ncbi.nlm.nih.gov/17603929/
- **[S4]** Role for glutathione peroxidase-4 in brain development and neuronal apoptosis: specific induction of enzyme expression in reactive astrocytes following brain injury - PubMed — https://pubmed.ncbi.nlm.nih.gov/17603929/
- **[S5]** Gpx4 ablation in adult mice results in a lethal phenotype accompanied by neuronal loss in brain - PubMed — https://pubmed.ncbi.nlm.nih.gov/22401858/
- **[S6]** 
            Ablation of ferroptosis regulator glutathione peroxidase 4 in forebrain neurons promotes cognitive impairment and neurodegeneration - PMC
         — https://pmc.ncbi.nlm.nih.gov/articles/PMC5312549/
- **[S7]** Neuronal selenoprotein expression is required for interneuron development and prevents seizures and neurodegeneration - PubMed — https://pubmed.ncbi.nlm.nih.gov/19890015/
- **[S8]** Mutations in the enzyme glutathione peroxidase 4 cause Sedaghatian-type spondylometaphyseal dysplasia - PubMed — https://pubmed.ncbi.nlm.nih.gov/24706940/
- **[S9]** A fin-loop-like structure in GPX4 underlies neuroprotection from ferroptosis - PubMed — https://pubmed.ncbi.nlm.nih.gov/41349546/
- **[S10]** GPX4 deficiency-dependent phospholipid peroxidation drives motor deficits of ALS - PubMed — https://pubmed.ncbi.nlm.nih.gov/36585109/
- **[S11]** 
            Reduced expression of the ferroptosis inhibitor GPx4 in multiple sclerosis and experimental autoimmune encephalomyelitis - PMC
         — https://pmc.ncbi.nlm.nih.gov/articles/PMC6347488/
- **[S12]** GPX4 at the Crossroads of Lipid Homeostasis and Ferroptosis - PubMed — https://pubmed.ncbi.nlm.nih.gov/30888116/
- **[S13]** Mutations in the enzyme glutathione peroxidase 4 cause Sedaghatian-type spondylometaphyseal dysplasia - PubMed — https://pubmed.ncbi.nlm.nih.gov/24706940/
- **[S14]** Role for glutathione peroxidase-4 in brain development and neuronal apoptosis: specific induction of enzyme expression in reactive astrocytes following brain injury - PubMed — https://pubmed.ncbi.nlm.nih.gov/17603929/
- **[S15]** GPX4 at the Crossroads of Lipid Homeostasis and Ferroptosis - PubMed — https://pubmed.ncbi.nlm.nih.gov/30888116/
- **[S16]** 
            Ablation of ferroptosis regulator glutathione peroxidase 4 in forebrain neurons promotes cognitive impairment and neurodegeneration - PMC
         — https://pmc.ncbi.nlm.nih.gov/articles/PMC5312549/
- **[S17]** Selenium Drives a Transcriptional Adaptive Program to Block Ferroptosis and Treat Stroke - PubMed — https://pubmed.ncbi.nlm.nih.gov/31056284/?utm_source=openai
- **[S18]** Role for glutathione peroxidase-4 in brain development and neuronal apoptosis: specific induction of enzyme expression in reactive astrocytes following brain injury - PubMed — https://pubmed.ncbi.nlm.nih.gov/17603929/
