## Supplementary material for "Understanding Substance Dependence: What Differentiates Addictive from Non-Addictive Drugs?": Folder for supplementary files: non-addictive-genes-claims-first100-029-rpl17-gene-protein-has-been-associated-with-regula-ca4d86f4d537.md

# non-addictive-genes-claims-first100-029-ca4d86f4d537

**Original claim:** RPL17 gene/protein has been associated with regulatory or homeostatic biological processes in neural systems.

## Claim

RPL17 gene/protein has been associated with regulatory or homeostatic biological processes in neural systems.

## Verdict

Verdict: Likely True

Broadly, **yes**—but the evidence is **limited and mostly indirect**. RPL17 has been linked to **neural developmental regulation** and to **synaptic/axonal translation machinery** in neural tissue, which are biologically relevant regulatory processes. However, there is **little direct evidence** that RPL17 itself is a dedicated mediator of **neural homeostasis** (for example, synaptic scaling, excitability set-point control, or circuit-level homeostatic plasticity) in mammals. ([pmc.ncbi.nlm.nih.gov](https://pmc.ncbi.nlm.nih.gov/articles/PMC2878125/)) [S1]

## Confidence

Medium

The claim is supportable if interpreted as a **broad association** with neural regulatory biology, but not if interpreted as a **well-established, specific homeostatic regulator** in the nervous system. Evidence comes from curated synaptic annotations, localization/proteomics, and songbird developmental studies rather than strong causal mammalian neurobiology. ([syngoportal.org](https://www.syngoportal.org/annotation_3602.html?utm_source=openai)) [S2]

## Supporting evidence

RPL17 is a canonical **60S ribosomal protein** encoded by a ubiquitously expressed human gene, but curated synaptic databases do place it in **neural/synaptic translation contexts**. In SynGO, RPL17 is annotated to **postsynaptic ribosome**, **presynaptic ribosome**, **translation at postsynapse**, and **translation at presynapse**. These annotations are based on synaptic proteomics plus established literature showing ribosomes and translation in synaptic compartments. ([syngoportal.org](https://www.syngoportal.org/gene_hgnc10307.html)) [S3]

There is also **direct developmental neural association** in vivo from zebra finch studies. In juvenile zebra finches, RPL17 mRNA was elevated in male song-control regions including **Area X**, **RA**, and the **ventral ventricular zone**, and the authors concluded the pattern was consistent with involvement in the **masculine development** of these nuclei and possibly in **neuron genesis and/or survival**. That is a genuine association with neural regulatory/developmental processes, even though it is species- and context-specific. ([pmc.ncbi.nlm.nih.gov](https://pmc.ncbi.nlm.nih.gov/articles/PMC2878125/)) [S4]

A follow-up zebra finch study strengthened that association by showing that **most BrdU-labeled newly generated cells** in song-system regions expressed RPL17, and the authors concluded that **RPL17, RPL37, and estradiol might all influence sexual differentiation** of the song system. Again, this is not homeostatic plasticity in the strict neuroscience sense, but it is a neural regulatory process. ([pubmed.ncbi.nlm.nih.gov](https://pubmed.ncbi.nlm.nih.gov/19373862/)) [S5]

RPL17 protein has also been observed in **regenerating sensory axons** together with other ribosomal proteins, rRNA, and translation factors. That finding supports the idea that RPL17 participates in **local axonal translation machinery**, which is relevant to neural growth and adaptive responses after injury. ([pubmed.ncbi.nlm.nih.gov](https://pubmed.ncbi.nlm.nih.gov/11717363/)) [S6]

More generally, the neural relevance of synaptic/axonal ribosomes is well established. Polyribosomes redistribute into dendritic spines during LTP, and axonal translatomes are dynamically regulated in developing and mature visual circuits. Because RPL17 is a constituent of these ribosomes, its presence in neural regulatory compartments is biologically plausible. ([pubmed.ncbi.nlm.nih.gov](https://pubmed.ncbi.nlm.nih.gov/12165474/?utm_source=openai)) [S7]

## Contradicting or limiting evidence

The strongest limitation is that **RPL17 is not known primarily as a neural regulator**. Authoritative gene resources describe it mainly as a **cytoplasmic 60S ribosomal subunit protein**, and NCBI notes **ubiquitous expression** across tissues. The Human Protein Atlas classifies it as having **low human brain regional specificity** and belonging to a **non-specific ribosome-related expression cluster**, which argues against a brain-specific regulatory role. ([ncbi.nlm.nih.gov](https://www.ncbi.nlm.nih.gov/gene/6139)) [S8]

The SynGO evidence is also more limited than it may first appear. The annotation rationale explicitly says that RPL17’s synaptic assignment is largely inferred because it is a **fixed ribosomal component**, and that **no functions of RPL17 are known independent from the ribosomal complex** in that context. In other words, current synaptic annotation supports that RPL17 is **present where synaptic translation occurs**, not that it is a unique control node for neural regulation. ([syngoportal.org](https://www.syngoportal.org/annotation_3553)) [S9]

Human genetics likewise does not currently point to a prominent neural phenotype. Recent work linked **RPL17 haploinsufficiency** to **Diamond-Blackfan anemia** and erythropoietic defects, with mechanistic effects on ribosome biogenesis and translation. That is important biology, but it is not strong evidence for a primary neural homeostatic function. ([pmc.ncbi.nlm.nih.gov](https://pmc.ncbi.nlm.nih.gov/articles/PMC11385091/)) [S10]

Finally, I did **not** find strong mammalian studies showing that manipulating RPL17 specifically alters classic neural homeostatic outcomes such as **synaptic scaling**, **excitability set-point regulation**, or **E/I balance maintenance**. So the claim is better supported for **association with neural regulatory contexts** than for **direct implication in neural homeostasis** per se. ([pubmed.ncbi.nlm.nih.gov](https://pubmed.ncbi.nlm.nih.gov/34813590/?utm_source=openai)) [S11]

## Mechanistic / biological rationale

A plausible mechanism exists, but it remains mostly **inferential**. Neural systems rely heavily on **local translation** in dendrites, spines, and axons for plasticity, development, and some forms of homeostatic adaptation. For example, translation-dependent pathways are required for homeostatic scaling in hippocampal neurons, and ribosomes redistribute into dendritic spines during synaptic strengthening. Therefore, a ribosomal protein such as RPL17 could influence neural regulation indirectly by shaping the capacity or properties of local protein synthesis. ([pubmed.ncbi.nlm.nih.gov](https://pubmed.ncbi.nlm.nih.gov/34813590/?utm_source=openai)) [S12]

At the molecular level, RPL17 is more than a generic label on the ribosome: studies of RPL17 deficiency show effects on **pre-rRNA processing**, **60S subunit maturation**, and the structure of the **peptide exit tunnel** region. Recent work on RPL17-associated Diamond-Blackfan anemia reported altered 5.8S rRNA processing and suggested that RPL17 directly contacts rRNA near the peptide exit tunnel, potentially affecting nascent-chain progression or chaperone interactions. That gives a credible route by which RPL17 perturbation could alter translation programs in any cell type, including neurons. ([ncbi.nlm.nih.gov](https://www.ncbi.nlm.nih.gov/gene/6139)) [S13]

Still, that mechanistic rationale should not be overstated. What is established is that **neuronal regulation often depends on ribosome-mediated local translation**; what is **not** yet established is that **RPL17 specifically** is a major dedicated regulator of neural homeostasis rather than one of many essential ribosomal subunits participating in that process. ([pubmed.ncbi.nlm.nih.gov](https://pubmed.ncbi.nlm.nih.gov/34813590/?utm_source=openai)) [S14]

## Bottom line

The claim is **likely true in a broad sense**: RPL17 has been associated with **neural regulatory biology**, especially **developmental regulation in songbird brain circuits** and **synaptic/axonal translation machinery**. But the evidence for a **specific homeostatic role** in neural systems is **sparse and indirect**. So the fairest summary is: **RPL17 is associated with neural regulatory contexts, mainly through its role as a ribosomal component in local translation, but it is not yet a well-established neural homeostasis gene/protein.** ([pmc.ncbi.nlm.nih.gov](https://pmc.ncbi.nlm.nih.gov/articles/PMC2878125/)) [S15]

## Cited sources

- **[S1]** 
            Sexually dimorphic expression of the genes encoding ribosomal proteins L17 and L37 in the song control nuclei of juvenile zebra finches - PMC
         — https://pmc.ncbi.nlm.nih.gov/articles/PMC2878125/
- **[S2]** SynGO - annotation details — https://www.syngoportal.org/annotation_3602.html?utm_source=openai
- **[S3]** SynGO - gene details — https://www.syngoportal.org/gene_hgnc10307.html
- **[S4]** 
            Sexually dimorphic expression of the genes encoding ribosomal proteins L17 and L37 in the song control nuclei of juvenile zebra finches - PMC
         — https://pmc.ncbi.nlm.nih.gov/articles/PMC2878125/
- **[S5]** Effects of estradiol on incorporation of new cells in the developing zebra finch song system: potential relationship to expression of ribosomal proteins L17 and L37 - PubMed — https://pubmed.ncbi.nlm.nih.gov/19373862/
- **[S6]** A functional role for intra-axonal protein synthesis during axonal regeneration from adult sensory neurons - PubMed — https://pubmed.ncbi.nlm.nih.gov/11717363/
- **[S7]** Polyribosomes redistribute from dendritic shafts into spines with enlarged synapses during LTP in developing rat hippocampal slices - PubMed — https://pubmed.ncbi.nlm.nih.gov/12165474/?utm_source=openai
- **[S8]** RPL17 ribosomal protein L17 [Homo sapiens (human)] - Gene - NCBI — https://www.ncbi.nlm.nih.gov/gene/6139
- **[S9]** SynGO - annotation details — https://www.syngoportal.org/annotation_3553
- **[S10]** 
            An atypical form of 60S ribosomal subunit in Diamond-Blackfan anemia linked to RPL17 variants - PMC
         — https://pmc.ncbi.nlm.nih.gov/articles/PMC11385091/
- **[S11]** Homeostatic scaling is driven by a translation-dependent degradation axis that recruits miRISC remodeling - PubMed — https://pubmed.ncbi.nlm.nih.gov/34813590/?utm_source=openai
- **[S12]** Homeostatic scaling is driven by a translation-dependent degradation axis that recruits miRISC remodeling - PubMed — https://pubmed.ncbi.nlm.nih.gov/34813590/?utm_source=openai
- **[S13]** RPL17 ribosomal protein L17 [Homo sapiens (human)] - Gene - NCBI — https://www.ncbi.nlm.nih.gov/gene/6139
- **[S14]** Homeostatic scaling is driven by a translation-dependent degradation axis that recruits miRISC remodeling - PubMed — https://pubmed.ncbi.nlm.nih.gov/34813590/?utm_source=openai
- **[S15]** 
            Sexually dimorphic expression of the genes encoding ribosomal proteins L17 and L37 in the song control nuclei of juvenile zebra finches - PMC
         — https://pmc.ncbi.nlm.nih.gov/articles/PMC2878125/
