## Supplementary material for "Understanding Substance Dependence: What Differentiates Addictive from Non-Addictive Drugs?": Folder for supplementary files: non-addictive-genes-claims-first100-030-hilpda-gene-protein-has-been-associated-with-regul-338d208e3977.md

# non-addictive-genes-claims-first100-030-338d208e3977

**Original claim:** HILPDA gene/protein has been associated with regulatory or homeostatic biological processes in neural systems.

## Claim

HILPDA gene/protein has been associated with regulatory or homeostatic biological processes in neural systems.

## Verdict

Verdict: Likely True

## Confidence

Medium

## Supporting evidence

HILPDA is expressed in multiple brain-relevant cell types. The Human Protein Atlas lists HILPDA as “cell type enhanced” in several CNS populations, including astrocytes, Bergmann glia, ependymal cells, oligodendrocytes, oligodendrocyte precursor cells, excitatory neurons, inhibitory neurons, other brain neurons, and microglia. That does not prove function, but it establishes that the gene is present in neural-system cell types where homeostatic regulation occurs. ([proteinatlas.org](https://www.proteinatlas.org/ENSG00000135245-HILPDA/single%2Bcell)) [S1]

There is direct evidence that HILPDA is induced in human astrocytes by a physiologically relevant homeostatic stressor: hypoxia. In primary human astrocytes exposed to hypoxia for 24 hours, HILPDA was among the significantly upregulated hypoxia-responsive transcripts, with reported fold-changes of about 4.4–6.1 in microarray probes. The same study found broad hypoxia-driven changes in metabolism and immune-response pathways in astrocytes, placing HILPDA within an astrocytic stress-adaptation program rather than an unrelated expression change. ([eprints.whiterose.ac.uk](https://eprints.whiterose.ac.uk/id/eprint/167666/1/Transcriptomic%20analysis%20human%20astrocytes%20hypoxia%20induced%20dysregulation%20Allen%20Int%20J%20Mol%20Sci%202020.pdf)) [S2]

The strongest functional neural evidence comes from a mouse sepsis-associated encephalopathy model. In that study, hippocampal astrocytic HILPDA increased during disease, and astrocyte-targeted HILPDA knockdown reduced astrocytic lipid-droplet accumulation, decreased conversion toward the neurotoxic A1 astrocyte phenotype, reduced abnormal neuronal membrane excitability, and improved spatial/contextual memory measures. This is fairly direct evidence that HILPDA can regulate a glial metabolic process with downstream consequences for neural circuit function and cognitive outcome. ([pmc.ncbi.nlm.nih.gov](https://pmc.ncbi.nlm.nih.gov/articles/PMC11099789/)) [S3]

Additional support comes from ischemic-stroke work on microglia. After brain ischemia in mice, sorted microglia upregulated a lipid-droplet biogenesis program that included Hilpda, and the same study linked lipid-laden microglia to impaired responses to ischemia in aged animals. This is not a HILPDA-perturbation experiment, so it is associative rather than causal, but it still places Hilpda within a neural immune/metabolic response relevant to brain tissue homeostasis after injury. ([pmc.ncbi.nlm.nih.gov](https://pmc.ncbi.nlm.nih.gov/articles/PMC9906381/)) [S4]

Broader nervous-system literature also supports the relevance of lipid-droplet regulation to neural homeostasis. Reviews of neuroglial lipid metabolism describe brain lipid handling as tightly linked to energy homeostasis, oxidative stress, neuroinflammation, and neuron–glia coupling, and note that glial lipid droplets can be protective by buffering excess or peroxidized lipids. Within that framework, HILPDA is discussed as a candidate regulator of lipid-droplet turnover in neural cells, especially astrocytes under hypoxia. ([cellandbioscience.biomedcentral.com](https://cellandbioscience.biomedcentral.com/articles/10.1186/s13578-022-00828-0)) [S5]

## Contradicting or limiting evidence

The evidence is not broad enough to say HILPDA is a well-established regulator of *general* neural homeostasis. Most direct mechanistic work on HILPDA has been done outside the nervous system—in hepatocytes, adipocytes, macrophages, and cancer models—where it regulates lipid storage and lipolysis. Those studies are biologically informative, but they do not by themselves prove a CNS-specific homeostatic role. ([pubmed.ncbi.nlm.nih.gov](https://pubmed.ncbi.nlm.nih.gov/29326160/?utm_source=openai)) [S6]

Within neural systems specifically, the evidence is concentrated in glia and in stress/pathology contexts such as hypoxia, sepsis-associated encephalopathy, and ischemic stroke. I did not find strong primary evidence showing that HILPDA is a canonical regulator of baseline synaptic homeostasis, neurotransmitter clearance, ion buffering, myelin maintenance, or developmental neural patterning under healthy conditions. A nervous-system review explicitly notes that it is still unclear whether HILPDA functions in neurons the way it does in hepatocytes, even though it is expressed in astrocytes and induced by hypoxia. ([frontiersin.org](https://www.frontiersin.org/journals/cell-and-developmental-biology/articles/10.3389/fcell.2022.863907/pdf)) [S7]

The stroke-related microglial data are also limited by causality: Hilpda was part of an induced transcriptional program, but the study did not directly manipulate Hilpda in microglia to show that it independently controls post-stroke recovery or inflammatory tone. So for microglia, the current literature supports association more than proof of mechanism. ([pmc.ncbi.nlm.nih.gov](https://pmc.ncbi.nlm.nih.gov/articles/PMC9906381/)) [S8]

## Mechanistic / biological rationale

HILPDA is a small lipid-droplet-associated protein induced by hypoxia and other metabolic stresses. Primary studies show it is a target of HIF-1 and PPAR pathways, localizes to lipid droplets/endoplasmic reticulum interfaces, inhibits adipose triglyceride lipase (ATGL), and can also promote DGAT1-dependent triglyceride storage. In plain terms, HILPDA shifts cells toward retaining neutral lipids in lipid droplets rather than rapidly hydrolyzing them. ([pubmed.ncbi.nlm.nih.gov](https://pubmed.ncbi.nlm.nih.gov/20624928/?utm_source=openai)) [S9]

That mechanism is plausibly relevant to neural homeostasis because glia are major lipid-buffering cells in the brain. Astrocytes and microglia help manage excess fatty acids, oxidized lipids, inflammatory lipid mediators, and metabolic stress; dysregulation of these processes is linked to neuroinflammation and neurodegeneration. Reviews of neuroglial lipid biology describe lipid droplets as hubs for energy storage, oxidative-stress buffering, and cell-stress adaptation in the brain. ([cellandbioscience.biomedcentral.com](https://cellandbioscience.biomedcentral.com/articles/10.1186/s13578-022-00828-0)) [S10]

Non-neural mechanistic studies strengthen this plausibility. In macrophages, Hilpda is crucial for neutral-lipid accumulation under hypoxia, supports resistance to lipid overload, limits reactive oxygen species after reoxygenation, and influences prostaglandin E2 production. In adipocytes, HILPDA participates in a negative-feedback loop that suppresses lipolysis and helps maintain lipid homeostasis under fatty-acid stress. These are classic homeostatic functions, and because microglia are tissue-resident macrophage-like cells and astrocytes also perform lipid-buffering roles, it is reasonable to infer a related stress-homeostatic role in neural tissue. That said, this last step is partly inference, not fully settled CNS-specific proof. ([pubmed.ncbi.nlm.nih.gov](https://pubmed.ncbi.nlm.nih.gov/28760743/?utm_source=openai)) [S11]

The astrocyte sepsis-encephalopathy study provides the best bridge from mechanism to neural phenotype: reducing astrocytic HILPDA reduced lipid-droplet accumulation and improved downstream neural function. That is consistent with HILPDA acting as a regulator of glial lipid homeostasis that secondarily affects neuronal excitability and cognition. ([pmc.ncbi.nlm.nih.gov](https://pmc.ncbi.nlm.nih.gov/articles/PMC11099789/)) [S12]

## Bottom line

The claim is **likely true**, but with an important qualifier: current evidence links HILPDA to **glial metabolic/stress-homeostatic regulation** in neural systems more than to broad, well-established “neural homeostasis” in the classic sense. The best-supported neural roles are in **astrocytes and microglia**, especially under **hypoxia, inflammation, ischemia, and sepsis-related brain injury**, where HILPDA is associated with lipid-droplet regulation and downstream effects on neuroinflammatory state, neuronal excitability, and cognitive outcome. Evidence for a major baseline role in healthy neurons or normal synaptic homeostasis remains limited. ([eprints.whiterose.ac.uk](https://eprints.whiterose.ac.uk/id/eprint/167666/1/Transcriptomic%20analysis%20human%20astrocytes%20hypoxia%20induced%20dysregulation%20Allen%20Int%20J%20Mol%20Sci%202020.pdf)) [S13]

## Cited sources

- **[S1]** Single cell type - HILPDA - The Human Protein Atlas — https://www.proteinatlas.org/ENSG00000135245-HILPDA/single%2Bcell
- **[S2]** Transcriptomic analysis of human astrocytes in vitro reveals hypoxia-induced mitochondrial dysfunction, modulation of metabolism, and dysregulation of the immune response — https://eprints.whiterose.ac.uk/id/eprint/167666/1/Transcriptomic%20analysis%20human%20astrocytes%20hypoxia%20induced%20dysregulation%20Allen%20Int%20J%20Mol%20Sci%202020.pdf
- **[S3]** 
            Astrocytic HILPDA promotes lipid droplets generation to drive cognitive dysfunction in mice with sepsis‐associated encephalopathy - PMC
         — https://pmc.ncbi.nlm.nih.gov/articles/PMC11099789/
- **[S4]** 
            Aged lipid‐laden microglia display impaired responses to stroke - PMC
         — https://pmc.ncbi.nlm.nih.gov/articles/PMC9906381/
- **[S5]** Lipid metabolism and storage in neuroglia: role in brain development and neurodegenerative diseases | Cell & Bioscience | Springer Nature Link — https://cellandbioscience.biomedcentral.com/articles/10.1186/s13578-022-00828-0
- **[S6]** Hypoxia-inducible lipid droplet-associated protein inhibits adipose triglyceride lipase - PubMed — https://pubmed.ncbi.nlm.nih.gov/29326160/?utm_source=openai
- **[S7]** fcell-2022-863907 1..12 — https://www.frontiersin.org/journals/cell-and-developmental-biology/articles/10.3389/fcell.2022.863907/pdf
- **[S8]** 
            Aged lipid‐laden microglia display impaired responses to stroke - PMC
         — https://pmc.ncbi.nlm.nih.gov/articles/PMC9906381/
- **[S9]** Hypoxia-inducible protein 2 is a novel lipid droplet protein and a specific target gene of hypoxia-inducible factor-1 - PubMed — https://pubmed.ncbi.nlm.nih.gov/20624928/?utm_source=openai
- **[S10]** Lipid metabolism and storage in neuroglia: role in brain development and neurodegenerative diseases | Cell & Bioscience | Springer Nature Link — https://cellandbioscience.biomedcentral.com/articles/10.1186/s13578-022-00828-0
- **[S11]** Hypoxia-inducible protein 2 Hig2/Hilpda mediates neutral lipid accumulation in macrophages and contributes to atherosclerosis in apolipoprotein E-deficient mice - PubMed — https://pubmed.ncbi.nlm.nih.gov/28760743/?utm_source=openai
- **[S12]** 
            Astrocytic HILPDA promotes lipid droplets generation to drive cognitive dysfunction in mice with sepsis‐associated encephalopathy - PMC
         — https://pmc.ncbi.nlm.nih.gov/articles/PMC11099789/
- **[S13]** Transcriptomic analysis of human astrocytes in vitro reveals hypoxia-induced mitochondrial dysfunction, modulation of metabolism, and dysregulation of the immune response — https://eprints.whiterose.ac.uk/id/eprint/167666/1/Transcriptomic%20analysis%20human%20astrocytes%20hypoxia%20induced%20dysregulation%20Allen%20Int%20J%20Mol%20Sci%202020.pdf
