## Supplementary material for "Understanding Substance Dependence: What Differentiates Addictive from Non-Addictive Drugs?": Folder for supplementary files: non-addictive-genes-claims-first100-031-fgfr2-gene-protein-has-been-associated-with-regula-06dfa6d8013e.md

# non-addictive-genes-claims-first100-031-06dfa6d8013e

**Original claim:** FGFR2 gene/protein has been associated with regulatory or homeostatic biological processes in neural systems.

## Claim

FGFR2 gene/protein has been associated with regulatory or homeostatic biological processes in neural systems.

## Verdict

Verdict: Highly Supported

Across multiple primary neuroscience studies, FGFR2 has been directly linked to regulatory functions in neural systems, including neural precursor self-renewal, cortical and hippocampal development, adult dentate neurogenesis, glial differentiation/positioning, astrocyte-neuron interaction, and oligodendrocyte/myelin regulation. Evidence for adult neural “homeostatic” roles is also present, though it is more context-dependent and comes mostly from rodent and injury/disease models rather than direct human physiology. ([pubmed.ncbi.nlm.nih.gov](https://pubmed.ncbi.nlm.nih.gov/20410112/)) [S1]

## Confidence

High

## Supporting evidence

A strong line of evidence comes from developmental forebrain studies. Conditional inactivation of **Fgfr2** in radial glial cells of the dorsal telencephalon reduced excitatory cortical neuron number, impaired medial prefrontal cortex formation, and was interpreted by the authors as reflecting reduced self-renewal of neuronal precursors in the ventricular zone. That is a direct association with a neural regulatory process rather than a mere expression correlation. ([pubmed.ncbi.nlm.nih.gov](https://pubmed.ncbi.nlm.nih.gov/20410112/)) [S2]

FGFR2 is also associated with regulation of neural stem/progenitor compartments and adult hippocampal maintenance. In a mouse study separating developmental from adult deletion, loss of **Fgfr2** reduced hippocampal volume when deleted embryonically, impaired adult spatial memory, and decreased immature/newly generated dentate gyrus neurons; importantly, deficits in short-term learning and new dentate neurons were still seen when **Fgfr2** was deleted only in adulthood. That supports an ongoing role in adult hippocampal neurogenesis, not just embryonic patterning. ([pubmed.ncbi.nlm.nih.gov](https://pubmed.ncbi.nlm.nih.gov/22541947/)) [S3]

FGFR2 has also been tied to glial regulatory programs in several neural cell types. In the developing cerebellum, conditional mouse studies found FGFR2 signaling was required for proper generation and positioning of Bergmann glia and for cell survival in the cerebellar primordium. Because Bergmann glia organize cerebellar layering and neuron-glia interactions, this is a clear neural regulatory role. ([pubmed.ncbi.nlm.nih.gov](https://pubmed.ncbi.nlm.nih.gov/24983448/)) [S4]

In postnatal brain, FGFR2 appears especially relevant to astroglial biology. One mouse study reported that FGFR2 is almost exclusively expressed in glial cells in postnatal brain, and that early postnatal astroglial loss of **Fgfr2** reduced astrocyte-neuron membrane contact and increased glial glutamine synthetase expression, alongside behavioral dysregulation. Those findings connect FGFR2 to astrocyte-dependent regulation of synaptic environment and neurotransmitter-cycle-related machinery, both of which are plausibly homeostatic functions. ([pubmed.ncbi.nlm.nih.gov](https://pubmed.ncbi.nlm.nih.gov/36906620/)) [S5]

A complementary anatomical study in rat brain found robust FGFR2 expression in astrocytes, especially in the hippocampus, cerebellum, and neurogenic regions such as the subventricular zone and dentate subgranular zone. The same study observed enhanced astrocytic FGFR2 expression around a perforant pathway lesion, suggesting participation in injury-responsive regulation and possibly repair-related homeostasis. ([pubmed.ncbi.nlm.nih.gov](https://pubmed.ncbi.nlm.nih.gov/16856175/)) [S6]

FGFR2 is also implicated in oligodendrocyte and myelin regulation. A mouse study reported that signaling through **FGFR2**, but not FGFR1, regulates myelin thickness via ERK1/2-MAPK and mTORC1-related signaling in the oligodendrocyte/myelin compartment. Since myelin thickness is a key determinant of axonal conduction and circuit stability, this is strong evidence that FGFR2 participates in regulatory maintenance of neural infrastructure. ([pubmed.ncbi.nlm.nih.gov](https://pubmed.ncbi.nlm.nih.gov/28193689/)) [S7]

Disease-model work reinforces a homeostatic interpretation. In experimental autoimmune encephalomyelitis, oligodendrocyte-specific deletion of **Fgfr2** led to a milder disease course, less myelin damage, greater axonal density, and lower inflammatory signaling. In a Parkinson’s disease model, astrocyte FGF7/FGFR2 autocrine signaling intensified astrocyte reactivity and neuroinflammation, and astrocytic **Fgfr2** knockdown was beneficial. These studies do not prove FGFR2 is always protective; instead, they show FGFR2 is mechanistically involved in regulating inflammatory and repair-related neural states. ([pubmed.ncbi.nlm.nih.gov](https://pubmed.ncbi.nlm.nih.gov/33103299/)) [S8]

## Contradicting or limiting evidence

The main limitation is not absence of evidence, but **context dependence**. FGFR2’s roles differ by cell type, developmental stage, and disease setting. For example, an older oligodendrocyte-specific knockout study found no obvious baseline defects in oligodendrocyte number, onset/extent of myelination, myelin ultrastructure, or myelin protein expression after **Fgfr2** loss alone. That tempers any claim that FGFR2 is universally required for all baseline glial homeostasis. ([pubmed.ncbi.nlm.nih.gov](https://pubmed.ncbi.nlm.nih.gov/17122059/)) [S9]

Also, much of the strongest evidence is from **mouse or rat** genetics, histology, and injury models rather than human interventional data. The adult-homeostasis angle is therefore somewhat less direct than the developmental-regulation angle. In other words, FGFR2’s role in **regulatory neural processes** is very well supported, while its role in steady-state adult neural **homeostasis** is supported but more inferential and model-dependent. ([pubmed.ncbi.nlm.nih.gov](https://pubmed.ncbi.nlm.nih.gov/20410112/)) [S10]

Finally, some pathways are not uniquely FGFR2-specific. In several neural contexts, FGF signaling involves overlap or compensation among FGFR family members, so not every FGF-related neural phenotype can be attributed solely to FGFR2 without direct receptor-specific manipulation. The strongest conclusions therefore come from the FGFR2-specific conditional knockout/knockdown studies above. ([pubmed.ncbi.nlm.nih.gov](https://pubmed.ncbi.nlm.nih.gov/20410112/)) [S11]

## Mechanistic / biological rationale

Biologically, the association makes sense. FGFR2 is a fibroblast growth factor receptor with tyrosine-kinase signaling properties, and neural studies link it to canonical growth-factor pathways that regulate cell fate, proliferation, survival, and differentiation. In cortical progenitors, FGFR2 promotes precursor self-renewal and proper excitatory neuron production; in hippocampus, it supports dentate neurogenesis and related learning phenotypes; in Bergmann glia, it affects generation and positioning; and in oligodendrocytes it signals through ERK1/2 and mTORC1-related mechanisms to control myelin growth. ([pubmed.ncbi.nlm.nih.gov](https://pubmed.ncbi.nlm.nih.gov/16856175/)) [S12]

In astrocytes, FGFR2-associated changes in astrocyte-neuron contact, glutamine synthetase levels, and inflammatory reactivity provide a plausible route by which FGFR2 can influence neural homeostasis more broadly—by modulating the extracellular milieu, neurotransmitter cycling, and the injury-response state of glia. That is especially relevant because astrocytes are central regulators of synaptic and metabolic stability in the CNS. ([pubmed.ncbi.nlm.nih.gov](https://pubmed.ncbi.nlm.nih.gov/36906620/)) [S13]

## Bottom line

Yes. The claim is well supported: **FGFR2 has repeatedly been associated with regulatory processes in neural systems**, and there is also meaningful evidence linking it to **homeostatic-like glial and myelin functions**, especially in astrocytes, oligodendrocytes, neurogenic niches, and injury/inflammation models. The strongest caveat is that most evidence is from animal experiments, and adult steady-state homeostasis is less directly established than developmental and injury-responsive regulation. ([pubmed.ncbi.nlm.nih.gov](https://pubmed.ncbi.nlm.nih.gov/20410112/)) [S14]

## Cited sources

- **[S1]** Fgfr2 is required for the development of the medial prefrontal cortex and its connections with limbic circuits - PubMed — https://pubmed.ncbi.nlm.nih.gov/20410112/
- **[S2]** Fgfr2 is required for the development of the medial prefrontal cortex and its connections with limbic circuits - PubMed — https://pubmed.ncbi.nlm.nih.gov/20410112/
- **[S3]** Learning and memory depend on fibroblast growth factor receptor 2 functioning in hippocampus - PubMed — https://pubmed.ncbi.nlm.nih.gov/22541947/
- **[S4]** FGF/FGFR2 signaling regulates the generation and correct positioning of Bergmann glia cells in the developing mouse cerebellum - PubMed — https://pubmed.ncbi.nlm.nih.gov/24983448/
- **[S5]** Neonatal loss of FGFR2 in astroglial cells affects locomotion, sociability, working memory, and glia-neuron interactions in mice - PubMed — https://pubmed.ncbi.nlm.nih.gov/36906620/
- **[S6]** Cytoarchitecture of fibroblast growth factor receptor 2 (FGFR-2) immunoreactivity in astrocytes of neurogenic and non-neurogenic regions of the young adult and aged rat brain - PubMed — https://pubmed.ncbi.nlm.nih.gov/16856175/
- **[S7]** Signaling by FGF Receptor 2, Not FGF Receptor 1, Regulates Myelin Thickness through Activation of ERK1/2-MAPK, Which Promotes mTORC1 Activity in an Akt-Independent Manner - PubMed — https://pubmed.ncbi.nlm.nih.gov/28193689/
- **[S8]** Oligodendrocyte-specific deletion of FGFR2 ameliorates MOG35-55 -induced EAE through ERK and Akt signalling - PubMed — https://pubmed.ncbi.nlm.nih.gov/33103299/
- **[S9]** Mice with conditional inactivation of fibroblast growth factor receptor-2 signaling in oligodendrocytes have normal myelin but display dramatic hyperactivity when combined with Cnp1 inactivation - PubMed — https://pubmed.ncbi.nlm.nih.gov/17122059/
- **[S10]** Fgfr2 is required for the development of the medial prefrontal cortex and its connections with limbic circuits - PubMed — https://pubmed.ncbi.nlm.nih.gov/20410112/
- **[S11]** Fgfr2 is required for the development of the medial prefrontal cortex and its connections with limbic circuits - PubMed — https://pubmed.ncbi.nlm.nih.gov/20410112/
- **[S12]** Cytoarchitecture of fibroblast growth factor receptor 2 (FGFR-2) immunoreactivity in astrocytes of neurogenic and non-neurogenic regions of the young adult and aged rat brain - PubMed — https://pubmed.ncbi.nlm.nih.gov/16856175/
- **[S13]** Neonatal loss of FGFR2 in astroglial cells affects locomotion, sociability, working memory, and glia-neuron interactions in mice - PubMed — https://pubmed.ncbi.nlm.nih.gov/36906620/
- **[S14]** Fgfr2 is required for the development of the medial prefrontal cortex and its connections with limbic circuits - PubMed — https://pubmed.ncbi.nlm.nih.gov/20410112/
