## Supplementary material for "Understanding Substance Dependence: What Differentiates Addictive from Non-Addictive Drugs?": Folder for supplementary files: non-addictive-genes-claims-first100-032-cadm3-gene-protein-has-been-associated-with-regula-8974242bec68.md

# non-addictive-genes-claims-first100-032-8974242bec68

**Original claim:** CADM3 gene/protein has been associated with regulatory or homeostatic biological processes in neural systems.

## Claim

CADM3 gene/protein has been associated with regulatory or homeostatic biological processes in neural systems. ([ncbi.nlm.nih.gov](https://www.ncbi.nlm.nih.gov/gene/57863)) [S1]

## Verdict

Verdict: Likely True

## Confidence

Medium. The evidence that CADM3 (also called Necl-1/SynCAM3) participates in **regulatory neural processes** is fairly strong across multiple experimental systems, especially for synapse formation, axon-glia interactions, myelination, and axonal domain organization. Evidence for **homeostasis** is present but narrower and is strongest in developmental retina models rather than adult mammalian neural homeostasis. ([pubmed.ncbi.nlm.nih.gov](https://pubmed.ncbi.nlm.nih.gov/38623325/)) [S2]

## Supporting evidence

CADM3 is a neural-enriched adhesion molecule, which makes a neural regulatory role biologically plausible from the outset. NCBI lists CADM3 as brain-biased, and the Human Protein Atlas reports high expression in neuropil/peripheral nerves with “synapse in neurons” as the brain cellular location. Early primary work also showed Necl-1/CADM3 is specifically expressed in neural tissue and localizes at contact sites among axons, terminals, and glial processes that form synapses, axon bundles, and myelinated axons. ([ncbi.nlm.nih.gov](https://www.ncbi.nlm.nih.gov/gene/57863)) [S3]

There is direct evidence that CADM3 regulates synaptic organization in a neural circuit. In a 2024 mouse retina study, Necl-1/CADM3 was localized to cones and type 4 OFF cone bipolar cells; knockout mice showed dislocated cone-bipolar synapses, abnormal horizontal-cell distribution, AMPA receptor mislocalization, altered short-wavelength visual signal transmission, and impaired optokinetic responses. The authors concluded that Necl-1 regulates cone synapse formation in the OFF cone pathway. ([pubmed.ncbi.nlm.nih.gov](https://pubmed.ncbi.nlm.nih.gov/38623325/)) [S4]

There is also substantial evidence for a regulatory role in myelination and axon-glia interaction. In peripheral nerve cocultures, axonal Necl-1/Cadm3 binds Schwann-cell Necl-4/Cadm4, and perturbing this adhesion system blocks Schwann-cell differentiation/myelination, identifying the pathway as important for initiation of myelination. Later genetic work showed that Cadm family proteins regulate myelination and the organization of myelinated axons, with Cadm3 identified as the main axonal ligand for glial Cadm4 in the PNS. ([pubmed.ncbi.nlm.nih.gov](https://pubmed.ncbi.nlm.nih.gov/17724124/)) [S5]

A 2008 mouse knockout study further showed that disrupting Necl-1 caused a **developmental delay** of axonal myelination in optic nerve and spinal cord, supporting a role in CNS myelination timing and axon-oligodendrocyte recognition/adhesion. That is a regulatory developmental phenotype within neural tissue, even if not a classic “homeostasis” phenotype. ([pubmed.ncbi.nlm.nih.gov](https://pubmed.ncbi.nlm.nih.gov/19036974/?utm_source=openai)) [S6]

CADM3 also shows more explicitly regulatory signaling behavior in vitro. In Schwann cell/DRG neuron myelinating cultures, knockdown of axonal Cadm3 promoted myelination, whereas overexpression of Cadm3 nearly abolished myelin segment formation; the extracellular domain interfered with ErbB3 and pro-myelinating PI3K/Akt signaling. The authors therefore proposed Cadm3 as a negative regulator of PNS myelination. ([pubmed.ncbi.nlm.nih.gov](https://pubmed.ncbi.nlm.nih.gov/27658374/?utm_source=openai)) [S7]

For the “homeostatic” part of the claim, there is at least one direct and relevant study in neural tissue. In developing zebrafish retina, homophilic Cadm3 interaction coordinated retinal progenitor proliferation, and the paper explicitly concluded that Cadm3 helps ensure **cell number homeostasis** of the developing neuroepithelium. That is bona fide homeostatic language applied to a neural developmental system. ([pmc.ncbi.nlm.nih.gov](https://pmc.ncbi.nlm.nih.gov/articles/PMC10082328/)) [S8]

Human genetics also supports neural functional relevance. Pathogenic CADM3 variants have been linked to Charcot-Marie-Tooth disease, and functional studies in cells and mice pointed to disrupted axon-glia interaction and abnormal axonal organization as the disease mechanism. This does not prove “homeostasis,” but it strengthens the case that CADM3 is functionally important in regulatory organization of neural systems rather than being a passive marker. ([pmc.ncbi.nlm.nih.gov](https://pmc.ncbi.nlm.nih.gov/articles/PMC8105037/)) [S9]

## Contradicting or limiting evidence

The main limitation is that much of the strongest functional evidence comes from **mouse, zebrafish, and in vitro systems**, not direct human mechanistic studies. The human evidence is largely genetic/disease-association based. ([pubmed.ncbi.nlm.nih.gov](https://pubmed.ncbi.nlm.nih.gov/38623325/)) [S10]

A second limitation is redundancy. Single Cadm3 knockout or loss can produce mild or even absent peripheral myelin phenotypes, whereas combined loss of multiple axonal Cadms reveals stronger abnormalities. This means CADM3 is functionally relevant, but not always singly essential, which weakens very strong causal wording if one were claiming a unique or dominant homeostatic role. ([pubmed.ncbi.nlm.nih.gov](https://pubmed.ncbi.nlm.nih.gov/33397712/)) [S11]

A third limitation is terminologic. Most papers describe CADM3 in terms of **cell adhesion, synapse formation, axon-glia recognition, myelination, and axonal organization**. Far fewer frame it explicitly as a regulator of neural “homeostasis,” especially in adult nervous systems. The clearest homeostasis evidence is developmental retinal cell-number homeostasis in zebrafish, which may not generalize to all neural systems or to adult neural homeostatic plasticity. ([pmc.ncbi.nlm.nih.gov](https://pmc.ncbi.nlm.nih.gov/articles/PMC10082328/)) [S12]

## Mechanistic / biological rationale

Mechanistically, CADM3 is well positioned to mediate regulatory functions because it is a calcium-independent cell-cell adhesion molecule concentrated in neural tissue and at neural contact sites. It can mediate homophilic adhesion and also heterophilic interactions with other nectin-like/Cadm proteins, especially the axon-glia pair Cadm3-Cadm4. Its cytoplasmic tail can associate with scaffolding proteins including Dlg3, Pals2, and CASK, and separate work showed it recruits protein 4.1N to the plasma membrane; together these interactions provide a route from extracellular adhesion to intracellular scaffolding/cytoskeletal organization. ([pubmed.ncbi.nlm.nih.gov](https://pubmed.ncbi.nlm.nih.gov/15741237/)) [S13]

That architecture fits the observed phenotypes: if CADM3 helps stabilize specific neuron-neuron or neuron-glia contacts, then altering CADM3 would be expected to affect synapse placement, receptor localization, axonal domain organization, and the timing or extent of myelination. The 2016 myelination study adds a signaling layer by linking Cadm3 to selective modulation of ErbB3/PI3K-Akt signaling, offering a plausible mechanism for how an adhesion molecule can act as a regulator rather than merely a structural glue. ([pubmed.ncbi.nlm.nih.gov](https://pubmed.ncbi.nlm.nih.gov/38623325/)) [S14]

The zebrafish retina study extends this logic to tissue-level homeostasis: homophilic Cadm3-mediated cell-cell interaction can convey local information about neighboring progenitor state and thereby tune proliferation to maintain appropriate neuroepithelial cell numbers. That is consistent with a broader model in which CADM3 participates in local regulatory feedback within neural tissues. ([pmc.ncbi.nlm.nih.gov](https://pmc.ncbi.nlm.nih.gov/articles/PMC10082328/)) [S15]

## Bottom line

The claim is **likely true**. CADM3 has repeatedly been linked to **regulatory processes in neural systems**, especially synapse formation, axon-glia adhesion, myelination, and organization of myelinated axons. There is also **some direct homeostasis-related evidence**, most notably Cadm3-dependent control of retinal neuroepithelial cell-number homeostasis in zebrafish. The caveat is that the “homeostatic” label is less broadly established than the “regulatory/developmental” one, and much of the strongest evidence comes from animal and cell models rather than direct human physiology. ([pubmed.ncbi.nlm.nih.gov](https://pubmed.ncbi.nlm.nih.gov/38623325/)) [S16]

## Cited sources

- **[S1]** CADM3 cell adhesion molecule 3 [Homo sapiens (human)] - Gene - NCBI — https://www.ncbi.nlm.nih.gov/gene/57863
- **[S2]** Necl-1/CADM3 regulates cone synapse formation in the mouse retina - PubMed — https://pubmed.ncbi.nlm.nih.gov/38623325/
- **[S3]** CADM3 cell adhesion molecule 3 [Homo sapiens (human)] - Gene - NCBI — https://www.ncbi.nlm.nih.gov/gene/57863
- **[S4]** Necl-1/CADM3 regulates cone synapse formation in the mouse retina - PubMed — https://pubmed.ncbi.nlm.nih.gov/38623325/
- **[S5]** Nectin-like proteins mediate axon Schwann cell interactions along the internode and are essential for myelination - PubMed — https://pubmed.ncbi.nlm.nih.gov/17724124/
- **[S6]** Disruption of Nectin-like 1 cell adhesion molecule leads to delayed axonal myelination in the CNS - PubMed — https://pubmed.ncbi.nlm.nih.gov/19036974/?utm_source=openai
- **[S7]** Cadm3 (Necl-1) interferes with the activation of the PI3 kinase/Akt signaling cascade and inhibits Schwann cell myelination in vitro - PubMed — https://pubmed.ncbi.nlm.nih.gov/27658374/?utm_source=openai
- **[S8]** 
            Homophilic interaction of cell adhesion molecule 3 coordinates retina neuroepithelial cell proliferation - PMC
         — https://pmc.ncbi.nlm.nih.gov/articles/PMC10082328/
- **[S9]** 
            A CADM3 variant causes Charcot-Marie-Tooth disease with marked upper limb involvement - PMC
         — https://pmc.ncbi.nlm.nih.gov/articles/PMC8105037/
- **[S10]** Necl-1/CADM3 regulates cone synapse formation in the mouse retina - PubMed — https://pubmed.ncbi.nlm.nih.gov/38623325/
- **[S11]** Differential Contribution of Cadm1-Cadm3 Cell Adhesion Molecules to Peripheral Myelinated Axons - PubMed — https://pubmed.ncbi.nlm.nih.gov/33397712/
- **[S12]** 
            Homophilic interaction of cell adhesion molecule 3 coordinates retina neuroepithelial cell proliferation - PMC
         — https://pmc.ncbi.nlm.nih.gov/articles/PMC10082328/
- **[S13]** Nectin-like molecule-1/TSLL1/SynCAM3: a neural tissue-specific immunoglobulin-like cell-cell adhesion molecule localizing at non-junctional contact sites of presynaptic nerve terminals, axons and glia cell processes - PubMed — https://pubmed.ncbi.nlm.nih.gov/15741237/
- **[S14]** Necl-1/CADM3 regulates cone synapse formation in the mouse retina - PubMed — https://pubmed.ncbi.nlm.nih.gov/38623325/
- **[S15]** 
            Homophilic interaction of cell adhesion molecule 3 coordinates retina neuroepithelial cell proliferation - PMC
         — https://pmc.ncbi.nlm.nih.gov/articles/PMC10082328/
- **[S16]** Necl-1/CADM3 regulates cone synapse formation in the mouse retina - PubMed — https://pubmed.ncbi.nlm.nih.gov/38623325/
