## Supplementary material for "Understanding Substance Dependence: What Differentiates Addictive from Non-Addictive Drugs?": Folder for supplementary files: non-addictive-genes-claims-first100-033-mknk2-gene-protein-has-been-associated-with-regula-67882e256024.md

# non-addictive-genes-claims-first100-033-67882e256024

**Original claim:** MKNK2 gene/protein has been associated with regulatory or homeostatic biological processes in neural systems.

## Claim

MKNK2 gene/protein has been associated with regulatory or homeostatic biological processes in neural systems.

## Verdict

Verdict: Likely True

The strongest support is for **regulatory roles** in neural systems—especially regulation of local/synaptic translation, synaptic proteome composition, nociceptor-related signaling, and microglial inflammatory state—rather than for a narrowly defined, experimentally established role in **classical neural homeostasis**. ([pubmed.ncbi.nlm.nih.gov](https://pubmed.ncbi.nlm.nih.gov/41688788/)) [S1]

## Confidence

Medium

There is credible and growing evidence that MKNK2 is involved in neural regulatory biology, but direct evidence specifically tying **MKNK2 alone** to canonical **homeostatic plasticity/homeostasis** is limited, and some neural phenotypes are better established for **MNK1/2 jointly** or are more strongly attributed to **MNK1** than MKNK2. ([pubmed.ncbi.nlm.nih.gov](https://pubmed.ncbi.nlm.nih.gov/41688788/)) [S2]

## Supporting evidence

MKNK2 is clearly present in neural tissues, which is a prerequisite for any neural regulatory role. The Human Protein Atlas reports MKNK2 RNA detected across all major human brain regions analyzed, with low regional specificity but broad distribution, and recent mouse work cited by the same field places Mknk2 mRNA in multiple neuronal populations and even in synaptic neuropil. ([proteinatlas.org](https://www.proteinatlas.org/ENSG00000099875-MKNK2/brain)) [S3]

The most direct evidence comes from a 2026 *Molecular Psychiatry* study examining **single-gene knockout mice**. In that work, loss of **MNK2** altered social/cognitive behavioral profiles and produced distinct changes in the **cortical synaptic proteome, transcriptome, and phosphoproteome**. Specifically, MNK2 deletion reduced expression and phosphorylation of synaptic proteins, and the authors concluded that MNK1 and MNK2 have specialized functions in synaptic compartments. That is strong evidence that MKNK2 is associated with **regulatory processes in neural systems**, particularly synaptic protein regulation. ([pubmed.ncbi.nlm.nih.gov](https://pubmed.ncbi.nlm.nih.gov/41688788/)) [S4]

The same study strengthens the association mechanistically: it states that local mRNA translation drives adaptive synaptic proteome changes, identifies MNK1/2 as key regulators of neuronal translation, and suggests that **MNK2 regulates translation of a pool of mRNAs including synaptic proteins**, potentially via mechanisms partly distinct from steady-state eIF4E phosphorylation. This directly places MKNK2 in neural translational regulation rather than merely as a passive brain-expressed kinase. ([pubmed.ncbi.nlm.nih.gov](https://pubmed.ncbi.nlm.nih.gov/41688788/)) [S5]

Human sensory-neuron data also support neural relevance. A 2023 *Neuroscience* paper found that **both MKNK1 and MKNK2 are expressed by nearly all neurons in human dorsal root and trigeminal ganglia**, with signal also in other cell types. The authors frame MNKs as important to translation via eIF4E and note prior implication of MNK signaling in regulation of nociceptor excitability and nociceptive behaviors. While that paper does not prove MKNK2-specific function in pain, it shows MKNK2 is positioned within human neural circuits where regulatory signaling is occurring. ([pubmed.ncbi.nlm.nih.gov](https://pubmed.ncbi.nlm.nih.gov/36764601/)) [S6]

There is also disease-context evidence in CNS immune cells. In a 2025 rat ischemic-stroke study, MKNK2 was expressed in neurons, astrocytes, oligodendrocytes, and microglia, with the most prominent post-injury upregulation in microglia. Knockdown of MKNK2 reduced HIF-1α signaling and shifted microglia away from pro-inflammatory M1 polarization. This is not “homeostasis” in the narrow synaptic-scaling sense, but it is a clear example of MKNK2 participating in **regulatory state control inside a neural system**. ([pmc.ncbi.nlm.nih.gov](https://pmc.ncbi.nlm.nih.gov/articles/PMC12417964/)) [S7]

More broadly, MNK-family literature supports a role in brain-regulatory processes such as synaptic plasticity, memory, and autism-related behaviors. A 2022 study found that Mnk1/2 double knockout impairs synaptic plasticity and memory, and linked this to the brain protein SYNGAP1 and mTORC1-related translational control. Because that work emphasizes shared MNK signaling and particularly highlights **Mnk1** as a downstream kinase for some effects, it is indirect for MKNK2 specifically—but it reinforces the biological plausibility that MKNK2 participates in neural regulatory translation programs. ([pmc.ncbi.nlm.nih.gov](https://pmc.ncbi.nlm.nih.gov/articles/PMC10411928/)) [S8]

## Contradicting or limiting evidence

The main limitation is that **direct MKNK2-specific evidence is still relatively sparse**. Many neural studies analyze **MNK1/2 together**, use double-knockout models, or focus on the shared MNK–eIF4E axis rather than isolating MKNK2. That means some of the neural-regulatory literature supports the MNK family more strongly than MKNK2 alone. ([pmc.ncbi.nlm.nih.gov](https://pmc.ncbi.nlm.nih.gov/articles/PMC10411928/)) [S9]

In sensory pain biology, the balance of evidence often points more strongly to **MKNK1** as the dominant isoform for regulating nociceptor excitability and nociceptive behavior. The human DRG/TG expression study explicitly notes that mouse knockout work suggests **Mknk1** is the key isoform in that setting, even though MKNK2 is co-expressed in human nociceptors. So neural association is present, but isoform-specific causality is not equally strong across all neural phenotypes. ([pubmed.ncbi.nlm.nih.gov](https://pubmed.ncbi.nlm.nih.gov/36764601/)) [S10]

The specific claim about **homeostatic** biological processes is also only partly supported. The literature retrieved here centers on **translation control, synaptic protein regulation, behavior, pain sensitization, and neuroinflammation**. I did not find comparably strong, explicit evidence that MKNK2 is a recognized regulator of classic **homeostatic plasticity**, synaptic scaling, or baseline neural homeostasis in the same direct way. ([pubmed.ncbi.nlm.nih.gov](https://pubmed.ncbi.nlm.nih.gov/41688788/)) [S11]

Historically, early knockout work also emphasized uncertainty about physiological function. In 2004, Mnk1/Mnk2 knockout experiments showed these kinases were essential for eIF4E phosphorylation but not for gross growth or development, and the physiological significance of eIF4E phosphorylation remained unclear. That does not negate newer neural findings, but it does argue against overstating how settled the biology is. ([pmc.ncbi.nlm.nih.gov](https://pmc.ncbi.nlm.nih.gov/articles/PMC444855/)) [S12]

## Mechanistic / biological rationale

MKNK2 encodes a MAPK-interacting serine/threonine kinase that lies downstream of ERK/p38 signaling and phosphorylates **eIF4E**, a core cap-binding translation factor. Older biochemical work indicates MNK2 has relatively high basal activity compared with MNK1, positioning it as a plausible constitutive regulator of translation in some contexts. ([pmc.ncbi.nlm.nih.gov](https://pmc.ncbi.nlm.nih.gov/articles/PMC444855/)) [S13]

In neurons, local and synaptic protein synthesis is a major way cells regulate synaptic composition, plasticity, and adaptive responses. Because MKNK2 is broadly present in brain tissue and synaptic compartments, and because MNK2 loss alters the synaptic proteome/phosphoproteome, the most plausible biological interpretation is that MKNK2 helps tune **which neural mRNAs are translated and/or phosphorylated at synapses**, thereby influencing circuit-relevant protein composition. ([proteinatlas.org](https://www.proteinatlas.org/ENSG00000099875-MKNK2/brain)) [S14]

Outside classical neurons, MKNK2 also appears capable of regulating **microglial state transitions** after injury, likely through translational control of HIF-1α-related inflammatory programs. In sensory neurons, MNK signaling is linked to local translation and excitability changes relevant to nociceptor sensitization. These mechanisms fit a broad definition of neural-system regulation, even if they are not all “homeostatic” in the strict textbook sense. ([pmc.ncbi.nlm.nih.gov](https://pmc.ncbi.nlm.nih.gov/articles/PMC12417964/)) [S15]

## Bottom line

Yes—**MKNK2 is reasonably associated with regulatory biological processes in neural systems**, especially **neuronal translation/synaptic protein regulation** and, in some contexts, **microglial inflammatory regulation**. The claim becomes weaker if interpreted to mean that MKNK2 has a well-established, specific role in **classical neural homeostasis/homeostatic plasticity**. So the fairest overall assessment is **Likely True**, with the caveat that the evidence is stronger for **neural regulation** than for **homeostasis per se**. ([pubmed.ncbi.nlm.nih.gov](https://pubmed.ncbi.nlm.nih.gov/41688788/)) [S16]

## Cited sources

- **[S1]** Distinct roles for MNK1 and MNK2 in social and cognitive behavior through kinase-specific regulation of the synaptic proteome and phosphoproteome - PubMed — https://pubmed.ncbi.nlm.nih.gov/41688788/
- **[S2]** Distinct roles for MNK1 and MNK2 in social and cognitive behavior through kinase-specific regulation of the synaptic proteome and phosphoproteome - PubMed — https://pubmed.ncbi.nlm.nih.gov/41688788/
- **[S3]** Brain tissue expression of MKNK2 - Summary - The Human Protein Atlas — https://www.proteinatlas.org/ENSG00000099875-MKNK2/brain
- **[S4]** Distinct roles for MNK1 and MNK2 in social and cognitive behavior through kinase-specific regulation of the synaptic proteome and phosphoproteome - PubMed — https://pubmed.ncbi.nlm.nih.gov/41688788/
- **[S5]** Distinct roles for MNK1 and MNK2 in social and cognitive behavior through kinase-specific regulation of the synaptic proteome and phosphoproteome - PubMed — https://pubmed.ncbi.nlm.nih.gov/41688788/
- **[S6]** MNK1 and MNK2 Expression in the Human Dorsal Root and Trigeminal Ganglion - PubMed — https://pubmed.ncbi.nlm.nih.gov/36764601/
- **[S7]** 
            KLF7 Blocks MKNK2/HIF‐1 Pathway‐Mediated M1 Microglia Polarization to Ameliorate Ischemic Stroke‐Induced Neurological Injury - PMC
         — https://pmc.ncbi.nlm.nih.gov/articles/PMC12417964/
- **[S8]** 
            Mnk1/2 kinases regulate memory and autism-related behaviours via Syngap1 - PMC
         — https://pmc.ncbi.nlm.nih.gov/articles/PMC10411928/
- **[S9]** 
            Mnk1/2 kinases regulate memory and autism-related behaviours via Syngap1 - PMC
         — https://pmc.ncbi.nlm.nih.gov/articles/PMC10411928/
- **[S10]** MNK1 and MNK2 Expression in the Human Dorsal Root and Trigeminal Ganglion - PubMed — https://pubmed.ncbi.nlm.nih.gov/36764601/
- **[S11]** Distinct roles for MNK1 and MNK2 in social and cognitive behavior through kinase-specific regulation of the synaptic proteome and phosphoproteome - PubMed — https://pubmed.ncbi.nlm.nih.gov/41688788/
- **[S12]** 
            Mnk2 and Mnk1 Are Essential for Constitutive and Inducible Phosphorylation of Eukaryotic Initiation Factor 4E but Not for Cell Growth or Development - PMC
         — https://pmc.ncbi.nlm.nih.gov/articles/PMC444855/
- **[S13]** 
            Mnk2 and Mnk1 Are Essential for Constitutive and Inducible Phosphorylation of Eukaryotic Initiation Factor 4E but Not for Cell Growth or Development - PMC
         — https://pmc.ncbi.nlm.nih.gov/articles/PMC444855/
- **[S14]** Brain tissue expression of MKNK2 - Summary - The Human Protein Atlas — https://www.proteinatlas.org/ENSG00000099875-MKNK2/brain
- **[S15]** 
            KLF7 Blocks MKNK2/HIF‐1 Pathway‐Mediated M1 Microglia Polarization to Ameliorate Ischemic Stroke‐Induced Neurological Injury - PMC
         — https://pmc.ncbi.nlm.nih.gov/articles/PMC12417964/
- **[S16]** Distinct roles for MNK1 and MNK2 in social and cognitive behavior through kinase-specific regulation of the synaptic proteome and phosphoproteome - PubMed — https://pubmed.ncbi.nlm.nih.gov/41688788/
