## Supplementary material for "Understanding Substance Dependence: What Differentiates Addictive from Non-Addictive Drugs?": Folder for supplementary files: non-addictive-genes-claims-first100-034-cirbp-gene-protein-has-been-associated-with-regula-c64e130abdf3.md

# non-addictive-genes-claims-first100-034-c64e130abdf3

**Original claim:** CIRBP gene/protein has been associated with regulatory or homeostatic biological processes in neural systems.

## Claim

CIRBP gene/protein has been associated with regulatory or homeostatic biological processes in neural systems.

## Verdict

Verdict: Likely True

There is solid **preclinical** evidence that CIRBP participates in neural regulatory/homeostatic processes, especially **sleep/circadian regulation** and **stress-responsive maintenance of neuronal integrity**. The strongest direct neural-homeostatic evidence is from mouse studies showing that Cirbp affects **cortical clock-gene responses to sleep deprivation** and **REM-sleep recovery**, i.e. sleep homeostasis. Additional animal and cell studies link CIRBP to regulation of **hippocampal synaptic structure**, **neural stem-cell proliferation under hypoxia**, and **neuronal iron/ferroptosis homeostasis**. However, most evidence is from rodents or cultured cells, not human CNS studies, and CIRBP can also be **pathogenic** in injured brain when released extracellularly. ([pubmed.ncbi.nlm.nih.gov](https://pubmed.ncbi.nlm.nih.gov/30720431/)) [S1]

## Confidence

High

Multiple independent studies directly connect CIRBP to neural regulatory phenomena, including **sleep homeostasis** and **post-transcriptional control of neural targets**, but the evidence base is still mostly experimental rather than clinical/human. ([pubmed.ncbi.nlm.nih.gov](https://pubmed.ncbi.nlm.nih.gov/30720431/)) [S2]

## Supporting evidence

CIRBP is, at baseline, a plausible regulator: the NCBI/Alliance annotation describes human CIRBP as an RNA-binding protein that **binds mRNA 3'-UTRs** and is involved in **mRNA stabilization** and **positive regulation of translation**. Those are classic molecular functions by which a gene can mediate homeostatic regulation in many tissues, including nervous tissue. ([ncbi.nlm.nih.gov](https://www.ncbi.nlm.nih.gov/gene/1153)) [S3]

The clearest direct evidence for a neural homeostatic role comes from sleep/circadian work in mice. In an eLife study, Cirbp expression in mouse brain rose during the sleep-dominant part of the day and fell during the wake-dominant interval; experimentally imposed sleep deprivation largely flattened that daily amplitude. In the same study, **Cirbp knockout** altered sleep-deprivation-induced cortical clock-gene responses and reduced REM rebound during recovery, with KO mice accruing only about half the extra REM sleep of wild-type mice. That is a direct association with a bona fide neural homeostatic process: **sleep homeostasis**. ([pubmed.ncbi.nlm.nih.gov](https://pubmed.ncbi.nlm.nih.gov/30720431/)) [S4]

CIRBP is also linked to circadian regulatory machinery at the post-transcriptional level. A mechanistic study found that Cirbp and Rbm3 are important regulators of **temperature-entrained circadian gene expression**; depletion of Cirbp reduced the amplitudes of core circadian genes, and PAR-CLIP/RNA-seq implicated binding near polyadenylation sites and control of **alternative polyadenylation**. This study was not primarily performed in neural tissue, so it is not by itself definitive for neural systems, but it strongly supports a regulatory role consistent with the mouse brain sleep/homeostasis findings. ([pubmed.ncbi.nlm.nih.gov](https://pubmed.ncbi.nlm.nih.gov/23792593/)) [S5]

Beyond sleep/circadian biology, CIRBP has been tied to neuronal structural maintenance. In a mouse hypobaric-hypoxia model, hippocampal dendritic spine density and cognition worsened under hypoxia, while PSD95—a key synaptic scaffolding protein—was shown to be **post-transcriptionally regulated by Cirbp through 3'-UTR binding**. Overexpressing Cirbp in hippocampus rescued abnormal PSD95 expression, reduced dendritic-spine injury, and improved cognitive readouts. This supports an association with regulatory/homeostatic processes that preserve synaptic architecture in neural tissue. ([pubmed.ncbi.nlm.nih.gov](https://pubmed.ncbi.nlm.nih.gov/34419133/)) [S6]

CIRBP has also been implicated in regulation of neural stem-cell behavior. In severe hypoxia, neural stem cells showed growth arrest, G0/G1 accumulation, and reduced Cyclin D1; CIRBP expression fell in parallel. Forced CIRBP expression restored NSC proliferation and Cyclin D1 under hypoxia, indicating a role in regulating cell-cycle homeostasis in a neural progenitor context. ([pubmed.ncbi.nlm.nih.gov](https://pubmed.ncbi.nlm.nih.gov/26927658/)) [S7]

More recently, a perinatal hypoxia study linked CIRBP to **neuronal iron/ferroptosis homeostasis**. In hypoxia-exposed hippocampus and neurons, the authors reported that CIRBP overexpression reduced ferroptotic injury and improved cognitive outcomes, with mechanistic evidence that CIRBP binds the **FTMT 3'-UTR** to stabilize its mRNA and promote translation. The same study explicitly framed hypoxia-induced CIRBP loss as important for **iron homeostasis** and nervous-system development. ([pubmed.ncbi.nlm.nih.gov](https://pubmed.ncbi.nlm.nih.gov/38388728/)) [S8]

## Contradicting or limiting evidence

The main limitation is that the evidence is overwhelmingly **preclinical**. The strongest papers are in mice, primary neurons, immortalized neuronal lines, or neural stem cells; direct evidence from human brain physiology is limited. So the claim is well supported as an **association in experimental neural systems**, but not yet strongly established in human neural homeostasis in vivo. ([pubmed.ncbi.nlm.nih.gov](https://pubmed.ncbi.nlm.nih.gov/30720431/)) [S9]

A second limitation is that many supportive findings arise under **perturbation** rather than normal steady state: sleep deprivation, hypobaric hypoxia, perinatal hypoxia, amyloid stress, ischemia, or trauma. That still counts as regulatory/homeostatic biology, but it means CIRBP is often being studied as a **stress-response regulator**, not as a constitutive master homeostat under ordinary conditions. ([pubmed.ncbi.nlm.nih.gov](https://pubmed.ncbi.nlm.nih.gov/30720431/)) [S10]

Importantly, CIRBP is **context-dependent** and not uniformly homeostatic. In cerebral ischemia, CIRP increased in mouse brain after MCAO; Cirbp deficiency reduced infarct volume, TNF-α induction, and microglial activation, and recombinant CIRP directly stimulated microglial TNF-α release and NF-κB signaling. The authors concluded that **extracellular CIRP is detrimental** in ischemic brain injury. ([pubmed.ncbi.nlm.nih.gov](https://pubmed.ncbi.nlm.nih.gov/24613680/)) [S11]

Similarly, in traumatic brain injury, neural-specific CIRP knockout reduced tissue loss, neuronal apoptosis, and glial activation, while extracellular CIRP promoted neuronal apoptosis via ER-stress signaling and correlated with inflammatory markers and injury biomarkers in mice and a small human patient sample. So CIRBP’s role depends heavily on **intracellular vs extracellular localization** and disease context. ([pubmed.ncbi.nlm.nih.gov](https://pubmed.ncbi.nlm.nih.gov/38817684/)) [S12]

## Mechanistic / biological rationale

Mechanistically, CIRBP is well positioned to regulate homeostatic processes because it is an RNA-binding protein with documented **3'-UTR binding**, **mRNA stabilization**, and **translational control** functions. Those mechanisms can tune protein output without requiring new transcription, which is exactly the kind of fast buffering used in homeostatic responses. ([ncbi.nlm.nih.gov](https://www.ncbi.nlm.nih.gov/gene/1153)) [S13]

In the sleep/circadian domain, the available data support a model in which CIRBP expression is influenced by behavioral state and temperature, then helps maintain the amplitude and proper response of **clock-gene expression** after sleep loss. That provides a plausible route by which CIRBP contributes to neural timing and recovery processes central to sleep homeostasis. ([pubmed.ncbi.nlm.nih.gov](https://pubmed.ncbi.nlm.nih.gov/30720431/)) [S14]

In neurons under stress, CIRBP appears to act by stabilizing specific transcripts relevant to structural or metabolic integrity. Examples include **PSD95** in hippocampal neurons, linking CIRBP to synaptic-spine maintenance, and **FTMT** in hypoxic neurons, linking CIRBP to iron handling and ferroptosis resistance. In neural stem cells, CIRBP restoration rescues **Cyclin D1** and proliferation under severe hypoxia, consistent with a role in preserving proliferative homeostasis in progenitor pools. ([pubmed.ncbi.nlm.nih.gov](https://pubmed.ncbi.nlm.nih.gov/34419133/)) [S15]

At the same time, when CIRP is released extracellularly after injury, it can function more like a danger signal than a homeostatic buffer, promoting microglial activation, inflammatory cytokine production, and neuronal death pathways. So the biology is not “CIRBP = always homeostatic”; rather, **intracellular CIRBP often looks regulatory/protective, whereas extracellular CIRP can be dysregulatory/pathogenic**. ([pubmed.ncbi.nlm.nih.gov](https://pubmed.ncbi.nlm.nih.gov/24613680/)) [S16]

## Bottom line

The claim is **likely true**. CIRBP has been associated—indeed experimentally linked—with **regulatory and homeostatic processes in neural systems**, most convincingly in **sleep homeostasis/cortical clock-gene regulation**, and also in **synaptic maintenance**, **neural stem-cell proliferation**, and **neuronal iron-stress control**. The caveat is that this evidence is mostly from **animal and cell models**, and CIRBP’s role is **highly context-dependent**, with extracellular CIRP becoming pro-inflammatory and injurious in brain disease models. ([pubmed.ncbi.nlm.nih.gov](https://pubmed.ncbi.nlm.nih.gov/30720431/)) [S17]

## Cited sources

- **[S1]** Cold-inducible RNA-binding protein (CIRBP) adjusts clock-gene expression and REM-sleep recovery following sleep deprivation - PubMed — https://pubmed.ncbi.nlm.nih.gov/30720431/
- **[S2]** Cold-inducible RNA-binding protein (CIRBP) adjusts clock-gene expression and REM-sleep recovery following sleep deprivation - PubMed — https://pubmed.ncbi.nlm.nih.gov/30720431/
- **[S3]** CIRBP cold inducible RNA binding protein [Homo sapiens (human)] - Gene - NCBI — https://www.ncbi.nlm.nih.gov/gene/1153
- **[S4]** Cold-inducible RNA-binding protein (CIRBP) adjusts clock-gene expression and REM-sleep recovery following sleep deprivation - PubMed — https://pubmed.ncbi.nlm.nih.gov/30720431/
- **[S5]** Cold-induced RNA-binding proteins regulate circadian gene expression by controlling alternative polyadenylation - PubMed — https://pubmed.ncbi.nlm.nih.gov/23792593/
- **[S6]** Cirbp-PSD95 axis protects against hypobaric hypoxia-induced aberrant morphology of hippocampal dendritic spines and cognitive deficits - PubMed — https://pubmed.ncbi.nlm.nih.gov/34419133/
- **[S7]** Involvement of Cold Inducible RNA-Binding Protein in Severe Hypoxia-Induced Growth Arrest of Neural Stem Cells In Vitro - PubMed — https://pubmed.ncbi.nlm.nih.gov/26927658/
- **[S8]** Cold-inducible RNA binding protein alleviates iron overload-induced neural ferroptosis under perinatal hypoxia insult - PubMed — https://pubmed.ncbi.nlm.nih.gov/38388728/
- **[S9]** Cold-inducible RNA-binding protein (CIRBP) adjusts clock-gene expression and REM-sleep recovery following sleep deprivation - PubMed — https://pubmed.ncbi.nlm.nih.gov/30720431/
- **[S10]** Cold-inducible RNA-binding protein (CIRBP) adjusts clock-gene expression and REM-sleep recovery following sleep deprivation - PubMed — https://pubmed.ncbi.nlm.nih.gov/30720431/
- **[S11]** Cold-inducible RNA-binding protein mediates neuroinflammation in cerebral ischemia - PubMed — https://pubmed.ncbi.nlm.nih.gov/24613680/
- **[S12]** Extracellular cold-inducible RNA-binding protein mediated neuroinflammation and neuronal apoptosis after traumatic brain injury - PubMed — https://pubmed.ncbi.nlm.nih.gov/38817684/
- **[S13]** CIRBP cold inducible RNA binding protein [Homo sapiens (human)] - Gene - NCBI — https://www.ncbi.nlm.nih.gov/gene/1153
- **[S14]** Cold-inducible RNA-binding protein (CIRBP) adjusts clock-gene expression and REM-sleep recovery following sleep deprivation - PubMed — https://pubmed.ncbi.nlm.nih.gov/30720431/
- **[S15]** Cirbp-PSD95 axis protects against hypobaric hypoxia-induced aberrant morphology of hippocampal dendritic spines and cognitive deficits - PubMed — https://pubmed.ncbi.nlm.nih.gov/34419133/
- **[S16]** Cold-inducible RNA-binding protein mediates neuroinflammation in cerebral ischemia - PubMed — https://pubmed.ncbi.nlm.nih.gov/24613680/
- **[S17]** Cold-inducible RNA-binding protein (CIRBP) adjusts clock-gene expression and REM-sleep recovery following sleep deprivation - PubMed — https://pubmed.ncbi.nlm.nih.gov/30720431/
