## Supplementary material for "Understanding Substance Dependence: What Differentiates Addictive from Non-Addictive Drugs?": Folder for supplementary files: non-addictive-genes-claims-first100-035-ca2-gene-protein-has-been-associated-with-regulato-97c5cde4b754.md

# non-addictive-genes-claims-first100-035-97c5cde4b754

**Original claim:** CA2 gene/protein has been associated with regulatory or homeostatic biological processes in neural systems.

## Claim

CA2 gene/protein has been associated with regulatory or homeostatic biological processes in neural systems.

## Verdict

Verdict: Likely True

## Confidence

High

## Supporting evidence

CA2 encodes carbonic anhydrase II (CA II), a highly active carbonic anhydrase that catalyzes reversible conversion between CO2 and bicarbonate/protons. That chemistry is directly relevant to acid-base regulation, bicarbonate handling, and ion-coupled transport. RefSeq/NCBI identifies CA2 as the gene for this enzyme, and neuroscience reviews describe carbonic anhydrases as important determinants of brain pH dynamics and neuronal function. ([ncbi.nlm.nih.gov](https://www.ncbi.nlm.nih.gov/gene/760)) [S1]

The strongest neural/homeostatic evidence is for **pH and bicarbonate homeostasis**. In mouse cortical astrocytes, NBCe1-mediated bicarbonate sensing was reported to be “supported by carbonic anhydrase II,” and the authors concluded this mechanism can alter both astrocyte intracellular pH and extracellular brain pH during acid/base disturbances. ([pubmed.ncbi.nlm.nih.gov](https://pubmed.ncbi.nlm.nih.gov/27981578/)) [S2]

A separate in vivo study of brain pH regulation showed that inhibiting carbonic anhydrase caused measurable extracellular acidification in cortex, and the authors concluded that bicarbonate buffering plus carbonic-anhydrase activity are essential for maintaining constant brain extracellular pH. While that experiment was not isoform-specific, the same paper notes high astrocytic expression of intracellular CA II and links CA II to the astrocytic bicarbonate-shuttle mechanism that stabilizes extracellular pH during neuronal activity. ([pmc.ncbi.nlm.nih.gov](https://pmc.ncbi.nlm.nih.gov/articles/PMC7545092/)) [S3]

CA II is also well positioned anatomically to participate in neural homeostasis. A human brain study localized CA II to oligodendrocytes, myelin, choroid plexus epithelium, a subset of mostly GABAergic neurons, and a few astrocytes, and concluded that CA II may participate in a broad range of biological processes in the developing and adult human brain. A CNS review likewise states that CA II is abundant in choroid plexus and mainly expressed in oligodendrocytes/myelin in normal brain. ([pubmed.ncbi.nlm.nih.gov](https://pubmed.ncbi.nlm.nih.gov/16825953/)) [S4]

The **choroid plexus / CSF** literature provides another homeostatic link. Reviews of CNS carbonic anhydrases state that CAs contribute to CSF formation and extracellular fluid/ion homeostasis, and specifically note that CA II is abundantly expressed in choroid plexus. In a 2022 rat study, CA2 was among the choroid-plexus carbonic anhydrases present, and acetazolamide lowered intracranial pressure and reduced a proxy of CSF secretion, supporting a bicarbonate-transport/CA-dependent role in this brain-fluid homeostatic system. ([pmc.ncbi.nlm.nih.gov](https://pmc.ncbi.nlm.nih.gov/articles/PMC7408524/)) [S5]

Loss-of-function evidence also points in the same direction. In CA II-deficient mice, oligodendrocytes appeared shrunken/degenerating and astrocytes were hypertrophic or swollen; the authors suggested that disturbed HCO3-/CO2 balance caused abnormal water and ion distribution in brain glial compartments. In humans, CA II deficiency syndrome is associated with progressive brain calcification and neurological manifestations, further indicating that normal CA2 function matters for CNS physiology, though these disease phenotypes are not by themselves proof of a specific “homeostatic pathway.” ([pubmed.ncbi.nlm.nih.gov](https://pubmed.ncbi.nlm.nih.gov/8229043/)) [S6]

Finally, curated functional annotations also support the claim at the “associated with” level. EMBL-EBI/PDBe lists CA2 with GO biological-process annotations including **neuron cellular homeostasis**, **regulation of intracellular pH**, **regulation of chloride transport**, and **positive regulation of GABAergic synaptic transmission**. These are database-level associations rather than single definitive experiments, but they are consistent with the primary literature. ([ebi.ac.uk](https://www.ebi.ac.uk/pdbe/entry/pdb/1bnq/biology)) [S7]

## Contradicting or limiting evidence

The main limitation is **specificity**: many papers discuss **carbonic anhydrases in general**, not CA2 alone, and some neural homeostatic roles are mediated mainly by other isoforms. For example, work on hippocampal interstitial pH buffering found that extracellular buffering in brain is attributable principally to membrane-bound CA IV, not CA II leaked from glia; knockout studies further supported roles for CA IV and CA XIV in extracellular pH regulation. So CA2 is associated with neural homeostasis, but it is not the sole or always dominant carbonic anhydrase for every neural compartment. ([pubmed.ncbi.nlm.nih.gov](https://pubmed.ncbi.nlm.nih.gov/11069930/)) [S8]

Likewise, for some neuronal excitability and seizure-related bicarbonate/GABA effects, CA VII appears especially important. In hippocampal pyramidal neurons, CA VII was described as the key age-dependent neuronal pH regulator for febrile seizures, although CA II was also present later and could contribute alongside CA VII in mature neurons. This means evidence for CA2 in neural regulation is real, but sometimes shared with or secondary to other isoforms. ([pubmed.ncbi.nlm.nih.gov](https://pubmed.ncbi.nlm.nih.gov/23881097/)) [S9]

Also, the claim uses the broad phrase “regulatory or homeostatic biological processes in neural systems.” The evidence is strongest for **acid-base, bicarbonate, ion/water, and CSF-related homeostasis**, and weaker for broader statements about neural-system regulation in general. ([pubmed.ncbi.nlm.nih.gov](https://pubmed.ncbi.nlm.nih.gov/27981578/)) [S10]

## Mechanistic / biological rationale

Mechanistically, CA II rapidly converts CO2 + H2O into HCO3- + H+, which lets cells buffer pH changes and feed bicarbonate-dependent transporters. In astrocytes, CA II can support NBCe1-dependent bicarbonate sensing and flux, helping stabilize intracellular and extracellular pH during neural activity or acid/base stress. ([pubmed.ncbi.nlm.nih.gov](https://pubmed.ncbi.nlm.nih.gov/27981578/)) [S11]

In choroid plexus epithelium, the same chemistry supports bicarbonate transport processes involved in CSF secretion and therefore intracranial fluid balance. In oligodendrocytes and glial compartments, disrupted CA II can plausibly disturb water/ion distribution because bicarbonate/CO2 balance is tightly coupled to osmotic and ionic handling. ([pubmed.ncbi.nlm.nih.gov](https://pubmed.ncbi.nlm.nih.gov/35768824/)) [S12]

This mechanistic logic matches CA II localization in oligodendrocytes, myelin, astrocytes, neurons, and choroid plexus, making the association with neural homeostatic biology biologically plausible rather than merely correlative. ([pubmed.ncbi.nlm.nih.gov](https://pubmed.ncbi.nlm.nih.gov/16825953/)) [S13]

## Bottom line

The claim is **likely true**. CA2/CA II has credible experimental and curated-support evidence linking it to **neural homeostatic/regulatory processes**, especially **brain pH regulation, bicarbonate sensing, ion/water balance in glia, and CSF secretion/homeostasis**. The main caveat is that these functions are **not unique to CA2**; other carbonic anhydrase isoforms, especially CA IV, CA XIV, and CA VII, also contribute importantly in different neural compartments. ([pubmed.ncbi.nlm.nih.gov](https://pubmed.ncbi.nlm.nih.gov/27981578/)) [S14]

## Cited sources

- **[S1]** CA2 carbonic anhydrase 2 [Homo sapiens (human)] - Gene - NCBI — https://www.ncbi.nlm.nih.gov/gene/760
- **[S2]** Bicarbonate sensing in mouse cortical astrocytes during extracellular acid/base disturbances - PubMed — https://pubmed.ncbi.nlm.nih.gov/27981578/
- **[S3]** 
            Astrocytes regulate brain extracellular pH via a neuronal activity-dependent bicarbonate shuttle - PMC
         — https://pmc.ncbi.nlm.nih.gov/articles/PMC7545092/
- **[S4]** Carbonic anhydrase II in the developing and adult human brain - PubMed — https://pubmed.ncbi.nlm.nih.gov/16825953/
- **[S5]** 
            The Expression of Carbonic Anhydrases II, IX and XII in Brain Tumors - PMC
         — https://pmc.ncbi.nlm.nih.gov/articles/PMC7408524/
- **[S6]** Glial cell abnormalities in the CNS of the carbonic anhydrase II deficient mutant mouse - PubMed — https://pubmed.ncbi.nlm.nih.gov/8229043/
- **[S7]** PDB 1bnq function and biology ‹ Protein Data Bank in Europe (PDBe) ‹ EMBL-EBI — https://www.ebi.ac.uk/pdbe/entry/pdb/1bnq/biology
- **[S8]** Interstitial carbonic anhydrase (CA) activity in brain is attributable to membrane-bound CA type IV - PubMed — https://pubmed.ncbi.nlm.nih.gov/11069930/
- **[S9]** Neuronal carbonic anhydrase VII provides GABAergic excitatory drive to exacerbate febrile seizures - PubMed — https://pubmed.ncbi.nlm.nih.gov/23881097/
- **[S10]** Bicarbonate sensing in mouse cortical astrocytes during extracellular acid/base disturbances - PubMed — https://pubmed.ncbi.nlm.nih.gov/27981578/
- **[S11]** Bicarbonate sensing in mouse cortical astrocytes during extracellular acid/base disturbances - PubMed — https://pubmed.ncbi.nlm.nih.gov/27981578/
- **[S12]** Acetazolamide modulates intracranial pressure directly by its action on the cerebrospinal fluid secretion apparatus - PubMed — https://pubmed.ncbi.nlm.nih.gov/35768824/
- **[S13]** Carbonic anhydrase II in the developing and adult human brain - PubMed — https://pubmed.ncbi.nlm.nih.gov/16825953/
- **[S14]** Bicarbonate sensing in mouse cortical astrocytes during extracellular acid/base disturbances - PubMed — https://pubmed.ncbi.nlm.nih.gov/27981578/
