## Supplementary material for "Understanding Substance Dependence: What Differentiates Addictive from Non-Addictive Drugs?": Folder for supplementary files: non-addictive-genes-claims-first100-036-mcf2l-gene-protein-has-been-associated-with-regula-27375250a369.md

# non-addictive-genes-claims-first100-036-27375250a369

**Original claim:** MCF2L gene/protein has been associated with regulatory or homeostatic biological processes in neural systems.

## Claim

**MCF2L gene/protein has been associated with regulatory or homeostatic biological processes in neural systems.**

## Verdict

Verdict: Likely True

MCF2L is **reasonably associated with regulatory processes in neural systems**, but the evidence is **stronger for neural regulation than for neural homeostasis**. The best direct data link MCF2L/DBS to **Schwann-cell migration downstream of neurotrophin signaling** and to **formation/stabilization of glutamatergic synapses in cortical neurons**. By contrast, evidence that MCF2L is a bona fide regulator of **neural homeostasis** in vivo is limited and mostly indirect. ([pmc.ncbi.nlm.nih.gov](https://pmc.ncbi.nlm.nih.gov/articles/PMC556009/)) [S1]

## Confidence

**Medium.**

The claim is broad enough to be supported by existing functional studies, but the literature is not extensive, much of it is from **cellular/rodent systems**, and the “homeostatic” part of the claim is not well established. ([pmc.ncbi.nlm.nih.gov](https://pmc.ncbi.nlm.nih.gov/articles/PMC556009/)) [S2]

## Supporting evidence

MCF2L encodes a **guanine nucleotide exchange factor (RhoGEF)** that interacts with GTP-bound **Rac1** and functions in **Rho/Rac signaling**, a pathway family heavily involved in cytoskeletal regulation, cell polarity, trafficking, and signaling responses. NCBI also reports broad expression of human MCF2L including **brain**, while the mouse ortholog is expressed in **brain** and shows broad **CNS** expression. Human Protein Atlas brain data classify MCF2L as **brain-enhanced** with low regional specificity across major brain regions, supporting biological plausibility for neural roles. ([ncbi.nlm.nih.gov](https://www.ncbi.nlm.nih.gov/gene/23263?utm_source=openai)) [S3]

The strongest direct neural-system evidence comes from **Schwann cells**. In a mechanistic PNAS study, the neurotrophin-3 receptor **TrkC** directly phosphorylated and activated **Dbs/MCF2L**; knockdown of Dbs in primary Schwann cells reduced **NT3-induced migration** by about 50%, and also reduced **Cdc42** and **JNK** pathway activation. The authors concluded that Dbs mediates the signaling pathway linking **NT3/TrkC to Schwann-cell migration**, a clearly regulatory process in the peripheral nervous system. ([pmc.ncbi.nlm.nih.gov](https://pmc.ncbi.nlm.nih.gov/articles/PMC556009/)) [S4]

That same Schwann-cell study further linked MCF2L to premyelinating biology: the authors noted that NT3-TrkC signaling can inhibit myelination and proposed that the **Dbs/Cdc42/JNK** pathway may be relevant to how Schwann cells migrate before myelination and possibly during remyelination after injury. This does not prove a homeostatic function, but it strengthens the case that MCF2L participates in **regulatory neural-glial programs** rather than being a purely non-neural factor. ([pmc.ncbi.nlm.nih.gov](https://pmc.ncbi.nlm.nih.gov/articles/PMC556009/)) [S5]

A second direct line of evidence comes from **cortical neurons**. In a PLOS One study, IL1RAPL1—an autism/intellectual-disability-associated synaptic protein—was shown to interact with **Mcf2l**. Knockdown of endogenous Mcf2l, or pharmacologic inhibition of **ROCK** downstream of **RhoA**, suppressed **IL1RAPL1-induced excitatory synapse formation** in cortical neurons. The same study found that Mcf2l knockdown diminished IL1RAPL1-dependent changes in **AMPA receptor subunit insertion**, and concluded that the **Mcf2l-RhoA-ROCK pathway mediates formation and stabilization of glutamatergic synapses**. That is a direct association with regulatory neural biology. ([pubmed.ncbi.nlm.nih.gov](https://pubmed.ncbi.nlm.nih.gov/23785489/)) [S6]

Even outside those dedicated neural studies, the broader literature recognizes MCF2L as relevant to neurotrophin-linked neural biology. The osteoarthritis GWAS paper identifying an MCF2L locus explicitly notes prior evidence that MCF2L regulates **neurotrophin-3-induced migration in Schwann cells**, tying the gene to the nerve growth factor family and neural signaling context. This is indirect support, but it shows the neural connection is not anecdotal. ([pmc.ncbi.nlm.nih.gov](https://pmc.ncbi.nlm.nih.gov/articles/PMC3169824/)) [S7]

## Contradicting or limiting evidence

The main limitation is that the evidence is **not strong for neural homeostasis per se**. The best-supported functions are **cell migration, signal transduction, synapse formation, and receptor trafficking**, which are regulatory processes, but not necessarily “homeostatic” processes in the classic sense of maintaining stable neural milieu, excitability, metabolism, or glial support over time. ([pmc.ncbi.nlm.nih.gov](https://pmc.ncbi.nlm.nih.gov/articles/PMC556009/)) [S8]

Most of the direct functional evidence is also **preclinical and reductionist**: primary Schwann-cell assays, cultured cortical neurons, and pathway perturbation experiments. I did not find strong evidence from human neurogenetic disorders, conditional neural knockout models, or large in vivo neuroscience studies showing that MCF2L is a major established regulator of CNS homeostasis. ([pmc.ncbi.nlm.nih.gov](https://pmc.ncbi.nlm.nih.gov/articles/PMC556009/)) [S9]

There are some **brain disease association** signals, but they are largely correlative. For example, an Alzheimer’s disease methylation/RNA study found **MCF2L** among the top differentially methylated regions in middle temporal gyrus. However, this does **not** establish causality or a specific homeostatic neural function; the same study did not find robust pathway enrichment after multiple-testing correction from the main methylation set, and the methylation-expression relationships highlighted other genes more clearly than MCF2L. ([pmc.ncbi.nlm.nih.gov](https://pmc.ncbi.nlm.nih.gov/articles/PMC10287777/)) [S10]

Likewise, human genetic association of MCF2L with **osteoarthritis** does not by itself prove a neural role. The neural relevance in that paper comes mostly from prior literature on Schwann-cell migration and NGF-family biology, not from direct demonstration that the OA-associated variant alters neural homeostasis. ([pmc.ncbi.nlm.nih.gov](https://pmc.ncbi.nlm.nih.gov/articles/PMC3169824/)) [S11]

## Mechanistic / biological rationale

Mechanistically, the claim is biologically plausible. MCF2L is a **Rho-family guanine nucleotide exchange factor**, so it sits upstream of small GTPase signaling that controls **actin remodeling, polarity, membrane dynamics, and signal propagation**. Those processes are central to how neural cells migrate, elaborate processes, traffic receptors, and build or stabilize synapses. ([ncbi.nlm.nih.gov](https://www.ncbi.nlm.nih.gov/gene/23263?utm_source=openai)) [S12]

In Schwann cells, the experimentally supported pathway is roughly **NT3 → TrkC → MCF2L/Dbs → Cdc42 → JNK**, promoting migration. In cortical neurons, the supported pathway is **IL1RAPL1 → Mcf2l → RhoA/ROCK**, influencing excitatory synapse formation and AMPA receptor trafficking/stabilization. These are clear **regulatory signaling roles within neural systems**. ([pmc.ncbi.nlm.nih.gov](https://pmc.ncbi.nlm.nih.gov/articles/PMC556009/)) [S13]

What remains uncertain is whether these pathway roles scale up to a broader role in **neural homeostasis**. Synapse stabilization and glial migration can contribute to homeostatic network maintenance in a broad sense, but the currently available literature supports that mainly as an **inference**, not as a directly demonstrated function of MCF2L. ([pmc.ncbi.nlm.nih.gov](https://pmc.ncbi.nlm.nih.gov/articles/PMC556009/)) [S14]

## Bottom line

The claim is **best judged as Likely True**. There is credible evidence that **MCF2L participates in regulatory neural processes**, especially **Schwann-cell migration** and **glutamatergic synapse formation/stabilization**. However, the evidence for a specific role in **neural homeostasis** is **limited and indirect**, so the claim is supported mainly on the “regulatory” half, not strongly on the “homeostatic” half. ([pmc.ncbi.nlm.nih.gov](https://pmc.ncbi.nlm.nih.gov/articles/PMC556009/)) [S15]

## Cited sources

- **[S1]** 
            The neurotrophin-3 receptor TrkC directly phosphorylates and activates the nucleotide exchange factor Dbs to enhance Schwann cell migration - PMC
         — https://pmc.ncbi.nlm.nih.gov/articles/PMC556009/
- **[S2]** 
            The neurotrophin-3 receptor TrkC directly phosphorylates and activates the nucleotide exchange factor Dbs to enhance Schwann cell migration - PMC
         — https://pmc.ncbi.nlm.nih.gov/articles/PMC556009/
- **[S3]** MCF2L MCF.2 cell line derived transforming sequence like [Homo sapiens (human)] - Gene - NCBI — https://www.ncbi.nlm.nih.gov/gene/23263?utm_source=openai
- **[S4]** 
            The neurotrophin-3 receptor TrkC directly phosphorylates and activates the nucleotide exchange factor Dbs to enhance Schwann cell migration - PMC
         — https://pmc.ncbi.nlm.nih.gov/articles/PMC556009/
- **[S5]** 
            The neurotrophin-3 receptor TrkC directly phosphorylates and activates the nucleotide exchange factor Dbs to enhance Schwann cell migration - PMC
         — https://pmc.ncbi.nlm.nih.gov/articles/PMC556009/
- **[S6]** IL1RAPL1 associated with mental retardation and autism regulates the formation and stabilization of glutamatergic synapses of cortical neurons through RhoA signaling pathway - PubMed — https://pubmed.ncbi.nlm.nih.gov/23785489/
- **[S7]** 
            A Variant in MCF2L Is Associated with Osteoarthritis - PMC
         — https://pmc.ncbi.nlm.nih.gov/articles/PMC3169824/
- **[S8]** 
            The neurotrophin-3 receptor TrkC directly phosphorylates and activates the nucleotide exchange factor Dbs to enhance Schwann cell migration - PMC
         — https://pmc.ncbi.nlm.nih.gov/articles/PMC556009/
- **[S9]** 
            The neurotrophin-3 receptor TrkC directly phosphorylates and activates the nucleotide exchange factor Dbs to enhance Schwann cell migration - PMC
         — https://pmc.ncbi.nlm.nih.gov/articles/PMC556009/
- **[S10]** 
            Integrated DNA Methylation/RNA Profiling in Middle Temporal Gyrus of Alzheimer’s Disease - PMC
         — https://pmc.ncbi.nlm.nih.gov/articles/PMC10287777/
- **[S11]** 
            A Variant in MCF2L Is Associated with Osteoarthritis - PMC
         — https://pmc.ncbi.nlm.nih.gov/articles/PMC3169824/
- **[S12]** MCF2L MCF.2 cell line derived transforming sequence like [Homo sapiens (human)] - Gene - NCBI — https://www.ncbi.nlm.nih.gov/gene/23263?utm_source=openai
- **[S13]** 
            The neurotrophin-3 receptor TrkC directly phosphorylates and activates the nucleotide exchange factor Dbs to enhance Schwann cell migration - PMC
         — https://pmc.ncbi.nlm.nih.gov/articles/PMC556009/
- **[S14]** 
            The neurotrophin-3 receptor TrkC directly phosphorylates and activates the nucleotide exchange factor Dbs to enhance Schwann cell migration - PMC
         — https://pmc.ncbi.nlm.nih.gov/articles/PMC556009/
- **[S15]** 
            The neurotrophin-3 receptor TrkC directly phosphorylates and activates the nucleotide exchange factor Dbs to enhance Schwann cell migration - PMC
         — https://pmc.ncbi.nlm.nih.gov/articles/PMC556009/
