## Supplementary material for "Understanding Substance Dependence: What Differentiates Addictive from Non-Addictive Drugs?": Folder for supplementary files: non-addictive-genes-claims-first100-037-arf1-gene-protein-has-been-associated-with-regulat-b3032215d9d5.md

# non-addictive-genes-claims-first100-037-b3032215d9d5

**Original claim:** ARF1 gene/protein has been associated with regulatory or homeostatic biological processes in neural systems.

## Claim

**ARF1 gene/protein has been associated with regulatory or homeostatic biological processes in neural systems.**

## Verdict

Verdict: Likely True

ARF1 is convincingly associated with **regulatory** processes in neural systems across several lines of evidence: neuronal migration, dendritic/Golgi polarization, synaptic receptor trafficking and long-term depression, dendrite pruning, neural stem-cell reactivation/regeneration, and Schwann-cell myelination. Human genetic data also link ARF1 dysfunction to neurodevelopmental brain phenotypes consistent with impaired neuronal migration. The claim is best supported if “regulatory” is emphasized; evidence specifically framed as **classical neural homeostasis** is more limited and indirect. ([pmc.ncbi.nlm.nih.gov](https://pmc.ncbi.nlm.nih.gov/articles/PMC3725416/)) [S1]

## Confidence

**High** — because multiple independent primary studies, in mammals and invertebrates, converge on neural regulatory roles for ARF1, and human ARF1-related disorder provides in vivo relevance. Confidence is not “maximal” only because the literature more often shows regulation of neural development/plasticity/maintenance than explicit “homeostatic” control. ([pmc.ncbi.nlm.nih.gov](https://pmc.ncbi.nlm.nih.gov/articles/PMC3725416/)) [S2]

## Supporting evidence

ARF1 is present in neural tissue and neural compartments relevant to regulation. In mouse brain development, ARF1-3 expression rises from embryonic to postnatal stages; ARF1 is found in cortical and hippocampal neuron cytoplasm and dendrites, as well as synaptosome fractions. A Human Protein Atlas summary also places ARF1 in a neuronal “synaptic function” expression cluster. This does not prove function by itself, but it supports biological plausibility in neural systems. ([pubmed.ncbi.nlm.nih.gov](https://pubmed.ncbi.nlm.nih.gov/38227057/)) [S3]

The strongest evidence in mature neurons is for **synaptic regulation**. In hippocampal neurons and organotypic slices, Rocca et al. showed that GTP-bound Arf1 binds PICK1, modulates Arp2/3-mediated actin polymerization, regulates GluA2-containing AMPA receptor trafficking and spine size, and is required for NMDA receptor-dependent long-term depression (LTD). Mutant Arf1 that cannot bind PICK1 reduced surface GluA2, produced smaller spines, and abolished LTD in slice recordings. That is a direct demonstration that ARF1 participates in a neural regulatory process. ([pmc.ncbi.nlm.nih.gov](https://pmc.ncbi.nlm.nih.gov/articles/PMC3725416/)) [S4]

ARF1 is also implicated in **neuronal migration and cortical development**. In embryonic mouse cortex, disruption of Arf1 activity impaired migration of electroporated cells toward the cortical plate and could disrupt neuroependymal integrity. Mechanistically, the study linked Filamin A and BIG2/ARFGEF2 to Arf1 activation during migration and adhesion-related processes. Human evidence aligns with this: a cohort study of individuals with de novo ARF1 variants found intellectual disability, microcephaly, seizures, and frequent brain MRI abnormalities consistent with neuronal migration disorders, including periventricular nodular heterotopia; several patient variants reduced ARF1 activity in vitro. ([pubmed.ncbi.nlm.nih.gov](https://pubmed.ncbi.nlm.nih.gov/24089482/)) [S5]

Beyond migration, ARF1 regulates **neuronal morphology and polarity**. In cultured hippocampal neurons, BIG2-ARF1-RhoA-mDia1 signaling was required for dendritic Golgi polarization, dendrite growth, and dendrite maintenance; constitutively active ARF1 rescued dendritic defects caused by ARFGEF2 loss. This places ARF1 in a pathway controlling the polarized organization of neurons, which is a classic regulatory function in neural cell biology. ([pubmed.ncbi.nlm.nih.gov](https://pubmed.ncbi.nlm.nih.gov/29455446/?utm_source=openai)) [S6]

Evidence from invertebrate neural systems broadens the picture. In Drosophila sensory neurons, Sec71 acting as a GEF for Arf1 was required for **dendrite pruning**, a conserved neural remodeling process. Separately, in quiescent Drosophila neural stem cells, Patronin functioned upstream of Arf1 and Msps to regulate acentrosomal microtubule assembly, neural stem-cell reactivation, regeneration, and E-cadherin targeting to neural stem cell–neuropil contacts. A recent study also found Arf1 and ARFGEF2/Sec71 important for **neuroblast polarity during asymmetric division**. Together these studies connect Arf1 to regulated remodeling, regeneration, and stem-cell state transitions in neural tissue. ([pubmed.ncbi.nlm.nih.gov](https://pubmed.ncbi.nlm.nih.gov/28420712/?utm_source=openai)) [S7]

ARF1’s relevance extends to the peripheral nervous system. In Schwann cell-specific knockout mice, loss of Arf1 reduced myelin protein zero localization and decreased myelin thickness, showing that BIG1/Arfgef1 and Arf1 regulate initiation of axonal myelination. Myelination is a maintenance-critical process for neural conduction and axonal protection, so this supports a role in neural system maintenance, even though the study focused on developmental initiation rather than abstract homeostasis. ([pubmed.ncbi.nlm.nih.gov](https://pubmed.ncbi.nlm.nih.gov/29740613/)) [S8]

## Contradicting or limiting evidence

The main limitation is definitional: the literature supports ARF1 most strongly as a regulator of **trafficking-dependent neural development, polarity, plasticity, and maintenance**, but relatively few papers describe ARF1 as a controller of “neural homeostasis” in the narrow sense of set-point stabilization or homeostatic plasticity. So the claim is stronger for **regulatory** processes than for explicitly demonstrated **homeostatic** ones. ([pmc.ncbi.nlm.nih.gov](https://pmc.ncbi.nlm.nih.gov/articles/PMC3725416/)) [S9]

Much of the functional evidence comes from **cultured neurons, mouse models, and Drosophila**, not from direct human mechanistic experiments in neural tissue. Human genetic data are supportive, but they mainly show disease association and variant effects on ARF1 activity rather than complete causal pathway mapping in human neurons. ([pmc.ncbi.nlm.nih.gov](https://pmc.ncbi.nlm.nih.gov/articles/PMC10579487/)) [S10]

The subcellular story is also nuanced. One mouse study detected ARF1 in synaptosome fractions and neuronal dendrites but did **not** find it in the core postsynaptic compartment, suggesting ARF1’s neural roles may be compartment-specific rather than a universal postsynaptic regulator. That limitation does not weaken the overall association, but it argues against overgeneralizing ARF1 as a broad synaptic scaffold protein. ([pubmed.ncbi.nlm.nih.gov](https://pubmed.ncbi.nlm.nih.gov/38227057/)) [S11]

Finally, even in ARF1-related human disorder, the exact pathogenic mechanism remains unresolved: the cohort paper notes uncertainty between haploinsufficiency and dominant-negative effects. So while ARF1 is clearly implicated in neurodevelopmental regulation, some mechanistic details are still unsettled. ([pmc.ncbi.nlm.nih.gov](https://pmc.ncbi.nlm.nih.gov/articles/PMC10579487/)) [S12]

## Mechanistic / biological rationale

ARF1 is a small membrane-associated GTPase that switches between GDP-bound and GTP-bound states and helps control vesicle formation, adaptor recruitment, cargo trafficking, and membrane-cytoskeleton coupling. In neural cells, those basic functions map naturally onto processes that require precise spatial control of membrane proteins and organelles. ([pmc.ncbi.nlm.nih.gov](https://pmc.ncbi.nlm.nih.gov/articles/PMC10579487/)) [S13]

Mechanistically, ARF1 can influence neural regulation through several linked routes:  
1. **Synaptic receptor trafficking and actin remodeling** via PICK1/Arp2/3, affecting AMPAR handling, spine morphology, and LTD. ([pmc.ncbi.nlm.nih.gov](https://pmc.ncbi.nlm.nih.gov/articles/PMC3725416/)) [S14]  
2. **Cell adhesion and migration** via BIG2/ARFGEF2-dependent activation and trafficking of adhesion-related cargo, relevant to cortical neuronal migration and neuroependymal integrity. ([pubmed.ncbi.nlm.nih.gov](https://pubmed.ncbi.nlm.nih.gov/24089482/)) [S15]  
3. **Golgi positioning and polarized growth** via BIG2-ARF1-RhoA-mDia1, affecting dendritic Golgi deployment and dendrite maintenance. ([pubmed.ncbi.nlm.nih.gov](https://pubmed.ncbi.nlm.nih.gov/29455446/?utm_source=openai)) [S16]  
4. **Membrane remodeling in glia** during Schwann-cell myelination, affecting myelin protein targeting and sheath formation. ([pubmed.ncbi.nlm.nih.gov](https://pubmed.ncbi.nlm.nih.gov/29740613/)) [S17]  
5. **Neural remodeling/stem-cell control** in Drosophila, where Arf1 contributes to dendrite pruning, neuroblast polarity, and quiescent neural stem-cell reactivation/regeneration. ([pubmed.ncbi.nlm.nih.gov](https://pubmed.ncbi.nlm.nih.gov/28420712/?utm_source=openai)) [S18]

These mechanisms make biological sense for a protein whose core job is to coordinate membrane trafficking with cytoskeletal organization. Neural systems depend heavily on exactly those functions for migration, polarity, synaptic remodeling, and maintenance. ([pmc.ncbi.nlm.nih.gov](https://pmc.ncbi.nlm.nih.gov/articles/PMC3725416/)) [S19]

## Bottom line

Yes—**ARF1 is credibly associated with regulatory biological processes in neural systems**, with evidence spanning neuronal migration, dendritic polarity, synaptic plasticity, dendrite pruning, neural stem-cell activation/regeneration, and peripheral myelination. The claim is **well supported if interpreted as “regulatory”**; support is **more limited if interpreted strictly as “directly proven neural homeostasis.”** ([pmc.ncbi.nlm.nih.gov](https://pmc.ncbi.nlm.nih.gov/articles/PMC3725416/)) [S20]

## Cited sources

- **[S1]** 
            The Small GTPase Arf1 Modulates Arp2/3-Mediated Actin Polymerization via PICK1 to Regulate Synaptic Plasticity - PMC
         — https://pmc.ncbi.nlm.nih.gov/articles/PMC3725416/
- **[S2]** 
            The Small GTPase Arf1 Modulates Arp2/3-Mediated Actin Polymerization via PICK1 to Regulate Synaptic Plasticity - PMC
         — https://pmc.ncbi.nlm.nih.gov/articles/PMC3725416/
- **[S3]** Expression analysis of type I ARF small GTPases ARF1-3 during mouse brain development - PubMed — https://pubmed.ncbi.nlm.nih.gov/38227057/
- **[S4]** 
            The Small GTPase Arf1 Modulates Arp2/3-Mediated Actin Polymerization via PICK1 to Regulate Synaptic Plasticity - PMC
         — https://pmc.ncbi.nlm.nih.gov/articles/PMC3725416/
- **[S5]** Filamin A regulates neuronal migration through brefeldin A-inhibited guanine exchange factor 2-dependent Arf1 activation - PubMed — https://pubmed.ncbi.nlm.nih.gov/24089482/
- **[S6]** BIG2-ARF1-RhoA-mDia1 Signaling Regulates Dendritic Golgi Polarization in Hippocampal Neurons - PubMed — https://pubmed.ncbi.nlm.nih.gov/29455446/?utm_source=openai
- **[S7]** Sec71 functions as a GEF for the small GTPase Arf1 to govern dendrite pruning of Drosophila sensory neurons - PubMed — https://pubmed.ncbi.nlm.nih.gov/28420712/?utm_source=openai
- **[S8]** BIG1/Arfgef1 and Arf1 regulate the initiation of myelination by Schwann cells in mice - PubMed — https://pubmed.ncbi.nlm.nih.gov/29740613/
- **[S9]** 
            The Small GTPase Arf1 Modulates Arp2/3-Mediated Actin Polymerization via PICK1 to Regulate Synaptic Plasticity - PMC
         — https://pmc.ncbi.nlm.nih.gov/articles/PMC3725416/
- **[S10]** 
            ARF1-related disorder: phenotypic and molecular spectrum - PMC
         — https://pmc.ncbi.nlm.nih.gov/articles/PMC10579487/
- **[S11]** Expression analysis of type I ARF small GTPases ARF1-3 during mouse brain development - PubMed — https://pubmed.ncbi.nlm.nih.gov/38227057/
- **[S12]** 
            ARF1-related disorder: phenotypic and molecular spectrum - PMC
         — https://pmc.ncbi.nlm.nih.gov/articles/PMC10579487/
- **[S13]** 
            ARF1-related disorder: phenotypic and molecular spectrum - PMC
         — https://pmc.ncbi.nlm.nih.gov/articles/PMC10579487/
- **[S14]** 
            The Small GTPase Arf1 Modulates Arp2/3-Mediated Actin Polymerization via PICK1 to Regulate Synaptic Plasticity - PMC
         — https://pmc.ncbi.nlm.nih.gov/articles/PMC3725416/
- **[S15]** Filamin A regulates neuronal migration through brefeldin A-inhibited guanine exchange factor 2-dependent Arf1 activation - PubMed — https://pubmed.ncbi.nlm.nih.gov/24089482/
- **[S16]** BIG2-ARF1-RhoA-mDia1 Signaling Regulates Dendritic Golgi Polarization in Hippocampal Neurons - PubMed — https://pubmed.ncbi.nlm.nih.gov/29455446/?utm_source=openai
- **[S17]** BIG1/Arfgef1 and Arf1 regulate the initiation of myelination by Schwann cells in mice - PubMed — https://pubmed.ncbi.nlm.nih.gov/29740613/
- **[S18]** Sec71 functions as a GEF for the small GTPase Arf1 to govern dendrite pruning of Drosophila sensory neurons - PubMed — https://pubmed.ncbi.nlm.nih.gov/28420712/?utm_source=openai
- **[S19]** 
            The Small GTPase Arf1 Modulates Arp2/3-Mediated Actin Polymerization via PICK1 to Regulate Synaptic Plasticity - PMC
         — https://pmc.ncbi.nlm.nih.gov/articles/PMC3725416/
- **[S20]** 
            The Small GTPase Arf1 Modulates Arp2/3-Mediated Actin Polymerization via PICK1 to Regulate Synaptic Plasticity - PMC
         — https://pmc.ncbi.nlm.nih.gov/articles/PMC3725416/
