## Supplementary material for "Understanding Substance Dependence: What Differentiates Addictive from Non-Addictive Drugs?": Folder for supplementary files: non-addictive-genes-claims-first100-038-pabpc1-gene-protein-has-been-associated-with-regul-3ac10f02d790.md

# non-addictive-genes-claims-first100-038-3ac10f02d790

**Original claim:** PABPC1 gene/protein has been associated with regulatory or homeostatic biological processes in neural systems.

## Claim

PABPC1 gene/protein has been associated with regulatory or homeostatic biological processes in neural systems.

## Verdict

Verdict: Likely True

PABPC1 is a well-established post-transcriptional regulator of mRNA stability and translation in general cell biology, and multiple studies place it in bona fide neural contexts, including dendritic/local translation in hippocampal neurons, neuroendocrine regulation of **Gnrh1** mRNA in hypothalamic neurons, neuronal RNA granules, and human/mouse neurodevelopmental phenotypes. The evidence is strongest for **regulatory** roles in neural systems; the case for **homeostatic** roles is real but narrower and more context-dependent. ([ncbi.nlm.nih.gov](https://www.ncbi.nlm.nih.gov/gene/26986)) [S1]

## Confidence

High

## Supporting evidence

PABPC1’s core molecular function is itself regulatory: the NCBI Gene record describes PABPC1 as a cytoplasmic poly(A)-binding protein that binds the 3′ poly(A) tail of mRNAs, promotes ribosome recruitment and translation initiation, and is also required for poly(A) shortening as the first step in mRNA decay. That establishes PABPC1 as a bona fide regulator of mRNA fate, making neural regulatory roles biologically plausible rather than speculative. ([ncbi.nlm.nih.gov](https://www.ncbi.nlm.nih.gov/gene/26986)) [S2]

In cultured rodent hippocampal neurons, **Pabpc1 mRNA** was found to be **dendritically localized**. In the same study, the authors emphasized that dendritically localized transcripts encoding translation machinery likely increase the translational capacity of stimulated synapses and help support long-lasting synaptic plasticity and synaptic tagging. This is direct evidence that the PABPC1 system is associated with local translational regulation in a classic neural-plasticity compartment. ([pmc.ncbi.nlm.nih.gov](https://pmc.ncbi.nlm.nih.gov/articles/PMC6675000/)) [S3]

The strongest direct “homeostatic/regulatory in neural tissue” evidence comes from the hypothalamus. In a mouse puberty study, MKRN3-mediated ubiquitination of PABPC1 reduced PABPC1 binding to poly(A) tails and to **Gnrh1** mRNA, shortened the **Gnrh1** poly(A) tail, and reduced PABPC1–EIF4G1 complex formation in hypothalamus. In **primary hypothalamic neurons**, loss of **Mkrn3** prolonged **Gnrh1** mRNA half-life, and the authors explicitly framed the MKRN3–PABPC1 axis as controlling the **stability, translation, and homeostasis** of **GNRH1** mRNA during puberty initiation. That is a direct neural-system example of PABPC1 participating in mRNA homeostasis. ([pmc.ncbi.nlm.nih.gov](https://pmc.ncbi.nlm.nih.gov/articles/PMC8053111/)) [S4]

PABPC1 also appears in neuronal translation granules. In mouse postnatal neocortex and differentiated neurons, JAKMIP1 co-immunoprecipitated with PABPC1, PABPC1 was present in translation-associated RNP granules, and JAKMIP1 colocalized with **PABPC1-positive granules** in differentiated neurons. Loss of Jakmip1 reduced nascent protein synthesis in soma and neurites and altered synaptic signaling and behavior, placing PABPC1 within a neuronal translational-regulatory complex linked to synaptic function. ([pubmed.ncbi.nlm.nih.gov](https://pubmed.ncbi.nlm.nih.gov/26627310/)) [S5]

Human genetics also supports neural relevance. A 2022 **Genetics in Medicine** study described children with **de novo PABPC1 variants** who had developmental delay, expressive speech delay, autistic features, and in some cases seizures/behavioral disorders. In utero electroporation experiments further showed that **Pabpc1 knockdown decreased proliferation of neural progenitor cells** in embryonic mouse brain, and several patient variants failed to rescue the defect. This does not prove “homeostasis” in the narrow physiological sense, but it strongly supports a role for PABPC1 in neural developmental regulation. ([pubmed.ncbi.nlm.nih.gov](https://pubmed.ncbi.nlm.nih.gov/35511136/)) [S6]

There is also neurodegeneration-linked evidence. In an SCA2 knock-in mouse model, PABPC1 showed tissue-specific changes in cortex and cerebellum, including progressive **insolubility/sequestration in cerebellum** and altered distribution in Purkinje cells. Separately, a mechanistic study showed that poly(A)-binding proteins act as RNA-dependent chaperones for **ATXN2**, regulating its condensation in **homeostasis and stress** and helping direct stress-granule behavior; because ATXN2 is implicated in ALS and spinocerebellar ataxia, this links PABPC1-like activity to neural stress-response regulation. ([pubmed.ncbi.nlm.nih.gov](https://pubmed.ncbi.nlm.nih.gov/22956915/)) [S7]

## Contradicting or limiting evidence

The biggest limitation is that **PABPC1 is not a neuron-specific gene**. Its canonical role is broad and housekeeping-like across many tissues and cell types, and a recent Nucleic Acids Research study reported that median **PABPC1 mRNA** levels vary widely across human tissues, with relatively low levels in brain compared with some non-brain tissues. So the best-supported statement is not that PABPC1 is a dedicated neural-homeostasis gene, but that a broadly acting RNA regulator is also deployed in neural regulatory programs. ([ncbi.nlm.nih.gov](https://www.ncbi.nlm.nih.gov/gene/26986)) [S8]

A second limitation is **isoform resolution**. Some neural studies discuss “PABP” or multiple cytoplasmic PABPs rather than isolating PABPC1 alone. Even the hypothalamic puberty paper studied both **PABPC1 and PABPC4** in parts of the mechanism, although it did directly assess endogenous PABPC1 binding to **Gnrh1** mRNA and PABPC1–EIF4G1 interaction in hypothalamus. That means some literature supports the broader PABP family in brain more strongly than PABPC1 alone. ([pmc.ncbi.nlm.nih.gov](https://pmc.ncbi.nlm.nih.gov/articles/PMC8053111/)) [S9]

A third limitation is that much of the mechanistic literature is **mouse, cultured-neuron, or cell-line based**, rather than direct demonstration of PABPC1 controlling normal neural homeostasis in humans. For example, FMRP–PABPC1 interaction data come from human neuronal-like SH-SY5Y cells, and some disease-mechanism work on PABPC1 and repeat-RNA degradation was performed in HeLa cells and patient-derived fibroblasts rather than neurons. Those data are relevant and supportive, but still indirect for normal human neural physiology. ([pmc.ncbi.nlm.nih.gov](https://pmc.ncbi.nlm.nih.gov/articles/PMC9753132/)) [S10]

## Mechanistic / biological rationale

Mechanistically, the claim makes sense. Neurons rely heavily on regulated mRNA stability and local translation, especially in dendrites, axons, synapses, and neuroendocrine cells. Because PABPC1 binds poly(A) tails, promotes translation initiation through EIF4G-associated closed-loop formation, and also influences deadenylation/decay, it is well positioned to act as a controller of RNA availability and protein synthesis in neural compartments. ([ncbi.nlm.nih.gov](https://www.ncbi.nlm.nih.gov/gene/26986)) [S11]

This rationale is reinforced by RNA-granule biology. In human neuronal cells, FMRP was shown to interact directly with **PABPC1**, and the study concluded that FMRP-bound mRNAs are stabilized and translationally repressed until signaling activates localized translation and decay. That places PABPC1 within neuron-relevant RNA storage/translation-control machinery. ([pmc.ncbi.nlm.nih.gov](https://pmc.ncbi.nlm.nih.gov/articles/PMC9753132/)) [S12]

PABPC-related stress-granule biology also fits a homeostatic model. The ATXN2 study showed that poly(A)-binding proteins can act as chaperones that prevent inappropriate condensation under baseline conditions and redirect mixing into stress granules under stress. Inference: even when not measured in neurons directly, such a mechanism is highly relevant to neural homeostasis because neurons are especially sensitive to defects in RNA granules, proteostasis, and translational stress responses. ([pmc.ncbi.nlm.nih.gov](https://pmc.ncbi.nlm.nih.gov/articles/PMC10318123/)) [S13]

## Bottom line

The claim is **likely true**. PABPC1 is fundamentally a broad RNA regulator, not a purely neural gene, but the literature supports its association with **neural regulatory processes** and provides at least some direct evidence for **neural mRNA homeostasis**, especially in hypothalamic neurons controlling **Gnrh1** expression. A precise summary would be: **PABPC1 is a general post-transcriptional regulator that also participates in neuronal/neuroendocrine RNA regulation, local translation, and stress-responsive RNA-granule biology in neural systems.** ([ncbi.nlm.nih.gov](https://www.ncbi.nlm.nih.gov/gene/26986)) [S14]

## Cited sources

- **[S1]** PABPC1 poly(A) binding protein cytoplasmic 1 [Homo sapiens (human)] - Gene - NCBI — https://www.ncbi.nlm.nih.gov/gene/26986
- **[S2]** PABPC1 poly(A) binding protein cytoplasmic 1 [Homo sapiens (human)] - Gene - NCBI — https://www.ncbi.nlm.nih.gov/gene/26986
- **[S3]** 
            Identification of Process-Localized mRNAs from Cultured Rodent Hippocampal Neurons - PMC
         — https://pmc.ncbi.nlm.nih.gov/articles/PMC6675000/
- **[S4]** 
            MKRN3-mediated ubiquitination of Poly(A)-binding proteins modulates the stability and translation of GNRH1 mRNA in mammalian puberty - PMC
         — https://pmc.ncbi.nlm.nih.gov/articles/PMC8053111/
- **[S5]** JAKMIP1, a Novel Regulator of Neuronal Translation, Modulates Synaptic Function and Autistic-like Behaviors in Mouse - PubMed — https://pubmed.ncbi.nlm.nih.gov/26627310/
- **[S6]** De novo variants in the PABP domain of PABPC1 lead to developmental delay - PubMed — https://pubmed.ncbi.nlm.nih.gov/35511136/
- **[S7]** ATXN2-CAG42 sequesters PABPC1 into insolubility and induces FBXW8 in cerebellum of old ataxic knock-in mice - PubMed — https://pubmed.ncbi.nlm.nih.gov/22956915/
- **[S8]** PABPC1 poly(A) binding protein cytoplasmic 1 [Homo sapiens (human)] - Gene - NCBI — https://www.ncbi.nlm.nih.gov/gene/26986
- **[S9]** 
            MKRN3-mediated ubiquitination of Poly(A)-binding proteins modulates the stability and translation of GNRH1 mRNA in mammalian puberty - PMC
         — https://pmc.ncbi.nlm.nih.gov/articles/PMC8053111/
- **[S10]** 
            Integrative Omics Indicate FMRP Sequesters mRNA From Translation and Deadenylation In Human Neuronal Cells - PMC
         — https://pmc.ncbi.nlm.nih.gov/articles/PMC9753132/
- **[S11]** PABPC1 poly(A) binding protein cytoplasmic 1 [Homo sapiens (human)] - Gene - NCBI — https://www.ncbi.nlm.nih.gov/gene/26986
- **[S12]** 
            Integrative Omics Indicate FMRP Sequesters mRNA From Translation and Deadenylation In Human Neuronal Cells - PMC
         — https://pmc.ncbi.nlm.nih.gov/articles/PMC9753132/
- **[S13]** 
            Poly(A)-binding protein is an ataxin-2 chaperone that regulates biomolecular condensates - PMC
         — https://pmc.ncbi.nlm.nih.gov/articles/PMC10318123/
- **[S14]** PABPC1 poly(A) binding protein cytoplasmic 1 [Homo sapiens (human)] - Gene - NCBI — https://www.ncbi.nlm.nih.gov/gene/26986
