## Supplementary material for "Understanding Substance Dependence: What Differentiates Addictive from Non-Addictive Drugs?": Folder for supplementary files: non-addictive-genes-claims-first100-039-baiap2-gene-protein-has-been-associated-with-regul-1eab43410e24.md

# non-addictive-genes-claims-first100-039-1eab43410e24

**Original claim:** BAIAP2 gene/protein has been associated with regulatory or homeostatic biological processes in neural systems.

## Claim

BAIAP2 gene/protein has been associated with regulatory or homeostatic biological processes in neural systems.

## Verdict

Verdict: Likely True

## Confidence

High

## Supporting evidence

BAIAP2 encodes IRSp53, a scaffold/adaptor protein that is strongly linked to **regulatory functions at excitatory synapses**. A peer-reviewed review summarizes IRSp53/BAIAP2 as an abundant postsynaptic-density component and an “important regulator” of actin-rich dendritic spines, with evidence connecting it to NMDA receptor regulation in the brain. ([pubmed.ncbi.nlm.nih.gov](https://pubmed.ncbi.nlm.nih.gov/26275848/)) [S1]

Primary mouse work provides direct functional evidence that BAIAP2 helps regulate neural signaling. In IRSp53/BAIAP2 knockout mice, investigators found a reduced AMPA/NMDA ratio, increased NMDA receptor-mediated miniature EPSC amplitude, enhanced hippocampal LTP, and impaired spatial/object learning. The authors concluded that IRSp53 is involved in regulation of NMDA receptor-mediated excitatory synaptic transmission, LTP, and learning/memory behaviors. That is a clear association with neural regulatory processes, even if not all of them are “homeostatic” in the narrow technical sense. ([pmc.ncbi.nlm.nih.gov](https://pmc.ncbi.nlm.nih.gov/articles/PMC6666077/)) [S2]

More recent mechanistic work strengthens this interpretation. In *Journal of Cell Biology* (2022), IRSp53/BAIAP2 was described as abundantly expressed in excitatory synapses and essential for synapse development and synaptic plasticity. The study showed that IRSp53 interacts with PSD-95 and Shank3 to promote postsynaptic-density formation and actin filament bundling, giving a concrete molecular basis for how BAIAP2 regulates synaptic structure and function. ([pubmed.ncbi.nlm.nih.gov](https://pubmed.ncbi.nlm.nih.gov/35819332/)) [S3]

BAIAP2 also shows **activity-dependent regulation within synapses**. In cultured hippocampal neurons, IRSp53 labeling at the postsynaptic density increased after depolarization or NMDA application, especially in the PSD pallium. This supports the idea that BAIAP2 dynamically participates in excitatory synapse reorganization under stimulated conditions rather than being a static structural component. ([pubmed.ncbi.nlm.nih.gov](https://pubmed.ncbi.nlm.nih.gov/29284046/)) [S4]

Cell-type-specific knockout data further support a role in maintaining regulated circuit properties. In 2020 work, excitatory-neuron deletion of IRSp53/BAIAP2 increased the excitatory/inhibitory synaptic transmission ratio in medial prefrontal cortex pyramidal neurons, while inhibitory-neuron deletion had much smaller effects. This suggests BAIAP2 contributes to the regulated balance of synaptic transmission in at least some cortical circuits. ([pubmed.ncbi.nlm.nih.gov](https://pubmed.ncbi.nlm.nih.gov/32116566/)) [S5]

Developmental studies also place BAIAP2 in neural regulatory biology. A 2024 *Development* paper found that Baiap2 knockdown in mouse cortex disrupted neuronal migration, morphogenesis, and differentiation, and a patient-associated BAIAP2 variant failed to rescue the migration defect. A 2025 *Epilepsia* study reported de novo missense variants in BAIAP2 in developmental and epileptic encephalopathies, with functional assays showing increased neuronal excitability due to enhanced excitatory synaptogenesis. These data extend BAIAP2’s role from synaptic regulation to broader regulation of neural development and excitability. ([pubmed.ncbi.nlm.nih.gov](https://pubmed.ncbi.nlm.nih.gov/38149472/)) [S6]

Authoritative database curation is consistent with this picture. The Mouse Genome Informatics resource lists Baiap2 phenotypes in the nervous system, notes that adult homozygous knockout mice “fail to regulate synaptic plasticity” and show hippocampus-associated learning deficits, and includes GO classifications spanning synapse and homeostatic process. Database annotations are not equivalent to a single decisive experiment, but they reflect aggregation of the published literature and support the claim’s broad framing. ([informatics.jax.org](https://www.informatics.jax.org/marker/MGI%3A2137336)) [S7]

## Contradicting or limiting evidence

The main limitation is that the strongest evidence supports **regulatory roles in neural systems**, not necessarily **classical homeostatic plasticity** in the narrow neuroscience sense (for example, canonical synaptic scaling or firing-rate homeostasis). The review literature and primary studies retrieved here emphasize dendritic spine development, NMDA receptor regulation, synapse development, excitatory transmission, and E/I balance more than direct tests of homeostatic scaling paradigms. ([pubmed.ncbi.nlm.nih.gov](https://pubmed.ncbi.nlm.nih.gov/26275848/)) [S8]

The evidence base is also weighted toward **mouse, neuronal culture, and molecular/cellular studies** rather than human clinical physiology. Human relevance is supported by neurodevelopmental-disorder genetics, but those studies mainly show that BAIAP2 variation can perturb neural development/excitability; they do not by themselves prove a normal homeostatic role in the intact human brain. ([pubmed.ncbi.nlm.nih.gov](https://pubmed.ncbi.nlm.nih.gov/38149472/)) [S9]

Finally, BAIAP2’s neural effects are not uniform across all neuron types or contexts. Expression and phenotypic impact appear strongest in glutamatergic/excitatory neurons, and some electrophysiologic/behavioral effects are sex- and cell-type-dependent. That argues against overgeneralizing BAIAP2 as a universal regulator of all neural homeostasis. ([pubmed.ncbi.nlm.nih.gov](https://pubmed.ncbi.nlm.nih.gov/32116566/)) [S10]

## Mechanistic / biological rationale

Mechanistically, BAIAP2/IRSp53 is well positioned to regulate neural structure and signaling. It is a multidomain protein with an I-BAR domain that binds/deforms membranes, a CRIB-related region that interacts with Cdc42/Rac signaling, an SH3 domain that binds cytoskeletal effectors, and a PDZ-binding motif that links to postsynaptic scaffolds such as PSD-95 and Shank. This domain architecture allows it to couple membrane shape, actin dynamics, and postsynaptic scaffold organization—exactly the kind of integration expected of a synaptic regulatory protein. ([pmc.ncbi.nlm.nih.gov](https://pmc.ncbi.nlm.nih.gov/articles/PMC6666077/)) [S11]

That mechanism is consistent with the phenotypes observed experimentally: altered dendritic spine morphology, abnormal NMDA receptor-mediated transmission, changes in LTP, altered E/I ratio, and activity-dependent recruitment to the PSD. In other words, BAIAP2 appears to help tune synaptic organization and excitatory signaling rather than merely serving as a passive marker of neurons. ([pmc.ncbi.nlm.nih.gov](https://pmc.ncbi.nlm.nih.gov/articles/PMC6666077/)) [S12]

There is also a plausible biochemical “on/off” control mechanism. Nature Communications work showed that phosphorylation-dependent binding of 14-3-3 proteins can lock IRSp53 in an autoinhibited conformation, reducing membrane binding and interactions with Cdc42/cytoskeletal effectors. This provides a credible molecular basis for dynamic regulation of BAIAP2 function in response to signaling state. ([nature.com](https://www.nature.com/articles/s41467-019-08317-8)) [S13]

## Bottom line

The claim is **likely true**. BAIAP2/IRSp53 is well supported as a **neural regulatory protein**, especially in excitatory synapses, where it influences dendritic spine organization, postsynaptic-density assembly, NMDA receptor-mediated transmission, synaptic plasticity, E/I balance, and aspects of neuronal development. ([pubmed.ncbi.nlm.nih.gov](https://pubmed.ncbi.nlm.nih.gov/26275848/)) [S14]

The weaker part of the claim is the word **“homeostatic.”** There is some indirect support—such as altered E/I balance, maintenance of normal NMDA signaling, and GO/database annotation to homeostatic process—but I did not find strong direct evidence that BAIAP2 has been specifically established as a core mediator of canonical neural homeostatic plasticity. So the safest formulation is: **BAIAP2 is strongly associated with regulatory processes in neural systems, and only more loosely/directly with neural homeostasis.** ([pubmed.ncbi.nlm.nih.gov](https://pubmed.ncbi.nlm.nih.gov/32116566/)) [S15]

## Cited sources

- **[S1]** IRSp53/BAIAP2 in dendritic spine development, NMDA receptor regulation, and psychiatric disorders - PubMed — https://pubmed.ncbi.nlm.nih.gov/26275848/
- **[S2]** 
            Enhanced NMDA Receptor-Mediated Synaptic Transmission, Enhanced Long-Term Potentiation, and Impaired Learning and Memory in Mice Lacking IRSp53 - PMC
         — https://pmc.ncbi.nlm.nih.gov/articles/PMC6666077/
- **[S3]** IRSp53 promotes postsynaptic density formation and actin filament bundling - PubMed — https://pubmed.ncbi.nlm.nih.gov/35819332/
- **[S4]** IRSp53 accumulates at the postsynaptic density under excitatory conditions - PubMed — https://pubmed.ncbi.nlm.nih.gov/29284046/
- **[S5]** IRSp53 Deletion in Glutamatergic and GABAergic Neurons and in Male and Female Mice Leads to Distinct Electrophysiological and Behavioral Phenotypes - PubMed — https://pubmed.ncbi.nlm.nih.gov/32116566/
- **[S6]** A lissencephaly-associated BAIAP2 variant causes defects in neuronal migration during brain development - PubMed — https://pubmed.ncbi.nlm.nih.gov/38149472/
- **[S7]** Baiap2 MGI Mouse Gene Detail - MGI:2137336 - brain-specific angiogenesis inhibitor 1-associated protein 2 — https://www.informatics.jax.org/marker/MGI%3A2137336
- **[S8]** IRSp53/BAIAP2 in dendritic spine development, NMDA receptor regulation, and psychiatric disorders - PubMed — https://pubmed.ncbi.nlm.nih.gov/26275848/
- **[S9]** A lissencephaly-associated BAIAP2 variant causes defects in neuronal migration during brain development - PubMed — https://pubmed.ncbi.nlm.nih.gov/38149472/
- **[S10]** IRSp53 Deletion in Glutamatergic and GABAergic Neurons and in Male and Female Mice Leads to Distinct Electrophysiological and Behavioral Phenotypes - PubMed — https://pubmed.ncbi.nlm.nih.gov/32116566/
- **[S11]** 
            Enhanced NMDA Receptor-Mediated Synaptic Transmission, Enhanced Long-Term Potentiation, and Impaired Learning and Memory in Mice Lacking IRSp53 - PMC
         — https://pmc.ncbi.nlm.nih.gov/articles/PMC6666077/
- **[S12]** 
            Enhanced NMDA Receptor-Mediated Synaptic Transmission, Enhanced Long-Term Potentiation, and Impaired Learning and Memory in Mice Lacking IRSp53 - PMC
         — https://pmc.ncbi.nlm.nih.gov/articles/PMC6666077/
- **[S13]** Mechanism of IRSp53 inhibition by 14-3-3 | Nature Communications — https://www.nature.com/articles/s41467-019-08317-8
- **[S14]** IRSp53/BAIAP2 in dendritic spine development, NMDA receptor regulation, and psychiatric disorders - PubMed — https://pubmed.ncbi.nlm.nih.gov/26275848/
- **[S15]** IRSp53 Deletion in Glutamatergic and GABAergic Neurons and in Male and Female Mice Leads to Distinct Electrophysiological and Behavioral Phenotypes - PubMed — https://pubmed.ncbi.nlm.nih.gov/32116566/
