## Supplementary material for "Understanding Substance Dependence: What Differentiates Addictive from Non-Addictive Drugs?": Folder for supplementary files: non-addictive-genes-claims-first100-040-agap2-gene-protein-has-been-associated-with-regula-23fa45b81495.md

# non-addictive-genes-claims-first100-040-23fa45b81495

**Original claim:** AGAP2 gene/protein has been associated with regulatory or homeostatic biological processes in neural systems.

## Claim

AGAP2 gene/protein has been associated with regulatory or homeostatic biological processes in neural systems.

## Verdict

Verdict: Likely True

The claim is supported **for neural regulatory functions**, especially when AGAP2 is considered at the **gene/isoform level**, because brain-enriched AGAP2 products such as **PIKE-L** are linked to neuronal survival, dendritogenesis, glutamate-receptor trafficking, long-term potentiation, excitotoxic resistance, and CNS myelination. The evidence is weaker for **direct, classical “homeostatic” roles** specifically, but the broader statement about neural regulatory/maintenance processes is well supported. ([ncbi.nlm.nih.gov](https://www.ncbi.nlm.nih.gov/gene/116986)) [S1]

## Confidence

Medium

Confidence is not low because there are multiple primary studies in neurons and mouse CNS showing AGAP2/PIKE involvement in core neural regulatory processes. It is not high because much of the strongest evidence is in **animal or cellular systems**, and because the neural literature is weighted toward the **PIKE-L/PIKE-S isoforms** rather than the more ubiquitous AGAP2/PIKE-A isoform; direct evidence for canonical neural “homeostasis” paradigms is less explicit than for synaptic signaling and cell-survival regulation. ([pmc.ncbi.nlm.nih.gov](https://pmc.ncbi.nlm.nih.gov/articles/PMC9818964/)) [S2]

## Supporting evidence

AGAP2 is plausibly a neural gene at baseline. In human reference resources, AGAP2 shows **biased expression in brain** in NCBI Gene, and the Human Protein Atlas classifies it as **tissue enriched in brain** and places it in a **“Brain - Synaptic signal transduction”** expression cluster with maximal confidence. NCBI also lists a reviewed human transcript variant encoding **PIKE-L**, described as participating in the **prevention of neuronal apoptosis**. ([ncbi.nlm.nih.gov](https://www.ncbi.nlm.nih.gov/gene/116986)) [S3]

Isoform biology is important here. A review of AGAP2/PIKE family biology notes that one gene yields multiple products, including **PIKE-L, AGAP2/PIKE-A, and PIKE-S**; the same source states that **PIKE-S and PIKE-L are brain-restricted**, whereas AGAP2/PIKE-A is not. Independent work likewise describes two human AGAP2 gene isoforms, with **isoform 1 encoding PIKE-L** and **isoform 2 encoding AGAP2/PIKE-A**. That means the strongest neural evidence belongs mainly to **brain-specific AGAP2 isoforms**, which still supports the gene-level claim. ([pmc.ncbi.nlm.nih.gov](https://pmc.ncbi.nlm.nih.gov/articles/PMC9818964/)) [S4]

Several primary studies show a direct role in **neuronal survival signaling**. In an early review, PIKE-L is described as coupling group I metabotropic glutamate receptors to PI3K and regulating **prevention of neuronal apoptosis**. A separate primary paper found that **netrin-1-mediated neuronal survival** depends on PIKE-L interaction with the dependence receptor **UNC5B**, with PIKE-L acting as a downstream survival effector in the nervous system. NCBI Gene also summarizes experimental evidence and GeneRIFs linking Agap2/Centg1 to **negative regulation of neuron apoptotic process** and neuronal protection. ([pubmed.ncbi.nlm.nih.gov](https://pubmed.ncbi.nlm.nih.gov/15951849/)) [S5]

In vivo developmental and maintenance phenotypes strengthen the case. A mouse study reported that PIKE deficiency causes a **thinner cortex, reduced neuronal density, smaller neuronal soma size**, impaired memory-related behavior, and increased vulnerability to neurotoxic injury, concluding that PIKE is important for **neuronal development, survival, and function** through PI3K/Akt signaling. Another study showed PIKE-deficient neurons are **hypersensitive to excitotoxic stimulation**, with higher calcium permeability, greater cell death after kainate exposure, and worse neurodegeneration in vivo. These are classic neural regulatory/protective phenotypes, even if not always labeled “homeostasis.” ([pmc.ncbi.nlm.nih.gov](https://pmc.ncbi.nlm.nih.gov/articles/PMC3551346/?utm_source=openai)) [S6]

There is also strong evidence for a synaptic regulatory role. In hippocampal/cortical neuron work, **PIKE-L was required for GluA2-containing AMPA receptor surface expression** and for **LTP-related trafficking**, linking NMDA-dependent signaling to AMPAR insertion and synaptic strengthening. The same paper concluded that PIKE is important from **LTP formation to neuronal survival**. A later review of PIKE in the brain likewise summarizes defective LTP, reduced surface GluA2, and memory impairment in PIKE-deficient animals. ([pmc.ncbi.nlm.nih.gov](https://pmc.ncbi.nlm.nih.gov/articles/PMC3199380/)) [S7]

Disease-model data point in the same direction. In Fragile X models, elevated PIKE contributed to abnormal PI3K signaling, dendritic spine density, synaptic plasticity, cortical network activity, and behavior; **genetic reduction of PIKE normalized molecular/cellular plasticity and improved behavior** in mouse and fly models. That does not prove AGAP2 is a human disease gene by itself, but it does show that changing AGAP2/PIKE dosage alters regulated neural phenotypes. ([pmc.ncbi.nlm.nih.gov](https://pmc.ncbi.nlm.nih.gov/articles/PMC4418204/)) [S8]

The neural association is not limited to neurons. A PNAS study found that **PIKE-L is CNS-specific**, expressed across multiple brain regions, and required for **oligodendroglial development and CNS myelination** via PI3K/Akt signaling. Myelination and remyelination are tissue-maintenance processes in the CNS, so this expands the evidence from synaptic regulation to broader neural system maintenance. ([pmc.ncbi.nlm.nih.gov](https://pmc.ncbi.nlm.nih.gov/articles/PMC3918755/)) [S9]

## Contradicting or limiting evidence

The main limitation is **terminology and scope**. The retrieved literature strongly supports AGAP2/PIKE in **regulatory neural processes** such as signaling, receptor trafficking, development, plasticity, and survival, but the evidence is less direct for AGAP2 as a dedicated mediator of **canonical neural homeostasis** in the narrow sense used for phenomena like homeostatic synaptic scaling. In the studies retrieved, the emphasis was on PI3K/Akt signaling, excitotoxic protection, dendritogenesis, LTP, and myelination. ([pmc.ncbi.nlm.nih.gov](https://pmc.ncbi.nlm.nih.gov/articles/PMC3199380/)) [S10]

A second limitation is **isoform ambiguity**. Some neural papers use the historical name **PIKE/Centg1** and focus on **PIKE-L** or PIKE-S, whereas the shorter **AGAP2/PIKE-A** isoform is more often discussed in cancer and non-neural contexts. So the statement is strongest when interpreted as referring to the **AGAP2 gene and its brain isoforms**, rather than every AGAP2 protein product equally. ([pmc.ncbi.nlm.nih.gov](https://pmc.ncbi.nlm.nih.gov/articles/PMC9818964/)) [S11]

Finally, the most compelling functional evidence comes from **mouse models, cultured neurons, and mechanistic cell biology**, not from controlled human clinical studies. Human data more clearly support **brain-enriched expression** than direct proof of neural-system function. ([ncbi.nlm.nih.gov](https://www.ncbi.nlm.nih.gov/gene/116986)) [S12]

## Mechanistic / biological rationale

Mechanistically, AGAP2-family proteins are well positioned to regulate neural systems because they combine **small-GTPase-related domains** with scaffolding/signaling interactions. NCBI lists AGAP2 functions related to **GTPase activity/activation**, **ionotropic glutamate receptor binding**, and **positive regulation of PI3K/AKT signaling**. In neural contexts, PIKE-L links extracellular cues such as **mGluR activation, netrin signaling, and glutamatergic activity** to intracellular **PI3K/Akt** pathways that support neuronal survival and synaptic responses. ([ncbi.nlm.nih.gov](https://www.ncbi.nlm.nih.gov/gene/216439)) [S13]

That mechanism plausibly explains the observed phenotypes: PI3K/Akt signaling supports **cell survival**, dendritic growth, and aspects of synaptic plasticity; PIKE-L also promotes **GluA2 surface trafficking**, which affects excitatory synaptic strength and resistance to excitotoxic calcium entry. In oligodendrocytes, the same signaling logic extends to **myelinogenesis**. Together, these functions fit a role in neural regulation and maintenance of stable function under developmental and activity-dependent demands. ([pmc.ncbi.nlm.nih.gov](https://pmc.ncbi.nlm.nih.gov/articles/PMC3199380/)) [S14]

## Bottom line

Overall, the claim is **likely true**. AGAP2—especially its brain-enriched isoforms **PIKE-L/PIKE-S**—has substantial evidence linking it to **regulatory processes in neural systems**, including neuronal survival, synaptic receptor trafficking, plasticity, dendritic development, excitotoxic protection, and myelination. The main caveat is that the literature more clearly supports **neural regulation/maintenance** than a narrowly defined, directly tested role in **classical neural homeostasis**. ([ncbi.nlm.nih.gov](https://www.ncbi.nlm.nih.gov/gene/116986)) [S15]

## Cited sources

- **[S1]** AGAP2 ArfGAP with GTPase domain, ankyrin repeat and PH domain 2 [Homo sapiens (human)] - Gene - NCBI — https://www.ncbi.nlm.nih.gov/gene/116986
- **[S2]** 
            Functional Role of AGAP2/PIKE-A in Fcγ Receptor-Mediated Phagocytosis - PMC
         — https://pmc.ncbi.nlm.nih.gov/articles/PMC9818964/
- **[S3]** AGAP2 ArfGAP with GTPase domain, ankyrin repeat and PH domain 2 [Homo sapiens (human)] - Gene - NCBI — https://www.ncbi.nlm.nih.gov/gene/116986
- **[S4]** 
            Functional Role of AGAP2/PIKE-A in Fcγ Receptor-Mediated Phagocytosis - PMC
         — https://pmc.ncbi.nlm.nih.gov/articles/PMC9818964/
- **[S5]** PIKE GTPase signaling and function - PubMed — https://pubmed.ncbi.nlm.nih.gov/15951849/
- **[S6]** Phosphoinositide 3-Kinase Enhancer Regulates Neuronal Dendritogenesis and Survival in Neocortex - PMC — https://pmc.ncbi.nlm.nih.gov/articles/PMC3551346/?utm_source=openai
- **[S7]** 
            PIKE-mediated PI3-kinase activity is required for AMPA receptor surface expression - PMC
         — https://pmc.ncbi.nlm.nih.gov/articles/PMC3199380/
- **[S8]** 
            Increased expression of the PI3K enhancer PIKE mediates deficits in synaptic plasticity and behavior in Fragile X syndrome - PMC
         — https://pmc.ncbi.nlm.nih.gov/articles/PMC4418204/
- **[S9]** 
            PIKE is essential for oligodendroglia development and CNS myelination - PMC
         — https://pmc.ncbi.nlm.nih.gov/articles/PMC3918755/
- **[S10]** 
            PIKE-mediated PI3-kinase activity is required for AMPA receptor surface expression - PMC
         — https://pmc.ncbi.nlm.nih.gov/articles/PMC3199380/
- **[S11]** 
            Functional Role of AGAP2/PIKE-A in Fcγ Receptor-Mediated Phagocytosis - PMC
         — https://pmc.ncbi.nlm.nih.gov/articles/PMC9818964/
- **[S12]** AGAP2 ArfGAP with GTPase domain, ankyrin repeat and PH domain 2 [Homo sapiens (human)] - Gene - NCBI — https://www.ncbi.nlm.nih.gov/gene/116986
- **[S13]** Agap2 ArfGAP with GTPase domain, ankyrin repeat and PH domain 2 [Mus musculus (house mouse)] - Gene - NCBI — https://www.ncbi.nlm.nih.gov/gene/216439
- **[S14]** 
            PIKE-mediated PI3-kinase activity is required for AMPA receptor surface expression - PMC
         — https://pmc.ncbi.nlm.nih.gov/articles/PMC3199380/
- **[S15]** AGAP2 ArfGAP with GTPase domain, ankyrin repeat and PH domain 2 [Homo sapiens (human)] - Gene - NCBI — https://www.ncbi.nlm.nih.gov/gene/116986
