## Supplementary material for "Understanding Substance Dependence: What Differentiates Addictive from Non-Addictive Drugs?": Folder for supplementary files: non-addictive-genes-claims-first100-041-nptx2-gene-protein-has-been-associated-with-regula-9bbba5e893f0.md

# non-addictive-genes-claims-first100-041-9bbba5e893f0

**Original claim:** NPTX2 gene/protein has been associated with regulatory or homeostatic biological processes in neural systems.

## Claim

NPTX2 gene/protein has been associated with regulatory or homeostatic biological processes in neural systems.

## Verdict

Verdict: Highly Supported

## Confidence

High

## Supporting evidence

NPTX2 encodes neuronal pentraxin-2, also called Narp/NP2, a synaptic protein expressed in brain and linked by authoritative reference resources to excitatory synapse formation, AMPA-receptor clustering, glutamatergic synapses, and chemical synaptic transmission. That basic annotation already places it in neural circuit regulation rather than being a merely structural or unrelated protein. ([ncbi.nlm.nih.gov](https://www.ncbi.nlm.nih.gov/gene/4885)) [S1]

The strongest direct evidence comes from primary mechanistic work showing that NPTX2/Narp participates in **homeostatic synaptic scaling**. In mouse/neuronal culture experiments, Chang et al. reported that Narp “regulates homeostatic scaling of excitatory synapses on parvalbumin interneurons,” with activity-dependent accumulation of Narp at excitatory synapses on PV interneurons and a requirement for Narp in activity-regulated changes in GluA4-mediated synaptic strength. That is a direct demonstration of a homeostatic regulatory role in neural circuitry. ([pmc.ncbi.nlm.nih.gov](https://pmc.ncbi.nlm.nih.gov/articles/PMC2949072/)) [S2]

Independent work places NPTX2 inside broader **circuit homeostasis programs**. In a Cell study on the activity-dependent transcription factor NPAS4, Nptx2 was identified as part of a cell-type-specific program whose shared goal is restricting overall network excitation. The authors explicitly describe Npas4-driven gene programs, including Nptx2, as achieving a “circuit-wide homeostatic response,” promoting increased excitation onto inhibitory neurons so that inhibition rises and network activity is restrained. ([pubmed.ncbi.nlm.nih.gov](https://pubmed.ncbi.nlm.nih.gov/24855953/)) [S3]

A second major line of evidence concerns **excitatory/inhibitory (E/I) balance via parvalbumin interneurons**. NPTX2 acts at excitatory synapses onto PV interneurons; in vivo deletion reduces pyramidal-neuron drive onto PV cells, and in Alzheimer-related work the authors frame this as a pyramidal neuron–NPTX2–PV interneuron **homeostatic mechanism**. In aging/AD-related human and mouse work, lower NPTX2 is linked to reduced GluA4 on PV interneurons and consequent circuit hyperexcitability, supporting a role in stabilizing inhibitory circuit recruitment. ([pmc.ncbi.nlm.nih.gov](https://pmc.ncbi.nlm.nih.gov/articles/PMC5404919/)) [S4]

Related developmental/circuit studies further show that neuronal pentraxins coordinate PV-interneuron maturation and circuit integration. In NPTX2/NPTXR-deficient mice, GluA4 protein at PV-interneuron synapses is markedly reduced despite preserved GluA4 mRNA, AMPA/NMDA ratios onto PV cells fall, feedforward inhibition is weakened, and network rhythms such as gamma oscillations are impaired. Those findings are consistent with NPTX2-family control of synaptic receptor stabilization and network-level regulation of excitation. ([pmc.ncbi.nlm.nih.gov](https://pmc.ncbi.nlm.nih.gov/articles/PMC4368480/)) [S5]

NPTX2 is also tied to **critical-period and sensory-plasticity regulation**. In juvenile visual cortex, brief monocular deprivation transiently lowers surface NPTX2; experimentally increasing NPTX2 prevents deprivation-induced pyramidal-to-PV disconnection and prevents ocular-dominance plasticity, whereas interfering with NPTX2 in adults enables juvenile-like ocular-dominance shifts. This shows NPTX2 functioning as a gatekeeper of plasticity-related circuit regulation, not merely a passive marker. ([pmc.ncbi.nlm.nih.gov](https://pmc.ncbi.nlm.nih.gov/articles/PMC8449314/)) [S6]

Evidence extends beyond synaptic scaling and critical periods into **extracellular matrix / perineuronal net regulation**. NPTX2 binds perineuronal-net glycosaminoglycans, is retained in PNN-rich fractions, and in neuronal culture enhances PNN formation. Because PNNs are major regulators of PV interneuron maturation, stability, and closure of heightened plasticity states, these data support an additional regulatory association of NPTX2 with neural homeostasis. ([pmc.ncbi.nlm.nih.gov](https://pmc.ncbi.nlm.nih.gov/articles/PMC6854953/)) [S7]

There is also evidence for a role in **immune-like homeostasis of synapses** in the adult brain. In a Science Translational Medicine study, Nptx2 bound complement C1q, limited classical complement pathway activity, and restrained microglia-mediated excitatory synapse loss. The authors concluded Nptx2 is, to their knowledge, the first neuronal protein shown to regulate this complement pathway in the adult brain. That is a strong example of regulatory control over synapse maintenance/homeostasis, albeit in a neurodegeneration-focused context. ([pmc.ncbi.nlm.nih.gov](https://pmc.ncbi.nlm.nih.gov/articles/PMC10467038/)) [S8]

Human data are more associative than causal but are directionally consistent. A 2022 review summarized neuronal pentraxins, including NPTX2, as proteins “involved in homeostatic synaptic plasticity” by recruiting postsynaptic receptors into synapses. In schizophrenia-related human CSF work, the authors further state that NPTX2 “plays an essential role in maintaining excitatory homeostasis by adaptively enhancing circuit inhibition,” and they observed diurnal CSF oscillation plus sleep-deprivation sensitivity, linking NPTX2 to dynamically regulated physiological circuit states. ([pubmed.ncbi.nlm.nih.gov](https://pubmed.ncbi.nlm.nih.gov/34460014/)) [S9]

Finally, NPTX2 has even been implicated in **circadian/homeostatic regulation of structural synaptic plasticity** in vivo. In zebrafish hypocretin neurons, rhythmic nptx2b expression modulated circadian synaptic changes, and arrhythmic overexpression increased synapse number and abolished rhythmicity. While this is not mammalian cortical homeostasis per se, it reinforces the general claim that NPTX2-family proteins are associated with neural regulatory/homeostatic processes. ([pubmed.ncbi.nlm.nih.gov](https://pubmed.ncbi.nlm.nih.gov/20920793/)) [S10]

## Contradicting or limiting evidence

The main limitation is that the best causal evidence is **preclinical**. Strong demonstrations of homeostatic scaling, E/I-balance regulation, critical-period gating, and complement control come mainly from mouse, rat, zebrafish, and culture systems rather than interventional human studies. Human evidence is mostly biomarker/correlation-based, especially in AD, FTD, and schizophrenia cohorts. ([pmc.ncbi.nlm.nih.gov](https://pmc.ncbi.nlm.nih.gov/articles/PMC2949072/)) [S11]

Also, not every phenotype is robust with NPTX2 loss alone. In the AD-focused review, Nptx2 knockout by itself did **not** produce marked PV-interneuron dysfunction in some gamma-oscillation assays, whereas combined loss with Nptxr produced much stronger effects. Likewise, some of the strongest circuit-deficit data come from NPTX2/NPTXR double mutants rather than NPTX2 single mutants. That means NPTX2 is important, but sometimes within a pentraxin complex rather than as a lone determinant. ([pmc.ncbi.nlm.nih.gov](https://pmc.ncbi.nlm.nih.gov/articles/PMC5404919/)) [S12]

Some proposed downstream roles are also context-dependent. For example, NPTX2 enhanced PNN formation in vitro, but prior adult in vivo observations did not show obvious WFA-labeled PNN changes in knockout mice, suggesting that certain extracellular-matrix effects may vary by developmental stage, region, or assay. ([pmc.ncbi.nlm.nih.gov](https://pmc.ncbi.nlm.nih.gov/articles/PMC6854953/)) [S13]

## Mechanistic / biological rationale

Mechanistically, NPTX2 is well positioned to regulate neural homeostasis because it is an **activity-induced extracellular synaptic organizer**. It is produced by excitatory neurons, trafficked/exocytosed at excitatory synapses, and helps cluster/stabilize AMPA receptors—especially GluA4-containing receptors—at synapses onto PV interneurons. Strengthening those excitatory inputs raises recruitment of inhibitory PV cells, which in turn restrains overall circuit excitation. That is a textbook negative-feedback/homeostatic architecture. ([pmc.ncbi.nlm.nih.gov](https://pmc.ncbi.nlm.nih.gov/articles/PMC2949072/)) [S14]

NPTX2 also appears to operate across multiple regulatory layers: synaptic receptor stabilization, critical-period plasticity gating, extracellular-matrix/PNN interactions, behavior- and sleep-linked trafficking, and complement-mediated protection from excess synapse pruning. These roles are mechanistically coherent because all affect how neural circuits maintain stable yet adaptable levels of excitation, inhibition, and connectivity. ([pmc.ncbi.nlm.nih.gov](https://pmc.ncbi.nlm.nih.gov/articles/PMC8449314/)) [S15]

## Bottom line

Yes. The claim is **highly supported**. NPTX2 is repeatedly linked—by primary experimental studies, reviews, and authoritative databases—to **homeostatic synaptic plasticity, regulation of excitatory drive onto inhibitory PV interneurons, E/I balance, critical-period plasticity, and synapse-maintenance mechanisms** in neural systems. The clearest causal evidence is from animal and cellular studies; in humans, the evidence is more associative/biomarker-based. But taken together, the literature strongly supports the statement that NPTX2 is associated with regulatory and homeostatic biological processes in the nervous system. ([ncbi.nlm.nih.gov](https://www.ncbi.nlm.nih.gov/gene/4885)) [S16]

## Cited sources

- **[S1]** NPTX2 neuronal pentraxin 2 [Homo sapiens (human)] - Gene - NCBI — https://www.ncbi.nlm.nih.gov/gene/4885
- **[S2]** 
            Narp regulates homeostatic scaling of excitatory synapses on Parvalbumin interneurons - PMC
         — https://pmc.ncbi.nlm.nih.gov/articles/PMC2949072/
- **[S3]** Npas4 regulates excitatory-inhibitory balance within neural circuits through cell-type-specific gene programs - PubMed — https://pubmed.ncbi.nlm.nih.gov/24855953/
- **[S4]** 
            NPTX2 and cognitive dysfunction in Alzheimer’s Disease - PMC
         — https://pmc.ncbi.nlm.nih.gov/articles/PMC5404919/
- **[S5]** 
            Pentraxins coordinate excitatory synapse maturation and circuit integration of parvalbumin interneurons - PMC
         — https://pmc.ncbi.nlm.nih.gov/articles/PMC4368480/
- **[S6]** 
            All-or-none disconnection of pyramidal inputs onto parvalbumin-positive interneurons gates ocular dominance plasticity - PMC
         — https://pmc.ncbi.nlm.nih.gov/articles/PMC8449314/
- **[S7]** 
            Neuronal Pentraxin 2 Binds PNNs and Enhances PNN Formation - PMC
         — https://pmc.ncbi.nlm.nih.gov/articles/PMC6854953/
- **[S8]** 
            The neuronal pentraxin Nptx2 regulates complement activity and restrains microglia-mediated synapse loss in neurodegeneration - PMC
         — https://pmc.ncbi.nlm.nih.gov/articles/PMC10467038/
- **[S9]** Neuronal pentraxins as biomarkers of synaptic activity: from physiological functions to pathological changes in neurodegeneration - PubMed — https://pubmed.ncbi.nlm.nih.gov/34460014/
- **[S10]** Circadian and homeostatic regulation of structural synaptic plasticity in hypocretin neurons - PubMed — https://pubmed.ncbi.nlm.nih.gov/20920793/
- **[S11]** 
            Narp regulates homeostatic scaling of excitatory synapses on Parvalbumin interneurons - PMC
         — https://pmc.ncbi.nlm.nih.gov/articles/PMC2949072/
- **[S12]** 
            NPTX2 and cognitive dysfunction in Alzheimer’s Disease - PMC
         — https://pmc.ncbi.nlm.nih.gov/articles/PMC5404919/
- **[S13]** 
            Neuronal Pentraxin 2 Binds PNNs and Enhances PNN Formation - PMC
         — https://pmc.ncbi.nlm.nih.gov/articles/PMC6854953/
- **[S14]** 
            Narp regulates homeostatic scaling of excitatory synapses on Parvalbumin interneurons - PMC
         — https://pmc.ncbi.nlm.nih.gov/articles/PMC2949072/
- **[S15]** 
            All-or-none disconnection of pyramidal inputs onto parvalbumin-positive interneurons gates ocular dominance plasticity - PMC
         — https://pmc.ncbi.nlm.nih.gov/articles/PMC8449314/
- **[S16]** NPTX2 neuronal pentraxin 2 [Homo sapiens (human)] - Gene - NCBI — https://www.ncbi.nlm.nih.gov/gene/4885
