## Supplementary material for "Understanding Substance Dependence: What Differentiates Addictive from Non-Addictive Drugs?": Folder for supplementary files: non-addictive-genes-claims-first100-042-sgk1-gene-protein-has-been-associated-with-regulat-c2717c6ddd79.md

# non-addictive-genes-claims-first100-042-c2717c6ddd79

**Original claim:** SGK1 gene/protein has been associated with regulatory or homeostatic biological processes in neural systems.

## Claim

SGK1 gene/protein has been associated with regulatory or homeostatic biological processes in neural systems.

## Verdict

Verdict: Likely True

SGK1 is well associated with **regulatory** processes in neural systems, including synaptic plasticity, glutamate receptor trafficking, intrinsic excitability, stress-responsive neural progenitor regulation, and oligodendrocyte remodeling. Evidence for a broad **homeostatic** role is also plausible, because SGK1 modulates ion channels, glutamate handling, and stress-feedback pathways that can stabilize neural function, but that part of the claim is usually inferred from mechanism rather than tested directly as “neural homeostasis.” Much of the evidence is from rodent or cell models rather than definitive human in vivo studies. ([learnmem.cshlp.org](https://learnmem.cshlp.org/content/13/2/114.full.pdf)) [S1]

## Confidence

High

There are multiple independent lines of evidence across neurons, neural progenitors, and glia, plus some human association data, all pointing to SGK1 as a neural regulatory kinase. Confidence is not maximal because the literature is more direct for **regulation/plasticity/excitability** than for the broader term **homeostasis**, and because much of the causal work is preclinical. ([pmc.ncbi.nlm.nih.gov](https://pmc.ncbi.nlm.nih.gov/articles/PMC3666742/)) [S2]

## Supporting evidence

SGK1 has been linked to **hippocampal synaptic plasticity and memory-related regulation**. A review of neuronal SGK1 summarized that hippocampal SGK1 expression rises after fear conditioning and enrichment, and that SGK1 affects long-term potentiation, dendrite growth, glutamate receptor regulation, glutamate transporter activity, and Kv-channel-dependent neuronal excitability. In a primary hippocampal LTP study, SGK phosphorylation increased after tetanization, dominant-negative SGK impaired the **expression** of LTP, and constitutively active SGK increased PSD-95 expression, supporting a direct role in neuronal plasticity rather than mere passive association. ([pmc.ncbi.nlm.nih.gov](https://pmc.ncbi.nlm.nih.gov/articles/PMC2988501/)) [S3]

SGK1 also regulates **glutamatergic synaptic transmission** in prefrontal cortex under stress. In rodent PFC neurons, acute stress or corticosterone increased synaptic and surface AMPAR/NMDAR expression through a **GR/SGK/Rab4** pathway, and blocking SGK prevented the stress-induced enhancement of working memory. A mechanistic JBC study showed corticosterone increases SGK-dependent phosphorylation of GDI at Ser-213, promoting GDI-Rab4 complex formation and Rab4-mediated recycling of AMPARs to the synaptic membrane. That is a clear example of SGK1 participating in neural regulatory circuitry controlling synaptic efficacy. ([pmc.ncbi.nlm.nih.gov](https://pmc.ncbi.nlm.nih.gov/articles/PMC3108461/)) [S4]

There is also evidence that SGK1 regulates **neural progenitor behavior and glucocorticoid signaling**. In human hippocampal progenitor cells, SGK1 mediated cortisol-induced reductions in proliferation and neuronal differentiation, acting both downstream of glucocorticoid receptor activation and upstream by enhancing GR phosphorylation and nuclear translocation. The same study found elevated SGK1 mRNA in the hippocampus of stressed rats and in peripheral blood of drug-free depressed patients, connecting SGK1 to stress-related regulation of neurogenesis in a human-relevant system. ([pmc.ncbi.nlm.nih.gov](https://pmc.ncbi.nlm.nih.gov/articles/PMC3666742/)) [S5]

A brain-specific neuronal isoform, **SGK1.1**, provides especially strong support for a role in regulating neural excitability. One study reported that SGK1.1 decreases neuronal **ASIC1** expression, a mechanism proposed to limit injury associated with ASIC1 activation during ischemia. Other studies showed SGK1.1 increases Kv7-family channel abundance and M-current, can partially rescue epilepsy-associated Kv7.2 loss-of-function mutations, and in transgenic mice reduces seizure severity and lowers hippocampal neuron excitability without grossly disrupting basal synaptic transmission. Those findings fit closely with a homeostatic/excitability-regulatory role. ([pubmed.ncbi.nlm.nih.gov](https://pubmed.ncbi.nlm.nih.gov/18334630/)) [S6]

SGK1 has also been implicated in **glial and white-matter regulation**, not just neuronal signaling. In a stress model, repeated corticosterone-dependent activation of the **PDK1-SGK1-NDRG1** pathway in oligodendrocytes increased adhesion-molecule expression and altered oligodendrocyte morphology in corpus callosum. A related study found that Sgk1 expression in adult rat brain under acute stress is dynamically regulated in oligodendrocytes, suggesting SGK1 participates in activity- and hormone-responsive neuron-glia interactions. These are regulatory processes within neural systems, even if not classic synaptic plasticity. ([pmc.ncbi.nlm.nih.gov](https://pmc.ncbi.nlm.nih.gov/articles/PMC3104997/)) [S7]

Human disease-associated data point in the same direction. In postmortem prefrontal cortex from PTSD subjects, SGK1 expression was markedly reduced, and experimentally decreasing SGK1 function in rat mPFC caused stress-related behavioral deficits, reduced dendritic spines, and altered electrophysiological properties. This does not prove a universal protective role, but it strongly supports the idea that SGK1 normally helps regulate neural structure and function. ([pmc.ncbi.nlm.nih.gov](https://pmc.ncbi.nlm.nih.gov/articles/PMC4623974/)) [S8]

## Contradicting or limiting evidence

The main limitation is that **direct human causal evidence is limited**. Much of the strongest evidence comes from cultured neurons, rodent stress models, slice physiology, or transgenic animals rather than human intervention studies. Human findings are mostly associative, such as altered SGK1 expression in depression- or PTSD-related contexts, so translation to general “neural homeostasis” should be stated cautiously. ([pmc.ncbi.nlm.nih.gov](https://pmc.ncbi.nlm.nih.gov/articles/PMC3666742/)) [S9]

SGK1’s effects are also **context-dependent and sometimes maladaptive**, which limits a simplistic interpretation of SGK1 as uniformly homeostatic. In human hippocampal progenitor cells, increased SGK1 mediated cortisol-induced suppression of proliferation and neuronal differentiation. A hippocampal study further reported that SGK1 activity can reduce dendritic spines in dorsal hippocampus. A recent review likewise describes SGK1 as having dual or even opposing roles across neurological conditions, including beneficial effects in some settings and harmful effects in others. ([pmc.ncbi.nlm.nih.gov](https://pmc.ncbi.nlm.nih.gov/articles/PMC3666742/)) [S10]

So, the strongest version of the claim is: **SGK1 is clearly associated with neural regulatory processes; its role in neural homeostasis is plausible and mechanistically supported, but often indirect and highly state-dependent.** ([pmc.ncbi.nlm.nih.gov](https://pmc.ncbi.nlm.nih.gov/articles/PMC2988501/)) [S11]

## Mechanistic / biological rationale

Mechanistically, SGK1 is a **stress- and PI3K/PDK1-responsive serine/threonine kinase** that can alter membrane protein trafficking, transcriptional responses, and cell-state transitions. In neurons, SGK1 promotes trafficking of AMPARs and NMDARs through **GDI/Rab4-dependent recycling**, which changes synaptic strength. It also regulates ion-channel systems such as Kv7/M-current and ASIC1, giving it a direct route to influence intrinsic excitability and resistance to overexcitation or injury. ([pmc.ncbi.nlm.nih.gov](https://pmc.ncbi.nlm.nih.gov/articles/PMC3108461/)) [S12]

In progenitor and stress-response biology, SGK1 forms part of a **glucocorticoid feedback module**: glucocorticoids induce SGK1, and SGK1 in turn enhances GR phosphorylation/nuclear translocation, thereby shaping transcriptional programs that affect proliferation and neuronal differentiation. In oligodendrocytes, SGK1 signals through **NDRG1 and adhesion molecules** to alter morphology and likely neuron-glia interaction states. Separately, the neuronal-function review notes SGK1 can stimulate glutamate transporters and regulate Kv channels, both of which are mechanistically compatible with maintaining excitatory balance. ([pmc.ncbi.nlm.nih.gov](https://pmc.ncbi.nlm.nih.gov/articles/PMC3666742/)) [S13]

Taken together, these pathways make biological sense for a protein involved in **regulating neural set-points, responsiveness, and adaptation**, even though the literature more often labels these effects as plasticity, stress signaling, or excitability control rather than explicitly “homeostasis.” ([pmc.ncbi.nlm.nih.gov](https://pmc.ncbi.nlm.nih.gov/articles/PMC3108461/)) [S14]

## Bottom line

The claim is **likely true**. SGK1 is convincingly associated with **regulatory** processes in neural systems—especially synaptic plasticity, receptor trafficking, excitability control, stress-responsive neurogenesis, and oligodendrocyte remodeling. Evidence for a broader **homeostatic** role is biologically credible but usually indirect and context-dependent, with SGK1 appearing protective in some settings and maladaptive in others. ([learnmem.cshlp.org](https://learnmem.cshlp.org/content/13/2/114.full.pdf)) [S15]

## Cited sources

- **[S1]** Untitled source — https://learnmem.cshlp.org/content/13/2/114.full.pdf
- **[S2]** 
            Role for the kinase SGK1 in stress, depression, and glucocorticoid effects on hippocampal neurogenesis - PMC
         — https://pmc.ncbi.nlm.nih.gov/articles/PMC3666742/
- **[S3]** 
            Significance of SGK1 in the regulation of neuronal function - PMC
         — https://pmc.ncbi.nlm.nih.gov/articles/PMC2988501/
- **[S4]** 
            Mechanisms for Acute Stress-Induced Enhancement of Glutamatergic Transmission and Working Memory - PMC
         — https://pmc.ncbi.nlm.nih.gov/articles/PMC3108461/
- **[S5]** 
            Role for the kinase SGK1 in stress, depression, and glucocorticoid effects on hippocampal neurogenesis - PMC
         — https://pmc.ncbi.nlm.nih.gov/articles/PMC3666742/
- **[S6]** A brain-specific SGK1 splice isoform regulates expression of ASIC1 in neurons - PubMed — https://pubmed.ncbi.nlm.nih.gov/18334630/
- **[S7]** 
            Plasma Corticosterone Activates SGK1 and Induces Morphological Changes in Oligodendrocytes in Corpus Callosum - PMC
         — https://pmc.ncbi.nlm.nih.gov/articles/PMC3104997/
- **[S8]** 
            Decreased SGK1 Expression and Function Contributes to Behavioral Deficits Induced by Traumatic Stress - PMC
         — https://pmc.ncbi.nlm.nih.gov/articles/PMC4623974/
- **[S9]** 
            Role for the kinase SGK1 in stress, depression, and glucocorticoid effects on hippocampal neurogenesis - PMC
         — https://pmc.ncbi.nlm.nih.gov/articles/PMC3666742/
- **[S10]** 
            Role for the kinase SGK1 in stress, depression, and glucocorticoid effects on hippocampal neurogenesis - PMC
         — https://pmc.ncbi.nlm.nih.gov/articles/PMC3666742/
- **[S11]** 
            Significance of SGK1 in the regulation of neuronal function - PMC
         — https://pmc.ncbi.nlm.nih.gov/articles/PMC2988501/
- **[S12]** 
            Mechanisms for Acute Stress-Induced Enhancement of Glutamatergic Transmission and Working Memory - PMC
         — https://pmc.ncbi.nlm.nih.gov/articles/PMC3108461/
- **[S13]** 
            Role for the kinase SGK1 in stress, depression, and glucocorticoid effects on hippocampal neurogenesis - PMC
         — https://pmc.ncbi.nlm.nih.gov/articles/PMC3666742/
- **[S14]** 
            Mechanisms for Acute Stress-Induced Enhancement of Glutamatergic Transmission and Working Memory - PMC
         — https://pmc.ncbi.nlm.nih.gov/articles/PMC3108461/
- **[S15]** Untitled source — https://learnmem.cshlp.org/content/13/2/114.full.pdf
