## Supplementary material for "Understanding Substance Dependence: What Differentiates Addictive from Non-Addictive Drugs?": Folder for supplementary files: non-addictive-genes-claims-first100-043-ptk2-gene-protein-has-been-associated-with-regulat-deae7b8858b6.md

# non-addictive-genes-claims-first100-043-deae7b8858b6

**Original claim:** PTK2 gene/protein has been associated with regulatory or homeostatic biological processes in neural systems.

## Claim

PTK2 gene/protein has been associated with regulatory or homeostatic biological processes in neural systems.

## Verdict

Verdict: Highly Supported

PTK2, which encodes focal adhesion kinase (FAK), is well supported as a regulator of multiple neural-system processes, especially neuronal migration, axon outgrowth/guidance, synapse formation/plasticity, and CNS myelination. The evidence is strongest for **regulatory** roles; evidence framed specifically as **neural homeostasis** is more limited and indirect. Because the claim uses “regulatory **or** homeostatic,” the overall claim is strongly supported. ([proteinatlas.org](https://www.proteinatlas.org/Q05397)) [S1]

## Confidence

High

This confidence is high because the conclusion rests on concordant evidence from curated reference resources plus several mechanistic studies in neural cells and mouse developmental/neurophysiology models, not on a single association study. The main caveat is that much of the strongest causal evidence is from animal and cell systems rather than direct human intervention studies. ([proteinatlas.org](https://www.proteinatlas.org/Q05397)) [S2]

## Supporting evidence

Authoritative annotation resources already place PTK2 in neural regulatory biology. The Human Protein Atlas page, citing UniProt function annotation, states that PTK2/FAK regulates axon growth, neuronal cell migration, axon branching, and synapse formation, and is required for normal nervous-system development. SynGO, a synapse-focused expert-curated ontology resource, places PTK2 at the postsynapse and links it to the biological process “regulation of modification of postsynaptic actin cytoskeleton,” which is a clear neural regulatory function. ([proteinatlas.org](https://www.proteinatlas.org/Q05397)) [S3]

Primary experimental studies support a direct role for PTK2/FAK in axon guidance and neurite regulation. In a 2004 *Nature Neuroscience* paper, the netrin receptor DCC was shown to interact with FAK; netrin-1 induced FAK phosphorylation, and disrupting FAK signaling abolished netrin-induced neurite outgrowth and attractive growth-cone turning. A companion study found that netrin-induced axon outgrowth and attraction were significantly reduced in neurons lacking FAK, tying PTK2 function to a classic neural guidance pathway rather than merely correlating it with neural tissue expression. ([pubmed.ncbi.nlm.nih.gov](https://pubmed.ncbi.nlm.nih.gov/15494733/)) [S4]

PTK2/FAK also has evidence in neuronal migration within the developing cortex. In *Journal of Neuroscience* (2011), FAK was reported to be cell-autonomously required for normal interaction of pyramidal neurons with radial glial fibers; loss of FAK disrupted migratory morphology, delayed radial migration, and increased tangential dispersion. That is a direct demonstration of a regulatory role in neural circuit assembly. ([pubmed.ncbi.nlm.nih.gov](https://pubmed.ncbi.nlm.nih.gov/21832197/)) [S5]

Beyond development, PTK2/FAK has evidence in synaptic regulation and systems-level neural function. In mouse hippocampal neurons, gain- and loss-of-function experiments showed FAK to be a critical regulator of hippocampal cell morphology; FAK inhibition affected miniature excitatory postsynaptic potentials and activity-dependent long-term potentiation, and the authors concluded that FAK mediates hippocampus-dependent spatial learning and memory in vivo. This places PTK2 squarely in neural regulatory processes relevant to synaptic plasticity and circuit function. ([pubmed.ncbi.nlm.nih.gov](https://pubmed.ncbi.nlm.nih.gov/21952616/)) [S6]

PTK2/FAK is also implicated in glial biology and myelination. In oligodendrocyte-specific inducible knockout mice, FAK loss just before active optic-nerve myelination reduced the number of myelinated fibers and decreased primary oligodendrocyte processes at postnatal day 14, leading the authors to conclude that FAK controls the efficiency and timing of early CNS myelination. That is another neural regulatory role, this time in a glial process essential for CNS function. ([pubmed.ncbi.nlm.nih.gov](https://pubmed.ncbi.nlm.nih.gov/19224576/)) [S7]

A review in *Current Opinion in Neurobiology* synthesizes these lines of evidence and describes FAK as a key intracellular coordinator of signaling pathways during neuronal motility and development, spanning events from neural induction to axon-dendrite arborization and synapse formation. While reviews are secondary evidence, this one is consistent with the primary literature above and supports the broader interpretation that PTK2 is embedded in neural regulatory programs. ([pubmed.ncbi.nlm.nih.gov](https://pubmed.ncbi.nlm.nih.gov/24705242/)) [S8]

## Contradicting or limiting evidence

The main limitation is not lack of evidence for neural regulation, but rather the **type** of evidence. Most strong causal studies I found are in cultured neurons, embryonic tissue, chick or mouse models, and conditional knockout systems. I did not find comparably strong direct human experimental evidence showing PTK2 manipulation alters neural homeostasis in vivo. So the claim is well supported biologically, but translational certainty to human physiology is somewhat less direct. ([pubmed.ncbi.nlm.nih.gov](https://pubmed.ncbi.nlm.nih.gov/15494733/)) [S9]

Also, the term **homeostatic** is somewhat broader than the evidence base here. The literature for PTK2/FAK is strongest for developmental and signaling regulation—migration, growth-cone behavior, synaptic plasticity, and myelination timing—rather than classic neural homeostasis endpoints such as ion buffering, neurotransmitter clearance, or network set-point stabilization. So if the claim had said PTK2 is specifically a major regulator of neural homeostasis, I would grade it more cautiously. ([pubmed.ncbi.nlm.nih.gov](https://pubmed.ncbi.nlm.nih.gov/15494733/)) [S10]

Another important qualifier is gene-family confusion. PTK2 encodes FAK, whereas PTK2B encodes Pyk2. PTK2B/Pyk2 has especially rich literature in forebrain neurons, synaptic plasticity, Alzheimer disease genetics, and activity-dependent signaling. That does **not** negate PTK2’s neural role, but it means some neuroscience discussions in the literature are really about PTK2B rather than PTK2 and should not be conflated. ([pubmed.ncbi.nlm.nih.gov](https://pubmed.ncbi.nlm.nih.gov/34690733/)) [S11]

Finally, PTK2’s neural effects are context-dependent rather than universal. For example, the 2011 migration study found FAK dispensable for glial-independent cortical interneuron migration but required for radial glia-guided pyramidal-neuron migration. That nuance argues for a regulated, pathway-specific role rather than a blanket requirement for every neural migratory process. ([pubmed.ncbi.nlm.nih.gov](https://pubmed.ncbi.nlm.nih.gov/21832197/)) [S12]

## Mechanistic / biological rationale

Mechanistically, PTK2/FAK is a non-receptor tyrosine kinase and scaffold that integrates signals from integrins, extracellular matrix contacts, growth factor receptors, GPCRs, and netrin receptors. Upon activation, it forms complexes with SRC-family kinases and engages PI3K-AKT, MAPK/ERK, and Rho-family GTPase signaling, all of which are central to cytoskeletal remodeling and adhesion dynamics. In neural cells, those signaling properties make it biologically plausible that PTK2 regulates growth-cone behavior, neurite extension, neuronal migration, axon branching, synapse formation, and postsynaptic actin remodeling. ([proteinatlas.org](https://www.proteinatlas.org/Q05397)) [S13]

The netrin/DCC studies give a concrete neural example of that mechanism: extracellular guidance cues activate DCC-associated FAK, which then helps translate an external cue into directional growth-cone movement and axon extension. The cortical-migration and myelination studies extend the same logic to neuron–glia and glia–matrix interactions, where FAK appears to regulate adhesive contacts and cytoskeletal organization needed for orderly neural development and maturation. ([pubmed.ncbi.nlm.nih.gov](https://pubmed.ncbi.nlm.nih.gov/15494733/)) [S14]

## Bottom line

Yes. PTK2/FAK is convincingly associated with **regulatory** biological processes in neural systems, including axon guidance, neuronal migration, synaptic regulation/plasticity, and CNS myelination. Evidence for a specifically **homeostatic** role is less direct, but the claim as written—“regulatory or homeostatic”—is well supported overall. ([proteinatlas.org](https://www.proteinatlas.org/Q05397)) [S15]

## Cited sources

- **[S1]** PTK2 protein expression summary - The Human Protein Atlas — https://www.proteinatlas.org/Q05397
- **[S2]** PTK2 protein expression summary - The Human Protein Atlas — https://www.proteinatlas.org/Q05397
- **[S3]** PTK2 protein expression summary - The Human Protein Atlas — https://www.proteinatlas.org/Q05397
- **[S4]** Focal adhesion kinase in netrin-1 signaling - PubMed — https://pubmed.ncbi.nlm.nih.gov/15494733/
- **[S5]** Focal adhesion kinase modulates radial glia-dependent neuronal migration through connexin-26 - PubMed — https://pubmed.ncbi.nlm.nih.gov/21832197/
- **[S6]** Focal adhesion kinase regulates neuronal growth, synaptic plasticity and hippocampus-dependent spatial learning and memory - PubMed — https://pubmed.ncbi.nlm.nih.gov/21952616/
- **[S7]** Focal adhesion kinase (FAK): A regulator of CNS myelination - PubMed — https://pubmed.ncbi.nlm.nih.gov/19224576/
- **[S8]** Focal adhesion kinase function in neuronal development - PubMed — https://pubmed.ncbi.nlm.nih.gov/24705242/
- **[S9]** Focal adhesion kinase in netrin-1 signaling - PubMed — https://pubmed.ncbi.nlm.nih.gov/15494733/
- **[S10]** Focal adhesion kinase in netrin-1 signaling - PubMed — https://pubmed.ncbi.nlm.nih.gov/15494733/
- **[S11]** The Non-receptor Tyrosine Kinase Pyk2 in Brain Function and Neurological and Psychiatric Diseases - PubMed — https://pubmed.ncbi.nlm.nih.gov/34690733/
- **[S12]** Focal adhesion kinase modulates radial glia-dependent neuronal migration through connexin-26 - PubMed — https://pubmed.ncbi.nlm.nih.gov/21832197/
- **[S13]** PTK2 protein expression summary - The Human Protein Atlas — https://www.proteinatlas.org/Q05397
- **[S14]** Focal adhesion kinase in netrin-1 signaling - PubMed — https://pubmed.ncbi.nlm.nih.gov/15494733/
- **[S15]** PTK2 protein expression summary - The Human Protein Atlas — https://www.proteinatlas.org/Q05397
