## Supplementary material for "Understanding Substance Dependence: What Differentiates Addictive from Non-Addictive Drugs?": Folder for supplementary files: non-addictive-genes-claims-first100-044-dgkz-gene-protein-has-been-associated-with-regulat-a15973748e22.md

# non-addictive-genes-claims-first100-044-a15973748e22

**Original claim:** DGKZ gene/protein has been associated with regulatory or homeostatic biological processes in neural systems.

## Claim

**DGKZ gene/protein has been associated with regulatory or homeostatic biological processes in neural systems.**

## Verdict

Verdict: Likely True

## Confidence

**High.**

## Supporting evidence

DGKZ encodes diacylglycerol kinase-ζ (DGKζ), and the protein is clearly relevant to neural tissue: Human Protein Atlas classifies **DGKZ RNA as tissue-enhanced in brain**, places it in a **“Brain - Neuronal signaling”** expression cluster, and reports protein expression as **most abundant in brain** among surveyed tissues. ([proteinatlas.org](https://www.proteinatlas.org/ENSG00000149091-DGKZ/tissue)) [S1]

The strongest direct evidence comes from synaptic biology studies in neurons and mice. In rat brain and cultured hippocampal neurons, DGKζ is enriched in synaptic fractions and the postsynaptic density, localizes mainly to dendrites and dendritic spines, and is targeted to excitatory synapses through interaction with the postsynaptic scaffold **PSD-95**. ([pmc.ncbi.nlm.nih.gov](https://pmc.ncbi.nlm.nih.gov/articles/PMC2683696/)) [S2]

Functionally, DGKζ behaves like a regulator of synaptic maintenance rather than a passive marker. In cultured hippocampal neurons, **DGKζ overexpression increased dendritic spine density**, while **DGKζ knockdown reduced spine density**; importantly, these effects required **DGKζ catalytic activity** and **PSD-95 binding**. In vivo, **Dgkz-knockout mice** showed reduced hippocampal spine density and reduced excitatory synaptic transmission, including lower mEPSC frequency and weaker evoked AMPA-mediated responses. ([pmc.ncbi.nlm.nih.gov](https://pmc.ncbi.nlm.nih.gov/articles/PMC2683696/)) [S3]

That same study addressed whether DGKζ acts in a maintenance/homeostasis-like role versus initial formation. Time-lapse imaging showed DGKζ knockdown mainly increased **spine disappearance**, with much less effect on spine appearance, leading the authors to conclude that **DGKζ is required for spine maintenance but not formation**. That is directly consistent with a role in maintaining stable synaptic architecture and excitatory signaling. ([pmc.ncbi.nlm.nih.gov](https://pmc.ncbi.nlm.nih.gov/articles/PMC2683696/)) [S4]

DGKζ has also been linked to regulation of neuronal morphogenesis. In cortical neurons and N1E-115 neuronal cells, DGKζ was found in cell bodies, neurites, and growth cones. Overexpression promoted neurite formation, while dominant-negative constructs impaired outgrowth; mechanistically, DGKζ formed a regulated complex with **syntrophin** and **Rac1** that the authors concluded **controls polarized outgrowth in neuronal cells**. This supports a broader regulatory role in neural development and structural organization. ([pubmed.ncbi.nlm.nih.gov](https://pubmed.ncbi.nlm.nih.gov/16055737/)) [S5]

There is also evidence connecting DGKζ to neuronal stress homeostasis. In hippocampal neurons, excitotoxic kainate stimulation caused DGKζ to shuttle from the nucleus to the cytoplasm, implicating it in early neuronal stress responses. A follow-up study found that cytoplasmic DGKζ facilitates **p53 anchoring/degradation**, and that **DGKζ-deficient brain has elevated p53** after kainate-induced seizures and even under basal conditions; the authors explicitly proposed that DGKζ helps control p53 function during **normal homeostasis and stress responses**. ([pubmed.ncbi.nlm.nih.gov](https://pubmed.ncbi.nlm.nih.gov/21362459/)) [S6]

A neuroscience review synthesizing these data likewise concluded that DGKζ is abundant in cerebellum/hippocampus/olfactory bulb and is involved in **spine maintenance**, with its enzymatic activity and postsynaptic localization being critical. ([pmc.ncbi.nlm.nih.gov](https://pmc.ncbi.nlm.nih.gov/articles/PMC4005014/)) [S7]

## Contradicting or limiting evidence

The main limitation is not that the claim is contradicted, but that the evidence base is **mostly preclinical**. The best functional data come from **rodent neurons, mouse knockouts, hippocampal slices, and neuronal cell models**, not from direct human in vivo experiments. ([pmc.ncbi.nlm.nih.gov](https://pmc.ncbi.nlm.nih.gov/articles/PMC2683696/)) [S8]

Also, the word **“homeostatic”** needs careful interpretation. There is good evidence that DGKζ supports **maintenance of dendritic spines**, **stability of excitatory transmission**, and **neuronal stress-response control**, but this is not the same as proving DGKZ is a central mediator of all forms of **classical homeostatic plasticity** such as synaptic scaling or firing-rate homeostasis. The strongest papers support a **maintenance/regulatory** role; the “homeostatic” label is therefore reasonable but somewhat broader than the direct experiments themselves. ([pmc.ncbi.nlm.nih.gov](https://pmc.ncbi.nlm.nih.gov/articles/PMC2683696/)) [S9]

There are also unresolved mechanistic details. The synaptic study noted that although the phenotype looked mainly postsynaptic, a **presynaptic contribution could not be fully excluded** in knockout animals. Reviews likewise state that some detailed brain-specific mechanisms remain unclear. ([pmc.ncbi.nlm.nih.gov](https://pmc.ncbi.nlm.nih.gov/articles/PMC2683696/)) [S10]

Human association data are indirect. For example, DGKZ has been discussed as a **candidate locus in schizophrenia**, and one study reported altered blood DGKZ expression in patients, but such findings do **not** establish a direct neural regulatory/homeostatic function. They only suggest potential clinical relevance. ([pubmed.ncbi.nlm.nih.gov](https://pubmed.ncbi.nlm.nih.gov/31087244/)) [S11]

## Mechanistic / biological rationale

DGKζ is a lipid-signaling enzyme that **phosphorylates diacylglycerol (DAG) to phosphatidic acid (PA)**. Because DAG activates effectors such as PKC/Ras-pathway components and PA has its own signaling roles, DGKζ acts as a biochemical switch that can locally terminate one signal and generate another. ([pubmed.ncbi.nlm.nih.gov](https://pubmed.ncbi.nlm.nih.gov/11257115/?utm_source=openai)) [S12]

That mechanism fits the neural data well. At excitatory synapses, PSD-95 recruits DGKζ to postsynaptic sites near receptors that generate DAG signaling, and DGKζ then helps control local DAG-to-PA conversion. The 2009 spine study explicitly proposed that this local lipid control is needed to maintain normal spine density and excitatory synaptic transmission. ([pmc.ncbi.nlm.nih.gov](https://pmc.ncbi.nlm.nih.gov/articles/PMC2683696/)) [S13]

DGKζ also interfaces with cytoskeletal regulators. Its interaction with **Rac1** and syntrophins provides a plausible route by which local lipid signaling could influence actin-dependent neurite extension and spine stability. ([pubmed.ncbi.nlm.nih.gov](https://pubmed.ncbi.nlm.nih.gov/16055737/)) [S14]

Under stress, DGKζ’s nucleocytoplasmic shuttling and effects on **p53** provide a second plausible “homeostatic” mechanism: not synaptic structure alone, but protection of neuronal survival pathways and restraint of inappropriate apoptotic signaling. ([pubmed.ncbi.nlm.nih.gov](https://pubmed.ncbi.nlm.nih.gov/21362459/)) [S15]

## Bottom line

Yes—**DGKZ/DGKζ has been associated with regulatory and maintenance-like processes in neural systems**, especially **postsynaptic spine maintenance, regulation of excitatory synaptic transmission, neurite outgrowth/polarity, and neuronal stress-response control**. The evidence is strongest in **rodent and cellular models**, so the claim is best judged **Likely True** rather than maximally established in humans. ([pmc.ncbi.nlm.nih.gov](https://pmc.ncbi.nlm.nih.gov/articles/PMC2683696/)) [S16]

## Cited sources

- **[S1]** Tissue expression of DGKZ - Summary - The Human Protein Atlas — https://www.proteinatlas.org/ENSG00000149091-DGKZ/tissue
- **[S2]** 
            Synaptic removal of diacylglycerol by DGKζ and PSD-95 regulates dendritic spine maintenance - PMC
         — https://pmc.ncbi.nlm.nih.gov/articles/PMC2683696/
- **[S3]** 
            Synaptic removal of diacylglycerol by DGKζ and PSD-95 regulates dendritic spine maintenance - PMC
         — https://pmc.ncbi.nlm.nih.gov/articles/PMC2683696/
- **[S4]** 
            Synaptic removal of diacylglycerol by DGKζ and PSD-95 regulates dendritic spine maintenance - PMC
         — https://pmc.ncbi.nlm.nih.gov/articles/PMC2683696/
- **[S5]** Regulation of neurite outgrowth in N1E-115 cells through PDZ-mediated recruitment of diacylglycerol kinase zeta - PubMed — https://pubmed.ncbi.nlm.nih.gov/16055737/
- **[S6]** Excitotoxicity by kainate-induced seizure causes diacylglycerol kinase ζ to shuttle from the nucleus to the cytoplasm in hippocampal neurons - PubMed — https://pubmed.ncbi.nlm.nih.gov/21362459/
- **[S7]** 
            Diacylglycerol kinase as a possible therapeutic target for neuronal diseases - PMC
         — https://pmc.ncbi.nlm.nih.gov/articles/PMC4005014/
- **[S8]** 
            Synaptic removal of diacylglycerol by DGKζ and PSD-95 regulates dendritic spine maintenance - PMC
         — https://pmc.ncbi.nlm.nih.gov/articles/PMC2683696/
- **[S9]** 
            Synaptic removal of diacylglycerol by DGKζ and PSD-95 regulates dendritic spine maintenance - PMC
         — https://pmc.ncbi.nlm.nih.gov/articles/PMC2683696/
- **[S10]** 
            Synaptic removal of diacylglycerol by DGKζ and PSD-95 regulates dendritic spine maintenance - PMC
         — https://pmc.ncbi.nlm.nih.gov/articles/PMC2683696/
- **[S11]** Expression analysis and genotyping of DGKZ: a GWAS-derived risk gene for schizophrenia - PubMed — https://pubmed.ncbi.nlm.nih.gov/31087244/
- **[S12]** Diacylglycerol kinase zeta regulates Ras activation by a novel mechanism - PubMed — https://pubmed.ncbi.nlm.nih.gov/11257115/?utm_source=openai
- **[S13]** 
            Synaptic removal of diacylglycerol by DGKζ and PSD-95 regulates dendritic spine maintenance - PMC
         — https://pmc.ncbi.nlm.nih.gov/articles/PMC2683696/
- **[S14]** Regulation of neurite outgrowth in N1E-115 cells through PDZ-mediated recruitment of diacylglycerol kinase zeta - PubMed — https://pubmed.ncbi.nlm.nih.gov/16055737/
- **[S15]** Excitotoxicity by kainate-induced seizure causes diacylglycerol kinase ζ to shuttle from the nucleus to the cytoplasm in hippocampal neurons - PubMed — https://pubmed.ncbi.nlm.nih.gov/21362459/
- **[S16]** 
            Synaptic removal of diacylglycerol by DGKζ and PSD-95 regulates dendritic spine maintenance - PMC
         — https://pmc.ncbi.nlm.nih.gov/articles/PMC2683696/
