## Supplementary material for "Understanding Substance Dependence: What Differentiates Addictive from Non-Addictive Drugs?": Folder for supplementary files: non-addictive-genes-claims-first100-045-rasd2-gene-protein-has-been-associated-with-regula-5fd896786c50.md

# non-addictive-genes-claims-first100-045-5fd896786c50

**Original claim:** RASD2 gene/protein has been associated with regulatory or homeostatic biological processes in neural systems.

## Claim

RASD2 gene/protein has been associated with regulatory or homeostatic biological processes in neural systems.

## Verdict

Verdict: Likely True

RASD2 encodes **Rhes** (Ras homolog enriched in striatum), a small GTPase with brain-enriched, especially striatal/putaminal, expression. Across multiple studies, Rhes has been linked to regulation of **dopaminergic signaling**, **cAMP/PKA signaling**, **D1/D2/A2A receptor-related responses**, **corticostriatal synaptic plasticity**, **mTOR signaling**, **autophagy/mitophagy**, and even **striatal iron homeostasis**. That is enough to support an association with regulatory processes in neural systems, and there is some direct support for homeostasis-related roles as well. However, much of the strongest evidence is from rodents and cell systems rather than direct human physiology, so the claim is better rated “Likely True” than “Highly Supported.” ([proteinatlas.org](https://www.proteinatlas.org/ENSG00000100302-RASD2/brain)) [S1]

## Confidence

Medium

Confidence is **medium** because the overall association is well supported, but the literature is concentrated in **striatal biology**, often in **mouse/rat or cell-based models**, and the precise normal physiological role of Rhes remains partly unresolved. Human evidence is strongest for expression pattern and genotype/phenotype association, not for direct causal functional studies in living human neural tissue. ([pmc.ncbi.nlm.nih.gov](https://pmc.ncbi.nlm.nih.gov/articles/PMC4707838/)) [S2]

## Supporting evidence

- **RASD2/Rhes is enriched in neural tissue, especially striatum/putamen.** The Human Protein Atlas classifies RASD2 as **tissue enriched in brain** and places it in a neuronal RNA-expression cluster. Human postmortem work also found **high RASD2 mRNA expression in putamen**, with additional expression in dorsolateral prefrontal cortex. ([proteinatlas.org](https://www.proteinatlas.org/ENSG00000100302-RASD2/brain)) [S3]

- **Loss-of-function studies support a role in striatal neural regulation.** In a knockout mouse study, Rhes disruption produced **behavioral abnormalities**, including increased anxiety and impaired motor coordination, supporting involvement in selected striatal functions rather than being a passive marker gene. ([pubmed.ncbi.nlm.nih.gov](https://pubmed.ncbi.nlm.nih.gov/15199135/)) [S4]

- **Rhes modulates dopamine-linked intracellular signaling.** A cell-based study showed that Rhes **reduced dopamine D1 receptor agonist-stimulated cAMP accumulation**, consistent with a role in GPCR signal regulation. In striatal neurons, another study reported that absence of Rhes altered **cAMP/PKA signaling** in both striatopallidal and striatonigral neurons, affecting dopaminergic behavioral responses. ([pubmed.ncbi.nlm.nih.gov](https://pubmed.ncbi.nlm.nih.gov/21374700/)) [S5]

- **Rhes regulates D2-receptor-related physiology in striatal cholinergic interneurons.** In rodents, Rhes was detected in striatal cholinergic interneurons, and Rhes knockout converted an expected **D2 receptor-mediated inhibitory response** into an abnormal excitatory response, indicating a bona fide role in regulating dopaminergic control of cholinergic excitability. ([pubmed.ncbi.nlm.nih.gov](https://pubmed.ncbi.nlm.nih.gov/25818655/)) [S6]

- **Rhes influences corticostriatal synaptic plasticity.** A detailed mouse study found that Rhes alters **striatal cAMP/PKA signaling** and **corticostriatal LTP** in a sex-dependent manner; in female knockouts, excessive A2AR/cAMP/PKA activity impaired LTP in A2AR/D2R-positive medium spiny neurons, and PKA inhibition restored plasticity. The same paper reported that hippocampal CA1 LTP was unaffected, suggesting a more selective striatal role. ([arpi.unipi.it](https://arpi.unipi.it/retrieve/e0d6c928-9183-fcf8-e053-d805fe0aa794/Ghiglieri_etal_Sci%20Reports_2015.pdf)) [S7]

- **RASD2 is connected to broader signaling cascades with clear regulatory significance.** A human/mouse translational study summarized evidence that Rasd2 affects neuronal processes tied to **cAMP/PKA** and **AKT** signaling and influences **D1R- and D2R-related processes** in striatal projection neurons and cholinergic interneurons. ([pmc.ncbi.nlm.nih.gov](https://pmc.ncbi.nlm.nih.gov/articles/PMC4707838/)) [S8]

- **Rhes is linked to homeostasis-relevant pathways beyond neurotransmission.** NCBI’s gene summary states that RASD2 functions as an **E3 ligase for SUMO attachment** and as an activator of **mTORC1**, a pathway relevant to axon growth, regeneration, and myelination. Experimentally, Rhes was shown to **bind and activate mTOR** in striatum. ([ncbi.nlm.nih.gov](https://www.ncbi.nlm.nih.gov/gene/23551)) [S9]

- **Rhes regulates autophagy and mitophagy, which are classic cellular homeostatic processes.** One study found that Rhes binds **Beclin-1** and activates **autophagy**; another found that Rhes is a **critical regulator of mitophagy via Nix**, affecting mitochondrial quality control and striatal vulnerability. These are direct links to neural homeostasis at the organelle/protein-turnover level. ([pubmed.ncbi.nlm.nih.gov](https://pubmed.ncbi.nlm.nih.gov/24324270/)) [S10]

- **There is also direct evidence tying Rhes to iron homeostasis in the striatum.** A neuroscience study reported that Rhes participates in iron uptake via DMT1-related machinery and concluded that Rhes may play a crucial role in **striatal iron homeostasis**. ([pubmed.ncbi.nlm.nih.gov](https://pubmed.ncbi.nlm.nih.gov/23999124/)) [S11]

- **RASD2 also shows human relevance beyond simple expression.** In humans, a noncoding RASD2 variant affecting mRNA levels was associated with **prefrontal gray matter volume** and **prefrontal/striatal activity during working memory**, which is consistent with a regulatory role in neural circuitry, though still indirect. ([pmc.ncbi.nlm.nih.gov](https://pmc.ncbi.nlm.nih.gov/articles/PMC4707838/)) [S12]

## Contradicting or limiting evidence

- **The normal physiological role is still not fully settled.** Even authors studying Rhes directly note that its role in striatal physiology “remains less understood,” despite growing evidence in pathology and signaling. That limits how confidently one can specify exactly *which* homeostatic functions are primary versus context-dependent. ([arpi.unipi.it](https://arpi.unipi.it/retrieve/e0d6c928-9183-fcf8-e053-d805fe0aa794/Ghiglieri_etal_Sci%20Reports_2015.pdf)) [S13]

- **Much of the strongest functional evidence is non-human.** The key mechanistic studies for dopamine signaling, synaptic plasticity, autophagy, mitophagy, and iron homeostasis are predominantly from **mouse, rat, or cultured-cell systems**, whereas human evidence is more about expression pattern and genetic association. This supports “associated with” but is weaker for strong claims about established human neural homeostatic function. ([pubmed.ncbi.nlm.nih.gov](https://pubmed.ncbi.nlm.nih.gov/25818655/)) [S14]

- **Disease-context results can point in different directions.** In some models, Rhes promotes pathologic processes such as mHTT toxicity or dyskinesia, and Rhes deletion is protective. Yet in a Huntington’s disease study with reduced striatal mTORC1 activity, **exogenous Rhes improved motor and neuropathologic outcomes**, implying that its effects may depend strongly on disease stage, pathway state, and experimental context. ([pubmed.ncbi.nlm.nih.gov](https://pubmed.ncbi.nlm.nih.gov/22179112/)) [S15]

- **The claim is broad, but the evidence is region-weighted.** Most direct evidence centers on **striatal neurons**, especially medium spiny neurons and cholinergic interneurons, rather than neural systems broadly across the whole brain. ([pubmed.ncbi.nlm.nih.gov](https://pubmed.ncbi.nlm.nih.gov/25818655/)) [S16]

## Mechanistic / biological rationale

Rhes is well positioned to serve regulatory and partly homeostatic roles because it is a **small signaling GTPase** enriched in striatal neurons and associated interneurons. Mechanistically, it can modulate **dopamine receptor-linked cAMP signaling**, likely through interactions with heterotrimeric G-protein pathways; this gives a plausible route for controlling neuronal excitability and neuromodulatory balance. ([pubmed.ncbi.nlm.nih.gov](https://pubmed.ncbi.nlm.nih.gov/21374700/)) [S17]

At the circuit level, altered Rhes changes **cAMP/PKA tone** and **A2AR/D2R-dependent corticostriatal plasticity**, showing how one molecule can influence the rules by which striatal synapses strengthen or fail to strengthen. That is a clear example of a regulatory role in neural information processing. ([arpi.unipi.it](https://arpi.unipi.it/retrieve/e0d6c928-9183-fcf8-e053-d805fe0aa794/Ghiglieri_etal_Sci%20Reports_2015.pdf)) [S18]

At the cell-biological level, Rhes also interfaces with **SUMOylation**, **mTORC1**, **autophagy**, **mitophagy**, and **iron trafficking**. Those pathways are central to protein quality control, energy/mitochondrial maintenance, and intracellular resource balance—core elements of cellular homeostasis. So even if the literature often frames Rhes in disease terms, the implicated mechanisms themselves are fundamentally regulatory/homeostatic. ([ncbi.nlm.nih.gov](https://www.ncbi.nlm.nih.gov/gene/23551)) [S19]

## Bottom line

The claim is **supported overall**: RASD2/Rhes has repeatedly been associated with **regulatory processes in neural systems**, especially in the **striatal dopaminergic and corticostriatal signaling/plasticity network**, and there is also meaningful evidence linking it to **homeostasis-related processes** such as **autophagy, mitophagy, and iron homeostasis**. The main caveat is that the strongest evidence comes from **rodent and cellular studies**, with **human evidence more indirect**. So the best evidence-based summary is: **RASD2/Rhes is very plausibly involved in neural regulatory processes and is reasonably associated with certain homeostatic processes, particularly in the striatum.** ([pubmed.ncbi.nlm.nih.gov](https://pubmed.ncbi.nlm.nih.gov/25818655/)) [S20]

## Cited sources

- **[S1]** https://www.proteinatlas.org/ENSG00000100302-RASD2/brain — https://www.proteinatlas.org/ENSG00000100302-RASD2/brain
- **[S2]** https://pmc.ncbi.nlm.nih.gov/articles/PMC4707838/ — https://pmc.ncbi.nlm.nih.gov/articles/PMC4707838/
- **[S3]** https://www.proteinatlas.org/ENSG00000100302-RASD2/brain — https://www.proteinatlas.org/ENSG00000100302-RASD2/brain
- **[S4]** https://pubmed.ncbi.nlm.nih.gov/15199135/ — https://pubmed.ncbi.nlm.nih.gov/15199135/
- **[S5]** https://pubmed.ncbi.nlm.nih.gov/21374700/ — https://pubmed.ncbi.nlm.nih.gov/21374700/
- **[S6]** https://pubmed.ncbi.nlm.nih.gov/25818655/ — https://pubmed.ncbi.nlm.nih.gov/25818655/
- **[S7]** https://arpi.unipi.it/retrieve/e0d6c928-9183-fcf8-e053-d805fe0aa794/Ghiglieri_etal_Sci%20Reports_2015.pdf — https://arpi.unipi.it/retrieve/e0d6c928-9183-fcf8-e053-d805fe0aa794/Ghiglieri_etal_Sci%20Reports_2015.pdf
- **[S8]** https://pmc.ncbi.nlm.nih.gov/articles/PMC4707838/ — https://pmc.ncbi.nlm.nih.gov/articles/PMC4707838/
- **[S9]** https://www.ncbi.nlm.nih.gov/gene/23551 — https://www.ncbi.nlm.nih.gov/gene/23551
- **[S10]** https://pubmed.ncbi.nlm.nih.gov/24324270/ — https://pubmed.ncbi.nlm.nih.gov/24324270/
- **[S11]** https://pubmed.ncbi.nlm.nih.gov/23999124/ — https://pubmed.ncbi.nlm.nih.gov/23999124/
- **[S12]** https://pmc.ncbi.nlm.nih.gov/articles/PMC4707838/ — https://pmc.ncbi.nlm.nih.gov/articles/PMC4707838/
- **[S13]** https://arpi.unipi.it/retrieve/e0d6c928-9183-fcf8-e053-d805fe0aa794/Ghiglieri_etal_Sci%20Reports_2015.pdf — https://arpi.unipi.it/retrieve/e0d6c928-9183-fcf8-e053-d805fe0aa794/Ghiglieri_etal_Sci%20Reports_2015.pdf
- **[S14]** https://pubmed.ncbi.nlm.nih.gov/25818655/ — https://pubmed.ncbi.nlm.nih.gov/25818655/
- **[S15]** https://pubmed.ncbi.nlm.nih.gov/22179112/ — https://pubmed.ncbi.nlm.nih.gov/22179112/
- **[S16]** https://pubmed.ncbi.nlm.nih.gov/25818655/ — https://pubmed.ncbi.nlm.nih.gov/25818655/
- **[S17]** https://pubmed.ncbi.nlm.nih.gov/21374700/ — https://pubmed.ncbi.nlm.nih.gov/21374700/
- **[S18]** https://arpi.unipi.it/retrieve/e0d6c928-9183-fcf8-e053-d805fe0aa794/Ghiglieri_etal_Sci%20Reports_2015.pdf — https://arpi.unipi.it/retrieve/e0d6c928-9183-fcf8-e053-d805fe0aa794/Ghiglieri_etal_Sci%20Reports_2015.pdf
- **[S19]** https://www.ncbi.nlm.nih.gov/gene/23551 — https://www.ncbi.nlm.nih.gov/gene/23551
- **[S20]** https://pubmed.ncbi.nlm.nih.gov/25818655/ — https://pubmed.ncbi.nlm.nih.gov/25818655/
