## Supplementary material for "Understanding Substance Dependence: What Differentiates Addictive from Non-Addictive Drugs?": Folder for supplementary files: non-addictive-genes-claims-first100-046-cdk18-gene-protein-has-been-associated-with-regula-4fe74f1d50b4.md

# non-addictive-genes-claims-first100-046-4fe74f1d50b4

**Original claim:** CDK18 gene/protein has been associated with regulatory or homeostatic biological processes in neural systems.

## Claim

CDK18 gene/protein has been associated with regulatory or homeostatic biological processes in neural systems.

## Verdict

Verdict: Likely True

CDK18 has **credible, though not extensive**, evidence linking it to **regulatory processes in the nervous system**, especially oligodendrocyte differentiation, myelination/remyelination biology, and tau-related neuronal pathology. The stronger part of the claim is **“regulatory processes in neural systems.”** The weaker part is **“homeostatic”** in the narrow neuroscience sense; I did not find strong direct evidence that CDK18 is an established regulator of canonical neural homeostasis such as synaptic scaling, excitability homeostasis, or neurotransmitter/ion homeostasis. ([pubmed.ncbi.nlm.nih.gov](https://pubmed.ncbi.nlm.nih.gov/31028571/)) [S1]

## Confidence

Medium

The confidence is medium because there is **at least one direct mechanistic CNS study** and multiple brain/pathology associations, but the literature is **sparse**, much of it is **non-human or disease-contextual**, and reviews still describe CDK18/PCTAIRE3 as an **understudied kinase** whose core physiological neuronal functions remain incompletely defined. ([pubmed.ncbi.nlm.nih.gov](https://pubmed.ncbi.nlm.nih.gov/31028571/)) [S2]

## Supporting evidence

A foundational point is that CDK18 is **expressed in the nervous system** rather than being a purely non-neural kinase. NCBI’s gene record cites the original transcript study showing that one CDK18/PCTAIRE3 isoform was detectable **throughout the human brain**, while another was more restricted to **subcortical basal ganglia nuclei and spinal cord**. The Human Protein Atlas likewise shows CDK18 RNA across major human brain regions, including cortex, hippocampal formation, basal ganglia, thalamus, cerebellum, and spinal cord. ([ncbi.nlm.nih.gov](https://www.ncbi.nlm.nih.gov/gene?Cmd=DetailsSearch&Db=gene&Term=5129)) [S3]

The strongest direct evidence for a **regulatory neural role** comes from oligodendrocyte biology. In a peer-reviewed study, CDK18 was reported to be **exclusively expressed in oligodendrocytes** among the neural cell types examined, to increase with **myelin formation during development**, and to rise during **remyelination** in a lysophosphatidylcholine demyelination model. Functionally, CDK18 overexpression promoted oligodendrocyte precursor cell (OPC) differentiation, while interference reduced differentiation, without altering proliferation or apoptosis. The same study linked this effect mechanistically to **ERK1/2 pathway activation**. That is direct evidence that CDK18 participates in a **regulatory biological process within the CNS**, namely control of glial differentiation relevant to myelin formation and repair. ([pubmed.ncbi.nlm.nih.gov](https://pubmed.ncbi.nlm.nih.gov/31028571/)) [S4]

This matters for the claim because OPC differentiation, myelination, and remyelination are not just developmental events; they are also part of how the CNS maintains and restores white-matter function after injury. So even if the literature does not explicitly frame CDK18 as a “homeostasis kinase,” it is reasonable to say it has been associated with **regulation of a neural maintenance/repair process**. That is still short of proving a broad homeostatic role, but it supports the claim’s regulatory half well. ([pubmed.ncbi.nlm.nih.gov](https://pubmed.ncbi.nlm.nih.gov/31028571/)) [S5]

There is also evidence linking CDK18 to **neuronal pathological regulation**, particularly tau biology. A human brain study in Alzheimer disease reported increased PCTAIRE3/CDK18 protein in **temporal cortex** and enrichment in **paired helical filaments**, with overexpression experiments suggesting CDK18 indirectly promotes phosphorylation of tau at **T231/S235**, sites altered early in AD pathogenesis. A later APP-dependent study similarly reported increased PCTAIRE-3 in human late-AD samples and interpreted the findings as implicating cell-cycle/transcriptional dysregulation in neurons. These findings do not establish normal homeostasis, but they do place CDK18 within **neural regulatory signaling relevant to disease biology**. ([pubmed.ncbi.nlm.nih.gov](https://pubmed.ncbi.nlm.nih.gov/16766195/)) [S6]

A further, weaker systems-level association comes from addiction-related transcriptomics. In rhesus macaques with chronic ethanol self-administration, **Cdk18** was identified as an intake-correlated hub in the **central amygdala** coexpression network, and the relevant modules were enriched for **synaptic and membrane** annotations. This does not prove that CDK18 itself regulates synapses or addiction circuitry, but it does show that CDK18 has been associated with neural transcriptional networks related to long-term alcohol exposure. ([pubmed.ncbi.nlm.nih.gov](https://pubmed.ncbi.nlm.nih.gov/28247455/)) [S7]

## Contradicting or limiting evidence

The main limitation is that **direct evidence is narrow**. The clearest mechanistic neural study I found is largely centered on **oligodendrocyte lineage cells** and experimental demyelination/remyelination, not broad neural homeostasis across neurons, circuits, or behavior. There is not much direct evidence that CDK18 controls canonical homeostatic processes such as synaptic homeostasis, firing-rate stabilization, sleep homeostasis, or calcium/ion homeostasis. ([pubmed.ncbi.nlm.nih.gov](https://pubmed.ncbi.nlm.nih.gov/31028571/)) [S8]

A second limitation is that some neural evidence is **pathology-associated rather than physiology-established**. The Alzheimer-related papers connect CDK18 to tau phosphorylation and disease-associated protein changes, but these findings are partly based on **overexpression** or disease tissue comparisons, which do not by themselves demonstrate that CDK18 normally performs a homeostatic role in healthy neural systems. ([pubmed.ncbi.nlm.nih.gov](https://pubmed.ncbi.nlm.nih.gov/16766195/)) [S9]

Third, reviews of the PCTAIRE kinase family explicitly emphasize how **understudied** these kinases are in the brain. One review described PCTKs as “forgotten brain kinases” and said their biological functions in the brain remain largely to be discovered; another review on CDK18/PCTK3 stated that the protein has received far less attention than other CDKs and that its functions are still being pieced together. That makes overconfident interpretation inappropriate. ([karger.com](https://karger.com/nsg/article-abstract/17/4/288/379073/PCTK-Proteins-The-Forgotten-Brain-Kinases)) [S10]

Finally, much of the better-established CDK18 literature is actually **non-neural**: genome integrity/replication-stress signaling, membrane cargo trafficking, and aquaporin-2 regulation in kidney cells are recurring themes. Those findings support CDK18 as a regulatory kinase in general, but they do not by themselves demonstrate neural homeostasis. ([pubmed.ncbi.nlm.nih.gov](https://pubmed.ncbi.nlm.nih.gov/33781185/)) [S11]

## Mechanistic / biological rationale

Biologically, the claim is plausible. CDK18 is a **serine/threonine cyclin-dependent kinase** in the PCTAIRE family, and reviews report activation by **cyclin A2** or **PKA-linked phosphorylation**. Kinases of this class often act as signal-integrating regulators rather than structural proteins, so a role in neural regulatory pathways is mechanistically credible. ([pubmed.ncbi.nlm.nih.gov](https://pubmed.ncbi.nlm.nih.gov/33781185/)) [S12]

In the strongest CNS mechanism currently available, CDK18 promotes OPC differentiation by increasing **ERK1/2 signaling**, which in turn elevates oligodendrocyte maturation markers such as MBP. Because oligodendrocyte differentiation is essential for **myelination and remyelination** in the CNS, CDK18 can reasonably be viewed as participating in a neural regulatory axis that affects tissue maintenance and repair. ([pubmed.ncbi.nlm.nih.gov](https://pubmed.ncbi.nlm.nih.gov/31028571/)) [S13]

In neurons, the tau literature suggests a different mechanism: CDK18 may influence the phosphorylation state of **tau**, apparently indirectly in available studies. Since tau phosphorylation state is relevant to neuronal cytoskeletal organization and neurodegenerative pathology, this provides an additional mechanistic route by which CDK18 could affect neural system regulation, though the evidence here is more disease-focused and less definitive for normal physiology. ([pubmed.ncbi.nlm.nih.gov](https://pubmed.ncbi.nlm.nih.gov/16766195/)) [S14]

## Bottom line

The claim is **best judged as Likely True**. CDK18 is **brain-expressed** and has **direct mechanistic evidence** for regulating a CNS process—especially **oligodendrocyte differentiation and myelination/remyelination-related biology**—with additional associations in **tau/Alzheimer-related neural signaling** and addiction-related brain transcriptomic networks. However, the evidence for a broad or canonical **neural homeostatic** role is still **limited and indirect**, so the strongest defensible wording is that CDK18 has been associated with **regulatory neural processes**, with only **partial support** for the “homeostatic” part of the claim. ([ncbi.nlm.nih.gov](https://www.ncbi.nlm.nih.gov/gene?Cmd=DetailsSearch&Db=gene&Term=5129)) [S15]

## Cited sources

- **[S1]** Cyclin-dependent Kinase 18 Promotes Oligodendrocyte Precursor Cell Differentiation through Activating the Extracellular Signal-Regulated Kinase Signaling Pathway - PubMed — https://pubmed.ncbi.nlm.nih.gov/31028571/
- **[S2]** Cyclin-dependent Kinase 18 Promotes Oligodendrocyte Precursor Cell Differentiation through Activating the Extracellular Signal-Regulated Kinase Signaling Pathway - PubMed — https://pubmed.ncbi.nlm.nih.gov/31028571/
- **[S3]** CDK18 cyclin dependent kinase 18 [Homo sapiens (human)] - Gene - NCBI — https://www.ncbi.nlm.nih.gov/gene?Cmd=DetailsSearch&Db=gene&Term=5129
- **[S4]** Cyclin-dependent Kinase 18 Promotes Oligodendrocyte Precursor Cell Differentiation through Activating the Extracellular Signal-Regulated Kinase Signaling Pathway - PubMed — https://pubmed.ncbi.nlm.nih.gov/31028571/
- **[S5]** Cyclin-dependent Kinase 18 Promotes Oligodendrocyte Precursor Cell Differentiation through Activating the Extracellular Signal-Regulated Kinase Signaling Pathway - PubMed — https://pubmed.ncbi.nlm.nih.gov/31028571/
- **[S6]** The regulation of tau phosphorylation by PCTAIRE 3: implications for the pathogenesis of Alzheimer's disease - PubMed — https://pubmed.ncbi.nlm.nih.gov/16766195/
- **[S7]** On the relationships in rhesus macaques between chronic ethanol consumption and the brain transcriptome - PubMed — https://pubmed.ncbi.nlm.nih.gov/28247455/
- **[S8]** Cyclin-dependent Kinase 18 Promotes Oligodendrocyte Precursor Cell Differentiation through Activating the Extracellular Signal-Regulated Kinase Signaling Pathway - PubMed — https://pubmed.ncbi.nlm.nih.gov/31028571/
- **[S9]** The regulation of tau phosphorylation by PCTAIRE 3: implications for the pathogenesis of Alzheimer's disease - PubMed — https://pubmed.ncbi.nlm.nih.gov/16766195/
- **[S10]** PCTK Proteins: The Forgotten Brain Kinases? | Neurosignals | Karger Publishers — https://karger.com/nsg/article-abstract/17/4/288/379073/PCTK-Proteins-The-Forgotten-Brain-Kinases
- **[S11]** Overview of PCTK3/CDK18: A Cyclin-Dependent Kinase Involved in Specific Functions in Post-Mitotic Cells - PubMed — https://pubmed.ncbi.nlm.nih.gov/33781185/
- **[S12]** Overview of PCTK3/CDK18: A Cyclin-Dependent Kinase Involved in Specific Functions in Post-Mitotic Cells - PubMed — https://pubmed.ncbi.nlm.nih.gov/33781185/
- **[S13]** Cyclin-dependent Kinase 18 Promotes Oligodendrocyte Precursor Cell Differentiation through Activating the Extracellular Signal-Regulated Kinase Signaling Pathway - PubMed — https://pubmed.ncbi.nlm.nih.gov/31028571/
- **[S14]** The regulation of tau phosphorylation by PCTAIRE 3: implications for the pathogenesis of Alzheimer's disease - PubMed — https://pubmed.ncbi.nlm.nih.gov/16766195/
- **[S15]** CDK18 cyclin dependent kinase 18 [Homo sapiens (human)] - Gene - NCBI — https://www.ncbi.nlm.nih.gov/gene?Cmd=DetailsSearch&Db=gene&Term=5129
