## Supplementary material for "Understanding Substance Dependence: What Differentiates Addictive from Non-Addictive Drugs?": Folder for supplementary files: non-addictive-genes-claims-first100-047-rasd1-gene-protein-has-been-associated-with-regula-ba2a07f778fb.md

# non-addictive-genes-claims-first100-047-ba2a07f778fb

**Original claim:** RASD1 gene/protein has been associated with regulatory or homeostatic biological processes in neural systems.

## Claim

RASD1 gene/protein has been associated with regulatory or homeostatic biological processes in neural systems.

## Verdict

Verdict: Highly Supported

Across the experimental neuroscience literature, **RASD1 (Dexras1)** is repeatedly linked to neural regulatory/homeostatic functions, especially in **circadian entrainment in the suprachiasmatic nucleus (SCN), neuronal iron handling, feedback control of NMDA receptor excitability, hypothalamic osmotic/stress-responsive transcription, and exercise-responsive hippocampal neurogenesis**. The main caveat is that this evidence is **mostly from rodent and cell-based studies rather than direct human clinical studies**. ([ncbi.nlm.nih.gov](https://www.ncbi.nlm.nih.gov/gene/51655)) [S1]

## Confidence

High

## Supporting evidence

RASD1 encodes **Dexras1**, a Ras-family small GTPase. NCBI describes it as an **activator of G-protein signaling** that can act as a **direct nucleotide exchange factor for Gi/Go proteins**, and notes interaction with the neuronal nitric oxide adaptor **CAPON/NOS1AP**. Early work in *Neuron* identified Dexras1 as a **brain-enriched** G protein that forms a ternary complex with **nNOS and CAPON**, and showed that Dexras1 is activated by **NMDA receptor-stimulated NO synthesis in cortical neurons**, placing it directly in neuronal signaling pathways. ([ncbi.nlm.nih.gov](https://www.ncbi.nlm.nih.gov/gene/51655)) [S2]

One of the strongest lines of evidence is **neuronal iron homeostasis**. In a primary mechanistic study, Cheah et al. reported that **NMDA receptor–NO transmission mediates neuronal iron homeostasis via Dexras1**, establishing a pathway from NMDA receptor activation to Dexras1-dependent regulation of iron uptake, and explicitly stating that glutamatergic neurotransmission has a physiologic role in regulating brain iron uptake. ([pmc.ncbi.nlm.nih.gov](https://pmc.ncbi.nlm.nih.gov/articles/PMC3150500/)) [S3]

That homeostatic theme was extended by later work showing that **lysosomal iron modulates NMDA receptor-mediated excitation via Dexras1**. In that study, Dexras1-dependent iron signaling fed back onto NMDA receptor function through PKC/Src signaling; the authors concluded that this pathway “may exert **homeostatic control of excitability**.” This is especially relevant to your wording because it directly connects RASD1/Dexras1 to a neural **feedback-regulatory** process. ([pubmed.ncbi.nlm.nih.gov](https://pubmed.ncbi.nlm.nih.gov/27080392/)) [S4]

Additional mechanistic support comes from a 2015 study showing that **PKA phosphorylation of Dexras1 suppresses iron trafficking by inhibiting S-nitrosylation**. That work further indicates that Dexras1 is not merely associated with iron flux, but is embedded in a **regulatory switch** in neurons in which phosphorylation counterbalances NO-dependent activation and thereby modulates iron influx and ROS-related signaling. ([pmc.ncbi.nlm.nih.gov](https://pmc.ncbi.nlm.nih.gov/articles/PMC4767167/?utm_source=openai)) [S5]

RASD1/Dexras1 is also well supported as a regulator of **circadian neural homeostasis/entrainment** in the SCN. In the original circadian knockout work, loss of Dexras1 altered photic resetting, enhanced nonphotic responses, changed free-running period, and reduced rhythm integrity; conversely, SCN-specific expression of constitutively active Dexras1 altered light responses. The authors concluded that Dexras1 has a **fundamental role in shaping phase-dependent responsiveness of the mammalian circadian clock**. ([pubmed.ncbi.nlm.nih.gov](https://pubmed.ncbi.nlm.nih.gov/17167088/)) [S6]

Follow-up studies reinforced that conclusion. Loss of Dexras1 caused **enhanced and phase-specific nonphotic phase shifts** and implicated Dexras1 in regulating multiple photic and nonphotic signal-transduction pathways involved in circadian entrainment. Another study found that under restricted feeding, Dexras1-deficient mice showed altered SCN PER1/PER2/pERK rhythms, indicating that Dexras1 helps buffer or regulate SCN responses to metabolic timing cues. In rats, Dexras1 expression itself was shown to be rhythmic in the SCN. ([pubmed.ncbi.nlm.nih.gov](https://pubmed.ncbi.nlm.nih.gov/21834641/)) [S7]

There is also direct evidence for a **homeostatic/stress-regulatory role in hypothalamic neurons**. Greenwood et al. showed that Rasd1 is expressed in **vasopressin neurons** of the PVN and SON, is induced by **hyperosmotic cues** and glucocorticoids, and modulates cAMP-PKA-CREB-dependent transcriptional responses. Their conclusion was that RASD1 abundance in vasopressin neurons is an important mechanism for **controlling transcriptional responses to stressors** in these hypothalamic nuclei. Because PVN/SON signaling is central to neuroendocrine control of water balance and stress, this is strong evidence for neural regulatory/homeostatic association. ([pubmed.ncbi.nlm.nih.gov](https://pubmed.ncbi.nlm.nih.gov/26739966/)) [S8]

A further example comes from the adult hippocampus. A 2018 study explicitly titled **“Dexras1 is a homeostatic regulator of exercise-dependent proliferation and cell survival in the hippocampal neurogenic niche”** showed that Dexras1 deletion suppresses exercise-induced progenitor proliferation, alters survival of mitotic and post-mitotic cells, and abolishes exercise-dependent ERK/MAPK-CREB activation and induction of pro-neurogenic genes such as **bdnf, trkB, and vegf-a**. This is direct evidence that the protein has been associated with **homeostatic regulation in a neural stem/progenitor system**. ([pubmed.ncbi.nlm.nih.gov](https://pubmed.ncbi.nlm.nih.gov/29593295/)) [S9]

## Contradicting or limiting evidence

The main limitation is **species/context**. The most persuasive data are from **mouse and rat models, cultured neurons, and ex vivo brain preparations**; I did not find comparable depth of direct evidence from human neural physiology or clinical intervention studies in the retrieved literature. So the claim is strongly supported for **experimental neural systems**, but less directly for human in vivo neurobiology. ([pubmed.ncbi.nlm.nih.gov](https://pubmed.ncbi.nlm.nih.gov/11086993/)) [S10]

Also, not every neural phenotype is broad or global. In one behavioral knockout study, Dexras1-null mice did **not** show altered baseline anxiety-like or depression-like behavior, and the behavioral effects were selective rather than uniformly disruptive. That does not argue against a regulatory role, but it does suggest that RASD1/Dexras1 is **not a universal controller of all neural homeostasis** and may act in pathway- and context-specific ways. ([pubmed.ncbi.nlm.nih.gov](https://pubmed.ncbi.nlm.nih.gov/26946266/)) [S11]

Finally, part of the Dexras1 literature emphasizes **pathophysiology**—for example NMDA excitotoxicity, neurodegeneration, optic neuritis, or injury-related iron dysregulation. Those studies support biological relevance, but some extrapolation is needed when moving from disease/injury paradigms to normal baseline homeostasis. ([pmc.ncbi.nlm.nih.gov](https://pmc.ncbi.nlm.nih.gov/articles/PMC3150500/)) [S12]

## Mechanistic / biological rationale

A coherent mechanism emerges across studies. In neurons, **NMDA receptor activation** stimulates **nNOS**, which via **CAPON/NOS1AP** couples NO signaling to **Dexras1**. Activated Dexras1 then interfaces with iron-transport machinery, including **DMT1-associated complexes**, to regulate neuronal iron influx. Because iron availability in turn feeds back onto NMDA receptor signaling and excitability, Dexras1 sits in a plausible **homeostatic feedback loop** linking synaptic activity, NO signaling, iron handling, oxidative stress risk, and receptor function. ([pubmed.ncbi.nlm.nih.gov](https://pubmed.ncbi.nlm.nih.gov/11086993/)) [S13]

In the SCN and hypothalamus, Dexras1 also modulates **cAMP/PKA/CREB** and **ERK/MAPK** pathways, both of which are canonical signal-integration systems for entrainment and stimulus-responsive gene expression. That gives a biologically plausible basis for its roles in **circadian phase regulation**, **osmotic/stress-responsive transcription**, and **activity-dependent neurogenesis**. ([pubmed.ncbi.nlm.nih.gov](https://pubmed.ncbi.nlm.nih.gov/17167088/)) [S14]

## Bottom line

Yes. The claim is **well supported** by preclinical neuroscience research. RASD1/Dexras1 has been associated with multiple **regulatory and homeostatic processes in neural systems**, including **neuronal iron homeostasis, feedback control of excitability, circadian entrainment, hypothalamic stress/osmotic response regulation, and hippocampal neurogenic niche homeostasis**. The key qualification is that this support comes **primarily from rodent and cellular studies**, not robust human clinical evidence. ([pmc.ncbi.nlm.nih.gov](https://pmc.ncbi.nlm.nih.gov/articles/PMC3150500/)) [S15]

## Cited sources

- **[S1]** RASD1 ras related dexamethasone induced 1 [Homo sapiens (human)] - Gene - NCBI — https://www.ncbi.nlm.nih.gov/gene/51655
- **[S2]** RASD1 ras related dexamethasone induced 1 [Homo sapiens (human)] - Gene - NCBI — https://www.ncbi.nlm.nih.gov/gene/51655
- **[S3]** 
            NMDA Receptor-Nitric Oxide Transmission Mediates Neuronal Iron Homeostasis via the GTPase Dexras1 - PMC
         — https://pmc.ncbi.nlm.nih.gov/articles/PMC3150500/
- **[S4]** Lysosomal iron modulates NMDA receptor-mediated excitation via small GTPase, Dexras1 - PubMed — https://pubmed.ncbi.nlm.nih.gov/27080392/
- **[S5]** PKA-mediated phosphorylation of Dexras1 suppresses iron trafficking by inhibiting S-nitrosylation - PMC — https://pmc.ncbi.nlm.nih.gov/articles/PMC4767167/?utm_source=openai
- **[S6]** The molecular gatekeeper Dexras1 sculpts the photic responsiveness of the mammalian circadian clock - PubMed — https://pubmed.ncbi.nlm.nih.gov/17167088/
- **[S7]** Loss of dexras1 alters nonphotic circadian phase shifts and reveals a role for the intergeniculate leaflet (IGL) in gene-targeted mice - PubMed — https://pubmed.ncbi.nlm.nih.gov/21834641/
- **[S8]** Rasd1, a small G protein with a big role in the hypothalamic response to neuronal activation - PubMed — https://pubmed.ncbi.nlm.nih.gov/26739966/
- **[S9]** Dexras1 is a homeostatic regulator of exercise-dependent proliferation and cell survival in the hippocampal neurogenic niche - PubMed — https://pubmed.ncbi.nlm.nih.gov/29593295/
- **[S10]** Dexras1: a G protein specifically coupled to neuronal nitric oxide synthase via CAPON - PubMed — https://pubmed.ncbi.nlm.nih.gov/11086993/
- **[S11]** Dexras1 a unique ras-GTPase interacts with NMDA receptor activity and provides a novel dissociation between anxiety, working memory and sensory gating - PubMed — https://pubmed.ncbi.nlm.nih.gov/26946266/
- **[S12]** 
            NMDA Receptor-Nitric Oxide Transmission Mediates Neuronal Iron Homeostasis via the GTPase Dexras1 - PMC
         — https://pmc.ncbi.nlm.nih.gov/articles/PMC3150500/
- **[S13]** Dexras1: a G protein specifically coupled to neuronal nitric oxide synthase via CAPON - PubMed — https://pubmed.ncbi.nlm.nih.gov/11086993/
- **[S14]** The molecular gatekeeper Dexras1 sculpts the photic responsiveness of the mammalian circadian clock - PubMed — https://pubmed.ncbi.nlm.nih.gov/17167088/
- **[S15]** 
            NMDA Receptor-Nitric Oxide Transmission Mediates Neuronal Iron Homeostasis via the GTPase Dexras1 - PMC
         — https://pmc.ncbi.nlm.nih.gov/articles/PMC3150500/
