## Supplementary material for "Understanding Substance Dependence: What Differentiates Addictive from Non-Addictive Drugs?": Folder for supplementary files: non-addictive-genes-claims-first100-048-rapgef5-gene-protein-has-been-associated-with-regu-07769dfb3c95.md

# non-addictive-genes-claims-first100-048-07769dfb3c95

**Original claim:** RAPGEF5 gene/protein has been associated with regulatory or homeostatic biological processes in neural systems.

## Claim

RAPGEF5 gene/protein has been associated with regulatory or homeostatic biological processes in neural systems.

## Verdict

Verdict: Likely True

## Confidence

Medium

## Supporting evidence

RAPGEF5 is clearly linked to the nervous system at the expression and annotation level. NCBI lists **broad human RAPGEF5 expression in brain**, and the Mouse Genome Informatics (MGI) resource shows **Rapgef5 expression in the mouse nervous system during development**. MGI also assigns Rapgef5 mutant-associated phenotype categories that include **nervous system** and **behavior/neurological**, and broad GO categories that include **system development**, **homeostatic process**, and **synapse**. These database-level associations do not prove mechanism on their own, but they support that RAPGEF5 is not a purely non-neural gene. ([ncbi.nlm.nih.gov](https://www.ncbi.nlm.nih.gov/gene/9771)) [S1]

More directly, a developmental neurobiology study reported that **mr-gef/Rapgef5 expression is regulated during differentiation of specific subsets of telencephalic neurons**. The paper’s summary states that the spatiotemporal expression profile in rodent ventral telencephalon is consistent with a role in the development of **young neurons, particularly GABAergic neurons**. That is direct evidence of association with a neural regulatory process, namely neuronal differentiation. ([sciencedirect.com](https://www.sciencedirect.com/science/article/abs/pii/S0165380603003018?utm_source=openai)) [S2]

A follow-up study strengthened that link. In mouse forebrain, **mr-gef was turned on in developing GABAergic neurons**, and its expression was **markedly reduced in Dlx1/2 double-mutant mice**, leading the authors to conclude that mr-gef and Dlx1/2 are part of a common pathway during **GABAergic neuronal development**. In adult/postmortem human cortex, the same study found altered layer-to-layer relationships in MR-GEF-expressing neurons in bipolar disorder and suggested that altered MR-GEF expression could influence **neurotransmission**. This is still associative, but it places RAPGEF5 within a neural developmental/signaling context rather than merely showing passive brain expression. ([pmc.ncbi.nlm.nih.gov](https://pmc.ncbi.nlm.nih.gov/articles/PMC2861006/)) [S3]

A later mouse/human embryogenesis paper also described RAPGEF5 as previously associated with **neural development** and reported expression in **discrete regions of the developing brain** and the broader **nervous system** in mouse and human embryos. The authors explicitly interpreted this as likely reflecting a **conserved developmental role**. ([sciencedirect.com](https://www.sciencedirect.com/science/article/abs/pii/S1567133X18301765)) [S4]

Mechanistically, RAPGEF5 has been shown to regulate **nuclear translocation of β-catenin** and thereby affect **Wnt target gene expression** in Xenopus embryos. Loss of Rapgef5 reduced nuclear β-catenin and reduced expression of the Wnt-responsive gene **foxj1**. While that experiment was not framed as a brain study, canonical **Wnt/β-catenin signaling is a major regulatory pathway in neural development**, and reviews describe it as important for **neural stem-cell proliferation/differentiation balance** and **neural stem-cell homeostasis**. So RAPGEF5 sits in a signaling axis with well-established neural regulatory relevance. ([pubmed.ncbi.nlm.nih.gov](https://pubmed.ncbi.nlm.nih.gov/29290587/)) [S5]

## Contradicting or limiting evidence

The main limitation is that **direct evidence for RAPGEF5 in neural homeostasis per se is sparse**. Most of the stronger data concern **developmental expression**, **neuronal differentiation**, or **embryonic Wnt signaling**, not classic adult neural homeostasis such as synaptic scaling, firing-rate homeostasis, ion/glial homeostasis, or maintenance of mature neural circuits. So the claim is stronger if “regulatory” is emphasized, and weaker if “homeostatic” is interpreted narrowly. ([sciencedirect.com](https://www.sciencedirect.com/science/article/abs/pii/S0165380603003018?utm_source=openai)) [S6]

Human evidence is also limited in strength. The bipolar-disorder paper is based on **postmortem association/correlation**, not causal intervention, and the embryogenesis paper mainly maps expression rather than proving neural function in mammals. Even the Wnt/β-catenin study demonstrating a RAPGEF5 mechanism was performed in an **embryonic left-right patterning** context rather than directly in neural tissue. ([pmc.ncbi.nlm.nih.gov](https://pmc.ncbi.nlm.nih.gov/articles/PMC2861006/)) [S7]

Database annotations add supportive context but should not be overread. MGI’s nervous-system phenotype/expression categories and broad GO groupings indicate that Rapgef5 is connected to neural biology, but such annotations are not equivalent to a dedicated mechanistic study showing RAPGEF5 maintains neural homeostasis. ([informatics.jax.org](https://informatics.jax.org/marker/MGI%3A2444365)) [S8]

## Mechanistic / biological rationale

RAPGEF5 is a **guanine nucleotide exchange factor (GEF)** for Rap-family small GTPases, including **RAP1A/RAP2A** and MRAS-related signaling. GEFs promote the active GTP-bound state of small GTPases, and Rap signaling is broadly involved in **cell adhesion, migration, polarity, and intracellular signaling**. Those are biologically plausible upstream controls for neuronal differentiation, migration, and circuit assembly. ([proteinatlas.org](https://www.proteinatlas.org/ENSG00000136237-RAPGEF5)) [S9]

RAPGEF5 appears unusual among RAPGEFs in that it has been reported to localize **specifically to the nucleus** in frog, mouse, and human cells. In the Xenopus study, RAPGEF5 was required for **β-catenin nuclear localization**, placing it at a key control point in canonical Wnt signaling. Because Wnt/β-catenin signaling helps govern **early vertebrate neural development** and the balance of **neural stem-cell proliferation versus differentiation**, RAPGEF5 has a credible mechanistic route into neural regulatory biology even if direct mammalian neural functional experiments remain limited. ([sciencedirect.com](https://www.sciencedirect.com/science/article/abs/pii/S1567133X18301765)) [S10]

The forebrain data fit that rationale. In developing telencephalon, mr-gef/Rapgef5 expression tracks with markers and transcriptional programs relevant to **GABAergic lineage differentiation and migration**, and Dlx1/2 loss substantially reduces expression. That pattern is consistent with RAPGEF5 acting in a **regulatory pathway** rather than being a generic housekeeping protein. The suggestion that altered cortical MR-GEF expression could affect **neurotransmission** is plausible, but should still be considered provisional. ([pmc.ncbi.nlm.nih.gov](https://pmc.ncbi.nlm.nih.gov/articles/PMC2861006/)) [S11]

## Bottom line

The claim is **likely true**, mainly because RAPGEF5 has been associated with **neural regulatory processes**: it is expressed in developing brain/nervous system, has been linked to **telencephalic and GABAergic neuronal differentiation**, and regulates a **Wnt/β-catenin pathway** that is well known to control neural development and neural stem-cell homeostasis. ([sciencedirect.com](https://www.sciencedirect.com/science/article/abs/pii/S1567133X18301765)) [S12]

However, the evidence is **much stronger for neural regulation/development than for direct neural homeostasis** in the strict adult physiological sense. So if the claim is interpreted broadly as “RAPGEF5 has been associated with regulatory biology in neural systems,” it is supported; if interpreted narrowly as “RAPGEF5 is an established neural homeostasis gene,” the evidence is not yet strong enough for that stronger statement. ([pmc.ncbi.nlm.nih.gov](https://pmc.ncbi.nlm.nih.gov/articles/PMC2861006/)) [S13]

## Cited sources

- **[S1]** RAPGEF5 Rap guanine nucleotide exchange factor 5 [Homo sapiens (human)] - Gene - NCBI — https://www.ncbi.nlm.nih.gov/gene/9771
- **[S2]** Expression of the guanine nucleotide exchange factor, mr-gef, is regulated during the differentiation of specific subsets of telencephalic neurons - ScienceDirect — https://www.sciencedirect.com/science/article/abs/pii/S0165380603003018?utm_source=openai
- **[S3]** 
            Expression of the Rap1 Guanine Nucleotide Exchange Factor, MR-GEF, Is Altered in Individuals with Bipolar Disorder - PMC
         — https://pmc.ncbi.nlm.nih.gov/articles/PMC2861006/
- **[S4]** Expression of the guanine nucleotide exchange factor, RAPGEF5, during mouse and human embryogenesis - ScienceDirect — https://www.sciencedirect.com/science/article/abs/pii/S1567133X18301765
- **[S5]** RAPGEF5 Regulates Nuclear Translocation of β-Catenin - PubMed — https://pubmed.ncbi.nlm.nih.gov/29290587/
- **[S6]** Expression of the guanine nucleotide exchange factor, mr-gef, is regulated during the differentiation of specific subsets of telencephalic neurons - ScienceDirect — https://www.sciencedirect.com/science/article/abs/pii/S0165380603003018?utm_source=openai
- **[S7]** 
            Expression of the Rap1 Guanine Nucleotide Exchange Factor, MR-GEF, Is Altered in Individuals with Bipolar Disorder - PMC
         — https://pmc.ncbi.nlm.nih.gov/articles/PMC2861006/
- **[S8]** Rapgef5 MGI Mouse Gene Detail - MGI:2444365 - Rap guanine nucleotide exchange factor (GEF) 5 — https://informatics.jax.org/marker/MGI%3A2444365
- **[S9]** RAPGEF5 protein expression summary - The Human Protein Atlas — https://www.proteinatlas.org/ENSG00000136237-RAPGEF5
- **[S10]** Expression of the guanine nucleotide exchange factor, RAPGEF5, during mouse and human embryogenesis - ScienceDirect — https://www.sciencedirect.com/science/article/abs/pii/S1567133X18301765
- **[S11]** 
            Expression of the Rap1 Guanine Nucleotide Exchange Factor, MR-GEF, Is Altered in Individuals with Bipolar Disorder - PMC
         — https://pmc.ncbi.nlm.nih.gov/articles/PMC2861006/
- **[S12]** Expression of the guanine nucleotide exchange factor, RAPGEF5, during mouse and human embryogenesis - ScienceDirect — https://www.sciencedirect.com/science/article/abs/pii/S1567133X18301765
- **[S13]** 
            Expression of the Rap1 Guanine Nucleotide Exchange Factor, MR-GEF, Is Altered in Individuals with Bipolar Disorder - PMC
         — https://pmc.ncbi.nlm.nih.gov/articles/PMC2861006/
