## Supplementary material for "Understanding Substance Dependence: What Differentiates Addictive from Non-Addictive Drugs?": Folder for supplementary files: non-addictive-genes-claims-first100-049-dync1li2-gene-protein-has-been-associated-with-reg-d7f3c418d382.md

# non-addictive-genes-claims-first100-049-d7f3c418d382

**Original claim:** DYNC1LI2 gene/protein has been associated with regulatory or homeostatic biological processes in neural systems.

## Claim

DYNC1LI2 gene/protein has been associated with regulatory or homeostatic biological processes in neural systems.

## Verdict

Verdict: Likely True

## Confidence

Medium

## Supporting evidence

DYNC1LI2 encodes cytoplasmic dynein 1 light intermediate chain 2 (LIC2), a subunit of the dynein-1 motor complex. Authoritative gene resources describe it as a protein-coding dynein component with broad brain expression; NCBI lists ubiquitous human brain expression, and the Human Protein Atlas shows substantial brain RNA signal. This does not prove function by itself, but it establishes that the gene is present in neural tissue at levels compatible with biologic relevance. ([ncbi.nlm.nih.gov](https://www.ncbi.nlm.nih.gov/gene/1783?utm_source=openai)) [S1]

The strongest direct evidence comes from an in vivo neocortical development study in rat. Gonçalves et al. found that LIC2/DYNC1LI2 has a specific role in the final phase of neuronal migration, terminal somal translocation (TST). In that study, LIC2 knockdown or expression of a dominant LIC2 fragment caused neurons to stall below the upper cortical plate, with elongated leading processes that still contacted the marginal zone while the soma failed to advance. The authors concluded that LIC2 contributes a distinct regulatory role during brain development, different from LIC1. That is a clear association with a regulatory process in a neural system. ([pmc.ncbi.nlm.nih.gov](https://pmc.ncbi.nlm.nih.gov/articles/PMC6400572/)) [S2]

A second line of evidence links DYNC1LI2 to neuronal polarity. In cultured hippocampal neurons, Yang et al. reported that phosphorylation of the polarity protein Par3 by Ndr kinases blocks Par3 interaction with dynein light intermediate chain 2 (Dlic2/DYNC1LI2), thereby helping localize Par3 asymmetrically and reinforce axon specification. Even though the study is framed around the Rassf5-Ndr-Par3 pathway, it directly implicates DYNC1LI2-containing dynein in the regulated transport events that establish neuronal polarity. ([pmc.ncbi.nlm.nih.gov](https://pmc.ncbi.nlm.nih.gov/articles/PMC4132390/?utm_source=openai)) [S3]

There is also newer but lower-certainty evidence from a 2025 bioRxiv preprint, which is explicitly marked in PubMed as not yet peer reviewed. That work reported that endfoot-localized Dync1li2 regulates radial glial basal morphology and affects interneuron organization during cortical development. Because it is a preprint, it should be treated as supportive rather than definitive, but it points in the same direction as the peer-reviewed neural-development studies above. ([pubmed.ncbi.nlm.nih.gov](https://pubmed.ncbi.nlm.nih.gov/41280023/)) [S4]

Mechanistically, DYNC1LI2 is not just a passive structural subunit. Recent work describes LIC1 and LIC2 as adaptor-binding subunits that define distinct dynein complexes with overlapping but non-identical functions, and the human dynein interactome work supports the broader concept that cargo-specific dynein regulation depends on such subunit/adaptor organization. That kind of molecular specialization makes the neural findings biologically coherent. ([pmc.ncbi.nlm.nih.gov](https://pmc.ncbi.nlm.nih.gov/articles/PMC11033342/?utm_source=openai)) [S5]

## Contradicting or limiting evidence

The main limitation is that the evidence is much stronger for **neural regulation/development** than for **neural homeostasis** specifically. I found direct neural evidence for neuronal migration, polarity, and cortical organization, but not strong peer-reviewed evidence showing that DYNC1LI2 is a major established regulator of synaptic homeostasis, firing-rate homeostasis, neurotransmitter homeostasis, or other classic neural homeostatic programs. ([pmc.ncbi.nlm.nih.gov](https://pmc.ncbi.nlm.nih.gov/articles/PMC6400572/)) [S6]

A separate study does show that DYNC1LI2 can promote cellular homeostasis by rescuing chaperone-mediated autophagy, reducing ER stress, correcting mitochondrial fragmentation, and improving cell survival in cystinotic fibroblasts and kidney proximal tubule cells. However, those are non-neural systems. So the homeostasis part of the claim is supported best at the general cellular level, not specifically within neurons or brain circuits. ([pubmed.ncbi.nlm.nih.gov](https://pubmed.ncbi.nlm.nih.gov/34643468/)) [S7]

The direct neural literature on DYNC1LI2 is also relatively sparse. Much of the dynein-neurology field centers on other dynein components such as DYNC1H1, BICD2, LIS1/PAFAH1B1, or the distinct intermediate-chain gene DYNC1I2. That matters because some neurodevelopmental disease papers that look visually similar in name are actually about **DYNC1I2**, not **DYNC1LI2**. The NCBI records confirm these are different genes. ([ncbi.nlm.nih.gov](https://www.ncbi.nlm.nih.gov/gene/1783?utm_source=openai)) [S8]

## Mechanistic / biological rationale

DYNC1LI2/LIC2 is part of cytoplasmic dynein-1, the major minus-end-directed microtubule motor used for long-range intracellular transport. LIC subunits help govern adaptor binding and thereby influence which cargos are transported and how dynein complexes are assembled functionally. Because neural cells are highly polarized and depend heavily on regulated trafficking of polarity proteins, RNAs, organelles, and signaling endosomes, a dynein subunit with cargo/adaptor specificity is well positioned to affect neural regulatory processes. ([pmc.ncbi.nlm.nih.gov](https://pmc.ncbi.nlm.nih.gov/articles/PMC11033342/?utm_source=openai)) [S9]

The neural studies fit that logic. In one context, DYNC1LI2-containing dynein regulates the final somal movement of migrating cortical neurons. In another, interaction with DYNC1LI2-containing dynein is part of the mechanism controlling Par3 distribution during axon specification. These are not generic housekeeping readouts; they are spatially and developmentally regulated neural processes. ([pmc.ncbi.nlm.nih.gov](https://pmc.ncbi.nlm.nih.gov/articles/PMC6400572/)) [S10]

A plausible but still inferential extension is that DYNC1LI2 may also contribute to neural homeostasis through trafficking-dependent proteostasis and organelle quality control. The cystinosis paper shows DYNC1LI2 can influence LAMP2A localization, chaperone-mediated autophagy, ER stress, mitochondrial morphology, and survival in non-neural cells. Since neurons are unusually dependent on autophagy, mitochondrial quality control, and long-distance retrograde transport, it is biologically plausible that analogous LIC2-dependent mechanisms help maintain neuronal homeostasis. But that specific neural-homeostasis role remains more inferred than directly demonstrated. ([pubmed.ncbi.nlm.nih.gov](https://pubmed.ncbi.nlm.nih.gov/34643468/)) [S11]

## Bottom line

DYNC1LI2 is **fairly well supported as being associated with regulatory processes in neural systems**, especially neuronal polarity, cortical development, and a specific phase of neuronal migration. The case that it is involved in **neural homeostasis** is more limited and mostly indirect, extrapolating from its dynein trafficking role and from non-neural cellular-homeostasis studies. So the overall claim is best judged **Likely True**: solid for neural regulation, suggestive but not yet strong for neural homeostasis specifically. ([pmc.ncbi.nlm.nih.gov](https://pmc.ncbi.nlm.nih.gov/articles/PMC6400572/)) [S12]

## Cited sources

- **[S1]** DYNC1LI2 dynein cytoplasmic 1 light intermediate chain 2 [Homo sapiens (human)] - Gene - NCBI — https://www.ncbi.nlm.nih.gov/gene/1783?utm_source=openai
- **[S2]** 
            Distinct roles for dynein light intermediate chains in neurogenesis, migration, and terminal somal translocation - PMC
         — https://pmc.ncbi.nlm.nih.gov/articles/PMC6400572/
- **[S3]** Rassf5 and Ndr kinases regulate neuronal polarity through Par3 phosphorylation in a novel pathway - PMC — https://pmc.ncbi.nlm.nih.gov/articles/PMC4132390/?utm_source=openai
- **[S4]** Subcellular transcriptome of radial glia reveals compartmentalized control of cortical development - PubMed — https://pubmed.ncbi.nlm.nih.gov/41280023/
- **[S5]** Dynein Light Intermediate Chains Exhibit Different Arginine Methylation Patterns - PMC — https://pmc.ncbi.nlm.nih.gov/articles/PMC11033342/?utm_source=openai
- **[S6]** 
            Distinct roles for dynein light intermediate chains in neurogenesis, migration, and terminal somal translocation - PMC
         — https://pmc.ncbi.nlm.nih.gov/articles/PMC6400572/
- **[S7]** DYNC1LI2 regulates localization of the chaperone-mediated autophagy receptor LAMP2A and improves cellular homeostasis in cystinosis - PubMed — https://pubmed.ncbi.nlm.nih.gov/34643468/
- **[S8]** DYNC1LI2 dynein cytoplasmic 1 light intermediate chain 2 [Homo sapiens (human)] - Gene - NCBI — https://www.ncbi.nlm.nih.gov/gene/1783?utm_source=openai
- **[S9]** Dynein Light Intermediate Chains Exhibit Different Arginine Methylation Patterns - PMC — https://pmc.ncbi.nlm.nih.gov/articles/PMC11033342/?utm_source=openai
- **[S10]** 
            Distinct roles for dynein light intermediate chains in neurogenesis, migration, and terminal somal translocation - PMC
         — https://pmc.ncbi.nlm.nih.gov/articles/PMC6400572/
- **[S11]** DYNC1LI2 regulates localization of the chaperone-mediated autophagy receptor LAMP2A and improves cellular homeostasis in cystinosis - PubMed — https://pubmed.ncbi.nlm.nih.gov/34643468/
- **[S12]** 
            Distinct roles for dynein light intermediate chains in neurogenesis, migration, and terminal somal translocation - PMC
         — https://pmc.ncbi.nlm.nih.gov/articles/PMC6400572/
