## Supplementary material for "Understanding Substance Dependence: What Differentiates Addictive from Non-Addictive Drugs?": Folder for supplementary files: non-addictive-genes-claims-first100-050-ngef-gene-protein-has-been-associated-with-regulat-b188de16eb32.md

# non-addictive-genes-claims-first100-050-b188de16eb32

**Original claim:** NGEF gene/protein has been associated with regulatory or homeostatic biological processes in neural systems.

## Claim

NGEF gene/protein has been associated with regulatory or homeostatic biological processes in neural systems.

## Verdict

Verdict: Highly Supported

NGEF, also called **ephexin-1/ARHGEF27**, is strongly linked to **regulatory neural processes**, especially **axon guidance, growth cone dynamics, dendritic spine remodeling, synapse organization, and neuromuscular junction maturation**. Official gene annotations also place it in ephrin receptor signaling and synapse-related processes, with brain-enriched expression and localization/activity in neural compartments such as growth cones and glutamatergic synapses. ([ncbi.nlm.nih.gov](https://www.ncbi.nlm.nih.gov/gene/25791)) [S1]

The **“homeostatic”** part of the claim is supported more narrowly: there is direct evidence for an **Ephexin-dependent presynaptic homeostatic signaling system in Drosophila NMJ**, but much less direct evidence that mammalian or human NGEF is a major neural homeostasis factor in the adult CNS. So the claim is best supported on the **regulatory** side, with more limited but real support on the **homeostatic** side. ([pubmed.ncbi.nlm.nih.gov](https://pubmed.ncbi.nlm.nih.gov/19249276/?utm_source=openai)) [S2]

## Confidence

High

Confidence is high because multiple independent experimental studies, including loss-of-function and signaling studies in neurons and animal models, converge on NGEF/Ephexin1 as a regulator of neural structure and signaling. Confidence is not absolute only because the specific term **homeostatic** is less directly established in mammalian systems than the broader term **regulatory**. ([pubmed.ncbi.nlm.nih.gov](https://pubmed.ncbi.nlm.nih.gov/11336673/)) [S3]

## Supporting evidence

NGEF is a protein-coding gene whose product is commonly called **ephexin-1**. NCBI lists it as **brain-biased** in expression and summarizes predicted involvement in the **ephrin receptor signaling pathway**, **regulation of actin cytoskeleton organization**, and **regulation of synapse organization**; it is also annotated as active in **glutamatergic synapse** and located in **growth cone**. The original characterization paper further reported that Ngef is predominantly expressed in brain, with strongest signal in the **caudate nucleus**. ([ncbi.nlm.nih.gov](https://www.ncbi.nlm.nih.gov/gene/25791)) [S4]

A foundational Cell paper identified ephexin as an **Eph receptor–interacting Rho-family guanine nucleotide exchange factor** and showed that ephrin-A stimulation modulates ephexin activity, shifting downstream GTPase signaling and altering cell morphology. In primary neurons, a mutant ephexin interfered with **ephrin-A–induced growth cone collapse**, directly tying the protein to a neural regulatory process governing axon navigation. ([pubmed.ncbi.nlm.nih.gov](https://pubmed.ncbi.nlm.nih.gov/11336673/)) [S5]

A follow-up Neuron study strengthened this by using **ephexin1-null mouse neurons** and chick RNA interference. It reported that ephexin1 is required for **normal axon outgrowth** and **ephrin-dependent axon repulsion**, and that tyrosine phosphorylation of ephexin1 is required for **growth cone collapse**. That is direct experimental evidence that NGEF regulates neural wiring behavior through signal-dependent cytoskeletal control. ([pubmed.ncbi.nlm.nih.gov](https://pubmed.ncbi.nlm.nih.gov/15848799/)) [S6]

Evidence also extends beyond early axon behavior into synaptic structure. In a Nature Neuroscience study, ephrin-A1/EphA4 signaling recruited and activated **Cdk5**, which then promoted ephexin1-dependent **RhoA** activation, causing **dendritic spine retraction** and reduced **mEPSC frequency** at hippocampal synapses. This places NGEF squarely in a regulatory pathway affecting **synaptic morphology and excitatory synaptic function**, both core neural regulatory phenomena. ([pubmed.ncbi.nlm.nih.gov](https://pubmed.ncbi.nlm.nih.gov/17143272/)) [S7]

NGEF/Ephexin1 also regulates peripheral synapse maturation. In mice, ephexin1 deficiency caused abnormal **acetylcholine receptor cluster maturation**, simplified NMJ morphology, impaired **neuromuscular transmission**, and muscle weakness; re-expression rescued structural defects. The authors concluded that ephexin1 is essential for **structural maturation and neurotransmission** at the neuromuscular junction. ([pmc.ncbi.nlm.nih.gov](https://pmc.ncbi.nlm.nih.gov/articles/PMC3718047/)) [S8]

More recent in vivo developmental work showed that ephexin1 is expressed in chick and mouse **lateral motor column neurons** during spinal motor axon extension, and that gain- and loss-of-function perturbations disrupt **motor axon trajectory selection**. This independently supports a conserved role in **Eph-mediated motor axon guidance**, another neural regulatory process. ([pubmed.ncbi.nlm.nih.gov](https://pubmed.ncbi.nlm.nih.gov/29363583/)) [S9]

For the homeostatic side specifically, a Drosophila study identified **ephexin** mutations that disrupt the **homeostatic enhancement of presynaptic release** after postsynaptic glutamate receptor impairment at the fly NMJ. The paper described a presynaptic homeostatic signaling system involving **Eph receptor, ephexin, Cdc42, and CaV2.1 calcium channels**. This is bona fide evidence for an Ephexin-family role in neural homeostatic regulation, though in an invertebrate model and at the NMJ rather than the mammalian CNS. ([pubmed.ncbi.nlm.nih.gov](https://pubmed.ncbi.nlm.nih.gov/19249276/?utm_source=openai)) [S10]

## Contradicting or limiting evidence

The main limitation is not that evidence is absent, but that it is **uneven across subclaims**. Direct evidence for NGEF/Ephexin1 in **regulatory neural processes** is strong, whereas direct evidence for **neural homeostasis** in mammalian or human systems is much thinner. Reviews mention synaptic homeostasis, but the clearest primary homeostatic experiment is in **Drosophila ephexin**, not human NGEF or mammalian CNS neurons. ([pmc.ncbi.nlm.nih.gov](https://pmc.ncbi.nlm.nih.gov/articles/PMC6406967/)) [S11]

A second limitation is that some official process annotations for human NGEF, such as **regulation of synapse pruning** or **negative regulation of dendritic spine morphogenesis**, are supported by mixed evidence codes including **IEA/ISS/IBA/TAS**, meaning some are inferred computationally, by homology, or from curated statements rather than direct human experimentation. Those annotations are useful and directionally consistent, but they are not all equivalent to direct mechanistic proof in humans. ([ncbi.nlm.nih.gov](https://www.ncbi.nlm.nih.gov/gene/25791)) [S12]

A third limitation is biological scope: much of the strongest evidence concerns **developmental wiring and synaptic remodeling** rather than broad, organism-level neural homeostasis. So if the intended claim were “NGEF is a central canonical homeostasis gene in adult mammalian neural systems,” the evidence would be weaker than for the broader statement given here. ([pubmed.ncbi.nlm.nih.gov](https://pubmed.ncbi.nlm.nih.gov/15848799/)) [S13]

## Mechanistic / biological rationale

Mechanistically, NGEF/Ephexin1 is well positioned to regulate neural processes because it is a **Rho-family GEF** that links extracellular receptor cues to intracellular cytoskeletal remodeling. It interacts with **EphA4/Eph receptors**, and upon ephrin stimulation its activity shifts toward **RhoA** relative to **Rac1/Cdc42**, which changes actin organization and thereby controls **growth cone motility**, **axon repulsion**, and cell shape. ([pubmed.ncbi.nlm.nih.gov](https://pubmed.ncbi.nlm.nih.gov/11336673/)) [S14]

At synapses, EphA4 recruits **Cdk5**, which phosphorylates ephexin1 and promotes further activation of RhoA signaling, leading to **dendritic spine retraction** and reduced excitatory synaptic activity. At the NMJ, ephexin1-dependent RhoA signaling helps stabilize and remodel **AChR clusters** and organize postsynaptic structure needed for efficient transmission. In Drosophila, Ephexin participates in a pathway that adjusts **presynaptic neurotransmitter release** when postsynaptic receptor function falls, providing a plausible mechanistic basis for homeostatic control. ([pubmed.ncbi.nlm.nih.gov](https://pubmed.ncbi.nlm.nih.gov/17143272/)) [S15]

Taken together, these mechanisms are exactly what one expects from a protein involved in **regulatory neural biology**: it transduces receptor activation into spatially restricted control of small GTPases, actin dynamics, synaptic structure, and signal output. ([pubmed.ncbi.nlm.nih.gov](https://pubmed.ncbi.nlm.nih.gov/11336673/)) [S16]

## Bottom line

Yes. **NGEF/Ephexin1 is clearly associated with regulatory biological processes in neural systems**, with strong evidence for roles in **axon guidance, growth cone collapse, dendritic spine remodeling, synapse organization, and neuromuscular junction maturation/function**. Evidence for a specifically **homeostatic** role exists, but is most direct in **Drosophila synaptic homeostasis** and is less established in mammalian/human neural systems. Overall, the claim is best judged **highly supported**, mainly because the regulatory-neural association is well demonstrated. ([ncbi.nlm.nih.gov](https://www.ncbi.nlm.nih.gov/gene/25791)) [S17]

## Cited sources

- **[S1]** NGEF neuronal guanine nucleotide exchange factor [Homo sapiens (human)] - Gene - NCBI — https://www.ncbi.nlm.nih.gov/gene/25791
- **[S2]** A presynaptic homeostatic signaling system composed of the Eph receptor, ephexin, Cdc42, and CaV2.1 calcium channels - PubMed — https://pubmed.ncbi.nlm.nih.gov/19249276/?utm_source=openai
- **[S3]** EphA receptors regulate growth cone dynamics through the novel guanine nucleotide exchange factor ephexin - PubMed — https://pubmed.ncbi.nlm.nih.gov/11336673/
- **[S4]** NGEF neuronal guanine nucleotide exchange factor [Homo sapiens (human)] - Gene - NCBI — https://www.ncbi.nlm.nih.gov/gene/25791
- **[S5]** EphA receptors regulate growth cone dynamics through the novel guanine nucleotide exchange factor ephexin - PubMed — https://pubmed.ncbi.nlm.nih.gov/11336673/
- **[S6]** Eph-dependent tyrosine phosphorylation of ephexin1 modulates growth cone collapse - PubMed — https://pubmed.ncbi.nlm.nih.gov/15848799/
- **[S7]** Cdk5 regulates EphA4-mediated dendritic spine retraction through an ephexin1-dependent mechanism - PubMed — https://pubmed.ncbi.nlm.nih.gov/17143272/
- **[S8]** 
            Ephexin1 is required for structural maturation and neurotransmission at the neuromuscular junction - PMC
         — https://pmc.ncbi.nlm.nih.gov/articles/PMC3718047/
- **[S9]** Ephexin1 Is Required for Eph-Mediated Limb Trajectory of Spinal Motor Axons - PubMed — https://pubmed.ncbi.nlm.nih.gov/29363583/
- **[S10]** A presynaptic homeostatic signaling system composed of the Eph receptor, ephexin, Cdc42, and CaV2.1 calcium channels - PubMed — https://pubmed.ncbi.nlm.nih.gov/19249276/?utm_source=openai
- **[S11]** 
            Emerging Roles of Ephexins in Physiology and Disease - PMC
         — https://pmc.ncbi.nlm.nih.gov/articles/PMC6406967/
- **[S12]** NGEF neuronal guanine nucleotide exchange factor [Homo sapiens (human)] - Gene - NCBI — https://www.ncbi.nlm.nih.gov/gene/25791
- **[S13]** Eph-dependent tyrosine phosphorylation of ephexin1 modulates growth cone collapse - PubMed — https://pubmed.ncbi.nlm.nih.gov/15848799/
- **[S14]** EphA receptors regulate growth cone dynamics through the novel guanine nucleotide exchange factor ephexin - PubMed — https://pubmed.ncbi.nlm.nih.gov/11336673/
- **[S15]** Cdk5 regulates EphA4-mediated dendritic spine retraction through an ephexin1-dependent mechanism - PubMed — https://pubmed.ncbi.nlm.nih.gov/17143272/
- **[S16]** EphA receptors regulate growth cone dynamics through the novel guanine nucleotide exchange factor ephexin - PubMed — https://pubmed.ncbi.nlm.nih.gov/11336673/
- **[S17]** NGEF neuronal guanine nucleotide exchange factor [Homo sapiens (human)] - Gene - NCBI — https://www.ncbi.nlm.nih.gov/gene/25791
