## Supplementary material for "Understanding Substance Dependence: What Differentiates Addictive from Non-Addictive Drugs?": Folder for supplementary files: non-addictive-genes-claims-first100-051-pfdn5-gene-protein-has-been-associated-with-regula-e6aeb3246207.md

# non-addictive-genes-claims-first100-051-e6aeb3246207

**Original claim:** PFDN5 gene/protein has been associated with regulatory or homeostatic biological processes in neural systems.

## Claim

PFDN5 gene/protein has been associated with regulatory or homeostatic biological processes in neural systems.

## Verdict

Verdict: Likely True

PFDN5 has credible experimental associations with neural **proteostasis/cytoskeletal homeostasis**, **neuronal development**, and **synaptic plasticity-related regulation**, especially in mouse, Drosophila, and neuronal cell models. However, the evidence is not yet strong enough to treat PFDN5 as a firmly established, human neural homeostasis gene on the same level as canonical synaptic or glial regulators. ([ncbi.nlm.nih.gov](https://www.ncbi.nlm.nih.gov/gene/5204?utm_source=openai)) [S1]

## Confidence

Medium

Confidence is **medium** because multiple studies support the claim, but the literature is still fairly limited and is dominated by non-human and mechanistic models rather than direct human genetic or clinical evidence. ([pubmed.ncbi.nlm.nih.gov](https://pubmed.ncbi.nlm.nih.gov/31505240/?utm_source=openai)) [S2]

## Supporting evidence

PFDN5 encodes prefoldin subunit 5, a component of the prefoldin co-chaperone complex that helps stabilize nascent polypeptides and promotes correct folding, especially of cytoskeletal substrates such as actin and tubulin. NCBI and review sources describe prefoldin as a core protein-folding/proteostasis factor, which is biologically relevant to neurons because they are highly dependent on long-lived cytoskeletal architecture and protein quality control. ([ncbi.nlm.nih.gov](https://www.ncbi.nlm.nih.gov/gene/5204?utm_source=openai)) [S3]

PFDN5 is also clearly present in the nervous system. The Human Protein Atlas classifies human PFDN5 as having **low tissue specificity** overall but shows that it is **detected in all brain regions** with **low regional specificity**; in single-cell data it is “cell type enhanced” in several brain-associated cell types, including excitatory neurons, inhibitory neurons, oligodendrocytes, astrocytes, microglia, ependymal cells, Bergmann glia, and oligodendrocyte progenitors. That pattern does not prove function, but it supports plausibility for neural roles. ([proteinatlas.org](https://www.proteinatlas.org/ENSG00000123349-PFDN5/brain)) [S4]

A direct association with a neural regulatory process comes from a mouse hippocampus study examining synaptic plasticity. In that work, PFDN5 was predominantly localized to hippocampal CA1–CA3 pyramidal layers; nicotine-induced LTP-like facilitation increased PFDN5 mRNA and protein, while a tau-transgenic Alzheimer model with impaired plasticity showed age-dependent decreases in hippocampal PFDN5. The authors concluded that PFDN5 levels change with synaptic plasticity and proposed it as an important component of that process. This is not definitive proof of causal regulation, but it is strong “association with neural regulation” evidence. ([pubmed.ncbi.nlm.nih.gov](https://pubmed.ncbi.nlm.nih.gov/31505240/)) [S5]

There is also stronger causal evidence in development. In a murine missense-mutant model, disruption of **Pfdn5** caused photoreceptor degeneration, central nervous system abnormalities, and abnormal sensory/neuronal development. The authors linked these phenotypes to reduced formation of microtubules and microfilaments, structures essential for cilia and neuronal architecture. This supports a role for PFDN5 in maintaining neural structural homeostasis during development. ([pubmed.ncbi.nlm.nih.gov](https://pubmed.ncbi.nlm.nih.gov/20956523/)) [S6]

More recent Drosophila work strengthens the case for a neural homeostatic function. In that study, Pfdn5 physically associated with axonal microtubules, loss of Pfdn5 caused neuromuscular junction defects and worsened tau-induced synaptic abnormalities, and neuronal overexpression reduced tau aggregation, neurodegeneration, and memory deficits. The paper explicitly framed Pfdn5 as controlling Tau toxicity via microtubule stability, which fits a neural homeostasis/proteostasis model. ([pubmed.ncbi.nlm.nih.gov](https://pubmed.ncbi.nlm.nih.gov/41533771/)) [S7]

Additional support comes from protein quality-control studies. In a JBC study, prefoldin localized with polyubiquitinated proteins, and prefoldin knockdown increased insoluble ubiquitinated protein accumulation under proteotoxic stress. Importantly, mice carrying an L110R mutation in MM-1α/PFD5/PFDN5 had increased polyubiquitinated protein aggregation in the brain, and the paper states that dysfunction of prefoldin can contribute to neurodegenerative disease. This is not a pure “neural signaling” role, but it is directly relevant to neural homeostasis because brain tissue showed impaired handling of damaged proteins. ([pubmed.ncbi.nlm.nih.gov](https://pubmed.ncbi.nlm.nih.gov/23946485/)) [S8]

Finally, neuronal cell data in Huntington-related models are consistent with a protective homeostatic role. In Neuro-2a neuronal cells, knockdown of **PFD5** disrupted prefoldin and increased aggregation of pathogenic huntingtin/polyQ species and cell death, while prefoldin overexpression suppressed aggregation. Because this study addresses the prefoldin complex rather than an isolated PFDN5-specific mechanism, it is somewhat indirect, but it still supports the idea that PFDN5-containing prefoldin participates in neuronal proteostasis. ([pubmed.ncbi.nlm.nih.gov](https://pubmed.ncbi.nlm.nih.gov/23720755/)) [S9]

## Contradicting or limiting evidence

The biggest limitation is that most evidence is **preclinical**. I did not identify strong human genetic evidence showing that PFDN5 mutations are a well-established cause of a primary neurologic disorder, nor strong clinical literature positioning PFDN5 as a canonical human neural homeostasis regulator. The evidence base is instead centered on mouse, fly, and neuronal cell models. ([pubmed.ncbi.nlm.nih.gov](https://pubmed.ncbi.nlm.nih.gov/20956523/)) [S10]

Also, PFDN5 is **not neural-specific**. Human Protein Atlas data show low tissue specificity and low brain regional specificity, which suggests it is better viewed as a broadly expressed intracellular chaperone/co-chaperone that also functions in neural tissue, rather than a dedicated neural regulator. ([proteinatlas.org](https://www.proteinatlas.org/ENSG00000123349-PFDN5/tissue)) [S11]

Some of the neural evidence is also more **correlative than causal**. For example, the hippocampal synaptic plasticity paper mainly shows expression changes with nicotine and with impaired plasticity in a tau model; that supports association, but does not by itself prove that endogenous PFDN5 is a master controller of plasticity. ([pubmed.ncbi.nlm.nih.gov](https://pubmed.ncbi.nlm.nih.gov/31505240/)) [S12]

Finally, part of the mechanistic literature concerns the **prefoldin complex as a whole** rather than PFDN5 uniquely. That means some claims about neuronal proteostasis are safest when phrased as “PFDN5-containing prefoldin is involved,” rather than attributing every effect to PFDN5 alone. ([pubmed.ncbi.nlm.nih.gov](https://pubmed.ncbi.nlm.nih.gov/23720755/)) [S13]

## Mechanistic / biological rationale

A plausible biological rationale is straightforward: prefoldin helps fold and maintain proteins central to **actin/tubulin cytoskeleton** and broader **protein quality control**. Neurons are unusually vulnerable to failures in these systems because of their polarized morphology, reliance on axonal transport, synaptic maintenance, and long lifespan. Thus, a chaperone/co-chaperone like PFDN5 is well positioned to influence neural homeostasis. ([ncbi.nlm.nih.gov](https://www.ncbi.nlm.nih.gov/gene/5204?utm_source=openai)) [S14]

The best neural-specific mechanism so far is **microtubule stabilization and suppression of toxic aggregation**. In Drosophila neurons, Pfdn5 associated with axonal microtubules and limited tau toxicity; loss of Pfdn5 worsened synaptic morphology and neurodegeneration, while overexpression improved memory deficits. That is a coherent homeostatic mechanism linking cytoskeletal maintenance to protection from neurotoxic proteinopathy. ([pubmed.ncbi.nlm.nih.gov](https://pubmed.ncbi.nlm.nih.gov/41533771/)) [S15]

A second mechanism is **proteostasis under stress**. Prefoldin/PFDN5 dysfunction increases ubiquitinated protein accumulation and brain aggregate burden in mutant mice, consistent with reduced capacity to maintain protein homeostasis in neural tissue. ([pubmed.ncbi.nlm.nih.gov](https://pubmed.ncbi.nlm.nih.gov/23946485/)) [S16]

There is also a broader **regulatory** angle: NCBI notes that PFDN5 may repress c-Myc transcriptional activity, and human studies show PFDN5/PFDN2 can influence co-transcriptional pre-mRNA splicing. Those functions are established mainly outside neuroscience, but they make it biologically plausible that PFDN5 could affect neural-state regulation beyond simple chaperoning. The current neural evidence for those nuclear/regulatory roles is still limited. ([ncbi.nlm.nih.gov](https://www.ncbi.nlm.nih.gov/gene/5204?utm_source=openai)) [S17]

## Bottom line

The claim is **likely true**. PFDN5 is associated with neural **regulatory/homeostatic** biology in several meaningful ways: expression in brain cell types, association with hippocampal synaptic plasticity, requirement for normal sensory/neuronal development, and protection of neurons from proteotoxic and tau-related stress. But the strongest evidence is still from model systems, and PFDN5 currently looks more like a **broad intracellular proteostasis/cytoskeletal factor with important neural roles** than a definitively established, human-specific neural homeostasis regulator. ([proteinatlas.org](https://www.proteinatlas.org/ENSG00000123349-PFDN5/single%2Bcell)) [S18]

## Cited sources

- **[S1]** PFDN5 prefoldin subunit 5 [Homo sapiens (human)] - Gene - NCBI — https://www.ncbi.nlm.nih.gov/gene/5204?utm_source=openai
- **[S2]** Changes in the expression of prefoldin subunit 5 depending on synaptic plasticity in the mouse hippocampus - PubMed — https://pubmed.ncbi.nlm.nih.gov/31505240/?utm_source=openai
- **[S3]** PFDN5 prefoldin subunit 5 [Homo sapiens (human)] - Gene - NCBI — https://www.ncbi.nlm.nih.gov/gene/5204?utm_source=openai
- **[S4]** Brain tissue expression of PFDN5 - Summary - The Human Protein Atlas — https://www.proteinatlas.org/ENSG00000123349-PFDN5/brain
- **[S5]** Changes in the expression of prefoldin subunit 5 depending on synaptic plasticity in the mouse hippocampus - PubMed — https://pubmed.ncbi.nlm.nih.gov/31505240/
- **[S6]** Prefoldin 5 is required for normal sensory and neuronal development in a murine model - PubMed — https://pubmed.ncbi.nlm.nih.gov/20956523/
- **[S7]** Prefoldin 5 is a microtubule-associated protein that suppresses Tau aggregation and neurotoxicity - PubMed — https://pubmed.ncbi.nlm.nih.gov/41533771/
- **[S8]** Prefoldin plays a role as a clearance factor in preventing proteasome inhibitor-induced protein aggregation - PubMed — https://pubmed.ncbi.nlm.nih.gov/23946485/
- **[S9]** Prefoldin protects neuronal cells from polyglutamine toxicity by preventing aggregation formation - PubMed — https://pubmed.ncbi.nlm.nih.gov/23720755/
- **[S10]** Prefoldin 5 is required for normal sensory and neuronal development in a murine model - PubMed — https://pubmed.ncbi.nlm.nih.gov/20956523/
- **[S11]** Tissue expression of PFDN5 - Summary - The Human Protein Atlas — https://www.proteinatlas.org/ENSG00000123349-PFDN5/tissue
- **[S12]** Changes in the expression of prefoldin subunit 5 depending on synaptic plasticity in the mouse hippocampus - PubMed — https://pubmed.ncbi.nlm.nih.gov/31505240/
- **[S13]** Prefoldin protects neuronal cells from polyglutamine toxicity by preventing aggregation formation - PubMed — https://pubmed.ncbi.nlm.nih.gov/23720755/
- **[S14]** PFDN5 prefoldin subunit 5 [Homo sapiens (human)] - Gene - NCBI — https://www.ncbi.nlm.nih.gov/gene/5204?utm_source=openai
- **[S15]** Prefoldin 5 is a microtubule-associated protein that suppresses Tau aggregation and neurotoxicity - PubMed — https://pubmed.ncbi.nlm.nih.gov/41533771/
- **[S16]** Prefoldin plays a role as a clearance factor in preventing proteasome inhibitor-induced protein aggregation - PubMed — https://pubmed.ncbi.nlm.nih.gov/23946485/
- **[S17]** PFDN5 prefoldin subunit 5 [Homo sapiens (human)] - Gene - NCBI — https://www.ncbi.nlm.nih.gov/gene/5204?utm_source=openai
- **[S18]** Single cell type - PFDN5 - The Human Protein Atlas — https://www.proteinatlas.org/ENSG00000123349-PFDN5/single%2Bcell
