## Supplementary material for "Understanding Substance Dependence: What Differentiates Addictive from Non-Addictive Drugs?": Folder for supplementary files: non-addictive-genes-claims-first100-052-thy1-gene-protein-has-been-associated-with-regulat-45a5bd6f8429.md

# non-addictive-genes-claims-first100-052-45a5bd6f8429

**Original claim:** THY1 gene/protein has been associated with regulatory or homeostatic biological processes in neural systems.

## Claim

THY1 gene/protein has been associated with regulatory or homeostatic biological processes in neural systems.

## Verdict

Verdict: Highly Supported

## Confidence

High

## Supporting evidence

Authoritative curated resources already place THY1 in neural regulatory biology. The reviewed UniProt entry for human THY1 states that it “may play a role in cell-cell or cell-ligand interactions during synaptogenesis and other events in the brain.” It also annotates THY1 to neural compartments including dendrite, growth cone, axolemma, glutamatergic synapse, presynapse, and postsynapse, and to biological processes such as cell-cell signaling, integrin-mediated signaling, negative regulation of axonogenesis, and negative regulation of neuron projection regeneration. Reactome repeats the synaptogenesis/brain-function annotation. ([rest.uniprot.org](https://rest.uniprot.org/uniprotkb/P04216.txt)) [S1]

There is now direct experimental evidence for a bona fide homeostatic role in neuron-glia communication. A 2025 *Glia* study using both whole-body and neuron-specific Thy1 knockout mice concluded that neuronal THY1 regulates astrocyte activation state and contributes to the “maintenance and restoration of tissue homeostasis in the brain.” In those mice, loss of neuronal THY1 led to a partially activated astrocyte phenotype, while under physiological conditions THY1-bearing neurons helped keep astrocytes in a nonreactive/quiescent state. ([pmc.ncbi.nlm.nih.gov](https://pmc.ncbi.nlm.nih.gov/articles/PMC12666997/)) [S2]

That same study further strengthens the “homeostatic” part of the claim mechanistically and phenotypically. The authors reported that deletion of Thy1 increased astrocytic intermediate-filament markers such as **Gfap** and **Vim**, and after cortical injury the absence of neuronal THY1 was associated with failure to terminate astrocyte activation appropriately. They explicitly interpret THY1 as important for restoring homeostatic conditions in the brain, especially under stressors such as aging or injury. ([pmc.ncbi.nlm.nih.gov](https://pmc.ncbi.nlm.nih.gov/articles/PMC12666997/)) [S3]

Older primary studies also support a broader regulatory role in neural systems. In a 2013 study, blocking the relevant neuroglial integrin pathway or interfering with Thy1-mediated neuron-astrocyte contact increased astrocytic CNTF expression; the authors concluded that neuronal Thy1 functions as a neuroglial CNTF repressor and that neuron-astroglia contact acts as a sensor enabling rapid neurotrophic responses. Because CNTF is a classic neurotrophic and injury-response factor, this is strong evidence that THY1 participates in regulation of trophic/homeostatic signaling between neurons and astrocytes. ([pubmed.ncbi.nlm.nih.gov](https://pubmed.ncbi.nlm.nih.gov/23693126/)) [S4]

Additional mechanistic work shows THY1 is involved in structural regulation of neuron-glia interactions. A 2017 study found that astrocyte-to-neuron signaling through integrin-engaged Thy-1 triggers neurite retraction via a CBP/Csk/Src/RhoA/ROCK pathway, directly linking THY1 to transcellular control of neuronal cytoskeleton remodeling. Earlier work also showed that astrocytic αVβ3 integrin is sufficient to trigger Thy-1-mediated inhibition of neurite outgrowth and retraction of neuronal processes, and a 1992 *Nature* paper reported that Thy-1 expression inhibits neurite outgrowth on mature astrocytes. These are clear examples of THY1 being associated with regulatory control of neural process growth and stabilization. ([pubmed.ncbi.nlm.nih.gov](https://pubmed.ncbi.nlm.nih.gov/27842221/)) [S5]

THY1 has also been associated with synaptic regulation. A classic study found Thy-1 in multiple populations of neuronal secretory vesicles and argued that it plays an important role in regulated neurotransmitter release at the synapse. Consistent with this, knockout studies reported regionally selective impairment of long-term potentiation in the dentate gyrus of Thy-1-null mice, and follow-up electrophysiology showed enhanced inhibitory synaptic bursts in dentate granule cells from knockout animals. Together, these findings support a role in regulating synaptic plasticity and inhibitory/excitatory network behavior, even if the effects are subtle and region-specific. ([pubmed.ncbi.nlm.nih.gov](https://pubmed.ncbi.nlm.nih.gov/9456327/?utm_source=openai)) [S6]

## Contradicting or limiting evidence

The main limitation is not absence of evidence, but uneven depth and context dependence. Even the recent *Glia* paper notes that although the Thy1 promoter is widely used in neuroscience, the endogenous role of THY1 in the brain had remained “largely unknown,” and the newer homeostatic interpretation is relatively recent. Earlier reviews similarly described the in vivo neural role of Thy-1 as elusive or lacking consensus. ([pubmed.ncbi.nlm.nih.gov](https://pubmed.ncbi.nlm.nih.gov/40936326/?utm_source=openai)) [S7]

Much of the strongest evidence comes from rodent systems, neuron-astrocyte co-cultures, injury models, or mechanistic cell biology rather than direct human clinical studies. That matters because the claim is broad enough to be true on association grounds, but the exact magnitude and generality of THY1’s role across all human neural circuits is still incompletely mapped. ([pmc.ncbi.nlm.nih.gov](https://pmc.ncbi.nlm.nih.gov/articles/PMC12666997/)) [S8]

The phenotype is also not one of catastrophic CNS failure when THY1 is removed. The 2025 study notes that young adult knockout mice can appear largely normal under baseline conditions, with age- or stress-dependent effects becoming more apparent. Likewise, earlier work found normal gross spatial learning despite dentate-gyrus LTP impairment, implying that THY1’s neural functions can be subtle, compensated, circuit-specific, or most evident under challenge conditions. ([pmc.ncbi.nlm.nih.gov](https://pmc.ncbi.nlm.nih.gov/articles/PMC12666997/)) [S9]

Finally, not every THY1-associated neural process is strictly “homeostatic” in the narrow physiological sense. Some evidence is more about synaptogenesis, neurite outgrowth, axonal regeneration restraint, or injury-related remodeling than about classic network homeostasis. So the strongest reading of the claim is that THY1 is associated with **regulatory** neural processes broadly, with the **homeostatic** label especially well supported for astrocyte quiescence/reactivity and tissue-restoration contexts. ([rest.uniprot.org](https://rest.uniprot.org/uniprotkb/P04216.txt)) [S10]

## Mechanistic / biological rationale

THY1 is a GPI-anchored surface glycoprotein enriched in brain tissue and localized to neuronal membranes and synaptic compartments. That positioning makes biological sense for a regulator of contact-dependent signaling rather than a purely structural marker. Its curated localization to dendrite, growth cone, and synaptic compartments is consistent with roles in neurite dynamics, synaptogenesis, and neuron-glia communication. ([rest.uniprot.org](https://rest.uniprot.org/uniprotkb/P04216.txt)) [S11]

Mechanistically, THY1 engages astrocytic integrins and related adhesion machinery in trans. Depending on context, this activates or modulates focal-adhesion and cytoskeletal signaling pathways including FAK/JNK, CBP/Csk/Src, RhoA/ROCK, and associated receptor clustering. Those pathways can alter astrocyte activation state, astrocyte migration, proliferation/apoptosis balance, CNTF expression, and neuronal neurite stability or retraction. This is exactly the kind of molecular architecture expected for a regulator of neural homeostasis and intercellular feedback. ([pubmed.ncbi.nlm.nih.gov](https://pubmed.ncbi.nlm.nih.gov/23693126/)) [S12]

In other words, THY1 is plausibly part of a membrane-level “set point” system: neurons expressing THY1 can restrain or tune astrocyte reactivity, while reciprocal astrocyte-integrin engagement can influence neuronal morphology and plasticity. That provides a coherent biological explanation for why THY1 is linked both to homeostatic astrocyte quiescence and to regulation of neurite/synaptic behavior. ([pmc.ncbi.nlm.nih.gov](https://pmc.ncbi.nlm.nih.gov/articles/PMC12666997/)) [S13]

## Bottom line

Yes. The claim is well supported. THY1 is associated with multiple **regulatory** processes in neural systems, including synaptogenesis-related cell interactions, regulation of axon/neurite growth and regeneration, synaptic plasticity, and neuron-astrocyte signaling. There is also now direct evidence that neuronal THY1 helps maintain astrocytes in a quiescent state and contributes to the **maintenance/restoration of brain tissue homeostasis**. The main caveat is that the strongest evidence is preclinical and context-dependent rather than broad human clinical proof. ([rest.uniprot.org](https://rest.uniprot.org/uniprotkb/P04216.txt)) [S14]

## Cited sources

- **[S1]** rest.uniprot.org — https://rest.uniprot.org/uniprotkb/P04216.txt
- **[S2]** 
            Neuronal THY1 Signaling Maintains Astrocytes in a Quiescent State - PMC
         — https://pmc.ncbi.nlm.nih.gov/articles/PMC12666997/
- **[S3]** 
            Neuronal THY1 Signaling Maintains Astrocytes in a Quiescent State - PMC
         — https://pmc.ncbi.nlm.nih.gov/articles/PMC12666997/
- **[S4]** Inhibition of a novel specific neuroglial integrin signaling pathway increases STAT3-mediated CNTF expression - PubMed — https://pubmed.ncbi.nlm.nih.gov/23693126/
- **[S5]** Astrocyte-to-neuron communication through integrin-engaged Thy-1/CBP/Csk/Src complex triggers neurite retraction via the RhoA/ROCK pathway - PubMed — https://pubmed.ncbi.nlm.nih.gov/27842221/
- **[S6]** Thy-1 is a component common to multiple populations of synaptic vesicles - PubMed — https://pubmed.ncbi.nlm.nih.gov/9456327/?utm_source=openai
- **[S7]** Neuronal THY1 Signaling Maintains Astrocytes in a Quiescent State - PubMed — https://pubmed.ncbi.nlm.nih.gov/40936326/?utm_source=openai
- **[S8]** 
            Neuronal THY1 Signaling Maintains Astrocytes in a Quiescent State - PMC
         — https://pmc.ncbi.nlm.nih.gov/articles/PMC12666997/
- **[S9]** 
            Neuronal THY1 Signaling Maintains Astrocytes in a Quiescent State - PMC
         — https://pmc.ncbi.nlm.nih.gov/articles/PMC12666997/
- **[S10]** rest.uniprot.org — https://rest.uniprot.org/uniprotkb/P04216.txt
- **[S11]** rest.uniprot.org — https://rest.uniprot.org/uniprotkb/P04216.txt
- **[S12]** Inhibition of a novel specific neuroglial integrin signaling pathway increases STAT3-mediated CNTF expression - PubMed — https://pubmed.ncbi.nlm.nih.gov/23693126/
- **[S13]** 
            Neuronal THY1 Signaling Maintains Astrocytes in a Quiescent State - PMC
         — https://pmc.ncbi.nlm.nih.gov/articles/PMC12666997/
- **[S14]** rest.uniprot.org — https://rest.uniprot.org/uniprotkb/P04216.txt
