## Supplementary material for "Understanding Substance Dependence: What Differentiates Addictive from Non-Addictive Drugs?": Folder for supplementary files: non-addictive-genes-claims-first100-053-rcan2-gene-protein-has-been-associated-with-regula-7fcaf20b4090.md

# non-addictive-genes-claims-first100-053-7fcaf20b4090

**Original claim:** RCAN2 gene/protein has been associated with regulatory or homeostatic biological processes in neural systems.

## Claim

RCAN2 gene/protein has been associated with regulatory or homeostatic biological processes in neural systems.

## Verdict

Verdict: Likely True

## Confidence

Medium

## Supporting evidence

RCAN2 is a bona fide regulator of the calcineurin pathway, and calcineurin is a major calcium-sensitive signaling node in the nervous system. The NCBI Gene entry describes RCAN2 as a member of the regulator-of-calcineurin family that binds calcineurin A and inhibits calcineurin-mediated NFAT signaling; it also notes broad expression in brain. In the original human isoform paper, the brain-specific/brain-enriched expression pattern was especially notable: the ZAKI-4/RCAN2 alpha transcript was reported as brain-exclusive, while beta transcripts were most abundant in brain among several tissues tested. ([ncbi.nlm.nih.gov](https://www.ncbi.nlm.nih.gov/gene/10231)) [S1]

There is direct evidence that RCAN2 is present across neural tissues and is regulated in development. A mouse brain mapping study found Rcan2 transcripts and proteins in every adult brain region examined, with region- and stage-specific distribution patterns; the authors concluded that the differential RCAN-family expression patterns “suggest selective functions” in the brain. In rats, ZAKI-4/RCAN2 mRNA was detected in cerebral cortex and cerebellum from late embryonic life onward, was widely expressed in neurons, and showed strong expression in regions including hippocampus and substantia nigra. ([pubmed.ncbi.nlm.nih.gov](https://pubmed.ncbi.nlm.nih.gov/17767500/)) [S2]

The strongest functional evidence for a neural homeostatic role comes from hypothalamic feeding regulation. In Rcan2 knockout mice, loss of Rcan2 reduced food intake and protected against age- and diet-induced obesity. The same study localized strong Rcan2 expression to ventromedial, dorsomedial, and paraventricular hypothalamic nuclei—classic neural centers governing energy balance—and found that fasting specifically increased hypothalamic Rcan2-3 mRNA, while Rcan2 deficiency blunted the hyperphagic response to starvation. The authors further reported that Rcan2 and leptin regulate body weight through different pathways, supporting a specific role in neural control of feeding homeostasis rather than a trivial downstream effect. ([pubmed.ncbi.nlm.nih.gov](https://pubmed.ncbi.nlm.nih.gov/21298050/)) [S3]

That hypothalamic/body-weight finding was partly reinforced in a later mouse study in females. That paper states that Rcan2 increases food intake, examines hypothalamic Rcan2 expression, and concludes that Rcan2 and estradiol independently regulate body weight through different mechanisms, with Rcan2 promoting intake and estradiol promoting expenditure. This is still metabolism-focused, but it is explicitly centered on hypothalamic neural circuits controlling a homeostatic behavior. ([pubmed.ncbi.nlm.nih.gov](https://pubmed.ncbi.nlm.nih.gov/28624805/)) [S4]

Taken together, these data support more than mere tissue expression: RCAN2 is brain-expressed, developmentally regulated in neurons, enriched in hypothalamic nuclei that govern homeostatic behaviors, and genetically linked in mice to regulation of food intake and starvation responses. That is enough to say RCAN2 has been associated with regulatory/homeostatic processes in neural systems. ([pubmed.ncbi.nlm.nih.gov](https://pubmed.ncbi.nlm.nih.gov/17767500/)) [S5]

## Contradicting or limiting evidence

The evidence base is not equally strong across all kinds of “neural homeostasis.” Most direct functional data are from rodent studies of hypothalamic energy-balance regulation, not from human neural physiology or from broad tests of synaptic/homeostatic plasticity across the CNS. So the claim is supported, but mainly in a specific neural-homeostatic domain rather than universally across neural systems. ([pubmed.ncbi.nlm.nih.gov](https://pubmed.ncbi.nlm.nih.gov/21298050/)) [S6]

There are also mechanistic uncertainties. One mouse study noted that Rcan2 brain distribution did not coincide well with calcineurin-rich regions such as hippocampus, arguing against RCAN2 being a simple universal proxy for calcineurin regulation throughout the brain. A later human obesity paper additionally summarized that there was no significant difference in hypothalamic calcineurin phosphatase activity between RCAN2-knockout and wild-type mice, suggesting that the biochemical mechanism in hypothalamic circuits may be more complex than straightforward global calcineurin inhibition. ([pmc.ncbi.nlm.nih.gov](https://pmc.ncbi.nlm.nih.gov/articles/PMC3029291/?utm_source=openai)) [S7]

Also, some of the neural evidence is associative or anatomical rather than causal. Expression in neurons, developmental regulation, or thyroid-hormone responsiveness in brain does not by itself prove a homeostatic function. Those data strengthen plausibility, but the causal case relies mostly on knockout mouse phenotypes in feeding/energy regulation. ([pubmed.ncbi.nlm.nih.gov](https://pubmed.ncbi.nlm.nih.gov/12102656/)) [S8]

## Mechanistic / biological rationale

The mechanistic rationale is biologically coherent. Calcineurin is highly expressed in brain and functions as a calcium-sensitive phosphatase involved in neuronal signaling, receptor regulation, synaptic plasticity, and calcium handling. Reviews of calcineurin in the nervous system describe it as participating in neuronal and glial processes relevant to homeostasis, including regulation of intracellular Ca2+ and synaptic function. Because RCAN2 binds calcineurin and inhibits its signaling output, RCAN2 is well positioned to act as an endogenous “brake” or tuning factor on calcium/calcineurin-dependent neural pathways. ([pmc.ncbi.nlm.nih.gov](https://pmc.ncbi.nlm.nih.gov/articles/PMC6062828/?utm_source=openai)) [S9]

RCAN2’s isoform biology strengthens that interpretation. The human alpha isoform is brain-specific, and thyroid hormone selectively induces that isoform while decreasing endogenous calcineurin activity in the experimental system used. In rat brain, RCAN2 expression changes during development and is altered by hypothyroid status in cortex. Those findings fit a model in which RCAN2 helps tune neural signaling states during development and endocrine adaptation, rather than serving only as a generic housekeeping protein. ([pubmed.ncbi.nlm.nih.gov](https://pubmed.ncbi.nlm.nih.gov/12102656/)) [S10]

The clearest homeostatic application is hypothalamic control of feeding. RCAN2 is concentrated in hypothalamic nuclei that coordinate appetite and energy balance, fasting upregulates the brain-predominant Rcan2-3 isoform in hypothalamus, and knockout mice show reduced food intake and reduced rebound feeding after starvation. That pattern is very consistent with involvement in neural set-point regulation of ingestive behavior. ([pubmed.ncbi.nlm.nih.gov](https://pubmed.ncbi.nlm.nih.gov/21298050/)) [S11]

## Bottom line

Yes—there is credible evidence that **RCAN2 is associated with regulatory/homeostatic processes in neural systems**, especially **hypothalamic regulation of food intake and body-weight homeostasis** in mice, with supporting brain-expression and developmental data in rodents and brain-enriched isoform data in humans. However, the strongest evidence is **not** for broad CNS homeostasis in general, and **direct human functional evidence remains limited**. So the fairest overall assessment is **Likely True**, not “Highly Supported.” ([pubmed.ncbi.nlm.nih.gov](https://pubmed.ncbi.nlm.nih.gov/21298050/)) [S12]

## Cited sources

- **[S1]** RCAN2 regulator of calcineurin 2 [Homo sapiens (human)] - Gene - NCBI — https://www.ncbi.nlm.nih.gov/gene/10231
- **[S2]** Differential expression of members of the RCAN family of calcineurin regulators suggests selective functions for these proteins in the brain - PubMed — https://pubmed.ncbi.nlm.nih.gov/17767500/
- **[S3]** Inactivation of the Rcan2 gene in mice ameliorates the age- and diet-induced obesity by causing a reduction in food intake - PubMed — https://pubmed.ncbi.nlm.nih.gov/21298050/
- **[S4]** Rcan2 and estradiol independently regulate body weight in female mice - PubMed — https://pubmed.ncbi.nlm.nih.gov/28624805/
- **[S5]** Differential expression of members of the RCAN family of calcineurin regulators suggests selective functions for these proteins in the brain - PubMed — https://pubmed.ncbi.nlm.nih.gov/17767500/
- **[S6]** Inactivation of the Rcan2 gene in mice ameliorates the age- and diet-induced obesity by causing a reduction in food intake - PubMed — https://pubmed.ncbi.nlm.nih.gov/21298050/
- **[S7]** Inactivation of the Rcan2 Gene in Mice Ameliorates the Age- and Diet-Induced Obesity by Causing a Reduction in Food Intake - PMC — https://pmc.ncbi.nlm.nih.gov/articles/PMC3029291/?utm_source=openai
- **[S8]** Novel human ZAKI-4 isoforms: hormonal and tissue-specific regulation and function as calcineurin inhibitors - PubMed — https://pubmed.ncbi.nlm.nih.gov/12102656/
- **[S9]** A Friend or Foe: Calcineurin across the Gamut of Neurological Disorders - PMC — https://pmc.ncbi.nlm.nih.gov/articles/PMC6062828/?utm_source=openai
- **[S10]** Novel human ZAKI-4 isoforms: hormonal and tissue-specific regulation and function as calcineurin inhibitors - PubMed — https://pubmed.ncbi.nlm.nih.gov/12102656/
- **[S11]** Inactivation of the Rcan2 gene in mice ameliorates the age- and diet-induced obesity by causing a reduction in food intake - PubMed — https://pubmed.ncbi.nlm.nih.gov/21298050/
- **[S12]** Inactivation of the Rcan2 gene in mice ameliorates the age- and diet-induced obesity by causing a reduction in food intake - PubMed — https://pubmed.ncbi.nlm.nih.gov/21298050/
