## Supplementary material for "Understanding Substance Dependence: What Differentiates Addictive from Non-Addictive Drugs?": Folder for supplementary files: non-addictive-genes-claims-first100-054-akt3-gene-protein-has-been-associated-with-regulat-a00df8ad858a.md

# non-addictive-genes-claims-first100-054-a00df8ad858a

**Original claim:** AKT3 gene/protein has been associated with regulatory or homeostatic biological processes in neural systems.

## Claim

AKT3 gene/protein has been associated with regulatory or homeostatic biological processes in neural systems.

## Verdict

Verdict: Likely True

AKT3 is strongly associated with **regulatory** processes in neural systems, especially brain growth, neural progenitor behavior, axon/neurite development, hippocampal neurogenesis, synaptic signaling, and cognition. The evidence for **neural homeostasis in the narrow sense** is less direct, but some studies suggest roles in steady-state protein synthesis and maintenance of normal signaling balance in brain circuits. Overall, the claim is supported, but mainly through **developmental and signaling-regulatory** evidence rather than classic homeostatic-plasticity studies. ([medlineplus.gov](https://medlineplus.gov/download/genetics/gene/akt3.pdf)) [S1]

## Confidence

High

Multiple independent lines of evidence support a neural regulatory role for AKT3: authoritative gene summaries, human genetics, mouse knockout models, and cellular/neurodevelopmental studies all converge on AKT3 as a brain-enriched PI3K-AKT-mTOR kinase with nonredundant functions in neural growth and signaling. Confidence is not maximal only because the “homeostatic” part of the claim is supported more indirectly than the “regulatory” part. ([medlineplus.gov](https://medlineplus.gov/download/genetics/gene/akt3.pdf)) [S2]

## Supporting evidence

AKT3 is unusually brain-focused compared with other AKT isoforms. MedlinePlus states that AKT3 is “most active in the nervous system” and is a key regulator of the PI3K-AKT-mTOR pathway, which controls protein synthesis, growth, proliferation, and cell survival; it specifically notes that studies suggest AKT3 plays a critical role in determining brain size. A review likewise summarizes that AKT3 is the predominant AKT isoform in adult mouse brain and is expressed at especially high levels in human fetal brain, supporting a specialized neural regulatory role. ([medlineplus.gov](https://medlineplus.gov/download/genetics/gene/akt3.pdf)) [S3]

Foundational mouse genetics strongly support AKT3 as a regulator of neural growth. In Akt3-null mice, brain size is selectively reduced by about 20%, with smaller and fewer brain cells and attenuated mTOR signaling in brain tissue; this is a fairly direct demonstration that AKT3 regulates neural growth-related processes rather than being a nonspecific housekeeping kinase. Human genetics points in the same direction: loss of AKT3 dosage is linked to microcephaly, while activating variants or increased dosage are linked to megalencephaly and other brain overgrowth disorders. ([pubmed.ncbi.nlm.nih.gov](https://pubmed.ncbi.nlm.nih.gov/15713641/)) [S4]

Human neurodevelopmental disease data further strengthen the association. Somatic activating AKT3 mutations have been found in hemimegalencephaly, and one study reported that AKT3 is the most abundant AKT paralog in brain during neurogenesis, with active AKT signaling enriched in cortical progenitor cells. Larger clinical series show that activating AKT3 mutations are associated with a spectrum of developmental brain malformations, including hemimegalencephaly, polymicrogyria, heterotopia, and diffuse megalencephaly. This is strong evidence that AKT3 participates in regulatory programs shaping neural tissue architecture. ([pubmed.ncbi.nlm.nih.gov](https://pubmed.ncbi.nlm.nih.gov/22500628/)) [S5]

Evidence also extends beyond embryonic brain growth into adult neural regulation. In adult mouse hippocampus, progenitor cells, neuroblasts, and mature newborn neurons in the dentate gyrus express Akt3 protein. Akt3 knockout reduced progenitor proliferation, neurite growth, telomerase activity, telomere length, TERT expression, and downstream mTOR effectors such as p70S6K and 4EBP1; the authors concluded that Akt3-mTOR signaling plays an important role in adult hippocampal neurogenesis. That places AKT3 in a bona fide neural regulatory process beyond gross morphogenesis. ([pubmed.ncbi.nlm.nih.gov](https://pubmed.ncbi.nlm.nih.gov/34077553/)) [S6]

Cellular neurodevelopment studies also implicate AKT3 in neurite and axon regulation. In cultured hippocampal neurons and embryonic brain slices, Akt3 knockdown reduced neurite length and branch points, and the authors concluded that Akt3 contributes to proper axon growth and branching during neural development. This is another direct neural regulatory phenotype. ([pubmed.ncbi.nlm.nih.gov](https://pubmed.ncbi.nlm.nih.gov/31471318/)) [S7]

There is also evidence for AKT3 in neural signaling and circuit-related function. AKT3 is expressed in hippocampal neurons and dendritic regions, and isoform-specific studies show AKT3 participates in hippocampal synaptic plasticity pathways, with distinct roles relative to AKT1. Separate knockout studies found that loss of Akt3 impairs spatial memory and long-lasting CA1 LTP via reduced mTOR-p70S6K/4EBP signaling, and other work found altered prefrontal/hippocampal cognitive function with deregulated AKT/mTORC2 signaling, suggesting AKT3 helps regulate signaling states important for normal circuit function. ([pmc.ncbi.nlm.nih.gov](https://pmc.ncbi.nlm.nih.gov/articles/PMC5722612/)) [S8]

## Contradicting or limiting evidence

The main limitation is that **direct evidence for “homeostatic” neural processes is limited**. Most of the literature located here concerns brain development, progenitor proliferation, protein synthesis, axon growth, synaptic plasticity, white-matter integrity, and neurodevelopmental disease—not classic neuronal homeostasis paradigms such as homeostatic synaptic scaling or intrinsic excitability control. So the claim is strongest if “regulatory” is taken broadly, and weaker if it is meant to assert a specific, well-established role in canonical neural homeostasis. ([pubmed.ncbi.nlm.nih.gov](https://pubmed.ncbi.nlm.nih.gov/15713641/)) [S9]

There is also some functional heterogeneity across synaptic studies. One hippocampal study found that single Akt3 knockout did **not** impair basal transmission, paired-pulse facilitation, early LTP, or CA1 late LTP, and proposed that AKT1 may compensate in that paradigm. By contrast, another study reported impaired long-LTP and spatial cognition in Akt3 knockout mice due to reduced mTOR signaling. These results are not irreconcilable, but they do show that AKT3’s role can be circuit-, assay-, or compensation-dependent rather than universal across all neural regulatory phenomena. ([pmc.ncbi.nlm.nih.gov](https://pmc.ncbi.nlm.nih.gov/articles/PMC5722612/)) [S10]

Finally, much of the strongest evidence comes from **mouse models** and **human pathogenic variants**, which are highly informative but do not fully define AKT3’s role in normal adult human neural physiology. The association is real, but the detailed boundaries of its role in day-to-day neural homeostasis remain incompletely mapped. ([pubmed.ncbi.nlm.nih.gov](https://pubmed.ncbi.nlm.nih.gov/28969385/)) [S11]

## Mechanistic / biological rationale

AKT3 encodes a serine/threonine kinase in the PI3K-AKT-mTOR pathway, a central signaling axis controlling translation, growth, proliferation, and survival. Because AKT3 is enriched in nervous tissue and prominent during fetal brain development, perturbing it would be expected to alter neural regulatory programs more strongly than perturbing some other AKT isoforms. ([medlineplus.gov](https://medlineplus.gov/download/genetics/gene/akt3.pdf)) [S12]

Mechanistically, loss of AKT3 lowers brain mTOR signaling and reduces phosphorylation of translational effectors such as S6, p70S6K, and 4EBP-family proteins; gain-of-function mutations increase kinase activity and drive excessive growth in the developing brain. In adult hippocampus, Akt3-mTOR signaling also influences TERT/telomerase-related biology, progenitor proliferation, and neurite growth. Together, these findings make biological sense for a molecule involved in regulating neural growth, progenitor pool behavior, and aspects of circuit plasticity/maintenance. ([pubmed.ncbi.nlm.nih.gov](https://pubmed.ncbi.nlm.nih.gov/15713641/)) [S13]

For the “homeostatic” angle specifically, the best support is indirect: one isoform-specific hippocampal study suggested AKT3 may be more important for **steady-state protein synthesis**, whereas AKT1 is more involved in activity-induced translation during CA1 L-LTP. That does not by itself prove a classic homeostatic role, but it is consistent with AKT3 helping maintain baseline neural biosynthetic/signaling set points. ([pmc.ncbi.nlm.nih.gov](https://pmc.ncbi.nlm.nih.gov/articles/PMC5722612/)) [S14]

## Bottom line

The claim is **likely true**. AKT3 is well associated with **regulatory biological processes in neural systems**, including brain growth, cortical development, neural progenitor proliferation, axon/neurite development, hippocampal neurogenesis, synaptic signaling, and cognitive circuit function. The evidence for **homeostatic** neural roles exists but is more indirect and much less developed than the evidence for developmental and signaling regulation. ([medlineplus.gov](https://medlineplus.gov/download/genetics/gene/akt3.pdf)) [S15]

## Cited sources

- **[S1]** AKT3 gene — https://medlineplus.gov/download/genetics/gene/akt3.pdf
- **[S2]** AKT3 gene — https://medlineplus.gov/download/genetics/gene/akt3.pdf
- **[S3]** AKT3 gene — https://medlineplus.gov/download/genetics/gene/akt3.pdf
- **[S4]** Role for Akt3/protein kinase Bgamma in attainment of normal brain size - PubMed — https://pubmed.ncbi.nlm.nih.gov/15713641/
- **[S5]** Somatic activation of AKT3 causes hemispheric developmental brain malformations - PubMed — https://pubmed.ncbi.nlm.nih.gov/22500628/
- **[S6]** Akt3-mTOR regulates hippocampal neurogenesis in adult mouse - PubMed — https://pubmed.ncbi.nlm.nih.gov/34077553/
- **[S7]** SIAH1 ubiquitin ligase mediates ubiquitination and degradation of Akt3 in neural development - PubMed — https://pubmed.ncbi.nlm.nih.gov/31471318/
- **[S8]** 
            AKT isoforms have distinct hippocampal expression and roles in synaptic plasticity - PMC
         — https://pmc.ncbi.nlm.nih.gov/articles/PMC5722612/
- **[S9]** Role for Akt3/protein kinase Bgamma in attainment of normal brain size - PubMed — https://pubmed.ncbi.nlm.nih.gov/15713641/
- **[S10]** 
            AKT isoforms have distinct hippocampal expression and roles in synaptic plasticity - PMC
         — https://pmc.ncbi.nlm.nih.gov/articles/PMC5722612/
- **[S11]** Mutations of AKT3 are associated with a wide spectrum of developmental disorders including extreme megalencephaly - PubMed — https://pubmed.ncbi.nlm.nih.gov/28969385/
- **[S12]** AKT3 gene — https://medlineplus.gov/download/genetics/gene/akt3.pdf
- **[S13]** Role for Akt3/protein kinase Bgamma in attainment of normal brain size - PubMed — https://pubmed.ncbi.nlm.nih.gov/15713641/
- **[S14]** 
            AKT isoforms have distinct hippocampal expression and roles in synaptic plasticity - PMC
         — https://pmc.ncbi.nlm.nih.gov/articles/PMC5722612/
- **[S15]** AKT3 gene — https://medlineplus.gov/download/genetics/gene/akt3.pdf
