## Supplementary material for "Understanding Substance Dependence: What Differentiates Addictive from Non-Addictive Drugs?": Folder for supplementary files: non-addictive-genes-claims-first100-055-lynx1-gene-protein-has-been-associated-with-regula-e326810af45e.md

# non-addictive-genes-claims-first100-055-e326810af45e

**Original claim:** LYNX1 gene/protein has been associated with regulatory or homeostatic biological processes in neural systems.

## Claim

LYNX1 gene/protein has been associated with regulatory or homeostatic biological processes in neural systems.

## Verdict

Verdict: Highly Supported

## Confidence

High

The literature supports this claim strongly. LYNX1 is not just loosely “associated” with neural regulation; multiple primary studies show that Lynx1 directly binds and modulates neuronal nicotinic acetylcholine receptors (nAChRs), alters receptor kinetics and assembly, constrains adult cortical plasticity, and helps balance neuronal activity with long-term neuronal survival. The main limitation is that most mechanistic evidence comes from rodent, cell, and ex vivo systems rather than direct human intervention studies. ([pubmed.ncbi.nlm.nih.gov](https://pubmed.ncbi.nlm.nih.gov/11906696/)) [S1]

## Supporting evidence

The foundational evidence is direct and mechanistic. The original discovery paper identified lynx1 as a brain-enriched Ly6/neurotoxin-family protein expressed in discrete neuronal populations including hippocampus, cortex, and cerebellum, and concluded it was a novel modulator of nAChRs with likely importance for regulation of cholinergic function in vivo. ([salilab.org](https://salilab.org/pdf/Miwa_Neuron_1999.pdf)) [S2]

That regulatory role was then shown experimentally. In a Neuron study, Ibañez-Tallon et al. demonstrated that Lynx1 colocalizes and physically associates with neuronal nAChRs, enhances desensitization, slows recovery from desensitization, and increases EC50 for agonist activation. Those are classic regulatory effects on receptor responsiveness rather than mere correlational observations. ([pubmed.ncbi.nlm.nih.gov](https://pubmed.ncbi.nlm.nih.gov/11906696/)) [S3]

Later work extended this from receptor kinetics to receptor biogenesis. Nichols et al. showed that Lynx1 acts not only at the plasma membrane but also in the endoplasmic reticulum, where it alters assembly of nascent α4 and β2 subunits and shifts α4β2 receptor stoichiometry toward the lower-sensitivity pentameric form. This is strong evidence that LYNX1 participates in regulatory control of cholinergic signaling at multiple levels, including receptor assembly and trafficking. ([pubmed.ncbi.nlm.nih.gov](https://pubmed.ncbi.nlm.nih.gov/25193667/?utm_source=openai)) [S4]

The in vivo evidence is especially relevant to the “homeostatic” part of the claim. In lynx1-null mice, loss of Lynx1 lowered the nicotine EC50 by about tenfold, reduced desensitization, elevated nicotine-evoked intracellular calcium, enhanced synaptic efficacy, and improved some learning and memory measures. But the same mice developed age-related vacuolating neurodegeneration that was worsened by nicotine and improved by removing nAChR subunits. The authors explicitly concluded that Lynx1 balances neuronal activity and survival in the CNS. That is a strong fit to a neural regulatory/homeostatic role. ([pubmed.ncbi.nlm.nih.gov](https://pubmed.ncbi.nlm.nih.gov/16950157/)) [S5]

A follow-up ultrastructural study reinforced this interpretation. Aged lynx1-null mice showed dorsal striatal degeneration, including loss of nerve fibers, disordered myelin, axoplasmic abnormalities, and neuronal loss. The study argued that Lynx1 is important for maintaining neuronal health during aging and that Lynx1 dosage may mediate a trade-off between neuroprotection and augmented learning. ([pmc.ncbi.nlm.nih.gov](https://pmc.ncbi.nlm.nih.gov/articles/PMC4265479/)) [S6]

LYNX1 also regulates systems-level plasticity. In adult visual cortex, Morishita et al. found that Lynx1 expression rises after the critical period and prevents adult plasticity; removing this “molecular brake” enhanced nAChR signaling and preserved ocular-dominance plasticity in adulthood. The paper specifically states that Lynx1 expression maintains stability of mature cortical networks in the presence of cholinergic innervation. ([pubmed.ncbi.nlm.nih.gov](https://pubmed.ncbi.nlm.nih.gov/21071629/)) [S7]

This plasticity-regulating role extends to structural plasticity. Deletion of Lynx1 in adult mouse visual cortex doubled baseline dendritic spine turnover, indicating that normal Lynx1 function actively limits ongoing adult cortical remodeling. That is again consistent with a regulatory/stabilizing role in neural circuit homeostasis. ([pubmed.ncbi.nlm.nih.gov](https://pubmed.ncbi.nlm.nih.gov/27605620/?utm_source=openai)) [S8]

There is also evidence beyond mouse cortex. In a human/rodent biochemical study, a water-soluble human Lynx1 preparation isolated multiple nAChR subunits from human cortical extracts and reduced nicotine-triggered ERK signaling in neuronal cell systems. The same study found protection against Aβ-induced cytotoxicity in cortical neurons, supporting the idea that Lynx1 can regulate cholinergic signaling and influence neuronal vulnerability in disease-relevant contexts. ([pubmed.ncbi.nlm.nih.gov](https://pubmed.ncbi.nlm.nih.gov/27460145/)) [S9]

Human disease-model data are directionally consistent, though less definitive. In fragile X syndrome iPSC-derived neural progenitors, reduced LYNX1 expression was linked to disturbed progenitor growth and interpreted as suggesting dysregulated cholinergic signaling. This does not prove a primary disease-driving role, but it is compatible with LYNX1 participating in neural regulatory programs in human cells. ([pubmed.ncbi.nlm.nih.gov](https://pubmed.ncbi.nlm.nih.gov/36506088/?utm_source=openai)) [S10]

## Contradicting or limiting evidence

The main limitation is that most strong evidence is from mice, cultured neurons, Xenopus/heterologous systems, or ex vivo tissue assays rather than direct human physiology or clinical studies. So the claim is highly supported for biological function in neural systems, but the human in vivo evidence remains limited. ([pubmed.ncbi.nlm.nih.gov](https://pubmed.ncbi.nlm.nih.gov/11906696/)) [S11]

Also, “homeostatic” is partly an inference from the data rather than always the exact term used by the authors. Primary papers more often describe Lynx1 as a “molecular brake,” a modulator of cholinergic tone, a maintainer of mature network stability, or a balancer of neuronal activity and survival. Those concepts are close to neural homeostasis, but not identical to formal homeostatic plasticity frameworks such as synaptic scaling. ([pubmed.ncbi.nlm.nih.gov](https://pubmed.ncbi.nlm.nih.gov/21071629/)) [S12]

Finally, Lynx1’s effects are context- and receptor-subtype-dependent, not uniformly inhibitory in every neural setting. For example, deletion of lynx1 reduced α6* nAChR function in some synaptosomal assays, and the authors concluded that lynx1 was necessary for normal α6* receptor function. So Lynx1 is best understood as a regulator of nicotinic signaling, not simply a universal suppressor. ([pmc.ncbi.nlm.nih.gov](https://pmc.ncbi.nlm.nih.gov/articles/PMC5716591/?utm_source=openai)) [S13]

If one extends the discussion to addiction, the evidence is more indirect. LYNX1 clearly modulates nicotine-relevant receptor systems, but the literature supports addiction relevance mainly through receptor physiology and mouse behavioral phenotypes, not through strong human causal evidence for addiction risk. ([pmc.ncbi.nlm.nih.gov](https://pmc.ncbi.nlm.nih.gov/articles/PMC5716591/?utm_source=openai)) [S14]

## Mechanistic / biological rationale

Mechanistically, Lynx1 is a GPI-anchored Ly6/prototoxin-family protein that binds nAChRs on neurons. It changes agonist sensitivity, accelerates desensitization, slows recovery from desensitization, and can alter receptor stoichiometry during assembly in the endoplasmic reticulum. These actions would be expected to tune cholinergic gain and prevent runaway receptor activation. ([pubmed.ncbi.nlm.nih.gov](https://pubmed.ncbi.nlm.nih.gov/11906696/)) [S15]

That receptor-level tuning plausibly explains the systems-level phenotypes. If Lynx1 dampens or fine-tunes nicotinic signaling, then removing it should increase cholinergic responsiveness, calcium entry, synaptic efficacy, and plasticity—which is what the mouse knockout studies observed. But excessive or poorly constrained nicotinic activity can also increase vulnerability to degeneration over time, which matches the aging and nicotine-exacerbated phenotypes in lynx1-null mice. ([pubmed.ncbi.nlm.nih.gov](https://pubmed.ncbi.nlm.nih.gov/16950157/)) [S16]

So the overall biological model is coherent: LYNX1 acts as an endogenous regulator of cholinergic receptor function that helps stabilize mature neural circuits and protect neuronal health, while also limiting some forms of experience-dependent plasticity. That is a classic trade-off between adaptability and stability, i.e., a homeostatic-style control function in neural systems. ([pubmed.ncbi.nlm.nih.gov](https://pubmed.ncbi.nlm.nih.gov/21071629/)) [S17]

## Bottom line

Yes. The claim is **highly supported**. Primary literature shows that LYNX1 directly regulates neuronal nicotinic receptor function, shapes cortical plasticity, and helps balance neural excitability with neuronal survival. The “homeostatic” wording is somewhat interpretive, but the underlying evidence for a neural regulatory/stabilizing role is strong. The main caveat is that the best evidence is mechanistic and preclinical rather than human clinical. ([pubmed.ncbi.nlm.nih.gov](https://pubmed.ncbi.nlm.nih.gov/11906696/)) [S18]

## Cited sources

- **[S1]** Novel modulation of neuronal nicotinic acetylcholine receptors by association with the endogenous prototoxin lynx1 - PubMed — https://pubmed.ncbi.nlm.nih.gov/11906696/
- **[S2]** 088464u017 — https://salilab.org/pdf/Miwa_Neuron_1999.pdf
- **[S3]** Novel modulation of neuronal nicotinic acetylcholine receptors by association with the endogenous prototoxin lynx1 - PubMed — https://pubmed.ncbi.nlm.nih.gov/11906696/
- **[S4]** Lynx1 shifts α4β2 nicotinic receptor subunit stoichiometry by affecting assembly in the endoplasmic reticulum - PubMed — https://pubmed.ncbi.nlm.nih.gov/25193667/?utm_source=openai
- **[S5]** The prototoxin lynx1 acts on nicotinic acetylcholine receptors to balance neuronal activity and survival in vivo - PubMed — https://pubmed.ncbi.nlm.nih.gov/16950157/
- **[S6]** 
            lynx1 supports neuronal health in mouse dorsal striatum during aging: an ultrastructural investigation - PMC
         — https://pmc.ncbi.nlm.nih.gov/articles/PMC4265479/
- **[S7]** Lynx1, a cholinergic brake, limits plasticity in adult visual cortex - PubMed — https://pubmed.ncbi.nlm.nih.gov/21071629/
- **[S8]** Lynx1 Limits Dendritic Spine Turnover in the Adult Visual Cortex - PubMed — https://pubmed.ncbi.nlm.nih.gov/27605620/?utm_source=openai
- **[S9]** Lynx1 and Aβ1-42 bind competitively to multiple nicotinic acetylcholine receptor subtypes - PubMed — https://pubmed.ncbi.nlm.nih.gov/27460145/
- **[S10]** Reduced LYNX1 expression in transcriptome of human iPSC-derived neural progenitors modeling fragile X syndrome - PubMed — https://pubmed.ncbi.nlm.nih.gov/36506088/?utm_source=openai
- **[S11]** Novel modulation of neuronal nicotinic acetylcholine receptors by association with the endogenous prototoxin lynx1 - PubMed — https://pubmed.ncbi.nlm.nih.gov/11906696/
- **[S12]** Lynx1, a cholinergic brake, limits plasticity in adult visual cortex - PubMed — https://pubmed.ncbi.nlm.nih.gov/21071629/
- **[S13]** Deletion of lynx1 reduces the function of α6* nicotinic receptors - PMC — https://pmc.ncbi.nlm.nih.gov/articles/PMC5716591/?utm_source=openai
- **[S14]** Deletion of lynx1 reduces the function of α6* nicotinic receptors - PMC — https://pmc.ncbi.nlm.nih.gov/articles/PMC5716591/?utm_source=openai
- **[S15]** Novel modulation of neuronal nicotinic acetylcholine receptors by association with the endogenous prototoxin lynx1 - PubMed — https://pubmed.ncbi.nlm.nih.gov/11906696/
- **[S16]** The prototoxin lynx1 acts on nicotinic acetylcholine receptors to balance neuronal activity and survival in vivo - PubMed — https://pubmed.ncbi.nlm.nih.gov/16950157/
- **[S17]** Lynx1, a cholinergic brake, limits plasticity in adult visual cortex - PubMed — https://pubmed.ncbi.nlm.nih.gov/21071629/
- **[S18]** Novel modulation of neuronal nicotinic acetylcholine receptors by association with the endogenous prototoxin lynx1 - PubMed — https://pubmed.ncbi.nlm.nih.gov/11906696/
