## Supplementary material for "Understanding Substance Dependence: What Differentiates Addictive from Non-Addictive Drugs?": Folder for supplementary files: non-addictive-genes-claims-first100-056-vwa1-gene-protein-has-been-associated-with-regulat-bf4cfb2020f2.md

# non-addictive-genes-claims-first100-056-bf4cfb2020f2

**Original claim:** VWA1 gene/protein has been associated with regulatory or homeostatic biological processes in neural systems.

## Claim

VWA1 gene/protein has been associated with regulatory or homeostatic biological processes in neural systems.

## Verdict

Verdict: Mixed Evidence

VWA1 **has been associated with neural-system biology**, especially in the **peripheral nervous system (PNS)**, where it is expressed in basement membrane structures and where loss of function disrupts peripheral nerve extracellular matrix organization and is linked to hereditary motor neuropathy/neuromyopathy. However, the evidence more strongly supports a **structural ECM maintenance/stabilization role** in peripheral nerve and motor-system development than a clearly established, direct role in **neural “regulatory” or “homeostatic” processes** in the usual neurobiological sense (for example, synaptic homeostasis, neuronal firing homeostasis, or canonical glial regulatory signaling). ([pubmed.ncbi.nlm.nih.gov](https://pubmed.ncbi.nlm.nih.gov/18314316/)) [S1]

## Confidence

Medium

## Supporting evidence

VWA1 encodes WARP, an extracellular-matrix protein. In a mouse reporter/immunostaining study, WARP was found in a **limited subset of basement membranes**, including the **vasculature of neural tissues** and **basement membrane structures of the peripheral nervous system**. This is direct evidence that VWA1 is associated with neural-associated tissue compartments rather than being purely non-neural. ([pubmed.ncbi.nlm.nih.gov](https://pubmed.ncbi.nlm.nih.gov/18314316/)) [S2]

Functional evidence is stronger in the PNS. In Vwa1-null mice, the peripheral nerve extracellular matrix was abnormal: collagen VI in sciatic nerve endoneurium was reduced/disorganized, adjacent Schwann-cell basement membranes showed abnormal focal fusion, and the authors concluded that WARP helps stabilize peripheral nerve matrix architecture. The same mice showed impaired **fine motor coordination** and altered **acute nociceptive response**, despite relatively preserved gross motor function. That supports a role in maintaining neural tissue integrity and function, albeit indirectly through ECM organization. ([pmc.ncbi.nlm.nih.gov](https://pmc.ncbi.nlm.nih.gov/articles/PMC2673271/)) [S3]

Human genetics further strengthens the neural association. In a 2021 Brain study, biallelic VWA1 variants were identified in 17 individuals from 15 families with **autosomal-recessive hereditary motor neuropathy**; the paper also noted high VWA1 expression in tibial nerve and absence of detectable protein from the major founder allele. In zebrafish, VWA1 loss caused reduced **motor-neuron axonal growth**, impaired **synaptic formation** in skeletal muscle, and reduced locomotor behavior. ([pubmed.ncbi.nlm.nih.gov](https://pubmed.ncbi.nlm.nih.gov/33559681/)) [S4]

An independent 2021 Brain paper found bi-allelic truncating VWA1 mutations in 15 affected individuals from six families with **neuromyopathy**, describing VWA1 as an ECM protein expressed in **muscle and peripheral nerve** and noting that lack of VWA1 had already been shown to compromise peripheral nerves in mouse models. ([pubmed.ncbi.nlm.nih.gov](https://pubmed.ncbi.nlm.nih.gov/33459760/?utm_source=openai)) [S5]

More recent clinical aggregation is consistent with that picture. A 2024 cohort/review of VWA1-related disease described a total of **34 individuals** with biallelic variants, further delineating a subtype of hereditary **axonal motor neuropathy** and confirming that VWA1-related pathology can mimic diverse neuromuscular conditions. This does not prove a homeostatic role by itself, but it reinforces that VWA1 loss repeatedly affects neural/motor system function in humans. ([pubmed.ncbi.nlm.nih.gov](https://pubmed.ncbi.nlm.nih.gov/39502942)) [S6]

There is also some evidence for a neural-adjacent vascular maintenance role. In the inner ear, WARP immunoreactivity delineated blood vessels in regions including the spiral and vestibular ganglia, and the authors suggested a role in maintaining vascular integrity there. Because vascular integrity is important for sensory-neural tissue maintenance, this is compatible with a homeostatic interpretation, though still indirect. ([pmc.ncbi.nlm.nih.gov](https://pmc.ncbi.nlm.nih.gov/articles/PMC3005006/?utm_source=openai)) [S7]

## Contradicting or limiting evidence

The main limitation is **how the claim is phrased**. Most of the evidence does **not** show that VWA1 is a direct regulator of neural homeostasis in neurons or glia. Instead, it shows that VWA1 is an **extracellular matrix component** whose loss perturbs peripheral nerve structure, motor axon development, and neuromuscular function. That is biologically relevant to neural maintenance, but it is not the same as demonstrating a primary role in canonical neural regulatory/homeostatic pathways. ([pmc.ncbi.nlm.nih.gov](https://pmc.ncbi.nlm.nih.gov/articles/PMC2673271/)) [S8]

The strongest mechanistic studies are in **mouse peripheral nerve** and **zebrafish developmental models**. There is little direct evidence that VWA1 regulates mature CNS neuronal activity, synaptic scaling, neurotransmitter balance, astrocyte homeostasis, microglial surveillance, or other classic neural homeostatic processes. The 2008 expression study found neural-tissue vasculature and PNS localization, but did not define a neural signaling function. ([pubmed.ncbi.nlm.nih.gov](https://pubmed.ncbi.nlm.nih.gov/18314316/)) [S9]

Some developmental data are even more indirect. In zebrafish craniofacial models, vwa1 loss altered cranial neural crest cell proliferation/condensation and FGF-pathway readouts, but these studies primarily concern **craniofacial chondrogenesis** rather than mature neural-system homeostasis. Neural crest biology is adjacent to neurodevelopment, yet this is not strong evidence for a homeostatic role in neural circuits. ([pmc.ncbi.nlm.nih.gov](https://pmc.ncbi.nlm.nih.gov/articles/PMC10137681/)) [S10]

Finally, some of the “regulatory” interpretation is inferential. For example, the inner-ear vascular paper suggested a role in vascular integrity based on localization, but that is weaker than direct intervention experiments demonstrating homeostatic control. ([pmc.ncbi.nlm.nih.gov](https://pmc.ncbi.nlm.nih.gov/articles/PMC3005006/?utm_source=openai)) [S11]

## Mechanistic / biological rationale

The most plausible mechanism is that VWA1/WARP acts as an **ECM-bridging/stabilizing protein**. Early biochemical work showed that WARP interacts with **perlecan**, and later work showed binding to **collagen VI**. In peripheral nerve, loss of WARP disrupts the relationship between Schwann-cell basement membranes and the surrounding collagenous matrix, producing an unstable endoneurial environment. ([pubmed.ncbi.nlm.nih.gov](https://pubmed.ncbi.nlm.nih.gov/16407285/?utm_source=openai)) [S12]

That mechanism fits known neurobiology: axons, Schwann cells, basement membranes, and pericellular/interstitial matrix form a structural-functional unit in peripheral nerve. If VWA1 helps organize that matrix, then VWA1 loss could plausibly impair **axon extension**, **Schwann-cell/axon separation**, **nerve conduction robustness**, and **neuromuscular junction formation**. The mouse and zebrafish phenotypes are consistent with exactly that sort of maintenance role. ([pmc.ncbi.nlm.nih.gov](https://pmc.ncbi.nlm.nih.gov/articles/PMC2673271/)) [S13]

A broader homeostatic interpretation is therefore biologically reasonable: maintaining extracellular matrix architecture is one way tissues preserve function under normal conditions. But the current evidence suggests VWA1 contributes to neural-system homeostasis **indirectly through extracellular matrix integrity**, especially in the PNS and perhaps specialized neurovascular structures, rather than serving as a well-established neural regulatory factor in its own right. ([pmc.ncbi.nlm.nih.gov](https://pmc.ncbi.nlm.nih.gov/articles/PMC2673271/)) [S14]

## Bottom line

VWA1 is **credibly associated with neural-system biology**, particularly **peripheral nerve structure, motor-neuron/axon development, and inherited motor neuropathy/neuromyopathy**. That supports a role in **maintenance/stabilization** of neural-associated extracellular matrix and possibly specialized neurovascular integrity. But the evidence for VWA1 as a **direct regulator of neural homeostatic or regulatory processes** is still limited. So the claim is **partly supported if interpreted broadly**, but **overstated if interpreted as a well-established direct neural homeostasis regulator**. ([pubmed.ncbi.nlm.nih.gov](https://pubmed.ncbi.nlm.nih.gov/18314316/)) [S15]

## Cited sources

- **[S1]** The extracellular matrix protein WARP is a novel component of a distinct subset of basement membranes - PubMed — https://pubmed.ncbi.nlm.nih.gov/18314316/
- **[S2]** The extracellular matrix protein WARP is a novel component of a distinct subset of basement membranes - PubMed — https://pubmed.ncbi.nlm.nih.gov/18314316/
- **[S3]** 
            Mice Lacking the Extracellular Matrix Protein WARP Develop Normally but Have Compromised Peripheral Nerve Structure and Function - PMC
         — https://pmc.ncbi.nlm.nih.gov/articles/PMC2673271/
- **[S4]** An ancestral 10-bp repeat expansion in VWA1 causes recessive hereditary motor neuropathy - PubMed — https://pubmed.ncbi.nlm.nih.gov/33559681/
- **[S5]** Bi-allelic truncating mutations in VWA1 cause neuromyopathy - PubMed — https://pubmed.ncbi.nlm.nih.gov/33459760/?utm_source=openai
- **[S6]** Autosomal recessive VWA1-related disorder: comprehensive analysis of phenotypic variability and genetic mutations - PubMed — https://pubmed.ncbi.nlm.nih.gov/39502942
- **[S7]** Immunocytochemical distribution of WARP (von Willebrand A domain-related protein) in the inner ear - PMC — https://pmc.ncbi.nlm.nih.gov/articles/PMC3005006/?utm_source=openai
- **[S8]** 
            Mice Lacking the Extracellular Matrix Protein WARP Develop Normally but Have Compromised Peripheral Nerve Structure and Function - PMC
         — https://pmc.ncbi.nlm.nih.gov/articles/PMC2673271/
- **[S9]** The extracellular matrix protein WARP is a novel component of a distinct subset of basement membranes - PubMed — https://pubmed.ncbi.nlm.nih.gov/18314316/
- **[S10]** 
            vwa1 Knockout in Zebrafish Causes Abnormal Craniofacial Chondrogenesis by Regulating FGF Pathway - PMC
         — https://pmc.ncbi.nlm.nih.gov/articles/PMC10137681/
- **[S11]** Immunocytochemical distribution of WARP (von Willebrand A domain-related protein) in the inner ear - PMC — https://pmc.ncbi.nlm.nih.gov/articles/PMC3005006/?utm_source=openai
- **[S12]** WARP is a novel multimeric component of the chondrocyte pericellular matrix that interacts with perlecan - PubMed — https://pubmed.ncbi.nlm.nih.gov/16407285/?utm_source=openai
- **[S13]** 
            Mice Lacking the Extracellular Matrix Protein WARP Develop Normally but Have Compromised Peripheral Nerve Structure and Function - PMC
         — https://pmc.ncbi.nlm.nih.gov/articles/PMC2673271/
- **[S14]** 
            Mice Lacking the Extracellular Matrix Protein WARP Develop Normally but Have Compromised Peripheral Nerve Structure and Function - PMC
         — https://pmc.ncbi.nlm.nih.gov/articles/PMC2673271/
- **[S15]** The extracellular matrix protein WARP is a novel component of a distinct subset of basement membranes - PubMed — https://pubmed.ncbi.nlm.nih.gov/18314316/
