## Supplementary material for "Understanding Substance Dependence: What Differentiates Addictive from Non-Addictive Drugs?": Folder for supplementary files: non-addictive-genes-claims-first100-057-rab11b-gene-protein-has-been-associated-with-regul-727078779331.md

# non-addictive-genes-claims-first100-057-727078779331

**Original claim:** RAB11B gene/protein has been associated with regulatory or homeostatic biological processes in neural systems.

## Claim

**Claim assessed:** *RAB11B gene/protein has been associated with regulatory or homeostatic biological processes in neural systems.*

## Verdict

Verdict: Likely True

RAB11B has credible, direct links to **regulatory functions in neural systems**, especially vesicle trafficking, synaptic vesicle cycling, exocytosis, cortical development, and ion-channel surface regulation in neurons or neuron-related tissues. Human genetics also links damaging **de novo** RAB11B variants to neurodevelopmental disorders with characteristic brain abnormalities. However, the evidence is stronger for **neural regulatory trafficking roles** than for a narrowly defined, experimentally demonstrated role in **neural homeostasis** per se. ([pubmed.ncbi.nlm.nih.gov](https://pubmed.ncbi.nlm.nih.gov/39883192/)) [S1]

## Confidence

**Medium.** Multiple independent lines of evidence support neural relevance of RAB11B, including mouse brain distribution studies, neuronal cell biology, auditory-neuron channel trafficking, and human disease genetics. Confidence is not higher because much of the broader “homeostasis” literature is at the **Rab11 family** level rather than isoform-specific to RAB11B, and explicit homeostasis-focused neural experiments for RAB11B remain limited. ([pubmed.ncbi.nlm.nih.gov](https://pubmed.ncbi.nlm.nih.gov/39883192/)) [S2]

## Supporting evidence

A recent mouse distribution study found that **RAB11B is abundant in brain**, is strongly present in **postmigratory neurons of the developing cortical plate**, remains expressed in most adult neurons, and likely makes **distinct contributions to cortical development and regulation of the synaptic vesicle cycle**. That is direct evidence that RAB11B is associated with regulatory processes in neural tissue, not just with generic housekeeping functions. ([pubmed.ncbi.nlm.nih.gov](https://pubmed.ncbi.nlm.nih.gov/39883192/)) [S3]

Earlier work in neuronal/neuroendocrine PC12 cells showed that **neuronal Rab11b** influences **Ca2+-regulated and constitutive exocytosis**. In the same study, Rab11 was enriched on **purified synaptic vesicles** from brain, alongside canonical synaptic vesicle markers. That places Rab11b in a core regulatory pathway for vesicle release and secretory control in neural cells. ([pubmed.ncbi.nlm.nih.gov](https://pubmed.ncbi.nlm.nih.gov/14627637/)) [S4]

RAB11B has also been tied to regulation of membrane excitability-related proteins in neural systems. In the cochlea, Rab11b physically interacted with the **BK (large-conductance Ca2+-activated K+) channel**, colocalized with BK in **auditory neurons** and related cochlear cells, and altered BK expression in gain- and loss-of-function assays. This supports a role in **ion-channel trafficking and membrane-protein regulation** in a specialized neural circuit. ([pubmed.ncbi.nlm.nih.gov](https://pubmed.ncbi.nlm.nih.gov/22935415/)) [S5]

Human genetics strongly supports neural importance. A 2017 study identified recurrent **de novo RAB11B missense variants** in children with severe intellectual disability, absent speech, epilepsy, hypotonia, and a shared brain MRI phenotype including reduced white matter volume, thin corpus callosum, cerebellar vermis hypoplasia, optic nerve hypoplasia, and mild ventriculomegaly. Functional work in that paper suggested the variants disrupt the GTP/GDP-binding pocket, mislocalize the protein, and alter interactions with regulators/effectors. ([pubmed.ncbi.nlm.nih.gov](https://pubmed.ncbi.nlm.nih.gov/29106825/)) [S6]

That association was reinforced in 2023, when seven additional individuals with **de novo** RAB11B variants were reported. Across the expanded cohort, the main manifestations included **intellectual disability, hypotonia, structural brain anomalies, visual impairment, and epilepsy**, further supporting RAB11B as a bona fide neurodevelopmental disease gene relevant to brain structure and function. ([pubmed.ncbi.nlm.nih.gov](https://pubmed.ncbi.nlm.nih.gov/37734130/)) [S7]

Authoritative databases are consistent with this picture. NCBI Gene lists RAB11B as involved in **exocytotic and endocytotic pathways**, notes an associated condition called **“neurodevelopmental disorder with ataxic gait, absent speech, and decreased cortical white matter,”** and includes Gene Ontology/functional annotations such as **endocytic recycling** and the **constitutive secretory pathway**. While database annotations are not the same as direct mechanistic proof, they align with the primary literature above. ([ncbi.nlm.nih.gov](https://www.ncbi.nlm.nih.gov/gene/9230)) [S8]

## Contradicting or limiting evidence

The claim becomes less certain if “homeostatic” is interpreted narrowly. I did **not** find strong isoform-specific human or mammalian studies showing that RAB11B directly mediates classic neural homeostatic processes such as **homeostatic synaptic scaling, firing-rate homeostasis, or glial/neuronal metabolic homeostasis**. Most direct evidence supports **regulatory trafficking** rather than explicit neural homeostasis. ([pubmed.ncbi.nlm.nih.gov](https://pubmed.ncbi.nlm.nih.gov/14627637/)) [S9]

RAB11B is also **not brain-specific** in current expression resources. The Human Protein Atlas classifies it as having **low tissue specificity** and **low human brain regional specificity**, with substantial expression across multiple non-neural tissues. So the evidence supports neural involvement, but not a uniquely neural protein. ([proteinatlas.org](https://www.proteinatlas.org/Q15907)) [S10]

A further limitation is that much of the neurobiology literature discusses **Rab11** broadly or focuses heavily on **RAB11A**, then extrapolates to family function. A review of Rab11 in neurodegenerative disease explicitly notes that closer examination of **Rab11b knockout** brain phenotypes is still needed and raises the possibility of compensation among family members. That makes some mechanistic inferences about RAB11B in neural homeostasis indirect. ([pmc.ncbi.nlm.nih.gov](https://pmc.ncbi.nlm.nih.gov/articles/PMC9726856/)) [S11]

## Mechanistic / biological rationale

RAB11B encodes a **small GTPase** in the Rab11 family that cycles between GDP- and GTP-bound states to control **recycling endosome trafficking**. In neural cells, this type of machinery is biologically well positioned to regulate the delivery, retrieval, and recycling of membrane proteins that shape synaptic transmission, neurite function, and compartment-specific signaling. ([pmc.ncbi.nlm.nih.gov](https://pmc.ncbi.nlm.nih.gov/articles/PMC9726856/)) [S12]

The neuron-specific data fit that mechanism. In developing and adult mouse brain, RAB11B is present in neurons and is proposed to contribute to **cortical development** and **synaptic vesicle cycle regulation**. In PC12 cells, Rab11b modulates exocytosis; in auditory tissue, it regulates BK-channel expression; and in human disease, pathogenic missense variants behave like functionally impaired Rab11b proteins with altered localization and regulator binding. Together, these findings make biological sense if RAB11B helps maintain proper **vesicular trafficking and membrane-protein composition** in neural cells. ([pubmed.ncbi.nlm.nih.gov](https://pubmed.ncbi.nlm.nih.gov/39883192/)) [S13]

So the strongest mechanistic interpretation is: **RAB11B is part of the intracellular trafficking machinery that regulates neural membrane and vesicle dynamics**, which can secondarily support neural stability and normal brain development. That is compatible with a role in neural homeostasis in a broad sense, but the direct evidence is better for **regulation** than for narrowly defined homeostatic control. ([pmc.ncbi.nlm.nih.gov](https://pmc.ncbi.nlm.nih.gov/articles/PMC9726856/)) [S14]

## Bottom line

RAB11B is **credibly associated with regulatory biological processes in neural systems**, especially neuronal trafficking, exocytosis, synaptic-vesicle-related functions, ion-channel regulation, and brain development. Human pathogenic variants further support neurological relevance. The claim is therefore **Likely True**, with the caveat that direct evidence for **explicit neural homeostasis** is still limited and less developed than the evidence for broader **neural regulatory trafficking functions**. ([pubmed.ncbi.nlm.nih.gov](https://pubmed.ncbi.nlm.nih.gov/39883192/)) [S15]

## Cited sources

- **[S1]** Distribution analysis of RAB11A and RAB11B, small GTP-binding proteins, in mice - PubMed — https://pubmed.ncbi.nlm.nih.gov/39883192/
- **[S2]** Distribution analysis of RAB11A and RAB11B, small GTP-binding proteins, in mice - PubMed — https://pubmed.ncbi.nlm.nih.gov/39883192/
- **[S3]** Distribution analysis of RAB11A and RAB11B, small GTP-binding proteins, in mice - PubMed — https://pubmed.ncbi.nlm.nih.gov/39883192/
- **[S4]** Divergent functions of neuronal Rab11b in Ca2+-regulated versus constitutive exocytosis - PubMed — https://pubmed.ncbi.nlm.nih.gov/14627637/
- **[S5]** The large conductance calcium-activated K(+) channel interacts with the small GTPase Rab11b - PubMed — https://pubmed.ncbi.nlm.nih.gov/22935415/
- **[S6]** Recurrent De Novo Mutations Disturbing the GTP/GDP Binding Pocket of RAB11B Cause Intellectual Disability and a Distinctive Brain Phenotype - PubMed — https://pubmed.ncbi.nlm.nih.gov/29106825/
- **[S7]** De Novo Variants in RAB11B Cause Various Degrees of Global Developmental Delay and Intellectual Disability in Children - PubMed — https://pubmed.ncbi.nlm.nih.gov/37734130/
- **[S8]** RAB11B RAB11B, member RAS oncogene family [Homo sapiens (human)] - Gene - NCBI — https://www.ncbi.nlm.nih.gov/gene/9230
- **[S9]** Divergent functions of neuronal Rab11b in Ca2+-regulated versus constitutive exocytosis - PubMed — https://pubmed.ncbi.nlm.nih.gov/14627637/
- **[S10]** RAB11B protein expression summary - The Human Protein Atlas — https://www.proteinatlas.org/Q15907
- **[S11]** 
            Rab11 and Its Role in Neurodegenerative Diseases - PMC
         — https://pmc.ncbi.nlm.nih.gov/articles/PMC9726856/
- **[S12]** 
            Rab11 and Its Role in Neurodegenerative Diseases - PMC
         — https://pmc.ncbi.nlm.nih.gov/articles/PMC9726856/
- **[S13]** Distribution analysis of RAB11A and RAB11B, small GTP-binding proteins, in mice - PubMed — https://pubmed.ncbi.nlm.nih.gov/39883192/
- **[S14]** 
            Rab11 and Its Role in Neurodegenerative Diseases - PMC
         — https://pmc.ncbi.nlm.nih.gov/articles/PMC9726856/
- **[S15]** Distribution analysis of RAB11A and RAB11B, small GTP-binding proteins, in mice - PubMed — https://pubmed.ncbi.nlm.nih.gov/39883192/
