## Supplementary material for "Understanding Substance Dependence: What Differentiates Addictive from Non-Addictive Drugs?": Folder for supplementary files: non-addictive-genes-claims-first100-058-pde4b-gene-protein-has-been-associated-with-regula-5c5ef9278518.md

# non-addictive-genes-claims-first100-058-5c5ef9278518

**Original claim:** PDE4B gene/protein has been associated with regulatory or homeostatic biological processes in neural systems.

## Claim

**PDE4B gene/protein has been associated with regulatory or homeostatic biological processes in neural systems.**

## Verdict

Verdict: Likely True

## Confidence

**High**

## Supporting evidence

PDE4B is an established **cAMP-specific phosphodiesterase** whose canonical function is to regulate intracellular cyclic nucleotide levels and signal transduction. NCBI’s RefSeq summary states that the encoded protein “regulates the cellular concentrations of cyclic nucleotides and thereby play[s] a role in signal transduction,” and reports broad expression including **brain**. ([ncbi.nlm.nih.gov](https://www.ncbi.nlm.nih.gov/gene/5142)) [S1]

Expression data support relevance to neural tissue rather than a purely peripheral role. In a quantitative survey of human tissues, PDE family mRNAs were measured across **10 brain regions, spinal cord, and dorsal root ganglia**, providing direct evidence that PDE4B is part of the human brain PDE landscape. ([pubmed.ncbi.nlm.nih.gov](https://pubmed.ncbi.nlm.nih.gov/20493887/)) [S2] Human Protein Atlas data further indicate that PDE4B RNA is **detected in all major sampled human brain regions**, including cerebral cortex, hippocampal formation, amygdala, basal ganglia, thalamus, hypothalamus, midbrain, cerebellum, spinal cord, white matter, and choroid plexus. ([proteinatlas.org](https://www.proteinatlas.org/ENSG00000184588-PDE4B/brain)) [S3]

There is also direct evidence that PDE4B participates in **regulation of synaptic plasticity-related signaling**. A primary study reported that **PDE4B3 expression changes during different phases of hippocampal long-term potentiation (LTP)**, a classic cellular model for learning and memory. ([pubmed.ncbi.nlm.nih.gov](https://pubmed.ncbi.nlm.nih.gov/12617967/?utm_source=openai)) [S4] A related study specifically linked **PDE4B and cAMP-level regulation** to LTP in rat hippocampal slices, including changes in a PSD-enriched fraction, tying PDE4B to subcellular control of plasticity-relevant signaling. ([pubmed.ncbi.nlm.nih.gov](https://pubmed.ncbi.nlm.nih.gov/15829230/?utm_source=openai)) [S5] A later review summarizes these data by stating that PDE4B3 was the first cAMP-specific phosphodiesterase associated with **control of LTP stages** in hippocampal neurons. ([pmc.ncbi.nlm.nih.gov](https://pmc.ncbi.nlm.nih.gov/articles/PMC7291338/)) [S6]

Functional perturbation studies reinforce a regulatory neural role. In a mouse model with **reduced PDE4B enzymatic activity**, investigators found increased hippocampal and amygdalar **pCREB/CREB**, and concluded that specific PDE4B inhibition **facilitates memory acquisition** and affects anxiety/fear-related behavior. That paper also notes broad brain distribution and prior links of PDE4B localization/expression to hippocampal plasticity. ([eprints.whiterose.ac.uk](https://eprints.whiterose.ac.uk/id/eprint/89082/8/Specific%20Inhibition%20of%20Phosphodiesterase-4B%20Results%20in%20Anxiolysis%20and%20Facilitates%20Memory%20Acquisition.pdf)) [S7] Conversely, **Pde4b knockout** mice showed **enhanced long-term depression and impaired reversal learning**, indicating that normal PDE4B activity contributes to proper tuning of plasticity and cognitive flexibility rather than simply acting as a generic suppressor. ([pubmed.ncbi.nlm.nih.gov](https://pubmed.ncbi.nlm.nih.gov/21458469/?utm_source=openai)) [S8]

Evidence also supports a role in **glial and neuroimmune regulation**, which is highly relevant to neural homeostasis. In microglia, proinflammatory cytokines increased PDE4 signaling, and PDE4B was observed in **activated but not resting microglia acutely following CNS injury**. ([pubmed.ncbi.nlm.nih.gov](https://pubmed.ncbi.nlm.nih.gov/22865690/)) [S9] The same review literature summarizes that PDE4B is the **primary PDE4 subfamily involved in the LPS response** in microglia/macrophages, that cytokine stimulation lowers microglial cAMP while increasing PDE4B activity, and that **PDE4B2 knockdown or PDE4 inhibition prevents the cAMP drop** after TNF-α stimulation. ([pmc.ncbi.nlm.nih.gov](https://pmc.ncbi.nlm.nih.gov/articles/PMC7291338/)) [S10]

Disease-model studies strengthen this point. In primary rat microglia, **amyloid-β upregulated Pde4B**, increasing TNF-α production; inhibiting PDE4B reduced TNF-α release by up to **70%**, implicating PDE4B directly in regulation of inflammatory output in a CNS-relevant cell type. ([pubmed.ncbi.nlm.nih.gov](https://pubmed.ncbi.nlm.nih.gov/15993984/)) [S11] After spinal cord injury, **Pde4b deletion** reduced inflammatory markers and activated microglia, increased markers associated with mature oligodendrocytes, improved white-matter sparing, and improved locomotor recovery. ([pubmed.ncbi.nlm.nih.gov](https://pubmed.ncbi.nlm.nih.gov/30557659/)) [S12] These findings are consistent with PDE4B influencing neuroinflammatory set-points and repair-related processes that bear on neural homeostasis. ([pubmed.ncbi.nlm.nih.gov](https://pubmed.ncbi.nlm.nih.gov/30557659/)) [S13]

At the molecular level, PDE4B also participates in **regulated cAMP signaling complexes** relevant to neural function. A landmark Science paper found that PDE4B is disrupted in a family with psychiatric illness and showed that **DISC1 interacts with PDE4B**, while elevated cAMP causes dissociation from DISC1 and **increases PDE4B activity**. The authors proposed a model in which DISC1 sequesters PDE4B in resting cells and releases it upon cAMP elevation. ([pubmed.ncbi.nlm.nih.gov](https://pubmed.ncbi.nlm.nih.gov/16293762/)) [S14] Because cAMP signaling is deeply involved in learning, memory, and mood, this is strong mechanistic evidence that PDE4B is part of neural regulatory machinery rather than a passive marker. ([pubmed.ncbi.nlm.nih.gov](https://pubmed.ncbi.nlm.nih.gov/16293762/)) [S15]

## Contradicting or limiting evidence

The main limitation is **semantic and evidentiary**, not that the claim is false. Much of the strongest evidence shows PDE4B in **regulation** of neural cAMP signaling, synaptic plasticity, and neuroinflammatory responses, but fewer studies explicitly frame PDE4B as a mediator of **“homeostasis”** in healthy neural systems. The clearest “homeostatic regulator” language I found for PDE4B comes from **dendritic cells**, not neurons or glia. ([frontiersin.org](https://www.frontiersin.org/journals/pharmacology/articles/10.3389/fphar.2022.833832/full?utm_source=openai)) [S16]

In addition, much of the mechanistic evidence comes from **rodent tissue, knockout models, or cultured glia**, rather than human interventional neuroscience. Human evidence is stronger for **expression** and **genetic/disease association** than for direct causal demonstration in living human neural systems. ([pubmed.ncbi.nlm.nih.gov](https://pubmed.ncbi.nlm.nih.gov/20493887/)) [S17]

The functional phenotype is also **not uniformly one-directional**. Reduced PDE4B activity can facilitate some memory-related outcomes, yet full knockout can impair reversal learning and alter LTD, implying that PDE4B’s role is best understood as **fine-tuning** neural signaling rather than simply promoting or suppressing function. ([eprints.whiterose.ac.uk](https://eprints.whiterose.ac.uk/id/eprint/89082/8/Specific%20Inhibition%20of%20Phosphodiesterase-4B%20Results%20in%20Anxiolysis%20and%20Facilitates%20Memory%20Acquisition.pdf)) [S18]

## Mechanistic / biological rationale

Biologically, the claim is plausible because PDE4B sits at a key control point in **cAMP signaling**, one of the major second-messenger systems in neurons and glia. PDE4B hydrolyzes cAMP, thereby shaping the magnitude, duration, and subcellular localization of downstream **PKA/CREB** signaling. ([ncbi.nlm.nih.gov](https://www.ncbi.nlm.nih.gov/gene/5142)) [S19]

In neurons, that logic fits well with the observed links to **LTP, memory acquisition, and CREB phosphorylation**. If PDE4B activity rises, cAMP signaling is curtailed; if PDE4B activity falls, cAMP-PKA-CREB signaling can be enhanced. The hippocampal LTP studies and PDE4B-mutant mouse data match this framework. ([pubmed.ncbi.nlm.nih.gov](https://pubmed.ncbi.nlm.nih.gov/12617967/?utm_source=openai)) [S20]

In glia, especially microglia, PDE4B appears to regulate how inflammatory stimuli are translated into **cytokine production** and altered cAMP tone. Because microglia help maintain synaptic environment, injury responses, and inflammatory balance in the CNS, PDE4B’s control of glial cAMP signaling provides a credible route by which it contributes to **regulatory and homeostasis-adjacent processes** in neural systems. ([pmc.ncbi.nlm.nih.gov](https://pmc.ncbi.nlm.nih.gov/articles/PMC7291338/)) [S21]

The DISC1 interaction offers an additional layer of regulation: PDE4B is not merely an enzyme floating in the cytosol, but part of **signalosomes/scaffolded complexes** whose activity changes with cellular state. That kind of compartmentalized control is exactly what would be expected for a protein involved in maintaining appropriate neural signaling dynamics. ([pubmed.ncbi.nlm.nih.gov](https://pubmed.ncbi.nlm.nih.gov/16293762/)) [S22]

## Bottom line

The claim is **well supported in its “regulatory” component** and **reasonably supported in a broader homeostasis-related sense**. PDE4B is broadly expressed in the brain, canonically regulates cAMP signal transduction, and has been tied experimentally to hippocampal synaptic plasticity, CREB-linked memory processes, microglial inflammatory regulation, and CNS injury responses. ([ncbi.nlm.nih.gov](https://www.ncbi.nlm.nih.gov/gene/5142)) [S23]

The main caution is that **explicit neural “homeostasis” language is less direct than the evidence for neural regulation**. So the most precise reading is: **PDE4B is clearly associated with regulatory biological processes in neural systems, and there is suggestive but less direct evidence that those roles contribute to neural homeostasis.** ([frontiersin.org](https://www.frontiersin.org/journals/pharmacology/articles/10.3389/fphar.2022.833832/full?utm_source=openai)) [S24]

## Cited sources

- **[S1]** PDE4B phosphodiesterase 4B [Homo sapiens (human)] - Gene - NCBI — https://www.ncbi.nlm.nih.gov/gene/5142
- **[S2]** Quantitative comparison of phosphodiesterase mRNA distribution in human brain and peripheral tissues - PubMed — https://pubmed.ncbi.nlm.nih.gov/20493887/
- **[S3]** Brain tissue expression of PDE4B - Summary - The Human Protein Atlas — https://www.proteinatlas.org/ENSG00000184588-PDE4B/brain
- **[S4]** Expression of the specific type IV phosphodiesterase gene PDE4B3 during different phases of long-term potentiation in single hippocampal slices of rats in vitro - PubMed — https://pubmed.ncbi.nlm.nih.gov/12617967/?utm_source=openai
- **[S5]** Phosphodiesterase 4B (PDE4B) and cAMP-level regulation within different tissue fractions of rat hippocampal slices during long-term potentiation in vitro - PubMed — https://pubmed.ncbi.nlm.nih.gov/15829230/?utm_source=openai
- **[S6]** 
            Phosphodiesterase 4B: Master Regulator of Brain Signaling - PMC
         — https://pmc.ncbi.nlm.nih.gov/articles/PMC7291338/
- **[S7]** Specific Inhibition of Phosphodiesterase-4B Results in Anxiolysis and Facilitates Memory Acquisition — https://eprints.whiterose.ac.uk/id/eprint/89082/8/Specific%20Inhibition%20of%20Phosphodiesterase-4B%20Results%20in%20Anxiolysis%20and%20Facilitates%20Memory%20Acquisition.pdf
- **[S8]** Enhanced long-term depression and impaired reversal learning in phosphodiesterase 4B-knockout (PDE4B-/-) mice - PubMed — https://pubmed.ncbi.nlm.nih.gov/21458469/?utm_source=openai
- **[S9]** Proinflammatory cytokine regulation of cyclic AMP-phosphodiesterase 4 signaling in microglia in vitro and following CNS injury - PubMed — https://pubmed.ncbi.nlm.nih.gov/22865690/
- **[S10]** 
            Phosphodiesterase 4B: Master Regulator of Brain Signaling - PMC
         — https://pmc.ncbi.nlm.nih.gov/articles/PMC7291338/
- **[S11]** The cAMP-specific phosphodiesterase 4B mediates Abeta-induced microglial activation - PubMed — https://pubmed.ncbi.nlm.nih.gov/15993984/
- **[S12]** Following spinal cord injury, PDE4B drives an acute, local inflammatory response and a chronic, systemic response exacerbated by gut dysbiosis and endotoxemia - PubMed — https://pubmed.ncbi.nlm.nih.gov/30557659/
- **[S13]** Following spinal cord injury, PDE4B drives an acute, local inflammatory response and a chronic, systemic response exacerbated by gut dysbiosis and endotoxemia - PubMed — https://pubmed.ncbi.nlm.nih.gov/30557659/
- **[S14]** DISC1 and PDE4B are interacting genetic factors in schizophrenia that regulate cAMP signaling - PubMed — https://pubmed.ncbi.nlm.nih.gov/16293762/
- **[S15]** DISC1 and PDE4B are interacting genetic factors in schizophrenia that regulate cAMP signaling - PubMed — https://pubmed.ncbi.nlm.nih.gov/16293762/
- **[S16]** Frontiers | PDE4B Is a Homeostatic Regulator of Cyclic AMP in Dendritic Cells — https://www.frontiersin.org/journals/pharmacology/articles/10.3389/fphar.2022.833832/full?utm_source=openai
- **[S17]** Quantitative comparison of phosphodiesterase mRNA distribution in human brain and peripheral tissues - PubMed — https://pubmed.ncbi.nlm.nih.gov/20493887/
- **[S18]** Specific Inhibition of Phosphodiesterase-4B Results in Anxiolysis and Facilitates Memory Acquisition — https://eprints.whiterose.ac.uk/id/eprint/89082/8/Specific%20Inhibition%20of%20Phosphodiesterase-4B%20Results%20in%20Anxiolysis%20and%20Facilitates%20Memory%20Acquisition.pdf
- **[S19]** PDE4B phosphodiesterase 4B [Homo sapiens (human)] - Gene - NCBI — https://www.ncbi.nlm.nih.gov/gene/5142
- **[S20]** Expression of the specific type IV phosphodiesterase gene PDE4B3 during different phases of long-term potentiation in single hippocampal slices of rats in vitro - PubMed — https://pubmed.ncbi.nlm.nih.gov/12617967/?utm_source=openai
- **[S21]** 
            Phosphodiesterase 4B: Master Regulator of Brain Signaling - PMC
         — https://pmc.ncbi.nlm.nih.gov/articles/PMC7291338/
- **[S22]** DISC1 and PDE4B are interacting genetic factors in schizophrenia that regulate cAMP signaling - PubMed — https://pubmed.ncbi.nlm.nih.gov/16293762/
- **[S23]** PDE4B phosphodiesterase 4B [Homo sapiens (human)] - Gene - NCBI — https://www.ncbi.nlm.nih.gov/gene/5142
- **[S24]** Frontiers | PDE4B Is a Homeostatic Regulator of Cyclic AMP in Dendritic Cells — https://www.frontiersin.org/journals/pharmacology/articles/10.3389/fphar.2022.833832/full?utm_source=openai
