## Supplementary material for "Understanding Substance Dependence: What Differentiates Addictive from Non-Addictive Drugs?": Folder for supplementary files: non-addictive-genes-claims-first100-059-kcnq2-gene-protein-has-been-associated-with-regula-ec84e392796e.md

# non-addictive-genes-claims-first100-059-ec84e392796e

**Original claim:** KCNQ2 gene/protein has been associated with regulatory or homeostatic biological processes in neural systems.

## Claim

KCNQ2 gene/protein has been associated with regulatory or homeostatic biological processes in neural systems.

## Verdict

Verdict: Highly Supported

## Confidence

High

## Supporting evidence

KCNQ2 is very well established as a neural regulatory gene/protein. The human NCBI Gene record annotates KCNQ2 as involved in **action potential**, **chemical synaptic transmission**, and **nervous system development**, and localizes it to the **axon initial segment**, **node of Ranvier**, plasma membrane, and voltage-gated potassium channel complexes—sites central to neural excitability control. ([ncbi.nlm.nih.gov](https://www.ncbi.nlm.nih.gov/gene/3785)) [S1]

The foundational evidence is strong and direct. In the original Science paper identifying KCNQ2/KCNQ3 as molecular correlates of the neuronal **M-current**, the authors stated that the M-current regulates subthreshold electrical excitability, firing properties, and responsiveness to synaptic input, and concluded that KCNQ2 contributes to this native current. That is a classic neural regulatory function. ([pubmed.ncbi.nlm.nih.gov](https://pubmed.ncbi.nlm.nih.gov/9836639/)) [S2]

Authoritative clinical summaries are consistent with this view. GeneReviews states that KCNQ2-encoded Kv7.2 subunits form potassium channels whose outward current reduces neuronal firing, especially repetitive firing; it also notes that suppression of KCNQ/M-current signaling increases neuronal excitability. MedlinePlus similarly explains that reduced or altered KCNQ2-dependent M-current leads to excessive neuronal excitability and neonatal seizures. ([ncbi.nlm.nih.gov](https://www.ncbi.nlm.nih.gov/books/NBK32534/)) [S3]

Multiple experimental systems show the same biology. In mouse neurons, Kcnq2 deletion or pathogenic loss-of-function variants reduce M-current and increase excitability; for example, ablation in cortical pyramidal neurons lowered rheobase and increased firing, while other mouse work showed reduced M-current density and impaired spike-frequency adaptation. In sympathetic and sensory neurons, KCNQ2 loss also increased excitability. ([pubmed.ncbi.nlm.nih.gov](https://pubmed.ncbi.nlm.nih.gov/28100740/)) [S4]

KCNQ2 is also linked to regulation at the circuit and synaptic level, not just single-cell firing. Presynaptic studies showed that KCNQ2-containing M-channels modulate release of norepinephrine, aspartate, and GABA from hippocampal nerve terminals, supporting a regulatory role in neurotransmission. Localization studies found KCNQ2 at key sites for neuronal network oscillations and synchronization, including axonal domains important for action-potential initiation and propagation. ([pmc.ncbi.nlm.nih.gov](https://pmc.ncbi.nlm.nih.gov/articles/PMC6729253/?utm_source=openai)) [S5]

Evidence for a **homeostatic** role is also present, though somewhat narrower than the excitability-regulation evidence. In PV+ interneuron-specific knockout mice, deletion of KCNQ2/3 or KCNQ2 led to **homeostatic potentiation of excitatory transmission** in pyramidal neurons and increased seizure susceptibility. A neurodevelopmental review likewise summarizes that precise regulation of Kv7.2/M-current is needed to maintain **neuronal network homeostasis**. ([pubmed.ncbi.nlm.nih.gov](https://pubmed.ncbi.nlm.nih.gov/30382937/)) [S6]

Human disease-model work points in the same direction. In iPSC-derived neurons modeling KCNQ2 encephalopathy, impaired M-current was associated with **dyshomeostatic modulation** of other potassium conductances, consistent with KCNQ2 participating in intrinsic excitability homeostasis and in compensatory network responses when perturbed. ([pmc.ncbi.nlm.nih.gov](https://pmc.ncbi.nlm.nih.gov/articles/PMC7864629/)) [S7]

## Contradicting or limiting evidence

The strongest evidence is for **regulation of neuronal excitability**, firing, and synaptic output. The “homeostatic” part of the claim is supported, but less directly and less uniformly. Much of the homeostasis language comes from network-adaptation studies, knockout models, or disease models showing compensatory changes after KCNQ2 dysfunction, rather than from a broad consensus that KCNQ2 is a canonical “homeostasis gene” in the same sense as dedicated homeostatic plasticity pathways. ([pubmed.ncbi.nlm.nih.gov](https://pubmed.ncbi.nlm.nih.gov/30382937/)) [S8]

There are also limits on cell-type generalization. A recent review notes that KCNQ2 expression is clearly neuronal, while evidence for a functional astrocyte role remains uncertain; the authors explicitly state that there is still no proof of a functional Kv7.2 role in astrocytes. So the evidence strongly supports neural regulatory roles in neurons, but not necessarily across all neural cell classes. ([pmc.ncbi.nlm.nih.gov](https://pmc.ncbi.nlm.nih.gov/articles/PMC7657400/)) [S9]

Some ontology annotations in major databases are traceable-author-statement or inferred annotations rather than direct single-experiment proof for every listed process. That does not weaken the core conclusion, but it means each individual annotated process should not be treated as equally direct evidence. ([ncbi.nlm.nih.gov](https://www.ncbi.nlm.nih.gov/gene/3785)) [S10]

## Mechanistic / biological rationale

The biology is coherent. KCNQ2 encodes Kv7.2, a voltage-gated potassium channel subunit that commonly assembles with KCNQ3 to form the neuronal **M-channel/M-current**. This current activates at subthreshold voltages, does not inactivate rapidly, and acts as a brake on depolarization. As a result, it stabilizes membrane potential, limits repetitive firing, shapes spike-frequency adaptation, and alters neuronal responsiveness to synaptic input. ([pubmed.ncbi.nlm.nih.gov](https://pubmed.ncbi.nlm.nih.gov/9836639/)) [S11]

Its subcellular localization fits that function. KCNQ2 is enriched at the **axon initial segment** and **nodes of Ranvier**, where action potentials are initiated and propagated. That positioning makes it well suited to regulate threshold, repetitive spiking, conduction behavior, oscillations, and synchronization. ([ncbi.nlm.nih.gov](https://www.ncbi.nlm.nih.gov/gene/3785)) [S12]

KCNQ2 is itself under physiological regulation by classic signaling pathways. GeneReviews notes dependence on G-protein-coupled receptor signaling; experimental studies show muscarinic signaling inhibits KCNQ current and regulates KCNQ2 phosphorylation via PKC, while calmodulin directly regulates KCNQ2 current and neuronal activity. Structural studies also show KCNQ2 is modulated by PIP2 and therapeutically relevant activators such as retigabine/CBD-related ligands. These features make KCNQ2 both a regulator of neural excitability and a target of broader regulatory networks. ([ncbi.nlm.nih.gov](https://www.ncbi.nlm.nih.gov/books/NBK32534/)) [S13]

## Bottom line

The claim is **well supported**. KCNQ2 is firmly associated with **regulatory processes in neural systems**, especially regulation of membrane excitability, action-potential firing, synaptic transmission, and network synchronization. Evidence for involvement in **homeostatic** neural processes also exists—particularly network and intrinsic-excitability homeostasis in knockout and disease models—but that part is somewhat more indirect and context-dependent than the core excitability-regulation evidence. Overall, the statement is accurate, especially if “regulatory or homeostatic” is interpreted broadly and neuron-focused. ([ncbi.nlm.nih.gov](https://www.ncbi.nlm.nih.gov/gene/3785)) [S14]

## Cited sources

- **[S1]** KCNQ2 potassium voltage-gated channel subfamily Q member 2 [Homo sapiens (human)] - Gene - NCBI — https://www.ncbi.nlm.nih.gov/gene/3785
- **[S2]** KCNQ2 and KCNQ3 potassium channel subunits: molecular correlates of the M-channel - PubMed — https://pubmed.ncbi.nlm.nih.gov/9836639/
- **[S3]** KCNQ2-Related Disorders - GeneReviews® - NCBI Bookshelf — https://www.ncbi.nlm.nih.gov/books/NBK32534/
- **[S4]** Epilepsy-Associated KCNQ2 Channels Regulate Multiple Intrinsic Properties of Layer 2/3 Pyramidal Neurons - PubMed — https://pubmed.ncbi.nlm.nih.gov/28100740/
- **[S5]** M Channels Containing KCNQ2 Subunits Modulate Norepinephrine, Aspartate, and GABA Release from Hippocampal Nerve Terminals - PMC — https://pmc.ncbi.nlm.nih.gov/articles/PMC6729253/?utm_source=openai
- **[S6]** Deletion of KCNQ2/3 potassium channels from PV+ interneurons leads to homeostatic potentiation of excitatory transmission - PubMed — https://pubmed.ncbi.nlm.nih.gov/30382937/
- **[S7]** 
            Dyshomeostatic modulation of Ca2+-activated K+ channels in a human neuronal model of KCNQ2 encephalopathy - PMC
         — https://pmc.ncbi.nlm.nih.gov/articles/PMC7864629/
- **[S8]** Deletion of KCNQ2/3 potassium channels from PV+ interneurons leads to homeostatic potentiation of excitatory transmission - PubMed — https://pubmed.ncbi.nlm.nih.gov/30382937/
- **[S9]** 
            The Role of Kv7.2 in Neurodevelopment: Insights and Gaps in Our Understanding - PMC
         — https://pmc.ncbi.nlm.nih.gov/articles/PMC7657400/
- **[S10]** KCNQ2 potassium voltage-gated channel subfamily Q member 2 [Homo sapiens (human)] - Gene - NCBI — https://www.ncbi.nlm.nih.gov/gene/3785
- **[S11]** KCNQ2 and KCNQ3 potassium channel subunits: molecular correlates of the M-channel - PubMed — https://pubmed.ncbi.nlm.nih.gov/9836639/
- **[S12]** KCNQ2 potassium voltage-gated channel subfamily Q member 2 [Homo sapiens (human)] - Gene - NCBI — https://www.ncbi.nlm.nih.gov/gene/3785
- **[S13]** KCNQ2-Related Disorders - GeneReviews® - NCBI Bookshelf — https://www.ncbi.nlm.nih.gov/books/NBK32534/
- **[S14]** KCNQ2 potassium voltage-gated channel subfamily Q member 2 [Homo sapiens (human)] - Gene - NCBI — https://www.ncbi.nlm.nih.gov/gene/3785
