## Supplementary material for "Understanding Substance Dependence: What Differentiates Addictive from Non-Addictive Drugs?": Folder for supplementary files: non-addictive-genes-claims-first100-060-ptk2b-gene-protein-has-been-associated-with-regula-9531d7be16ac.md

# non-addictive-genes-claims-first100-060-9531d7be16ac

**Original claim:** PTK2B gene/protein has been associated with regulatory or homeostatic biological processes in neural systems.

## Claim

PTK2B gene/protein has been associated with regulatory or homeostatic biological processes in neural systems.

## Verdict

Verdict: Likely True

PTK2B (encoding the kinase Pyk2) is well associated with **neural regulatory processes**, especially calcium-dependent signaling, synaptic transmission/plasticity, postsynaptic organization, and activity-dependent regulation of ion channels and receptors. The evidence for a specifically **homeostatic** role is more limited and often indirect, but the broader “regulatory or homeostatic” wording is supported overall. ([ncbi.nlm.nih.gov](https://www.ncbi.nlm.nih.gov/gene/2185)) [S1]

## Confidence

High.

Confidence is high because the association is supported by multiple independent lines of evidence: curated gene/protein resources, synapse-focused ontology annotations, and several primary neuroscience studies in hippocampal and striatal systems. The main reason this is not rated higher is that “homeostatic” is less directly established than “regulatory,” especially in healthy human neural tissue. ([ncbi.nlm.nih.gov](https://www.ncbi.nlm.nih.gov/gene/2185)) [S2]

## Supporting evidence

- **Authoritative gene resources link PTK2B to neuronal regulation.** NCBI’s human PTK2B summary states that the gene encodes a cytoplasmic tyrosine kinase involved in **calcium-induced regulation of ion channels** and MAPK signaling, and may act as an intermediate between neurotransmitter/neuropeptide signaling and downstream pathways that regulate **neuronal activity**. The original 1995 Nature paper similarly reported that PYK2 is highly expressed in the CNS and activated by rises in intracellular Ca²⁺, with effects on ion-channel function and MAPK signaling. ([ncbi.nlm.nih.gov](https://www.ncbi.nlm.nih.gov/gene/2185)) [S3]

- **PTK2B is clearly situated in brain and synaptic contexts.** The Human Protein Atlas classifies PTK2B in a brain RNA expression cluster labeled **“Neurons - Mixed function (mainly)”**, with detection across all major brain regions. SynGO places PTK2B at the **postsynapse**, **postsynaptic density**, and **presynapse**, and annotates it to biological processes including **postsynaptic modulation of chemical synaptic transmission** and **regulation of postsynaptic density assembly**. NCBI/GO also lists PTK2B as part of the **NMDA selective glutamate receptor complex** and involved in positive regulation of glutamatergic synaptic transmission. ([proteinatlas.org](https://www.proteinatlas.org/ENSG00000120899-PTK2B/brain)) [S4]

- **Primary studies show PTK2B/Pyk2 regulates hippocampal synaptic plasticity.** In a 2001 *Neuron* study, Pyk2 acted upstream of Src to enhance NMDA receptor function in CA1 neurons, and blocking Pyk2 prevented induction of long-term potentiation (LTP). A 2010 *Journal of Neuroscience* paper then showed that Pyk2 also regulates hippocampal **long-term depression (LTD)**. Together, these are classic neural regulatory processes rather than incidental associations. ([pubmed.ncbi.nlm.nih.gov](https://pubmed.ncbi.nlm.nih.gov/11239437/)) [S5]

- **Genetic loss-of-function studies support a regulatory role in synapse structure and function.** In 2017, Pyk2-deficient mice showed impaired hippocampal-dependent memory and CA1 LTP, along with changes in **NMDA receptors, PSD-95, and dendritic spines**. The same work concluded that Pyk2 has essential roles in spine structure and synaptic function. This strongly supports PTK2B as a regulator of neural circuit properties, especially at excitatory synapses. ([pubmed.ncbi.nlm.nih.gov](https://pubmed.ncbi.nlm.nih.gov/28555636/)) [S6]

- **Regulatory effects extend beyond hippocampal CA1 to other neural systems.** A 2021 study in striatum found that Pyk2 stabilizes dendrites and spines of medium spiny neurons, and that Pyk2 loss disrupts behavioral flexibility. That is consistent with a role in maintaining structural/synaptic integrity of neural circuits, not just acute signaling. ([pubmed.ncbi.nlm.nih.gov](https://pubmed.ncbi.nlm.nih.gov/34943950/)) [S7]

- **Mechanistic receptor/channel studies fit a neural regulatory role.** Pyk2 has been shown to mediate receptor-driven potentiation of NMDA receptor currents in cortical neurons via the Pyk2/Src pathway, and more recently to participate in α1-adrenergic receptor → PKC → Pyk2 → Src signaling that enhances **CaV1.2 L-type calcium channel activity** and promotes a form of hippocampal LTP. These findings place PTK2B directly in the regulation of neuronal excitability and synaptic strength. ([pmc.ncbi.nlm.nih.gov](https://pmc.ncbi.nlm.nih.gov/articles/PMC6758206/)) [S8]

- **There is also suggestive evidence for roles related to intracellular calcium homeostasis.** In hippocampal neurons, Pyk2 has been implicated in regulating ER-mitochondria contact sites (MAMs) and calcium handling, with authors proposing that Pyk2 is important for calcium regulation in a physiological neuronal context. This is not definitive proof of a broad “homeostatic” role, but it supports the idea that PTK2B contributes to cellular processes that help stabilize neuronal signaling states. ([pmc.ncbi.nlm.nih.gov](https://pmc.ncbi.nlm.nih.gov/articles/PMC8909471/)) [S9]

## Contradicting or limiting evidence

- **“Homeostatic” is less directly supported than “regulatory.”** The strongest evidence is for PTK2B in **synaptic regulation/plasticity**, ion-channel modulation, receptor signaling, and maintenance of synaptic structures. Direct demonstrations in canonical neural **homeostatic plasticity** paradigms are comparatively sparse. Some papers discuss “synaptic homeostasis” or calcium regulation, but often in disease contexts rather than normal physiological homeostasis. ([pmc.ncbi.nlm.nih.gov](https://pmc.ncbi.nlm.nih.gov/articles/PMC8909471/)) [S10]

- **Much of the strongest evidence is from rodents or cultured neurons, not direct human functional experiments.** Human evidence is strongest for expression, ontology placement, and disease association; causal functional evidence in neural systems comes mainly from mouse knockouts, slice electrophysiology, and neuronal cultures. ([proteinatlas.org](https://www.proteinatlas.org/ENSG00000120899-PTK2B/brain)) [S11]

- **Some functional effects appear context- or protocol-dependent.** For example, a later study reported that CA1 LTP in Pyk2 knockout mice was impaired with a high-frequency stimulation protocol but not with theta-burst stimulation, suggesting that PTK2B’s role in plasticity is real but not uniform across all induction conditions. ([pubmed.ncbi.nlm.nih.gov](https://pubmed.ncbi.nlm.nih.gov/34381140/?utm_source=openai)) [S12]

- **PTK2B is not neuron-specific.** NCBI notes broad tissue expression, and PTK2B has substantial non-neural roles in immune, vascular, and adhesion-related signaling. So while it is clearly associated with neural regulation, it is not a dedicated neural-homeostasis gene in the narrow sense. ([ncbi.nlm.nih.gov](https://www.ncbi.nlm.nih.gov/gene/2185)) [S13]

## Mechanistic / biological rationale

Biologically, PTK2B/Pyk2 is a good fit for a neural regulatory molecule because it is a **Ca²⁺-responsive non-receptor tyrosine kinase** activated by membrane depolarization, nicotinic acetylcholine receptor stimulation, PKC activation, and other conditions that raise intracellular calcium. Once activated, it can engage **Src-family kinases**, modulate **NMDA receptor** and **voltage-gated calcium channel** function, influence **MAPK** signaling, and alter postsynaptic structural elements such as **PSD-95** and dendritic spines. SynGO/GO annotations placing PTK2B at pre/postsynaptic sites and in postsynaptic modulation are consistent with these experimental data. This makes PTK2B mechanistically plausible as a regulator of neural signal integration, synaptic strength, and aspects of structural stability; a broader homeostatic role remains plausible but less directly established. ([pubmed.ncbi.nlm.nih.gov](https://pubmed.ncbi.nlm.nih.gov/7544443/)) [S14]

## Bottom line

The claim is **likely true**. PTK2B/Pyk2 is convincingly associated with **regulatory biological processes in neural systems**—notably calcium-dependent control of ion channels, NMDA-receptor signaling, LTP/LTD, postsynaptic organization, and dendritic spine/neurite stability. The evidence for a specifically **homeostatic** role exists but is more limited and indirect than the evidence for synaptic and signaling regulation. ([ncbi.nlm.nih.gov](https://www.ncbi.nlm.nih.gov/gene/2185)) [S15]

## Cited sources

- **[S1]** PTK2B protein tyrosine kinase 2 beta [Homo sapiens (human)] - Gene - NCBI — https://www.ncbi.nlm.nih.gov/gene/2185
- **[S2]** PTK2B protein tyrosine kinase 2 beta [Homo sapiens (human)] - Gene - NCBI — https://www.ncbi.nlm.nih.gov/gene/2185
- **[S3]** PTK2B protein tyrosine kinase 2 beta [Homo sapiens (human)] - Gene - NCBI — https://www.ncbi.nlm.nih.gov/gene/2185
- **[S4]** Brain tissue expression of PTK2B - Summary - The Human Protein Atlas — https://www.proteinatlas.org/ENSG00000120899-PTK2B/brain
- **[S5]** CAKbeta/Pyk2 kinase is a signaling link for induction of long-term potentiation in CA1 hippocampus - PubMed — https://pubmed.ncbi.nlm.nih.gov/11239437/
- **[S6]** Pyk2 modulates hippocampal excitatory synapses and contributes to cognitive deficits in a Huntington's disease model - PubMed — https://pubmed.ncbi.nlm.nih.gov/28555636/
- **[S7]** Pyk2 Stabilizes Striatal Medium Spiny Neuron Structure and Striatal-Dependent Action - PubMed — https://pubmed.ncbi.nlm.nih.gov/34943950/
- **[S8]** 
            Metabotropic Glutamate Receptor 1-Induced Upregulation of NMDA Receptor Current: Mediation through the Pyk2/Src-Family Kinase Pathway in Cortical Neurons - PMC
         — https://pmc.ncbi.nlm.nih.gov/articles/PMC6758206/
- **[S9]** 
            Pyk2 Regulates MAMs and Mitochondrial Dynamics in Hippocampal Neurons - PMC
         — https://pmc.ncbi.nlm.nih.gov/articles/PMC8909471/
- **[S10]** 
            Pyk2 Regulates MAMs and Mitochondrial Dynamics in Hippocampal Neurons - PMC
         — https://pmc.ncbi.nlm.nih.gov/articles/PMC8909471/
- **[S11]** Brain tissue expression of PTK2B - Summary - The Human Protein Atlas — https://www.proteinatlas.org/ENSG00000120899-PTK2B/brain
- **[S12]** Pyk2 in dorsal hippocampus plays a selective role in spatial memory and synaptic plasticity - PubMed — https://pubmed.ncbi.nlm.nih.gov/34381140/?utm_source=openai
- **[S13]** PTK2B protein tyrosine kinase 2 beta [Homo sapiens (human)] - Gene - NCBI — https://www.ncbi.nlm.nih.gov/gene/2185
- **[S14]** Protein tyrosine kinase PYK2 involved in Ca(2+)-induced regulation of ion channel and MAP kinase functions - PubMed — https://pubmed.ncbi.nlm.nih.gov/7544443/
- **[S15]** PTK2B protein tyrosine kinase 2 beta [Homo sapiens (human)] - Gene - NCBI — https://www.ncbi.nlm.nih.gov/gene/2185
