## Supplementary material for "Understanding Substance Dependence: What Differentiates Addictive from Non-Addictive Drugs?": Folder for supplementary files: non-addictive-genes-claims-first100-061-pacsin1-gene-protein-has-been-associated-with-regu-fea2ea82ec5e.md

# non-addictive-genes-claims-first100-061-fea2ea82ec5e

**Original claim:** PACSIN1 gene/protein has been associated with regulatory or homeostatic biological processes in neural systems.

## Claim

PACSIN1 gene/protein has been associated with regulatory or homeostatic biological processes in neural systems.

## Verdict

Verdict: Highly Supported

## Confidence

High

## Supporting evidence

PACSIN1 is well established in the neuroscience literature as a neuron-enriched membrane-remodeling/endocytic adaptor involved in **regulation** of multiple neural processes, especially receptor trafficking, synaptic vesicle recycling, synaptic plasticity, and neuronal morphogenesis. A recent review summarized that, in the CNS, PACSIN1 regulates neuromorphogenesis, receptor trafficking, glutamatergic neurotransmission, and synaptic plasticity, and concluded that its function is important for dendritic-tree organization and efficient synaptic signaling. Rodent gene annotations are concordant, listing roles such as positive regulation of dendrite development and regulation of postsynaptic neurotransmitter receptor internalization. ([pmc.ncbi.nlm.nih.gov](https://pmc.ncbi.nlm.nih.gov/articles/PMC9285741/)) [S1]

On the **presynaptic/regulatory** side, PACSIN1 has strong evidence for controlling synaptic vesicle recycling. A lamprey synapse study found that perturbing syndapin/PACSIN impairs vesicle recycling during intense stimulation and proposed a role in bulk endocytosis at high release rates. In mouse, syndapin I/PACSIN1 knockout caused abnormalities in synaptic vesicle formation, reduced excitatory and inhibitory synaptic transmission, altered hippocampal network activity, and spontaneous generalized seizures. Together, these are classic examples of a protein participating in neural regulatory machinery that stabilizes synaptic function under activity load. ([pmc.ncbi.nlm.nih.gov](https://pmc.ncbi.nlm.nih.gov/articles/PMC6670478/?utm_source=openai)) [S2]

On the **postsynaptic/regulatory** side, PACSIN1 directly regulates glutamate receptor trafficking. For AMPA receptors, PACSIN1 was shown to control NMDA receptor-triggered GluA2 internalization and recycling in hippocampal neurons, linking endocytic and recycling machineries needed for dynamic AMPAR trafficking; PACSIN-PICK1 signaling was also required for activity-dependent AMPAR internalization and cerebellar long-term depression. For NMDA receptors, PACSIN1/syndapin1 mediates endocytosis and synaptic removal of NR3A-containing NMDARs, with evidence that this helps regulate receptor abundance/subunit composition during synaptic maturation. These findings squarely place PACSIN1 in neural regulatory processes governing synaptic composition and plasticity. ([nature.com](https://www.nature.com/articles/srep31070)) [S3]

There is also meaningful evidence for a **homeostatic/inhibitory-balance** connection. In the KCC2 interactome study, KCC2 was described as essential for establishing the neuronal chloride gradient required for hyperpolarizing inhibition. PACSIN1 was identified as a major KCC2 interactor, and functional experiments showed that reducing PACSIN1 increased KCC2 expression and shifted neuronal chloride handling toward stronger inhibition, while PACSIN1 overexpression reduced KCC2. Because KCC2 is central to chloride homeostasis and inhibitory synaptic stability, this provides direct evidence that PACSIN1 is associated not just with “regulation” broadly, but with a bona fide homeostasis-relevant neural process. ([pmc.ncbi.nlm.nih.gov](https://pmc.ncbi.nlm.nih.gov/articles/PMC5640428/)) [S4]

PACSIN1 is also implicated in **structural regulation** of neurons. CK2-dependent phosphorylation of PACSIN1 promotes Rac1-dependent dendritic spine formation and is described by the authors as required for dendritic spine adaptation in synaptic plasticity. Separately, PACSIN1 binds Tau and regulates axonal elongation/branching by modulating microtubule dynamics. These are additional regulatory roles in neural development and maintenance rather than isolated biochemical associations. ([pmc.ncbi.nlm.nih.gov](https://pmc.ncbi.nlm.nih.gov/articles/PMC3611001/)) [S5]

## Contradicting or limiting evidence

The main limitation is that most direct evidence comes from **cellular and animal models**—cultured hippocampal neurons, mouse knockouts, lamprey synapses, and biochemical interactome studies—rather than from human in vivo physiology or clinical intervention studies. The human-disease relevance is plausible and in some cases suggested by proteomic/genetic observations, but the strongest evidence for PACSIN1’s functions remains experimental rather than clinical. ([pmc.ncbi.nlm.nih.gov](https://pmc.ncbi.nlm.nih.gov/articles/PMC9285741/)) [S6]

A second limitation is that the word **“homeostatic”** is somewhat broader than what many PACSIN1 papers explicitly test. The literature more directly demonstrates roles in receptor trafficking, endocytosis, synaptic plasticity, inhibitory strength, and neuronal morphology. Calling these “regulatory” is straightforward; calling them “homeostatic” is best supported when referring to chloride/inhibitory balance via KCC2, and somewhat more inferential when referring to synaptic balance in general. That inference is reasonable, but it is still an inference from mechanistic data rather than a large literature specifically labeling PACSIN1 as a canonical “homeostatic plasticity” factor. ([pmc.ncbi.nlm.nih.gov](https://pmc.ncbi.nlm.nih.gov/articles/PMC5640428/)) [S7]

There is also some **context dependence** across neural systems. For example, in the Drosophila larval neuromuscular junction literature summarized in the PACSIN review, syndapin was reported to be dispensable for presynaptic synaptic vesicle endocytosis and instead localized postsynaptically. So PACSIN-family functions are not identical across all species/synapse types, which cautions against overgeneralizing any one mechanism. ([pmc.ncbi.nlm.nih.gov](https://pmc.ncbi.nlm.nih.gov/articles/PMC9285741/)) [S8]

## Mechanistic / biological rationale

Mechanistically, PACSIN1 is well suited to regulate neural homeostasis-like processes because it combines **membrane-shaping** and **protein-scaffolding** functions. PACSIN1 contains an F-BAR domain and an SH3 domain; the F-BAR domain is associated with membrane binding/bending, while the SH3 domain supports interactions with partners such as dynamin and actin regulators. In AMPAR trafficking experiments, the F-BAR and SH3 domains were both required for NMDAR-dependent receptor internalization/recycling, consistent with PACSIN1 acting as a membrane-deformation adaptor that couples endocytosis to cytoskeletal remodeling. ([ncbi.nlm.nih.gov](https://www.ncbi.nlm.nih.gov/gene/29993)) [S9]

The same logic extends to synaptic stability. By controlling endocytosis/recycling of AMPARs and NMDARs, PACSIN1 can influence synaptic strength and receptor composition. By regulating KCC2 abundance, it can modulate chloride extrusion and therefore the efficacy of inhibitory transmission. By participating in dendritic spine formation through CK2/BDNF/Rac1 signaling, and in axonal microtubule dynamics via Tau, it also affects the structural substrate on which synaptic regulation occurs. In short, PACSIN1 sits at an intersection of membrane traffic, cytoskeletal control, receptor availability, and inhibitory set-point regulation—exactly the kinds of mechanisms expected for neural regulatory and partially homeostatic functions. ([nature.com](https://www.nature.com/articles/srep31070)) [S10]

## Bottom line

The claim is **accurate**. PACSIN1 is strongly associated with **regulatory biological processes in neural systems**, including synaptic vesicle recycling, glutamate receptor trafficking, synaptic plasticity, inhibitory-strength regulation via KCC2, and neuronal morphogenesis. The **homeostatic** part of the claim is also supportable, especially through PACSIN1’s regulation of KCC2/chloride-dependent inhibition, though that label is somewhat broader and less directly tested than its receptor-trafficking and synaptic-regulation roles. Overall, the evidence is strong enough for **Highly Supported**. ([pmc.ncbi.nlm.nih.gov](https://pmc.ncbi.nlm.nih.gov/articles/PMC9285741/)) [S11]

## Cited sources

- **[S1]** 
            PACSIN proteins in vivo: Roles in development and physiology - PMC
         — https://pmc.ncbi.nlm.nih.gov/articles/PMC9285741/
- **[S2]** Perturbation of Syndapin/PACSIN Impairs Synaptic Vesicle Recycling Evoked by Intense Stimulation - PMC — https://pmc.ncbi.nlm.nih.gov/articles/PMC6670478/?utm_source=openai
- **[S3]** PACSIN1 regulates the dynamics of AMPA receptor trafficking | Scientific Reports — https://www.nature.com/articles/srep31070
- **[S4]** 
            Native KCC2 interactome reveals PACSIN1 as a critical regulator of synaptic inhibition - PMC
         — https://pmc.ncbi.nlm.nih.gov/articles/PMC5640428/
- **[S5]** 
            Casein Kinase 2 Phosphorylation of Protein Kinase C and Casein Kinase 2 Substrate in Neurons (PACSIN) 1 Protein Regulates Neuronal Spine Formation - PMC
         — https://pmc.ncbi.nlm.nih.gov/articles/PMC3611001/
- **[S6]** 
            PACSIN proteins in vivo: Roles in development and physiology - PMC
         — https://pmc.ncbi.nlm.nih.gov/articles/PMC9285741/
- **[S7]** 
            Native KCC2 interactome reveals PACSIN1 as a critical regulator of synaptic inhibition - PMC
         — https://pmc.ncbi.nlm.nih.gov/articles/PMC5640428/
- **[S8]** 
            PACSIN proteins in vivo: Roles in development and physiology - PMC
         — https://pmc.ncbi.nlm.nih.gov/articles/PMC9285741/
- **[S9]** PACSIN1 protein kinase C and casein kinase substrate in neurons 1 [Homo sapiens (human)] - Gene - NCBI — https://www.ncbi.nlm.nih.gov/gene/29993
- **[S10]** PACSIN1 regulates the dynamics of AMPA receptor trafficking | Scientific Reports — https://www.nature.com/articles/srep31070
- **[S11]** 
            PACSIN proteins in vivo: Roles in development and physiology - PMC
         — https://pmc.ncbi.nlm.nih.gov/articles/PMC9285741/
