## Supplementary material for "Understanding Substance Dependence: What Differentiates Addictive from Non-Addictive Drugs?": Folder for supplementary files: non-addictive-genes-claims-first100-062-scn1b-gene-protein-has-been-associated-with-regula-f43a9d66ecc0.md

# non-addictive-genes-claims-first100-062-f43a9d66ecc0

**Original claim:** SCN1B gene/protein has been associated with regulatory or homeostatic biological processes in neural systems.

## Claim

SCN1B gene/protein has been associated with regulatory or homeostatic biological processes in neural systems. ([ncbi.nlm.nih.gov](https://www.ncbi.nlm.nih.gov/gene/6324)) [S1]

## Verdict

Verdict: Highly Supported

## Confidence

High

## Supporting evidence

SCN1B encodes the **sodium channel regulatory subunit beta-1** (β1/β1B), and authoritative gene resources already annotate it to regulatory neural-related processes, including **regulation of membrane potential**, **neuronal action potential propagation**, and **positive regulation of neuron projection development**. Those annotations alone are not the strongest evidence, but they are consistent with a large experimental literature. ([amigo.geneontology.org](https://amigo.geneontology.org/amigo/gene_product/UniProtKB%3AQ07699)) [S2]

The strongest support comes from experimental neuroscience studies showing that SCN1B regulates **neuronal excitability**. In dorsal root ganglion neurons, Scn1b deletion altered sodium and potassium currents and made neurons hyperexcitable; the authors concluded that Scn1b regulates electrical excitability in vivo. ([pubmed.ncbi.nlm.nih.gov](https://pubmed.ncbi.nlm.nih.gov/21555511/)) [S3]

SCN1B also regulates **brain circuit development and synaptic organization**, not just acute ion-channel behavior. In Scn1b-null mice, β1 loss altered dentate granule cell proliferation/localization, caused hippocampal and cerebellar pathfinding abnormalities, and produced age-dependent changes in synaptic transmission and excitability; the authors concluded that Scn1b is required for establishment of normal electrical activity in the brain and plays an important role in development of neuronal synaptic circuits and excitability in the CNS. ([pmc.ncbi.nlm.nih.gov](https://pmc.ncbi.nlm.nih.gov/articles/PMC3549092/)) [S4]

Independent work showed that β1-mediated signaling promotes **neurite outgrowth** and contributes to **postnatal CNS development**. In vivo, Scn1b-null mice had defective cerebellar granule neuron axon extension/fasciculation and abnormal corticospinal tract pathfinding, supporting a regulatory role in neural patterning and microorganization. ([pubmed.ncbi.nlm.nih.gov](https://pubmed.ncbi.nlm.nih.gov/18354028/)) [S5]

A recent synthesis of the epilepsy/channelopathy literature states that β1/β1B subunits are developmentally regulated Ig-CAMs and ion-channel modulators that play critical roles in **regulation of excitability**, and that their functions include modulation of sodium, potassium, and calcium channels, regulation of gene expression after proteolytic cleavage, and regulation of cell adhesion and neuronal patterning. The same review summarizes altered excitability, altered axon pathfinding, altered α-subunit expression, and delayed maturation of GABAergic signaling in Scn1b-null brains. ([ncbi.nlm.nih.gov](https://www.ncbi.nlm.nih.gov/books/NBK609884/)) [S6]

Evidence also extends from single-cell physiology to **network-level regulation**. A 2023 study found that Scn1b deletion reduced sodium-current heterogeneity between pyramidal neurons, increased spike-timing correlations, and altered the coupling requirements for network synchronization, concluding that sodium-current heterogeneity regulated by Scn1b is a potent regulator of spike-pattern diversity and network function. ([nature.com](https://www.nature.com/articles/s41598-023-36036-0)) [S7]

Finally, disease genetics is consistent with biological importance in neural regulation. Reviews summarize that SCN1B variants are linked to developmental and epileptic encephalopathies, and a recent JCI Insight study notes that Scn1b-null mice model SCN1B-linked Dravet syndrome with seizures, altered pathfinding, and altered Purkinje/granule cell excitability. ([ncbi.nlm.nih.gov](https://www.ncbi.nlm.nih.gov/books/NBK609884/)) [S8]

## Contradicting or limiting evidence

The main limitation is **specificity**: the evidence strongly supports SCN1B in **regulatory** neural processes, but support for a narrowly defined role in **homeostatic** neural biology is more indirect. I did not find strong primary evidence placing SCN1B as a canonical mediator of classic “homeostatic plasticity” frameworks in the same way that term is often used in systems neuroscience. Instead, the literature mainly supports roles in excitability control, developmental patterning, synaptic function, and network synchronization. ([ncbi.nlm.nih.gov](https://www.ncbi.nlm.nih.gov/books/NBK609884/)) [S9]

The phenotype is also **cell-type and context dependent**. Reviews note that many Scn1b-null neuronal populations show altered excitability, but some do not; dentate granule neurons, CA1 pyramidal neurons, and some hippocampal interneurons can show little measurable excitability change under the tested conditions. Effects can differ by neuron type, developmental stage, and whether the model is full knockout versus specific disease variant. ([pmc.ncbi.nlm.nih.gov](https://pmc.ncbi.nlm.nih.gov/articles/PMC5856584/)) [S10]

There is also a **human-evidence gap** for fine-grained biological-process claims. Human genetics strongly links SCN1B to epilepsy syndromes, but much of the mechanistic evidence for “regulatory processes in neural systems” comes from mouse, slice, or cellular studies rather than direct human neurophysiology. ([ncbi.nlm.nih.gov](https://www.ncbi.nlm.nih.gov/books/NBK609884/)) [S11]

## Mechanistic / biological rationale

SCN1B is well positioned mechanistically to influence neural regulatory processes because β1 is both an **auxiliary ion-channel regulator** and an **immunoglobulin superfamily cell-adhesion molecule**. It modulates voltage-gated sodium-channel trafficking and gating, can influence potassium and calcium channel function, and can undergo regulated intramembrane proteolysis that contributes to downstream gene-expression control. These are classic mechanisms by which neurons regulate excitability and circuit assembly. ([ncbi.nlm.nih.gov](https://www.ncbi.nlm.nih.gov/books/NBK609884/)) [S12]

At the subcellular level, β1 is enriched at the **axon initial segment** in many neurons, where action-potential initiation is controlled. Loss of Scn1b changes the complement of α-subunits at this domain, such as reduced Nav1.6 and compensatory increases in Nav1.1 in cerebellar granule neurons, supporting a role in fine-tuning excitability to cell- and compartment-specific requirements. ([ncbi.nlm.nih.gov](https://www.ncbi.nlm.nih.gov/books/NBK609884/)) [S13]

Beyond acute firing, β1-mediated adhesion/signaling helps regulate **neurite outgrowth, axon fasciculation, migration, and neuronal patterning**. Those developmental processes can secondarily shape synaptic connectivity and excitability. Indeed, abnormal pathfinding appears early in Scn1b-null mice, before overt epileptiform activity in some paradigms. ([pubmed.ncbi.nlm.nih.gov](https://pubmed.ncbi.nlm.nih.gov/18354028/)) [S14]

There is also a plausible bridge to a broader “homeostatic” interpretation: Scn1b affects GABAergic maturation and network synchronization, and network models suggest it helps determine the dynamic range over which neuronal populations resist synchronization. That is relevant to neural stability, though it is still better described as **regulation of excitability and circuit stability** than as established homeostatic plasticity per se. ([ncbi.nlm.nih.gov](https://www.ncbi.nlm.nih.gov/books/NBK609884/)) [S15]

## Bottom line

The claim is **well supported**, especially if “regulatory” is interpreted broadly. SCN1B/β1 has strong experimental associations with **regulation of neuronal excitability, membrane potential-related processes, neurite outgrowth, axon pathfinding, synaptic circuit development, GABAergic maturation, and network synchronization** in neural systems. ([ncbi.nlm.nih.gov](https://www.ncbi.nlm.nih.gov/gene/6324)) [S16]

The only important caveat is wording: the evidence is **strongest for neural regulatory roles**, whereas a specific role in **canonical homeostatic neural processes** is less directly established. So the claim is best read as true primarily because SCN1B is clearly associated with **regulatory** biological processes in neural systems. ([ncbi.nlm.nih.gov](https://www.ncbi.nlm.nih.gov/books/NBK609884/)) [S17]

## Cited sources

- **[S1]** SCN1B sodium voltage-gated channel beta subunit 1 [Homo sapiens (human)] - Gene - NCBI — https://www.ncbi.nlm.nih.gov/gene/6324
- **[S2]** AmiGO 2: Gene Product Details for UniProtKB:Q07699 — https://amigo.geneontology.org/amigo/gene_product/UniProtKB%3AQ07699
- **[S3]** Na+ channel Scn1b gene regulates dorsal root ganglion nociceptor excitability in vivo - PubMed — https://pubmed.ncbi.nlm.nih.gov/21555511/
- **[S4]** 
            Abnormal neuronal patterning occurs during early postnatal brain development of Scn1b-null mice and precedes hyperexcitability - PMC
         — https://pmc.ncbi.nlm.nih.gov/articles/PMC3549092/
- **[S5]** Voltage-gated Na+ channel beta1 subunit-mediated neurite outgrowth requires Fyn kinase and contributes to postnatal CNS development in vivo - PubMed — https://pubmed.ncbi.nlm.nih.gov/18354028/
- **[S6]** Sodium Channelopathies in Human and Animal Models of Epilepsy and Neurodevelopmental Disorders - Jasper's Basic Mechanisms of the Epilepsies - NCBI Bookshelf — https://www.ncbi.nlm.nih.gov/books/NBK609884/
- **[S7]** Heterogeneity of voltage gated sodium current density between neurons decorrelates spiking and suppresses network synchronization in Scn1b null mouse models | Scientific Reports — https://www.nature.com/articles/s41598-023-36036-0
- **[S8]** Sodium Channelopathies in Human and Animal Models of Epilepsy and Neurodevelopmental Disorders - Jasper's Basic Mechanisms of the Epilepsies - NCBI Bookshelf — https://www.ncbi.nlm.nih.gov/books/NBK609884/
- **[S9]** Sodium Channelopathies in Human and Animal Models of Epilepsy and Neurodevelopmental Disorders - Jasper's Basic Mechanisms of the Epilepsies - NCBI Bookshelf — https://www.ncbi.nlm.nih.gov/books/NBK609884/
- **[S10]** 
            Voltage-gated sodium channel β subunits: The power outside the pore in brain development and disease - PMC
         — https://pmc.ncbi.nlm.nih.gov/articles/PMC5856584/
- **[S11]** Sodium Channelopathies in Human and Animal Models of Epilepsy and Neurodevelopmental Disorders - Jasper's Basic Mechanisms of the Epilepsies - NCBI Bookshelf — https://www.ncbi.nlm.nih.gov/books/NBK609884/
- **[S12]** Sodium Channelopathies in Human and Animal Models of Epilepsy and Neurodevelopmental Disorders - Jasper's Basic Mechanisms of the Epilepsies - NCBI Bookshelf — https://www.ncbi.nlm.nih.gov/books/NBK609884/
- **[S13]** Sodium Channelopathies in Human and Animal Models of Epilepsy and Neurodevelopmental Disorders - Jasper's Basic Mechanisms of the Epilepsies - NCBI Bookshelf — https://www.ncbi.nlm.nih.gov/books/NBK609884/
- **[S14]** Voltage-gated Na+ channel beta1 subunit-mediated neurite outgrowth requires Fyn kinase and contributes to postnatal CNS development in vivo - PubMed — https://pubmed.ncbi.nlm.nih.gov/18354028/
- **[S15]** Sodium Channelopathies in Human and Animal Models of Epilepsy and Neurodevelopmental Disorders - Jasper's Basic Mechanisms of the Epilepsies - NCBI Bookshelf — https://www.ncbi.nlm.nih.gov/books/NBK609884/
- **[S16]** SCN1B sodium voltage-gated channel beta subunit 1 [Homo sapiens (human)] - Gene - NCBI — https://www.ncbi.nlm.nih.gov/gene/6324
- **[S17]** Sodium Channelopathies in Human and Animal Models of Epilepsy and Neurodevelopmental Disorders - Jasper's Basic Mechanisms of the Epilepsies - NCBI Bookshelf — https://www.ncbi.nlm.nih.gov/books/NBK609884/
