## Supplementary material for "Understanding Substance Dependence: What Differentiates Addictive from Non-Addictive Drugs?": Folder for supplementary files: non-addictive-genes-claims-first100-063-pam-gene-protein-has-been-associated-with-regulato-8399ca238eac.md

# non-addictive-genes-claims-first100-063-8399ca238eac

**Original claim:** PAM gene/protein has been associated with regulatory or homeostatic biological processes in neural systems.

## Claim

PAM gene/protein has been associated with regulatory or homeostatic biological processes in neural systems. PAM here refers to **peptidylglycine α-amidating monooxygenase**, the conserved enzyme encoded by the **PAM** gene. ([pmc.ncbi.nlm.nih.gov](https://pmc.ncbi.nlm.nih.gov/articles/PMC9177522/)) [S1]

## Verdict

Verdict: Highly Supported

Multiple lines of evidence support this claim. PAM is broadly expressed in the nervous system, is required for maturation of many amidated neuropeptides, and perturbing PAM in animal models changes synaptic transmission, neuronal copper homeostasis, and behaviors linked to neural regulation such as anxiety, fear conditioning, seizure sensitivity, and thermoregulation. ([pubmed.ncbi.nlm.nih.gov](https://pubmed.ncbi.nlm.nih.gov/1729435/)) [S2]

## Confidence

High

The evidence base is strong at the **molecular**, **cellular**, and **animal physiology/behavior** levels. The main caveat is that direct **human CNS causal evidence** is much thinner than the rodent and mechanistic literature. ([pubmed.ncbi.nlm.nih.gov](https://pubmed.ncbi.nlm.nih.gov/20648645/)) [S3]

## Supporting evidence

PAM is a core neuropeptide-processing enzyme, not a peripheral incidental factor. It is the **only known enzyme** that catalyzes peptide C-terminal α-amidation, and this modification is often necessary for peptide stability, receptor binding, and full bioactivity. Reviews further note that mice lacking PAM die by mid-gestation, underscoring its biological importance. ([pmc.ncbi.nlm.nih.gov](https://pmc.ncbi.nlm.nih.gov/articles/PMC9177522/)) [S4]

PAM is clearly present in neural tissue. A classic rat brain study found **PAM mRNA and immunoreactivity in all major brain areas except cerebellum**, with particularly high expression in hypothalamic magnocellular neurons, hippocampus, and olfactory cortex. The authors also noted that regions rich in amidated neuropeptides showed high PAM expression, linking PAM to peptidergic neural regulation. ([pubmed.ncbi.nlm.nih.gov](https://pubmed.ncbi.nlm.nih.gov/1729435/)) [S5]

The connection to neural regulation is strengthened by the fact that PAM is essential for synthesis of **amidated neuropeptides**, and many such peptides participate in physiological regulation. A review focused on nervous-system function concluded that reduced PAM expression and mild copper deficiency produce similar abnormalities in mice in processes known to be neuropeptide-regulated, including **thermal regulation, seizure sensitivity, and anxiety-like behavior**. ([pubmed.ncbi.nlm.nih.gov](https://pubmed.ncbi.nlm.nih.gov/20648645/)) [S6]

There is also direct evidence that PAM influences synaptic physiology. In **Pam+/−** mice, the amygdala shows abnormal emotional behavior together with altered synaptic properties: anxiety-like behavior, impaired contextual and cued fear conditioning, deficient long-term potentiation at thalamic afferent synapses, and altered excitatory and GABAergic signaling. Importantly, PAM protein was detected in both pyramidal neurons and local GABAergic interneurons in the basolateral amygdala, tying the phenotype to neural circuit function rather than only systemic endocrine effects. ([pubmed.ncbi.nlm.nih.gov](https://pubmed.ncbi.nlm.nih.gov/20943906/)) [S7]

PAM has also been linked specifically to **neural homeostasis of copper**, a biologically meaningful homeostatic process because copper is required for normal synaptic function. In a region-specific mouse study, Pam heterozygosity caused selective copper deficiency in the amygdala, reduced expression of copper-handling genes such as **Atox1** and **Atp7a**, and altered catecholamine metabolism. The authors explicitly concluded that this demonstrates a role for PAM, beyond amidation, in **regulating copper homeostasis in brain**. ([pubmed.ncbi.nlm.nih.gov](https://pubmed.ncbi.nlm.nih.gov/24032518/)) [S8]

Cell-type-specific genetics provides additional support. Deleting Pam in **excitatory forebrain neurons** altered behaviors dependent on cortical/hippocampal/amygdalar circuits, including novel object recognition, Barnes maze search strategy, and social behavior. The study also noted that without PAM, excitatory neurons in amygdala and hippocampus cannot generate biologically active forms of several amidated transmitters/neuromodulators, including hypocretins, PACAP, neuropeptide Y, and cholecystokinin. ([pmc.ncbi.nlm.nih.gov](https://pmc.ncbi.nlm.nih.gov/articles/PMC7858240/)) [S9]

Taken together, these data show much more than mere expression. They connect PAM to **regulatory neural signaling**, **synaptic plasticity**, and at least one explicit neural **homeostatic** process: copper handling in defined brain regions. ([pubmed.ncbi.nlm.nih.gov](https://pubmed.ncbi.nlm.nih.gov/20943906/)) [S10]

## Contradicting or limiting evidence

The strongest limitation is translational rather than conceptual. Most of the decisive evidence comes from **rodent models**, neuroendocrine cell systems, and biochemical studies, not from direct human interventional or neuropathological studies. ([pubmed.ncbi.nlm.nih.gov](https://pubmed.ncbi.nlm.nih.gov/1729435/)) [S11]

Direct human neural data do exist but are limited and mostly associative. For example, one older study reported increased PAM activity in cerebrospinal fluid from patients with multiple sclerosis, but that does not establish a general causal role for PAM in neural homeostasis across humans. ([pubmed.ncbi.nlm.nih.gov](https://pubmed.ncbi.nlm.nih.gov/7606087/?utm_source=openai)) [S12]

Also, some of the “regulatory/homeostatic” inference is indirect. PAM often acts upstream by enabling maturation of many different amidated peptides, so some observed neural phenotypes may reflect combined downstream effects of altered neuropeptide signaling rather than a single dedicated “homeostasis pathway.” That still supports association with neural regulatory processes, but it limits pathway-level specificity. ([pubmed.ncbi.nlm.nih.gov](https://pubmed.ncbi.nlm.nih.gov/20648645/)) [S13]

Finally, part of the mechanistic literature concerns **neuroendocrine** or non-neural cells, secretory granules, hypoxia responses, or cilia. Those findings are biologically relevant and plausible, but not all of them demonstrate a nervous-system-specific homeostatic role on their own. ([pmc.ncbi.nlm.nih.gov](https://pmc.ncbi.nlm.nih.gov/articles/PMC30969/)) [S14]

## Mechanistic / biological rationale

The core mechanism is straightforward: PAM catalyzes the two-step conversion of glycine-extended peptide precursors into **α-amidated bioactive peptides**. Because amidation can increase receptor affinity, reduce pH sensitivity, and improve peptide stability, loss or reduction of PAM predictably weakens signaling by many neuropeptides. That alone makes PAM highly relevant to neural regulation. ([pmc.ncbi.nlm.nih.gov](https://pmc.ncbi.nlm.nih.gov/articles/PMC9177522/)) [S15]

This is especially important because numerous amidated neuropeptides participate in neural and systemic homeostatic control. A recent study on hypoxia states explicitly that many peptides involved in **homeostatic control** require PAM for biosynthesis, and it further shows that physiologic hypoxia can inhibit PAM monooxygenase activity and that PAM expression can alter gene-expression programs. ([pubmed.ncbi.nlm.nih.gov](https://pubmed.ncbi.nlm.nih.gov/34061983/)) [S16]

PAM also appears to have functions beyond its catalytic luminal domains. Its **cytosolic domain** participates in trafficking and signaling. In neuroendocrine cells, PAM influences regulated secretion, secretory granule maturation, and communication between luminal conditions and cytosolic trafficking machinery; a soluble cytosolic fragment can even reach the nucleus and affect gene expression. Those mechanisms make it biologically plausible that PAM modulates neuronal secretory pathways and metal-handling programs, not just peptide chemistry. ([pmc.ncbi.nlm.nih.gov](https://pmc.ncbi.nlm.nih.gov/articles/PMC30969/)) [S17]

The copper link is mechanistically compelling. PAM is a **copper-dependent cuproenzyme**, and its relationship with ATP7A/Atox1 provides a credible route by which altered PAM levels could disturb copper delivery in neurons, affecting synaptic function and plasticity. The amygdala studies fit this model well: altered PAM dosage, altered copper-handling gene expression, altered local copper status, and altered synaptic physiology all co-occur. ([pubmed.ncbi.nlm.nih.gov](https://pubmed.ncbi.nlm.nih.gov/20648645/)) [S18]

## Bottom line

Yes. The claim is **highly supported**: PAM is strongly associated with **regulatory and homeostatic biology in neural systems**, especially through its indispensable role in amidated neuropeptide production and through evidence linking PAM to **synaptic transmission**, **emotional/fear circuitry**, **thermoregulatory and seizure-related phenotypes**, and **brain copper homeostasis**. The main qualifier is that the clearest causal data come from animal and mechanistic studies rather than extensive human CNS studies. ([pubmed.ncbi.nlm.nih.gov](https://pubmed.ncbi.nlm.nih.gov/20648645/)) [S19]

## Cited sources

- **[S1]** 
            Peptidylglycine α-amidating monooxygenase as a therapeutic target or biomarker - PMC
         — https://pmc.ncbi.nlm.nih.gov/articles/PMC9177522/
- **[S2]** Expression of peptidylglycine alpha-amidating monooxygenase (EC 1.14.17.3) in the rat central nervous system - PubMed — https://pubmed.ncbi.nlm.nih.gov/1729435/
- **[S3]** Peptidylgycine α-amidating monooxygenase and copper: a gene-nutrient interaction critical to nervous system function - PubMed — https://pubmed.ncbi.nlm.nih.gov/20648645/
- **[S4]** 
            Peptidylglycine α-amidating monooxygenase as a therapeutic target or biomarker - PMC
         — https://pmc.ncbi.nlm.nih.gov/articles/PMC9177522/
- **[S5]** Expression of peptidylglycine alpha-amidating monooxygenase (EC 1.14.17.3) in the rat central nervous system - PubMed — https://pubmed.ncbi.nlm.nih.gov/1729435/
- **[S6]** Peptidylgycine α-amidating monooxygenase and copper: a gene-nutrient interaction critical to nervous system function - PubMed — https://pubmed.ncbi.nlm.nih.gov/20648645/
- **[S7]** Haploinsufficiency in peptidylglycine alpha-amidating monooxygenase leads to altered synaptic transmission in the amygdala and impaired emotional responses - PubMed — https://pubmed.ncbi.nlm.nih.gov/20943906/
- **[S8]** Peptidylglycine α-amidating monooxygenase heterozygosity alters brain copper handling with region specificity - PubMed — https://pubmed.ncbi.nlm.nih.gov/24032518/
- **[S9]** 
            Cell-type specific knockout of peptidylglycine α-amidating monooxygenase reveals specific behavioral roles in excitatory forebrain neurons and cardiomyocytes - PMC
         — https://pmc.ncbi.nlm.nih.gov/articles/PMC7858240/
- **[S10]** Haploinsufficiency in peptidylglycine alpha-amidating monooxygenase leads to altered synaptic transmission in the amygdala and impaired emotional responses - PubMed — https://pubmed.ncbi.nlm.nih.gov/20943906/
- **[S11]** Expression of peptidylglycine alpha-amidating monooxygenase (EC 1.14.17.3) in the rat central nervous system - PubMed — https://pubmed.ncbi.nlm.nih.gov/1729435/
- **[S12]** Increased peptidylglycine alpha-amidating monooxygenase activity in cerebrospinal fluid of patients with multiple sclerosis - PubMed — https://pubmed.ncbi.nlm.nih.gov/7606087/?utm_source=openai
- **[S13]** Peptidylgycine α-amidating monooxygenase and copper: a gene-nutrient interaction critical to nervous system function - PubMed — https://pubmed.ncbi.nlm.nih.gov/20648645/
- **[S14]** 
            Signaling Mediated by the Cytosolic Domain of Peptidylglycine α-Amidating Monooxygenase - PMC
         — https://pmc.ncbi.nlm.nih.gov/articles/PMC30969/
- **[S15]** 
            Peptidylglycine α-amidating monooxygenase as a therapeutic target or biomarker - PMC
         — https://pmc.ncbi.nlm.nih.gov/articles/PMC9177522/
- **[S16]** Multiple roles for peptidylglycine α-amidating monooxygenase in the response to hypoxia - PubMed — https://pubmed.ncbi.nlm.nih.gov/34061983/
- **[S17]** 
            Signaling Mediated by the Cytosolic Domain of Peptidylglycine α-Amidating Monooxygenase - PMC
         — https://pmc.ncbi.nlm.nih.gov/articles/PMC30969/
- **[S18]** Peptidylgycine α-amidating monooxygenase and copper: a gene-nutrient interaction critical to nervous system function - PubMed — https://pubmed.ncbi.nlm.nih.gov/20648645/
- **[S19]** Peptidylgycine α-amidating monooxygenase and copper: a gene-nutrient interaction critical to nervous system function - PubMed — https://pubmed.ncbi.nlm.nih.gov/20648645/
