## Supplementary material for "Understanding Substance Dependence: What Differentiates Addictive from Non-Addictive Drugs?": Folder for supplementary files: non-addictive-genes-claims-first100-064-cers1-gene-protein-has-been-associated-with-regula-18c2cad2d4bc.md

# non-addictive-genes-claims-first100-064-18c2cad2d4bc

**Original claim:** CERS1 gene/protein has been associated with regulatory or homeostatic biological processes in neural systems.

## Claim

CERS1 gene/protein has been associated with regulatory or homeostatic biological processes in neural systems.

## Verdict

Verdict: Likely True

## Confidence

High

## Supporting evidence

- **CERS1/CerS1 is strongly enriched in the nervous system, especially neurons.** A major review of mammalian ceramide synthases reports that CerS1 is mainly expressed in brain and, within brain tissue, in most neurons with little to no expression in white matter cells; it is also slightly upregulated postnatally, consistent with a role in neuronal membrane/sphingolipid biology. Independent mouse work found CerS1 immunosignals in neurons but not other brain cell types. ([pmc.ncbi.nlm.nih.gov](https://pmc.ncbi.nlm.nih.gov/articles/PMC2858252/)) [S1]

- **Loss of CERS1 disrupts brain sphingolipid steady state and causes neurodegeneration in mice.** In the flincher/toppler mouse mutants, loss of CerS1 activity reduced brain sphingolipid biosynthesis, caused major changes in steady-state sphingolipids and sphingoid bases, and led to cerebellar ataxia, Purkinje cell degeneration, lipofuscin accumulation, and ubiquitinated protein deposits. The authors explicitly concluded that loss of CerS1 “dramatically impacts many aspects of sphingolipid homeostasis,” which is direct evidence tying CERS1 to neural lipid/protein homeostasis. ([pubmed.ncbi.nlm.nih.gov](https://pubmed.ncbi.nlm.nih.gov/21625621/?utm_source=openai)) [S2]

- **A second mouse knockout model linked neuronal CerS1 to developmental regulation and neural function.** In Cers1-deficient mice, investigators found cerebellar foliation defects, progressive cerebellar shrinkage, neuronal apoptosis, and marked reductions in gangliosides and myelin-associated glycoprotein in cerebellum and forebrain. Behavioral deficits included impaired exploration, locomotion, and motor coordination. The paper concluded that CerS1-derived ceramide has an essential function in the **regulation of cerebellar development and neurodevelopmentally regulated behavior**. ([pubmed.ncbi.nlm.nih.gov](https://pubmed.ncbi.nlm.nih.gov/23074226/)) [S3]

- **Human genetics supports clinical relevance in neural systems.** A homozygous CERS1 mutation was identified in siblings with progressive myoclonus epilepsy and dementia; the mutation decreased C18-ceramide levels, and CerS1 downregulation in a neuroblastoma line triggered ER-stress and pro-apoptotic pathways. Additional case reports linked CERS1 deficiency to progressive myoclonic epilepsy with prominent ataxia. This supports an association between CERS1 dysfunction and failure of neural cellular stability/survival in humans. ([pubmed.ncbi.nlm.nih.gov](https://pubmed.ncbi.nlm.nih.gov/24782409/)) [S4]

- **Rescue experiments argue that CerS1 is tied to neuronal homeostasis, not just a disease marker.** In Cers1-deficient mice, ectopic neuronal expression of CerS2 almost completely suppressed pathology even though it did **not** restore C18 ceramide; instead, it normalized long-chain bases (LCBs). The study concluded that high LCB accumulation is strongly associated with neuronal death, implying that CerS1 normally contributes to neuronal survival by maintaining sphingolipid metabolite balance. ([pubmed.ncbi.nlm.nih.gov](https://pubmed.ncbi.nlm.nih.gov/27162368/)) [S5]

- **More recent work connects CerS1 to mitochondrial quality-control pathways in brain.** A 2019 study showed that cellular stress drives trafficking of CerS1 from the ER to the outer mitochondrial membrane via p17/PERMIT, producing mitochondrial ceramide and inducing mitophagy; genetic ablation of p17/PERMIT reduced this response in brain tissue. A 2023 study extended this to neurons and cerebellum in vivo: CerS1-dependent mitophagy shaped brain metabolite responses, and impaired CerS1/p17/PERMIT/PARKIN signaling led to damaged-mitochondria accumulation and age-dependent sensorimotor abnormalities. That is a plausible and fairly direct form of neural homeostatic regulation. ([pubmed.ncbi.nlm.nih.gov](https://pubmed.ncbi.nlm.nih.gov/31535025/)) [S6]

## Contradicting or limiting evidence

- **The claim is broad, while the evidence is more specific.** The strongest data support roles in **neuronal sphingolipid balance, mitochondrial quality control, cerebellar development, and neuronal survival**. There are fewer studies explicitly framing CERS1 as a general “homeostatic regulator” across neural systems in normal physiology. Much of the evidence comes from knockout mutants, rare Mendelian disease, or stress paradigms rather than routine baseline homeostasis in healthy humans. ([pmc.ncbi.nlm.nih.gov](https://pmc.ncbi.nlm.nih.gov/articles/PMC3098191/?utm_source=openai)) [S7]

- **Mechanistic attribution remains partly unresolved.** The CerS2-rescue study showed that pathology could be largely corrected without restoring neuronal C18 ceramide itself, suggesting that the harmful effect of CERS1 loss may be driven substantially by buildup of upstream long-chain bases rather than simply loss of one CerS1-made lipid species. So the exact homeostatic variable controlled by CERS1 is still being worked out. ([pubmed.ncbi.nlm.nih.gov](https://pubmed.ncbi.nlm.nih.gov/27162368/)) [S8]

- **Some mechanistic studies are indirect for neural physiology.** Important work on CerS1 turnover, stress signaling, and mitophagy used cancer or transformed cell systems, which are useful mechanistically but do not perfectly model mature neurons in vivo. Likewise, a recent review notes that understanding of sphingolipid regulation in physiology and disease is still incomplete. ([pubmed.ncbi.nlm.nih.gov](https://pubmed.ncbi.nlm.nih.gov/19393694/)) [S9]

## Mechanistic / biological rationale

- **Biochemically, CERS1 encodes ceramide synthase 1, the major neuronal enzyme for making C18 ceramide.** Because ceramide is the backbone for many complex sphingolipids, neuron-enriched CerS1 is well positioned to regulate membrane composition, signaling, and organelle function in neurons. ([pmc.ncbi.nlm.nih.gov](https://pmc.ncbi.nlm.nih.gov/articles/PMC2858252/)) [S10]

- **At the membrane level, CerS1 helps maintain neuronal sphingolipid composition.** In Cers1-deficient mice, gangliosides fell markedly and myelin-associated glycoprotein expression dropped, implying that CerS1-derived lipids help stabilize neuron–oligodendrocyte interactions and support cerebellar structure/function. This is consistent with a homeostatic role in preserving neural membrane organization and intercellular support. ([pubmed.ncbi.nlm.nih.gov](https://pubmed.ncbi.nlm.nih.gov/23074226/)) [S11]

- **At the metabolite level, CerS1 appears to buffer potentially toxic long-chain base accumulation.** When CerS1 is absent, long-chain bases rise; restoring their levels rescues neuronal pathology even without restoring the normal C18-ceramide profile. That pattern argues that CerS1 participates in metabolic homeostasis needed for neurite integrity and neuronal survival. ([pubmed.ncbi.nlm.nih.gov](https://pubmed.ncbi.nlm.nih.gov/27162368/)) [S12]

- **At the organelle level, CerS1 participates in stress-responsive mitophagy.** Under stress, CerS1 can be trafficked from ER to mitochondria, where CerS1-generated ceramide helps recruit mitophagic machinery. In brain/cerebellum models, failure of this pathway was linked to damaged-mitochondria accumulation and age-related sensorimotor defects, fitting a mitochondrial quality-control/homeostasis model. ([pubmed.ncbi.nlm.nih.gov](https://pubmed.ncbi.nlm.nih.gov/31535025/)) [S13]

- **CERS1 itself is regulated by signaling pathways, reinforcing its placement in regulatory networks.** CerS1 undergoes rapid proteasomal turnover and is controlled by p38 MAPK and PKC, with direct consequences for ceramide homeostasis. While these data are not neuron-specific, they provide a mechanistic basis for how CerS1 levels could be dynamically regulated in stressed neural cells. ([pubmed.ncbi.nlm.nih.gov](https://pubmed.ncbi.nlm.nih.gov/19393694/)) [S14]

## Bottom line

The claim is **likely true**. CERS1 is not just a generic lipid enzyme; the literature links it to **neuron-enriched sphingolipid regulation, maintenance of metabolite balance, cerebellar development, neuronal survival, and mitochondrial quality-control pathways** in brain-relevant models and rare human disease. The evidence is strongest for **neural lipid/metabolic homeostasis and developmental regulation**, and somewhat less direct for a broad systems-level “homeostatic” role across all neural contexts. ([pmc.ncbi.nlm.nih.gov](https://pmc.ncbi.nlm.nih.gov/articles/PMC3098191/?utm_source=openai)) [S15]

## Cited sources

- **[S1]** 
            Mammalian Ceramide Synthases - PMC
         — https://pmc.ncbi.nlm.nih.gov/articles/PMC2858252/
- **[S2]** A deficiency of ceramide biosynthesis causes cerebellar purkinje cell neurodegeneration and lipofuscin accumulation - PubMed — https://pubmed.ncbi.nlm.nih.gov/21625621/?utm_source=openai
- **[S3]** Ablation of neuronal ceramide synthase 1 in mice decreases ganglioside levels and expression of myelin-associated glycoprotein in oligodendrocytes - PubMed — https://pubmed.ncbi.nlm.nih.gov/23074226/
- **[S4]** Impairment of ceramide synthesis causes a novel progressive myoclonus epilepsy - PubMed — https://pubmed.ncbi.nlm.nih.gov/24782409/
- **[S5]** Ectopic expression of ceramide synthase 2 in neurons suppresses neurodegeneration induced by ceramide synthase 1 deficiency - PubMed — https://pubmed.ncbi.nlm.nih.gov/27162368/
- **[S6]** Mitochondrial protein import is regulated by p17/PERMIT to mediate lipid metabolism and cellular stress - PubMed — https://pubmed.ncbi.nlm.nih.gov/31535025/
- **[S7]** A Deficiency of Ceramide Biosynthesis Causes Cerebellar Purkinje Cell Neurodegeneration and Lipofuscin Accumulation - PMC — https://pmc.ncbi.nlm.nih.gov/articles/PMC3098191/?utm_source=openai
- **[S8]** Ectopic expression of ceramide synthase 2 in neurons suppresses neurodegeneration induced by ceramide synthase 1 deficiency - PubMed — https://pubmed.ncbi.nlm.nih.gov/27162368/
- **[S9]** Ceramide synthase 1 is regulated by proteasomal mediated turnover - PubMed — https://pubmed.ncbi.nlm.nih.gov/19393694/
- **[S10]** 
            Mammalian Ceramide Synthases - PMC
         — https://pmc.ncbi.nlm.nih.gov/articles/PMC2858252/
- **[S11]** Ablation of neuronal ceramide synthase 1 in mice decreases ganglioside levels and expression of myelin-associated glycoprotein in oligodendrocytes - PubMed — https://pubmed.ncbi.nlm.nih.gov/23074226/
- **[S12]** Ectopic expression of ceramide synthase 2 in neurons suppresses neurodegeneration induced by ceramide synthase 1 deficiency - PubMed — https://pubmed.ncbi.nlm.nih.gov/27162368/
- **[S13]** Mitochondrial protein import is regulated by p17/PERMIT to mediate lipid metabolism and cellular stress - PubMed — https://pubmed.ncbi.nlm.nih.gov/31535025/
- **[S14]** Ceramide synthase 1 is regulated by proteasomal mediated turnover - PubMed — https://pubmed.ncbi.nlm.nih.gov/19393694/
- **[S15]** A Deficiency of Ceramide Biosynthesis Causes Cerebellar Purkinje Cell Neurodegeneration and Lipofuscin Accumulation - PMC — https://pmc.ncbi.nlm.nih.gov/articles/PMC3098191/?utm_source=openai
