## Supplementary material for "Understanding Substance Dependence: What Differentiates Addictive from Non-Addictive Drugs?": Folder for supplementary files: non-addictive-genes-claims-first100-065-ca11-gene-protein-has-been-associated-with-regulat-e3699aca4bb8.md

# non-addictive-genes-claims-first100-065-e3699aca4bb8

**Original claim:** CA11 gene/protein has been associated with regulatory or homeostatic biological processes in neural systems.

## Claim

CA11 gene/protein has been associated with regulatory or homeostatic biological processes in neural systems.

## Verdict

Verdict: Likely True

CA11 is fairly well associated with **neural regulatory biology**, especially through its strong CNS-enriched expression, developmental expression in neuroprogenitors and migrating neurons, and its identification as a **neurexin-associated synaptic protein**. However, evidence for a **direct, well-established homeostatic role** in healthy neural circuits is still limited. So the broad claim is supportable, but mainly on the **regulatory/synaptic** side rather than on a firmly demonstrated neural homeostasis role. ([pubmed.ncbi.nlm.nih.gov](https://pubmed.ncbi.nlm.nih.gov/10350627/)) [S1]

## Confidence

Medium

The evidence base is real but uneven: CA11 is reproducibly linked to the brain and synaptic/neurexin biology, yet its precise physiological function remains incompletely defined, and much of the strongest mechanistic work has focused more directly on the closely related protein CA10 than on CA11 itself. ([pubmed.ncbi.nlm.nih.gov](https://pubmed.ncbi.nlm.nih.gov/28154140/)) [S2]

## Supporting evidence

Human CA11 was originally reported to be **strongly expressed in brain**, especially in the **cerebellum, cerebral cortex, and putamen**, and the authors concluded this pattern suggests a general role in the human CNS. Independent modern expression resources also classify CA11 as **brain-enriched** and place it in a brain RNA-expression cluster labeled **“synaptic signal transduction.”** ([pubmed.ncbi.nlm.nih.gov](https://pubmed.ncbi.nlm.nih.gov/10350627/)) [S3]

Developmental neuroanatomy studies strengthen that association. In human brain tissue, CA11 protein was found in **neuronal cell bodies, neurites, and astrocytes** in selected brain regions; in fetal brain it was present in **neuroprogenitor cells in the subventricular zone** and later in **migrating neural cells** moving toward cortex. Those are classic contexts of neural regulation and organization, even if they do not by themselves prove a specific molecular function. ([pubmed.ncbi.nlm.nih.gov](https://pubmed.ncbi.nlm.nih.gov/12044474/)) [S4]

The strongest mechanistic association is with **neurexins**, major presynaptic organizers of synapse formation and function. In mouse brain, proteomic isolation of neurexin-1 complexes identified **CA10 and CA11** as co-associated proteins. The Sterky et al. study then showed direct neurexin binding for CA10 and argued that **CA10 and, by extension, CA11** act as endogenous neurexin ligands; a later review summarized CA10/CA11 as neurexin-binding partners whose **cis interactions in the secretory pathway enable neurexin trafficking**. Because neurexins are central regulators of synaptic connectivity and signaling, this places CA11 within a plausible neural regulatory network. ([pubmed.ncbi.nlm.nih.gov](https://pubmed.ncbi.nlm.nih.gov/28154140/)) [S5]

Authoritative mouse annotation databases also summarize Car11 in explicitly regulatory neural terms. The Jackson Laboratory/Alliance-derived GO summary for mouse **Car11** states that it **enables neurexin family protein binding**, is **involved in regulation of synaptic vesicle exocytosis**, and acts within processes including **calcium ion import**; it is also annotated as active in the **Schaffer collateral–CA1 synapse**. Database annotations should not outweigh primary experiments, but they are consistent with the synaptic-regulatory interpretation. ([informatics.jax.org](https://www.informatics.jax.org/go/marker/MGI%3A1309489)) [S6]

There is also disease-context evidence consistent with a regulatory role in neural systems. In glioma models, CA11 and CA10 were described as **secreted by depolarized neurons**, and CA11 knockdown promoted more aggressive glioma behavior, leading the authors to conclude that CA11/CA10 **negatively regulate neuronal activity-dependent glioma growth**. That is not the same as normal neuronal homeostasis, but it does show CA11 participating in neuron-derived regulatory signaling in a CNS microenvironment. ([pmc.ncbi.nlm.nih.gov](https://pmc.ncbi.nlm.nih.gov/articles/PMC6487704/)) [S7]

A more indirect “homeostatic” clue comes from a heterologous-system study showing that catalytically inactive CARPs, including **CA11**, can enhance **MCT1-mediated lactate transport**, with the authors proposing a noncatalytic “proton antenna” role. Since lactate shuttling is relevant to brain metabolic support, this offers a conceivable homeostatic mechanism, though it has not been firmly demonstrated for CA11 in intact neural tissue. ([pubmed.ncbi.nlm.nih.gov](https://pubmed.ncbi.nlm.nih.gov/31033227/)) [S8]

## Contradicting or limiting evidence

The main limitation is that **CA11’s precise physiological role is still poorly defined**. A review focused on CARPs VIII/X/XI states that these proteins are predominantly expressed in brain, but their **precise roles are poorly known**. That makes any strong claim about CA11’s specific neural function premature. ([pubmed.ncbi.nlm.nih.gov](https://pubmed.ncbi.nlm.nih.gov/23294106/?utm_source=openai)) [S9]

Also, the best mechanistic synaptic paper directly dissected **CA10** much more extensively than CA11. CA11 was identified in the neurexin complex and is repeatedly discussed as a likely homologous partner, but several functional conclusions for CA11 are at least partly **inferred from CA10 homology and shared binding behavior**, rather than demonstrated by CA11-specific knockout or rescue experiments in normal neurons. ([pubmed.ncbi.nlm.nih.gov](https://pubmed.ncbi.nlm.nih.gov/28154140/)) [S10]

Importantly, there is little evidence that CA11 itself performs classic carbonic-anhydrase-like **acid-base homeostasis** in the nervous system. CA11 lacks the key zinc-binding histidines required for catalytic CA activity, and reviews classify it as **catalytically inactive**. So while other carbonic anhydrases clearly contribute to pH regulation, that should **not** be assumed for CA11. ([pubmed.ncbi.nlm.nih.gov](https://pubmed.ncbi.nlm.nih.gov/10350627/)) [S11]

The MCT1/lactate-transport result is also limited by model system: it was shown in **Xenopus oocytes**, not in neurons or intact brain tissue. Thus, it supports a possible homeostatic mechanism but remains **indirect** for neural systems. ([pubmed.ncbi.nlm.nih.gov](https://pubmed.ncbi.nlm.nih.gov/31033227/)) [S12]

## Mechanistic / biological rationale

A biologically coherent model is that CA11 is an **acatalytic, brain-enriched carbonic-anhydrase-related protein** that participates in **synaptic protein networks** rather than in classical CO2 hydration chemistry. Its association with **neurexins** is particularly meaningful because neurexins organize synaptic adhesion, transmitter release machinery, and synapse specification. If CA11 helps neurexin trafficking, folding, or assembly of trans-synaptic complexes, then CA11 would reasonably count as being involved in **regulatory processes in neural systems**. ([pubmed.ncbi.nlm.nih.gov](https://pubmed.ncbi.nlm.nih.gov/28154140/)) [S13]

Its fetal expression in **neuroprogenitors** and migrating neurons also fits a developmental-regulatory role, potentially in neural differentiation or circuit assembly. That interpretation has been suggested in the CARP literature, though it remains more hypothesis-generating than definitive. ([pubmed.ncbi.nlm.nih.gov](https://pubmed.ncbi.nlm.nih.gov/12044474/)) [S14]

A possible, more homeostasis-oriented mechanism is that CA11 may noncatalytically facilitate **MCT1-dependent lactate/proton transport**. If that occurs in vivo in brain, it could contribute to metabolic support or pH-coupled transport relevant to neuron-glia interactions. But at present this remains a **plausible extrapolation**, not a firmly established neural function. ([pubmed.ncbi.nlm.nih.gov](https://pubmed.ncbi.nlm.nih.gov/31033227/)) [S15]

## Bottom line

The claim is **best judged Likely True**: CA11 is clearly associated with the **nervous system** and with **regulatory/synaptic biology**, especially via brain-enriched expression, developmental neural expression, and neurexin-linked synaptic mechanisms. What is **not** yet well established is a direct, specific role for CA11 in **neural homeostasis** in the strict sense. So: **regulatory association yes; strong homeostatic proof not yet.** ([pubmed.ncbi.nlm.nih.gov](https://pubmed.ncbi.nlm.nih.gov/10350627/)) [S16]

## Cited sources

- **[S1]** cDNA sequence, mRNA expression, and chromosomal localization of human carbonic anhydrase-related protein, CA-RP XI - PubMed — https://pubmed.ncbi.nlm.nih.gov/10350627/
- **[S2]** Carbonic anhydrase-related protein CA10 is an evolutionarily conserved pan-neurexin ligand - PubMed — https://pubmed.ncbi.nlm.nih.gov/28154140/
- **[S3]** cDNA sequence, mRNA expression, and chromosomal localization of human carbonic anhydrase-related protein, CA-RP XI - PubMed — https://pubmed.ncbi.nlm.nih.gov/10350627/
- **[S4]** Developmental expression of carbonic anhydrase-related proteins VIII, X, and XI in the human brain - PubMed — https://pubmed.ncbi.nlm.nih.gov/12044474/
- **[S5]** Carbonic anhydrase-related protein CA10 is an evolutionarily conserved pan-neurexin ligand - PubMed — https://pubmed.ncbi.nlm.nih.gov/28154140/
- **[S6]** Gene Ontology Classifications — https://www.informatics.jax.org/go/marker/MGI%3A1309489
- **[S7]** 
            CA10 and CA11 negatively regulate neuronal activity‐dependent growth of gliomas - PMC
         — https://pmc.ncbi.nlm.nih.gov/articles/PMC6487704/
- **[S8]** Catalytically inactive carbonic anhydrase-related proteins enhance transport of lactate by MCT1 - PubMed — https://pubmed.ncbi.nlm.nih.gov/31033227/
- **[S9]** An update on carbonic anhydrase-related proteins VIII, X and XI - PubMed — https://pubmed.ncbi.nlm.nih.gov/23294106/?utm_source=openai
- **[S10]** Carbonic anhydrase-related protein CA10 is an evolutionarily conserved pan-neurexin ligand - PubMed — https://pubmed.ncbi.nlm.nih.gov/28154140/
- **[S11]** cDNA sequence, mRNA expression, and chromosomal localization of human carbonic anhydrase-related protein, CA-RP XI - PubMed — https://pubmed.ncbi.nlm.nih.gov/10350627/
- **[S12]** Catalytically inactive carbonic anhydrase-related proteins enhance transport of lactate by MCT1 - PubMed — https://pubmed.ncbi.nlm.nih.gov/31033227/
- **[S13]** Carbonic anhydrase-related protein CA10 is an evolutionarily conserved pan-neurexin ligand - PubMed — https://pubmed.ncbi.nlm.nih.gov/28154140/
- **[S14]** Developmental expression of carbonic anhydrase-related proteins VIII, X, and XI in the human brain - PubMed — https://pubmed.ncbi.nlm.nih.gov/12044474/
- **[S15]** Catalytically inactive carbonic anhydrase-related proteins enhance transport of lactate by MCT1 - PubMed — https://pubmed.ncbi.nlm.nih.gov/31033227/
- **[S16]** cDNA sequence, mRNA expression, and chromosomal localization of human carbonic anhydrase-related protein, CA-RP XI - PubMed — https://pubmed.ncbi.nlm.nih.gov/10350627/
