## Supplementary material for "Understanding Substance Dependence: What Differentiates Addictive from Non-Addictive Drugs?": Folder for supplementary files: non-addictive-genes-claims-first100-066-ubl5-gene-protein-has-been-associated-with-regulat-1805f5643e4e.md

# non-addictive-genes-claims-first100-066-1805f5643e4e

**Original claim:** UBL5 gene/protein has been associated with regulatory or homeostatic biological processes in neural systems.

## Claim

UBL5 gene/protein has been associated with regulatory or homeostatic biological processes in neural systems. ([ncbi.nlm.nih.gov](https://www.ncbi.nlm.nih.gov/gene/59286)) [S1]

## Verdict

Verdict: Likely True

## Confidence

Medium

## Supporting evidence

UBL5 is the human **ubiquitin-like 5** gene, and **“beacon”** is a recognized alias for the same gene/protein in NCBI records. NCBI also annotates UBL5 as involved in **mRNA splicing via the spliceosome** and **positive regulation of protein localization to mitochondrion**, which are regulatory/homeostatic cellular functions. ([ncbi.nlm.nih.gov](https://www.ncbi.nlm.nih.gov/gene/59286)) [S2]

There is direct **brain and hypothalamic association** data. The Human Protein Atlas reports UBL5 RNA expression across multiple human brain regions, including the **hypothalamus**, and analogous datasets are also available for mouse brain. ([proteinatlas.org](https://www.proteinatlas.org/ENSG00000198258-UBL5/brain)) [S3]

The strongest neural-system evidence comes from the earlier **“beacon” neuroendocrine literature**. In the original sand-rat study, beacon/UBL5 was identified from **hypothalamic mRNA**, and hypothalamic beacon expression correlated with body fat. Importantly, **intracerebroventricular beacon infusion increased food intake, body weight, and hypothalamic NPY expression**, directly linking this molecule to central regulation of energy balance. ([pubmed.ncbi.nlm.nih.gov](https://pubmed.ncbi.nlm.nih.gov/11078442/)) [S4]

Follow-up neuroanatomical studies localized beacon/UBL5 protein to classic **homeostatic hypothalamic nuclei**. In mouse, beacon mRNA was detected in hypothalamus, and immunoreactivity was concentrated in the **supraoptic** and **paraventricular** nuclei, median eminence, pituitary stalk, and posterior pituitary; nearly all oxytocin-immunoreactive neurons and a subset of vasopressin neurons in PVH/SO were beacon-positive. The authors also note that human UBL5 has the exact same sequence as beacon. ([pubmed.ncbi.nlm.nih.gov](https://pubmed.ncbi.nlm.nih.gov/12932856/)) [S5]

A rat study similarly found beacon-like immunoreactivity in multiple hypothalamic regions involved in feeding and neuroendocrine control, including the **paraventricular, supraoptic, suprachiasmatic, arcuate, and preoptic/lateral hypothalamic areas**, and concluded that the peptide may influence biological activities associated with the **hypothalamic-pituitary axis**. ([pubmed.ncbi.nlm.nih.gov](https://pubmed.ncbi.nlm.nih.gov/11755265/?utm_source=openai)) [S6]

Human relevance is supported by a postmortem study showing that beacon-like/UBL5 immunoreactivity is **highly expressed in human hypothalamus**, including paraventricular and supraoptic nuclei, and is also present in hippocampus, thalamus, and some neocortical areas. That study found altered hypothalamic expression in schizophrenia and in a rat model, which at minimum shows that UBL5 is a measurable brain-associated protein in human neural tissue. ([pubmed.ncbi.nlm.nih.gov](https://pubmed.ncbi.nlm.nih.gov/18191906/?utm_source=openai)) [S7]

Beyond the nervous system, modern UBL5 biology strongly supports a **regulatory/homeostatic** profile. In *C. elegans*, ubl-5 is required for induction of mitochondrial chaperone genes during the **mitochondrial unfolded protein response (UPRmt)** and helps animals cope with mitochondrial stress. ([pubmed.ncbi.nlm.nih.gov](https://pubmed.ncbi.nlm.nih.gov/16816413/)) [S8] In mammalian cells, UBL5 has been shown to be a **stress-responsive survival regulator** in the **ER stress–PERK** pathway, where its depletion promotes apoptosis and its overexpression protects against ER stress-induced cell death. ([pubmed.ncbi.nlm.nih.gov](https://pubmed.ncbi.nlm.nih.gov/37315790/)) [S9] Human-cell studies also show UBL5 is essential for **pre-mRNA splicing**, another core regulatory process. ([pubmed.ncbi.nlm.nih.gov](https://pubmed.ncbi.nlm.nih.gov/25092792/)) [S10]

Taken together, these data support the claim that UBL5 has been associated with **neural-system regulatory/homeostatic processes**, especially in the **hypothalamic/neuroendocrine control of energy balance**, while also having broader conserved roles in stress adaptation and gene-expression control. ([pubmed.ncbi.nlm.nih.gov](https://pubmed.ncbi.nlm.nih.gov/11078442/)) [S11]

## Contradicting or limiting evidence

The evidence is **not broad or definitive across all neural systems**. Most direct nervous-system studies are relatively old and focus mainly on **hypothalamic/neuroendocrine** contexts rather than synaptic plasticity, circuit homeostasis, or pan-neuronal physiology. ([pubmed.ncbi.nlm.nih.gov](https://pubmed.ncbi.nlm.nih.gov/11078442/)) [S12]

Also, much of the strongest modern mechanistic work on UBL5 concerns **non-neural systems**: spliceosome function in HeLa cells, mitochondrial-stress signaling in worms and human cell lines, Fanconi-anemia pathway biology, and ER-stress survival in hepatoma or other cultured cells. These studies strongly establish UBL5 as a regulator of cellular homeostasis, but they do **not by themselves prove a neuron-specific homeostatic role**. ([pubmed.ncbi.nlm.nih.gov](https://pubmed.ncbi.nlm.nih.gov/25092792/)) [S13]

Human genetic evidence tying UBL5 to obesity-related phenotypes is also **inconsistent**. NCBI lists both positive metabolic-association papers and a study concluding that UBL5 is unlikely to play a major role in susceptibility to **early-onset obesity**; that obesity genetics paper specifically found **no evidence for association** in obese children. ([ncbi.nlm.nih.gov](https://www.ncbi.nlm.nih.gov/gene/59286)) [S14]

The schizophrenia-related brain findings are **observational** and do not establish that UBL5 drives disease or neural homeostasis. Likewise, transcriptomic differences involving UBL5 in cortical pyramidal neurons from schizophrenia brains show association, not mechanism. ([pubmed.ncbi.nlm.nih.gov](https://pubmed.ncbi.nlm.nih.gov/18191906/)) [S15]

## Mechanistic / biological rationale

Biologically, the claim is plausible. UBL5 is a conserved ubiquitin-like protein with roles in **RNA splicing**, **mitochondrial stress responses**, and **ER-stress survival**. Those processes are central to maintaining cellular proteostasis, mitochondrial integrity, and gene-expression fidelity. ([ncbi.nlm.nih.gov](https://www.ncbi.nlm.nih.gov/gene/59286)) [S16]

That matters for neural systems because hypothalamic and other neurons are highly dependent on **mitochondrial function**, **secretory capacity**, and **tight transcriptional/splicing control** to sustain homeostatic outputs. The direct localization of beacon/UBL5 to oxytocin- and vasopressin-rich hypothalamic nuclei, plus its effects on **food intake** and **NPY expression**, fit a model in which UBL5 participates in **neuroendocrine energy-homeostasis regulation**. ([pubmed.ncbi.nlm.nih.gov](https://pubmed.ncbi.nlm.nih.gov/12932856/)) [S17]

A broader inference is that UBL5’s established cellular-homeostasis roles could support neuronal resilience as well, but that extrapolation remains **indirect** because direct neuron-specific mechanistic experiments are limited. ([pubmed.ncbi.nlm.nih.gov](https://pubmed.ncbi.nlm.nih.gov/37315790/)) [S18]

## Bottom line

The claim is **likely true**, but with an important qualifier: the best direct evidence links UBL5/“beacon” to **hypothalamic and neuroendocrine regulation of energy balance**, not to a broadly established role in all neural homeostasis. UBL5 is clearly a conserved regulator of stress-response and gene-expression pathways, and it is expressed in brain tissue, including hypothalamus. However, **direct mechanistic evidence in neurons remains limited**, so the strongest formulation is that UBL5 has been associated with **neural-system homeostatic/regulatory processes, especially hypothalamic energy-balance pathways**. ([pubmed.ncbi.nlm.nih.gov](https://pubmed.ncbi.nlm.nih.gov/11078442/)) [S19]

## Cited sources

- **[S1]** UBL5 ubiquitin like 5 [Homo sapiens (human)] - Gene - NCBI — https://www.ncbi.nlm.nih.gov/gene/59286
- **[S2]** UBL5 ubiquitin like 5 [Homo sapiens (human)] - Gene - NCBI — https://www.ncbi.nlm.nih.gov/gene/59286
- **[S3]** Brain tissue expression of UBL5 - Summary - The Human Protein Atlas — https://www.proteinatlas.org/ENSG00000198258-UBL5/brain
- **[S4]** Beacon: a novel gene involved in the regulation of energy balance - PubMed — https://pubmed.ncbi.nlm.nih.gov/11078442/
- **[S5]** Beacon/ubiquitin-like 5-immunoreactivity in the hypothalamus and pituitary of the mouse - PubMed — https://pubmed.ncbi.nlm.nih.gov/12932856/
- **[S6]** Beacon-like immunoreactivity in the hypothalamus of Sprague-Dawley rats - PubMed — https://pubmed.ncbi.nlm.nih.gov/11755265/?utm_source=openai
- **[S7]** Beacon-like/ubiquitin-5-like immunoreactivity is highly expressed in human hypothalamus and increased in haloperidol-treated schizophrenics and a rat model of schizophrenia - PubMed — https://pubmed.ncbi.nlm.nih.gov/18191906/?utm_source=openai
- **[S8]** Ubiquitin-like protein 5 positively regulates chaperone gene expression in the mitochondrial unfolded protein response - PubMed — https://pubmed.ncbi.nlm.nih.gov/16816413/
- **[S9]** Ubiquitin-like protein 5 is a novel player in the UPR-PERK arm and ER stress-induced cell death - PubMed — https://pubmed.ncbi.nlm.nih.gov/37315790/
- **[S10]** UBL5 is essential for pre-mRNA splicing and sister chromatid cohesion in human cells - PubMed — https://pubmed.ncbi.nlm.nih.gov/25092792/
- **[S11]** Beacon: a novel gene involved in the regulation of energy balance - PubMed — https://pubmed.ncbi.nlm.nih.gov/11078442/
- **[S12]** Beacon: a novel gene involved in the regulation of energy balance - PubMed — https://pubmed.ncbi.nlm.nih.gov/11078442/
- **[S13]** UBL5 is essential for pre-mRNA splicing and sister chromatid cohesion in human cells - PubMed — https://pubmed.ncbi.nlm.nih.gov/25092792/
- **[S14]** UBL5 ubiquitin like 5 [Homo sapiens (human)] - Gene - NCBI — https://www.ncbi.nlm.nih.gov/gene/59286
- **[S15]** Beacon-like/ubiquitin-5-like immunoreactivity is highly expressed in human hypothalamus and increased in haloperidol-treated schizophrenics and a rat model of schizophrenia - PubMed — https://pubmed.ncbi.nlm.nih.gov/18191906/
- **[S16]** UBL5 ubiquitin like 5 [Homo sapiens (human)] - Gene - NCBI — https://www.ncbi.nlm.nih.gov/gene/59286
- **[S17]** Beacon/ubiquitin-like 5-immunoreactivity in the hypothalamus and pituitary of the mouse - PubMed — https://pubmed.ncbi.nlm.nih.gov/12932856/
- **[S18]** Ubiquitin-like protein 5 is a novel player in the UPR-PERK arm and ER stress-induced cell death - PubMed — https://pubmed.ncbi.nlm.nih.gov/37315790/
- **[S19]** Beacon: a novel gene involved in the regulation of energy balance - PubMed — https://pubmed.ncbi.nlm.nih.gov/11078442/
