## Supplementary material for "Understanding Substance Dependence: What Differentiates Addictive from Non-Addictive Drugs?": Folder for supplementary files: non-addictive-genes-claims-first100-067-nisch-gene-protein-has-been-associated-with-regula-85c9ee6c2017.md

# non-addictive-genes-claims-first100-067-85c9ee6c2017

**Original claim:** NISCH gene/protein has been associated with regulatory or homeostatic biological processes in neural systems.

## Claim

NISCH gene/protein has been associated with regulatory or homeostatic biological processes in neural systems.

## Verdict

Verdict: Likely True

NISCH encodes **nischarin**, and the available literature supports its involvement in several **regulatory neural processes**, including **neuronal migration**, **neurite/dendrite outgrowth**, **dendritic spine morphogenesis**, and **astrocytic glutamate transporter trafficking**. The best evidence for a specifically **homeostatic** neural role is a study showing nischarin regulates **GLT-1/EAAT2 surface trafficking and glutamate uptake in astrocytes**, a process central to extracellular glutamate control and protection from excitotoxicity. However, the evidence base is still **mostly preclinical** and does **not clearly establish NISCH as a major mediator of canonical neural homeostasis** such as firing-rate homeostasis or synaptic scaling in humans. ([ncbi.nlm.nih.gov](https://www.ncbi.nlm.nih.gov/gene/11188)) [S1]

## Confidence

Medium

Confidence is **medium** because multiple independent papers place nischarin in neural regulatory pathways, but most evidence comes from **rodent tissue, cultured neurons/astrocytes, and genetic perturbation models**, with comparatively limited direct human functional data. ([pubmed.ncbi.nlm.nih.gov](https://pubmed.ncbi.nlm.nih.gov/40262669/)) [S2]

## Supporting evidence

Human and vertebrate reference resources indicate that **NISCH is expressed in the nervous system**. NCBI’s RefSeq summary states that in neural tissue NISCH has been found to affect **neuronal growth and differentiation**, and the Human Protein Atlas classifies NISCH as **brain-enhanced**, with RNA detected across all major surveyed human brain regions. ([ncbi.nlm.nih.gov](https://www.ncbi.nlm.nih.gov/gene/11188)) [S3]

Experimental work in rodents supports a neural developmental/regulatory role. A 2013 study reported that nischarin is expressed in multiple regions and cell types of the adult rat brain and concluded that it is a **key regulatory molecule controlling neuronal migration**. A newer 2025 study found that nischarin expression increases in rat cortex and hippocampus during postnatal development and that suppressing nischarin enhances neuronal migration while altering the **PAK1/LIMK1/cofilin** pathway, consistent with a role in shaping the normal developmental trajectory of the brain. ([pmc.ncbi.nlm.nih.gov](https://pmc.ncbi.nlm.nih.gov/articles/PMC3546990/?utm_source=openai)) [S4]

A separate neuronal study showed that **knockdown of nischarin promotes neurite outgrowth and dendrite elongation** in neuronal cells and primary cortical neurons. In that work, nischarin interacted with **PAK1/2** and negatively regulated their phosphorylation, indicating that nischarin acts as a brake on neuronal process extension rather than being merely a passive marker of neural tissue. ([pubmed.ncbi.nlm.nih.gov](https://pubmed.ncbi.nlm.nih.gov/26670864/)) [S5]

The strongest direct evidence for a **homeostatic-type neural function** comes from astrocytes. A 2022 study identified nischarin as a **physiological binding partner of GLT-1** in astrocytes and showed that glutamate drives **nischarin-dependent GLT-1 internalization**. Nisch knockout increased astrocytic GLT-1 surface density, increased transporter current, and was associated with **enhanced neuroprotection**. Because astrocytic GLT-1/EAAT2 is a principal mechanism for clearing extracellular glutamate and preventing excitotoxicity, this places nischarin in a bona fide regulatory pathway relevant to **glutamate homeostasis** in the CNS. ([pmc.ncbi.nlm.nih.gov](https://pmc.ncbi.nlm.nih.gov/articles/PMC9010640/)) [S6]

There is also evidence that NISCH influences **synaptic structure and cognition-related neural phenotypes**. A 2023 study at a psychiatric GWAS locus implicated NISCH as a candidate risk gene and found that increasing NISCH expression altered **dendritic spine morphogenesis** and impaired **spatial working memory** in mice; the same paper reported higher NISCH mRNA in postmortem brains from patients with psychiatric disorders than in controls. These findings strengthen the case that NISCH participates in neural regulatory biology, even if disease causality remains incomplete. ([pmc.ncbi.nlm.nih.gov](https://pmc.ncbi.nlm.nih.gov/articles/PMC10347724/)) [S7]

Older in vivo work further connected nischarin to **brainstem signaling regulation**. In rats, knockdown of nischarin in the rostral ventrolateral medulla largely abolished **rilmenidine-evoked ERK1/2 phosphorylation and hypotension**, suggesting nischarin participates in central neural signaling pathways relevant to autonomic regulation. ([pmc.ncbi.nlm.nih.gov](https://pmc.ncbi.nlm.nih.gov/articles/PMC2704025/)) [S8]

## Contradicting or limiting evidence

The main limitation is that the evidence is **not broad enough to claim NISCH is a well-established master regulator of neural homeostasis**. In the literature reviewed, nischarin’s neural roles are described mainly in the contexts of **migration, neurite growth, spine morphology, receptor/transporter trafficking, and injury-related models**, rather than the classic neuroscience meanings of homeostasis such as **firing-rate homeostasis, synaptic scaling, or network set-point control**. That makes the claim supportable mainly through the words **“associated with”** and **“regulatory,”** and only more cautiously through **“homeostatic.”** ([pubmed.ncbi.nlm.nih.gov](https://pubmed.ncbi.nlm.nih.gov/26670864/)) [S9]

Another limitation is **species and model dependence**. Much of the functional evidence comes from **rat or mouse studies**, cultured neurons, Neuro-2a/PC12-like systems, or overexpression/knockdown approaches. Those models are valuable, but they do not by themselves prove the same magnitude or context of function in the human brain. ([pubmed.ncbi.nlm.nih.gov](https://pubmed.ncbi.nlm.nih.gov/26670864/)) [S10]

Even the human psychiatric-risk study is appropriately cautious: its authors note that whether NISCH-driven spine abnormalities are **sufficient** to produce psychiatric symptoms remains undetermined, and that the relevant **neural circuits** are still unclear. ([pmc.ncbi.nlm.nih.gov](https://pmc.ncbi.nlm.nih.gov/articles/PMC10347724/)) [S11]

## Mechanistic / biological rationale

Mechanistically, nischarin is an **intracellular scaffold/adaptor protein** linked to membrane-associated signaling, endosomal trafficking, and cytoskeletal regulation. NCBI describes it as a cytosolic, inner-plasma-membrane-associated imidazoline-1 receptor protein, and experimental studies show it can influence signaling systems tied to **integrins, Rac/PAK/LIMK/cofilin, and receptor/transporter trafficking**. ([ncbi.nlm.nih.gov](https://www.ncbi.nlm.nih.gov/gene/11188)) [S12]

That biology fits neural regulation well. In neurons, **actin cytoskeleton remodeling** underlies migration, neurite extension, and dendritic spine shape. A protein that restrains or tunes **PAK/LIMK/cofilin signaling** could plausibly regulate where neurons migrate, how fast neurites extend, and how spines mature. The migration and neurite papers are therefore mechanistically coherent rather than isolated observations. ([pubmed.ncbi.nlm.nih.gov](https://pubmed.ncbi.nlm.nih.gov/26670864/)) [S13]

In astrocytes, the rationale is even closer to a homeostatic function. GLT-1/EAAT2 is the major astrocytic glutamate transporter responsible for keeping extracellular glutamate in check. Since nischarin promotes **GLT-1 internalization**, it can tune how much transporter remains at the cell surface and therefore how strongly astrocytes buffer synaptic glutamate. That is a direct route by which NISCH could influence **neural excitability stability and protection from excitotoxic stress**. ([pmc.ncbi.nlm.nih.gov](https://pmc.ncbi.nlm.nih.gov/articles/PMC9010640/)) [S14]

## Bottom line

Overall, the claim is **likely true**. NISCH/nischarin has credible experimental links to **regulatory processes in neural systems**, especially **neuronal migration**, **neurite/dendrite growth**, **dendritic spine morphogenesis**, and **astrocytic glutamate transporter regulation**. The strongest **homeostatic** angle is its role in controlling **astrocyte GLT-1 trafficking and glutamate clearance**, not yet a broad demonstration of classical neural homeostatic plasticity. So the safest summary is: **NISCH is supported as a neural regulatory gene/protein, with some evidence for participation in glutamatergic homeostatic control, but the homeostasis claim is narrower and less mature than the general regulatory claim.** ([ncbi.nlm.nih.gov](https://www.ncbi.nlm.nih.gov/gene/11188)) [S15]

## Cited sources

- **[S1]** NISCH nischarin [Homo sapiens (human)] - Gene - NCBI — https://www.ncbi.nlm.nih.gov/gene/11188
- **[S2]** Spatiotemporal expression of Nischarin in developing rat brain mediates neuronal migration via the PAK1/LIMK1/cofilin pathway - PubMed — https://pubmed.ncbi.nlm.nih.gov/40262669/
- **[S3]** NISCH nischarin [Homo sapiens (human)] - Gene - NCBI — https://www.ncbi.nlm.nih.gov/gene/11188
- **[S4]** Nischarin Is Differentially Expressed in Rat Brain and Regulates Neuronal Migration - PMC — https://pmc.ncbi.nlm.nih.gov/articles/PMC3546990/?utm_source=openai
- **[S5]** Inhibition of Nischarin Expression Promotes Neurite Outgrowth through Regulation of PAK Activity - PubMed — https://pubmed.ncbi.nlm.nih.gov/26670864/
- **[S6]** 
            The non-adrenergic imidazoline-1 receptor protein nischarin is a key regulator of astrocyte glutamate uptake - PMC
         — https://pmc.ncbi.nlm.nih.gov/articles/PMC9010640/
- **[S7]** 
            Identification of a psychiatric risk gene NISCH at 3p21.1 GWAS locus mediating dendritic spine morphogenesis and cognitive function - PMC
         — https://pmc.ncbi.nlm.nih.gov/articles/PMC10347724/
- **[S8]** 
            Inhibition of Nischarin Expression Attenuates Rilmenidine-Evoked Hypotension and Phosphorylated Extracellular Signal-Regulated Kinase 1/2 Production in the Rostral Ventrolateral Medulla of Rats - PMC
         — https://pmc.ncbi.nlm.nih.gov/articles/PMC2704025/
- **[S9]** Inhibition of Nischarin Expression Promotes Neurite Outgrowth through Regulation of PAK Activity - PubMed — https://pubmed.ncbi.nlm.nih.gov/26670864/
- **[S10]** Inhibition of Nischarin Expression Promotes Neurite Outgrowth through Regulation of PAK Activity - PubMed — https://pubmed.ncbi.nlm.nih.gov/26670864/
- **[S11]** 
            Identification of a psychiatric risk gene NISCH at 3p21.1 GWAS locus mediating dendritic spine morphogenesis and cognitive function - PMC
         — https://pmc.ncbi.nlm.nih.gov/articles/PMC10347724/
- **[S12]** NISCH nischarin [Homo sapiens (human)] - Gene - NCBI — https://www.ncbi.nlm.nih.gov/gene/11188
- **[S13]** Inhibition of Nischarin Expression Promotes Neurite Outgrowth through Regulation of PAK Activity - PubMed — https://pubmed.ncbi.nlm.nih.gov/26670864/
- **[S14]** 
            The non-adrenergic imidazoline-1 receptor protein nischarin is a key regulator of astrocyte glutamate uptake - PMC
         — https://pmc.ncbi.nlm.nih.gov/articles/PMC9010640/
- **[S15]** NISCH nischarin [Homo sapiens (human)] - Gene - NCBI — https://www.ncbi.nlm.nih.gov/gene/11188
