## Supplementary material for "Understanding Substance Dependence: What Differentiates Addictive from Non-Addictive Drugs?": Folder for supplementary files: non-addictive-genes-claims-first100-068-nkx6-2-gene-protein-has-been-associated-with-regul-dc25d0a270a4.md

# non-addictive-genes-claims-first100-068-dc25d0a270a4

**Original claim:** NKX6-2 gene/protein has been associated with regulatory or homeostatic biological processes in neural systems.

## Claim

NKX6-2 gene/protein has been associated with regulatory or homeostatic biological processes in neural systems.

## Verdict

Verdict: Highly Supported

NKX6-2 is well documented as a **homeobox transcription factor with repressor activity** in the nervous system, especially in **ventral neural patterning**, **oligodendrocyte maturation**, **myelin-related gene regulation**, and **maintenance of myelin/paranodal structure**. Human genetics also links loss of NKX6-2 to **hypomyelinating neurologic disease**, reinforcing that its neural regulatory role is biologically important rather than merely correlative. ([proteinatlas.org](https://www.proteinatlas.org/ENSG00000148826-NKX6-2)) [S1]

## Confidence

High

The evidence is convergent across **expression studies**, **mouse loss-of-function experiments**, **developmental neurobiology**, and **human Mendelian disease**. The only important qualifier is that the strongest “homeostatic” evidence is mainly about **myelin maintenance/homeostasis**, not broad neural homeostasis in the synaptic-plasticity sense. ([pmc.ncbi.nlm.nih.gov](https://pmc.ncbi.nlm.nih.gov/articles/PMC2807475/)) [S2]

## Supporting evidence

- **NKX6-2 is itself a regulatory protein.** Human Protein Atlas / UniProt-based annotation describes NKX6-2 as a **transcription factor with repressor activity**, involved in regulation of **axon-glial interactions at myelin paranodes**; it binds a defined DNA motif and, in oligodendrocytes, binds the **MBP** and **PLP1** promoter regions. Entrez/Alliance-based summary on the same page also links it to **regulation of myelination**, **negative regulation of transcription**, and **neurogenesis**. ([proteinatlas.org](https://www.proteinatlas.org/ENSG00000148826-NKX6-2)) [S3]

- **It has a direct neural developmental regulatory role.** In a classic mouse study, Nkx6.2 and Nkx6.1 were described as **transcriptional repressors** expressed in overlapping ventral spinal cord progenitor domains; altered Nkx6 activity changed interneuron and motor-neuron specification, supporting a model in which these proteins direct neural cell fates by **actively suppressing alternative transcriptional programs**. ([sciencedirect.com](https://www.sciencedirect.com/science/article/pii/S0896627301004123)) [S4]

- **Its neural expression pattern is highly consistent with oligodendrocyte-related regulation.** Postnatal mouse work showed Nkx6.2 is **specifically expressed in mature oligodendrocytes** in spinal cord and brain, rather than broadly across neural cell types, and appears downstream of Nkx2.2 during oligodendrocyte lineage progression. ([pmc.ncbi.nlm.nih.gov](https://pmc.ncbi.nlm.nih.gov/articles/PMC2807475/)) [S5]

- **Nkx6.2 contributes to postnatal myelin maintenance, not just embryonic patterning.** In Nkx6.2-null mice, investigators found **motor coordination and CNS conduction defects**, abnormal paranodes, ectopic axoglial junctions, and age-related instability of paranodal dimensions. The study concluded that Nkx6.2 sits genetically upstream of **cell-adhesion** and **cytoskeletal-remodeling** programs at paranodes, including altered NF155, contactin, and stathmin-1 expression. That is strong evidence for a role in neural structural maintenance/homeostasis at the axon-glial interface. ([pmc.ncbi.nlm.nih.gov](https://pmc.ncbi.nlm.nih.gov/articles/PMC6730372/)) [S6]

- **Human disease genetics strongly supports biological relevance in CNS myelin regulation.** A *Brain* study identified biallelic inactivating **NKX6-2** variants causing severe **hypomyelinating leukodystrophy**, and described NKX6-2 as having a main role in **oligodendrocyte differentiation** and **regulation of myelin-specific gene expression**. ([academic.oup.com](https://academic.oup.com/brain/article/140/10/2550/4085303)) [S7]

- A separate human genetics study on progressive spastic ataxia/hypomyelination concluded that the syndrome confirms a **non-redundant role of NKX6-2 in myelin homeostasis in the central nervous system** and placed NKX6-2 within a broader **myelin regulation pathway**. That directly supports the “homeostatic” part of the claim, albeit in a fairly specific myelin-centered sense. ([pmc.ncbi.nlm.nih.gov](https://pmc.ncbi.nlm.nih.gov/articles/PMC5473715/)) [S8]

- **Additional neural-system regulatory effects have been reported outside white matter alone.** In mouse spinal cord, Nkx6.2 loss altered **locomotor pattern** and **spinal locomotor network activity**, likely through defective survival of a subset of limb-innervating motor neurons and abnormal migration/distribution of V2a interneurons. The authors concluded Nkx6.2 regulates development of spinal neuronal populations and formation of locomotor circuits. ([pubmed.ncbi.nlm.nih.gov](https://pubmed.ncbi.nlm.nih.gov/31969659/)) [S9]

- **Cortical interneuron lineage data also point toward a regulatory developmental role.** Fate-mapping and loss-of-function work found that Nkx6-2-expressing telencephalic progenitors give rise to multiple cortical interneuron subtypes, with a small, partially penetrant loss of one interneuron subtype in Nkx6-2-null mice. ([pubmed.ncbi.nlm.nih.gov](https://pubmed.ncbi.nlm.nih.gov/19363146/)) [S10]

## Contradicting or limiting evidence

An early Nkx6.2 knockout study reported that mice were **viable and fertile without obvious gross CNS histologic defects**, and that myelin gene expression appeared normal in the mutant brain and spinal cord. The authors interpreted this as evidence that Nkx6.2 may have **redundant** developmental functions under some conditions. ([pubmed.ncbi.nlm.nih.gov](https://pubmed.ncbi.nlm.nih.gov/11390667/)) [S11]

Some later neural phenotypes are **modest or selective**, not universal. For example, cortical interneuron effects were reported as **small and partially penetrant**, and in spinal cord not all neuronal subpopulations were equally affected by Nkx6.2 loss. ([pubmed.ncbi.nlm.nih.gov](https://pubmed.ncbi.nlm.nih.gov/19363146/)) [S12]

Also, the strongest evidence for “homeostasis” is **narrowly centered on CNS myelin/paranodal maintenance**. There is little direct evidence that NKX6-2 is a major regulator of broader neural homeostatic processes such as **synaptic scaling**, **firing-rate homeostasis**, or **redox/metabolic homeostasis** across neurons in general. So the claim is best interpreted as true mainly in the context of **neural development and myelin/axon-glial stability**. ([pmc.ncbi.nlm.nih.gov](https://pmc.ncbi.nlm.nih.gov/articles/PMC6730372/)) [S13]

## Mechanistic / biological rationale

Mechanistically, NKX6-2 fits the claim because it is a **sequence-specific homeobox DNA-binding transcription factor** with **repressor activity**. In developmental contexts, Nkx6-family repression helps neural progenitors adopt one fate while suppressing competing transcriptional programs. ([proteinatlas.org](https://www.proteinatlas.org/ENSG00000148826-NKX6-2)) [S14]

In oligodendrocytes, NKX6-2 appears positioned to regulate both **myelin gene expression** and **axon-glial structural organization**. It has been linked to **MBP** and **PLP1** promoter binding, and loss of Nkx6.2 disrupts expression of paranodal/structural effectors such as **NF155**, **contactin**, and **stathmin-1**, which plausibly explains impaired paranode organization, abnormal conduction, and hypomyelination phenotypes. ([proteinatlas.org](https://www.proteinatlas.org/ENSG00000148826-NKX6-2)) [S15]

This makes biological sense: a transcriptional repressor expressed in **mature oligodendrocytes** and **remyelinating oligodendrocytes** would be expected to help coordinate the gene programs needed for **myelin assembly, maintenance, and remodeling**. Human biallelic loss-of-function phenotypes provide strong in vivo confirmation that this regulatory role is clinically meaningful. ([pmc.ncbi.nlm.nih.gov](https://pmc.ncbi.nlm.nih.gov/articles/PMC2807475/)) [S16]

## Bottom line

Yes. NKX6-2 is **strongly associated with regulatory biological processes in neural systems**, and there is also credible evidence for a **myelin-centered homeostatic/maintenance role** in the CNS. The safest precise phrasing is: **NKX6-2 is a neural transcriptional regulator involved in ventral neural patterning, oligodendrocyte differentiation, myelin gene regulation, and maintenance of paranodal/myelin integrity**. ([sciencedirect.com](https://www.sciencedirect.com/science/article/pii/S0896627301004123)) [S17]

## Cited sources

- **[S1]** NKX6-2 protein expression summary - The Human Protein Atlas — https://www.proteinatlas.org/ENSG00000148826-NKX6-2
- **[S2]** 
            Co-localization of Nkx6.2 and Nkx2.2 Homeodomain Proteins in Differentiated Myelinating Oligodendrocytes - PMC
         — https://pmc.ncbi.nlm.nih.gov/articles/PMC2807475/
- **[S3]** NKX6-2 protein expression summary - The Human Protein Atlas — https://www.proteinatlas.org/ENSG00000148826-NKX6-2
- **[S4]** Different Levels of Repressor Activity Assign Redundant and Specific Roles to Nkx6 Genes in Motor Neuron and Interneuron Specification - ScienceDirect — https://www.sciencedirect.com/science/article/pii/S0896627301004123
- **[S5]** 
            Co-localization of Nkx6.2 and Nkx2.2 Homeodomain Proteins in Differentiated Myelinating Oligodendrocytes - PMC
         — https://pmc.ncbi.nlm.nih.gov/articles/PMC2807475/
- **[S6]** 
            CNS Myelin Paranodes Require Nkx6-2 Homeoprotein Transcriptional Activity for Normal Structure - PMC
         — https://pmc.ncbi.nlm.nih.gov/articles/PMC6730372/
- **[S7]** Untitled source — https://academic.oup.com/brain/article/140/10/2550/4085303
- **[S8]** 
            Mutations in NKX6-2 Cause Progressive Spastic Ataxia and Hypomyelination - PMC
         — https://pmc.ncbi.nlm.nih.gov/articles/PMC5473715/
- **[S9]** Onecut-dependent Nkx6.2 transcription factor expression is required for proper formation and activity of spinal locomotor circuits - PubMed — https://pubmed.ncbi.nlm.nih.gov/31969659/
- **[S10]** Characterization of Nkx6-2-derived neocortical interneuron lineages - PubMed — https://pubmed.ncbi.nlm.nih.gov/19363146/
- **[S11]** Mice lacking the Nkx6.2 (Gtx) homeodomain transcription factor develop and reproduce normally - PubMed — https://pubmed.ncbi.nlm.nih.gov/11390667/
- **[S12]** Characterization of Nkx6-2-derived neocortical interneuron lineages - PubMed — https://pubmed.ncbi.nlm.nih.gov/19363146/
- **[S13]** 
            CNS Myelin Paranodes Require Nkx6-2 Homeoprotein Transcriptional Activity for Normal Structure - PMC
         — https://pmc.ncbi.nlm.nih.gov/articles/PMC6730372/
- **[S14]** NKX6-2 protein expression summary - The Human Protein Atlas — https://www.proteinatlas.org/ENSG00000148826-NKX6-2
- **[S15]** NKX6-2 protein expression summary - The Human Protein Atlas — https://www.proteinatlas.org/ENSG00000148826-NKX6-2
- **[S16]** 
            Co-localization of Nkx6.2 and Nkx2.2 Homeodomain Proteins in Differentiated Myelinating Oligodendrocytes - PMC
         — https://pmc.ncbi.nlm.nih.gov/articles/PMC2807475/
- **[S17]** Different Levels of Repressor Activity Assign Redundant and Specific Roles to Nkx6 Genes in Motor Neuron and Interneuron Specification - ScienceDirect — https://www.sciencedirect.com/science/article/pii/S0896627301004123
