## Supplementary material for "Understanding Substance Dependence: What Differentiates Addictive from Non-Addictive Drugs?": Folder for supplementary files: non-addictive-genes-claims-first100-069-fyn-gene-protein-has-been-associated-with-regulato-3e6094741ae5.md

# non-addictive-genes-claims-first100-069-3e6094741ae5

**Original claim:** FYN gene/protein has been associated with regulatory or homeostatic biological processes in neural systems.

## Claim

FYN gene/protein has been associated with regulatory or homeostatic biological processes in neural systems.

## Verdict

Verdict: Likely True

FYN is well supported as a regulator of multiple neural biological processes, including synaptic receptor regulation, synaptic plasticity, oligodendrocyte maturation, and CNS myelination. The claim is strongest for **regulatory** roles; evidence for a specifically named **homeostatic** role exists mainly by inference from receptor trafficking, excitation/inhibition tuning, and myelin maintenance rather than from studies explicitly labeling FYN as a core “homeostasis” gene. ([pmc.ncbi.nlm.nih.gov](https://pmc.ncbi.nlm.nih.gov/articles/PMC4713834/)) [S1]

## Confidence

High

Multiple independent lines of evidence converge: expression/localization data place FYN in neural cell types and synaptic compartments; mouse loss- and gain-of-function studies show effects on myelination and synaptic plasticity; and human iPSC-derived neuron work shows Fyn-dependent regulation of NMDA receptor function. The main uncertainty is the breadth of the term “homeostatic,” not whether FYN participates in neural regulation. ([proteinatlas.org](https://www.proteinatlas.org/ENSG00000010810-FYN)) [S2]

## Supporting evidence

**Neural expression and compartment association.** Human Protein Atlas data place **FYN** in a brain-related “neuronal signaling” expression cluster and show enrichment in neural-supporting cell populations including astrocytes and oligodendrocyte-lineage cells. Complementary NCBI/Alliance annotations for the rodent ortholog place Fyn in dendrites, glutamatergic synapses, and postsynaptic density, which are classic sites of neural signal regulation. ([proteinatlas.org](https://www.proteinatlas.org/ENSG00000010810-FYN)) [S3]

**Broad review-level synthesis from nervous-system studies.** A focused review of Fyn in the nervous system summarizes transgenic mouse work showing that Fyn has been implicated in neuronal migration, myelination, synaptic plasticity, and regulation of excitatory and inhibitory receptors. That review also notes that Fyn is expressed in both neurons and glia and that its expression/activity rise during development, supporting an established role in neural regulatory biology rather than an isolated disease-only association. ([pmc.ncbi.nlm.nih.gov](https://pmc.ncbi.nlm.nih.gov/articles/PMC4713834/)) [S4]

**Strong animal evidence for glial regulation and myelination.** In a classic knockout study, **fyn−/−** mice showed reduced oligodendrocyte number in corpus callosum, reduced forebrain myelin content, and reduced myelin sheaths; kinase-dead Fyn mutants showed a similar myelination deficit, arguing that Fyn kinase activity is required for normal CNS myelination. This is direct evidence that Fyn regulates a major neural-support process with clear developmental and maintenance implications. ([pmc.ncbi.nlm.nih.gov](https://pmc.ncbi.nlm.nih.gov/articles/PMC6762613/)) [S5]

**Mechanistically linked myelination signaling.** A later study found that BDNF/TrkB signaling uses Fyn as an intermediate kinase in oligodendrocytes: inhibiting Src-family activity blocked the promyelinating effect of BDNF, overexpressing Fyn promoted ERK1/2 phosphorylation and myelination, and kinase-dead Fyn prevented BDNF’s promyelinating effect. That makes Fyn not just correlated with, but functionally embedded in, a neural regulatory pathway controlling oligodendrocyte maturation/myelination. ([pubmed.ncbi.nlm.nih.gov](https://pubmed.ncbi.nlm.nih.gov/26449489/)) [S6]

**Synaptic plasticity evidence in vivo.** The same nervous-system review summarizes that Fyn knockout mice have impaired hippocampal long-term potentiation (LTP) under weak tetanic stimulation and impaired spatial memory, while constitutively active Fyn lowers the threshold for LTP induction. A rescue study then showed that reintroducing Fyn in adult forebrain neurons restored Schaffer collateral LTP in fyn-deficient mice, supporting a direct role in the molecular regulation of synaptic plasticity rather than only a secondary developmental effect. ([pmc.ncbi.nlm.nih.gov](https://pmc.ncbi.nlm.nih.gov/articles/PMC4713834/)) [S7]

**Regulation of excitatory and inhibitory receptor trafficking/signaling.** The review further describes Fyn phosphorylation of NMDA receptor subunit GluN2B/NR2B and of the GABA\(_A\) receptor γ2 subunit. In that synthesis, Fyn-dependent phosphorylation of NR2B Y1472 is linked to reduced clathrin-mediated endocytosis and altered receptor complex signaling, while phosphorylation of GABA\(_A\) receptor γ2 is linked to reduced endocytosis and enhanced synaptic inhibition. Those are textbook examples of neural regulatory processes and are relevant to synaptic stability and network balance. ([pmc.ncbi.nlm.nih.gov](https://pmc.ncbi.nlm.nih.gov/articles/PMC4713834/)) [S8]

**Human-neuron evidence.** In human iPSC-derived neurons, Fyn-interfering peptide suppressed NMDA receptor currents, and the authors concluded that Fyn regulates GluN2B-dominant NMDAR function in human neurons. They explicitly interpreted this as conservation of a core molecular process related to synaptic metaplasticity. This is important because it moves the evidence beyond rodent-only systems. ([pmc.ncbi.nlm.nih.gov](https://pmc.ncbi.nlm.nih.gov/articles/PMC4819183/)) [S9]

## Contradicting or limiting evidence

The biggest limitation is **terminology**. Much of the literature describes FYN in terms of **synaptic plasticity, receptor regulation, neuronal migration, oligodendrocyte maturation, and myelination**, not explicitly as a canonical “homeostasis” factor. So the statement is more secure if read as “FYN is associated with neural regulatory processes,” and somewhat less secure if read as “FYN is a central, directly demonstrated controller of neural homeostasis.” ([pmc.ncbi.nlm.nih.gov](https://pmc.ncbi.nlm.nih.gov/articles/PMC4713834/)) [S10]

There is also evidence of **context dependence and redundancy**. The review notes that LTP deficits in Fyn knockout mice can be masked earlier in life by compensatory Src-family signaling, implying that Fyn is important but not always uniquely required. Likewise, the myelination phenotype is regionally selective: forebrain/corpus callosum is strongly affected, whereas spinal cord myelination is relatively preserved in the knockout. That argues against an overly broad claim that Fyn uniformly governs all neural regulatory processes. ([pmc.ncbi.nlm.nih.gov](https://pmc.ncbi.nlm.nih.gov/articles/PMC4713834/)) [S11]

Finally, direct **human in vivo** evidence remains limited. The strongest human-relevant functional evidence here comes from iPSC-derived neurons rather than from interventional studies in human brains. So the association is biologically credible and experimentally supported, but much of the causal evidence still comes from mouse and cell-culture systems. ([pmc.ncbi.nlm.nih.gov](https://pmc.ncbi.nlm.nih.gov/articles/PMC4819183/)) [S12]

## Mechanistic / biological rationale

Fyn is a **Src-family non-receptor tyrosine kinase**, a protein class well suited to regulatory signaling because it can rapidly modify downstream proteins by phosphorylation. In the nervous system, Fyn activity is itself controlled by regulatory tyrosines: phosphorylation at the C-terminal inhibitory site favors an inactive conformation, whereas phosphorylation in the activation loop promotes catalytic activity and protein interactions. This is exactly the sort of architecture expected for a molecular regulator rather than a static structural protein. ([pmc.ncbi.nlm.nih.gov](https://pmc.ncbi.nlm.nih.gov/articles/PMC4713834/)) [S13]

In neurons, Fyn helps regulate **synaptic receptor composition and signaling strength**. Review-level evidence indicates that Fyn phosphorylation of NMDA receptor components can stabilize synaptic NMDARs and alter downstream signaling, while phosphorylation of GABA\(_A\) receptor components can reduce receptor endocytosis and enhance inhibitory transmission. Conceptually, those actions could support network stability by tuning excitation and inhibition, which is why a “homeostatic” interpretation is biologically plausible even when many papers use the word “regulatory” or “plasticity” instead. ([pmc.ncbi.nlm.nih.gov](https://pmc.ncbi.nlm.nih.gov/articles/PMC4713834/)) [S14]

In oligodendrocytes, Fyn appears to integrate extracellular cues such as **BDNF/TrkB**, adhesion/integrin signals, and intracellular ERK/cytoskeletal/MBP-mRNA pathways to drive maturation and myelin production. Because myelin is critical for conduction timing and circuit reliability, this provides a strong rationale for why Fyn would be relevant to neural system regulation and maintenance. ([pubmed.ncbi.nlm.nih.gov](https://pubmed.ncbi.nlm.nih.gov/26449489/)) [S15]

## Bottom line

The claim is **Likely True**. FYN is convincingly associated with **neural regulatory processes**, especially synaptic plasticity/receptor regulation and oligodendrocyte-dependent myelination. Calling it a participant in broader neural **homeostatic** biology is reasonable, but that part is somewhat more inferential and less directly named in the literature than its regulatory roles. ([pmc.ncbi.nlm.nih.gov](https://pmc.ncbi.nlm.nih.gov/articles/PMC4713834/)) [S16]

## Cited sources

- **[S1]** 
            Fyn in Neurodevelopment and Ischemic Brain Injury - PMC
         — https://pmc.ncbi.nlm.nih.gov/articles/PMC4713834/
- **[S2]** FYN protein expression summary - The Human Protein Atlas — https://www.proteinatlas.org/ENSG00000010810-FYN
- **[S3]** FYN protein expression summary - The Human Protein Atlas — https://www.proteinatlas.org/ENSG00000010810-FYN
- **[S4]** 
            Fyn in Neurodevelopment and Ischemic Brain Injury - PMC
         — https://pmc.ncbi.nlm.nih.gov/articles/PMC4713834/
- **[S5]** 
            A Unique Role for Fyn in CNS Myelination - PMC
         — https://pmc.ncbi.nlm.nih.gov/articles/PMC6762613/
- **[S6]** Fyn is an intermediate kinase that BDNF utilizes to promote oligodendrocyte myelination - PubMed — https://pubmed.ncbi.nlm.nih.gov/26449489/
- **[S7]** 
            Fyn in Neurodevelopment and Ischemic Brain Injury - PMC
         — https://pmc.ncbi.nlm.nih.gov/articles/PMC4713834/
- **[S8]** 
            Fyn in Neurodevelopment and Ischemic Brain Injury - PMC
         — https://pmc.ncbi.nlm.nih.gov/articles/PMC4713834/
- **[S9]** 
            Fyn Kinase regulates GluN2B subunit-dominant NMDA receptors in human induced pluripotent stem cell-derived neurons - PMC
         — https://pmc.ncbi.nlm.nih.gov/articles/PMC4819183/
- **[S10]** 
            Fyn in Neurodevelopment and Ischemic Brain Injury - PMC
         — https://pmc.ncbi.nlm.nih.gov/articles/PMC4713834/
- **[S11]** 
            Fyn in Neurodevelopment and Ischemic Brain Injury - PMC
         — https://pmc.ncbi.nlm.nih.gov/articles/PMC4713834/
- **[S12]** 
            Fyn Kinase regulates GluN2B subunit-dominant NMDA receptors in human induced pluripotent stem cell-derived neurons - PMC
         — https://pmc.ncbi.nlm.nih.gov/articles/PMC4819183/
- **[S13]** 
            Fyn in Neurodevelopment and Ischemic Brain Injury - PMC
         — https://pmc.ncbi.nlm.nih.gov/articles/PMC4713834/
- **[S14]** 
            Fyn in Neurodevelopment and Ischemic Brain Injury - PMC
         — https://pmc.ncbi.nlm.nih.gov/articles/PMC4713834/
- **[S15]** Fyn is an intermediate kinase that BDNF utilizes to promote oligodendrocyte myelination - PubMed — https://pubmed.ncbi.nlm.nih.gov/26449489/
- **[S16]** 
            Fyn in Neurodevelopment and Ischemic Brain Injury - PMC
         — https://pmc.ncbi.nlm.nih.gov/articles/PMC4713834/
