## Supplementary material for "Understanding Substance Dependence: What Differentiates Addictive from Non-Addictive Drugs?": Folder for supplementary files: non-addictive-genes-claims-first100-070-pink1-gene-protein-has-been-associated-with-regula-4d5d7ef6fcd3.md

# non-addictive-genes-claims-first100-070-4d5d7ef6fcd3

**Original claim:** PINK1 gene/protein has been associated with regulatory or homeostatic biological processes in neural systems.

## Claim

PINK1 gene/protein has been associated with regulatory or homeostatic biological processes in neural systems. ([pmc.ncbi.nlm.nih.gov](https://pmc.ncbi.nlm.nih.gov/articles/PMC7071653/)) [S1]

## Verdict

Verdict: Highly Supported

## Confidence

High

## Supporting evidence

The claim is strongly supported by multiple lines of evidence spanning human genetics, human neuron models, animal studies, and mechanistic reviews. In humans, **biallelic PINK1 mutations are an established cause of young-onset Parkinson disease**, and neuropathology in reported cases shows characteristic **neuronal loss in the substantia nigra**. GeneReviews also summarizes evidence for altered synaptic transmission and synaptic plasticity in PINK1-linked disease models, which directly ties PINK1 to neural regulatory processes rather than to a generic cellular stress phenotype alone. ([ncbi.nlm.nih.gov](https://www.ncbi.nlm.nih.gov/sites/books/NBK26472/)) [S2]

At the cellular level, PINK1 is one of the best-studied regulators of **mitochondrial quality control**, a core homeostatic process that is especially important in neurons because of their high energetic demand and extended morphology. Reviews of the PINK1/Parkin pathway describe PINK1 accumulation on damaged mitochondria, phosphorylation of ubiquitin/Parkin, and downstream control of mitochondrial fission, cargo selection, and autophagic clearance. These reviews explicitly frame PINK1/Parkin as “core organizers” of mitochondrial quality control and mitochondrial surveillance in the nervous system. ([pmc.ncbi.nlm.nih.gov](https://pmc.ncbi.nlm.nih.gov/articles/PMC7071653/)) [S3]

This association is not limited to abstract pathway diagrams. In **primary mammalian hippocampal and dopaminergic neurons**, manipulation of the PINK1/Parkin pathway alters mitochondrial dynamics, dendritic spine number, and neuronal vulnerability to excitotoxicity. In that study, PINK1 overexpression or knockdown had opposite effects on spine formation, and the authors concluded that altered PINK1/Parkin activity affects **synaptic morphogenesis** in neurons. That is direct evidence for a regulatory role in neural structure/function. ([pmc.ncbi.nlm.nih.gov](https://pmc.ncbi.nlm.nih.gov/articles/PMC3140825/)) [S4]

Additional neuronal evidence comes from studies of dendrites and spines. In mouse cortical neurons and brain tissue, **PINK1 deficiency simplified dendritic architecture, reduced synaptic input, and lowered spine density/maturation**, and these defects were reversible by restoring human PINK1 or downstream mediators. Independent work also showed that PINK1 interacts with PKA/VCP-p47 signaling to promote dendritic arborization, further supporting a role in maintaining neuronal architecture. ([pmc.ncbi.nlm.nih.gov](https://pmc.ncbi.nlm.nih.gov/articles/PMC9581559/)) [S5]

PINK1 has also been linked to **neuronal calcium and energetic homeostasis**, which are canonical homeostatic functions in excitable cells. In PINK1-depleted or knockout cells, investigators found reduced mitochondrial calcium uptake after physiological stimulation, impaired mitochondrial ATP synthesis, and altered cytosolic calcium extrusion; the energetic defect was also confirmed in the brain of PINK1-knockout mice. A more recent study further linked the PINK1/Parkin pathway to regulation of ER/cytosolic calcium homeostasis through CISD1 and IP3R signaling. ([pubmed.ncbi.nlm.nih.gov](https://pubmed.ncbi.nlm.nih.gov/21385841/)) [S6]

Evidence also extends beyond neurons to **glia**, which are central to neural homeostasis. A PINK1-deficiency study in glial systems found abnormal innate immune responses and increased inflammation-induced nitric oxide-mediated neuron death. A later review summarized that PINK1-deficient astrocytes show impaired mitochondrial function, higher ROS/NO-related responses, and altered cytokine signaling, while microglia show reduced IL-10 secretion. These findings support a role for PINK1 in **neuroimmune regulation** and in maintaining a glial environment that protects neurons. ([pmc.ncbi.nlm.nih.gov](https://pmc.ncbi.nlm.nih.gov/articles/PMC5762685/?utm_source=openai)) [S7]

There is also evidence for non-canonical, non-mitophagic neural regulation by **processed/cleaved PINK1**. Reviews and primary studies report that cytosolic or cleaved PINK1 can regulate neuronal development, neurite/dendrite maintenance, and neuronal plasticity through **PKA-CREB-BDNF** signaling. In cultured primary neurons, endogenous PINK1 activation increased PKA activity, CREB activation, BDNF production/release, and TRKB signaling, supporting a plausible role in neuronal differentiation and plasticity. ([frontiersin.org](https://www.frontiersin.org/articles/10.3389/fnins.2021.769331/full)) [S8]

Finally, newer human-cell work strengthens the neural-homeostasis association. In **PINK1-deficient human iPSC-derived dopaminergic neurons**, investigators found markedly reduced mitochondrial membrane potential, reduced mitophagy, reduced mitochondrial biogenesis, lower PGC-1α, and lower TH expression, with selective effects in TH-positive neurons. That is highly relevant evidence that PINK1 helps maintain mitochondrial and dopaminergic neuronal homeostasis in a human neural context. ([pmc.ncbi.nlm.nih.gov](https://pmc.ncbi.nlm.nih.gov/articles/PMC10327008/)) [S9]

## Contradicting or limiting evidence

The main limitation is not whether PINK1 is associated with neural homeostasis—it clearly is—but **how universal or central that role is in every neural context**. For example, in vivo mouse reporter work found that **basal mitophagy in several tissues, including dopaminergic neurons and microglia, can occur independently of Pink1**, even though stress-evoked Parkin activation remained PINK1-dependent. This means PINK1 is not required for all forms of basal mitochondrial turnover in the nervous system. ([pmc.ncbi.nlm.nih.gov](https://pmc.ncbi.nlm.nih.gov/articles/PMC5807059/)) [S10]

Likewise, some neuron studies have suggested that classical depolarization-triggered PINK1/Parkin mitophagy may behave differently in neurons than in transformed cell lines. One mammalian neuron study explicitly noted that it did **not** observe convincing mitophagy in cultured hippocampal neurons under its conditions and suggested that additional neuron-specific mechanisms may regulate mitochondrial quality control. ([pmc.ncbi.nlm.nih.gov](https://pmc.ncbi.nlm.nih.gov/articles/PMC3140825/)) [S11]

There is also an important species/context caveat. A primate study reported that **PINK1 loss caused neurodegeneration without obvious changes in mitochondrial protein expression or morphology**, and the authors argued that PINK1 kinase activity affecting broader protein phosphorylation may be more critical for neuronal survival in primates than the classical mitochondrial-homeostasis model alone. This does not contradict the claim, but it means the biology may be broader than “PINK1 = mitophagy” and that rodent/non-neuronal models do not capture everything. ([pubmed.ncbi.nlm.nih.gov](https://pubmed.ncbi.nlm.nih.gov/34800266/)) [S12]

So the limiting evidence mainly narrows the interpretation: PINK1 is very clearly associated with neural regulatory/homeostatic processes, but **the exact dominant process depends on cell type, species, and stress state**. ([pmc.ncbi.nlm.nih.gov](https://pmc.ncbi.nlm.nih.gov/articles/PMC5807059/)) [S13]

## Mechanistic / biological rationale

A strong biological rationale exists for PINK1 involvement in neural homeostasis. Under mitochondrial damage, PINK1 accumulates on the outer mitochondrial membrane and activates Parkin by phosphorylating ubiquitin and Parkin, initiating selective handling of dysfunctional mitochondria. Downstream consequences include changes in fission/fusion balance, ubiquitination of outer-membrane substrates, recruitment of autophagy receptors, and mitochondrial turnover or remodeling. These are classic organelle homeostasis functions. ([pmc.ncbi.nlm.nih.gov](https://pmc.ncbi.nlm.nih.gov/articles/PMC7071653/)) [S14]

In neurons, that matters especially because synapses and long neurites require local ATP production, calcium buffering, and controlled mitochondrial trafficking. Consistent with this, PINK1 influences mitochondrial dynamics in hippocampal and dopaminergic neurons, affects dendritic spine morphology, and modulates calcium handling and ATP maintenance. These functions fit a homeostatic model in which PINK1 helps keep neuronal energetics, signaling, and structure within functional bounds. ([pmc.ncbi.nlm.nih.gov](https://pmc.ncbi.nlm.nih.gov/articles/PMC3140825/)) [S15]

PINK1 also appears to have **non-canonical signaling roles**. Cleaved/cytosolic PINK1 can activate PKA-linked pathways that promote dendritic arborization and BDNF-associated neuronal development/plasticity. In glia, PINK1 restrains excessive inflammatory signaling and thereby helps preserve a neuroprotective environment. Thus, the mechanistic picture is broader than mitochondrial disposal alone: PINK1 participates in **organelle quality control, calcium/energy regulation, dendrite/synapse maintenance, trophic signaling, and inflammatory restraint** in neural cells. ([pmc.ncbi.nlm.nih.gov](https://pmc.ncbi.nlm.nih.gov/articles/PMC6377406/)) [S16]

## Bottom line

Yes. The statement is **highly supported**. PINK1 is robustly associated with multiple **regulatory/homeostatic processes in neural systems**, especially mitochondrial quality control, calcium/energy homeostasis, dendritic and synaptic maintenance, and glial inflammatory regulation. The strongest caveat is that its role is **context-dependent**: not every form of basal mitophagy or every neuronal phenotype is strictly PINK1-dependent, and primate data suggest broader kinase-signaling roles beyond classical mitophagy. But on the core claim—that PINK1 is associated with regulatory or homeostatic biology in neural systems—the evidence is strong and convergent. ([ncbi.nlm.nih.gov](https://www.ncbi.nlm.nih.gov/sites/books/NBK26472/)) [S17]

## Cited sources

- **[S1]** 
            PINK1 and Parkin mitochondrial quality control: a source of regional vulnerability in Parkinson’s disease - PMC
         — https://pmc.ncbi.nlm.nih.gov/articles/PMC7071653/
- **[S2]** PINK1 Type of Young-Onset Parkinson Disease - GeneReviews® - NCBI Bookshelf — https://www.ncbi.nlm.nih.gov/sites/books/NBK26472/
- **[S3]** 
            PINK1 and Parkin mitochondrial quality control: a source of regional vulnerability in Parkinson’s disease - PMC
         — https://pmc.ncbi.nlm.nih.gov/articles/PMC7071653/
- **[S4]** 
            The PINK1/Parkin pathway regulates mitochondrial dynamics and function in mammalian hippocampal and dopaminergic neurons - PMC
         — https://pmc.ncbi.nlm.nih.gov/articles/PMC3140825/
- **[S5]** 
            Endogenous PTEN-Induced Kinase 1 Regulates Dendritic Architecture and Spinogenesis - PMC
         — https://pmc.ncbi.nlm.nih.gov/articles/PMC9581559/
- **[S6]** Depletion of PINK1 affects mitochondrial metabolism, calcium homeostasis and energy maintenance - PubMed — https://pubmed.ncbi.nlm.nih.gov/21385841/
- **[S7]** Lack of PINK1 alters glia innate immune responses and enhances inflammation-induced, nitric oxide-mediated neuron death - PMC — https://pmc.ncbi.nlm.nih.gov/articles/PMC5762685/?utm_source=openai
- **[S8]** Frontiers | Role of Cleaved PINK1 in Neuronal Development, Synaptogenesis, and Plasticity: Implications for Parkinson’s Disease — https://www.frontiersin.org/articles/10.3389/fnins.2021.769331/full
- **[S9]** 
            Defects in Mitochondrial Biogenesis Drive Mitochondrial Alterations in PINK1-deficient Human Dopamine Neurons - PMC
         — https://pmc.ncbi.nlm.nih.gov/articles/PMC10327008/
- **[S10]** 
            Basal Mitophagy Occurs Independently of PINK1 in Mouse Tissues of High Metabolic Demand - PMC
         — https://pmc.ncbi.nlm.nih.gov/articles/PMC5807059/
- **[S11]** 
            The PINK1/Parkin pathway regulates mitochondrial dynamics and function in mammalian hippocampal and dopaminergic neurons - PMC
         — https://pmc.ncbi.nlm.nih.gov/articles/PMC3140825/
- **[S12]** PINK1 kinase dysfunction triggers neurodegeneration in the primate brain without impacting mitochondrial homeostasis - PubMed — https://pubmed.ncbi.nlm.nih.gov/34800266/
- **[S13]** 
            Basal Mitophagy Occurs Independently of PINK1 in Mouse Tissues of High Metabolic Demand - PMC
         — https://pmc.ncbi.nlm.nih.gov/articles/PMC5807059/
- **[S14]** 
            PINK1 and Parkin mitochondrial quality control: a source of regional vulnerability in Parkinson’s disease - PMC
         — https://pmc.ncbi.nlm.nih.gov/articles/PMC7071653/
- **[S15]** 
            The PINK1/Parkin pathway regulates mitochondrial dynamics and function in mammalian hippocampal and dopaminergic neurons - PMC
         — https://pmc.ncbi.nlm.nih.gov/articles/PMC3140825/
- **[S16]** 
            PINK1 Interacts with VCP/p97 and Activates PKA to Promote NSFL1C/p47 Phosphorylation and Dendritic Arborization in Neurons - PMC
         — https://pmc.ncbi.nlm.nih.gov/articles/PMC6377406/
- **[S17]** PINK1 Type of Young-Onset Parkinson Disease - GeneReviews® - NCBI Bookshelf — https://www.ncbi.nlm.nih.gov/sites/books/NBK26472/
