## Supplementary material for "Understanding Substance Dependence: What Differentiates Addictive from Non-Addictive Drugs?": Folder for supplementary files: non-addictive-genes-claims-first100-071-hibch-gene-protein-has-been-associated-with-regula-4cc626c9df6c.md

# non-addictive-genes-claims-first100-071-4cc626c9df6c

**Original claim:** HIBCH gene/protein has been associated with regulatory or homeostatic biological processes in neural systems.

## Claim

HIBCH gene/protein has been associated with regulatory or homeostatic biological processes in neural systems.

## Verdict

Verdict: Mixed Evidence

HIBCH is **clearly associated with neural disease and neural vulnerability**, but the evidence mostly points to an **indirect metabolic/mitochondrial housekeeping role** rather than a well-established, primary role as a neural regulatory or homeostatic factor. Curated gene annotations emphasize **mitochondrial hydrolase activity** and **L-valine catabolism**, not canonical neural homeostasis pathways. Still, human genetic disease, brain imaging, treatment response, and one disease-context experimental study do support an association with **neuronal stability/metabolic homeostasis** in the nervous system. ([ncbi.nlm.nih.gov](https://www.ncbi.nlm.nih.gov/gene/26275)) [S1]

## Confidence

Medium

The basic enzymatic function of HIBCH is well established, and the human neurologic phenotype of HIBCH deficiency is well documented. What is less certain is whether that should be counted as a bona fide **“regulatory or homeostatic biological process in neural systems”** versus a more general mitochondrial metabolic role whose failure secondarily injures the brain. ([ncbi.nlm.nih.gov](https://www.ncbi.nlm.nih.gov/gene/26275)) [S2]

## Supporting evidence

HIBCH encodes a **mitochondrial 3-hydroxyisobutyryl-CoA hydrolase** involved in valine catabolism; NCBI’s curated function/process display highlights **3-hydroxyisobutyryl-CoA hydrolase activity** and involvement in the **L-valine catabolic process**. That places HIBCH in core metabolic maintenance rather than neurotransmission or synaptic signaling per se. ([ncbi.nlm.nih.gov](https://www.ncbi.nlm.nih.gov/gene/26275?utm_source=openai)) [S3]

HIBCH is expressed in the brain. Human Protein Atlas data show **low tissue specificity**, **low regional specificity**, and detection across **all major brain regions**; in brain RNA clustering it falls into an **oligodendrocyte-associated mixed-function cluster** rather than a sharply neuron-specific pattern. This is consistent with a broadly required metabolic enzyme that could contribute to neural cell homeostasis. ([proteinatlas.org](https://www.proteinatlas.org/ENSG00000198130-HIBCH/brain)) [S4]

Human loss-of-function evidence strongly links HIBCH to neurologic integrity. GeneReviews states that HIBCH deficiency commonly presents in infancy or childhood with **developmental delay/regression, hypotonia, seizures, movement disorders, vision impairment, episodes of neurologic deterioration**, and characteristic **brain MRI lesions** involving the basal ganglia, white matter, cerebellum, and brainstem. ([ncbi.nlm.nih.gov](https://www.ncbi.nlm.nih.gov/books/NBK619246)) [S5]

Early case reports likewise showed that HIBCH deficiency causes **progressive infantile neurodegeneration** with Leigh-like basal ganglia changes. In the 2007 report, both affected patients had hypotonia, feeding problems, motor delay, and neurologic regression, with deficient HIBCH enzyme activity and little to no HIBCH protein. ([pmc.ncbi.nlm.nih.gov](https://pmc.ncbi.nlm.nih.gov/articles/PMC1785315/)) [S6]

Subsequent work expanded the neurologic association. A 2020 multicenter study of ECHS1/HIBCH-related disease found that HIBCH defects were recurrent causes of **Leigh syndrome-like neurologic disease**, with **basal ganglia lesions** and genotype-phenotype correlations across a large pooled cohort. ([pubmed.ncbi.nlm.nih.gov](https://pubmed.ncbi.nlm.nih.gov/32677093/)) [S7]

There is also some more direct “neural stability/homeostasis” language in experimental work. In a presynaptic proteomics study of juvenile neuronal ceroid lipofuscinosis (CLN3 disease), HIBCH protein was reduced in vulnerable synaptic fractions in mouse thalamus and in postmortem human JNCL cortex. In the accompanying Drosophila screen, HIBCH knockdown **enhanced CLN3-dependent neurodegeneration**, leading the authors to conclude that HIBCH can modify **neuronal stability** in that disease context. ([pmc.ncbi.nlm.nih.gov](https://pmc.ncbi.nlm.nih.gov/articles/PMC5622084/)) [S8]

Clinical response to pathway-directed treatment also supports a role in maintaining neural metabolic stability. Case reports and series describe **clinical and neuroradiologic improvement** after a **valine-restricted diet** in some HIBCH-deficient patients, including improvement in dystonia and brain lesions. ([pubmed.ncbi.nlm.nih.gov](https://pubmed.ncbi.nlm.nih.gov/31679561/)) [S9]

## Contradicting or limiting evidence

The strongest limitation is that **HIBCH is not primarily annotated as a neural regulatory/homeostatic gene**. The best-curated functional sources emphasize **mitochondrial metabolism** and **valine catabolism**, not synaptic homeostasis, neuronal signaling, glial regulation, neurotransmitter balance, or other classic neural regulatory programs. ([ncbi.nlm.nih.gov](https://www.ncbi.nlm.nih.gov/gene/26275)) [S10]

Brain expression alone is not enough to establish a neural homeostatic function. The Human Protein Atlas profile shows HIBCH is **broadly expressed** with **low specificity** across tissues and brain regions, which fits a general housekeeping enzyme more than a dedicated neural regulator. ([proteinatlas.org](https://www.proteinatlas.org/ENSG00000198130-HIBCH/brain)) [S11]

The most direct “neuronal stability” evidence is limited and contextual. In the CLN3 study, HIBCH manipulation alone **did not** cause degeneration in flies without the CLN3 disease background; the authors explicitly interpreted HIBCH as a modifier of neuronal stability **in a CLN3-specific context**, not as a standalone general controller of neuronal homeostasis. ([pmc.ncbi.nlm.nih.gov](https://pmc.ncbi.nlm.nih.gov/articles/PMC5622084/)) [S12]

Mechanistic cell studies linking HIBCH loss to oxidative-metabolic defects were not performed in neurons. A 2025 mitochondrial CRISPR screen showed that **HIBCH knockout reduced oxygen consumption and cellular fitness**, but these experiments were done in **HeLa cells**, and the authors noted that such findings may be **cell-specific** and may not fully represent quiescent cells such as neurons or muscle. ([embopress.org](https://www.embopress.org/doi/10.1038/s44319-025-00459-9)) [S13]

So, the evidence is strongest for this narrower statement: **HIBCH is a mitochondrial valine-catabolism enzyme whose deficiency disrupts neural metabolic integrity and can contribute to neurodegeneration**. That is weaker than saying HIBCH is an established neural homeostatic regulator. ([ncbi.nlm.nih.gov](https://www.ncbi.nlm.nih.gov/books/NBK619246)) [S14]

## Mechanistic / biological rationale

HIBCH catalyzes a key step in valine degradation. Older biochemical work in human liver found high activity of methacrylyl-CoA hydratase and HIB-CoA hydrolase, suggesting these steps are important for handling the **potentially toxic intermediate methacrylyl-CoA**. ([journals.lww.com](https://journals.lww.com/hep/abstract/1996/12000/the_valine_catabolic_pathway_in_human_liver_.14.aspx)) [S15]

GeneReviews and case literature indicate that HIBCH deficiency leads to buildup of **3-hydroxyisobutyryl-CoA / hydroxy-C4-carnitine** and upstream **methacrylyl-CoA-derived toxic intermediates**. These reactive metabolites can form adducts with thiol-containing compounds, plausibly impairing mitochondrial enzymes and redox buffering systems. ([ncbi.nlm.nih.gov](https://www.ncbi.nlm.nih.gov/books/NBK619246)) [S16]

The 2013 HIBCH report proposed that depletion of **cysteine, glutathione, and thioredoxin** by methacrylyl-CoA could cause oxidative injury and secondary respiratory-chain dysfunction, helping explain the Leigh-like neurologic phenotype. ([pmc.ncbi.nlm.nih.gov](https://pmc.ncbi.nlm.nih.gov/articles/PMC4222069/?utm_source=openai)) [S17]

More recent functional work supports this broader mechanism. In a mitochondrial genetic screen, disruption at the ECHS1/HIBCH steps of valine catabolism impaired **oxygen consumption**, decreased **protein lipoylation**, reduced **PDH activity**, and compromised **oxidative metabolism/cellular fitness**. The authors proposed that reactive valine intermediates damage sulfur-dependent mitochondrial processes. ([embopress.org](https://www.embopress.org/doi/10.1038/s44319-025-00459-9)) [S18]

Because the brain is highly dependent on mitochondrial ATP production and is especially vulnerable to failures in oxidative metabolism, it is biologically plausible that HIBCH contributes to **neural homeostasis indirectly through metabolic maintenance** rather than through direct neural signaling or transcriptional regulation. ([embopress.org](https://www.embopress.org/doi/10.1038/s44319-025-00459-9)) [S19]

## Bottom line

HIBCH is **not well supported as a canonical neural regulatory/homeostatic gene** in the usual sense. Its established role is as a **mitochondrial valine-catabolism enzyme**. However, there is meaningful evidence that HIBCH is **associated with neural metabolic stability and neuronal vulnerability**, because HIBCH deficiency causes Leigh-like neurodegeneration, some patients improve with valine restriction, and one disease-model study identified HIBCH as a **modifier of neuronal stability** in context. Overall, the claim is best rated **Mixed Evidence**: **supported indirectly through neural metabolic homeostasis and disease association, but not strongly supported as a direct, primary neural homeostatic regulator.** ([ncbi.nlm.nih.gov](https://www.ncbi.nlm.nih.gov/gene/26275)) [S20]

## Cited sources

- **[S1]** HIBCH 3-hydroxyisobutyryl-CoA hydrolase [Homo sapiens (human)] - Gene - NCBI — https://www.ncbi.nlm.nih.gov/gene/26275
- **[S2]** HIBCH 3-hydroxyisobutyryl-CoA hydrolase [Homo sapiens (human)] - Gene - NCBI — https://www.ncbi.nlm.nih.gov/gene/26275
- **[S3]** HIBCH 3-hydroxyisobutyryl-CoA hydrolase [Homo sapiens (human)] - Gene - NCBI — https://www.ncbi.nlm.nih.gov/gene/26275?utm_source=openai
- **[S4]** Brain tissue expression of HIBCH - Summary - The Human Protein Atlas — https://www.proteinatlas.org/ENSG00000198130-HIBCH/brain
- **[S5]** 3-Hydroxyisobutyryl-CoA Hydrolase Deficiency - GeneReviews® - NCBI Bookshelf — https://www.ncbi.nlm.nih.gov/books/NBK619246
- **[S6]** 
            Mutations in the Gene Encoding 3-Hydroxyisobutyryl-CoA Hydrolase Results in Progressive Infantile Neurodegeneration - PMC
         — https://pmc.ncbi.nlm.nih.gov/articles/PMC1785315/
- **[S7]** Delineating the neurological phenotype in children with defects in the ECHS1 or HIBCH gene - PubMed — https://pubmed.ncbi.nlm.nih.gov/32677093/
- **[S8]** 
            Proteomic mapping of differentially vulnerable pre-synaptic populations identifies regulators of neuronal stability in vivo - PMC
         — https://pmc.ncbi.nlm.nih.gov/articles/PMC5622084/
- **[S9]** A therapeutic regimen for 3-hydroxyisobutyryl-CoA hydrolase deficiency with exercise-induced dystonia - PubMed — https://pubmed.ncbi.nlm.nih.gov/31679561/
- **[S10]** HIBCH 3-hydroxyisobutyryl-CoA hydrolase [Homo sapiens (human)] - Gene - NCBI — https://www.ncbi.nlm.nih.gov/gene/26275
- **[S11]** Brain tissue expression of HIBCH - Summary - The Human Protein Atlas — https://www.proteinatlas.org/ENSG00000198130-HIBCH/brain
- **[S12]** 
            Proteomic mapping of differentially vulnerable pre-synaptic populations identifies regulators of neuronal stability in vivo - PMC
         — https://pmc.ncbi.nlm.nih.gov/articles/PMC5622084/
- **[S13]** Time-resolved mitochondrial screen identifies regulatory components of oxidative metabolism | EMBO Reports | Springer Nature Link — https://www.embopress.org/doi/10.1038/s44319-025-00459-9
- **[S14]** 3-Hydroxyisobutyryl-CoA Hydrolase Deficiency - GeneReviews® - NCBI Bookshelf — https://www.ncbi.nlm.nih.gov/books/NBK619246
- **[S15]** 
	Hepatology
 — https://journals.lww.com/hep/abstract/1996/12000/the_valine_catabolic_pathway_in_human_liver_.14.aspx
- **[S16]** 3-Hydroxyisobutyryl-CoA Hydrolase Deficiency - GeneReviews® - NCBI Bookshelf — https://www.ncbi.nlm.nih.gov/books/NBK619246
- **[S17]** HIBCH mutations can cause Leigh-like disease with combined deficiency of multiple mitochondrial respiratory chain enzymes and pyruvate dehydrogenase - PMC — https://pmc.ncbi.nlm.nih.gov/articles/PMC4222069/?utm_source=openai
- **[S18]** Time-resolved mitochondrial screen identifies regulatory components of oxidative metabolism | EMBO Reports | Springer Nature Link — https://www.embopress.org/doi/10.1038/s44319-025-00459-9
- **[S19]** Time-resolved mitochondrial screen identifies regulatory components of oxidative metabolism | EMBO Reports | Springer Nature Link — https://www.embopress.org/doi/10.1038/s44319-025-00459-9
- **[S20]** HIBCH 3-hydroxyisobutyryl-CoA hydrolase [Homo sapiens (human)] - Gene - NCBI — https://www.ncbi.nlm.nih.gov/gene/26275
