## Supplementary material for "Understanding Substance Dependence: What Differentiates Addictive from Non-Addictive Drugs?": Folder for supplementary files: non-addictive-genes-claims-first100-072-mapk10-gene-protein-has-been-associated-with-regul-e63c844af216.md

# non-addictive-genes-claims-first100-072-e63c844af216

**Original claim:** MAPK10 gene/protein has been associated with regulatory or homeostatic biological processes in neural systems.

## Claim

MAPK10 gene/protein has been associated with regulatory or homeostatic biological processes in neural systems.

## Verdict

Verdict: Highly Supported

MAPK10 encodes JNK3, a neuron-enriched MAP kinase with substantial evidence linking it to **regulatory functions in neural systems**, including neuronal development, neurite growth/regeneration, synaptic organization, receptor trafficking, synaptic plasticity, and stress-response signaling in neurons. There is also **direct, though more context-specific, evidence for homeostatic roles**, especially in hypothalamic circuits controlling energy balance under metabolic stress. ([pmc.ncbi.nlm.nih.gov](https://pmc.ncbi.nlm.nih.gov/articles/PMC7464089/)) [S1]

## Confidence

High

The evidence comes from multiple levels: curated expression resources, human genetic observations, mouse loss-of-function studies, primary neuronal culture experiments, and review synthesis of the pathway literature. The main caveat is that “homeostatic” evidence is narrower than “regulatory” evidence and is strongest in specific neural contexts rather than across all neural systems. ([proteinatlas.org](https://www.proteinatlas.org/ENSG00000109339-MAPK10/brain)) [S2]

## Supporting evidence

- **Neuron-enriched / brain-focused expression:** JNK3 expression is much more restricted than JNK1/2 and is reported mainly in brain, heart, and testis; in neural tissue it is found in post-mitotic neurons and across multiple brain regions. Human Protein Atlas data also place MAPK10 in a neuronal “synaptic function” expression cluster and show broad detection across major brain regions. ([pmc.ncbi.nlm.nih.gov](https://pmc.ncbi.nlm.nih.gov/articles/PMC7464089/)) [S3]

- **Human neurodevelopmental association:** De novo MAPK10/JNK3 truncations in two unrelated patients were reported in association with intellectual disability and seizures, with the authors concluding this highlights a role for JNK3 in human brain development. A later review further notes that partial loss of JNK3 function alters regulation of postsynaptic proteins and is correlated with cognitive disorders. ([pubmed.ncbi.nlm.nih.gov](https://pubmed.ncbi.nlm.nih.gov/23329067/)) [S4]

- **Neuronal development / regeneration:** In primary midbrain dopaminergic neuron cultures, JNK3 knockdown reduced neurite outgrowth, impaired regeneration after lesion, and lowered neuronal viability; rescue by ectopic JNK3 expression supported a direct functional role. Review synthesis also describes JNK3 as participating in physiological neuronal processes such as development, regeneration, and differentiation/reprogramming. ([pmc.ncbi.nlm.nih.gov](https://pmc.ncbi.nlm.nih.gov/articles/PMC3178779/)) [S5]

- **Synaptic regulation and plasticity:** JNK signaling in neurons is linked to dendritic spine plasticity and stability, and review evidence specifically states that JNK3-mediated phosphorylation of PSD-95 can down-regulate AMPA and NMDA receptors, thereby negatively regulating synaptic plasticity. A disease-model study in primary neurons and APP/PS1-derived neurons found that suppressing JNK3 reduced AMPA receptor endocytosis and improved synaptic readouts, consistent with JNK3 acting as a regulator of postsynaptic signaling. ([pmc.ncbi.nlm.nih.gov](https://pmc.ncbi.nlm.nih.gov/articles/PMC7600688/)) [S6]

- **Homeostatic neural function in metabolic circuits:** In a well-cited eLife study, JNK3 deficiency in AgRP neurons increased excitatory signaling and promoted hyperphagia during high-fat feeding; the authors concluded that JNK3 is a key mediator of the **homeostatic regulation of energy balance in response to metabolic stress**. That is a direct example of MAPK10/JNK3 participating in a homeostatic neural process. ([elifesciences.org](https://elifesciences.org/articles/10031?utm_source=openai)) [S7]

- **Additional regulatory physiology:** Review evidence also describes JNK3 involvement in neuronal cell-volume regulatory signaling under osmotic challenge and in modulation of axonal branching/filopodial dynamics, both consistent with regulatory control of neuronal state and structure. ([pmc.ncbi.nlm.nih.gov](https://pmc.ncbi.nlm.nih.gov/articles/PMC7464089/)) [S8]

## Contradicting or limiting evidence

- **Stronger for “regulatory” than for broad “homeostatic”:** The literature more clearly supports MAPK10/JNK3 as a regulator of neural signaling, plasticity, development, and stress responses than as a universal neural homeostasis factor. Direct “homeostatic” demonstrations are comparatively fewer and are especially clear in hypothalamic metabolic circuits rather than across the entire nervous system. ([pmc.ncbi.nlm.nih.gov](https://pmc.ncbi.nlm.nih.gov/articles/PMC7600688/)) [S9]

- **Many studies are from animal or cellular systems:** Key evidence comes from mouse knockouts, cultured neurons, and disease/stress models. That is strong for biological association, but it means the breadth of MAPK10’s roles in normal human neural homeostasis is still not fully mapped. ([pmc.ncbi.nlm.nih.gov](https://pmc.ncbi.nlm.nih.gov/articles/PMC3178779/)) [S10]

- **Functional redundancy complicates interpretation:** Single Jnk3 knockout mice reportedly show no gross early structural brain abnormality, while stronger developmental phenotypes emerge with combined JNK isoform perturbation. This suggests redundancy with JNK1/JNK2 and means absence of a dramatic baseline phenotype does not imply lack of regulatory function. ([pmc.ncbi.nlm.nih.gov](https://pmc.ncbi.nlm.nih.gov/articles/PMC7464089/)) [S11]

- **A lot of MAPK10/JNK3 literature is pathology-centered:** ischemia, neurodegeneration, excitotoxicity, and apoptosis are major themes. Those studies still demonstrate regulatory roles in neurons, but they do not always establish that MAPK10 is a constitutive homeostatic controller in the same way as classic ion-homeostasis or synaptic-scaling genes. ([pmc.ncbi.nlm.nih.gov](https://pmc.ncbi.nlm.nih.gov/articles/PMC7600688/)) [S12]

## Mechanistic / biological rationale

MAPK10/JNK3 is a stress-activated serine/threonine kinase within the JNK MAPK cascade. Upstream kinases such as MKK4/MKK7 activate it, while scaffold proteins such as JIP-1 and β-arrestin2 help localize signaling modules to distinct neuronal compartments, including cytoplasm, nucleus, axon growth cones, and pre/postsynaptic regions. That compartmentalization is exactly what would be expected for a protein that helps regulate neural-state transitions rather than merely being a passive marker. ([pmc.ncbi.nlm.nih.gov](https://pmc.ncbi.nlm.nih.gov/articles/PMC7464089/)) [S13]

Mechanistically, JNK3 can regulate neurons at multiple levels:  
1. **Transcriptional regulation** via phosphorylation of c-Jun/AP-1 pathway components, influencing stress-response and survival programs.  
2. **Structural regulation** of neurites/axons and regeneration-related programs.  
3. **Synaptic regulation** via PSD-95-associated signaling and effects on AMPA/NMDA receptor abundance or trafficking, which changes synaptic strength and plasticity. ([pharos.nih.gov](https://pharos.nih.gov/targets/MAPK10?utm_source=openai)) [S14]

For the “homeostatic” part of the claim, the most convincing mechanism is in the hypothalamus: metabolic stress activates neuronal JNK3 signaling in AgRP circuits, altering excitatory transmission and thereby helping stabilize energy-balance responses to dietary challenge. That is a genuine homeostatic function, albeit in a specific physiological domain. ([elifesciences.org](https://elifesciences.org/articles/10031?utm_source=openai)) [S15]

## Bottom line

Yes—this claim is well supported. MAPK10/JNK3 is convincingly associated with **regulatory biological processes in neural systems**, including development, neurite regeneration, synaptic organization, receptor trafficking, and synaptic plasticity. Evidence for **homeostatic** roles also exists, most clearly in hypothalamic control of energy balance during metabolic stress, but is more context-specific than the broader regulatory evidence. ([pmc.ncbi.nlm.nih.gov](https://pmc.ncbi.nlm.nih.gov/articles/PMC7464089/)) [S16]

## Cited sources

- **[S1]** 
            Biological Properties of JNK3 and Its Function in Neurons, Astrocytes, Pancreatic β-Cells and Cardiovascular Cells - PMC
         — https://pmc.ncbi.nlm.nih.gov/articles/PMC7464089/
- **[S2]** Brain tissue expression of MAPK10 - Summary - The Human Protein Atlas — https://www.proteinatlas.org/ENSG00000109339-MAPK10/brain
- **[S3]** 
            Biological Properties of JNK3 and Its Function in Neurons, Astrocytes, Pancreatic β-Cells and Cardiovascular Cells - PMC
         — https://pmc.ncbi.nlm.nih.gov/articles/PMC7464089/
- **[S4]** Characterisation of de novo MAPK10/JNK3 truncation mutations associated with cognitive disorders in two unrelated patients - PubMed — https://pubmed.ncbi.nlm.nih.gov/23329067/
- **[S5]** 
            JNK Isoforms Differentially Regulate Neurite Growth and Regeneration in Dopaminergic Neurons In Vitro - PMC
         — https://pmc.ncbi.nlm.nih.gov/articles/PMC3178779/
- **[S6]** 
            JNK3 as Therapeutic Target and Biomarker in Neurodegenerative and Neurodevelopmental Brain Diseases - PMC
         — https://pmc.ncbi.nlm.nih.gov/articles/PMC7600688/
- **[S7]** Excitatory transmission onto AgRP neurons is regulated by cJun NH2-terminal kinase 3 in response to metabolic stress | eLife — https://elifesciences.org/articles/10031?utm_source=openai
- **[S8]** 
            Biological Properties of JNK3 and Its Function in Neurons, Astrocytes, Pancreatic β-Cells and Cardiovascular Cells - PMC
         — https://pmc.ncbi.nlm.nih.gov/articles/PMC7464089/
- **[S9]** 
            JNK3 as Therapeutic Target and Biomarker in Neurodegenerative and Neurodevelopmental Brain Diseases - PMC
         — https://pmc.ncbi.nlm.nih.gov/articles/PMC7600688/
- **[S10]** 
            JNK Isoforms Differentially Regulate Neurite Growth and Regeneration in Dopaminergic Neurons In Vitro - PMC
         — https://pmc.ncbi.nlm.nih.gov/articles/PMC3178779/
- **[S11]** 
            Biological Properties of JNK3 and Its Function in Neurons, Astrocytes, Pancreatic β-Cells and Cardiovascular Cells - PMC
         — https://pmc.ncbi.nlm.nih.gov/articles/PMC7464089/
- **[S12]** 
            JNK3 as Therapeutic Target and Biomarker in Neurodegenerative and Neurodevelopmental Brain Diseases - PMC
         — https://pmc.ncbi.nlm.nih.gov/articles/PMC7600688/
- **[S13]** 
            Biological Properties of JNK3 and Its Function in Neurons, Astrocytes, Pancreatic β-Cells and Cardiovascular Cells - PMC
         — https://pmc.ncbi.nlm.nih.gov/articles/PMC7464089/
- **[S14]** Pharos : Target Details - MAPK10 — https://pharos.nih.gov/targets/MAPK10?utm_source=openai
- **[S15]** Excitatory transmission onto AgRP neurons is regulated by cJun NH2-terminal kinase 3 in response to metabolic stress | eLife — https://elifesciences.org/articles/10031?utm_source=openai
- **[S16]** 
            Biological Properties of JNK3 and Its Function in Neurons, Astrocytes, Pancreatic β-Cells and Cardiovascular Cells - PMC
         — https://pmc.ncbi.nlm.nih.gov/articles/PMC7464089/
