## Supplementary material for "Understanding Substance Dependence: What Differentiates Addictive from Non-Addictive Drugs?": Folder for supplementary files: non-addictive-genes-claims-first100-073-mast3-gene-protein-has-been-associated-with-regula-63c70742522f.md

# non-addictive-genes-claims-first100-073-63c70742522f

**Original claim:** MAST3 gene/protein has been associated with regulatory or homeostatic biological processes in neural systems.

## Claim

MAST3 gene/protein has been associated with regulatory or homeostatic biological processes in neural systems.

## Verdict

Verdict: Likely True

The best available evidence supports **association with neural regulatory processes**, especially phosphorylation-dependent signaling in neurons. The evidence is **weaker for a broad, explicitly “homeostatic” role**; that part is mostly inferential rather than directly demonstrated. ([pubmed.ncbi.nlm.nih.gov](https://pubmed.ncbi.nlm.nih.gov/28167675/)) [S1]

## Confidence

Medium

Confidence is **medium** because there is solid mechanistic and human genetic evidence tying MAST3 to neuronal signaling regulation, but the literature is still relatively small, concentrated in a few pathways and disease cohorts, and does not yet show a broad, independently replicated role in neural homeostasis across systems. ([pubmed.ncbi.nlm.nih.gov](https://pubmed.ncbi.nlm.nih.gov/28167675/)) [S2]

## Supporting evidence

MAST3 is expressed in brain regions relevant to neuronal signaling. In a rodent expression study of the MAST kinase family, **MAST3 showed selective expression in the striatum and cerebral cortex**, supporting a neural role rather than a purely ubiquitous housekeeping function. A later human epilepsy study further reported that **MAST3 expression is largely restricted to excitatory cortical neurons late in prenatal development and postnatally**, strengthening the case that its major disease-relevant actions are neuronal. ([pubmed.ncbi.nlm.nih.gov](https://pubmed.ncbi.nlm.nih.gov/18206861/)) [S3]

The strongest direct evidence for a **regulatory neural function** comes from studies of the **ARPP-16/PP2A pathway in striatal neurons**. Andrade et al. showed that MAST3 phosphorylates **ARPP-16 at Ser46** in vitro and in vivo, and that this converts ARPP-16 into a **selective inhibitor of PP2A heterotrimers**. In striatum, Ser46 phosphorylation was high at baseline, consistent with **basal control of PP2A activity in striatal neurons**. Conditional loss of ARPP-16/19 altered phosphorylation of PP2A substrates including **DARPP-32, AKT, and ERK**, and also changed motivated behavior and the acute locomotor response to cocaine. That is direct evidence that MAST3 sits in a neuronal regulatory circuit affecting intracellular signaling and behavior. ([pubmed.ncbi.nlm.nih.gov](https://pubmed.ncbi.nlm.nih.gov/28167675/)) [S4]

A follow-up study showed that this is not just a linear pathway but a **switch-like regulatory module**. PKA phosphorylation of ARPP-16 at Ser88 suppresses MAST3-driven Ser46 phosphorylation, and PKA can also inhibit MAST3 itself; together these interactions create a **cAMP-regulated switch in PP2A inhibition**. In the authors’ discussion, they explicitly contrast meiosis with **non-dividing neurons in striatum**, where **MAST3 is basally active** and helps maintain constitutive PP2A inhibition until cAMP/PKA signaling shifts the system. This is a classic example of a neural regulatory mechanism controlling phosphorylation balance. ([pmc.ncbi.nlm.nih.gov](https://pmc.ncbi.nlm.nih.gov/articles/PMC5515580/)) [S5]

The broader relevance of this pathway to neuronal function is biologically plausible because PP2A is a major neuronal phosphatase. A review of cAMP-regulated phosphatases in brain notes that phosphorylation/dephosphorylation balance is critical for **neuronal development, synaptic transmission, plasticity, ionic conductance, brain function, and behavior**, and specifically discusses ARPP-16/PP2A as part of this network. That does not prove MAST3 is itself a master homeostatic protein, but it places the known MAST3 pathway inside well-established neural regulatory architecture. ([pmc.ncbi.nlm.nih.gov](https://pmc.ncbi.nlm.nih.gov/articles/PMC6433392/)) [S6]

Human genetics also supports a neural regulatory role. In a cohort of individuals with developmental and epileptic encephalopathy, **de novo missense variants in the MAST3 serine/threonine kinase domain** were associated with epilepsy, and tested variants generally produced **increased phosphorylation of the MAST3 substrate ARPP-16**, suggesting a **gain-of-function** effect on signaling. Because the same study localized MAST3 mainly to **cortical excitatory neurons**, the disease data fit a model in which abnormal MAST3-mediated regulation perturbs neuronal excitability/signaling. ([pubmed.ncbi.nlm.nih.gov](https://pubmed.ncbi.nlm.nih.gov/34185323/)) [S7]

Additional, but weaker, supportive evidence comes from neurodevelopmental studies. A 2022 genotype-phenotype paper reported that **MAST3 expression rises during fetal brain development and remains high postnatally in several brain regions**, is co-expressed with known developmental and epileptic encephalopathy genes, and that zebrafish mast3 crispants showed **central nervous system morphological abnormalities**. These findings support association with neural developmental regulation, though they are less definitive mechanistically than the striatal PP2A work. ([pmc.ncbi.nlm.nih.gov](https://pmc.ncbi.nlm.nih.gov/articles/PMC8790505/)) [S8]

## Contradicting or limiting evidence

The main limitation is that **direct evidence for “homeostatic” neural function is sparse**. The strongest papers show MAST3 regulating phosphatase signaling in specific neuronal contexts, but they do **not** directly demonstrate canonical homeostatic processes such as synaptic scaling, firing-rate homeostasis, ion homeostasis, or proteostatic homeostasis. So the claim is better supported if interpreted as **“regulatory”** rather than strongly **“homeostatic.”** ([pubmed.ncbi.nlm.nih.gov](https://pubmed.ncbi.nlm.nih.gov/28167675/)) [S9]

The mechanistic literature is also **narrow**. Most direct functional work centers on one pathway—**MAST3 → ARPP-16 → PP2A**—mainly in striatal neurons or heterologous cells. There is much less direct in vivo evidence for MAST3 across other neural cell types, circuits, or behaviors. Independent replication is still limited. ([pubmed.ncbi.nlm.nih.gov](https://pubmed.ncbi.nlm.nih.gov/28167675/)) [S10]

Some supportive studies are **association-heavy or exploratory**. For example, developmental-expression analyses, co-expression with other disease genes, and zebrafish morphology studies support relevance to the nervous system, but they do not by themselves prove that MAST3 directly executes a neural regulatory or homeostatic program. Likewise, disease association with epilepsy shows importance to neural function, but disease linkage alone does not specify mechanism. ([pubmed.ncbi.nlm.nih.gov](https://pubmed.ncbi.nlm.nih.gov/34185323/)) [S11]

Finally, MAST3 is not exclusively a neural protein: it has also been implicated in immune/inflammatory pathways and cancer-related signaling in non-neural systems. That does not argue against a neural role, but it does mean the current evidence does **not** establish MAST3 as a uniquely neural homeostasis factor. ([pmc.ncbi.nlm.nih.gov](https://pmc.ncbi.nlm.nih.gov/articles/PMC2574831/?utm_source=openai)) [S12]

## Mechanistic / biological rationale

MAST3 encodes a **serine/threonine kinase** with a PDZ-containing architecture typical of proteins that organize signaling complexes. In neurons, its best-defined function is to control the phosphorylation state of **ARPP-16**, which in turn regulates **PP2A**, a major phosphatase involved in neuronal signaling. Because neurons rely heavily on tightly tuned phosphorylation/dephosphorylation for receptor signaling, synaptic plasticity, and excitability, a kinase that sets PP2A tone is inherently part of a **regulatory control system**. ([pubmed.ncbi.nlm.nih.gov](https://pubmed.ncbi.nlm.nih.gov/28167675/)) [S13]

That same logic explains the disease genetics. If pathogenic MAST3 variants increase ARPP-16 phosphorylation in **cortical excitatory neurons**, they could shift PP2A-dependent signaling away from its normal set point, altering phosphorylation of downstream substrates and biasing networks toward hyperexcitability or abnormal development. This is a biologically coherent mechanism for epilepsy and neurodevelopmental phenotypes, even if the field has not yet fully mapped the downstream cascade in human brain. ([pubmed.ncbi.nlm.nih.gov](https://pubmed.ncbi.nlm.nih.gov/34185323/)) [S14]

## Bottom line

The claim is **likely true**, with an important qualifier: **MAST3 is clearly associated with regulatory signaling processes in neural systems**, especially phospho-signaling control in neurons, but the evidence for a broad, explicitly **homeostatic** role is still limited and indirect. If you mean **“MAST3 participates in neural regulatory pathways that help set signaling balance,”** the literature supports that. If you mean **“MAST3 is an established mediator of neural homeostasis writ large,”** the evidence is not yet strong enough for that stronger wording. ([pubmed.ncbi.nlm.nih.gov](https://pubmed.ncbi.nlm.nih.gov/28167675/)) [S15]

## Cited sources

- **[S1]** ARPP-16 Is a Striatal-Enriched Inhibitor of Protein Phosphatase 2A Regulated by Microtubule-Associated Serine/Threonine Kinase 3 (Mast 3 Kinase) - PubMed — https://pubmed.ncbi.nlm.nih.gov/28167675/
- **[S2]** ARPP-16 Is a Striatal-Enriched Inhibitor of Protein Phosphatase 2A Regulated by Microtubule-Associated Serine/Threonine Kinase 3 (Mast 3 Kinase) - PubMed — https://pubmed.ncbi.nlm.nih.gov/28167675/
- **[S3]** Expression of the MAST family of serine/threonine kinases - PubMed — https://pubmed.ncbi.nlm.nih.gov/18206861/
- **[S4]** ARPP-16 Is a Striatal-Enriched Inhibitor of Protein Phosphatase 2A Regulated by Microtubule-Associated Serine/Threonine Kinase 3 (Mast 3 Kinase) - PubMed — https://pubmed.ncbi.nlm.nih.gov/28167675/
- **[S5]** 
            Reciprocal regulation of ARPP-16 by PKA and MAST3 kinases provides a cAMP-regulated switch in protein phosphatase 2A inhibition - PMC
         — https://pmc.ncbi.nlm.nih.gov/articles/PMC5515580/
- **[S6]** 
            cAMP regulation of protein phosphatases PP1 and PP2A in brain - PMC
         — https://pmc.ncbi.nlm.nih.gov/articles/PMC6433392/
- **[S7]** Pathogenic MAST3 Variants in the STK Domain Are Associated with Epilepsy - PubMed — https://pubmed.ncbi.nlm.nih.gov/34185323/
- **[S8]** 
            The Role of Microtubule Associated Serine/Threonine Kinase 3 Variants in Neurodevelopmental Diseases: Genotype-Phenotype Association - PMC
         — https://pmc.ncbi.nlm.nih.gov/articles/PMC8790505/
- **[S9]** ARPP-16 Is a Striatal-Enriched Inhibitor of Protein Phosphatase 2A Regulated by Microtubule-Associated Serine/Threonine Kinase 3 (Mast 3 Kinase) - PubMed — https://pubmed.ncbi.nlm.nih.gov/28167675/
- **[S10]** ARPP-16 Is a Striatal-Enriched Inhibitor of Protein Phosphatase 2A Regulated by Microtubule-Associated Serine/Threonine Kinase 3 (Mast 3 Kinase) - PubMed — https://pubmed.ncbi.nlm.nih.gov/28167675/
- **[S11]** Pathogenic MAST3 Variants in the STK Domain Are Associated with Epilepsy - PubMed — https://pubmed.ncbi.nlm.nih.gov/34185323/
- **[S12]** MAST3: a Novel IBD Risk Factor that Modulates TLR4 Signaling - PMC — https://pmc.ncbi.nlm.nih.gov/articles/PMC2574831/?utm_source=openai
- **[S13]** ARPP-16 Is a Striatal-Enriched Inhibitor of Protein Phosphatase 2A Regulated by Microtubule-Associated Serine/Threonine Kinase 3 (Mast 3 Kinase) - PubMed — https://pubmed.ncbi.nlm.nih.gov/28167675/
- **[S14]** Pathogenic MAST3 Variants in the STK Domain Are Associated with Epilepsy - PubMed — https://pubmed.ncbi.nlm.nih.gov/34185323/
- **[S15]** ARPP-16 Is a Striatal-Enriched Inhibitor of Protein Phosphatase 2A Regulated by Microtubule-Associated Serine/Threonine Kinase 3 (Mast 3 Kinase) - PubMed — https://pubmed.ncbi.nlm.nih.gov/28167675/
