## Supplementary material for "Understanding Substance Dependence: What Differentiates Addictive from Non-Addictive Drugs?": Folder for supplementary files: non-addictive-genes-claims-first100-074-ralgds-gene-protein-has-been-associated-with-regul-f412d412170c.md

# non-addictive-genes-claims-first100-074-f412d412170c

**Original claim:** RALGDS gene/protein has been associated with regulatory or homeostatic biological processes in neural systems.

## Claim

**RALGDS gene/protein has been associated with regulatory or homeostatic biological processes in neural systems.**

## Verdict

Verdict: Likely True

This claim is **best supported if “regulatory” is interpreted broadly**—for example, regulation of receptor trafficking and synaptic signaling in neurons. I found **direct experimental evidence** linking RalGDS to regulation of neuronal group I metabotropic glutamate receptor (mGluR1/5) endocytosis, plus expression/annotation evidence placing RALGDS in nervous tissue. However, I found **much weaker direct evidence for a specifically “homeostatic” role** in the canonical neuroscience sense (such as synaptic scaling, firing-rate homeostasis, or network set-point stabilization). ([pubmed.ncbi.nlm.nih.gov](https://pubmed.ncbi.nlm.nih.gov/15470141/)) [S1]

## Confidence

**Medium.**

The main reason confidence is not high is that the neural evidence is **real but relatively sparse and narrow**, and much of the broader functional literature on RalGDS is outside the nervous system. ([pubmed.ncbi.nlm.nih.gov](https://pubmed.ncbi.nlm.nih.gov/15470141/)) [S2]

## Supporting evidence

RALGDS is a **guanine nucleotide exchange factor (GEF)** for the small GTPases **RalA and RalB**, and the original cloning paper reported that its mRNA and protein were found broadly across tissues examined. More recent mouse/human gene resources also place the gene in nervous tissue: NCBI’s mouse gene summary states that **Ralgds is expressed in the nervous system and retina**, while Human Protein Atlas brain pages show measurable expression across multiple brain regions, including cortex, hippocampal formation, hypothalamus, basal ganglia, spinal cord, white matter, and others; in hypothalamus, HPA/GTEx values are in the tens to ~100 nTPM range depending on dataset/subregion. This does not prove neural function by itself, but it does establish that RALGDS is present in neural tissues where such functions are possible. ([pubmed.ncbi.nlm.nih.gov](https://pubmed.ncbi.nlm.nih.gov/8094051/)) [S3]

The strongest direct neural evidence comes from **Bhattacharya et al. (J Neurosci, 2004)**. In that study, the authors showed that **Ral, RalGDS, and PLD2 are constitutively associated with group I mGluRs (mGluR1a and mGluR5a)** and that this complex regulates **constitutive mGluR endocytosis**. Importantly, the work was not limited to heterologous cells: the paper reports receptor internalization in **neuronal cultures**, colocalization in **axons and dendrites**, and co-immunoprecipitation of endogenous components from **whole-brain lysates**. Because group I mGluRs are core regulators of excitatory synaptic signaling, this is a bona fide association of RalGDS with a **regulatory biological process in neural systems**. ([pubmed.ncbi.nlm.nih.gov](https://pubmed.ncbi.nlm.nih.gov/15470141/)) [S4]

Model-organism curation is directionally consistent with that conclusion. MGI lists mouse **Ralgds** as expressed in the **nervous system** and places it in broad Gene Ontology categories that include **signaling**, **establishment of localization**, **homeostatic process**, and the cellular component **synapse**. Those annotations should be interpreted cautiously—some are predicted or aggregated rather than direct neural functional experiments—but they are consistent with RALGDS being part of signaling/trafficking machinery relevant to neural regulation. ([informatics.jax.org](https://www.informatics.jax.org/marker/MGI%3A107485)) [S5]

There is also useful **indirect neural support** from the downstream pathway. In mouse synaptosomes, manipulating **Ral** signaling altered **PKC-enhanced glutamate release**, suppressed refilling of the **readily releasable pool** of synaptic vesicles, and implicated Ral interaction with the **Sec6/8 exocyst complex** in modulation of synaptic strength. Since RalGDS is a principal upstream GEF for Ral, these data make biological sense of how RalGDS could influence neural regulatory processes, even though that particular study did **not** manipulate RalGDS directly. ([pubmed.ncbi.nlm.nih.gov](https://pubmed.ncbi.nlm.nih.gov/11865051/)) [S6]

## Contradicting or limiting evidence

The key limitation is that **direct neural evidence for RalGDS itself is limited**. The 2004 mGluR paper is strong evidence for a neuronal regulatory association, but the experimental emphasis is on the **Ral/RalGDS/PLD2 complex** and on receptor trafficking; it does **not** establish a broad role for RalGDS across neural homeostasis, neural development, or behavior. In other words, the literature supports **a specific neural regulatory role**, not a deeply validated general neural-homeostatic role. ([pubmed.ncbi.nlm.nih.gov](https://pubmed.ncbi.nlm.nih.gov/15470141/)) [S7]

A second limitation is that much of the functional literature on RalGDS is **outside neuroscience**. Examples include **endothelial exocytosis**, where RalGDS knockdown inhibited regulated Weibel-Palade body release; **skin carcinogenesis**, where RalGDS knockout mice were viable and developmentally normal but resistant to tumor formation; and **cardiac autophagy/hypertrophy**, where RalGDS mediated stress-induced autophagic remodeling. These studies reinforce that RalGDS is a broadly acting trafficking/signaling regulator, but they also show that the best-established functions are not neural-specific. ([pubmed.ncbi.nlm.nih.gov](https://pubmed.ncbi.nlm.nih.gov/18417737/?utm_source=openai)) [S8]

I also did **not find strong direct evidence** in the retrieved literature for RALGDS in canonical neural homeostasis paradigms such as **synaptic scaling, homeostatic intrinsic plasticity, excitation/inhibition set-point control, or circuit-level activity stabilization**. So if the claim is interpreted narrowly as “RALGDS is a known neural homeostasis gene,” the evidence would be too thin for a stronger verdict. This is why the claim lands at **Likely True** rather than **Highly Supported**. ([pubmed.ncbi.nlm.nih.gov](https://pubmed.ncbi.nlm.nih.gov/15470141/)) [S9]

## Mechanistic / biological rationale

Mechanistically, the claim is plausible. RALGDS is a **RalA/RalB GEF** and Ras/Rap effector involved in **intracellular transport and membrane trafficking**. In neurons, regulatory control of receptor endocytosis, vesicle docking, and exocyst-dependent trafficking is central to how synapses adjust signaling strength and maintain stable communication. A molecule that couples upstream Ras-family signaling to **Ral/exocyst trafficking modules** is therefore well positioned to influence neural regulatory processes. ([proteinatlas.org](https://www.proteinatlas.org/ENSG00000160271-RALGDS)) [S10]

The best-supported neural mechanism is **group I mGluR trafficking**: Bhattacharya et al. propose that an **mGluR-scaffolded RalGDS/Ral/PLD2 complex** provides an alternative pathway for constitutive receptor internalization. Since receptor trafficking is a major determinant of synaptic responsiveness, that is a credible route by which RalGDS could contribute to neural regulation. Separately, non-neural studies show RalGDS can link **Ca²⁺/cAMP/calmodulin** inputs to exocytic or autophagic trafficking programs, which strengthens the biological plausibility of analogous regulatory roles in neurons, even if those roles remain incompletely tested. ([pubmed.ncbi.nlm.nih.gov](https://pubmed.ncbi.nlm.nih.gov/15470141/)) [S11]

## Bottom line

**Yes, with caveats:** RALGDS has been associated with **regulatory processes in neural systems**, most directly through **neuronal mGluR trafficking/endocytosis** and supporting evidence of brain expression. But the evidence for a **specific homeostatic role** in neural systems is **limited and indirect**. So the claim is **Likely True** if read broadly as “RALGDS participates in neural regulatory biology,” but it is **not yet strongly established** as a dedicated neural homeostasis factor. ([pubmed.ncbi.nlm.nih.gov](https://pubmed.ncbi.nlm.nih.gov/15470141/)) [S12]

## Cited sources

- **[S1]** Ral and phospholipase D2-dependent pathway for constitutive metabotropic glutamate receptor endocytosis - PubMed — https://pubmed.ncbi.nlm.nih.gov/15470141/
- **[S2]** Ral and phospholipase D2-dependent pathway for constitutive metabotropic glutamate receptor endocytosis - PubMed — https://pubmed.ncbi.nlm.nih.gov/15470141/
- **[S3]** Characterization of a guanine nucleotide dissociation stimulator for a ras-related GTPase - PubMed — https://pubmed.ncbi.nlm.nih.gov/8094051/
- **[S4]** Ral and phospholipase D2-dependent pathway for constitutive metabotropic glutamate receptor endocytosis - PubMed — https://pubmed.ncbi.nlm.nih.gov/15470141/
- **[S5]** Ralgds MGI Mouse Gene Detail - MGI:107485 - ral guanine nucleotide dissociation stimulator — https://www.informatics.jax.org/marker/MGI%3A107485
- **[S6]** Ral-GTPase influences the regulation of the readily releasable pool of synaptic vesicles - PubMed — https://pubmed.ncbi.nlm.nih.gov/11865051/
- **[S7]** Ral and phospholipase D2-dependent pathway for constitutive metabotropic glutamate receptor endocytosis - PubMed — https://pubmed.ncbi.nlm.nih.gov/15470141/
- **[S8]** Guanine exchange factor RalGDS mediates exocytosis of Weibel-Palade bodies from endothelial cells - PubMed — https://pubmed.ncbi.nlm.nih.gov/18417737/?utm_source=openai
- **[S9]** Ral and phospholipase D2-dependent pathway for constitutive metabotropic glutamate receptor endocytosis - PubMed — https://pubmed.ncbi.nlm.nih.gov/15470141/
- **[S10]** RALGDS protein expression summary - The Human Protein Atlas — https://www.proteinatlas.org/ENSG00000160271-RALGDS
- **[S11]** Ral and phospholipase D2-dependent pathway for constitutive metabotropic glutamate receptor endocytosis - PubMed — https://pubmed.ncbi.nlm.nih.gov/15470141/
- **[S12]** Ral and phospholipase D2-dependent pathway for constitutive metabotropic glutamate receptor endocytosis - PubMed — https://pubmed.ncbi.nlm.nih.gov/15470141/
