## Supplementary material for "Understanding Substance Dependence: What Differentiates Addictive from Non-Addictive Drugs?": Folder for supplementary files: non-addictive-genes-claims-first100-075-agap3-gene-protein-has-been-associated-with-regula-d0ab524e5811.md

# non-addictive-genes-claims-first100-075-d0ab524e5811

**Original claim:** AGAP3 gene/protein has been associated with regulatory or homeostatic biological processes in neural systems.

## Claim

AGAP3 gene/protein has been associated with regulatory or homeostatic biological processes in neural systems.

## Verdict

Verdict: Likely True

## Confidence

High

## Supporting evidence

AGAP3 is directly tied to **neural regulatory processes** in authoritative gene annotations and experimental neuroscience literature. The NCBI Gene RefSeq summary describes AGAP3 as “an essential component” of the NMDA receptor signaling complex that mediates long-term potentiation (LTP) by linking NMDA receptor activation to AMPA receptor trafficking, and notes substantial brain expression. ([ncbi.nlm.nih.gov](https://www.ncbi.nlm.nih.gov/gene/116988)) [S1]

The strongest direct evidence comes from a 2013 *Journal of Neuroscience* study in rat neurons. That study found AGAP3/CRAG enriched in postsynaptic density fractions, colocalized mainly with PSD95 rather than the presynaptic marker vGlut, and therefore localized to postsynaptic sites in the adult brain. Functionally, AGAP3 knockdown increased basal Ras and Arf6 activity, indicating that AGAP3 normally regulates these signaling pathways. The same study further showed that AGAP3 controls basal and activity-dependent GluA1/AMPA receptor trafficking and is required for normal chemically induced LTP-related signaling. ([pmc.ncbi.nlm.nih.gov](https://pmc.ncbi.nlm.nih.gov/articles/PMC3728681/)) [S2]

That same experimental paper supports a specifically **regulatory** role rather than mere correlation. AGAP3 acted as an Arf6 GAP, negatively regulating Arf6 and Ras activity under basal conditions and influencing surface GluA1 abundance. The authors concluded that AGAP3 links NMDA receptor activation to AMPA receptor trafficking, separating effects on basal receptor trafficking from activity-dependent trafficking through distinct signaling domains. This is a classic synaptic regulatory function in excitatory neural systems. ([pmc.ncbi.nlm.nih.gov](https://pmc.ncbi.nlm.nih.gov/articles/PMC3728681/)) [S3]

Independent authoritative annotation is consistent with those experiments. On the NCBI Gene page, GO/GOA annotations place AGAP3 in the **glutamatergic synapse** and **postsynaptic density** and annotate it as involved in **regulation of postsynaptic membrane neurotransmitter receptor levels**. While these GO lines are not all based on direct assay, they align well with the experimental receptor-trafficking data. ([ncbi.nlm.nih.gov](https://www.ncbi.nlm.nih.gov/gene/116988)) [S4]

Evidence also extends beyond acute synaptic signaling. A 2021 mouse study on the short AGAP3 splice variant **CRAG/Centaurin-γ3** reported that CRAG is highly expressed in the dentate gyrus, promotes neuronal cell survival and antioxidant defense during neuronal development, and that forebrain-specific deficiency caused immature dentate gyri with excess immature neurons, fewer mature neurons, and hyperactivity. Those findings support a role in regulating neuronal maturation and stress-responsive processes in a neural circuit context. ([pubmed.ncbi.nlm.nih.gov](https://pubmed.ncbi.nlm.nih.gov/33811862/)) [S5]

There is also suggestive evidence for activity-dependent regulation in memory-related neural plasticity. A 2023 cross-species study found increased Agap3 transcription in dorsal hippocampus after novelty-associated memory paradigms and proposed AGAP3 as a candidate plasticity-related protein for initial memory consolidation, though the authors explicitly said further work is needed to confirm function. ([molecularbrain.biomedcentral.com](https://molecularbrain.biomedcentral.com/articles/10.1186/s13041-023-01056-4)) [S6]

## Contradicting or limiting evidence

The main limitation is that evidence for **homeostatic** function in the narrow neuroscience sense is much weaker than evidence for general **regulatory** function. The best-supported AGAP3 literature is about NMDA/AMPA receptor trafficking, Ras/ERK-Arf6 signaling, synaptic plasticity, neuronal stress responses, and neuronal maturation. I did not find strong direct evidence that AGAP3 is a canonical mediator of **homeostatic synaptic scaling** or broader neural homeostasis comparable to the better-established homeostatic-plasticity literature for other postsynaptic signaling proteins. This is an inference from the scope of the cited AGAP3 literature. ([pmc.ncbi.nlm.nih.gov](https://pmc.ncbi.nlm.nih.gov/articles/PMC3728681/)) [S7]

A second limitation is that much of the strongest mechanistic evidence comes from **rodent primary neurons or mouse models**, not human interventional studies. The NCBI summary and GO annotations are useful, but some GO entries are marked IEA/IBA/ISO rather than direct experimental assay, so they should be treated as supportive rather than definitive on their own. ([ncbi.nlm.nih.gov](https://www.ncbi.nlm.nih.gov/gene/116988)) [S8]

A third limitation is isoform specificity. Some relevant neural evidence concerns **CRAG**, a short splice variant of AGAP3, especially for antioxidant defense, proteostasis, and dentate gyrus maturation. Those findings strengthen the broader claim that the AGAP3 locus/protein products participate in neural regulatory biology, but they do not prove that full-length AGAP3 performs every CRAG function. ([pubmed.ncbi.nlm.nih.gov](https://pubmed.ncbi.nlm.nih.gov/33811862/)) [S9]

Finally, the memory-consolidation study is associative and hypothesis-generating. It shows Agap3 transcription rises with novelty-associated learning conditions, but the authors themselves state that the exact contribution of AGAP3 to initial memory consolidation remains to be established. ([molecularbrain.biomedcentral.com](https://molecularbrain.biomedcentral.com/articles/10.1186/s13041-023-01056-4)) [S10]

## Mechanistic / biological rationale

AGAP3’s domain structure fits a regulatory synaptic role. NCBI and NCBI’s conserved-domain resource describe AGAP3 as containing a GTPase-like domain, pleckstrin homology domain, ArfGAP domain, and ankyrin repeats, and as part of the NMDA receptor complex that regulates Arf6 and Ras/ERK signaling while linking NMDA receptor activation to AMPA receptor trafficking. That architecture is exactly what one would expect for a signaling/scaffolding regulator of postsynaptic receptor dynamics. ([ncbi.nlm.nih.gov](https://www.ncbi.nlm.nih.gov/gene/116988)) [S11]

Biologically, this makes sense because tuning synaptic AMPA receptor insertion/removal is a core mechanism for adjusting excitatory synaptic strength. In neurons, AGAP3 appears to restrain basal Ras and Arf6 activity, regulate steady-state AMPA receptor trafficking, and participate in activity-triggered receptor changes during LTP. Those are regulatory processes that help shape excitatory synapse function and stability, even if they are not yet firmly established as “homeostatic scaling” per se. ([pmc.ncbi.nlm.nih.gov](https://pmc.ncbi.nlm.nih.gov/articles/PMC3728681/)) [S12]

For the CRAG splice variant, the proposed mechanism is more stress-response/homeostasis-like: nuclear localization, activation of SRF/c-Fos/AP-1 pathways, facilitation of proteasomal handling of misfolded/polyglutamine proteins, and antioxidant defense. In neuronal cells and mouse models, that provides a plausible route by which the AGAP3 locus could contribute to neuronal proteostasis and developmental resilience. ([pubmed.ncbi.nlm.nih.gov](https://pubmed.ncbi.nlm.nih.gov/21832068/)) [S13]

## Bottom line

The claim is **likely true**. AGAP3 is well associated with **regulatory biological processes in neural systems**, especially postsynaptic signaling, Ras/ERK and Arf6 regulation, AMPA receptor trafficking, and synaptic plasticity. There is also supportive evidence from the CRAG splice variant for neuronal stress-response and maturation roles. However, the evidence is much stronger for **synaptic regulation/plasticity** than for a specific, established role in **neural homeostasis** or canonical homeostatic plasticity. ([ncbi.nlm.nih.gov](https://www.ncbi.nlm.nih.gov/gene/116988)) [S14]

## Cited sources

- **[S1]** AGAP3 ArfGAP with GTPase domain, ankyrin repeat and PH domain 3 [Homo sapiens (human)] - Gene - NCBI — https://www.ncbi.nlm.nih.gov/gene/116988
- **[S2]** 
            AGAP3 and Arf6 Regulate Trafficking of AMPA Receptors and Synaptic Plasticity - PMC
         — https://pmc.ncbi.nlm.nih.gov/articles/PMC3728681/
- **[S3]** 
            AGAP3 and Arf6 Regulate Trafficking of AMPA Receptors and Synaptic Plasticity - PMC
         — https://pmc.ncbi.nlm.nih.gov/articles/PMC3728681/
- **[S4]** AGAP3 ArfGAP with GTPase domain, ankyrin repeat and PH domain 3 [Homo sapiens (human)] - Gene - NCBI — https://www.ncbi.nlm.nih.gov/gene/116988
- **[S5]** Forebrain-specific deficiency of the GTPase CRAG/Centaurin-γ3 leads to immature dentate gyri and hyperactivity in mice - PubMed — https://pubmed.ncbi.nlm.nih.gov/33811862/
- **[S6]** Novelty-induced memory consolidation is accompanied by increased Agap3 transcription: a cross-species study | Molecular Brain | Springer Nature Link — https://molecularbrain.biomedcentral.com/articles/10.1186/s13041-023-01056-4
- **[S7]** 
            AGAP3 and Arf6 Regulate Trafficking of AMPA Receptors and Synaptic Plasticity - PMC
         — https://pmc.ncbi.nlm.nih.gov/articles/PMC3728681/
- **[S8]** AGAP3 ArfGAP with GTPase domain, ankyrin repeat and PH domain 3 [Homo sapiens (human)] - Gene - NCBI — https://www.ncbi.nlm.nih.gov/gene/116988
- **[S9]** Forebrain-specific deficiency of the GTPase CRAG/Centaurin-γ3 leads to immature dentate gyri and hyperactivity in mice - PubMed — https://pubmed.ncbi.nlm.nih.gov/33811862/
- **[S10]** Novelty-induced memory consolidation is accompanied by increased Agap3 transcription: a cross-species study | Molecular Brain | Springer Nature Link — https://molecularbrain.biomedcentral.com/articles/10.1186/s13041-023-01056-4
- **[S11]** AGAP3 ArfGAP with GTPase domain, ankyrin repeat and PH domain 3 [Homo sapiens (human)] - Gene - NCBI — https://www.ncbi.nlm.nih.gov/gene/116988
- **[S12]** 
            AGAP3 and Arf6 Regulate Trafficking of AMPA Receptors and Synaptic Plasticity - PMC
         — https://pmc.ncbi.nlm.nih.gov/articles/PMC3728681/
- **[S13]** CRMP5-associated GTPase (CRAG) protein protects neuronal cells against cytotoxicity of expanded polyglutamine protein partially via c-Fos-dependent activator protein-1 activation - PubMed — https://pubmed.ncbi.nlm.nih.gov/21832068/
- **[S14]** AGAP3 ArfGAP with GTPase domain, ankyrin repeat and PH domain 3 [Homo sapiens (human)] - Gene - NCBI — https://www.ncbi.nlm.nih.gov/gene/116988
