## Supplementary material for "Understanding Substance Dependence: What Differentiates Addictive from Non-Addictive Drugs?": Folder for supplementary files: non-addictive-genes-claims-first100-076-rab6a-gene-protein-has-been-associated-with-regula-2c32315fb028.md

# non-addictive-genes-claims-first100-076-2c32315fb028

**Original claim:** RAB6A gene/protein has been associated with regulatory or homeostatic biological processes in neural systems.

## Claim

RAB6A gene/protein has been associated with regulatory or homeostatic biological processes in neural systems.

## Verdict

Verdict: Likely True

## Confidence

Medium

## Supporting evidence

RAB6A is a Rab-family small GTPase localized to the Golgi apparatus that regulates both retrograde and anterograde membrane trafficking, and it is expressed in brain tissue; NCBI lists human **RAB6A** as ubiquitously expressed with substantial brain expression, and mouse **Rab6a** as expressed in the nervous system and adult cortex. ([ncbi.nlm.nih.gov](https://www.ncbi.nlm.nih.gov/gene/5870)) [S1]

Direct neural-system evidence comes from developmental mouse studies. In the embryonic cortex, **RAB6A/B double knockout** disrupted apical localization of the polarity determinant **Crumbs3**, caused retraction of apical radial glial processes, increased progenitor delamination/ectopic division, and the authors concluded that the RAB6-dynein-LIS1 pathway is required for maintenance of **neuroepithelial integrity**. That is a clear association with regulatory processes governing neural stem-cell polarity and tissue organization. ([pubmed.ncbi.nlm.nih.gov](https://pubmed.ncbi.nlm.nih.gov/35979738/)) [S2]

A second mouse CNS study strengthened this point. In **CNS-specific Rab6a/b double-knockout mice**, loss of Rab6 caused severe neocortical and cerebellar dysplasia; in cultured neurons it led to abnormal accumulation of **synaptic vesicle precursors** near the Golgi, impaired axonal extension, loss of **axon-dendrite polarity**, and expansion of somatic lysosomes. These findings connect Rab6-family signaling to neuronal polarity and intracellular trafficking systems that are central to stable neuronal organization. ([pubmed.ncbi.nlm.nih.gov](https://pubmed.ncbi.nlm.nih.gov/38830762/)) [S3]

There is also evidence tying **RAB6A specifically** to glial elements of neural systems. In mouse and human brain, Rab6A immunostaining was reported as a **pan-astrocytic marker**, strongly present in peripheral astrocyte processes and excluded from microglia, oligodendrocytes, and NG2 cells. The study was largely descriptive, but it supports a brain-specific association of RAB6A with astrocytic trafficking architecture relevant to glia-neuron signaling. ([pmc.ncbi.nlm.nih.gov](https://pmc.ncbi.nlm.nih.gov/articles/PMC7824777/)) [S4]

More recent neuron-focused trafficking work links **GFP-RAB6A** to transport of biosynthetic **LAMP1-positive** carriers in axons. That study showed RAB6A colocalization with carriers that enter the axon and replenish axonal lysosomes. Because lysosomal trafficking is a core component of cellular maintenance, this provides additional evidence that RAB6A is associated with homeostasis-relevant trafficking in neurons, though this specific paper was more mechanistic/observational than a clean RAB6A-loss-of-function test. ([pubmed.ncbi.nlm.nih.gov](https://pubmed.ncbi.nlm.nih.gov/40016183/)) [S5]

Evolutionary data support the broader point. In **C. elegans**, Rab6 homologs **RAB-6.1** and **RAB-6.2** regulate retrograde recycling of the AMPA-receptor subunit **GLR-1** in neurons; mutants fail to recycle GLR-1 properly and show behavioral defects. While not human/mouse RAB6A per se, these studies reinforce a conserved Rab6-family role in neural cargo regulation. ([pubmed.ncbi.nlm.nih.gov](https://pubmed.ncbi.nlm.nih.gov/26891225/)) [S6]

## Contradicting or limiting evidence

The main limitation is **isoform specificity**. Much of the strongest mammalian neural evidence comes from **Rab6a/Rab6b double knockout** rather than selective RAB6A loss alone, so the data show a robust role for the **Rab6 family in neural systems**, but only partially isolate how much of that effect is uniquely attributable to RAB6A. ([pubmed.ncbi.nlm.nih.gov](https://pubmed.ncbi.nlm.nih.gov/35979738/)) [S7]

Relatedly, neuronal isoform expression complicates interpretation. A presynaptic study notes that mammals have a ubiquitous **Rab6A** and a more **brain-prominent Rab6B**; that same work found Rab6B to be the dominant Rab6 isoform in brain. Conversely, the astrocyte study found Rab6A strongly in astrocytes and reported that **RAB6B is hardly expressed in astrocytes if at all**. So, in brain tissue, “Rab6 in neural systems” often reflects a division of labor between **RAB6A in astroglia** and **RAB6B in neurons**, rather than a uniformly defined RAB6A neuronal function. ([pmc.ncbi.nlm.nih.gov](https://pmc.ncbi.nlm.nih.gov/articles/PMC7360120/)) [S8]

Also, the evidence for **homeostasis** is less direct than the evidence for **regulation/polarity/trafficking**. I found strong support that Rab6 pathways regulate neural polarity, vesicle trafficking, receptor recycling, and neuroepithelial integrity, but much less direct evidence that **RAB6A** is an established mediator of canonical neural homeostatic programs such as firing-rate homeostasis, synaptic scaling, or calcium homeostasis. The available literature located here is centered more on trafficking and structural regulation than on classical homeostatic plasticity. ([pubmed.ncbi.nlm.nih.gov](https://pubmed.ncbi.nlm.nih.gov/35979738/)) [S9]

Finally, disease-linked observations are mostly associative. In Alzheimer disease brains, Rab6 was increased in hippocampus, entorhinal cortex, and temporal cortex and correlated with the ER-stress marker **BiP**; this suggests involvement in protein-quality-control responses, but it does not prove that altered Rab6 is causally homeostatic or protective. ([pubmed.ncbi.nlm.nih.gov](https://pubmed.ncbi.nlm.nih.gov/17573808/)) [S10]

## Mechanistic / biological rationale

Mechanistically, the claim is biologically plausible. RAB6A is a Golgi-associated Rab GTPase that controls targeting/fusion of transport carriers and regulates traffic from endosomes/Golgi back to the ER as well as export toward the plasma membrane. Independent cell-biology work further links Rab6 to Golgi trafficking pathways required for **Golgi homeostasis**. In neural cells, where polarized long-distance trafficking is unusually demanding, defects in this machinery would be expected to disrupt polarity, receptor delivery, lysosome distribution, and secretory balance. ([ncbi.nlm.nih.gov](https://www.ncbi.nlm.nih.gov/gene/5870)) [S11]

That expectation matches the neural phenotypes observed experimentally: impaired apical transport of polarity components in radial glia, defective synaptic-vesicle-precursor trafficking, loss of axon-dendrite polarity, and altered lysosomal distribution in neurons. These are not just generic transport defects; in neural tissue they bear directly on regulatory organization and maintenance of cell state. ([pubmed.ncbi.nlm.nih.gov](https://pubmed.ncbi.nlm.nih.gov/35979738/)) [S12]

There is also a plausible **proteostasis/homeostasis** angle. The Alzheimer study explicitly frames Rab6 as part of retrograde Golgi-ER trafficking that may serve as a **post-ER quality-control system**, and it found Rab6 upregulation correlated with ER stress in affected brain regions. Separately, systems-level work on Alzheimer-associated protein networks identified **RAB6A** as a central node potentially helping direct metastable proteins toward endosomal-lysosomal degradation. This does not prove a normal physiologic homeostatic role in healthy neurons, but it supports the idea that RAB6A sits in pathways relevant to neural proteostasis. ([pubmed.ncbi.nlm.nih.gov](https://pubmed.ncbi.nlm.nih.gov/17573808/)) [S13]

## Bottom line

The claim is **likely true**, especially if “regulatory or homeostatic” is interpreted broadly. There is good evidence that **RAB6A/Rab6-family pathways** participate in **regulatory neural processes** such as neural progenitor polarity, neuroepithelial integrity, axon-dendrite polarity, and trafficking of synaptic or lysosomal cargo. Evidence for a more specific role in **neural homeostasis** exists but is more indirect, often tied to lysosomal/proteostatic trafficking or disease-associated quality-control responses, and is frequently not cleanly separable from **Rab6B** or family-wide redundancy. ([pubmed.ncbi.nlm.nih.gov](https://pubmed.ncbi.nlm.nih.gov/35979738/)) [S14]

## Cited sources

- **[S1]** RAB6A RAB6A, member RAS oncogene family [Homo sapiens (human)] - Gene - NCBI — https://www.ncbi.nlm.nih.gov/gene/5870
- **[S2]** RAB6 and dynein drive post-Golgi apical transport to prevent neuronal progenitor delamination - PubMed — https://pubmed.ncbi.nlm.nih.gov/35979738/
- **[S3]** Rab6-Mediated Polarized Transport of Synaptic Vesicle Precursors Is Essential for the Establishment of Neuronal Polarity and Brain Formation - PubMed — https://pubmed.ncbi.nlm.nih.gov/38830762/
- **[S4]** 
            Rab6A as a Pan-Astrocytic Marker in Mouse and Human Brain, and Comparison with Other Glial Markers (GFAP, GS, Aldh1L1, SOX9) - PMC
         — https://pmc.ncbi.nlm.nih.gov/articles/PMC7824777/
- **[S5]** Spatiotemporal proteomics reveals the biosynthetic lysosomal membrane protein interactome in neurons - PubMed — https://pubmed.ncbi.nlm.nih.gov/40016183/
- **[S6]** RAB-6.1 and RAB-6.2 Promote Retrograde Transport in C. elegans - PubMed — https://pubmed.ncbi.nlm.nih.gov/26891225/
- **[S7]** RAB6 and dynein drive post-Golgi apical transport to prevent neuronal progenitor delamination - PubMed — https://pubmed.ncbi.nlm.nih.gov/35979738/
- **[S8]** 
            ELKS1 Captures Rab6-Marked Vesicular Cargo in Presynaptic Nerve Terminals - PMC
         — https://pmc.ncbi.nlm.nih.gov/articles/PMC7360120/
- **[S9]** RAB6 and dynein drive post-Golgi apical transport to prevent neuronal progenitor delamination - PubMed — https://pubmed.ncbi.nlm.nih.gov/35979738/
- **[S10]** Rab6 is increased in Alzheimer's disease brain and correlates with endoplasmic reticulum stress - PubMed — https://pubmed.ncbi.nlm.nih.gov/17573808/
- **[S11]** RAB6A RAB6A, member RAS oncogene family [Homo sapiens (human)] - Gene - NCBI — https://www.ncbi.nlm.nih.gov/gene/5870
- **[S12]** RAB6 and dynein drive post-Golgi apical transport to prevent neuronal progenitor delamination - PubMed — https://pubmed.ncbi.nlm.nih.gov/35979738/
- **[S13]** Rab6 is increased in Alzheimer's disease brain and correlates with endoplasmic reticulum stress - PubMed — https://pubmed.ncbi.nlm.nih.gov/17573808/
- **[S14]** RAB6 and dynein drive post-Golgi apical transport to prevent neuronal progenitor delamination - PubMed — https://pubmed.ncbi.nlm.nih.gov/35979738/
