## Supplementary material for "Understanding Substance Dependence: What Differentiates Addictive from Non-Addictive Drugs?": Folder for supplementary files: non-addictive-genes-claims-first100-077-rab7a-gene-protein-has-been-associated-with-regula-7eb098c11c37.md

# non-addictive-genes-claims-first100-077-7eb098c11c37

**Original claim:** RAB7A gene/protein has been associated with regulatory or homeostatic biological processes in neural systems.

## Claim

RAB7A gene/protein has been associated with regulatory or homeostatic biological processes in neural systems.

## Verdict

Verdict: Highly Supported

Multiple independent lines of evidence support this claim. In human genetics, **RAB7A mutations cause Charcot-Marie-Tooth type 2B (CMT2B), an axonal peripheral neuropathy**, and RAB7A is expressed in sensory and motor neurons. In experimental neural models, RAB7A is linked to **neurotrophin receptor trafficking/signaling, axonal transport, autophagosome-lysosome fusion, mitophagy, lysosomal function, and neuronal survival/degeneration**. ([pubmed.ncbi.nlm.nih.gov](https://pubmed.ncbi.nlm.nih.gov/12545426/?utm_source=openai)) [S1]

## Confidence

High

## Supporting evidence

**1) Human disease genetics directly connect RAB7A to neural biology.**  
The original disease-gene studies showed that missense mutations in **RAB7A cause CMT2B neuropathy**, and the authors reported RAB7 expression in **sensory and motor neurons**. GeneReviews likewise describes normal Rab7a as essential for intracellular membrane trafficking and notes that abnormal Rab7a can lead to lysosomal malfunction and impaired neurite outgrowth. This is strong evidence that RAB7A is not merely a generic trafficking protein, but one with clinically relevant roles in the nervous system. ([pubmed.ncbi.nlm.nih.gov](https://pubmed.ncbi.nlm.nih.gov/12545426/?utm_source=openai)) [S2]

**2) Neuron-focused literature links RAB7A to regulation of trophic signaling and neuronal health.**  
A recent review of Rab7 in neurons describes Rab7-positive signaling endosomes as part of the machinery that balances **local axonal signaling, retrograde transport, signal termination, and nuclear survival signaling**. In that framework, proper Rab7 effector balance helps maintain TrkA trafficking and neuronal health; dysregulation is proposed to impair survival signaling in CMT2B. ([pubmed.ncbi.nlm.nih.gov](https://pubmed.ncbi.nlm.nih.gov/37759799/)) [S3]

**3) Primary neuron experiments show that altered Rab7 function disrupts neuronal regulatory pathways.**  
In rat dorsal root ganglion neurons and PC12-derived neuronal cells, CMT2B-associated Rab7 mutants caused **abnormal axonal transport**, reduced TrkA surface levels/NGF binding, impaired NGF-induced ERK signaling, and produced **axonal degeneration**. These are classic regulatory neuronal processes—growth factor trafficking, survival signaling, and axonal maintenance—rather than incidental associations. ([pubmed.ncbi.nlm.nih.gov](https://pubmed.ncbi.nlm.nih.gov/23616551/)) [S4]

**4) RAB7A is tied to neural homeostasis through autophagy-lysosome biology.**  
Autophagy is a well-established neuronal homeostatic pathway: neurons are highly dependent on constitutive autophagy to maintain proteome and organelle quality, and loss of neuronal autophagy promotes neurodegeneration. Against that background, studies specifically implicate Rab7/RAB7A in late autophagic steps, especially autophagosome maturation and lysosome fusion. ([pubmed.ncbi.nlm.nih.gov](https://pubmed.ncbi.nlm.nih.gov/27251616/)) [S5]

**5) A CNS mouse study linked Rab7-dependent lysosome regulation to neuronal survival and homeostasis.**  
In neurons, the Rab7 effector **WDR91** was shown to be required for controlled Rab7-dependent lysosome fusion. CNS-specific loss of Wdr91 caused **lysosome enlargement/dysfunction, autophagic cargo accumulation, behavioral abnormalities, and marked neuronal loss**. Although the manipulated gene there was Wdr91, the study directly frames the phenotype as a defect in **Rab7-dependent maintenance of late endosome/lysosome homeostasis** in neurons. ([pubmed.ncbi.nlm.nih.gov](https://pubmed.ncbi.nlm.nih.gov/34028500/)) [S6]

**6) Direct neural-cell evidence shows RAB7A supports mitochondrial and lysosomal homeostasis.**  
In N2a neuroblastoma cells exposed to a prion peptide stressor, RAB7A was reported to mediate **mitophagosome formation, autophagosome-lysosome fusion, lysosomal integrity, and mitochondrial homeostasis**; RAB7A depletion impaired mitophagy and increased neuronal apoptosis. This is direct evidence that RAB7A can participate in homeostatic quality-control processes in a neural cell context. ([pubmed.ncbi.nlm.nih.gov](https://pubmed.ncbi.nlm.nih.gov/36449254/)) [S7]

**7) Newer disease-model work also supports a role in neuronal autophagic regulation.**  
A 2025 study found that activating Rab7 promoted **retrograde axonal autophagosome transport** and restored autophagic flux in Alzheimer-model neurons/cells, supporting the broader idea that Rab7-regulated degradative trafficking is functionally important in neurons. ([pubmed.ncbi.nlm.nih.gov](https://pubmed.ncbi.nlm.nih.gov/40490761/)) [S8]

## Contradicting or limiting evidence

**The main limitation is not whether an association exists, but how specifically to define it.**  
Most strong evidence supports **subcellular homeostasis/regulation in neurons**—endolysosomal trafficking, neurotrophin signaling, autophagic flux, lysosome function, mitochondrial quality control, and axonal maintenance. There is much less direct evidence that RAB7A is a regulator of **systems-level neural homeostasis** in the sense of circuit homeostatic plasticity or excitability control. ([pubmed.ncbi.nlm.nih.gov](https://pubmed.ncbi.nlm.nih.gov/37759799/)) [S9]

**Some evidence is indirect or model-dependent.**  
Several studies rely on cultured cells, neuron-like N2a cells, fibroblasts, or disease models rather than normal intact mammalian neural circuits. That still supports the claim as written, but it means the strongest conclusion is about **cellular/organellar regulation in neural systems**, not every possible meaning of “homeostatic” in neuroscience. ([pubmed.ncbi.nlm.nih.gov](https://pubmed.ncbi.nlm.nih.gov/36449254/)) [S10]

**There is also mechanistic disagreement about how disease-causing Rab7 mutations act.**  
Some work argues CMT2B mutations cause **misregulated activation/uncontrolled nucleotide exchange**, whereas other work in Drosophila supports a **partial loss-of-function/dosage-sensitive** model. This disagreement affects fine mechanistic interpretation, but not the broader conclusion that Rab7/RAB7A is deeply involved in neuronal regulatory/homeostatic pathways. ([pubmed.ncbi.nlm.nih.gov](https://pubmed.ncbi.nlm.nih.gov/20028791/)) [S11]

## Mechanistic / biological rationale

RAB7A encodes a **small Rab-family GTPase** that regulates **late endosome maturation and trafficking from late endosomes to lysosomes**. That role places it at a key control point for cargo degradation, organelle positioning, and membrane fusion. ([ncbi.nlm.nih.gov](https://www.ncbi.nlm.nih.gov/gene/7879)) [S12]

In neurons, these functions are especially important because neurons are polarized, long-lived, and heavily reliant on constitutive degradative quality control. Neuronal autophagy is itself considered a core homeostatic process, particularly in axons, where cargo must often be formed distally and transported retrogradely for maturation and degradation. ([pubmed.ncbi.nlm.nih.gov](https://pubmed.ncbi.nlm.nih.gov/27251616/)) [S13]

Rab7-positive endosomes also intersect with **neurotrophin receptor trafficking**. The neuron-focused Rab7 review describes Rab7 as helping position signaling endosomes between recycling, retrograde transport, maturation, and lysosomal degradation; in that way, Rab7 can shape the balance between **local signaling and nuclear survival signaling**. ([pubmed.ncbi.nlm.nih.gov](https://pubmed.ncbi.nlm.nih.gov/37759799/)) [S14]

Rab7/RAB7A is also mechanistically tied to **autophagosome-lysosome fusion and mitophagy**, which explains why disturbed Rab7 function can lead to lysosomal dysfunction, cargo accumulation, mitochondrial stress, and neuronal degeneration. ([pubmed.ncbi.nlm.nih.gov](https://pubmed.ncbi.nlm.nih.gov/34028500/)) [S15]

## Bottom line

Yes—**RAB7A is strongly associated with regulatory and homeostatic processes in neural systems**. The best-supported interpretation is that RAB7A contributes to **neuronal intracellular homeostasis**, especially through **endolysosomal trafficking, neurotrophin-signaling endosome regulation, autophagic/mitophagic flux, lysosomal function, and axonal maintenance**. The evidence is strongest for **cellular and organellar homeostasis in neurons**, rather than for circuit-level homeostatic plasticity specifically. ([pubmed.ncbi.nlm.nih.gov](https://pubmed.ncbi.nlm.nih.gov/12545426/?utm_source=openai)) [S16]

## Cited sources

- **[S1]** Mutations in the small GTP-ase late endosomal protein RAB7 cause Charcot-Marie-Tooth type 2B neuropathy - PubMed — https://pubmed.ncbi.nlm.nih.gov/12545426/?utm_source=openai
- **[S2]** Mutations in the small GTP-ase late endosomal protein RAB7 cause Charcot-Marie-Tooth type 2B neuropathy - PubMed — https://pubmed.ncbi.nlm.nih.gov/12545426/?utm_source=openai
- **[S3]** Regulation of Endosomal Trafficking by Rab7 and Its Effectors in Neurons: Clues from Charcot-Marie-Tooth 2B Disease - PubMed — https://pubmed.ncbi.nlm.nih.gov/37759799/
- **[S4]** Defective axonal transport of Rab7 GTPase results in dysregulated trophic signaling - PubMed — https://pubmed.ncbi.nlm.nih.gov/23616551/
- **[S5]** Compartment-Specific Regulation of Autophagy in Primary Neurons - PubMed — https://pubmed.ncbi.nlm.nih.gov/27251616/
- **[S6]** The Rab7 effector WDR91 promotes autophagy-lysosome degradation in neurons by regulating lysosome fusion - PubMed — https://pubmed.ncbi.nlm.nih.gov/34028500/
- **[S7]** RAB7A GTPase Is Involved in Mitophagosome Formation and Autophagosome-Lysosome Fusion in N2a Cells Treated with the Prion Protein Fragment 106-126 - PubMed — https://pubmed.ncbi.nlm.nih.gov/36449254/
- **[S8]** AP2A1 activates Rab7 to promote axonal autophagosome transport and slow the progression of Alzheimer's disease - PubMed — https://pubmed.ncbi.nlm.nih.gov/40490761/
- **[S9]** Regulation of Endosomal Trafficking by Rab7 and Its Effectors in Neurons: Clues from Charcot-Marie-Tooth 2B Disease - PubMed — https://pubmed.ncbi.nlm.nih.gov/37759799/
- **[S10]** RAB7A GTPase Is Involved in Mitophagosome Formation and Autophagosome-Lysosome Fusion in N2a Cells Treated with the Prion Protein Fragment 106-126 - PubMed — https://pubmed.ncbi.nlm.nih.gov/36449254/
- **[S11]** Disease mutations in Rab7 result in unregulated nucleotide exchange and inappropriate activation - PubMed — https://pubmed.ncbi.nlm.nih.gov/20028791/
- **[S12]** RAB7A RAB7A, member RAS oncogene family [Homo sapiens (human)] - Gene - NCBI — https://www.ncbi.nlm.nih.gov/gene/7879
- **[S13]** Compartment-Specific Regulation of Autophagy in Primary Neurons - PubMed — https://pubmed.ncbi.nlm.nih.gov/27251616/
- **[S14]** Regulation of Endosomal Trafficking by Rab7 and Its Effectors in Neurons: Clues from Charcot-Marie-Tooth 2B Disease - PubMed — https://pubmed.ncbi.nlm.nih.gov/37759799/
- **[S15]** The Rab7 effector WDR91 promotes autophagy-lysosome degradation in neurons by regulating lysosome fusion - PubMed — https://pubmed.ncbi.nlm.nih.gov/34028500/
- **[S16]** Mutations in the small GTP-ase late endosomal protein RAB7 cause Charcot-Marie-Tooth type 2B neuropathy - PubMed — https://pubmed.ncbi.nlm.nih.gov/12545426/?utm_source=openai
