## Supplementary material for "Understanding Substance Dependence: What Differentiates Addictive from Non-Addictive Drugs?": Folder for supplementary files: non-addictive-genes-claims-first100-078-pde2a-gene-protein-has-been-associated-with-regula-8b043b549477.md

# non-addictive-genes-claims-first100-078-8b043b549477

**Original claim:** PDE2A gene/protein has been associated with regulatory or homeostatic biological processes in neural systems.

## Claim

PDE2A gene/protein has been associated with regulatory or homeostatic biological processes in neural systems.

## Verdict

Verdict: Highly Supported

PDE2A is well associated with **regulatory functions in neural systems**. Across gene annotation resources, neuroanatomical mapping, electrophysiology, pharmacology, and rare human genetics, PDE2A is repeatedly implicated in regulating neuronal cyclic-nucleotide signaling, neurotransmitter release, synaptic plasticity, pain processing, and circuit-level function. The evidence for **“regulatory”** roles is strong. The evidence for **“homeostatic”** roles is more indirect, usually inferred from PDE2A’s role in constraining cAMP/cGMP signaling microdomains rather than from experiments explicitly designed around formal neural homeostasis paradigms. ([ncbi.nlm.nih.gov](https://www.ncbi.nlm.nih.gov/gene/5138)) [S1]

## Confidence

High

## Supporting evidence

PDE2A encodes a dual-substrate phosphodiesterase that hydrolyzes both cAMP and cGMP, and the current NCBI gene summary lists it as involved in **regulation of signal transduction**. NCBI also reports substantial brain expression. That alone does not prove a neural regulatory role, but it is consistent with one and fits the much more specific functional literature. ([ncbi.nlm.nih.gov](https://www.ncbi.nlm.nih.gov/gene/5138)) [S2]

Neuroanatomical studies provide strong localization evidence. In a cross-species immunohistochemical study, PDE2A protein was reported to be **most abundant in brain relative to peripheral tissues**, with prominent expression in cortex, hippocampus, amygdala, habenula, basal ganglia, and interpeduncular nucleus. A rat-brain mapping study likewise found PDE2A mRNA in cortical and hippocampal pyramidal neurons, dentate granule cells, and striatal medium spiny neurons, with protein concentrated especially in **axons and nerve terminals**. The authors explicitly concluded that PDE2A likely modulates cortical, hippocampal, and striatal networks and may regulate **axonal excitability or transmitter release**. ([pubmed.ncbi.nlm.nih.gov](https://pubmed.ncbi.nlm.nih.gov/19506089/)) [S3]

There is also direct functional evidence that PDE2A regulates synaptic signaling. In the medial habenula–interpeduncular nucleus pathway, natriuretic peptides were shown to **block glutamate release** through activation of PDE2A, reducing presynaptic PKA signaling; this effect was reversed by PDE2A inhibition. That is a direct demonstration of PDE2A participating in presynaptic regulation of neurotransmission in a defined neural circuit. ([pubmed.ncbi.nlm.nih.gov](https://pubmed.ncbi.nlm.nih.gov/23045693/?utm_source=openai)) [S4]

A separate study in striatal medium spiny neurons showed that NO/cGMP signaling suppresses transient cAMP signals through **PDE2 activation**, with effects extending downstream to PKA. This places PDE2A/PDE2 at a key control point for cyclic-nucleotide cross-talk in striatal neurons, a canonical regulatory role in neural signaling. ([pubmed.ncbi.nlm.nih.gov](https://pubmed.ncbi.nlm.nih.gov/24302895/)) [S5]

PDE2 inhibition also alters higher-order neural plasticity phenotypes. In cultured neurons and hippocampal slices, selective PDE2 inhibition increased neuronal cGMP, enhanced **long-term potentiation**, and improved performance in rodent memory tasks without altering basal synaptic transmission. That pattern strongly supports a role in regulating activity-dependent synaptic plasticity rather than simply causing nonspecific excitation. ([pubmed.ncbi.nlm.nih.gov](https://pubmed.ncbi.nlm.nih.gov/15555642/)) [S6]

Additional disease-model work points in the same direction. In a chronic-stress model, PDE2 inhibition reversed stress-induced deficits in learning and memory, improved CA1 dendritic branching and spine density, and normalized signaling molecules linked to plasticity, including CREB and BDNF-related pathways. After cortical stroke, PDE2A inhibition improved motor recovery and increased axonal projections and functional connectivity in peri-infarct cortex. These data extend PDE2A’s regulatory association from synapses to structural and circuit remodeling. ([pubmed.ncbi.nlm.nih.gov](https://pubmed.ncbi.nlm.nih.gov/25442113/)) [S7]

PDE2A has also been linked to sensory processing in the nervous system. In mouse spinal cord, PDE2A was localized to neurons in the superficial dorsal horn, was upregulated after peripheral inflammation, and pharmacologic inhibition altered inflammatory pain behavior while increasing spinal cAMP. This is another direct example of PDE2A regulating neural signaling outputs in vivo. ([pubmed.ncbi.nlm.nih.gov](https://pubmed.ncbi.nlm.nih.gov/24758774/)) [S8]

Human genetics further strengthens the association. A homozygous loss-of-function PDE2A variant was associated with **early-onset hereditary chorea**, and the mutant protein showed markedly reduced enzymatic activity. A later study found additional biallelic PDE2A variants causing syndromic paroxysmal dyskinesia with developmental or cognitive impairment, sometimes with EEG abnormalities or epilepsy. These are rare disorders, but they support the idea that intact PDE2A function is important for regulation of human neural circuit physiology, especially in striatal systems. ([pubmed.ncbi.nlm.nih.gov](https://pubmed.ncbi.nlm.nih.gov/29392776/)) [S9]

Reviews of the field summarize the same picture: PDE2/PDE2A is enriched in forebrain/limbic regions and is increasingly viewed as relevant to cognition, mood, pain, and neurodegenerative or neuropsychiatric conditions because of its control over cAMP/cGMP signaling in the CNS. ([pubmed.ncbi.nlm.nih.gov](https://pubmed.ncbi.nlm.nih.gov/28956338/)) [S10]

## Contradicting or limiting evidence

The main limitation is semantic and evidentiary: most studies support PDE2A as a **regulator of neural signaling**, but relatively few explicitly frame it as a mediator of **neural homeostasis**. So if “homeostatic” is interpreted narrowly—e.g., classic firing-rate homeostasis, synaptic scaling, or glial-neuronal homeostatic maintenance—the evidence is thinner and more inferential. ([pubmed.ncbi.nlm.nih.gov](https://pubmed.ncbi.nlm.nih.gov/23000621/)) [S11]

A second limitation is that much of the strongest causal evidence comes from **rodent, slice, or pharmacologic** studies rather than human interventional studies. Human evidence exists, but it is mostly from rare genetic disorders rather than broad population or clinical-trial data. ([pubmed.ncbi.nlm.nih.gov](https://pubmed.ncbi.nlm.nih.gov/15555642/)) [S12]

PDE2A’s effects are also **context-dependent**, not uniformly protective. For example, inhibiting PDE2A may improve cognition or post-stroke connectivity in some models, but in spinal cord inflammatory pain models it increased nociceptive behavior. That argues for PDE2A as a signaling regulator or buffer rather than a universally beneficial factor. ([pubmed.ncbi.nlm.nih.gov](https://pubmed.ncbi.nlm.nih.gov/36163142/)) [S13]

Finally, PDE2A is not neural-specific. It is expressed in multiple tissues, and mouse whole-body loss of PDE2A is embryonically lethal, showing important systemic roles. That broad biology does not weaken the neural association, but it does mean PDE2A is part of wider regulatory physiology, not an exclusively neural homeostasis gene. ([ncbi.nlm.nih.gov](https://www.ncbi.nlm.nih.gov/gene/5138)) [S14]

## Mechanistic / biological rationale

The mechanistic rationale is strong. PDE2A hydrolyzes both cAMP and cGMP, and structural work showed that **cGMP binding to its GAF regulatory domain activates catalytic activity**. This gives PDE2A a built-in way to couple cGMP signals to suppression of cAMP signaling. In neurons, that makes PDE2A a plausible “brake” or constraint on second-messenger cascades. ([pubmed.ncbi.nlm.nih.gov](https://pubmed.ncbi.nlm.nih.gov/12271124/)) [S15]

Its localization supports that function. PDE2A is enriched in axons, terminals, and specific neuronal populations in cortex, hippocampus, striatum, habenula, and spinal dorsal horn, placing it exactly where control of local cyclic-nucleotide microdomains would affect neurotransmitter release, excitability, and plasticity. ([pubmed.ncbi.nlm.nih.gov](https://pubmed.ncbi.nlm.nih.gov/23000621/)) [S16]

Functionally, the literature shows this mechanism operating in neural systems: PDE2A links natriuretic peptide/cGMP signaling to reduced presynaptic PKA and transmitter release, and links NO/cGMP to suppression of transient cAMP/PKA signaling in striatal neurons. Those are classic examples of regulatory signal integration. From a biological standpoint, such compartmentalized control is very compatible with a **homeostatic buffering** role, even if most papers do not use that exact term. ([pubmed.ncbi.nlm.nih.gov](https://pubmed.ncbi.nlm.nih.gov/23045693/?utm_source=openai)) [S17]

## Bottom line

Yes—this claim is well supported. PDE2A is strongly associated with **regulatory biological processes in neural systems**, including control of cyclic-nucleotide signaling, presynaptic neurotransmitter release, synaptic plasticity, pain processing, and circuit function. The case for **homeostatic** involvement is plausible and mechanistically sensible, but the literature more directly supports **neural regulation** than narrowly defined neural homeostasis. ([pubmed.ncbi.nlm.nih.gov](https://pubmed.ncbi.nlm.nih.gov/23000621/)) [S18]

## Cited sources

- **[S1]** PDE2A phosphodiesterase 2A [Homo sapiens (human)] - Gene - NCBI — https://www.ncbi.nlm.nih.gov/gene/5138
- **[S2]** PDE2A phosphodiesterase 2A [Homo sapiens (human)] - Gene - NCBI — https://www.ncbi.nlm.nih.gov/gene/5138
- **[S3]** Immunohistochemical localization of phosphodiesterase 2A in multiple mammalian species - PubMed — https://pubmed.ncbi.nlm.nih.gov/19506089/
- **[S4]** Natriuretic peptides block synaptic transmission by activating phosphodiesterase 2A and reducing presynaptic PKA activity - PubMed — https://pubmed.ncbi.nlm.nih.gov/23045693/?utm_source=openai
- **[S5]** The NO/cGMP pathway inhibits transient cAMP signals through the activation of PDE2 in striatal neurons - PubMed — https://pubmed.ncbi.nlm.nih.gov/24302895/
- **[S6]** Inhibition of phosphodiesterase 2 increases neuronal cGMP, synaptic plasticity and memory performance - PubMed — https://pubmed.ncbi.nlm.nih.gov/15555642/
- **[S7]** Inhibition of phosphodiesterase 2 reverses impaired cognition and neuronal remodeling caused by chronic stress - PubMed — https://pubmed.ncbi.nlm.nih.gov/25442113/
- **[S8]** Phosphodiesterase 2A localized in the spinal cord contributes to inflammatory pain processing - PubMed — https://pubmed.ncbi.nlm.nih.gov/24758774/
- **[S9]** A homozygous loss-of-function mutation in PDE2A associated to early-onset hereditary chorea - PubMed — https://pubmed.ncbi.nlm.nih.gov/29392776/
- **[S10]** The Role of Phosphodiesterase-2 in Psychiatric and Neurodegenerative Disorders - PubMed — https://pubmed.ncbi.nlm.nih.gov/28956338/
- **[S11]** The distribution of phosphodiesterase 2A in the rat brain - PubMed — https://pubmed.ncbi.nlm.nih.gov/23000621/
- **[S12]** Inhibition of phosphodiesterase 2 increases neuronal cGMP, synaptic plasticity and memory performance - PubMed — https://pubmed.ncbi.nlm.nih.gov/15555642/
- **[S13]** PDE2A Inhibition Enhances Axonal Sprouting, Functional Connectivity, and Recovery after Stroke - PubMed — https://pubmed.ncbi.nlm.nih.gov/36163142/
- **[S14]** PDE2A phosphodiesterase 2A [Homo sapiens (human)] - Gene - NCBI — https://www.ncbi.nlm.nih.gov/gene/5138
- **[S15]** The two GAF domains in phosphodiesterase 2A have distinct roles in dimerization and in cGMP binding - PubMed — https://pubmed.ncbi.nlm.nih.gov/12271124/
- **[S16]** The distribution of phosphodiesterase 2A in the rat brain - PubMed — https://pubmed.ncbi.nlm.nih.gov/23000621/
- **[S17]** Natriuretic peptides block synaptic transmission by activating phosphodiesterase 2A and reducing presynaptic PKA activity - PubMed — https://pubmed.ncbi.nlm.nih.gov/23045693/?utm_source=openai
- **[S18]** The distribution of phosphodiesterase 2A in the rat brain - PubMed — https://pubmed.ncbi.nlm.nih.gov/23000621/
