## Supplementary material for "Understanding Substance Dependence: What Differentiates Addictive from Non-Addictive Drugs?": Folder for supplementary files: non-addictive-genes-claims-first100-079-ngfr-gene-protein-has-been-associated-with-regulat-1b66e2c5684e.md

# non-addictive-genes-claims-first100-079-1b66e2c5684e

**Original claim:** NGFR gene/protein has been associated with regulatory or homeostatic biological processes in neural systems.

## Claim

NGFR gene/protein has been associated with regulatory or homeostatic biological processes in neural systems.

## Verdict

Verdict: Highly Supported

NGFR encodes the p75 neurotrophin receptor (p75NTR; TNFRSF16/CD271), a receptor that is extensively implicated in regulation of neural development, neuronal survival versus apoptosis, axon growth and pruning, neurogenesis, synaptic plasticity, and some explicitly homeostatic neural functions. The “regulatory” part of the claim is very strongly supported. The “homeostatic” part is also supported, but more specifically in circadian regulation, hypothalamic feeding responses, and hippocampal synaptic plasticity, with most direct evidence coming from animal and cell studies rather than human intervention data. ([pubmed.ncbi.nlm.nih.gov](https://pubmed.ncbi.nlm.nih.gov/26109945/)) [S1]

## Confidence

High

This conclusion is supported by convergent evidence from curated gene annotations, reviews, and multiple primary studies across development, adult neurogenesis, circuit maturation, and neural plasticity. The main limitation is that much of the strongest mechanistic evidence is preclinical. ([ncbi.nlm.nih.gov](https://www.ncbi.nlm.nih.gov/gene/4804)) [S2]

## Supporting evidence

Human NGFR gene annotations at NCBI/GO already place the gene in biologic processes directly relevant to neural regulation, including **nervous system development**, **Rho protein signal transduction**, **circadian regulation of gene expression**, and **presynaptic modulation of chemical synaptic transmission**. These annotations are not all equally strong—some are direct-assay or traceable-author statements, others are computational/electronic—but they are directionally consistent with NGFR being tied to regulatory and homeostatic neural biology. ([ncbi.nlm.nih.gov](https://www.ncbi.nlm.nih.gov/gene/4804)) [S3]

High-level reviews describe p75NTR as a multifunctional regulator rather than a passive marker. A 2015 review states that p75NTR has emerged as a target for control of neurotrophin activity because of evidence for **regulation of neural plasticity in the mature nervous system**, **promotion of adult neurogenesis**, and control of **neuron survival** and other processes supporting neural function. An earlier review likewise summarized p75NTR as participating in a “diverse array” of cellular responses including **apoptosis, neurite outgrowth, and myelination**, with downstream signaling through NF-κB, JNK, and Rac. ([pubmed.ncbi.nlm.nih.gov](https://pubmed.ncbi.nlm.nih.gov/26109945/)) [S4]

Direct developmental evidence is strong. In a conditional mouse knockout, deleting p75NTR from embryonic neural progenitors increased progenitor apoptosis, reduced Tbr2-positive progenitors, halved neurogenesis, and led to fewer cortical pyramidal neurons, interneurons, basal forebrain cholinergic neurons, and striatal neurons in adulthood. That is a clear demonstration that NGFR/p75NTR regulates neural progenitor survival and proper nervous-system formation. ([pubmed.ncbi.nlm.nih.gov](https://pubmed.ncbi.nlm.nih.gov/31488566/)) [S5]

Adult neurogenesis data point in the same direction. In the subventricular zone, p75NTR-positive cells were reported to account for the neurosphere-producing precursor population, and p75NTR-deficient mice showed markedly reduced neurogenic potential and fewer neuroblasts in vivo. In the dentate gyrus, p75NTR was expressed in proliferating progenitors, and knockout mice showed a large decrease in BrdU-positive cells, supporting a role in proliferation and/or early maturation of neural and glial lineages. ([pubmed.ncbi.nlm.nih.gov](https://pubmed.ncbi.nlm.nih.gov/17494700/)) [S6]

NGFR/p75NTR also regulates circuit formation and plasticity. In cortical parvalbumin interneurons, p75NTR expression was shown to restrain the maturation of inhibitory connectivity; conditional loss caused precocious perisomatic innervation and perineuronal-net formation, while activating p75NTR in adults destabilized PV connectivity and restored cortical plasticity. In the hippocampus, p75NTR was described as a negative regulator of structural and functional plasticity, and knockout mice were resistant to age-related changes in synaptic plasticity and memory; the authors explicitly concluded that p75NTR is important for maintaining homeostasis of hippocampal synaptic plasticity. ([pubmed.ncbi.nlm.nih.gov](https://pubmed.ncbi.nlm.nih.gov/30936240/)) [S7]

The receptor is also tied to axon-growth control and structural refinement. A classic Neuron study showed that p75NTR regulates RhoA in a ligand-dependent manner: p75NTR activated RhoA, neurotrophin binding abolished that activation, and p75NTR mutation retarded axonal outgrowth in vivo. A separate Nature Neuroscience study identified **developmental axon pruning mediated by BDNF-p75NTR-dependent axon degeneration**, directly linking NGFR to activity-dependent circuit refinement. ([pubmed.ncbi.nlm.nih.gov](https://pubmed.ncbi.nlm.nih.gov/10595511/)) [S8]

For the specifically **homeostatic** side of the claim, there are two notable examples. First, in the hypothalamus, p75NTR in AgRP neurons was reported to be **necessary for homeostatic feeding and food anticipation**, indicating a role in behavioral responses to energy deficit. Second, p75NTR was shown to be a CLOCK/BMAL1-regulated oscillating gene in the suprachiasmatic nucleus (SCN), and loss of p75NTR altered oscillation of clock genes in the SCN and other tissues; the authors concluded it plays an important role in maintenance of clock and metabolic gene oscillation. Those are bona fide homeostatic/regulatory neural functions. ([pmc.ncbi.nlm.nih.gov](https://pmc.ncbi.nlm.nih.gov/articles/PMC7056271/)) [S9]

## Contradicting or limiting evidence

The main caveat is not absence of evidence, but **context dependence**. p75NTR can support survival in some neural contexts and apoptosis in others. Reviews emphasize that p75NTR’s outputs depend on ligand form, cell state, and binding partners such as Trk receptors, sortilin, and the Nogo receptor. A primary study in neocortical subplate neurons even found that p75NTR promoted survival rather than death, contrary to its better-known pro-apoptotic role in other settings. So NGFR is clearly regulatory, but not in a single-direction or uniformly “homeostatic” way. ([pubmed.ncbi.nlm.nih.gov](https://pubmed.ncbi.nlm.nih.gov/26109945/)) [S10]

Also, some homeostatic effects appear conditional rather than baseline-universal. In the AgRP-neuron study, p75NTR deletion did **not** alter locomotor activity or metabolism under normal ad libitum chow conditions; its role became evident under homeostatic challenge such as fasting/scheduled feeding. That supports a role in adaptive homeostatic response, but not necessarily a constitutive requirement for all baseline neural homeostasis. ([pmc.ncbi.nlm.nih.gov](https://pmc.ncbi.nlm.nih.gov/articles/PMC7056271/)) [S11]

A further limitation is translational. Most direct functional evidence comes from mouse genetics, rat tissue, or cultured neurons. That is strong for biological plausibility and association, but it is not the same as direct proof that NGFR governs neural homeostasis in humans at the organismal level. In addition, some annotation-level evidence in NCBI Gene/GO is computational rather than experimentally direct. ([ncbi.nlm.nih.gov](https://www.ncbi.nlm.nih.gov/gene/4804)) [S12]

There is also some stage specificity. In the dentate-gyrus literature summarized by Bernabeu and Longo, prior work found fewer early BrdU-positive cells in p75NTR-null mice but no difference in BrdU/NeuN double-labeled cells at six weeks, suggesting that p75NTR’s effects may be strongest on early proliferation or maturation rather than on all later outcomes. ([link.springer.com](https://link.springer.com/article/10.1186/1471-2202-11-136)) [S13]

## Mechanistic / biological rationale

Mechanistically, NGFR/p75NTR is well positioned to regulate neural processes because it is a signaling hub rather than a simple ligand-binding receptor. It belongs to the TNF receptor superfamily, lacks intrinsic catalytic activity, and exerts its effects through receptor complexes and adaptor proteins. Reviews and primary studies show that p75NTR can:  
- enhance TrkA/TrkB/TrkC-mediated trophic signaling and neuronal survival,  
- partner with sortilin to bind proneurotrophins and trigger apoptosis, and  
- partner with Nogo receptor complexes to regulate RhoA-dependent growth-cone collapse, neurite retraction, and spine-related structural remodeling. ([pubmed.ncbi.nlm.nih.gov](https://pubmed.ncbi.nlm.nih.gov/26109945/)) [S14]

This signaling logic explains why NGFR repeatedly appears in regulatory and homeostatic neural phenomena. By modulating PI3K-Akt survival signaling, NF-κB/JNK stress pathways, and RhoA-dependent cytoskeletal dynamics, p75NTR can influence progenitor maintenance, axon extension versus retraction, developmental pruning, synaptic remodeling, and timing of circuit maturation. Its role in SCN clock-gene oscillation and hypothalamic feeding circuits further fits the idea that NGFR can participate in neural systems that stabilize physiology across time and metabolic state. ([pubmed.ncbi.nlm.nih.gov](https://pubmed.ncbi.nlm.nih.gov/31736712/)) [S15]

## Bottom line

Yes. The claim is well supported. NGFR/p75NTR is strongly associated with **regulatory** biological processes in neural systems, including neural development, progenitor survival, axon dynamics, circuit refinement, and synaptic plasticity. There is also credible, mechanistically grounded evidence for **homeostatic** roles in specific neural contexts such as circadian regulation, hypothalamic feeding responses, and hippocampal synaptic plasticity. The main qualification is that these homeostatic roles are context-specific and supported mostly by preclinical rather than direct human functional studies. ([pubmed.ncbi.nlm.nih.gov](https://pubmed.ncbi.nlm.nih.gov/26109945/)) [S16]

## Cited sources

- **[S1]** The p75 neurotrophin receptor: at the crossroad of neural repair and death - PubMed — https://pubmed.ncbi.nlm.nih.gov/26109945/
- **[S2]** NGFR nerve growth factor receptor [Homo sapiens (human)] - Gene - NCBI — https://www.ncbi.nlm.nih.gov/gene/4804
- **[S3]** NGFR nerve growth factor receptor [Homo sapiens (human)] - Gene - NCBI — https://www.ncbi.nlm.nih.gov/gene/4804
- **[S4]** The p75 neurotrophin receptor: at the crossroad of neural repair and death - PubMed — https://pubmed.ncbi.nlm.nih.gov/26109945/
- **[S5]** The p75 neurotrophin receptor is required for the survival of neuronal progenitors and normal formation of the basal forebrain, striatum, thalamus and neocortex - PubMed — https://pubmed.ncbi.nlm.nih.gov/31488566/
- **[S6]** p75 neurotrophin receptor expression defines a population of BDNF-responsive neurogenic precursor cells - PubMed — https://pubmed.ncbi.nlm.nih.gov/17494700/
- **[S7]** p75 Neurotrophin Receptor Activation Regulates the Timing of the Maturation of Cortical Parvalbumin Interneuron Connectivity and Promotes Juvenile-like Plasticity in Adult Visual Cortex - PubMed — https://pubmed.ncbi.nlm.nih.gov/30936240/
- **[S8]** Neurotrophin binding to the p75 receptor modulates Rho activity and axonal outgrowth - PubMed — https://pubmed.ncbi.nlm.nih.gov/10595511/
- **[S9]** 
            The p75 neurotrophin receptor in AgRP neurons is necessary for homeostatic feeding and food anticipation - PMC
         — https://pmc.ncbi.nlm.nih.gov/articles/PMC7056271/
- **[S10]** The p75 neurotrophin receptor: at the crossroad of neural repair and death - PubMed — https://pubmed.ncbi.nlm.nih.gov/26109945/
- **[S11]** 
            The p75 neurotrophin receptor in AgRP neurons is necessary for homeostatic feeding and food anticipation - PMC
         — https://pmc.ncbi.nlm.nih.gov/articles/PMC7056271/
- **[S12]** NGFR nerve growth factor receptor [Homo sapiens (human)] - Gene - NCBI — https://www.ncbi.nlm.nih.gov/gene/4804
- **[S13]** The p75 neurotrophin receptor is expressed by adult mouse dentate progenitor cells and regulates neuronal and non-neuronal cell genesis | BMC Neuroscience | Springer Nature Link — https://link.springer.com/article/10.1186/1471-2202-11-136
- **[S14]** The p75 neurotrophin receptor: at the crossroad of neural repair and death - PubMed — https://pubmed.ncbi.nlm.nih.gov/26109945/
- **[S15]** The p75 Neurotrophin Receptor Facilitates TrkB Signaling and Function in Rat Hippocampal Neurons - PubMed — https://pubmed.ncbi.nlm.nih.gov/31736712/
- **[S16]** The p75 neurotrophin receptor: at the crossroad of neural repair and death - PubMed — https://pubmed.ncbi.nlm.nih.gov/26109945/
