## Supplementary material for "Understanding Substance Dependence: What Differentiates Addictive from Non-Addictive Drugs?": Folder for supplementary files: non-addictive-genes-claims-first100-080-reep5-gene-protein-has-been-associated-with-regula-b19860f2fe0a.md

# non-addictive-genes-claims-first100-080-b19860f2fe0a

**Original claim:** REEP5 gene/protein has been associated with regulatory or homeostatic biological processes in neural systems.

## Claim

REEP5 gene/protein has been associated with regulatory or homeostatic biological processes in neural systems.

## Verdict

Verdict: Likely True

## Confidence

Medium

## Supporting evidence

REEP5 is expressed in nervous-system tissue, including brain, and authoritative gene resources annotate it as being involved in **endoplasmic reticulum (ER) membrane organization** and **regulation of intracellular transport**. NCBI lists substantial brain expression for human REEP5 and summarizes it as localized to the ER tubular network with predicted roles in ER membrane organization and intracellular transport. ([ncbi.nlm.nih.gov](https://www.ncbi.nlm.nih.gov/gene/7905)) [S1]

Independent expression resources also place REEP5 in a brain-expression cluster labeled **“Neurons & Synapses – Synaptic function (mainly)”**, which is not proof of function by itself but does support neural-system association. ([proteinatlas.org](https://www.proteinatlas.org/ENSG00000129625-REEP5)) [S2]

Although a 2014 Brain Research paper focused mainly on REEP1/REEP2, it reported that **REEP5 mRNA is highly expressed in superior cervical and stellate sympathetic ganglia**, indicating enrichment in peripheral neural tissue. ([pubmed.ncbi.nlm.nih.gov](https://pubmed.ncbi.nlm.nih.gov/24355597/)) [S3]

At the mechanistic level, REEP5 belongs to the REEP/DP1/Yop1p family of ER-shaping proteins. A neuroscience review on neuronal ER morphology states that REEP proteins help shape curved ER tubules, and specifically notes that **REEP5 interacts with reticulons to form and stabilize ER membrane curvature**. The same review emphasizes that controlled ER organization and turnover are important for neuronal health and function. ([mdpi.com](https://www.mdpi.com/2073-4409/10/5/970)) [S4]

There is also disease-context evidence from neural tissue. In an Alzheimer’s disease-related mouse/human pathology study, REEP5 immunostaining marked **dystrophic neurites** and abnormal accumulations of tubular ER around amyloid plaques, linking REEP5-positive ER structures to disrupted neuritic homeostasis in brain tissue. ([pubmed.ncbi.nlm.nih.gov](https://pubmed.ncbi.nlm.nih.gov/26619807/)) [S5]

More recently, in a dopaminergic neurodegeneration model caused by **WDR45 deficiency**, REEP5 was used to identify abnormal **tubular ER-positive axonal enlargements** in striatal dopaminergic projections, again associating REEP5 with axonal ER organization in a neural pathology context. ([pubmed.ncbi.nlm.nih.gov](https://pubmed.ncbi.nlm.nih.gov/39183331/)) [S6]

Finally, outside the nervous system but still relevant to the biology of “homeostasis,” cell-biological studies show that REEP5 can regulate organelle organization. A 2024 study found that REEP5 interacts with MFN1/2 to mediate ER-mitochondria tethering, mitochondrial distribution, and mitochondrial ROS levels, all classic homeostatic processes. ([pubmed.ncbi.nlm.nih.gov](https://pubmed.ncbi.nlm.nih.gov/39133213/)) [S7] A separate commentary on REEP5 biology concluded that REEP5 is centrally involved in SR/ER organization and cellular stress responses. ([pubmed.ncbi.nlm.nih.gov](https://pubmed.ncbi.nlm.nih.gov/32548572/)) [S8]

## Contradicting or limiting evidence

The biggest limitation is that **direct causal evidence in neurons is limited**. Much of the strongest mechanistic work on REEP5 homeostatic function comes from **U2OS, MEFs, HEK293T, or cardiac myocyte systems**, not from central neurons or intact neural circuits. ([pubmed.ncbi.nlm.nih.gov](https://pubmed.ncbi.nlm.nih.gov/39133213/)) [S9]

The neural evidence is therefore often **associational** rather than definitive: brain/sympathetic-ganglion expression, placement in a neuron/synapse-enriched expression cluster, and localization to pathological neurites or axonal enlargements. Those findings support association with neural ER organization, but they do **not** yet show that REEP5 is a proven regulator of synaptic homeostasis, firing-rate homeostasis, neurotransmitter balance, or other canonical neural homeostatic programs. ([pubmed.ncbi.nlm.nih.gov](https://pubmed.ncbi.nlm.nih.gov/24355597/)) [S10]

Also, compared with other REEP family members such as **REEP1**, which has clearer links to hereditary spastic paraplegia and neuron-specific ER biology, REEP5 has much less direct human neurogenetic evidence. Reviews of the REEP family discuss neurological disease prominently, but REEP5-specific neural disease causality remains less established. ([pubmed.ncbi.nlm.nih.gov](https://pubmed.ncbi.nlm.nih.gov/36191880/)) [S11]

## Mechanistic / biological rationale

Biologically, the claim is plausible. Neurons rely heavily on **distributed ER networks** that extend into axons and dendrites, where they support membrane trafficking, protein handling, lipid metabolism, organelle contact sites, and calcium-related signaling. Reviews of neuronal ER biology emphasize that the shape and maintenance of ER tubules are integral to neuronal function and vulnerability in neurodegeneration. ([mdpi.com](https://www.mdpi.com/2073-4409/10/5/970)) [S12]

REEP5 fits this framework because it is an **ER tubule-shaping protein**. NCBI and review-level sources place it in ER membrane organization, and the neuronal ER review specifically describes REEP5/DP1 as a reticulon partner that stabilizes membrane curvature. ([ncbi.nlm.nih.gov](https://www.ncbi.nlm.nih.gov/gene/7905)) [S13]

That matters for homeostasis because ER architecture influences **intracellular transport**, **ER-mitochondria contact sites**, and downstream control of **redox state/ROS**. The 2024 REEP5-MFN1/2 study showed that altering REEP5 changes mitochondrial positioning and ROS production, which are broadly homeostatic functions and are especially relevant in long neuronal processes where organelle distribution is critical. This is an inference from general cell biology plus neural expression/pathology data, not a fully established neuron-specific mechanism. ([pubmed.ncbi.nlm.nih.gov](https://pubmed.ncbi.nlm.nih.gov/39133213/)) [S14]

The pathology studies are consistent with that model: when axonal or neuritic ER becomes abnormal in neurodegenerative settings, REEP5-positive tubular ER structures accumulate in dystrophic neurites or damaged dopaminergic axons. That does not prove REEP5 is the driver, but it strengthens the association between REEP5-marked ER organization and neural structural homeostasis. ([pubmed.ncbi.nlm.nih.gov](https://pubmed.ncbi.nlm.nih.gov/26619807/)) [S15]

## Bottom line

The claim is **likely true**, but mainly in a **qualified** sense. REEP5 is clearly associated with neural tissue, ER membrane organization, and organelle-level homeostatic processes such as intracellular transport, ER architecture, ER-mitochondria coupling, and redox regulation. Neural pathology studies also place REEP5 in abnormal neurites and axons. ([ncbi.nlm.nih.gov](https://www.ncbi.nlm.nih.gov/gene/7905)) [S16]

However, the field has **not yet produced strong neuron-specific causal evidence** showing that REEP5 is a major established regulator of neural homeostasis in the same way some other ER-shaping proteins are. So the safest interpretation is: **yes, REEP5 has been associated with regulatory/homeostatic biology in neural systems, but much of that evidence is indirect, mechanistic, or pathology-linked rather than definitive functional proof in neurons.** ([mdpi.com](https://www.mdpi.com/2073-4409/10/5/970)) [S17]

## Cited sources

- **[S1]** REEP5 receptor accessory protein 5 [Homo sapiens (human)] - Gene - NCBI — https://www.ncbi.nlm.nih.gov/gene/7905
- **[S2]** REEP5 protein expression summary - The Human Protein Atlas — https://www.proteinatlas.org/ENSG00000129625-REEP5
- **[S3]** REEP1 and REEP2 proteins are preferentially expressed in neuronal and neuronal-like exocytotic tissues - PubMed — https://pubmed.ncbi.nlm.nih.gov/24355597/
- **[S4]** Morphological Heterogeneity of the Endoplasmic Reticulum within Neurons and Its Implications in Neurodegeneration | MDPI — https://www.mdpi.com/2073-4409/10/5/970
- **[S5]** Dysfunctional tubular endoplasmic reticulum constitutes a pathological feature of Alzheimer's disease - PubMed — https://pubmed.ncbi.nlm.nih.gov/26619807/
- **[S6]** Pathological characteristics of axons and alterations of proteomic and lipidomic profiles in midbrain dopaminergic neurodegeneration induced by WDR45-deficiency - PubMed — https://pubmed.ncbi.nlm.nih.gov/39183331/
- **[S7]** Dynamic interaction of REEP5-MFN1/2 enables mitochondrial hitchhiking on tubular ER - PubMed — https://pubmed.ncbi.nlm.nih.gov/39133213/
- **[S8]** Towards understanding the role of Receptor Expression Enhancing Protein 5 (REEP5) in cardiac muscle and beyond - PubMed — https://pubmed.ncbi.nlm.nih.gov/32548572/
- **[S9]** Dynamic interaction of REEP5-MFN1/2 enables mitochondrial hitchhiking on tubular ER - PubMed — https://pubmed.ncbi.nlm.nih.gov/39133213/
- **[S10]** REEP1 and REEP2 proteins are preferentially expressed in neuronal and neuronal-like exocytotic tissues - PubMed — https://pubmed.ncbi.nlm.nih.gov/24355597/
- **[S11]** The REEP family of proteins: Molecular targets and role in pathophysiology - PubMed — https://pubmed.ncbi.nlm.nih.gov/36191880/
- **[S12]** Morphological Heterogeneity of the Endoplasmic Reticulum within Neurons and Its Implications in Neurodegeneration | MDPI — https://www.mdpi.com/2073-4409/10/5/970
- **[S13]** REEP5 receptor accessory protein 5 [Homo sapiens (human)] - Gene - NCBI — https://www.ncbi.nlm.nih.gov/gene/7905
- **[S14]** Dynamic interaction of REEP5-MFN1/2 enables mitochondrial hitchhiking on tubular ER - PubMed — https://pubmed.ncbi.nlm.nih.gov/39133213/
- **[S15]** Dysfunctional tubular endoplasmic reticulum constitutes a pathological feature of Alzheimer's disease - PubMed — https://pubmed.ncbi.nlm.nih.gov/26619807/
- **[S16]** REEP5 receptor accessory protein 5 [Homo sapiens (human)] - Gene - NCBI — https://www.ncbi.nlm.nih.gov/gene/7905
- **[S17]** Morphological Heterogeneity of the Endoplasmic Reticulum within Neurons and Its Implications in Neurodegeneration | MDPI — https://www.mdpi.com/2073-4409/10/5/970
