## Supplementary material for "Understanding Substance Dependence: What Differentiates Addictive from Non-Addictive Drugs?": Folder for supplementary files: non-addictive-genes-claims-first100-081-diras2-gene-protein-has-been-associated-with-regul-ca42362afcab.md

# non-addictive-genes-claims-first100-081-ca42362afcab

**Original claim:** DIRAS2 gene/protein has been associated with regulatory or homeostatic biological processes in neural systems.

## Claim

DIRAS2 gene/protein has been associated with regulatory or homeostatic biological processes in neural systems.

## Verdict

Verdict: Likely True

## Confidence

Medium

## Supporting evidence

DIRAS2 is clearly a brain-enriched gene/protein, which is a prerequisite for the claim’s “in neural systems” part. In the original biochemical characterization of Di-Ras2, Kontani et al. reported that human Di-Ras2 is specifically expressed in heart and brain and has unusual small-GTPase properties, including low intrinsic GTPase activity and failure to activate canonical Raf/MAPK signaling like classical Ras proteins. They also proposed it may regulate cell morphogenesis in a distinct way. ([pubmed.ncbi.nlm.nih.gov](https://pubmed.ncbi.nlm.nih.gov/12194967/)) [S1]

More direct neural evidence comes from a dedicated brain-expression study. Grünewald et al. found that DIRAS2 expression in human brain was highest in hippocampus and cerebral cortex, increased strongly during mouse brain development, and was detected in glutamatergic and catecholaminergic neurons. That places DIRAS2 in neuronal populations involved in synaptic signaling and circuit regulation rather than in generic non-neural tissue. ([pubmed.ncbi.nlm.nih.gov](https://pubmed.ncbi.nlm.nih.gov/29488099/)) [S2]

There is also experimental evidence that DIRAS2 influences gene-regulatory programs relevant to neural development. In murine hippocampal primary cells, shRNA knockdown of Diras2 altered expression of 1,612 genes, with enrichment for biological processes involved in neurodevelopment; 33 selected expression changes were validated by qPCR. The authors concluded that Diras2 affects numerous genes and molecular pathways relevant to neurodevelopmental processes. That is a direct association with regulatory biological processes in neural cells, even if it does not fully define the causal pathway. ([pubmed.ncbi.nlm.nih.gov](https://pubmed.ncbi.nlm.nih.gov/30623719/)) [S3]

Large-scale synaptic proteomics also links Diras2 to synaptic specialization. In a study profiling 18 synapse types across brain regions and cell types, Taft et al. identified Diras2 among proteins enriched with excitatory/VGlut1-associated synaptic communities, noting that Diras2 had not previously been annotated as synaptic. This does not prove function by itself, but it strongly associates DIRAS2 with molecular organization of excitatory synapses, which are core regulatory units of neural signaling. ([pmc.ncbi.nlm.nih.gov](https://pmc.ncbi.nlm.nih.gov/articles/PMC10686415/)) [S4]

Human genetic and neurophysiologic studies further support relevance to neural regulation, though more indirectly. Reif et al. reported association of DIRAS2 variants/haplotypes with adult ADHD and related traits, and discussed its brain-specific expression and possible role in serotonergic regulation. Separately, Grünewald et al. showed that an ADHD-associated promoter polymorphism increased reporter-gene expression and was associated with altered NoGo anteriorization, interpreted as an effect on prefrontal response control. These studies do not establish biochemical function, but they do connect DIRAS2 variation to neural control systems and cortical inhibitory processing. ([pmc.ncbi.nlm.nih.gov](https://pmc.ncbi.nlm.nih.gov/articles/PMC3176568/)) [S5]

Finally, an evolutionarily related line of evidence supports a synaptic regulatory role for the Di-Ras family. In *C. elegans*, the Di-Ras homolog drn-1 was expressed specifically in neurons and functioned with EPAC-1 to control acetylcholine release at neuromuscular junctions. Because this is a homolog study in an invertebrate, it should be treated as indirect support, but it is biologically consistent with a conserved role in regulating synaptic activity. ([tohoku.elsevierpure.com](https://tohoku.elsevierpure.com/en/publications/neuronally-expressed-ras-family-gtpase-di-ras-modulates-synaptic-)) [S6]

## Contradicting or limiting evidence

The main limitation is that DIRAS2’s precise neural function remains poorly defined. Even the adult ADHD association paper stated that “almost nothing is known” about DIRAS2 function and emphasized that relevant molecular cascades remain elusive. The promoter-polymorphism study likewise said that mechanisms linking DIRAS2 to ADHD require further clarification. So while association exists, mechanistic certainty is limited. ([pmc.ncbi.nlm.nih.gov](https://pmc.ncbi.nlm.nih.gov/articles/PMC3176568/)) [S7]

The strongest mammalian neural evidence is still mostly associative or perturbational rather than definitive. Expression in selected neuron types, altered transcriptomes after knockdown, and synaptic-proteome enrichment all suggest involvement in neural regulatory biology, but none alone proves that DIRAS2 is a core homeostatic regulator in vivo. Notably, I did not find a peer-reviewed mammalian study showing that selective DIRAS2 loss or gain directly disrupts neural homeostasis, synaptic scaling, firing-rate homeostasis, neurotransmitter homeostasis, or a similarly canonical homeostatic process. ([pubmed.ncbi.nlm.nih.gov](https://pubmed.ncbi.nlm.nih.gov/29488099/)) [S8]

The human psychiatric genetics evidence is also not decisive for biology on its own. ADHD associations can indicate relevance to neural regulation, but they do not prove a specific molecular role, and the original association paper itself noted caveats such as possible false positives and incomplete replication across samples. ([pmc.ncbi.nlm.nih.gov](https://pmc.ncbi.nlm.nih.gov/articles/PMC3176568/)) [S9]

Some mechanistic studies of DIRAS2 are outside neural systems, especially in cancer biology and autophagy. Those studies show DIRAS2 can regulate signaling and autophagy in non-neural contexts, but they do not directly establish neural homeostatic function. They are therefore better viewed as mechanistic clues than direct evidence for the present claim. ([pmc.ncbi.nlm.nih.gov](https://pmc.ncbi.nlm.nih.gov/articles/PMC5959337/?utm_source=openai)) [S10]

## Mechanistic / biological rationale

A plausible biological rationale exists. DIRAS2 encodes a small Ras-family GTPase, and small GTPases commonly regulate intracellular signaling, cell morphology, trafficking, and differentiation. DIRAS2 is unusual because it has low GTPase activity, appears predominantly GTP-bound, and does not signal like canonical Ras through Raf/MAPK. That unusual biochemistry suggests it may act as a specialized signaling modulator rather than a standard growth-pathway switch. ([pubmed.ncbi.nlm.nih.gov](https://pubmed.ncbi.nlm.nih.gov/12194967/)) [S11]

In brain, this is complemented by a specific regulatory interaction: endogenous Di-Ras2 from rat brain cytosol co-purified with SmgGDS, and SmgGDS reduced Di-Ras2’s affinity for guanine nucleotides while stabilizing the protein after synthesis. That is direct evidence that Di-Ras2 itself is subject to biochemical regulation in neural tissue, and it provides a concrete mechanism by which its signaling state could be controlled in neurons. ([pubmed.ncbi.nlm.nih.gov](https://pubmed.ncbi.nlm.nih.gov/26149690/)) [S12]

Its spatial pattern also fits a regulatory neural role. High expression in hippocampus and cortex, developmental upregulation, and localization to glutamatergic/catecholaminergic neurons are all consistent with involvement in circuit maturation, synaptic signaling, or activity-dependent neuronal specialization. The synaptic-proteome data placing Diras2 in excitatory synaptic protein communities further support this interpretation. ([pubmed.ncbi.nlm.nih.gov](https://pubmed.ncbi.nlm.nih.gov/29488099/)) [S13]

Still, the “homeostatic” part of the claim is less secure than the “regulatory” part. I found evidence that DIRAS2 is associated with neural regulatory processes such as neurodevelopment, neuronal signaling, synaptic organization, and prefrontal inhibitory control, but not strong direct evidence that it is a defined homeostatic factor in the same sense as genes controlling synaptic scaling, calcium homeostasis, or neurotransmitter buffering. The best homeostasis-adjacent support is indirect: synaptic-proteome localization, neuronal expression, and homolog data showing modulation of transmitter release. ([pubmed.ncbi.nlm.nih.gov](https://pubmed.ncbi.nlm.nih.gov/30623719/)) [S14]

## Bottom line

The claim is **likely true**, mainly because DIRAS2 has been linked by multiple primary studies to **neuronal expression, neurodevelopmental gene regulation, excitatory synaptic molecular organization, and neural control phenotypes**. ([pubmed.ncbi.nlm.nih.gov](https://pubmed.ncbi.nlm.nih.gov/29488099/)) [S15]

However, the evidence supports **association with neural regulatory processes** much more strongly than it supports a **specific, established role in neural homeostasis**. In other words: DIRAS2 is reasonably associated with regulatory biology in neural systems, but its exact mechanistic role remains incompletely characterized, and direct proof of a canonical homeostatic function is still sparse. ([pmc.ncbi.nlm.nih.gov](https://pmc.ncbi.nlm.nih.gov/articles/PMC3176568/)) [S16]

## Cited sources

- **[S1]** Di-Ras, a distinct subgroup of ras family GTPases with unique biochemical properties - PubMed — https://pubmed.ncbi.nlm.nih.gov/12194967/
- **[S2]** Expression of the ADHD candidate gene Diras2 in the brain - PubMed — https://pubmed.ncbi.nlm.nih.gov/29488099/
- **[S3]** Knockdown of the ADHD Candidate Gene Diras2 in Murine Hippocampal Primary Cells - PubMed — https://pubmed.ncbi.nlm.nih.gov/30623719/
- **[S4]** 
            The proteomic landscape of synaptic diversity across brain regions and cell types - PMC
         — https://pmc.ncbi.nlm.nih.gov/articles/PMC10686415/
- **[S5]** 
            DIRAS2 is Associated with Adult ADHD, Related Traits, and Co-Morbid Disorders - PMC
         — https://pmc.ncbi.nlm.nih.gov/articles/PMC3176568/
- **[S6]** 
        Neuronally expressed Ras-family GTPase Di-Ras modulates synaptic activity in Caenorhabditis elegans
      -  Tohoku University — https://tohoku.elsevierpure.com/en/publications/neuronally-expressed-ras-family-gtpase-di-ras-modulates-synaptic-
- **[S7]** 
            DIRAS2 is Associated with Adult ADHD, Related Traits, and Co-Morbid Disorders - PMC
         — https://pmc.ncbi.nlm.nih.gov/articles/PMC3176568/
- **[S8]** Expression of the ADHD candidate gene Diras2 in the brain - PubMed — https://pubmed.ncbi.nlm.nih.gov/29488099/
- **[S9]** 
            DIRAS2 is Associated with Adult ADHD, Related Traits, and Co-Morbid Disorders - PMC
         — https://pmc.ncbi.nlm.nih.gov/articles/PMC3176568/
- **[S10]** RAS-related GTPases DIRAS1 and DIRAS2 induce autophagic cancer cell death and are required for autophagy in murine ovarian cancer cells - PMC — https://pmc.ncbi.nlm.nih.gov/articles/PMC5959337/?utm_source=openai
- **[S11]** Di-Ras, a distinct subgroup of ras family GTPases with unique biochemical properties - PubMed — https://pubmed.ncbi.nlm.nih.gov/12194967/
- **[S12]** Di-Ras2 Protein Forms a Complex with SmgGDS Protein in Brain Cytosol in Order to Be in a Low Affinity State for Guanine Nucleotides - PubMed — https://pubmed.ncbi.nlm.nih.gov/26149690/
- **[S13]** Expression of the ADHD candidate gene Diras2 in the brain - PubMed — https://pubmed.ncbi.nlm.nih.gov/29488099/
- **[S14]** Knockdown of the ADHD Candidate Gene Diras2 in Murine Hippocampal Primary Cells - PubMed — https://pubmed.ncbi.nlm.nih.gov/30623719/
- **[S15]** Expression of the ADHD candidate gene Diras2 in the brain - PubMed — https://pubmed.ncbi.nlm.nih.gov/29488099/
- **[S16]** 
            DIRAS2 is Associated with Adult ADHD, Related Traits, and Co-Morbid Disorders - PMC
         — https://pmc.ncbi.nlm.nih.gov/articles/PMC3176568/
