## Supplementary material for "Understanding Substance Dependence: What Differentiates Addictive from Non-Addictive Drugs?": Folder for supplementary files: non-addictive-genes-claims-first100-082-clk1-gene-protein-has-been-associated-with-regulat-d95de27a799b.md

# non-addictive-genes-claims-first100-082-d95de27a799b

**Original claim:** CLK1 gene/protein has been associated with regulatory or homeostatic biological processes in neural systems.

## Claim

**Claim assessed:** “CLK1 gene/protein has been associated with regulatory or homeostatic biological processes in neural systems.”

For clarity, I interpret **CLK1** here as the human **CDC-like kinase 1 (HGNC: CLK1)**, a nuclear dual-specificity kinase involved in splicing regulation. This is distinct from the model-organism **clk-1/COQ7** mitochondrial gene, which uses similar nomenclature but is a different protein. ([ncbi.nlm.nih.gov](https://www.ncbi.nlm.nih.gov/gene/1195)) [S1]

## Verdict

Verdict: Likely True

The claim is **likely true**, with an important nuance: the strongest evidence links **CDC-like kinase 1 (CLK1)** to **regulatory RNA-processing/splicing functions** that are relevant to neural biology, including **neuronal differentiation** and regulation of **tau/MAPT splicing** in the mammalian nervous system. Evidence for **homeostatic** roles exists as well, but it is strongest at the level of **general cellular splicing homeostasis under stress**, not as a well-established, dedicated **neural-system homeostasis** factor in vivo. ([ncbi.nlm.nih.gov](https://www.ncbi.nlm.nih.gov/gene/1195)) [S2]

## Confidence

**Medium.**

Confidence is not high because the literature clearly supports **regulatory** roles and neural relevance, but more direct evidence for **physiological neural homeostasis** in intact brains or neural circuits is limited. Much of the evidence comes from **cell lines, splicing assays, and disease-oriented inference**, rather than definitive in vivo neural homeostasis studies. ([pmc.ncbi.nlm.nih.gov](https://pmc.ncbi.nlm.nih.gov/articles/PMC359226/)) [S3]

## Supporting evidence

CLK1 is a well-established **splicing-factor kinase**. NCBI summarizes that human CLK1 encodes a dual-specificity kinase that, in the nucleus, phosphorylates **serine/arginine-rich (SR) proteins** involved in pre-mRNA processing and may indirectly govern splice-site selection. That alone places CLK1 squarely in a **regulatory** biological role. ([ncbi.nlm.nih.gov](https://www.ncbi.nlm.nih.gov/gene/1195)) [S4]

CLK1 is also present in brain tissue. The Human Protein Atlas reports CLK1 RNA as **detected in all** analyzed human brain regions and places it in a brain-expression cluster associated mainly with **oligodendrocytes**. However, it also classifies CLK1 as having **low tissue specificity**, indicating it is not brain-specific. This supports neural association, while also showing it is part of broadly used cellular regulatory machinery. ([proteinatlas.org](https://www.proteinatlas.org/ENSG00000013441-CLK1/brain)) [S5]

A direct neural-system link comes from a primary study in **PC12 neuron-like cells**: inducible expression of CLK caused **morphological differentiation** and **neurite elaboration**, mimicking several nerve growth factor–dependent events. That is direct evidence that CLK family activity can regulate neuron-like differentiation programs. ([pmc.ncbi.nlm.nih.gov](https://pmc.ncbi.nlm.nih.gov/articles/PMC359226/)) [S6]

A second strong neural link is **tau/MAPT splicing**. In a Mol Cell Neurosci study, Hartmann et al. reported that tau transcript splicing is regulated in the **mammalian nervous system**, and that **exon 10 usage is regulated by CLK1, CLK2, CLK3, and CLK4** through phosphorylation of SR proteins. Because MAPT exon 10 choice is a canonical neuron-relevant splicing event with implications for tauopathy, this is strong evidence that CLK1 is associated with a **regulatory process in neural systems**. ([pubmed.ncbi.nlm.nih.gov](https://pubmed.ncbi.nlm.nih.gov/11461155/)) [S7]

Secondary sources are consistent with this. A review on CLKs notes that CLKs regulate **TAU** splicing and that tau is important for microtubule stabilization and thereby affects processes including **neuronal activity**. Another review focused on Alzheimer’s disease describes CLK1 as important in RNA-splicing regulation through phosphorylation of SR proteins and discusses its proposed relevance to neurodegenerative pathophysiology. ([pmc.ncbi.nlm.nih.gov](https://pmc.ncbi.nlm.nih.gov/articles/PMC7593917/)) [S8]

There is also bona fide **homeostatic** evidence for CLK1, although it is not neural-specific. In a J Cell Biol study, stress promoted maturation of intron-retaining **Clk1/4** RNAs, and the resulting CLK proteins rephosphorylated SR proteins after stress. The authors explicitly proposed a **“ReSCUE” model for homeostatic control of SR protein phosphorylation**. This supports the claim’s “homeostatic” language, even if the demonstrated system was general cellular stress/splicing homeostasis rather than a defined neural homeostasis pathway. ([pubmed.ncbi.nlm.nih.gov](https://pubmed.ncbi.nlm.nih.gov/21949414/)) [S9]

Finally, CLK1 appears to be part of self-regulatory and broader signaling networks: NCBI and review sources describe **autoregulation of CLK1 by alternative splicing**, interaction with **SRPK1**, and effects on spliceosome assembly. These features fit a role in maintaining regulated RNA-processing states rather than being a passive housekeeping enzyme. ([ncbi.nlm.nih.gov](https://www.ncbi.nlm.nih.gov/gene/1195)) [S10]

## Contradicting or limiting evidence

The main limitation is **specificity**. CLK1 is **ubiquitously expressed** and has **low tissue specificity**, so the evidence does **not** support calling it a neural-specific regulator. It is better described as a broadly expressed splicing kinase with neural relevance. ([ncbi.nlm.nih.gov](https://www.ncbi.nlm.nih.gov/gene/1195)) [S11]

A second limitation is that the strongest “homeostatic” evidence is for **general SR-protein phosphorylation homeostasis under stress**, not for a clearly defined role in **neural homeostasis** such as synaptic scaling, excitability homeostasis, neurotransmitter balance, or glial homeostatic programs. The literature here is indirect. ([pubmed.ncbi.nlm.nih.gov](https://pubmed.ncbi.nlm.nih.gov/21949414/)) [S12]

A third limitation is that some neural evidence is **model-dependent**. The PC12 differentiation result is informative, but PC12 is a pheochromocytoma-derived model, not an intact nervous system. Likewise, much of the tau evidence comes from splicing assays, cotransfection systems, and disease-oriented mechanistic work rather than direct physiology in healthy neural circuits. ([pmc.ncbi.nlm.nih.gov](https://pmc.ncbi.nlm.nih.gov/articles/PMC359226/)) [S13]

Even recent developmental reviews remain cautious: one review states that although evidence suggests CLKs are key splicing regulators associated with **neural and neuroectodermal differentiation phenotypes**, the **developmental functions of CLKs are mostly unknown**. That is a meaningful caveat against overstating the claim. ([pmc.ncbi.nlm.nih.gov](https://pmc.ncbi.nlm.nih.gov/articles/PMC10952393/)) [S14]

## Mechanistic / biological rationale

Mechanistically, the claim is biologically plausible. CLK1 phosphorylates **SR splicing factors**, altering their activity, localization, and splice-site selection behavior. NCBI describes CLK1 as a regulator of pre-mRNA processing, and review data indicate that CLK1 forms a functional system with **SRPK1** to mobilize hyperphosphorylated SR proteins for efficient spliceosome assembly. ([ncbi.nlm.nih.gov](https://www.ncbi.nlm.nih.gov/gene/1195)) [S15]

That matters especially in neural systems because neurons rely heavily on **alternative splicing** to generate cell-type-specific and developmentally regulated isoforms. The tau/MAPT example is especially relevant: tau exon 10 is a regulated event in the mammalian nervous system, and CLK-family kinases influence that exon’s inclusion through phosphorylation-dependent control of splicing factors. Since tau isoform balance affects neuronal microtubule biology, this provides a mechanistic bridge from CLK1 activity to neural regulation. ([pubmed.ncbi.nlm.nih.gov](https://pubmed.ncbi.nlm.nih.gov/11461155/)) [S16]

The homeostatic angle comes from CLK1’s role in restoring splicing-factor phosphorylation after stress. In the ReSCUE model, stress causes SR-protein dephosphorylation; stress-responsive maturation of stored intron-retaining **Clk1/4** transcripts then produces kinase that rephosphorylates SR proteins, helping restore the system. That is a genuine homeostatic feedback concept, albeit shown in a general cell-biological context. ([pubmed.ncbi.nlm.nih.gov](https://pubmed.ncbi.nlm.nih.gov/21949414/)) [S17]

## Bottom line

The statement is **reasonable but should be phrased carefully**. For **human CDC-like kinase 1**, the literature supports that CLK1 is associated with **regulatory biological processes relevant to neural systems**, especially **RNA splicing**, **neuronal differentiation-like responses**, and **neuron-relevant tau/MAPT isoform regulation**. Evidence for **homeostatic** function also exists, but it is best described as **cellular splicing homeostasis under stress**, not yet as a strongly established **neural homeostasis regulator** in vivo. So the overall claim is best judged **Likely True**, with the caveat that the **regulatory** part is better supported than the specifically **neural homeostatic** part. ([ncbi.nlm.nih.gov](https://www.ncbi.nlm.nih.gov/gene/1195)) [S18]

## Cited sources

- **[S1]** CLK1 CDC like kinase 1 [Homo sapiens (human)] - Gene - NCBI — https://www.ncbi.nlm.nih.gov/gene/1195
- **[S2]** CLK1 CDC like kinase 1 [Homo sapiens (human)] - Gene - NCBI — https://www.ncbi.nlm.nih.gov/gene/1195
- **[S3]** 
            The dual-specificity CLK kinase induces neuronal differentiation of PC12 cells - PMC
         — https://pmc.ncbi.nlm.nih.gov/articles/PMC359226/
- **[S4]** CLK1 CDC like kinase 1 [Homo sapiens (human)] - Gene - NCBI — https://www.ncbi.nlm.nih.gov/gene/1195
- **[S5]** Brain tissue expression of CLK1 - Summary - The Human Protein Atlas — https://www.proteinatlas.org/ENSG00000013441-CLK1/brain
- **[S6]** 
            The dual-specificity CLK kinase induces neuronal differentiation of PC12 cells - PMC
         — https://pmc.ncbi.nlm.nih.gov/articles/PMC359226/
- **[S7]** Regulation of alternative splicing of human tau exon 10 by phosphorylation of splicing factors - PubMed — https://pubmed.ncbi.nlm.nih.gov/11461155/
- **[S8]** 
            Cdc-Like Kinases (CLKs): Biology, Chemical Probes, and Therapeutic Potential - PMC
         — https://pmc.ncbi.nlm.nih.gov/articles/PMC7593917/
- **[S9]** Stress-responsive maturation of Clk1/4 pre-mRNAs promotes phosphorylation of SR splicing factor - PubMed — https://pubmed.ncbi.nlm.nih.gov/21949414/
- **[S10]** CLK1 CDC like kinase 1 [Homo sapiens (human)] - Gene - NCBI — https://www.ncbi.nlm.nih.gov/gene/1195
- **[S11]** CLK1 CDC like kinase 1 [Homo sapiens (human)] - Gene - NCBI — https://www.ncbi.nlm.nih.gov/gene/1195
- **[S12]** Stress-responsive maturation of Clk1/4 pre-mRNAs promotes phosphorylation of SR splicing factor - PubMed — https://pubmed.ncbi.nlm.nih.gov/21949414/
- **[S13]** 
            The dual-specificity CLK kinase induces neuronal differentiation of PC12 cells - PMC
         — https://pmc.ncbi.nlm.nih.gov/articles/PMC359226/
- **[S14]** 
            Functions of SRPK, CLK and DYRK kinases in stem cells, development, and human developmental disorders - PMC
         — https://pmc.ncbi.nlm.nih.gov/articles/PMC10952393/
- **[S15]** CLK1 CDC like kinase 1 [Homo sapiens (human)] - Gene - NCBI — https://www.ncbi.nlm.nih.gov/gene/1195
- **[S16]** Regulation of alternative splicing of human tau exon 10 by phosphorylation of splicing factors - PubMed — https://pubmed.ncbi.nlm.nih.gov/11461155/
- **[S17]** Stress-responsive maturation of Clk1/4 pre-mRNAs promotes phosphorylation of SR splicing factor - PubMed — https://pubmed.ncbi.nlm.nih.gov/21949414/
- **[S18]** CLK1 CDC like kinase 1 [Homo sapiens (human)] - Gene - NCBI — https://www.ncbi.nlm.nih.gov/gene/1195
