## Supplementary material for "Understanding Substance Dependence: What Differentiates Addictive from Non-Addictive Drugs?": Folder for supplementary files: non-addictive-genes-claims-first100-083-pdp1-gene-protein-has-been-associated-with-regulat-70b6c800c011.md

# non-addictive-genes-claims-first100-083-70b6c800c011

**Original claim:** PDP1 gene/protein has been associated with regulatory or homeostatic biological processes in neural systems.

## Claim

**“PDP1 gene/protein has been associated with regulatory or homeostatic biological processes in neural systems.”**

## Verdict

Verdict: Likely True

## Confidence

**Medium**

## Supporting evidence

A key caveat is that **“PDP1” is nomenclaturally ambiguous across species**. In humans, **PDP1** is the mitochondrial **pyruvate dehydrogenase phosphatase catalytic subunit 1**, which activates the pyruvate dehydrogenase complex (PDC) by dephosphorylation. In *Drosophila*, **Pdp1** is a **PAR-domain bZIP transcription factor**, a different protein class with a well-established role in circadian clock neurons. Because the claim is broad and says “gene/protein” rather than specifying species, both literatures are relevant. ([medlineplus.gov](https://medlineplus.gov/genetics/gene/pdp1)) [S1]

For **human/mammalian PDP1**, there is real neural-system association, though it is mostly **metabolic-regulatory** rather than synaptic. Human Protein Atlas data classify PDP1 RNA as **“tissue enhanced (brain)”** and show human brain regional enrichment in **basal ganglia, cerebral cortex, and olfactory region**. A clinical review of PDP1 deficiency also notes that both PDP1 and PDP2 are highly expressed in brain, with PDP1 protein reported at about **7-fold higher than PDP2** there. ([v20.proteinatlas.org](https://v20.proteinatlas.org/ENSG00000164951-PDP1)) [S2]

There is also direct animal evidence that PDP1 participates in **brain metabolic regulation**. In a rat traumatic brain injury study, PDP1 mRNA was reported as **highly expressed in cerebral cortex, hippocampus, and thalamus**, and its expression changed dynamically after injury. The authors linked those changes to PDH regulation, pyruvate/lactate handling, and neuronal vulnerability, which is consistent with a role in neural metabolic homeostasis. ([pubmed.ncbi.nlm.nih.gov](https://pubmed.ncbi.nlm.nih.gov/22884618/)) [S3]

Related brain-cell work supports the same direction. A review on PDK/PDP regulation in the nervous system states that the **PDK/PDP system regulates PDC activity**, contributes to the distinct metabolic phenotypes of **neurons and astrocytes**, and influences **neurometabolic coupling** and monocarboxylate shuttling between those cell types. The same review specifically states that **brain PDK2 and PDP1** may be important for controlling PDH phosphorylation/dephosphorylation under normal conditions and after traumatic brain injury. A primary study in cultured rat brain cells likewise found that pyruvate metabolism is differentially regulated in neurons versus astrocytes through PDC phosphorylation state. ([pmc.ncbi.nlm.nih.gov](https://pmc.ncbi.nlm.nih.gov/articles/PMC3520047/)) [S4]

Human genetics also supports neural relevance, albeit indirectly. MedlinePlus states that pathogenic **PDP1** variants can cause **pyruvate dehydrogenase deficiency** with **delayed development and neurological problems**, and emphasizes that the **brain is especially dependent** on this energy-producing pathway. That does not by itself prove a specific homeostatic neural function, but it does strongly support biological importance in the nervous system. ([medlineplus.gov](https://medlineplus.gov/genetics/gene/pdp1)) [S5]

For ***Drosophila* Pdp1**, the evidence is much stronger and more directly “regulatory/homeostatic” in a neural sense. A landmark *Cell* paper showed that **VRI and PDP1 directly regulate dClock expression** and that the **clock stops in a Pdp1 null mutant**, identifying Pdp1 as an **essential clock gene** in the fly circadian system. Since circadian timing governs sleep/activity organization and temporal homeostasis in neural circuits, this is a direct association with neural regulatory biology. ([pubmed.ncbi.nlm.nih.gov](https://pubmed.ncbi.nlm.nih.gov/12581523/)) [S6]

Curated model-organism resources are consistent with that interpretation. FlyBase summarizes *Drosophila* **Pdp1** as a PAR-domain transcription factor that **“regulates gene expression in muscles and in circadian clock neurons.”** A review on *Drosophila* sleep/circadian biology likewise describes a **second feedback loop formed by PDP1 and Vrille** that drives **Clk mRNA cycling**. ([flybase.org](https://flybase.org/reports/FBgn0016694)) [S7]

## Contradicting or limiting evidence

The main limitation is that for **human PDP1**, the evidence is mostly about **mitochondrial fuel regulation**, not classic neural homeostasis constructs such as **synaptic scaling, ion homeostasis, excitability set-points, neurotransmitter release control, or circuit-level homeostatic plasticity**. I did not find strong evidence that human PDP1 is a canonical regulator of those neural processes specifically. The association is therefore **real but indirect**, centered on energy metabolism in neural tissue rather than on a dedicated neuronal homeostasis pathway. ([medlineplus.gov](https://medlineplus.gov/genetics/gene/pdp1)) [S8]

Even in flies, the exact mechanistic placement of Pdp1 within the circadian system has been refined over time. A later study found that reducing or overexpressing **PDP1ε** in clock cells **did not abolish Clk mRNA cycling or core oscillator function**, arguing that PDP1ε may be **more important for clock output/behavioral rhythmicity** than for the core oscillator itself. So the broad claim that Pdp1 is tied to neural regulation is still supported, but the precise mechanism is more nuanced than the earliest model suggested. ([pubmed.ncbi.nlm.nih.gov](https://pubmed.ncbi.nlm.nih.gov/17344391/)) [S9]

A second limitation is cross-species interpretation: the **human PDP1 phosphatase** and the ***Drosophila* Pdp1 transcription factor** are not homologous in function in the simple sense of “same protein doing the same neural job.” The strongest “neural regulatory” evidence comes from the fly clock-gene literature, whereas the strongest human evidence is metabolic and disease-associated. ([medlineplus.gov](https://medlineplus.gov/genetics/gene/pdp1)) [S10]

## Mechanistic / biological rationale

For **human/mammalian PDP1**, the rationale is straightforward: PDP1 **dephosphorylates and activates PDH/PDC**, increasing conversion of **pyruvate to acetyl-CoA** and thereby supporting mitochondrial ATP production. Because neural tissue is highly energy dependent, and because neuron–astrocyte interactions rely on tightly regulated carbohydrate oxidation and lactate/pyruvate handling, a PDP1-dependent control point is biologically plausible as part of **neural metabolic homeostasis**. Injury-induced changes in brain PDP1 expression further support that interpretation. ([medlineplus.gov](https://medlineplus.gov/genetics/gene/pdp1)) [S11]

For ***Drosophila* Pdp1**, the mechanism is transcriptional. Pdp1 is a **PAR-domain bZIP transcription factor** expressed in **circadian clock neurons**, where it participates in the transcriptional feedback architecture that shapes circadian timing and behavioral rhythmicity. Because circadian systems are fundamental regulators of sleep-wake organization, daily physiology, and stability of neural output over time, this is a clear example of a **regulatory/homeostatic process in a neural system**. ([pubmed.ncbi.nlm.nih.gov](https://pubmed.ncbi.nlm.nih.gov/12581523/)) [S12]

## Bottom line

The claim is **likely true**, but with an important qualification. If you mean ***Drosophila* Pdp1**, the evidence is **strong** that it participates in **neural regulatory processes**, especially circadian clock function and behavioral rhythmicity. If you mean **human PDP1**, the evidence is still supportive, but mostly through **brain-enriched expression, neural metabolic regulation, neurometabolic coupling, injury response, and neurological disease phenotypes**, rather than through direct evidence for classic synaptic or circuit homeostatic control. Overall, PDP1 has been associated with **regulatory/homeostatic biology in neural systems**, but the strength and exact meaning of that association depend heavily on **which PDP1** you mean. ([pubmed.ncbi.nlm.nih.gov](https://pubmed.ncbi.nlm.nih.gov/12581523/)) [S13]

## Cited sources

- **[S1]** PDP1 gene: MedlinePlus Genetics — https://medlineplus.gov/genetics/gene/pdp1
- **[S2]** PDP1 protein expression summary - The Human Protein Atlas — https://v20.proteinatlas.org/ENSG00000164951-PDP1
- **[S3]** Pyruvate dehydrogenase phosphatase1 mRNA expression is divergently and dynamically regulated between rat cerebral cortex, hippocampus and thalamus after traumatic brain injury: a potential biomarker of TBI-induced hyper- and hypo-glycaemia and neuronal vulnerability - PubMed — https://pubmed.ncbi.nlm.nih.gov/22884618/
- **[S4]** 
            Pyruvate Dehydrogenase Kinases in the Nervous System: Their Principal Functions in Neuronal-glial Metabolic Interaction and Neuro-metabolic Disorders - PMC
         — https://pmc.ncbi.nlm.nih.gov/articles/PMC3520047/
- **[S5]** PDP1 gene: MedlinePlus Genetics — https://medlineplus.gov/genetics/gene/pdp1
- **[S6]** vrille, Pdp1, and dClock form a second feedback loop in the Drosophila circadian clock - PubMed — https://pubmed.ncbi.nlm.nih.gov/12581523/
- **[S7]**         
    FlyBase Gene Report: Dmel\Pdp1
 — https://flybase.org/reports/FBgn0016694
- **[S8]** PDP1 gene: MedlinePlus Genetics — https://medlineplus.gov/genetics/gene/pdp1
- **[S9]** PDP1epsilon functions downstream of the circadian oscillator to mediate behavioral rhythms - PubMed — https://pubmed.ncbi.nlm.nih.gov/17344391/
- **[S10]** PDP1 gene: MedlinePlus Genetics — https://medlineplus.gov/genetics/gene/pdp1
- **[S11]** PDP1 gene: MedlinePlus Genetics — https://medlineplus.gov/genetics/gene/pdp1
- **[S12]** vrille, Pdp1, and dClock form a second feedback loop in the Drosophila circadian clock - PubMed — https://pubmed.ncbi.nlm.nih.gov/12581523/
- **[S13]** vrille, Pdp1, and dClock form a second feedback loop in the Drosophila circadian clock - PubMed — https://pubmed.ncbi.nlm.nih.gov/12581523/
