## Supplementary material for "Understanding Substance Dependence: What Differentiates Addictive from Non-Addictive Drugs?": Folder for supplementary files: non-addictive-genes-claims-first100-084-dclk1-gene-protein-has-been-associated-with-regula-45fb59760238.md

# non-addictive-genes-claims-first100-084-45fb59760238

**Original claim:** DCLK1 gene/protein has been associated with regulatory or homeostatic biological processes in neural systems.

## Claim

DCLK1 gene/protein has been associated with regulatory or homeostatic biological processes in neural systems.

## Verdict

Verdict: Likely True

## Confidence

Medium

## Supporting evidence

DCLK1 is a neuron-associated microtubule-binding serine/threonine kinase whose authoritative NCBI gene summary lists roles in neuronal migration, retrograde transport, neuronal apoptosis, and neurogenesis. Its domain architecture is well suited to regulatory functions in neurons: two N-terminal doublecortin domains bind microtubules, while the C-terminal kinase domain can modulate downstream signaling. ([ncbi.nlm.nih.gov](https://www.ncbi.nlm.nih.gov/gene/9201)) [S1]

The strongest evidence is from developmental neurobiology. In cortical progenitors, DCLK was shown to regulate bipolar mitotic spindle formation and M-phase progression through a dynein-dependent mechanism; knockdown disrupted spindle structure, altered cell-cycle progression, and changed neural progenitor fate during corticogenesis. This is direct evidence that DCLK1-family products participate in regulatory control of neurogenesis rather than merely serving as passive markers. ([pubmed.ncbi.nlm.nih.gov](https://pubmed.ncbi.nlm.nih.gov/16387637/)) [S2]

Related work on the DCLK splice variant DCL found expression in mitotic cells and radial glia during early corticogenesis. Knockdown induced spindle collapse, reduced proliferative-zone cell numbers, and disrupted radial glial processes, supporting a role for the DCLK1 locus in regulating precursor division and radial scaffold stability in the developing brain. ([pubmed.ncbi.nlm.nih.gov](https://pubmed.ncbi.nlm.nih.gov/17313568/)) [S3]

DCLK1 is also implicated in neuronal migration and axon formation. A cortical-neuron study identified MAP7D1 as a DCLK1 substrate and showed that DCLK1-dependent phosphorylation of MAP7D1 promotes callosal axon elongation. Other genetic studies found that Dclk and Dcx cooperate in murine hippocampal development, and DCLK1 has been linked to fiber tract decussation and neuronal migration, again supporting a regulatory role in neural circuit formation. ([pubmed.ncbi.nlm.nih.gov](https://pubmed.ncbi.nlm.nih.gov/27503845/)) [S4]

Evidence extends beyond embryonic development into postmitotic neurons. In cultured and in vivo neurons, DCLK proteins localize to dendritic and postsynaptic compartments, regulate dendritic remodeling, and influence synapse maturation. One study concluded that DCLK proteins promote distal dendritic growth while suppressing synapse maturation and PSD-95 accumulation, indicating active regulation of structural plasticity rather than a housekeeping role. ([pmc.ncbi.nlm.nih.gov](https://pmc.ncbi.nlm.nih.gov/articles/PMC4017031/)) [S5]

A later conditional mouse study likewise placed DCLK1 in postnatal circuit regulation, but with a different directional effect: deleting microtubule-binding DCLK1 isoforms in pyramidal neurons reduced spine density and mature mushroom spines in prefrontal cortex, supporting a role in spine growth and maturation. Although the direction of effect differs from the earlier study, both studies agree that DCLK1 regulates dendritic spine/synapse biology in neural tissue. ([sciencedirect.com](https://www.sciencedirect.com/science/article/abs/pii/S0306452222004390?utm_source=openai)) [S6]

Mechanistically linked transport data strengthen the case. DCLK1 remains expressed in mature neurons, labels dendritic microtubules, and is required for KIF1-dependent dense-core vesicle trafficking and dendrite development. More broadly, DCX/DCLK1-deficient neurons show impaired KIF1A-mediated transport of synaptic vesicle cargo, which is a plausible route by which DCLK1 could regulate neuronal connectivity and plasticity. ([pmc.ncbi.nlm.nih.gov](https://pmc.ncbi.nlm.nih.gov/articles/PMC4741305/)) [S7]

There is also evidence for relevance in the mature diseased brain. In mouse and human neuron models of synucleinopathy, DCLK1 knockdown reduced total and phosphorylated α-synuclein and lessened neurotoxicity, showing that DCLK1 can regulate protein handling and vulnerability in mature neurons. That is not definitive proof of normal physiological homeostasis, but it does support involvement in neural regulatory processes beyond development. ([pubmed.ncbi.nlm.nih.gov](https://pubmed.ncbi.nlm.nih.gov/31748376/)) [S8]

Human association data are supportive but weaker than the animal/mechanistic work. DCLK1 variants have been associated with verbal memory/general cognition and with schizophrenia/ADHD-related phenotypes, consistent with a role in neurodevelopment and synaptic plasticity. These studies support neural relevance, though they do not by themselves prove molecular function. ([pubmed.ncbi.nlm.nih.gov](https://pubmed.ncbi.nlm.nih.gov/19844571/?utm_source=openai)) [S9]

## Contradicting or limiting evidence

The main limitation is that **homeostatic** function in the strict sense is not strongly established. Most direct evidence concerns neurogenesis, migration, axon/dendrite growth, and synapse/spine remodeling. Those are clearly regulatory neural processes, but they are not the same as classic neural homeostasis such as firing-rate homeostasis, neurotransmitter balance, or glial metabolic homeostasis. Evidence specifically using that framework is sparse. ([pubmed.ncbi.nlm.nih.gov](https://pubmed.ncbi.nlm.nih.gov/16387637/)) [S10]

Single-gene loss can also produce surprisingly mild gross phenotypes, implying redundancy. Targeted deletion of Dclk showed no appreciable hippocampal developmental defect in one mouse study, and a later report noted normal gross brain architecture, cortical lamination, and progenitor migration in global DCLK1 knockout mice. Stronger phenotypes often emerge when DCLK1 is examined with DCX or in specific cell types, stages, or compartments. That means DCLK1 is involved, but not always singly essential. ([academic.oup.com](https://academic.oup.com/cercor/article-pdf/16/suppl_1/i69/17295647/bhk005.pdf?utm_source=openai)) [S11]

Another limitation is inconsistency in the direction of synaptic effects. The 2013 study argued DCLK1 suppresses synapse maturation and helps maintain dendritic structural plasticity, whereas the 2023 conditional-mouse study found DCLK1 facilitates spine growth and maturation in prefrontal pyramidal neurons. This likely reflects isoform-, developmental-stage-, cell-type-, or brain-region-specific biology, but it lowers confidence about the precise role at synapses. ([pmc.ncbi.nlm.nih.gov](https://pmc.ncbi.nlm.nih.gov/articles/PMC4017031/)) [S12]

Human evidence is also mostly associative. GWAS/cognitive studies and psychiatric associations are hypothesis-supporting, but they do not demonstrate that DCLK1 directly maintains neural homeostasis in humans. ([pubmed.ncbi.nlm.nih.gov](https://pubmed.ncbi.nlm.nih.gov/19844571/?utm_source=openai)) [S13]

## Mechanistic / biological rationale

A regulatory neural role for DCLK1 is biologically plausible. Its doublecortin domains bind microtubules and regulate polymerization, while its kinase domain can modify substrates and its linker/tail regions support protein interactions. An eLife study showed that DCLK1 autophosphorylation tunes its microtubule-binding affinity by preventing aberrant phosphorylation in the doublecortin domains. That kind of autoregulation is exactly what one would expect from a protein controlling cytoskeletal dynamics in neurons. ([pubmed.ncbi.nlm.nih.gov](https://pubmed.ncbi.nlm.nih.gov/34310279/)) [S14]

At the cellular level, DCLK1-family proteins influence mitotic spindle behavior in neural progenitors, KIF1/KIF1A-dependent cargo transport, axon elongation via MAP7D1 phosphorylation, and dendritic/spine architecture in postmitotic neurons. DCLK proteins are also found in postsynaptic compartments and can interact with synaptic scaffolding-related systems such as PSD proteins and NrCAM. Together, these functions provide a coherent mechanism by which DCLK1 could regulate neuronal morphogenesis, connectivity, and aspects of plasticity or stability. ([pubmed.ncbi.nlm.nih.gov](https://pubmed.ncbi.nlm.nih.gov/16387637/)) [S15]

## Bottom line

The claim is **likely true** if interpreted broadly: DCLK1 is well associated with **regulatory** biological processes in neural systems, especially neurogenesis, progenitor division, neuronal migration, axon/dendrite development, and synaptic/spine remodeling. ([ncbi.nlm.nih.gov](https://www.ncbi.nlm.nih.gov/gene/9201)) [S16]

What is **less well supported** is a specific claim that DCLK1 has an established role in **neural homeostasis** in the narrow physiological sense. The literature supports developmental and structural regulation much more strongly than canonical homeostatic control. So the safest conclusion is: **DCLK1 is clearly linked to neural regulatory processes; evidence for explicit neural homeostatic roles is presently indirect or limited.** ([pubmed.ncbi.nlm.nih.gov](https://pubmed.ncbi.nlm.nih.gov/31748376/)) [S17]

## Cited sources

- **[S1]** DCLK1 doublecortin like kinase 1 [Homo sapiens (human)] - Gene - NCBI — https://www.ncbi.nlm.nih.gov/gene/9201
- **[S2]** Doublecortin-like kinase controls neurogenesis by regulating mitotic spindles and M phase progression - PubMed — https://pubmed.ncbi.nlm.nih.gov/16387637/
- **[S3]** Doublecortin-like, a microtubule-associated protein expressed in radial glia, is crucial for neuronal precursor division and radial process stability - PubMed — https://pubmed.ncbi.nlm.nih.gov/17313568/
- **[S4]** DCLK1 phosphorylates the microtubule-associated protein MAP7D1 to promote axon elongation in cortical neurons - PubMed — https://pubmed.ncbi.nlm.nih.gov/27503845/
- **[S5]** 
            Doublecortin-like kinase enhances dendritic remodelling and negatively regulates synapse maturation - PMC
         — https://pmc.ncbi.nlm.nih.gov/articles/PMC4017031/
- **[S6]** Doublecortin-Like Kinase 1 Facilitates Dendritic Spine Growth of Pyramidal Neurons in Mouse Prefrontal Cortex - ScienceDirect — https://www.sciencedirect.com/science/article/abs/pii/S0306452222004390?utm_source=openai
- **[S7]** 
            Microtubule‐binding protein doublecortin‐like kinase 1 (DCLK1) guides kinesin‐3‐mediated cargo transport to dendrites - PMC
         — https://pmc.ncbi.nlm.nih.gov/articles/PMC4741305/
- **[S8]** Doublecortin-like Kinase 1 Regulates α-Synuclein Levels and Toxicity - PubMed — https://pubmed.ncbi.nlm.nih.gov/31748376/
- **[S9]** Variants in doublecortin- and calmodulin kinase like 1, a gene up-regulated by BDNF, are associated with memory and general cognitive abilities - PubMed — https://pubmed.ncbi.nlm.nih.gov/19844571/?utm_source=openai
- **[S10]** Doublecortin-like kinase controls neurogenesis by regulating mitotic spindles and M phase progression - PubMed — https://pubmed.ncbi.nlm.nih.gov/16387637/
- **[S11]** Doublecortin and Doublecortin-Like Kinase 1 Genes Cooperate in Murine Hippocampal Development | Cerebral Cortex | Oxford Academic — https://academic.oup.com/cercor/article-pdf/16/suppl_1/i69/17295647/bhk005.pdf?utm_source=openai
- **[S12]** 
            Doublecortin-like kinase enhances dendritic remodelling and negatively regulates synapse maturation - PMC
         — https://pmc.ncbi.nlm.nih.gov/articles/PMC4017031/
- **[S13]** Variants in doublecortin- and calmodulin kinase like 1, a gene up-regulated by BDNF, are associated with memory and general cognitive abilities - PubMed — https://pubmed.ncbi.nlm.nih.gov/19844571/?utm_source=openai
- **[S14]** Autoregulatory control of microtubule binding in doublecortin-like kinase 1 - PubMed — https://pubmed.ncbi.nlm.nih.gov/34310279/
- **[S15]** Doublecortin-like kinase controls neurogenesis by regulating mitotic spindles and M phase progression - PubMed — https://pubmed.ncbi.nlm.nih.gov/16387637/
- **[S16]** DCLK1 doublecortin like kinase 1 [Homo sapiens (human)] - Gene - NCBI — https://www.ncbi.nlm.nih.gov/gene/9201
- **[S17]** Doublecortin-like Kinase 1 Regulates α-Synuclein Levels and Toxicity - PubMed — https://pubmed.ncbi.nlm.nih.gov/31748376/
