## Supplementary material for "Understanding Substance Dependence: What Differentiates Addictive from Non-Addictive Drugs?": Folder for supplementary files: non-addictive-genes-claims-first100-085-rab11a-gene-protein-has-been-associated-with-regul-84ce77242dcf.md

# non-addictive-genes-claims-first100-085-84ce77242dcf

**Original claim:** RAB11A gene/protein has been associated with regulatory or homeostatic biological processes in neural systems.

## Claim

**Claim:** *RAB11A gene/protein has been associated with regulatory or homeostatic biological processes in neural systems.*

## Verdict

Verdict: Likely True

## Confidence

**High**

## Supporting evidence

RAB11A is well established as a recycling-endosome small GTPase, and the neural literature consistently links Rab11/RAB11A-dependent trafficking to processes that regulate receptor availability, synaptic plasticity, neurite morphology, and sensory-cell integrity. A 2020 review focused on the nervous system states that Rab11-mediated trafficking helps “maintain neurological function,” including transport of integrins, Trk receptors, and AMPA receptors to support axon/dendrite regeneration and synaptic plasticity. A separate review of neuronal AMPA-receptor trafficking describes Rab11-positive recycling endosomes as part of the controlled machinery that governs basal receptor recycling, long-term potentiation (LTP), and long-term depression (LTD), emphasizing that tight spatiotemporal regulation of Rab activity is critical for proper neuronal function. ([pubmed.ncbi.nlm.nih.gov](https://pubmed.ncbi.nlm.nih.gov/32329391/)) [S1]

There is also direct experimental evidence in neurons. In hippocampal neurons, BDNF increased GTP-bound Rab11 in dendrites, and Rab11/Myosin Vb signaling promoted dendritic branching by regulating recycling endosome dynamics and receptor accumulation in dendrites. In adult rat brain, suppression of Rab11 reduced dendritic arborization, impaired LTP induction, and worsened hippocampal-dependent spatial memory, indicating that Rab11-dependent recycling endosomes regulate key aspects of synaptic structure and function. ([pmc.ncbi.nlm.nih.gov](https://pmc.ncbi.nlm.nih.gov/articles/PMC3684039/)) [S2]

Synaptic receptor trafficking is one of the clearest “regulatory” neural roles. In mammalian neurons, LTP-associated AMPA-receptor insertion and spine growth require recycling endosome mobilization through a Myosin Vb/Rab11-FIP2 mechanism; disrupting this pathway abolishes LTP-related exocytosis from recycling endosomes and prevents AMPAR insertion and spine enlargement. Complementing this, a Neuron study showed that endocytic trafficking and recycling maintain a mobile surface pool of AMPA receptors required for synaptic potentiation. Together, these findings place Rab11-linked trafficking in the control loop that regulates excitatory synaptic strength. ([pubmed.ncbi.nlm.nih.gov](https://pubmed.ncbi.nlm.nih.gov/18984164/)) [S3]

Evidence specifically implicating **Rab11a** rather than only the Rab11 family is also available. In a neuronal model, Rab11a knockdown reduced neuronal differentiation markers and neurite-like process elongation; similar effects were seen in primary cortical neurons, while Rab11b knockdown did not significantly alter neuronal morphology in that study. This supports a specific role for Rab11a in neuronal morphological differentiation, a developmental regulatory process. ([pubmed.ncbi.nlm.nih.gov](https://pubmed.ncbi.nlm.nih.gov/38390945/)) [S4]

Rab11a has additionally been linked to regulation of neurotransmitter-receptor recycling in neuron-like cells. In PC12 cells expressing the CNS-relevant M4 muscarinic acetylcholine receptor, dominant-negative Rab11a impaired receptor recycling after agonist-induced internalization. The authors explicitly note that the balance between receptor internalization, recycling, and degradation determines neuronal responsiveness to ligand, making Rab11a part of a regulatory system that tunes signaling responsiveness. ([pmc.ncbi.nlm.nih.gov](https://pmc.ncbi.nlm.nih.gov/articles/PMC6757814/)) [S5]

Association with neural-system integrity extends beyond central synapses. In the mammalian organ of Corti, Rab11a was reported to be essential for development and integrity of stereocilia and kinocilia, linking trafficking to polarity-regulatory components and proper assembly of the mechanotransduction apparatus in hair cells. In peripheral glia, Rab11a knockdown in Schwann cells reduced tunneling nanotube formation, increased apoptosis, reduced migration, and secondarily decreased axonal outgrowth from co-cultured dorsal root ganglion neurons. These are not classical “homeostatic plasticity” studies, but they do support roles in maintaining ordered neural-cell architecture and intercellular support functions. ([pubmed.ncbi.nlm.nih.gov](https://pubmed.ncbi.nlm.nih.gov/37225424/)) [S6]

Human genetics also supports neural relevance. Recent clinical and experimental work reported de novo RAB11A mutation-associated neurodevelopmental disorder with intellectual disability and brain anomalies, and rab11a deficiency in zebrafish reduced CNS/forebrain/midbrain size. This does not by itself prove a homeostatic role, but it strengthens the conclusion that RAB11A is important for normal neural-system regulation and maintenance. ([pmc.ncbi.nlm.nih.gov](https://pmc.ncbi.nlm.nih.gov/articles/PMC12434753/)) [S7]

## Contradicting or limiting evidence

The main limitation is **specificity**. A substantial portion of the neural literature refers to “Rab11” or Rab11-positive recycling endosomes without cleanly separating **RAB11A** from **RAB11B**. That means some of the strongest mechanistic synaptic-plasticity data support the Rab11 pathway broadly, but do not always prove that RAB11A alone is the decisive isoform in every neural context. ([pubmed.ncbi.nlm.nih.gov](https://pubmed.ncbi.nlm.nih.gov/28628388/)) [S8]

A second limitation is the wording **“homeostatic biological processes.”** Direct evidence that RAB11A specifically mediates canonical neural homeostasis measures—such as firing-rate homeostasis or textbook “homeostatic synaptic scaling”—is limited. The strongest evidence is for receptor recycling, dendritic morphogenesis, synaptic plasticity, endosomal trafficking, and structural integrity, which are regulatory and maintenance-related processes, but not always labeled “homeostasis” in the narrow neurophysiology sense. ([pubmed.ncbi.nlm.nih.gov](https://pubmed.ncbi.nlm.nih.gov/28628388/)) [S9]

A third limitation is that some explicit “homeostasis” evidence for Rab11a comes from non-neural systems. For example, Rab11a knockout perturbs endosome-lysosome pathway homeostasis in HeLa cells, showing a central role in endolysosomal biogenesis and recycling. That finding gives a strong mechanistic rationale for neural relevance, but it is still an indirect extrapolation when applied to neurons. ([pubmed.ncbi.nlm.nih.gov](https://pubmed.ncbi.nlm.nih.gov/30981667/)) [S10]

Finally, human evidence remains relatively sparse compared with the large cell-biological literature. Neurodevelopmental associations are emerging, but most mechanistic support still comes from cultured neurons, neuron-like cells, rodent studies, zebrafish, or invertebrate disease models. ([pmc.ncbi.nlm.nih.gov](https://pmc.ncbi.nlm.nih.gov/articles/PMC12434753/)) [S11]

## Mechanistic / biological rationale

Biologically, the claim is plausible because RAB11A is a membrane-trafficking regulator that cycles between GDP- and GTP-bound states and organizes recycling endosome function. Active Rab11 recruits effectors such as Rab11-FIPs and Myosin Vb to sort internalized cargo away from degradation and back toward the plasma membrane or specific subcellular compartments. In neurons, where signaling depends on tight control of receptor number and localization, this kind of trafficking machinery is inherently regulatory. ([pubmed.ncbi.nlm.nih.gov](https://pubmed.ncbi.nlm.nih.gov/28628388/)) [S12]

In neural systems, that translates into several maintenance-like functions: preserving surface pools of AMPA receptors needed for synaptic potentiation, allowing activity-dependent insertion/removal of receptors during LTP/LTD, supporting BDNF-sensitive dendritic branching, and maintaining specialized apical structures in sensory hair cells. More broadly, Rab11a’s role in endosome-lysosome homeostasis provides a credible route by which its dysfunction could disturb neuronal proteostasis, receptor turnover, and organelle balance. ([sciencedirect.com](https://www.sciencedirect.com/science/article/pii/S0896627309004000)) [S13]

## Bottom line

The claim is **supported overall**: RAB11A/Rab11a has been repeatedly associated with **regulatory** processes in neural systems, especially receptor recycling, synaptic plasticity, dendritic morphology, and sensory-cell structural integrity. The evidence for **homeostatic** roles is real but often indirect or broader than strict “homeostatic plasticity” definitions, and some neural studies do not cleanly distinguish RAB11A from RAB11B. So the most defensible conclusion is **Likely True**, not stronger. ([pubmed.ncbi.nlm.nih.gov](https://pubmed.ncbi.nlm.nih.gov/32329391/)) [S14]

## Cited sources

- **[S1]** Rab11-mediated recycling endosome role in nervous system development and neurodegenerative diseases - PubMed — https://pubmed.ncbi.nlm.nih.gov/32329391/
- **[S2]** 
            BDNF Regulates Rab11-Mediated Recycling Endosome Dynamics to Induce Dendritic Branching - PMC
         — https://pmc.ncbi.nlm.nih.gov/articles/PMC3684039/
- **[S3]** Myosin Vb mobilizes recycling endosomes and AMPA receptors for postsynaptic plasticity - PubMed — https://pubmed.ncbi.nlm.nih.gov/18984164/
- **[S4]** Rab11a Controls Cell Shape via C9orf72 Protein: Possible Relationships to Frontotemporal Dementia/Amyotrophic Lateral Sclerosis (FTDALS) Type 1 - PubMed — https://pubmed.ncbi.nlm.nih.gov/38390945/
- **[S5]** 
            Rab11a and Myosin Vb Regulate Recycling of the M4Muscarinic Acetylcholine Receptor - PMC
         — https://pmc.ncbi.nlm.nih.gov/articles/PMC6757814/
- **[S6]** Rab11a Is Essential for the Development and Integrity of the Stereocilia and Kinocilia in the Mammalian Organ of Corti - PubMed — https://pubmed.ncbi.nlm.nih.gov/37225424/
- **[S7]** 
            A de novo mutation in RAB11A is associated with neurodevelopmental disorder accompanied by variable multisystem abnormalities - PMC
         — https://pmc.ncbi.nlm.nih.gov/articles/PMC12434753/
- **[S8]** Coordination of AMPA receptor trafficking by Rab GTPases - PubMed — https://pubmed.ncbi.nlm.nih.gov/28628388/
- **[S9]** Coordination of AMPA receptor trafficking by Rab GTPases - PubMed — https://pubmed.ncbi.nlm.nih.gov/28628388/
- **[S10]** A role for Rab11 in the homeostasis of the endosome-lysosomal pathway - PubMed — https://pubmed.ncbi.nlm.nih.gov/30981667/
- **[S11]** 
            A de novo mutation in RAB11A is associated with neurodevelopmental disorder accompanied by variable multisystem abnormalities - PMC
         — https://pmc.ncbi.nlm.nih.gov/articles/PMC12434753/
- **[S12]** Coordination of AMPA receptor trafficking by Rab GTPases - PubMed — https://pubmed.ncbi.nlm.nih.gov/28628388/
- **[S13]** Endocytic Trafficking and Recycling Maintain a Pool of Mobile Surface AMPA Receptors Required for Synaptic Potentiation - ScienceDirect — https://www.sciencedirect.com/science/article/pii/S0896627309004000
- **[S14]** Rab11-mediated recycling endosome role in nervous system development and neurodegenerative diseases - PubMed — https://pubmed.ncbi.nlm.nih.gov/32329391/
