## Supplementary material for "Understanding Substance Dependence: What Differentiates Addictive from Non-Addictive Drugs?": Folder for supplementary files: non-addictive-genes-claims-first100-086-pik3r1-gene-protein-has-been-associated-with-regul-f42a37be4023.md

# non-addictive-genes-claims-first100-086-f42a37be4023

**Original claim:** PIK3R1 gene/protein has been associated with regulatory or homeostatic biological processes in neural systems.

## Claim

PIK3R1 gene/protein has been associated with regulatory or homeostatic biological processes in neural systems. ([medlineplus.gov](https://medlineplus.gov/genetics/gene/pik3r1/)) [S1]

## Verdict

Verdict: Likely True

PIK3R1 is credibly linked to several **regulatory functions in neural systems**—including neural progenitor proliferation/differentiation, synaptic plasticity, neuronal survival, and hypothalamic control of metabolic homeostasis. However, the evidence is stronger for **specific neural regulatory roles** than for a single broadly defined “neural homeostasis” function, and some studies implicate the wider PI3K pathway rather than PIK3R1 alone. ([pubmed.ncbi.nlm.nih.gov](https://pubmed.ncbi.nlm.nih.gov/29953879/)) [S2]

## Confidence

Medium

The confidence is medium because there is direct experimental evidence in neural cells and animal models, but the claim is broad and part of the literature attributes effects to **class I PI3K signaling generally**, with PIK3R1 sometimes one component among several isoforms/subunits. Human in vivo evidence for normal neural homeostasis is more limited than the animal/cellular evidence. ([pmc.ncbi.nlm.nih.gov](https://pmc.ncbi.nlm.nih.gov/articles/PMC11335278/)) [S3]

## Supporting evidence

PIK3R1 encodes the p85α regulatory subunit of class IA PI3K, and authoritative databases place it in neural-relevant contexts. NCBI lists brain expression for human PIK3R1, while the Human Protein Atlas reports that it is detected across all major brain regions and clusters with neuronal-expression programs. KEGG maps PIK3R1 to the **neurotrophin signaling pathway** and **cholinergic synapse**, both clearly relevant to neural regulation. ([ncbi.nlm.nih.gov](https://www.ncbi.nlm.nih.gov/gene/5295?utm_source=openai)) [S4]

There is direct evidence that PIK3R1/p85 regulates **neural development**. In a mouse corticogenesis study, overexpression of p85 inhibited neural progenitor proliferation, inhibited neuronal differentiation, and prolonged cell-cycle duration in the developing cerebral cortex. That is a straightforward example of a neural regulatory role. ([pubmed.ncbi.nlm.nih.gov](https://pubmed.ncbi.nlm.nih.gov/29953879/)) [S5]

There is also direct evidence for a role in **synaptic regulation and plasticity**, which are core homeostatic/regulatory processes in neural circuits. A 2024 hippocampal study found that the PI3K regulatory subunit **p85α** was specifically required for activity-dependent cofilin recruitment into dendritic spines during LTP-like stimulation, and that this linked p85α to actin remodeling and synaptic potentiation. The same paper notes prior behavioral and cognitive deficits in p85α-deficient mice, supporting functional relevance in brain physiology. ([pmc.ncbi.nlm.nih.gov](https://pmc.ncbi.nlm.nih.gov/articles/PMC11335278/)) [S6]

Older in vivo behavioral work is directionally consistent: mice lacking p85α showed hyperactivity, object-recognition memory deficits, anxiety-related behavior, and prior reductions in PI3K activity associated with losses of synapses and myelinated axons in brain regions. While behavioral phenotypes are indirect, they fit with a role in maintaining normal neural circuit function. ([pubmed.ncbi.nlm.nih.gov](https://pubmed.ncbi.nlm.nih.gov/18538520/)) [S7]

PIK3R1 is also implicated in **neuronal control of systemic homeostasis** through the hypothalamus. In POMC-neuron-specific Pik3r1 knockout mice, altered PI3K signaling changed insulin sensitivity and glucose regulation; the authors concluded that PI3K activity in POMC neurons modulates hepatic insulin sensitivity and glucose homeostasis. That is not merely neural relevance—it is a direct example of a neuronal population using Pik3r1-dependent signaling to regulate homeostatic physiology. ([pmc.ncbi.nlm.nih.gov](https://pmc.ncbi.nlm.nih.gov/articles/PMC2775989/)) [S8]

Glia-neuron homeostatic support is another supportive line. In an astrocyte-mediated neuroprotection study, PI3K signaling identified through **PI3K (p85) immunoprecipitates** was required for astrocyte-conditioned-media neuroprotection, and the authors proposed a PI3K-dependent signaling complex mediating neuronal survival. This supports association with protective, homeostasis-like interactions in neural tissue, even though the study is framed at the PI3K/p85 complex level rather than exclusively by gene symbol. ([pmc.ncbi.nlm.nih.gov](https://pmc.ncbi.nlm.nih.gov/articles/PMC7948738/?utm_source=openai)) [S9]

## Contradicting or limiting evidence

A key limitation is **specificity**. Much of the neural literature is on the **PI3K-AKT pathway as a whole**, not always on PIK3R1 uniquely. Even in the hippocampal synaptic-plasticity work, p85β could exert opposite or compensatory effects, indicating that the outcome depends on relative regulatory-subunit composition rather than only PIK3R1 abundance. ([pmc.ncbi.nlm.nih.gov](https://pmc.ncbi.nlm.nih.gov/articles/PMC11335278/)) [S10]

Likewise, in hypothalamic POMC neurons, prior work cited by the same study showed that concurrent loss of p85α and p85β abolished responses to leptin/insulin, implying redundancy among regulatory subunits. So while Pik3r1 clearly matters, its role can be partially buffered or modified by other PI3K components. ([pmc.ncbi.nlm.nih.gov](https://pmc.ncbi.nlm.nih.gov/articles/PMC2775989/)) [S11]

Another limitation is that the word **“homeostatic”** is broader than the strongest evidence base. The most direct support is for neurodevelopmental regulation, synaptic plasticity, neuronal survival, and hypothalamic metabolic control. Evidence for PIK3R1 as a dedicated regulator of, for example, microglial steady-state homeostasis is more indirect and often inferred from class I PI3K signaling generally rather than from PIK3R1-specific perturbation. ([pmc.ncbi.nlm.nih.gov](https://pmc.ncbi.nlm.nih.gov/articles/PMC8627624/)) [S12]

Human evidence is also thinner than the animal/cellular literature. There are human disease associations and pathway annotations, but fewer clean human physiological studies showing that normal neural homeostasis depends specifically on PIK3R1. ([ncbi.nlm.nih.gov](https://www.ncbi.nlm.nih.gov/gene/5295?utm_source=openai)) [S13]

## Mechanistic / biological rationale

Mechanistically, PIK3R1 encodes a **regulatory subunit** that couples activated receptors to class IA PI3K signaling. That places it upstream of pathways controlling cell survival, metabolism, cytoskeletal dynamics, and growth-factor responses. In neural contexts, those are exactly the kinds of processes that underlie developmental regulation, synaptic remodeling, and tissue maintenance. ([medlineplus.gov](https://medlineplus.gov/genetics/gene/pik3r1/)) [S14]

In developing cortex, altered p85 levels can change progenitor proliferation and differentiation, plausibly by shifting PI3K/AKT-dependent control of cell cycle and maturation programs. In hippocampal neurons, p85α helps recruit cofilin into spines and influences Rac1/Cdc42-linked actin remodeling, providing a concrete mechanism for regulating synaptic structure and potentiation. ([pubmed.ncbi.nlm.nih.gov](https://pubmed.ncbi.nlm.nih.gov/29953879/)) [S15]

In hypothalamic neurons, PI3K signaling downstream of insulin/leptin receptors influences glucose regulation and body-weight responses, offering a mechanistic bridge between neural signaling and organismal homeostasis. In astrocyte-neuron systems, PI3K/p85-dependent signaling can support neuronal survival, which is consistent with a homeostatic support role for glia. ([pmc.ncbi.nlm.nih.gov](https://pmc.ncbi.nlm.nih.gov/articles/PMC2775989/)) [S16]

More broadly, reviews of neural PI3K biology describe the pathway as central to neuronal metabolism, tissue homeostasis, and neural stem/progenitor maintenance, while microglial reviews show PI3K-AKT signaling is used by receptors governing microglial behavior during homeostasis and neuroinflammation. Because PIK3R1 encodes a major regulatory node in that pathway and is expressed in neurons and glia, the overall biological rationale is strong, though not always PIK3R1-exclusive. ([pmc.ncbi.nlm.nih.gov](https://pmc.ncbi.nlm.nih.gov/articles/PMC6321294/)) [S17]

## Bottom line

The claim is **likely true**. PIK3R1 is not just a generic signaling gene with incidental brain expression; it has been tied experimentally to **neural progenitor regulation, synaptic plasticity, neuronal survival, and hypothalamic control of metabolic homeostasis**. The main caveat is that some evidence is pathway-level or isoform-shared rather than uniquely attributable to PIK3R1 in every neural context. ([pubmed.ncbi.nlm.nih.gov](https://pubmed.ncbi.nlm.nih.gov/29953879/)) [S18]

## Cited sources

- **[S1]** PIK3R1 gene: MedlinePlus Genetics — https://medlineplus.gov/genetics/gene/pik3r1/
- **[S2]** The effect of P85 on neuronal proliferation and differentiation during development of mouse cerebral cortex - PubMed — https://pubmed.ncbi.nlm.nih.gov/29953879/
- **[S3]** 
            PI3K couples long-term synaptic potentiation with cofilin recruitment and actin polymerization in dendritic spines via its regulatory subunit p85α - PMC
         — https://pmc.ncbi.nlm.nih.gov/articles/PMC11335278/
- **[S4]** PIK3R1 phosphoinositide-3-kinase regulatory subunit 1 [Homo sapiens (human)] - Gene - NCBI — https://www.ncbi.nlm.nih.gov/gene/5295?utm_source=openai
- **[S5]** The effect of P85 on neuronal proliferation and differentiation during development of mouse cerebral cortex - PubMed — https://pubmed.ncbi.nlm.nih.gov/29953879/
- **[S6]** 
            PI3K couples long-term synaptic potentiation with cofilin recruitment and actin polymerization in dendritic spines via its regulatory subunit p85α - PMC
         — https://pmc.ncbi.nlm.nih.gov/articles/PMC11335278/
- **[S7]** Hyperactivity, memory deficit and anxiety-related behaviors in mice lacking the p85alpha subunit of phosphoinositide-3 kinase - PubMed — https://pubmed.ncbi.nlm.nih.gov/18538520/
- **[S8]** 
            Phosphatidyl Inositol 3-Kinase Signaling in Hypothalamic Proopiomelanocortin Neurons Contributes to the Regulation of Glucose Homeostasis - PMC
         — https://pmc.ncbi.nlm.nih.gov/articles/PMC2775989/
- **[S9]** An endogenous PI3K interactome promoting astrocyte-mediated neuroprotection identifies a novel association with RNA-binding protein ZC3H14 - PMC — https://pmc.ncbi.nlm.nih.gov/articles/PMC7948738/?utm_source=openai
- **[S10]** 
            PI3K couples long-term synaptic potentiation with cofilin recruitment and actin polymerization in dendritic spines via its regulatory subunit p85α - PMC
         — https://pmc.ncbi.nlm.nih.gov/articles/PMC11335278/
- **[S11]** 
            Phosphatidyl Inositol 3-Kinase Signaling in Hypothalamic Proopiomelanocortin Neurons Contributes to the Regulation of Glucose Homeostasis - PMC
         — https://pmc.ncbi.nlm.nih.gov/articles/PMC2775989/
- **[S12]** 
            Dysregulated phosphoinositide 3-kinase signaling in microglia: shaping chronic neuroinflammation - PMC
         — https://pmc.ncbi.nlm.nih.gov/articles/PMC8627624/
- **[S13]** PIK3R1 phosphoinositide-3-kinase regulatory subunit 1 [Homo sapiens (human)] - Gene - NCBI — https://www.ncbi.nlm.nih.gov/gene/5295?utm_source=openai
- **[S14]** PIK3R1 gene: MedlinePlus Genetics — https://medlineplus.gov/genetics/gene/pik3r1/
- **[S15]** The effect of P85 on neuronal proliferation and differentiation during development of mouse cerebral cortex - PubMed — https://pubmed.ncbi.nlm.nih.gov/29953879/
- **[S16]** 
            Phosphatidyl Inositol 3-Kinase Signaling in Hypothalamic Proopiomelanocortin Neurons Contributes to the Regulation of Glucose Homeostasis - PMC
         — https://pmc.ncbi.nlm.nih.gov/articles/PMC2775989/
- **[S17]** 
            PI3K Signaling in Neurons: A Central Node for the Control of Multiple Functions - PMC
         — https://pmc.ncbi.nlm.nih.gov/articles/PMC6321294/
- **[S18]** The effect of P85 on neuronal proliferation and differentiation during development of mouse cerebral cortex - PubMed — https://pubmed.ncbi.nlm.nih.gov/29953879/
