## Supplementary material for "Understanding Substance Dependence: What Differentiates Addictive from Non-Addictive Drugs?": Folder for supplementary files: non-addictive-genes-claims-first100-087-dnajb6-gene-protein-has-been-associated-with-regul-9ba0691127cd.md

# non-addictive-genes-claims-first100-087-9ba0691127cd

**Original claim:** DNAJB6 gene/protein has been associated with regulatory or homeostatic biological processes in neural systems.

## Claim

**DNAJB6 gene/protein has been associated with regulatory or homeostatic biological processes in neural systems.**

## Verdict

Verdict: Likely True

## Confidence

**Medium**

## Supporting evidence

DNAJB6 encodes a DnaJ/Hsp40 family co-chaperone. Authoritative gene and expression resources describe it as a molecular chaperone involved in protein folding/complex assembly, and note a possible role in polyglutamine aggregation in specific neurons. Independent human brain data show DNAJB6 is broadly detected across brain regions and is found mainly in neurons and oligodendrocytes, supporting genuine relevance to neural tissue rather than a purely non-neural function. ([ncbi.nlm.nih.gov](https://www.ncbi.nlm.nih.gov/gene/10049?utm_source=openai)) [S1]

The strongest neural association is with **proteostasis**—a homeostatic process that maintains protein folding, trafficking, and degradation. In patient-derived neural progenitor/neuronal models of Huntington’s disease and spinocerebellar ataxia type 3, DNAJB6 expression drops during differentiation into neurons, and that drop tracks with increased neuronal susceptibility to amyloid-like polyglutamine aggregation. In the same study, increasing DNAJB6 in neurons antagonized glutamate-induced aggregation, while reducing DNAJB6 in progenitors promoted spontaneous aggregation. ([pubmed.ncbi.nlm.nih.gov](https://pubmed.ncbi.nlm.nih.gov/32268123/?utm_source=openai)) [S2]

Tau-related work points in the same direction. In a human neuroblastoma model, knockdown of DNAJB6 increased tau aggregation, whereas overexpression of the cytoplasmic isoform DNAJB6b reduced tau aggregation and promoted tau clearance; the J-domain was required for this protective effect. The same paper also reported age-dependent reduction of DNAJB6 expression alongside increased tau aggregation in hippocampal tissue from a tauopathy mouse model. That is a fairly direct association between DNAJB6 and maintenance of neuronal protein homeostasis. ([pubmed.ncbi.nlm.nih.gov](https://pubmed.ncbi.nlm.nih.gov/38110916/)) [S3]

Evidence is also substantial in synucleinopathy models. DNAJB6 was reported to suppress α-synuclein aggregation together with Hsp70, and earlier neuropathology work found DnaJB6 in Lewy bodies and highly upregulated in parkinsonian astrocytes. In a rat Parkinson’s disease model, overexpression of DNAJB6b reduced α-syn aggregation, dopaminergic neuron loss, and motor deficits. These findings place DNAJB6 within neural stress-response and proteostatic pathways that buffer toxic protein accumulation. ([pmc.ncbi.nlm.nih.gov](https://pmc.ncbi.nlm.nih.gov/articles/PMC5567236/)) [S4]

More recent ALS/FTD work extends the association beyond amyloid proteins. A 2025 study found that DNAJB6 reduced toxicity and insoluble species of FUS, TDP-43, and hnRNPA1, colocalized with FUS-containing granules in hiPSC-derived motor neurons, and prevented motor neuron loss plus microglial activation in a mouse FUS-ALS model. This supports a broader role for DNAJB6 in maintaining neural proteome stability under proteotoxic stress. ([pubmed.ncbi.nlm.nih.gov](https://pubmed.ncbi.nlm.nih.gov/41271702/)) [S5]

There is also evidence for **non-cell-autonomous** neural protection. In a Drosophila Huntington’s disease model, astrocyte-specific expression of DNAJB6 delayed neurodegeneration and extended lifespan even when the toxic polyQ protein was expressed only in neurons. That supports an association with glia-neuron homeostatic interactions, not only neuron-intrinsic quality control. ([pubmed.ncbi.nlm.nih.gov](https://pubmed.ncbi.nlm.nih.gov/30408590/)) [S6]

## Contradicting or limiting evidence

The main limitation is that most of the neural evidence is from **disease, stress, or overexpression/knockdown models**, not from normal healthy nervous system physiology. The literature I found is dominated by aggregation-prone proteins—polyQ, tau, α-synuclein, FUS, TDP-43, hnRNPA1—rather than classic neural homeostatic processes such as synaptic scaling, firing-rate homeostasis, neurotransmitter balance, or circuit stabilization. So the claim is well supported for **neural proteostasis**, but much less directly for broader neural homeostasis. ([pubmed.ncbi.nlm.nih.gov](https://pubmed.ncbi.nlm.nih.gov/32268123/?utm_source=openai)) [S7]

There is also evidence that DNAJB6’s effects are **context-dependent rather than uniformly protective**. In primary neurons, overexpression of DNAJB6 induced cell death in the absence of proteotoxic stress, yet was protective when mutant huntingtin was co-expressed. That suggests DNAJB6 participates in neural regulatory pathways, but not in a simple “more DNAJB6 is always homeostatic” manner. ([pubmed.ncbi.nlm.nih.gov](https://pubmed.ncbi.nlm.nih.gov/26476842/)) [S8]

Cell-type breadth is another qualifier. Although DNAJB6 is broadly present across brain regions, a 2024 human brain study found it mainly in neurons and oligodendrocytes, with only sparse presence in astrocytes and microglia. The same study did not find a significant change in the proportion of DNAJB6-positive oligodendrocytes in multiple system atrophy versus controls. So DNAJB6 is clearly neural-relevant, but current evidence does not support a universal role across all neural cell types or all disease contexts. ([pubmed.ncbi.nlm.nih.gov](https://pubmed.ncbi.nlm.nih.gov/39228066/)) [S9]

## Mechanistic / biological rationale

Biologically, the claim makes sense because **proteostasis is itself a homeostatic process**. Reviews define the proteostasis network as the system that regulates protein synthesis, folding, transport, and degradation to maintain proteome integrity and cellular homeostasis; HSP40/DNAJ proteins are a core part of that network. DNAJB6 fits this framework as a J-domain co-chaperone that partners with Hsp70 and regulates client-protein handling. ([pubmed.ncbi.nlm.nih.gov](https://pubmed.ncbi.nlm.nih.gov/28400444/?utm_source=openai)) [S10]

Mechanistic studies show that DNAJB6 can directly restrain pathogenic protein self-assembly. It suppresses Aβ42 amyloid formation at substoichiometric ratios by binding growing aggregates, cooperates with Hsp70 to suppress α-synuclein aggregation, and requires its J-domain to prevent tau aggregation. In the ALS/FTD context, DNAJB6 co-phase-separates with FUS condensates and keeps them in a looser, gel-like, less fibrillization-prone state. Together, these findings provide a coherent mechanism for why DNAJB6 would be associated with neural homeostatic regulation: it helps preserve protein quality control in long-lived, aggregation-vulnerable cells such as neurons. ([pmc.ncbi.nlm.nih.gov](https://pmc.ncbi.nlm.nih.gov/articles/PMC4223311/)) [S11]

## Bottom line

Overall, the claim is **likely true**. DNAJB6 is convincingly associated with **neural proteostasis/protein-quality-control pathways** in neurons, oligodendrocytes, and in some glial contexts, and multiple studies show that it modulates aggregation, toxicity, and cell survival in neural disease models. However, the evidence is strongest for **stress-responsive proteostatic homeostasis** in neurodegeneration-related settings, not for broad everyday neural homeostasis or canonical synaptic regulatory functions. ([pubmed.ncbi.nlm.nih.gov](https://pubmed.ncbi.nlm.nih.gov/39228066/)) [S12]

## Cited sources

- **[S1]** DNAJB6 DnaJ heat shock protein family (Hsp40) member B6 [Homo sapiens (human)] - Gene - NCBI — https://www.ncbi.nlm.nih.gov/gene/10049?utm_source=openai
- **[S2]** DNAJB6, a Key Factor in Neuronal Sensitivity to Amyloidogenesis - PubMed — https://pubmed.ncbi.nlm.nih.gov/32268123/?utm_source=openai
- **[S3]** The HSP40 family chaperone isoform DNAJB6b prevents neuronal cells from tau aggregation - PubMed — https://pubmed.ncbi.nlm.nih.gov/38110916/
- **[S4]** 
            The molecular chaperones DNAJB6 and Hsp70 cooperate to suppress α-synuclein aggregation - PMC
         — https://pmc.ncbi.nlm.nih.gov/articles/PMC5567236/
- **[S5]** Multiplex neurodegeneration proteotoxicity platform reveals DNAJB6 promotes non-toxic FUS condensate gelation and inhibits neurotoxicity - PubMed — https://pubmed.ncbi.nlm.nih.gov/41271702/
- **[S6]** Astrocytic expression of the chaperone DNAJB6 results in non-cell autonomous protection in Huntington's disease - PubMed — https://pubmed.ncbi.nlm.nih.gov/30408590/
- **[S7]** DNAJB6, a Key Factor in Neuronal Sensitivity to Amyloidogenesis - PubMed — https://pubmed.ncbi.nlm.nih.gov/32268123/?utm_source=openai
- **[S8]** Cell and Context-Dependent Effects of the Heat Shock Protein DNAJB6 on Neuronal Survival - PubMed — https://pubmed.ncbi.nlm.nih.gov/26476842/
- **[S9]** DNAJB6 is expressed in neurons and oligodendrocytes of the human brain - PubMed — https://pubmed.ncbi.nlm.nih.gov/39228066/
- **[S10]** Shaping proteostasis at the cellular, tissue, and organismal level - PubMed — https://pubmed.ncbi.nlm.nih.gov/28400444/?utm_source=openai
- **[S11]** 
            Interaction of the Molecular Chaperone DNAJB6 with Growing Amyloid-beta 42 (Aβ42) Aggregates Leads to Sub-stoichiometric Inhibition of Amyloid Formation - PMC
         — https://pmc.ncbi.nlm.nih.gov/articles/PMC4223311/
- **[S12]** DNAJB6 is expressed in neurons and oligodendrocytes of the human brain - PubMed — https://pubmed.ncbi.nlm.nih.gov/39228066/
