## Supplementary material for "Understanding Substance Dependence: What Differentiates Addictive from Non-Addictive Drugs?": Folder for supplementary files: non-addictive-genes-claims-first100-088-ddr1-gene-protein-has-been-associated-with-regulat-aacd706d37ee.md

# non-addictive-genes-claims-first100-088-aacd706d37ee

**Original claim:** DDR1 gene/protein has been associated with regulatory or homeostatic biological processes in neural systems.

## Claim

DDR1 gene/protein has been associated with regulatory or homeostatic biological processes in neural systems.

## Verdict

Verdict: Highly Supported

## Confidence

High

## Supporting evidence

DDR1 is clearly expressed in neural tissues and has been linked to multiple **regulatory** processes in the nervous system, especially in glia. Human Protein Atlas brain RNA data detect DDR1 across all major human brain regions, and several primary studies localize DDR1 to oligodendrocytes, myelin, and in some datasets astrocytes/endothelial cells. In adult human cortex, DDR1 mRNA and protein were identified in myelin and oligodendrocyte cell bodies, with expression correlating positively with myelin-associated genes such as **MAG** and **OLIG2**. ([proteinatlas.org](https://www.proteinatlas.org/ENSG00000204580-DDR1/brain)) [S1]

The strongest functional evidence is for **oligodendrocyte differentiation, myelination, and remyelination**, which are quintessential regulatory processes for neural integrity and conduction. A 2008 study found DDR1 upregulated during oligodendrocyte differentiation in vitro and during remyelination in the cuprizone mouse model, specifically in oligodendrocyte-lineage cells rather than microglia or reactive astrocytes. ([pubmed.ncbi.nlm.nih.gov](https://pubmed.ncbi.nlm.nih.gov/18836851/)) [S2]

More direct causal evidence came in 2023: in mouse CNS, Ddr1 was selectively upregulated in **newly differentiated oligodendrocytes**, and Ddr1 knockout delayed oligodendrocyte maturation, reduced axonal myelination, caused apparent motor dysfunction, and impaired myelin repair after lysolecithin-induced demyelination. That study also linked the phenotype to altered **ERK** signaling rather than AKT signaling. This is strong evidence that DDR1 participates in regulatory programs that maintain or restore white-matter function. ([pubmed.ncbi.nlm.nih.gov](https://pubmed.ncbi.nlm.nih.gov/37373466/)) [S3]

Additional mechanistic support comes from in vitro oligodendrocyte work showing that **collagen IV**, a known DDR1 ligand, induces DDR1 phosphorylation in human oligodendroglial cells and accelerates branching/maturation of primary oligodendrocytes. That fits a model in which DDR1 helps translate extracellular-matrix cues into glial maturation programs. ([pubmed.ncbi.nlm.nih.gov](https://pubmed.ncbi.nlm.nih.gov/36675255/)) [S4]

DDR1 has also been implicated in **neuronal morphogenesis**. In an early cerebellar study, DDR1 was detected in developing CNS regions, and dominant-negative DDR1 impaired neurite extension and parallel fiber extension of cerebellar granule neurons. This supports a regulatory role in neuronal process outgrowth, though this line of evidence is older and less extensive than the oligodendrocyte literature. ([pubmed.ncbi.nlm.nih.gov](https://pubmed.ncbi.nlm.nih.gov/10970885/)) [S5]

Beyond developmental regulation, DDR1 has been associated with processes relevant to **neural homeostasis under stress or injury**. In cultured microglia, collagen activated DDR1 and drove inflammatory outputs including nitric oxide, iNOS, COX-2, CD40, and MMP-9; blocking DDR1 blunted this response. In a rat ischemia-reperfusion model, DDR1 knockdown reduced phospho-DDR1, lowered MMP-9, decreased infarct volume, and lessened blood-brain barrier disruption. These findings place DDR1 in regulation of neuroinflammation and BBB integrity—both core components of CNS homeostasis, especially during injury. ([pubmed.ncbi.nlm.nih.gov](https://pubmed.ncbi.nlm.nih.gov/17969104/)) [S6]

Human transcriptomic studies further strengthen the “associated with regulatory processes” part of the claim. In developing and adult human brain, DDR1 coexpression modules were enriched for **oligodendrocyte-related and myelination-related genes**, and another large adult dorsolateral prefrontal cortex analysis placed DDR1 transcripts in modules enriched for **astrocyte/OPC proliferation**, **oligodendrocyte differentiation/myelination**, and even **neuronal synaptic transmission**. Coexpression alone does not prove function, but it independently supports DDR1’s embedding in neural regulatory networks. ([pubmed.ncbi.nlm.nih.gov](https://pubmed.ncbi.nlm.nih.gov/34323026/)) [S7]

Finally, disease-model work links DDR1 to proteostatic and inflammatory control in neurodegeneration. In mouse models challenged with α-synuclein, partial or complete DDR1 deletion or inhibition increased autophagy and reduced inflammatory and neurotoxic protein readouts. That suggests DDR1 can modulate pathways relevant to CNS homeostasis, although these data are mainly preclinical. ([pubmed.ncbi.nlm.nih.gov](https://pubmed.ncbi.nlm.nih.gov/32776088/?utm_source=openai)) [S8]

## Contradicting or limiting evidence

The main limitation is that **most causal evidence is preclinical**. The best data come from mouse knockouts, rat stroke models, and cell culture; human evidence is largely based on expression, colocalization, and coexpression rather than interventional studies. So while association is strong, definitive proof of DDR1’s role in normal human neural homeostasis is still incomplete. ([pubmed.ncbi.nlm.nih.gov](https://pubmed.ncbi.nlm.nih.gov/37373466/)) [S9]

There is also some **cell-type ambiguity**. Some studies emphasize oligodendrocytes and myelin, whereas others find associations with astrocytes, type 2 microglia, endothelial cells, or mixed-function brain clusters. For example, human cortical tissue work found DDR1 in myelin plus some astrocytes and endothelial cells, while transcriptomic analyses suggested that some schizophrenia-related DDR1 signals may be more astrocytic than oligodendroglial. The Human Protein Atlas also assigns DDR1 to a broad astrocyte/mixed-function RNA cluster rather than a purely oligodendrocyte-specific one. This does not weaken the overall claim, but it means DDR1’s neural role is probably **context-, cell-state-, and isoform-dependent** rather than restricted to one cell type. ([pubmed.ncbi.nlm.nih.gov](https://pubmed.ncbi.nlm.nih.gov/20380825/)) [S10]

The term **“homeostatic”** needs some caution. DDR1 is convincingly involved in regulatory neural processes, especially myelination/remyelination and injury responses. But direct evidence that DDR1 is a constitutive, central homeostatic controller in the healthy adult CNS is thinner than the evidence for developmental regulation and pathological response. A review on biological barriers explicitly notes that barrier-related DDR1 studies remain scarce and that mechanisms need further clarification. ([pubmed.ncbi.nlm.nih.gov](https://pubmed.ncbi.nlm.nih.gov/35665520/)) [S11]

Some neurodegeneration studies also use **kinase inhibitors with off-target activity**. For instance, nilotinib inhibits DDR1 but also Abl, even if reported CNS levels in those studies were argued to be more compatible with DDR1 inhibition. So the autophagy/neuroprotection evidence is supportive but not perfectly DDR1-specific. ([pmc.ncbi.nlm.nih.gov](https://pmc.ncbi.nlm.nih.gov/articles/PMC7566445/)) [S12]

## Mechanistic / biological rationale

DDR1 is a **collagen-activated receptor tyrosine kinase**, making it well positioned to convert extracellular-matrix signals into intracellular regulatory responses. In neural systems, that is biologically plausible because extracellular matrix composition helps govern oligodendrocyte maturation, axon-glia interactions, inflammation, and barrier behavior. ([sciencedirect.com](https://www.sciencedirect.com/science/article/pii/S0167488919300928)) [S13]

In oligodendrocytes, the emerging model is that DDR1 is expressed most strongly in **newly differentiated / early myelinating states**, where it helps drive maturation and myelin formation. The 2023 mouse knockout work links loss of Ddr1 to delayed differentiation and dysregulated ERK signaling; the collagen IV study shows ligand-dependent DDR1 phosphorylation and accelerated oligodendrocyte branching. Together, these findings support a mechanism in which DDR1 acts as an ECM-responsive regulator of oligodendrocyte state transitions and myelin repair. ([pubmed.ncbi.nlm.nih.gov](https://pubmed.ncbi.nlm.nih.gov/37373466/)) [S14]

In microglia and vascular injury contexts, DDR1 appears to couple collagen exposure to inflammatory signaling through **p38 MAPK, JNK, NF-κB**, and **MMP-9**, which can influence BBB permeability and tissue damage. In neurodegeneration models, DDR1 inhibition has been associated with increased autophagy and reduced toxic protein burden and inflammation, suggesting a role in balancing proteostasis and inflammatory tone. ([pubmed.ncbi.nlm.nih.gov](https://pubmed.ncbi.nlm.nih.gov/17969104/)) [S15]

## Bottom line

Yes—this claim is well supported. DDR1 has been repeatedly associated with **regulatory biological processes in neural systems**, most strongly **oligodendrocyte differentiation, myelination, remyelination, and neurite outgrowth**, and it is also implicated in **microglial inflammatory signaling, blood-brain barrier regulation, and autophagy-related responses** in neural injury and neurodegeneration models. The caveat is that the most convincing evidence is **preclinical**, and DDR1’s exact role in **baseline adult neural homeostasis** remains less firmly established than its role in developmental and injury-response regulation. ([pubmed.ncbi.nlm.nih.gov](https://pubmed.ncbi.nlm.nih.gov/37373466/)) [S16]

## Cited sources

- **[S1]** Brain tissue expression of DDR1 - Summary - The Human Protein Atlas — https://www.proteinatlas.org/ENSG00000204580-DDR1/brain
- **[S2]** Discoidin domain receptor 1, a tyrosine kinase receptor, is upregulated in an experimental model of remyelination and during oligodendrocyte differentiation in vitro - PubMed — https://pubmed.ncbi.nlm.nih.gov/18836851/
- **[S3]** Evidence That DDR1 Promotes Oligodendrocyte Differentiation during Development and Myelin Repair after Injury - PubMed — https://pubmed.ncbi.nlm.nih.gov/37373466/
- **[S4]** DDR1 and Its Ligand, Collagen IV, Are Involved in In Vitro Oligodendrocyte Maturation - PubMed — https://pubmed.ncbi.nlm.nih.gov/36675255/
- **[S5]** Discoidin domain receptor 1 functions in axon extension of cerebellar granule neurons - PubMed — https://pubmed.ncbi.nlm.nih.gov/10970885/
- **[S6]** Discoidin domain receptor 1 mediates collagen-induced inflammatory activation of microglia in culture - PubMed — https://pubmed.ncbi.nlm.nih.gov/17969104/
- **[S7]** Coexpression of the discoidin domain receptor 1 gene with oligodendrocyte-related and schizophrenia risk genes in the developing and adult human brain - PubMed — https://pubmed.ncbi.nlm.nih.gov/34323026/
- **[S8]** Discoidin Domain Receptor 1 is a therapeutic target for neurodegenerative diseases - PubMed — https://pubmed.ncbi.nlm.nih.gov/32776088/?utm_source=openai
- **[S9]** Evidence That DDR1 Promotes Oligodendrocyte Differentiation during Development and Myelin Repair after Injury - PubMed — https://pubmed.ncbi.nlm.nih.gov/37373466/
- **[S10]** Expression of the tyrosine kinase discoidin domain receptor 1 (DDR1) in human central nervous system myelin - PubMed — https://pubmed.ncbi.nlm.nih.gov/20380825/
- **[S11]** Discoidin domain receptor 1 may be involved in biological barrier homeostasis - PubMed — https://pubmed.ncbi.nlm.nih.gov/35665520/
- **[S12]** 
            Discoidin Domain Receptor 1 is a therapeutic target for neurodegenerative diseases - PMC
         — https://pmc.ncbi.nlm.nih.gov/articles/PMC7566445/
- **[S13]** Expression of DDR1 in the CNS and in myelinating oligodendrocytes - ScienceDirect — https://www.sciencedirect.com/science/article/pii/S0167488919300928
- **[S14]** Evidence That DDR1 Promotes Oligodendrocyte Differentiation during Development and Myelin Repair after Injury - PubMed — https://pubmed.ncbi.nlm.nih.gov/37373466/
- **[S15]** Discoidin domain receptor 1 mediates collagen-induced inflammatory activation of microglia in culture - PubMed — https://pubmed.ncbi.nlm.nih.gov/17969104/
- **[S16]** Evidence That DDR1 Promotes Oligodendrocyte Differentiation during Development and Myelin Repair after Injury - PubMed — https://pubmed.ncbi.nlm.nih.gov/37373466/
