## Supplementary material for "Understanding Substance Dependence: What Differentiates Addictive from Non-Addictive Drugs?": Folder for supplementary files: non-addictive-genes-claims-first100-089-sh3gl2-gene-protein-has-been-associated-with-regul-18cc0b20423f.md

# non-addictive-genes-claims-first100-089-18cc0b20423f

**Original claim:** SH3GL2 gene/protein has been associated with regulatory or homeostatic biological processes in neural systems.

## Claim

SH3GL2 gene/protein has been associated with regulatory or homeostatic biological processes in neural systems.

## Verdict

Verdict: Highly Supported

SH3GL2 encodes endophilin-A1, a brain-enriched membrane-remodeling/adaptor protein with substantial experimental evidence linking it to regulation of synaptic vesicle endocytosis, receptor trafficking, dendritic spine structure/plasticity, inhibitory synapse organization, excitation-inhibition balance, and activity-coupled synaptic autophagy/proteostasis in neural tissue. The strongest evidence comes from animal and neuronal cell studies, with supportive human genetic association data. ([ncbi.nlm.nih.gov](https://www.ncbi.nlm.nih.gov/gene/6456)) [S1]

## Confidence

High

The core statement is broad and well supported across multiple independent lines of evidence: curated gene annotations, classic synaptic trafficking studies, postsynaptic plasticity studies, newer inhibitory-synapse/E-I balance work, and synaptic autophagy papers. Confidence is slightly tempered because much of the mechanistic evidence is preclinical rather than from direct human functional studies. ([ncbi.nlm.nih.gov](https://www.ncbi.nlm.nih.gov/gene/20404)) [S2]

## Supporting evidence

SH3GL2 is a plausible neural regulatory gene on expression grounds alone. The human NCBI Gene record lists SH3GL2 as **biased in brain** expression, and the original discovery paper reported that SH3GL2 is **preferentially expressed in the brain/CNS**. The mouse ortholog is annotated as expressed in the central and peripheral nervous system and active in synaptic compartments including glutamatergic synapse and presynaptic cytosol. ([ncbi.nlm.nih.gov](https://www.ncbi.nlm.nih.gov/gene/6456)) [S3]

Curated functional annotations also support a regulatory neural role. The mouse gene summary states that Sh3gl2 is involved in **cellular response to brain-derived neurotrophic factor stimulus, dendrite extension, postsynaptic actin cytoskeleton organization, regulation of receptor internalization, and synaptic vesicle endocytosis**. The GO evidence table further includes experimentally supported annotations for **postsynaptic actin cytoskeleton organization** and **regulation of receptor internalization**, plus synaptic-vesicle-endocytosis-related annotations. Human GO annotations similarly include **cellular response to BDNF, central nervous system development, dendrite extension, and postsynaptic actin cytoskeleton organization**, although some of these are inference-based rather than direct human assays. ([ncbi.nlm.nih.gov](https://www.ncbi.nlm.nih.gov/gene/20404)) [S4]

The best-established function of endophilin-A1/SH3GL2 is in **synaptic vesicle endocytosis/recycling**, which is fundamentally regulatory for neurotransmission. In *C. elegans*, endophilin was shown to be required for synaptic vesicle recycling and to localize/stabilize synaptojanin at synapses. In mammalian neurons, loss of endophilin caused accumulation of clathrin-coated vesicles at synapses, impaired synaptic transmission, and severe neurological phenotypes including epilepsy and neurodegeneration, supporting a conserved role in regulating vesicle retrieval/uncoating at nerve terminals. Additional work showed endophilin is needed for ultrafast endocytosis and clathrin uncoating at synapses. ([pubmed.ncbi.nlm.nih.gov](https://pubmed.ncbi.nlm.nih.gov/14622579/)) [S5]

There is also strong evidence for a **postsynaptic regulatory role**. A 2015 study found that endophilin-A1 regulates dendritic spine morphogenesis and stability through interaction with p140Cap and actin-remodeling machinery. A later study showed that loss of endophilin-A1 in hippocampal CA1 neurons impairs AMPA-receptor-mediated synaptic transmission, long-term potentiation, and memory-related structural plasticity, while endophilin-A1 promotes actin polymerization in dendritic spines during synaptic potentiation. A 2021 study further linked endophilin-A1 to Ca²⁺/calmodulin-triggered acute spine enlargement during LTP. These are clear examples of SH3GL2 participating in regulatory processes in neural circuits. ([pubmed.ncbi.nlm.nih.gov](https://pubmed.ncbi.nlm.nih.gov/25771685/)) [S6]

Most directly relevant to “homeostatic” neural function, a 2024 study showed that endophilin-A1 localizes to inhibitory postsynapses, interacts with **gephyrin**, promotes organization of the inhibitory postsynaptic density, and stabilizes synaptic **GABA\_A receptor** clustering. In mice, loss of endophilin-A1 weakened inhibitory synaptic transmission, disrupted **excitation/inhibition balance**, and increased seizure susceptibility. That is a direct demonstration that SH3GL2 contributes to a core neural homeostatic/regulatory process. ([pubmed.ncbi.nlm.nih.gov](https://pubmed.ncbi.nlm.nih.gov/41036704/)) [S7]

SH3GL2 is also implicated in **synaptic autophagy and proteostasis**, which are canonical homeostatic processes. A recent mechanistic paper reported that Endophilin-A/SH3GL2 helps couple neuronal activity and calcium influx to **autophagy induction at synapses**. The authors explicitly interpret this pathway as supporting **synaptic homeostasis and protein quality control** during intense neuronal activity, and showed that a Parkinson’s disease risk variant in SH3GL2 disrupts this calcium-responsive autophagy mechanism. ([pmc.ncbi.nlm.nih.gov](https://pmc.ncbi.nlm.nih.gov/articles/PMC11062392/)) [S8]

More broadly, endophilin-A deficiency in brain was reported to cause **dysregulation of autophagy and the ubiquitin-proteasome system**, with the authors concluding that the FBXO32-endophilin network helps maintain **neuronal health and protein homeostasis**. Although that study concerns endophilin-A deficiency at the family level rather than SH3GL2 alone, it reinforces the idea that this protein family participates in neural homeostatic maintenance. ([pmc.ncbi.nlm.nih.gov](https://pmc.ncbi.nlm.nih.gov/articles/PMC5080600/)) [S9]

Human genetics adds supportive, though less mechanistic, evidence. A large Parkinson’s disease GWAS meta-analysis nominated **SH3GL2** as a PD risk-locus candidate gene and noted its likely role in synaptic vesicle endocytosis; later work connected a PD-risk SH3GL2 variant to defective synaptic autophagy. These findings do not prove normal physiology by themselves, but they are consistent with SH3GL2 acting in neural regulatory/homeostatic pathways relevant to neurodegeneration. ([pmc.ncbi.nlm.nih.gov](https://pmc.ncbi.nlm.nih.gov/articles/PMC5812477/)) [S10]

## Contradicting or limiting evidence

I did not find strong evidence contradicting the claim. The main limitations are about **scope and evidentiary level**, not direction. Much of the mechanistic evidence comes from **mouse, fly, worm, and cultured-neuron** systems rather than direct human physiology experiments. Thus, the existence of a neural regulatory role is well supported, but the exact extent of that role in humans is inferred partly from conserved biology and genetics. ([pubmed.ncbi.nlm.nih.gov](https://pubmed.ncbi.nlm.nih.gov/14622579/)) [S11]

Some curated annotations for SH3GL2 are also based on **electronic inference, sequence similarity, or orthology**, especially in the human record. The strongest parts of the case therefore come from the primary experimental literature on synaptic vesicle recycling, dendritic spine plasticity, inhibitory postsynaptic organization, and synaptic autophagy, rather than from annotation databases alone. ([ncbi.nlm.nih.gov](https://www.ncbi.nlm.nih.gov/gene/20404)) [S12]

Also, the term **“homeostatic”** deserves precision. For SH3GL2, the most direct homeostasis evidence is in **E/I balance** and **activity-linked synaptic autophagy/protein quality control**. Other roles, such as synaptic vesicle endocytosis or spine morphogenesis, are clearly **regulatory** but may be described as “homeostatic” only indirectly or in a broader systems sense. ([pubmed.ncbi.nlm.nih.gov](https://pubmed.ncbi.nlm.nih.gov/41036704/)) [S13]

## Mechanistic / biological rationale

Endophilin-A1 is a BAR-SH3 domain protein well suited to regulate neural membrane trafficking. Its BAR domain senses/induces membrane curvature, while its SH3 domain binds partners such as dynamin and synaptojanin, allowing it to help coordinate coated-pit constriction, vesicle uncoating, and synaptic vesicle recycling. That directly links SH3GL2 to regulation of neurotransmitter release machinery. ([pubmed.ncbi.nlm.nih.gov](https://pubmed.ncbi.nlm.nih.gov/22099461/)) [S14]

At the postsynaptic side, endophilin-A1 recruits p140Cap/cortactin-associated actin-remodeling machinery to dendritic spines and is required for normal spine morphogenesis, stabilization, and activity-dependent spine enlargement during LTP. This provides a mechanistic basis for its effects on synaptic strength, AMPAR-mediated transmission, and memory-related plasticity. ([pubmed.ncbi.nlm.nih.gov](https://pubmed.ncbi.nlm.nih.gov/25771685/)) [S15]

At inhibitory synapses, endophilin-A1 binds gephyrin and promotes plasma-membrane tethering plus actin-supported stabilization of inhibitory postsynaptic components, including GABA\_A receptors. This offers a direct mechanism by which SH3GL2 can help maintain circuit-level **E/I balance**, a classic homeostatic property of neural systems. ([pubmed.ncbi.nlm.nih.gov](https://pubmed.ncbi.nlm.nih.gov/41036704/)) [S16]

Finally, during intense neuronal activity, calcium-dependent redistribution of Endophilin-A/SH3GL2 promotes local synaptic autophagosome formation. That links neuronal firing to turnover of damaged proteins and organelles, providing a plausible mechanism for its role in **synaptic homeostasis, proteostasis, and neuroprotection**. ([pmc.ncbi.nlm.nih.gov](https://pmc.ncbi.nlm.nih.gov/articles/PMC11062392/)) [S17]

## Bottom line

Yes. The claim is **highly supported**: SH3GL2/endophilin-A1 is well associated with **regulatory neural processes** and has credible direct evidence for **homeostatic roles**, especially in **excitation-inhibition balance** and **activity-dependent synaptic autophagy/protein quality control**. The caveat is that most mechanistic proof comes from experimental model systems rather than direct human functional studies. ([ncbi.nlm.nih.gov](https://www.ncbi.nlm.nih.gov/gene/20404)) [S18]

## Cited sources

- **[S1]** SH3GL2 SH3 domain containing GRB2 like 2, endophilin A1 [Homo sapiens (human)] - Gene - NCBI — https://www.ncbi.nlm.nih.gov/gene/6456
- **[S2]** Sh3gl2 SH3-domain GRB2-like 2 [Mus musculus (house mouse)] - Gene - NCBI — https://www.ncbi.nlm.nih.gov/gene/20404
- **[S3]** SH3GL2 SH3 domain containing GRB2 like 2, endophilin A1 [Homo sapiens (human)] - Gene - NCBI — https://www.ncbi.nlm.nih.gov/gene/6456
- **[S4]** Sh3gl2 SH3-domain GRB2-like 2 [Mus musculus (house mouse)] - Gene - NCBI — https://www.ncbi.nlm.nih.gov/gene/20404
- **[S5]** Endophilin is required for synaptic vesicle endocytosis by localizing synaptojanin - PubMed — https://pubmed.ncbi.nlm.nih.gov/14622579/
- **[S6]** Endophilin A1 regulates dendritic spine morphogenesis and stability through interaction with p140Cap - PubMed — https://pubmed.ncbi.nlm.nih.gov/25771685/
- **[S7]** Endophilin A1 facilitates organization of the GABAergic postsynaptic machinery to maintain excitation-inhibition balance - PubMed — https://pubmed.ncbi.nlm.nih.gov/41036704/
- **[S8]** 
            Endophilin-A/SH3GL2 calcium switch for synaptic autophagy induction is impaired by a Parkinson’s risk variant - PMC
         — https://pmc.ncbi.nlm.nih.gov/articles/PMC11062392/
- **[S9]** 
            Endophilin-A Deficiency Induces the Foxo3a-Fbxo32 Network in the Brain and Causes Dysregulation of Autophagy and the Ubiquitin-Proteasome System - PMC
         — https://pmc.ncbi.nlm.nih.gov/articles/PMC5080600/
- **[S10]** 
            A meta-analysis of genome-wide association studies identifies 17 new Parkinson’s disease risk loci - PMC
         — https://pmc.ncbi.nlm.nih.gov/articles/PMC5812477/
- **[S11]** Endophilin is required for synaptic vesicle endocytosis by localizing synaptojanin - PubMed — https://pubmed.ncbi.nlm.nih.gov/14622579/
- **[S12]** Sh3gl2 SH3-domain GRB2-like 2 [Mus musculus (house mouse)] - Gene - NCBI — https://www.ncbi.nlm.nih.gov/gene/20404
- **[S13]** Endophilin A1 facilitates organization of the GABAergic postsynaptic machinery to maintain excitation-inhibition balance - PubMed — https://pubmed.ncbi.nlm.nih.gov/41036704/
- **[S14]** Recruitment of endophilin to clathrin-coated pit necks is required for efficient vesicle uncoating after fission - PubMed — https://pubmed.ncbi.nlm.nih.gov/22099461/
- **[S15]** Endophilin A1 regulates dendritic spine morphogenesis and stability through interaction with p140Cap - PubMed — https://pubmed.ncbi.nlm.nih.gov/25771685/
- **[S16]** Endophilin A1 facilitates organization of the GABAergic postsynaptic machinery to maintain excitation-inhibition balance - PubMed — https://pubmed.ncbi.nlm.nih.gov/41036704/
- **[S17]** 
            Endophilin-A/SH3GL2 calcium switch for synaptic autophagy induction is impaired by a Parkinson’s risk variant - PMC
         — https://pmc.ncbi.nlm.nih.gov/articles/PMC11062392/
- **[S18]** Sh3gl2 SH3-domain GRB2-like 2 [Mus musculus (house mouse)] - Gene - NCBI — https://www.ncbi.nlm.nih.gov/gene/20404
