## Supplementary material for "Understanding Substance Dependence: What Differentiates Addictive from Non-Addictive Drugs?": Folder for supplementary files: non-addictive-genes-claims-first100-090-ndufa11-gene-protein-has-been-associated-with-regu-e873291a90db.md

# non-addictive-genes-claims-first100-090-e873291a90db

**Original claim:** NDUFA11 gene/protein has been associated with regulatory or homeostatic biological processes in neural systems.

## Claim

NDUFA11 gene/protein has been associated with regulatory or homeostatic biological processes in neural systems.

## Verdict

Verdict: Likely True

NDUFA11 **has been associated** with neural-system homeostasis/regulation, but mostly **indirectly through mitochondrial complex I integrity, bioenergetic stability, oxidative-stress control, and neuronal survival/excitability under stress**, rather than as a classic neuron-specific signaling or transcriptional regulator. The strongest neural-specific evidence comes from recent ischemic-stroke and epilepsy studies showing altered NDUFA11/NDUFS1 complex status in neural models, plus a stroke study in primary cortical neurons where increasing Ndufa11 improved survival and electrophysiological recovery. Broader cell and organism studies strongly support a homeostatic role in mitochondrial respiration and ultrastructure, which is biologically relevant to neural tissue. ([ncbi.nlm.nih.gov](https://www.ncbi.nlm.nih.gov/gene/126328?utm_source=openai)) [S1]

## Confidence

Medium

The overall direction of evidence supports the claim, but the literature is still relatively sparse for **direct, normal-physiology neural homeostasis**. Much of the evidence is from mitochondrial cell biology, rare human disease, and recent disease-model studies rather than extensive neuron-specific loss/gain-of-function work in vivo. ([pmc.ncbi.nlm.nih.gov](https://pmc.ncbi.nlm.nih.gov/articles/PMC3839705/)) [S2]

## Supporting evidence

**1) Core molecular function is explicitly homeostatic at the mitochondrial level.** NDUFA11 encodes a subunit of membrane-bound mitochondrial complex I. In human cells, suppressing NDUFA11 impaired complex I assembly, reduced complex-I-linked oxygen consumption, and fragmented the mitochondrial network; the authors concluded that NDUFA11 behaves like an intrinsic assembly factor important for building the membrane arm of complex I. This is direct evidence that NDUFA11 supports mitochondrial respiratory stability rather than being a passive marker. ([ncbi.nlm.nih.gov](https://www.ncbi.nlm.nih.gov/gene/126328?utm_source=openai)) [S3]

**2) An animal model shows NDUFA11 is required for balanced respiration, mitochondrial morphology, and organismal health.** In *C. elegans*, loss or reduction of the NDUFA11 homolog destabilized complex I and supercomplexes, perturbed respiration, triggered compensatory complex II activity with potential ROS consequences, and caused abnormal cristae morphology. The authors explicitly interpreted NDUF-11 as required for “balanced respiration” and mitochondrial structural maintenance. Those are canonical homeostatic processes, even though this was not a neural-specific study. ([pubmed.ncbi.nlm.nih.gov](https://pubmed.ncbi.nlm.nih.gov/34106255/?utm_source=openai)) [S4]

**3) Human mutations cause disorders with clear neurologic involvement.** A 2008 case series identified a deleterious splice-site mutation in NDUFA11 in six patients from three unrelated families with isolated complex I deficiency. The reported phenotypes included encephalocardiomyopathy and fatal infantile lactic acidemia, placing NDUFA11 in the set of mitochondrial disease genes with brain involvement. That does not by itself prove a regulatory role, but it strongly links NDUFA11 dysfunction to failure of neural-system metabolic maintenance. ([pubmed.ncbi.nlm.nih.gov](https://pubmed.ncbi.nlm.nih.gov/18306244/)) [S5]

**4) There is now direct neural-model evidence in ischemic stroke.** A 2025 study found NDUFA11 expression was markedly reduced in blood from ischemic-stroke patients, and reduced in both in vitro and in vivo stroke models; it also reported fewer NDUFS1–NDUFA11 complexes in oxygen-glucose deprivation/reperfusion and MCAO models, with the proteins detected in neurons and cortex. This is a disease-context association between NDUFA11 and neuronal injury/homeostatic failure. ([pubmed.ncbi.nlm.nih.gov](https://pubmed.ncbi.nlm.nih.gov/39877656/)) [S6]

**5) A follow-up mechanistic stroke paper strengthens causality in neurons.** In a 2025 study of 865 ischemic-stroke patients with type 2 diabetes, metformin use was associated with milder neurologic deficits. In primary cortical neurons, metformin increased survival after OGD/R, restored action-potential amplitude, upregulated Ndufa11, increased the Ndufs1–Ndufa11 complex, and suppressed disulfidptosis-related changes. Importantly, lentiviral Ndufa11 overexpression recapitulated these protective effects, including improved neuronal survival and electrophysiology. That is strong evidence that Ndufa11 can participate in maintaining neural cellular function under stress. ([pubmed.ncbi.nlm.nih.gov](https://pubmed.ncbi.nlm.nih.gov/41236076/)) [S7]

**6) Another neural disease model shows a similar pattern.** In epilepsy, a 2025 study identified NDUFA11 among disulfidptosis-related molecules and found decreased NDUFS1–NDUFA11 complex formation in seizure models. This is not proof of a physiological homeostatic role in healthy brain, but it independently associates NDUFA11 with neural injury/stability pathways across a second CNS disorder. ([sciencedirect.com](https://www.sciencedirect.com/science/article/pii/S0969996125000051)) [S8]

**7) Brain association is also seen at the systems/genetic level.** Human atlas data show NDUFA11 has low tissue specificity and falls into a “cellular respiration” expression cluster, with expression detected across multiple brain regions. In addition, a genetic-imaging study in Alzheimer’s disease identified NDUFA11 variants associated with hippocampal subfield volumes. These are supportive but lower-strength associations. ([proteinatlas.org](https://www.proteinatlas.org/ENSG00000174886-NDUFA11/tissue)) [S9]

## Contradicting or limiting evidence

**Most evidence is not about a dedicated neural regulatory program.** NDUFA11 is best characterized as a **complex I structural/assembly component**, not as a canonical neural regulator such as a neurotransmitter receptor, ion-channel subunit, transcription factor, or synaptic homeostasis protein. So if the claim is interpreted narrowly—meaning a gene specifically known to regulate neural homeostasis as its primary function—the evidence is thinner. ([ncbi.nlm.nih.gov](https://www.ncbi.nlm.nih.gov/gene/126328?utm_source=openai)) [S10]

**Neural-specific data are still limited and heavily disease-focused.** The stroke and epilepsy papers are recent and mostly study injury states, disulfidptosis-related complexes, and biomarker behavior. They do not yet establish that NDUFA11 is a central regulator of normal neural homeostasis in healthy animals. ([pubmed.ncbi.nlm.nih.gov](https://pubmed.ncbi.nlm.nih.gov/41236076/)) [S11]

**Some supporting findings are associative rather than mechanistically decisive.** The hippocampal-volume study is genetic association; the stroke biomarker paper includes bioinformatic and validation data; and broad brain-expression atlases establish plausibility, not function. These support the claim’s “associated with” wording, but they are not definitive on their own. ([pmc.ncbi.nlm.nih.gov](https://pmc.ncbi.nlm.nih.gov/articles/PMC10915996/)) [S12]

**I did not find strong evidence that NDUFA11 directly regulates classic neural homeostatic domains** such as synaptic scaling, neurotransmitter release set-points, glial homeostatic signaling, or activity-dependent circuit stabilization in normal brain tissue. The current literature is more convincingly about mitochondrial maintenance with neural consequences. ([pmc.ncbi.nlm.nih.gov](https://pmc.ncbi.nlm.nih.gov/articles/PMC3839705/)) [S13]

## Mechanistic / biological rationale

Mechanistically, the literature supports NDUFA11 as a stabilizer of complex I and respiratory supercomplex organization. When NDUFA11 is reduced, complex I assembly falters, oxygen consumption falls, mitochondrial ultrastructure becomes abnormal, and ROS-related stress can rise. Inference: because neurons are highly dependent on mitochondrial respiration, a gene that preserves complex I integrity would reasonably contribute to neural homeostasis, especially during metabolic stress. ([pmc.ncbi.nlm.nih.gov](https://pmc.ncbi.nlm.nih.gov/articles/PMC3839705/)) [S14]

The neural papers fit that mechanism. In ischemic models, reduced NDUFA11 or reduced NDUFS1–NDUFA11 complex formation tracks with neuronal injury, while raising Ndufa11 improves neuronal survival and action-potential amplitude. That pattern is consistent with NDUFA11 supporting a **bioenergetic homeostasis layer** that helps neurons maintain viability and electrophysiological function under ischemic stress. ([pubmed.ncbi.nlm.nih.gov](https://pubmed.ncbi.nlm.nih.gov/41236076/)) [S15]

So biologically, NDUFA11’s neural relevance appears to be **mitochondrial and metabolic first**, with downstream effects on neuronal survival, excitability, and injury responses, rather than direct control of neural signaling pathways. ([ncbi.nlm.nih.gov](https://www.ncbi.nlm.nih.gov/gene/126328?utm_source=openai)) [S16]

## Bottom line

The claim is **likely true**, with an important qualifier: NDUFA11 is **not primarily established as a neuron-specific regulatory gene**, but the literature does associate it with **homeostatic functions relevant to neural systems** through mitochondrial complex I assembly/stability, respiratory balance, oxidative-stress handling, and protection of neurons during ischemic or seizure-related stress. The evidence is strongest for **mitochondrial homeostasis with neural consequences**, and weaker for a direct, specialized role in normal neural regulatory circuits. ([pmc.ncbi.nlm.nih.gov](https://pmc.ncbi.nlm.nih.gov/articles/PMC3839705/)) [S17]

## Cited sources

- **[S1]** NDUFA11 NADH:ubiquinone oxidoreductase subunit A11 [Homo sapiens (human)] - Gene - NCBI — https://www.ncbi.nlm.nih.gov/gene/126328?utm_source=openai
- **[S2]** 
            Assembly factors for the membrane arm of human complex I - PMC
         — https://pmc.ncbi.nlm.nih.gov/articles/PMC3839705/
- **[S3]** NDUFA11 NADH:ubiquinone oxidoreductase subunit A11 [Homo sapiens (human)] - Gene - NCBI — https://www.ncbi.nlm.nih.gov/gene/126328?utm_source=openai
- **[S4]** Maintenance of complex I and its supercomplexes by NDUF-11 is essential for mitochondrial structure, function and health - PubMed — https://pubmed.ncbi.nlm.nih.gov/34106255/?utm_source=openai
- **[S5]** Mitochondrial complex I deficiency caused by a deleterious NDUFA11 mutation - PubMed — https://pubmed.ncbi.nlm.nih.gov/18306244/
- **[S6]** NDUFA11 may be the disulfidptosis-related biomarker of ischemic stroke based on integrated bioinformatics, clinical samples, and experimental analyses - PubMed — https://pubmed.ncbi.nlm.nih.gov/39877656/
- **[S7]** Mechanisms of metformin in alleviating neurological deficits in stroke patients: Involving Ndufa11-mediated mitochondrial protection, enhanced neuronal electrophysiology, and inhibition of disulfidptosis - PubMed — https://pubmed.ncbi.nlm.nih.gov/41236076/
- **[S8]** Identifying disulfidptosis-related biomarkers in epilepsy based on integrated bioinformatics and experimental analyses - ScienceDirect — https://www.sciencedirect.com/science/article/pii/S0969996125000051
- **[S9]** Tissue expression of NDUFA11 - Summary - The Human Protein Atlas — https://www.proteinatlas.org/ENSG00000174886-NDUFA11/tissue
- **[S10]** NDUFA11 NADH:ubiquinone oxidoreductase subunit A11 [Homo sapiens (human)] - Gene - NCBI — https://www.ncbi.nlm.nih.gov/gene/126328?utm_source=openai
- **[S11]** Mechanisms of metformin in alleviating neurological deficits in stroke patients: Involving Ndufa11-mediated mitochondrial protection, enhanced neuronal electrophysiology, and inhibition of disulfidptosis - PubMed — https://pubmed.ncbi.nlm.nih.gov/41236076/
- **[S12]** 
            Genetic architecture of hippocampus subfields volumes in Alzheimer’s disease - PMC
         — https://pmc.ncbi.nlm.nih.gov/articles/PMC10915996/
- **[S13]** 
            Assembly factors for the membrane arm of human complex I - PMC
         — https://pmc.ncbi.nlm.nih.gov/articles/PMC3839705/
- **[S14]** 
            Assembly factors for the membrane arm of human complex I - PMC
         — https://pmc.ncbi.nlm.nih.gov/articles/PMC3839705/
- **[S15]** Mechanisms of metformin in alleviating neurological deficits in stroke patients: Involving Ndufa11-mediated mitochondrial protection, enhanced neuronal electrophysiology, and inhibition of disulfidptosis - PubMed — https://pubmed.ncbi.nlm.nih.gov/41236076/
- **[S16]** NDUFA11 NADH:ubiquinone oxidoreductase subunit A11 [Homo sapiens (human)] - Gene - NCBI — https://www.ncbi.nlm.nih.gov/gene/126328?utm_source=openai
- **[S17]** 
            Assembly factors for the membrane arm of human complex I - PMC
         — https://pmc.ncbi.nlm.nih.gov/articles/PMC3839705/
