## Supplementary material for "Understanding Substance Dependence: What Differentiates Addictive from Non-Addictive Drugs?": Folder for supplementary files: non-addictive-genes-claims-first100-091-vegfa-gene-protein-has-been-associated-with-regula-4e20710745a8.md

# non-addictive-genes-claims-first100-091-4e20710745a8

**Original claim:** VEGFA gene/protein has been associated with regulatory or homeostatic biological processes in neural systems.

## Claim

**VEGFA gene/protein has been associated with regulatory or homeostatic biological processes in neural systems.** ([pmc.ncbi.nlm.nih.gov](https://pmc.ncbi.nlm.nih.gov/articles/PMC11884128/)) [S1]

## Verdict

Verdict: Highly Supported

Across developmental, adult, and injury-related neuroscience literature, VEGFA is repeatedly implicated in **regulatory functions** in neural systems, including neuronal migration, axon guidance, synaptic transmission/plasticity, neurogenesis, and neural stem-cell maintenance. Evidence for a specifically **homeostatic** role is strongest in the **neurovascular unit / blood-brain barrier milieu** and in **maintenance of adult hippocampal stem-cell pools**, though this role is context-dependent because excess or reactive VEGFA can also destabilize barrier integrity. ([pmc.ncbi.nlm.nih.gov](https://pmc.ncbi.nlm.nih.gov/articles/PMC11884128/)) [S2]

## Confidence

**High.** The claim is broad and uses the weak causal language “has been associated with,” and that standard is met by multiple reviews plus diverse primary studies in mice, zebrafish, and human neural cell systems. The main limitation is that much of the strongest mechanistic evidence is preclinical rather than direct human in vivo evidence. ([pubmed.ncbi.nlm.nih.gov](https://pubmed.ncbi.nlm.nih.gov/20885857/?utm_source=openai)) [S3]

## Supporting evidence

1. **Reviews consistently place VEGFA in neural regulatory biology, not just angiogenesis.** A major review in *Neuron Glia Biology* states that VEGF/VEGFA is critical in the developing and adult nervous system and discusses roles in neurogenesis, neuronal migration, glia, and neuronal health. A recent 2025 review likewise summarizes VEGFA effects on neuronal development, synaptic excitability/transmission/plasticity, dendritic structure, and neuroprotection. ([pubmed.ncbi.nlm.nih.gov](https://pubmed.ncbi.nlm.nih.gov/20885857/?utm_source=openai)) [S4]

2. **Adult hippocampal plasticity and neurogenesis are regulated by VEGFA signaling.** In a classic *Nature Genetics* study, enriched environment and spatial learning increased hippocampal VEGF, hippocampal VEGF gene transfer roughly doubled neurogenesis and improved cognition, and inhibition of KDR/VEGFR2 impaired basal neurogenesis and learning. That is direct evidence for VEGFA as a regulator of adult neural plasticity-related processes. ([pubmed.ncbi.nlm.nih.gov](https://pubmed.ncbi.nlm.nih.gov/15258583/)) [S5]

3. **VEGFA modulates synaptic function and LTP in mature neural circuits.** In a conditional mouse study, reversible VEGF overexpression increased memory and dentate gyrus LTP, whereas blocking endogenous VEGF impaired memory and abolished LTP, even at time points too early for newly generated neurons to explain the effects. This strongly supports a role in ongoing regulation of mature neuronal plasticity. ([pubmed.ncbi.nlm.nih.gov](https://pubmed.ncbi.nlm.nih.gov/21385942/)) [S6]

4. **Direct neuronal VEGF-VEGFR2 signaling has been demonstrated.** A 2016 study found that VEGF increases NMDA receptor-mediated postsynaptic responses, promotes new synapse formation, and that neural-cell VEGFR2 silencing impairs hippocampal synaptic plasticity and emotional-memory consolidation. The authors explicitly concluded that VEGF signaling in neurons via VEGFR2 is required for proper synaptic function. ([pubmed.ncbi.nlm.nih.gov](https://pubmed.ncbi.nlm.nih.gov/26728568/)) [S7]

5. **VEGFA is linked to maintenance of adult neural stem cells, a homeostatic-like function.** A 2015 paper reported that adult hippocampal neural stem/progenitor cells secrete VEGF and that this contributes to maintenance of the stem-cell pool in the adult subgranular zone. A 2025 follow-up mechanistic study further concluded that intracrine VEGF-VEGFR2 signaling in adult dentate gyrus neural stem cells prevents stem-cell exhaustion and supports their proximity to local blood vessels. ([pmc.ncbi.nlm.nih.gov](https://pmc.ncbi.nlm.nih.gov/articles/PMC4386397/)) [S8]

6. **VEGFA regulates neural development beyond vascular growth.** VEGFA/NRP1 signaling promotes migration, survival, and axon guidance in specific neuronal populations, including facial branchiomotor neurons, GnRH neurons, and retinal ganglion cell axons. In the developing forebrain, vascular-derived Vegfa altered cortical interneuron numbers, intracortical distribution, and proximity to blood vessels, indicating a direct developmental regulatory role in neural circuit assembly. ([pmc.ncbi.nlm.nih.gov](https://pmc.ncbi.nlm.nih.gov/articles/PMC4302834/)) [S9]

7. **VEGFA participates in neurovascular organization relevant to brain homeostasis.** BBB-forming and fenestrated brain capillaries are crucial for region-specific neural function and brain homeostasis. A 2023 eLife study identified Vegfa as part of the ligand interplay shaping fenestrated CNS vascular beds, and a review of blood-neural barriers emphasizes that these barriers preserve CNS homeostasis by controlling molecular flux. Together, these data support association of VEGFA with neural-system homeostatic regulation at the neurovascular interface. ([pmc.ncbi.nlm.nih.gov](https://pmc.ncbi.nlm.nih.gov/articles/PMC10229134/)) [S10]

8. **Human neural-cell evidence also points in the same direction.** In human neural progenitor-derived cultures, endogenous VEGFA/VEGFR2 signaling was present, VEGFA promoted neurite growth and neuronal differentiation, and inhibition of VEGFA signaling with sFlt-1 reduced these neurogenic effects. This is not in vivo proof, but it supports conservation of VEGFA’s neural regulatory role in human cells. ([pubmed.ncbi.nlm.nih.gov](https://pubmed.ncbi.nlm.nih.gov/38700092/)) [S11]

## Contradicting or limiting evidence

1. **Most strong mechanistic evidence is preclinical.** The clearest causal data come from mouse, zebrafish, and in vitro systems; direct human in vivo evidence for VEGFA as a homeostatic neural regulator remains limited. ([pubmed.ncbi.nlm.nih.gov](https://pubmed.ncbi.nlm.nih.gov/15258583/)) [S12]

2. **Not all observed neural effects are necessarily direct.** Reviews note that VEGFA acts on endothelial cells, glia, and neurons, and disentangling direct neuronal signaling from indirect vascular or niche-mediated effects is often difficult. That matters especially for claims about “neural homeostasis,” where some effects may be mediated through the neurovascular unit rather than neurons alone. ([pubmed.ncbi.nlm.nih.gov](https://pubmed.ncbi.nlm.nih.gov/20885857/?utm_source=openai)) [S13]

3. **VEGFA is context-dependent and can oppose homeostasis when dysregulated.** In inflammatory CNS disease models, astrocyte-derived VEGF-A disrupts claudin-5/occludin, increases BBB permeability, and worsens immune infiltration and tissue injury. So VEGFA is not uniformly homeostatic; physiologic versus pathological context is crucial. ([pmc.ncbi.nlm.nih.gov](https://pmc.ncbi.nlm.nih.gov/articles/PMC3386814/)) [S14]

4. **Some structural effects are stage-specific rather than universal maintenance functions.** The 2025 review notes that blocking VEGFA does not affect already established mature dendrites and may even increase spine formation after neuronal maturation, suggesting VEGFA’s roles vary by developmental stage, cell type, and readout. ([pmc.ncbi.nlm.nih.gov](https://pmc.ncbi.nlm.nih.gov/articles/PMC11884128/)) [S15]

## Mechanistic / biological rationale

VEGFA is a pleiotropic ligand that signals through **VEGFR2/KDR**, **VEGFR1/FLT1**, and **neuropilin-1 (NRP1)**, with receptor usage varying by cell type and developmental context. In neurons, NRP1-dependent VEGFA signaling helps guide migration and axon projection in select populations. In hippocampal circuits, VEGFA-VEGFR2 signaling enhances NMDA receptor-related synaptic responses, supports LTP, and influences memory-related plasticity. In adult neural stem cells, VEGFA can function through an intracrine VEGF-VEGFR2 mechanism to preserve the stem-cell pool and vascular association. At the systems level, VEGFA also shapes neurovascular architecture and barrier specialization, which are key determinants of the chemical milieu needed for neural stability and function. ([pmc.ncbi.nlm.nih.gov](https://pmc.ncbi.nlm.nih.gov/articles/PMC4302834/)) [S16]

## Bottom line

The claim is **well supported**. VEGFA is not just an angiogenic factor; it has substantial literature linking it to **regulation of neural development, synaptic plasticity, cognition, neurogenesis, and neural stem-cell maintenance**, and it also contributes to **neurovascular processes that support brain homeostasis**. The main caveat is that “homeostatic” is more strongly supported for **stem-cell maintenance and neurovascular/barrier regulation** than for all neural contexts, and excessive VEGFA can become pathogenic rather than stabilizing. ([pubmed.ncbi.nlm.nih.gov](https://pubmed.ncbi.nlm.nih.gov/15258583/)) [S17]

## Cited sources

- **[S1]** 
            Beyond vessels: unraveling the impact of VEGFs on neuronal functions and structure - PMC
         — https://pmc.ncbi.nlm.nih.gov/articles/PMC11884128/
- **[S2]** 
            Beyond vessels: unraveling the impact of VEGFs on neuronal functions and structure - PMC
         — https://pmc.ncbi.nlm.nih.gov/articles/PMC11884128/
- **[S3]** VEGF in the nervous system - PubMed — https://pubmed.ncbi.nlm.nih.gov/20885857/?utm_source=openai
- **[S4]** VEGF in the nervous system - PubMed — https://pubmed.ncbi.nlm.nih.gov/20885857/?utm_source=openai
- **[S5]** VEGF links hippocampal activity with neurogenesis, learning and memory - PubMed — https://pubmed.ncbi.nlm.nih.gov/15258583/
- **[S6]** Reversible modulations of neuronal plasticity by VEGF - PubMed — https://pubmed.ncbi.nlm.nih.gov/21385942/
- **[S7]** A critical role for VEGF and VEGFR2 in NMDA receptor synaptic function and fear-related behavior - PubMed — https://pubmed.ncbi.nlm.nih.gov/26728568/
- **[S8]** 
            Adult hippocampal neural stem and progenitor cells regulate the neurogenic niche by secreting VEGF - PMC
         — https://pmc.ncbi.nlm.nih.gov/articles/PMC4386397/
- **[S9]** 
            VEGF189 binds NRP1 and is sufficient for VEGF/NRP1-dependent neuronal patterning in the developing brain - PMC
         — https://pmc.ncbi.nlm.nih.gov/articles/PMC4302834/
- **[S10]** 
            Local angiogenic interplay of Vegfc/d and Vegfa controls brain region-specific emergence of fenestrated capillaries - PMC
         — https://pmc.ncbi.nlm.nih.gov/articles/PMC10229134/
- **[S11]** sFlt-1 impairs neurite growth and neuronal differentiation in SH-SY5Y cells and human neurons - PubMed — https://pubmed.ncbi.nlm.nih.gov/38700092/
- **[S12]** VEGF links hippocampal activity with neurogenesis, learning and memory - PubMed — https://pubmed.ncbi.nlm.nih.gov/15258583/
- **[S13]** VEGF in the nervous system - PubMed — https://pubmed.ncbi.nlm.nih.gov/20885857/?utm_source=openai
- **[S14]** 
            Astrocyte-derived VEGF-A drives blood-brain barrier disruption in CNS inflammatory disease - PMC
         — https://pmc.ncbi.nlm.nih.gov/articles/PMC3386814/
- **[S15]** 
            Beyond vessels: unraveling the impact of VEGFs on neuronal functions and structure - PMC
         — https://pmc.ncbi.nlm.nih.gov/articles/PMC11884128/
- **[S16]** 
            VEGF189 binds NRP1 and is sufficient for VEGF/NRP1-dependent neuronal patterning in the developing brain - PMC
         — https://pmc.ncbi.nlm.nih.gov/articles/PMC4302834/
- **[S17]** VEGF links hippocampal activity with neurogenesis, learning and memory - PubMed — https://pubmed.ncbi.nlm.nih.gov/15258583/
