## Supplementary material for "Understanding Substance Dependence: What Differentiates Addictive from Non-Addictive Drugs?": Folder for supplementary files: non-addictive-genes-claims-first100-092-kcnj13-gene-protein-has-been-associated-with-regul-1d611d80a9a4.md

# non-addictive-genes-claims-first100-092-1d611d80a9a4

**Original claim:** KCNJ13 gene/protein has been associated with regulatory or homeostatic biological processes in neural systems.

## Claim

KCNJ13 gene/protein has been associated with regulatory or homeostatic biological processes in neural systems.

## Verdict

Verdict: Likely True

## Confidence

High

## Supporting evidence

KCNJ13 encodes the inwardly rectifying potassium channel Kir7.1. The strongest evidence tying it to neural homeostasis comes from the retina, where Kir7.1 is concentrated in the apical membrane of retinal pigment epithelium (RPE), a support tissue that is essential for nourishing and maintaining the **neural retina**. Multiple studies report that Kir7.1 helps move K\(^+\) from RPE cells into the subretinal space and supports normal retinal electrophysiology. Loss or suppression of Kcnj13/Kir7.1 in mouse RPE causes reduced ERG responses, photoreceptor degeneration, and impaired survival of adjacent retinal neurons, all consistent with a homeostatic support role for neural tissue rather than a purely structural role. ([pmc.ncbi.nlm.nih.gov](https://pmc.ncbi.nlm.nih.gov/articles/PMC6707361/)) [S1]

Older and newer physiology studies converge on the same point: Kir7.1 is a major component of the apical RPE K\(^+\) conductance and is positioned to respond to light-evoked changes in subretinal K\(^+\). In bovine RPE, Kir7.1 was identified as the dominant apical Kir subunit and was proposed to regulate K\(^+\) transport and the electrical response of the RPE to changes in subretinal K\(^+\) concentration. A recent mouse study further concluded that Kir7.1 is crucial for buffering changes in subretinal K\(^+\) during light stimulation because of its unusual dependence on extracellular K\(^+\). ([pubmed.ncbi.nlm.nih.gov](https://pubmed.ncbi.nlm.nih.gov/12824269/?utm_source=openai)) [S2]

Human-cell evidence also supports a homeostatic/supportive role in the retinal neural environment. In CRISPR-edited human iPSC-derived RPE, KCNJ13 deletion significantly reduced phagocytic uptake of photoreceptor outer segments and lowered expression of phagocytosis-related genes. Because outer-segment clearance is a classic RPE maintenance function required for photoreceptor health, this links KCNJ13 to a bona fide regulatory/homeostatic process in a neural system. Separate iPSC-RPE work found that KCNJ13 loss increases oxidative-stress vulnerability and cell death, again pointing to a role in maintaining a stable retinal support environment. ([pubmed.ncbi.nlm.nih.gov](https://pubmed.ncbi.nlm.nih.gov/32437550/)) [S3]

Evidence is not limited to the eye. In the brain’s choroid plexus epithelium, Kir7.1 is prominently expressed with Na\(^+\)/K\(^+\)-ATPase and NKCC1. A 2025 mouse study found that conditional loss or altered K\(^+\)-dependence of Kir7.1 changed cerebrospinal fluid (CSF) K\(^+\) concentration, and concluded that Kir7.1 has an important role in regulating CSF K\(^+\). Because CSF is continuous with brain interstitial fluid, the authors noted that Kir7.1-mediated control of CSF K\(^+\) could influence neuronal excitability. Earlier localization studies had already shown Kir7.1 enrichment in choroid plexus epithelium, supporting a long-standing link between KCNJ13 and CNS ionic homeostasis. ([pubmed.ncbi.nlm.nih.gov](https://pubmed.ncbi.nlm.nih.gov/41212743/)) [S4]

There is also direct CNS glial evidence. In mouse optic nerve, Kcnj13/Kir7.1 transcripts were found at levels comparable to Kir4.1, a canonical glial K\(^+\) channel. Patch-clamp and pharmacology showed that Kir7.1 contributes substantially to oligodendrocyte K\(^+\) conductance, and blocking Kir7.1 compromised oligodendrocyte survival and worsened injury during oxygen-glucose deprivation. The authors concluded that Kir7.1 helps maintain oligodendroglial integrity and likely supports ion/water homeostasis in white matter. That is a direct association with a classic neural homeostatic process. ([pmc.ncbi.nlm.nih.gov](https://pmc.ncbi.nlm.nih.gov/articles/PMC7166203/)) [S5]

Additional neural-regulatory evidence comes from hypothalamic neurons. MC4R signaling has been shown to couple to Kir7.1 in hypothalamic PVN neurons, altering membrane potential and firing independently of canonical G\(_s\)/cAMP signaling. Later work further supported Kcnj13 expression in PVN neurons and its role in regulating neuronal firing and feeding-related circuitry. This is more about neuromodulatory regulation than tissue homeostasis per se, but it still supports the claim that KCNJ13 participates in regulatory processes in neural systems. ([pubmed.ncbi.nlm.nih.gov](https://pubmed.ncbi.nlm.nih.gov/25600267/)) [S6]

## Contradicting or limiting evidence

The evidence base is strongest for **supportive epithelia and glia** that maintain neural environments, especially RPE and choroid plexus, rather than for a broad, well-established role in neurons throughout the human CNS. In fact, one mouse developmental/pathology study concluded that later ocular expression becomes centered on the RPE with no or lower expression in the neuroretina, implying that much of the neural relevance in the eye is indirect—through support of photoreceptors—rather than through strong expression in retinal neurons themselves. ([pubmed.ncbi.nlm.nih.gov](https://pubmed.ncbi.nlm.nih.gov/35096838/)) [S7]

Likewise, although adult mouse brain studies reported widespread Kir7.1 immunostaining in neurons and glia, those same authors explicitly stated that the physiological function of Kir7.1 in neurons and glia remains unresolved. So expression data and plausible inference extend beyond retina/choroid plexus, but definitive functional proof across the CNS is still incomplete. ([pmc.ncbi.nlm.nih.gov](https://pmc.ncbi.nlm.nih.gov/articles/PMC6794205/)) [S8]

There is also some mechanistic nuance within the retina. A 2025 mutant-mouse study supported the idea that Kir7.1 is crucial for buffering subretinal K\(^+\), but found that selectively disrupting this buffering property altered the ERG c-wave without clearly impairing a- or b-waves. The authors inferred that disease from KCNJ13 mutations may involve additional functions—such as outer-segment recycling or other RPE processes—beyond K\(^+\) buffering alone. That does not contradict a homeostatic role, but it means “subretinal K\(^+\) homeostasis” is probably not the whole story. ([pubmed.ncbi.nlm.nih.gov](https://pubmed.ncbi.nlm.nih.gov/41247777/)) [S9]

Finally, much of the strongest mechanistic evidence comes from mouse tissue, ex vivo electrophysiology, and iPSC-derived RPE models rather than direct human in vivo neurophysiology. Human genetics clearly links KCNJ13 to retinal disease, but direct demonstration of broader human brain homeostatic phenotypes remains limited. ([pubmed.ncbi.nlm.nih.gov](https://pubmed.ncbi.nlm.nih.gov/21763485/?utm_source=openai)) [S10]

## Mechanistic / biological rationale

Kir7.1’s biophysical properties fit a homeostatic role. Unlike many other Kir channels, it has low single-channel conductance and unusually weak or atypical dependence on extracellular K\(^+\), making it well suited to provide a relatively steady background K\(^+\) conductance. In RPE and choroid plexus, Kir7.1 colocalizes with Na\(^+\)/K\(^+\)-ATPase; this architecture supports “K\(^+\) recycling,” helps set membrane potential, and sustains transepithelial ion transport. In practical terms, that means KCNJ13 is positioned to stabilize extracellular ionic conditions in compartments that directly affect photoreceptor excitability and CSF composition. ([pmc.ncbi.nlm.nih.gov](https://pmc.ncbi.nlm.nih.gov/articles/PMC4594557/)) [S11]

In the retina, light changes subretinal K\(^+\) concentration. Kir7.1 in RPE is thought to buffer those shifts, support fluid transport, and preserve the microenvironment around photoreceptor outer segments. Loss of KCNJ13 disrupts this environment and is associated with reduced ERG responses, photoreceptor degeneration, impaired phagocytosis of outer segments, and increased oxidative-stress susceptibility in RPE. Together, those effects are exactly what one would expect from failure of a homeostatic support mechanism in a neural tissue. ([pmc.ncbi.nlm.nih.gov](https://pmc.ncbi.nlm.nih.gov/articles/PMC6707361/)) [S12]

In the choroid plexus and brain, the rationale is similar: Kir7.1 contributes to the epithelial membrane potential and to control of CSF K\(^+\), a parameter that can influence neuronal excitability. In glia and some neurons, available data further suggest roles in K\(^+\)/water handling and excitability control. Kir7.1 is also modulated by signaling systems including MC4R-related pathways and steroid signaling, reinforcing that it participates in regulated neural-support physiology rather than passive ion leak alone. ([pubmed.ncbi.nlm.nih.gov](https://pubmed.ncbi.nlm.nih.gov/41212743/)) [S13]

## Bottom line

The claim is **likely true**. KCNJ13/Kir7.1 is convincingly associated with **homeostatic regulation in neural-support compartments**, especially the retinal pigment epithelium and choroid plexus, where it helps maintain ionic conditions relevant to photoreceptors and the brain. There is also supportive evidence for roles in oligodendrocytes and hypothalamic neuronal excitability. The main caveat is that the best evidence is concentrated in retina/choroid plexus and in animal or cell-model systems, so broader CNS-wide functions in humans are less firmly established. ([pmc.ncbi.nlm.nih.gov](https://pmc.ncbi.nlm.nih.gov/articles/PMC6707361/)) [S14]

## Cited sources

- **[S1]** 
            Conditional Loss of Kcnj13 in the Retinal Pigment Epithelium Causes Photoreceptor Degeneration - PMC
         — https://pmc.ncbi.nlm.nih.gov/articles/PMC6707361/
- **[S2]** Expression and localization of the inwardly rectifying potassium channel Kir7.1 in native bovine retinal pigment epithelium - PubMed — https://pubmed.ncbi.nlm.nih.gov/12824269/?utm_source=openai
- **[S3]** KCNJ13 Gene Deletion Impairs Cell Alignment and Phagocytosis in Retinal Pigment Epithelium Derived from Human-Induced Pluripotent Stem Cells - PubMed — https://pubmed.ncbi.nlm.nih.gov/32437550/
- **[S4]** Role of the Choroid Plexus Kir7.1 Channel in the Regulation of Mouse Cerebrospinal Fluid K+ Concentration - PubMed — https://pubmed.ncbi.nlm.nih.gov/41212743/
- **[S5]** 
            A critical role for the inward rectifying potassium channel Kir7.1 in oligodendrocytes of the mouse optic nerve - PMC
         — https://pmc.ncbi.nlm.nih.gov/articles/PMC7166203/
- **[S6]** G-protein-independent coupling of MC4R to Kir7.1 in hypothalamic neurons - PubMed — https://pubmed.ncbi.nlm.nih.gov/25600267/
- **[S7]** Retinal Development and Pathophysiology in Kcnj13 Knockout Mice - PubMed — https://pubmed.ncbi.nlm.nih.gov/35096838/
- **[S8]** 
            Glial and neuronal expression of the Inward Rectifying Potassium Channel Kir7.1 in the adult mouse brain - PMC
         — https://pmc.ncbi.nlm.nih.gov/articles/PMC6794205/
- **[S9]** Role of Kir7.1 K+ channel in retinal pigment epithelium probed in a Kir7.1-M125R-expressing mutant mouse - PubMed — https://pubmed.ncbi.nlm.nih.gov/41247777/
- **[S10]** Recessive mutations in KCNJ13, encoding an inwardly rectifying potassium channel subunit, cause leber congenital amaurosis - PubMed — https://pubmed.ncbi.nlm.nih.gov/21763485/?utm_source=openai
- **[S11]** 
            Focus on Kir7.1: physiology and channelopathy - PMC
         — https://pmc.ncbi.nlm.nih.gov/articles/PMC4594557/
- **[S12]** 
            Conditional Loss of Kcnj13 in the Retinal Pigment Epithelium Causes Photoreceptor Degeneration - PMC
         — https://pmc.ncbi.nlm.nih.gov/articles/PMC6707361/
- **[S13]** Role of the Choroid Plexus Kir7.1 Channel in the Regulation of Mouse Cerebrospinal Fluid K+ Concentration - PubMed — https://pubmed.ncbi.nlm.nih.gov/41212743/
- **[S14]** 
            Conditional Loss of Kcnj13 in the Retinal Pigment Epithelium Causes Photoreceptor Degeneration - PMC
         — https://pmc.ncbi.nlm.nih.gov/articles/PMC6707361/
