## Supplementary material for "Understanding Substance Dependence: What Differentiates Addictive from Non-Addictive Drugs?": Folder for supplementary files: non-addictive-genes-claims-first100-093-slco2b1-gene-protein-has-been-associated-with-regu-aae31c2459fe.md

# non-addictive-genes-claims-first100-093-aae31c2459fe

**Original claim:** SLCO2B1 gene/protein has been associated with regulatory or homeostatic biological processes in neural systems.

## Claim

SLCO2B1 gene/protein has been associated with regulatory or homeostatic biological processes in neural systems. ([pubmed.ncbi.nlm.nih.gov](https://pubmed.ncbi.nlm.nih.gov/25132355/)) [S1]

## Verdict

Verdict: Likely True

## Confidence

Medium. ([pubmed.ncbi.nlm.nih.gov](https://pubmed.ncbi.nlm.nih.gov/25132355/)) [S2]

## Supporting evidence

There is solid evidence that **SLCO2B1/OATP2B1 is present in neural barrier and neural-immune compartments**, which are central to CNS homeostasis. In human tissue, OATP2B1 was localized to **endothelial cells of brain capillaries** in cortex, cerebellum, and hippocampus, supporting a role at the blood-brain barrier (BBB). Independent transcriptomic and proteomic studies found **SLCO2B1 enriched in human brain microvessels**, and a recent human BBB proteomics study reported that **SLCO2B1 is the predominantly expressed SLCO transporter in the adult human BBB**. ([pubmed.ncbi.nlm.nih.gov](https://pubmed.ncbi.nlm.nih.gov/25132355/)) [S3]

The most direct “homeostasis-like” evidence comes from **mouse microglia**. A 2022 study identified Slco2b1 as a **heme importer** that increases cellular iron availability. In brain single-cell datasets, Slco2b1 expression was described as **highly restricted to microglia**; primary microglia from Slco2b1-knockout mice showed a marked loss of heme-analog uptake. Importantly, RNA-seq of knockout microglia showed enrichment of **inflammation-related gene ontology terms** and upregulation of disease-associated microglial markers such as **Apoe** and **Clec7a**, linking Slco2b1 to microglial heme/iron handling and inflammatory state regulation. Since microglia are core cells for maintenance of the neural environment, this is strong evidence that Slco2b1 is associated with neural homeostatic biology, at least in mice. ([pmc.ncbi.nlm.nih.gov](https://pmc.ncbi.nlm.nih.gov/articles/PMC9356996/)) [S4]

There is also evidence connecting OATP2B1 to **transport of neuroactive substrates** relevant to CNS regulation. Human OATP2B1 transports sulfated steroids including **DHEAS** and **pregnenolone sulfate** in heterologous systems, and brain-focused reviews identify OATP2B1 as one of the most plausible transporters involved in **neurosteroid traffic across the BBB**. Neurosteroid transport is biologically relevant because these compounds influence neuronal excitability, development, cognition, and stress-related homeostasis. ([pubmed.ncbi.nlm.nih.gov](https://pubmed.ncbi.nlm.nih.gov/16908597/?utm_source=openai)) [S5]

Older BBB work also supports involvement of OATP-family transporters in neuroactive steroid handling. In a mouse BBB model, **DHEAS efflux across the BBB** was demonstrated, and uptake into brain endothelial cells was inhibited by digoxin, implicating an **oatp2-mediated** component. Separately, rat BBB perfusion studies showed rapid uptake and metabolism of **DHEAS and pregnenolone sulfate** at the BBB, with the authors concluding that circulating steroid sulfates can serve as hormone precursors at the BBB, with implications for **neurogenesis, neuronal survival, learning, and memory**. While these studies do not prove human OATP2B1 is the sole physiological transporter, they support a broader association between OATP-family steroid transport and neural regulatory biology. ([pubmed.ncbi.nlm.nih.gov](https://pubmed.ncbi.nlm.nih.gov/11032880/)) [S6]

A further neural-system link comes from retina, which is neural tissue. In the 2015 human retina/brain study, OATP2B1 and OATP1A2 were investigated as transporters for **substance P** and **vasoactive intestinal peptide (VIP)**. The authors concluded that expression of these transporters in retinal neurons implies a role in **reuptake of peptide neurotransmitters**, and that the high expression of OATP2B1 in brain capillaries supports an important role in CNS substance penetration across the BBB. This is not definitive proof of a homeostatic role for OATP2B1 in the brain parenchyma, but it is a published association with neural signaling-regulatory substrates. ([pubmed.ncbi.nlm.nih.gov](https://pubmed.ncbi.nlm.nih.gov/25132355/)) [S7]

## Contradicting or limiting evidence

The main limitation is that much of the evidence is **indirect or compartment-specific**. For the human CNS, the strongest data are mostly about **expression/localization at the BBB** and transporter capability, rather than direct demonstration that SLCO2B1 regulates a specific neural homeostatic phenotype in vivo. Several reviews phrase its role cautiously as “may be involved” in transport of neurosteroids or neuropeptides across the BBB. ([pmc.ncbi.nlm.nih.gov](https://pmc.ncbi.nlm.nih.gov/articles/PMC8038419/?utm_source=openai)) [S8]

A notable recent challenge concerns **pregnenolone sulfate**. A 2025 study using humanized/knockout rats and an in vitro human BBB model reported that **PregS permeation at the BBB was independent of OATP2B1**, concluding that OATP2B1 does not mediate PregS uptake across the BBB in that system. This does not negate all neural-homeostasis associations for SLCO2B1, but it weakens one specific proposed mechanism for neurosteroid homeostasis. ([pubmed.ncbi.nlm.nih.gov](https://pubmed.ncbi.nlm.nih.gov/40070341/)) [S9]

The BBB expression literature has also had some **methodological inconsistency**. Earlier quantitative human BBB proteomics found many OATP-family transporters below quantification in humans, whereas later, more sensitive or differently designed studies detected and quantified SLCO2B1 and even identified it as the predominant SLCO transporter in adult human BBB. That pattern suggests the signal is probably real, but it also means the exact abundance and physiological weight of SLCO2B1 at the BBB have not been entirely settled across platforms. ([pubmed.ncbi.nlm.nih.gov](https://pubmed.ncbi.nlm.nih.gov/21291474/?utm_source=openai)) [S10]

Finally, the strongest direct homeostatic evidence—microglial heme uptake and inflammatory-state effects after Slco2b1 deletion—is from **mouse**, not human, tissue. Extrapolation to human neural homeostasis is plausible but still an inference. ([pmc.ncbi.nlm.nih.gov](https://pmc.ncbi.nlm.nih.gov/articles/PMC9356996/)) [S11]

## Mechanistic / biological rationale

A biologically coherent rationale exists for why SLCO2B1 could participate in neural homeostasis. First, at the **BBB**, transporters help maintain the CNS milieu by controlling exchange of endogenous molecules between blood and brain. Because OATP2B1 is enriched in brain microvessels and can transport **steroid sulfates** and other organic anions, it is well positioned to influence the local availability of neuroactive compounds at this interface. ([pmc.ncbi.nlm.nih.gov](https://pmc.ncbi.nlm.nih.gov/articles/PMC3906042/)) [S12]

Second, the relevant substrates themselves have neural regulatory functions. **DHEAS** and **pregnenolone sulfate** are neuroactive steroids implicated in cognition, neurogenesis, neuronal excitability, and stress-related homeostasis; rat BBB studies further suggest that steroid sulfates can be converted at the BBB into free steroids that may act as hormone precursors for the brain. Thus, even if OATP2B1 is only one part of a larger transport/metabolism network, association with these substrates gives a plausible route by which SLCO2B1 could contribute to neural regulatory processes. ([pubmed.ncbi.nlm.nih.gov](https://pubmed.ncbi.nlm.nih.gov/28665486/)) [S13]

Third, in **microglia**, Slco2b1-mediated heme uptake offers a direct homeostasis mechanism. Heme and iron handling are tightly linked to oxidative stress and inflammatory tone. In the 2022 study, Slco2b1 loss reduced microglial heme uptake and shifted microglial transcription toward inflammatory programs; the authors explicitly proposed that microglial heme uptake through SLCO2B1 could help protect neurons and astrocytes from oxidative heme damage under conditions such as intracranial hemorrhage. That is a credible homeostatic model, though still not fully proven in disease physiology. ([pmc.ncbi.nlm.nih.gov](https://pmc.ncbi.nlm.nih.gov/articles/PMC9356996/)) [S14]

## Bottom line

Overall, the claim is **likely true**. SLCO2B1 is credibly associated with neural-system regulatory/homeostatic biology through: **(1)** expression in human brain capillary endothelium, **(2)** proposed transport of neuroactive steroid and peptide substrates at neural barriers, and **(3)** especially compelling mouse evidence that Slco2b1 regulates **microglial heme/iron handling and inflammatory state**, which are core components of neural homeostasis. However, the evidence is uneven: the strongest causal data are in mouse microglia, while human CNS evidence is more often expression- and transport-based than phenotype-based, and at least one recent study disputes a specific proposed role in pregnenolone sulfate BBB transport. ([pubmed.ncbi.nlm.nih.gov](https://pubmed.ncbi.nlm.nih.gov/25132355/)) [S15]

## Cited sources

- **[S1]** Differential cellular expression of organic anion transporting peptides OATP1A2 and OATP2B1 in the human retina and brain: implications for carrier-mediated transport of neuropeptides and neurosteriods in the CNS - PubMed — https://pubmed.ncbi.nlm.nih.gov/25132355/
- **[S2]** Differential cellular expression of organic anion transporting peptides OATP1A2 and OATP2B1 in the human retina and brain: implications for carrier-mediated transport of neuropeptides and neurosteriods in the CNS - PubMed — https://pubmed.ncbi.nlm.nih.gov/25132355/
- **[S3]** Differential cellular expression of organic anion transporting peptides OATP1A2 and OATP2B1 in the human retina and brain: implications for carrier-mediated transport of neuropeptides and neurosteriods in the CNS - PubMed — https://pubmed.ncbi.nlm.nih.gov/25132355/
- **[S4]** 
            Metabolic-scale gene activation screens identify SLCO2B1 as a heme transporter that enhances cellular iron availability - PMC
         — https://pmc.ncbi.nlm.nih.gov/articles/PMC9356996/
- **[S5]** Modification of OATP2B1-mediated transport by steroid hormones - PubMed — https://pubmed.ncbi.nlm.nih.gov/16908597/?utm_source=openai
- **[S6]** Blood-brain barrier is involved in the efflux transport of a neuroactive steroid, dehydroepiandrosterone sulfate, via organic anion transporting polypeptide 2 - PubMed — https://pubmed.ncbi.nlm.nih.gov/11032880/
- **[S7]** Differential cellular expression of organic anion transporting peptides OATP1A2 and OATP2B1 in the human retina and brain: implications for carrier-mediated transport of neuropeptides and neurosteriods in the CNS - PubMed — https://pubmed.ncbi.nlm.nih.gov/25132355/
- **[S8]** Disease-Induced Modulation of Drug Transporters at the Blood–Brain Barrier Level - PMC — https://pmc.ncbi.nlm.nih.gov/articles/PMC8038419/?utm_source=openai
- **[S9]** Pregnenolone Sulfate Permeation at the Blood-Brain Barrier is Independent of OATP2B1-In Vivo and In Vitro Insights - PubMed — https://pubmed.ncbi.nlm.nih.gov/40070341/
- **[S10]** Quantitative targeted absolute proteomics of human blood-brain barrier transporters and receptors - PubMed — https://pubmed.ncbi.nlm.nih.gov/21291474/?utm_source=openai
- **[S11]** 
            Metabolic-scale gene activation screens identify SLCO2B1 as a heme transporter that enhances cellular iron availability - PMC
         — https://pmc.ncbi.nlm.nih.gov/articles/PMC9356996/
- **[S12]** 
            Profiling Solute Carrier Transporters in the Human Blood-Brain Barrier - PMC
         — https://pmc.ncbi.nlm.nih.gov/articles/PMC3906042/
- **[S13]** Uptake and metabolism of sulphated steroids by the blood-brain barrier in the adult male rat - PubMed — https://pubmed.ncbi.nlm.nih.gov/28665486/
- **[S14]** 
            Metabolic-scale gene activation screens identify SLCO2B1 as a heme transporter that enhances cellular iron availability - PMC
         — https://pmc.ncbi.nlm.nih.gov/articles/PMC9356996/
- **[S15]** Differential cellular expression of organic anion transporting peptides OATP1A2 and OATP2B1 in the human retina and brain: implications for carrier-mediated transport of neuropeptides and neurosteriods in the CNS - PubMed — https://pubmed.ncbi.nlm.nih.gov/25132355/
