## Supplementary material for "Understanding Substance Dependence: What Differentiates Addictive from Non-Addictive Drugs?": Folder for supplementary files: non-addictive-genes-claims-first100-094-cd47-gene-protein-has-been-associated-with-regulat-2dd59f721651.md

# non-addictive-genes-claims-first100-094-2dd59f721651

**Original claim:** CD47 gene/protein has been associated with regulatory or homeostatic biological processes in neural systems.

## Claim

CD47 gene/protein has been associated with regulatory or homeostatic biological processes in neural systems. ([pmc.ncbi.nlm.nih.gov](https://pmc.ncbi.nlm.nih.gov/articles/PMC8271744/)) [S1]

## Verdict

Verdict: Highly Supported

Multiple independent peer-reviewed studies and reviews support that CD47 is involved in neural regulatory/homeostatic processes, especially by limiting microglia-mediated phagocytosis of synapses and myelin, thereby helping maintain circuit integrity and tissue balance in the CNS. The strongest evidence is preclinical (mouse, monkey, and cell-culture), with additional human relevance from expression data and disease-tissue studies. ([pubmed.ncbi.nlm.nih.gov](https://pubmed.ncbi.nlm.nih.gov/30308165/)) [S2]

## Confidence

High

## Supporting evidence

CD47 is expressed in the nervous system, including neurons, and at lower levels in oligodendrocytes; a recent human MS pathology paper cites Human Protein Atlas and Brain RNA-seq data showing high neuronal CD47 expression. A 2024 review also notes that CD47 isoforms 3 and 4 are expressed predominantly in neural tissue. ([pmc.ncbi.nlm.nih.gov](https://pmc.ncbi.nlm.nih.gov/articles/PMC11339711/)) [S3]

The best direct evidence for a homeostatic neural role comes from developmental synaptic pruning. In the mouse retinogeniculate system, Lehrman et al. showed that CD47 protects synapses from excess microglia-mediated pruning during development. CD47 was enriched during the pruning period, while SIRPα was primarily expressed by microglia at P5-P10; deletion of CD47 disrupted normal activity-dependent engulfment and caused persistent synapse loss/overpruning. ([pubmed.ncbi.nlm.nih.gov](https://pubmed.ncbi.nlm.nih.gov/30308165/)) [S4]

This finding has been reinforced by later work. A Nature Communications study reported that loss of microglial SIRPα promoted synaptic pruning in preclinical neurodegeneration models and increased synaptic loss and cognitive impairment; the same study also found reduced synaptic CD47 on less-active neurons, consistent with CD47 acting as a protective “don’t eat me” signal for synapses. ([pubmed.ncbi.nlm.nih.gov](https://pubmed.ncbi.nlm.nih.gov/33795678/)) [S5]

High-level reviews now treat CD47-SIRPα as part of the machinery that preserves synaptic homeostasis. An Annual Review of Neuroscience article states that CD47 on neuronal membranes signals to microglial SIRPα to suppress phagocytosis of synaptic material, and that loss of either protein causes excess engulfment and overpruning. A recent review of microglial phagocytic mechanisms makes the same point and connects this developmental mechanism to disease. ([annualreviews.org](https://www.annualreviews.org/doi/pdf/10.1146/annurev-neuro-070918-050306)) [S6]

CD47 is also associated with white-matter/myelin homeostasis. In vitro work showed that CD47 on myelin down-regulates myelin phagocytosis by microglia and macrophages via SIRPα; blocking CD47 or SIRPα increased myelin engulfment. This is a clear regulatory function in neural tissue maintenance, even if context-dependent. ([pubmed.ncbi.nlm.nih.gov](https://pubmed.ncbi.nlm.nih.gov/21401967/)) [S7]

Human disease-tissue studies are consistent with this homeostatic interpretation. In cortical tissue from multiple sclerosis cases, investigators explicitly stated that control of microglial activity through CD47-SIRPα has been implicated in brain homeostasis, and they examined altered expression of this axis in MS gray and white matter. ([pubmed.ncbi.nlm.nih.gov](https://pubmed.ncbi.nlm.nih.gov/39175944/)) [S8]

Aging-primate data also support a neural homeostatic role. In aged monkey cortex, reduced CD47-PSD95 colocalization and reduced neuronal CD47 expression were associated with patterns consistent with increased vulnerability to microglia-mediated synapse loss and cognitive impairment. That is still associative, but it extends the CD47-homeostasis link beyond rodents and development. ([pubmed.ncbi.nlm.nih.gov](https://pubmed.ncbi.nlm.nih.gov/37989825/)) [S9]

Beyond phagocytic regulation, older neuronal studies suggest CD47 can regulate neuronal development directly. In cultured cortical neurons, CD47 overexpression enhanced dendritic outgrowth, synaptic protein expression, glutamate release, and calcium oscillations, while knockdown reduced MAPK activity and synaptic protein levels. In hippocampal neurons, CD47 loss impaired dendritic and axonal development, and rescue/overexpression restored branching through Rac/Cdc42-related signaling. These are developmental-regulatory rather than classic “homeostasis” findings, but they still support the claim’s broader wording about regulatory biological processes in neural systems. ([pubmed.ncbi.nlm.nih.gov](https://pubmed.ncbi.nlm.nih.gov/15297459/)) [S10]

## Contradicting or limiting evidence

The strongest limitation is that most evidence is preclinical. Much of the causal evidence comes from mouse knockouts, viral overexpression, or cultured neurons/microglia, while human evidence is mostly expression-based or pathological association rather than direct intervention. ([pubmed.ncbi.nlm.nih.gov](https://pubmed.ncbi.nlm.nih.gov/30308165/)) [S11]

CD47’s neural role is also clearly context-dependent, not uniformly protective. In intracerebral hemorrhage models, blocking CD47 accelerated hematoma clearance by microglia/macrophages and reduced brain swelling, neuronal loss, and neurological deficits. Similarly, CD47 knockout improved recovery after spinal cord injury. These findings suggest that while CD47 can preserve intact neural structures, it can also impede beneficial debris clearance after injury. ([pubmed.ncbi.nlm.nih.gov](https://pubmed.ncbi.nlm.nih.gov/31084334/)) [S12]

There is also evidence that both insufficient and excessive CD47-SIRPα signaling may be harmful in some pathological settings. In a mouse model of perioperative neurocognitive disorder, reduced CD47/SIRPα was linked to excessive synapse engulfment, yet experimental SIRPα overexpression also failed to rescue cognition and could worsen total engulfment through increased microglial burden; the authors concluded that both too low and too high levels could be detrimental. ([frontiersin.org](https://www.frontiersin.org/journals/neuroscience/articles/10.3389/fnins.2022.788675/full)) [S13]

Some older cell-culture work even found that forced neuronal CD47 expression could induce caspase-dependent apoptosis in cultured cortical neurons under certain conditions, indicating that CD47 signaling is not simply homeostatic in all neural contexts. ([pubmed.ncbi.nlm.nih.gov](https://pubmed.ncbi.nlm.nih.gov/12124426/)) [S14]

So the evidence does **not** support a simplistic claim that CD47 is always beneficial in the nervous system. Rather, it supports that CD47 is a regulator of neural homeostasis and remodeling whose effects depend on cell type, ligand/receptor pairing, developmental stage, and injury/disease context. ([pmc.ncbi.nlm.nih.gov](https://pmc.ncbi.nlm.nih.gov/articles/PMC8271744/)) [S15]

## Mechanistic / biological rationale

The clearest mechanism is the canonical CD47-SIRPα “don’t eat me” pathway. In neural tissue, neuronal or myelin CD47 binds SIRPα on microglia/macrophages, suppressing phagocytosis. In practice, this acts as a brake on elimination of synaptic material and myelin, helping preserve appropriate synapses and avoid unnecessary engulfment. ([annualreviews.org](https://www.annualreviews.org/doi/pdf/10.1146/annurev-neuro-070918-050306)) [S16]

This is biologically plausible as a homeostatic counterweight to “eat me” systems such as complement. Reviews of neuroimmune crosstalk and microglial pruning describe CD47 alongside other neuronal signals that restrain microglial activation and phagocytosis, thereby contributing to brain immune balance and synaptic stability. ([jneuroinflammation.biomedcentral.com](https://jneuroinflammation.biomedcentral.com/articles/10.1186/1742-2094-9-155)) [S17]

CD47 also has noncanonical, cell-intrinsic signaling roles. A 2024 review summarizes links to PI3K/AKT, MAPK/ERK, nitric oxide, stress-response, apoptosis, and metabolic pathways, and notes that neural-predominant isoforms exist. Older neuronal studies are consistent with these signaling effects, showing CD47-dependent regulation of MAPK, Rac, and Cdc42 pathways during neurite and dendrite development. ([nrf.com.au](https://www.nrf.com.au/uploads/documents/Publications/BT/Cell%20autonomous%20functions%20of%20CD47%20in%20regulating%20cellular%20plasticity%20and%20metabolic%20plasticity.pdf)) [S18]

Taken together, CD47 fits well as a regulator of neural homeostasis in two senses: first, as an intercellular checkpoint limiting inappropriate microglial engulfment; second, as a cell-intrinsic modulator of neuronal growth/survival programs. The first role is much better established than the second. ([pubmed.ncbi.nlm.nih.gov](https://pubmed.ncbi.nlm.nih.gov/30308165/)) [S19]

## Bottom line

The claim is well supported. CD47 is convincingly associated with regulatory and homeostatic processes in neural systems, most strongly through CD47-SIRPα signaling that restrains microglial pruning of synapses and phagocytosis of myelin. There is also supporting evidence for roles in neuronal development and signaling. The main caveat is that the evidence is mostly from animal and cell studies, and CD47’s effects are context-dependent: what preserves healthy synapses or myelin can, after injury, also slow beneficial debris clearance. ([pubmed.ncbi.nlm.nih.gov](https://pubmed.ncbi.nlm.nih.gov/30308165/)) [S20]

## Cited sources

- **[S1]** 
            CD47 in the Brain and Neurodegeneration: An Update on the Role in Neuroinflammatory Pathways - PMC
         — https://pmc.ncbi.nlm.nih.gov/articles/PMC8271744/
- **[S2]** CD47 Protects Synapses from Excess Microglia-Mediated Pruning during Development - PubMed — https://pubmed.ncbi.nlm.nih.gov/30308165/
- **[S3]** 
            Cortical CD200–CD200R and CD47–SIRPα expression is associated with multiple sclerosis pathology - PMC
         — https://pmc.ncbi.nlm.nih.gov/articles/PMC11339711/
- **[S4]** CD47 Protects Synapses from Excess Microglia-Mediated Pruning during Development - PubMed — https://pubmed.ncbi.nlm.nih.gov/30308165/
- **[S5]** Loss of microglial SIRPα promotes synaptic pruning in preclinical models of neurodegeneration - PubMed — https://pubmed.ncbi.nlm.nih.gov/33795678/
- **[S6]** Untitled source — https://www.annualreviews.org/doi/pdf/10.1146/annurev-neuro-070918-050306
- **[S7]** Myelin down-regulates myelin phagocytosis by microglia and macrophages through interactions between CD47 on myelin and SIRPα (signal regulatory protein-α) on phagocytes - PubMed — https://pubmed.ncbi.nlm.nih.gov/21401967/
- **[S8]** Cortical CD200-CD200R and CD47-SIRPα expression is associated with multiple sclerosis pathology - PubMed — https://pubmed.ncbi.nlm.nih.gov/39175944/
- **[S9]** Immune proteins C1q and CD47 may contribute to aberrant microglia-mediated synapse loss in the aging monkey brain that is associated with cognitive impairment - PubMed — https://pubmed.ncbi.nlm.nih.gov/37989825/
- **[S10]** Neuronal roles of the integrin-associated protein (IAP/CD47) in developing cortical neurons - PubMed — https://pubmed.ncbi.nlm.nih.gov/15297459/
- **[S11]** CD47 Protects Synapses from Excess Microglia-Mediated Pruning during Development - PubMed — https://pubmed.ncbi.nlm.nih.gov/30308165/
- **[S12]** Enhancement of Hematoma Clearance With CD47 Blocking Antibody in Experimental Intracerebral Hemorrhage - PubMed — https://pubmed.ncbi.nlm.nih.gov/31084334/
- **[S13]** Frontiers | Anomalous Levels of CD47/Signal Regulatory Protein Alpha in the Hippocampus Lead to Excess Microglial Engulfment in Mouse Model of Perioperative Neurocognitive Disorders — https://www.frontiersin.org/journals/neuroscience/articles/10.3389/fnins.2022.788675/full
- **[S14]** Expression of CD47/integrin-associated protein induces death of cultured cerebral cortical neurons - PubMed — https://pubmed.ncbi.nlm.nih.gov/12124426/
- **[S15]** 
            CD47 in the Brain and Neurodegeneration: An Update on the Role in Neuroinflammatory Pathways - PMC
         — https://pmc.ncbi.nlm.nih.gov/articles/PMC8271744/
- **[S16]** Untitled source — https://www.annualreviews.org/doi/pdf/10.1146/annurev-neuro-070918-050306
- **[S17]** Neuroimmune crosstalk in the central nervous system and its significance for neurological diseases | Journal of Neuroinflammation | Springer Nature Link — https://jneuroinflammation.biomedcentral.com/articles/10.1186/1742-2094-9-155
- **[S18]** Cell autonomous functions of CD47 in regulating cellular plasticity and metabolic plasticity — https://www.nrf.com.au/uploads/documents/Publications/BT/Cell%20autonomous%20functions%20of%20CD47%20in%20regulating%20cellular%20plasticity%20and%20metabolic%20plasticity.pdf
- **[S19]** CD47 Protects Synapses from Excess Microglia-Mediated Pruning during Development - PubMed — https://pubmed.ncbi.nlm.nih.gov/30308165/
- **[S20]** CD47 Protects Synapses from Excess Microglia-Mediated Pruning during Development - PubMed — https://pubmed.ncbi.nlm.nih.gov/30308165/
