## Supplementary material for "Understanding Substance Dependence: What Differentiates Addictive from Non-Addictive Drugs?": Folder for supplementary files: non-addictive-genes-claims-first100-095-hbs1l-gene-protein-has-been-associated-with-regula-33ce0cdeb1c1.md

# non-addictive-genes-claims-first100-095-33ce0cdeb1c1

**Original claim:** HBS1L gene/protein has been associated with regulatory or homeostatic biological processes in neural systems.

## Claim

**Claim:** *HBS1L gene/protein has been associated with regulatory or homeostatic biological processes in neural systems.*

## Verdict

Verdict: Likely True

HBS1L is **not primarily known as a neuron-specific regulator**, but there is credible evidence that it is associated with **regulatory processes in neural tissues**, especially **cerebellar development** and the **retina/visual system**, through its role in **ribosome rescue and mRNA surveillance**. The evidence for **adult neural homeostasis** is more limited and less direct than the evidence for **neurodevelopmental and visual-system regulation**. ([pmc.ncbi.nlm.nih.gov](https://pmc.ncbi.nlm.nih.gov/articles/PMC8075583/)) [S1]

## Confidence

**Medium.**

The association is supported by multiple mechanistic and animal studies plus rare human genetic observations, but the human evidence base is still small and the strongest data are concentrated in **developmental** and **retinal/visual** contexts rather than broad, well-established nervous-system homeostasis in adults. ([academic.oup.com](https://academic.oup.com/biomedgerontology/article/80/4/glae304/7932361)) [S2]

## Supporting evidence

HBS1L encodes a **translational GTPase** involved in **rescue of stalled ribosomes** and **no-go decay / mRNA surveillance**, which are core regulatory processes controlling translation quality and proteostasis. Authoritative gene ontology annotations list Hbs1l as involved in **“nuclear-transcribed mRNA catabolic process, no-go decay”** and **“rescue of stalled ribosome,”** and mechanistic reviews describe the HBS1L-PELO complex as promoting dissociation of stalled ribosomes and thereby enabling downstream quality-control pathways. ([informatics.jax.org](https://www.informatics.jax.org/go/marker/MGI%3A1891704?utm_source=openai)) [S3]

In the nervous system specifically, a strong mouse study found that **Hbs1l is expressed in multiple brain cell types** and that **conditional deletion during cerebellar development** disrupts normal neurodevelopment. In that model, Hbs1l loss impaired **cell-cycle exit** of ventricular-zone progenitors, reduced formation of **Purkinje cells**, **interneuron precursors**, **rhombic-lip derivatives**, and **oligodendroglial progenitors**, leading the authors to conclude that Hbs1l is required for generation of multiple cerebellar cell types. That is a direct association with a **regulatory biological process in a neural system**. ([pmc.ncbi.nlm.nih.gov](https://pmc.ncbi.nlm.nih.gov/articles/PMC8075583/)) [S4]

Human genetic observations also point in the same direction. Biallelic pathogenic HBS1L variants have been reported in a child with **growth restriction, neurodevelopmental delay, hypotonia, microcephaly, and retinal abnormalities**, and follow-up ophthalmologic work showed impaired retinal function with abnormal rod- and cone-mediated responses. A review of ribosome-associated quality control and neurologic disease likewise summarizes HBS1L variants as an ultra-rare cause of neurodevelopmental phenotypes with retinal involvement. ([pubmed.ncbi.nlm.nih.gov](https://pubmed.ncbi.nlm.nih.gov/30707697/)) [S5]

The retina provides especially relevant evidence because it is neural tissue. In Hbs1l-deficient mice, investigators observed **retinal thinning**, **photoreceptor apoptosis**, and broad proteomic changes, with downregulated proteins enriched for **phototransduction**, **cilium assembly**, and **photoreceptor cell development**. The same work showed Hbs1l is broadly expressed during retinal development and relatively enriched in photoreceptors/bipolar cells at certain stages. These findings link HBS1L to regulation of protein-expression programs needed for retinal neural development and function. ([pubmed.ncbi.nlm.nih.gov](https://pubmed.ncbi.nlm.nih.gov/38966981/)) [S6]

An additional recent Drosophila study provides a particularly clean mechanistic neural example: loss of the HBS1L homolog **Hbs1** reduced **ATF4** expression, caused **vision defects** on electroretinography, and produced lamina-layer abnormalities consistent with defective synapse formation. Depleting Hbs1 specifically in **lamina neurons** reproduced the phenotype, and restoring ATF4 partially rescued it. The authors concluded that Hbs1-Pelo-mediated **translation regulation of ATF4 in neurons** is required for visual-system development and function. That is direct evidence of HBS1L-family involvement in a **regulatory neural process**. ([pubmed.ncbi.nlm.nih.gov](https://pubmed.ncbi.nlm.nih.gov/41279977/)) [S7]

## Contradicting or limiting evidence

The main limitation is that **direct human evidence is sparse**. The currently cited human literature involves a **very small number of affected individuals**, and one review explicitly notes that the full physiological function of HBS1L remains incompletely understood and that further work is needed to define downstream consequences of dysfunction. So the claim is better supported as an **association** than as a settled, broad principle of neural biology. ([academic.oup.com](https://academic.oup.com/biomedgerontology/article/80/4/glae304/7932361)) [S8]

Also, the neural phenotype appears to be **context- and stage-specific**, not universal across all neurons. In the cerebellum, Hbs1l was required for **initial production** of granule-cell precursors, but was reported to be **dispensable for later cell-cycle progression** of those precursors in the external granule layer. The same study also found that loss of Hbs1l did **not** affect survival of **terminally differentiated granule cells** in older mice, indicating that Hbs1l is not simply a universal maintenance factor for all mature neurons. ([pmc.ncbi.nlm.nih.gov](https://pmc.ncbi.nlm.nih.gov/articles/PMC8075583/)) [S9]

Finally, some “homeostatic” language in the literature comes from **general cellular** or **yeast** contexts rather than direct mammalian neural experiments. For example, retinal investigators summarize prior work linking HBS1/HBS1L to oxidative-stress resilience and endoplasmic-reticulum homeostasis, but those are not equivalent to proving a broad role in **neural homeostasis** across the mammalian CNS. ([pmc.ncbi.nlm.nih.gov](https://pmc.ncbi.nlm.nih.gov/articles/PMC10614867/)) [S10]

## Mechanistic / biological rationale

Biologically, the association makes sense. HBS1L partners with **PELO** and **ABCE1** to detect and resolve **stalled ribosomes**, enabling degradation of faulty mRNAs and aberrant nascent proteins. Reviews describe this as a central part of **ribosome-associated quality control**, a system needed to prevent proteotoxic stress and maintain translation fidelity. Loss of HBS1L function can therefore perturb **proteostasis**, **mRNA surveillance**, and translational regulation. ([pmc.ncbi.nlm.nih.gov](https://pmc.ncbi.nlm.nih.gov/articles/PMC8487456/)) [S11]

Experimental data are consistent with that mechanism. In HBS1L-deficient human and mouse cells, loss of HBS1L is associated with **PELO depletion**, **80S monosome accumulation**, ribosome pausing, and altered translation-related signaling including increased **mTOR/4E-BP** pathway readouts. In the retina, HBS1L deficiency also reduced **EDF1**, a ribosome-collision response factor, alongside many proteins needed for photoreceptor structure and visual signaling. ([pubmed.ncbi.nlm.nih.gov](https://pubmed.ncbi.nlm.nih.gov/30707697/)) [S12]

In neural tissue, such translation-quality-control defects are plausibly consequential because neurons and photoreceptors are highly dependent on tight control of local and global protein synthesis. The Drosophila visual-system study adds a specific example: Hbs1-Pelo promotes correct translation reinitiation of **ATF4**, a stress-responsive transcription factor, and this regulation is required for lamina-neuron function and visual output. The retinal mouse work further argues that photoreceptors may be especially vulnerable because of their heavy demand for continual synthesis and renewal of outer-segment proteins. This last point is partly an inference from the tissue’s biology plus the observed phenotype, but it is well aligned with the published data. ([pubmed.ncbi.nlm.nih.gov](https://pubmed.ncbi.nlm.nih.gov/41279977/)) [S13]

## Bottom line

Overall, the claim is **likely true**. HBS1L has credible links to **regulatory neural processes**, especially **cerebellar neurogenesis** and **retinal/visual-system development and function**, through its core role in **ribosome rescue and translational quality control**. What is **less firmly established** is a broad, well-demonstrated role in **adult neural homeostasis** across the nervous system. So the safest evidence-based wording is: **HBS1L is associated with neural developmental/regulatory processes, and it may contribute to neural homeostatic mechanisms via translational quality control, but the homeostasis claim is currently more limited than the developmental one.** ([pmc.ncbi.nlm.nih.gov](https://pmc.ncbi.nlm.nih.gov/articles/PMC8075583/)) [S14]

## Cited sources

- **[S1]** 
            Defects in translation-dependent quality control pathways lead to convergent molecular and neurodevelopmental pathology - PMC
         — https://pmc.ncbi.nlm.nih.gov/articles/PMC8075583/
- **[S2]** Relationship Between Ribosome-Associated Quality Control and Neurological Disorders | The Journals of Gerontology: Series A | Oxford Academic — https://academic.oup.com/biomedgerontology/article/80/4/glae304/7932361
- **[S3]** Gene Ontology Classifications — https://www.informatics.jax.org/go/marker/MGI%3A1891704?utm_source=openai
- **[S4]** 
            Defects in translation-dependent quality control pathways lead to convergent molecular and neurodevelopmental pathology - PMC
         — https://pmc.ncbi.nlm.nih.gov/articles/PMC8075583/
- **[S5]** Mammalian Hbs1L deficiency causes congenital anomalies and developmental delay associated with Pelota depletion and 80S monosome accumulation - PubMed — https://pubmed.ncbi.nlm.nih.gov/30707697/
- **[S6]** HBS1L deficiency causes retinal dystrophy in a child and in a mouse model associated with defective development of photoreceptor cells - PubMed — https://pubmed.ncbi.nlm.nih.gov/38966981/
- **[S7]** Translation regulation of ATF4 by the termination complex Hbs1-Pelo is required for visual system development and function - PubMed — https://pubmed.ncbi.nlm.nih.gov/41279977/
- **[S8]** Relationship Between Ribosome-Associated Quality Control and Neurological Disorders | The Journals of Gerontology: Series A | Oxford Academic — https://academic.oup.com/biomedgerontology/article/80/4/glae304/7932361
- **[S9]** 
            Defects in translation-dependent quality control pathways lead to convergent molecular and neurodevelopmental pathology - PMC
         — https://pmc.ncbi.nlm.nih.gov/articles/PMC8075583/
- **[S10]** 
            Genetic deficiency of ribosomal rescue factor HBS1L causes retinal dystrophy associated with Pelota and EDF1 depletion - PMC
         — https://pmc.ncbi.nlm.nih.gov/articles/PMC10614867/
- **[S11]** 
            Detecting and Rescuing Stalled Ribosomes - PMC
         — https://pmc.ncbi.nlm.nih.gov/articles/PMC8487456/
- **[S12]** Mammalian Hbs1L deficiency causes congenital anomalies and developmental delay associated with Pelota depletion and 80S monosome accumulation - PubMed — https://pubmed.ncbi.nlm.nih.gov/30707697/
- **[S13]** Translation regulation of ATF4 by the termination complex Hbs1-Pelo is required for visual system development and function - PubMed — https://pubmed.ncbi.nlm.nih.gov/41279977/
- **[S14]** 
            Defects in translation-dependent quality control pathways lead to convergent molecular and neurodevelopmental pathology - PMC
         — https://pmc.ncbi.nlm.nih.gov/articles/PMC8075583/
