## Supplementary material for "Understanding Substance Dependence: What Differentiates Addictive from Non-Addictive Drugs?": Folder for supplementary files: non-addictive-genes-claims-first100-096-arhgef2-gene-protein-has-been-associated-with-regu-cf9aabbedc5e.md

# non-addictive-genes-claims-first100-096-cf9aabbedc5e

**Original claim:** ARHGEF2 gene/protein has been associated with regulatory or homeostatic biological processes in neural systems.

## Claim

ARHGEF2 gene/protein (also called GEF-H1/Lfc) has been associated with regulatory or homeostatic biological processes in neural systems. ([pmc.ncbi.nlm.nih.gov](https://pmc.ncbi.nlm.nih.gov/articles/PMC5428974/)) [S1]

## Verdict

Verdict: Highly Supported

Multiple independent lines of evidence support **regulatory roles** for ARHGEF2 in neural systems, especially in **neurogenesis, mitotic spindle orientation of neural precursors, neuronal migration, neurite/axon remodeling, dendritic spine development, and neural tube morphogenesis**. The evidence for **classic adult neural homeostasis** per se is much thinner, but the claim is phrased broadly as “regulatory or homeostatic,” and the regulatory component is well supported. ([pmc.ncbi.nlm.nih.gov](https://pmc.ncbi.nlm.nih.gov/articles/PMC5428974/)) [S2]

## Confidence

High

The claim is broad and is backed by human genetics, mouse studies, Xenopus developmental work, and cultured-neuron experiments. Confidence is high for **regulatory roles in neural biology**, but not specifically for **homeostatic maintenance in the mature nervous system**. ([pubmed.ncbi.nlm.nih.gov](https://pubmed.ncbi.nlm.nih.gov/28453519/?utm_source=openai)) [S3]

## Supporting evidence

- **Human neurodevelopmental disease evidence:** A homozygous loss-of-function ARHGEF2 mutation was reported in affected children with **intellectual disability, mild microcephaly, and midbrain-hindbrain malformation**. In the same study, ARHGEF2 loss perturbed **progenitor differentiation**, altered **mitotic spindle orientation**, reduced **RhoA/ROCK/MLC signaling**, and disrupted **precerebellar neuron migration**, directly tying ARHGEF2 to regulated neural developmental processes. ([pubmed.ncbi.nlm.nih.gov](https://pubmed.ncbi.nlm.nih.gov/28453519/?utm_source=openai)) [S4]

- **Expression and function in neural precursors:** In mouse neocortex, ARHGEF2 was reported to be expressed in **neural precursors and immature neurons**. Its downregulation kept radial precursors cycling and inhibited neurogenesis, consistent with a role in regulating the balance between progenitor maintenance and neuronal differentiation. ([pmc.ncbi.nlm.nih.gov](https://pmc.ncbi.nlm.nih.gov/articles/PMC5428974/)) [S5]

- **Cortical differentiation mechanism:** In a CRISPR knockout mouse model, loss of Arhgef2 reduced brain size and decreased cortical **Mettl14** expression and total **m6A RNA methylation**. The downregulated m6A-marked transcripts were enriched for biological processes including **neurogenesis** and **synapse organization**, and included **Npdc1** and **Cend1**, genes linked to neural differentiation and cell-cycle exit. Rescue experiments supported this pathway functionally. ([pubmed.ncbi.nlm.nih.gov](https://pubmed.ncbi.nlm.nih.gov/34142067/?utm_source=openai)) [S6]

- **Dendritic spine regulation:** In cultured neurons, GEF-H1 localized to **postsynaptic density/spines** and regulated **dendritic spine density and length**. Knockdown decreased neuronal RhoA activity, increased spine density/length, and pharmacologic inhibition of **RhoA/ROCK** blocked the effects of GEF-H1 overexpression, showing that ARHGEF2 regulates synaptic structural development through a defined signaling cascade. ([pmc.ncbi.nlm.nih.gov](https://pmc.ncbi.nlm.nih.gov/articles/PMC2638734/)) [S7]

- **Neurite/axon remodeling and neuronal polarity:** In developing neurons, GEF-H1 knockdown abolished **NT-3-induced minor neurite retraction** and the associated increase in **RhoA activity**, supporting a role in the regulation of neurite remodeling and polarity-related signaling. ([pmc.ncbi.nlm.nih.gov](https://pmc.ncbi.nlm.nih.gov/articles/PMC5484694/)) [S8]

- **Neural tube morphogenesis:** In Xenopus embryos, GEF-H1 depletion interfered with **cell intercalation** and **apical constriction** in neuroectoderm, producing **neural tube defects**. This places ARHGEF2 in regulated morphogenetic programs of the developing nervous system. ([pmc.ncbi.nlm.nih.gov](https://pmc.ncbi.nlm.nih.gov/articles/PMC4038946/)) [S9]

Taken together, these data show that ARHGEF2 is not merely expressed in neural tissue; it is functionally implicated in several **regulated neural-system processes** across development and neuronal structural organization. ([pmc.ncbi.nlm.nih.gov](https://pmc.ncbi.nlm.nih.gov/articles/PMC5428974/)) [S10]

## Contradicting or limiting evidence

- The strongest evidence is for **developmental and morphoregulatory functions**, not for “homeostasis” in the narrower sense of maintaining stable neural activity, synaptic scaling, or adult circuit set points. I did not find strong direct evidence that ARHGEF2 is a canonical mediator of mature neural homeostasis. ([pmc.ncbi.nlm.nih.gov](https://pmc.ncbi.nlm.nih.gov/articles/PMC8185223/)) [S11]

- Some neural findings are **model-specific**. For example, Xenopus knockdown caused neural tube closure defects, but the Xenopus study itself notes that **mice lacking GEF-H1/Lfc reportedly do not show obvious neural tube abnormalities**, suggesting functional redundancy or species/context dependence. ([pmc.ncbi.nlm.nih.gov](https://pmc.ncbi.nlm.nih.gov/articles/PMC4038946/)) [S12]

- Human evidence is important but still **limited in scope**, centering largely on a rare recessive neurodevelopmental disorder rather than broad epidemiologic or clinical evidence across common neurologic conditions. ([pubmed.ncbi.nlm.nih.gov](https://pubmed.ncbi.nlm.nih.gov/28453519/?utm_source=openai)) [S13]

## Mechanistic / biological rationale

ARHGEF2 is a **microtubule-associated, RhoA-specific guanine nucleotide exchange factor** that links **microtubule dynamics to actin cytoskeleton regulation**. That biology makes it well suited to control neural processes that depend on cell shape, polarity, movement, and structural remodeling. ([pmc.ncbi.nlm.nih.gov](https://pmc.ncbi.nlm.nih.gov/articles/PMC5428974/)) [S14]

In neural precursor cells, this mechanism plausibly explains why ARHGEF2 affects **mitotic spindle orientation** and thus the balance between **self-renewal and differentiation**. In the Ravindran study, ARHGEF2 deficiency shifted spindle orientation and perturbed cell fate decisions, consistent with a regulator of developmental patterning. ([pmc.ncbi.nlm.nih.gov](https://pmc.ncbi.nlm.nih.gov/articles/PMC5428974/)) [S15]

In neurons and neuroepithelium, ARHGEF2 acts through **RhoA/ROCK/myosin-related pathways** to influence **dendritic spine morphology**, **neurite retraction**, **apical constriction**, and **migration**. Those are classic regulatory processes in neural development and structural plasticity. ([pmc.ncbi.nlm.nih.gov](https://pmc.ncbi.nlm.nih.gov/articles/PMC2638734/)) [S16]

A newer cortical study also suggests ARHGEF2 can regulate neural differentiation through a less expected route: by affecting **Mettl14-dependent m6A RNA methylation** and downstream differentiation-associated transcripts such as **Npdc1** and **Cend1**. This broadens the mechanism from pure cytoskeletal control to **gene-expression/post-transcriptional regulation** in neural development. ([pmc.ncbi.nlm.nih.gov](https://pmc.ncbi.nlm.nih.gov/articles/PMC8185223/)) [S17]

## Bottom line

Yes—this claim is **well supported**, especially if “regulatory” is interpreted in the usual biological sense. ARHGEF2 has substantial evidence linking it to **regulated neural developmental and structural processes**, including neurogenesis, precursor fate control, neuronal migration, neurite remodeling, dendritic spine development, and neural tube morphogenesis. The weaker part is **homeostasis** specifically in the mature nervous system; that is not nearly as directly established. ([pmc.ncbi.nlm.nih.gov](https://pmc.ncbi.nlm.nih.gov/articles/PMC5428974/)) [S18]

## Cited sources

- **[S1]** 
            Homozygous ARHGEF2 mutation causes intellectual disability and midbrain-hindbrain malformation - PMC
         — https://pmc.ncbi.nlm.nih.gov/articles/PMC5428974/
- **[S2]** 
            Homozygous ARHGEF2 mutation causes intellectual disability and midbrain-hindbrain malformation - PMC
         — https://pmc.ncbi.nlm.nih.gov/articles/PMC5428974/
- **[S3]** Homozygous ARHGEF2 mutation causes intellectual disability and midbrain-hindbrain malformation - PubMed — https://pubmed.ncbi.nlm.nih.gov/28453519/?utm_source=openai
- **[S4]** Homozygous ARHGEF2 mutation causes intellectual disability and midbrain-hindbrain malformation - PubMed — https://pubmed.ncbi.nlm.nih.gov/28453519/?utm_source=openai
- **[S5]** 
            Homozygous ARHGEF2 mutation causes intellectual disability and midbrain-hindbrain malformation - PMC
         — https://pmc.ncbi.nlm.nih.gov/articles/PMC5428974/
- **[S6]** Arhgef2 regulates neural differentiation in the cerebral cortex through mRNA m6A-methylation of Npdc1 and Cend1 - PubMed — https://pubmed.ncbi.nlm.nih.gov/34142067/?utm_source=openai
- **[S7]** 
            AMPA receptor and GEF-H1/Lfc complex regulates dendritic spine development through RhoA signaling cascade - PMC
         — https://pmc.ncbi.nlm.nih.gov/articles/PMC2638734/
- **[S8]** 
            Discovery of long-range inhibitory signaling to ensure single axon formation - PMC
         — https://pmc.ncbi.nlm.nih.gov/articles/PMC5484694/
- **[S9]** 
            GEF-H1 functions in apical constriction and cell intercalations and is essential for vertebrate neural tube closure - PMC
         — https://pmc.ncbi.nlm.nih.gov/articles/PMC4038946/
- **[S10]** 
            Homozygous ARHGEF2 mutation causes intellectual disability and midbrain-hindbrain malformation - PMC
         — https://pmc.ncbi.nlm.nih.gov/articles/PMC5428974/
- **[S11]** 
            Arhgef2 regulates neural differentiation in the cerebral cortex through mRNA m6A-methylation of Npdc1 and Cend1 - PMC
         — https://pmc.ncbi.nlm.nih.gov/articles/PMC8185223/
- **[S12]** 
            GEF-H1 functions in apical constriction and cell intercalations and is essential for vertebrate neural tube closure - PMC
         — https://pmc.ncbi.nlm.nih.gov/articles/PMC4038946/
- **[S13]** Homozygous ARHGEF2 mutation causes intellectual disability and midbrain-hindbrain malformation - PubMed — https://pubmed.ncbi.nlm.nih.gov/28453519/?utm_source=openai
- **[S14]** 
            Homozygous ARHGEF2 mutation causes intellectual disability and midbrain-hindbrain malformation - PMC
         — https://pmc.ncbi.nlm.nih.gov/articles/PMC5428974/
- **[S15]** 
            Homozygous ARHGEF2 mutation causes intellectual disability and midbrain-hindbrain malformation - PMC
         — https://pmc.ncbi.nlm.nih.gov/articles/PMC5428974/
- **[S16]** 
            AMPA receptor and GEF-H1/Lfc complex regulates dendritic spine development through RhoA signaling cascade - PMC
         — https://pmc.ncbi.nlm.nih.gov/articles/PMC2638734/
- **[S17]** 
            Arhgef2 regulates neural differentiation in the cerebral cortex through mRNA m6A-methylation of Npdc1 and Cend1 - PMC
         — https://pmc.ncbi.nlm.nih.gov/articles/PMC8185223/
- **[S18]** 
            Homozygous ARHGEF2 mutation causes intellectual disability and midbrain-hindbrain malformation - PMC
         — https://pmc.ncbi.nlm.nih.gov/articles/PMC5428974/
